# A machine learning-based approach to identify reliable gold standards for protein complex composition prediction

**DOI:** 10.1101/2023.10.25.564023

**Authors:** Pengcheng Yang, Youngwoo Lee, Daniel B. Szymanski, Jun Xie

## Abstract

Co-Fractionation Mass Spectrometry (CFMS) enables the discovery of protein complexes and the systems-level analyses of multimer dynamics that facilitate responses to environmental and developmental conditions. A major challenge in the CFMS analyses, and other omics approaches in general, is to conduct validation experiments at scale and develop precise methods to evaluate the performance of the analyses. For protein complex composition predictions, CORUM is commonly used as a source of known complexes; however, the subunit pools in cell extracts are very rarely in the assumed fully assembled states. Therefore, a fundamental conflict exists between the assumed multimerization of the CORUM “gold standards” and the CFMS experimental datasets to be evaluated. In this paper, we develop a machine learning-based “small world” data analysis method. This method uses size exclusion chromatography profiles of predicted CORUM complex subunits to identify relatively rare instances of fully assembled complexes, as well as bona fide stable CORUM subcomplexes. Our method involves a two-stage machine learning approach that is designed to leverage evolutionarily conserved sequences among CORUM subunits and integrate it with size exclusion chromatography profile data from CFMS experiments. The generated gold standards are evaluated by both statistical significance and size comparison between calculated and predicted complexes. We expect these gold standards to serve as improved benchmarks to assess the overall reliability of CFMS-based protein complex composition predictions.

## Introduction

Co-fractionation mass spectrometry (CFMS) is a high-throughput, mass spectrometry-based protein quantification coupled with biochemical fractionation methods to analyze protein complex compositions under non-denaturing conditions. This “guilt by association” method was initially used to predict protein organelle localization based on co-elution with known marker proteins (1) and was subsequently applied to predict protein interactors (2, 3). The technique has evolved from the principle that proteins present in a stable complex co-migrate independent of the separation method used. In plant systems, CFMS has been extensively employed for the determination of apparent masses, localization, and compositions of protein complexes across a wide variety of species and tissue types, including leaves, roots, and flowers (4–9), as well as in organelles like chloroplasts (10) and mitochondria (11, 12). CFMS has broad use as a valuable tool to analyze protein complex dynamics, including circadian changes (13), protein-ligand interactions (14, 15), and multimerization variants across plant species (16). A remaining challenge is to determine the appropriate data types and profile analysis methods to generate the most accurate protein complex composition predictions (17, 18). A major impediment to progress in this area is the lack of a large set of reliable gold standards to evaluate prediction accuracies.

Protein complex prediction methods vary wildly among different studies. Many are achieved by using multiple metrics from external data sources that could inform multimerization behaviors. Examples include using mRNA co-expression or co-citation information via machine learning classifiers (3, 9, 18–20) or integration of existing protein interaction predictions from orthogonal approaches (4–6, 16, 21–24). There is no clear-cut agreement on the effectiveness of those strategies for protein complex predictions (18). All approaches rely on known protein complexes as gold standards from a reference database, like the CORUM mammalian protein complex database (25). Gold standard protein complex datasets are critical because they define accuracy measures, e.g., the precision and recall, of protein complex predictions. Inaccurate gold standards result in a wrong validation dataset, misleading the prediction model to unreliable predictions.

The CORUM protein complexes are widely used as assumed gold standards to evaluate protein complex predictions (18). CORUM complex subunits can be identified across species using InParanoid (26) or EggNOG (27) that accurately map orthologs between species (9, 20, 22). During the evaluation of complex predictions, pairs of subunits in the reference CORUM complexes comprise a positive set of protein-protein interactions. Negative interactions are created from proteins not present in the positive interaction set. These positive and negative sets have been used for training and testing computation methods of protein complex predictions (9, 18, 20, 28). This approach assumes that CORUM complexes are fully assembled in the cell extracts that are used as the input for CFMS experiments. However, in our previous CFMS experiments (6, 16, 22, 23), many CORUM complex subunits have an apparent mass similar to their theoretical monomeric mass. Furthermore, in several recent publications, weak correlations were reported among subunits of CORUM complexes in the CFMS datasets (6, 16). Pang et al. (18) displayed probability density plots of the Pearson correlations between fractionation profiles of shared subunits of CORUM human complexes (Figure 2 in (18)), in which the density plots have a mode around 0 correlation values. A similarly low correlation among subunits of CORUM complexes was reported in Extended Data Figure 2 of Wan et al. (9, 18, 20, 28). These plots and our experimental data indicate that the most abundant cellular pools of CORUM complex subunits, which are what are primarily detected in LC/MS, are not necessarily in the fully assembled state. In other words, subunits of orthologous proteins from a CORUM complex do not always co-elute, suggesting that CORUM complexes are unlikely to be reliable gold standards for protein complex predictions.

In this paper, we aim to demonstrate the issue with the current practice of using CORUM as knowns and to develop a machine learning method for identifying reliable CORUM complexes or subcomplexes. These can subsequently be used as gold standards for CFMS analysis. More specifically, we collect plant proteins that are orthologous to subunits of a CORUM complex. Next, we conduct a “small world” analysis, examining the CFMS elution profile patterns of every orthocomplex one by one. In this context, the “small world” analysis refers to the implementation of our machine learning method within each orthocomplex.

Subcomplex predictions are generated using a robust unsupervised machine learning method, namely self-organizing map (SOM). We use a limited set of well-known rice complexes to train the algorithm, taking advantage of its unsupervised nature. Following the unsupervised learning, two additional validation steps are employed to enhance the subcomplex predictions. Firstly, statistical significance analysis is performed on the predictions using Monte Carlo simulations. Secondly, we compare the apparent mass of a predicted subcomplex with the calculated mass derived from its subunits.

This multifaceted procedure identifies a gold standard, defined as a CORUM complex or subcomplex that is composed of subunits with analogous CFMS elution profiles, is statistically significant, and has consistent apparent mass and calculated mass. The machine learning approach is applied to identify gold standards for two species, rice and Arabidopsis, providing evidence of its generalizability. Additionally, we applied our gold standards identified here, along with the metrics of intactness and purity, to demonstrate the method’s potential for evaluating a new set of de novo protein complex predictions.

In summary, the machine learning-based approach is designed to tackle the issues of the current practice of using CORUM as gold standards. The small-world analysis is developed to integrate evolutionarily conserved information from CORUM with co-elution profile data from CFMS experiments. These two information sources complement each other, and their integration enables the prediction of accurate gold standards.

## Materials and Methods

### External SEC data acquisition and filtering

In this study, we used a published rice CFMS dataset, which was previously used for protein complex predictions (16). The data were downloaded from the <u>Supplemental Data Sets S2</u> and <u>S3</u> in the publication (16). The dataset contains two replicates of size exclusion chromatography (SEC) fraction profiles and protein apparent mass (*M*_app_: measured mass from SEC experiments), monomeric mass (*M*_mono_), and multimerization state (*R*_app_: multimerization determinator, which is defined as a ratio of *M*_app_ to *M*_mono_) (4, 29). We applied data filtering criteria, consistent with established guidelines from the existing literature (16, 22, 23), to discern and select reliable protein profiles for the discovery of gold standard complexes. These filtering methods have been shown to be robust for large scale multimerization analyses conducted using many different tissue types and species (4, 6, 16, 22, 23). More specifically, reproducible protein profiles were selected if their elution peak locations between the two replicates were within 2-SEC fractions. This criterion was justified by the consistent observation that about 90% of peaks were within this 2-SEC range across the two replicates of the SEC experiment (16, 22, 23). If the reproducible peak was located at the first (void) fraction measured by blue dextran (4, 23) in all replicates, it was defined as an unresolvable peak and removed from the dataset. When a protein profile had more than one reproducible peak, we deconvolved these peaks into individual peaks that represent different multimerization states (6, 16, 23). Additionally, if a peak was separated from a multiple-peak protein, it was treated as an individual profile in the following data analysis if its mean *M*_app_ from the two replicates was less than 850 kDa. The criterion of less than 850 kDa indicates the peak in at least one replicate is a resolvable peak over the SEC column (16). These individual peak profiles were annotated with a numerical suffix appended to the protein name to indicate peak numbers for each of the two replicates in the Supplemental Tables.

### Interkingdom ortholog mapping for CORUM orthocomplex assignment

It is commonly assumed that those large, evolutionarily conserved complexes carry out similar functions in all eukaryotic cells. Given that many of these complexes are involved in essential functions, their presence in all cells is anticipated. Additionally, we assume similarity in protein assemblies and stoichiometries across different species. To infer human-to-rice orthologs (Figrue 1A), human proteome search database (9606.fasta; Uniprot version) was directly obtained from InParanoid 8 website (https://inparanoidb.sbc.su.se/), and rice proteome (Osativa_323_v7.0.protein_primaryTranscriptOnly.fa; MSU version) was downloaded from Phytozome V12 database (https://phytozome-next.jgi.doe.gov/). We chose to use InParanoid (26), software Version 4.1, to infer orthologs between human and rice species, as InParanoid is a tool and database that robustly define orthology relationships across kingdoms. The inferred orthologs were reported in Supplemental Table S1A. To identify rice orthologous complexes (orthocomplexes) corresponding to CORUM complexes, human protein complexes were downloaded from the CORUM database (http://mips.helmholtz-muenchen.de/corum/).

According to the ortholog groups specified in Supplemental Table S1A, the subunits in the human protein complexes were converted to their corresponding rice orthologs (Supplemental Table S2). A plant species has experienced polyploidization and gene duplication, creating different levels of genetic redundancies across species (30). This genome-wide complexity makes the ortholog assignment challenging. It is common that multiple rice orthologs/paralogs were mapped to a single human ortholog (Figure 1A). In such cases, we treated all rice paralogs as members of a rice orthocomplex but considered them a single “ortho-paralog” group to calculate the rice proteome overlap with the human CORUM complexes. The subunit coverage of a rice orthocomplex is defined as the ratio of the number of subunits with inferred orthologs to the total number of subunits of a CORUM complex. Those complexes with a subunit coverage greater than or equal to 2/3 were chosen as high-coverage rice orthocomplexes for gold standard complex predictions.

**Figure 1.**
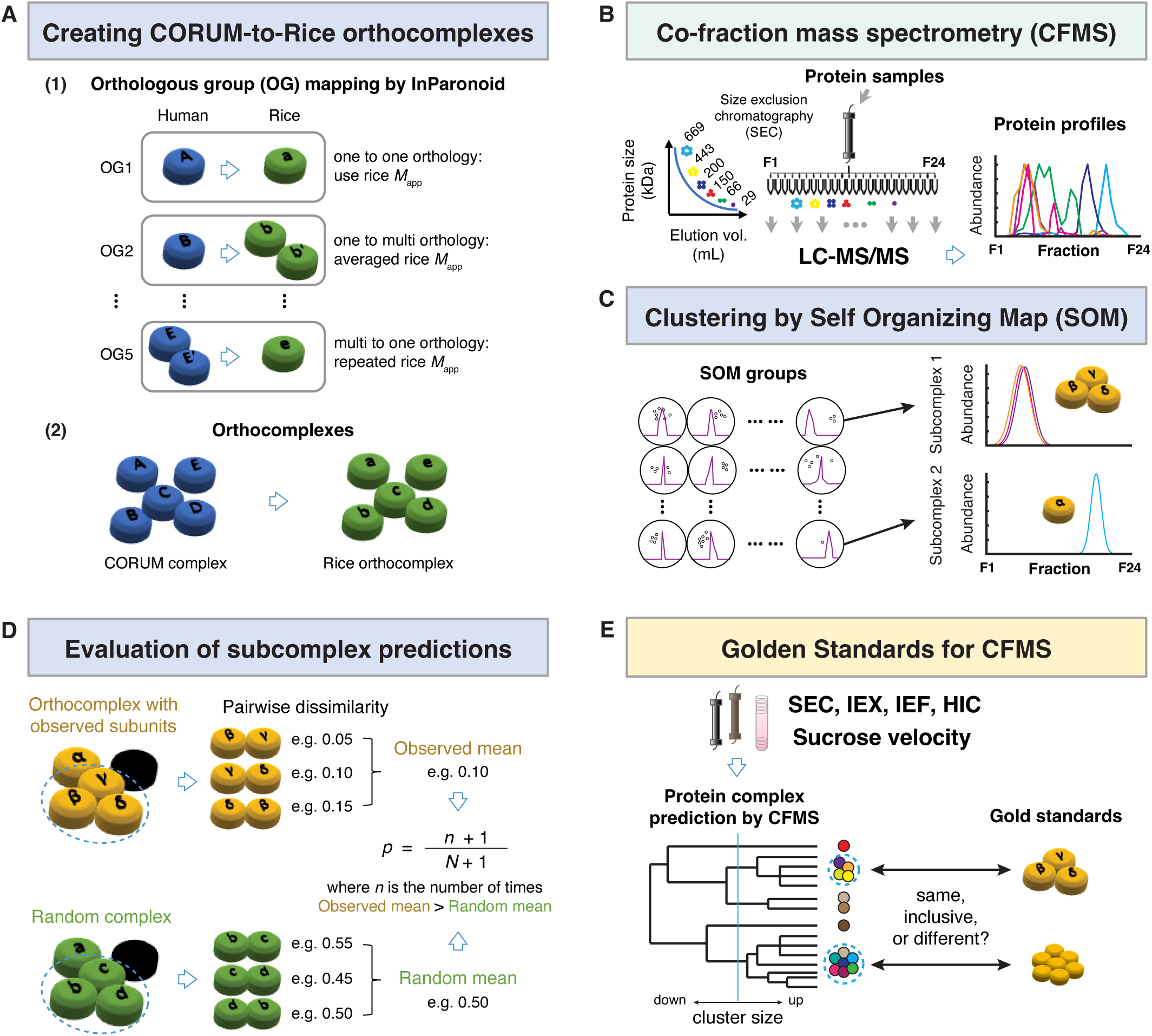
Identification of bona-fide gold standards to evaluate prediction accuracies in co-fractionation mass spectrometry-based protein complex discovery in plant species. **A**, Human and rice proteins were assigned into orthologous groups (OGs) using InParanoid algorithm (25), and rice orthocomplexes (green) were built from CORUM complexes (blue) based on the OGs. **B,** SEC profiles in CFMS analysis were used in the identification of gold standard complexes. **C,** SOM was used to cluster protein profiles generated in B. Each code/group (outside circle) contains a group profile (purple) representing profiles of all protein members (dots). **D,** Subcomplex predictions in SEC datasets are evaluated via a statistical bootstrap *p*-value calculation. Among experimentally detected subunits (yellow) in a rice orthocomplex, subunits with similar SEC profiles are clustered in a subcomplex (dotted line). Profile similarity scores are calculated between all possible pairs of subunits in the subcomplex and then are averaged to get the mean dissimilarity of the subcomplex. At the same time, the same number of proteins observed in the orthocomplex are sampled from randomly generated plant orthocomplex (green). The random mean is calculated as mean dissimilarity for pairs of proteins in the random subcomplex. The *p*-value for each subcomplex is calculated as the fraction of times the observed mean is larger than the random mean. **E,** Predicted gold standards can evaluate protein complex prediction results by CFMS performed with any types of biochemical separations.

### Integration of rice orthocomplexes into rice SEC data

After assigning rice orthocomplexes through the mapping of human CORUM complexes, we integrated this orthocomplex information with the experimental SEC profile data. We identified useful rice orthocomplexes that contained at least two subunits in the SEC profile data. Subsequently, we refined these orthocomplexes by eliminating redundant orthocomplexes consisting of identical rice subunits, smaller orthocomplexes that were subsets of larger ones, and orthocomplexes comprising solely rice subunits mapped to a single human ortholog.

### Experimental Design and Statistical Rationale

#### Distance metric

In the small-world analysis, we utilized a distance metric that combines two similarity measurements of a pair of proteins in an orthocomplex, i.e., the correlation of the fractionation profiles between the pair and the distance of peak fraction locations between the pair. First, a weighted cross-correlation (*WCC*) between a pair of protein profiles was calculated by the following formula (31)

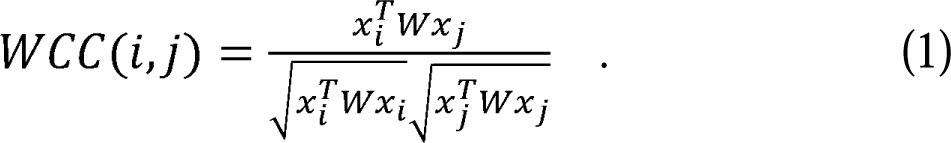

Here, *x_i_* and *x_j_* were two column vectors representing the SEC profiles of the two proteins in the orthocomplex. The weight matrix, denoted as *W*, was constructed with ones on the main diagonal, but elements on the sub-diagonal and the super-diagonal decreased proportionally as they moved away from the main diagonal. All elements outside of the sub- and super-diagonals were set to zero. We set the bandwidth of *W* to be 2 and used a weighting function of (1 − distance/3). That is, there were 2 sub-diagonals and 2 super-diagonals in the weight matrix *W*, and the weights on the first sub/super-diagonals were 2/3, while the weights on the second sub/super-diagonals were 1/3. This choice of parameters enables *WCC* to extract the similarity information across the peak profiles within a neighborhood of 2 fractions. A distance (*WCCd*) based on the weighted cross-correlation for a pair of proteins was given by

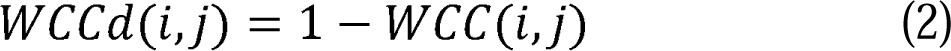

Another distance metric between the pair of proteins was calculated from their corresponding peak locations as follows:

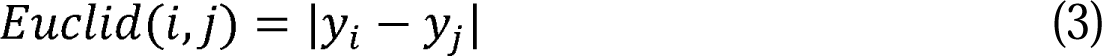

where *y_i_* and *y_j_* were the peak fraction locations of a pair of proteins, obtained using the Gaussian peak fitting algorithm (6). These peak locations had been standardized by dividing them by the largest peak location of a complex, ensuring they fall within the range of 0 to 1 for the Euclidean distance calculation in Formula (3). Both distance measurements were computed for the two replicates of the SEC data. An overall distance between two proteins was defined as

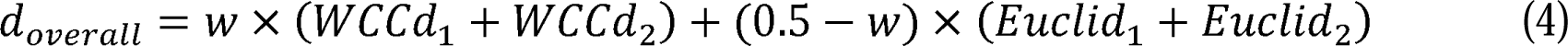

where *WCCd_1_* and *WCCd_2_* were the weighted cross-correlation distance, and *Euclid_1_* and *Euclid_2_* were the Euclidean distance between the peak locations of the pair of proteins. There were two distances indicated by the subscripts 1 and 2 for the two SEC data replicates. The combination weight *w* was obtained by model training described in the following.

#### Pre-clustering evaluation of orthocomplexes

Before conducting an unsupervised clustering analysis as described next, we first assessed whether an orthocomplex formed a single cluster. This step also helped us evaluate which orthocomplexes were potentially fully assembled CORUM complexes, as a fully assembled orthocomplex must be a single cluster. More specifically, in cases where an orthocomplex consisted of only two subunits in the SEC profile data, these small orthocomplexes were excluded from the subsequent clustering analysis and were labeled as “only 2 proteins in the complex before clustering” in the result tables. However, as later defined, these instances were subjected to a bootstrap *p*-value calculation to determine if the two subunits had the potential to form a complex. If the resulting *p*-value fell below the 5% threshold, the two subunits had a significantly small distance and hence were predicted to form a single cluster.

For orthocomplexes with more than two subunits from the SEC data, we assessed whether all subunits exhibited similar profiles, which was indicated by significantly small pairwise distance metrics. Two distance metrics, defined in Formula (2) and Formula (3), were utilized for this evaluation. If the distances calculated for all subunit pairs within an orthocomplex fell below the 5th percentile threshold of an empirical distribution, we inferred that all subunits within that orthocomplex were part of the same complex. The empirical distribution was constructed by calculating pairwise distances among all proteins within the SEC data. We systematically examined both distance metrics, and any orthocomplexes that met the 5th percentile criterion for either distance metric were predicted to form a single cluster. In Supplemental Table S3, column T, those meeting the *WCCd* distance criterion were labeled as “all proteins defined in the complex by similarity before clustering”, while those meeting the Euclidean distance criterion were labeled as “all proteins defined in the complex by Gaussian peak distance before clustering”.

Note that the 5% threshold for claiming statistical significance is applied after confirming that two protein subunits belong to the same CORUM orthocomplex, as our method involves CORUM ortholog mapping initially. Since the likelihood of two random proteins being in the same CORUM orthocomplex is expected to be low, the actual *p*-value threshold for statistical significance is much lower than 5%. Furthermore, some of these orthocomplexes were potentially fully assembled CORUM orthologs, as our analysis indicated they formed a unified complex that could not be separated. To rigorously confirm the full assembly, we further examined whether these inseparable orthocomplexes consisted of all CORUM subunits as well as if the apparent mass observed in the SEC experiment agreed with the calculated mass summing over all subunits (more details in the size evaluation section below). Our confirmation of orthocomplexes as fully assembled was substantiated by satisfying these additional two criteria.

#### Two-stage clustering algorithm for subcomplex prediction

Only a very small number of rice orthocomplexes were found to be fully assembled. For the majority of the orthocomplexes, we conducted a small-world clustering analysis on each one individually. The small-world analysis employed a two-stage clustering algorithm with the distance metric defined in Formula (4). Our two-stage procedure is composed of a clustering step by self-organizing map (SOM) algorithm (32) and a cluster merging step by the affinity propagation (AP) algorithm (33). SOM is a machine learning method for clustering analysis. It is a specific type of neural network model that is designed to represent SEC profiles using robust features. As a result, the method exhibits resistance to random noises and possesses the advantage of robustness against data measurement errors or missing values. Employing the SOM algorithm, the subunits of a rice orthocomplex were clustered, leading to the formation of distinct subgroups. Importantly, these distinct subgroups were predicted to represent subcomplexes within the larger orthocomplex.

To yield the optimal final clusters, a relatively large number of clusters was initially selected and then followed by the merging of resulting clusters using the AP algorithm. For this two-step clustering analysis, we fine-tuned three parameters of the clustering algorithms, including the weight w on *d_overall_* in Formula (4), the number of clusters in SOM, and the merging threshold for the AP algorithm. Choosing tuning parameters for machine learning algorithms is widely acknowledged as a challenging task. The standard procedure involves simulation experiments with validation data containing known information. Specifically, we started with a large number of clusters in SOM, close to the number of subunits in an orthocomplex. The weight w on *d_overall_* was chosen from a grid between 0 to 0.5 (the range of w as indicated in Formula (4)) with an increment of 0.05. The merging threshold of the AP algorithm ranged from 0.7 to 0.95, with larger values leading to more fine-grained clusters during the merging process that ends up with more clusters.

We used four well-known rice complexes (19S proteasome, 20S proteasome, 14-3-3 hetero/homooligomers, and the exosome) to determine the three tuning parameters (Supplemental Table S3B). We set these parameters to ensure that subunits for each of the four known complexes were clustered together. These known complexes are part of four CORUM orthocomplexes with the following IDs: PA700−20S−PA28 complex (CORUM ID: 193), RAF1−MAP2K1−YWHAE complex (CORUM ID: 5873), HSF1−YWHAE complex (CORUM ID: 2145), and Exosome (CORUM ID: 789), as indicated in Supplemental Table S3. Finally, the obtained tuning parameters were set up as the default parameter values in our computation package for subcomplex discoveries in rice as well as Arabidopsis.

The trained two-stage clustering algorithm was implemented across the rice orthocomplexes one by one. For each rice orthocomplex, the two-stage clustering approach could yield clusters with multiple members or a single member. Clusters with multiple members were designated as subcomplexes, while single-member clusters were labelled as “singletons”. In cases where all members of a predicted subcomplex were mapped to a single human ortholog, it was called an “ortho-paralog subcomplex”. Altogether, the results were labeled as “subcomplex i”, “subcomplex (ortho-paralog) i”, and “singleton i”, where i is a subcomplex identification, in the Supplemental Tables and Figures.

Furthermore, based on the external input data, proteins with small *R*_app_ values, specifically *R*_app_ ≤ 1.6, were classified as predicted monomers (16, 22, 23). This somewhat arbitrary threshold is applied to reduce the frequency of false positive golden standard predictions and could be set to a higher value if desired. The criterion of *R*_app_ ≤ 1.6 was used to exclude potential dimers from the pool of putative monomers. When our algorithm identified proteins with *R*_app_ ≤ 1.6 as singletons, we deemed these monomer predictions accurate. This is because proteins with the small range of *R*_app_ were anticipated to be monomers, and the clustering algorithm affirmed this anticipation.

#### Statistical significance and size evaluation of identified subcomplexes

The subcomplexes, as well as the potentially fully assembled complexes, identified through the small-world analysis, were evaluated using statistical *p*-values and a comparison of apparent mass (*M*_app_) and calculated mass (*M*_calc_) to identify the gold standards (Figure 1D). First, a bootstrap *p*-value for a subcomplex/complex was calculated using Monte Carlo simulations. The mean distance for pairs of proteins in a subcomplex was calculated using the overall distance, *d_overall_*, in Formula (4). A random complex was generated by randomly sampling the same number of proteins in the identified subcomplex from all SEC profiles in the CORUM orthocomplex profile dataset. Protein subunits in the random complex were not related, hence their distance *d_overall_* followed a pure random distribution. The mean distance for pairs of proteins present in the random subcomplex was calculated. We ran the random sampling for a large number of times. The *p*-value was calculated as the fraction of times the random mean distance was smaller than the observed mean distance of the identified subcomplex (Figure 1D). Additionally, false discovery rates (FDR) for the *p*-values were calculated to address the multiple testing issue associated with identifying hundreds of subcomplexes.

A random complex was generated by randomly selecting protein subunits from all proteins with SEC profiles. Because of the randomization process, an empirical distribution derived from the random complexes was not affected by factors such as protein abundance. On the other hand, the empirical distribution was affected by the size of the complex. In the case of small random complexes, such as those of size 2, 3, or 4, the random distribution exhibits a stretched shape. When a random complex contains more than 5 subunits, the empirical distribution was more similar to a Gaussian curve (Supplemental Figure S4).

For the *M*_app_ and *M*_calc_ comparison, the *M*_calc_ of an identified subcomplex/complex was obtained by summing the monomeric mass values of all subunits in the given complex. For an orthocomplex that contained multiple rice paralogs mapping to a single human ortholog, their *M*_mono_ values were averaged in calculating the complex *M*_calc_ (Figure 1A). Notably, information regarding the stoichiometry of subunits within a complex is rarely available (25). Also, the purpose of this paper is to determine which ones form fully assembled or sub complexes with a stoichiometry that is identical to human. Thus, we manually curated the most common stoichiometry information in the RCSB Protein Data Bank (RCSB PDB) for the selected rice orthocomplexes (Supplemental Table S3, columns X and Y) and incorporated this stoichiometry information when calculating *M*_calc_.

For each subcomplex/complex with *p*-value < 0.05, its *M*_app_ was compared with its calculated mass *M*_calc_ (Supplemental Table S4). Identified subcomplexes/complexes with similar *M*_app_ and *M*_calc_ were deemed gold standards. More specifically, *M*_app_ for a subcomplex/complex was given by

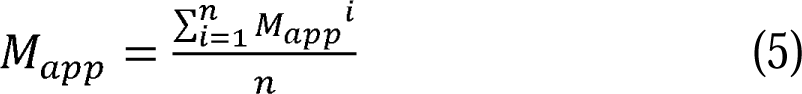

where *n* was the number of subunits in the subcomplex/complex, and *M*_app_*^i^* was the apparent mass of a subunit. Let *M*_calc_ denote the sum of the monomeric masses of subunits within the identified subcomplex/complex. We determined a subcomplex/complex with *p*-value < 0.05 as a gold standard if it satisfied the following criterion:

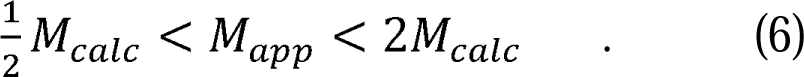

This criterion signified that the relative difference between the SEC experiment mass (*M*_app_) and the mass calculated from the identified subcomplex/complex subunit composition (*M*_calc_) was constrained within a factor of 1. In other words, subcomplexes/complexes with *p*-value < 0.05 that exhibited matching *M*_app_ and *M*_calc_ values, as defined by this criterion, were selected as gold standards. Notably, some of these gold standards originated from the pre-clustering complexes. When their *M*_app_ and *M*_calc_ values matched, they were confirmed as fully assembled CORUM orthocomplexes.

#### Use of gold standards to evaluate de novo complex predictions

We used the selected gold standards to evaluate protein complex predictions. Typical evaluations in the literature were merely based on positive and negative sets of protein-protein interactions (PPI). The use of false positive and false negative, or equivalently precision and recall, to assess CFMS data is limited conceptually to the inference of PPIs. We consider this practice inadequate because the information of PPI is rooted in pairwise interactions, while the information of a protein complex should be from a group of subunits. In other words, the evaluation of complex prediction should be grounded in the composition of the complex, a group of multiple subunits, not pairwise links.

To overcome the limitation with the use of precision and recall, we defined two complex-centric metrics: intactness and purity. Intactness and purity were used to measure the similarity between two sets of protein complexes, one set of gold standards and the other set of *de novo* predictions. Let *C* denote a list of de novo protein complex predictions,

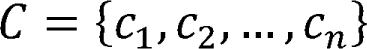

where *c_i_* is a predicted complex with multiple subunits. Analogously, let *G* denote our list of gold standards,

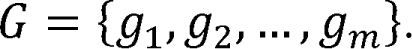

Without loss of generality, we assume *G* is a subset of *C*, given that gold standards typically comprise a smaller set compared to the set of proteins to be predicted. In instances where the gold standard set is not entirely encompassed within the larger set of proteins for complex predictions, we will employ *G* ∩ *C* as the gold standard set. For each gold standard protein complex, *g_i_* ∈ G, we define the intactness and purity as

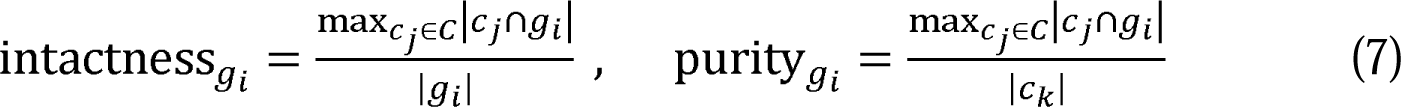

where *c_k_* is the protein complex in *C* with the most overlaps with *g_i_*, and | | denotes the number of proteins of the corresponding set. Note that when the maximum overlap between a gold standard complex and the predicted complexes, max_*cj}c*_❘*c_j_* ∩ *g_i_*❘, involves only one subunit, the calculation of intactness is not meaningful. In such cases, the corresponding gold standard is completely split into different predicted complexes; thus, the intactness values are denoted as “Not Defined” (as in Supplemental Table S5).

The intactness metric quantifies how well the predicted complex captures the entirety of the proteins that are present in the gold standard complex. A high intactness value for a specific gold standard complex indicates that the predicted complex adequately represents the entire protein composition of the gold standard. In contrast, a lower intactness value suggests that the predicted complex might only partially capture the proteins within the gold standard complex. The purity metric assesses the specificity of a predicted protein complex with respect to a particular gold standard complex. It evaluates the proportion of proteins within the predicted complex that are also part of the gold standard complex. A higher purity value for a specific gold standard complex indicates that the predicted complex primarily comprises proteins characteristic of that gold standard. Conversely, a lower purity value implies that the predicted complex contains proteins beyond the gold standard complex.

The size of the gold standard complexes affects both the intactness and purity metrics. Larger gold standards require more accurate predictions to achieve high intactness values, while smaller gold standards would attain high intactness values more easily. Conversely, larger gold standards could lead to higher purity values due to the potential for greater protein overlaps. These two metrics collectively offer an evaluation of the accuracy of protein complex predictions.

### Dimerization prediction by AlphaFold Multimer on COSMIC^2^

To run the AlphaFold Multimer software package v2.2.0 (34), the COSMIC^2^ cloud platform was used (35). Protein sequences for each dimeric subcomplex were obtained from the rice proteome file *Osativa_323_v7.0.protein.fa* (36) in Phytozome V12 (37), and then searched against the full database (full_dbs) as default. Ranking confidence scores, a weighted combination of interface predicted Template Modeling score (ipTM) and predicted Template Modeling score (pTM), were used as model confidence metrics. The averaged model confidence score of the top 5 predicted models was used to evaluate predicted dimeric subcomplexes.

### Arabidopsis subcomplex predictions using the two-stage clustering

The identical analysis pipeline was employed to identify Arabidopsis subcomplexes. We replicated all analysis steps using an Arabidopsis CFMS dataset (23), including CFMS data quality filtering, ortholog mapping for CORUM complexes in Arabidopsis, and the two-stage clustering analysis on the Arabidopsis SEC profile data. Crucially, we applied the same set of tuning parameter values for the clustering algorithm in Arabidopsis subcomplex predictions.

### Statistical tests and data analysis

Statistical analysis was performed using R version 4.2.0 (38) on RStudio 2022.07.1 (39). The Flexible Self-Organizing Maps in Kohonen 3.0 package for R (40) and the APCluster package for R (41) were implemented for the SOM and AP algorithms, respectively. Gaussian fitting code (https://github.com/dlchenstat/Gaussian-fitting) was run on MATLAB (R2022a). Microsoft Excel on Office 365 for Mac was used to organize and display the analyzed data.

## Results

Figure 1 presents an overview of the workflow to identify true gold standards by integrating the CORUM database with the SEC profile data from rice tissue extracts. The overall scheme was to identify rice orthocomplex subunits based on subunit overlap in the plant and animal kingdoms and sequence similarity between the individual complex subunits. The SEC profile data from rice were mined for reproducible *M*_app_ measurements for the orthocomplex subunits, and the developed machine learning method was applied to group similar protein profiles from the SEC data in order to identify fully- and partially-assembled complexes in the rice cell extract. The predicted subcomplexes/complexes were further evaluated using statistical significance and size comparison between calculated mass and measured apparent mass. The validated subcomplexes/complexes were defined as gold standards. These gold standards serve as reliable references for the evaluation and analysis of future protein complex predictions.

### Generating rice orthocomplexes from CORUM

To specifically analyze known complex and construct a reference library of putative gold standards genes, we identified CORUM orthologs in rice using the InParanoid algorithm (26). InParanoid search compared 20,834 human protein sequences with 42,160 proteins in rice. The algorithm assigned 5,363 human proteins and 8,178 rice proteins into 3,131 distinct orthologous groups at the whole proteome level (Supplemental Table S1A). The higher number of orthologs from rice was due to an elevated gene copy number compared to humans (30). We next created predicted rice orthocomplexes from CORUM human complex compositions (25) using the ortholog dataset generated above. As discussed in the ortholog mapping section of the Methods and Materials, there were complicated scenarios when we assigned orthologs between human and rice. Frequently, multiple rice orthologs/paralogs were mapped to a single human ortholog, and vice versa (Figure 1A(1)). We constructed a CORUM orthocomplex by including all rice paralogs that were orthologous to each human subunit in the corresponding CORUM complex. Among the 3,047 human complexes curated in CORUM, 1,964 (64.5% of total CORUM complexes) had at least one subunit orthologous to one or more rice proteins, 920 out of the 1964 rice orthocomplexes exhibited subunit coverage greater than or equal to 2/3. Four hundred and thirty-six (14.3% of total CORUM complexes) of the 1,964 rice orthocomplexes displayed completely conserved subunit compositions and are expected to have very similar core functions in the cell (Figure 2A; Supplemental Figure S1; Supplemental Table S2).

**Figure 2.**
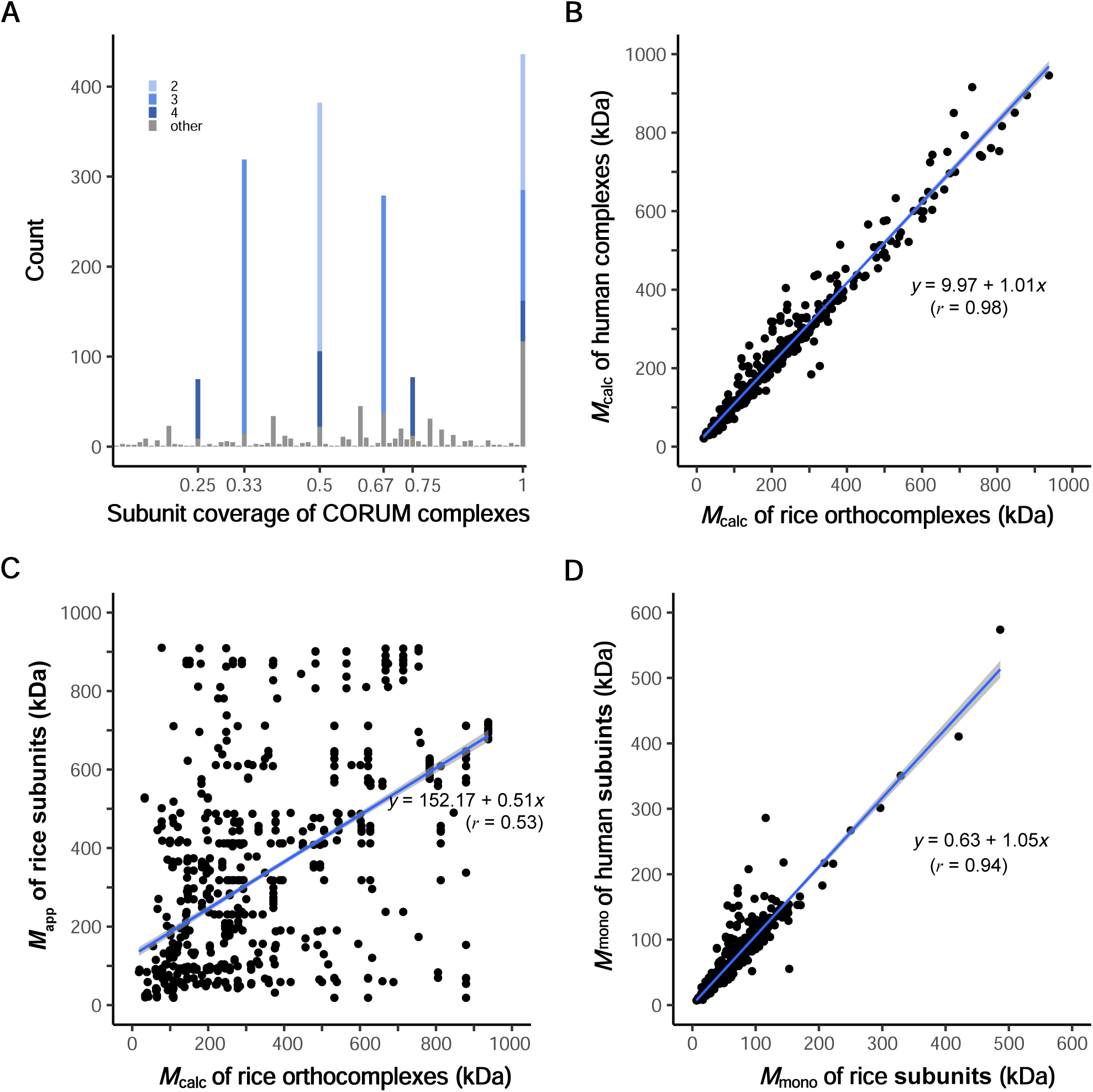
Assumed CORUM gold standard complexes do not exist in a fully assembled state in plant species. **A**, Genomic level subunit coverages of CORUM complexes to rice orthocomplexes. The coverages are defined as the ratio of the number of subunits in a rice orthocomplex to the number of subunits in its orthologous human CORUM complexes. The genome coverages of 1964 rice orthocomplexes were calculated and plotted at different subunit coverages. The sharp peaks are influenced by the number of members within the CORUM complexes. The various colors within each peak represent the relative proportions of different complex sizes covered. **B,** Conserved predicted masses of human complexes and rice orthocomplexes. *M*_calc_ of 436 rice orthocomplexes with 100% subunit coverage to CORUM complexes were plotted. *M*_app_ values of the one-to-multi rice orthologs were averaged in *M*_calc_ calculation (Figure 1A). **C,** CORUM complex subunits rarely exist in a fully assembled complex. A scatter plot presents protein complex conservation in size between CORUM complexes and rice orthocomplexes. *M*_app_ values of subunits of the 258 rice orthocomplexes were obtained from the reference rice CFMS datasets. **D,** Conserved masses of human CORUM subunits and rice orthocomplex subunits. A scatter plot shows conserved *M*_mono_ values between human and rice subunits that assemble into the known complexes. The rice orthocomplexes with 100% subunit coverage to CORUM complexes were plotted.

To assess variability in the sizes of the rice and human orthologs, we compared predicted protein complexes within the 436 human and rice orthocomplexes that shared 100% subunit coverage. The *M*_calc_ value is determined by summing the monomeric masses of individual subunits within an orthocomplex. In Figure 2B, we illustrate that rice and human orthocomplexes exhibit similar complex sizes when considering *M*_calc_ (*r* = 0.986 and slope =1.01), indicating that most complex possess similar masses. Among those 436 orthocomplexes, 258 had detected subunit(s) in the SEC datasets. However, when *M*_calc_ values of these highly conserved rice orthocomplexes were compared to their subunit *M*_app_ values measured in the rice SEC experiments, low correlation and a regression line with slope much less than 1 were found (Figure 2C, *r* = 0.53 and *R*^2^ = 0.28, slope = 0.51). The lack of correlation and the low slope in Figure 2C indicate that the *M*_app_ of rice orthocomplexes from SEC experiments do not align with their calculated sizes. The low correlation shown in Figure 2C was not due to differences in *M*_mono_ of subunits in these 258 orthocomplexes, as we found a strong positive correlation between *M*_mono_ values for this subset of human and rice orthocomplex subunits (Figure 2D).

Furthermore, 25 proteins from the rice orthocomplexes with 100% subunit coverage had *R*_app_ ≤ 1.6 and hence were considered as likely monomers. The existence of monomeric subunit pools and partially assembled complexes can explain the large number of data points falling well below the diagonal in Figure 2C. Data points above the diagonal may reflect novel complexes in which CORUM orthocomplexes and/or subcomplexes interact with unknown proteins. These results are consistent with previous observations (6, 18, 20, 22) and indicate that CORUM subunits detected in CFMS experiments rarely agree with the predicted mass of the fully assembled state.

### Gold standard predictions

To identify reliable rice orthocomplexes, we extracted reproducible protein elution profiles from the reference rice SEC datasets (16). There were 3,426 proteins present in both of the two SEC replicates, and 197 had multiple peaks that arose when the protein existed in multiple multimerization states. We deconvolved the multiple peaks to generate 350 reproducible peaks, and a total of 2,618 protein subunits with *R*_app_ > 1 were used as the rice SEC reference profiles. We further curated rice orthocomplexes with a subunit coverage greater than or equal to 2/3 in the rice SEC profiles. Among the 920 orthocomplexes with subunit coverage greater than or equal to 2/3, 531 orthocomplexes have at least one rice subunit detected by the rice SEC reference profiles, and 287 out of 531 orthocomplexes have at least two rice subunits detected by the rice SEC reference profiles. After eliminating redundant orthocomplexes, 103 rice orthocomplexes were selected by integration of rice orthocomplexes into the rice SEC data for further analyses. The small-world analysis was performed across the 103 rice orthocomplexes one by one to predict the composition of subcomplexes. The results of the small-world analysis, including essential details such as cluster composition, singletons, statistical significance, and size evaluation, were reported in Supplemental Table S3 and illustrated in Supplemental Figures S2 and S3. More specifically, in Supplemental Figure S2, we plotted *M*_app_ versus *M*_calc_ of the identified subcomplexes after the small-world analysis, demonstrating the existence of partially assembled complexes in the rice cell extract. Additionally, SEC profile plots illustrating co-elution patterns of the small-world analysis results were generated in Supplemental Figure S3.

During the process of small-world analysis, subunits within each rice orthocomplex were clustered based on their profiles and distances. The outcome of this analysis included 162 subcomplexes/complexes from the 103 rice orthocomplexes (Figure 3A; Supplemental Figures S2 and S3; Supplemental Table S3). Random sampling was conducted a large number of times, specifically 134,350 times, which is equal to 50 times the total number of the rice SEC reference profiles in our dataset. Supplemental Figure S4 displays the random empirical distributions for various subcomplex sizes. The *p*-values and FDR values for the 162 subcomplexes/complexes were reported in Supplemental Table S3, with 112 of them having *p*-values less than 5% and the corresponding FDR values less than 7%. After removing redundancies from the list of 112, and discarding potential monomers (*R*_app_ ≤ 1.6), we identified 79 unique subcomplexes/complexes with small *p*-values (Supplemental Table S4).

**Figure 3.**
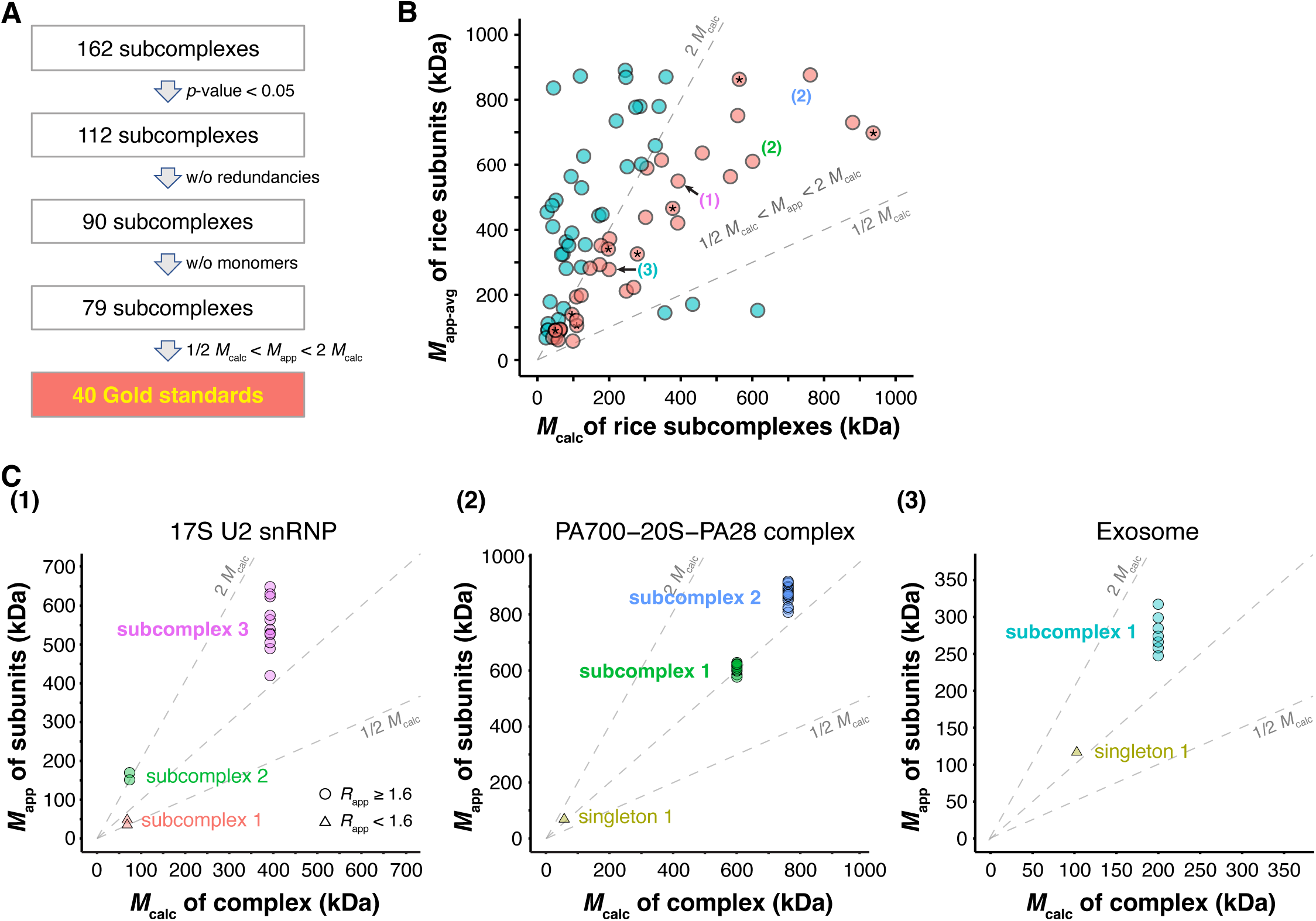
Useful gold standards near the diagonal. **A**, A process flow to identify gold standards from the small world analysis result. **B,** Gold standard subcomplexes/complexes of rice with matched *M*_calc_ and *M*_app-avg_ are rendered in pink. Asterisk (*) indicates fully assembled CORUM orthocomplexes. Numbers in parentheses point out predicted subcomplexes present in the corresponding panel in Figure 3C. **C,** *M*_calc_ values of predicted subcomplexes and *M*_app_ values of subunits of the subcomplexes. Gold standard subcomplexes (*p*-values = 0.0) were highlighted using bold text font.

We proceeded to compare their *M*_app_ and *M*_calc_ values using the criterion defined in Formula (6) to this set of 79 subcomplexes/complexes (Figure 3B; Supplemental Table S4). In comparison to Figure 2C, Figure 3B exhibits considerably fewer data points below the diagonal. This is because a subcomplex comprises fewer subunits than the CORUM orthocomplex. Therefore, the reduction in *M*_calc_ values brought them closer to the *M*_app_ values obtained from the SEC experiment. This result indicates that our algorithm successfully identified more reliable subcomplex formations. In Figure 3B, those subcomplexes/complexes with elevated *M*_app_ greater than two times *M*_calc_ might be attributable to undetected or unknown subunits in the complexes with non-spherical shapes, or unknown subunit stoichiometries. Among the set of 79 subcomplexes/complexes, 40 demonstrated substantial agreements between *M*_app_ and *M*_calc_, as shown in Figure 3B, meeting the criterion of Formula (6). These 40 subcomplexes or complexes were stable subcomplexes/complexes supported by both statistically significant *p*-values and consistent apparent masses (Figure 3B and 3C). They serve as gold standards to evaluate CFMS predictions. Additionally, within these 40 gold standards, our algorithm identified 8 fully assembled CORUM orthocomplexes. These fully assembled complexes met the criteria of containing all CORUM subunits, being statistically significant, and having similar *M*_app_ and *M*_calc_ values according to Formula (6). A summary list of these 40 gold standards is provided in Table 1, with additional details in Supplemental Table S5.

**Table 1.** Predicted subcomplexes that could be used as gold standards to evaluate CFMS based protein complex predictions.

To further justify the machine learning approach, we applied the method to randomly shuffled rice proteins. Specifically, we randomly permuted the protein IDs among the total of 42,160 rice proteins. This permutation generates a random dataset where a SEC profile (if available) does not correspond to its true protein but is instead assigned to a random protein. We conducted the entire process of the small-world analysis on the shuffled data, including CORUM ortholog mapping, SOM clustering analysis, statistical significance testing, and comparison of the *M*_app_ and *M*_calc_ values. Among multiple random simulations with shuffled data, we typically identify either no gold standard or one gold standard.

### Confirmation of monomer identification by the algorithm

Within the set of 103 CORUM orthocomplexes, there existed a list of 50 proteins with small *R*_app_ values (*R*_app_ ≤ 1.6), likely being monomers or multimers with a restricted type of binding partner. This list, derived directly from the original input data, served as prior information to evaluate and validate the accuracy of our algorithm in discerning monomeric proteins within orthocomplexes. None of the 50 potential monomers were predicted to form subcomplexes. Our analysis yielded three types of prediction results. Firstly, 14 were predicted as singletons with significantly large distance from the rest of the proteins within their respective CORUM orthocomplexes (Supplemental Table S6A). Secondly, our algorithm accurately identified 9 putative monomers, where all rice subunits were mapped to a single human ortholog (Supplemental Table S6B). Lastly, the remaining 27 putative monomers were clustered into different subcomplexes by our algorithm, but these subcomplexes had large *p*-values, indicating they were not expected to form discrete complexes.

### Structural validation of RNA polymerase II subcomplexes

Many novel subcomplexes were identified through our two-stage clustering approach in this study. Next, we used RNA pol II as an example to provide structural support for the consistency of our subcomplex predictions with existing knowledge. The dynamic assembly of RNA polymerase II complex (POL II) with general transcription factors into transcription preinitiation complexes is central to transcriptional control (42). Because the solved structure of RNA pol II in rice plants is not available, we mapped all rice detected subunits of rice ortho-RNA pol II complex over the human RNA pol II (PDB: 6O9L) in this analysis. The rice reference CFMS dataset included reproducible protein elution profiles for 12 subunits of the POL II complex (Figure 4A). RPB3, RPB9, and RBP11a had two reproducible, resolvable peaks. Our clustering assigned the POL II subunits into four different subcomplexes based on their elution profiles and reproducible peaks (Figure 4B). While subunits assigned in the subcomplexes 1, 3, and 4 showed similar profiles in both replicates (Supplemental Figure S3), TFIIB and RPB9 in the subcomplex 2 had a peak at around fractions 18-19. Thus, each subcomplex prediction was evaluated using a statistical bootstrap *p*-value calculation (Supplemental Table S3). As shown in the profiles, all the subunits within the subcomplexes 1, 3, and 4 had predicted *p*-values less than 0.01. Our analysis predicts that the low and high mass peaks of RBP3 and RBP11 correspond to a heterodimer and subcomplex 3, respectively. The prediction for the subcomplex 2 was insignificant (*p*-value = 0.56), and both subunits were predicted to be monomeric based on *R*_app_ ≤ 1.6.

**Figure 4.**
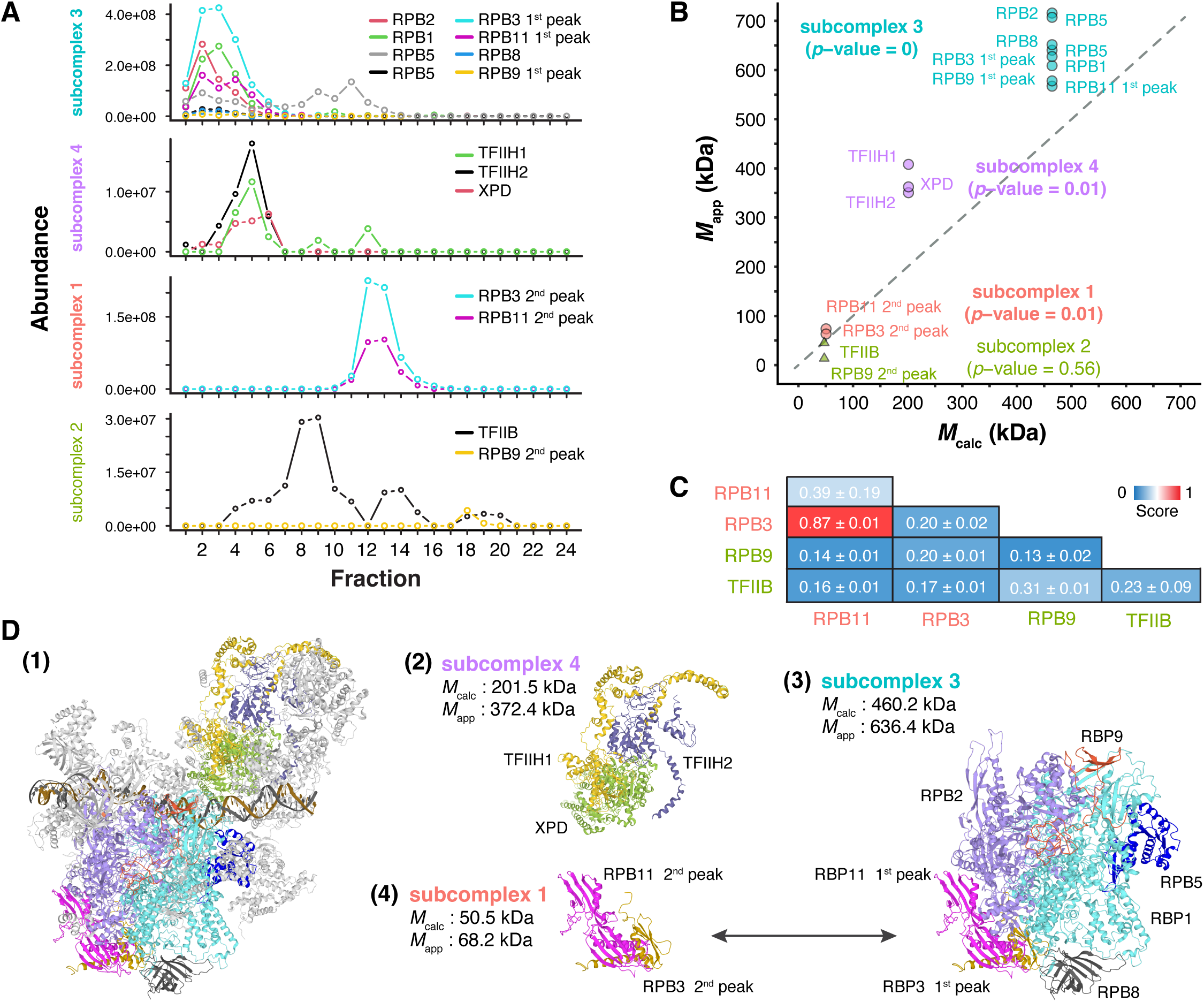
Validation of subcomplexes predicted in CORUM RNA polymerase II complex. **A**, Protein elution profiles of subunits in each predicted subcomplex. Profiles in another replicate can be found in the Supplemental Figure S3. **B,** *M*_calc_ values of predicted subcomplexes and *M*_app_ values of subunits of the subcomplexes. Circles indicate multimers, while triangles mean monomer (*R*_app_ ≤ 1.6). **C,** Dimerization prediction between subcomplex subunits by AlphaFold Multimer (34). AlphaFold Multimer was run on COSMIC^2^ to predict top5 models (35). Each value in the table is the mean of ranking confidence score (ipTM + pTM) ± standard deviation. **D,** Structural validation of predicted subcomplexes. Predicted subcomplexes are searched in RCSB Protein Data Bank (https://www.rcsb.org/). The structure of the fully assembled CORUM RNA polymerase II complex is available (PDB: 6O9L). (1) Undetected subunits in the CFMS dataset are colored gray. (2) – (4) Subcomplexes were predicted in B. *M*_calc_ and *M*_app_ of predicted rice subcomplexes are summarized right next to the subcomplex structures.

We also compared the *M*_calc_ values of predicted subcomplexes and *M*_app_ values of subunits of the subcomplexes to evaluate the predictions (Figure 4B). In addition to their significant *p*-values, *M*_app_ values of predicted subcomplex subunits were plotted nearby, supporting the presence of those three significant subcomplexes in the reference datasets. Similar to those with *M*_app-avg_ > 2 *M*_calc_, the slight skewness toward elevated *M*_app_ might be due to undetected or unknown proteins in the complex with non-spherical shapes. TFIIB in both replicates had similar peak locations (subcomplex 2 Fractions 12-15) to RPB3 and RBP11 peaks in the subcomplex 1 (Figure 4A and Supplemental Figure S3), indicating a potential association of TFIIB to the subcomplex 1.

Due to the lack of solved structures for rice POL II, we used AlphaFold Multimer (34) to test for potential direct interactions among the subunits present in subcomplexes 1 and 2. The mean of the confidence scores from the top 5 predicted models were calculated for the 10 possible combinations (Figure 4C). The heteromerization between RPB3 and RBP11 possessed the highest model confidence (ranking-confidence score: 0.87 ± 0.01), supporting the predicted subcomplex 1. All other heteromeric and homomeric models for TFIIB were also predicted with very low ranking-confidence scores (0.16 ± 0.01 ∼ 0.31 ± 0.01), indicating TFIIB may not be retained as a stable complex with other RNA pol II subunits in the cell extracts. The predicted three subcomplexes were also structurally validated (Figure 4D). The Cryo-EM solved structure of PIC (42) was downloaded from the Protein Data Bank (PDB: 6O9L). All the subunits in the predicted three subcomplexes are mapped onto the holoenzyme structure in a spatially plausible configuration (Figure 4D(1)). Subunits of the subcomplex 4 were assigned to the TFIIH complex (Figure 4D(2)), while those of the subcomplex 3 were assigned to the POL II complex (Figure 4D(3)). Each of them was also supported by the solved structures (PDB: 6DRD & 7NVW). The second peaks of RPB3 and RBP11 assembled into the subcomplex 1 with size consensus *M*_calc_ and *M*_app_. RPB3 and RPB11 heteromerization has been shown in Arabidopsis (43), and the subcomplex is at the core of POLII assembly (44). Our data are consistent with a model in which the RPB3 and RPB11 heteromerization occurs prior to the association with RPB10 and RPB12 to form the RPB3 subcomplex (45, 46). In summary, this example provides structural validation supporting our subcomplex predictions, pointing to the existence of discrete RNA Pol II subcomplexes in the cell. It seems common that a holocomplex assembly is a regulated event, and CFMS data can provide clues about the path through which this occurs. Interestingly, these abundant sub-complexes may not reside in the nucleus. The cytosolic fraction analyzed in this study is not enriched in abundant nuclear proteins like histones (6, 22), and cytosolic assembly of core RNA POL II has been demonstrated in a human cell line (45–47). Therefore, these subcomplexes could reflect entities with distinct functions in the cytosol and/or protein complexes that cycle between nuclear and non-nuclear localizations.

Importantly, it is worth noting that our methods and results are not specific to plants. Partial assembly of CORUM complexes likely occurs in many organisms, and our methods can be directly applied to detect it in other species.

### Use of gold standards to evaluate a global protein complex prediction

The gold standards were used to evaluate the global prediction results in the rice reference protein complex datasets (16). These datasets were generated using replicate coelution profiles from SEC plus replicates from ion exchange chromatography (IEX), which is an orthogonal separation method of SEC. In the published protein complex prediction, the resulting dendrogram at a specific cluster number determines the compositions of predicted complexes. The specific cluster number is selected based on the resolution of the data and maximized to decrease false positives (23). Of our 40 gold standards, 34 were present in the reference SEC and IEX data used for the protein complex predictions, and they consisted of multiple subunits. Consequently, we employed them to assess the overall reliability of 1,000 predicted protein complexes derived from the rice dataset (16). The assessment was performed using the intactness and purity metrics, as defined in Formula (7). As previously explained in the Material and Methods section, these metrics are more suitable for the evaluation of complex predictions. In a reliable prediction, gold standard complexes should exhibit high intactness and purity. On the other hand, these two metrics are affected by the size of the protein complexes (see Materials and Methods). Figure 5A-D display histograms and density plots of the intactness and purity values, comparing our gold standard subcomplexes and their fully assembled CORUM counterparts. In the assessment of specific global protein complex predictions, Figure 5A and 5B clearly show that subcomplex-based gold standards give high intactness values with the density curve leaning toward the right-hand side. On the other hand, Figure 5C and 5D show that the histogram and density curve of subcomplex-based gold standards overlap with the plots from the CORUM-based reference complexes with comparable purity values. As our gold standards tend to be smaller than the CORUM full complexes, and since the size of reference complexes impacts the intactness and purity values in opposite directions, it is essential to consider these two metrics together. The higher intactness values and similar purity values collectively indicate that the global predicted protein complexes align better with our gold standards than with the CORUM full complexes.

**Figure 5.**
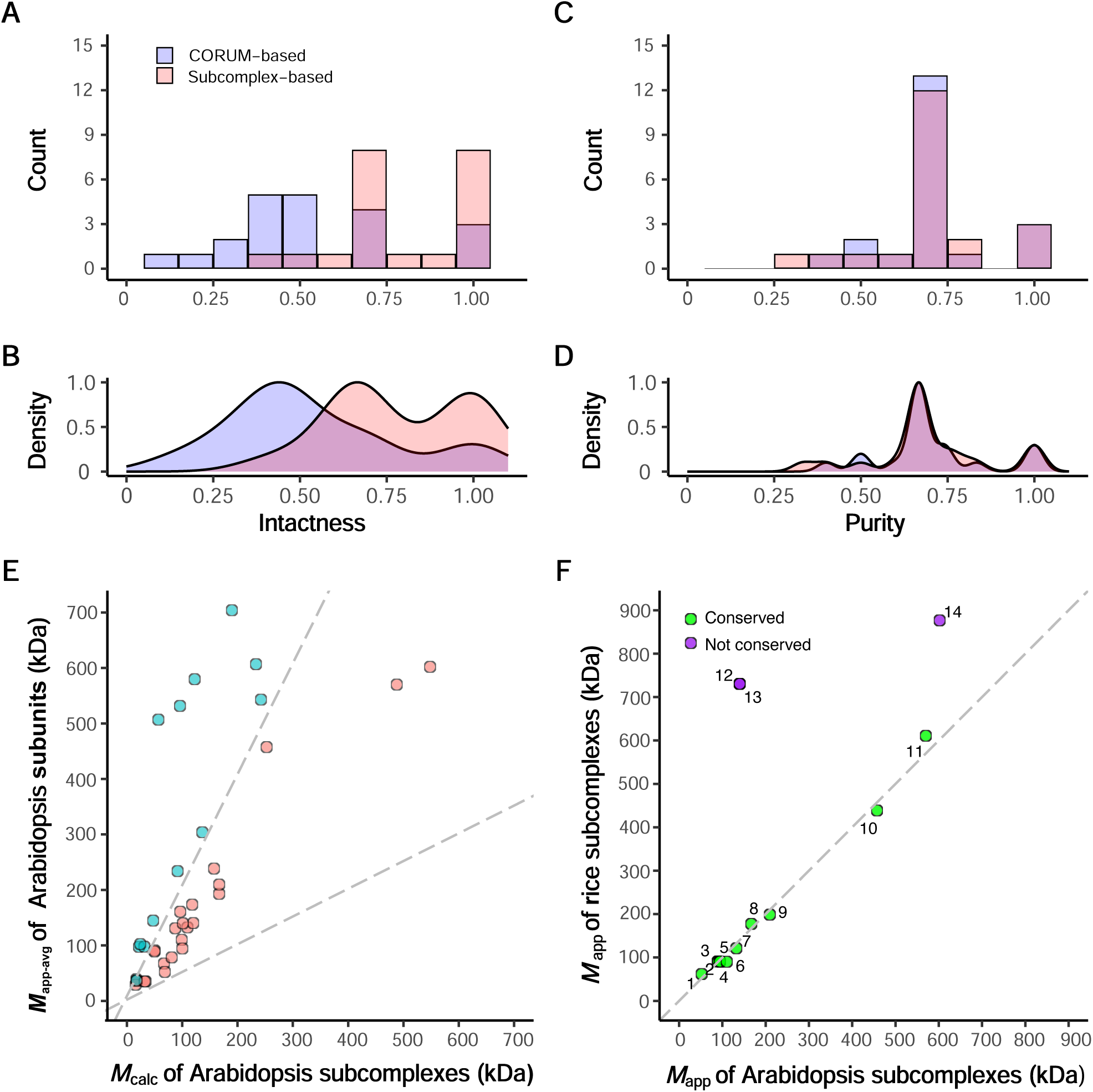
The validated subcomplexes and CORUM complexes were used as standards to evaluate a global prediction of protein complex composition in rice. The global predictions of rice complexes (16) were re-evaluated with respect to the intactness (A, B) and purity (C, D) of assumed fully assembled CORUM orthocomplexes (slate blue) or the validated gold standards defined in this study (pink). A and C display histograms representing the intactness and purity, respectively, while B and D depict the corresponding density plots. E, Gold standard subcomplexes/complexes of Arabidopsis with matched *M*_calc_ and *M*_app-avg_ are rendered in pink. F, Conservation of rice and Arabidopsis subcomplexes. *M*_app_ values of rice gold standards and *M*_app_ values of their orthologous Arabidopsis gold standards were plotted. The conserved gold standards (green) and non-conserved ones (purple) are reported in Supplemental Table S10.

Our gold standards, which are essentially predicted knowns and validated by statistical significance and matched size masses from SEC experiments (*M*_app_) and the calculation (*M*_calc_), have shown robust performance in evaluating protein complex predictions. While higher intactness or purity values are typically deemed optimal with the use of confirmed known complexes as references, our gold standards affirm their utility when computationally validated. We anticipate that the gold standards identified here will serve as improved benchmarks to assess future protein complex predictions.

### Gold standard discoveries in Arabidopsis

To determine if the developed methods were robust, we applied the same pipeline for identifying gold standards to another species, Arabidopsis. The analysis steps, as demonstrated in Figure 1, were replicated. More specifically, the InParanoid software was run to find the orthologs from 20,834 human protein sequences and 35,368 Arabidopsis protein sequences. The algorithm assigned 5,674 human proteins and 10,774 Arabidopsis proteins into 3,227 distinct orthologous groups (Supplemental Table S1B). Among the 3,047 human complexes curated in CORUM, 1,985 (65.1% of total CORUM complexes) had at least one subunit orthologous to one or more Arabidopsis proteins. Nine hundred and seventy of the 1,985 Arabidopsis orthocomplexes exhibited subunit coverage greater than or equal to 2/3 (Supplemental Table S7).

We used a previously published Arabidopsis SEC dataset from the Arabidopsis protein complex predictions of soluble leaf proteins based on parallel SEC and IEX separations (23). Then we applied the same filtering criteria used in the rice subcomplex prediction to find reproducible protein elution profiles. A total of 1,738 protein subunits with *R*_app_ > 1 were used as the Arabidopsis SEC reference profiles. Among the 970 orthocomplexes with subunit coverage greater than or equal to 2/3, 289 orthocomplexes had at least one Arabidopsis subunit detected by the Arabidopsis SEC reference profiles, and 184 out of 289 orthocomplexes had at least two Arabidopsis subunits detected by the Arabidopsis SEC reference profiles. After eliminating redundant orthocomplexes, as done with the rice data, 63 orthocomplexes (Supplemental Table S8) remained for further analysis for gold standard discovery. As observed for rice in Figure 2C and as reported for Arabidopsis microsomal proteins (6), there was poor agreement between *M*_app_ of Arabidopsis subunits and *M*_calc_ of the assumed fully assembled Arabidopsis orthocomplexes (Supplemental Figure S5).

We applied the same tuning parameters used for the two-stage clustering algorithm with the rice dataset to predict gold standards in Arabidopsis. We identified 23 Arabidopsis gold standards with statistically significant *p*-values and similar *M*_app_ and *M*_calc_ values that satisfied the criterion in Formula (6) (Figure 5E and Supplemental Table S9). To analyze the extent to which subcomplexes were conserved in a monocot and dicot species, we compared the 40 rice subcomplexes and 23 Arabidopsis subcomplexes. A stringent comparison requiring the complete overlap of all subunits between an Arabidopsis subcomplex and its corresponding rice subcomplex was limited to only 5 predicted gold standards. This is largely due to non-overlapping proteome coverages obtained from different tissue types and different mass spectrometers. As an alternative approach, we compared *M*_app_ values of subcomplexes predicted in both species for cases in which at least one conserved subunit was detected (Figure 5F and Supplemental Table S10). Eleven subcomplexes were represented in the datasets from both species and showed the same/similar *M*_app_, suggesting conserved assemblies. This result showcases the generalizability of our gold standard prediction algorithm in species that diverged more than 100 million years ago. On the other hand, the agreement was not absolute as three rice orthologs had an elevated *M*_app_ compared to their Arabidopsis counterparts (Figure 5F).

## Discussion

Protein correlation profiling or CFMS is a powerful technique that enables one to make open-ended predictions about the multimerization state (2, 4–7, 13, 22, 48), composition (3, 9, 16, 20, 23, 49), and localization of endogenous protein complexes (7, 10–12). As the sensitivity of mass spectrometers and the sophistication through which subcellular fractions can be analyzed improve (50, 51), the ability of CFMS link in vitro biochemistry with cell function will only increase. A major limitation in the CFMS field and a knowledge gap in biology in general is the lack of data on the true multimerization states of proteins in the cell. For CFMS analyses, known complexes are needed to evaluate the accuracy of large-scale predictions, and CORUM has served that purpose. CORUM is a useful database of fully assembled complexes that have a known function in the cell. However, this does not mean that the fully assembled complex is the most abundant state of the subunits in cells, nor does it exclude the possibility that individual subunits or subcomplexes have functions that are independent of the fully assembled complex. In plant cell extracts, CORUM orthocomplexes are rarely fully assembled (6, 22), and to our knowledge, this has not been addressed directly in non-plant systems. The lack of agreement cannot be explained as an artifactual disassembly that occurs during cell lysis or extended chromatographic separations because many of the CORUM orthologs have apparent masses that greatly exceed that of the fully assembled CORUM complex (Figure 2C and Supplemental Figure S5). It seems likely that many CORUM subunits/subcomplexes/fully assembled complexes have unknown or moonlighting functions that involve physical interactions with unknown proteins/complexes. Many CORUM subunit orthologs show a reduced apparent mass compared to the predicted fully assembled complex (Figure 2C). This could reflect instances in which assembly status may differ between subcellular locations (Figure 4) or those in which full assembly is a highly regulated event, and the fully assembled state is a low abundance pool that escapes detection in CFMS pipelines that can only detect major peaks. Regardless of the explanation, and because of the fundamental importance of reliable gold standards, the possibility of widespread conflicts between the CORUM-predicted multimerization states and those measured using SEC and crude cell extracts needs to be systematically examined.

This paper describes robust experimental methods and statistical approaches that can be broadly adopted to identify a more reliable set of gold standards. These reliable standards more accurately reflect the multimerization states of CORUM orthologs in cell extracts (Figure 5A-D). Genome sequencing and useful algorithms like InParanoid and EggNOG make it possible to accurately map orthologs among diverse species (9, 16, 20, 22, 23). The complex discovery of knowns is based on standard SEC chromatography, label-free protein quantification methods, and data filtering criteria that can be adapted to any experimental system. The robust two-stage clustering algorithm was developed using a small number of known rice complexes, and the exact settings used to predict reliable rice complexes were also used successfully in Arabidopsis (Figures 5). Evolutionarily conserved multimeric assemblies have been reported previously (9, 20), and this phenomenon seems to also apply to many of the gold standards described here (Figure 5F). It may be some of these complexes share similar paths to full assembly and the manner in which subunits are distributed among differing multimerization states could encode information for the cell. Importantly gold standards cannot be blindly transferred across species or kingdom lines (Figure 5F), and orthologs and paralogs frequently display multimerization variability (16, 22), so care must be taken to systematically build databases of useful gold standards.

Having databases of more accurate gold standards will fundamentally improve how CFMS predictions are evaluated but may not help to reduce a certain type of errors (Figure 5C-D). Chance co-elution is a major confounding effect because the complexity of input sample is too high and the distribution of *M*_app_ is skewed toward relatively small complexes (17, 18, 23). Some benefit could be gained by using gold standards to optimally tune the number of groups or clusters in a CFMS prediction (16, 23), but this will not solve the problem. Although more expensive and time consuming in terms of the increased sample numbers required, a major benefit could be obtained by conducting IEX and SEC separations in series and increasing the upstream fractionation of the input material. We expect improved mass spectrometry instrumentation and computational tools to enable ever more powerful CFMS-based analyses of protein multimerization complexity and dynamics.

## Data and materials availability

The source code and sample input data for the small-world analysis are publicly available on GitHub (https://github.com/yangpengchengstat/R-code-S4_Class-protein-clustering-based-on-data-integration-of-corum-and-inparanoid.git). The package at the github link contains comprehensive information on running the code, description of the input data, and steps of performing hyperparameter tuning.

## Supporting information

Table 1 Predicted gold standards for CFMS predictions

Supplemental Table S1 Human to plant ortholog mappings

Supplemental Table S2 Rice CORUM orthocomplexes

Supplemental Table S3 Rice orthocomplexes and the subcomplex prediction results by the small world analysis

Supplemental Table S4 Significant rice subcomplexes with small p-values

Supplemental Table S5 Predicted rice gold standards

Supplemental Table S6 Predicted rice monomers

Supplemental Table S7 Arabidopsis CORUM orthocomplexes

Supplemental Table S8 Arabidopsis orthocomplexes and the subcomplex prediction results by the small world analysis

Supplemental Table S9. Predicted Arabidopsis gold standards

Supplemental Table S10 Conserved and nonconserved gold standards between rice and Arabidopsis

Supplemental Figure S1 Subunit coverages of rice orthocomplexes

Supplemental Figure S2 Rice subcomplex identification by small world analysis(Mapp vs Mcalc plots)

Supplemental Figure S3 Rice protein SEC profiles illustrating small-world analysis results

Supplemental Figure S4 Histograms of mean of pair-wised doverall from random simulations of rice orthocomplexes

Supplemental Figure S5 Subunit coverages of Arabidopsis orthocomplexes

Supplemental Figure S6 Arabidopsis subcomplex identification by small world analysis(Mapp vs Mcalc plots)

Supplemental Figure S7 Arabidopsis protein SEC profiles illustrating small-world analysis results

## Acknowledgments

Publication of this article was funded in part by Purdue University Libraries Open Access Publishing Fund.

## Abbreviations

CFMS: co-fractionation mass spectrometry
CORUM: the comprehensive resource of mammalian protein complexes database
SEC: size exclusion chromatography
IEX: ion exchange chromatography
*M*_app_: protein apparent mass
*M*_mono_: protein monomeric mass
*M*_calc_: protein calculated mass
*R*_app_: multimerization state or determinator
*WCC*: weighted cross-correlation
*Euclid*: Euclidean distance
*d*: distance between two proteins
SOM: self-organizing map
AP: affinity propagation

## Supplemental Figures

Supplemental Figure S1. Subunit coverages of rice orthocomplexes.

Supplemental Figure S2. Rice subcomplex identification by small world analysis (*M*_app_ vs *M*_calc_ plots).

Supplemental Figure S3. Rice protein SEC profiles illustrating small-world analysis results.

Supplemental Figure S4. Histograms of mean of pair-wised *d_overall_* from random simulations of rice orthocomplexes.

Supplemental Figure S5. Subunit coverages of Arabidopsis orthocomplexes.

Supplemental Figure S6. Arabidopsis subcomplex identification by small world analysis (*M*_app_ vs *M*_calc_ plots).

Supplemental Figure S7. Arabidopsis protein SEC profiles illustrating small-world analysis results.

## Supplemental Tables

Supplemental Table S1. Human to plant ortholog mappings.

Supplemental Table S2. Rice CORUM orthocomplexes.

Supplemental Table S3. Rice orthocomplexes and the subcomplex prediction results by the small world analysis.

Supplemental Table S4. Significant rice subcomplexes with small *p*-values.

Supplemental Table S5. Predicted rice gold standards.

Supplemental Table S6. Predicted rice monomers.

Supplemental Table S7. Arabidopsis CORUM orthocomplexes.

Supplemental Table S8. Arabidopsis orthocomplexes and the subcomplex prediction results by the small world analysis.

Supplemental Table S9. Predicted Arabidopsis gold .

Supplemental Table S10. Conserved and non-conserved gold standards between rice and Arabidopsis.

