## Supplementary material for "A machine learning-based approach to identify reliable gold standards for protein complex composition prediction": Table 1 Predicted gold standards for CFMS predictions

Table 1. Predicted gold standards to evaluate CFMS-based protein complex predictions.

| CORUM complex | Rice subcomplex IDs  (# of subunits orthocomplex) | Subcomplex prediction *p*-value | *M*calc of subcomplex (kDa) | *M*app-avg of subcomplex (kDa) | # of CORUM subunits predicted  /  total # of CORUM subunits |
| --- | --- | --- | --- | --- | --- |
| 17S U2 snRNP | subcomplex 3 (11) | 0.000008 | 392.7 | 550.0 | 9 / 33 |
| Gamma-BAR-AP1 complex | all proteins defined in the complex by Gaussian peak distance before clustering (3) | 0.004744 | 391.6 | 421.3 | 5 / 9 |
| APPBP1-UBA3 complex | only 2 proteins in the complex before clustering (2) | 0.000947 | 108.7 | 121.3 | 2 / 2 |
| RNA polymerase II holoenzyme complex | subcomplex 1 (2) | 0.010076 | 50.5 | 68.2 | 2 / 24 |
| RNA polymerase II holoenzyme complex | subcomplex 3 (8) | 0.000008 | 460.2 | 636.4 | 7 / 24 |
| RNA polymerase II holoenzyme complex | subcomplex 4 (3) | 0.011642 | 201.5 | 372.4 | 3 / 24 |
| C complex spliceosome | subcomplex 2 (5) | 0.000252 | 880.0 | 730.4 | 5 / 80 |
| CAPZalpha-CAPZbeta complex | only 2 proteins in the complex before clustering (2) | 0.030328 | 61.7 | 89.5 | 2 / 2 |
| CCT complex (chaperonin containing TCP1 complex) | all proteins defined in the complex by Gaussian peak distance before clustering (10) | 0.000008 | 937.1 | 698.2 | 8 / 8 |
| Prefoldin complex | all proteins defined in the complex by Gaussian peak distance before clustering (6) | 0.000008 | 99.1 | 136.8 | 6 / 6 |
| SEC23-SEC24 adaptor complex | all proteins defined in the complex by Gaussian peak distance before clustering (3) | 0.000069 | 197.6 | 341.0 | 2 / 2 |
| CSA-POLIIa complex | subcomplex 1 (6) | 0.000008 | 301.8 | 438.7 | 6 / 13 |
| CUL4A-DDB1-RBBP5 complex | subcomplex 1 (3) | 0.031703 | 278.6 | 325.9 | 3 / 3 |
| EIF2B1-EIF2B2-EIF2B3-EIF2B4-EIF2B5 complex | all proteins defined in the complex by similarity before clustering (5) | 0.000008 | 563.4 | 863.2 | 5 / 5 |
| Elongator holo complex | all proteins defined in the complex by Gaussian peak distance before clustering (3) | 0.000122 | 305.4 | 590.2 | 3 / 4 |
| ESCRT-III complex | only 2 proteins in the complex before clustering (2) | 0.013789 | 98.8 | 57.9 | 4 / 10 |
| Exosome | subcomplex 1 (7) | 0.000008 | 199.9 | 278.1 | 7 / 11 |
| FIB-associated protein complex | subcomplex (ortho-paralog) 2 (4) | 0.000023 | 49.8 | 90.1 | 1 / 6 |
| HCF-1 complex | subcomplex 1 (6) | 0.000008 | 177.3 | 352.0 | 2 / 18 |
| KAT2A-Oxoglutarate dehydrogenase complex | subcomplex (ortho-paralog) 1 (2) | 0.000122 | 108.8 | 193.6 | 1 / 4 |
| KAT2A-Oxoglutarate dehydrogenase complex | subcomplex 1 (3) | 0.002040 | 109.5 | 114.8 | 2 / 4 |
| LSm1-7 complex | subcomplex 2 (4) | 0.000642 | 46.6 | 78.3 | 4 / 7 |
| LSm2-8 complex | subcomplex 2 (5) | 0.000863 | 63.1 | 94.6 | 5 / 7 |
| MCM2-MCM4-MCM6-MCM7 complex | all proteins defined in the complex by similarity before clustering (4) | 0.000046 | 377.6 | 466.3 | 4 / 4 |
| Membrane protein complex (VCP, UFD1L, SEC61B) | subcomplex (ortho-paralog) 2 (2) | 0.000405 | 538.7 | 563.7 | 1 / 3 |
| mRNA decay complex (UPF1, UPF2, UPF3B, DCP2, XRN1, XRN2, EXOSC2, EXOSC4, EXOSC10, PARN) | subcomplex 1 (3) | 0.004362 | 173.7 | 293.0 | 3 / 10 |
| p400-associated complex | subcomplex 1 (4) | 0.001597 | 346.2 | 614.7 | 3 / 6 |
| PA700-20S-PA28 complex | subcomplex 1 (15) | 0.000008 | 600.7 | 610.7 | 11 / 36 |
| PA700-20S-PA28 complex | subcomplex 2 (21) | 0.000008 | 761.6 | 876.6 | 15 / 36 |
| Parvulin-associated pre-rRNP complex | subcomplex (ortho-paralog) 1 (3) | 0.000290 | 49.7 | 91.1 | 1 / 12 |
| RAF1-PPP2-PIN1 complex | subcomplex 1 (2) | 0.003018 | 121.9 | 198.7 | 2 / 5 |
| Retromer complex (SNX1, SNX2, VPS35, VPS29, VPS26B) | all proteins defined in the complex by similarity before clustering (3) | 0.002559 | 248.0 | 211.7 | 4 / 5 |
| Ribosome, cytoplasmic | subcomplex 2 (2) | 0.001604 | 42.3 | 68.5 | 2 / 80 |
| SNW1 complex | subcomplex 1 (3) | 0.042039 | 268.6 | 223.1 | 2 / 18 |
| SNW1 complex | subcomplex 2 (4) | 0.002040 | 559.1 | 751.7 | 3 / 18 |
| SNW1 complex | subcomplex 3 (5) | 0.000008 | 50.1 | 90.8 | 1 / 18 |
| Spliceosome | subcomplex 1 (3) | 0.000733 | 64.3 | 93.8 | 4 / 143 |
| Spliceosome | subcomplex 2 (2) | 0.010122 | 57.5 | 61.8 | 2 / 143 |
| TBCD-ARL2-tubulin (beta -TBCE complex) | subcomplex 1 (2) | 0.000970 | 146.7 | 282.0 | 2 / 4 |
