## Supplementary figures and images for "A machine learning-based approach to identify reliable gold standards for protein complex composition prediction"

### Supplemental Figure S1 Subunit coverages of rice orthocomplexes

**Supplemental Figure S1.** Subunit coverages of rice orthocomplexes

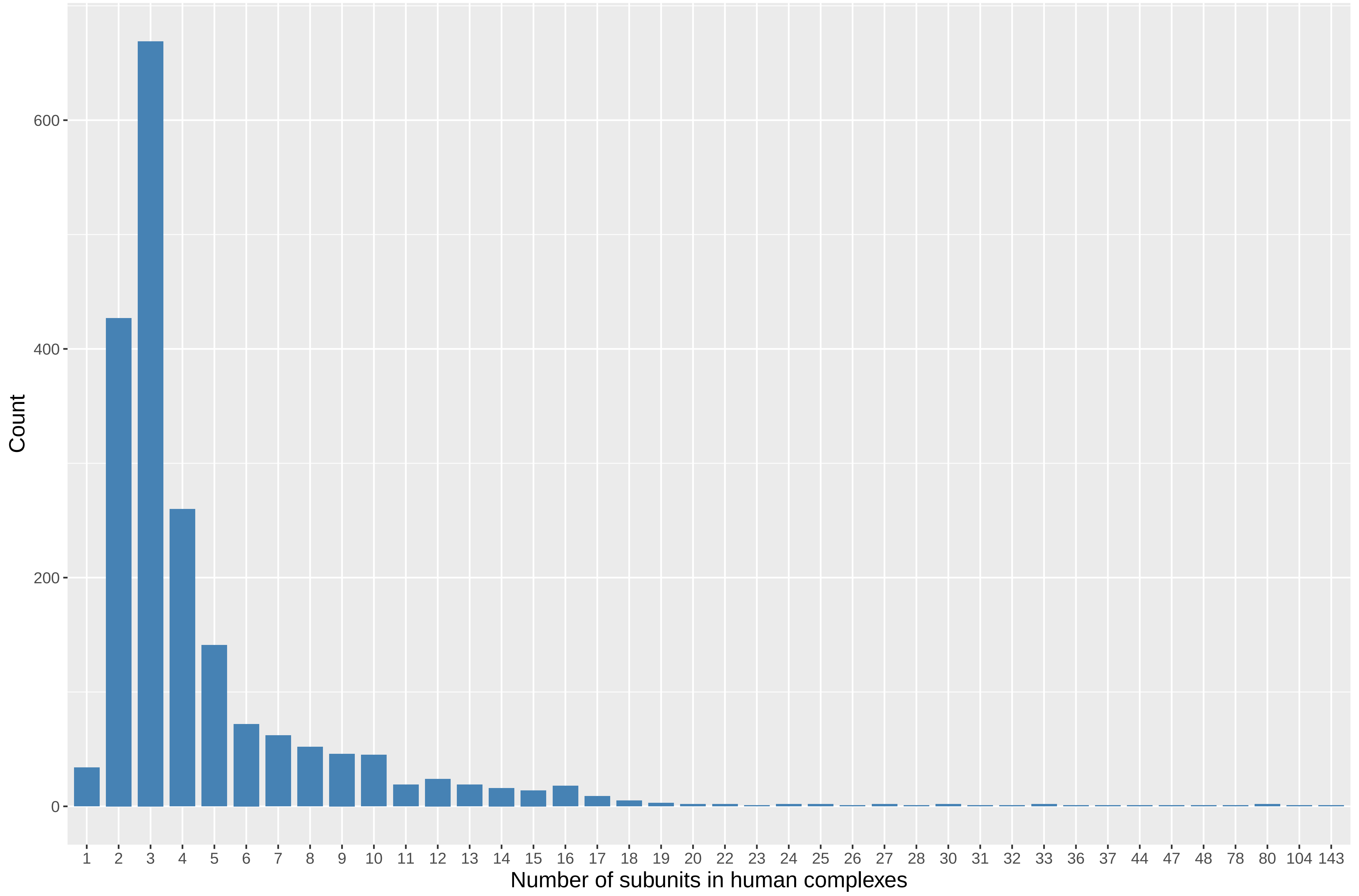

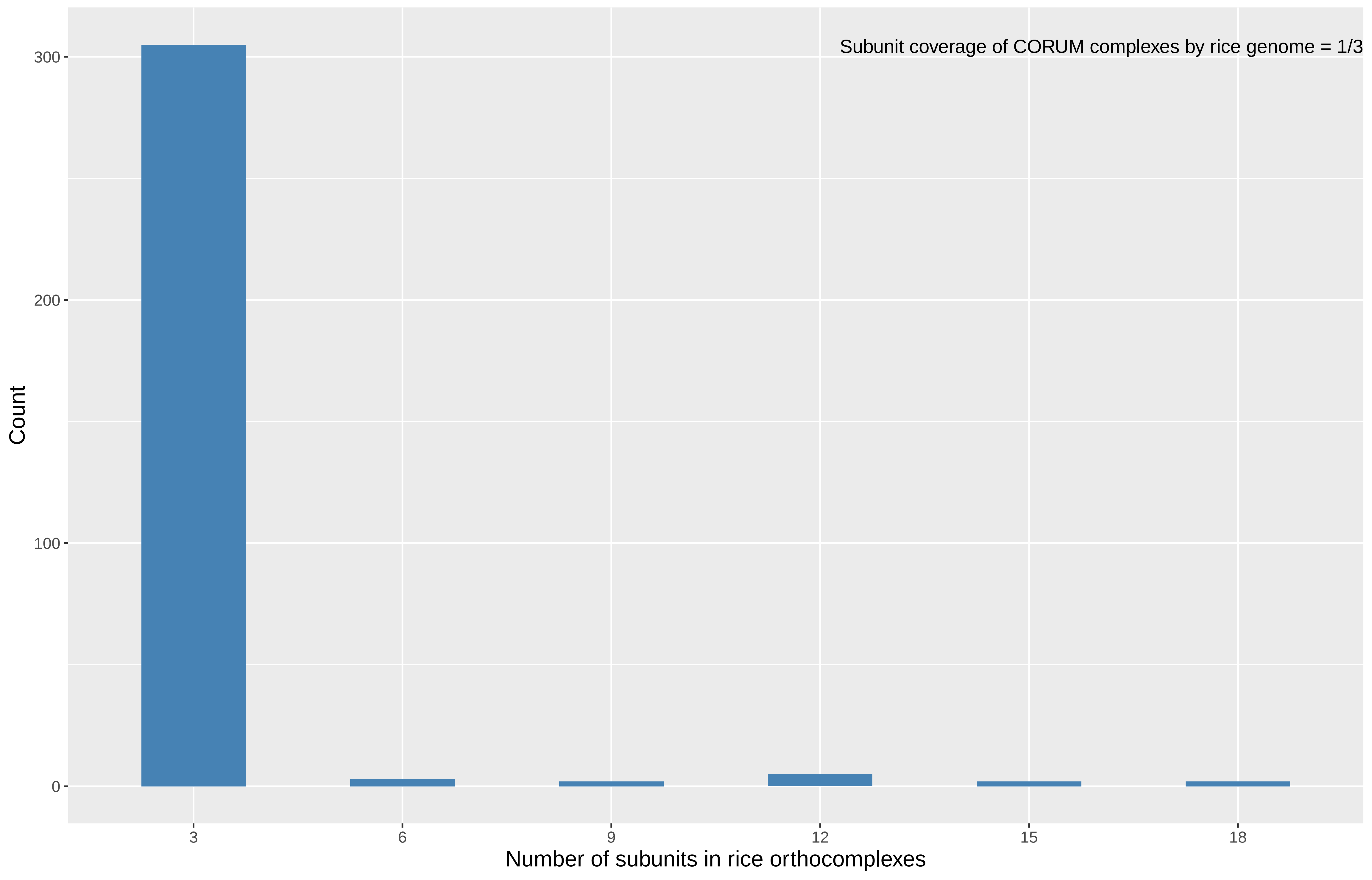

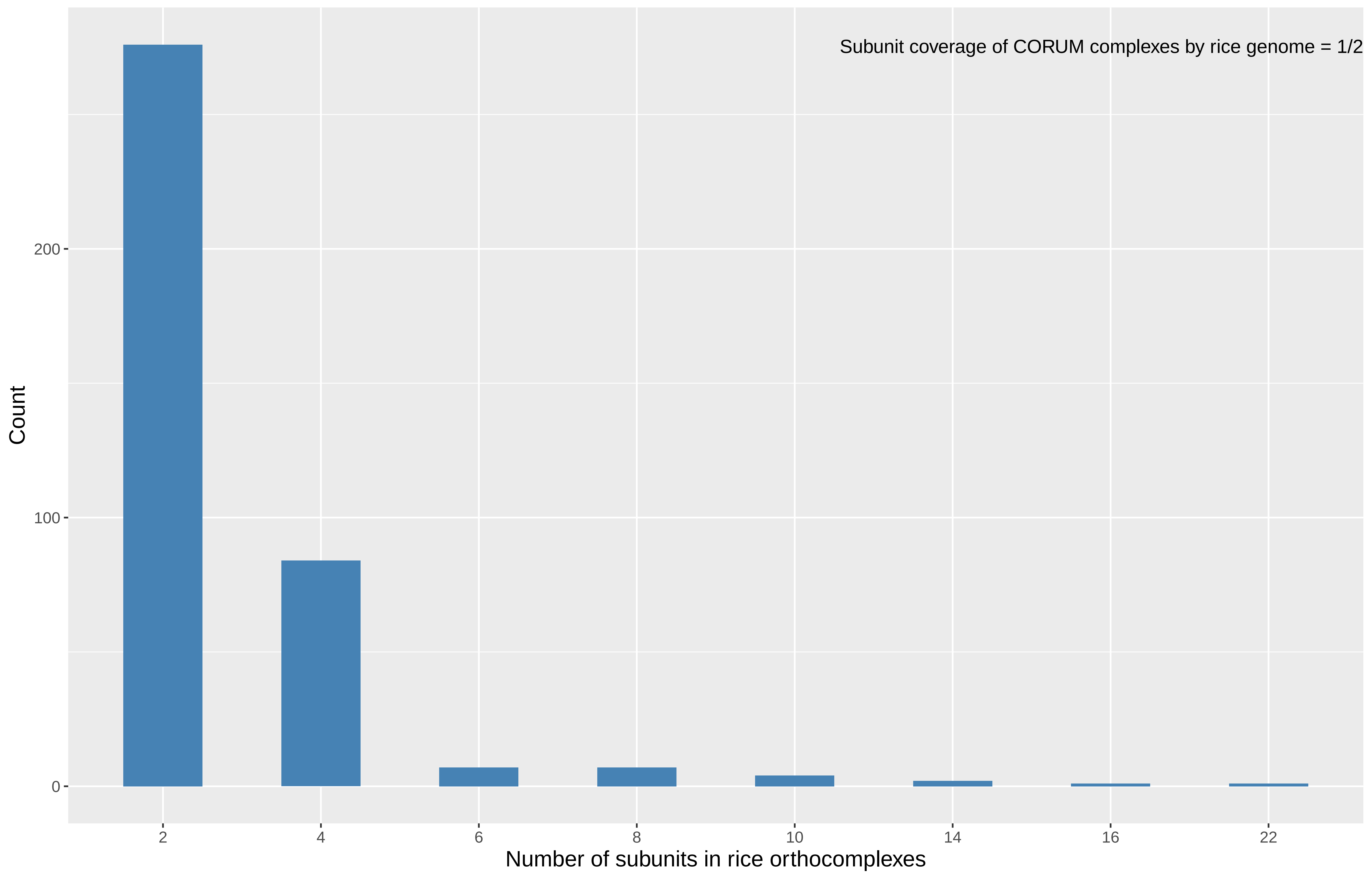

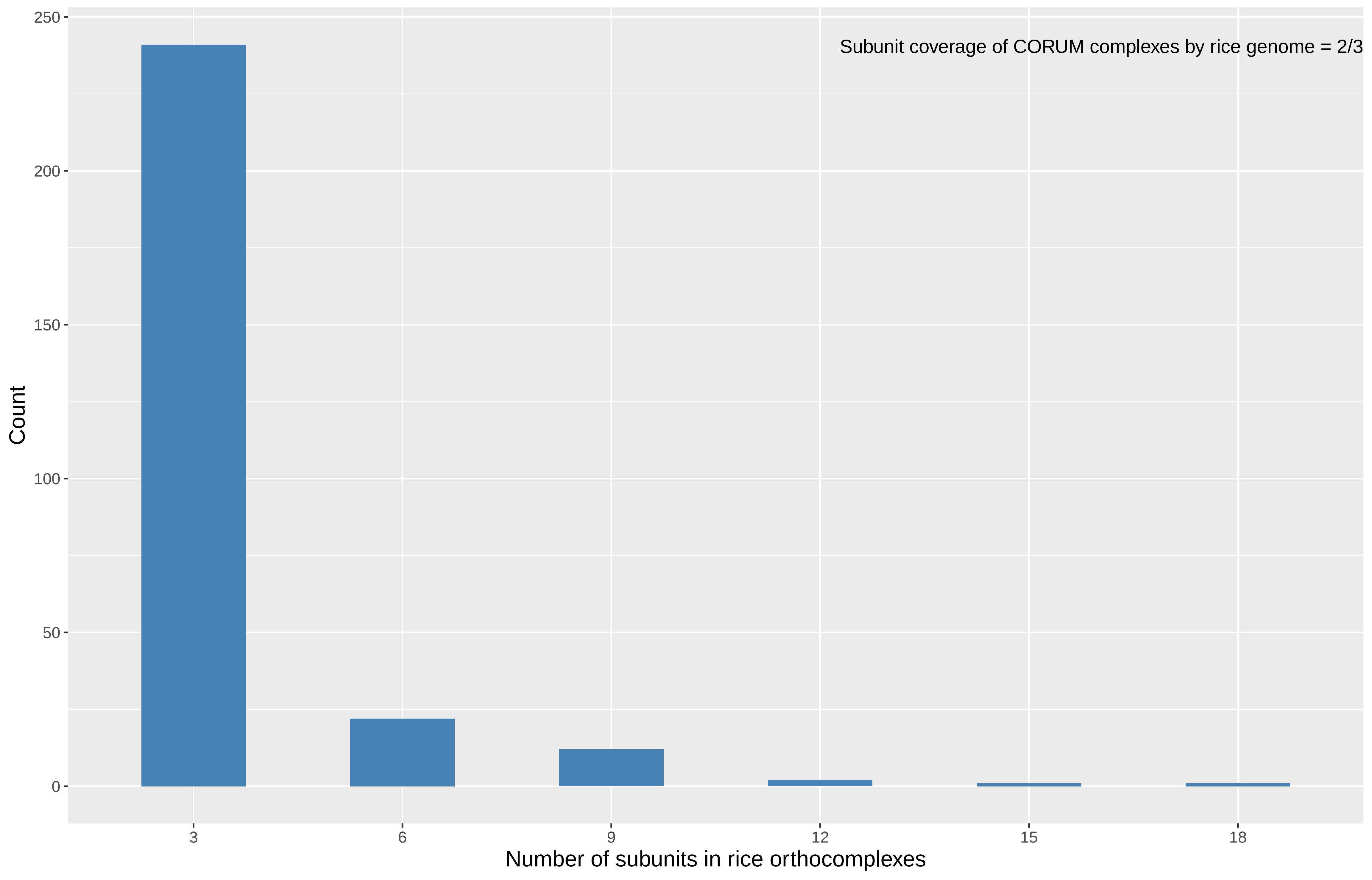

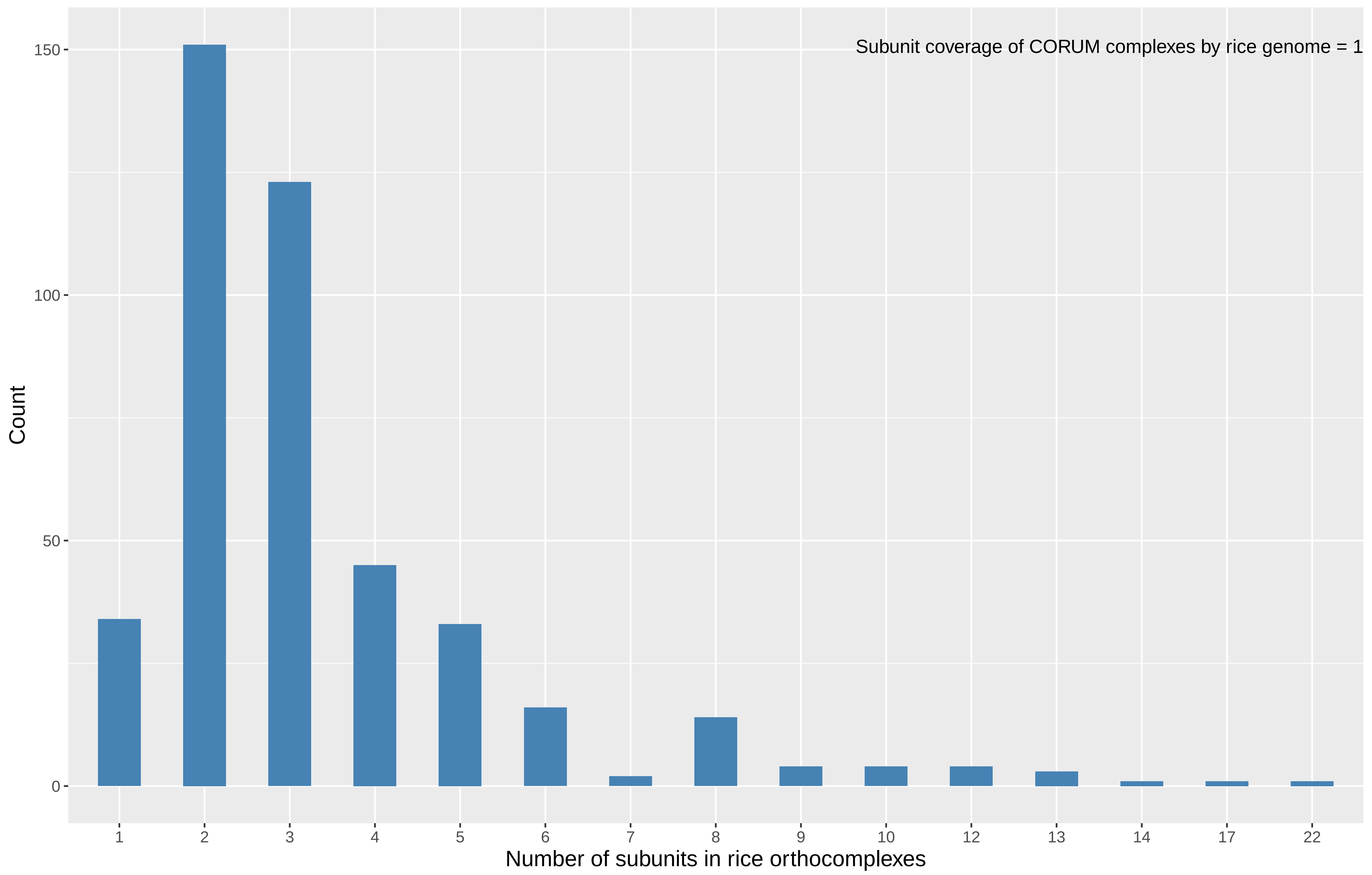

### Supplemental Figure S5 Subunit coverages of Arabidopsis orthocomplexes

**Supplemental Figure S5.** Subunit coverages of Arabidopsis orthocomplexes

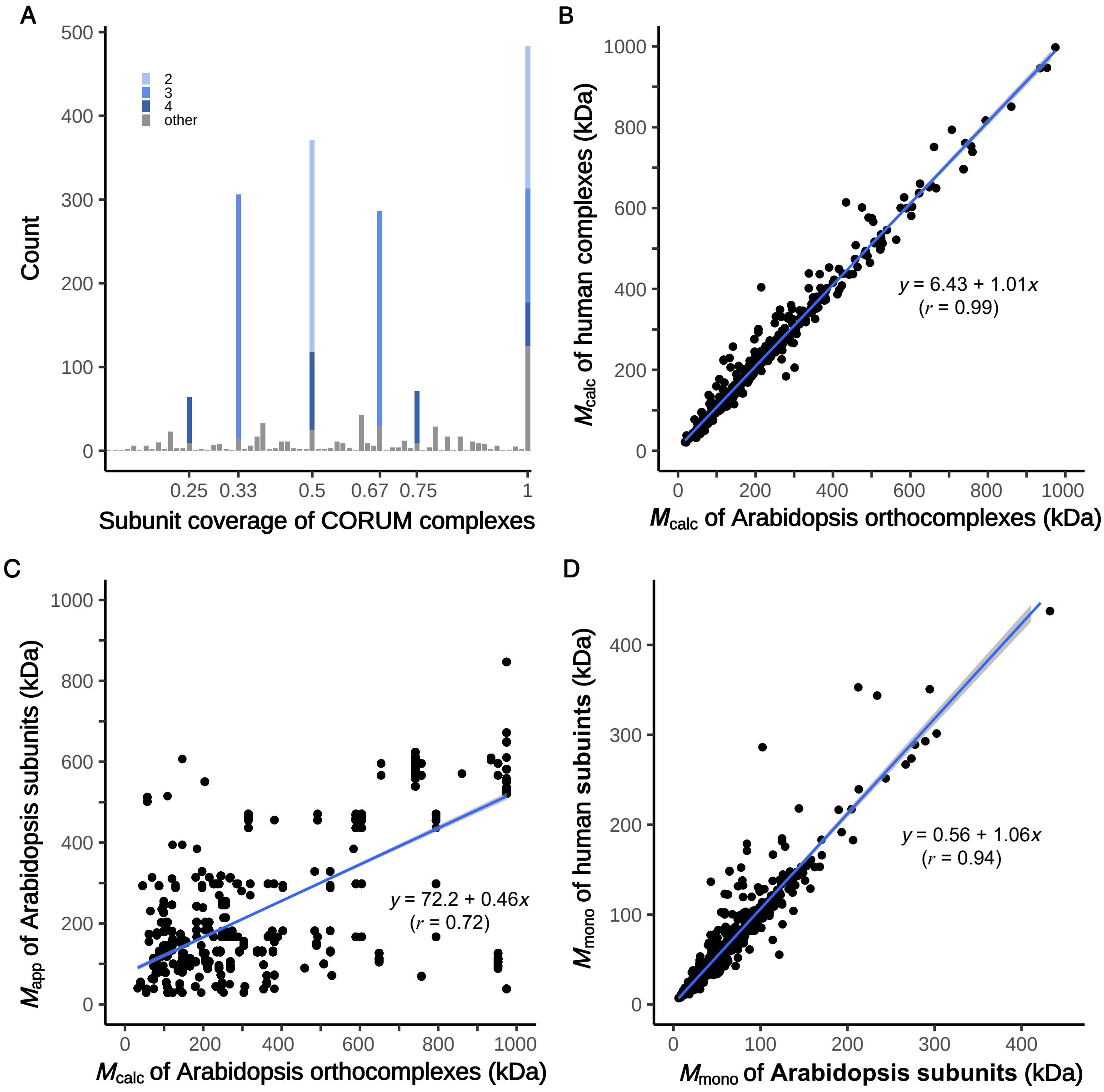

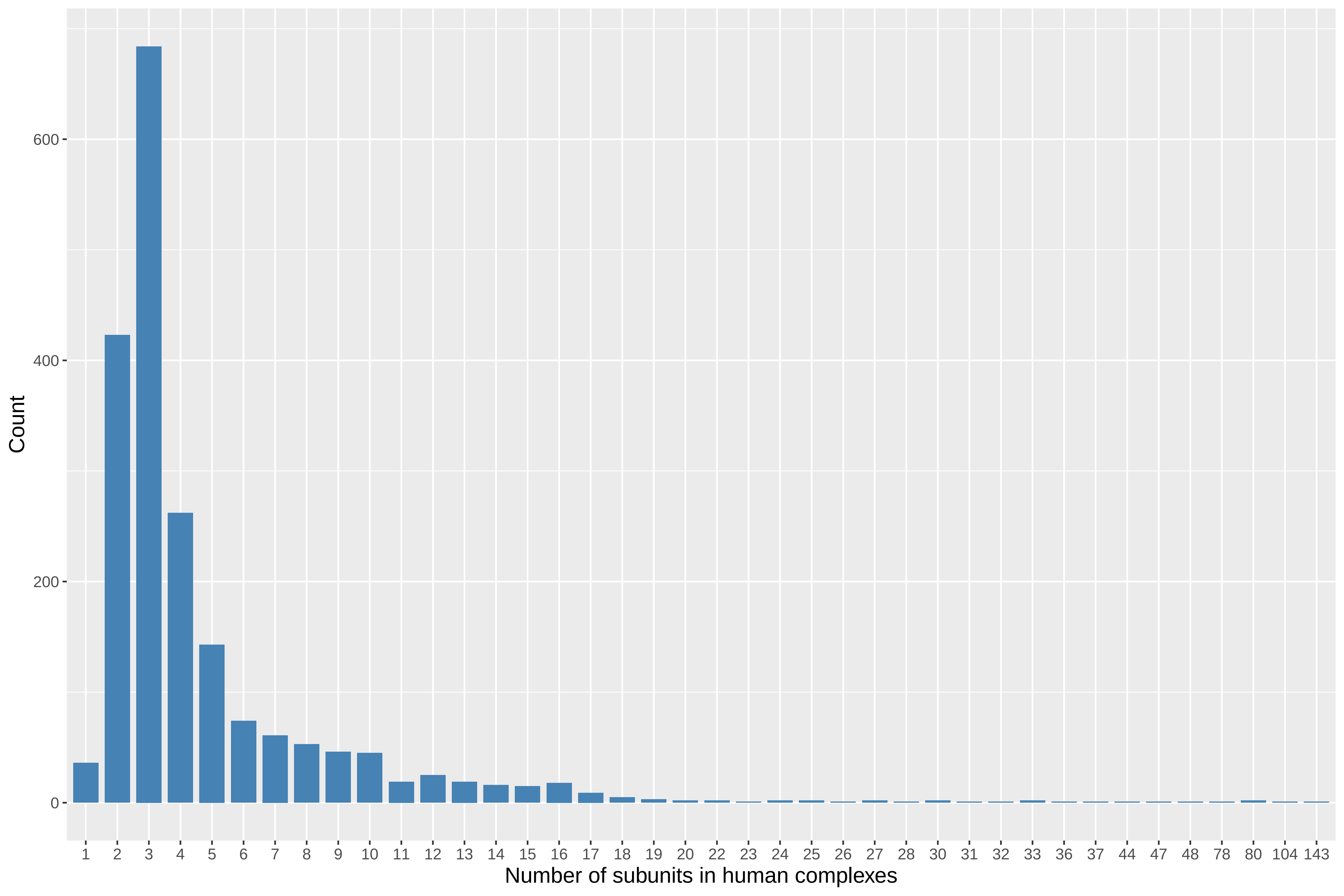

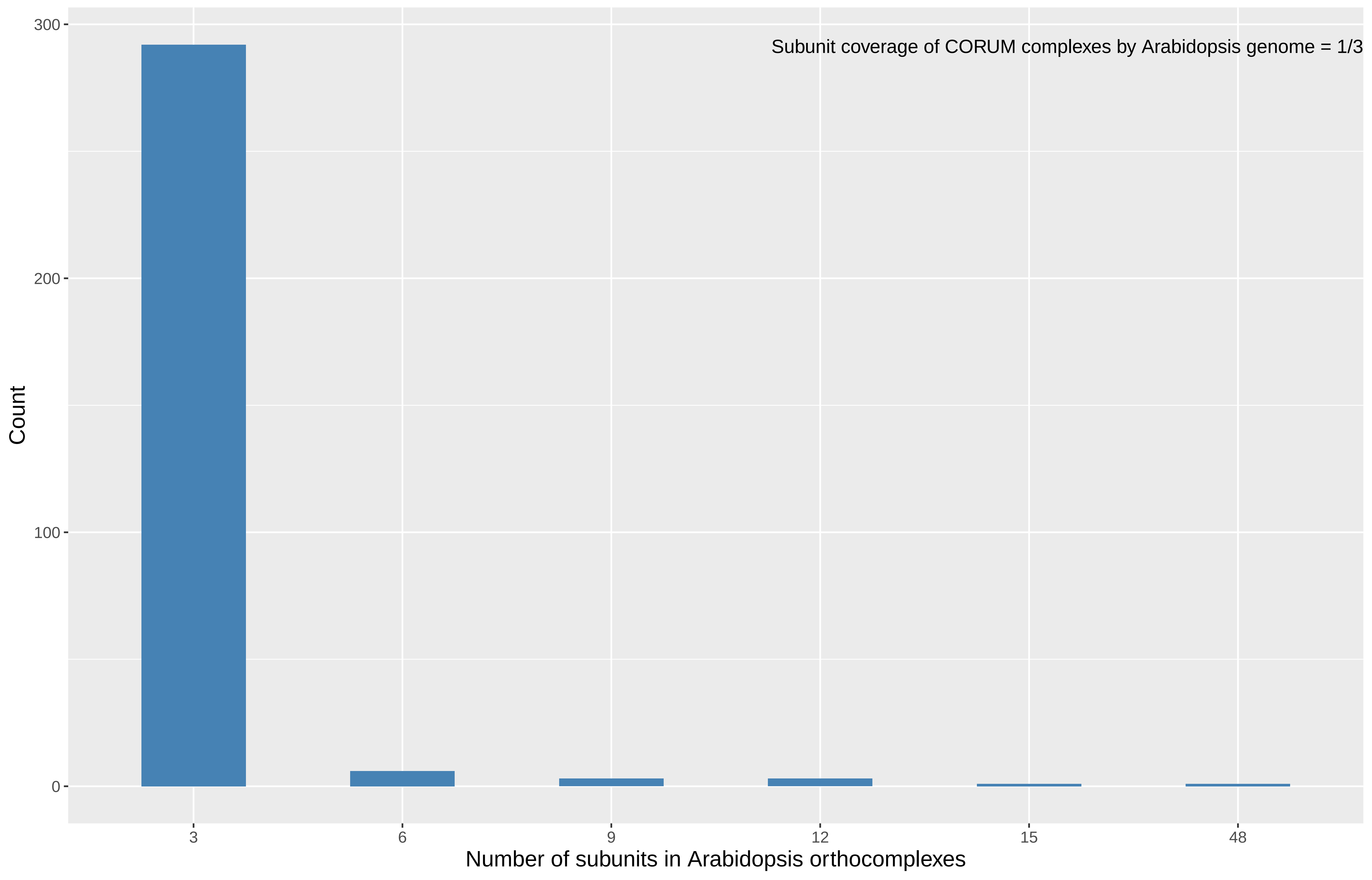

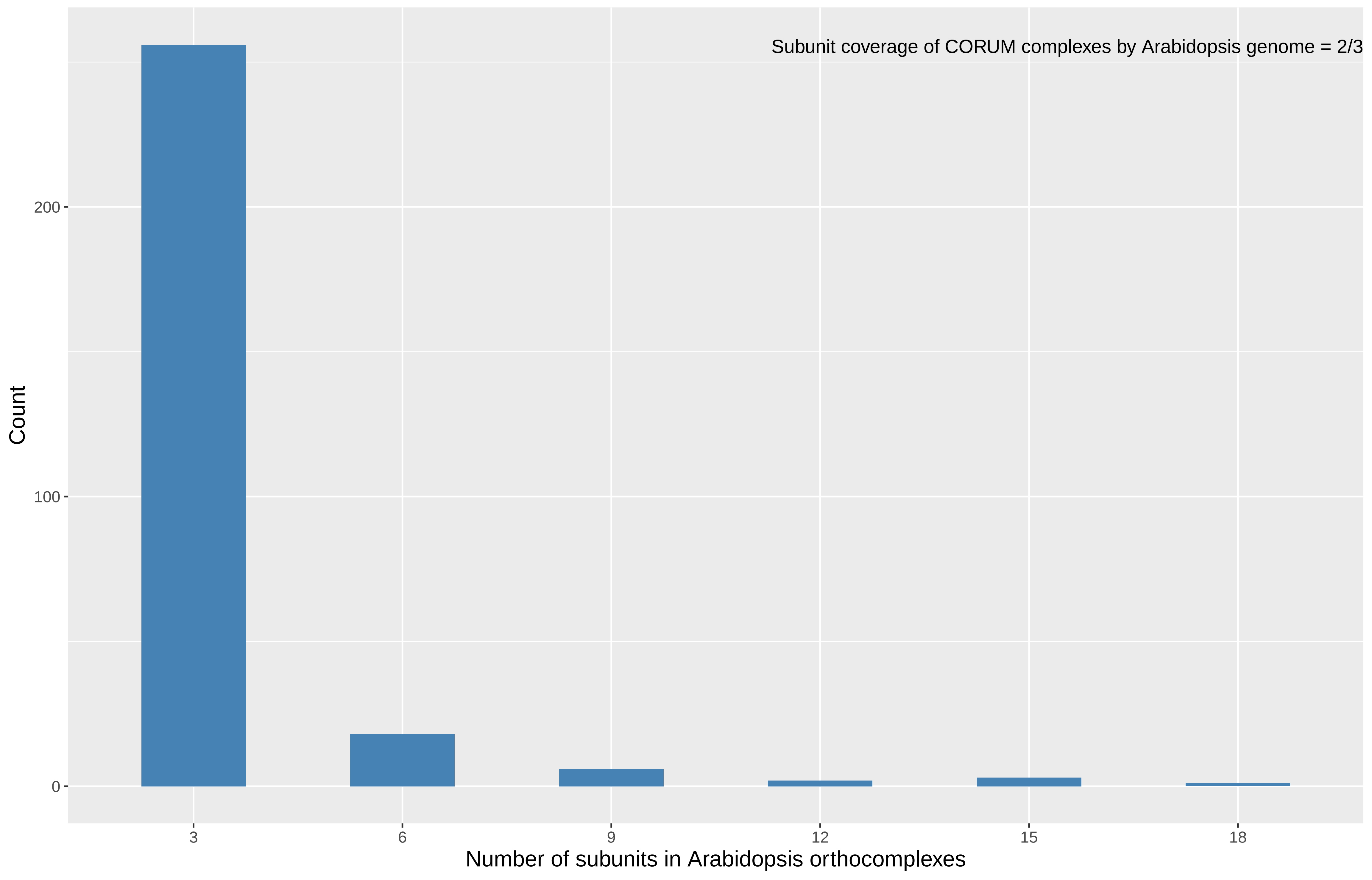

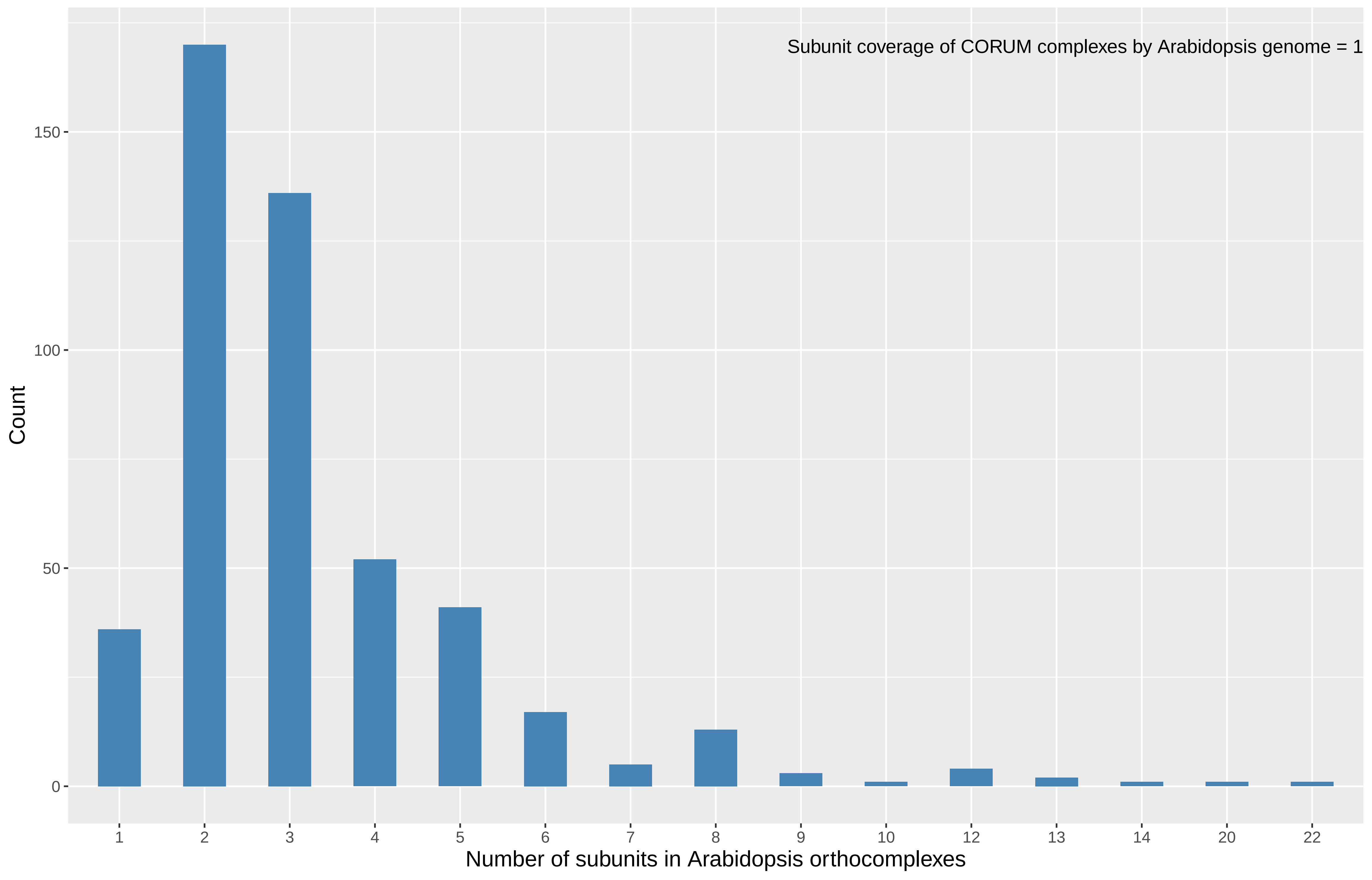
