## Supplemental Figure S2 Rice subcomplex identification by small world analysis(Mapp vs Mcalc plots) for "A machine learning-based approach to identify reliable gold standards for protein complex composition prediction"

**Supplemental Figure S2.** Rice subcomplex identification by small world analysis ( $M_{app}$ vs $M_{calc}$ )

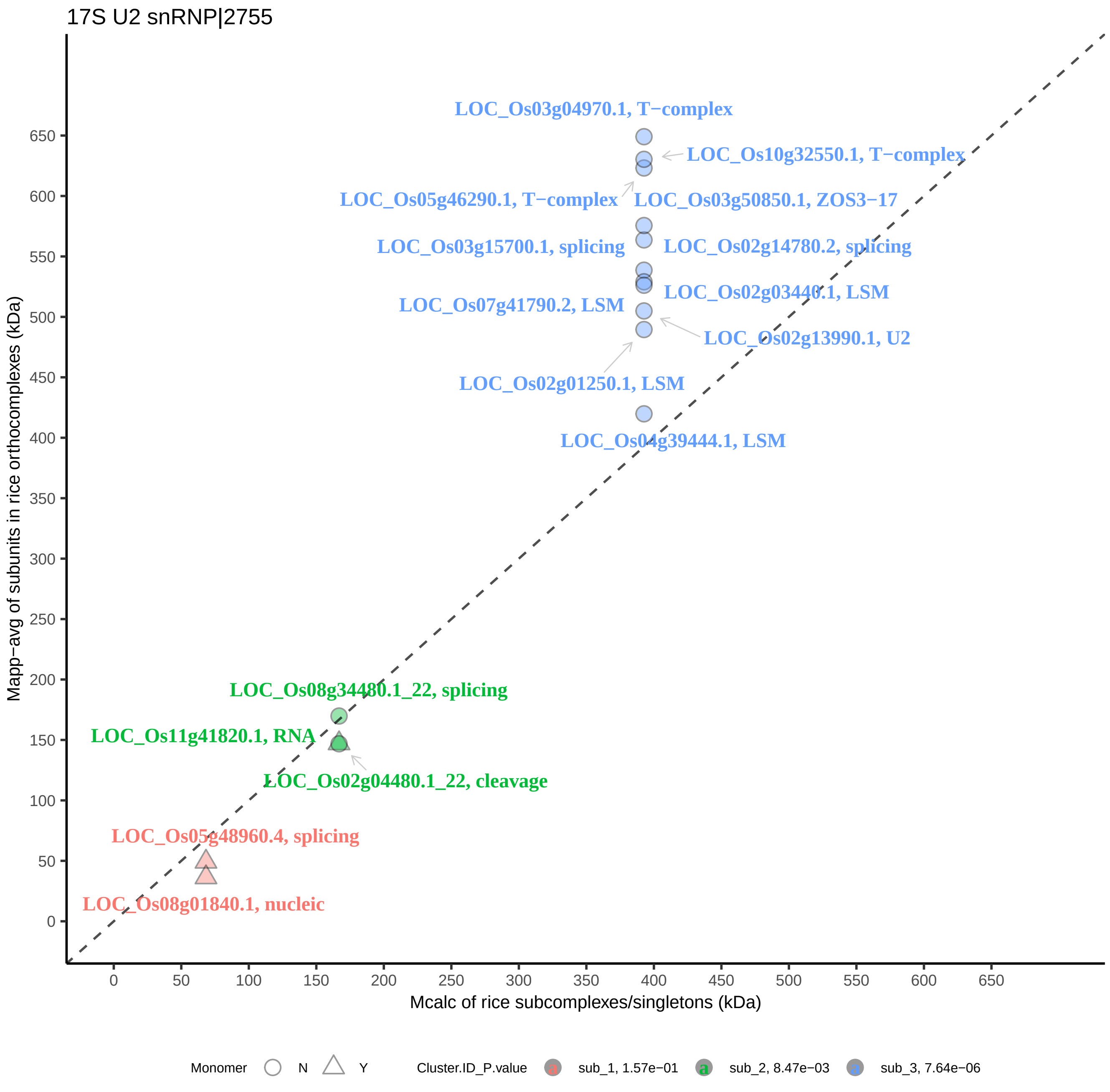

17S U2 snRNP|2755
LOC_Os03g04970.1, T−complex
LOC_Os10g32550.1, T−complex
LOC_Os05g46290.1, T−complex
LOC_Os03g50850.1, ZOS3−17
LOC_Os03g15700.1, splicing
LOC_Os02g14780.2, splicing
LOC_Os02g03440.1, LSM
LOC_Os07g41790.2, LSM
LOC_Os02g13990.1, U2
LOC_Os02g01250.1, LSM
LOC_Os04g39444.1, LSM
LOC_Os08g34480.1_22, splicing
LOC_Os11g41820.1, RNA
LOC_Os02g04480.1_22, cleavage
LOC_Os05g48960.4, splicing
LOC_Os08g01840.1, nucleic
Mapp−avg of subunits in rice orthocomplexes (kDa)
Mcalc of rice subcomplexes/singletons (kDa)
Monomer N Y
Cluster.ID_P.value
sub_1, 1.57e−01
sub_2, 8.47e−03
sub_3, 7.64e−06

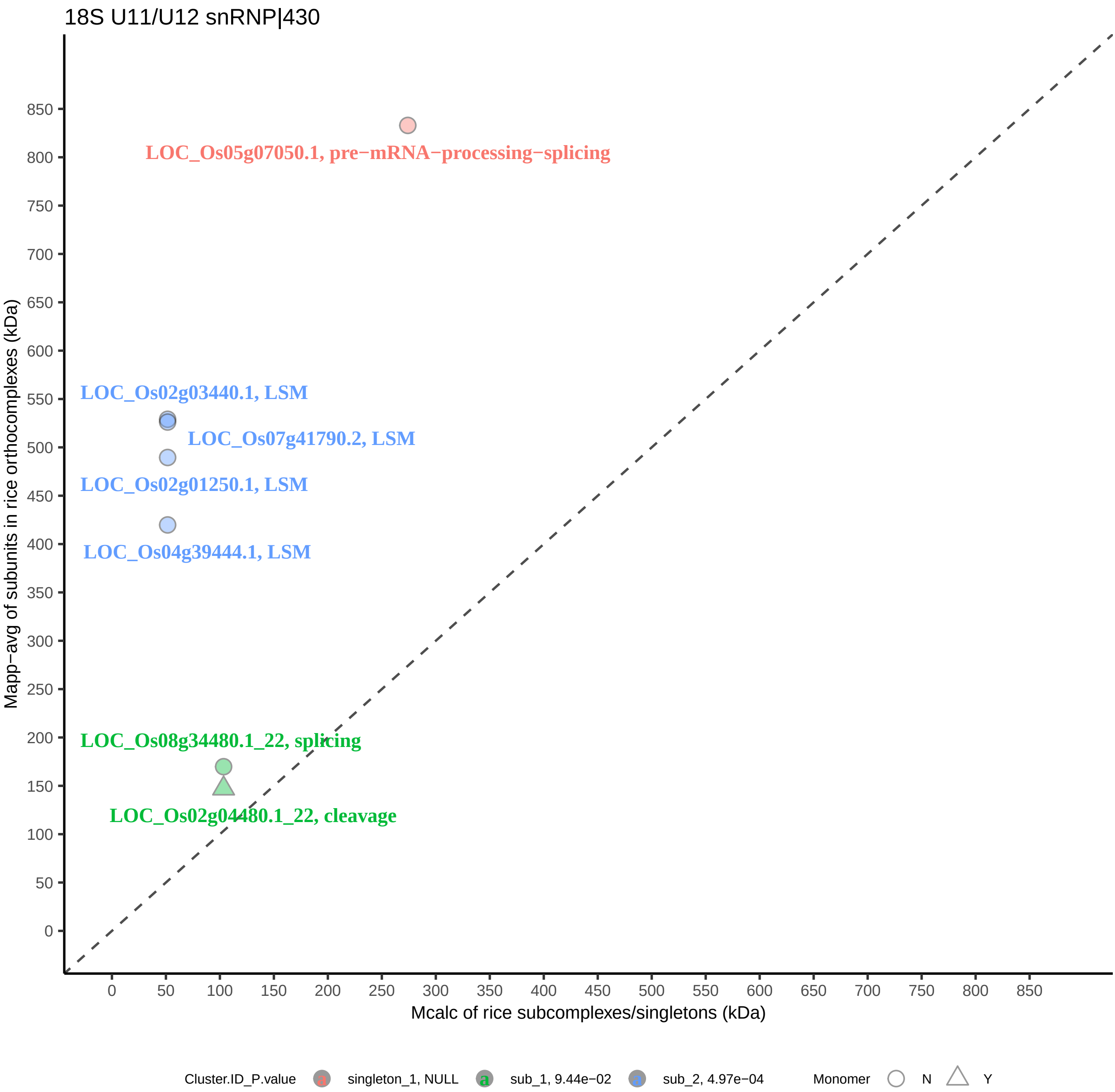
18S U11/U12 snRNP|430
LOC_Os05g07050.1, pre−mRNA−processing−splicing
LOC_Os02g03440.1, LSM
LOC_Os07g41790.2, LSM
LOC_Os02g01250.1, LSM
LOC_Os04g39444.1, LSM
LOC_Os08g34480.1_22, splicing
LOC_Os02g04480.1_22, cleavage
Mapp−avg of subunits in rice orthocomplexes (kDa)
Mcalc of rice subcomplexes/singletons (kDa)
0
50
100
150
200
250
300
350
400
450
500
550
600
650
700
750
800
850
Cluster.ID_P.value
singleton_1, NULL
sub_1, 9.44e−02
sub_2, 4.97e−04
Monomer
N
Y

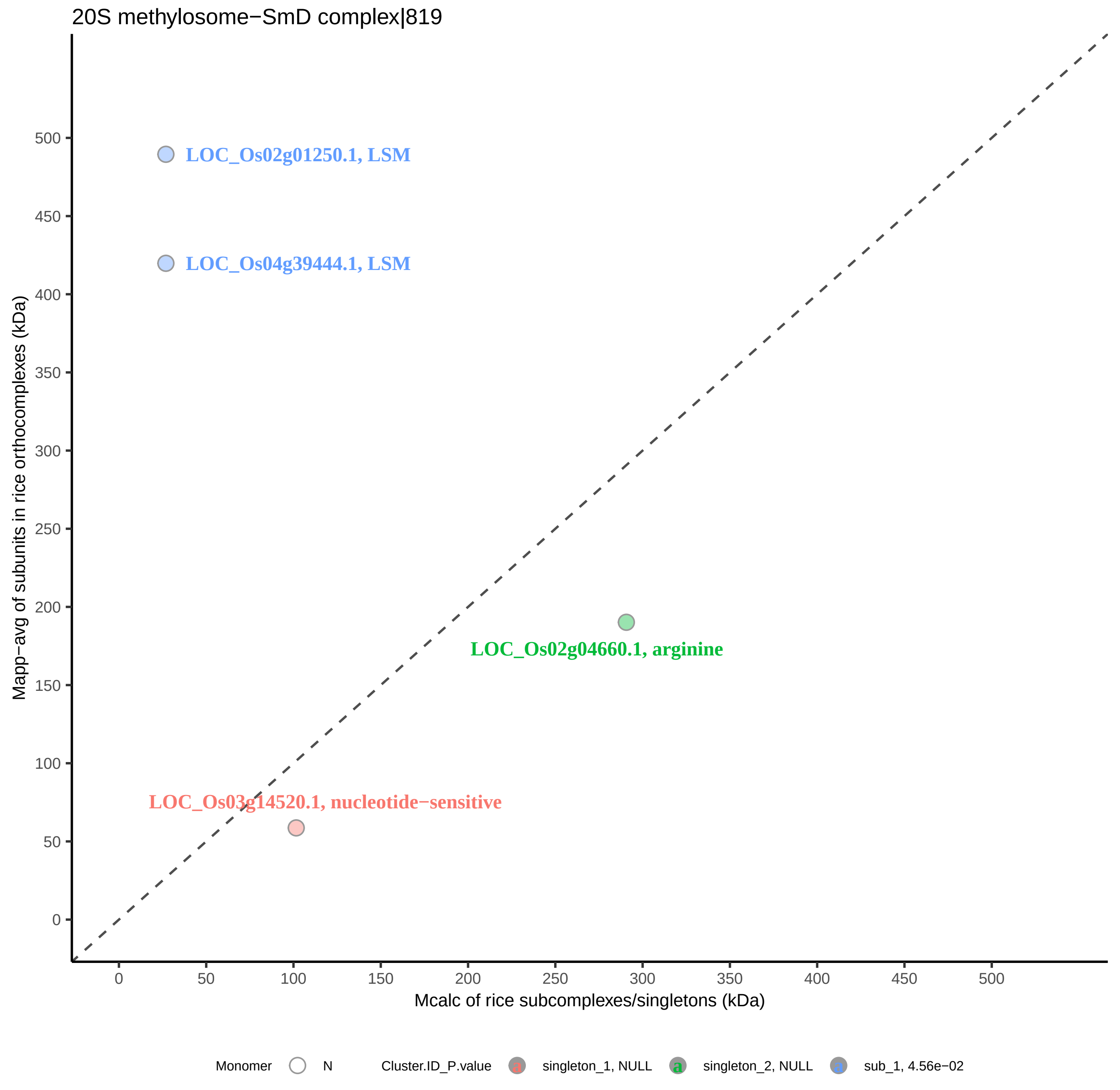
20S methylosome−SmD complex|819
LOC_Os02g01250.1, LSM
LOC_Os04g39444.1, LSM
LOC_Os02g04660.1, arginine
LOC_Os03g14520.1, nucleotide−sensitive
Mapp−avg of subunits in rice orthocomplexes (kDa)
Mcalc of rice subcomplexes/singletons (kDa)
0
50
100
150
200
250
300
350
400
450
500
Monomer
N
Cluster.ID_P.value
singleton_1, NULL
singleton_2, NULL
sub_1, 4.56e−02

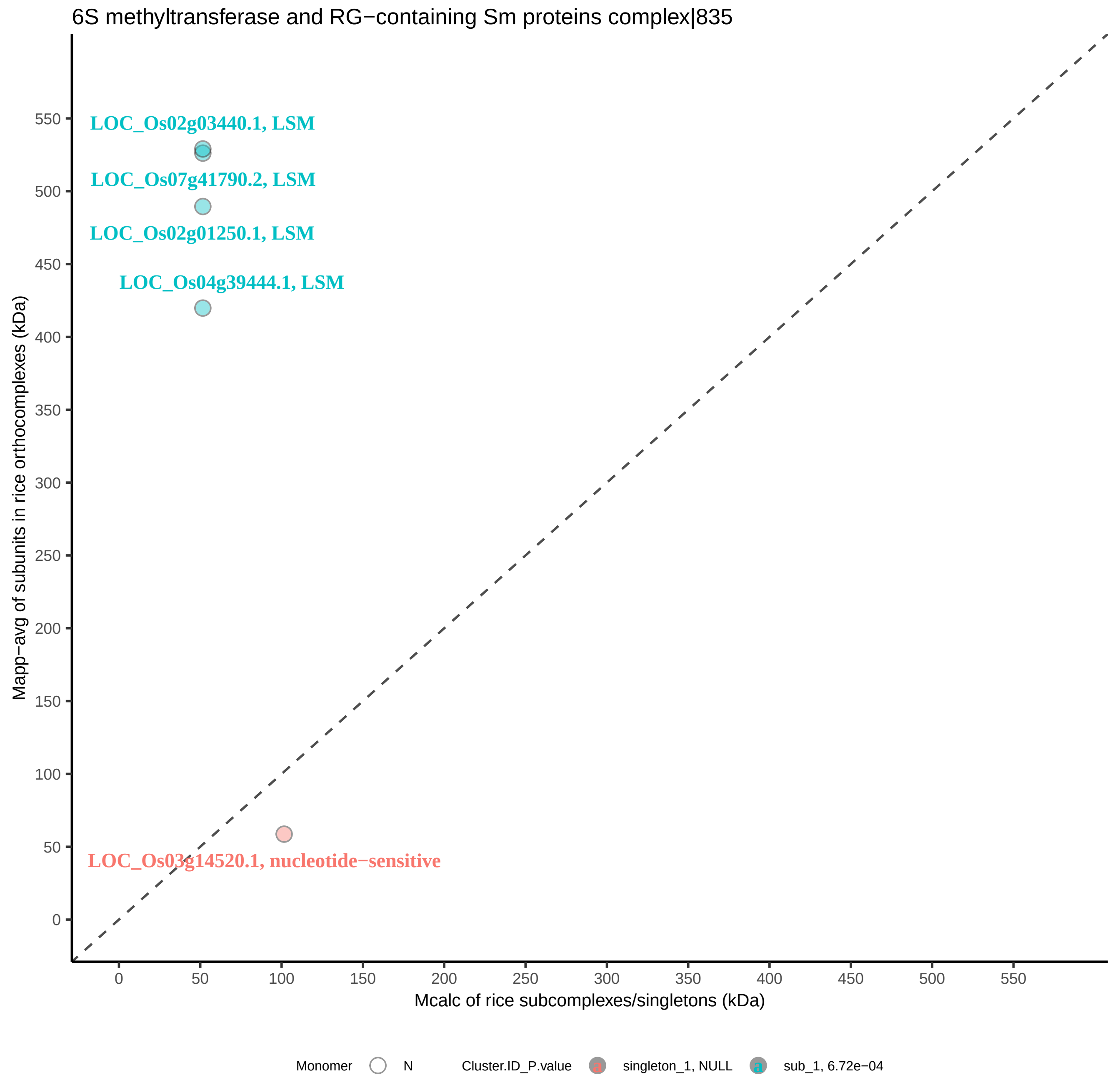

6S methyltransferase and RG−containing Sm proteins complex|835
LOC_Os02g03440.1, LSM
LOC_Os07g41790.2, LSM
LOC_Os02g01250.1, LSM
LOC_Os04g39444.1, LSM
LOC_Os03g14520.1, nucleotide−sensitive
Mapp−avg of subunits in rice orthocomplexes (kDa)
Mcalc of rice subcomplexes/singletons (kDa)
0
50
100
150
200
250
300
350
400
450
500
550
Monomer
N
Cluster.ID_P.value
singleton_1, NULL
sub_1, 6.72e−04

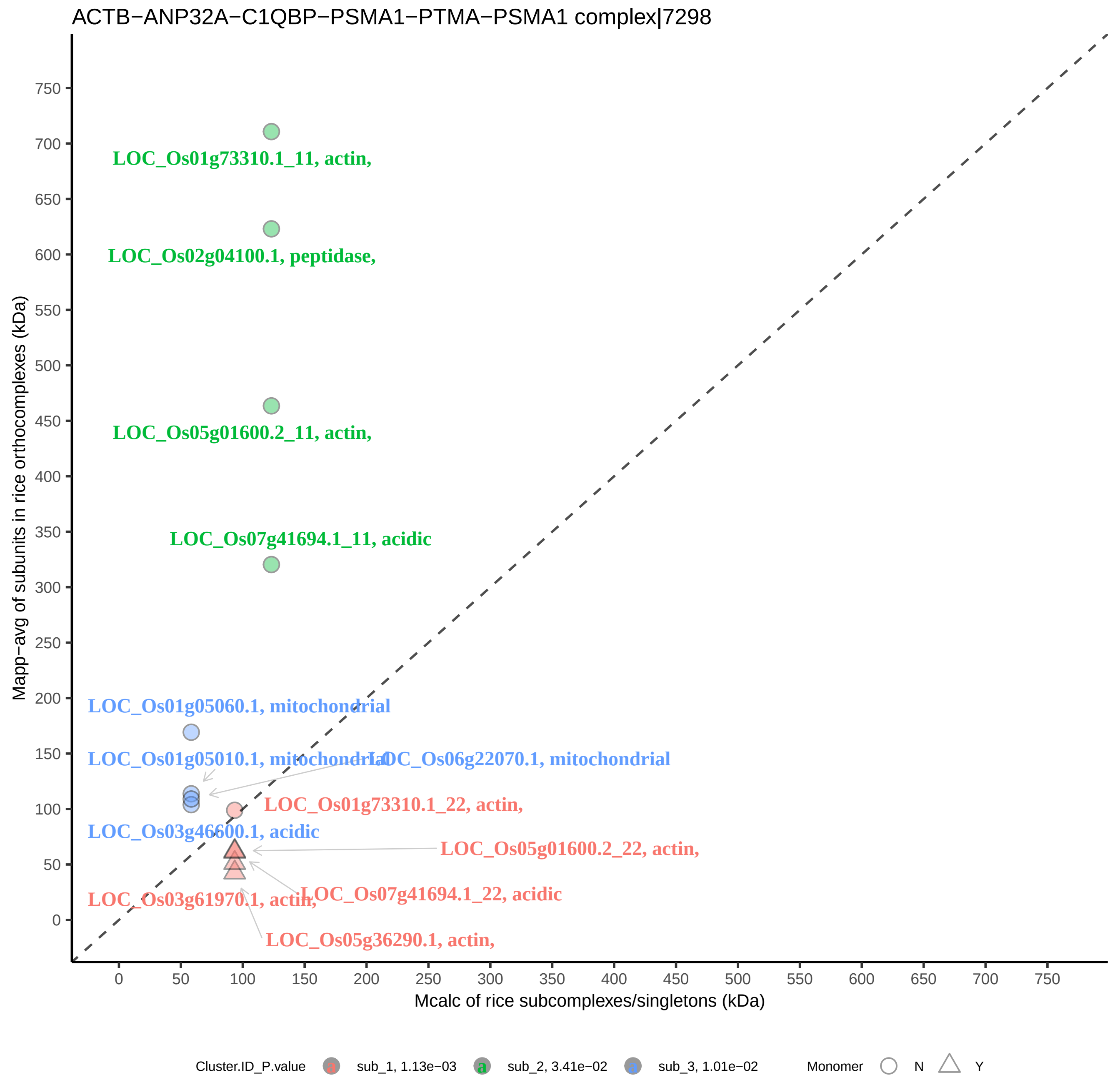

ACTB−ANP32A−C1QBP−PSMA1−PTMA−PSMA1 complex|7298
Mapp−avg of subunits in rice orthocomplexes (kDa)
Mcalc of rice subcomplexes/singletons (kDa)
0
50
100
150
200
250
300
350
400
450
500
550
600
650
700
750
LOC_Os01g73310.1_11, actin,
LOC_Os02g04100.1, peptidase,
LOC_Os05g01600.2_11, actin,
LOC_Os07g41694.1_11, acidic
LOC_Os01g05060.1, mitochondrial
LOC_Os01g05010.1, mitochondrial
LOC_Os06g22070.1, mitochondrial
LOC_Os01g73310.1_22, actin,
LOC_Os03g46600.1, acidic
LOC_Os05g01600.2_22, actin,
LOC_Os07g41694.1_22, acidic
LOC_Os03g61970.1, actin,
LOC_Os05g36290.1, actin,
Cluster.ID_P.value
sub_1, 1.13e−03
sub_2, 3.41e−02
sub_3, 1.01e−02
Monomer
N
Y

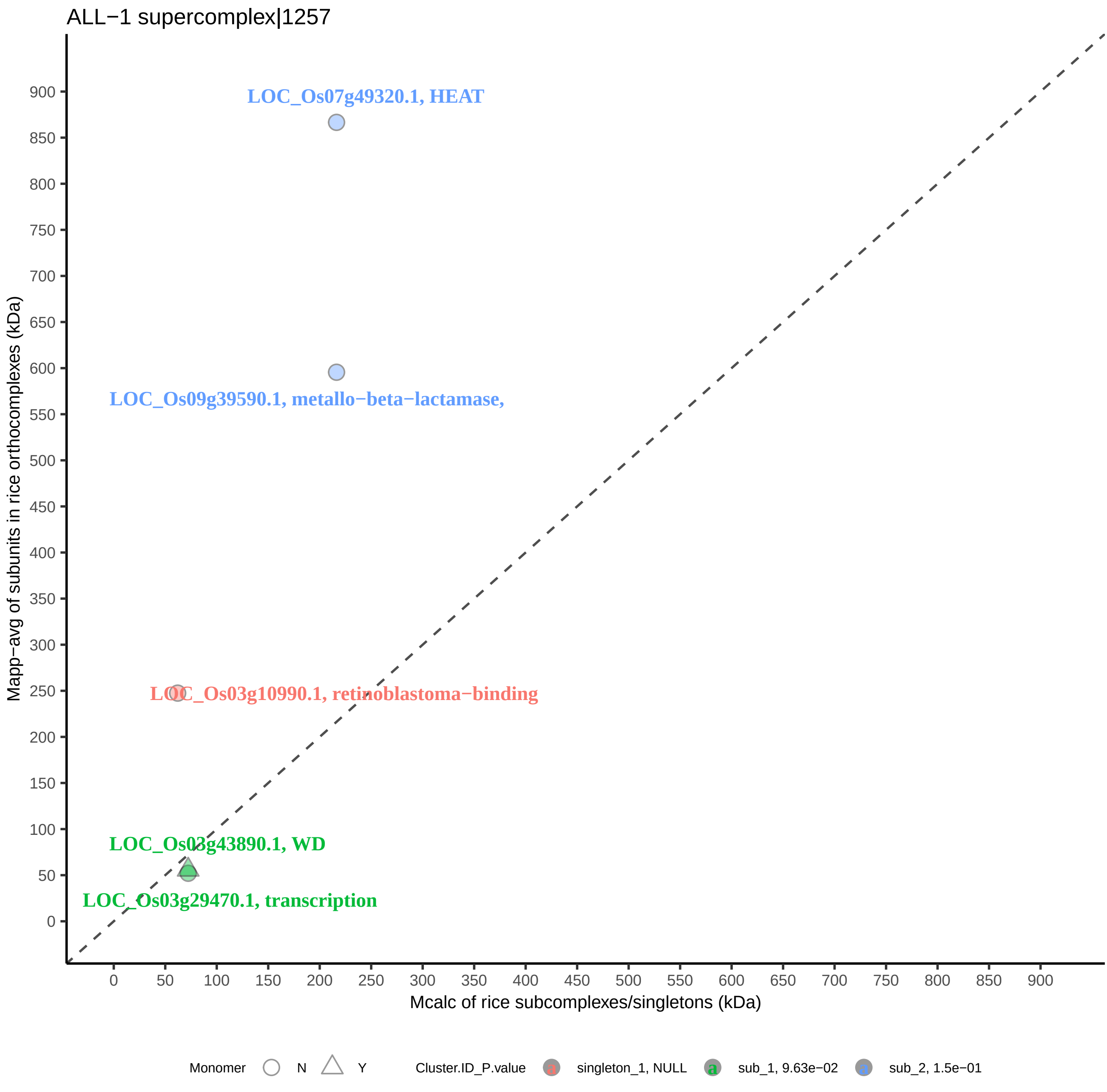

ALL−1 supercomplex|1257
LOC_Os07g49320.1, HEAT
LOC_Os09g39590.1, metallo−beta−lactamase,
LOC_Os03g10990.1, retinoblastoma−binding
LOC_Os03g43890.1, WD
LOC_Os03g29470.1, transcription
Mapp−avg of subunits in rice orthocomplexes (kDa)
Mcalc of rice subcomplexes/singletons (kDa)
0
50
100
150
200
250
300
350
400
450
500
550
600
650
700
750
800
850
900
Monomer
N
Y
Cluster.ID_P.value
singleton_1, NULL
sub_1, 9.63e−02
sub_2, 1.5e−01

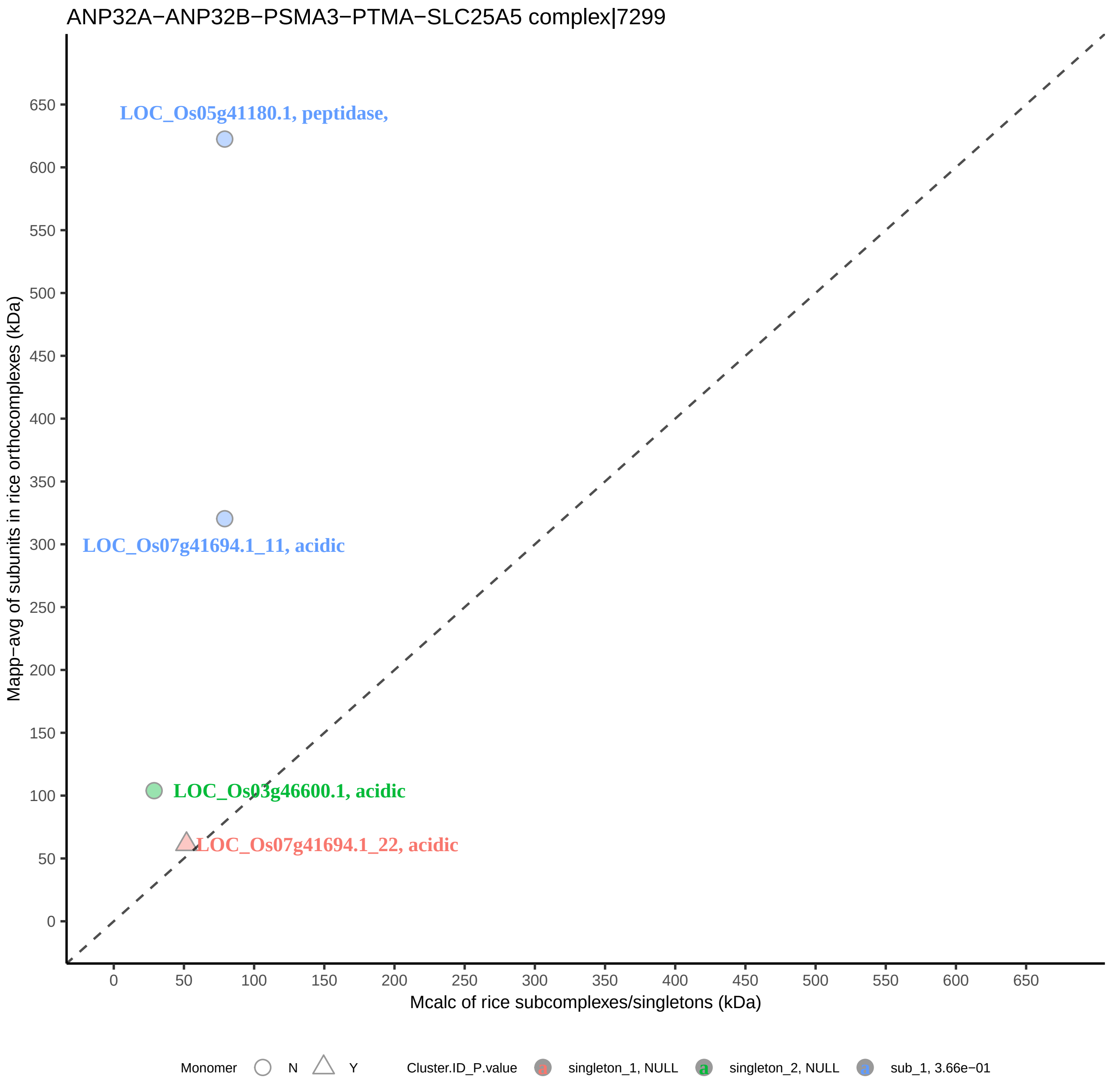
ANP32A−ANP32B−PSMA3−PTMA−SLC25A5 complex|7299
LOC_Os05g41180.1, peptidase,
LOC_Os07g41694.1_11, acidic
LOC_Os03g46600.1, acidic
LOC_Os07g41694.1_22, acidic
Mapp−avg of subunits in rice orthocomplexes (kDa)
Mcalc of rice subcomplexes/singletons (kDa)
0
50
100
150
200
250
300
350
400
450
500
550
600
650
Monomer
N
Y
Cluster.ID_P.value
singleton_1, NULL
singleton_2, NULL
sub_1, 3.66e−01

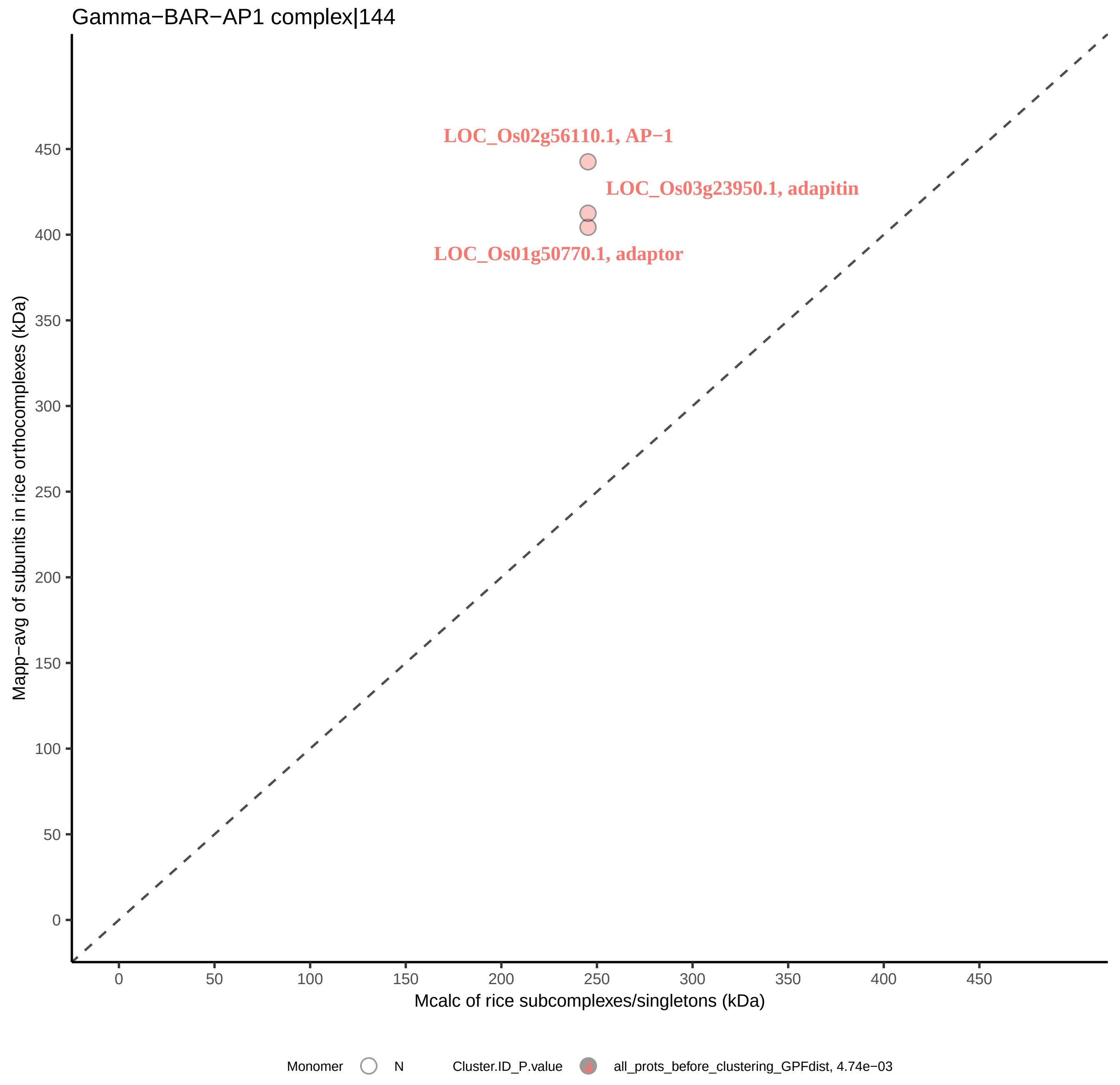

Gamma−BAR−AP1 complex|144
LOC_Os02g56110.1, AP−1
LOC_Os03g23950.1, adapitin
LOC_Os01g50770.1, adaptor
Mapp−avg of subunits in rice orthocomplexes (kDa)
Mcalc of rice subcomplexes/singletons (kDa)
0
50
100
150
200
250
300
350
400
450
Monomer
N
Cluster.ID_P.value
all_prots_before_clustering_GPFdist, 4.74e−03

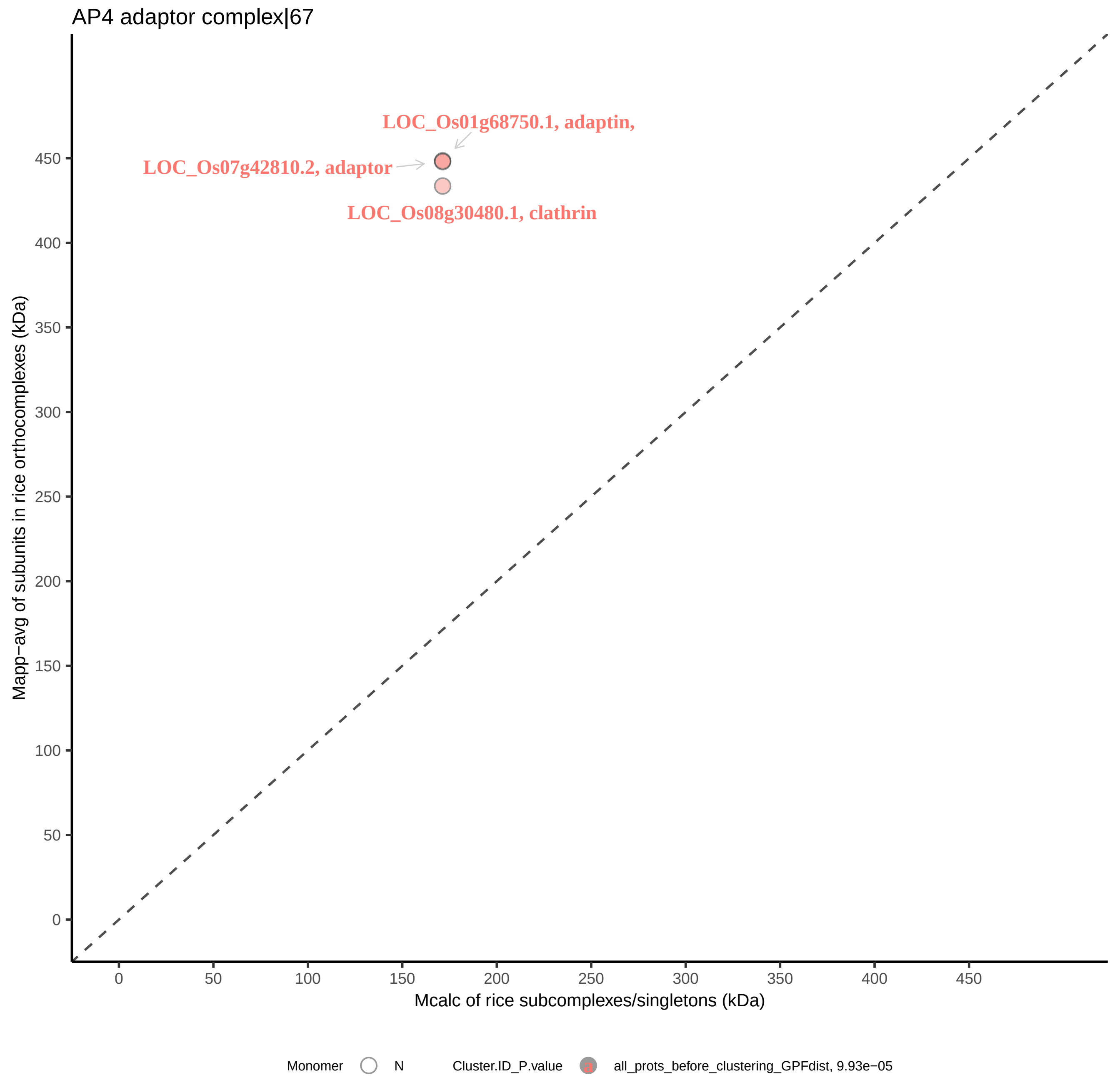

AP4 adaptor complex|67
LOC_Os01g68750.1, adaptin,
LOC_Os07g42810.2, adaptor
LOC_Os08g30480.1, clathrin
Mapp−avg of subunits in rice orthocomplexes (kDa)
Mcalc of rice subcomplexes/singletons (kDa)
0
50
100
150
200
250
300
350
400
450
Monomer
N
Cluster.ID_P.value
all_prots_before_clustering_GPFdist, 9.93e−05

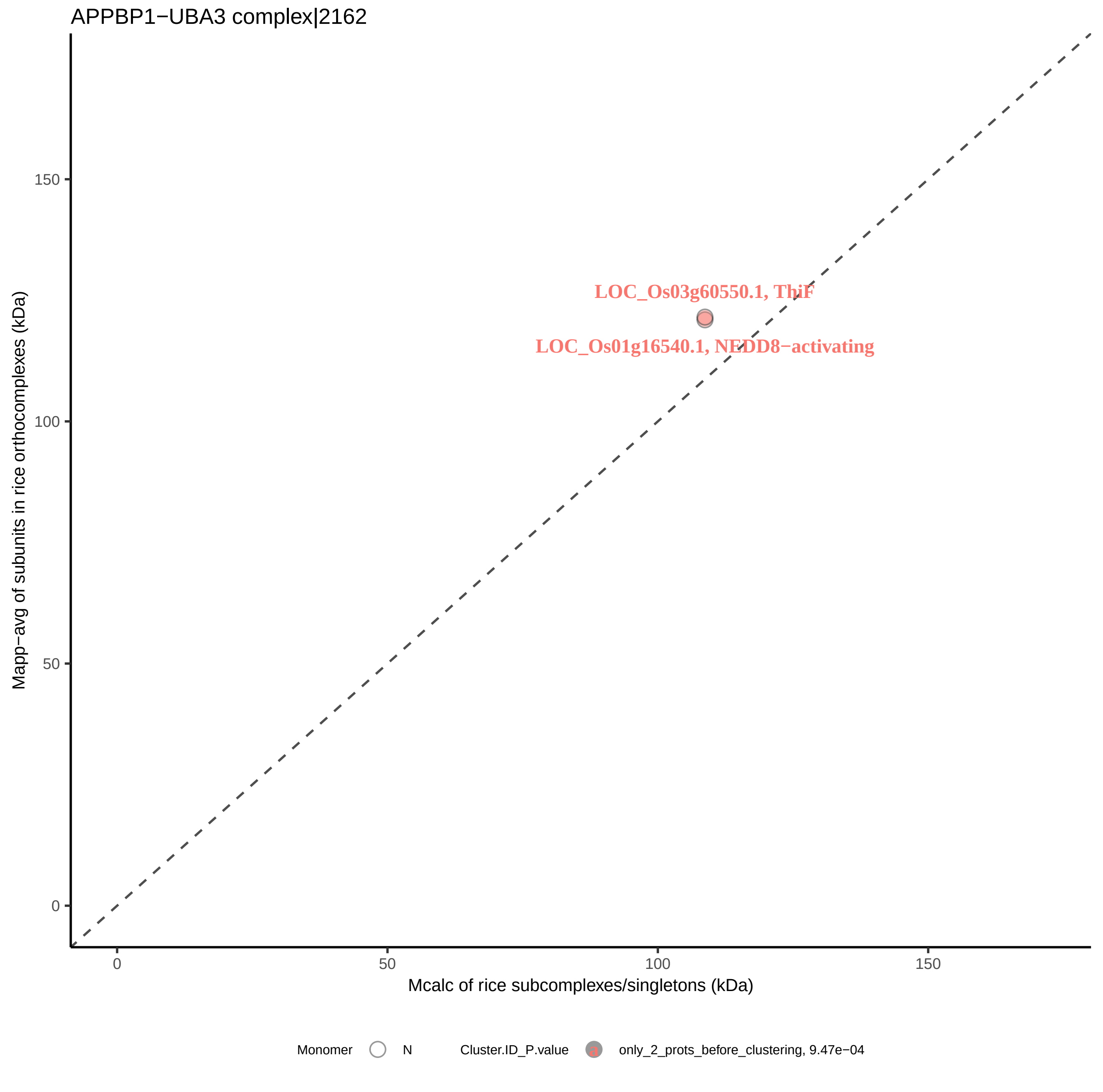

APPBP1−UBA3 complex|2162
LOC_Os03g60550.1, ThiF
LOC_Os01g16540.1, NEDD8−activating
Mapp−avg of subunits in rice orthocomplexes (kDa)
Mcalc of rice subcomplexes/singletons (kDa)
0
50
100
150
Monomer
N
Cluster.ID_P.value
only_2_prots_before_clustering, 9.47e−04

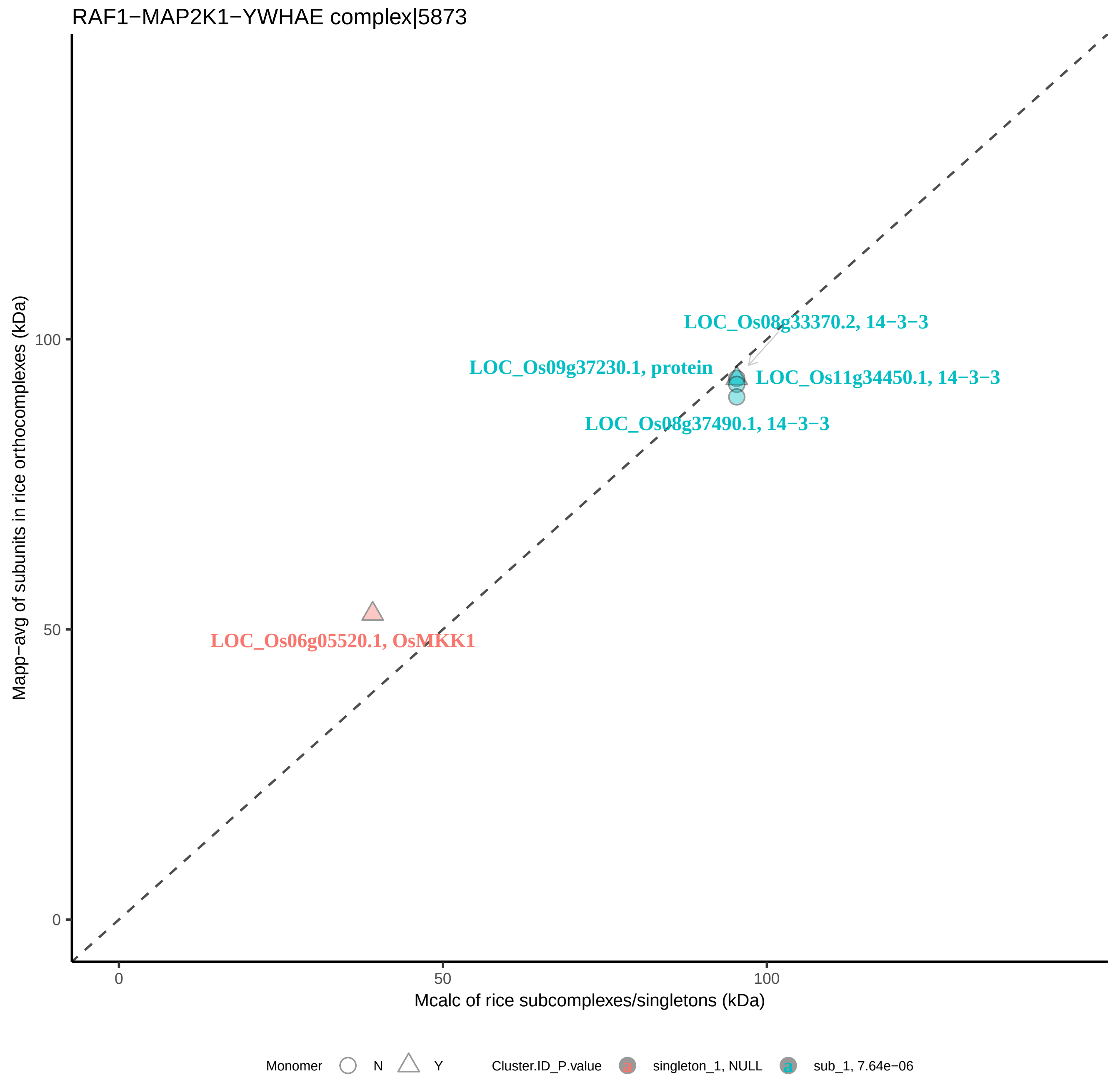

RAF1−MAP2K1−YWHAE complex|5873
LOC_Os08g33370.2, 14−3−3
LOC_Os09g37230.1, protein
LOC_Os11g34450.1, 14−3−3
LOC_Os08g37490.1, 14−3−3
LOC_Os06g05520.1, OsMKK1
Mapp−avg of subunits in rice orthocomplexes (kDa)
Mcalc of rice subcomplexes/singletons (kDa)
0
50
100
0
50
100
Monomer
N
Y
Cluster.ID_P.value
singleton_1, NULL
sub_1, 7.64e−06

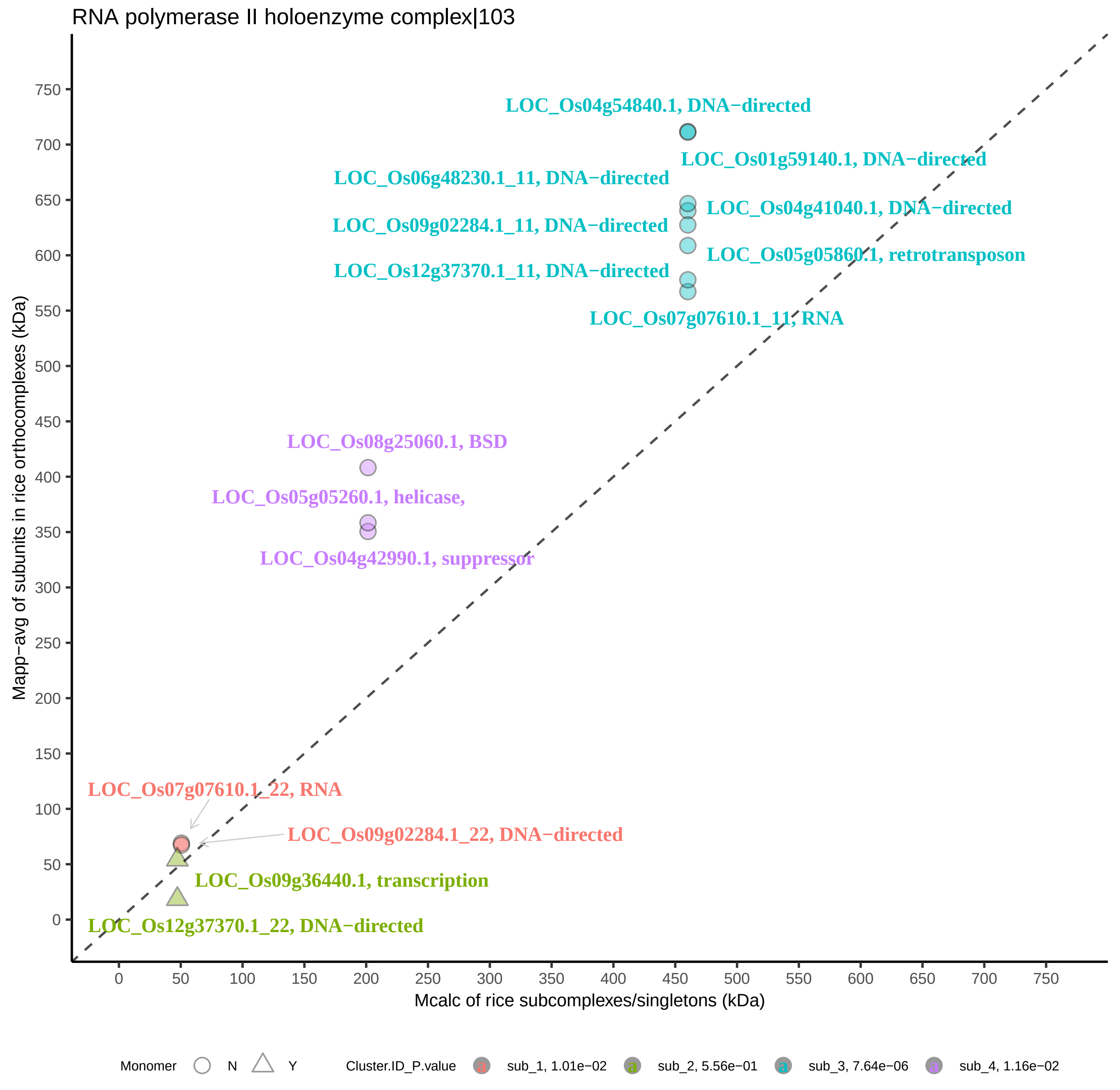
RNA polymerase II holoenzyme complex|103
LOC_Os04g54840.1, DNA−directed
LOC_Os01g59140.1, DNA−directed
LOC_Os06g48230.1_11, DNA−directed
LOC_Os04g41040.1, DNA−directed
LOC_Os09g02284.1_11, DNA−directed
LOC_Os05g05860.1, retrotransposon
LOC_Os12g37370.1_11, DNA−directed
LOC_Os07g07610.1_11, RNA
LOC_Os08g25060.1, BSD
LOC_Os05g05260.1, helicase,
LOC_Os04g42990.1, suppressor
LOC_Os07g07610.1_22, RNA
LOC_Os09g02284.1_22, DNA−directed
LOC_Os09g36440.1, transcription
LOC_Os12g37370.1_22, DNA−directed
Mapp−avg of subunits in rice orthocomplexes (kDa)
Mcalc of rice subcomplexes/singletons (kDa)
0 50 100 150 200 250 300 350 400 450 500 550 600 650 700 750
Monomer N Y
Cluster.ID_P.value
sub_1, 1.01e−02
sub_2, 5.56e−01
sub_3, 7.64e−06
sub_4, 1.16e−02

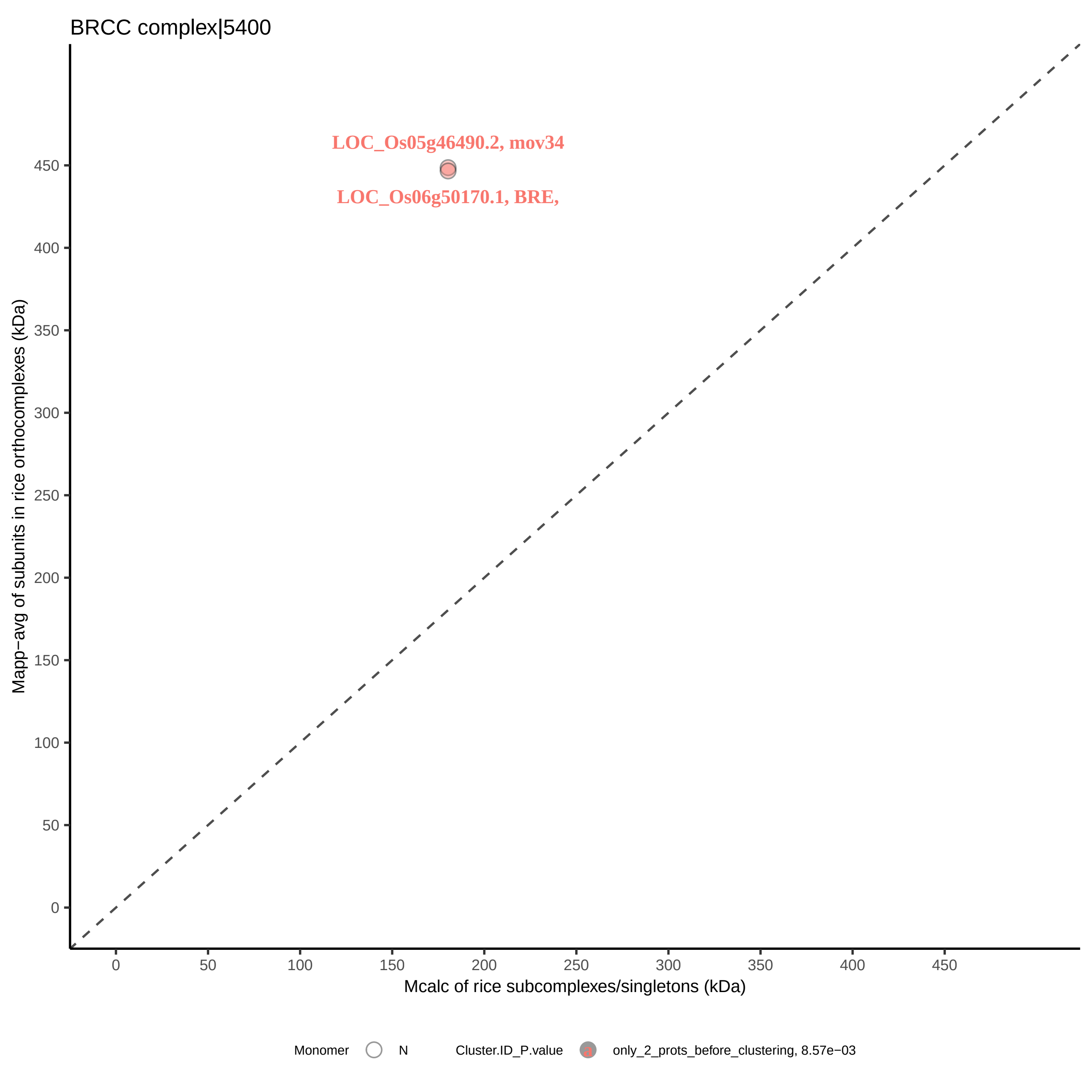

BRCC complex|5400
LOC_Os05g46490.2, mov34
LOC_Os06g50170.1, BRE,
Mapp−avg of subunits in rice orthocomplexes (kDa)
0
50
100
150
200
250
300
350
400
450
Mcalc of rice subcomplexes/singletons (kDa)
Monomer
N
Cluster.ID_P.value
only_2_prots_before_clustering, 8.57e−03

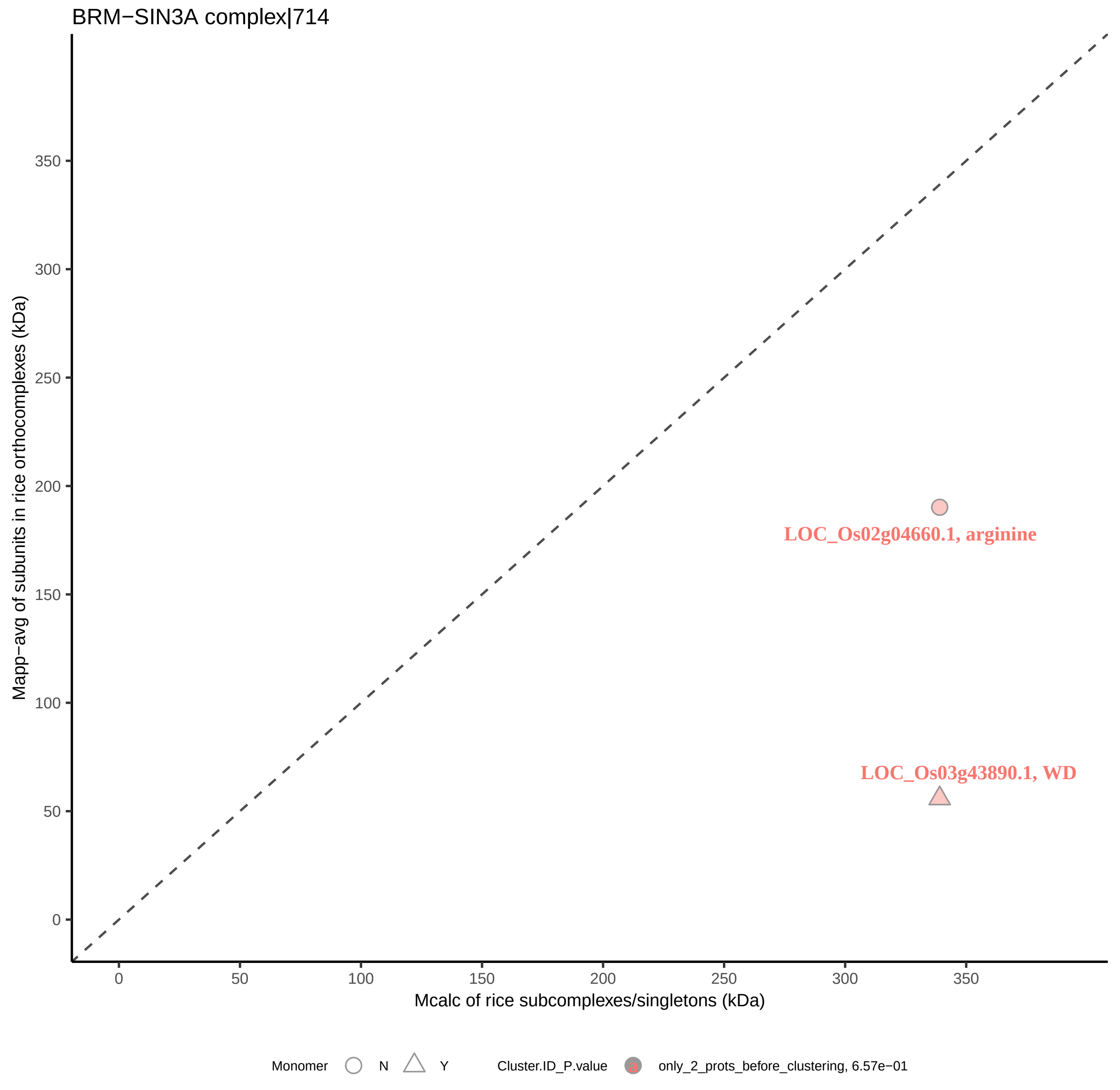
BRM−SIN3A complex|714
Mapp−avg of subunits in rice orthocomplexes (kDa)
350
300
250
200
150
100
50
0
LOC_Os02g04660.1, arginine
LOC_Os03g43890.1, WD
0
50
100
150
200
250
300
350
Mcalc of rice subcomplexes/singletons (kDa)
Monomer
N
Y
Cluster.ID_P.value
only_2_prots_before_clustering, 6.57e−01

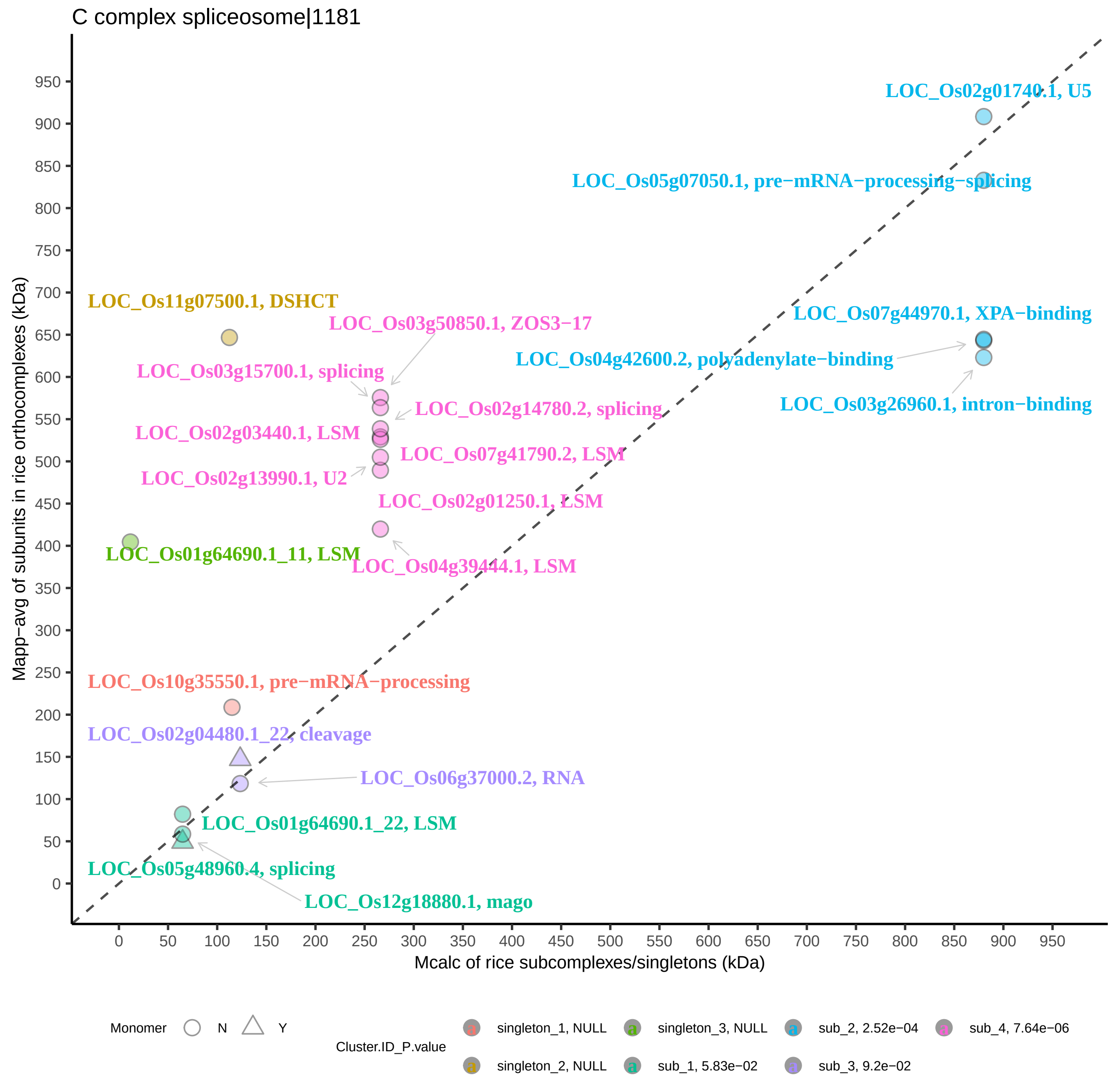
C complex spliceosome|1181
Mapp−avg of subunits in rice orthocomplexes (kDa)
Mcalc of rice subcomplexes/singletons (kDa)
0
50
100
150
200
250
300
350
400
450
500
550
600
650
700
750
800
850
900
950
LOC_Os02g01740.1, U5
LOC_Os05g07050.1, pre−mRNA−processing−splicing
LOC_Os07g44970.1, XPA−binding
LOC_Os04g42600.2, polyadenylate−binding
LOC_Os03g26960.1, intron−binding
LOC_Os11g07500.1, DSHCT
LOC_Os03g50850.1, ZOS3−17
LOC_Os03g15700.1, splicing
LOC_Os02g14780.2, splicing
LOC_Os02g03440.1, LSM
LOC_Os07g41790.2, LSM
LOC_Os02g13990.1, U2
LOC_Os02g01250.1, LSM
LOC_Os01g64690.1_11, LSM
LOC_Os04g39444.1, LSM
LOC_Os10g35550.1, pre−mRNA−processing
LOC_Os02g04480.1_22, cleavage
LOC_Os06g37000.2, RNA
LOC_Os01g64690.1_22, LSM
LOC_Os05g48960.4, splicing
LOC_Os12g18880.1, mago
Monomer
N
Y
Cluster.ID_P.value
singleton_1, NULL
singleton_2, NULL
singleton_3, NULL
sub_1, 5.83e−02
sub_2, 2.52e−04
sub_3, 9.2e−02
sub_4, 7.64e−06

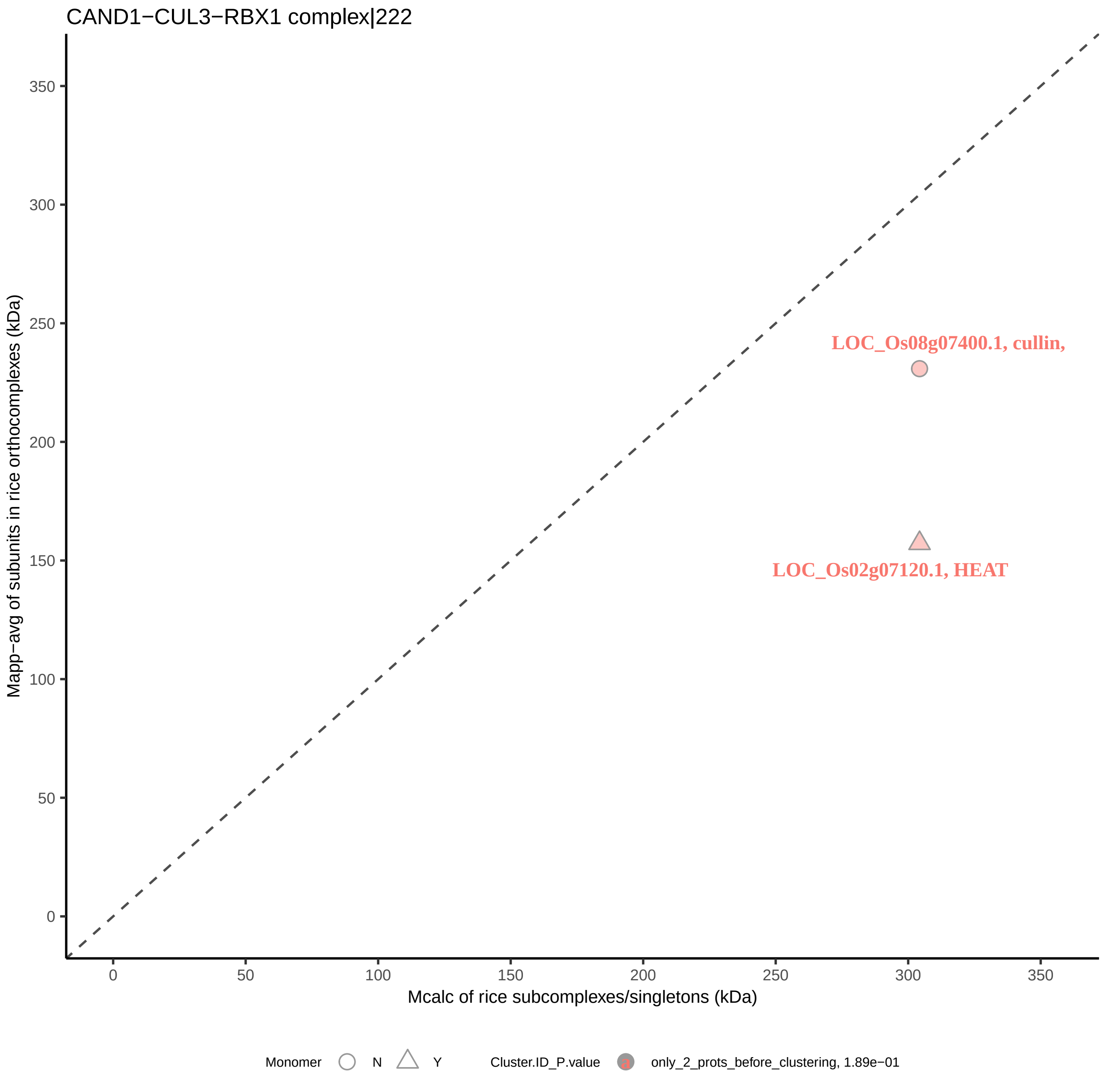

CAND1−CUL3−RBX1 complex|222
Mapp−avg of subunits in rice orthocomplexes (kDa)
350
300
250
200
150
100
50
0
LOC_Os08g07400.1, cullin,
LOC_Os02g07120.1, HEAT
0
50
100
150
200
250
300
350
Mcalc of rice subcomplexes/singletons (kDa)
Monomer
N
Y
Cluster.ID_P.value
a
only_2_prots_before_clustering, 1.89e−01

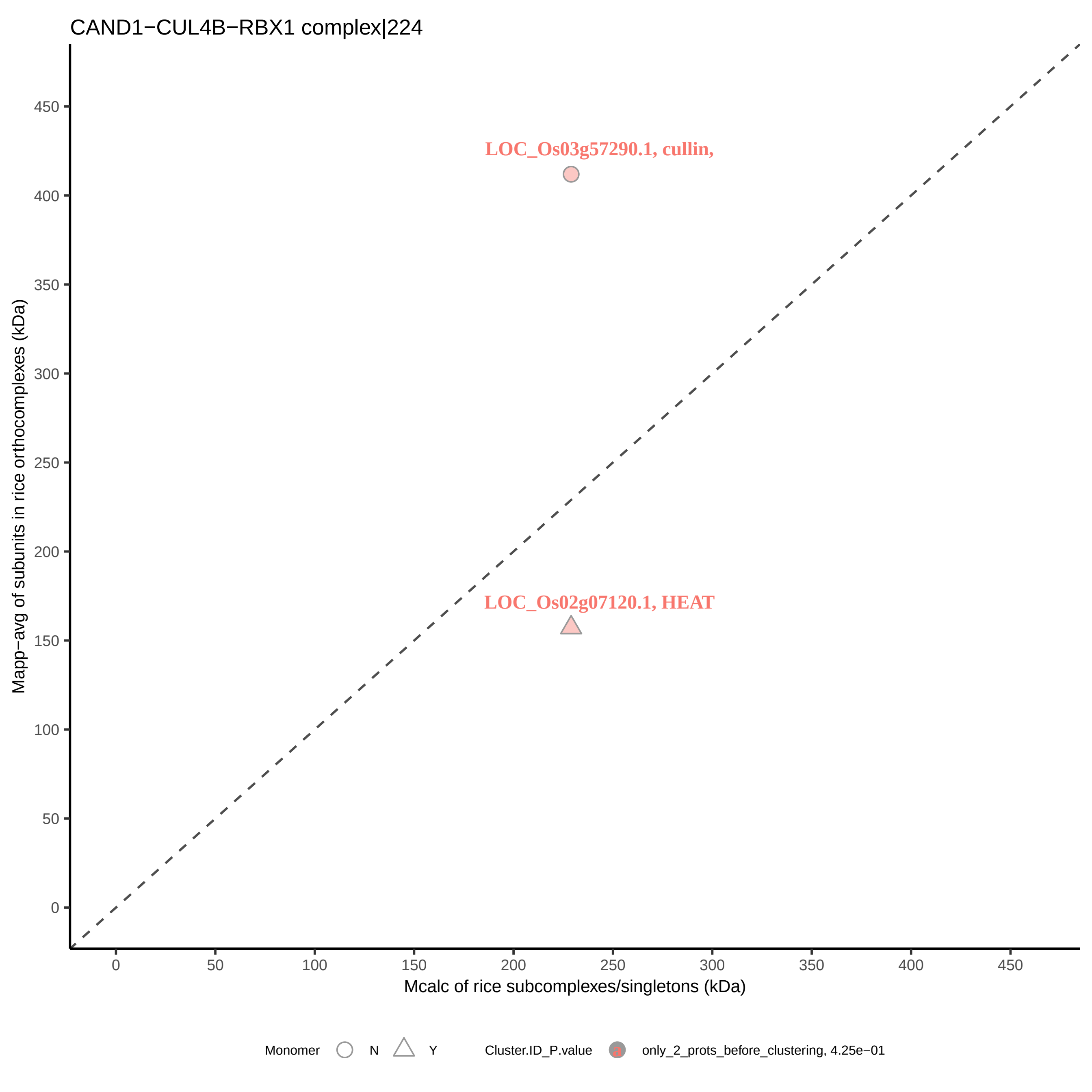
CAND1−CUL4B−RBX1 complex|224
LOC_Os03g57290.1, cullin,
LOC_Os02g07120.1, HEAT
Mapp−avg of subunits in rice orthocomplexes (kDa)
Mcalc of rice subcomplexes/singletons (kDa)
0
50
100
150
200
250
300
350
400
450
Monomer
N
Y
Cluster.ID_P.value
only_2_prots_before_clustering, 4.25e−01

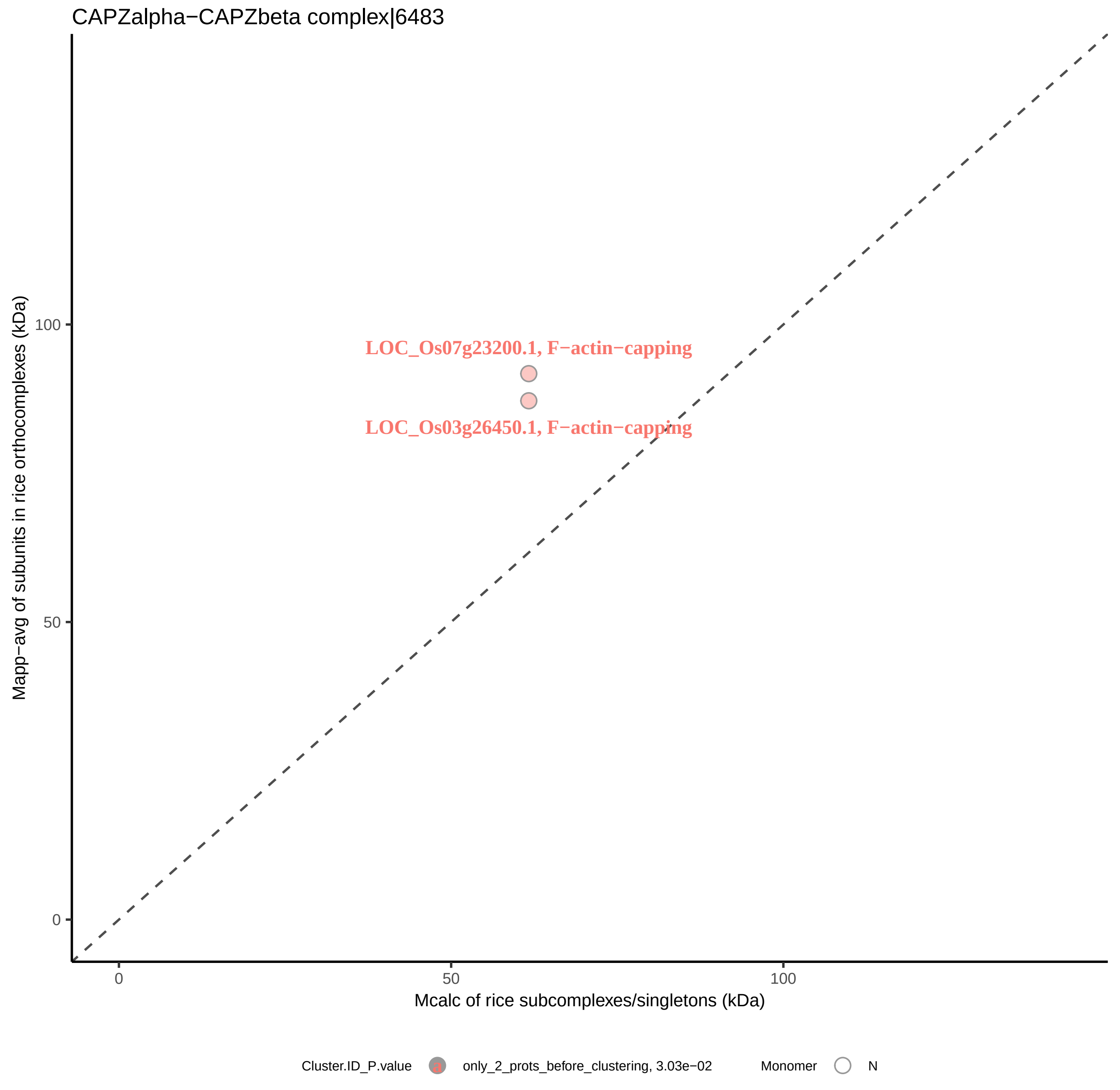
CAPZalpha−CAPZbeta complex|6483
LOC_Os07g23200.1, F−actin−capping
LOC_Os03g26450.1, F−actin−capping
100
50
0
0
50
100
Mapp−avg of subunits in rice orthocomplexes (kDa)
Mcalc of rice subcomplexes/singletons (kDa)
Cluster.ID_P.value
only_2_prots_before_clustering, 3.03e−02
Monomer
N

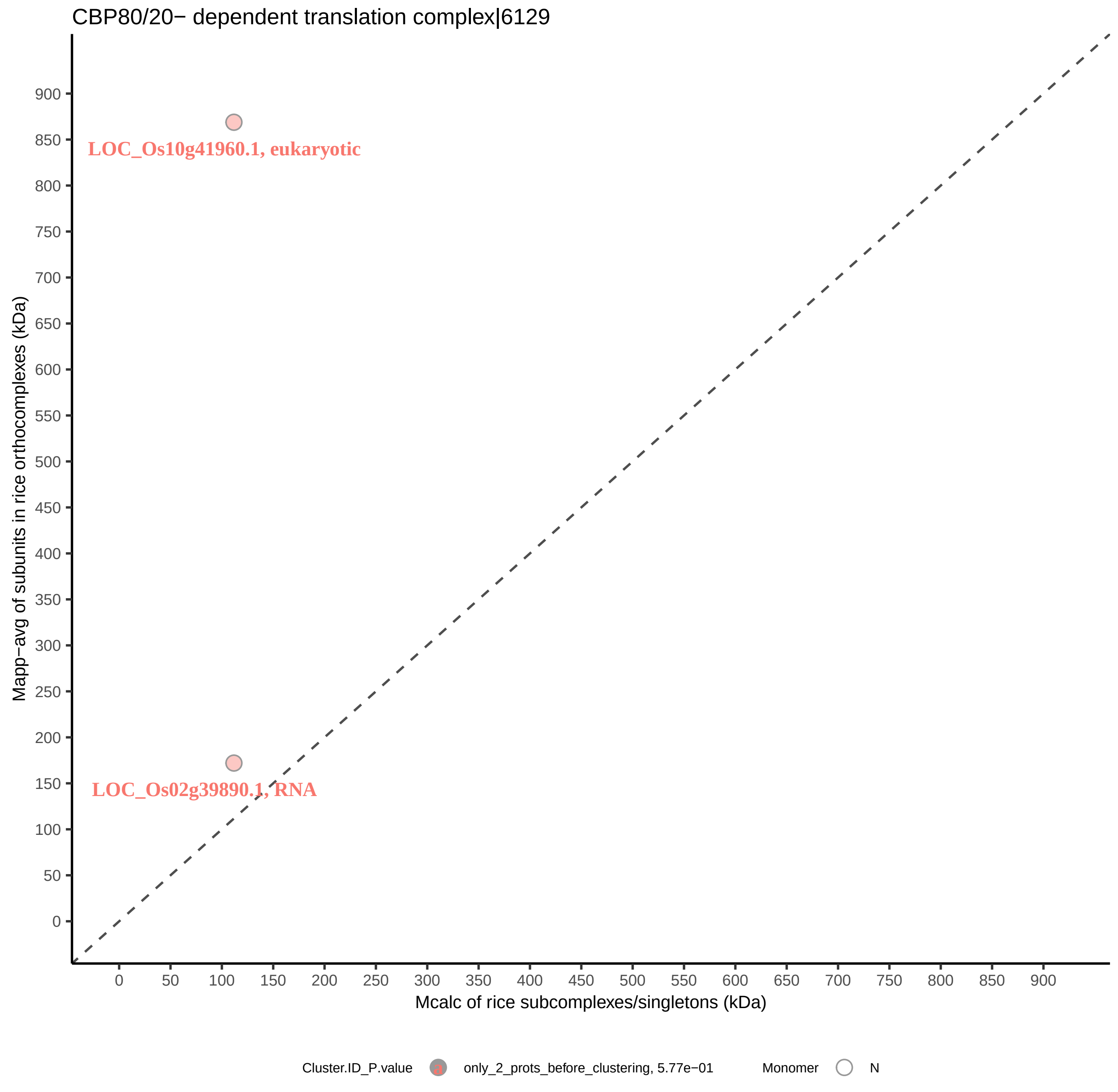
CBP80/20− dependent translation complex|6129
LOC_Os10g41960.1, eukaryotic
LOC_Os02g39890.1, RNA
Mapp−avg of subunits in rice orthocomplexes (kDa)
Mcalc of rice subcomplexes/singletons (kDa)
0
50
100
150
200
250
300
350
400
450
500
550
600
650
700
750
800
850
900
Cluster.ID_P.value
only_2_prots_before_clustering, 5.77e−01
Monomer
N

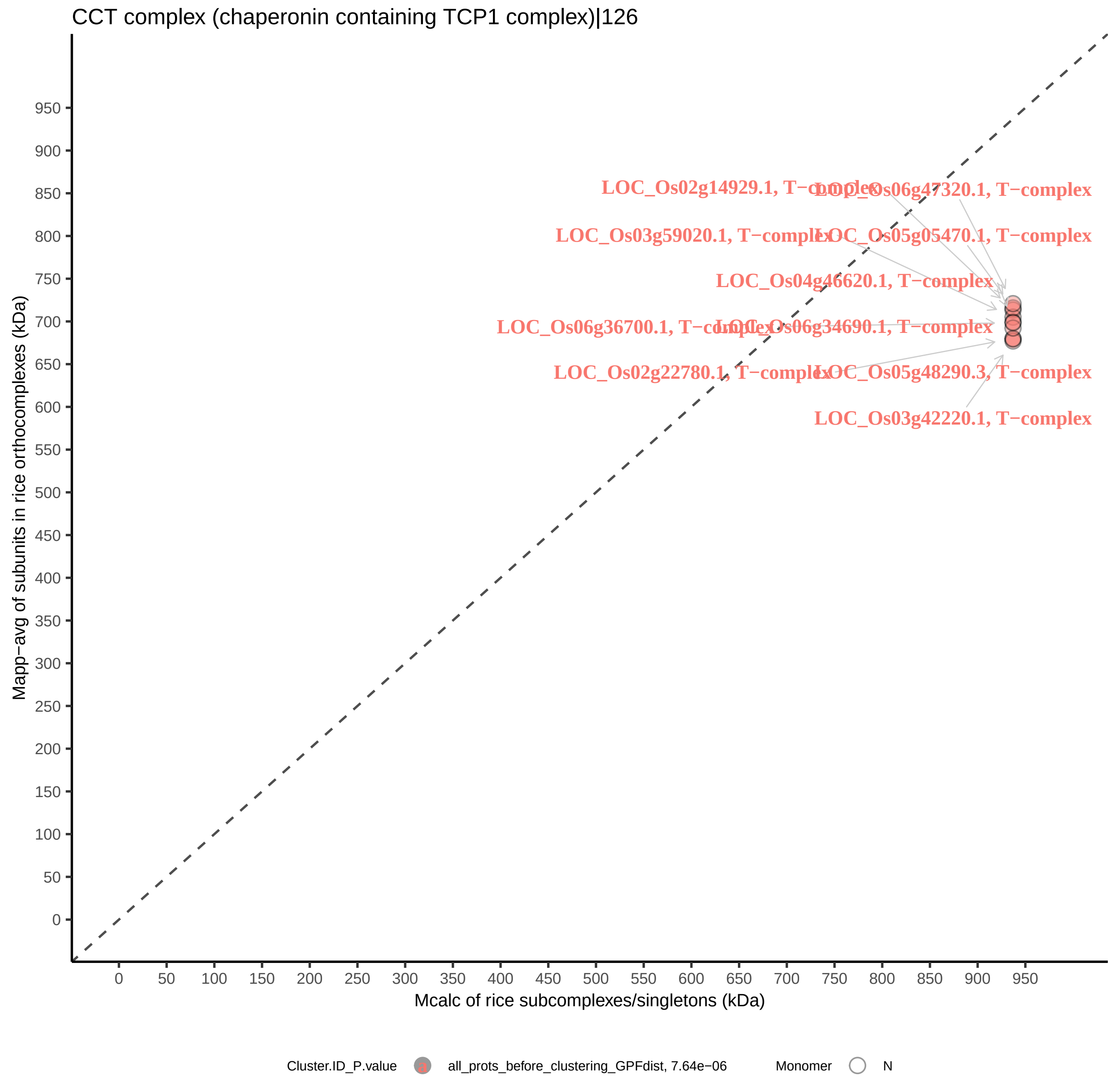
CCT complex (chaperonin containing TCP1 complex)|126
Mapp−avg of subunits in rice orthocomplexes (kDa)
Mcalc of rice subcomplexes/singletons (kDa)
0
50
100
150
200
250
300
350
400
450
500
550
600
650
700
750
800
850
900
950
LOC_Os02g14929.1, T−complex
LOC_Os06g47320.1, T−complex
LOC_Os03g59020.1, T−complex
LOC_Os05g05470.1, T−complex
LOC_Os04g46620.1, T−complex
LOC_Os06g36700.1, T−complex
LOC_Os06g34690.1, T−complex
LOC_Os02g22780.1, T−complex
LOC_Os05g48290.3, T−complex
LOC_Os03g42220.1, T−complex
Cluster.ID_P.value
all_prots_before_clustering_GPFdist, 7.64e−06
Monomer
N

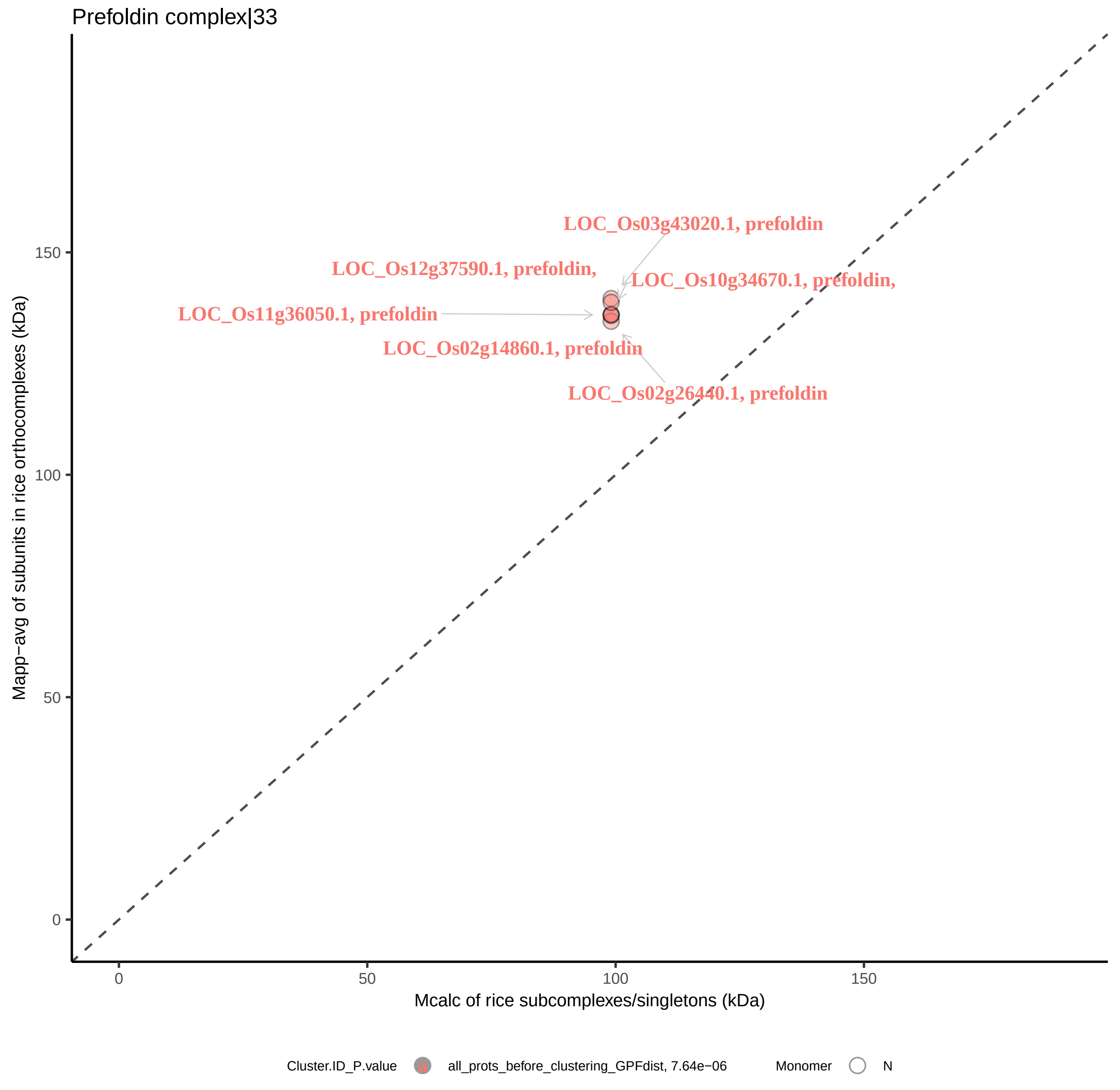

Prefoldin complex|33
LOC_Os03g43020.1, prefoldin
LOC_Os12g37590.1, prefoldin,
LOC_Os10g34670.1, prefoldin,
LOC_Os11g36050.1, prefoldin
LOC_Os02g14860.1, prefoldin
LOC_Os02g26440.1, prefoldin
Mapp−avg of subunits in rice orthocomplexes (kDa)
150
100
50
0
0
50
100
150
Mcalc of rice subcomplexes/singletons (kDa)
Cluster.ID_P.value
all_prots_before_clustering_GPFdist, 7.64e−06
Monomer
N

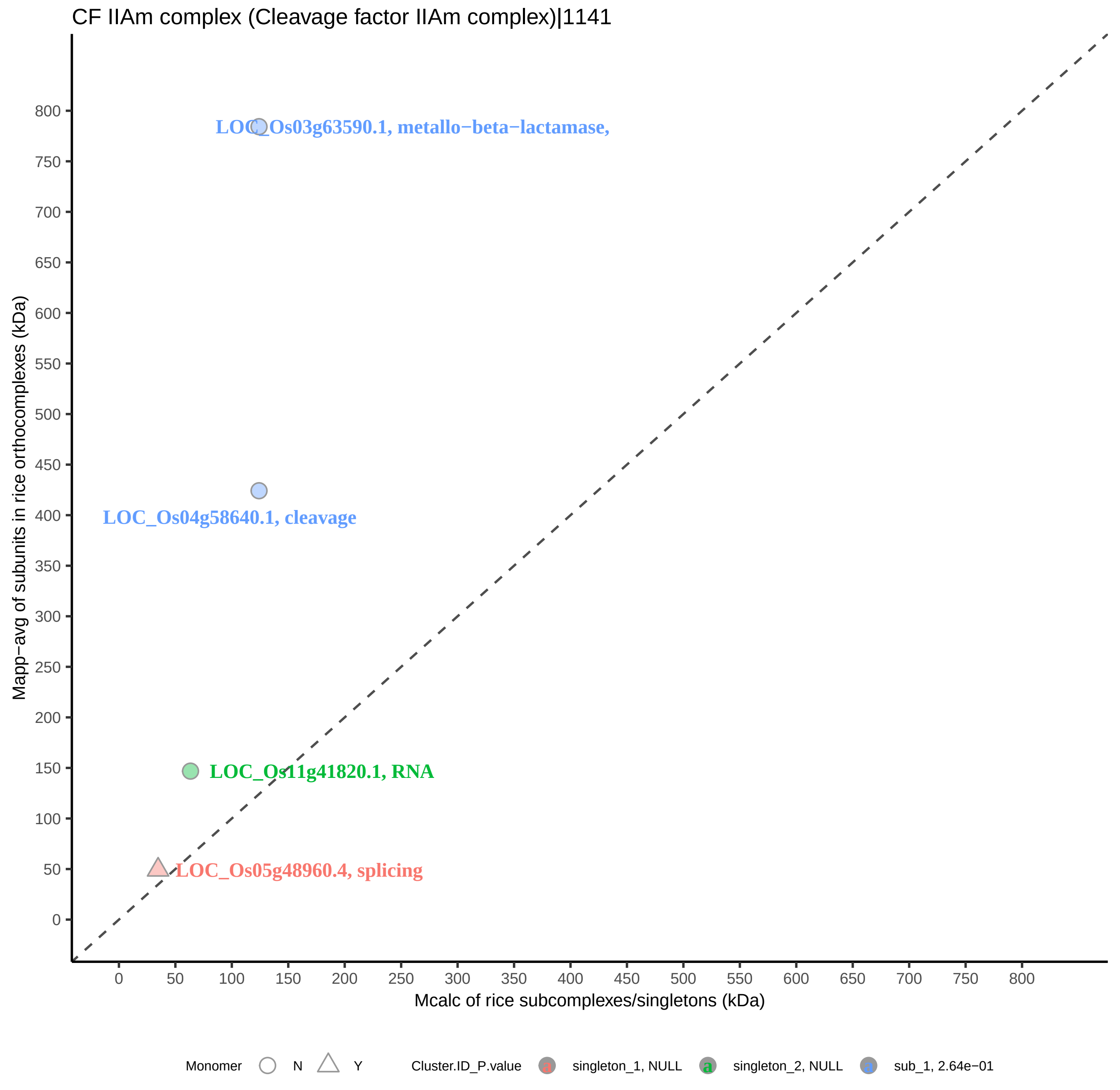
CF IIAm complex (Cleavage factor IIAm complex)|1141
LOC_Os03g63590.1, metallo−beta−lactamase,
LOC_Os04g58640.1, cleavage
LOC_Os11g41820.1, RNA
LOC_Os05g48960.4, splicing
Mapp−avg of subunits in rice orthocomplexes (kDa)
Mcalc of rice subcomplexes/singletons (kDa)
0
50
100
150
200
250
300
350
400
450
500
550
600
650
700
750
800
Monomer
N
Y
Cluster.ID_P.value
a
singleton_1, NULL
a
singleton_2, NULL
a
sub_1, 2.64e−01

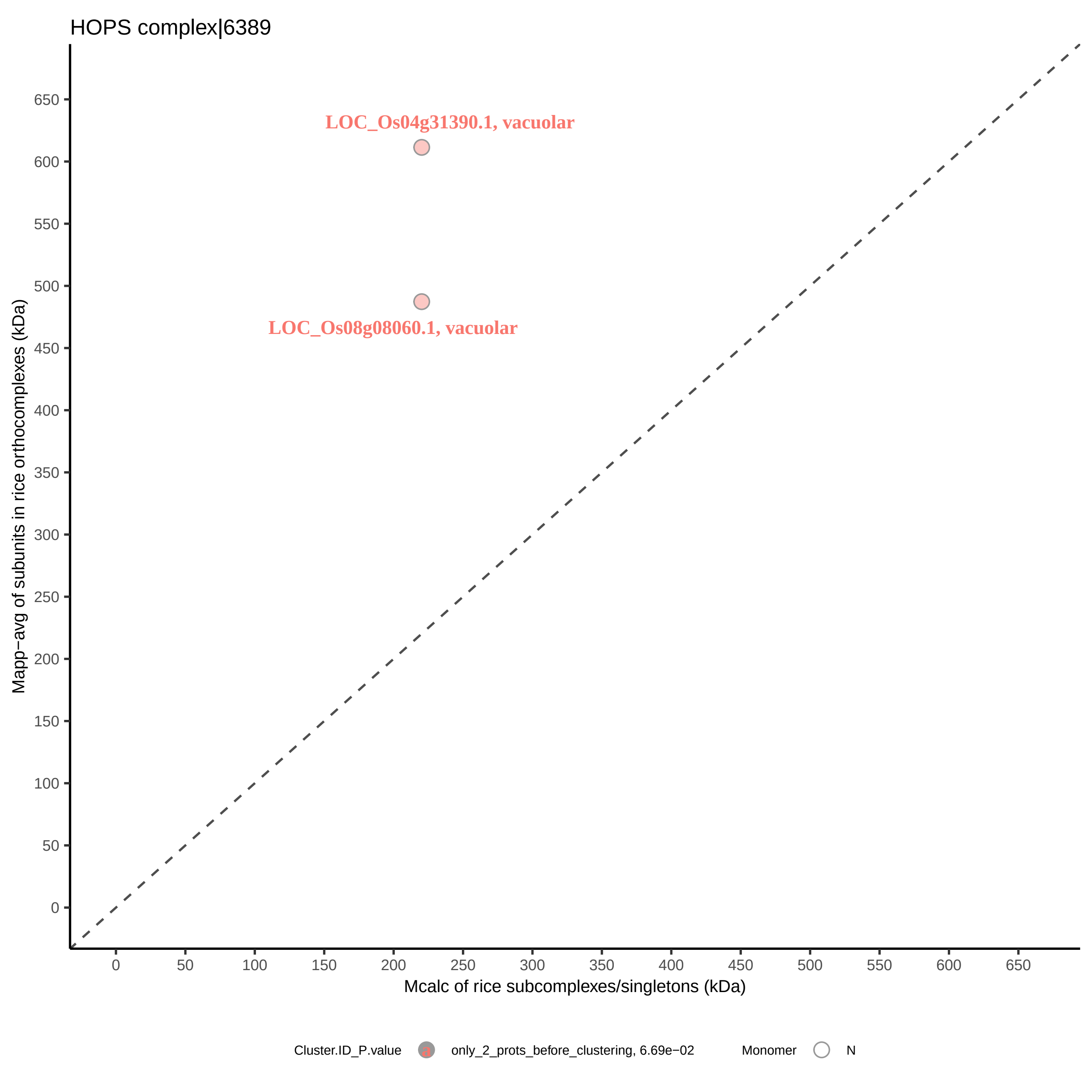
HOPS complex|6389
LOC_Os04g31390.1, vacuolar
LOC_Os08g08060.1, vacuolar
Mapp−avg of subunits in rice orthocomplexes (kDa)
Mcalc of rice subcomplexes/singletons (kDa)
0
50
100
150
200
250
300
350
400
450
500
550
600
650
Cluster.ID_P.value
only_2_prots_before_clustering, 6.69e−02
Monomer
N

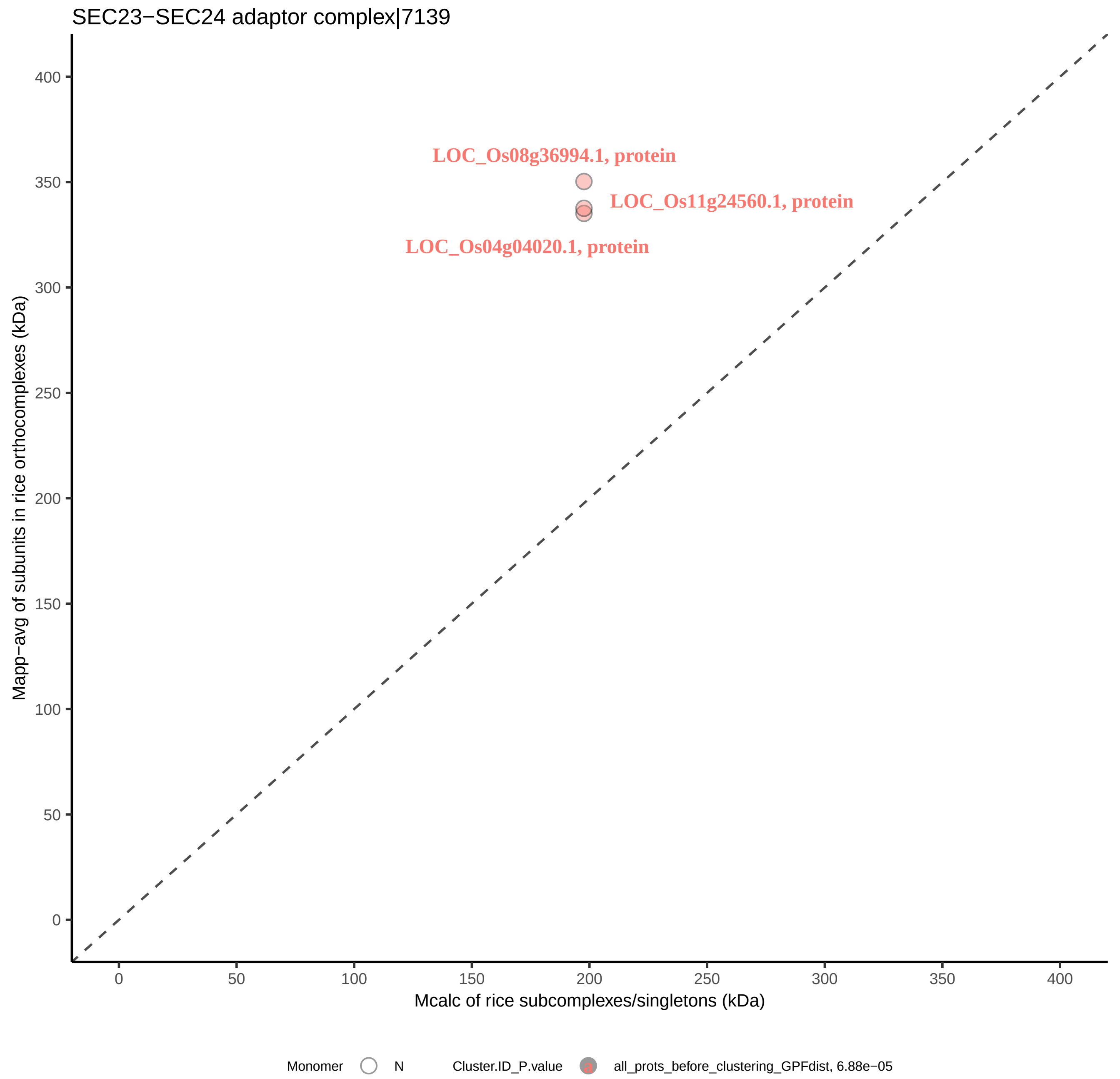
SEC23−SEC24 adaptor complex|7139
LOC_Os08g36994.1, protein
LOC_Os11g24560.1, protein
LOC_Os04g04020.1, protein
Mapp−avg of subunits in rice orthocomplexes (kDa)
Mcalc of rice subcomplexes/singletons (kDa)
0
50
100
150
200
250
300
350
400
Monomer
N
Cluster.ID_P.value
all_prots_before_clustering_GPFdist, 6.88e−05

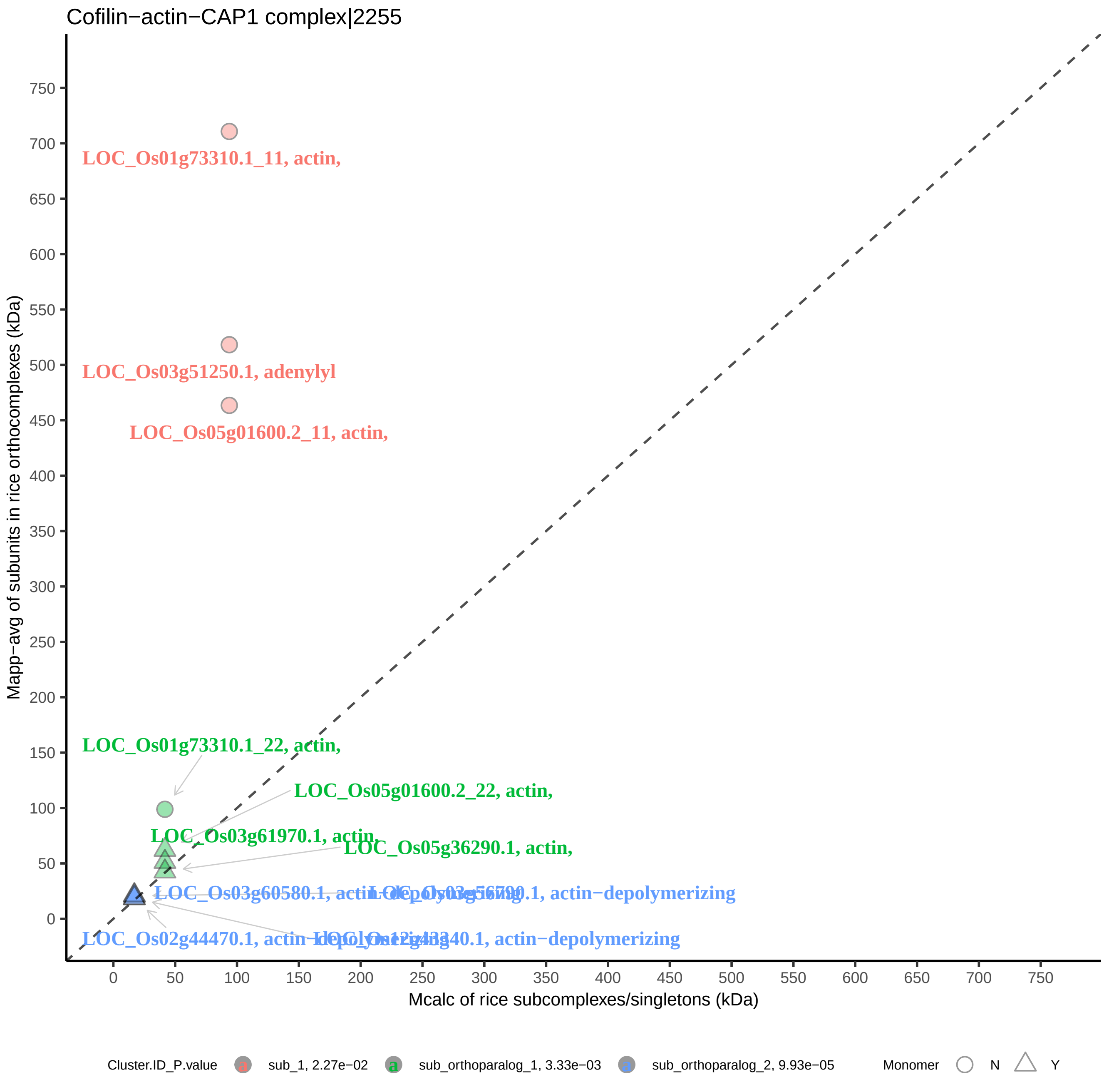

Cofilin−actin−CAP1 complex|2255
LOC_Os01g73310.1_11, actin,
LOC_Os03g51250.1, adenylyl
LOC_Os05g01600.2_11, actin,
LOC_Os01g73310.1_22, actin,
LOC_Os05g01600.2_22, actin,
LOC_Os03g61970.1, actin,
LOC_Os05g36290.1, actin,
LOC_Os03g60580.1, actin−depolymerizing
LOC_Os03g56790.1, actin−depolymerizing
LOC_Os02g44470.1, actin−depolymerizing
LOC_Os12g43340.1, actin−depolymerizing
Mapp−avg of subunits in rice orthocomplexes (kDa)
Mcalc of rice subcomplexes/singletons (kDa)
0
50
100
150
200
250
300
350
400
450
500
550
600
650
700
750
Cluster.ID_P.value
sub_1, 2.27e−02
sub_orthoparalog_1, 3.33e−03
sub_orthoparalog_2, 9.93e−05
Monomer
N
Y

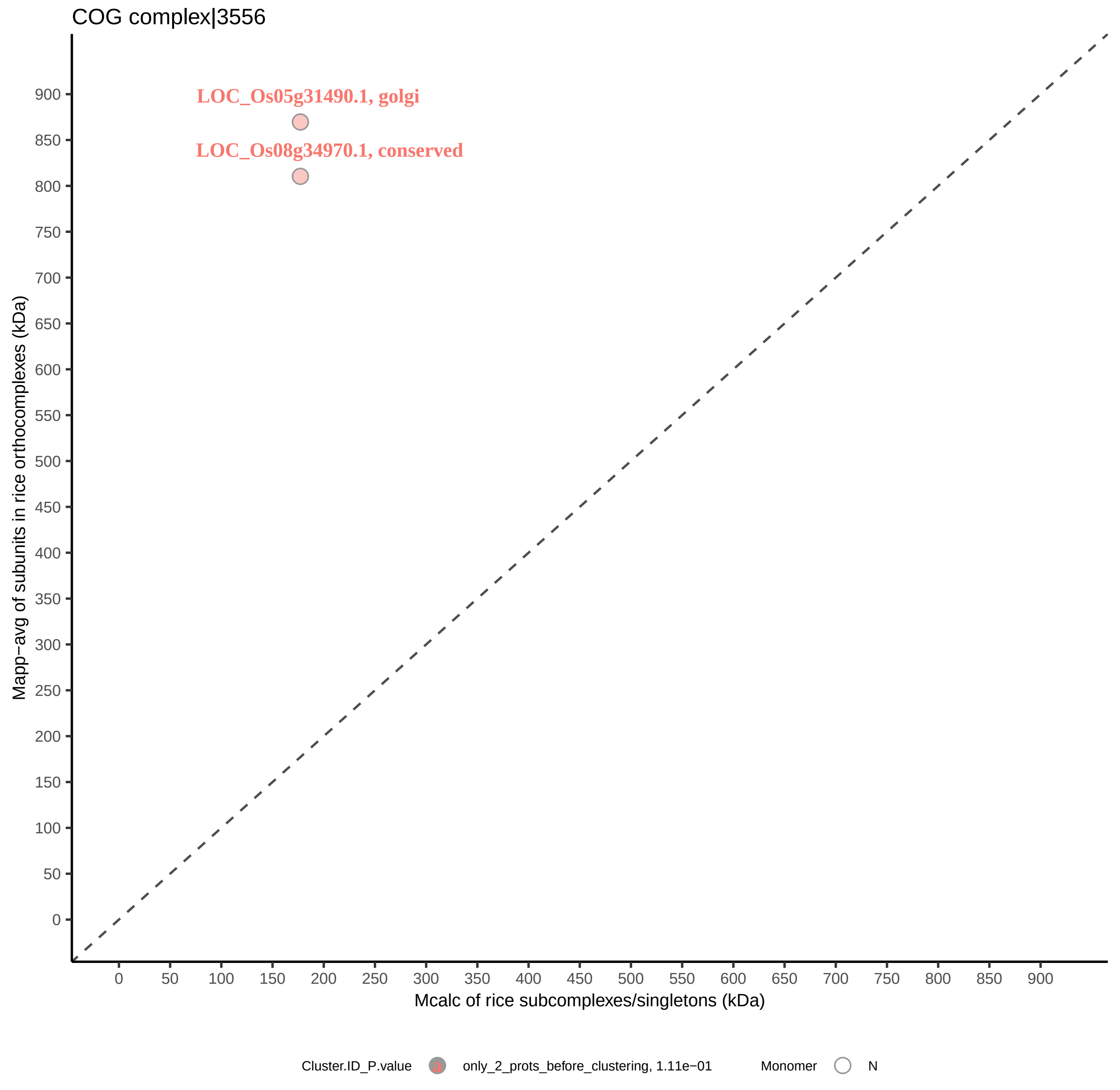

COG complex|3556
LOC_Os05g31490.1, golgi
LOC_Os08g34970.1, conserved
Mapp−avg of subunits in rice orthocomplexes (kDa)
Mcalc of rice subcomplexes/singletons (kDa)
0
50
100
150
200
250
300
350
400
450
500
550
600
650
700
750
800
850
900
Cluster.ID_P.value
only_2_prots_before_clustering, 1.11e−01
Monomer
N

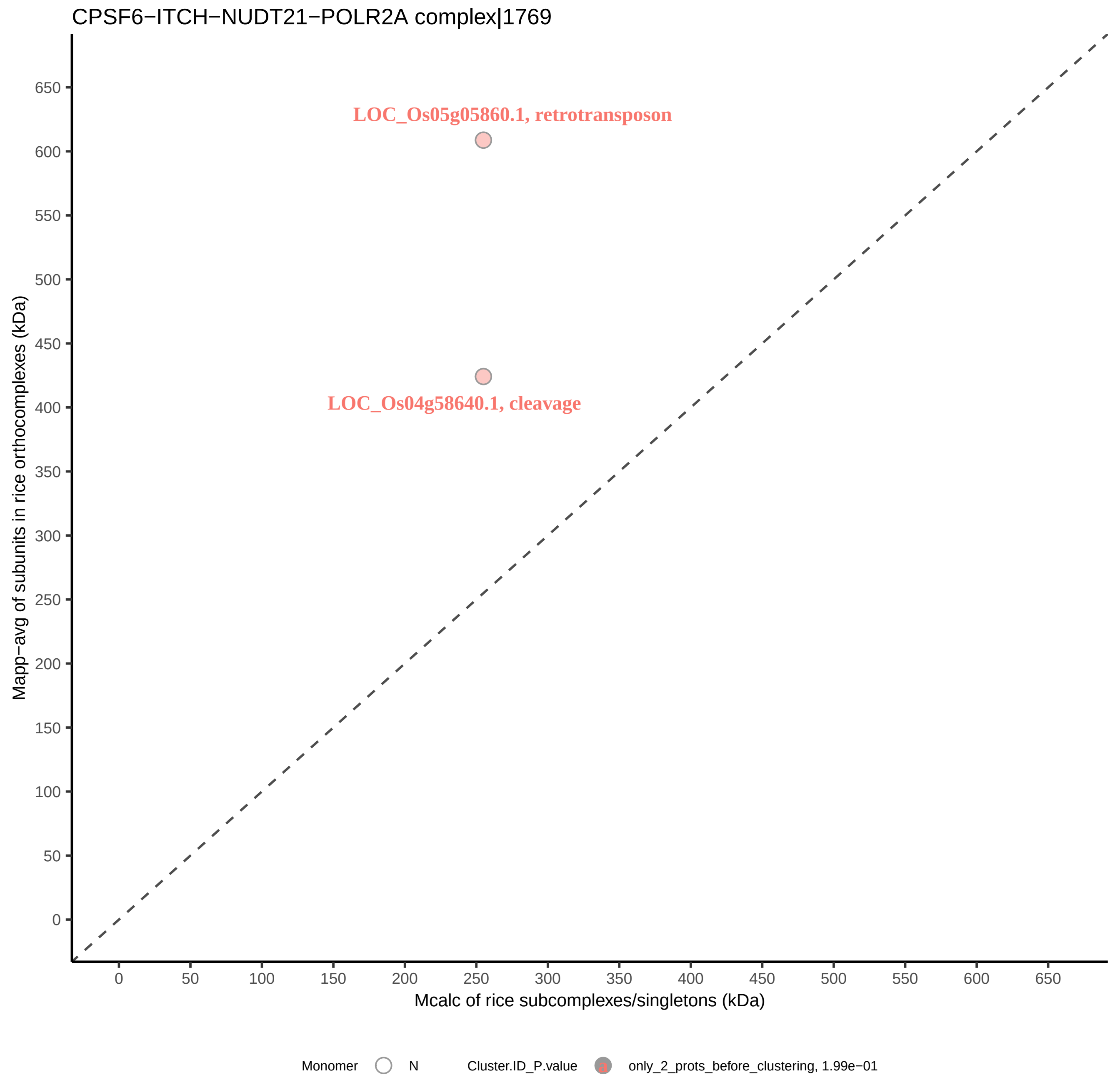

CPSF6−ITCH−NUDT21−POLR2A complex|1769
LOC_Os05g05860.1, retrotransposon
LOC_Os04g58640.1, cleavage
Mapp−avg of subunits in rice orthocomplexes (kDa)
Mcalc of rice subcomplexes/singletons (kDa)
0
50
100
150
200
250
300
350
400
450
500
550
600
650
Monomer
N
Cluster.ID_P.value
only_2_prots_before_clustering, 1.99e−01

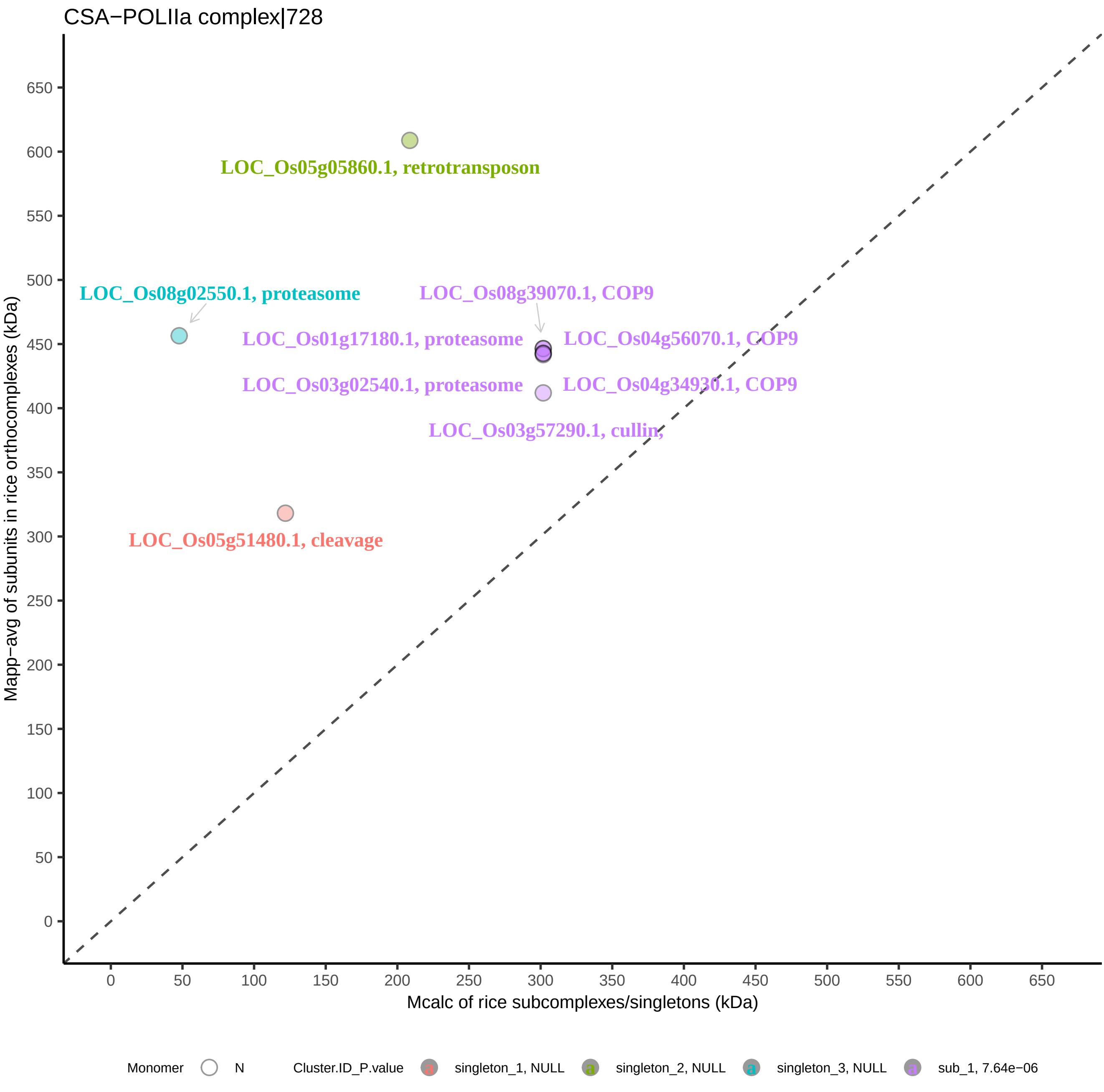
CSA−POLIIa complex|728
LOC_Os05g05860.1, retrotransposon
LOC_Os08g02550.1, proteasome
LOC_Os08g39070.1, COP9
LOC_Os01g17180.1, proteasome
LOC_Os04g56070.1, COP9
LOC_Os03g02540.1, proteasome
LOC_Os04g34930.1, COP9
LOC_Os03g57290.1, cullin,
LOC_Os05g51480.1, cleavage
Mapp−avg of subunits in rice orthocomplexes (kDa)
Mcalc of rice subcomplexes/singletons (kDa)
0
50
100
150
200
250
300
350
400
450
500
550
600
650
Monomer
N
Cluster.ID_P.value
singleton_1, NULL
singleton_2, NULL
singleton_3, NULL
sub_1, 7.64e−06

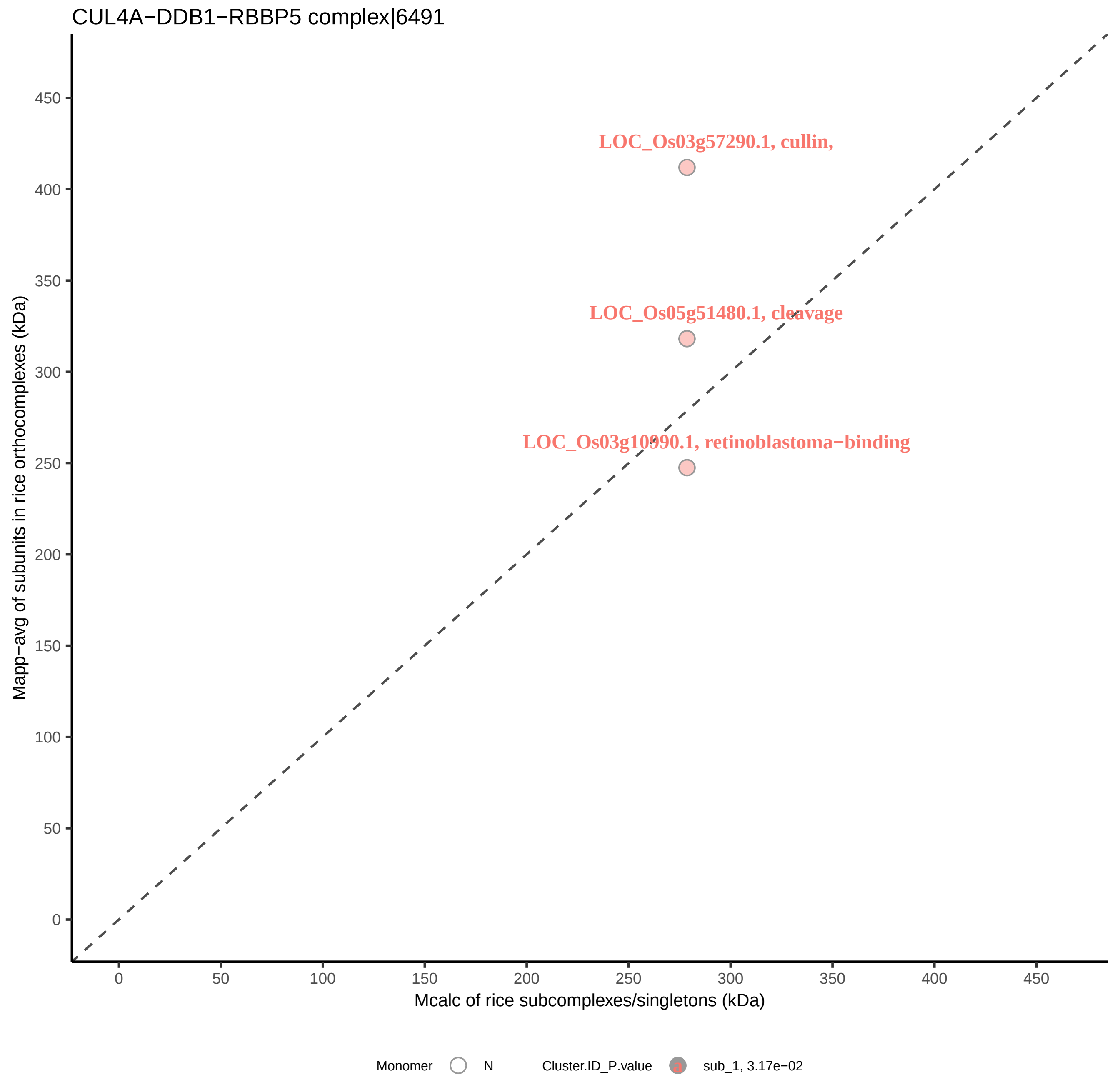
CUL4A−DDB1−RBBP5 complex|6491
LOC_Os03g57290.1, cullin,
LOC_Os05g51480.1, cleavage
LOC_Os03g10990.1, retinoblastoma−binding
Mapp−avg of subunits in rice orthocomplexes (kDa)
0
50
100
150
200
250
300
350
400
450
Mcalc of rice subcomplexes/singletons (kDa)
Monomer
N
Cluster.ID_P.value
sub_1, 3.17e−02

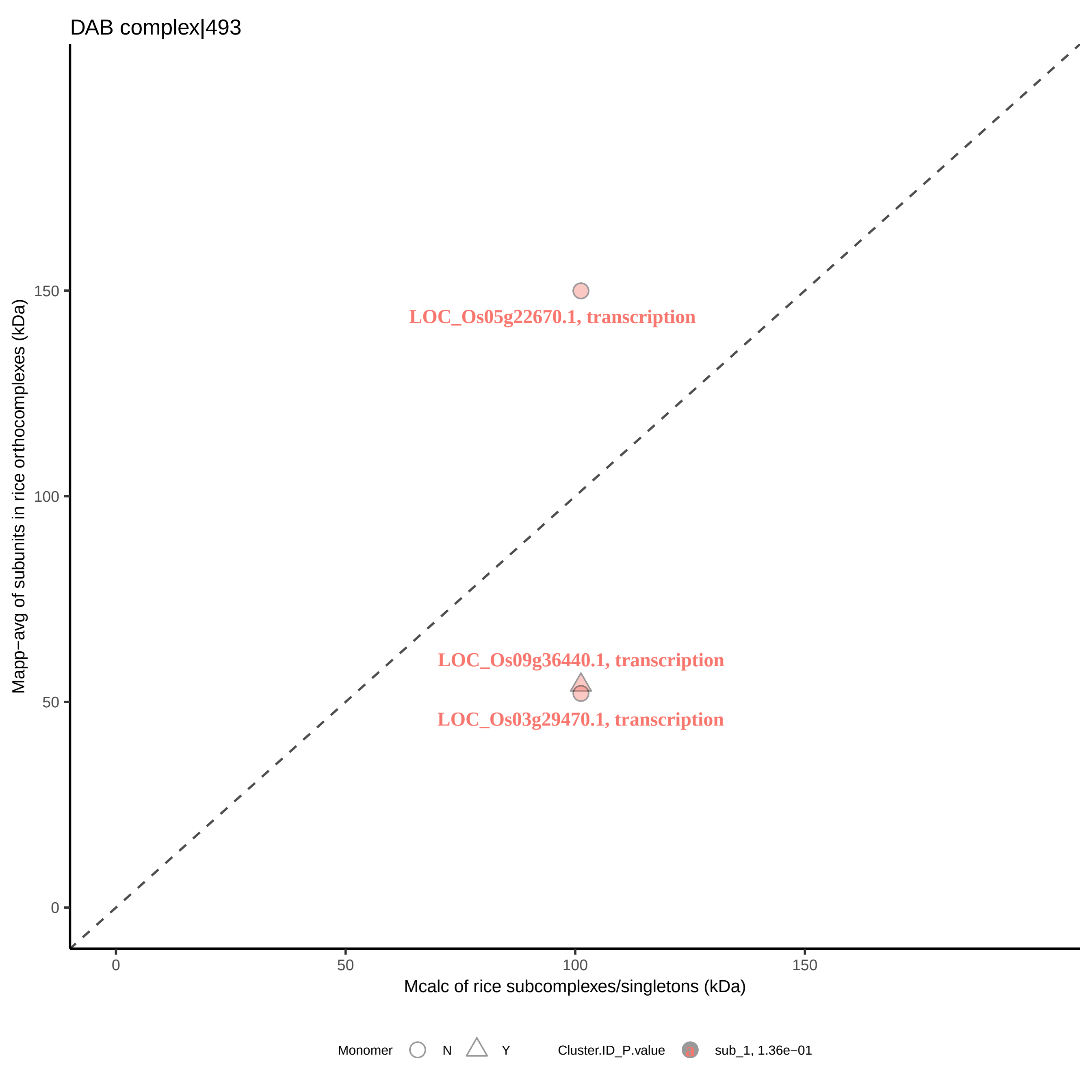
DAB complex|493
LOC_Os05g22670.1, transcription
LOC_Os09g36440.1, transcription
LOC_Os03g29470.1, transcription
Mapp−avg of subunits in rice orthocomplexes (kDa)
0
50
100
150
Mcalc of rice subcomplexes/singletons (kDa)
Monomer
N
Y
Cluster.ID_P.value
a
sub_1, 1.36e−01

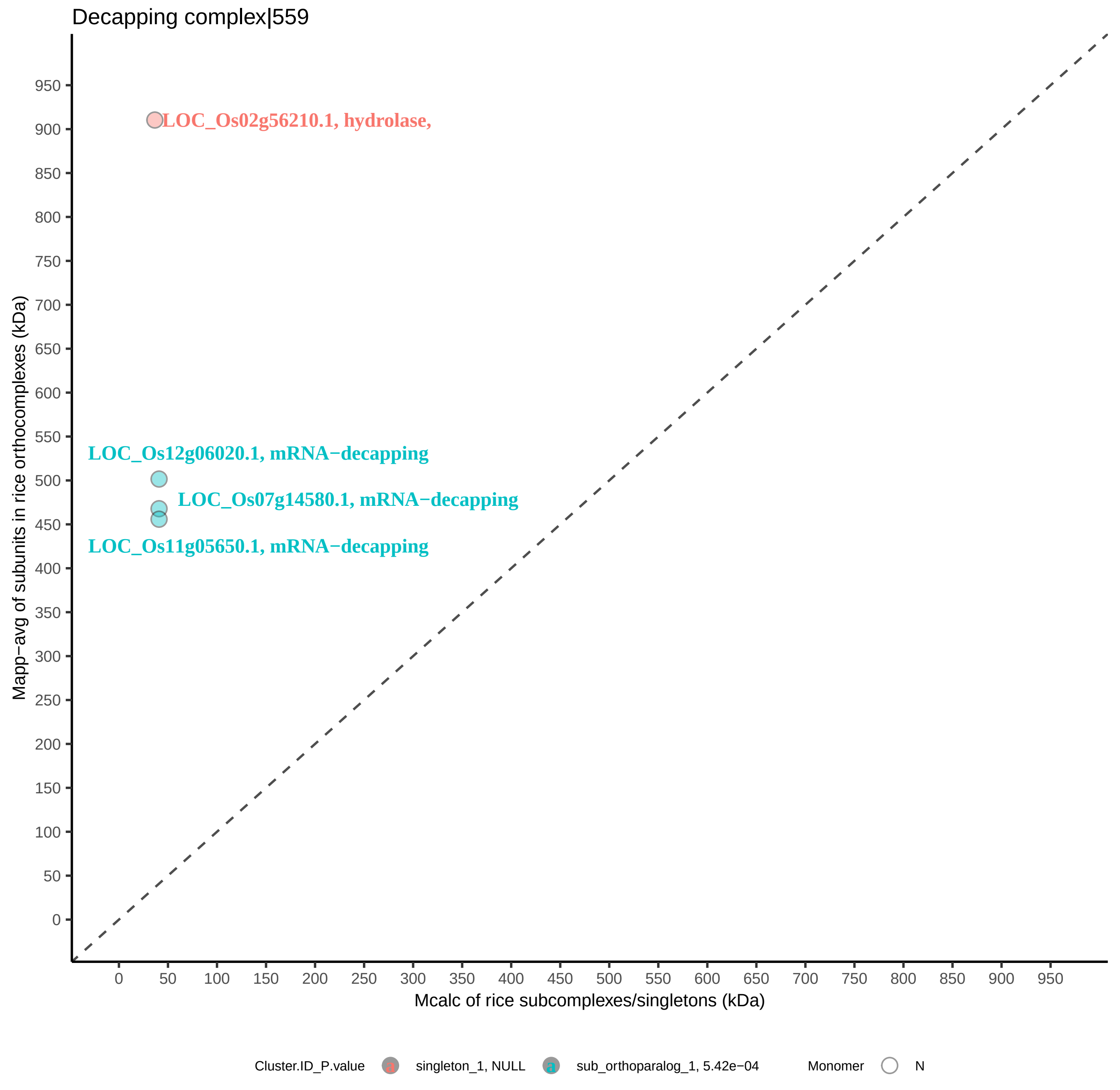
Decapping complex|559
LOC_Os02g56210.1, hydrolase,
LOC_Os12g06020.1, mRNA−decapping
LOC_Os07g14580.1, mRNA−decapping
LOC_Os11g05650.1, mRNA−decapping
Mapp−avg of subunits in rice orthocomplexes (kDa)
Mcalc of rice subcomplexes/singletons (kDa)
Cluster.ID_P.value
singleton_1, NULL
sub_orthoparalog_1, 5.42e−04
Monomer
N

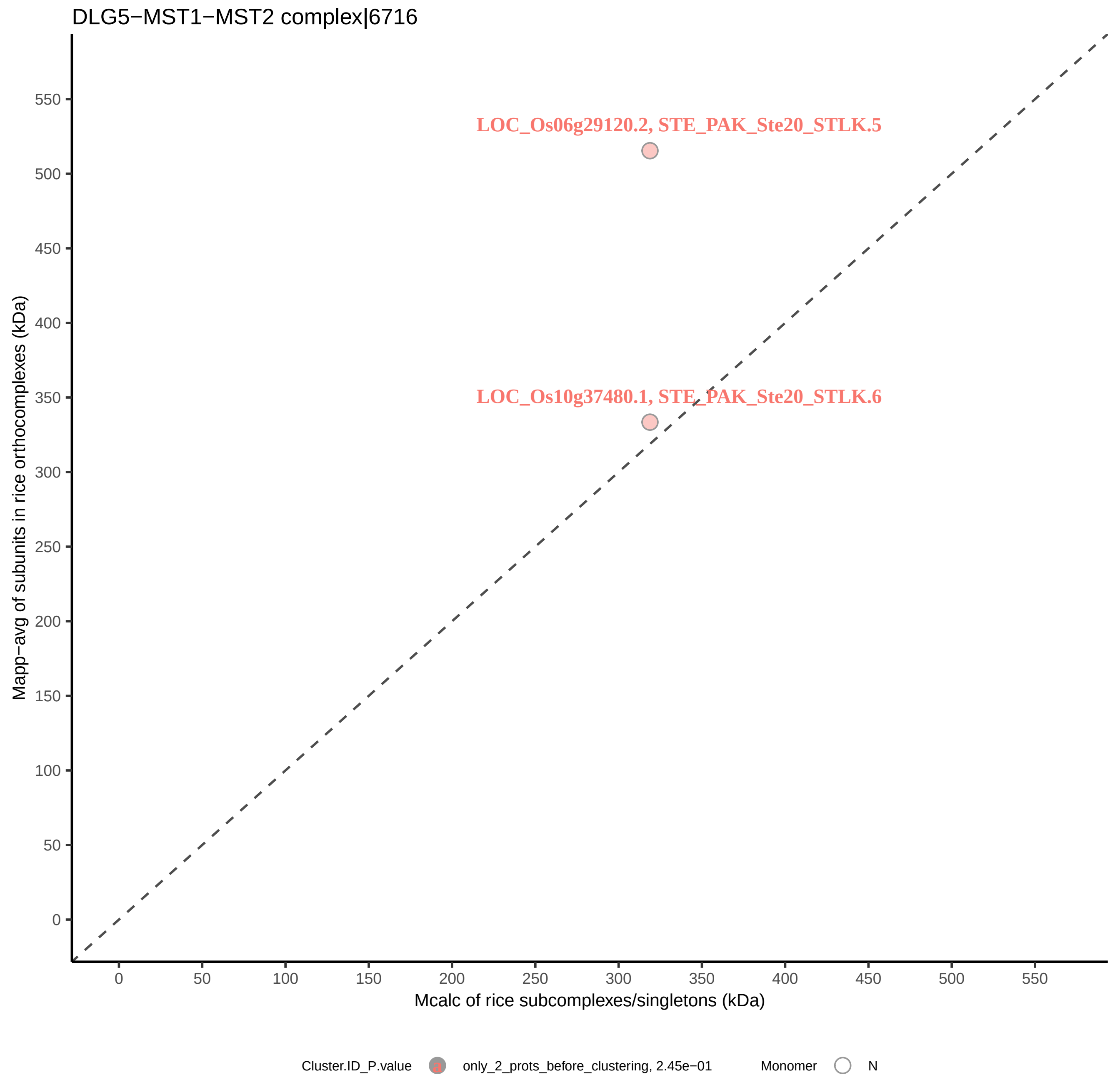
DLG5−MST1−MST2 complex|6716
LOC_Os06g29120.2, STE_PAK_Ste20_STLK.5
LOC_Os10g37480.1, STE_PAK_Ste20_STLK.6
Mapp−avg of subunits in rice orthocomplexes (kDa)
Mcalc of rice subcomplexes/singletons (kDa)
0
50
100
150
200
250
300
350
400
450
500
550
Cluster.ID_P.value
only_2_prots_before_clustering, 2.45e−01
Monomer
N

DNA synthesome complex (17 subunits)|1111

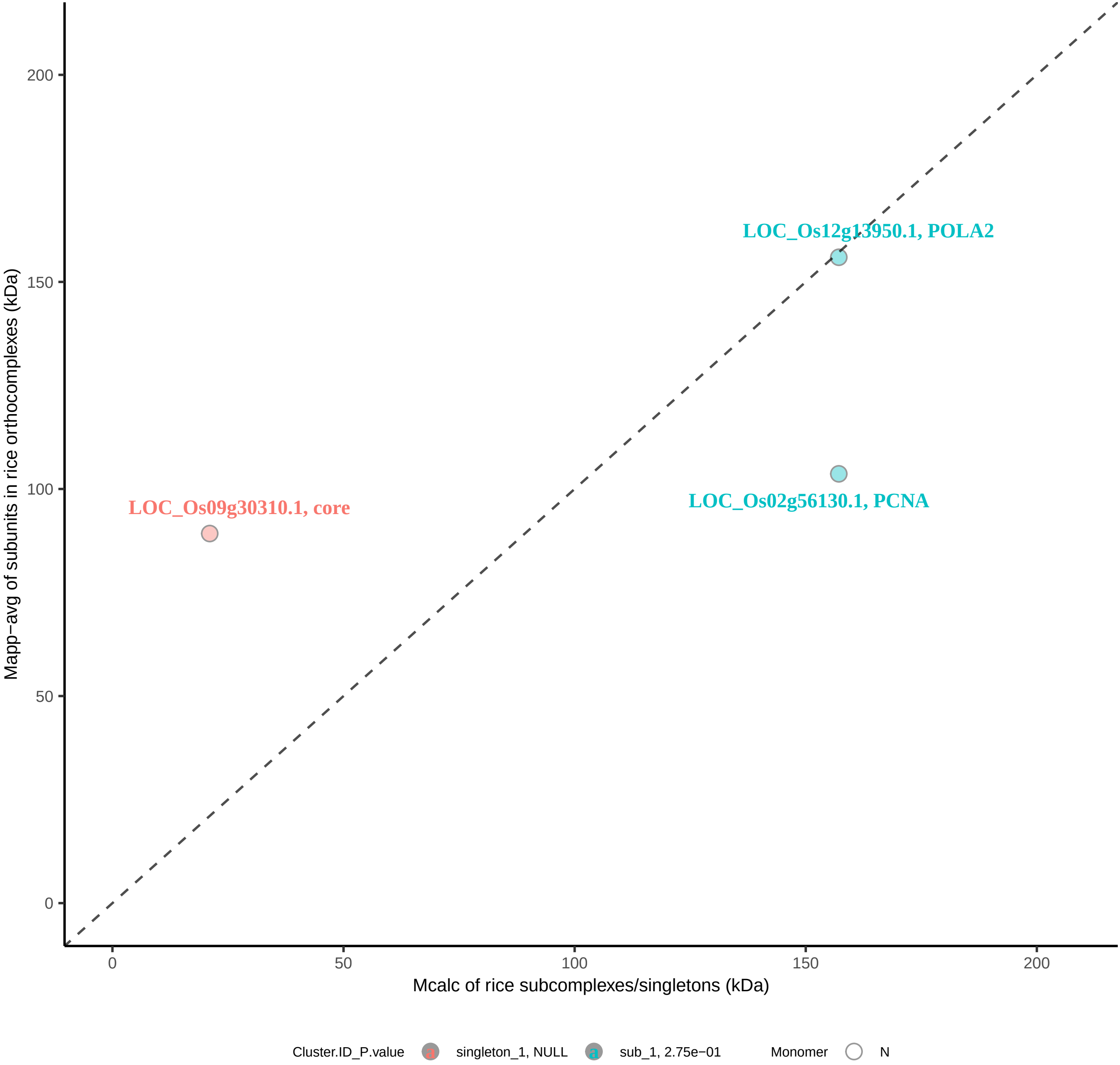

DNJC3−DNAJB1−HSPA8 complex|6396

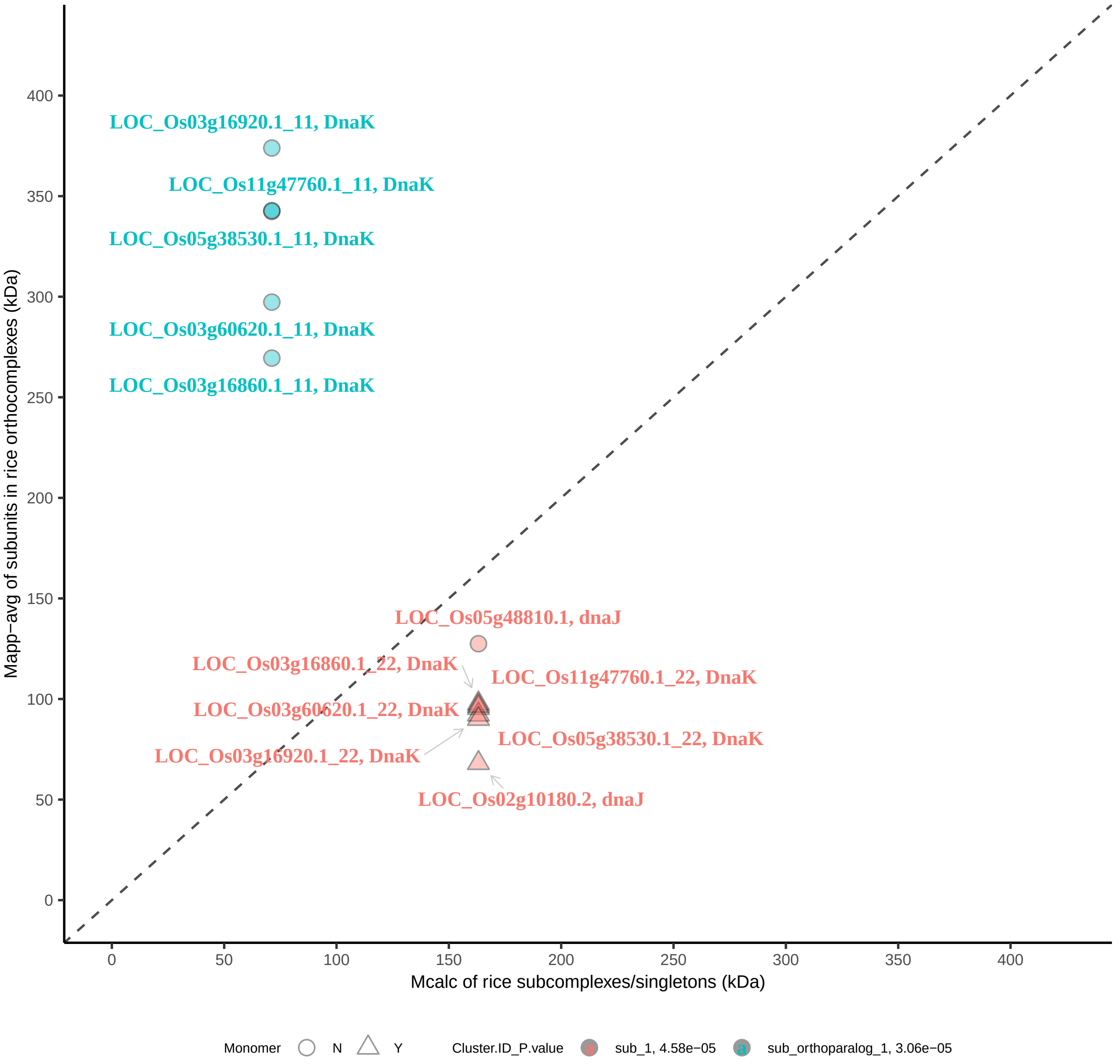

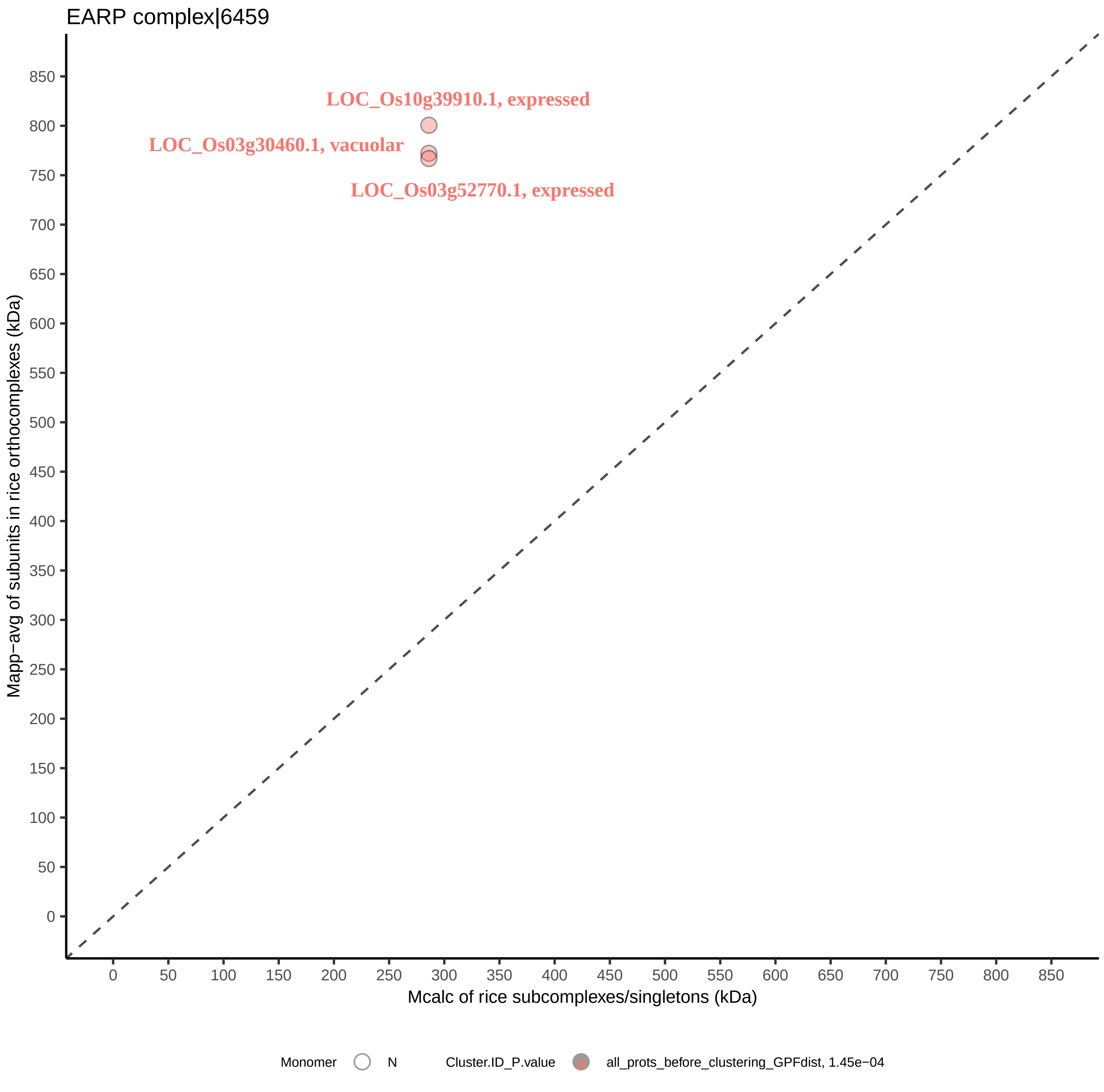

EARP complex|6459
LOC_Os10g39910.1, expressed
LOC_Os03g30460.1, vacuolar
LOC_Os03g52770.1, expressed
Mapp−avg of subunits in rice orthocomplexes (kDa)
Mcalc of rice subcomplexes/singletons (kDa)
0
50
100
150
200
250
300
350
400
450
500
550
600
650
700
750
800
850
Monomer
N
Cluster.ID_P.value
all_prots_before_clustering_GPFdist, 1.45e−04

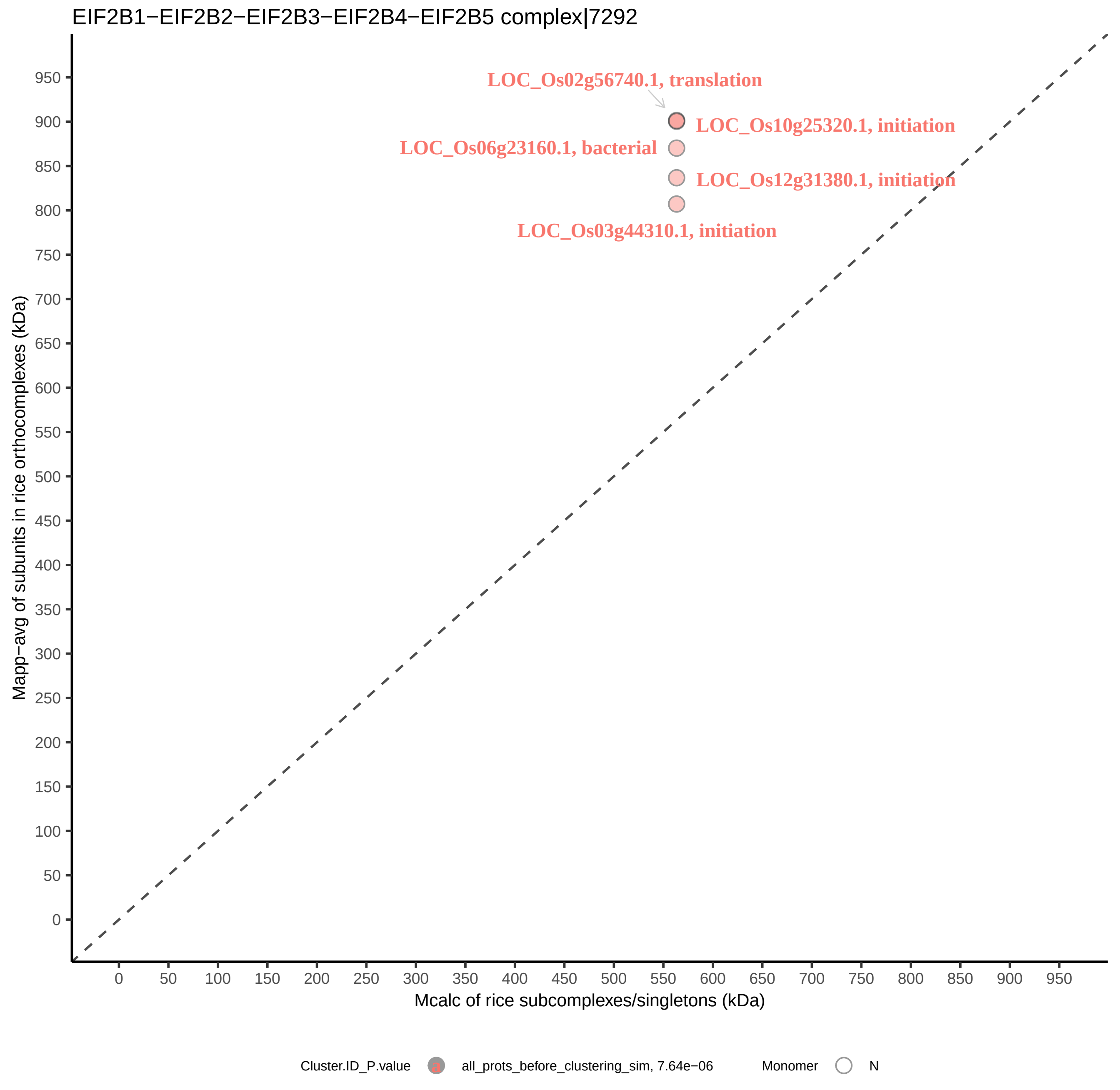

EIF2B1−EIF2B2−EIF2B3−EIF2B4−EIF2B5 complex|7292
LOC_Os02g56740.1, translation
LOC_Os10g25320.1, initiation
LOC_Os06g23160.1, bacterial
LOC_Os12g31380.1, initiation
LOC_Os03g44310.1, initiation
Mapp−avg of subunits in rice orthocomplexes (kDa)
Mcalc of rice subcomplexes/singletons (kDa)
0
50
100
150
200
250
300
350
400
450
500
550
600
650
700
750
800
850
900
950
Cluster.ID_P.value
a
all_prots_before_clustering_sim, 7.64e−06
Monomer
N

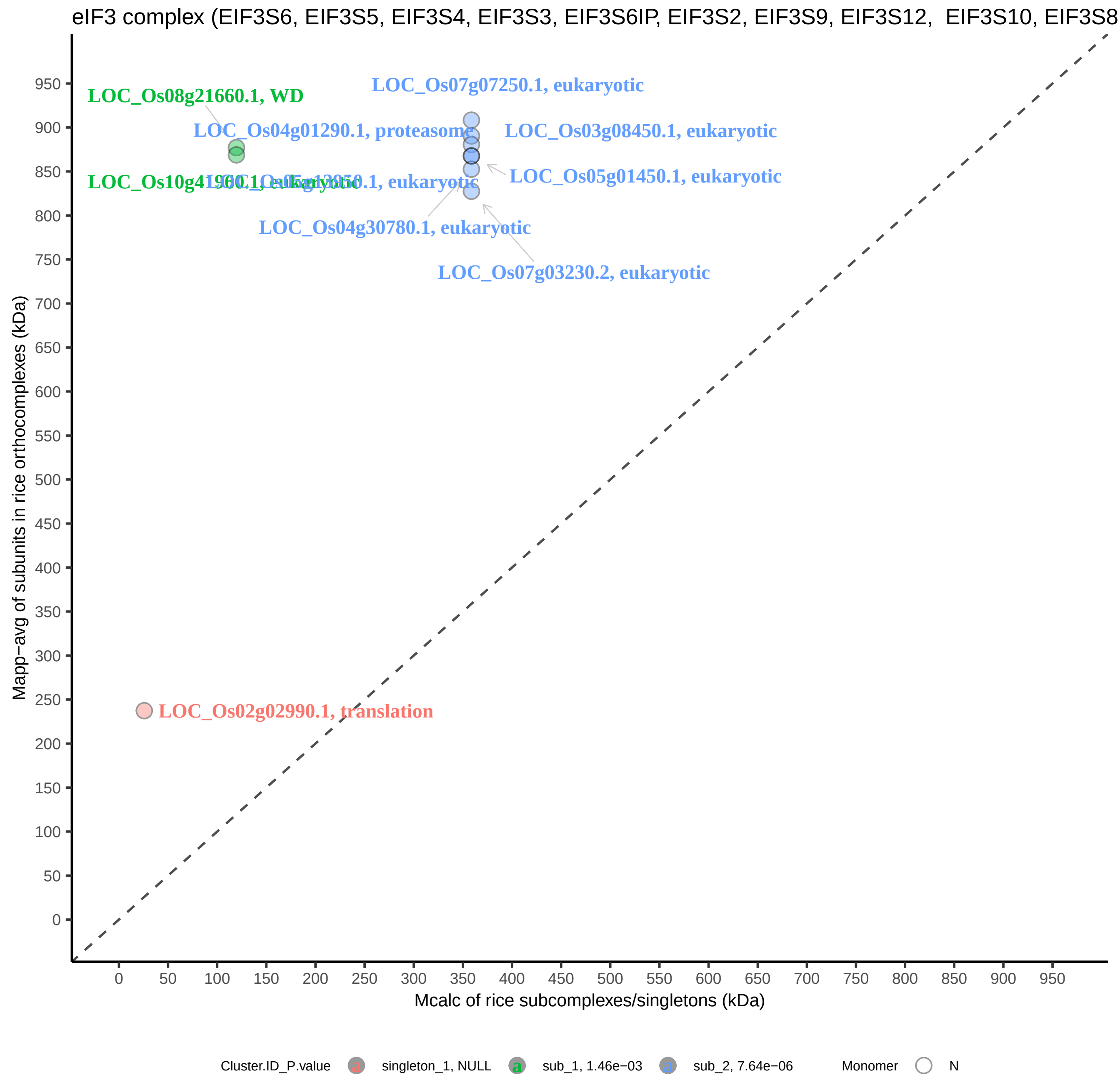

eIF3 complex (EIF3S6, EIF3S5, EIF3S4, EIF3S3, EIF3S6IP, EIF3S2, EIF3S9, EIF3S12, EIF3S10, EIF3S8
LOC_Os08g21660.1, WD
LOC_Os07g07250.1, eukaryotic
LOC_Os04g01290.1, proteasome
LOC_Os03g08450.1, eukaryotic
LOC_Os05g01450.1, eukaryotic
LOC_Os10g41960.1, eukaryotic
LOC_Os05g13950.1, eukaryotic
LOC_Os04g30780.1, eukaryotic
LOC_Os07g03230.2, eukaryotic
LOC_Os02g02990.1, translation
Mapp−avg of subunits in rice orthocomplexes (kDa)
Mcalc of rice subcomplexes/singletons (kDa)
0
50
100
150
200
250
300
350
400
450
500
550
600
650
700
750
800
850
900
950
Cluster.ID_P.value
singleton_1, NULL
sub_1, 1.46e−03
sub_2, 7.64e−06
Monomer
N

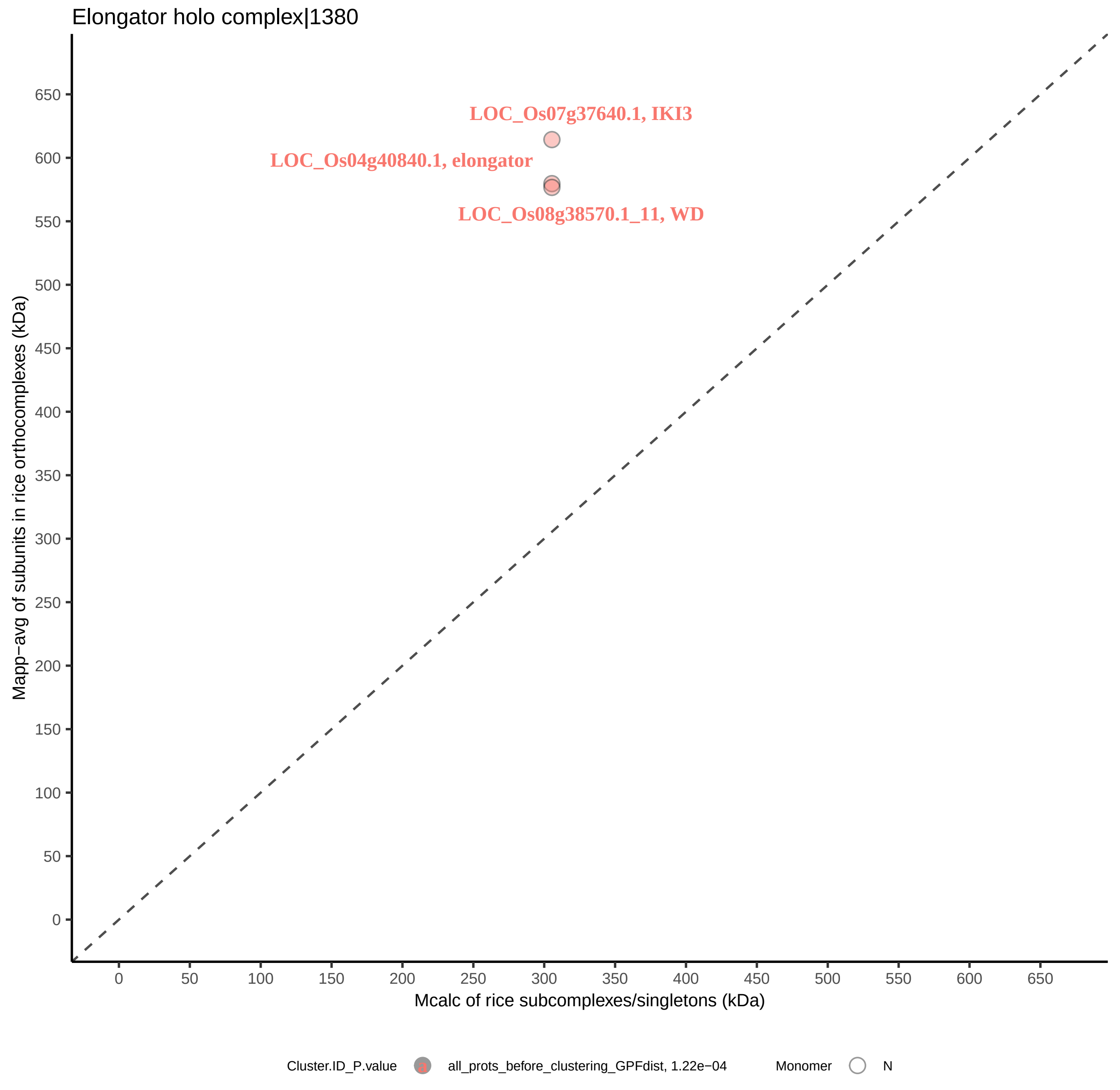
Elongator holo complex|1380
LOC_Os07g37640.1, IKI3
LOC_Os04g40840.1, elongator
LOC_Os08g38570.1_11, WD
Mapp−avg of subunits in rice orthocomplexes (kDa)
Mcalc of rice subcomplexes/singletons (kDa)
0
50
100
150
200
250
300
350
400
450
500
550
600
650
Cluster.ID_P.value
all_prots_before_clustering_GPFdist, 1.22e−04
Monomer
N

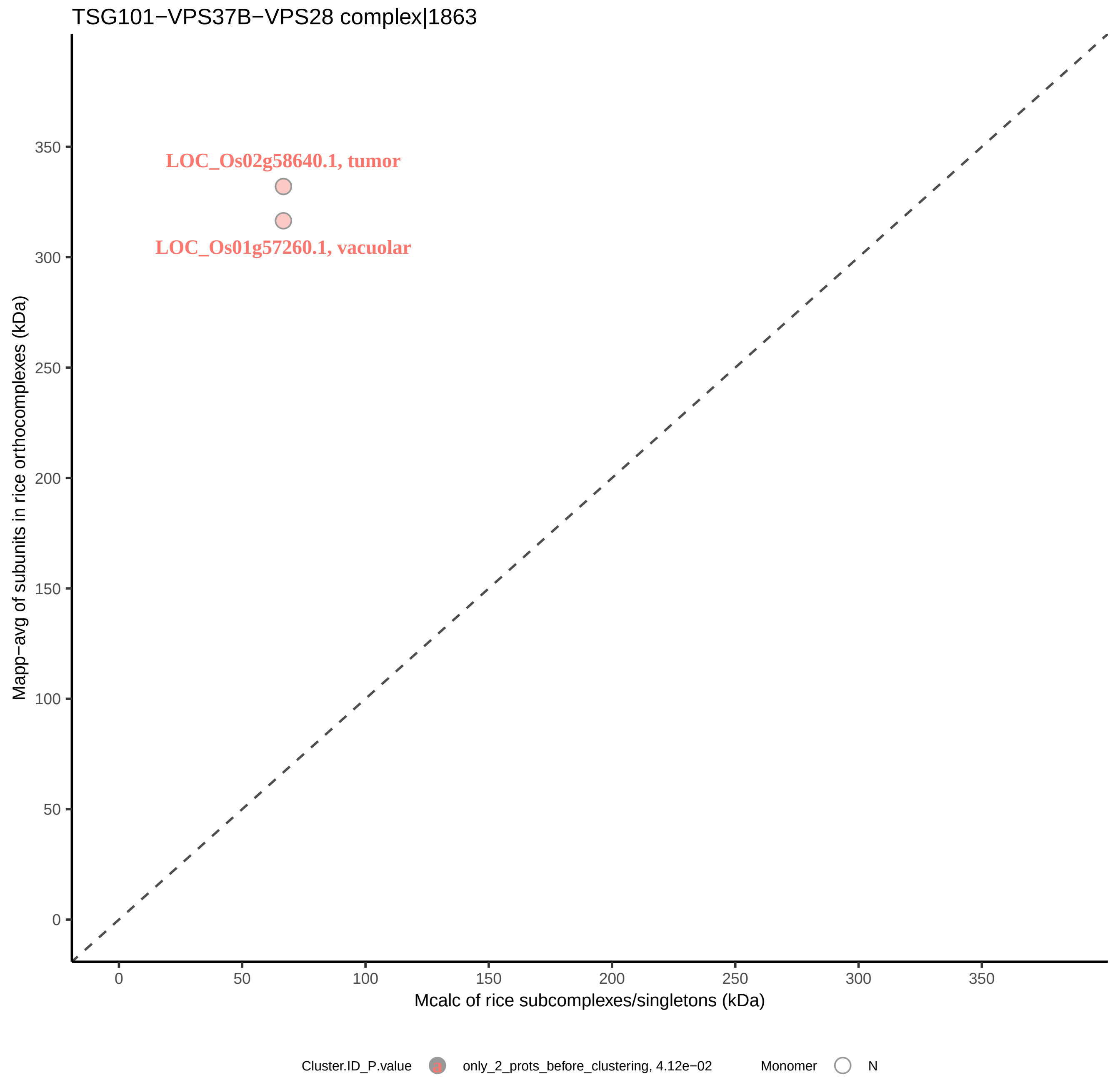
TSG101−VPS37B−VPS28 complex|1863
LOC_Os02g58640.1, tumor
LOC_Os01g57260.1, vacuolar
Mapp−avg of subunits in rice orthocomplexes (kDa)
0
50
100
150
200
250
300
350
Mcalc of rice subcomplexes/singletons (kDa)
0
50
100
150
200
250
300
350
Cluster.ID_P.value
only_2_prots_before_clustering, 4.12e−02
Monomer
N

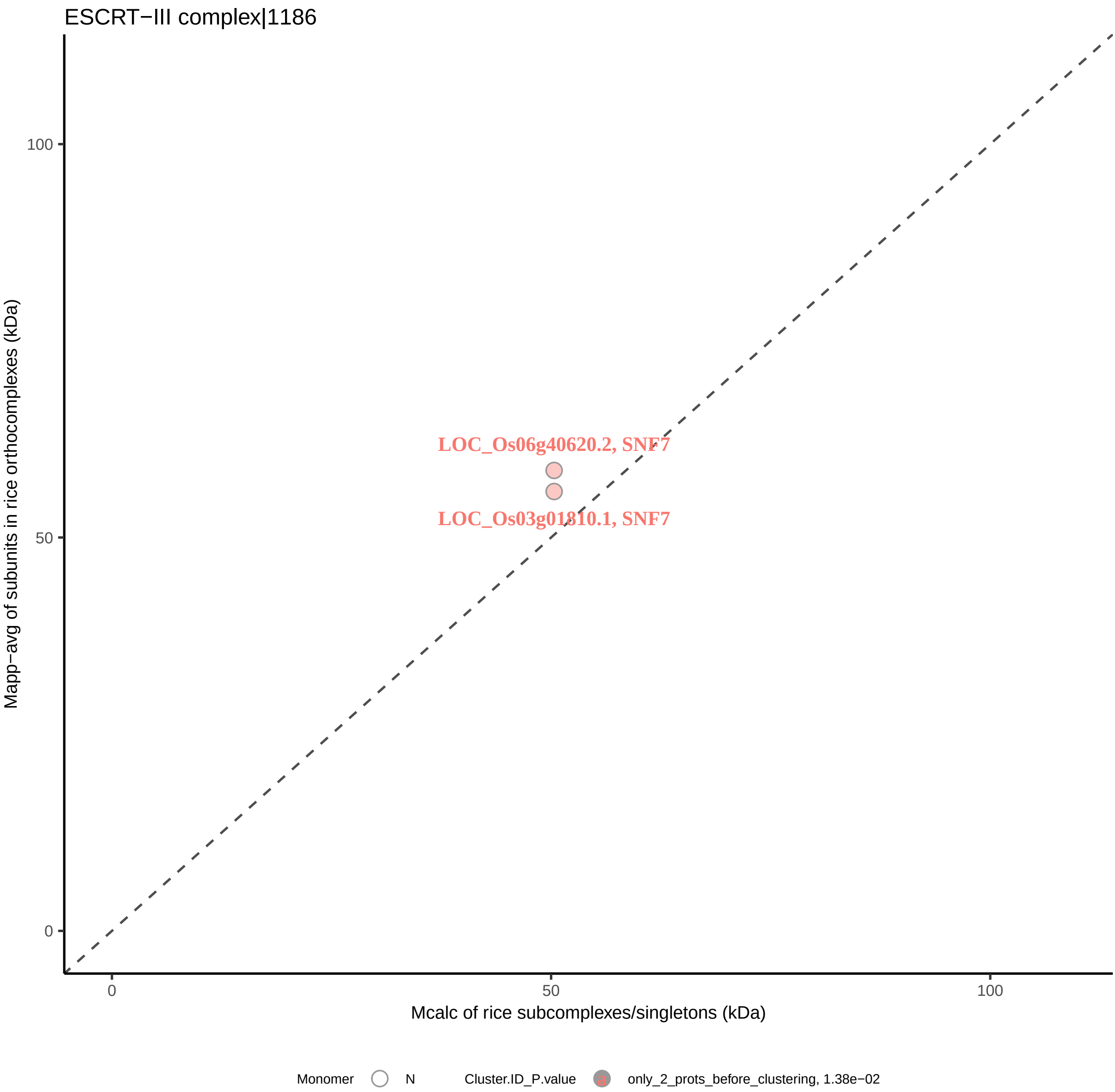
ESCRT−III complex|1186
Mapp−avg of subunits in rice orthocomplexes (kDa)
100
50
0
LOC_Os06g40620.2, SNF7
LOC_Os03g01810.1, SNF7
0
50
100
Mcalc of rice subcomplexes/singletons (kDa)
Monomer
N
Cluster.ID_P.value
only_2_prots_before_clustering, 1.38e−02

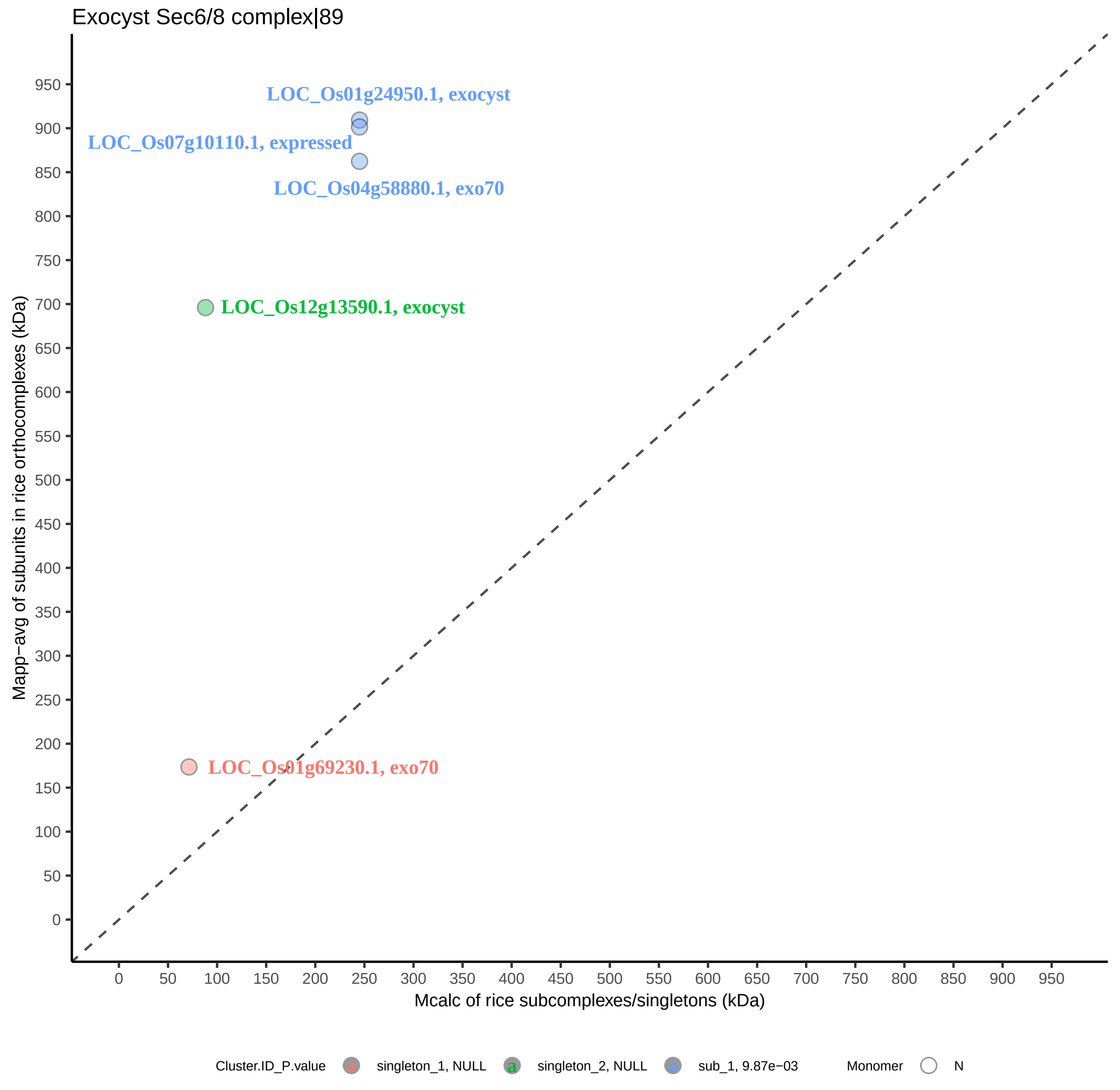

Exocyst Sec6/8 complex|89
LOC_Os01g24950.1, exocyst
LOC_Os07g10110.1, expressed
LOC_Os04g58880.1, exo70
LOC_Os12g13590.1, exocyst
LOC_Os01g69230.1, exo70
Mapp−avg of subunits in rice orthocomplexes (kDa)
Mcalc of rice subcomplexes/singletons (kDa)
0
50
100
150
200
250
300
350
400
450
500
550
600
650
700
750
800
850
900
950
Cluster.ID_P.value
singleton_1, NULL
singleton_2, NULL
sub_1, 9.87e−03
Monomer
N

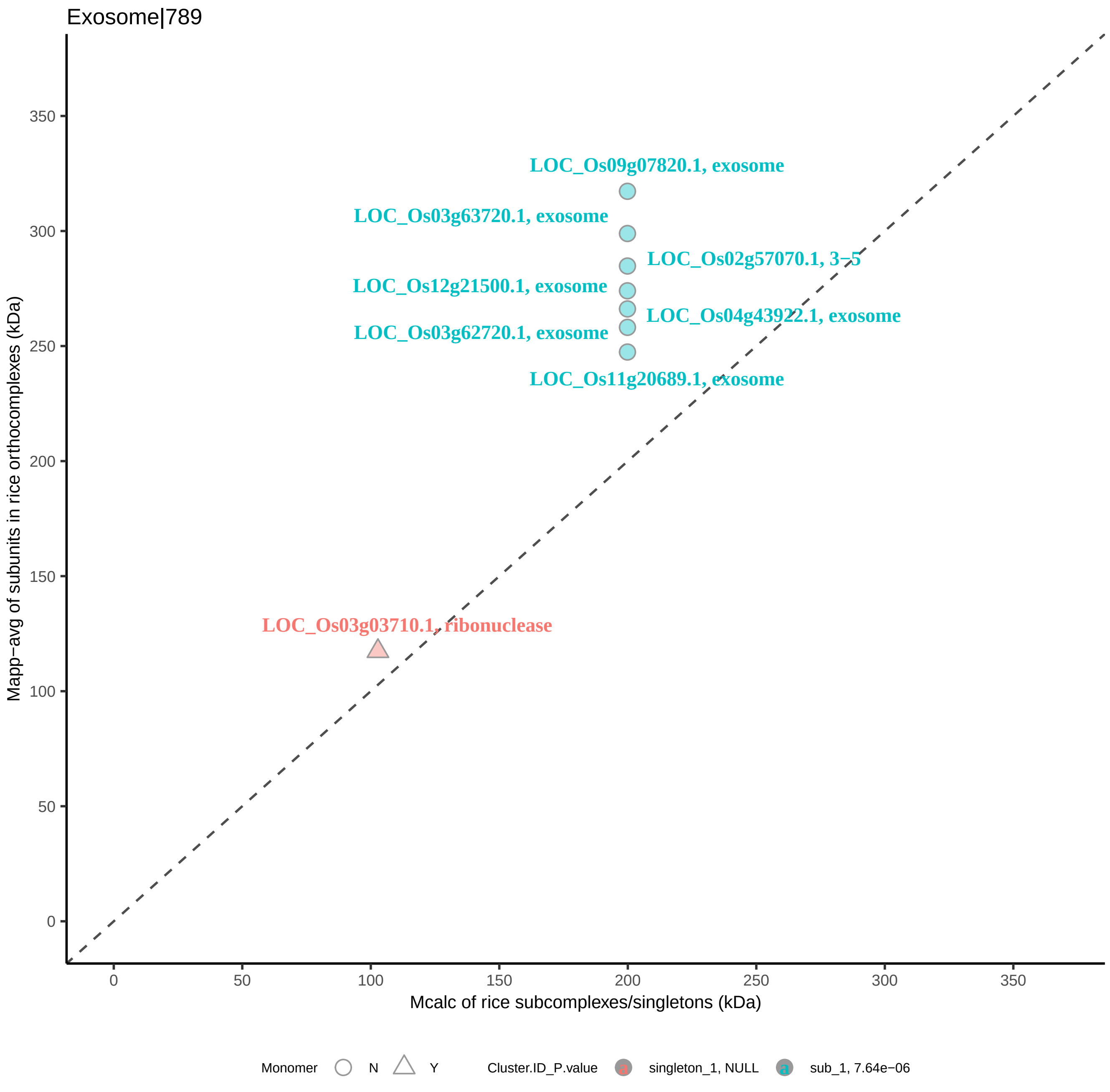

Exosome|789
LOC_Os09g07820.1, exosome
LOC_Os03g63720.1, exosome
LOC_Os02g57070.1, 3−5
LOC_Os12g21500.1, exosome
LOC_Os04g43922.1, exosome
LOC_Os03g62720.1, exosome
LOC_Os11g20689.1, exosome
LOC_Os03g03710.1, ribonuclease
Mapp−avg of subunits in rice orthocomplexes (kDa)
Mcalc of rice subcomplexes/singletons (kDa)
0
50
100
150
200
250
300
350
Monomer
N
Y
Cluster.ID_P.value
singleton_1, NULL
sub_1, 7.64e−06

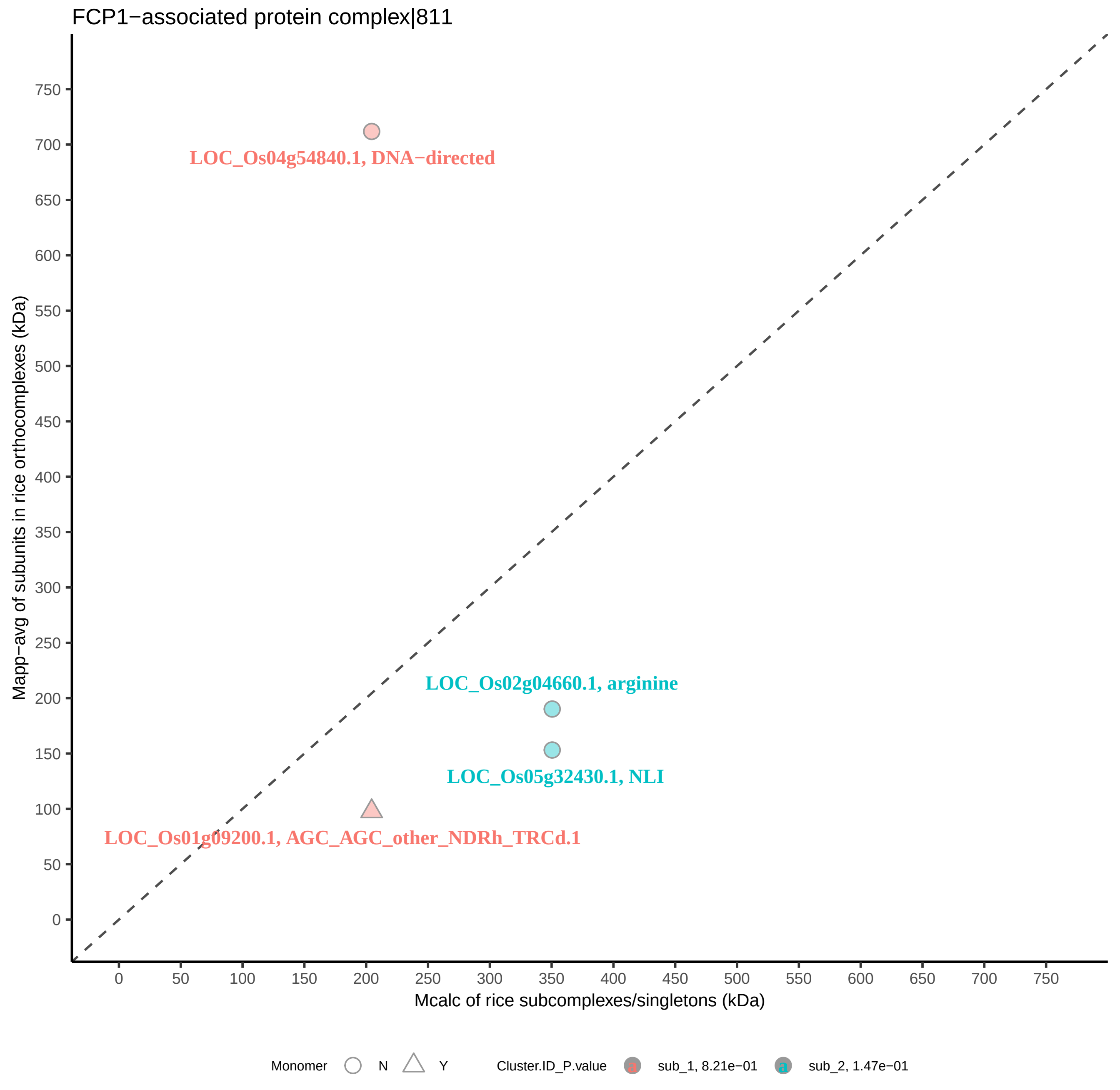
FCP1−associated protein complex|811
LOC_Os04g54840.1, DNA−directed
LOC_Os02g04660.1, arginine
LOC_Os05g32430.1, NLI
LOC_Os01g09200.1, AGC_AGC_other_NDRh_TRCd.1
Mapp−avg of subunits in rice orthocomplexes (kDa)
Mcalc of rice subcomplexes/singletons (kDa)
0
50
100
150
200
250
300
350
400
450
500
550
600
650
700
750
Monomer
N
Y
Cluster.ID_P.value
sub_1, 8.21e−01
sub_2, 1.47e−01

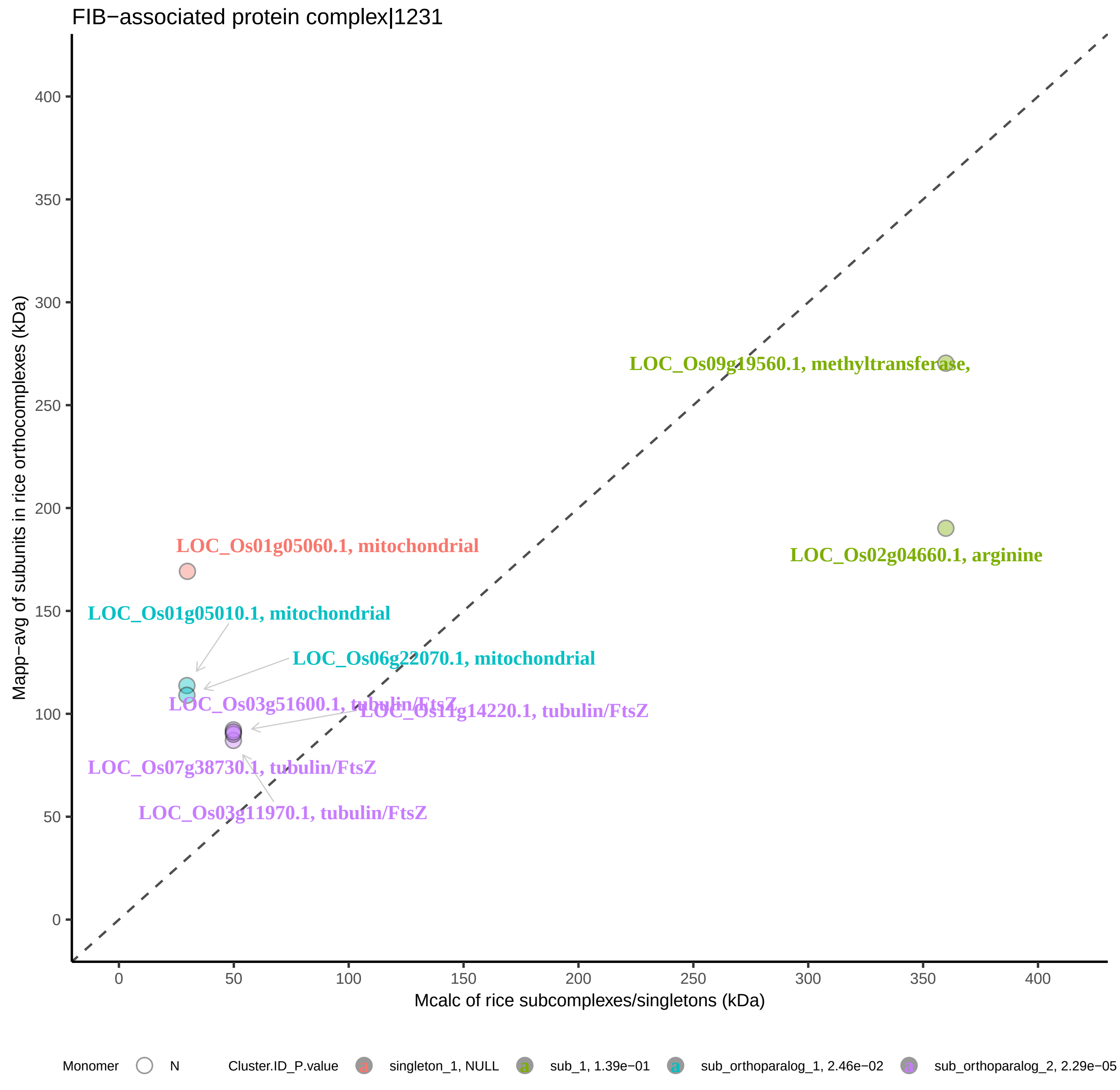
FIB−associated protein complex|1231
LOC_Os09g19560.1, methyltransferase,
LOC_Os02g04660.1, arginine
LOC_Os01g05060.1, mitochondrial
LOC_Os01g05010.1, mitochondrial
LOC_Os06g22070.1, mitochondrial
LOC_Os03g51600.1, tubulin/FtsZ
LOC_Os11g14220.1, tubulin/FtsZ
LOC_Os07g38730.1, tubulin/FtsZ
LOC_Os03g11970.1, tubulin/FtsZ
Mapp−avg of subunits in rice orthocomplexes (kDa)
Mcalc of rice subcomplexes/singletons (kDa)
0
50
100
150
200
250
300
350
400
Monomer
N
Cluster.ID_P.value
singleton_1, NULL
sub_1, 1.39e−01
sub_orthoparalog_1, 2.46e−02
sub_orthoparalog_2, 2.29e−05

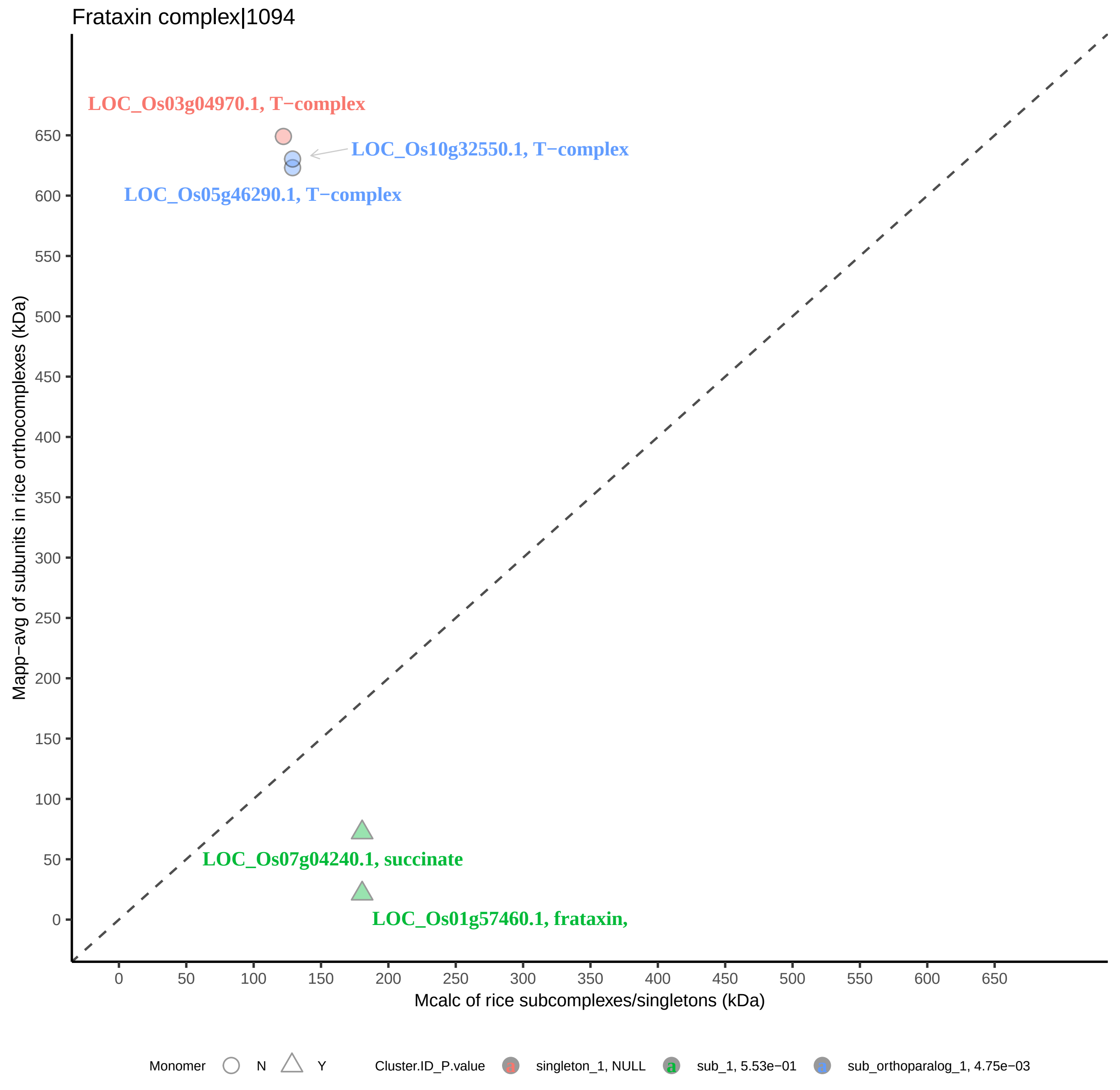

Frataxin complex|1094
LOC_Os03g04970.1, T−complex
LOC_Os10g32550.1, T−complex
LOC_Os05g46290.1, T−complex
LOC_Os07g04240.1, succinate
LOC_Os01g57460.1, frataxin,
Mapp−avg of subunits in rice orthocomplexes (kDa)
0
50
100
150
200
250
300
350
400
450
500
550
600
650
Mcalc of rice subcomplexes/singletons (kDa)
0
50
100
150
200
250
300
350
400
450
500
550
600
650
Monomer
N
Y
Cluster.ID_P.value
singleton_1, NULL
sub_1, 5.53e−01
sub_orthoparalog_1, 4.75e−03

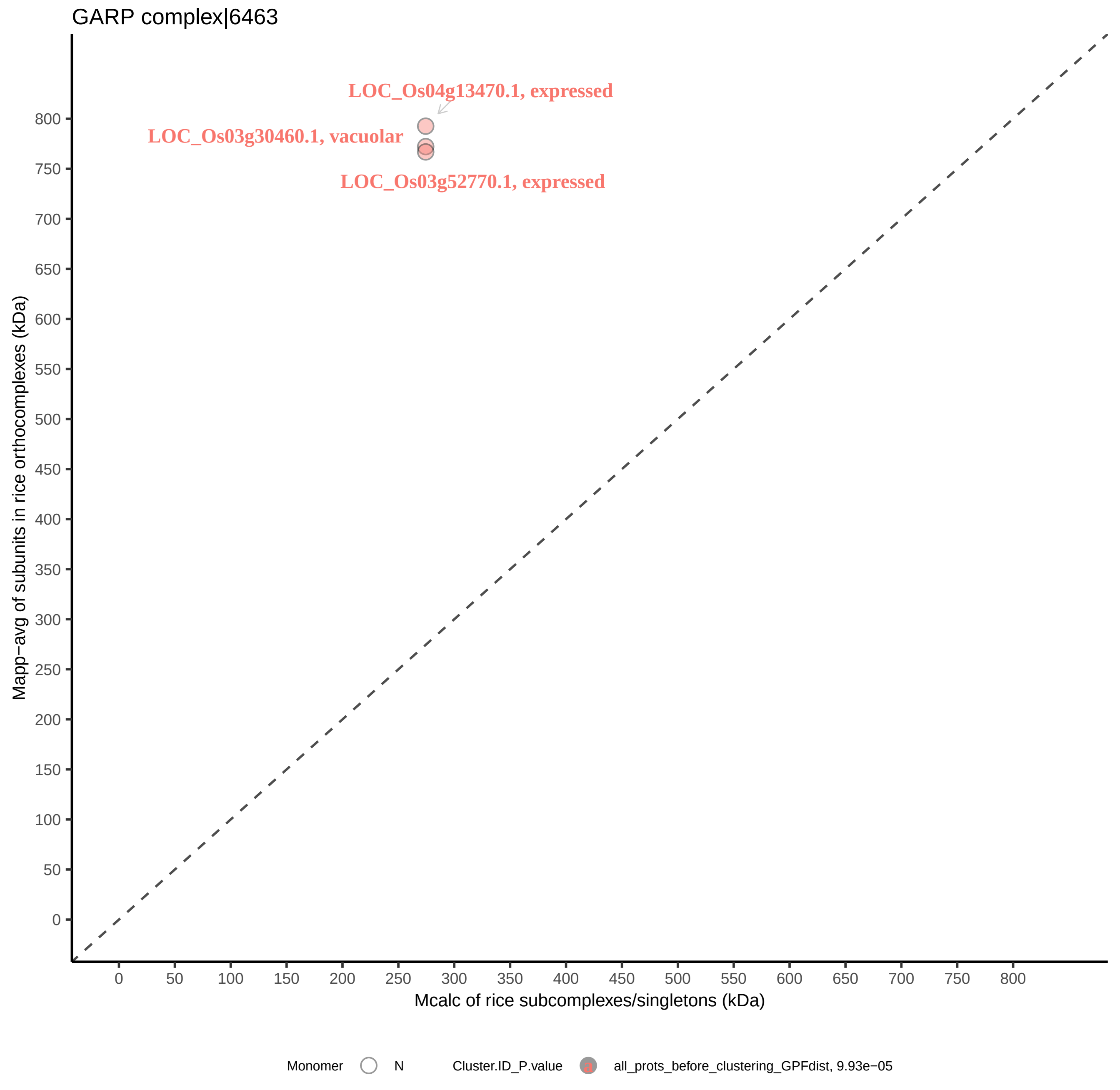

GARP complex|6463
LOC_Os04g13470.1, expressed
LOC_Os03g30460.1, vacuolar
LOC_Os03g52770.1, expressed
Mapp−avg of subunits in rice orthocomplexes (kDa)
800
750
700
650
600
550
500
450
400
350
300
250
200
150
100
50
0
0
50
100
150
200
250
300
350
400
450
500
550
600
650
700
750
800
Mcalc of rice subcomplexes/singletons (kDa)
Monomer
N
Cluster.ID_P.value
all_prots_before_clustering_GPFdist, 9.93e−05

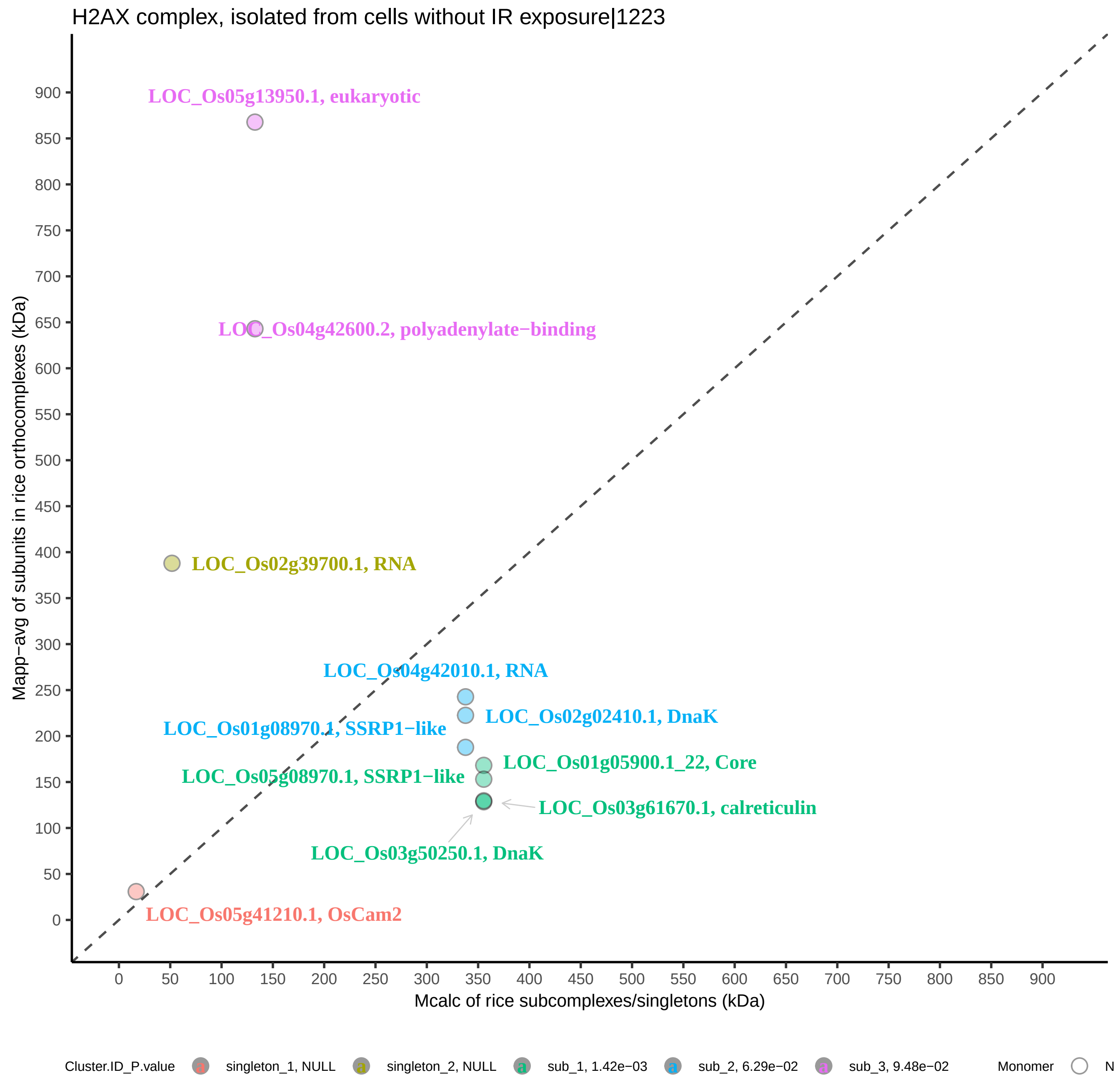
H2AX complex, isolated from cells without IR exposure|1223
LOC_Os05g13950.1, eukaryotic
LOC_Os04g42600.2, polyadenylate−binding
LOC_Os02g39700.1, RNA
LOC_Os04g42010.1, RNA
LOC_Os02g02410.1, DnaK
LOC_Os01g08970.1, SSRP1−like
LOC_Os01g05900.1_22, Core
LOC_Os05g08970.1, SSRP1−like
LOC_Os03g61670.1, calreticulin
LOC_Os03g50250.1, DnaK
LOC_Os05g41210.1, OsCam2
Mapp−avg of subunits in rice orthocomplexes (kDa)
Mcalc of rice subcomplexes/singletons (kDa)
0
50
100
150
200
250
300
350
400
450
500
550
600
650
700
750
800
850
900
Cluster.ID_P.value
singleton_1, NULL
singleton_2, NULL
sub_1, 1.42e−03
sub_2, 6.29e−02
sub_3, 9.48e−02
Monomer

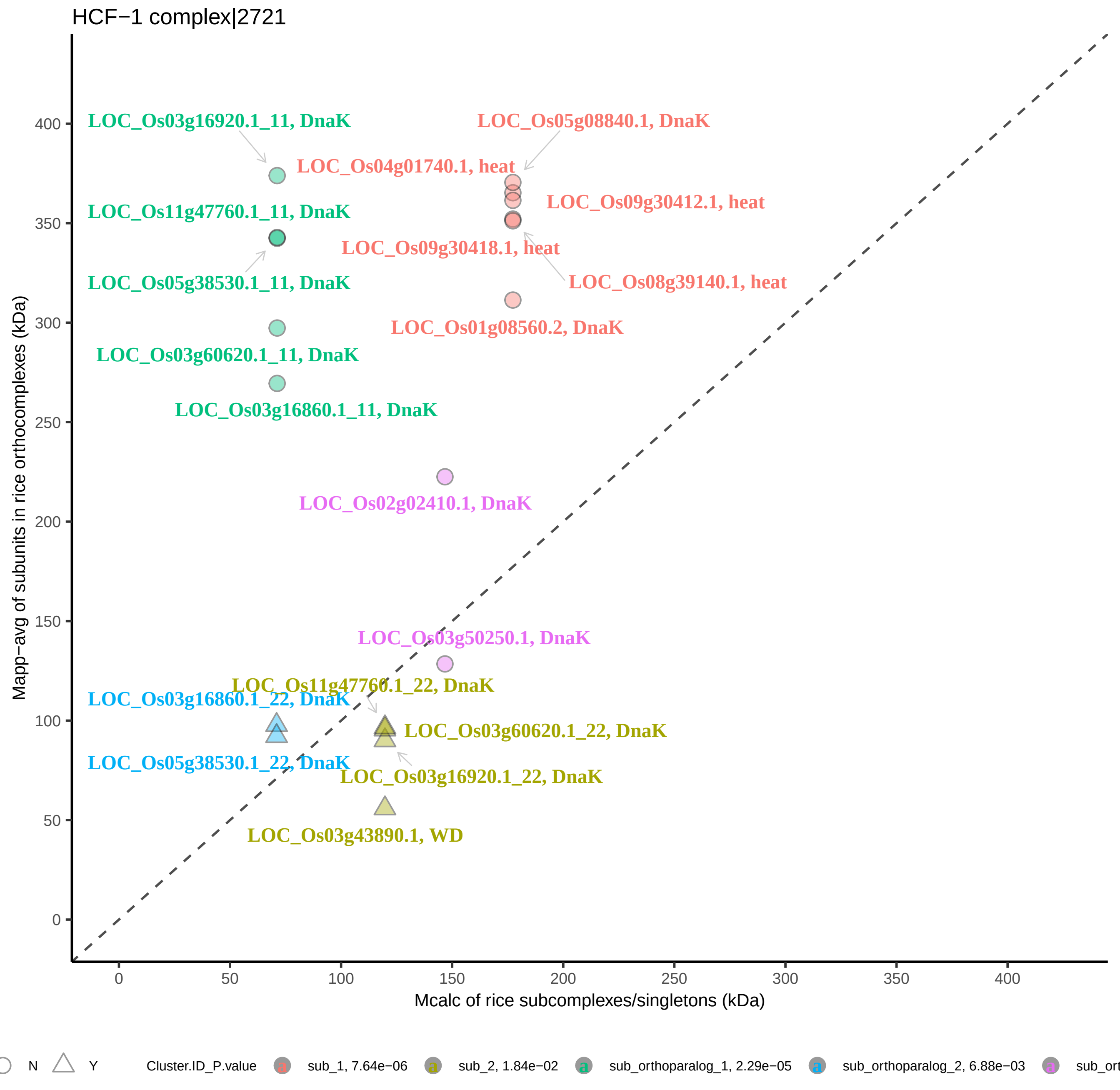

HCF−1 complex|2721
LOC_Os03g16920.1_11, DnaK
LOC_Os05g08840.1, DnaK
LOC_Os04g01740.1, heat
LOC_Os09g30412.1, heat
LOC_Os11g47760.1_11, DnaK
LOC_Os09g30418.1, heat
LOC_Os05g38530.1_11, DnaK
LOC_Os08g39140.1, heat
LOC_Os01g08560.2, DnaK
LOC_Os03g60620.1_11, DnaK
LOC_Os03g16860.1_11, DnaK
LOC_Os02g02410.1, DnaK
LOC_Os03g50250.1, DnaK
LOC_Os11g47760.1_22, DnaK
LOC_Os03g16860.1_22, DnaK
LOC_Os03g60620.1_22, DnaK
LOC_Os05g38530.1_22, DnaK
LOC_Os03g16920.1_22, DnaK
LOC_Os03g43890.1, WD
Mapp−avg of subunits in rice orthocomplexes (kDa)
Mcalc of rice subcomplexes/singletons (kDa)
0
50
100
150
200
250
300
350
400
N
Y
Cluster.ID_P.value
sub_1, 7.64e−06
sub_2, 1.84e−02
sub_orthoparalog_1, 2.29e−05
sub_orthoparalog_2, 6.88e−03

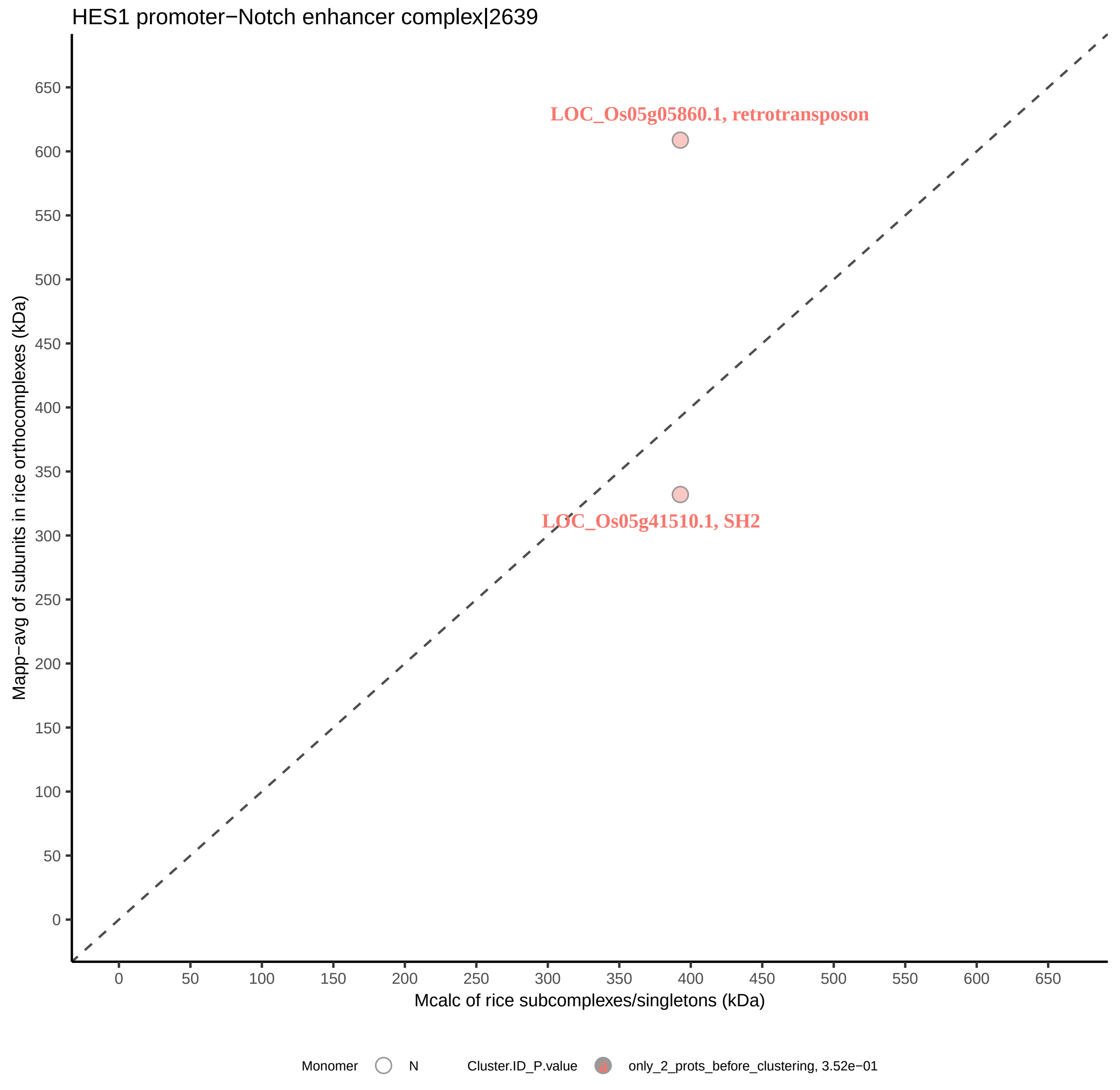

HES1 promoter−Notch enhancer complex|2639
LOC_Os05g05860.1, retrotransposon
LOC_Os05g41510.1, SH2
Mapp−avg of subunits in rice orthocomplexes (kDa)
Mcalc of rice subcomplexes/singletons (kDa)
0
50
100
150
200
250
300
350
400
450
500
550
600
650
Monomer
N
Cluster.ID_P.value
a
only_2_prots_before_clustering, 3.52e−01

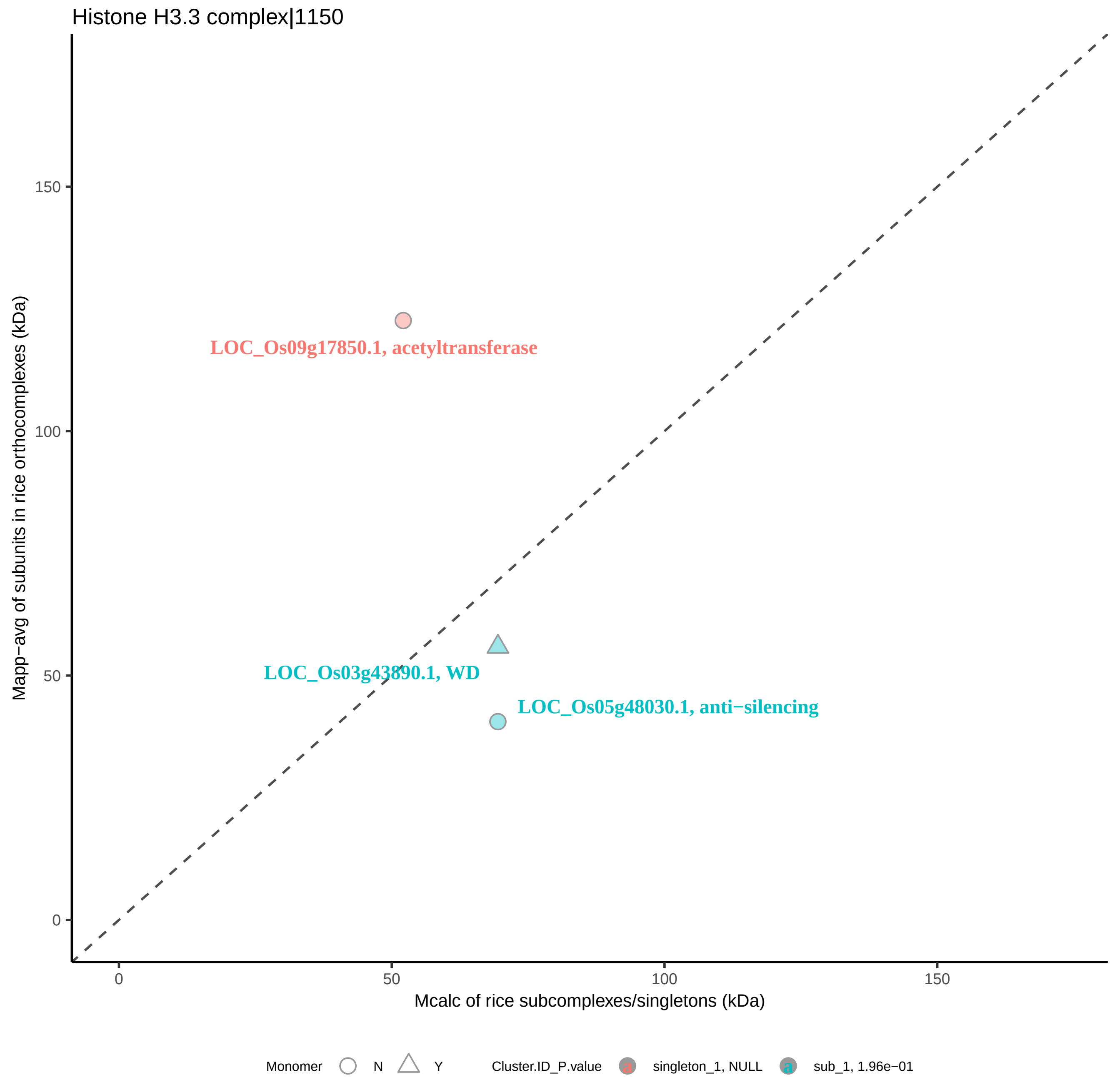

Histone H3.3 complex|1150
Mapp−avg of subunits in rice orthocomplexes (kDa)
Mcalc of rice subcomplexes/singletons (kDa)
0
50
100
150
LOC_Os09g17850.1, acetyltransferase
LOC_Os03g43890.1, WD
LOC_Os05g48030.1, anti−silencing
Monomer
N
Y
Cluster.ID_P.value
singleton_1, NULL
sub_1, 1.96e−01

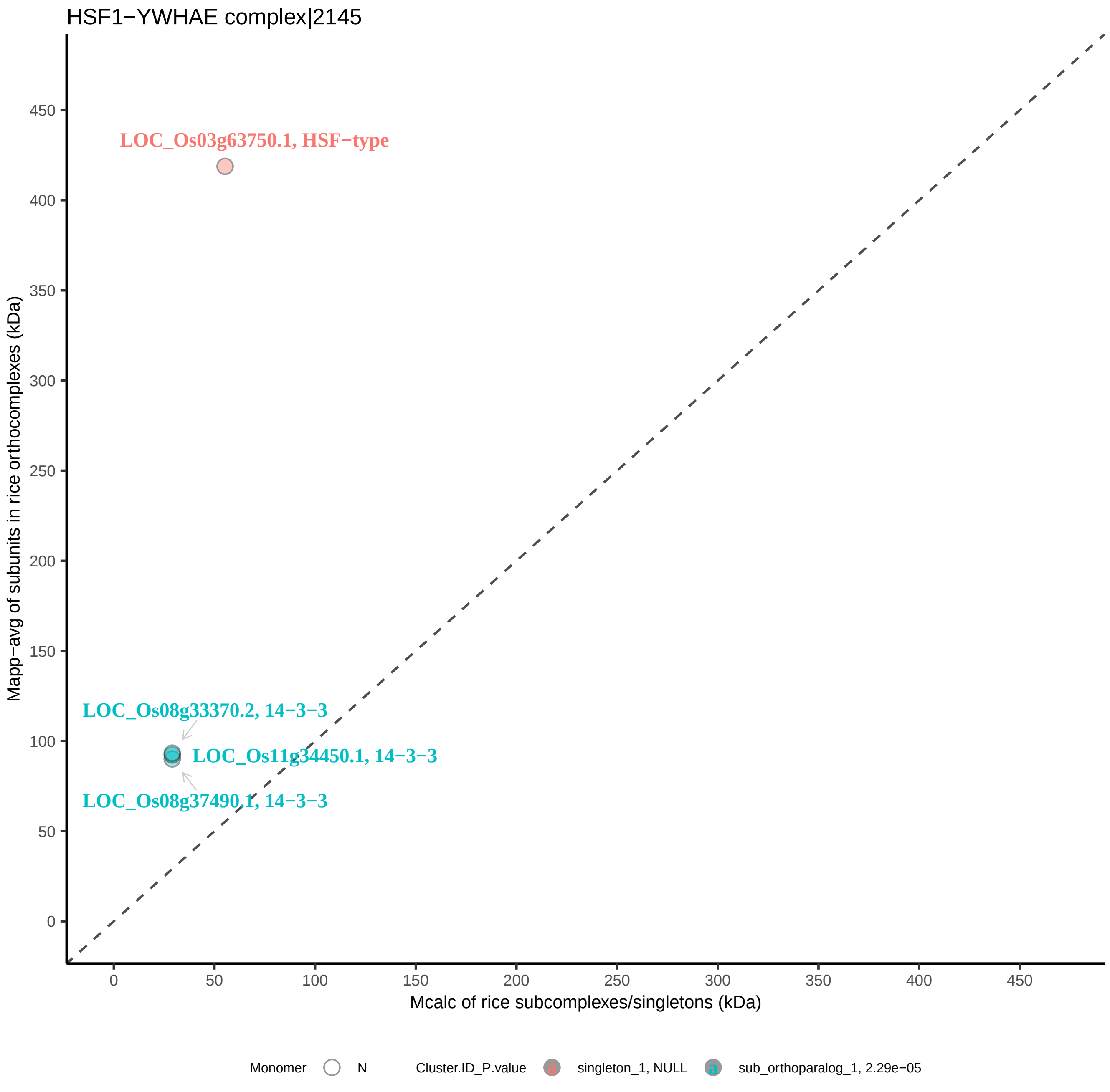
HSF1−YWHAE complex|2145
LOC_Os03g63750.1, HSF−type
LOC_Os08g33370.2, 14−3−3
LOC_Os11g34450.1, 14−3−3
LOC_Os08g37490.1, 14−3−3
Mapp−avg of subunits in rice orthocomplexes (kDa)
Mcalc of rice subcomplexes/singletons (kDa)
0
50
100
150
200
250
300
350
400
450
Monomer
N
Cluster.ID_P.value
singleton_1, NULL
sub_orthoparalog_1, 2.29e−05

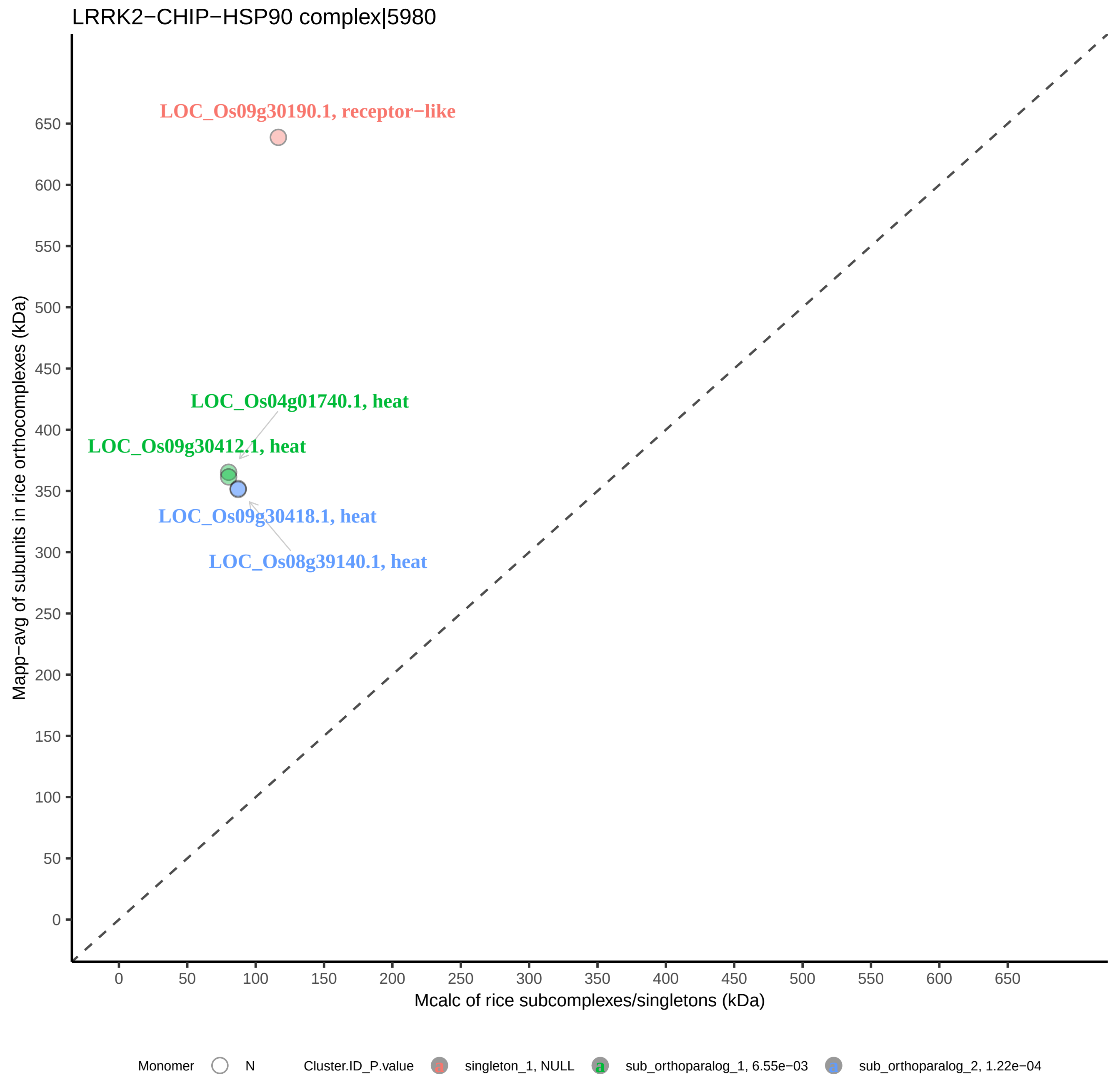

LRRK2−CHIP−HSP90 complex|5980
LOC_Os09g30190.1, receptor−like
LOC_Os04g01740.1, heat
LOC_Os09g30412.1, heat
LOC_Os09g30418.1, heat
LOC_Os08g39140.1, heat
Mapp−avg of subunits in rice orthocomplexes (kDa)
Mcalc of rice subcomplexes/singletons (kDa)
0
50
100
150
200
250
300
350
400
450
500
550
600
650
Monomer
N
Cluster.ID_P.value
singleton_1, NULL
sub_orthoparalog_1, 6.55e−03
sub_orthoparalog_2, 1.22e−04

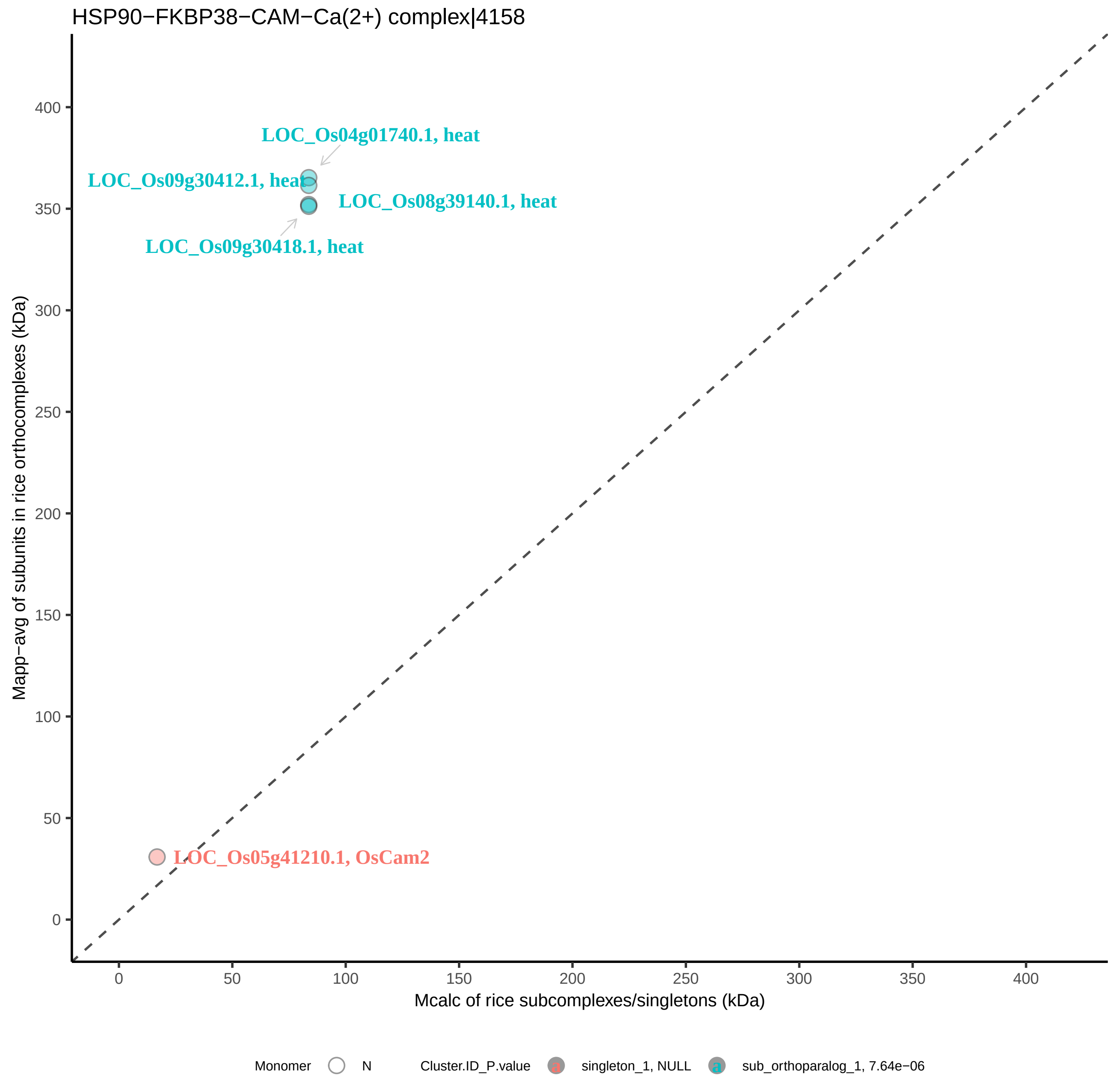

HSP90−FKBP38−CAM−Ca(2+) complex|4158
LOC_Os04g01740.1, heat
LOC_Os09g30412.1, heat
LOC_Os08g39140.1, heat
LOC_Os09g30418.1, heat
LOC_Os05g41210.1, OsCam2
Mapp−avg of subunits in rice orthocomplexes (kDa)
Mcalc of rice subcomplexes/singletons (kDa)
0
50
100
150
200
250
300
350
400
Monomer
N
Cluster.ID_P.value
singleton_1, NULL
sub_orthoparalog_1, 7.64e−06

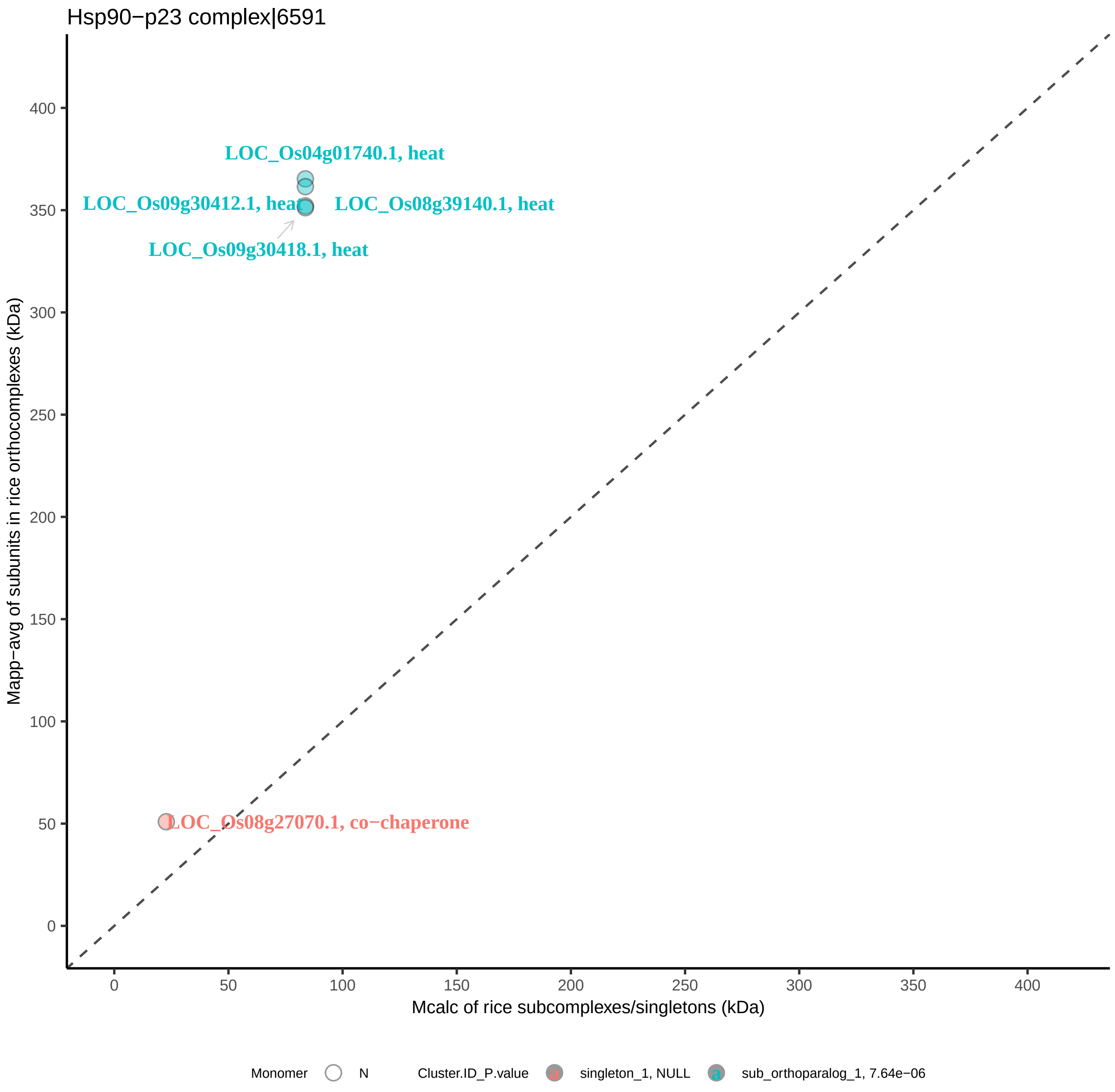
Hsp90−p23 complex|6591
LOC_Os04g01740.1, heat
LOC_Os09g30412.1, heat
LOC_Os08g39140.1, heat
LOC_Os09g30418.1, heat
LOC_Os08g27070.1, co−chaperone
Mapp−avg of subunits in rice orthocomplexes (kDa)
Mcalc of rice subcomplexes/singletons (kDa)
0
50
100
150
200
250
300
350
400
Monomer
N
Cluster.ID_P.value
singleton_1, NULL
sub_orthoparalog_1, 7.64e−06

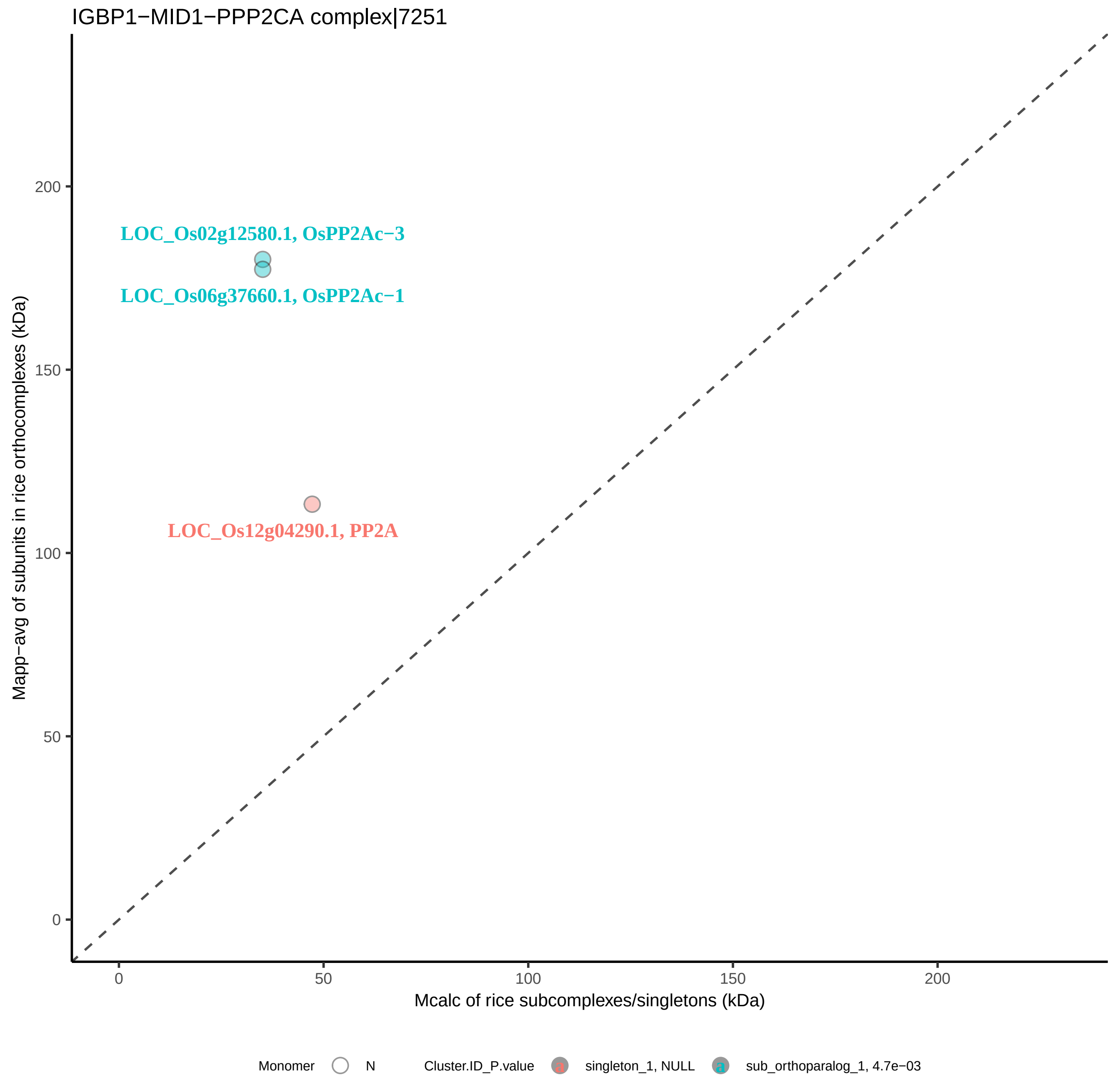

IGBP1−MID1−PPP2CA complex|7251
LOC_Os02g12580.1, OsPP2Ac−3
LOC_Os06g37660.1, OsPP2Ac−1
LOC_Os12g04290.1, PP2A
Mapp−avg of subunits in rice orthocomplexes (kDa)
Mcalc of rice subcomplexes/singletons (kDa)
0
50
100
150
200
Monomer
N
Cluster.ID_P.value
singleton_1, NULL
sub_orthoparalog_1, 4.7e−03

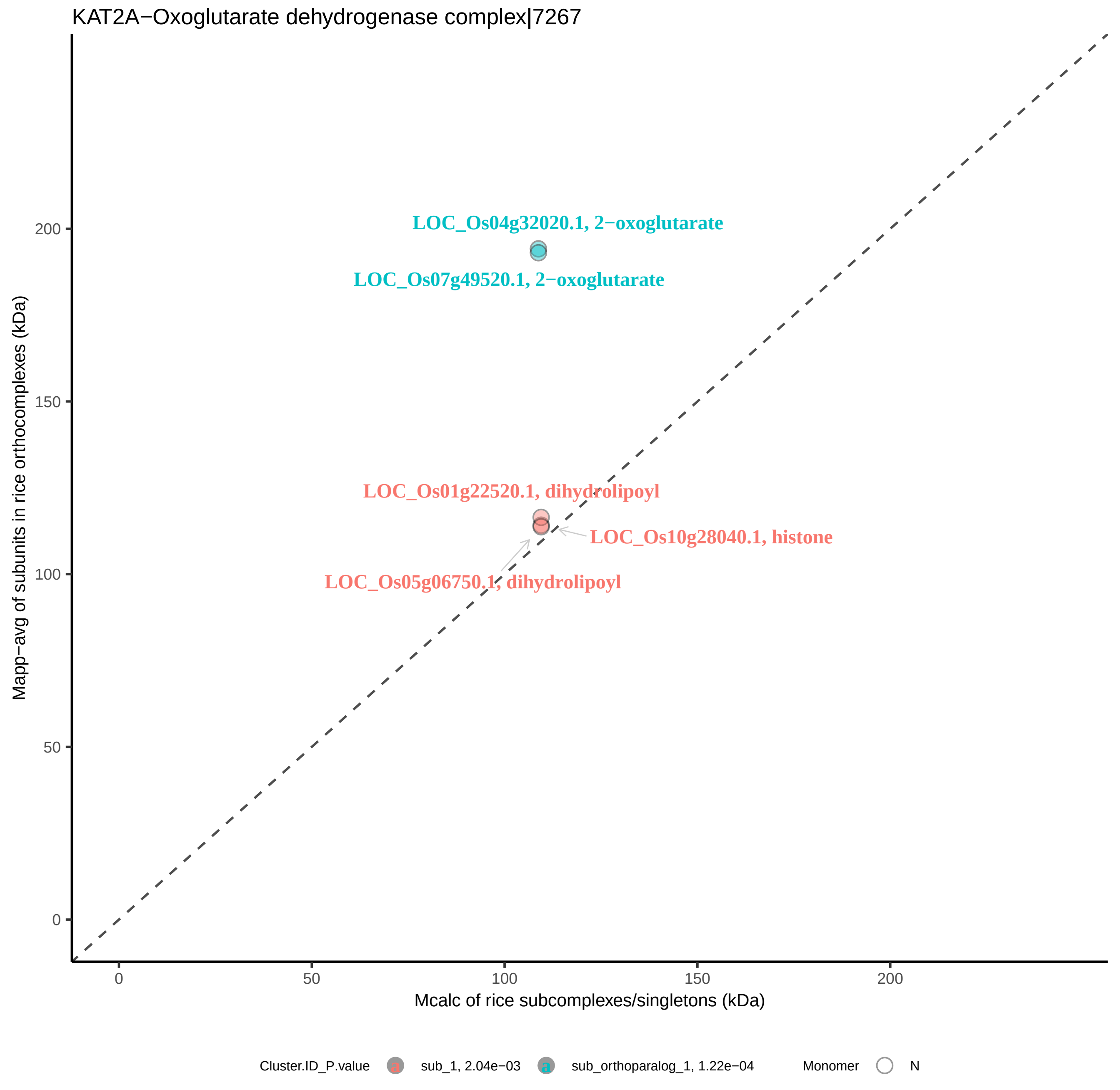
KAT2A−Oxoglutarate dehydrogenase complex|7267
LOC_Os04g32020.1, 2−oxoglutarate
LOC_Os07g49520.1, 2−oxoglutarate
LOC_Os01g22520.1, dihydrolipoyl
LOC_Os10g28040.1, histone
LOC_Os05g06750.1, dihydrolipoyl
Mapp−avg of subunits in rice orthocomplexes (kDa)
Mcalc of rice subcomplexes/singletons (kDa)
0
50
100
150
200
Cluster.ID_P.value
sub_1, 2.04e−03
sub_orthoparalog_1, 1.22e−04
Monomer
N

LARP1−PABPC1−RYDEN complex|7563

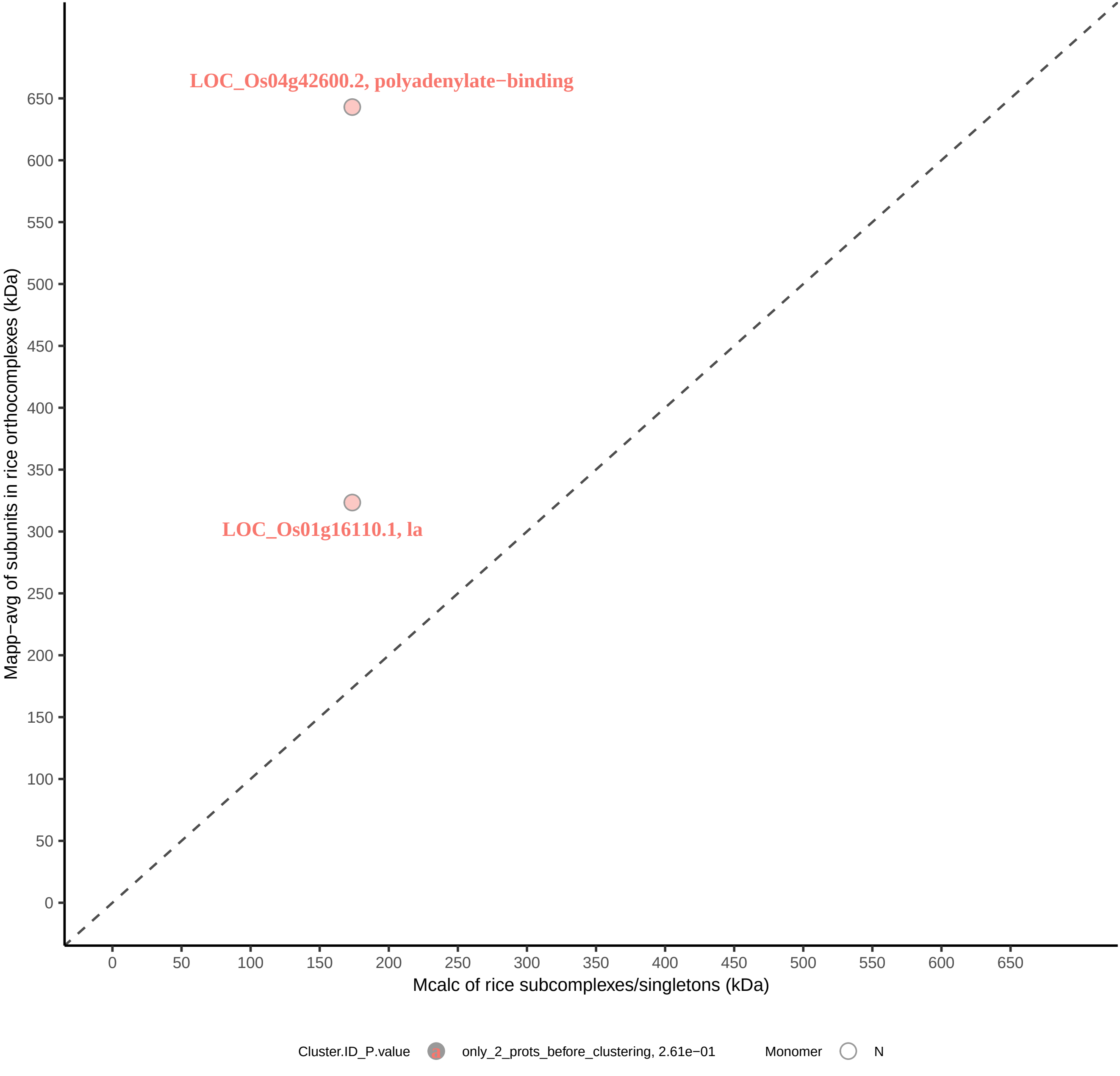

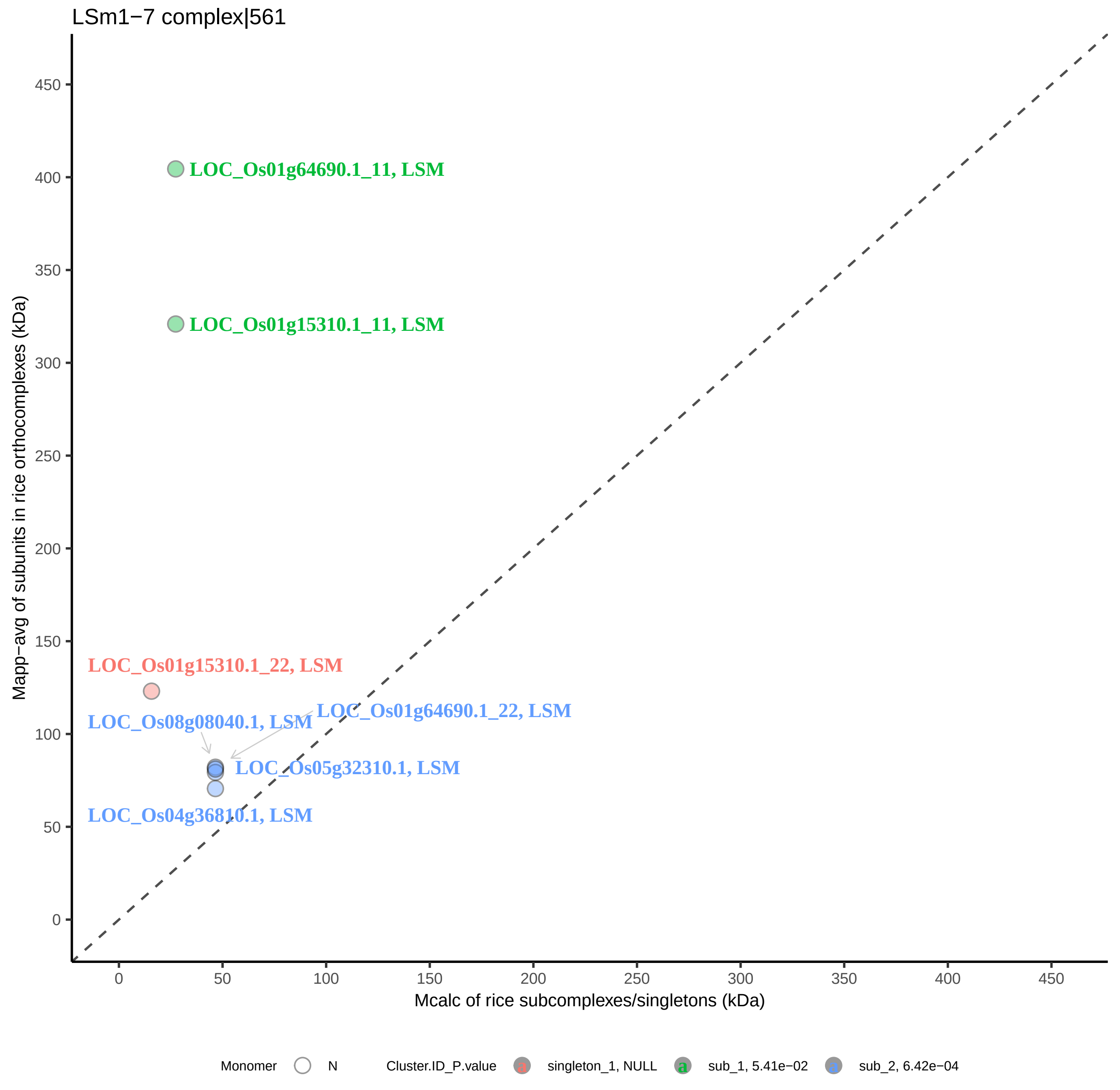
LSm1−7 complex|561
LOC_Os01g64690.1_11, LSM
LOC_Os01g15310.1_11, LSM
LOC_Os01g15310.1_22, LSM
LOC_Os01g64690.1_22, LSM
LOC_Os08g08040.1, LSM
LOC_Os05g32310.1, LSM
LOC_Os04g36810.1, LSM
Mapp−avg of subunits in rice orthocomplexes (kDa)
Mcalc of rice subcomplexes/singletons (kDa)
0
50
100
150
200
250
300
350
400
450
Monomer
N
Cluster.ID_P.value
singleton_1, NULL
sub_1, 5.41e−02
sub_2, 6.42e−04

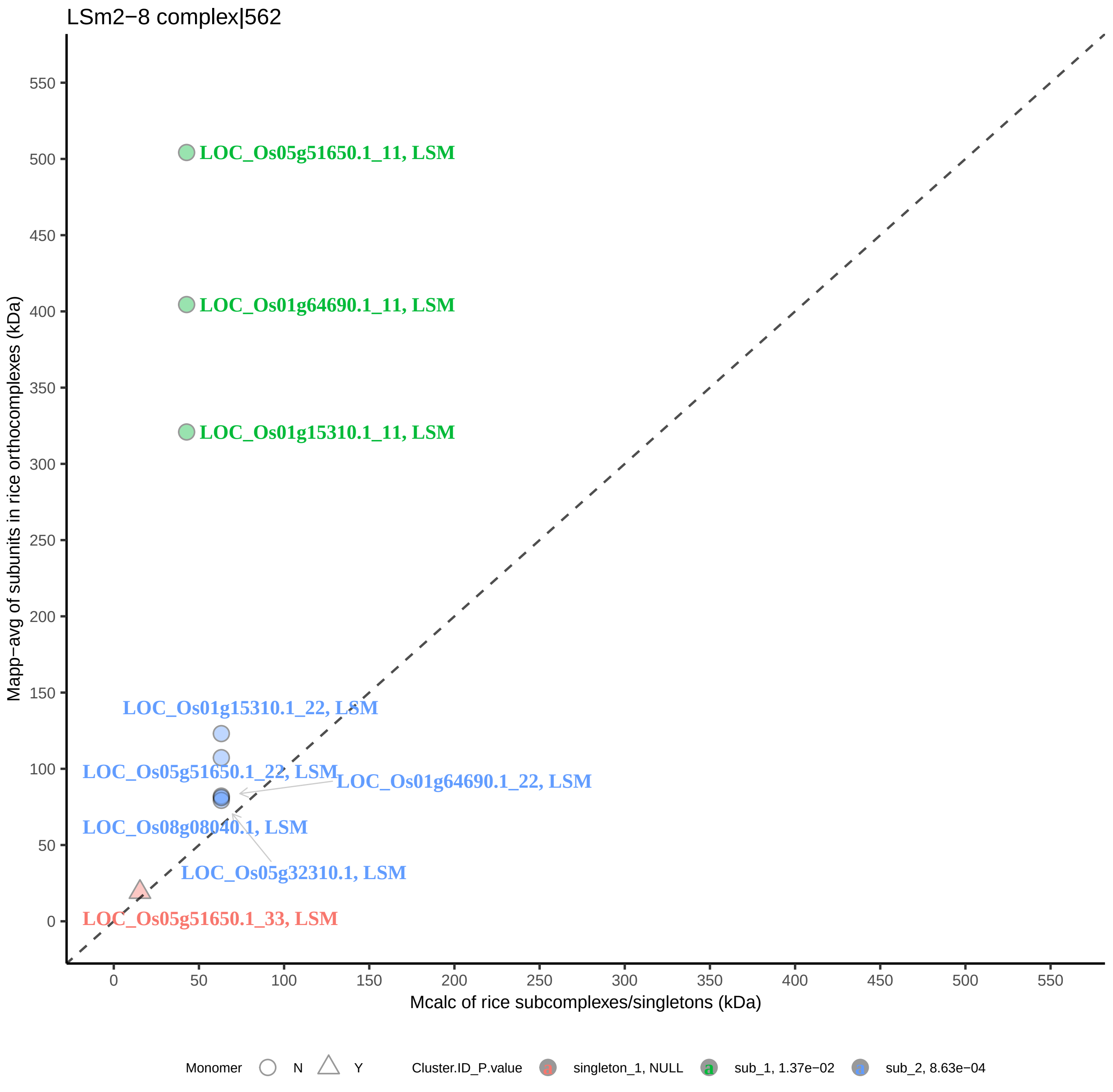
LSm2−8 complex|562
LOC_Os05g51650.1_11, LSM
LOC_Os01g64690.1_11, LSM
LOC_Os01g15310.1_11, LSM
LOC_Os01g15310.1_22, LSM
LOC_Os05g51650.1_22, LSM
LOC_Os01g64690.1_22, LSM
LOC_Os08g08040.1, LSM
LOC_Os05g32310.1, LSM
LOC_Os05g51650.1_33, LSM
Mapp−avg of subunits in rice orthocomplexes (kDa)
Mcalc of rice subcomplexes/singletons (kDa)
0
50
100
150
200
250
300
350
400
450
500
550
Monomer N Y
Cluster.ID_P.value singleton_1, NULL sub_1, 1.37e−02 sub_2, 8.63e−04

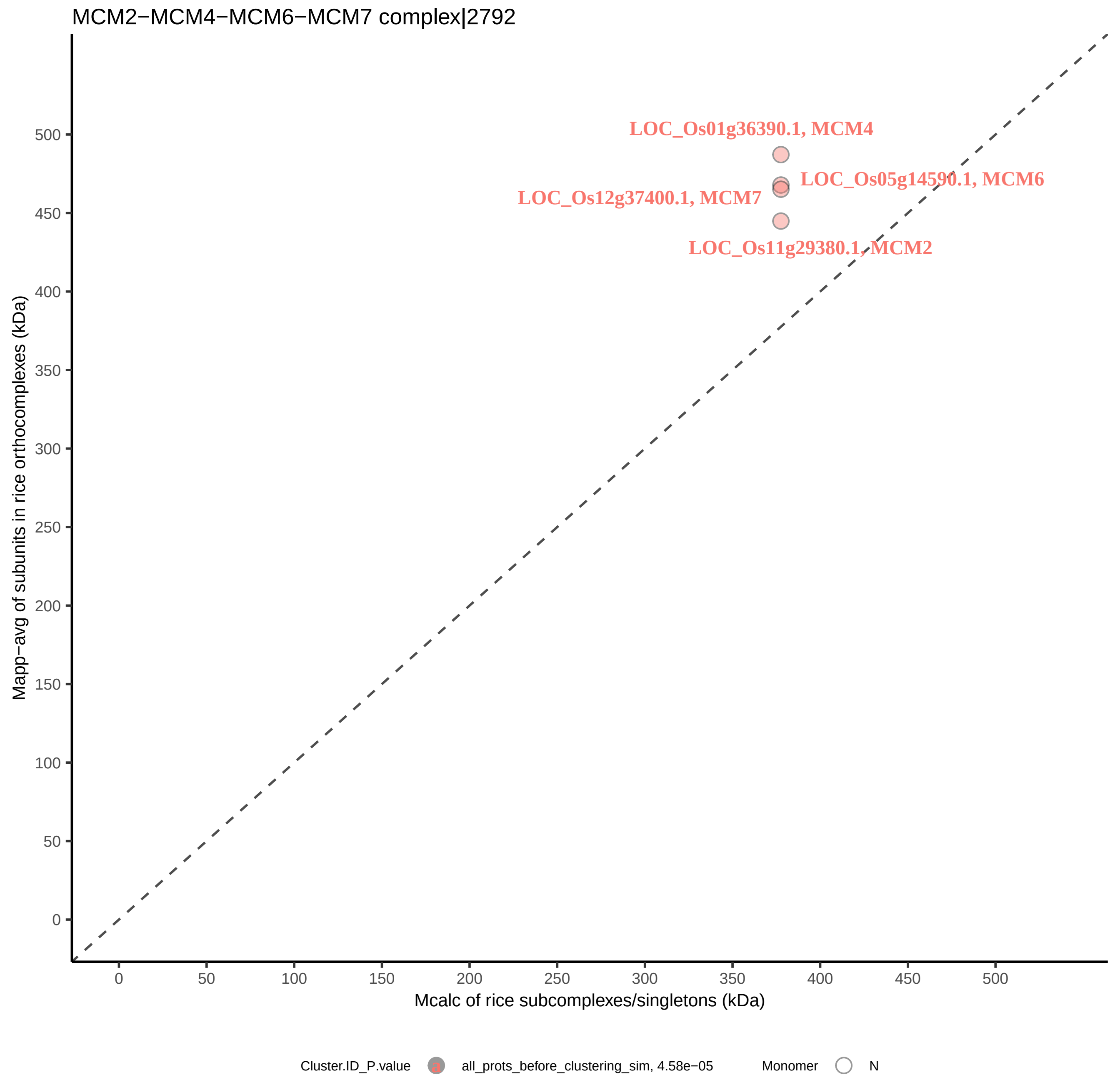

MCM2−MCM4−MCM6−MCM7 complex|2792
LOC_Os01g36390.1, MCM4
LOC_Os05g14590.1, MCM6
LOC_Os12g37400.1, MCM7
LOC_Os11g29380.1, MCM2
Mapp−avg of subunits in rice orthocomplexes (kDa)
Mcalc of rice subcomplexes/singletons (kDa)
0
50
100
150
200
250
300
350
400
450
500
Cluster.ID_P.value
all_prots_before_clustering_sim, 4.58e−05
Monomer
N

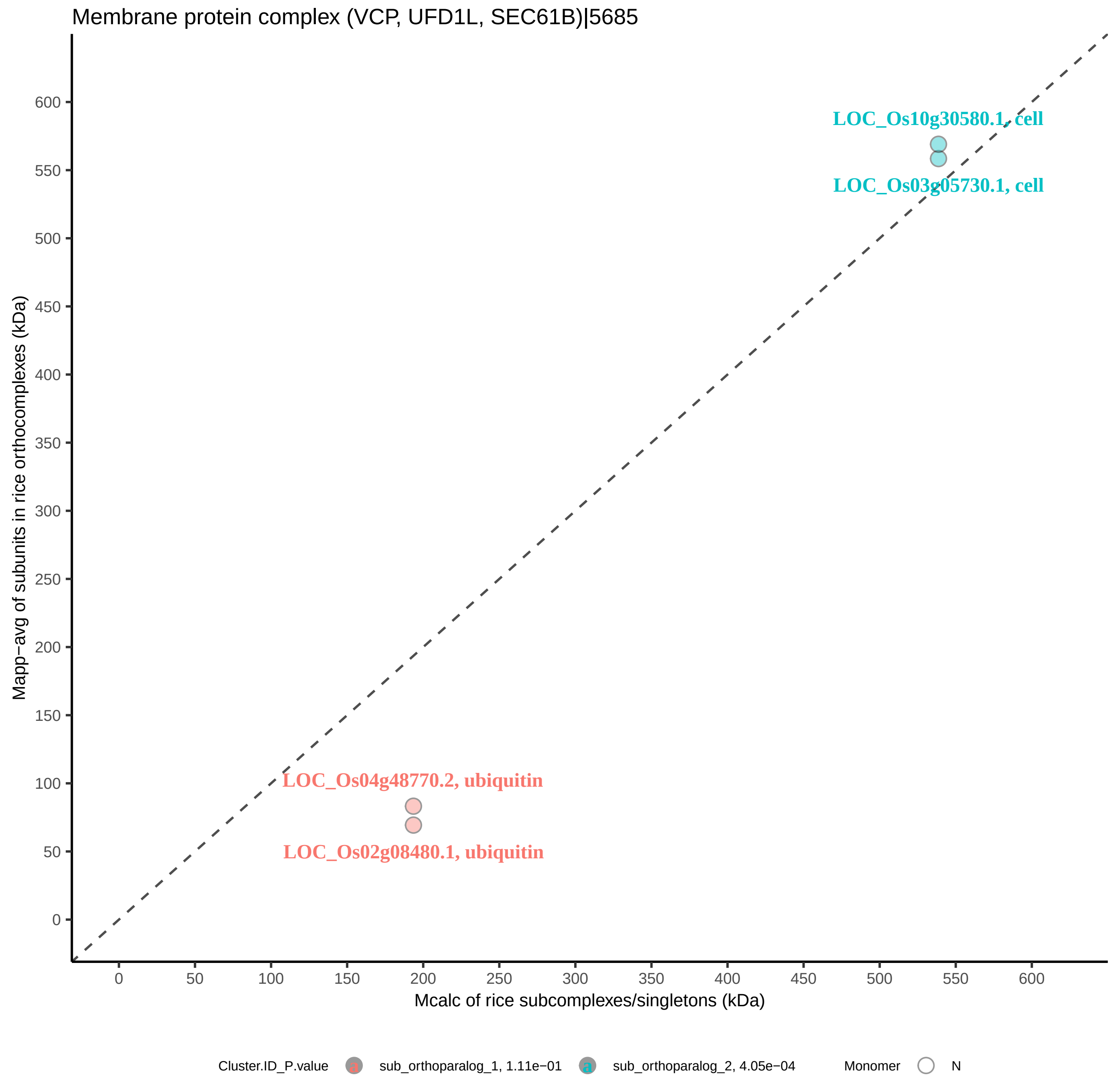
Membrane protein complex (VCP, UFD1L, SEC61B)|5685
LOC_Os10g30580.1, cell
LOC_Os03g05730.1, cell
LOC_Os04g48770.2, ubiquitin
LOC_Os02g08480.1, ubiquitin
Mapp−avg of subunits in rice orthocomplexes (kDa)
Mcalc of rice subcomplexes/singletons (kDa)
0
50
100
150
200
250
300
350
400
450
500
550
600
Cluster.ID_P.value
sub_orthoparalog_1, 1.11e−01
sub_orthoparalog_2, 4.05e−04
Monomer
N

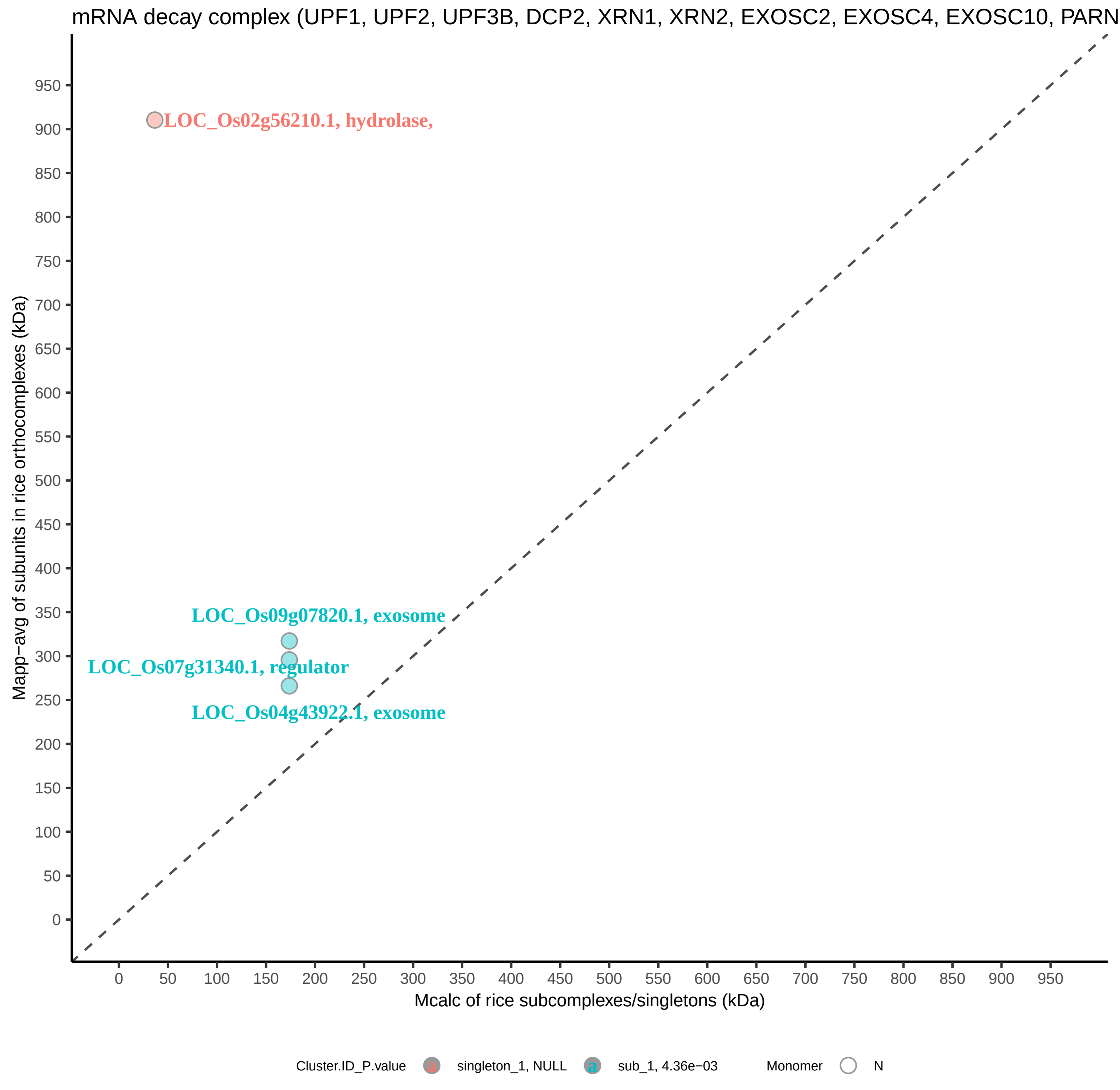

mRNA decay complex (UPF1, UPF2, UPF3B, DCP2, XRN1, XRN2, EXOSC2, EXOSC4, EXOSC10, PARN
LOC_Os02g56210.1, hydrolase,
LOC_Os09g07820.1, exosome
LOC_Os07g31340.1, regulator
LOC_Os04g43922.1, exosome
Mapp−avg of subunits in rice orthocomplexes (kDa)
Mcalc of rice subcomplexes/singletons (kDa)
0
50
100
150
200
250
300
350
400
450
500
550
600
650
700
750
800
850
900
950
Cluster.ID_P.value
singleton_1, NULL
sub_1, 4.36e−03
Monomer
N

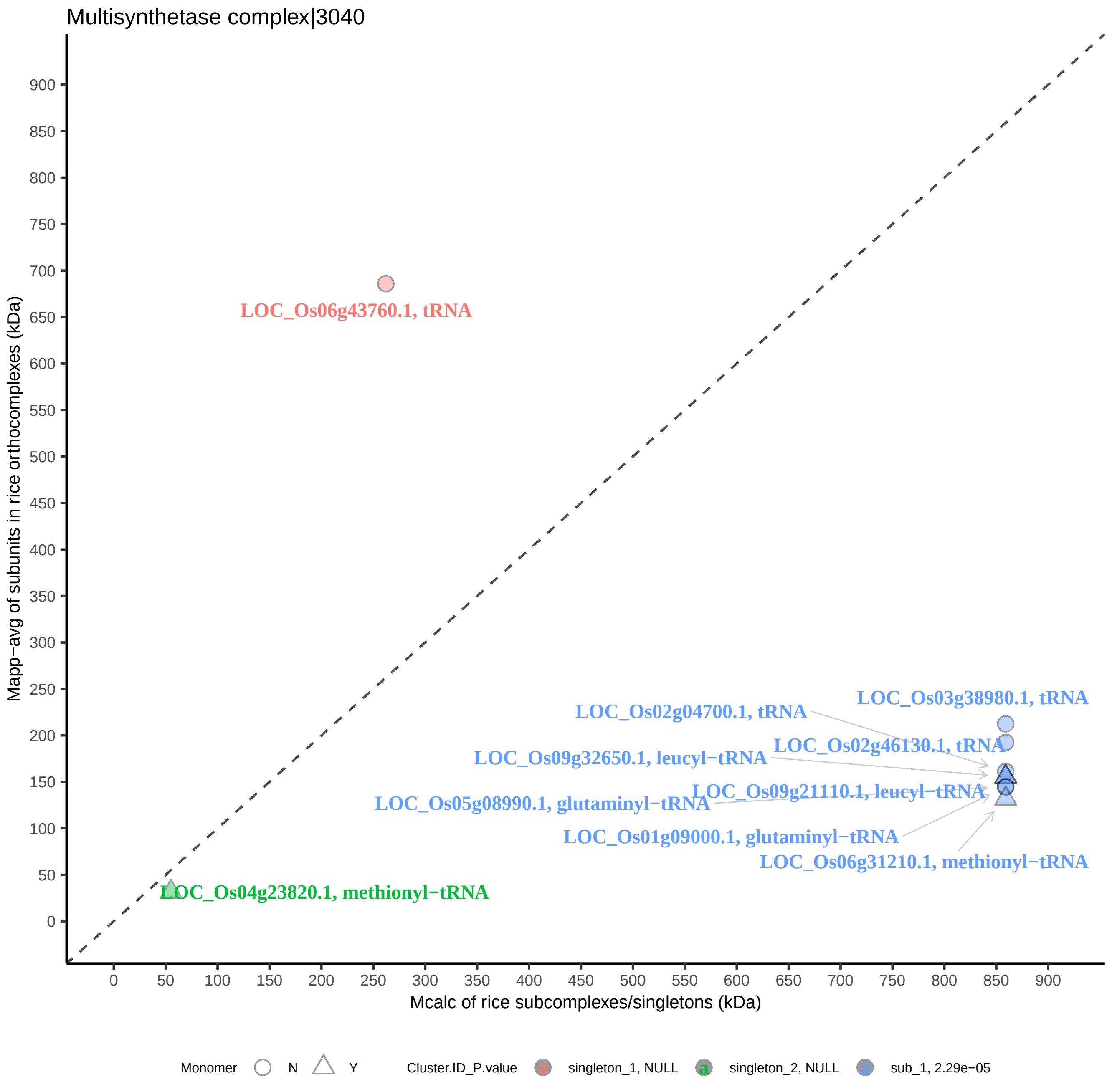
Multisynthetase complex|3040
Mapp−avg of subunits in rice orthocomplexes (kDa)
Mcalc of rice subcomplexes/singletons (kDa)
0
50
100
150
200
250
300
350
400
450
500
550
600
650
700
750
800
850
900
LOC_Os06g43760.1, tRNA
LOC_Os03g38980.1, tRNA
LOC_Os02g04700.1, tRNA
LOC_Os02g46130.1, tRNA
LOC_Os09g32650.1, leucyl−tRNA
LOC_Os09g21110.1, leucyl−tRNA
LOC_Os05g08990.1, glutaminyl−tRNA
LOC_Os01g09000.1, glutaminyl−tRNA
LOC_Os06g31210.1, methionyl−tRNA
LOC_Os04g23820.1, methionyl−tRNA
Monomer
N
Y
Cluster.ID_P.value
singleton_1, NULL
singleton_2, NULL
sub_1, 2.29e−05

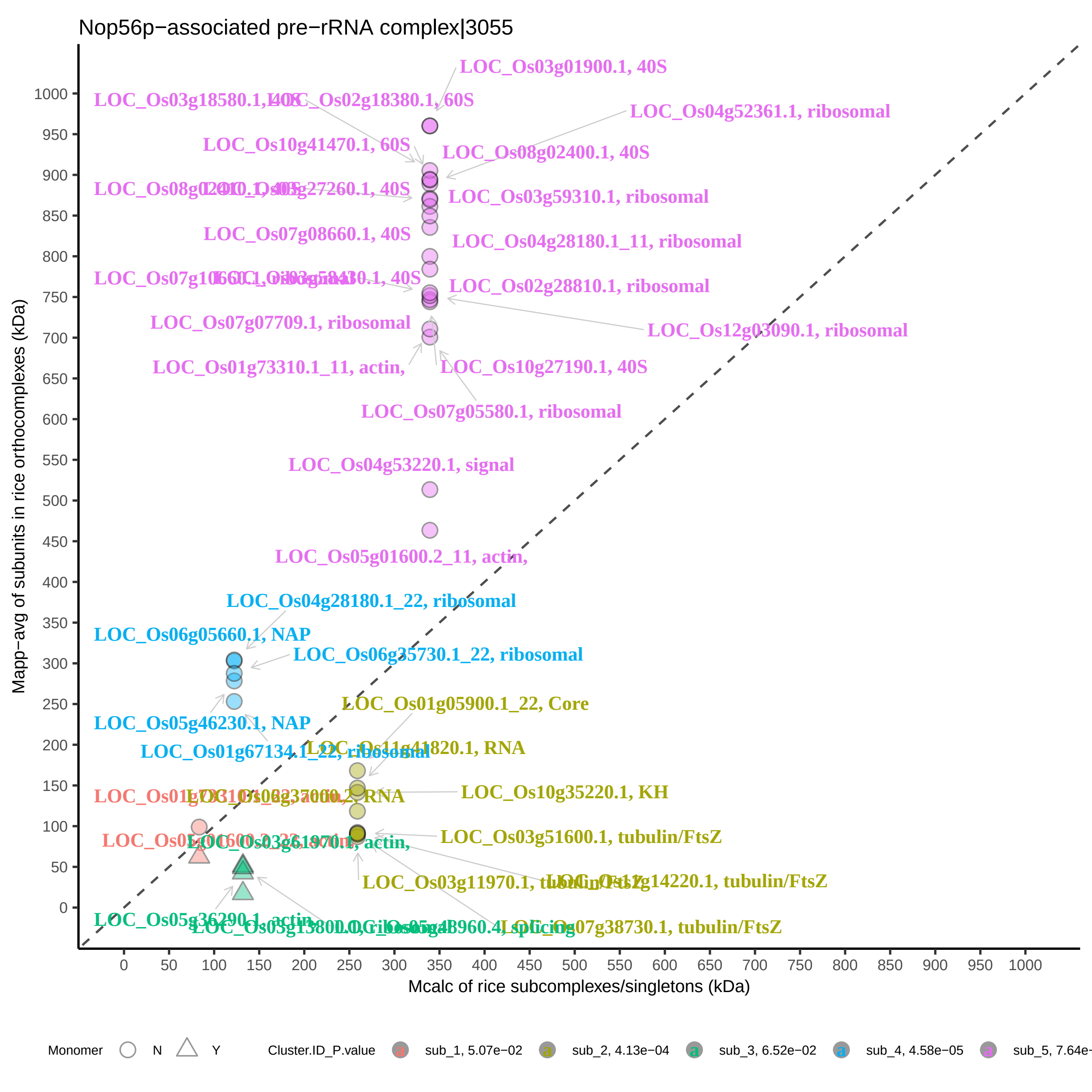

Nop56p−associated pre−rRNA complex|3055
LOC_Os03g01900.1, 40S
LOC_Os02g18380.1, 60S
LOC_Os04g52361.1, ribosomal
LOC_Os10g41470.1, 60S
LOC_Os08g02400.1, 40S
LOC_Os03g59310.1, ribosomal
LOC_Os07g08660.1, 40S
LOC_Os04g28180.1_11, ribosomal
LOC_Os02g28810.1, ribosomal
LOC_Os07g07709.1, ribosomal
LOC_Os12g03090.1, ribosomal
LOC_Os01g73310.1_11, actin,
LOC_Os10g27190.1, 40S
LOC_Os07g05580.1, ribosomal
LOC_Os04g53220.1, signal
LOC_Os05g01600.2_11, actin,
LOC_Os04g28180.1_22, ribosomal
LOC_Os06g05660.1, NAP
LOC_Os06g35730.1_22, ribosomal
LOC_Os01g05900.1_22, Core
LOC_Os05g46230.1, NAP
LOC_Os11g41820.1, RNA
LOC_Os10g35220.1, KH
LOC_Os03g51600.1, tubulin/FtsZ
LOC_Os03g11970.1, tubulin/FtsZ
LOC_Os07g38730.1, tubulin/FtsZ
LOC_Os05g36290.1, actin,
Mapp−avg of subunits in rice orthocomplexes (kDa)
Mcalc of rice subcomplexes/singletons (kDa)
Monomer
N
Y
Cluster.ID_P.value
sub_1, 5.07e−02
sub_2, 4.13e−04
sub_3, 6.52e−02
sub_4, 4.58e−05

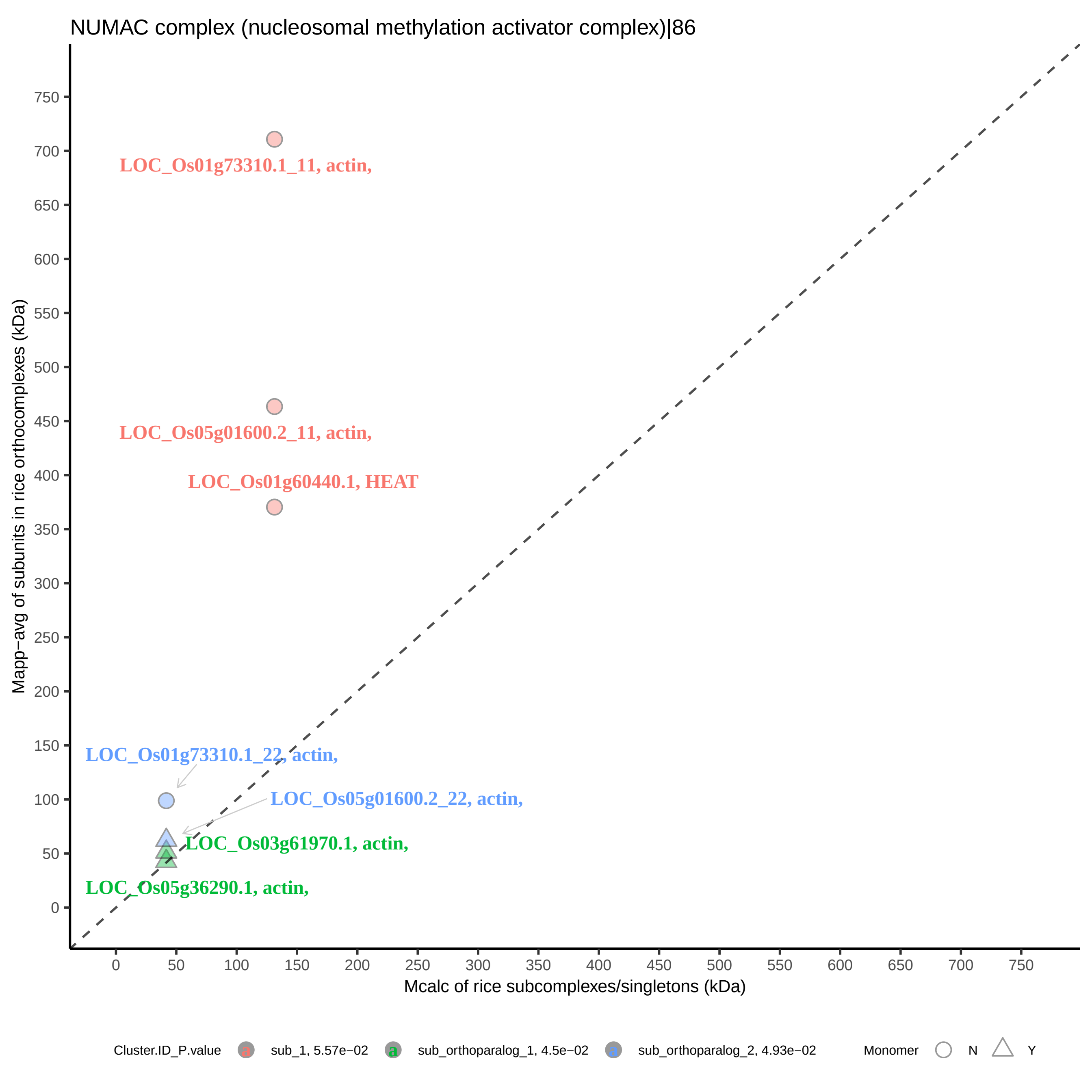

NUMAC complex (nucleosomal methylation activator complex)|86
LOC_Os01g73310.1_11, actin,
LOC_Os05g01600.2_11, actin,
LOC_Os01g60440.1, HEAT
LOC_Os01g73310.1_22, actin,
LOC_Os05g01600.2_22, actin,
LOC_Os03g61970.1, actin,
LOC_Os05g36290.1, actin,
Mapp−avg of subunits in rice orthocomplexes (kDa)
Mcalc of rice subcomplexes/singletons (kDa)
0 50 100 150 200 250 300 350 400 450 500 550 600 650 700 750
Cluster.ID_P.value
sub_1, 5.57e−02
sub_orthoparalog_1, 4.5e−02
sub_orthoparalog_2, 4.93e−02
Monomer
N
Y

OTUB1−UBC13−MMS2 complex|7051

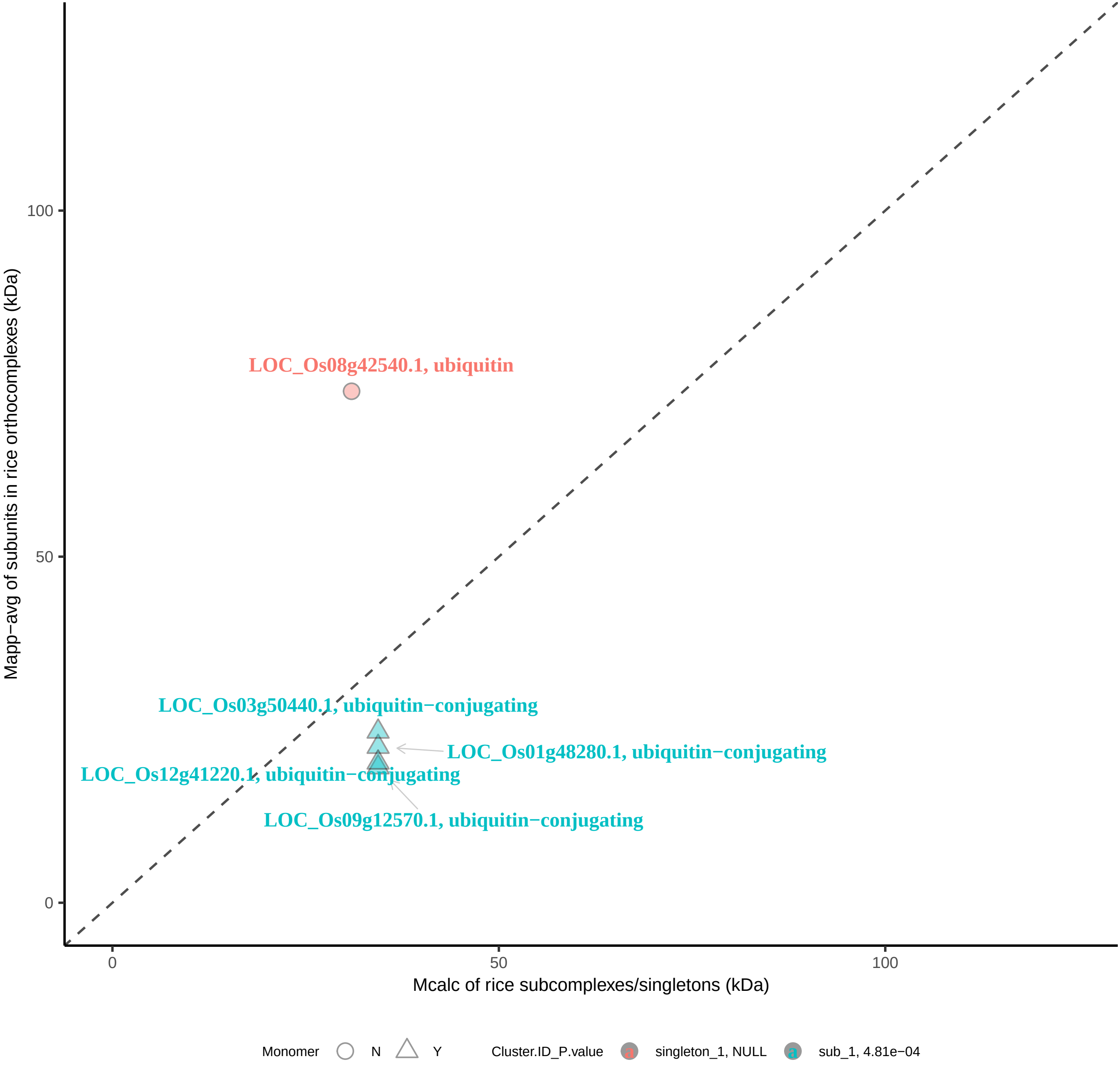

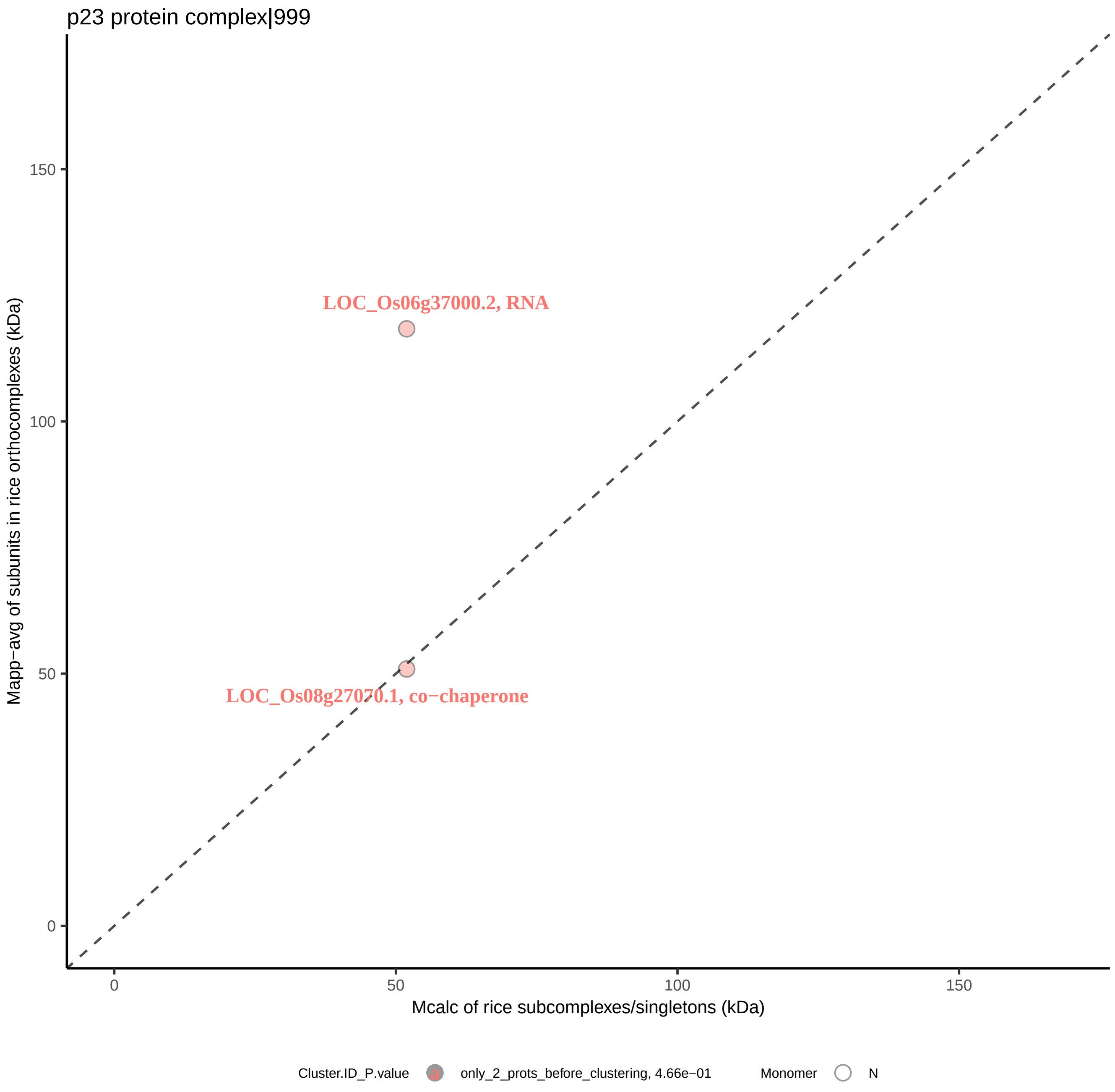

p23 protein complex|999
LOC_Os06g37000.2, RNA
LOC_Os08g27070.1, co−chaperone
150
100
50
0
Mapp−avg of subunits in rice orthocomplexes (kDa)
0
50
100
150
Mcalc of rice subcomplexes/singletons (kDa)
Cluster.ID_P.value
only_2_prots_before_clustering, 4.66e−01
Monomer
N

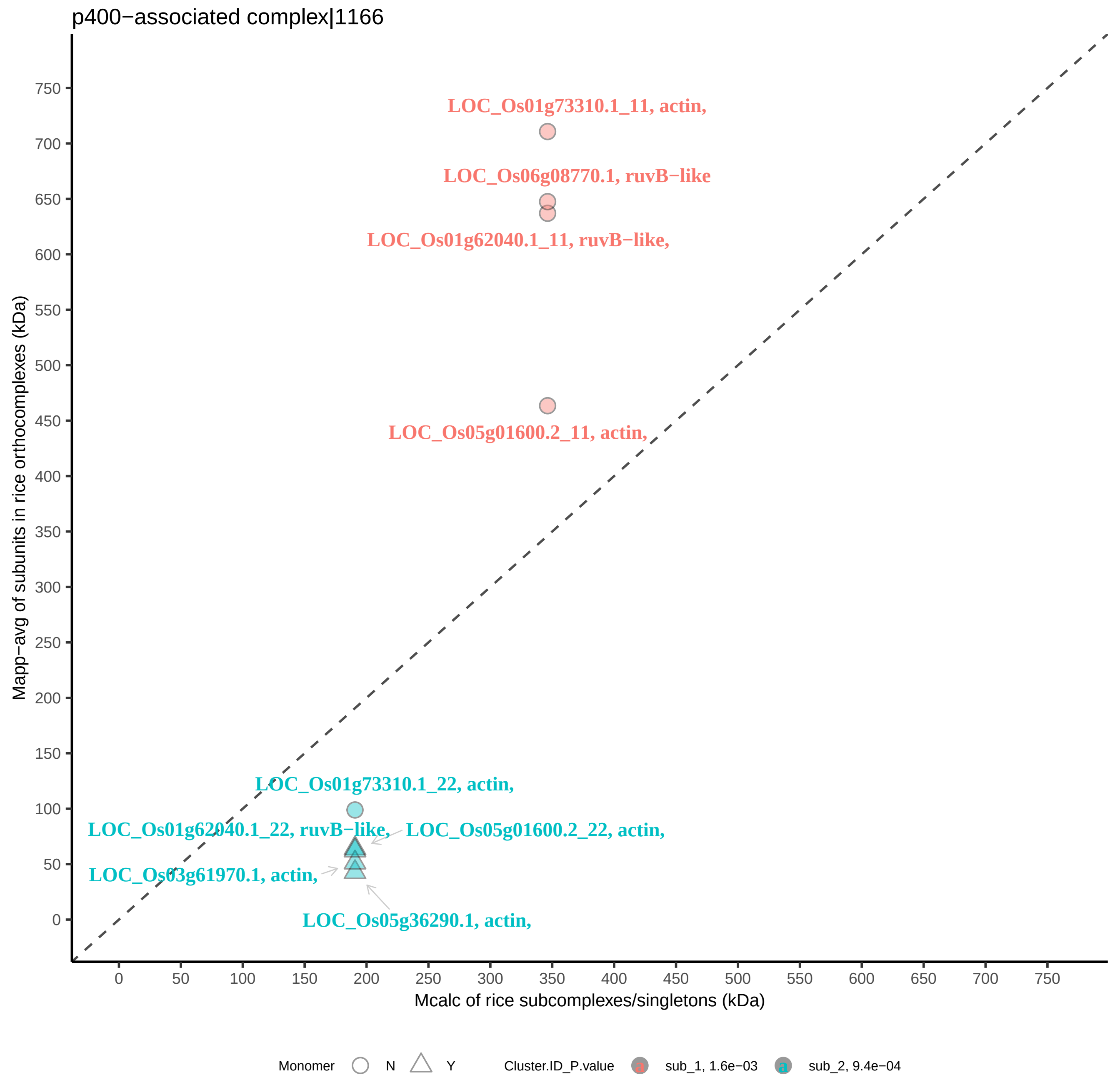

p400−associated complex|1166
LOC_Os01g73310.1_11, actin,
LOC_Os06g08770.1, ruvB−like
LOC_Os01g62040.1_11, ruvB−like,
LOC_Os05g01600.2_11, actin,
LOC_Os01g73310.1_22, actin,
LOC_Os01g62040.1_22, ruvB−like,
LOC_Os05g01600.2_22, actin,
LOC_Os03g61970.1, actin,
LOC_Os05g36290.1, actin,
Mapp−avg of subunits in rice orthocomplexes (kDa)
Mcalc of rice subcomplexes/singletons (kDa)
Monomer
N
Y
Cluster.ID_P.value
sub_1, 1.6e−03
sub_2, 9.4e−04

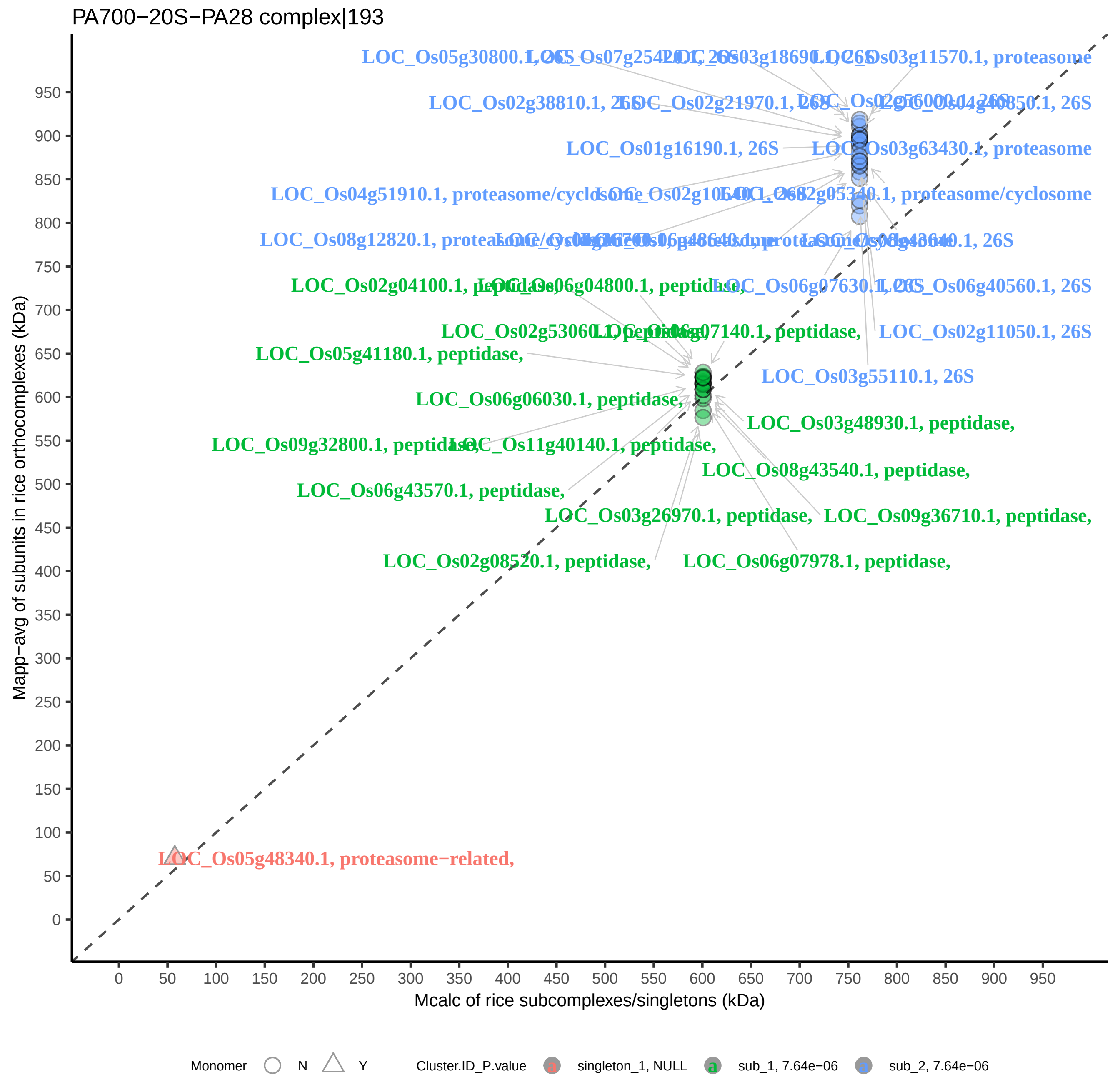
PA700−20S−PA28 complex|193
Mapp−avg of subunits in rice orthocomplexes (kDa)
Mcalc of rice subcomplexes/singletons (kDa)
LOC_Os05g48340.1, proteasome−related,
LOC_Os05g41180.1, peptidase,
LOC_Os06g06030.1, peptidase,
LOC_Os06g43570.1, peptidase,
LOC_Os02g08520.1, peptidase,
LOC_Os06g07978.1, peptidase,
LOC_Os09g36710.1, peptidase,
LOC_Os08g43540.1, peptidase,
LOC_Os03g48930.1, peptidase,
LOC_Os03g26970.1, peptidase,
LOC_Os11g40140.1, peptidase,
LOC_Os02g11050.1, 26S
LOC_Os03g55110.1, 26S
LOC_Os06g40560.1, 26S
LOC_Os01g16190.1, 26S
LOC_Os03g63430.1, proteasome
LOC_Os02g38810.1, 26S
LOC_Os03g11570.1, proteasome
Monomer N Y Cluster.ID_P.value singleton_1, NULL sub_1, 7.64e−06 sub_2, 7.64e−06

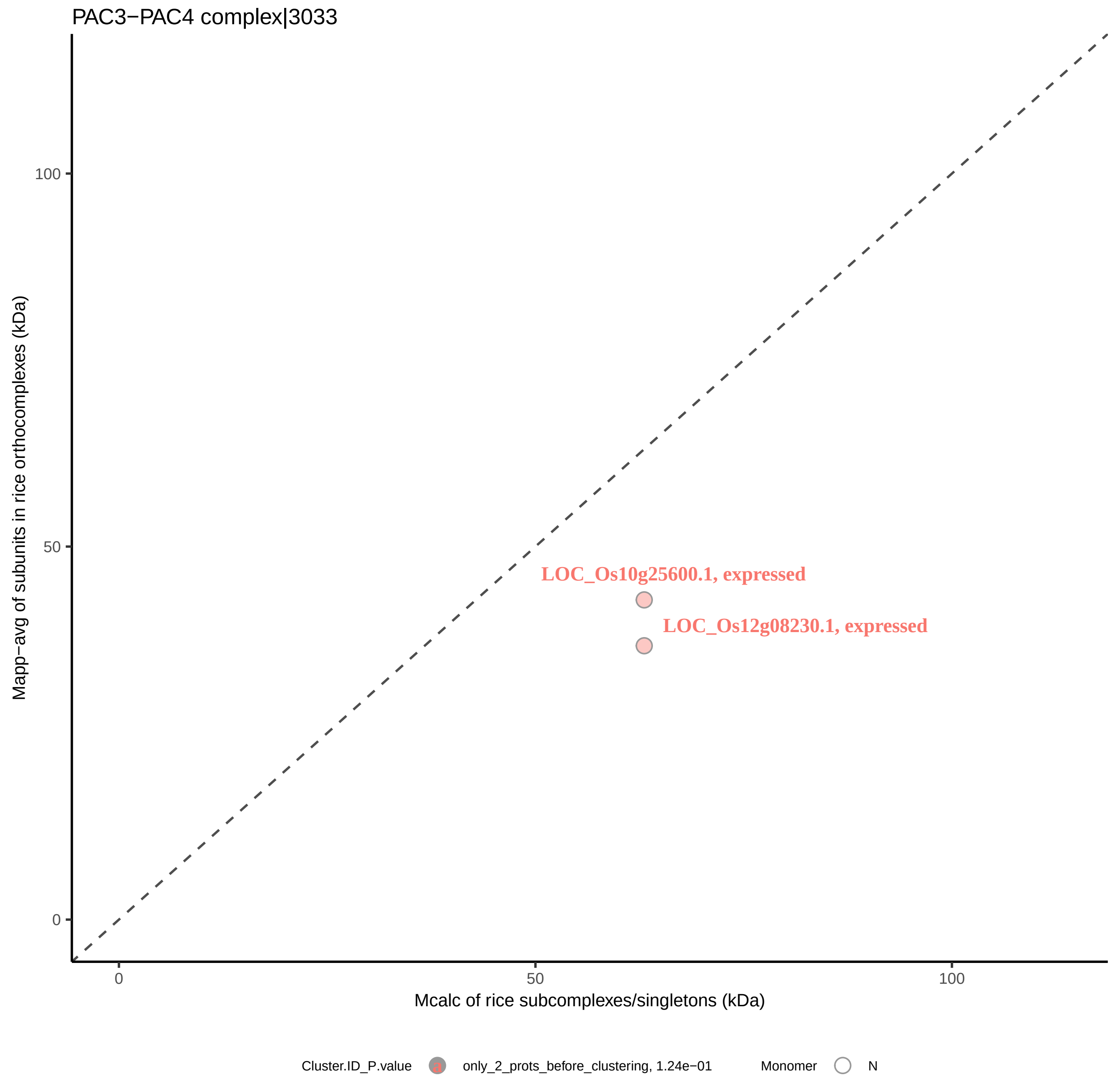

PAC3−PAC4 complex|3033
Mapp−avg of subunits in rice orthocomplexes (kDa)
100
50
0
LOC_Os10g25600.1, expressed
LOC_Os12g08230.1, expressed
0
50
100
Mcalc of rice subcomplexes/singletons (kDa)
Cluster.ID_P.value
only_2_prots_before_clustering, 1.24e−01
Monomer
N

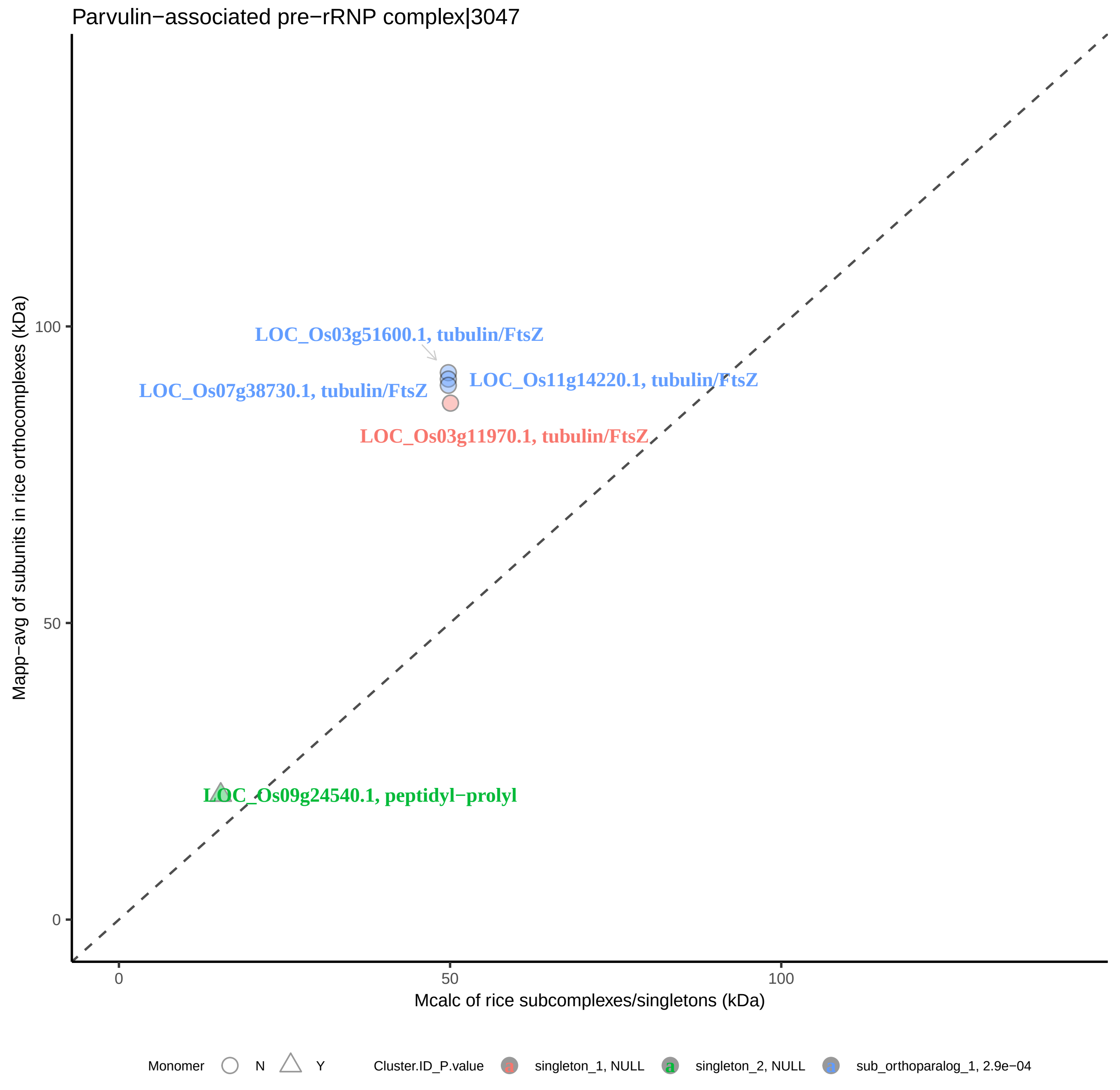

Parvulin−associated pre−rRNP complex|3047
LOC_Os03g51600.1, tubulin/FtsZ
LOC_Os11g14220.1, tubulin/FtsZ
LOC_Os07g38730.1, tubulin/FtsZ
LOC_Os03g11970.1, tubulin/FtsZ
LOC_Os09g24540.1, peptidyl−prolyl
Mapp−avg of subunits in rice orthocomplexes (kDa)
100
50
0
0
50
100
Mcalc of rice subcomplexes/singletons (kDa)
Monomer
N
Y
Cluster.ID_P.value
singleton_1, NULL
singleton_2, NULL
sub_orthoparalog_1, 2.9e−04

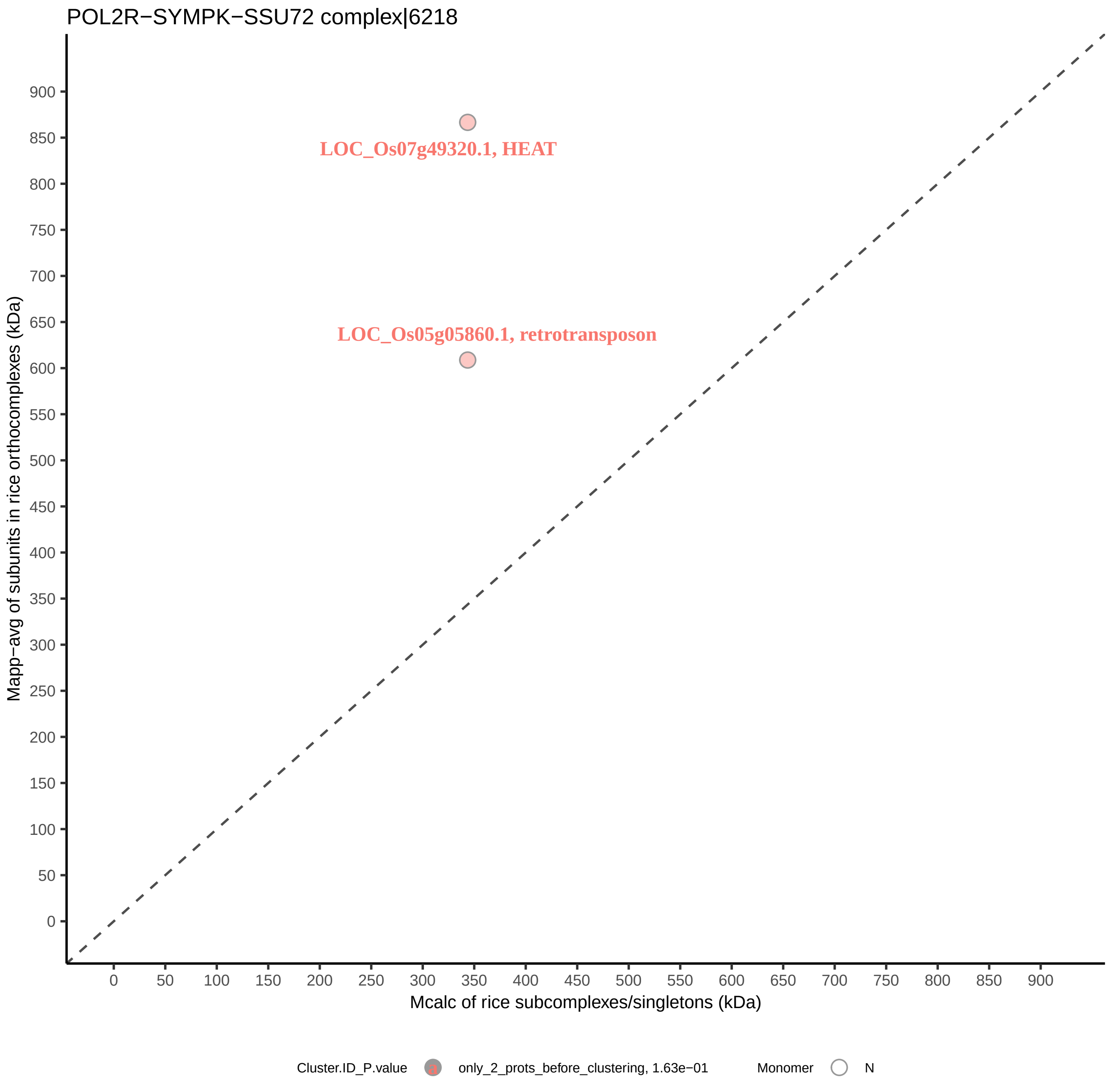

POL2R−SYMPK−SSU72 complex|6218
LOC_Os07g49320.1, HEAT
LOC_Os05g05860.1, retrotransposon
Mapp−avg of subunits in rice orthocomplexes (kDa)
Mcalc of rice subcomplexes/singletons (kDa)
0
50
100
150
200
250
300
350
400
450
500
550
600
650
700
750
800
850
900
Cluster.ID_P.value
only_2_prots_before_clustering, 1.63e−01
Monomer
N

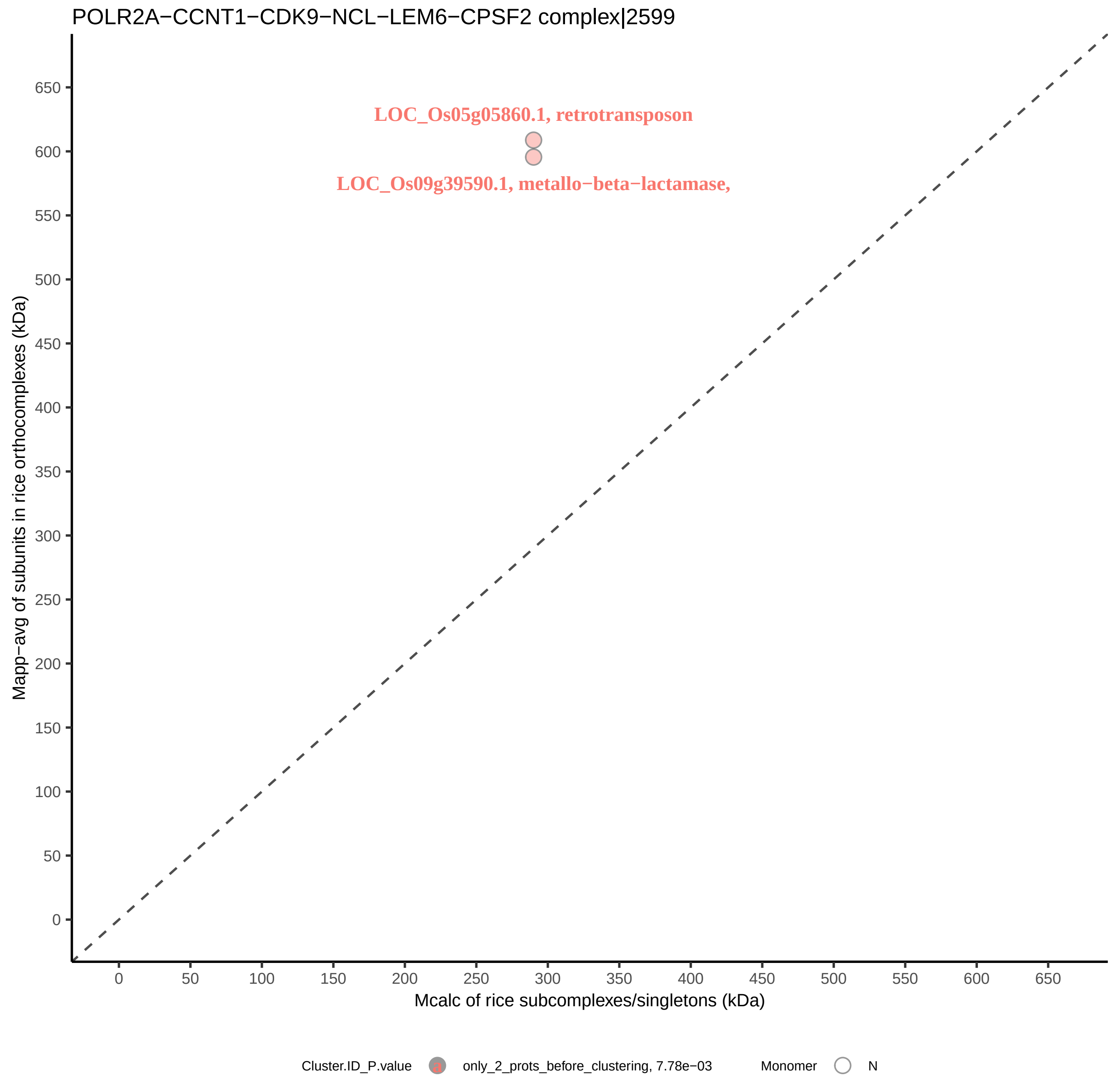

POLR2A−CCNT1−CDK9−NCL−LEM6−CPSF2 complex|2599
LOC_Os05g05860.1, retrotransposon
LOC_Os09g39590.1, metallo−beta−lactamase,
Mapp−avg of subunits in rice orthocomplexes (kDa)
Mcalc of rice subcomplexes/singletons (kDa)
0
50
100
150
200
250
300
350
400
450
500
550
600
650
Cluster.ID_P.value
only_2_prots_before_clustering, 7.78e−03
Monomer
N

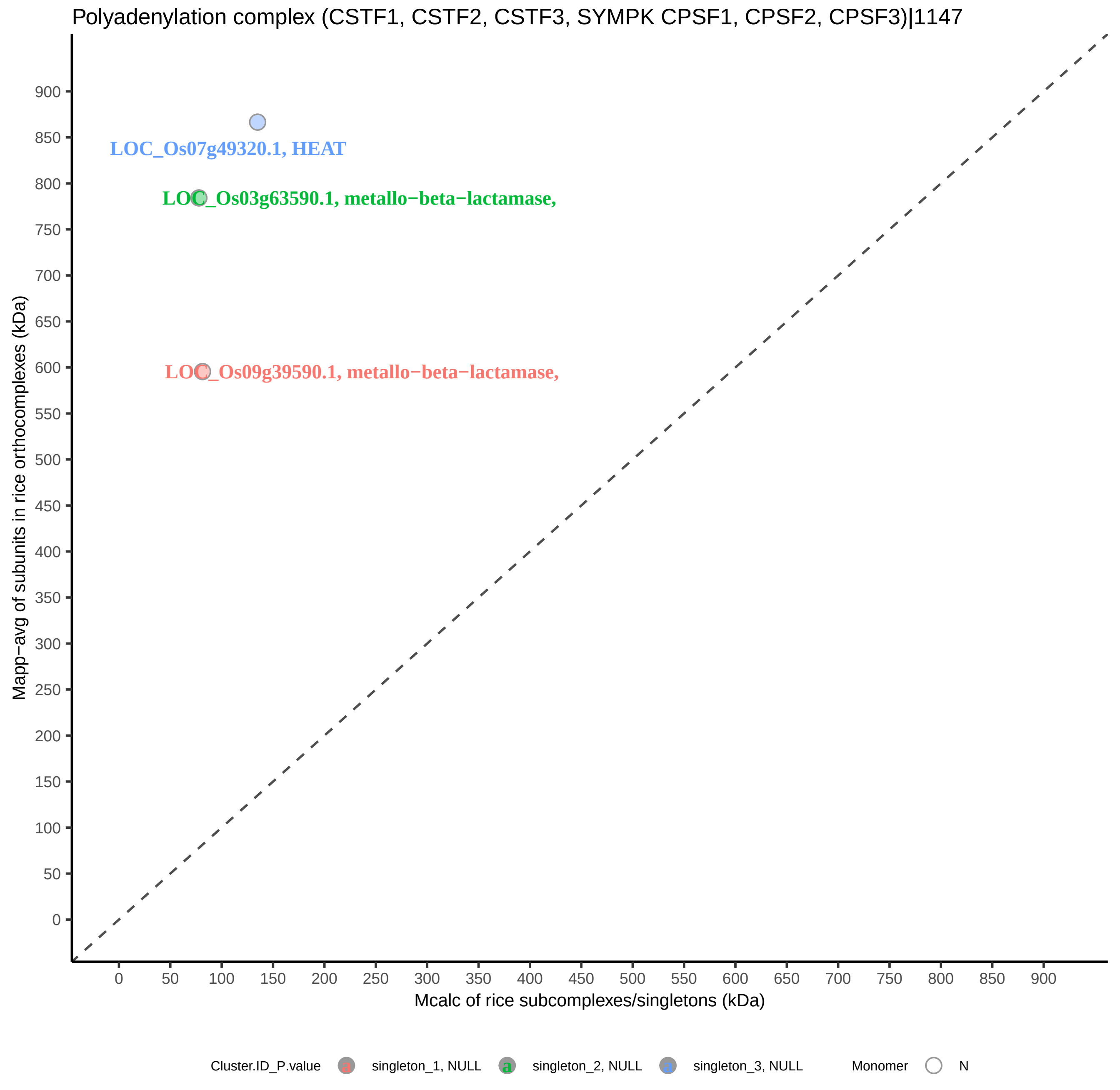
Polyadenylation complex (CSTF1, CSTF2, CSTF3, SYMPK CPSF1, CPSF2, CPSF3)|1147
LOC_Os07g49320.1, HEAT
LOC_Os03g63590.1, metallo−beta−lactamase,
LOC_Os09g39590.1, metallo−beta−lactamase,
Mapp−avg of subunits in rice orthocomplexes (kDa)
Mcalc of rice subcomplexes/singletons (kDa)
0
50
100
150
200
250
300
350
400
450
500
550
600
650
700
750
800
850
900
Cluster.ID_P.value
singleton_1, NULL
singleton_2, NULL
singleton_3, NULL
Monomer
N

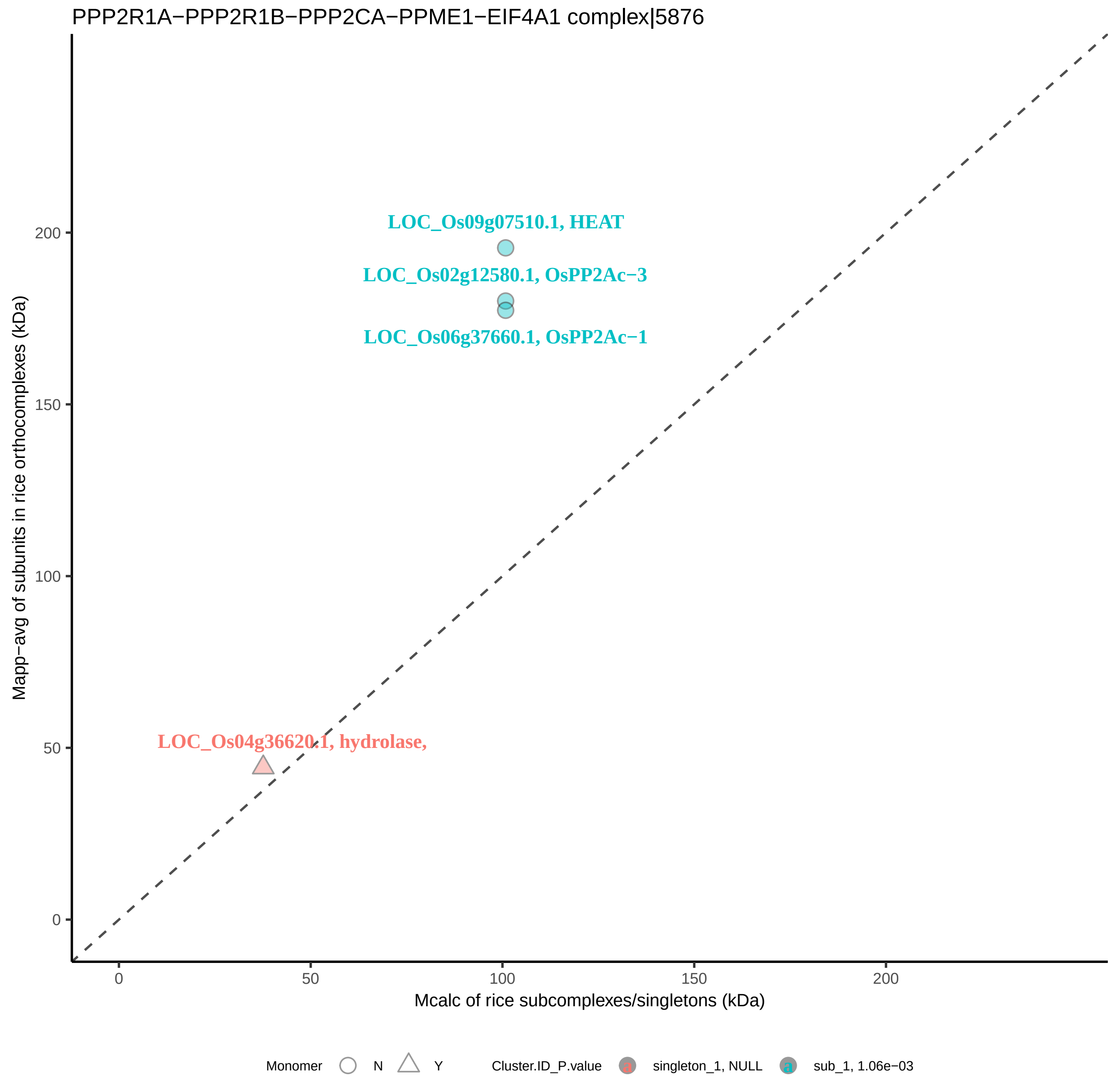

PPP2R1A−PPP2R1B−PPP2CA−PPME1−EIF4A1 complex|5876
LOC_Os09g07510.1, HEAT
LOC_Os02g12580.1, OsPP2Ac−3
LOC_Os06g37660.1, OsPP2Ac−1
LOC_Os04g36620.1, hydrolase,
Mapp−avg of subunits in rice orthocomplexes (kDa)
Mcalc of rice subcomplexes/singletons (kDa)
0
50
100
150
200
Monomer
N
Y
Cluster.ID_P.value
singleton_1, NULL
sub_1, 1.06e−03

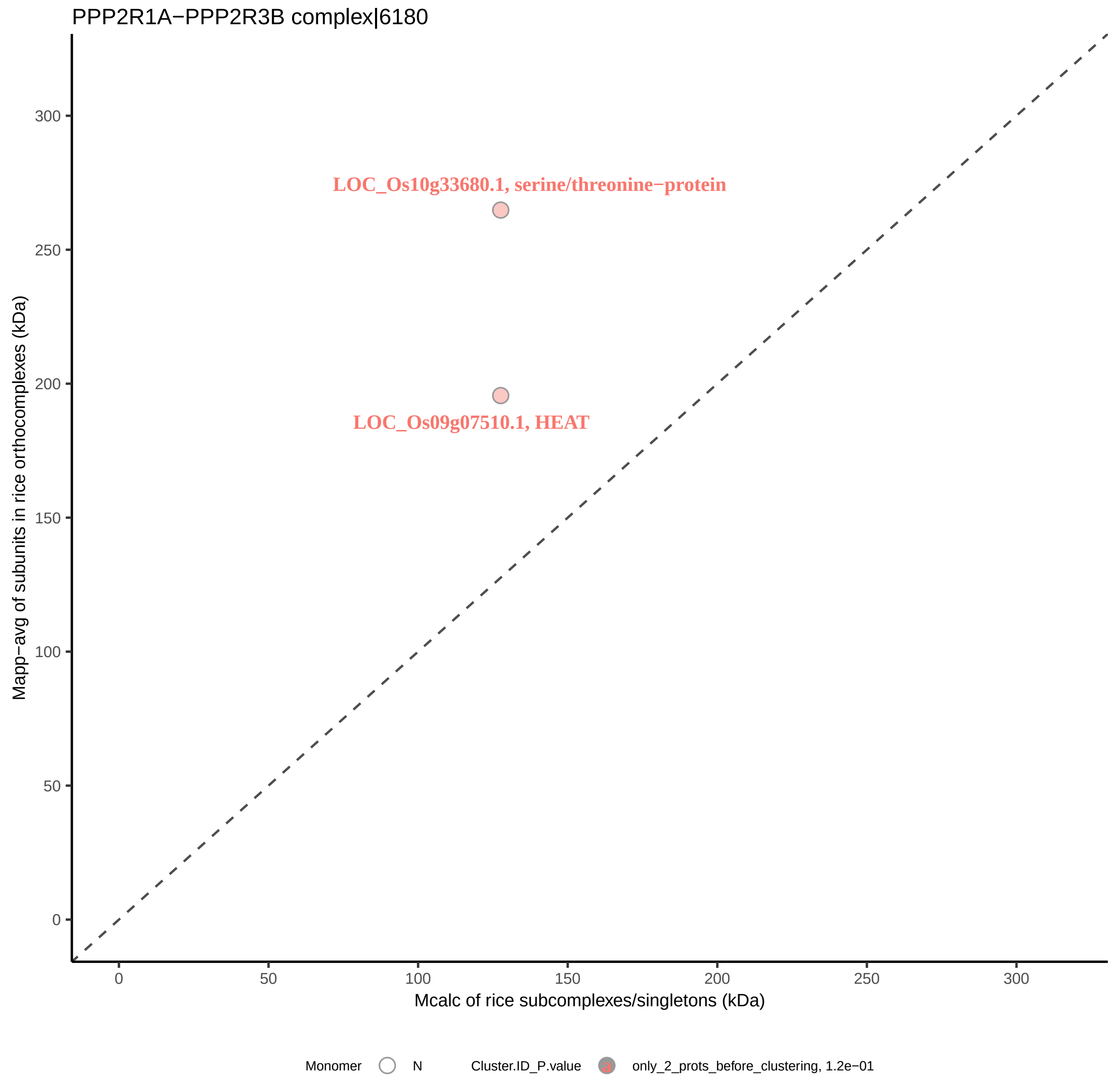

PPP2R1A−PPP2R3B complex|6180
LOC_Os10g33680.1, serine/threonine−protein
LOC_Os09g07510.1, HEAT
Mapp−avg of subunits in rice orthocomplexes (kDa)
Mcalc of rice subcomplexes/singletons (kDa)
0
50
100
150
200
250
300
Monomer
N
Cluster.ID_P.value
a
only_2_prots_before_clustering, 1.2e−01

Pre−initiation complex (PIC)|5736

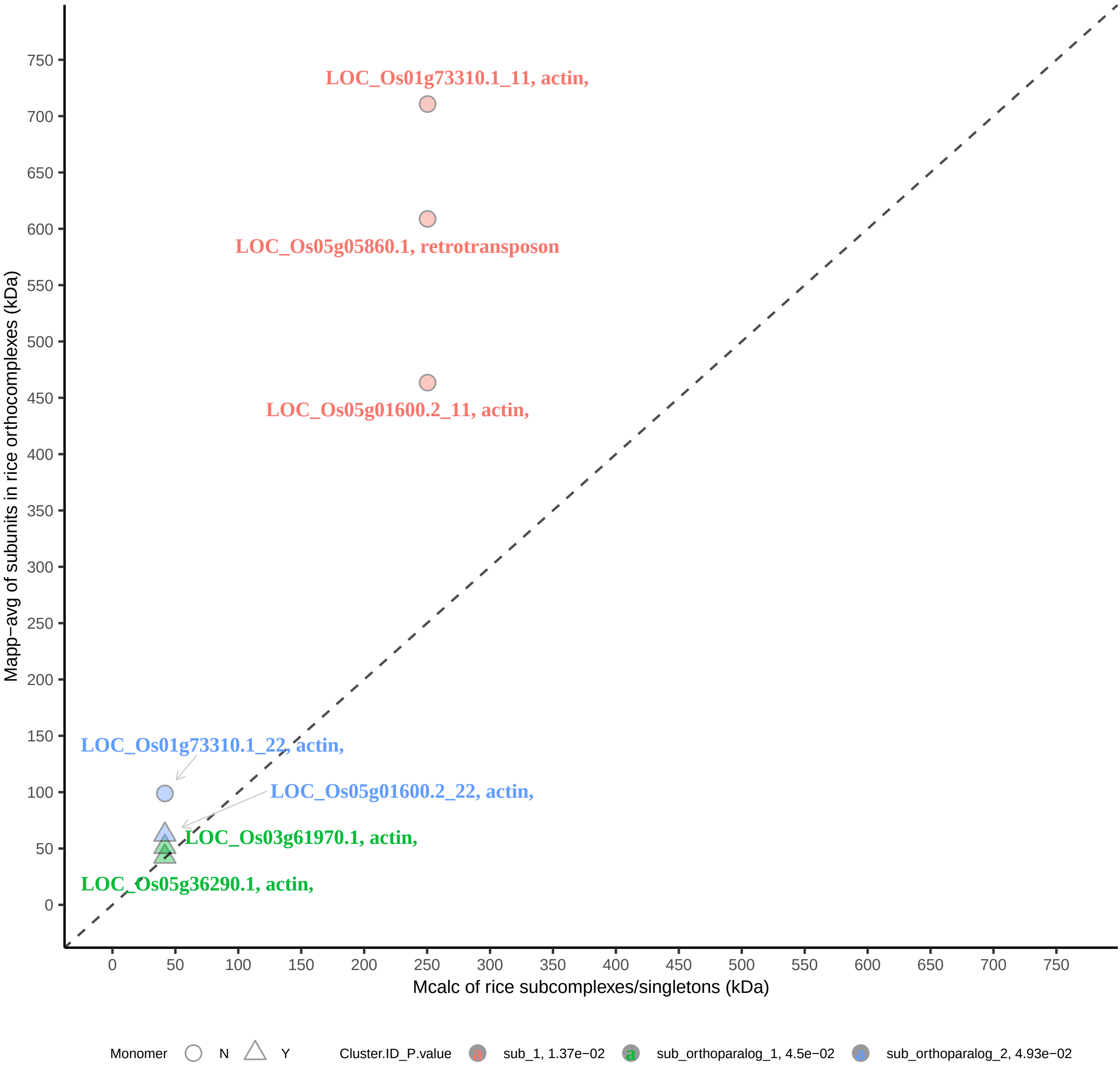

PRKAA2−PRKAB2−PRKAG3 complex|6153

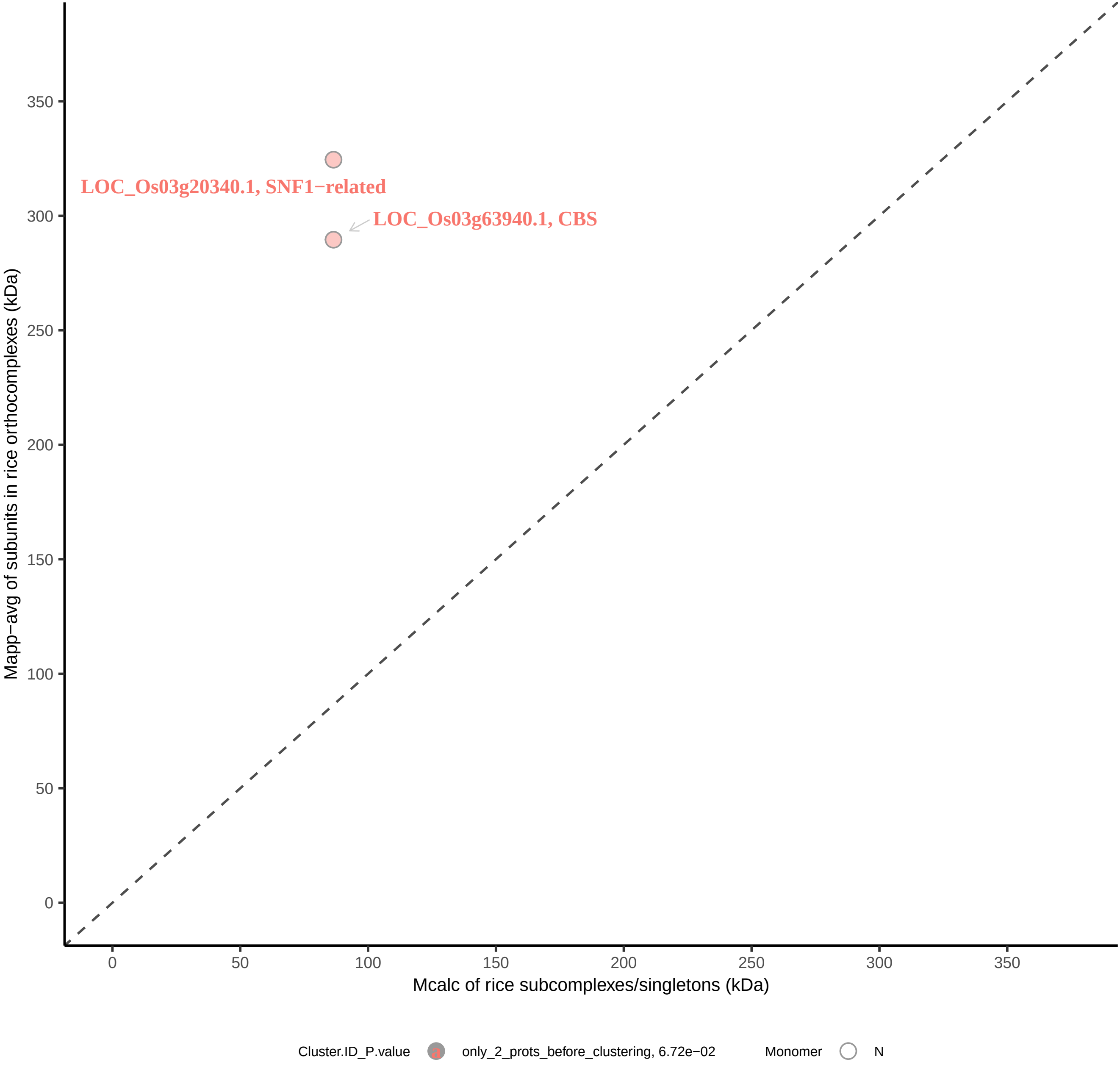

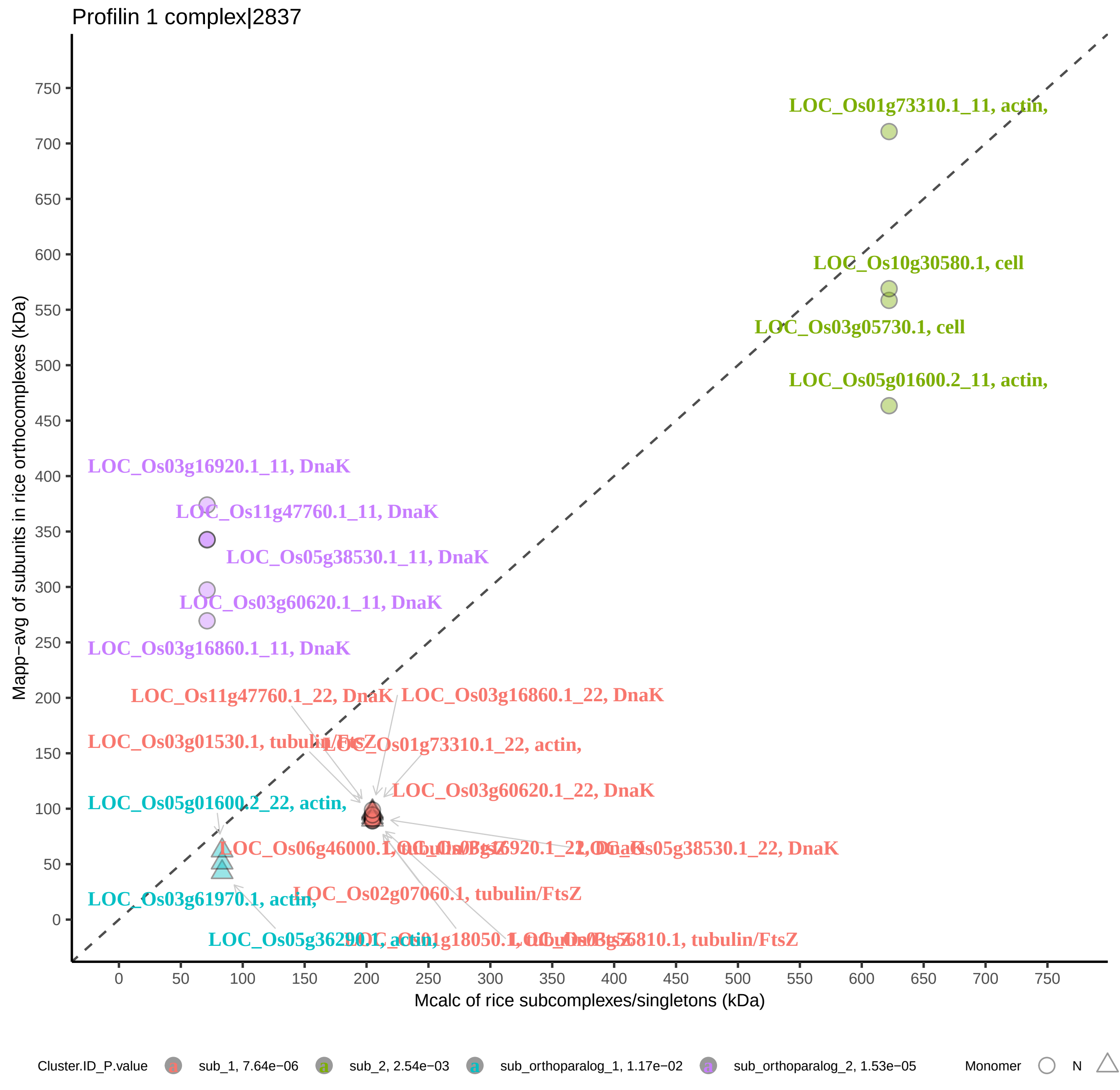

Profilin 1 complex|2837
LOC_Os01g73310.1_11, actin,
LOC_Os10g30580.1, cell
LOC_Os03g05730.1, cell
LOC_Os05g01600.2_11, actin,
LOC_Os03g16920.1_11, DnaK
LOC_Os11g47760.1_11, DnaK
LOC_Os05g38530.1_11, DnaK
LOC_Os03g60620.1_11, DnaK
LOC_Os03g16860.1_11, DnaK
LOC_Os11g47760.1_22, DnaK
LOC_Os03g16860.1_22, DnaK
LOC_Os03g01530.1, tubulin/FtsZ
LOC_Os01g73310.1_22, actin,
LOC_Os03g60620.1_22, DnaK
LOC_Os05g01600.2_22, actin,
LOC_Os06g46000.1, tubulin/FtsZ
LOC_Os03g16920.1_22, DnaK
LOC_Os05g38530.1_22, DnaK
LOC_Os02g07060.1, tubulin/FtsZ
LOC_Os03g61970.1, actin,
LOC_Os05g36290.1, actin,
LOC_Os01g18050.1, tubulin/FtsZ
LOC_Os03g56810.1, tubulin/FtsZ
Mapp−avg of subunits in rice orthocomplexes (kDa)
Mcalc of rice subcomplexes/singletons (kDa)
Cluster.ID_P.value
sub_1, 7.64e−06
sub_2, 2.54e−03
sub_orthoparalog_1, 1.17e−02
sub_orthoparalog_2, 1.53e−05
Monomer
N

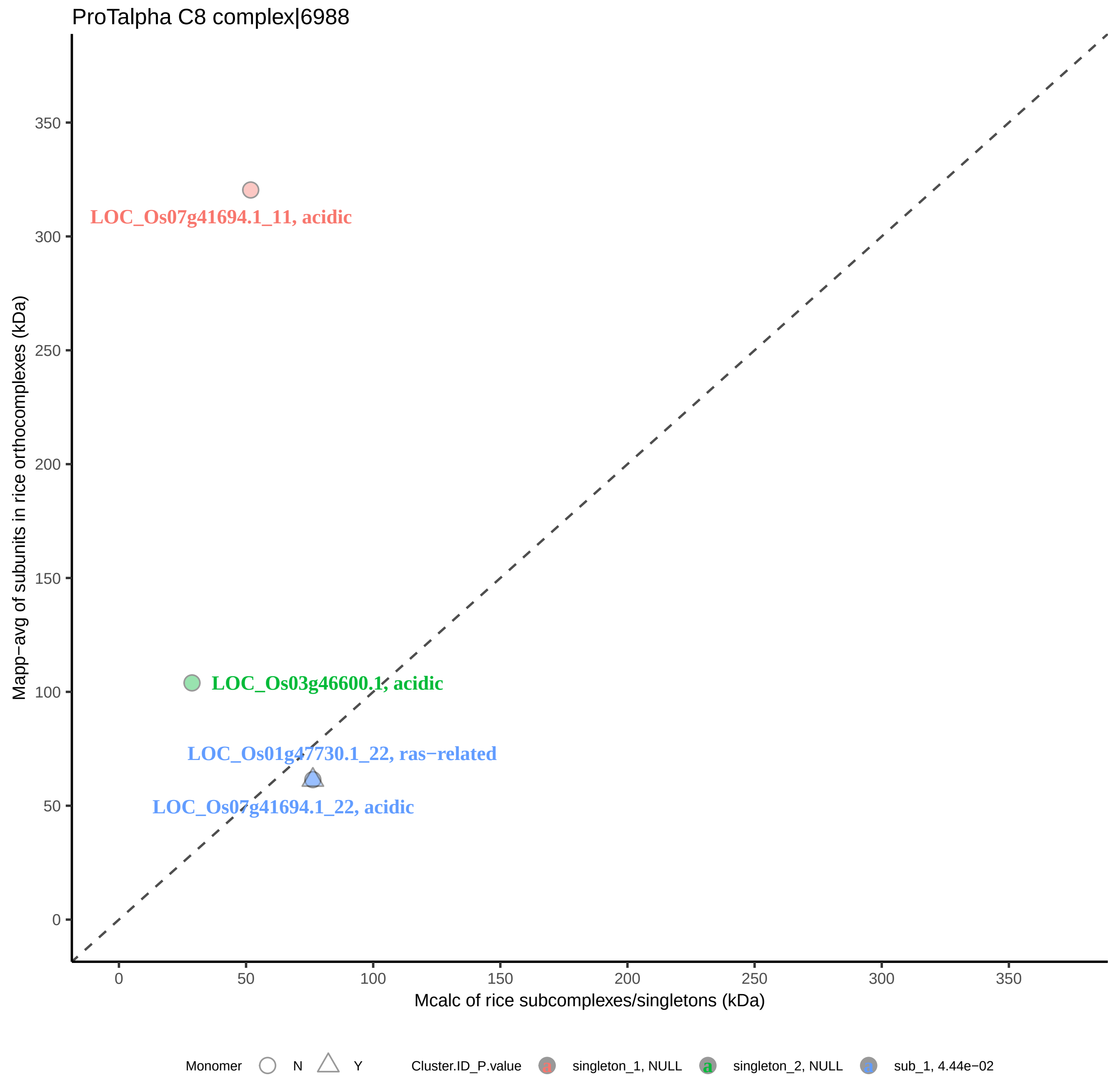

ProTalpha C8 complex|6988
LOC_Os07g41694.1_11, acidic
LOC_Os03g46600.1, acidic
LOC_Os01g47730.1_22, ras−related
LOC_Os07g41694.1_22, acidic
Mapp−avg of subunits in rice orthocomplexes (kDa)
Mcalc of rice subcomplexes/singletons (kDa)
0
50
100
150
200
250
300
350
Monomer
N
Y
Cluster.ID_P.value
singleton_1, NULL
singleton_2, NULL
sub_1, 4.44e−02

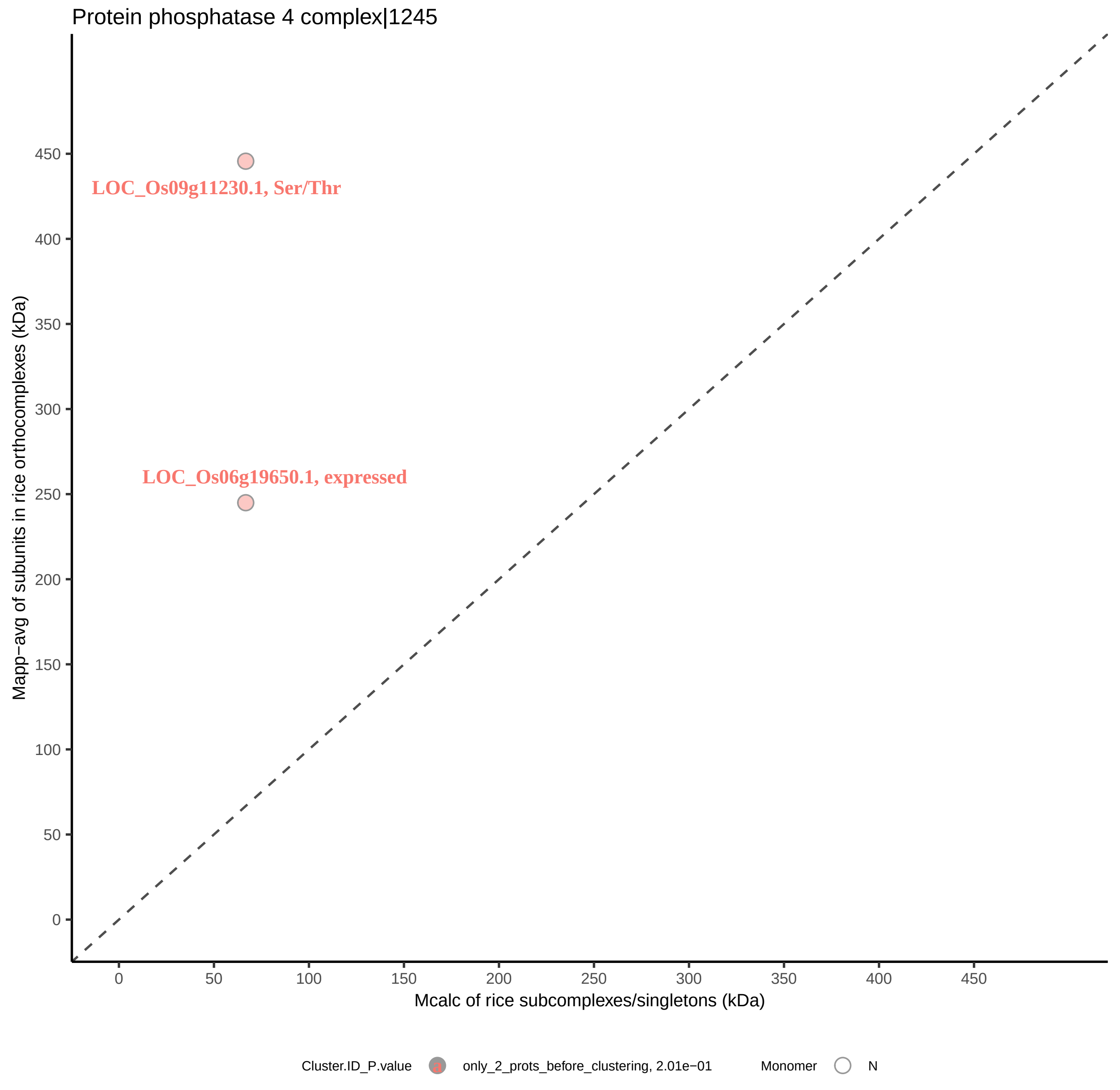

Protein phosphatase 4 complex|1245
LOC_Os09g11230.1, Ser/Thr
LOC_Os06g19650.1, expressed
Mapp−avg of subunits in rice orthocomplexes (kDa)
0
50
100
150
200
250
300
350
400
450
Mcalc of rice subcomplexes/singletons (kDa)
Cluster.ID_P.value
a
only_2_prots_before_clustering, 2.01e−01
Monomer
N

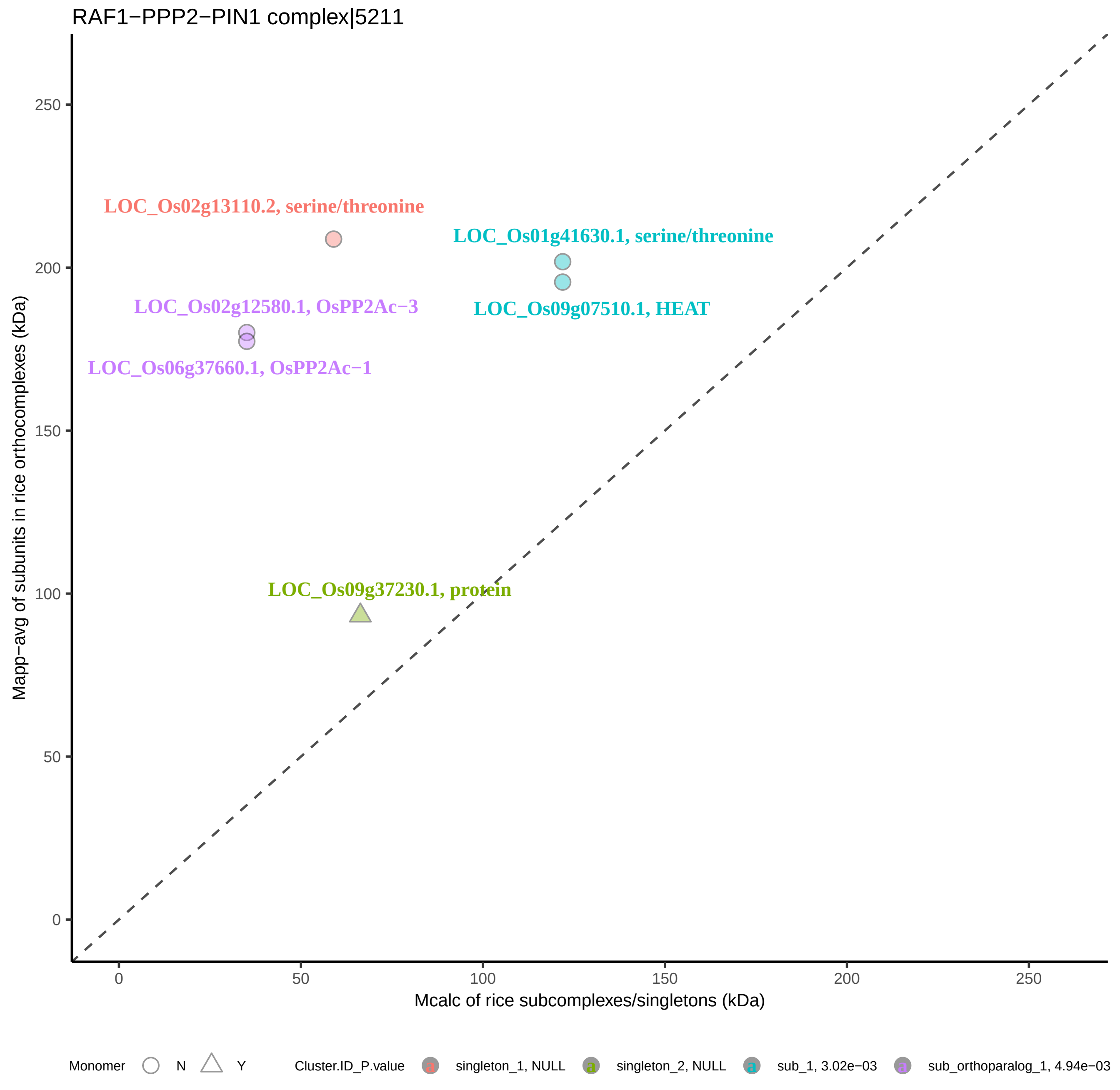

RAF1−PPP2−PIN1 complex|5211
LOC_Os02g13110.2, serine/threonine
LOC_Os01g41630.1, serine/threonine
LOC_Os02g12580.1, OsPP2Ac−3
LOC_Os09g07510.1, HEAT
LOC_Os06g37660.1, OsPP2Ac−1
LOC_Os09g37230.1, protein
Mapp−avg of subunits in rice orthocomplexes (kDa)
Mcalc of rice subcomplexes/singletons (kDa)
0
50
100
150
200
250
Monomer
N
Y
Cluster.ID_P.value
singleton_1, NULL
singleton_2, NULL
sub_1, 3.02e−03
sub_orthoparalog_1, 4.94e−03

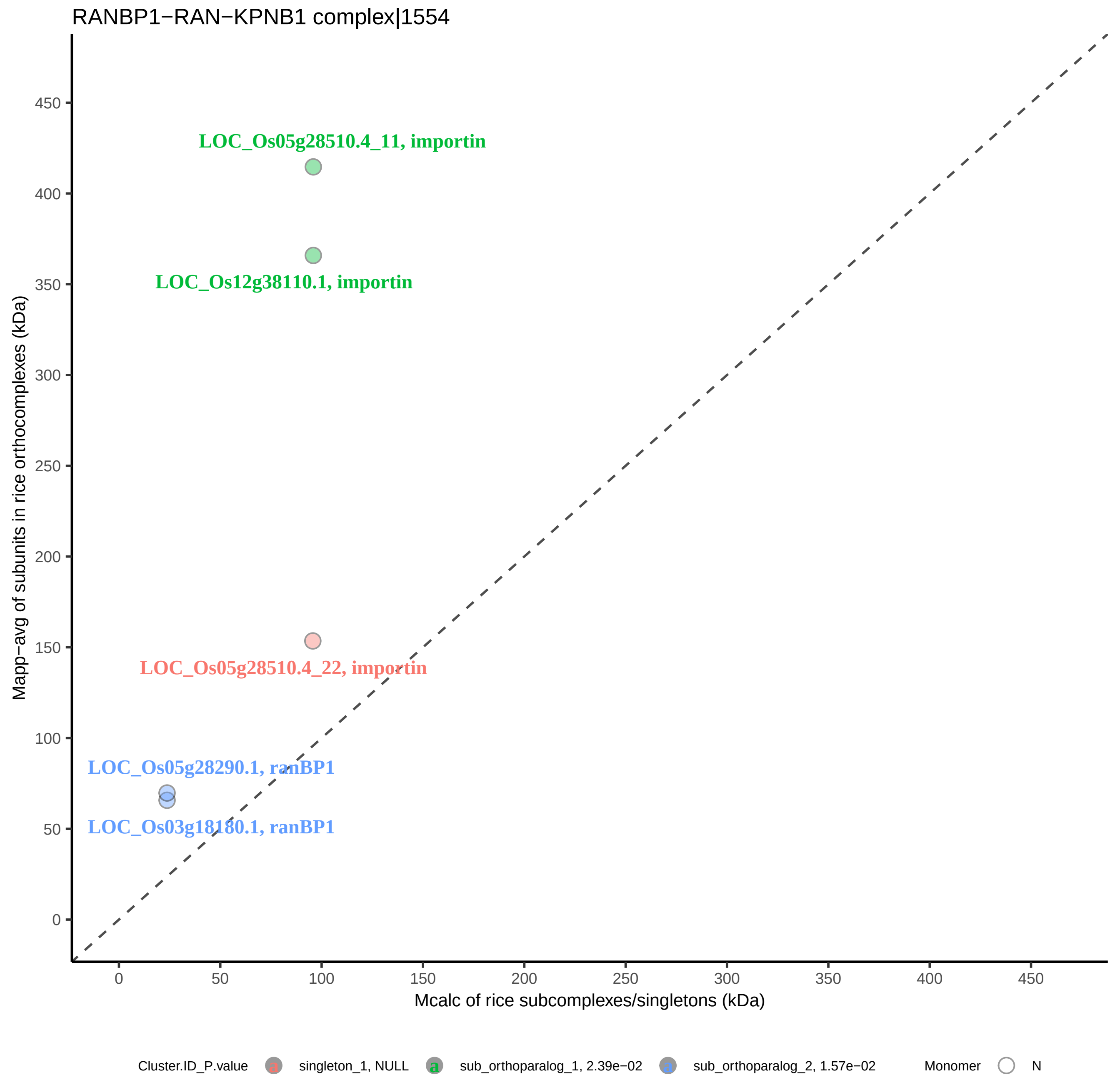
RANBP1−RAN−KPNB1 complex|1554
LOC_Os05g28510.4_11, importin
LOC_Os12g38110.1, importin
LOC_Os05g28510.4_22, importin
LOC_Os05g28290.1, ranBP1
LOC_Os03g18180.1, ranBP1
Mapp−avg of subunits in rice orthocomplexes (kDa)
Mcalc of rice subcomplexes/singletons (kDa)
0
50
100
150
200
250
300
350
400
450
Cluster.ID_P.value
singleton_1, NULL
sub_orthoparalog_1, 2.39e−02
sub_orthoparalog_2, 1.57e−02
Monomer
N

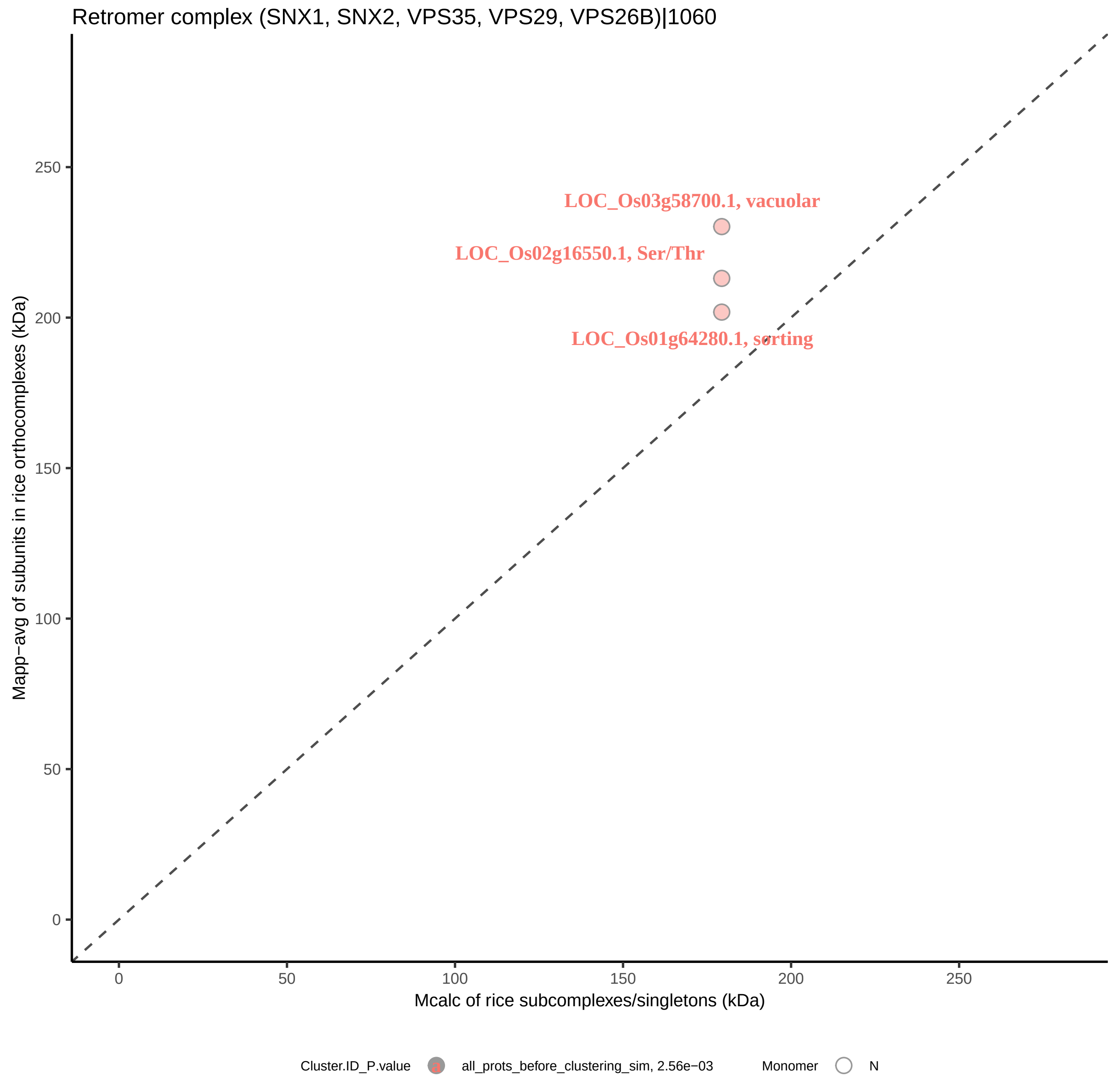

Retromer complex (SNX1, SNX2, VPS35, VPS29, VPS26B)|1060
LOC_Os03g58700.1, vacuolar
LOC_Os02g16550.1, Ser/Thr
LOC_Os01g64280.1, sorting
Mapp−avg of subunits in rice orthocomplexes (kDa)
0
50
100
150
200
250
Mcalc of rice subcomplexes/singletons (kDa)
0
50
100
150
200
250
Cluster.ID_P.value
all_prots_before_clustering_sim, 2.56e−03
Monomer
N

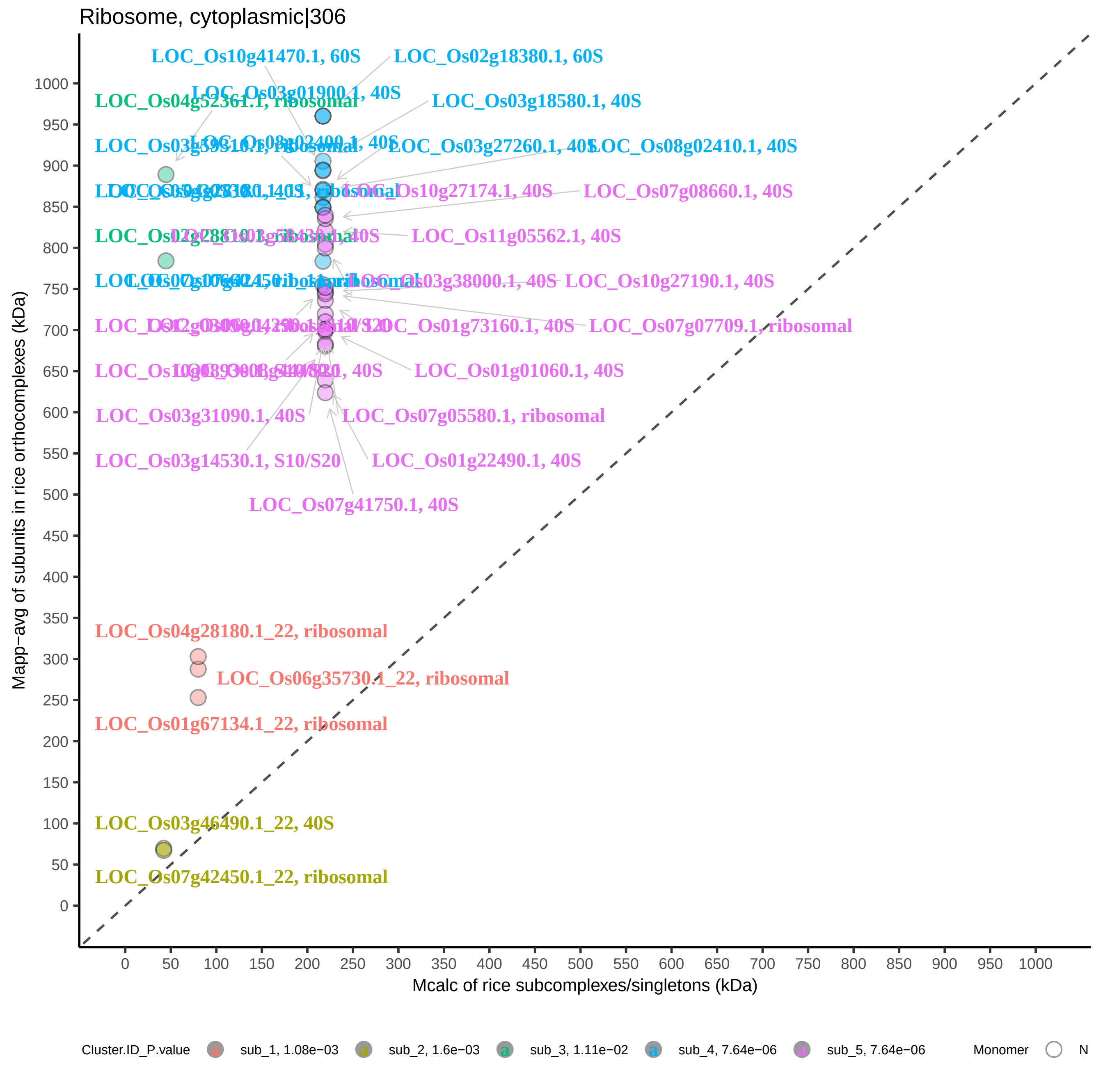

Ribosome, cytoplasmic|306
LOC_Os10g41470.1, 60S
LOC_Os02g18380.1, 60S
LOC_Os03g01900.1, 40S
LOC_Os04g52361.1, ribosomal
LOC_Os03g18580.1, 40S
LOC_Os08g02400.1, 40S
LOC_Os03g27260.1, 40S
LOC_Os08g02410.1, 40S
LOC_Os10g27174.1, 40S
LOC_Os07g08660.1, 40S
LOC_Os11g05562.1, 40S
LOC_Os03g38000.1, 40S
LOC_Os10g27190.1, 40S
LOC_Os01g73160.1, 40S
LOC_Os07g07709.1, ribosomal
LOC_Os01g01060.1, 40S
LOC_Os03g31090.1, 40S
LOC_Os07g05580.1, ribosomal
LOC_Os03g14530.1, S10/S20
LOC_Os01g22490.1, 40S
LOC_Os07g41750.1, 40S
LOC_Os04g28180.1_22, ribosomal
LOC_Os06g35730.1_22, ribosomal
LOC_Os01g67134.1_22, ribosomal
LOC_Os03g46490.1_22, 40S
LOC_Os07g42450.1_22, ribosomal
Mapp−avg of subunits in rice orthocomplexes (kDa)
Mcalc of rice subcomplexes/singletons (kDa)
Cluster.ID_P.value
sub_1, 1.08e−03
sub_2, 1.6e−03
sub_3, 1.11e−02
sub_4, 7.64e−06
sub_5, 7.64e−06
Monomer
N

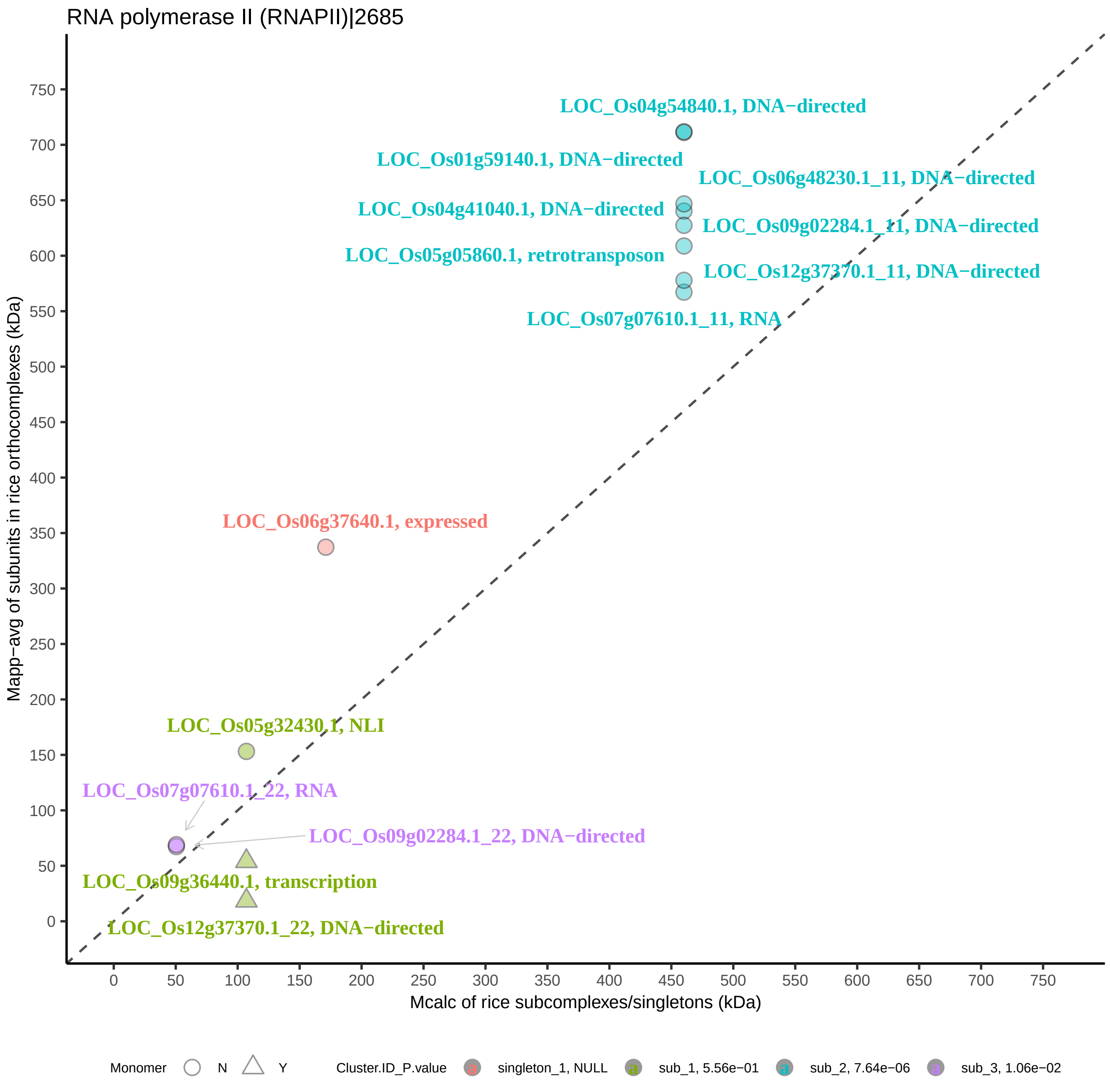

RNA polymerase II (RNAPII)|2685
Mapp−avg of subunits in rice orthocomplexes (kDa)
Mcalc of rice subcomplexes/singletons (kDa)
LOC_Os04g54840.1, DNA−directed
LOC_Os01g59140.1, DNA−directed
LOC_Os06g48230.1_11, DNA−directed
LOC_Os04g41040.1, DNA−directed
LOC_Os09g02284.1_11, DNA−directed
LOC_Os05g05860.1, retrotransposon
LOC_Os12g37370.1_11, DNA−directed
LOC_Os07g07610.1_11, RNA
LOC_Os06g37640.1, expressed
LOC_Os05g32430.1, NLI
LOC_Os07g07610.1_22, RNA
LOC_Os09g02284.1_22, DNA−directed
LOC_Os09g36440.1, transcription
LOC_Os12g37370.1_22, DNA−directed
Monomer N Y
Cluster.ID_P.value singleton_1, NULL sub_1, 5.56e−01 sub_2, 7.64e−06 sub_3, 1.06e−02

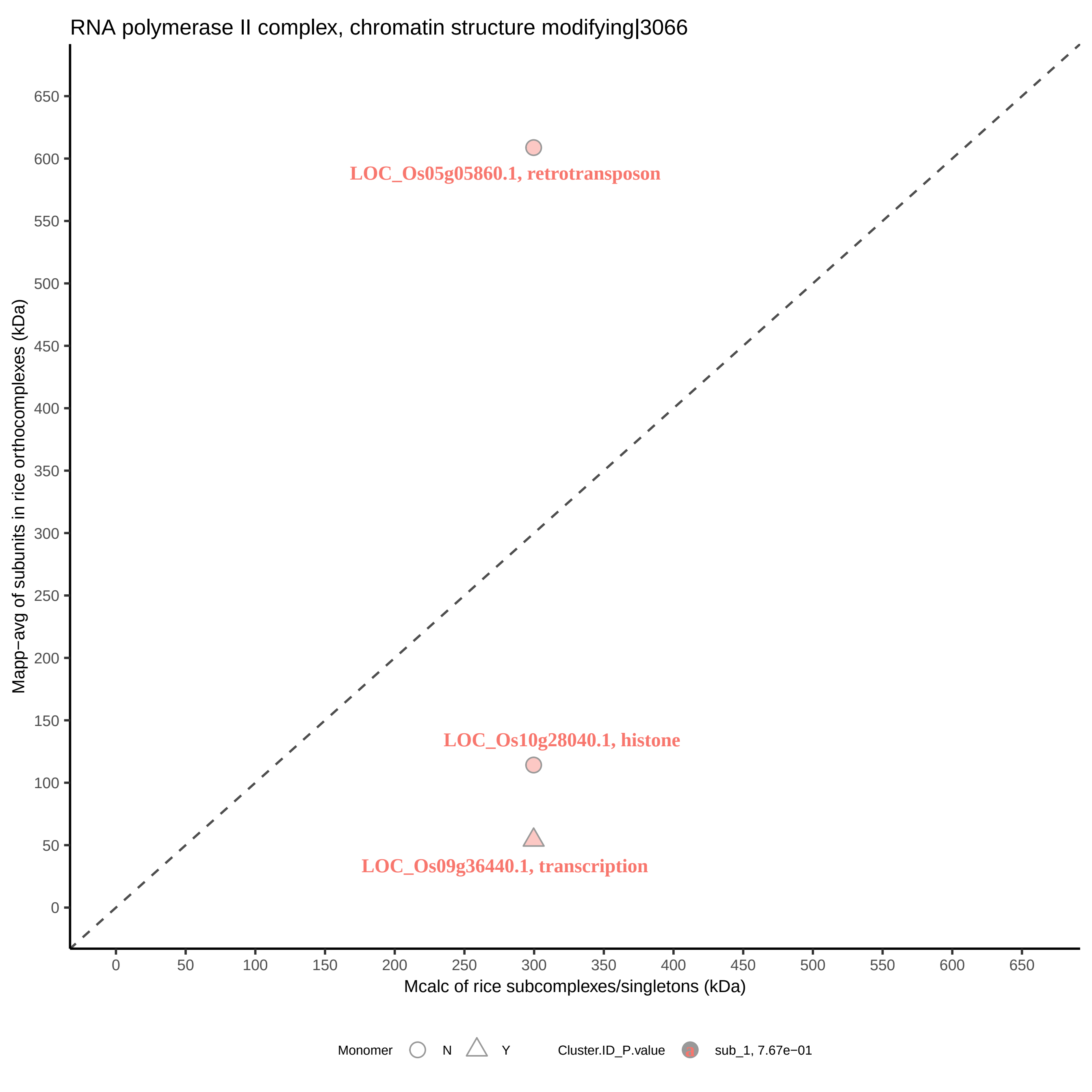
RNA polymerase II complex, chromatin structure modifying|3066
Mapp−avg of subunits in rice orthocomplexes (kDa)
Mcalc of rice subcomplexes/singletons (kDa)
LOC_Os05g05860.1, retrotransposon
LOC_Os10g28040.1, histone
LOC_Os09g36440.1, transcription
0
50
100
150
200
250
300
350
400
450
500
550
600
650
Monomer
N
Y
Cluster.ID_P.value
sub_1, 7.67e−01

RNA−induced silencing complex, RISC|3032

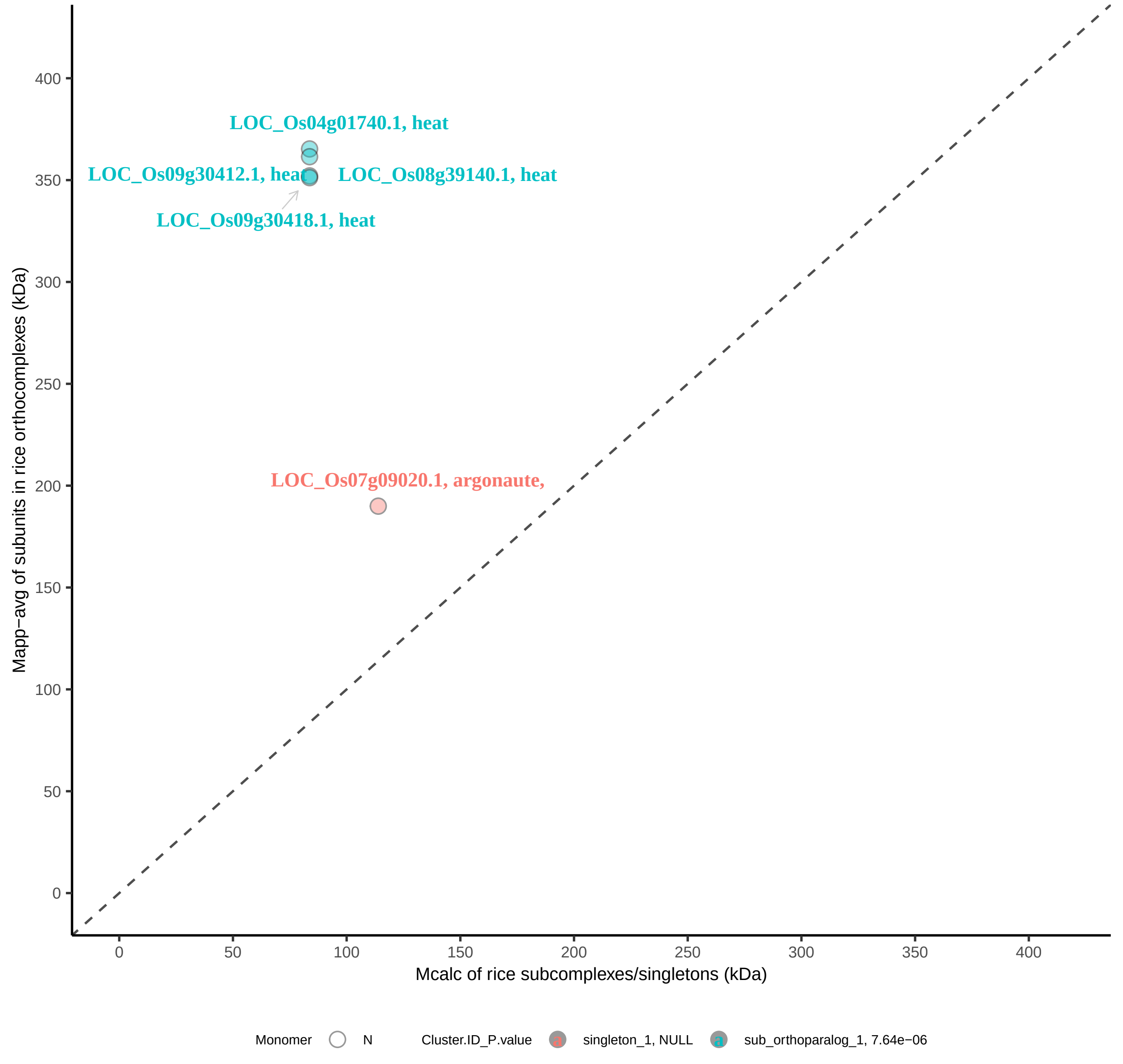

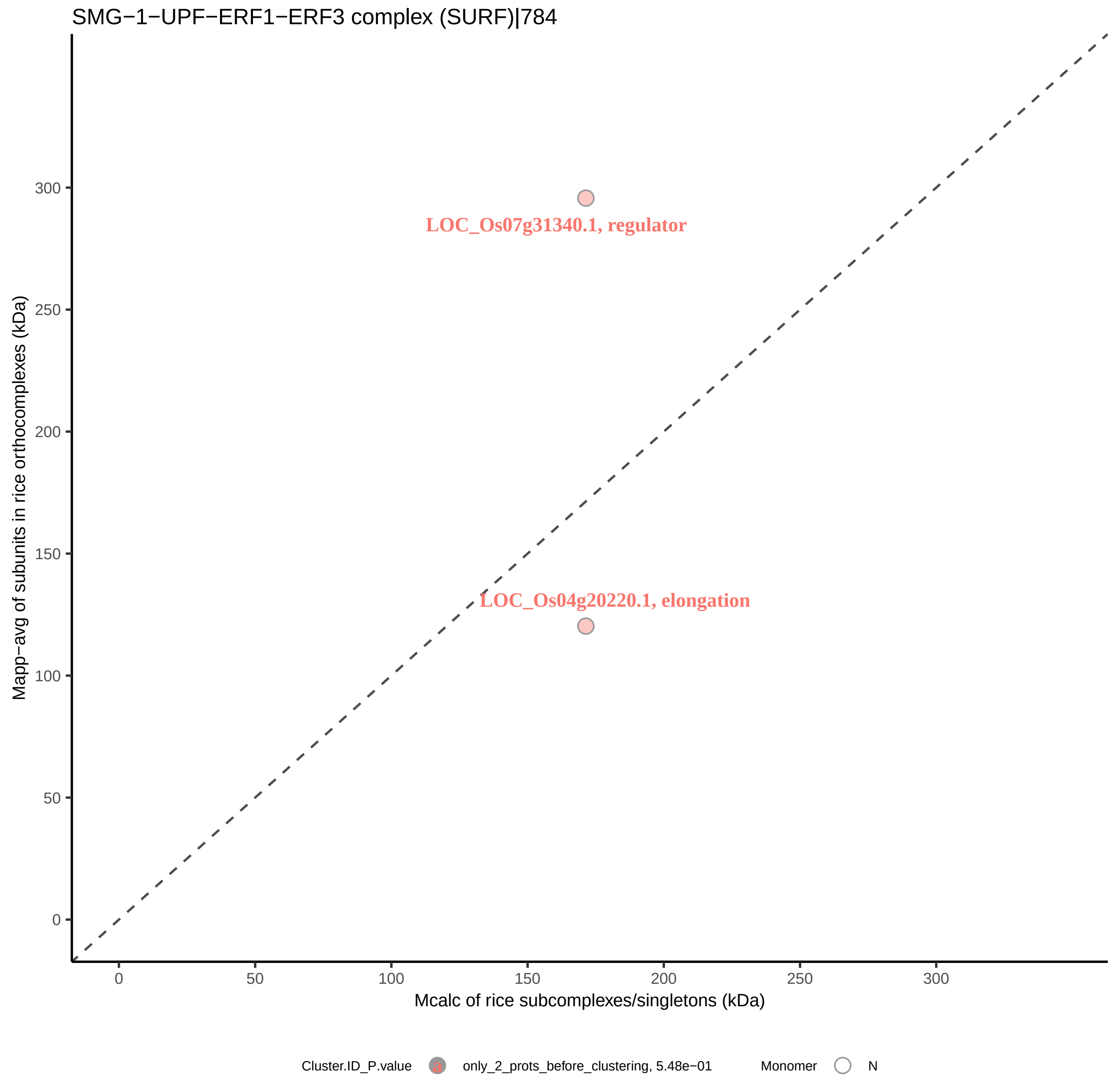

SMG−1−UPF−ERF1−ERF3 complex (SURF)|784
LOC_Os07g31340.1, regulator
LOC_Os04g20220.1, elongation
Mapp−avg of subunits in rice orthocomplexes (kDa)
Mcalc of rice subcomplexes/singletons (kDa)
0
50
100
150
200
250
300
Cluster.ID_P.value
only_2_prots_before_clustering, 5.48e−01
Monomer
N

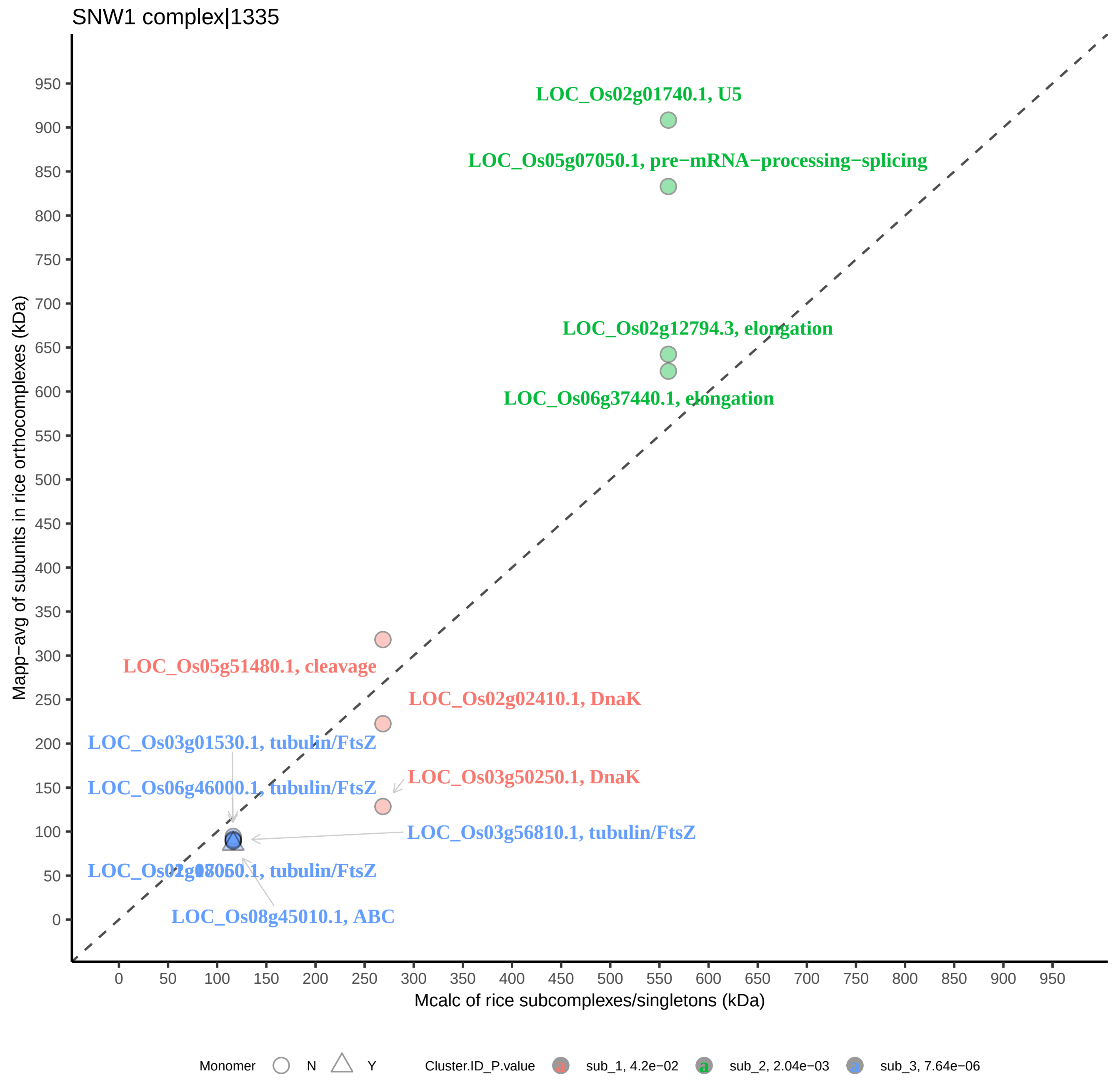
SNW1 complex|1335
LOC_Os02g01740.1, U5
LOC_Os05g07050.1, pre−mRNA−processing−splicing
LOC_Os02g12794.3, elongation
LOC_Os06g37440.1, elongation
LOC_Os05g51480.1, cleavage
LOC_Os02g02410.1, DnaK
LOC_Os03g01530.1, tubulin/FtsZ
LOC_Os06g46000.1, tubulin/FtsZ
LOC_Os03g50250.1, DnaK
LOC_Os03g56810.1, tubulin/FtsZ
LOC_Os08g45010.1, ABC
Mapp−avg of subunits in rice orthocomplexes (kDa)
Mcalc of rice subcomplexes/singletons (kDa)
0
50
100
150
200
250
300
350
400
450
500
550
600
650
700
750
800
850
900
950
Monomer
N
Y
Cluster.ID_P.value
sub_1, 4.2e−02
sub_2, 2.04e−03
sub_3, 7.64e−06

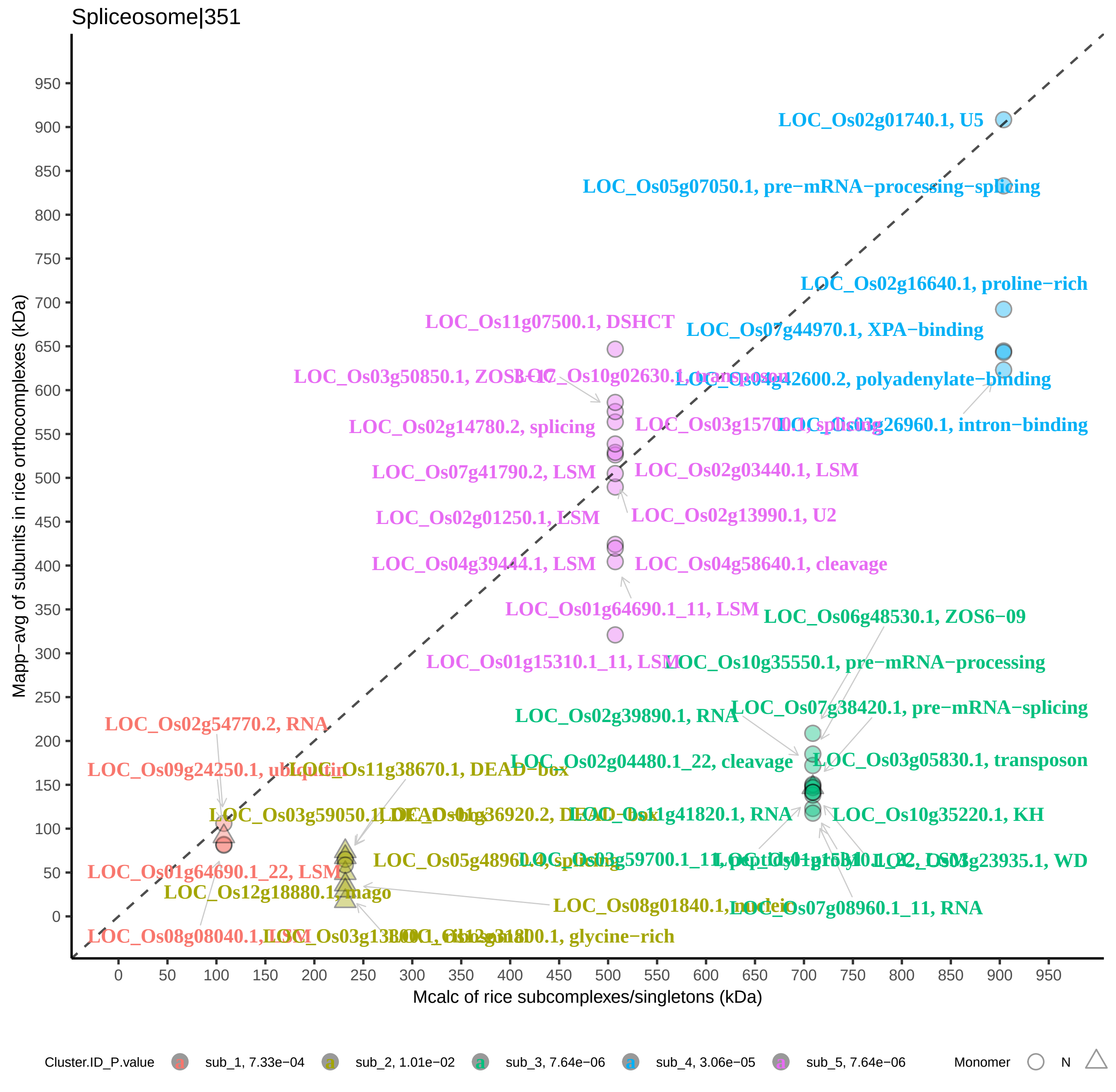
Spliceosome|351
Mapp−avg of subunits in rice orthocomplexes (kDa)
Mcalc of rice subcomplexes/singletons (kDa)
0
50
100
150
200
250
300
350
400
450
500
550
600
650
700
750
800
850
900
950
LOC_Os02g01740.1, U5
LOC_Os05g07050.1, pre−mRNA−processing−splicing
LOC_Os02g16640.1, proline−rich
LOC_Os07g44970.1, XPA−binding
LOC_Os04g42600.2, polyadenylate−binding
LOC_Os03g26960.1, intron−binding
LOC_Os11g07500.1, DSHCT
LOC_Os02g14780.2, splicing
LOC_Os07g41790.2, LSM
LOC_Os02g03440.1, LSM
LOC_Os02g13990.1, U2
LOC_Os02g01250.1, LSM
LOC_Os04g39444.1, LSM
LOC_Os04g58640.1, cleavage
LOC_Os01g64690.1_11, LSM
LOC_Os06g48530.1, ZOS6−09
LOC_Os10g35550.1, pre−mRNA−processing
LOC_Os07g38420.1, pre−mRNA−splicing
LOC_Os02g39890.1, RNA
LOC_Os02g04480.1_22, cleavage
LOC_Os03g05830.1, transposon
LOC_Os10g35220.1, KH
LOC_Os07g08960.1_11, RNA
LOC_Os02g54770.2, RNA
LOC_Os01g64690.1_22, LSM
LOC_Os12g18880.1, mago
LOC_Os08g01840.1, nuclear
glycine−rich
Cluster.ID_P.value
sub_1, 7.33e−04
sub_2, 1.01e−02
sub_3, 7.64e−06
sub_4, 3.06e−05
sub_5, 7.64e−06
Monomer
N

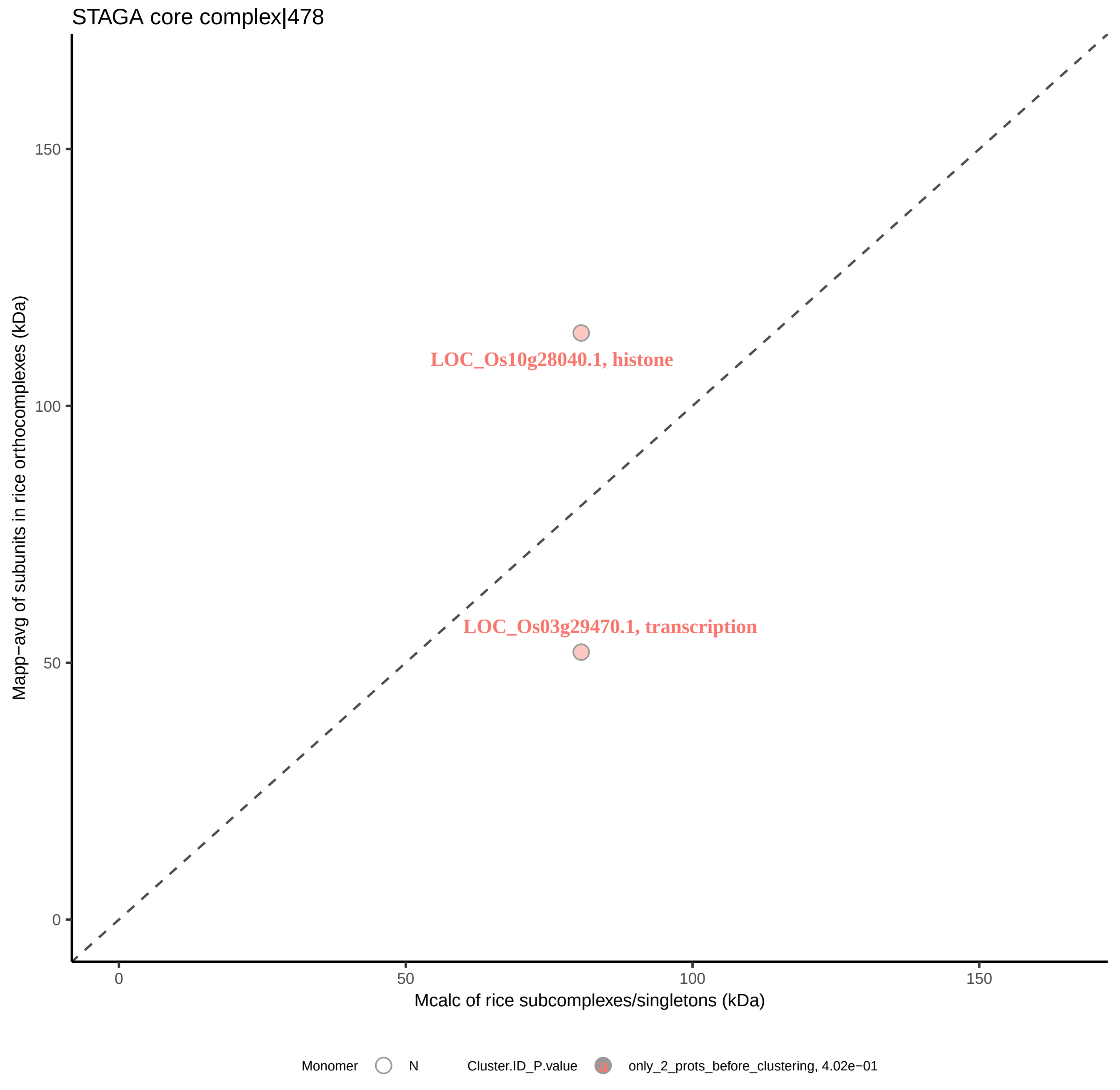
STAGA core complex|478
Mapp−avg of subunits in rice orthocomplexes (kDa)
150
100
50
0
LOC_Os10g28040.1, histone
LOC_Os03g29470.1, transcription
0
50
100
150
Mcalc of rice subcomplexes/singletons (kDa)
Monomer
N
Cluster.ID_P.value
only_2_prots_before_clustering, 4.02e−01

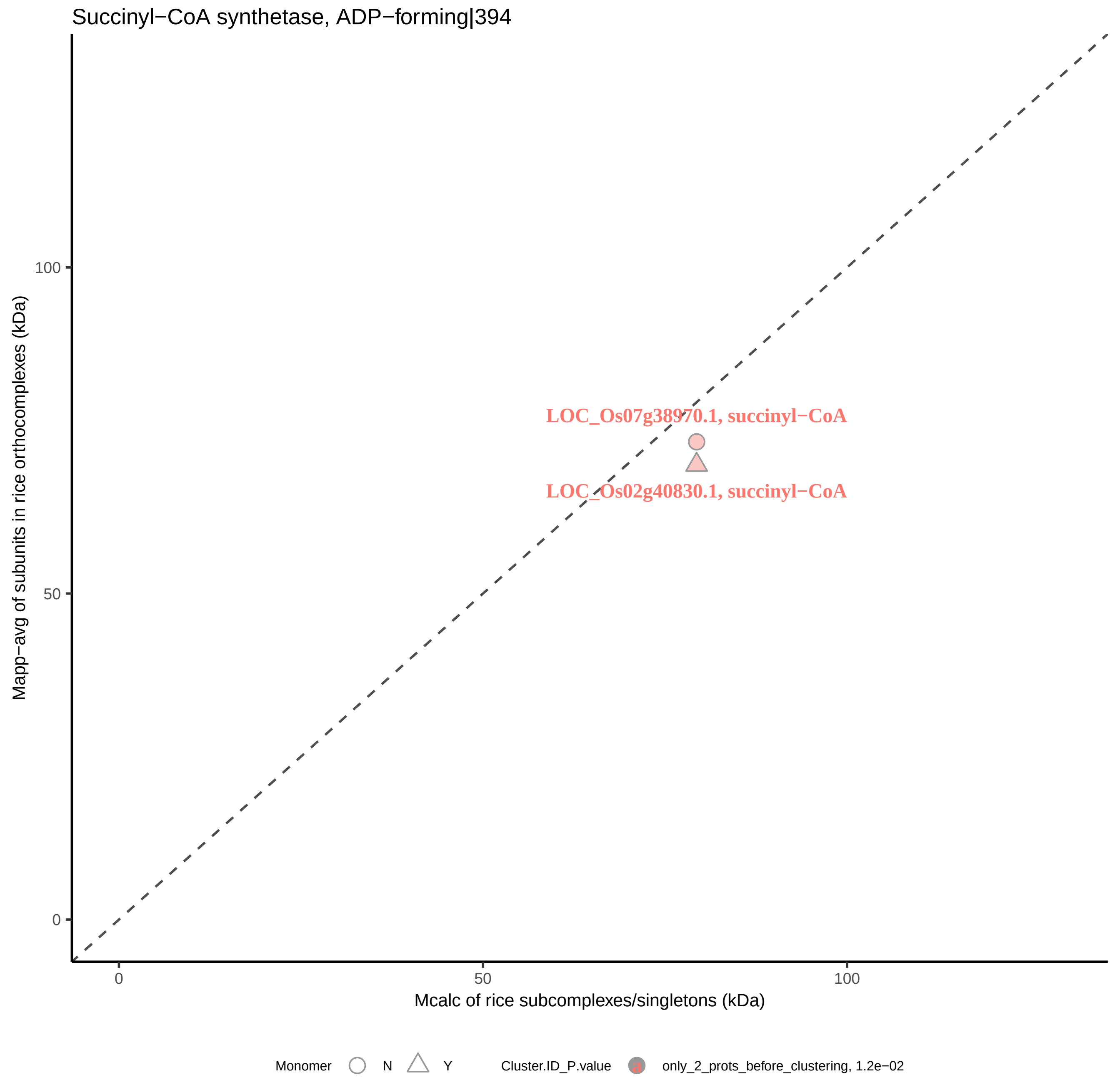

Succinyl−CoA synthetase, ADP−forming|394
Mapp−avg of subunits in rice orthocomplexes (kDa)
100
50
0
LOC_Os07g38970.1, succinyl−CoA
LOC_Os02g40830.1, succinyl−CoA
0
50
100
Mcalc of rice subcomplexes/singletons (kDa)
Monomer
N
Y
Cluster.ID_P.value
a
only_2_prots_before_clustering, 1.2e−02

TBCD−ARL2−tubulin(beta)−TBCE complex|6715

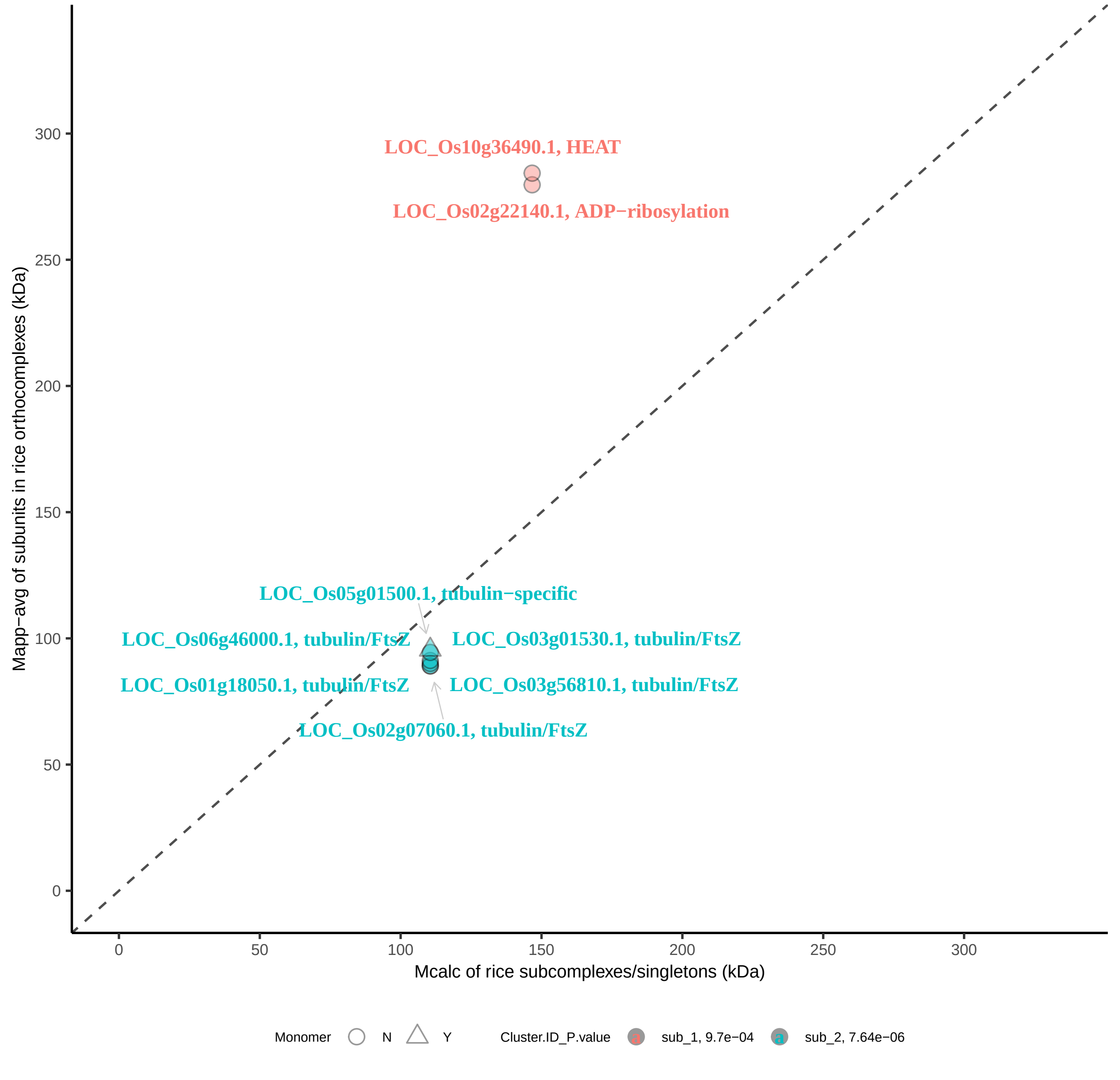

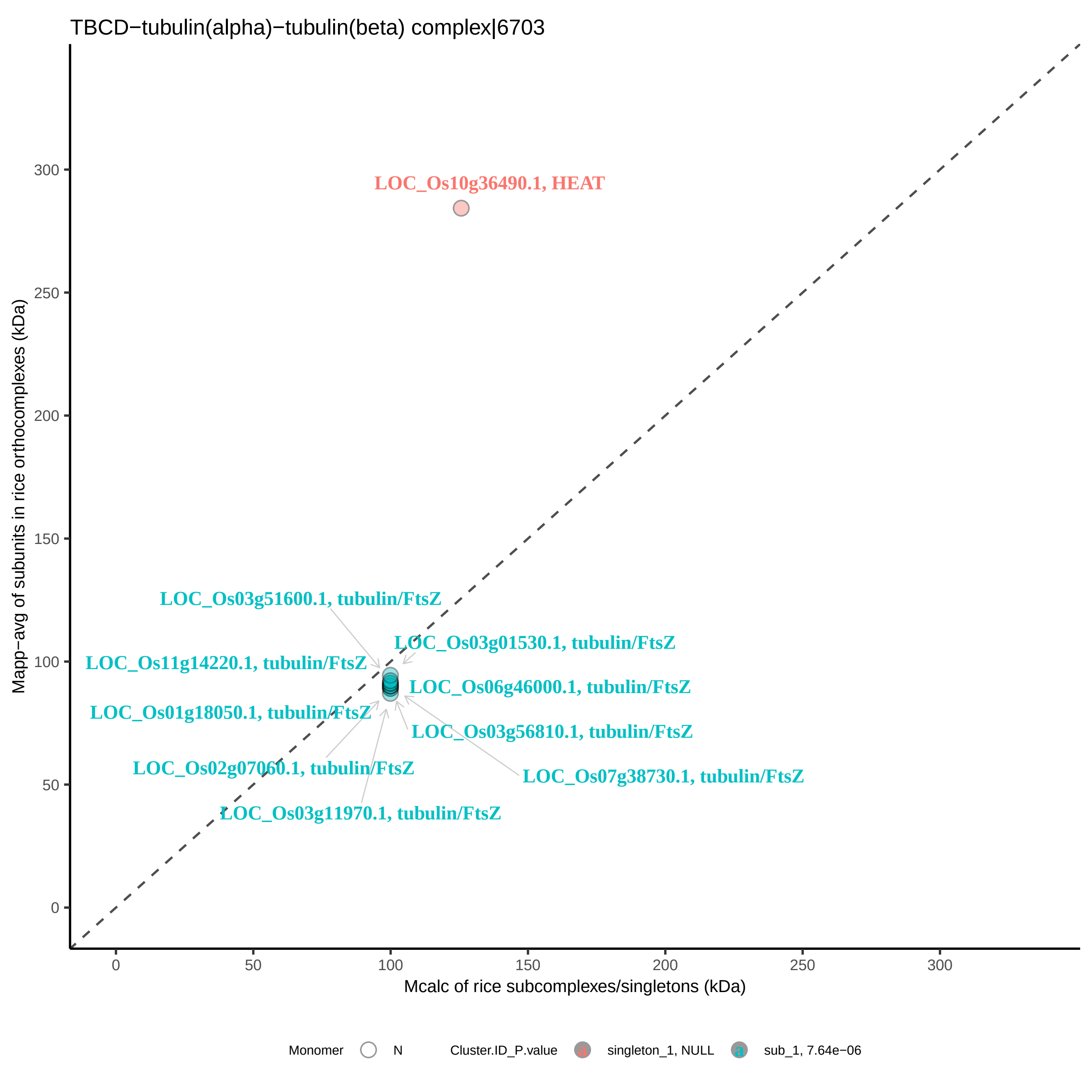

TBCD−tubulin(alpha)−tubulin(beta) complex|6703
LOC_Os10g36490.1, HEAT
LOC_Os03g51600.1, tubulin/FtsZ
LOC_Os03g01530.1, tubulin/FtsZ
LOC_Os11g14220.1, tubulin/FtsZ
LOC_Os06g46000.1, tubulin/FtsZ
LOC_Os01g18050.1, tubulin/FtsZ
LOC_Os03g56810.1, tubulin/FtsZ
LOC_Os02g07060.1, tubulin/FtsZ
LOC_Os07g38730.1, tubulin/FtsZ
LOC_Os03g11970.1, tubulin/FtsZ
Mapp−avg of subunits in rice orthocomplexes (kDa)
Mcalc of rice subcomplexes/singletons (kDa)
0
50
100
150
200
250
300
Monomer
N
Cluster.ID_P.value
singleton_1, NULL
sub_1, 7.64e−06

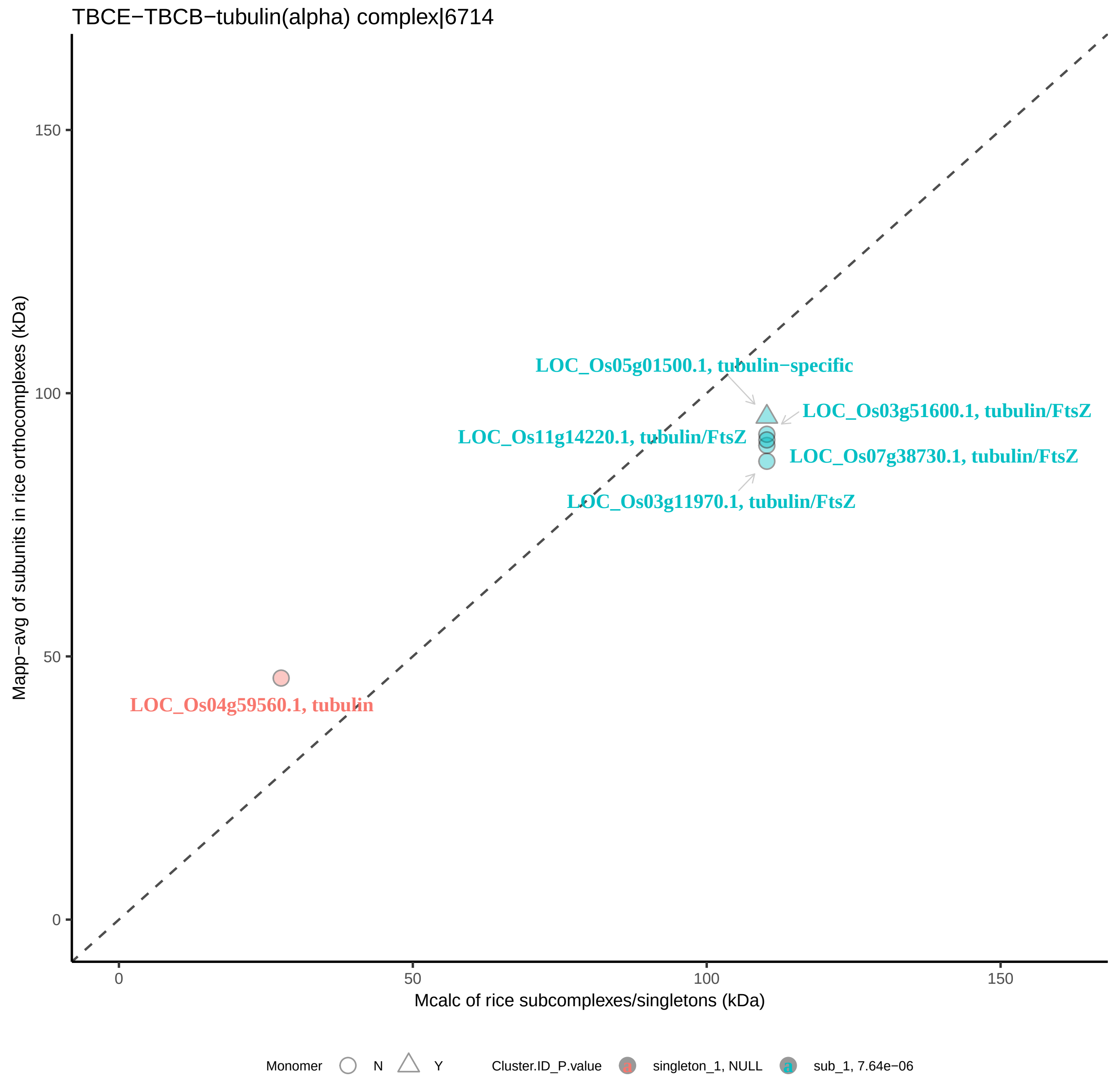

TBCE−TBCB−tubulin(alpha) complex|6714
Mapp−avg of subunits in rice orthocomplexes (kDa)
Mcalc of rice subcomplexes/singletons (kDa)
0
50
100
150
LOC_Os05g01500.1, tubulin−specific
LOC_Os03g51600.1, tubulin/FtsZ
LOC_Os11g14220.1, tubulin/FtsZ
LOC_Os07g38730.1, tubulin/FtsZ
LOC_Os03g11970.1, tubulin/FtsZ
LOC_Os04g59560.1, tubulin
Monomer
N
Y
Cluster.ID_P.value
singleton_1, NULL
sub_1, 7.64e−06

TFIIH transcription factor complex|107

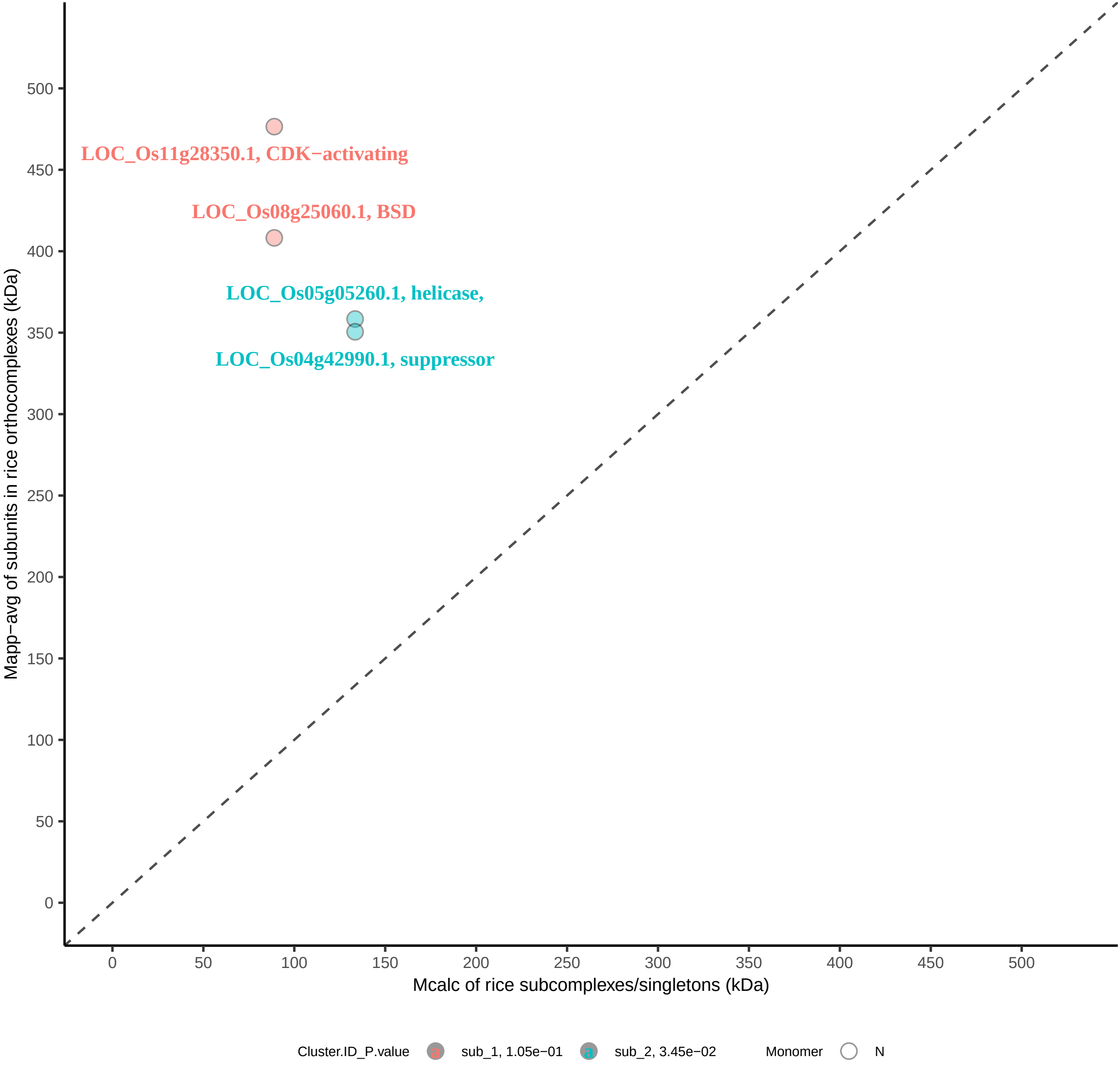

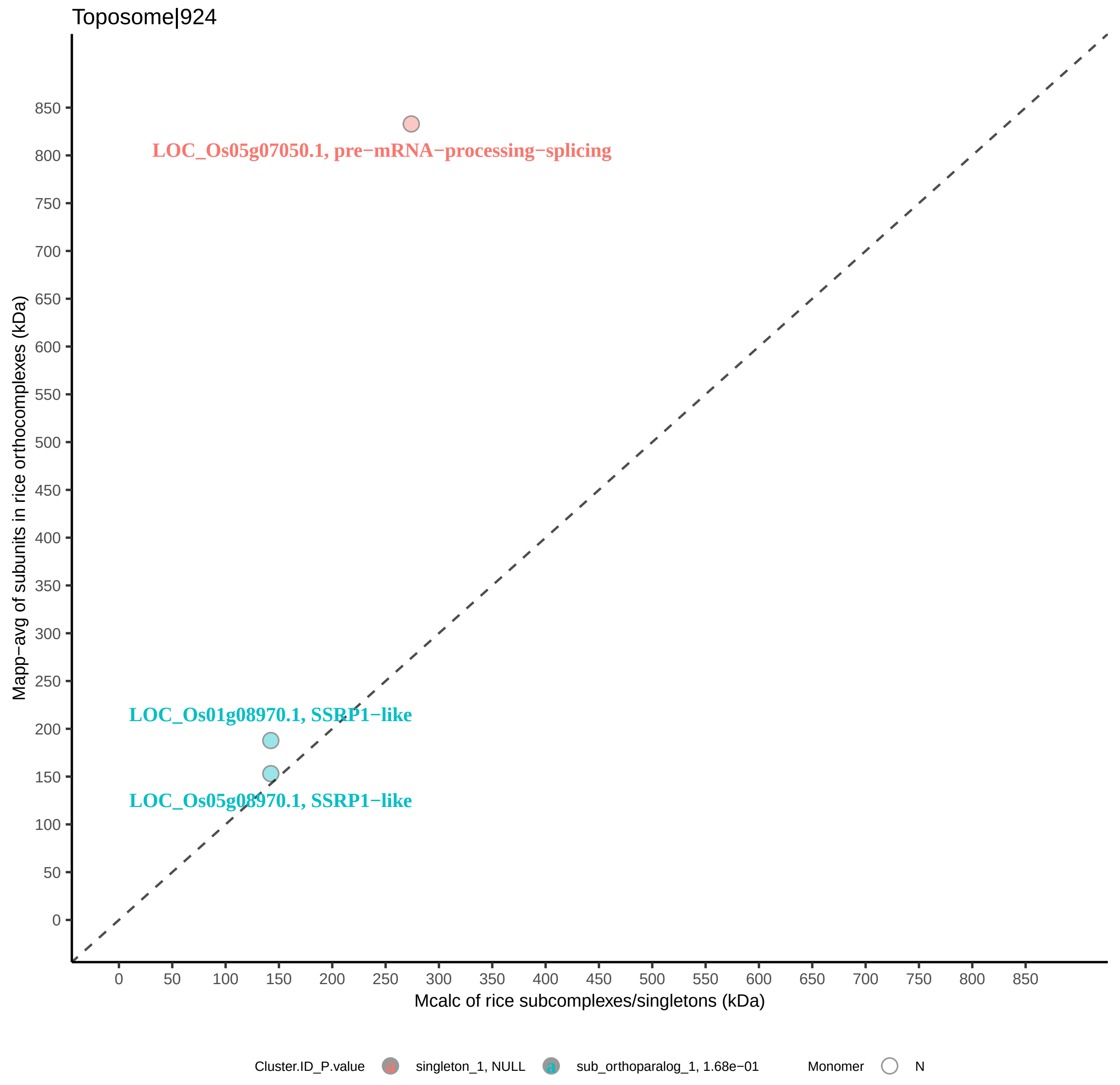

Toposome|924
LOC_Os05g07050.1, pre−mRNA−processing−splicing
LOC_Os01g08970.1, SSRP1−like
LOC_Os05g08970.1, SSRP1−like
Mapp−avg of subunits in rice orthocomplexes (kDa)
Mcalc of rice subcomplexes/singletons (kDa)
0
50
100
150
200
250
300
350
400
450
500
550
600
650
700
750
800
850
Cluster.ID_P.value
singleton_1, NULL
sub_orthoparalog_1, 1.68e−01
Monomer
N

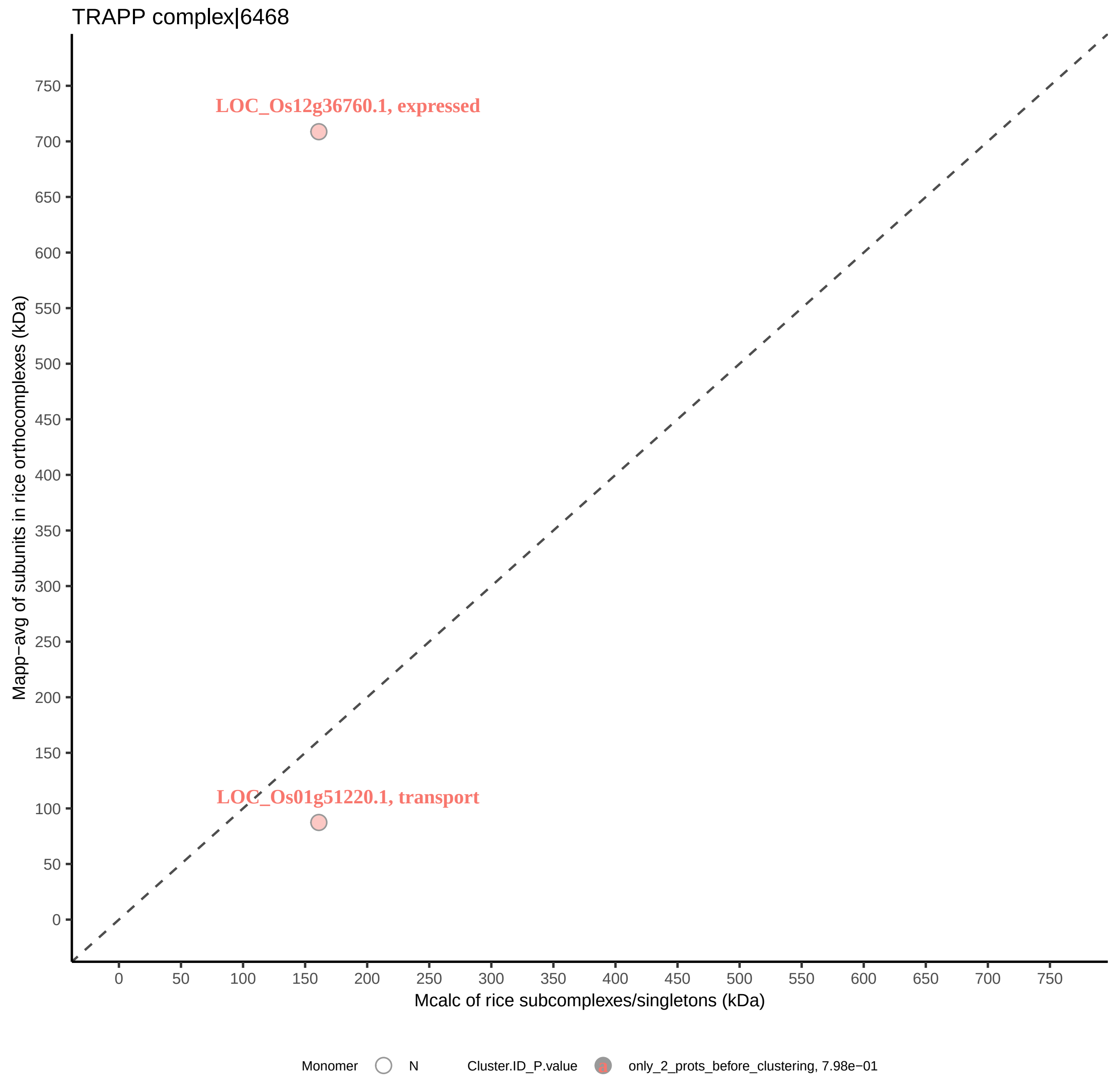
TRAPP complex|6468
LOC_Os12g36760.1, expressed
LOC_Os01g51220.1, transport
Mapp−avg of subunits in rice orthocomplexes (kDa)
Mcalc of rice subcomplexes/singletons (kDa)
0
50
100
150
200
250
300
350
400
450
500
550
600
650
700
750
Monomer
N
Cluster.ID_P.value
only_2_prots_before_clustering, 7.98e−01

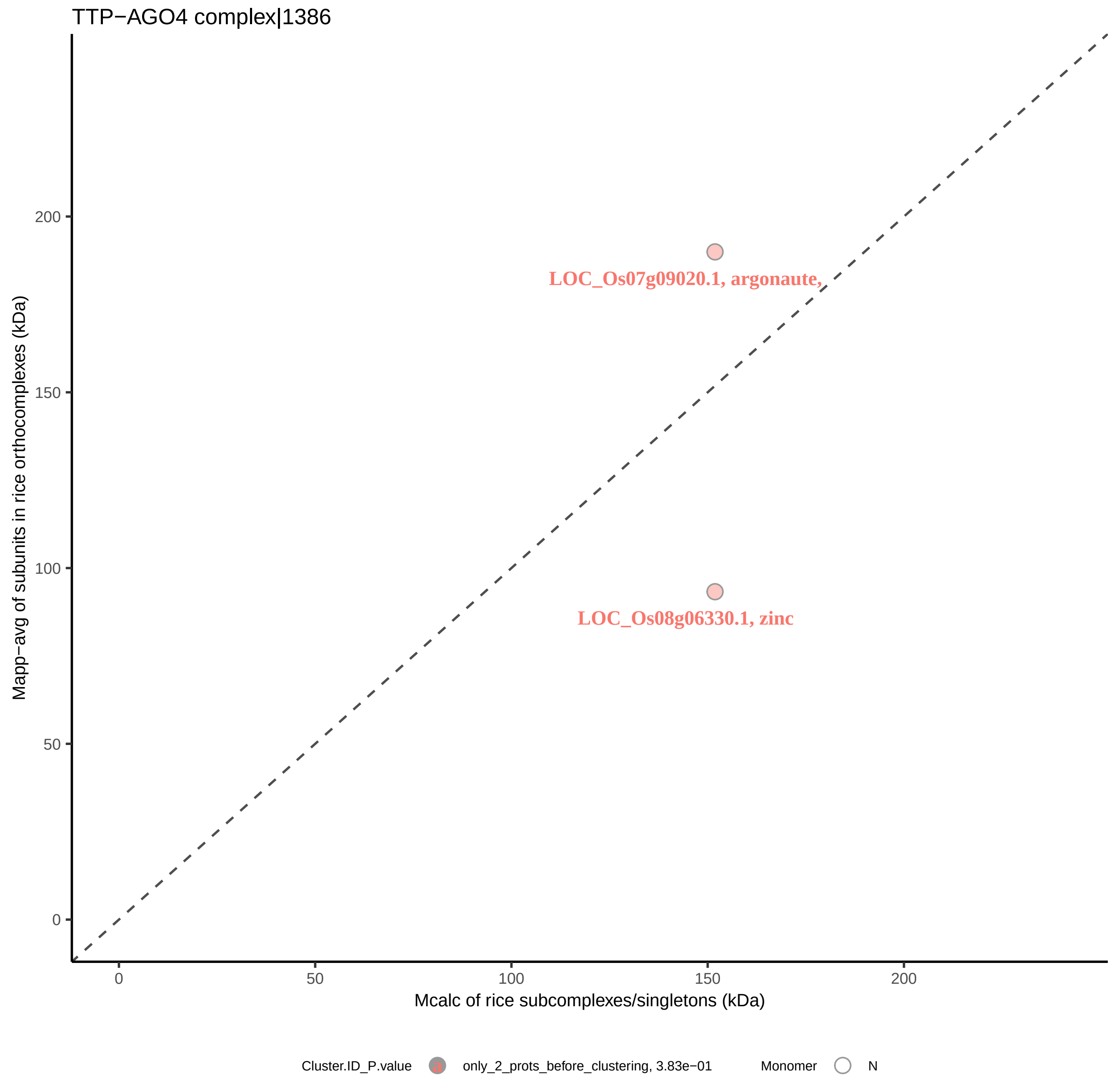

TTP−AGO4 complex|1386
Mapp−avg of subunits in rice orthocomplexes (kDa)
200
150
100
50
0
LOC_Os07g09020.1, argonaute,
LOC_Os08g06330.1, zinc
0
50
100
150
200
Mcalc of rice subcomplexes/singletons (kDa)
Cluster.ID_P.value
a
only_2_prots_before_clustering, 3.83e−01
Monomer
N

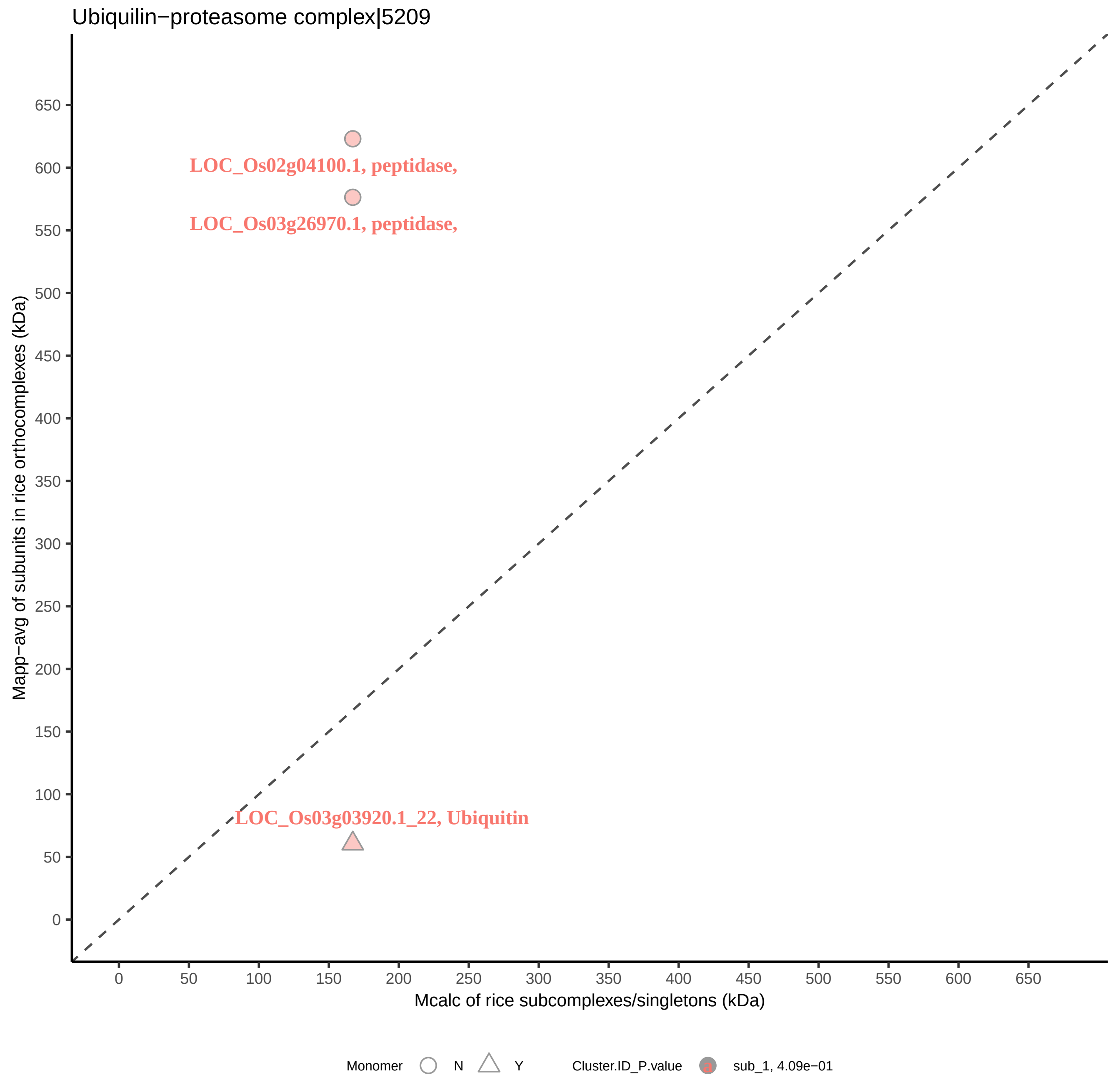

Ubiquilin−proteasome complex|5209
LOC_Os02g04100.1, peptidase,
LOC_Os03g26970.1, peptidase,
LOC_Os03g03920.1_22, Ubiquitin
Mapp−avg of subunits in rice orthocomplexes (kDa)
Mcalc of rice subcomplexes/singletons (kDa)
Monomer
N
Y
Cluster.ID_P.value
sub_1, 4.09e−01

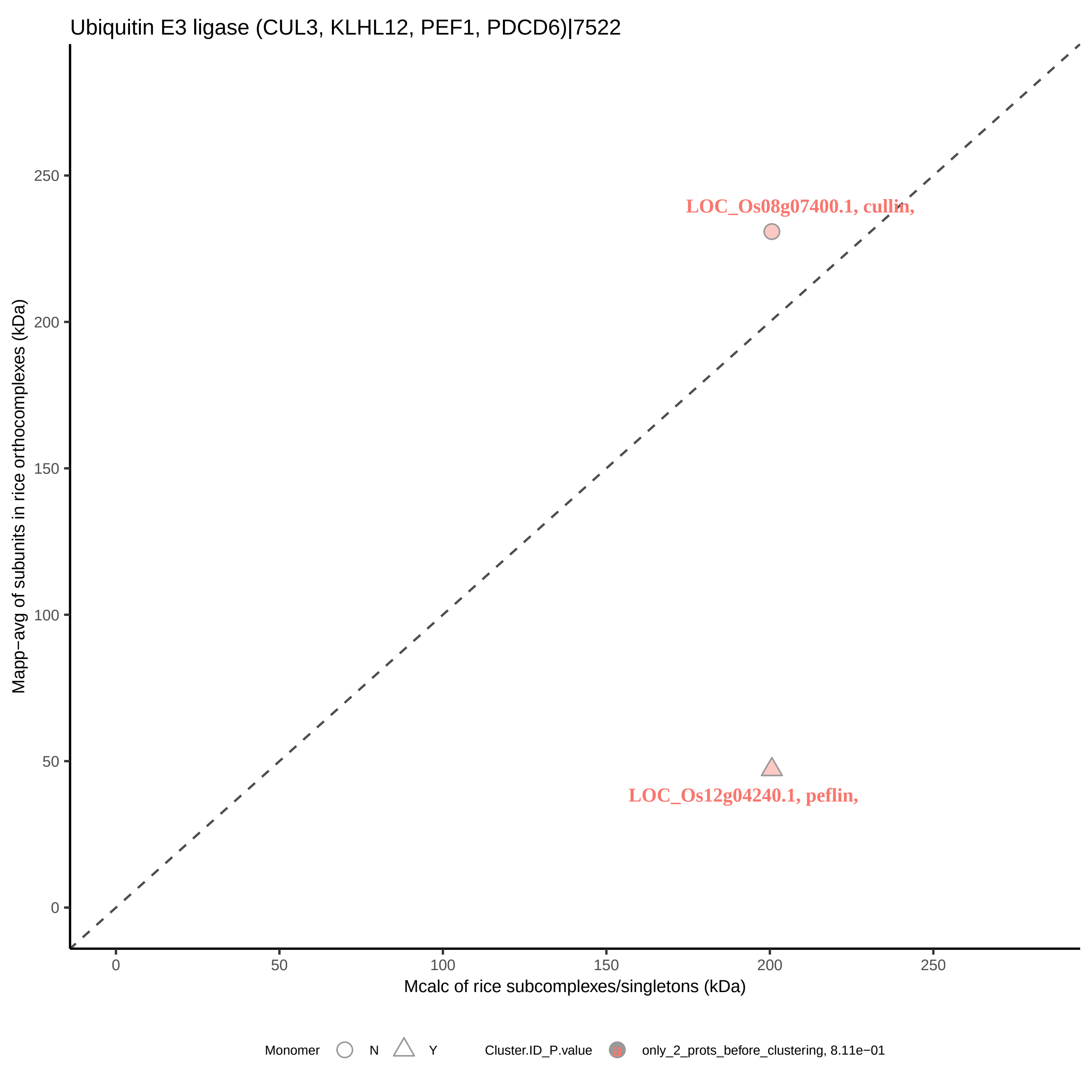

Ubiquitin E3 ligase (CUL3, KLHL12, PEF1, PDCD6)|7522
Mapp−avg of subunits in rice orthocomplexes (kDa)
Mcalc of rice subcomplexes/singletons (kDa)
0
50
100
150
200
250
LOC_Os08g07400.1, cullin,
LOC_Os12g04240.1, peflin,
Monomer
N
Y
Cluster.ID_P.value
only_2_prots_before_clustering, 8.11e−01

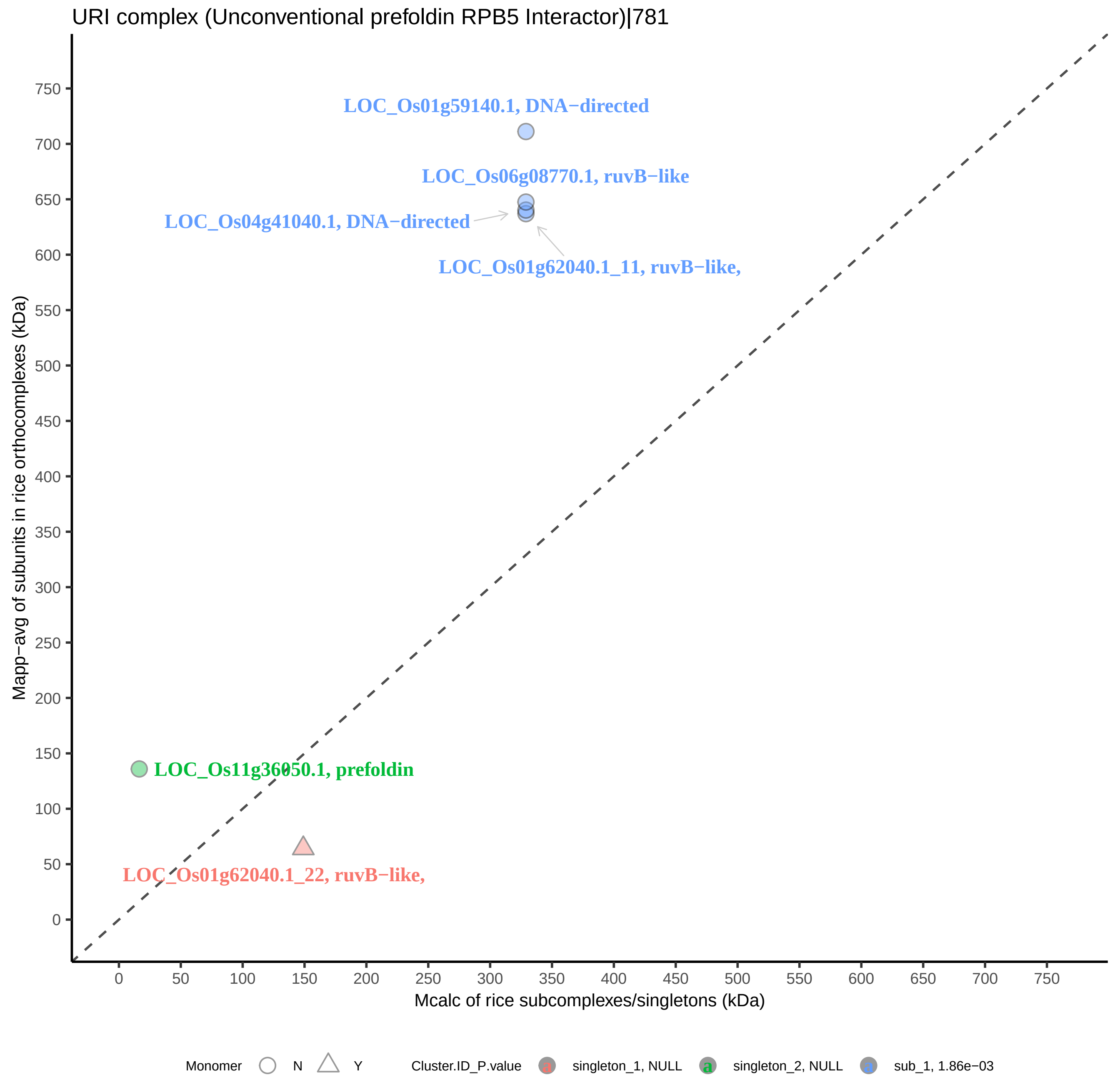
URI complex (Unconventional prefoldin RPB5 Interactor)|781
LOC_Os01g59140.1, DNA−directed
LOC_Os06g08770.1, ruvB−like
LOC_Os04g41040.1, DNA−directed
LOC_Os01g62040.1_11, ruvB−like,
LOC_Os11g36050.1, prefoldin
LOC_Os01g62040.1_22, ruvB−like,
Mapp−avg of subunits in rice orthocomplexes (kDa)
Mcalc of rice subcomplexes/singletons (kDa)
0
50
100
150
200
250
300
350
400
450
500
550
600
650
700
750
Monomer
N
Y
Cluster.ID_P.value
singleton_1, NULL
singleton_2, NULL
sub_1, 1.86e−03
