## Supplemental Figure S3 Rice protein SEC profiles illustrating small-world analysis results for "A machine learning-based approach to identify reliable gold standards for protein complex composition prediction"

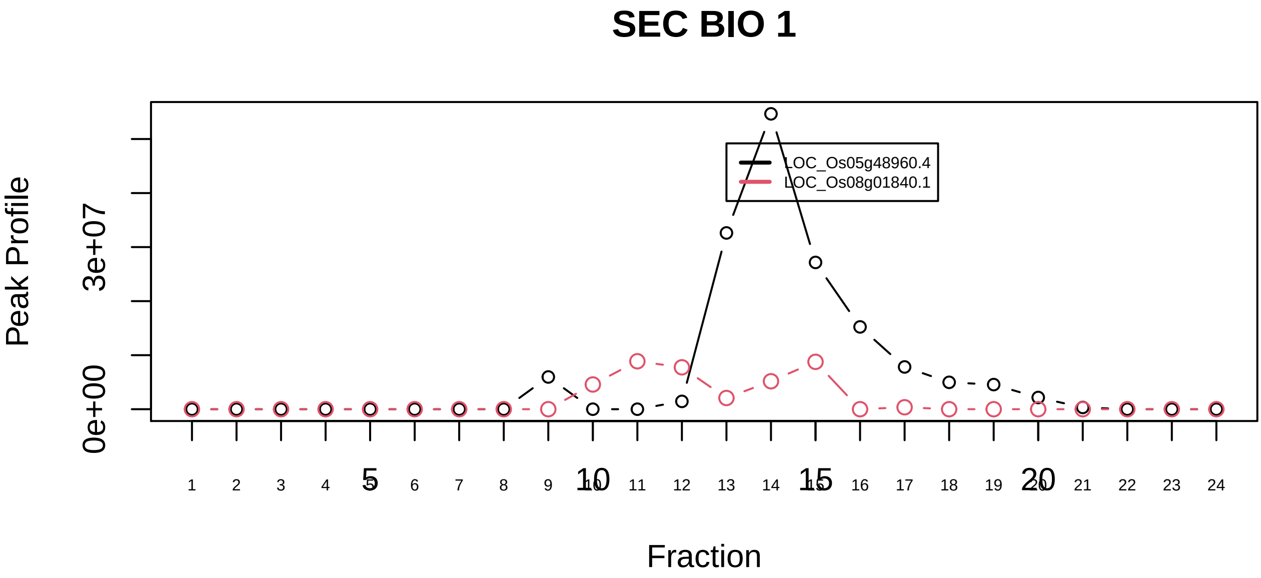

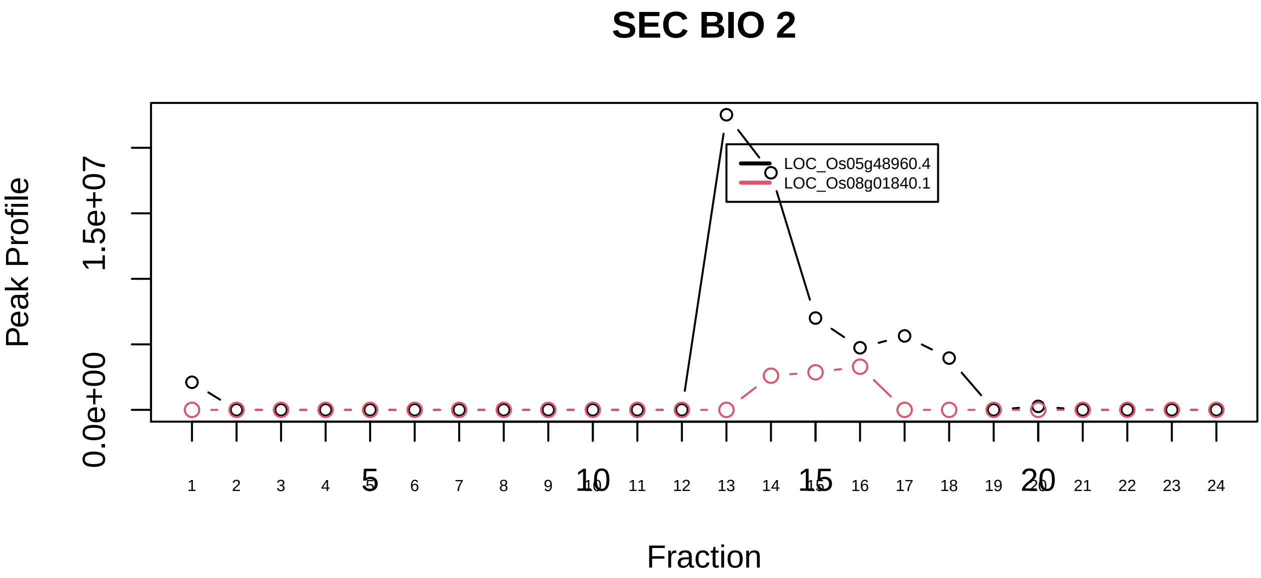

17S U2 snRNP|2755−−sub_1

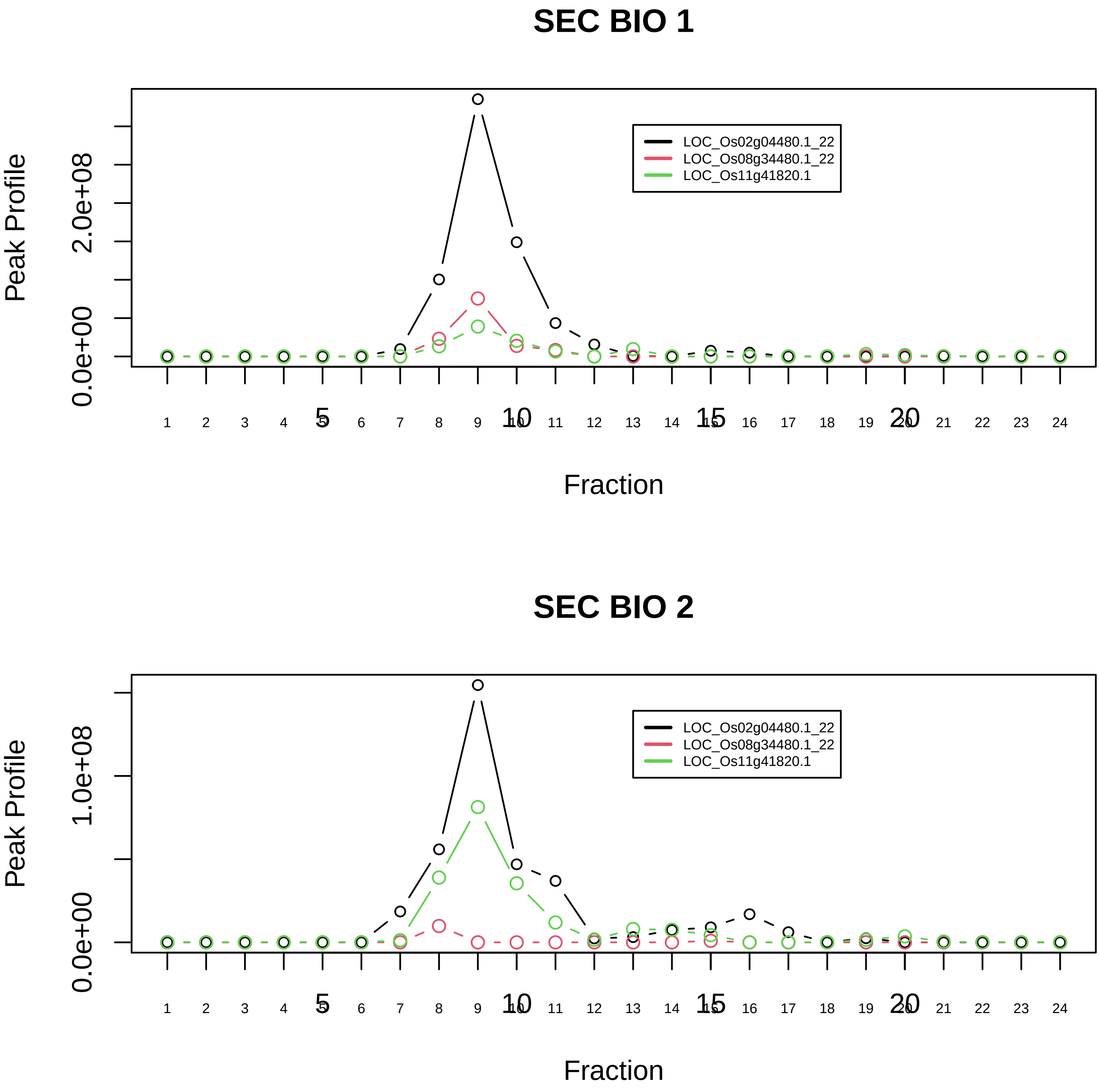

17S U2 snRNP|2755−−sub_2

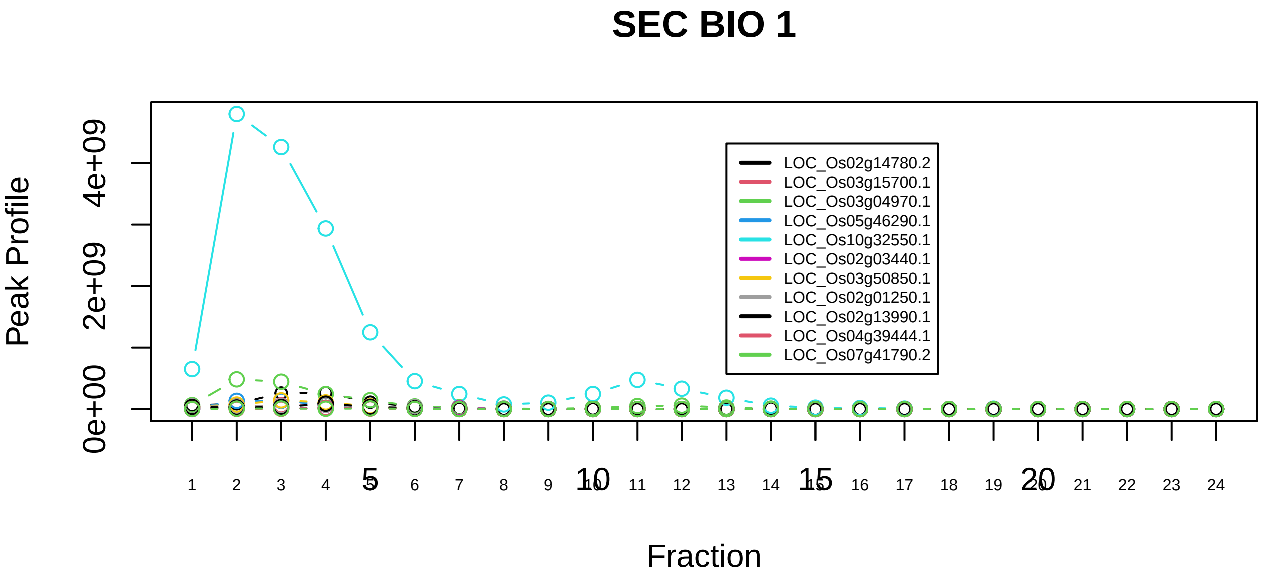

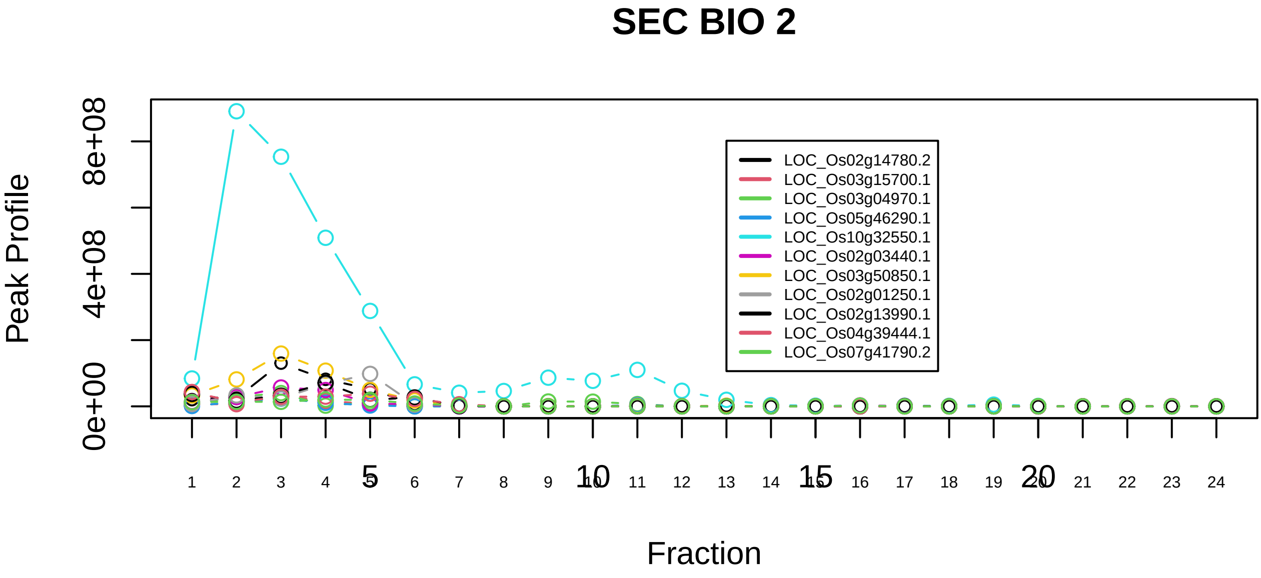

17S U2 snRNP|2755−−sub_3

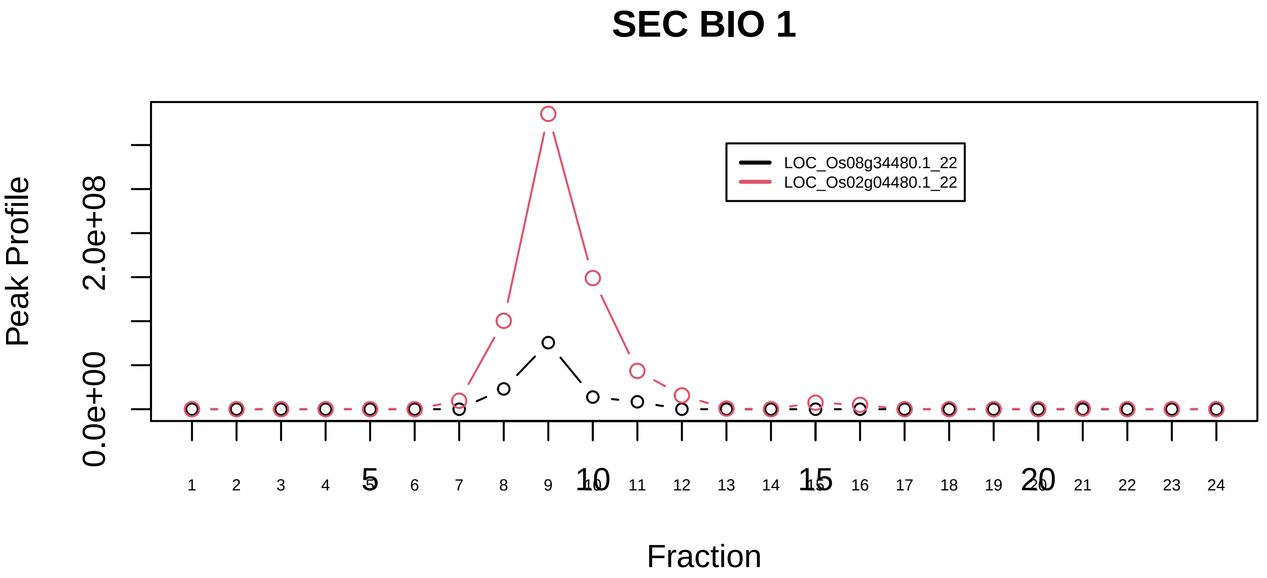

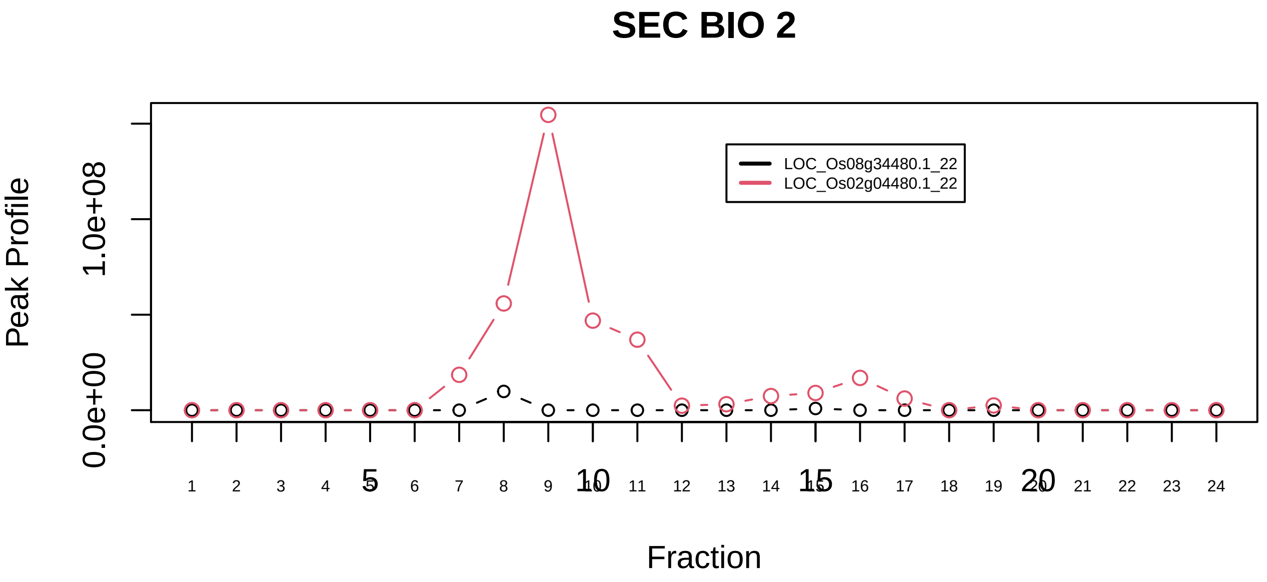

18S U11/U12 snRNP|430−−sub_1

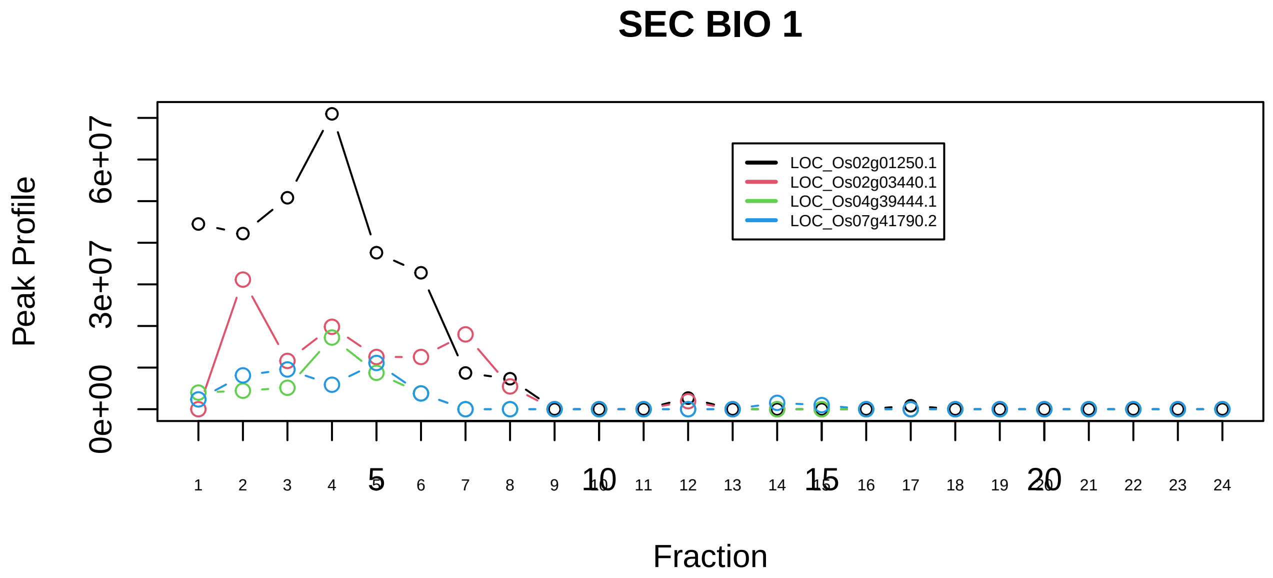

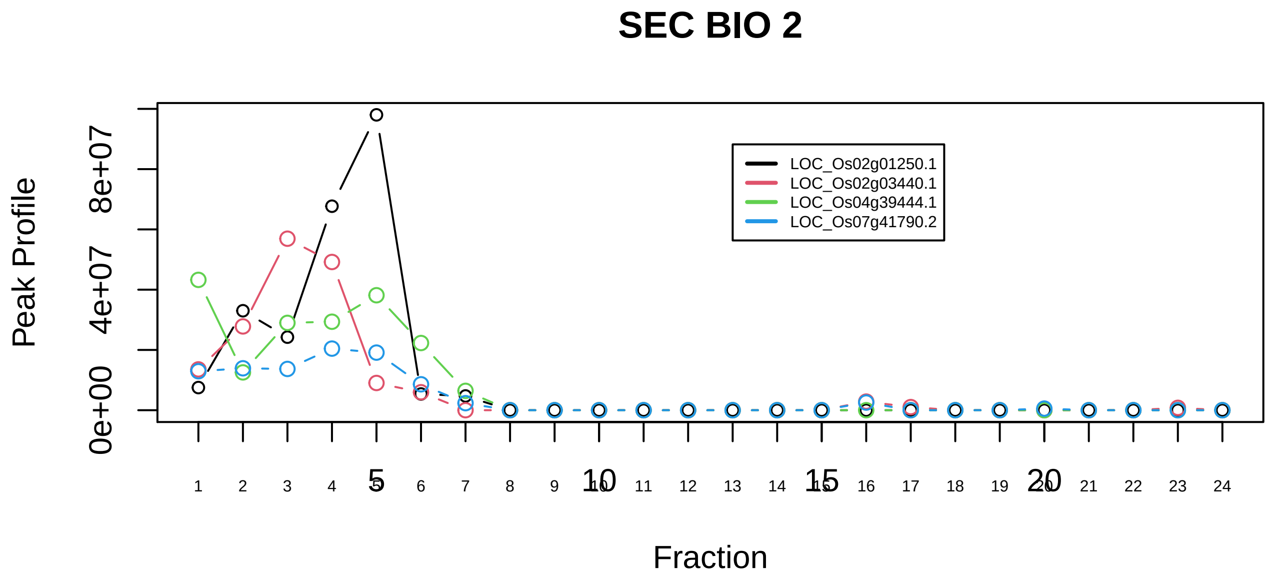

18S U11/U12 snRNP|430−−sub_2

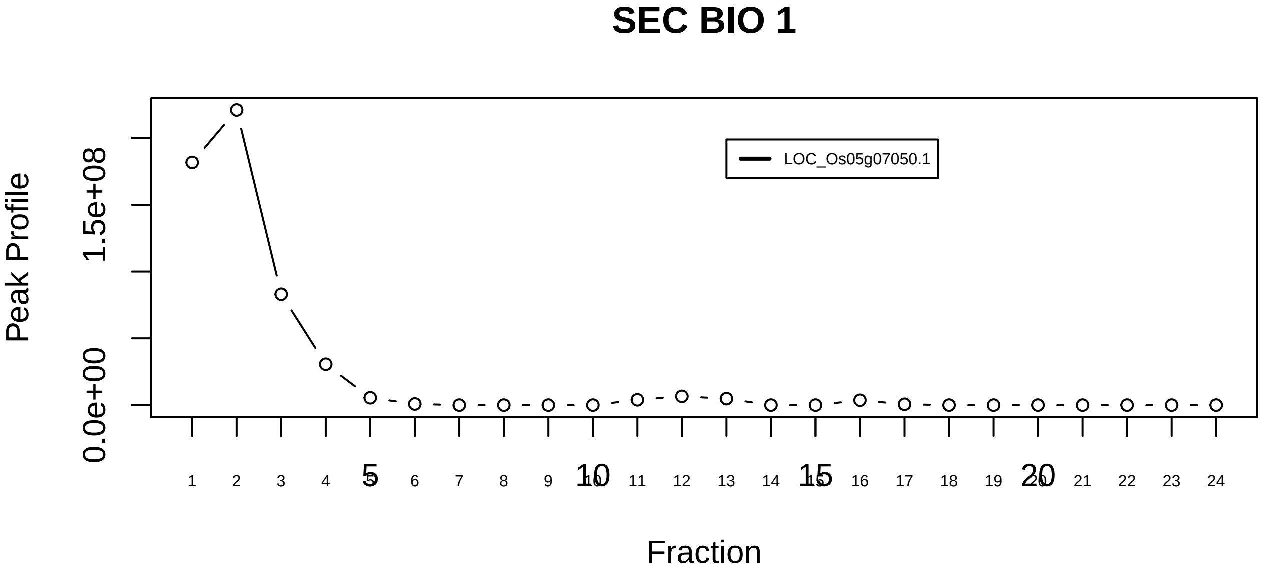

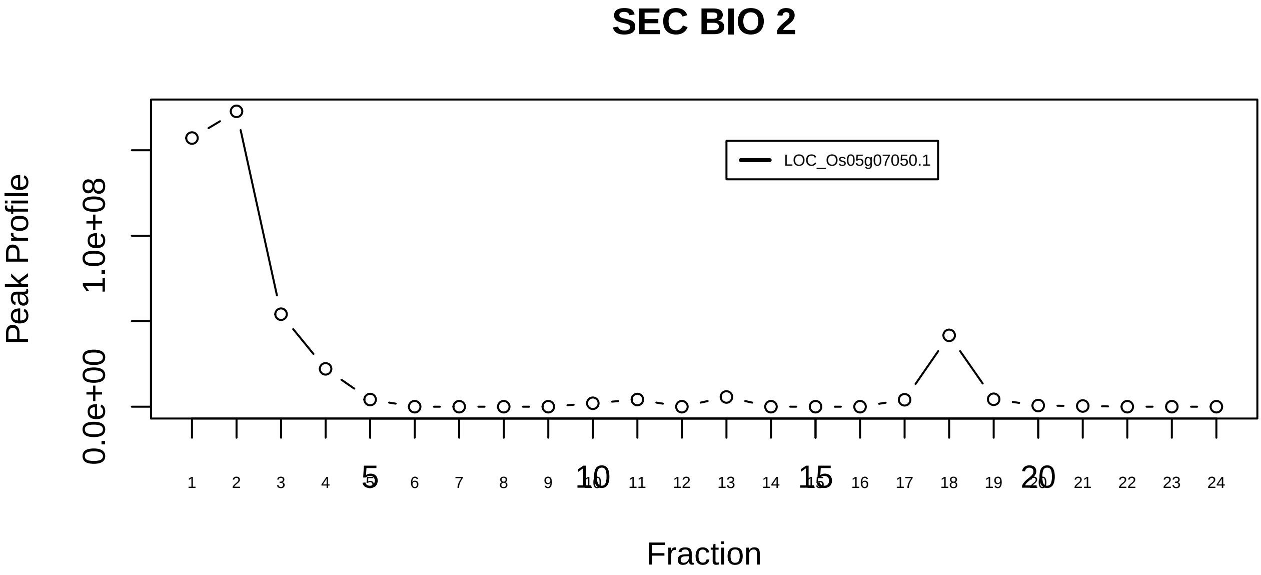

18S U11/U12 snRNP|430−−singleton_1

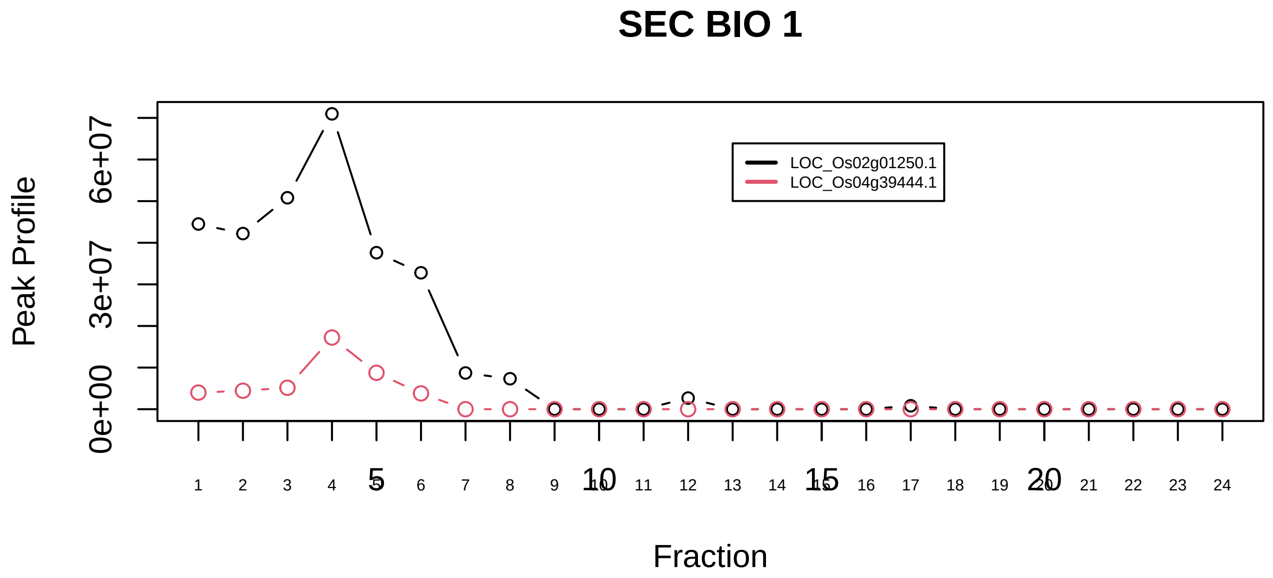

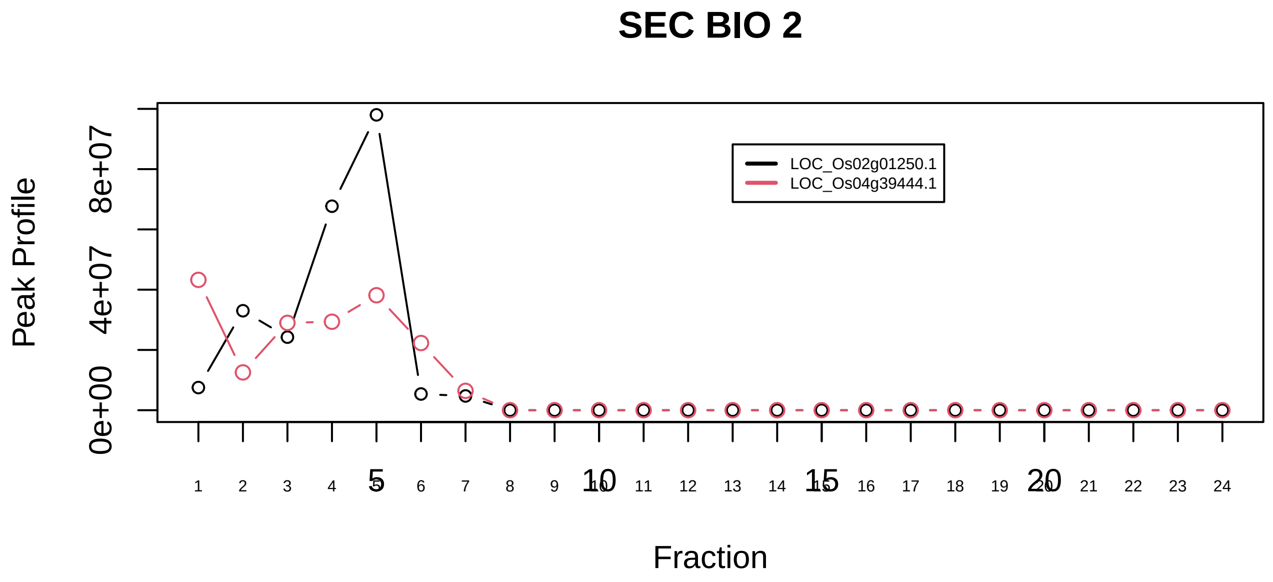

20S methylosome−SmD complex|819−−sub_1

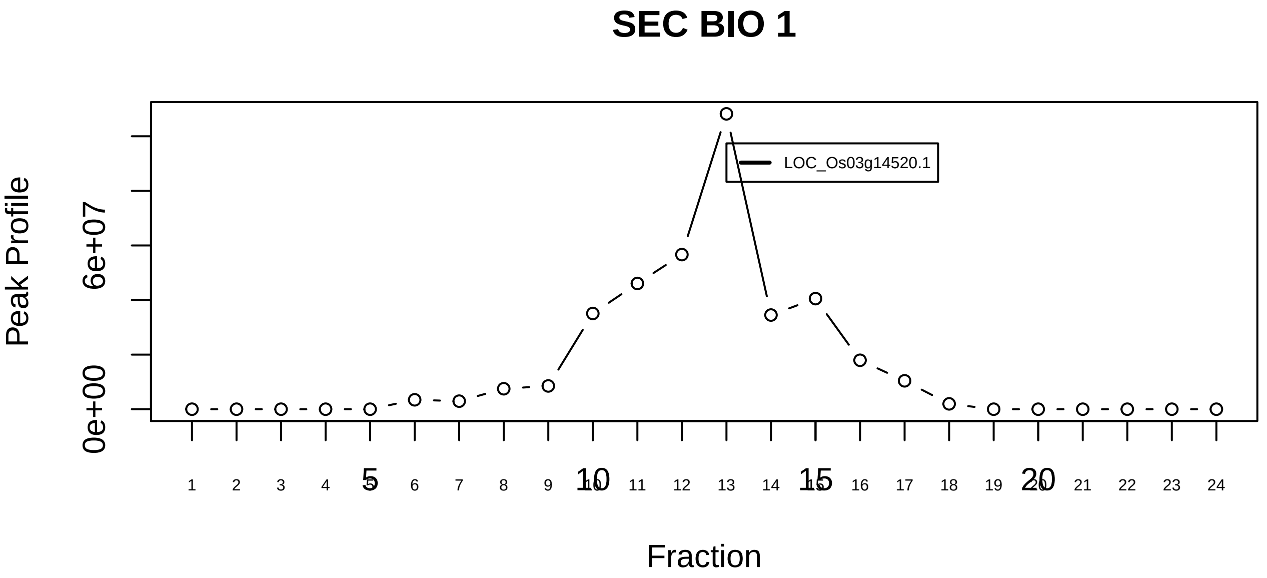

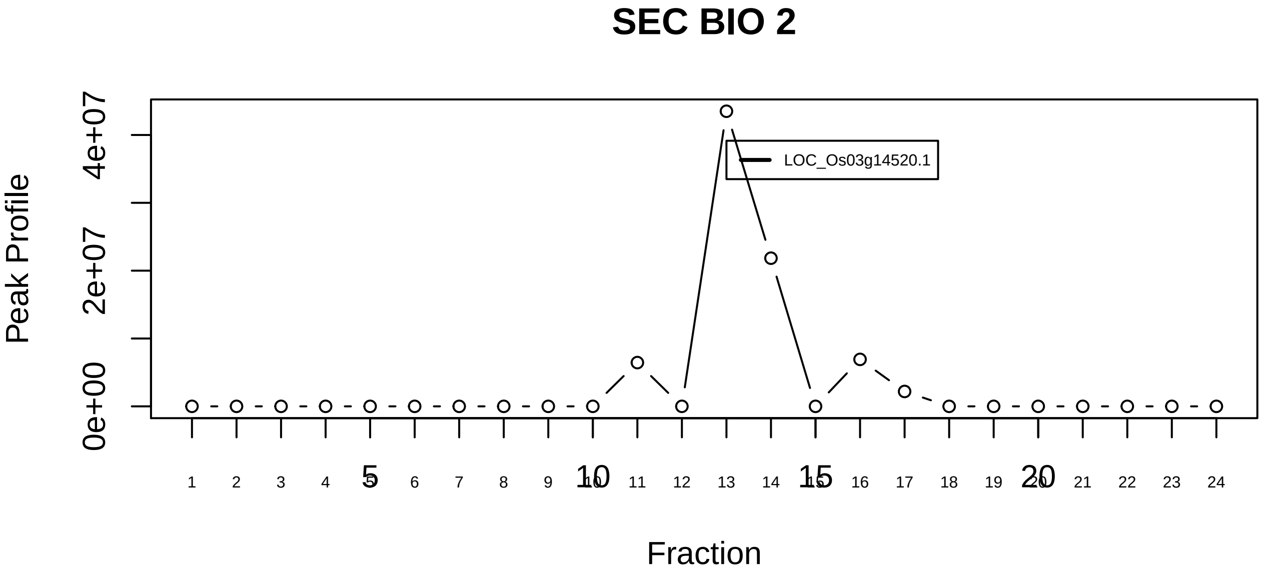

20S methylosome−SmD complex|819−−singleton_1

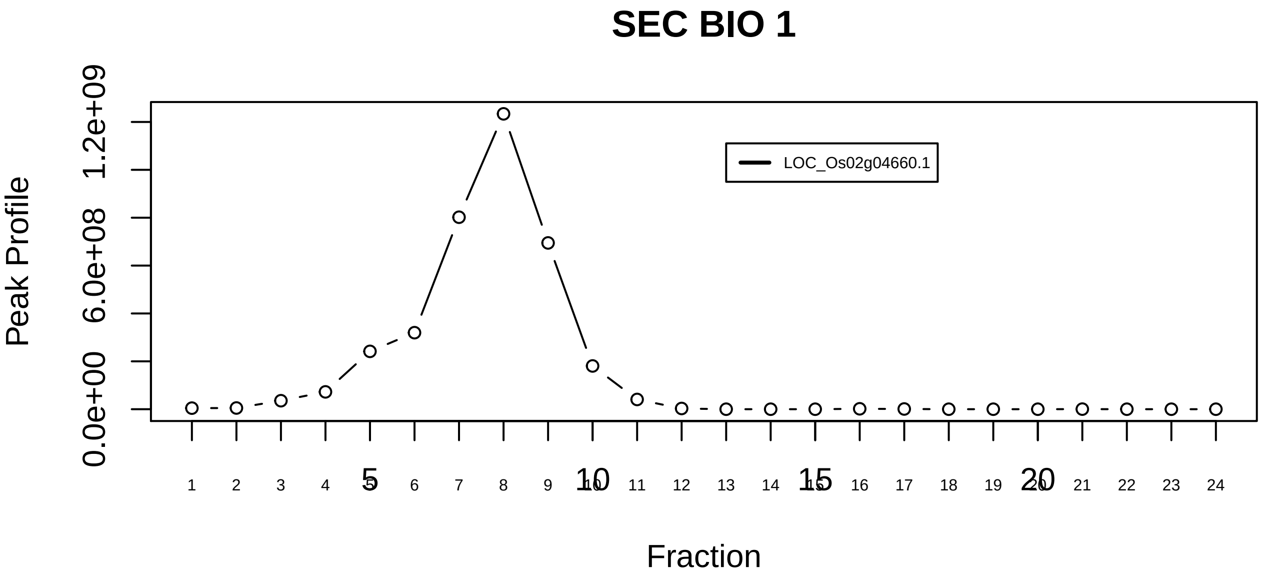

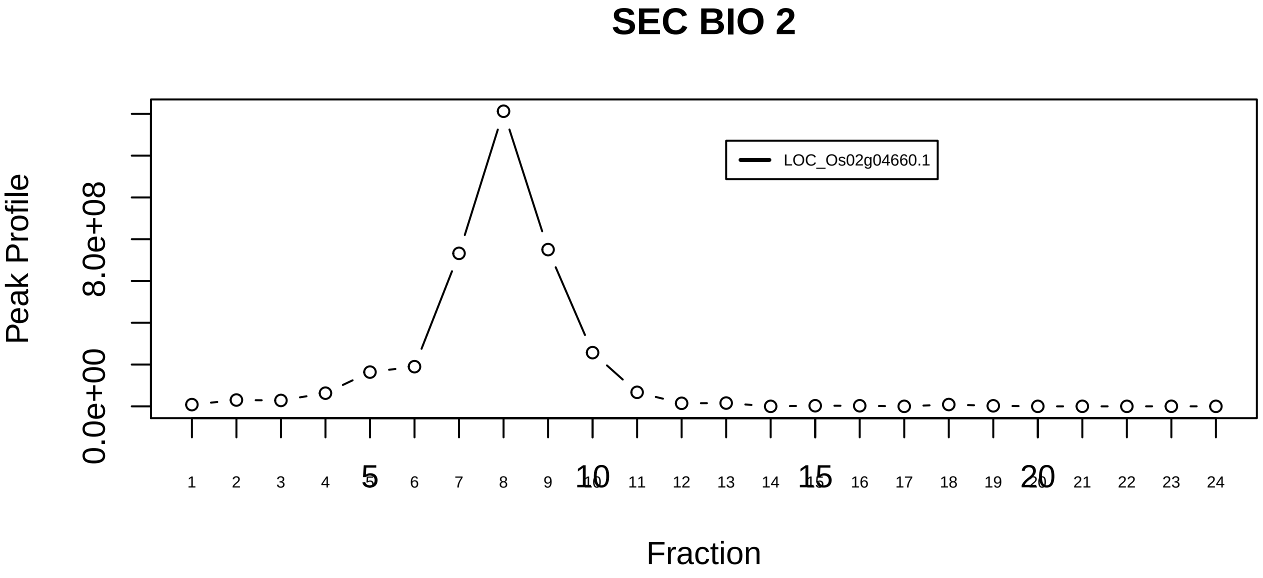

20S methylosome−SmD complex|819−−singleton_2

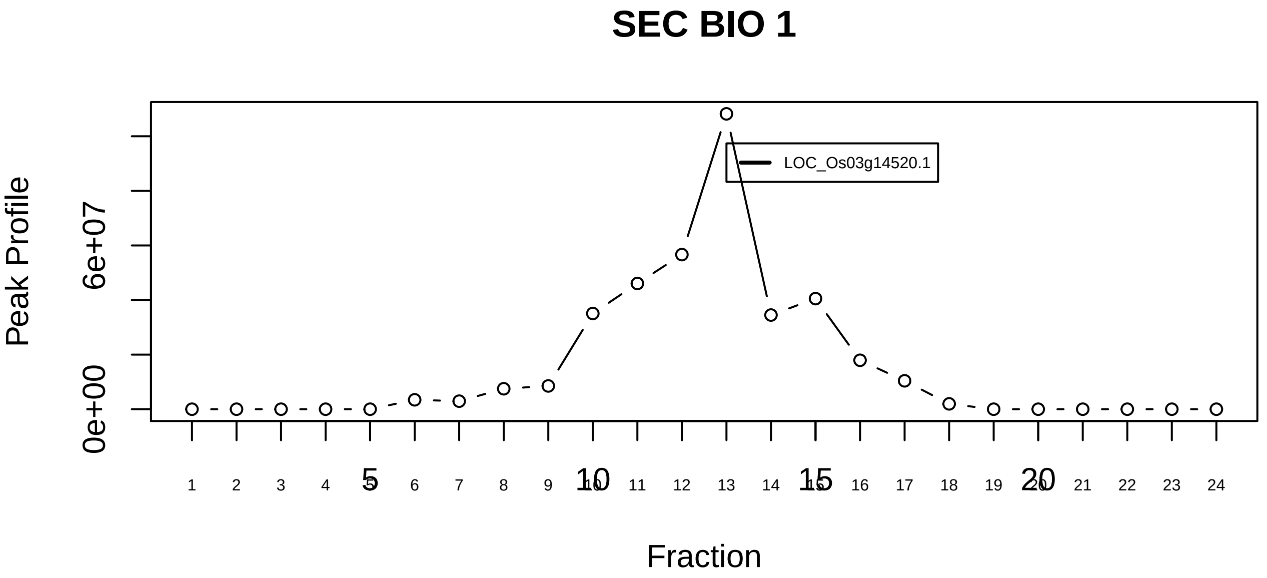

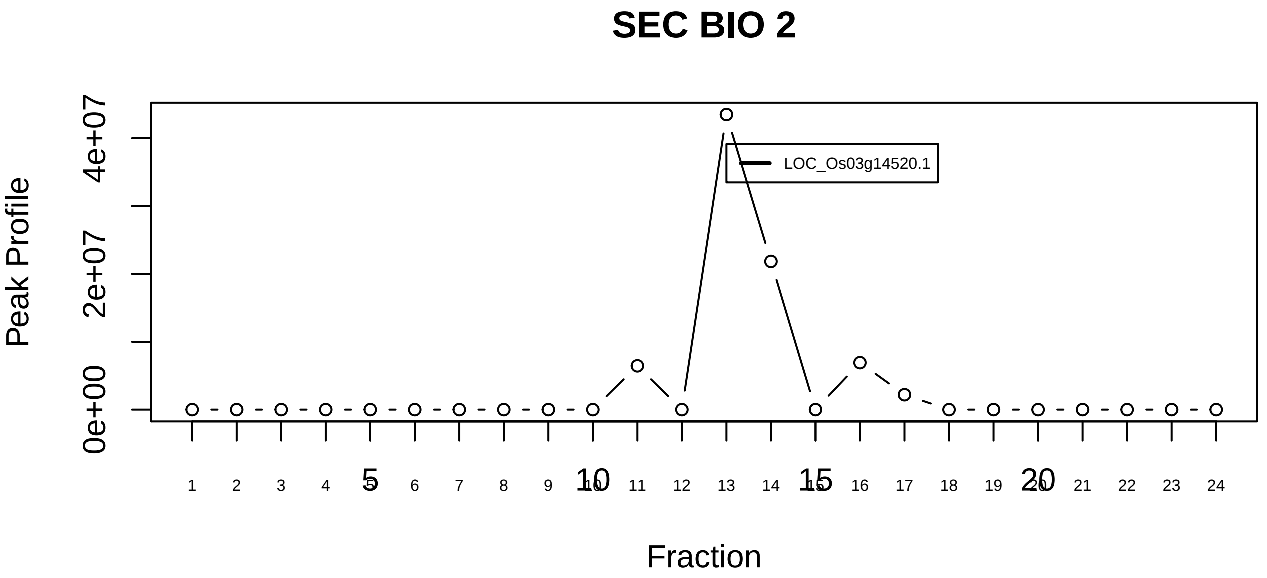

6S methyltransferase and RG−containing Sm proteins complex|835−−singleton_1

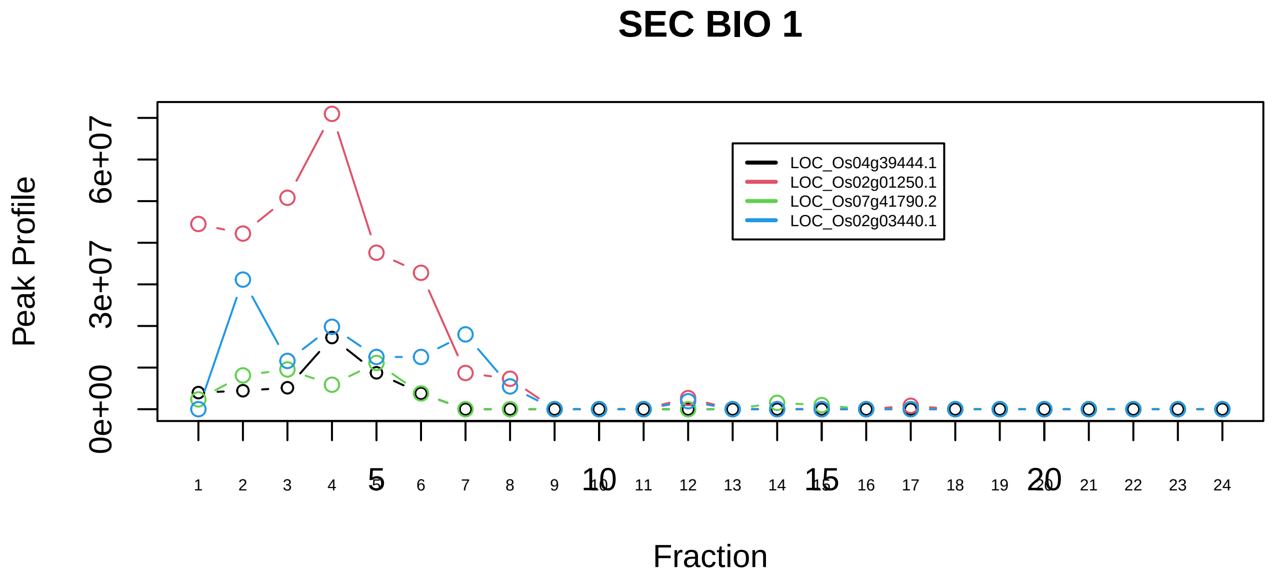

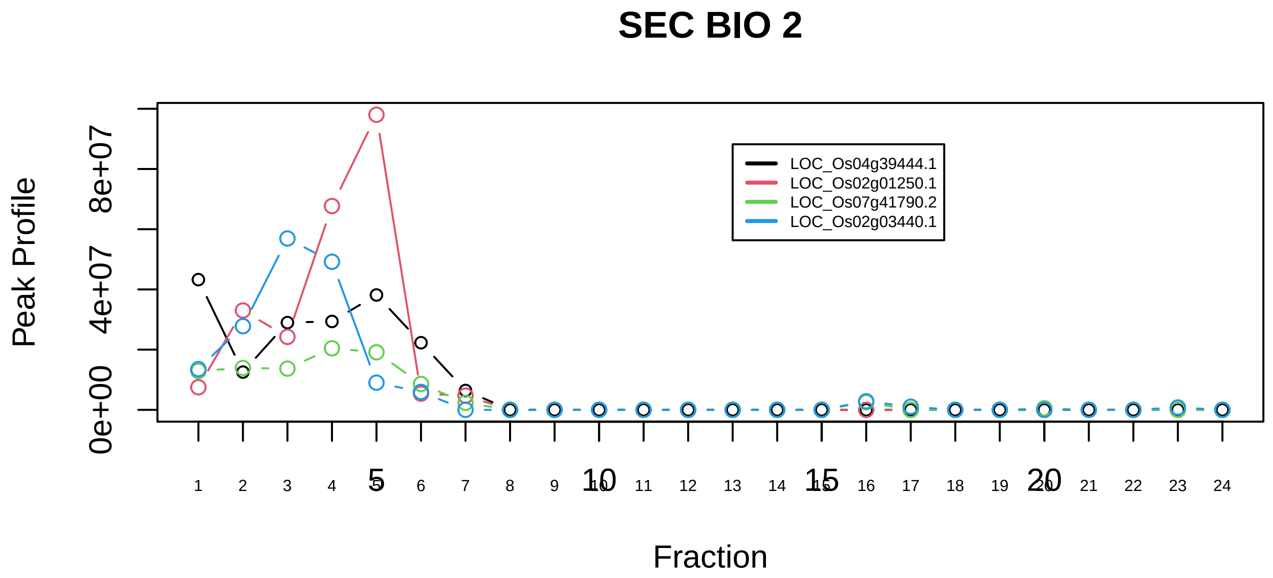

6S methyltransferase and RG−containing Sm proteins complex|835−−sub_1

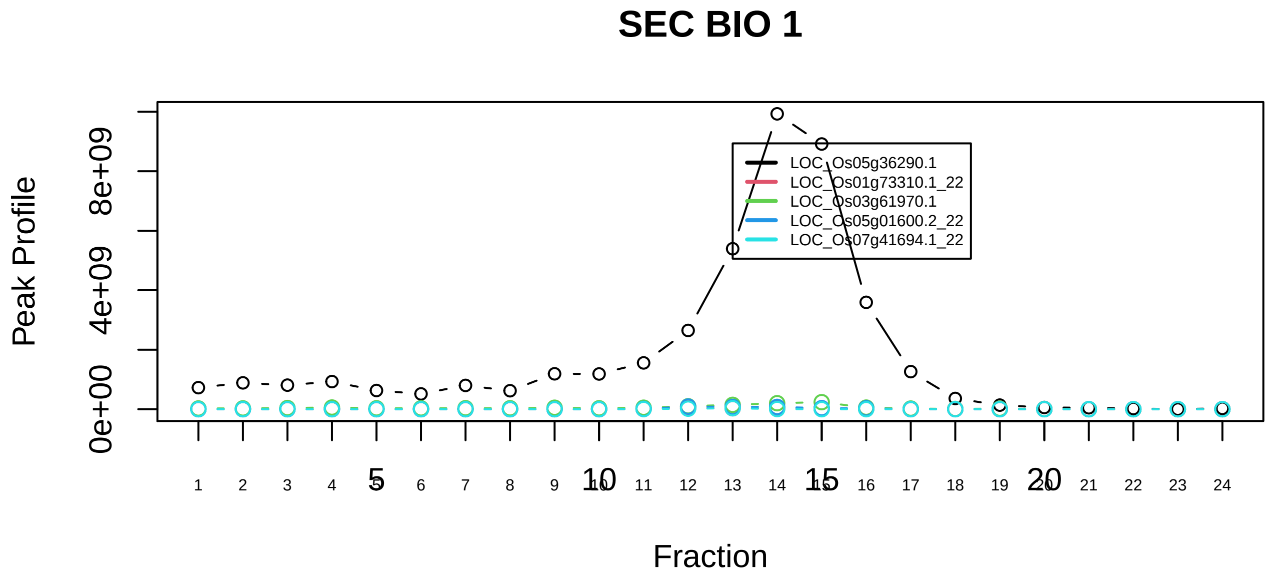

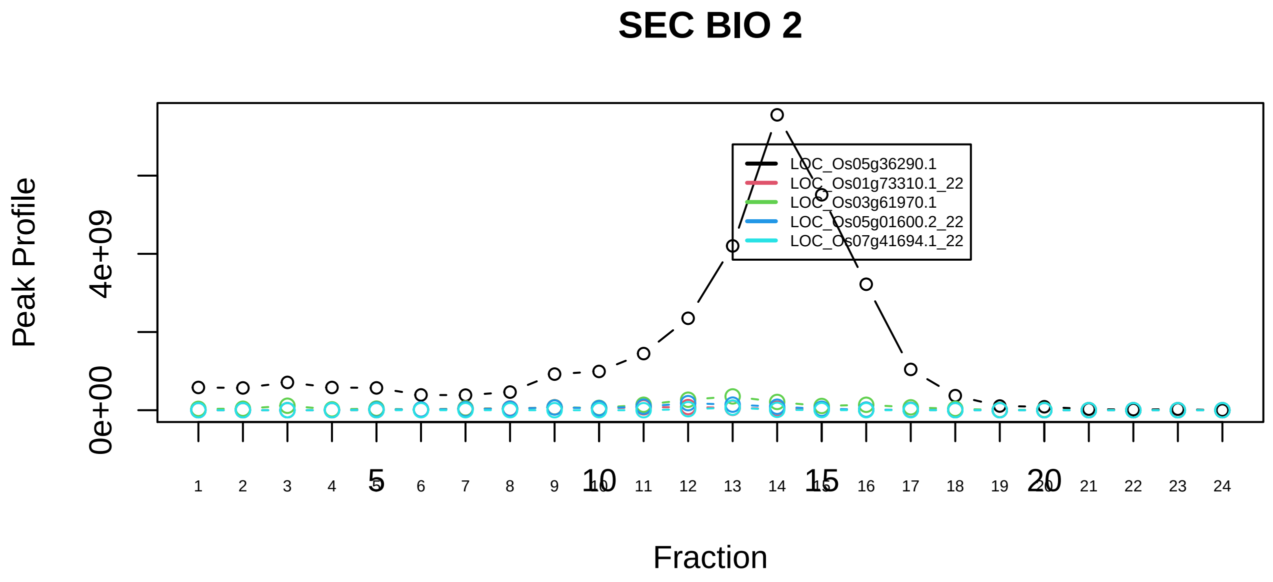

ACTB−ANP32A−C1QBP−PSMA1−PTMA−PSMA1 complex|7298−−sub_1

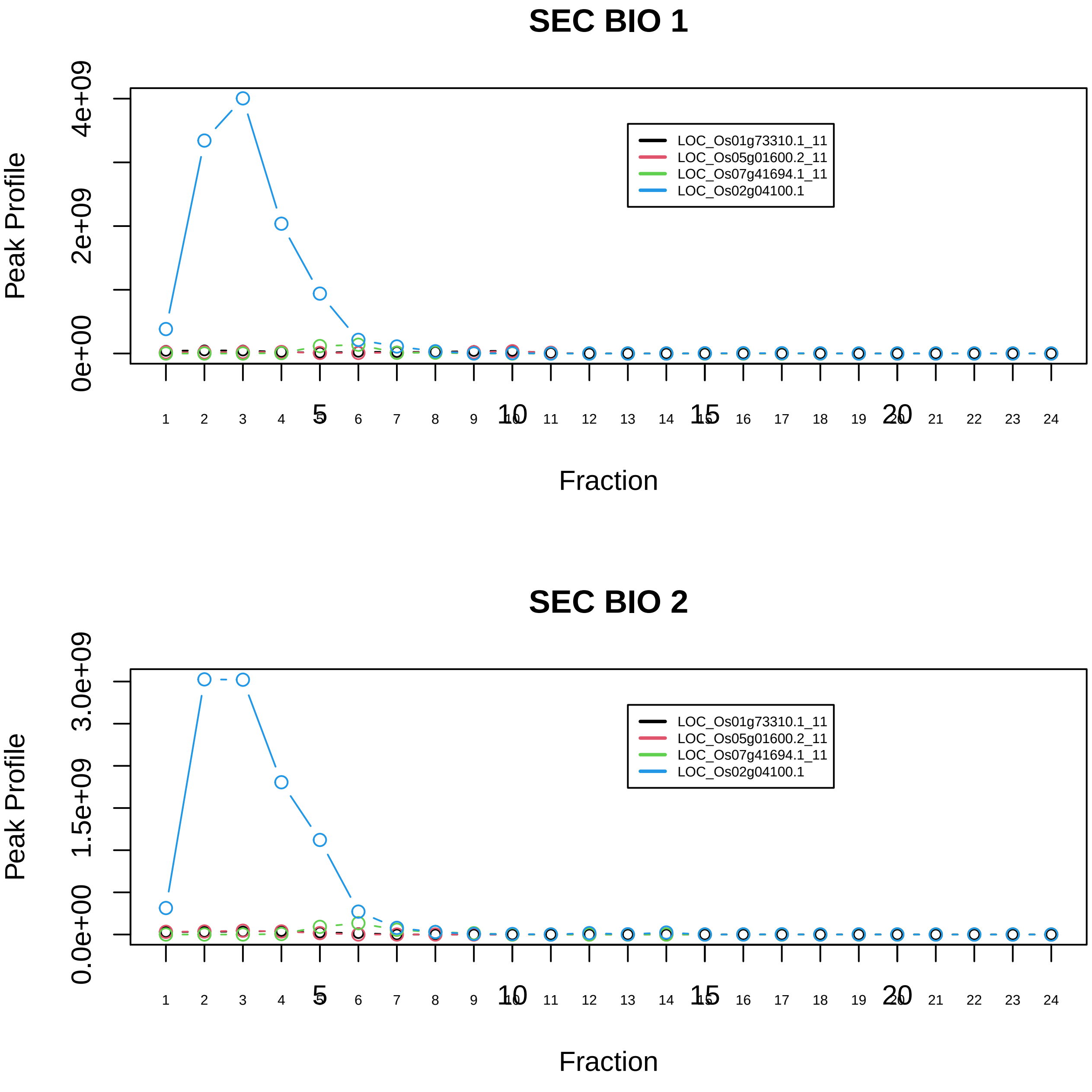

ACTB−ANP32A−C1QBP−PSMA1−PTMA−PSMA1 complex|7298−−sub_2

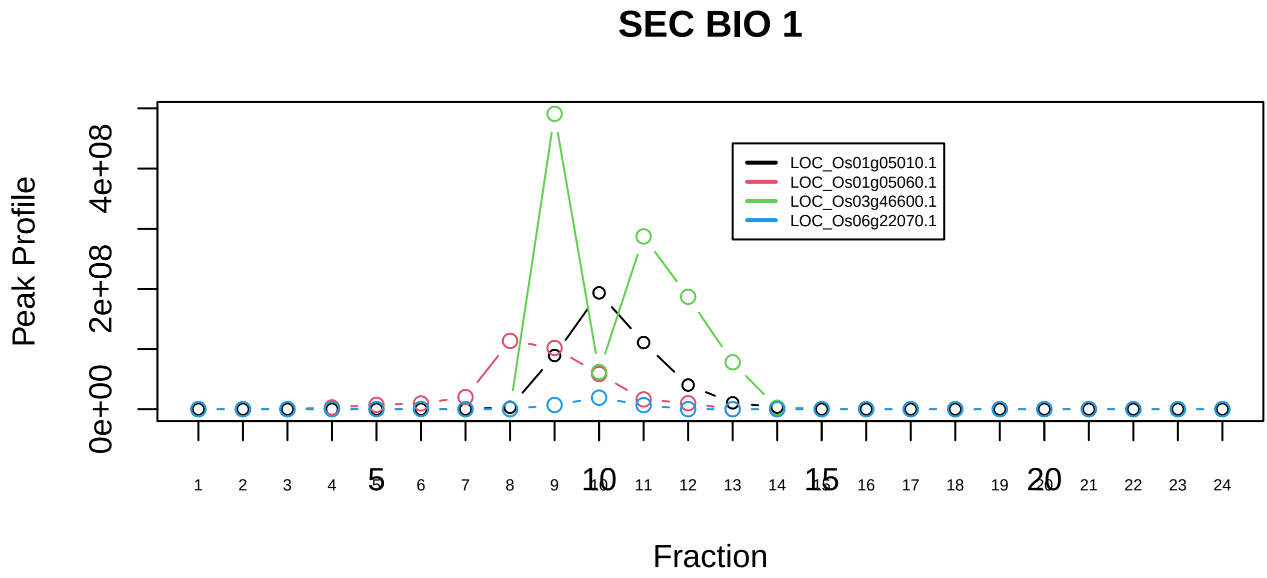

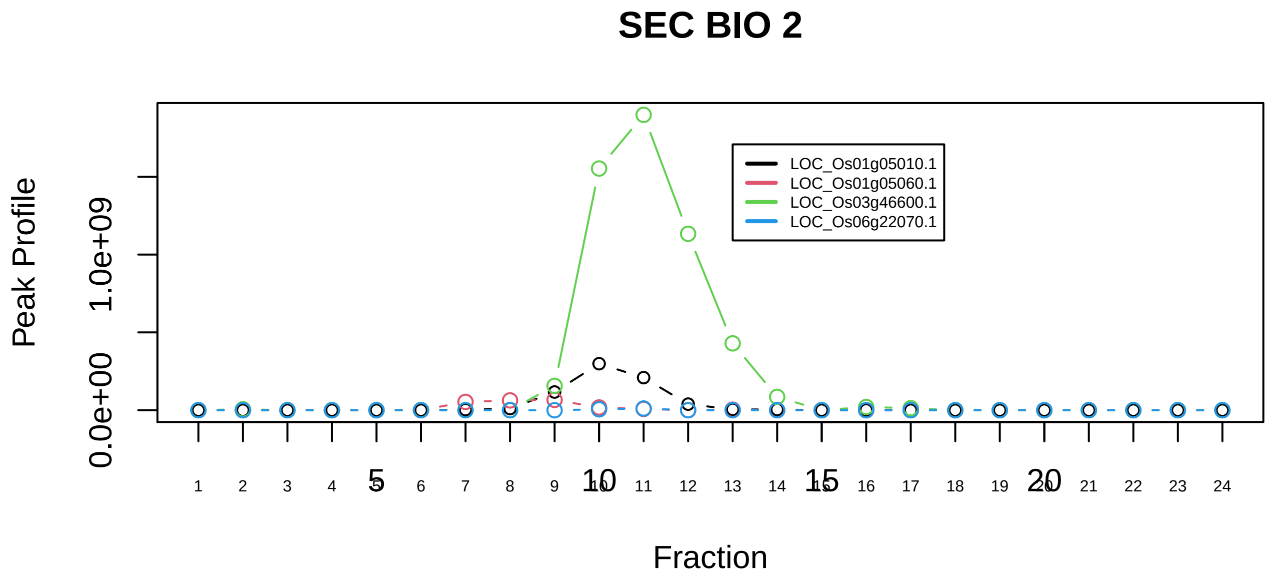

ACTB−ANP32A−C1QBP−PSMA1−PTMA−PSMA1 complex|7298−−sub_3

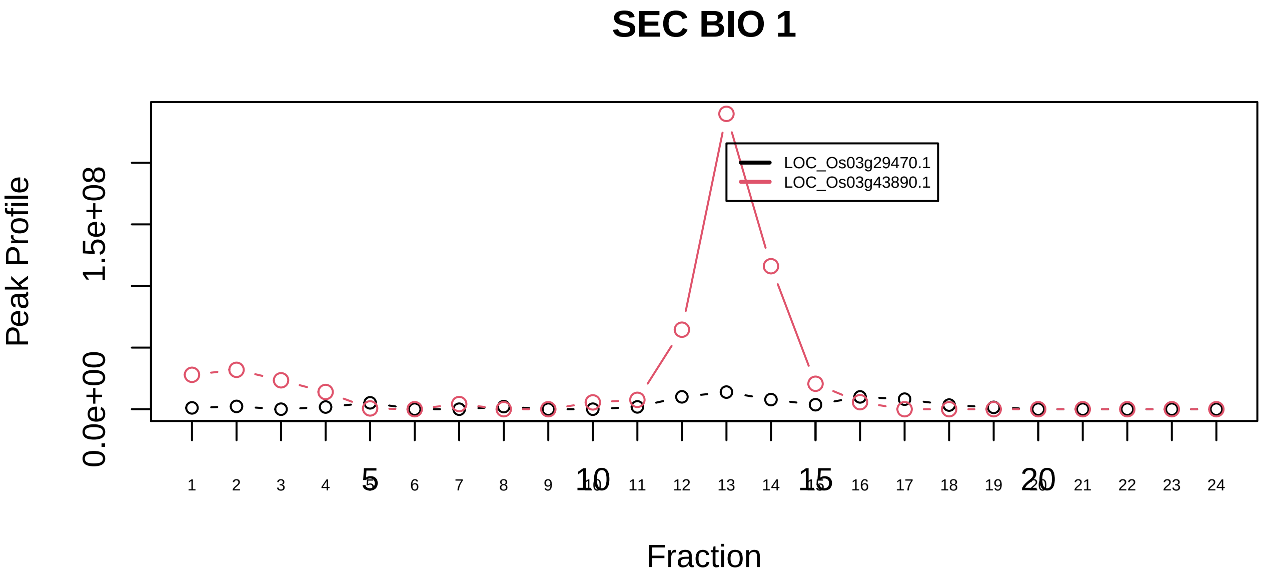

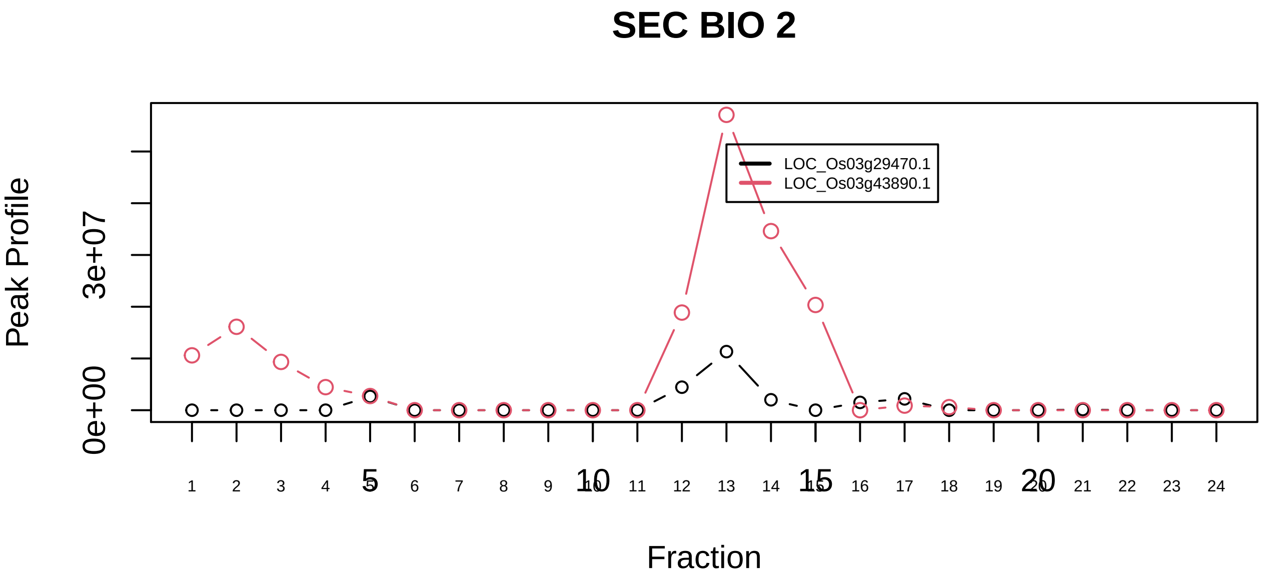

ALL−1 supercomplex|1257−−sub_1

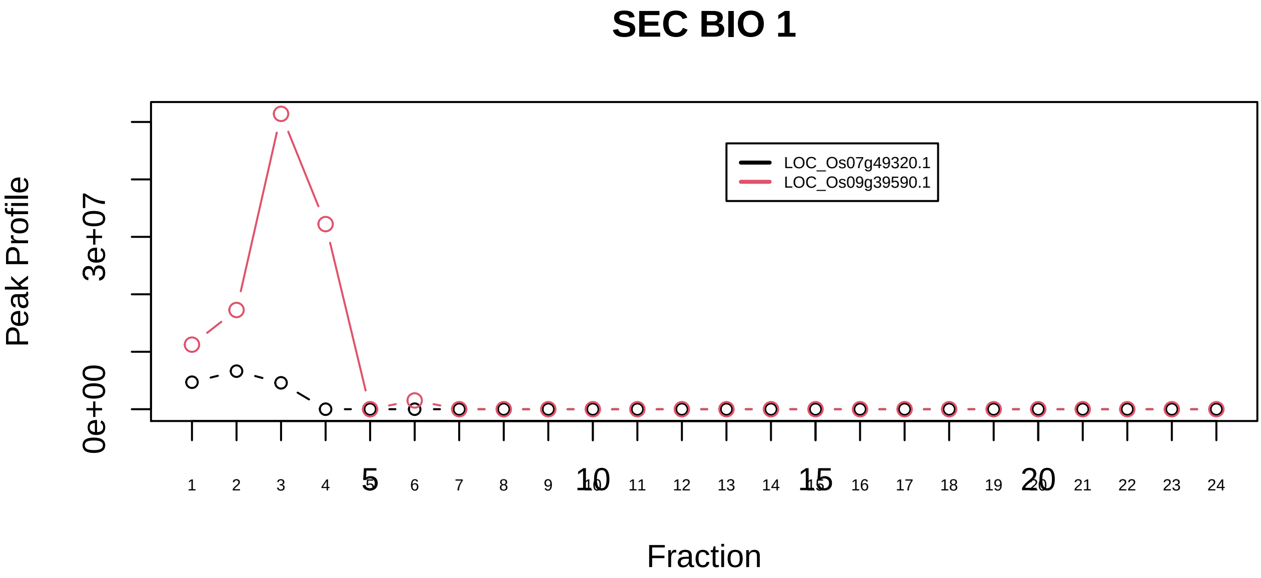

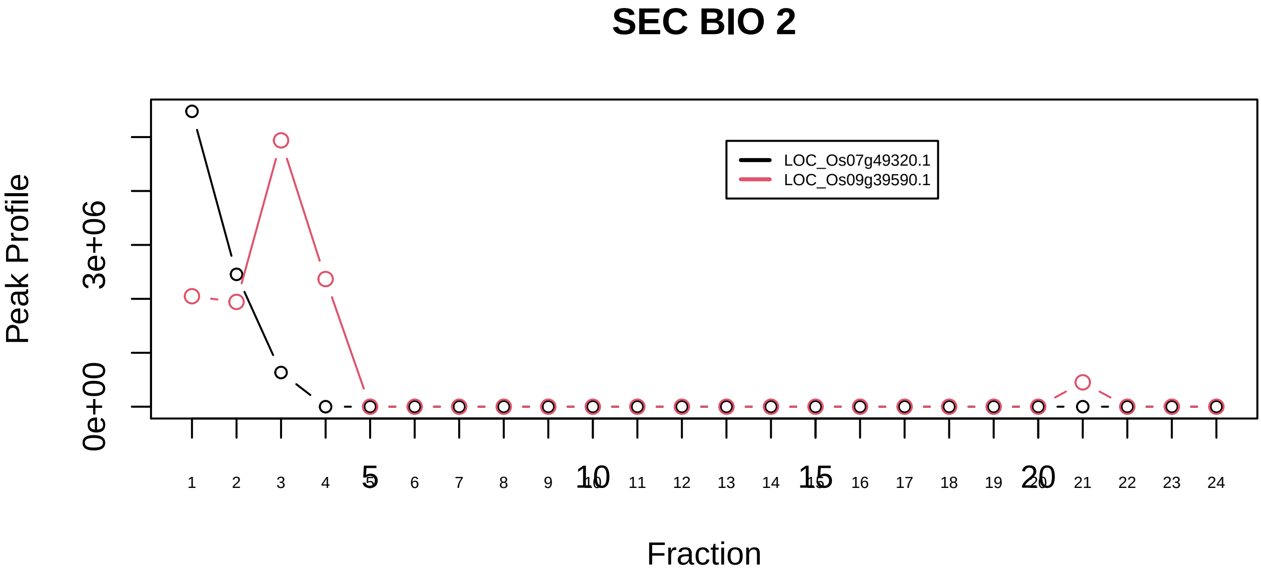

ALL−1 supercomplex|1257−−sub_2

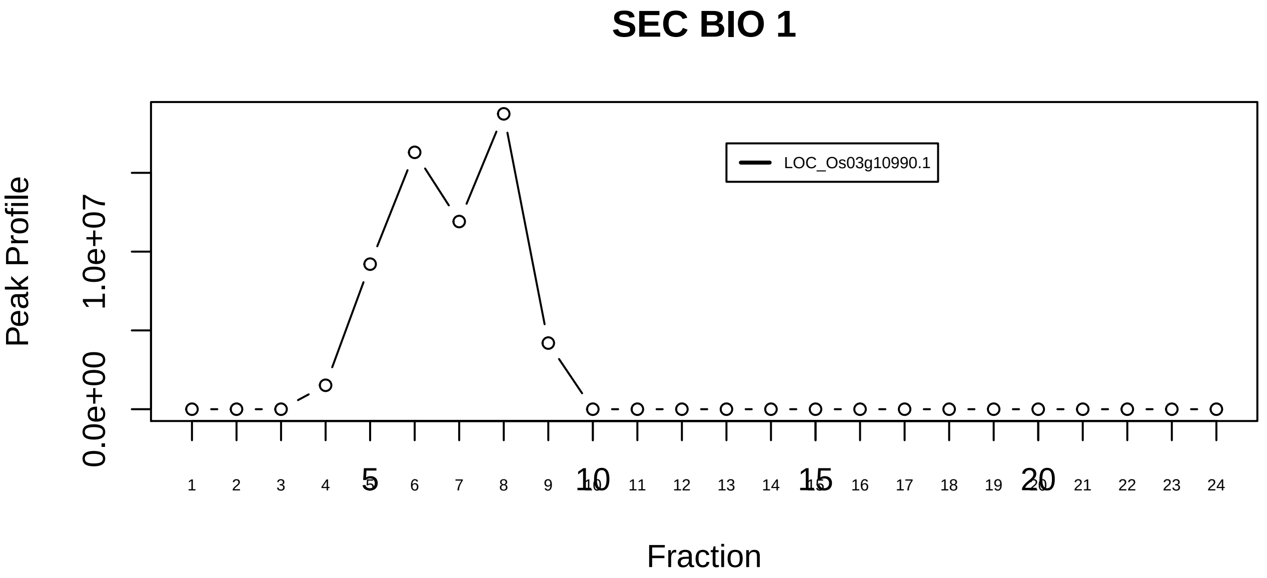

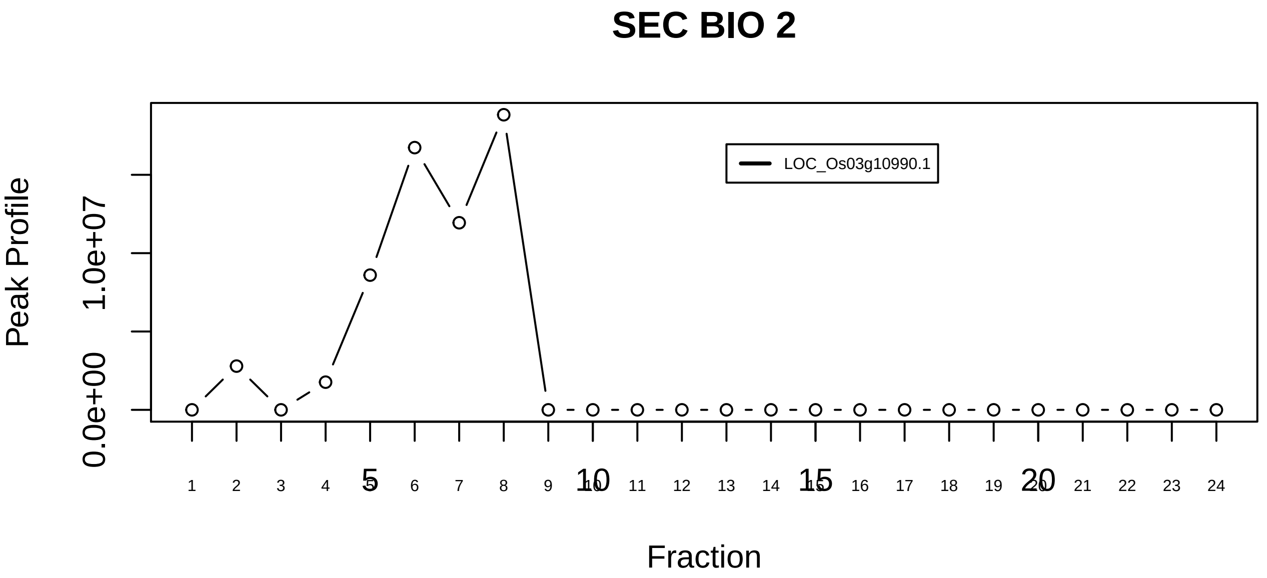

ALL−1 supercomplex|1257−−singleton_1

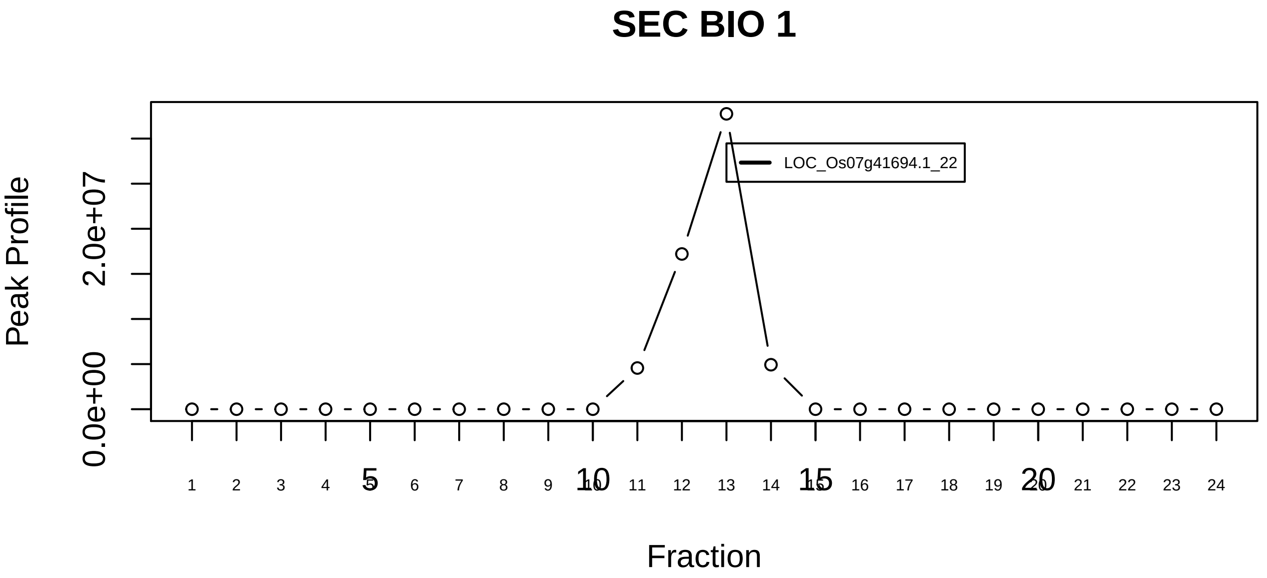
SEC BIO 1
Peak Profile
0.0e+00
2.0e+07
LOC_Os07g41694.1_22
Fraction

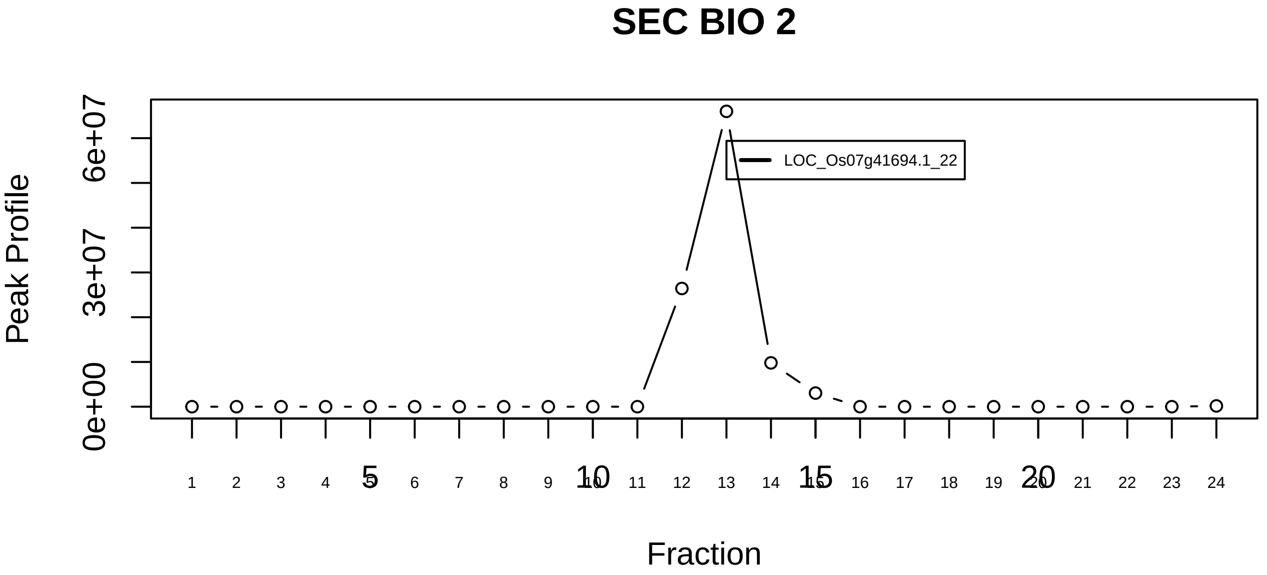
SEC BIO 2
Peak Profile
0e+00
3e+07
6e+07
LOC_Os07g41694.1_22
Fraction

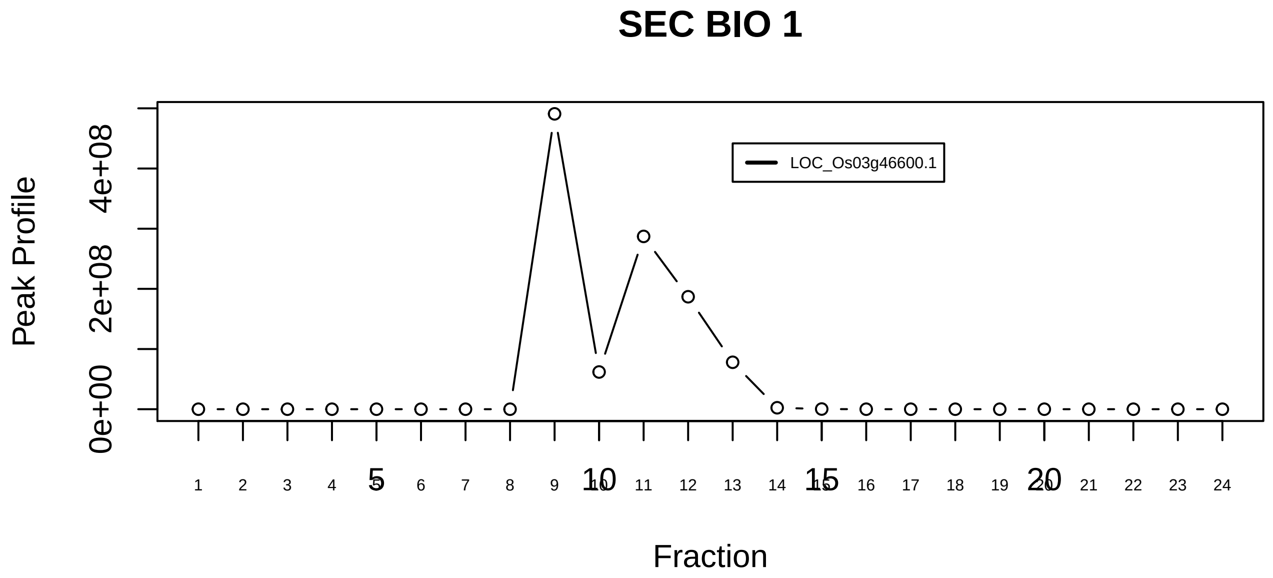

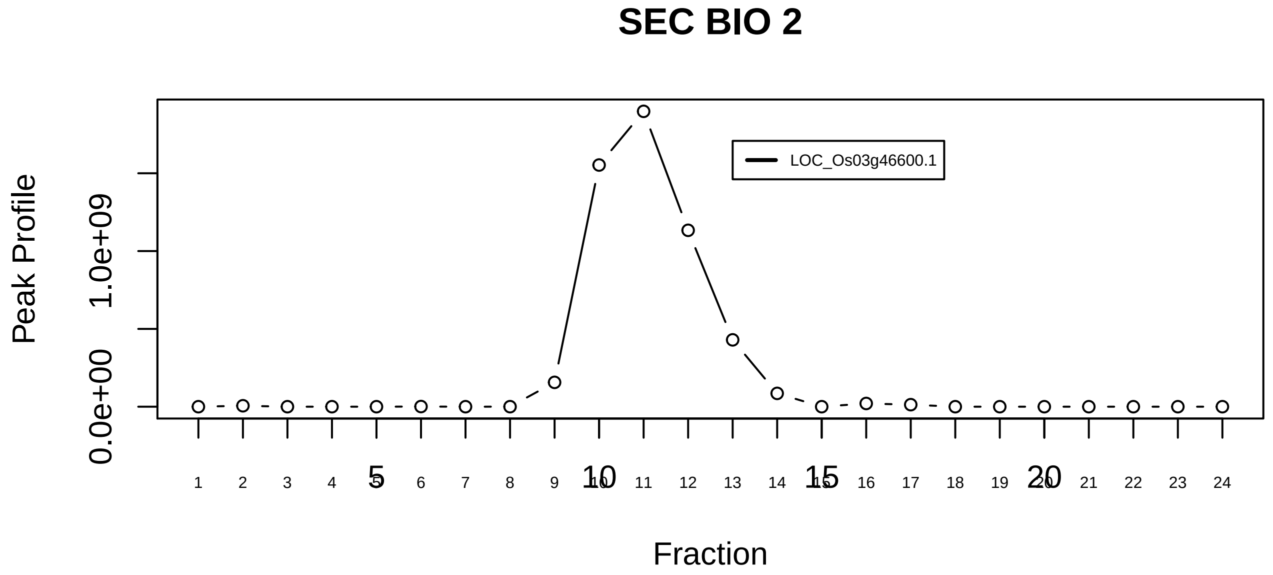

ANP32A−ANP32B−PSMA3−PTMA−SLC25A5 complex|7299−−singleton_2

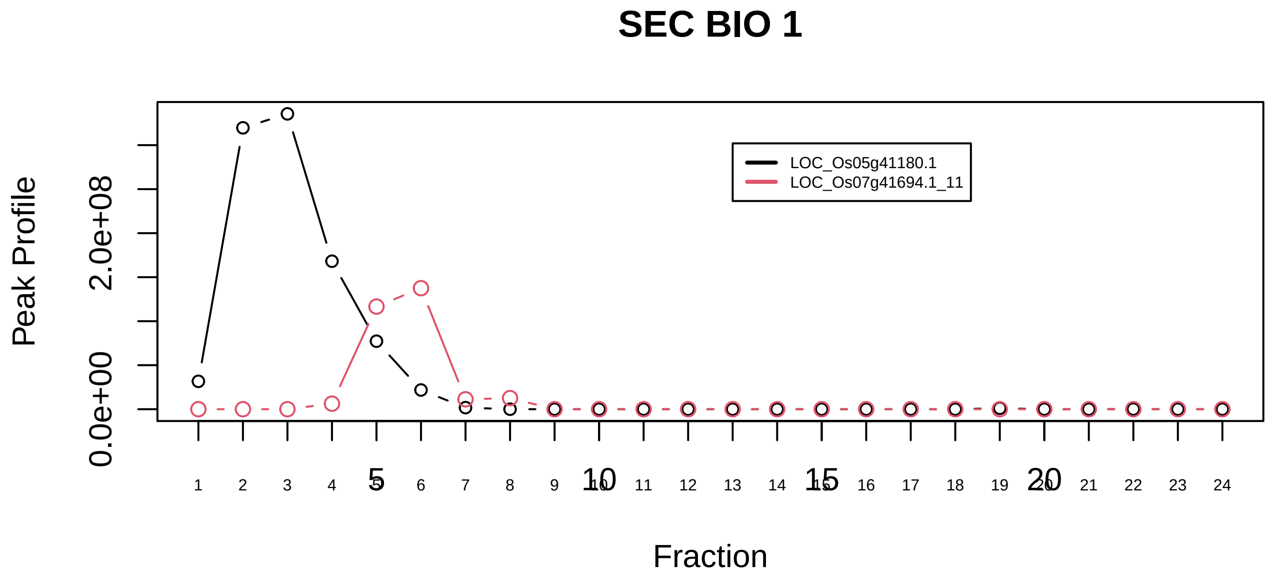

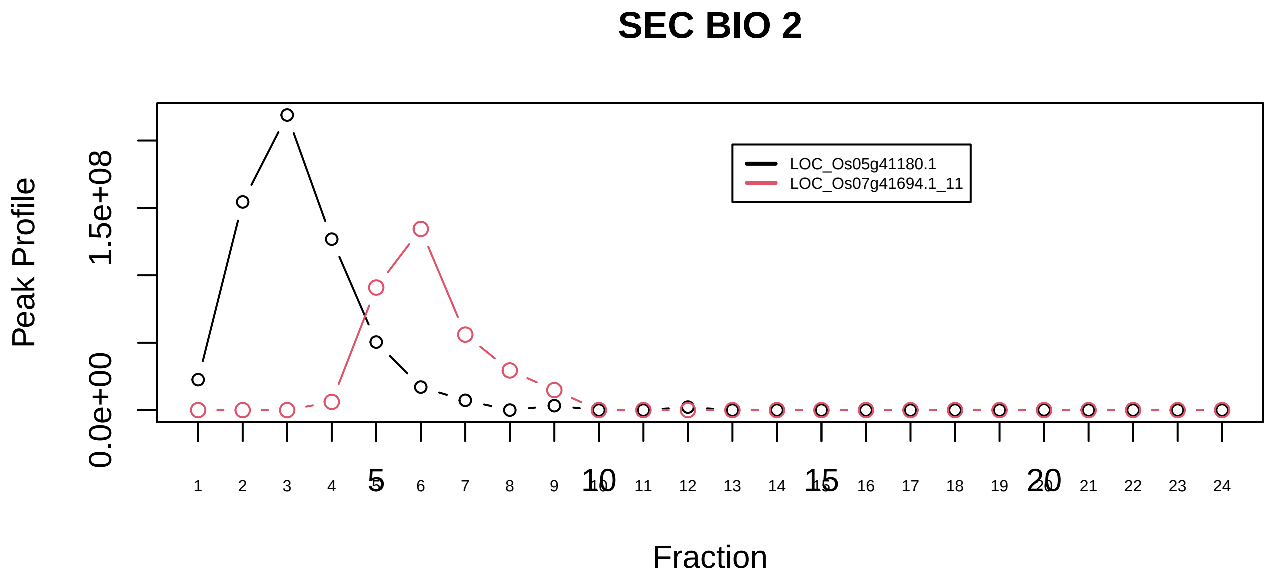

ANP32A−ANP32B−PSMA3−PTMA−SLC25A5 complex|7299−−sub_1

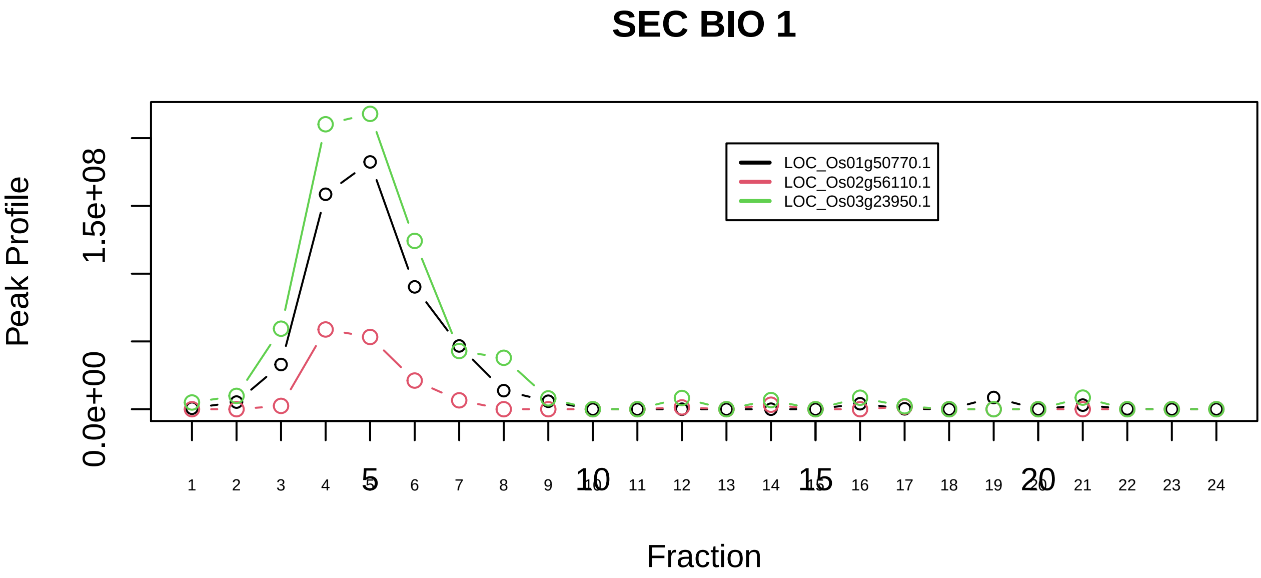

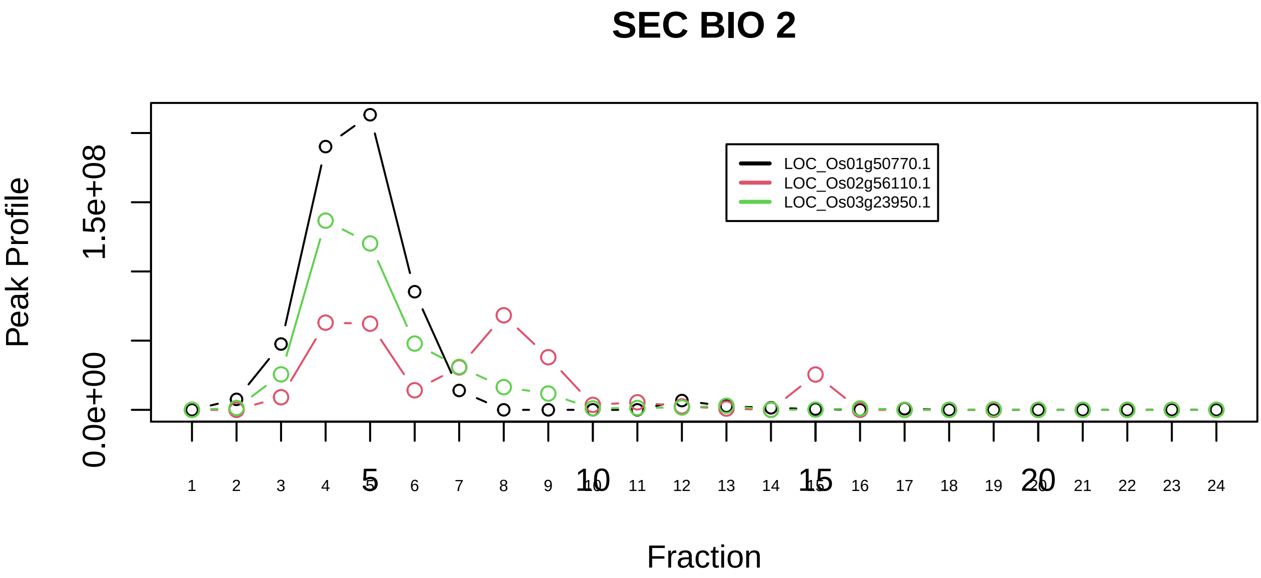

Gamma−BAR−AP1 complex|144−−all_prots_before_clustering_GPFdist

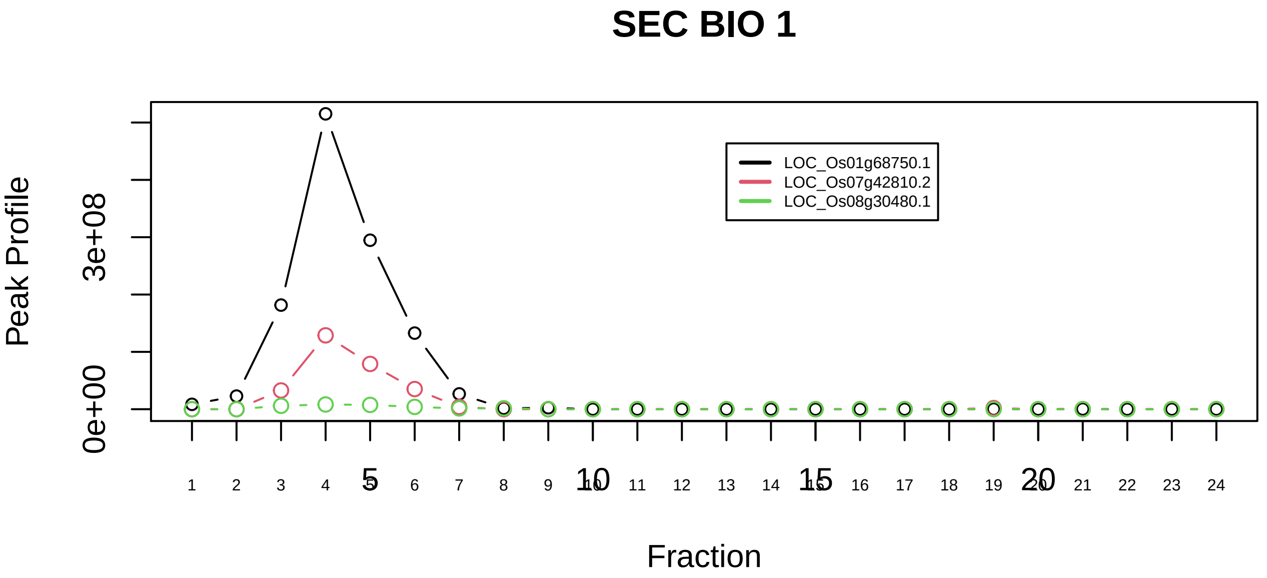

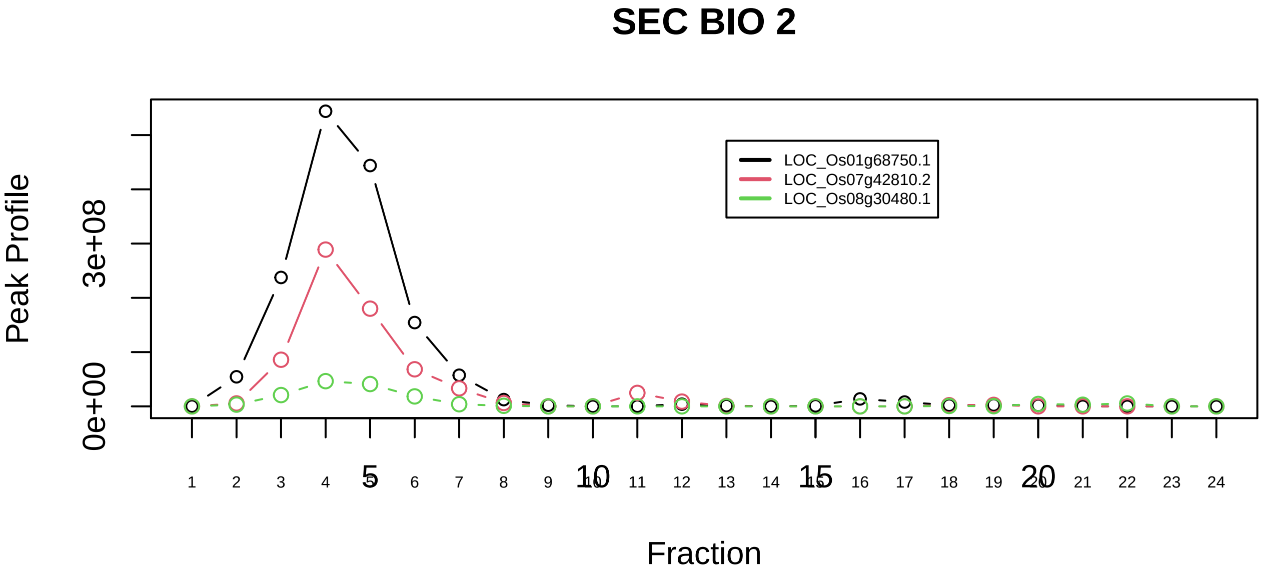

AP4 adaptor complex|67−−all_prots_before_clustering_GPFdist

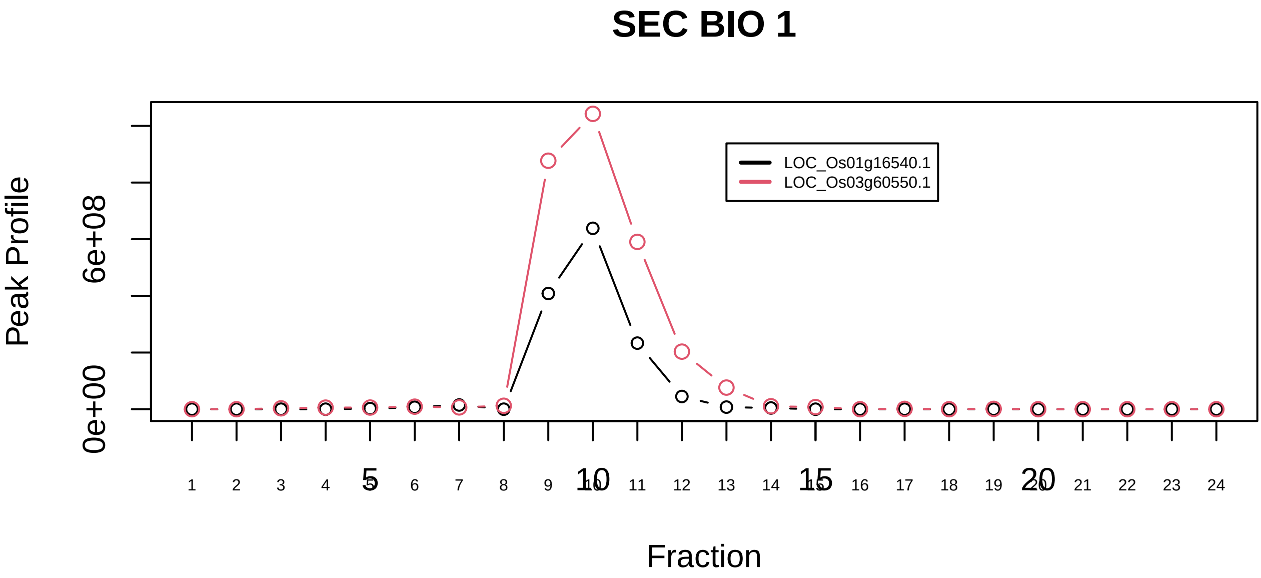

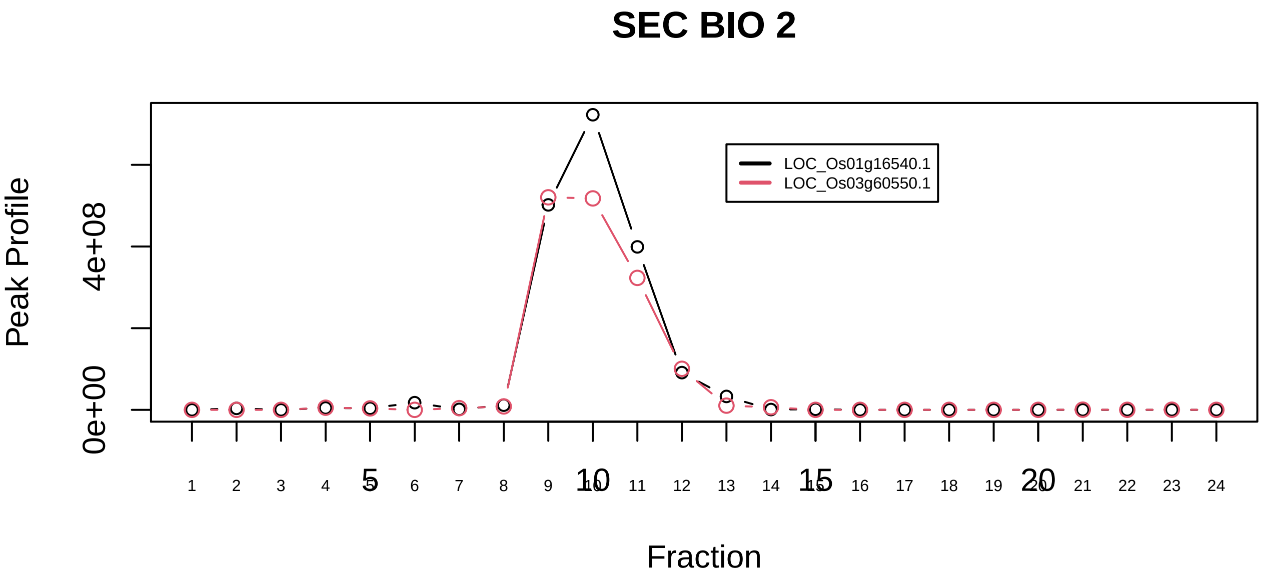

APPBP1−UBA3 complex|2162−−only_2_prots_before_clustering

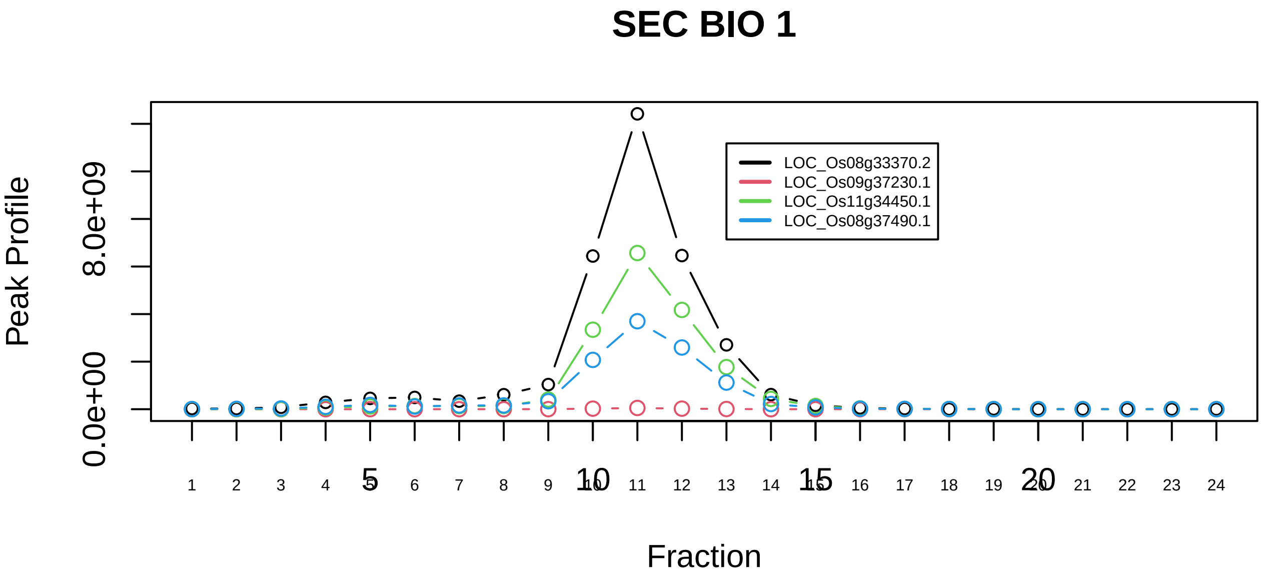

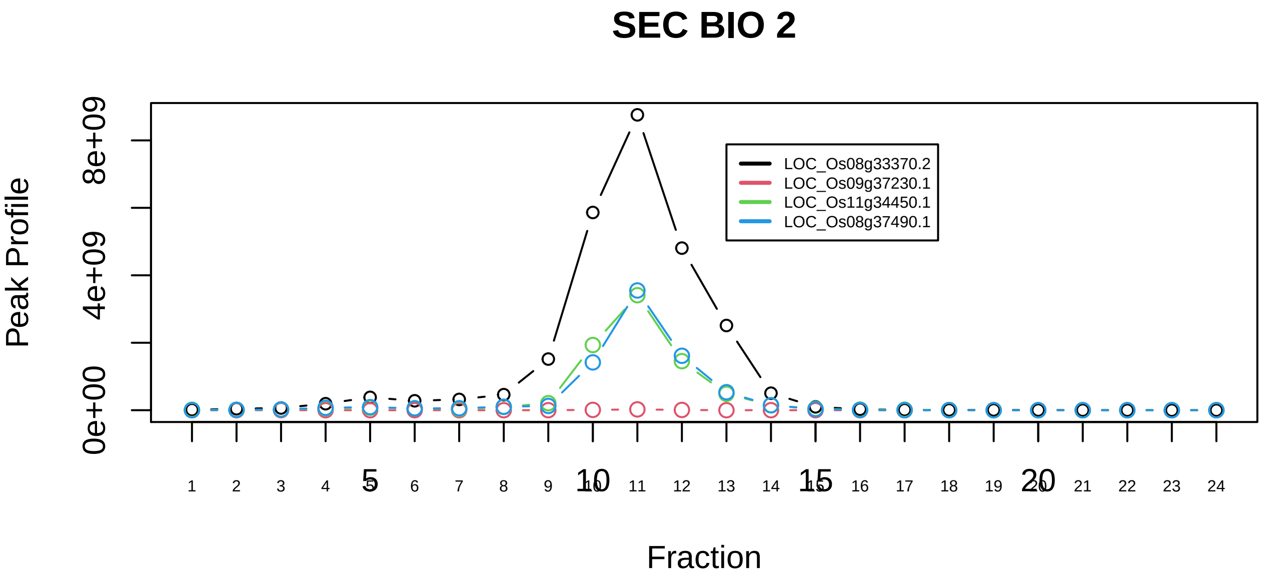

RAF1−MAP2K1−YWHAE complex|5873−−sub_1

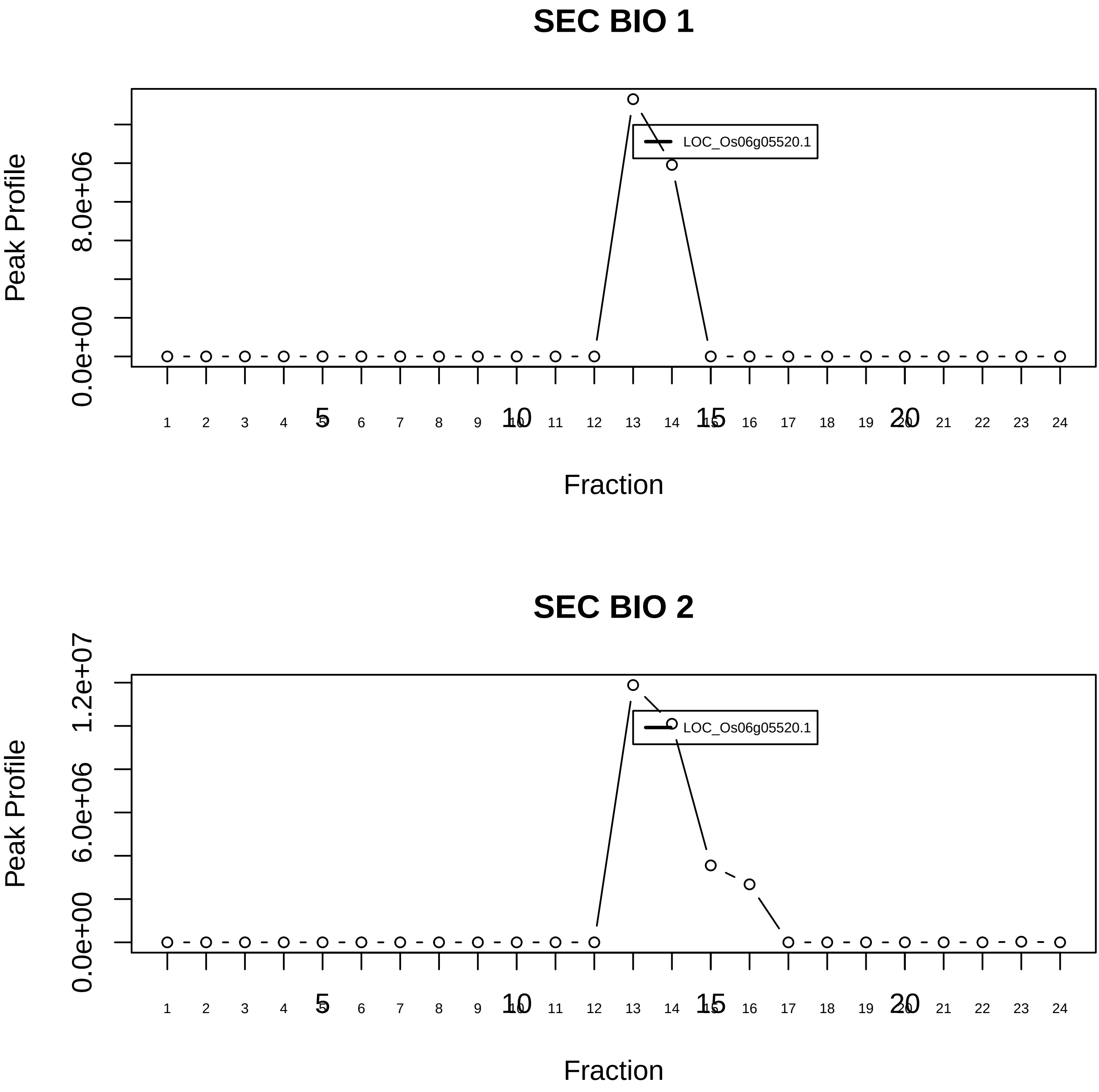

RAF1−MAP2K1−YWHAE complex|5873−−singleton_1

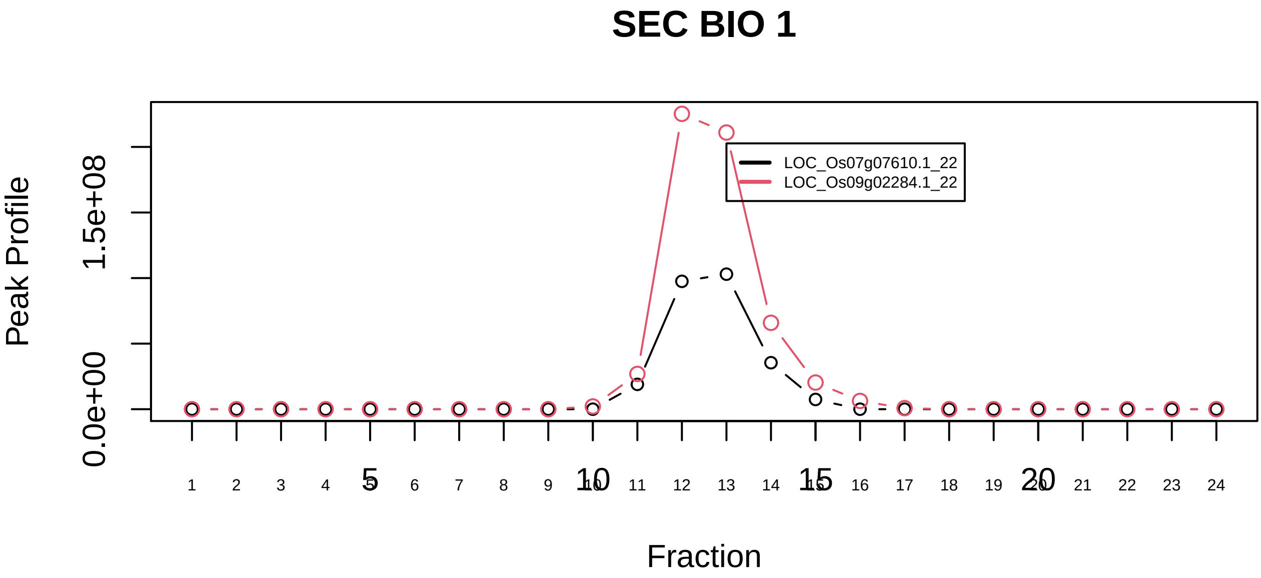

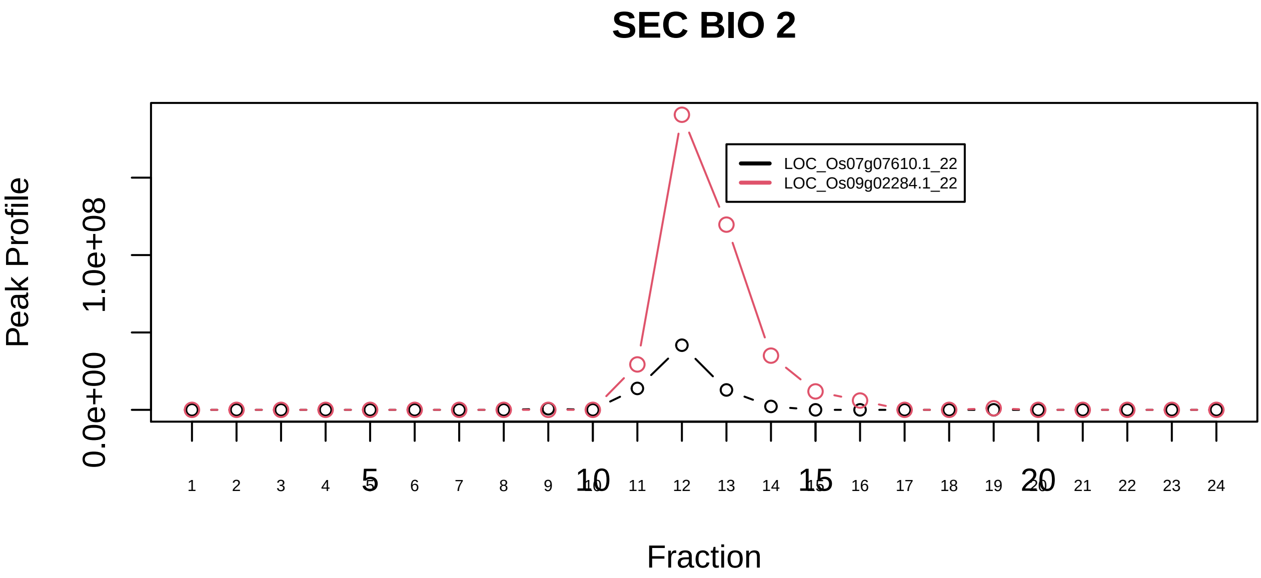

RNA polymerase II holoenzyme complex|103−−sub_1

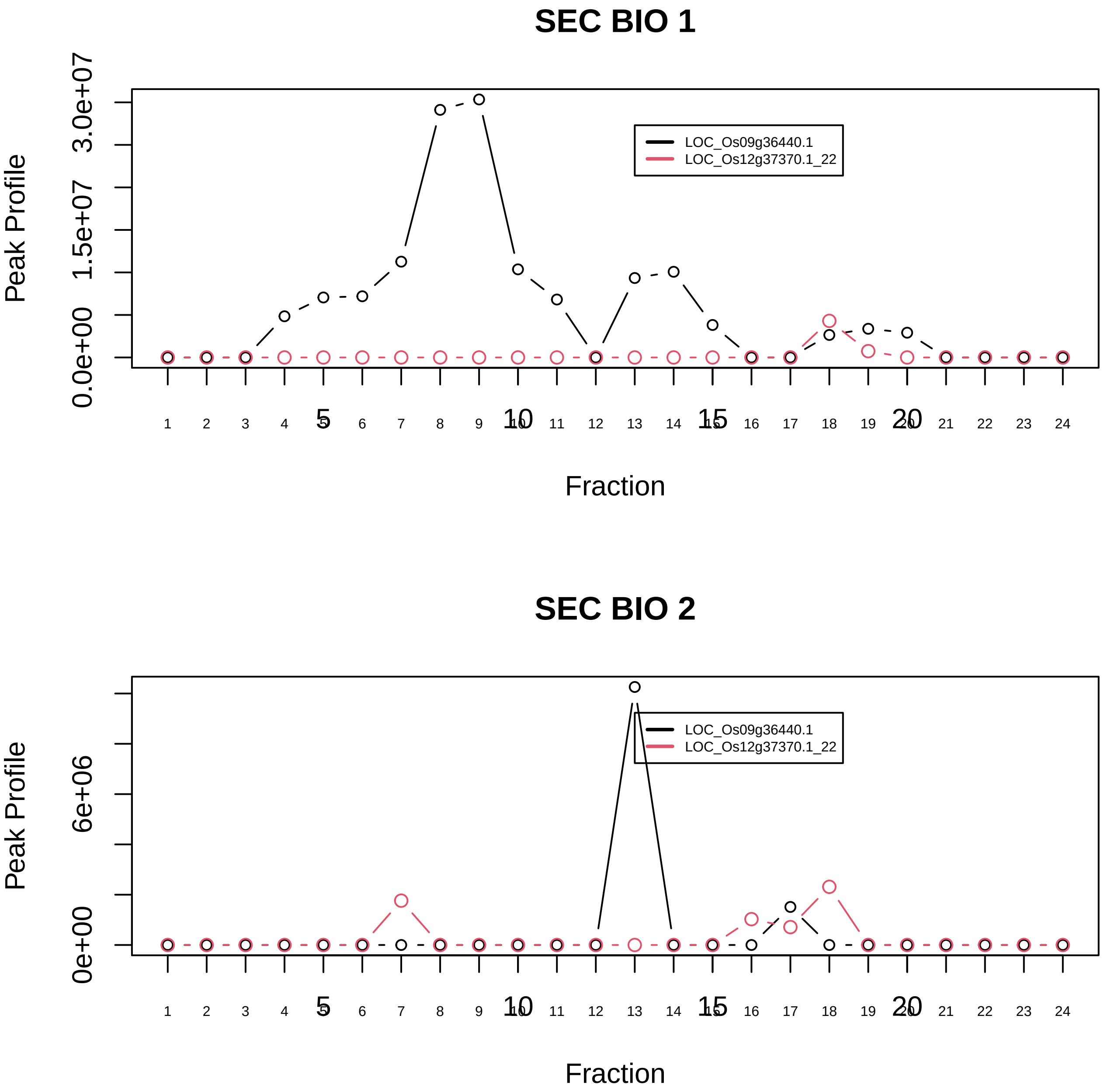

RNA polymerase II holoenzyme complex|103−−sub_2

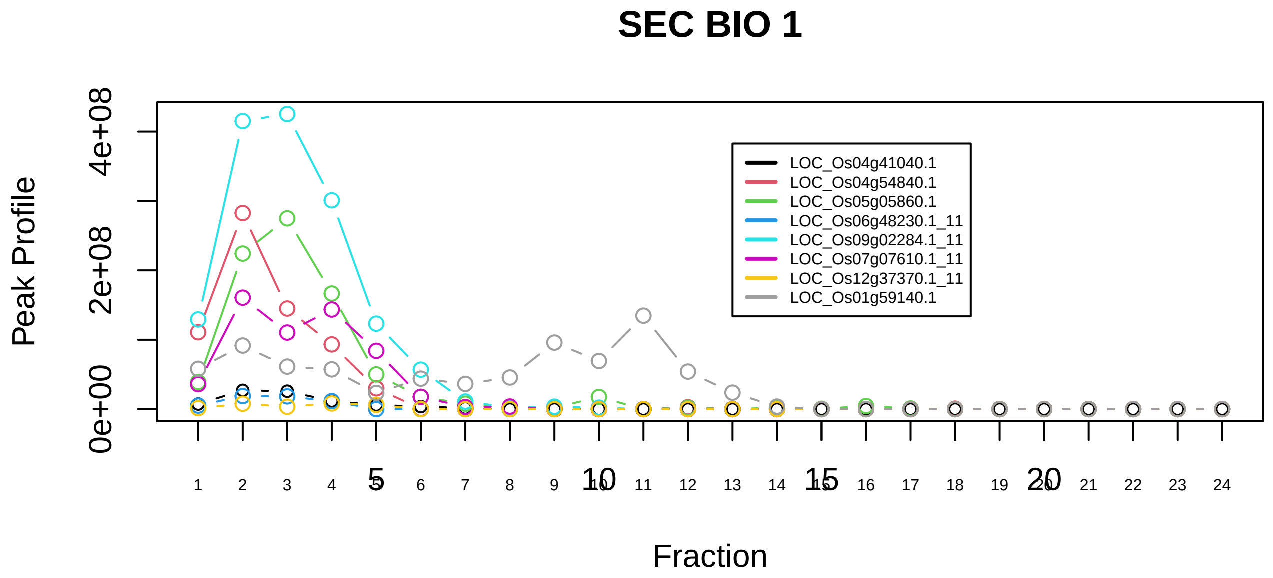

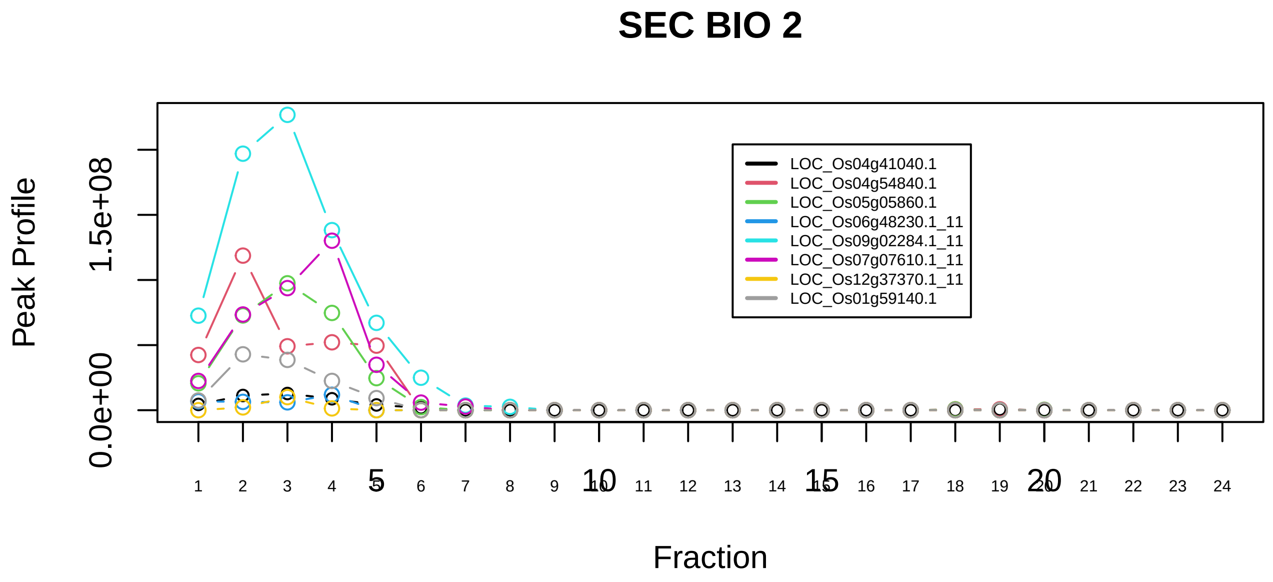

RNA polymerase II holoenzyme complex|103−−sub_3

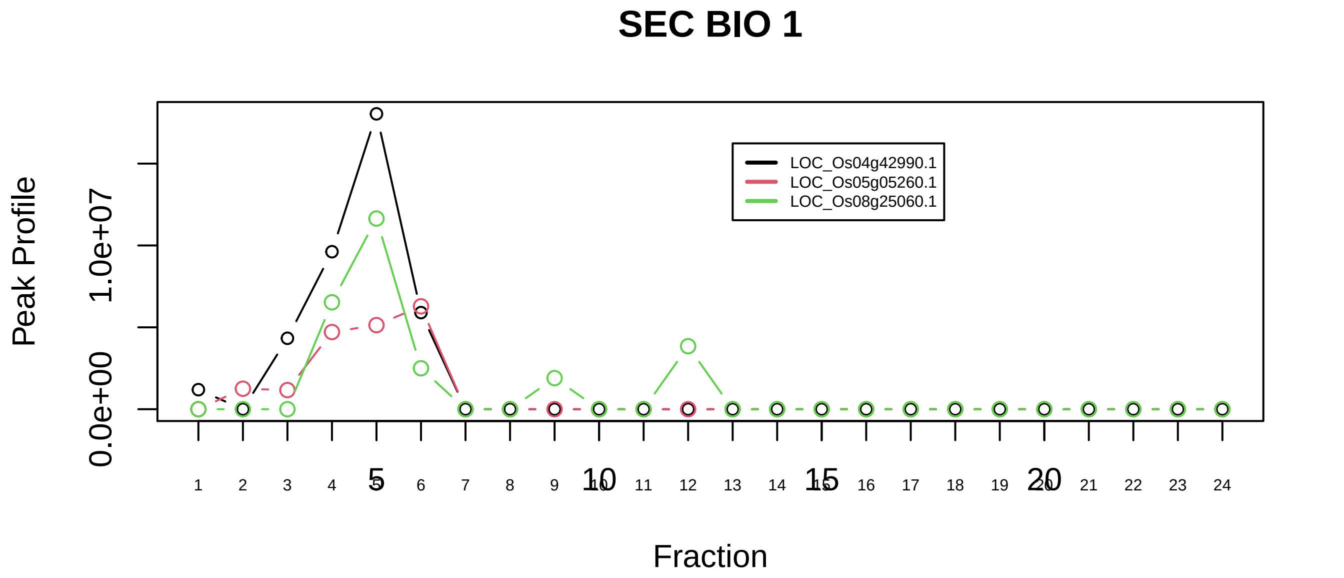

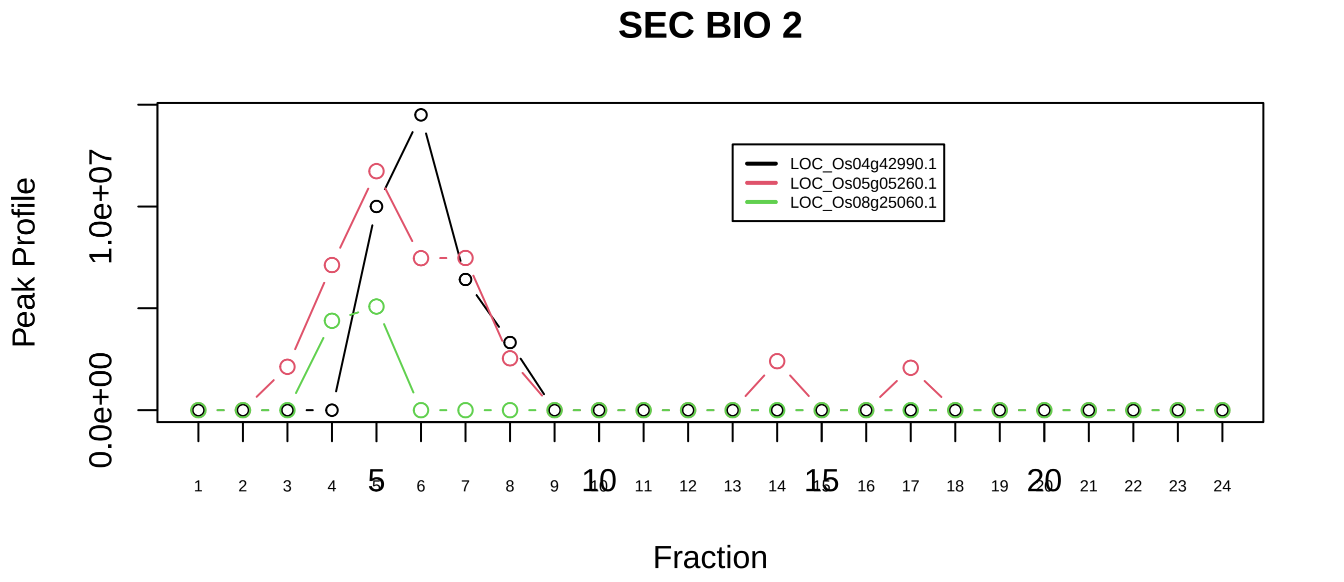

RNA polymerase II holoenzyme complex|103−−sub_4

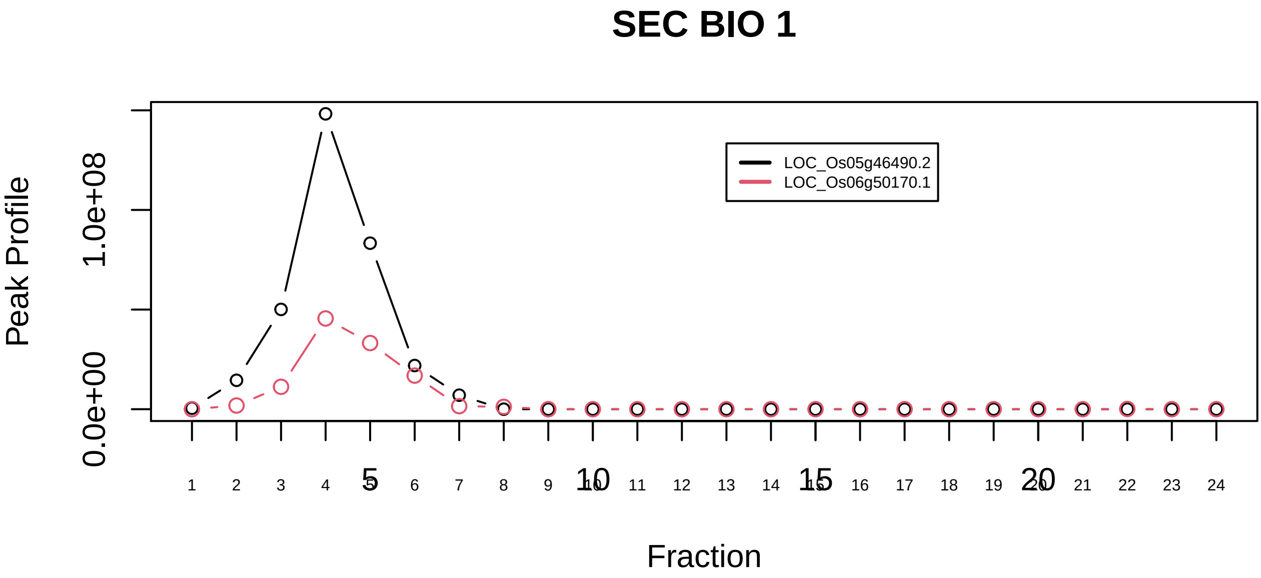

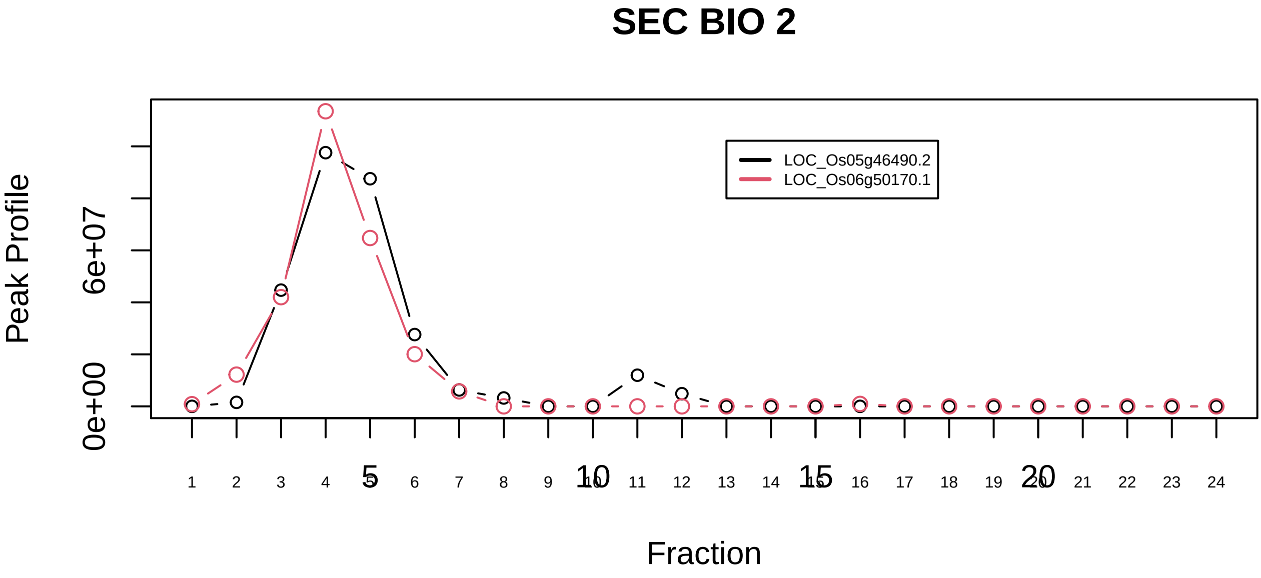

BRCC complex|5400−−only_2_prots_before_clustering

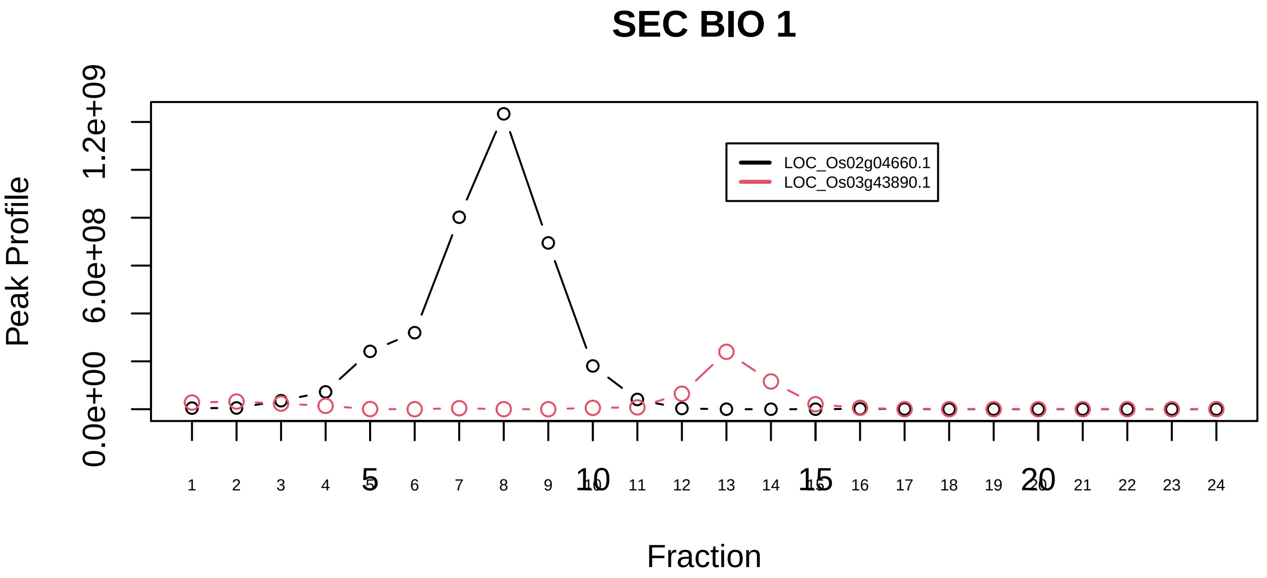

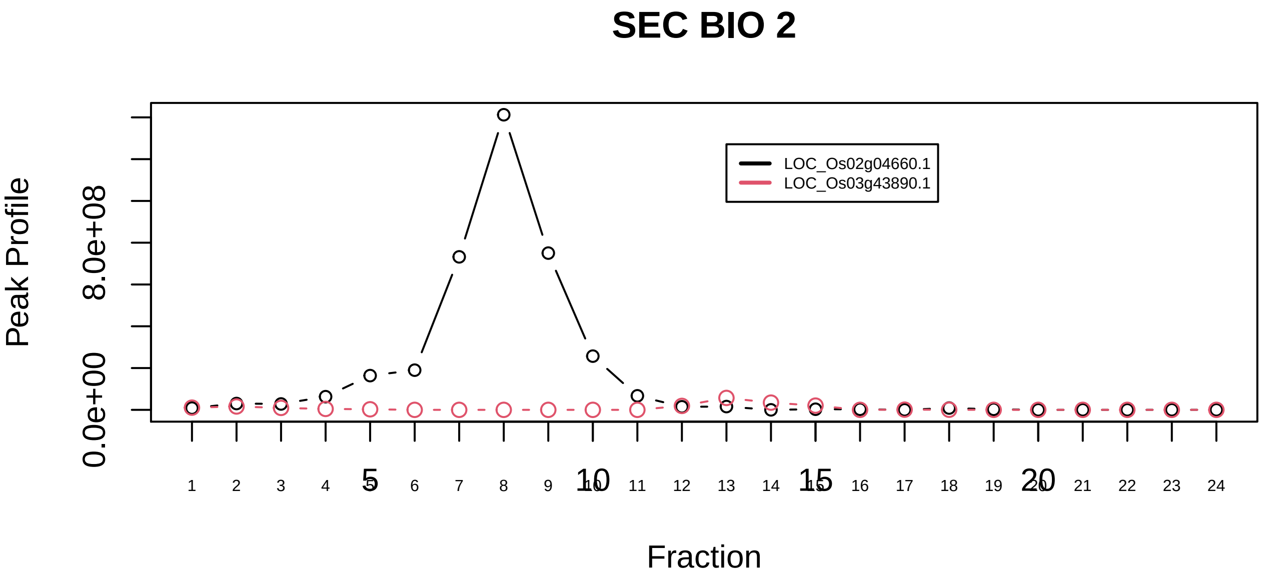

BRM−SIN3A complex|714−−only_2_prots_before_clustering

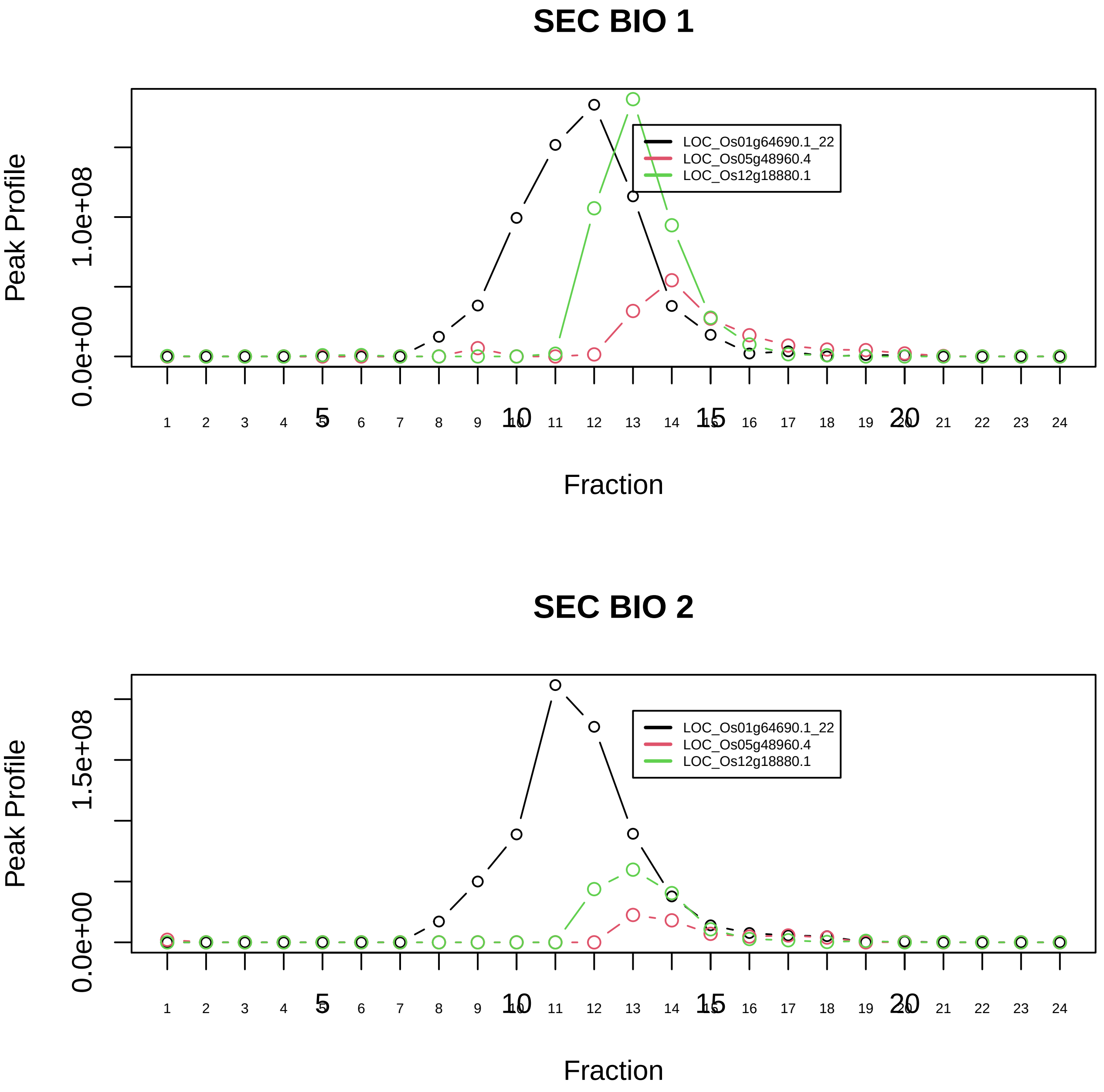

C complex spliceosome|1181−−sub_1

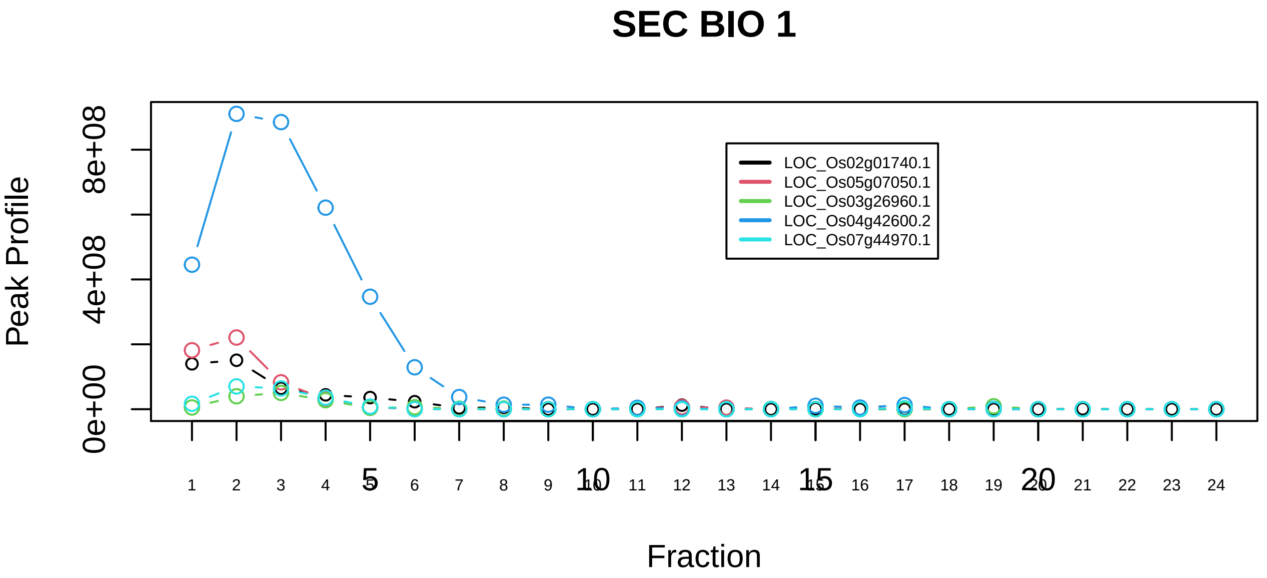

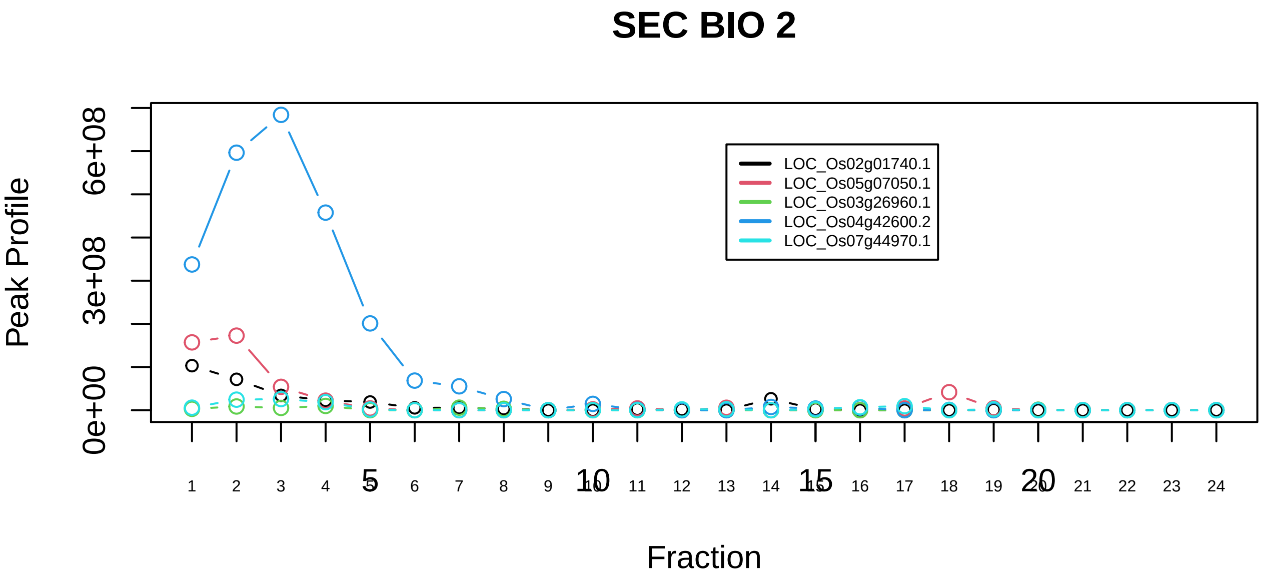

C complex spliceosome|1181−−sub_2

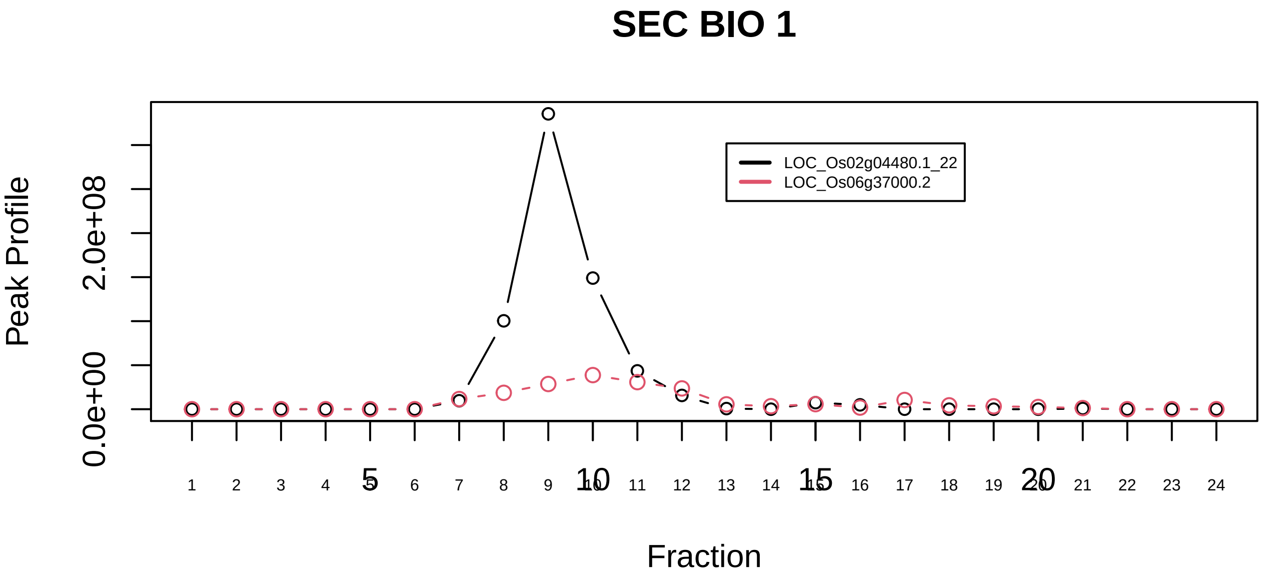

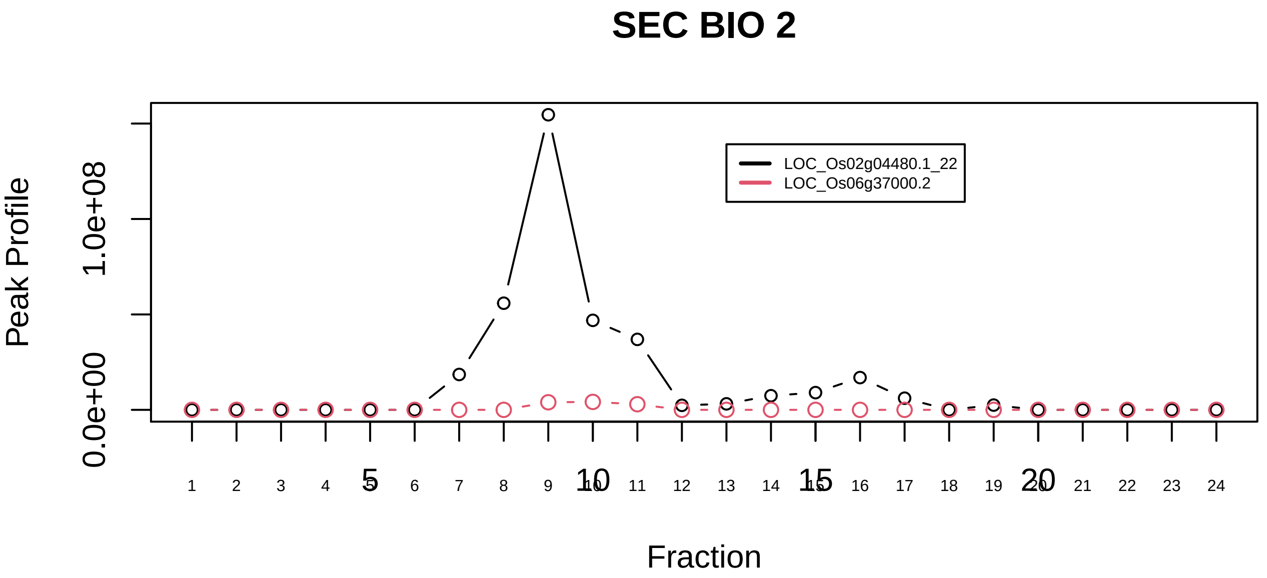

C complex spliceosome|1181−−sub_3

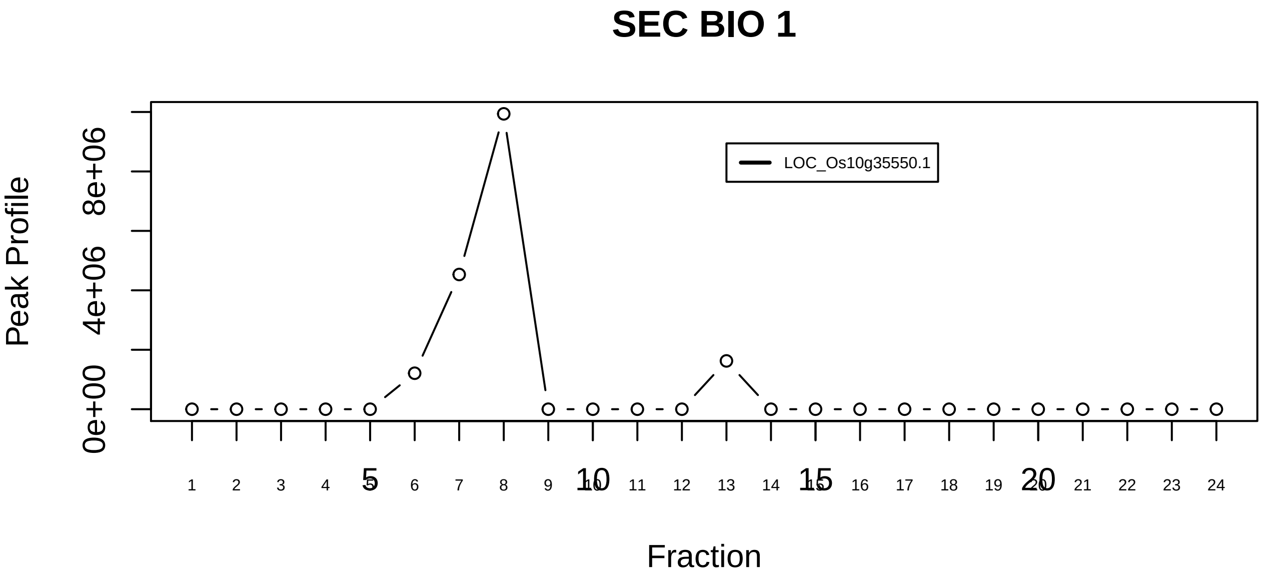

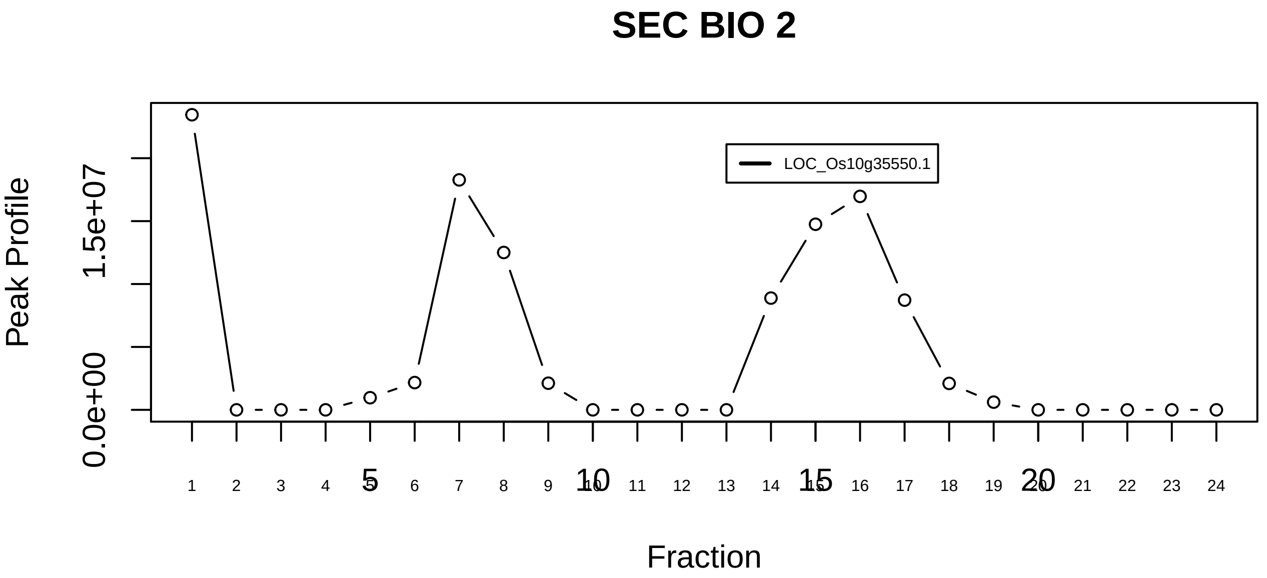

C complex spliceosome|1181−−singleton_1

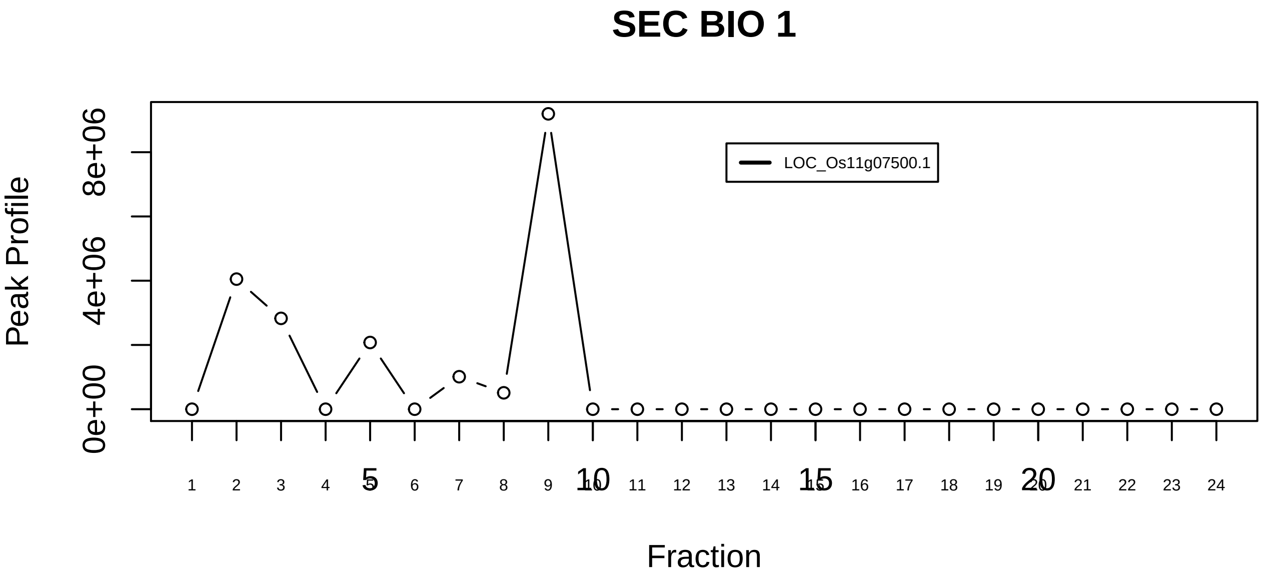

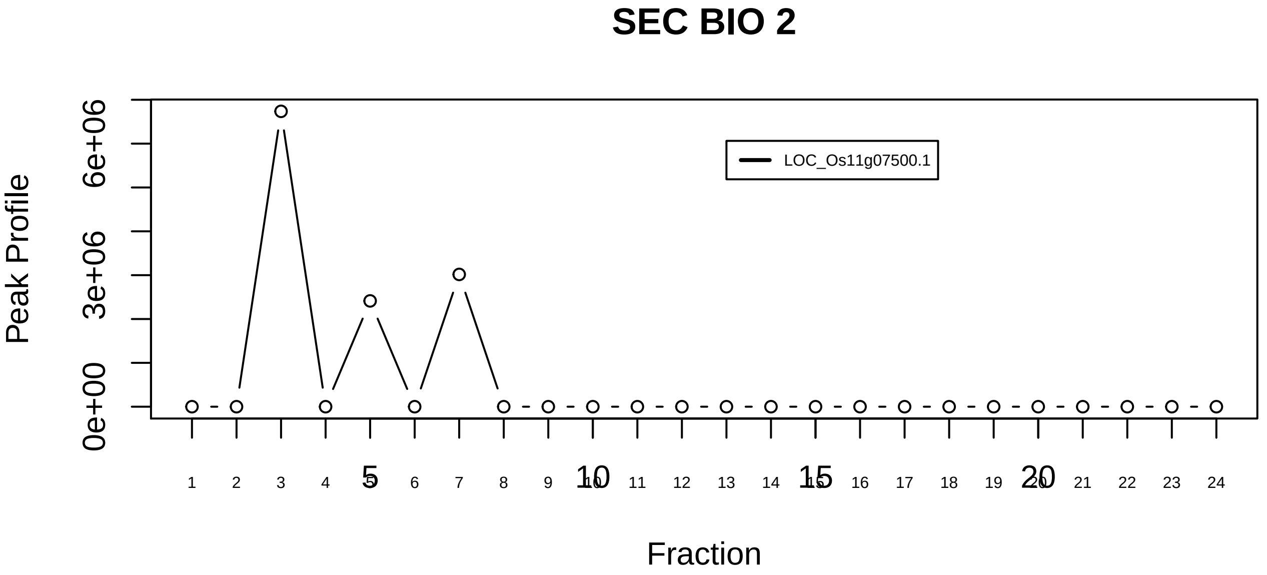

C complex spliceosome|1181−−singleton_2

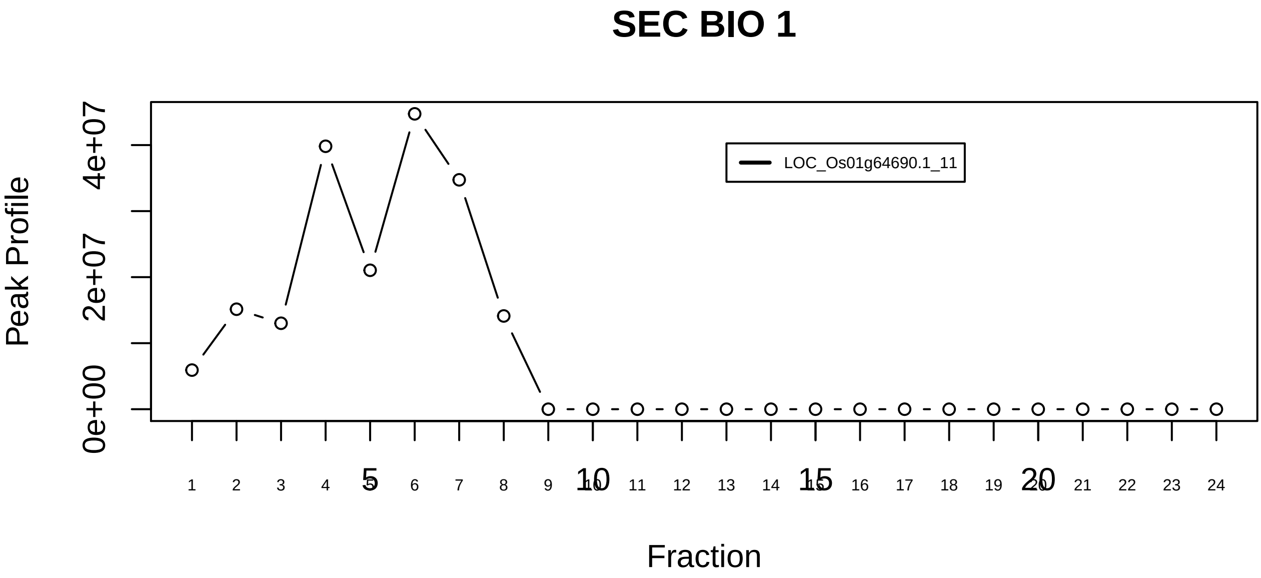

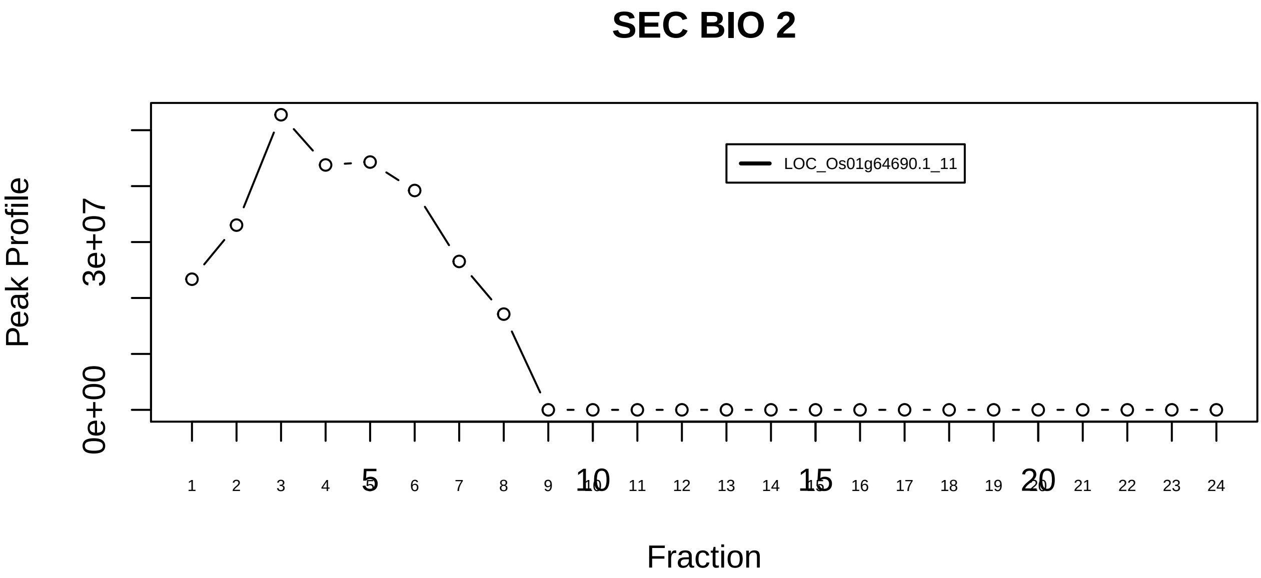

C complex spliceosome|1181−−singleton_3

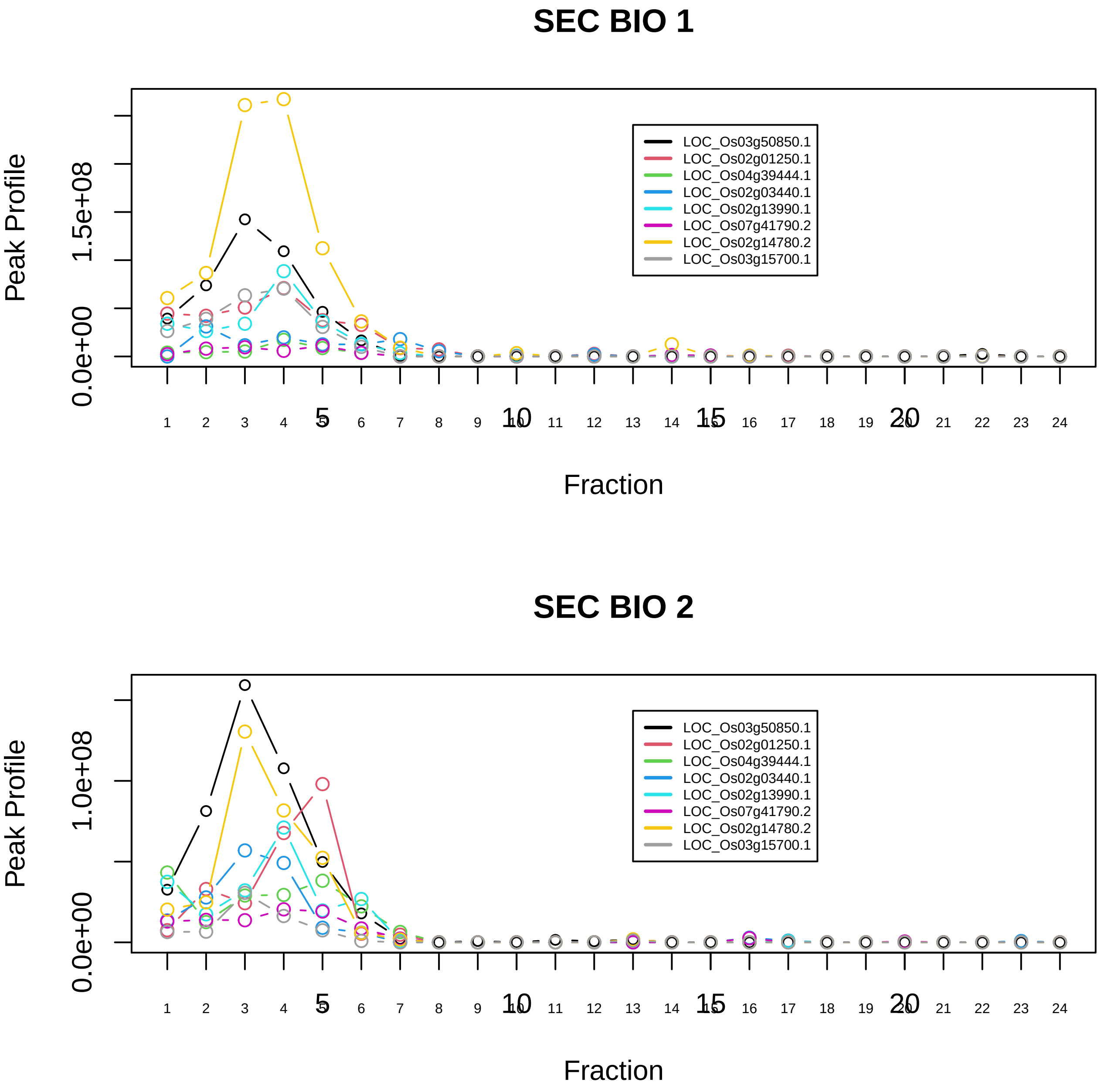

C complex spliceosome|1181−−sub_4

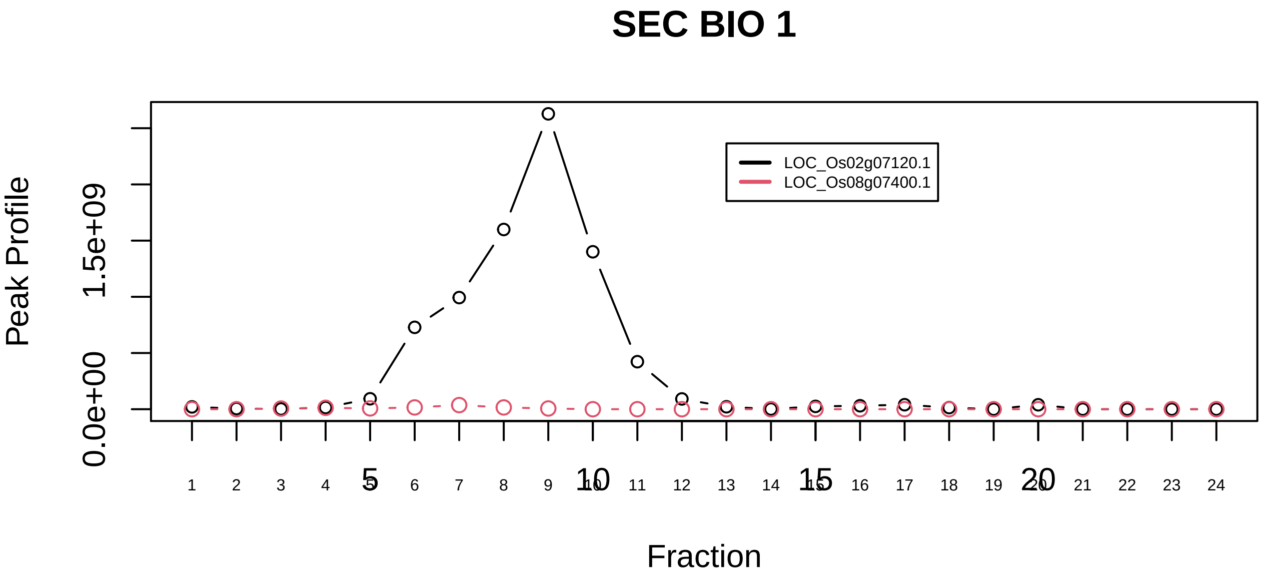

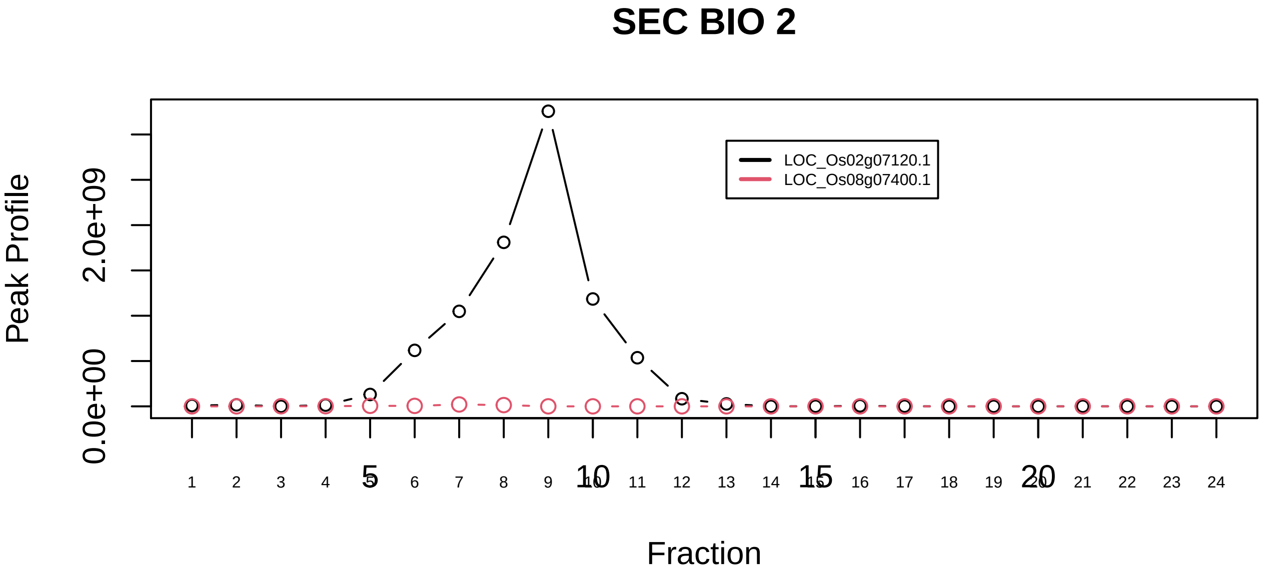

CAND1−CUL3−RBX1 complex|222−−only_2_prots_before_clustering

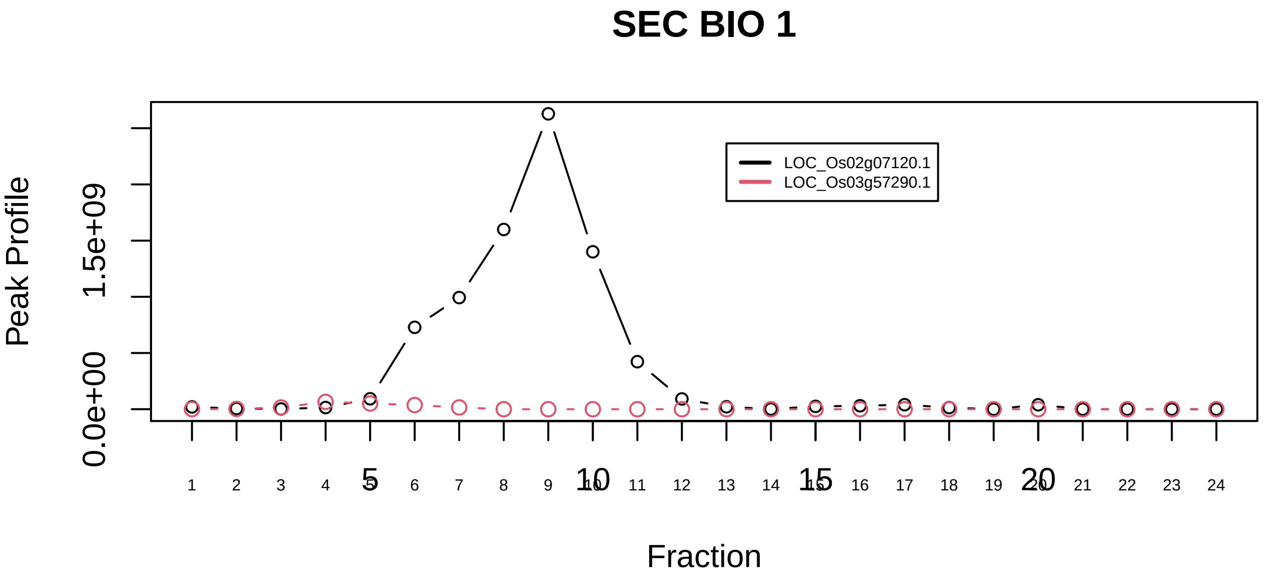

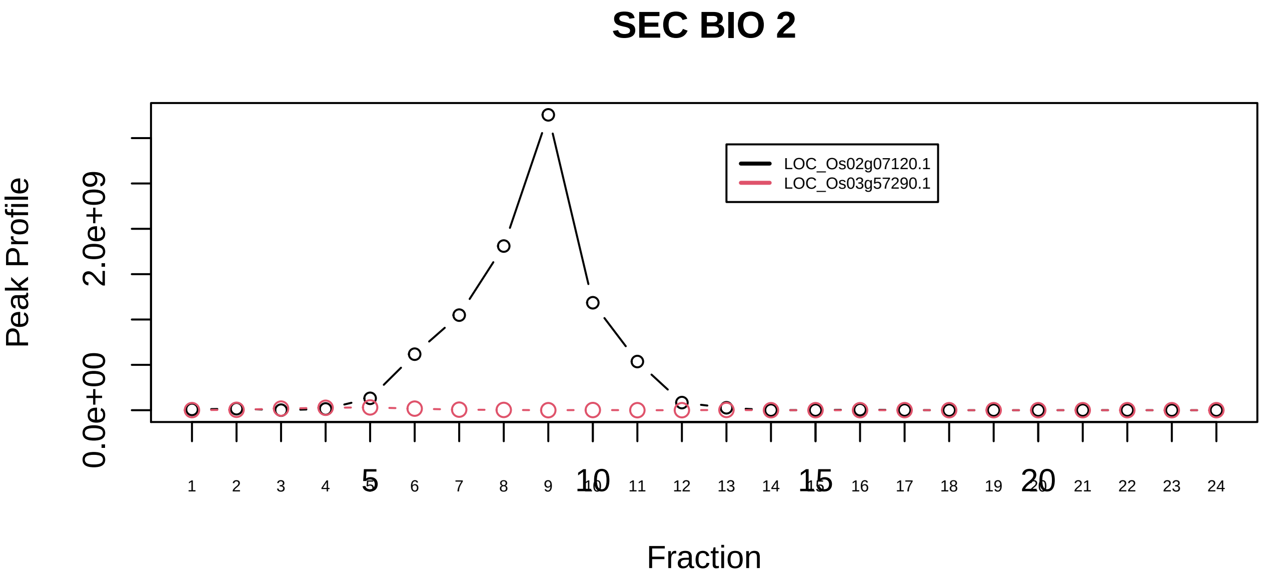

CAND1−CUL4B−RBX1 complex|224−−only_2_prots_before_clustering

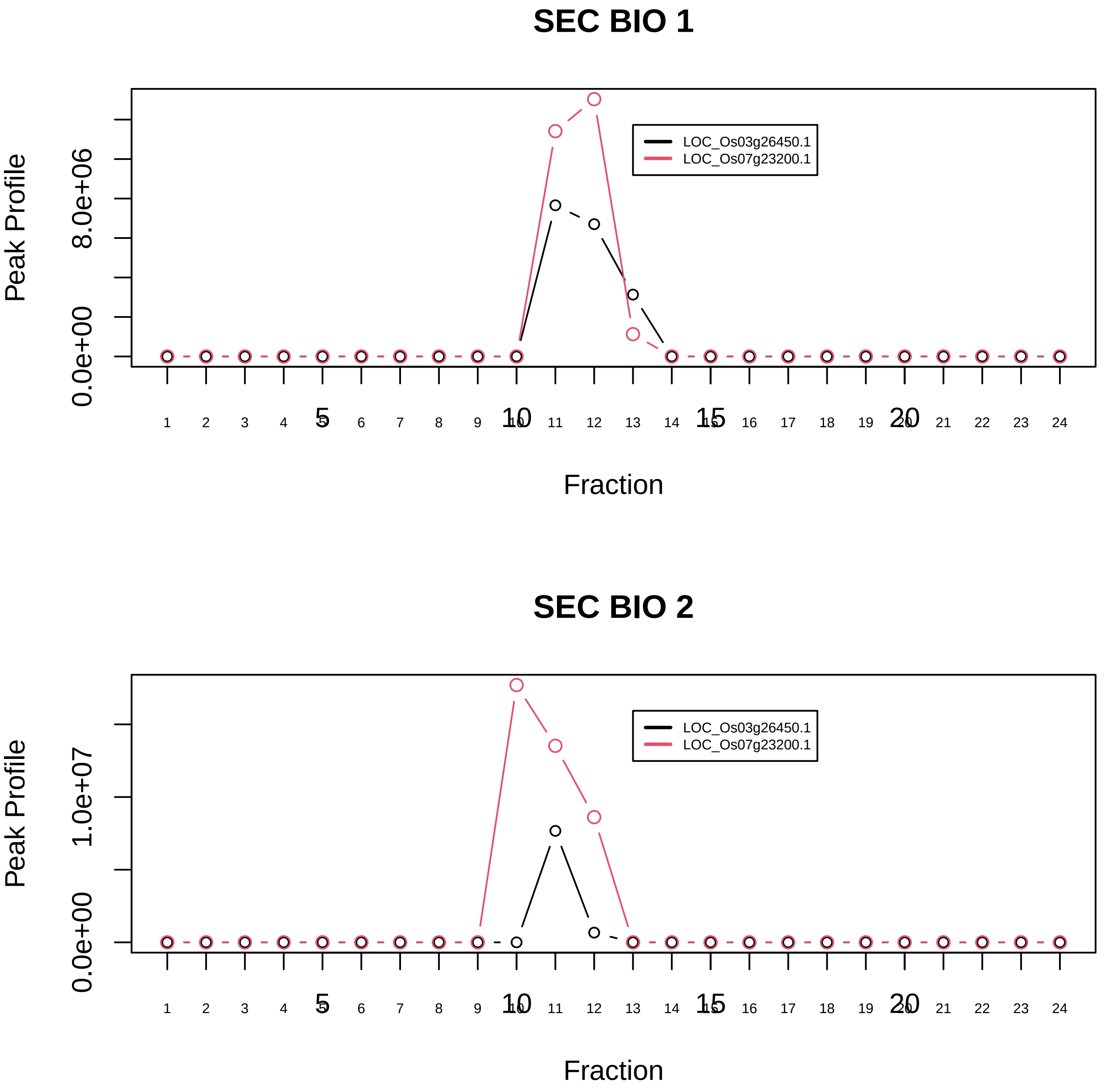

CAPZalpha−CAPZbeta complex|6483−−only_2_prots_before_clustering

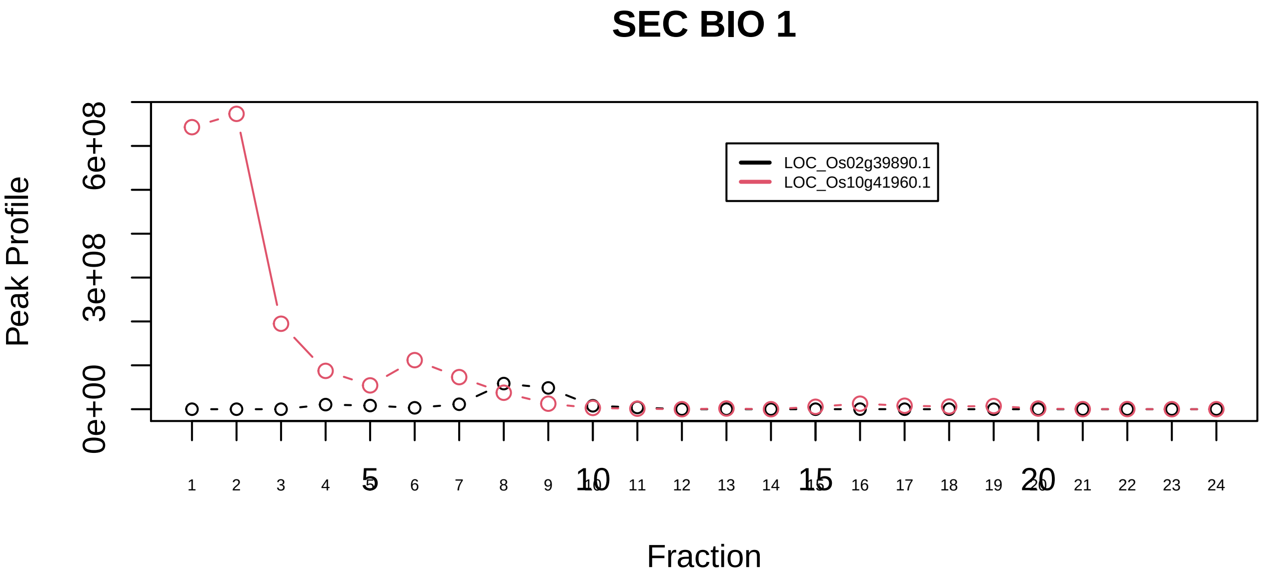

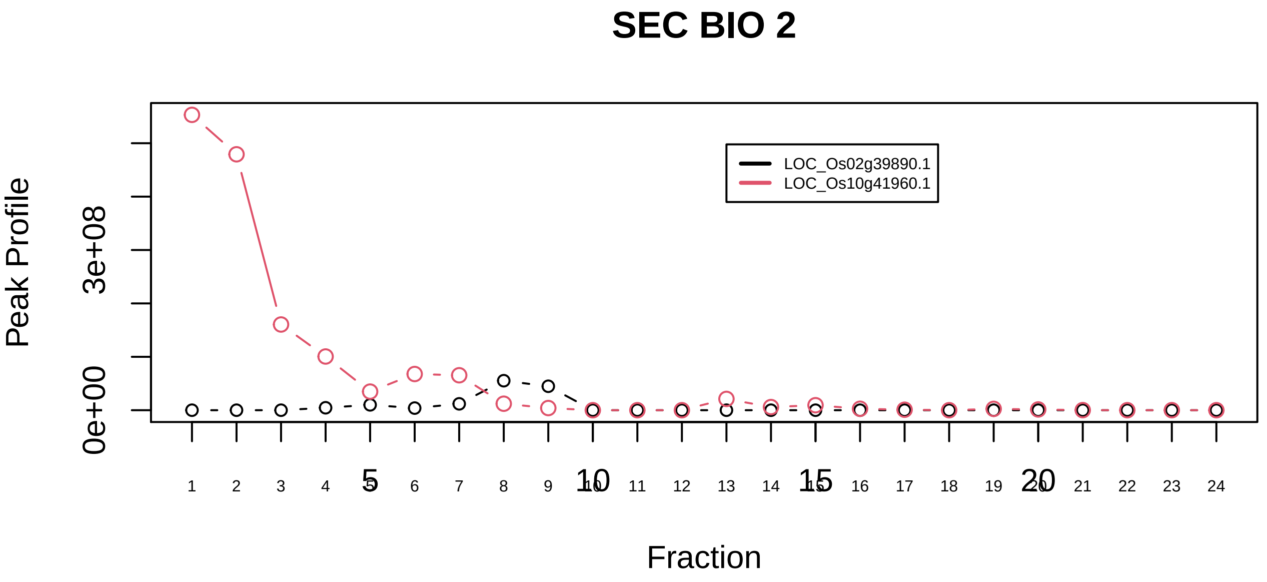

CBP80/20− dependent translation complex|6129−−only_2_prots_before_clustering

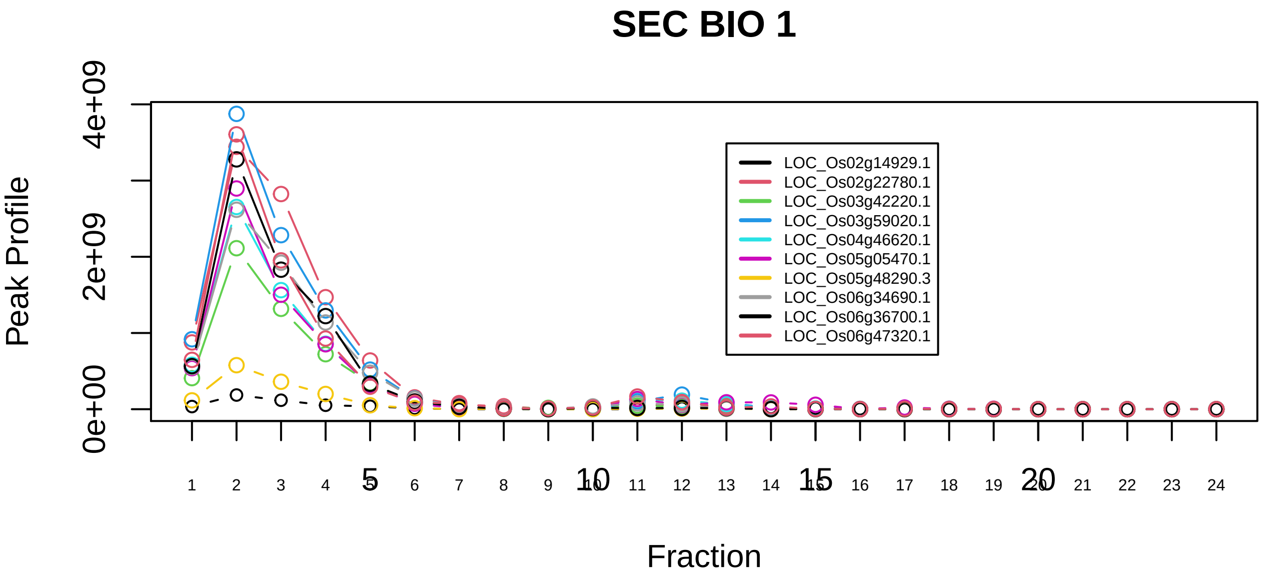

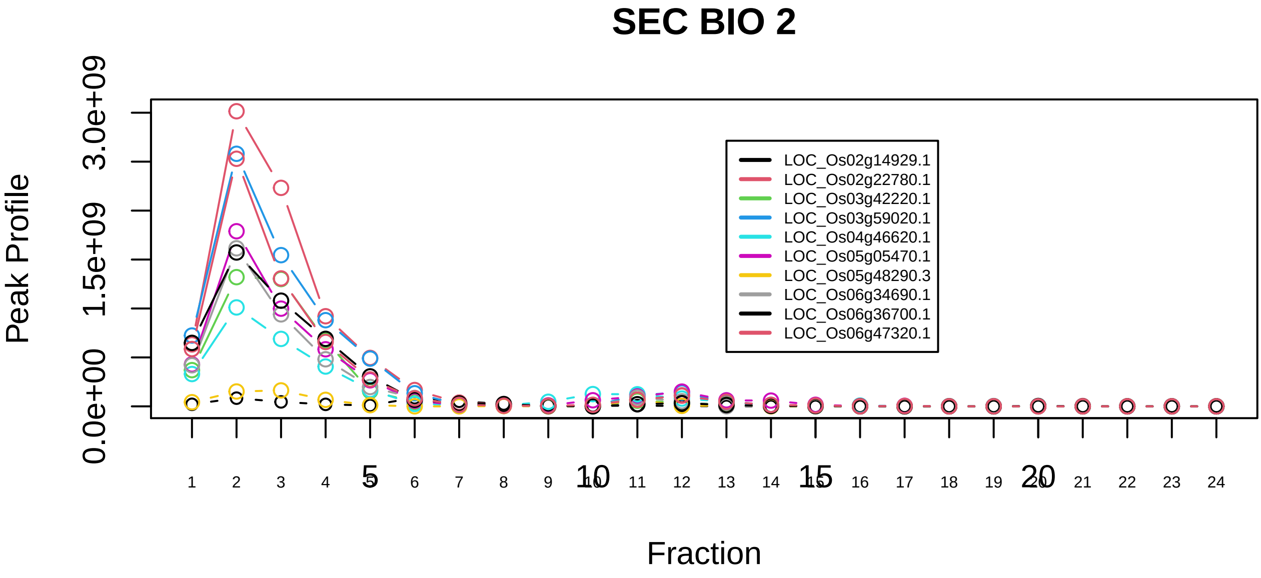

CCT complex (chaperonin containing TCP1 complex)|126−−all_prots_before_clustering_GPFdist

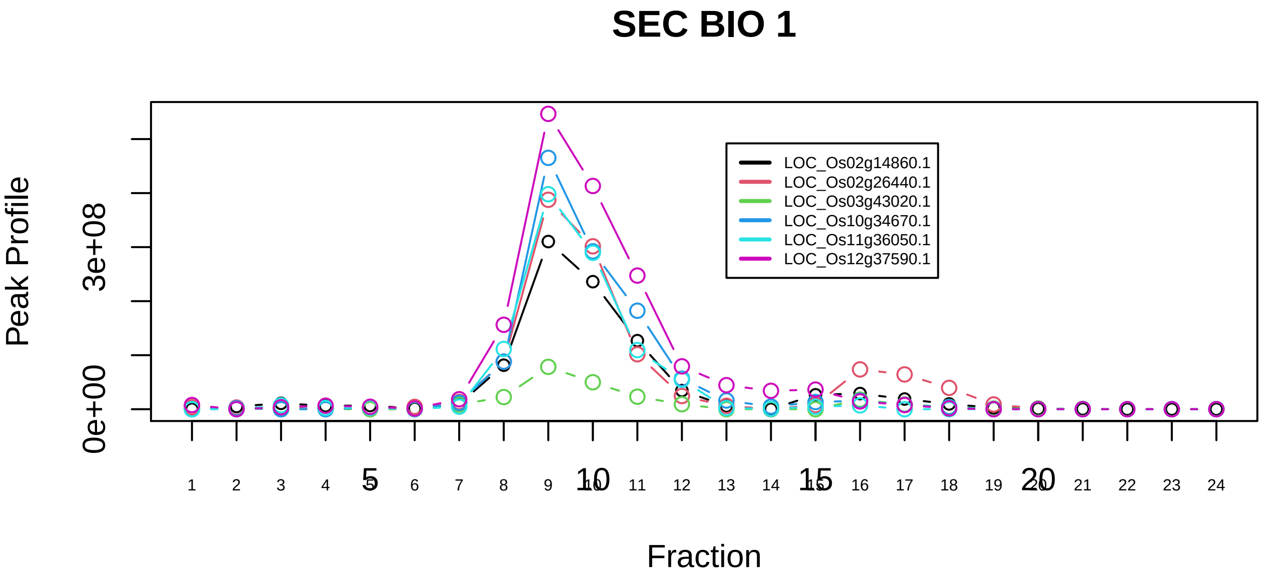

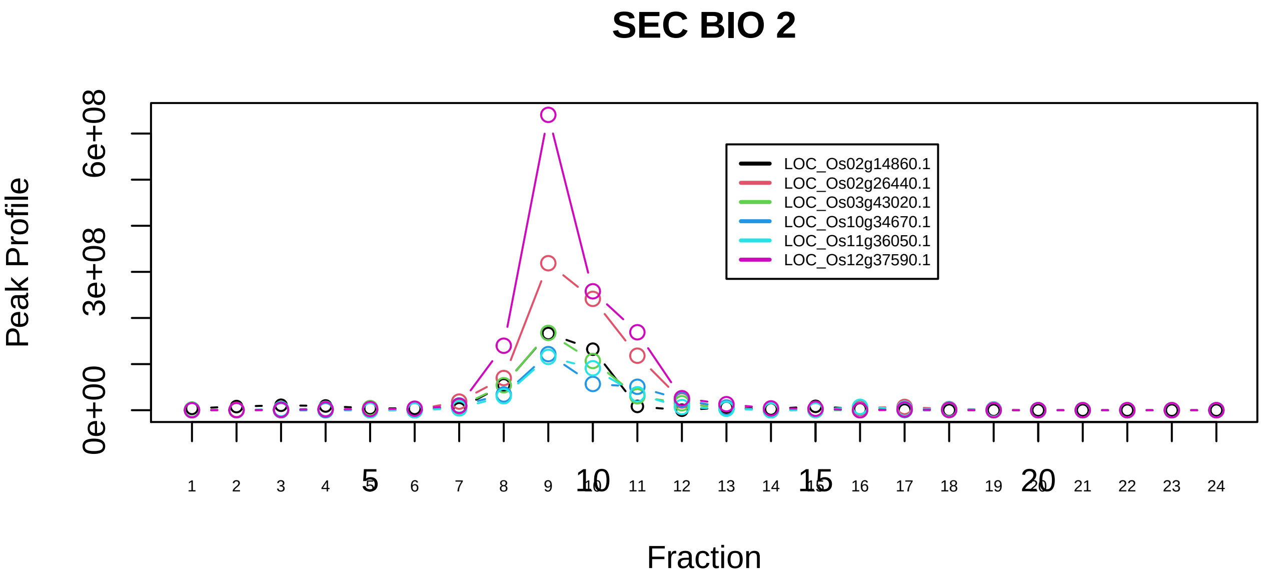

Prefoldin complex|33−−all_prots_before_clustering_GPFdist

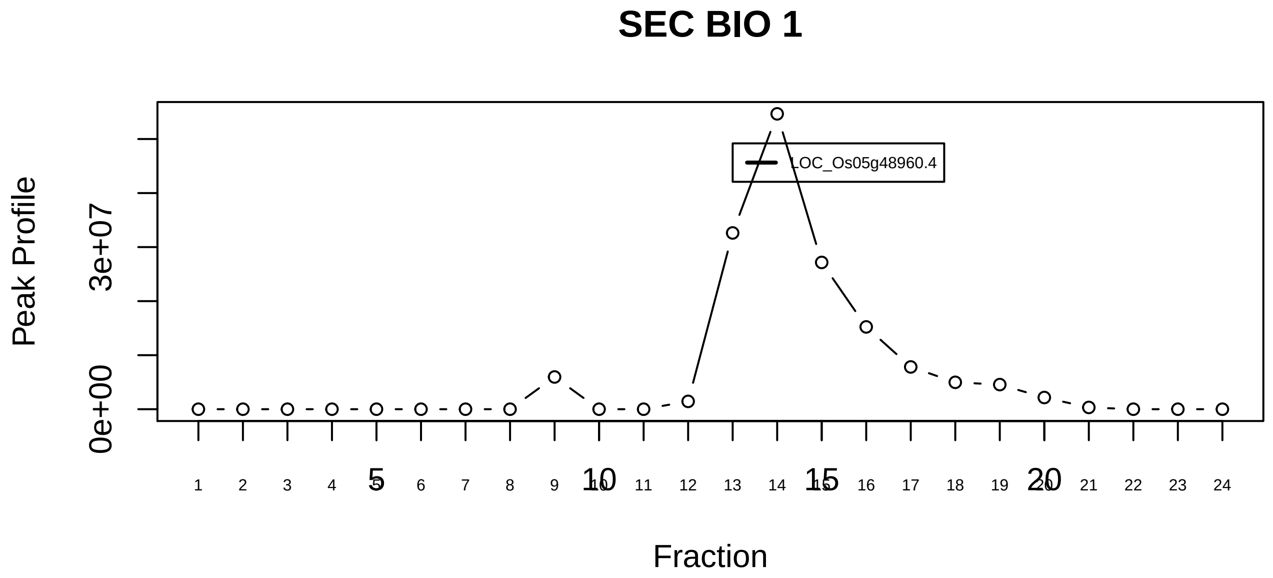

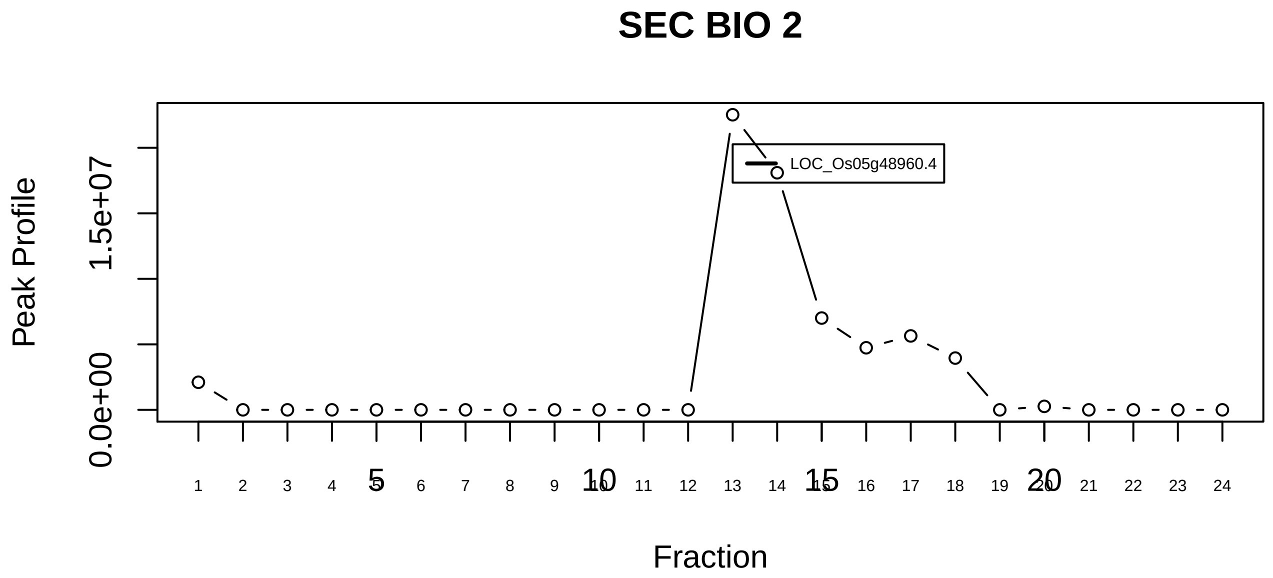

CF IIAm complex (Cleavage factor IIAm complex)|1141−−singleton_1

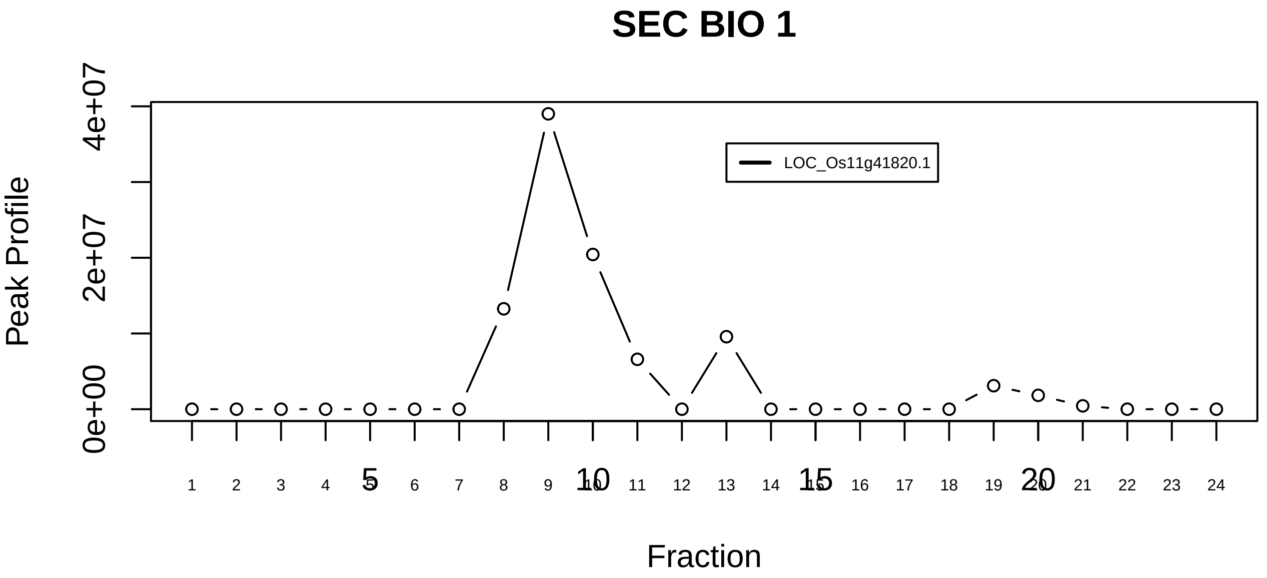

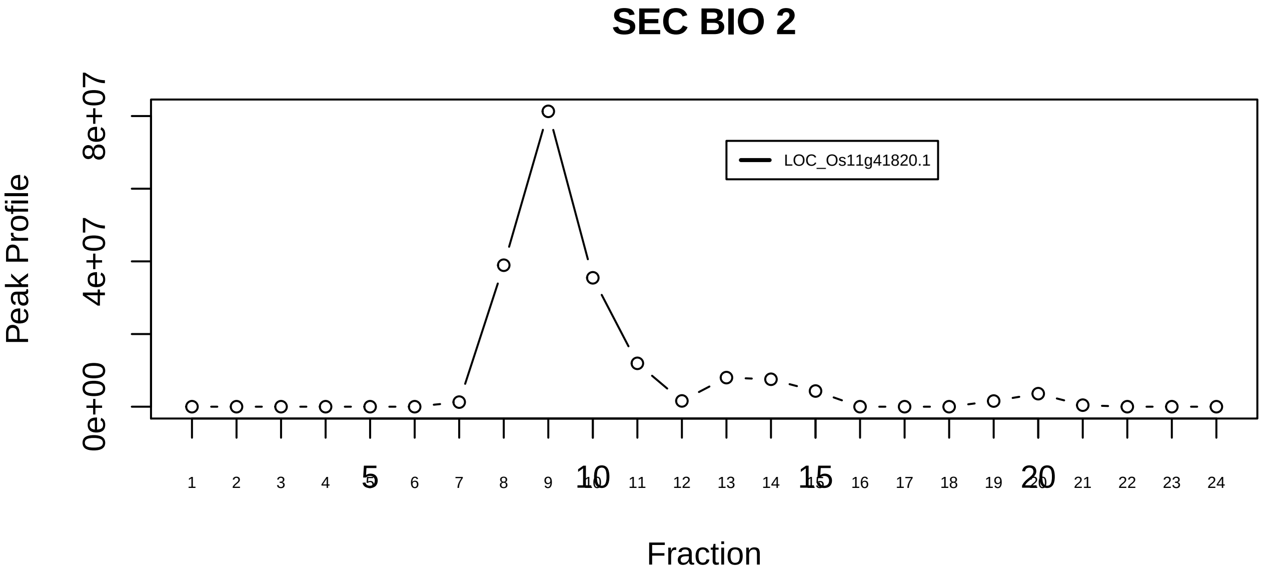

CF IIAm complex (Cleavage factor IIAm complex)|1141−−singleton_2

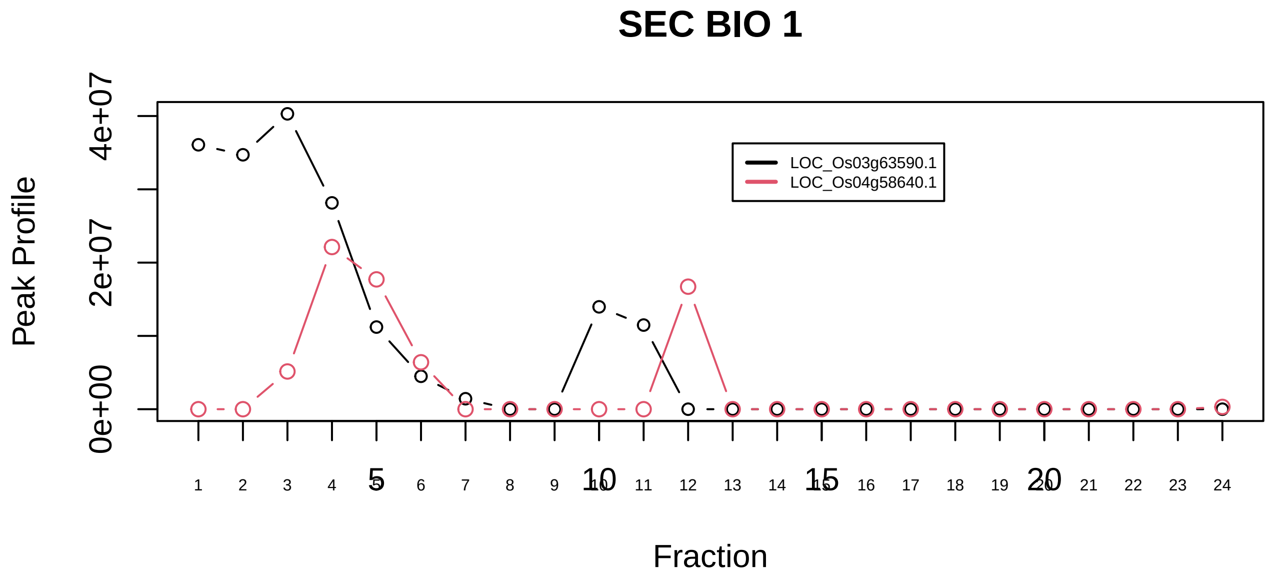

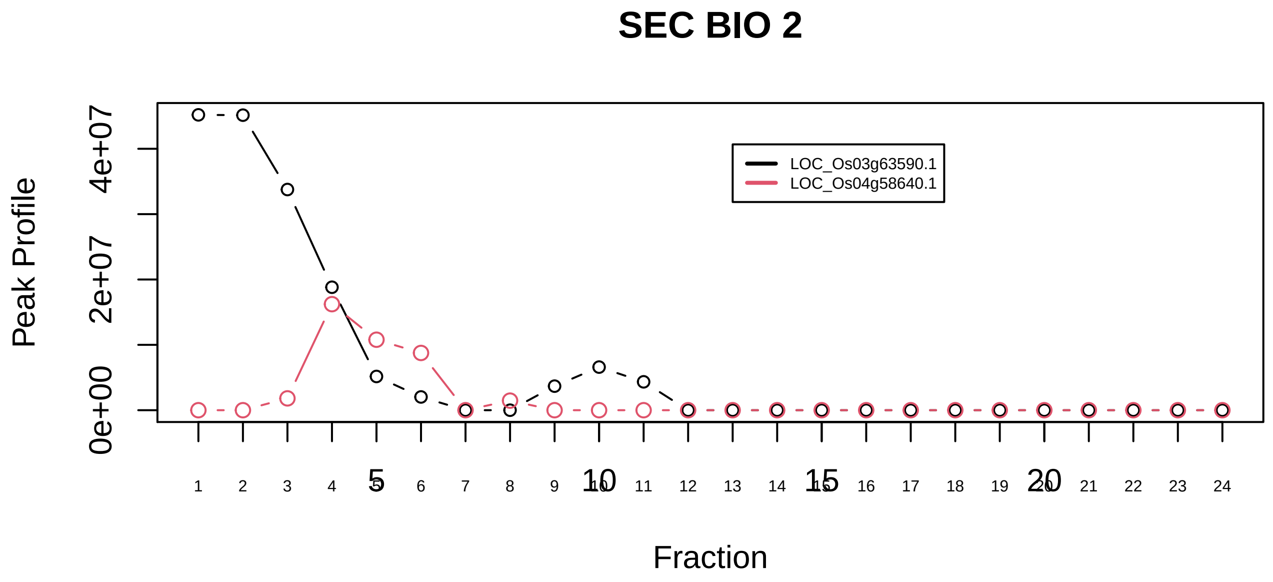

CF IIAm complex (Cleavage factor IIAm complex)|1141−−sub_1

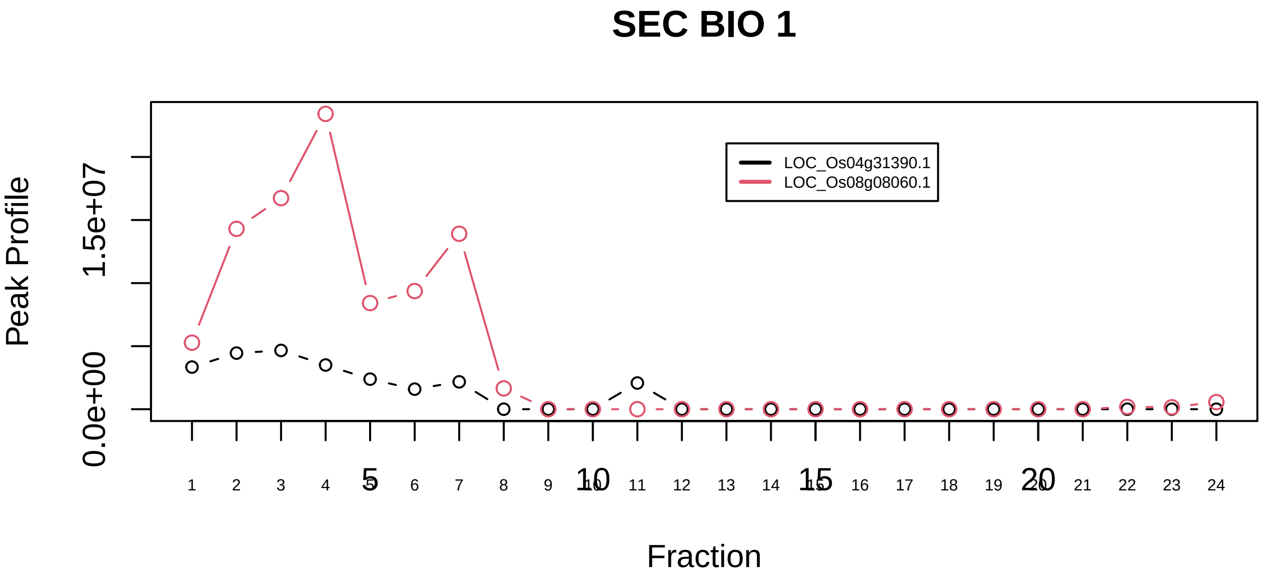

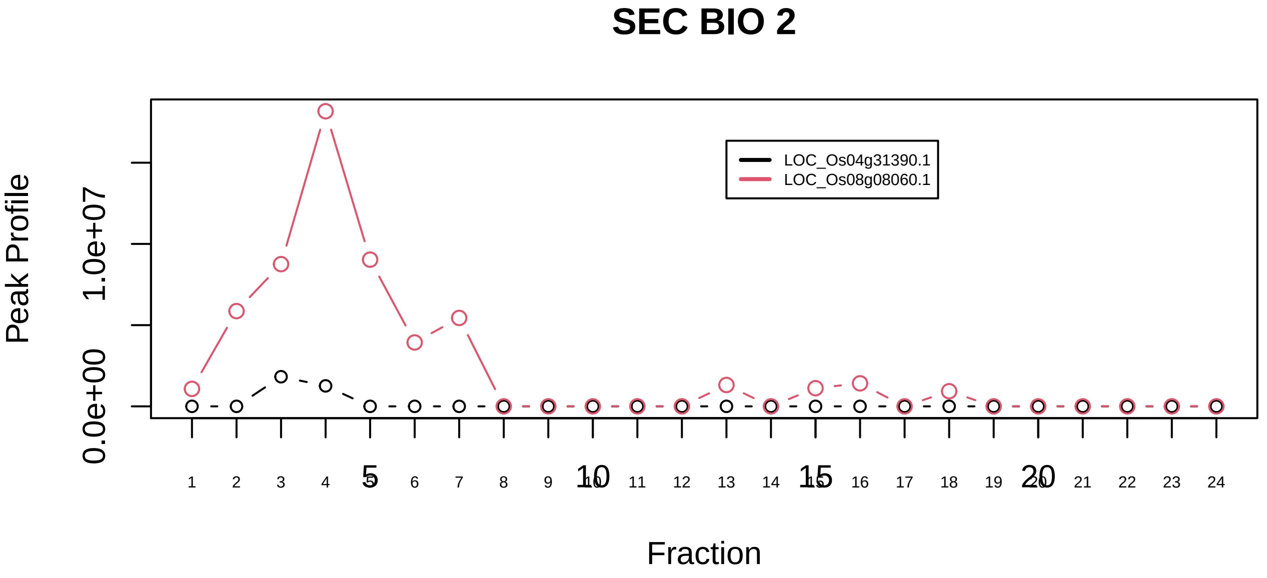

HOPS complex|6389−−only_2_prots_before_clustering

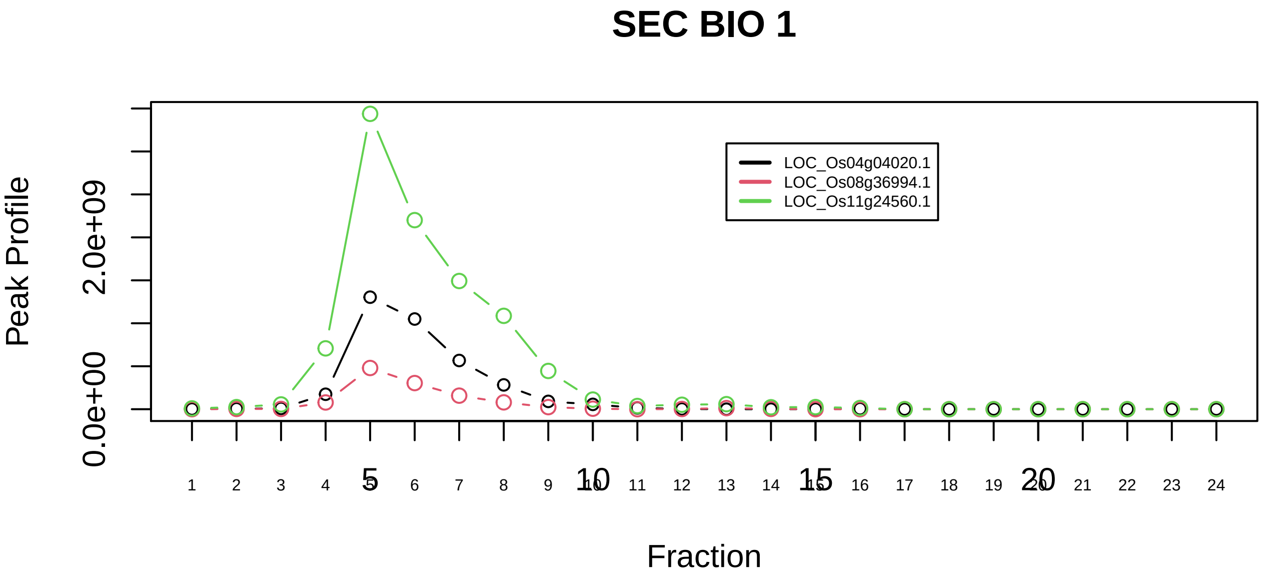

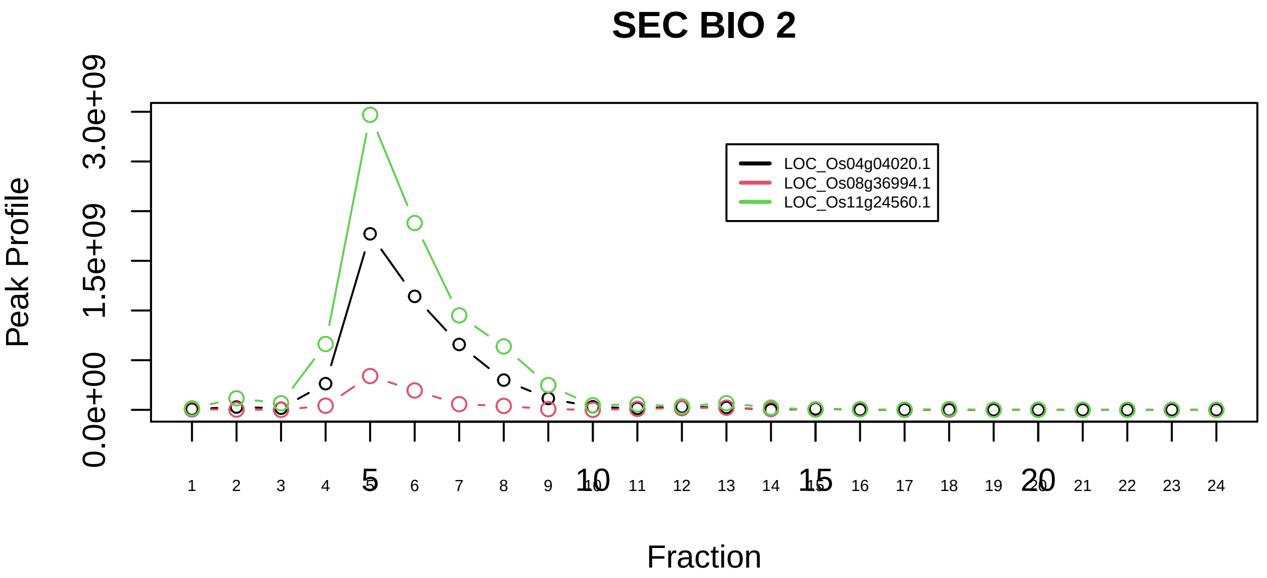

SEC23−SEC24 adaptor complex|7139−−all_prots_before_clustering_GPFdist

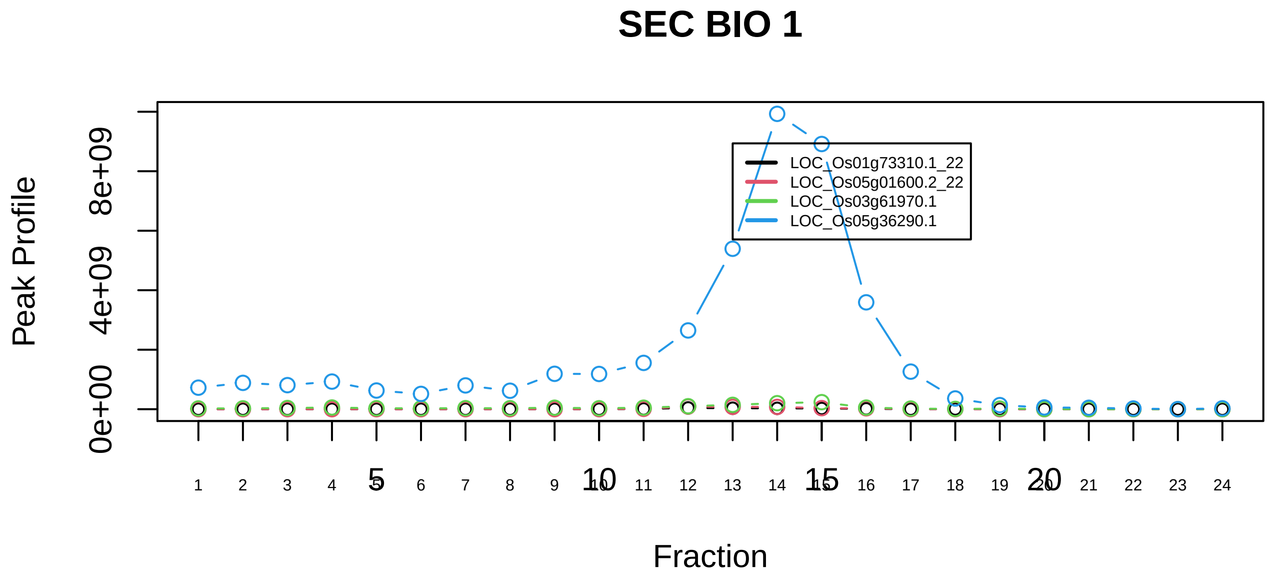

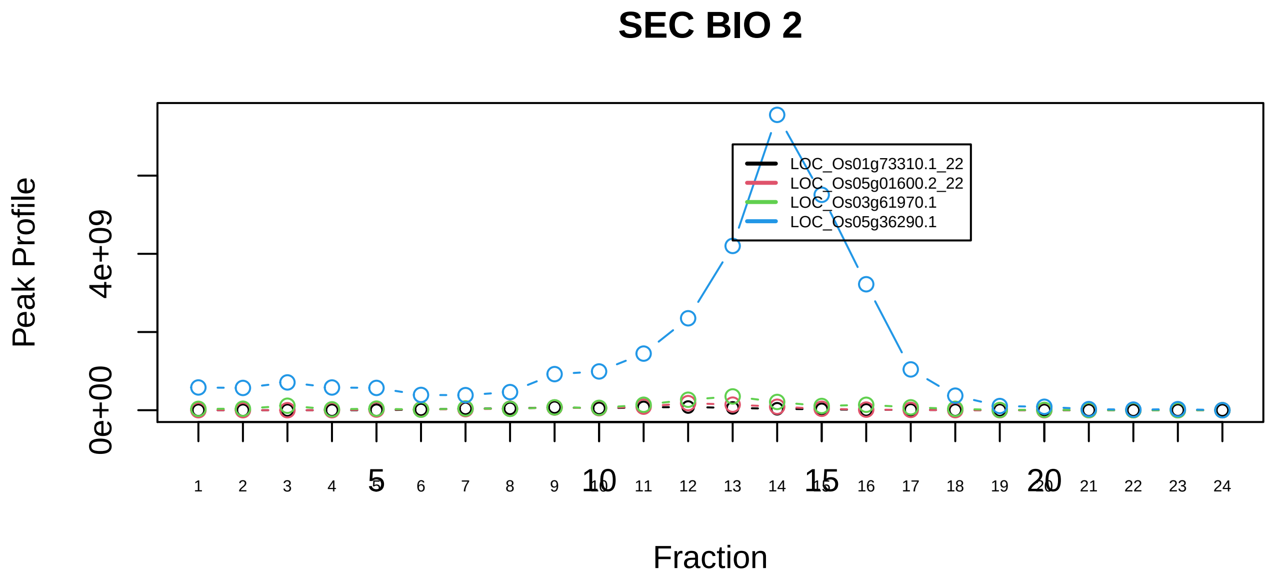

Cofilin−actin−CAP1 complex|2255−−sub_orthoparalog_1

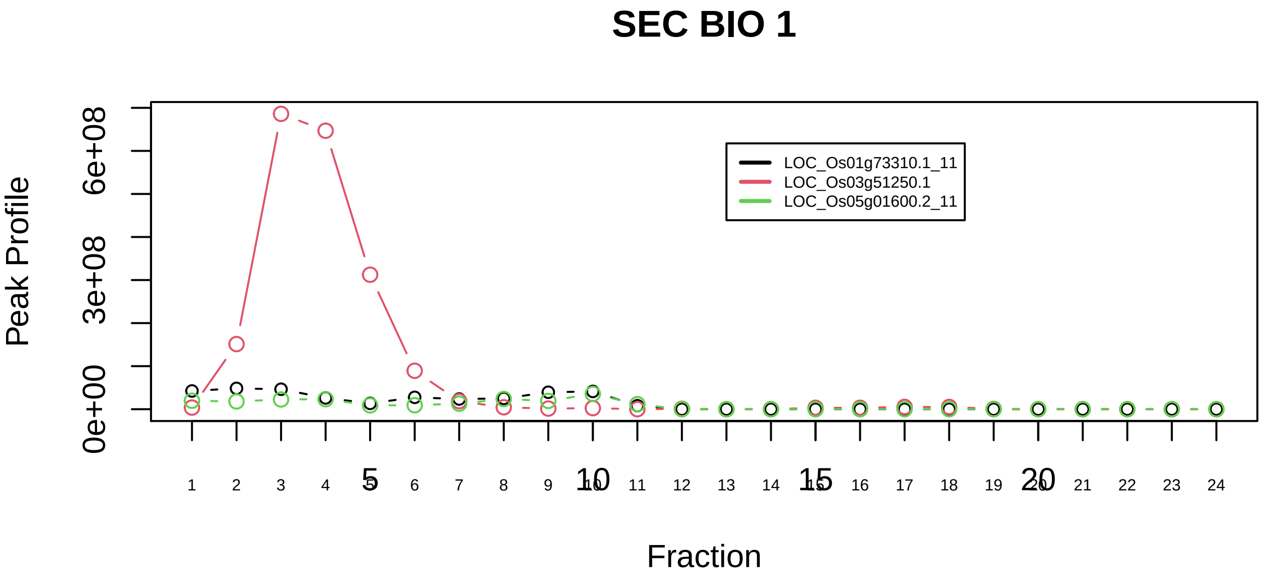

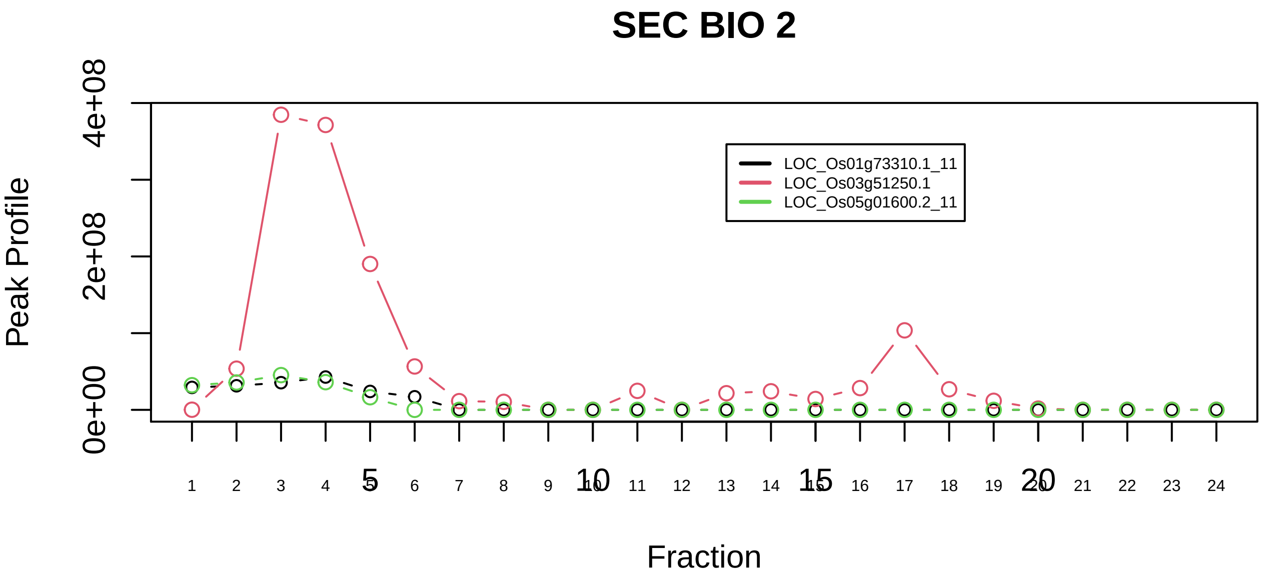

Cofilin−actin−CAP1 complex|2255−−sub_1

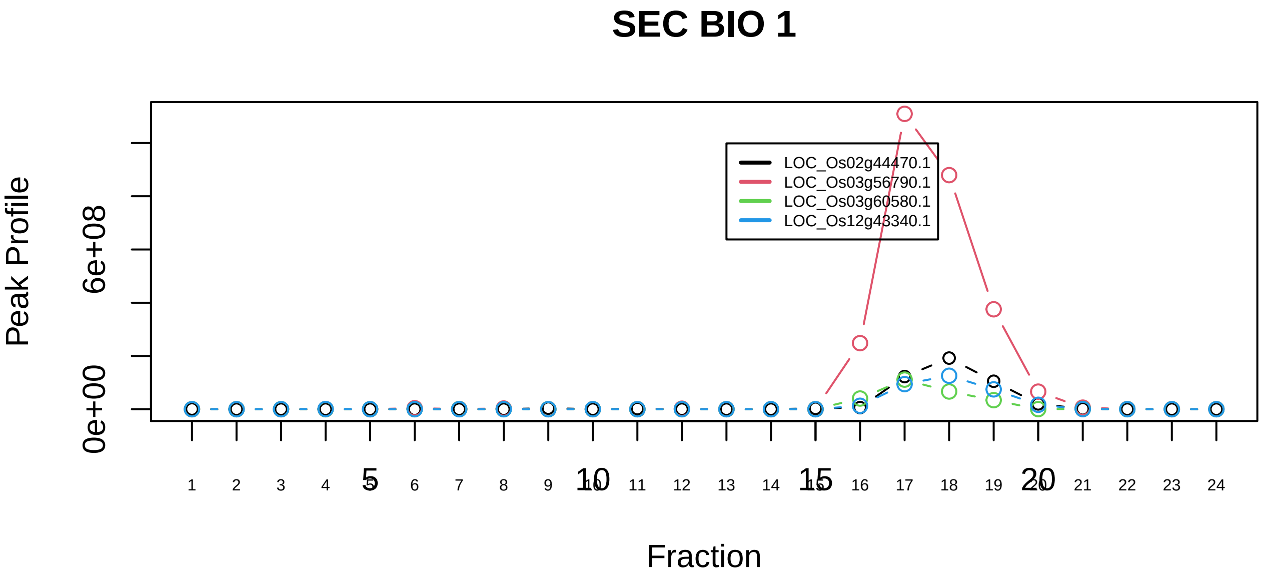

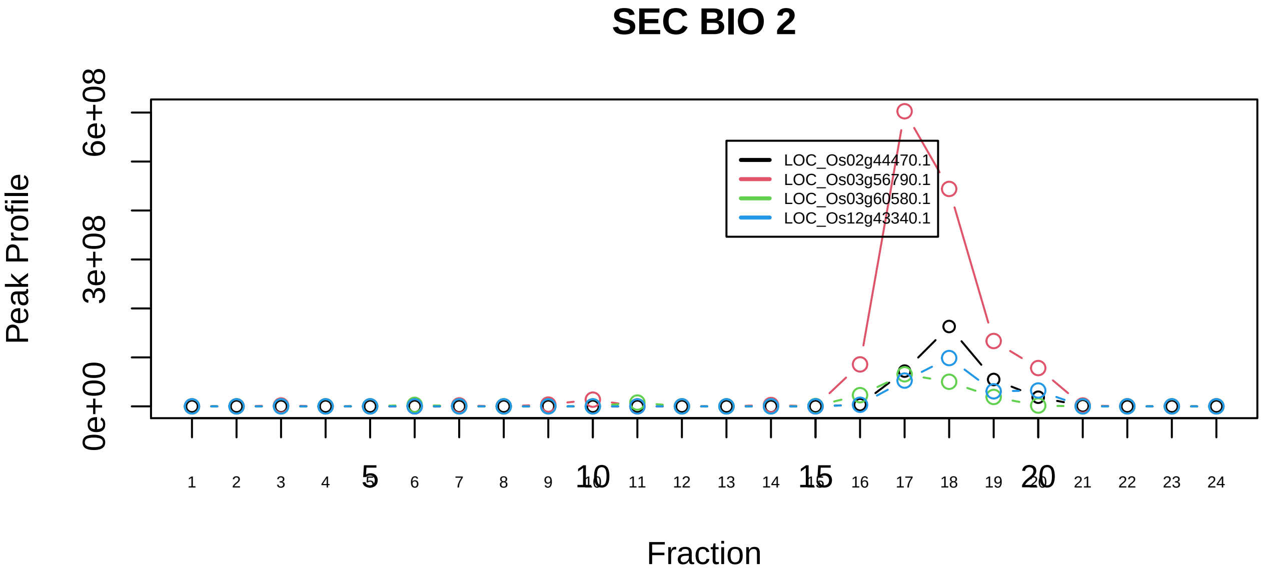

Cofilin−actin−CAP1 complex|2255−−sub_orthoparalog_2

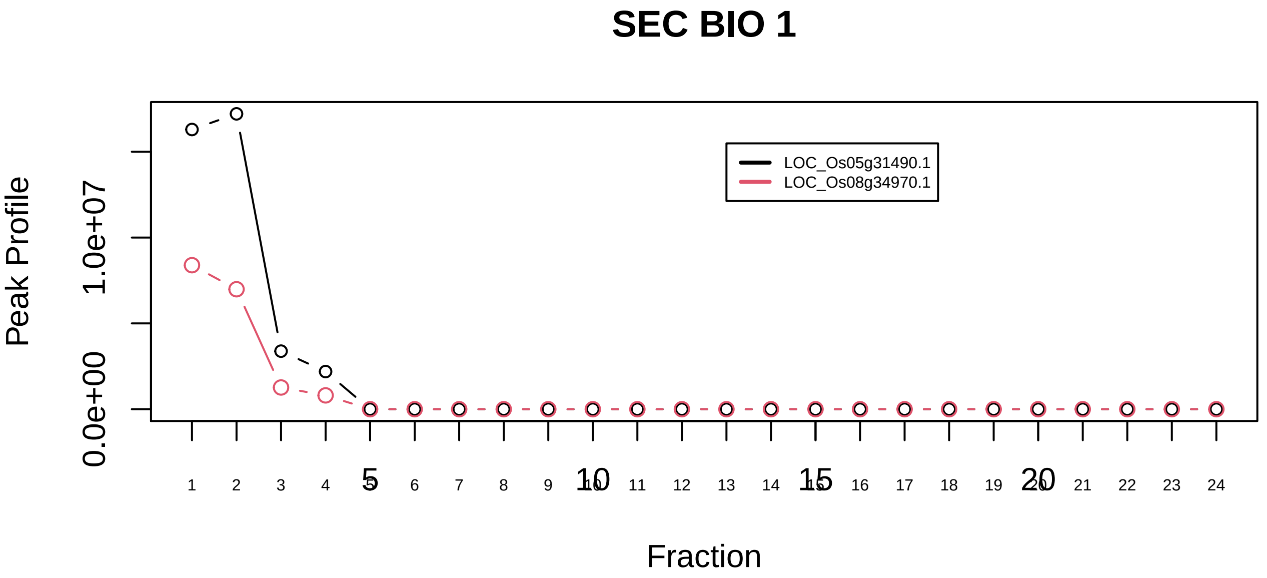

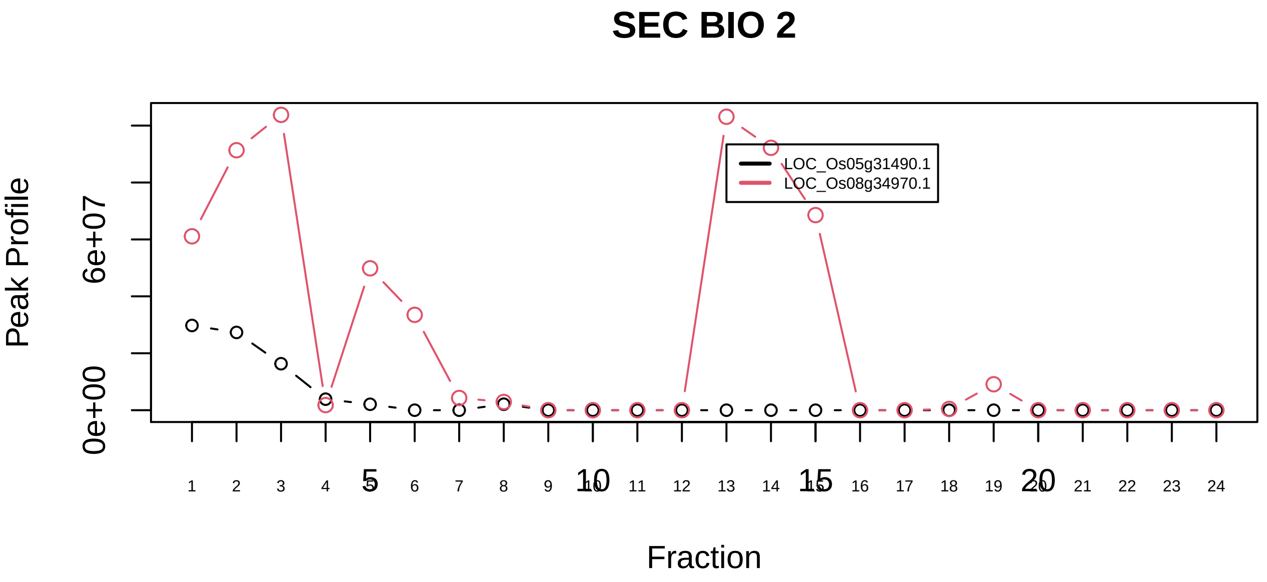

COG complex|3556−−only_2_prots_before_clustering

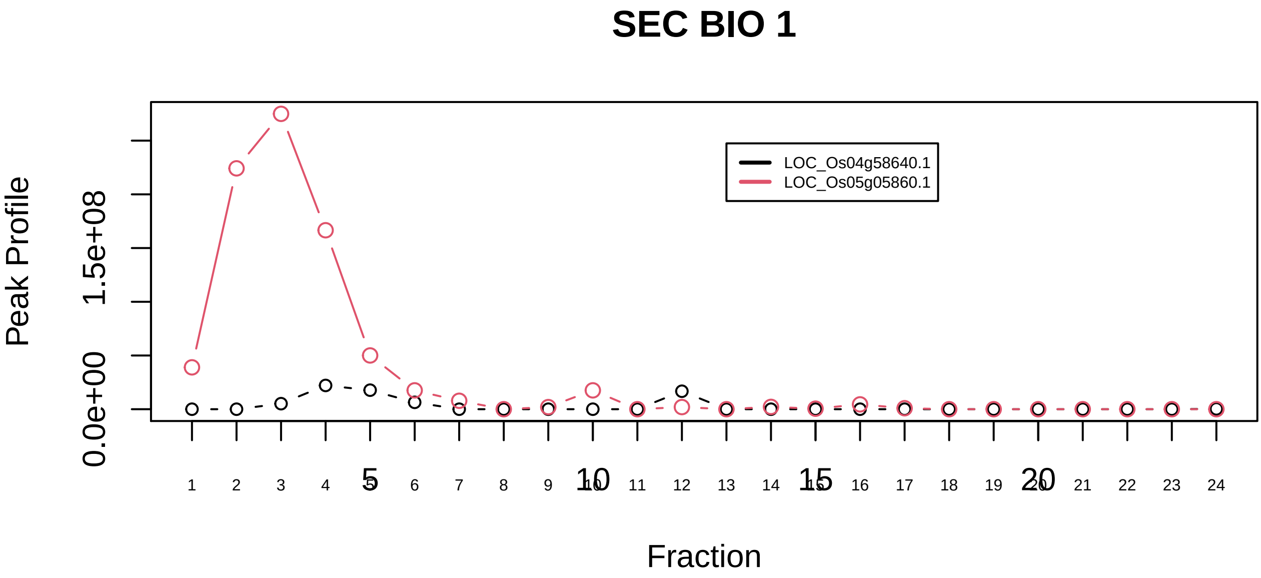

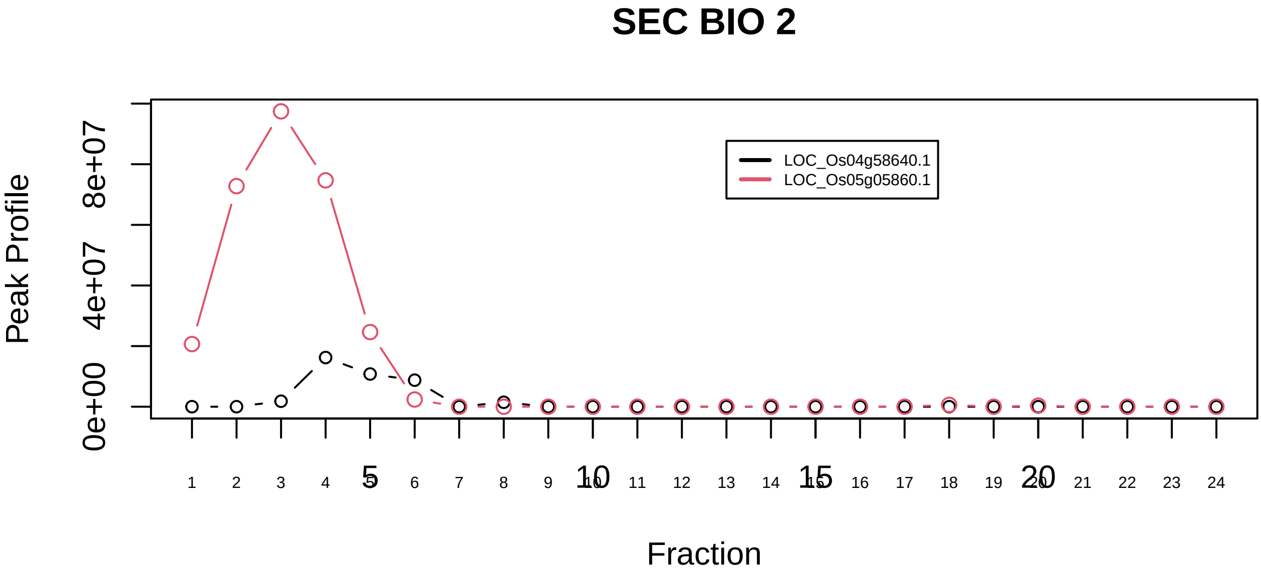

CPSF6−ITCH−NUDT21−POLR2A complex|1769−−only_2_prots_before_clustering

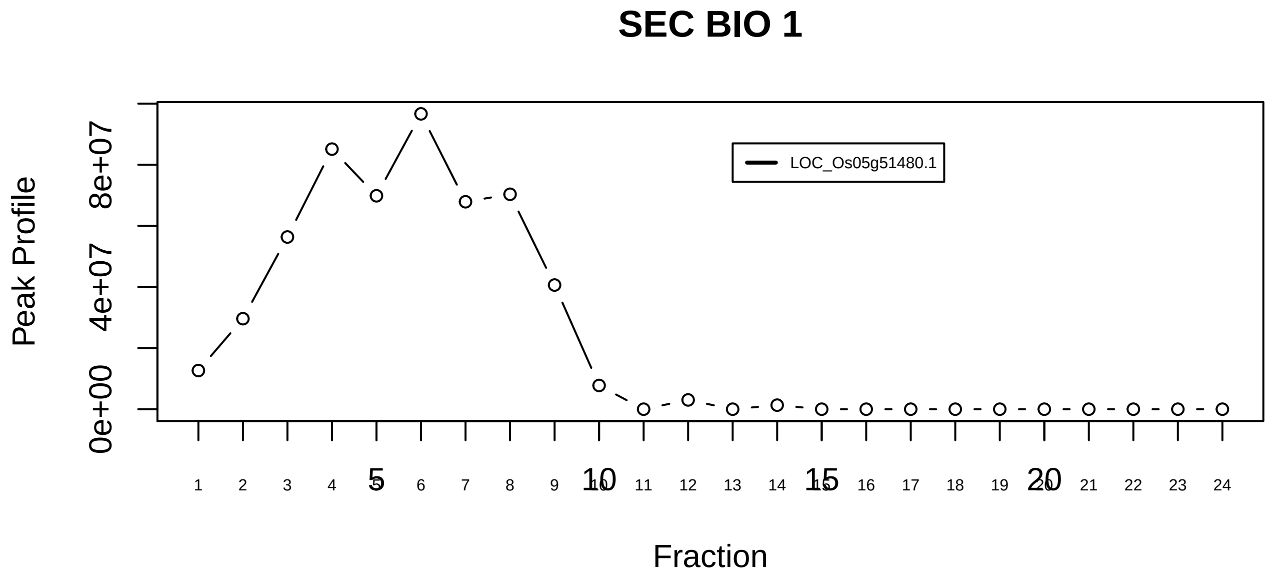

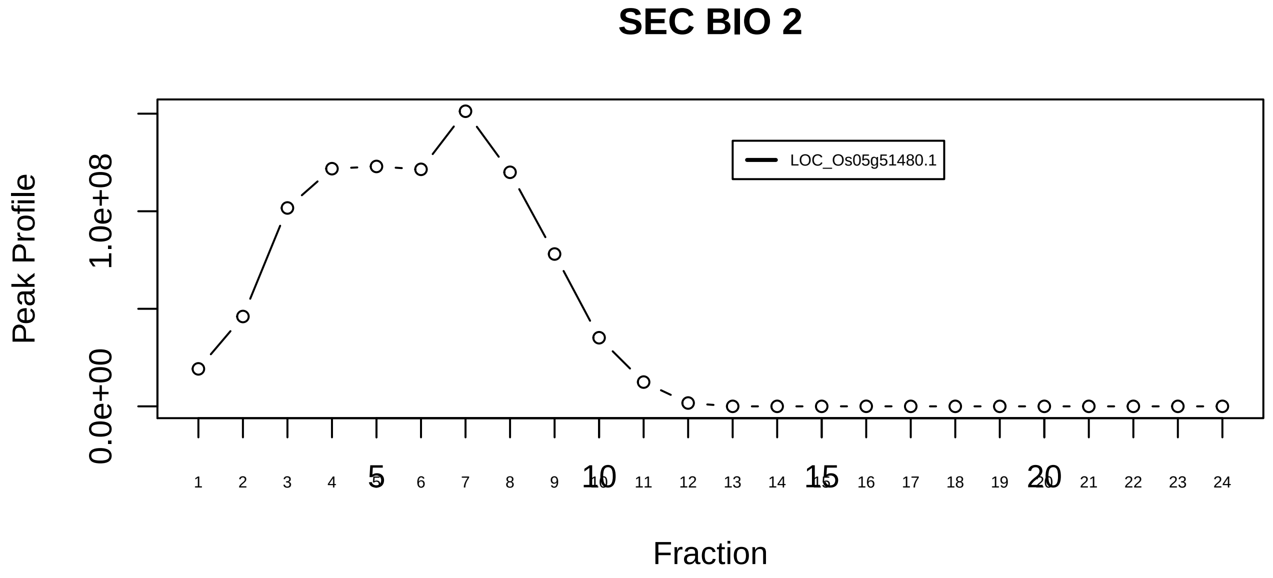

CSA−POLIIa complex|728−−singleton_1

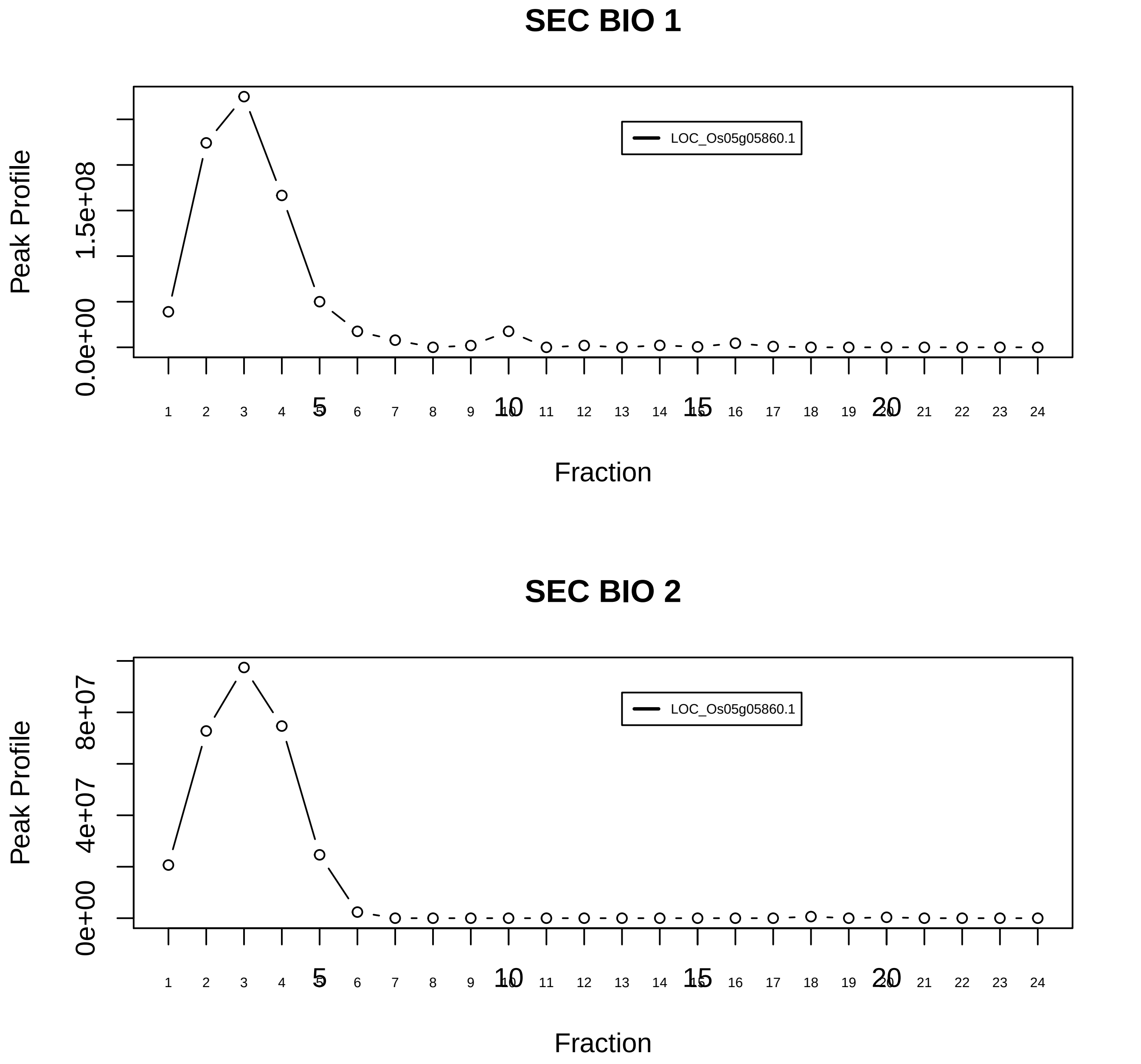
SEC BIO 1
LOC_Os05g05860.1
Peak Profile
0.0e+00
1.5e+08
Fraction
1 2 3 4 5 6 7 8 9 10 11 12 13 14 15 16 17 18 19 20 21 22 23 24
5
10
15
20
SEC BIO 2
LOC_Os05g05860.1
Peak Profile
0e+00
4e+07
8e+07
Fraction

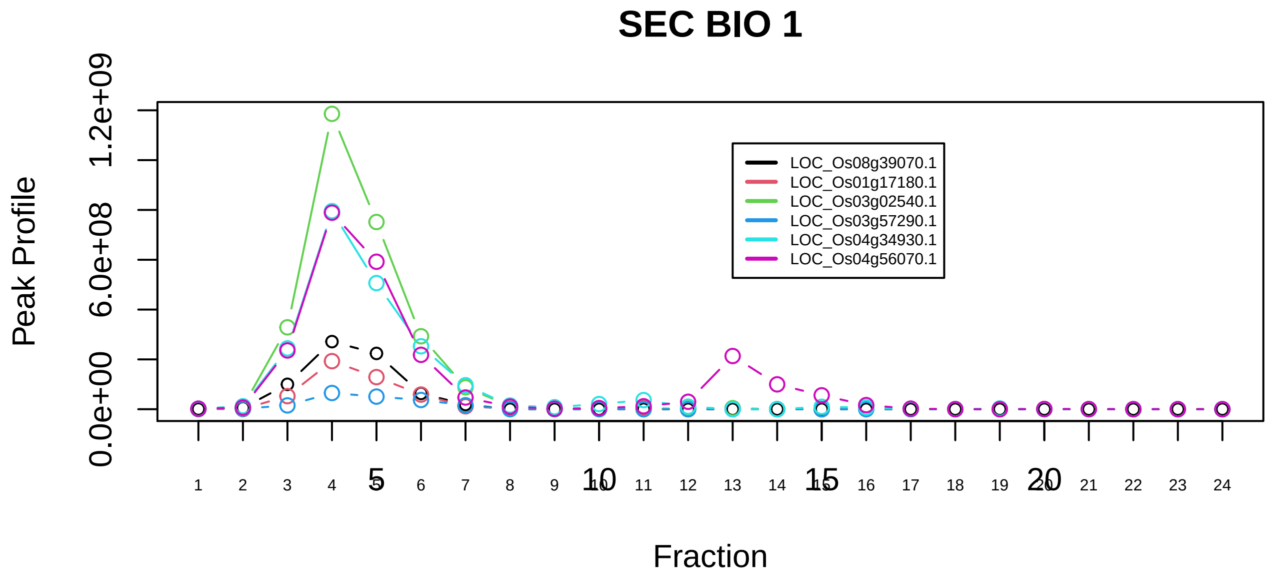

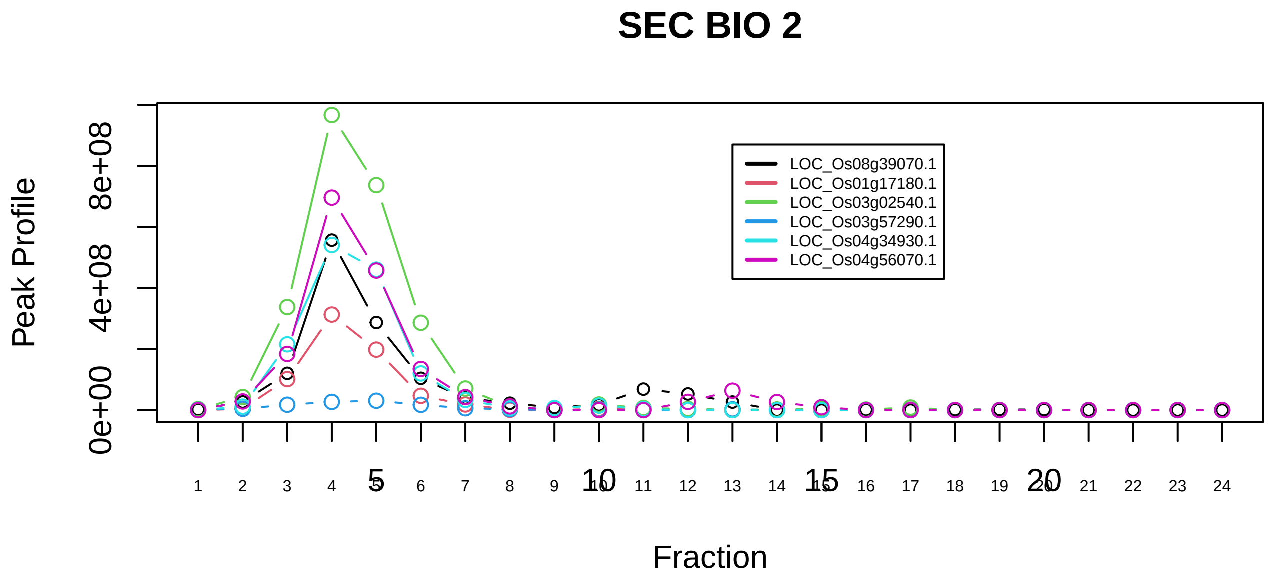

CSA−POLIIa complex|728−−sub_1

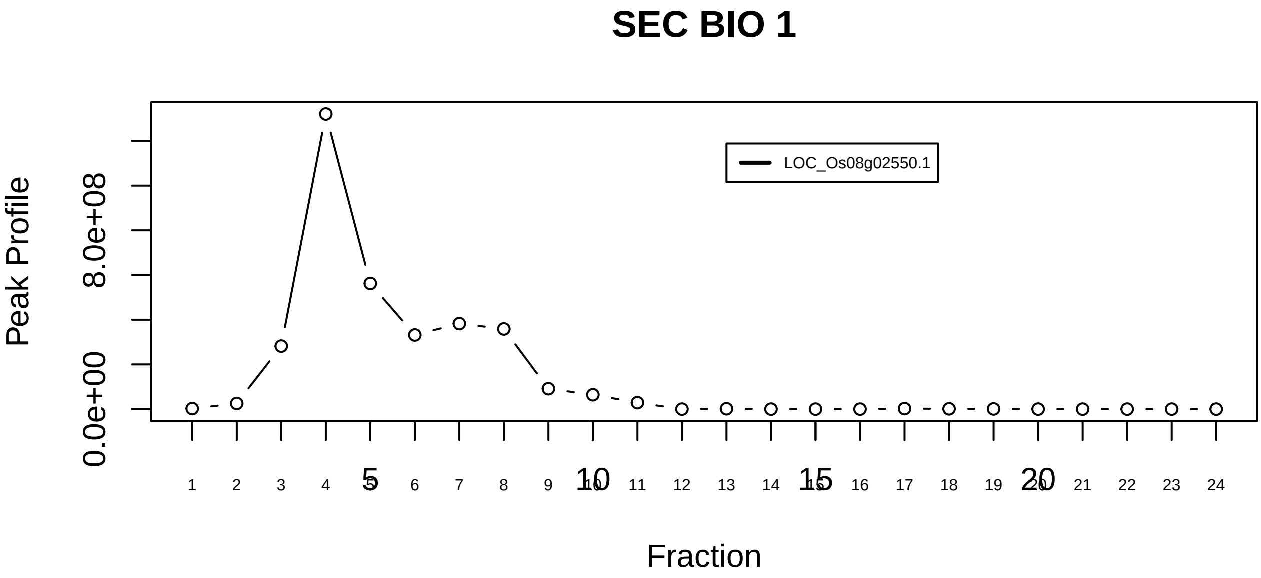

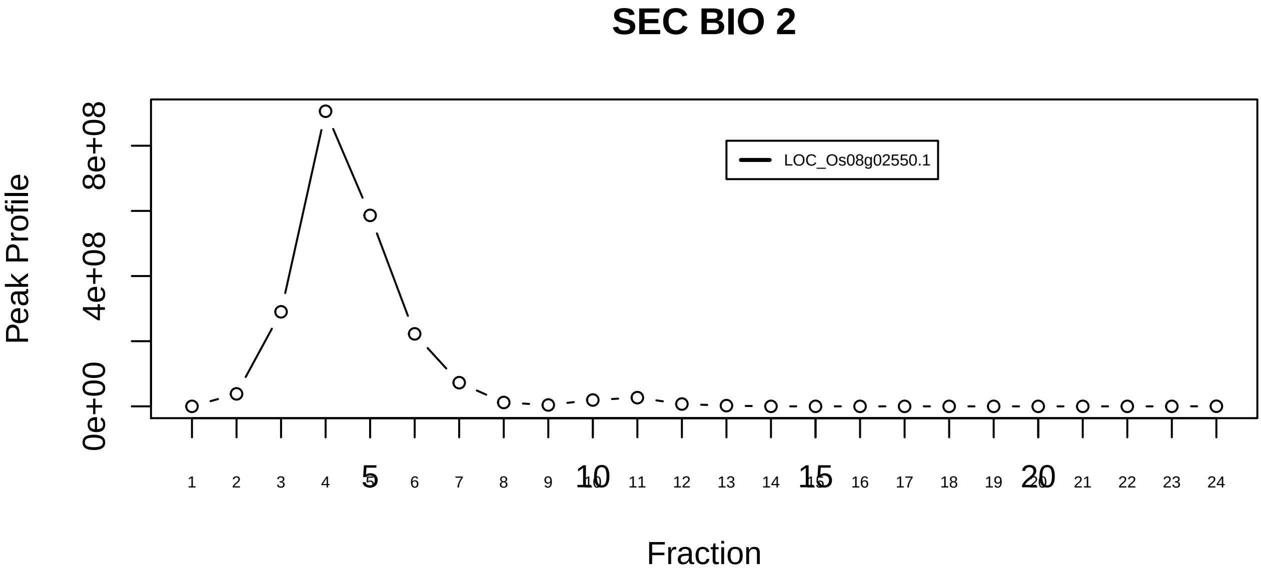

CSA−POLIIa complex|728−−singleton_3

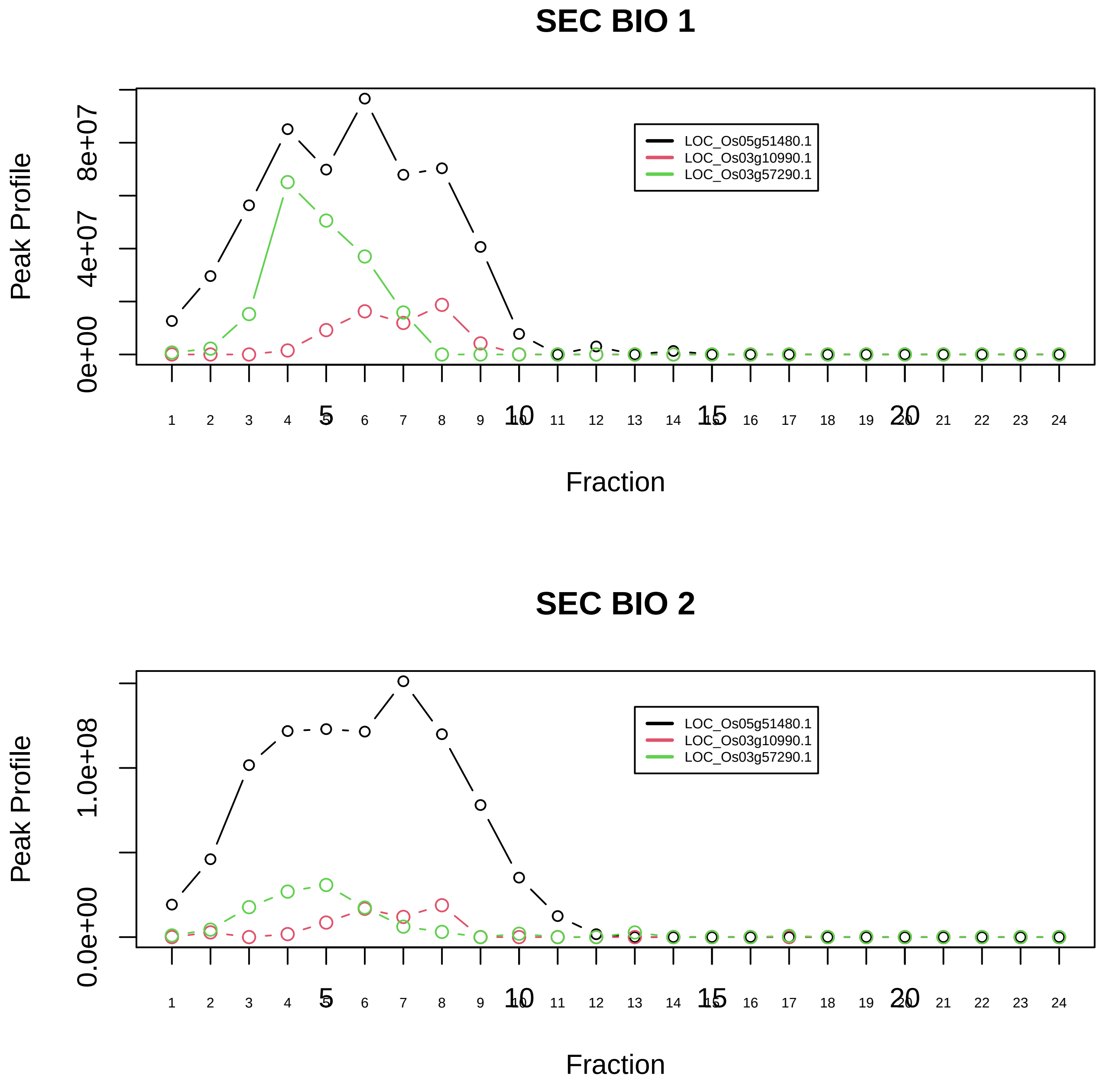

CUL4A−DDB1−RBBP5 complex|6491−−sub_1

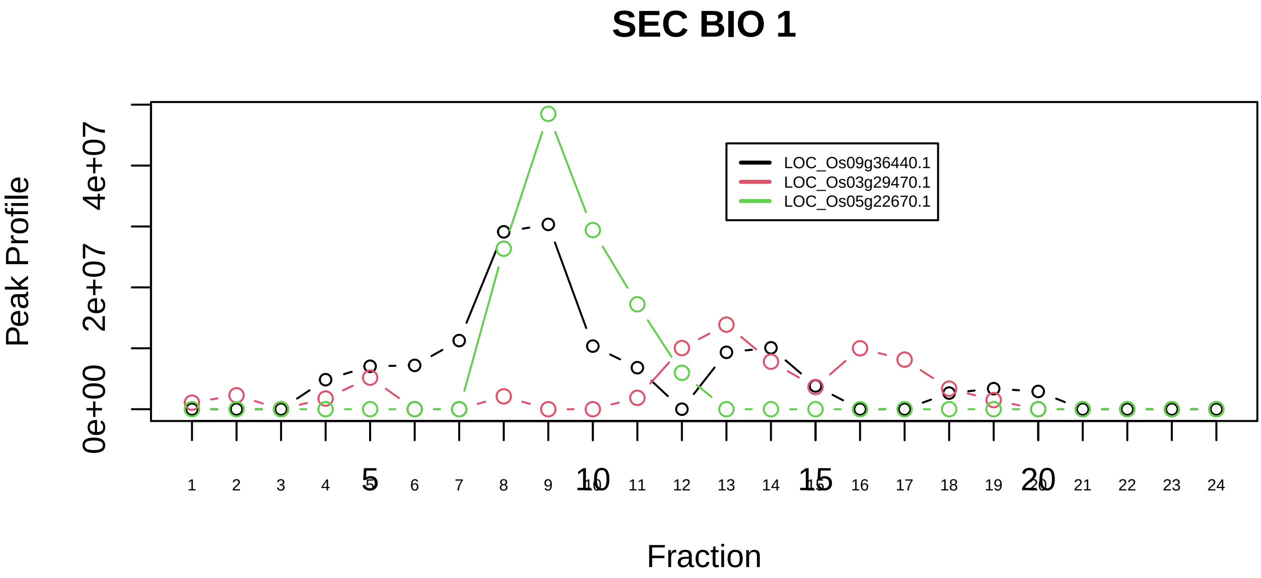

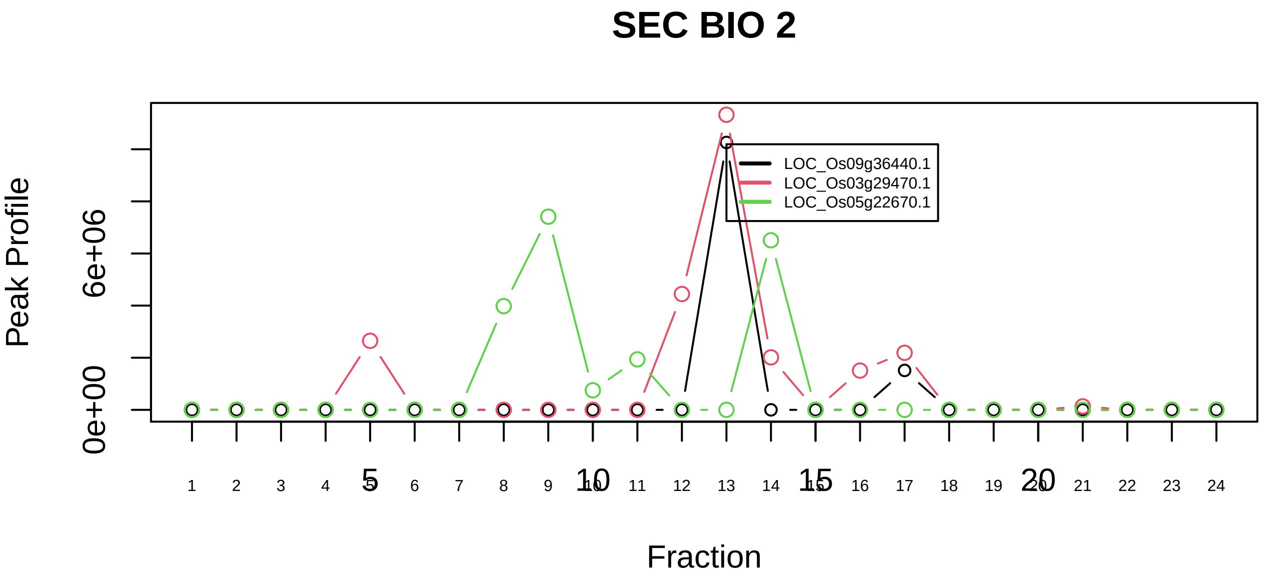

DAB complex|493−−sub_1

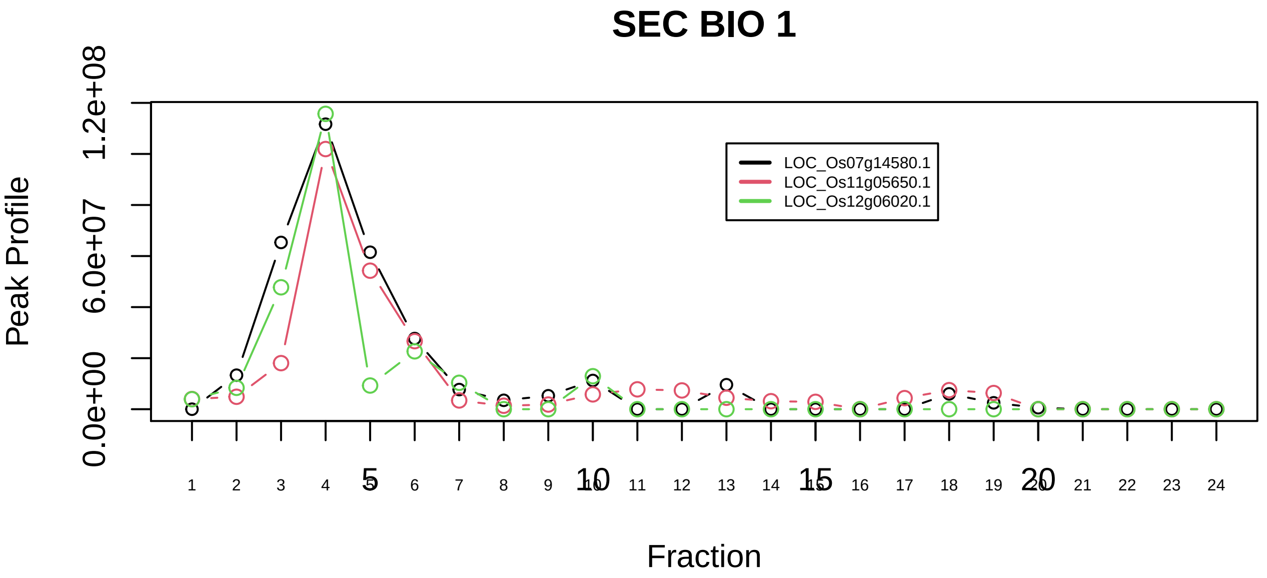

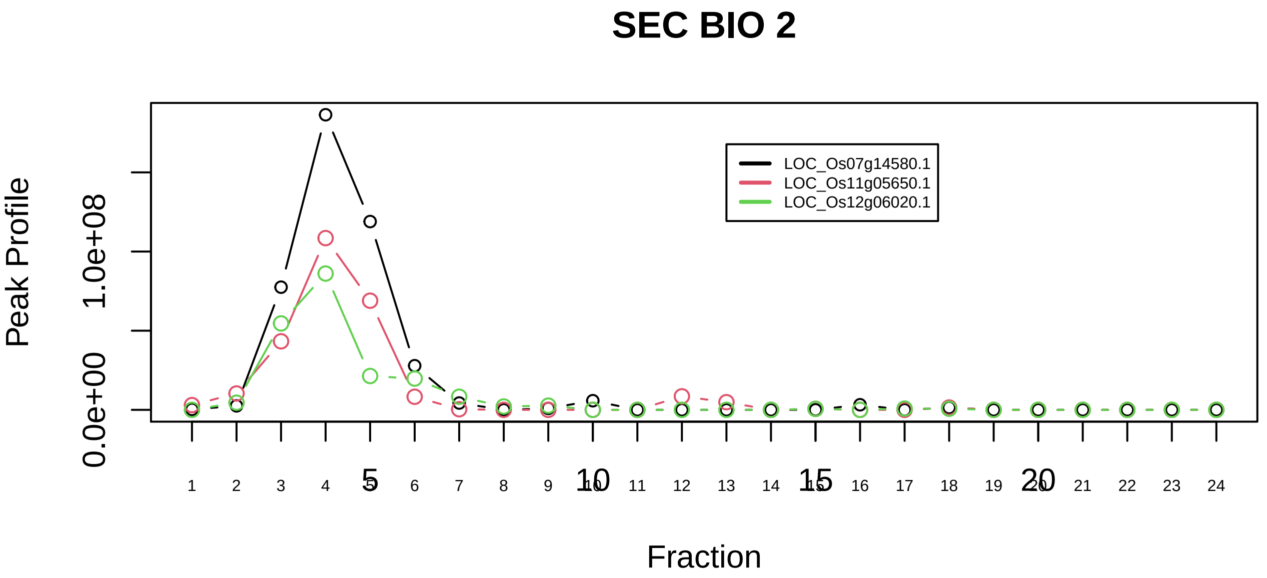

Decapping complex|559−−sub_orthoparalog_1

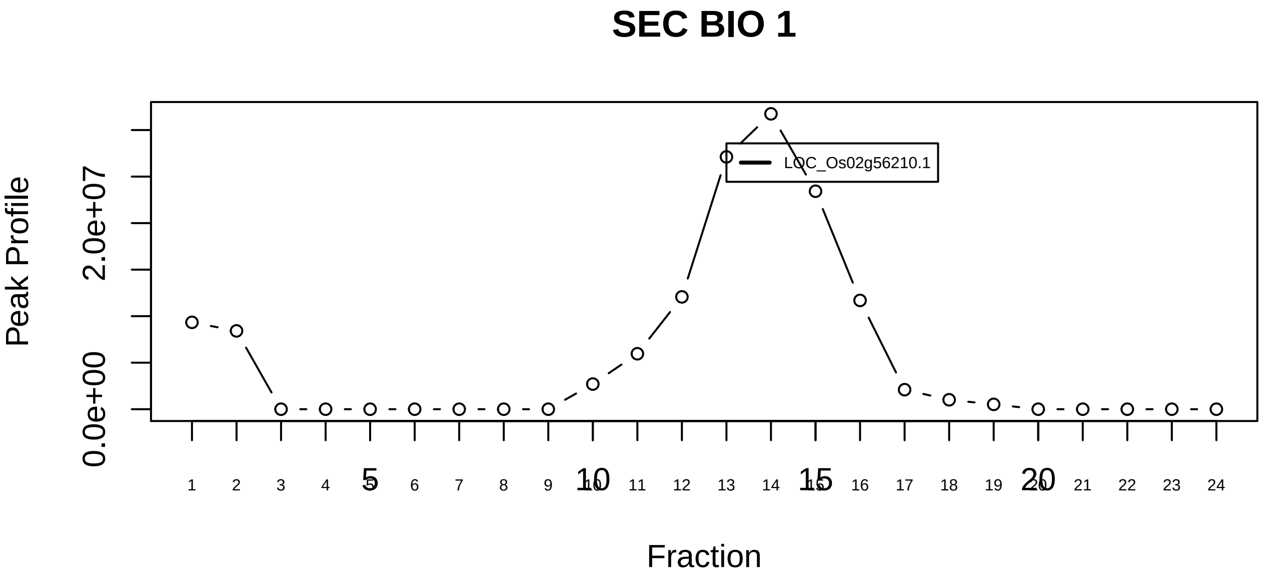

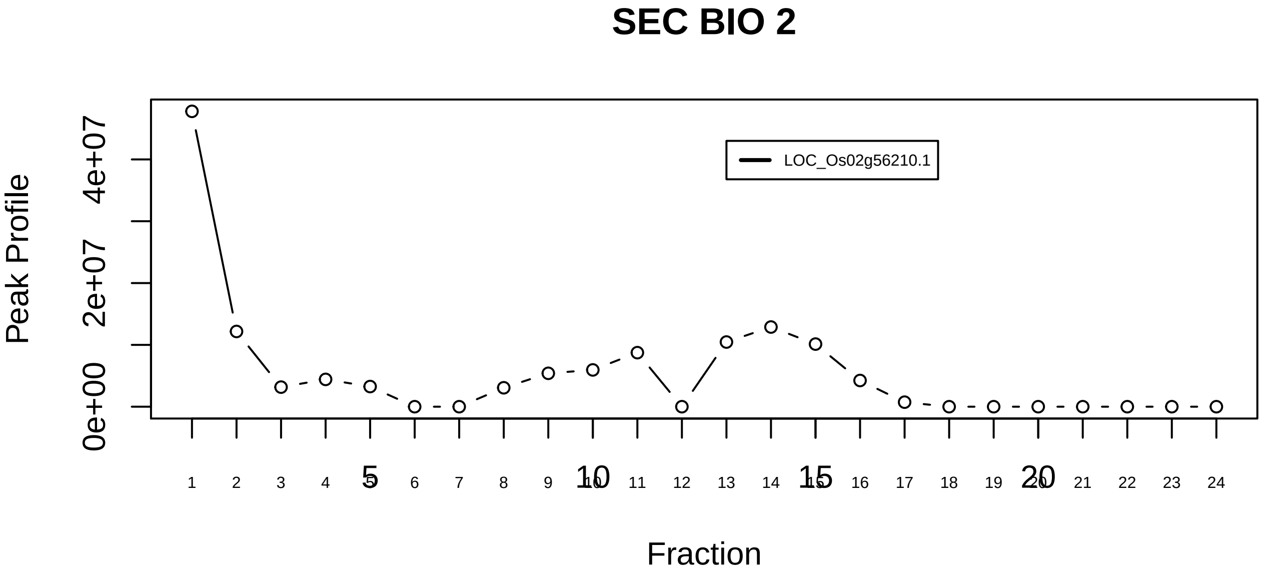

Decapping complex|559−−singleton_1

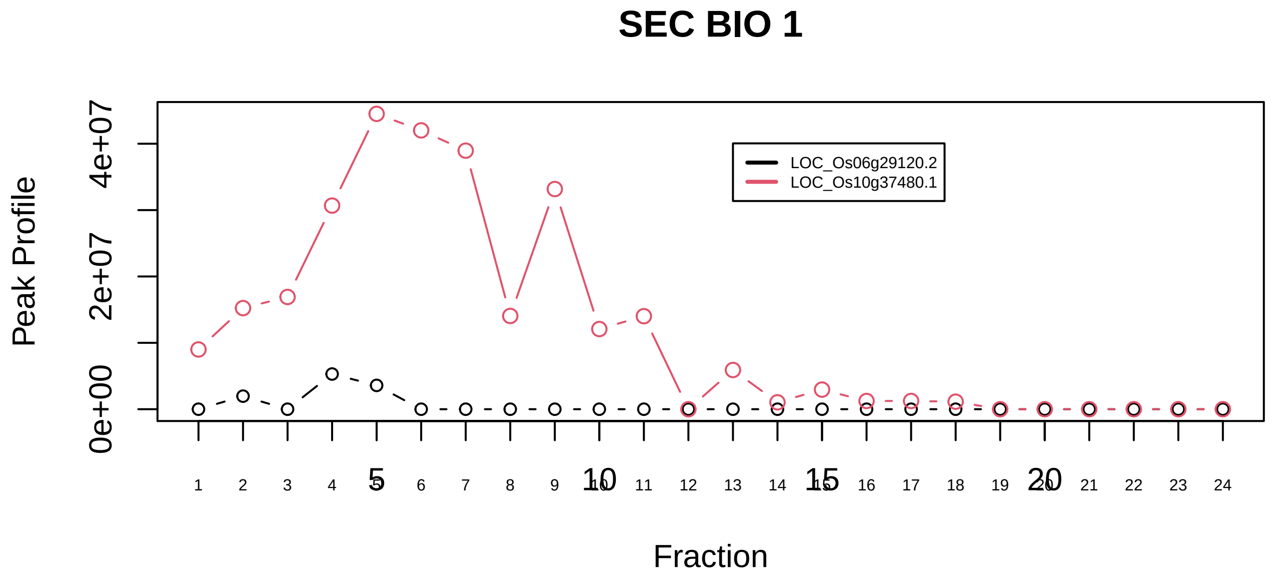

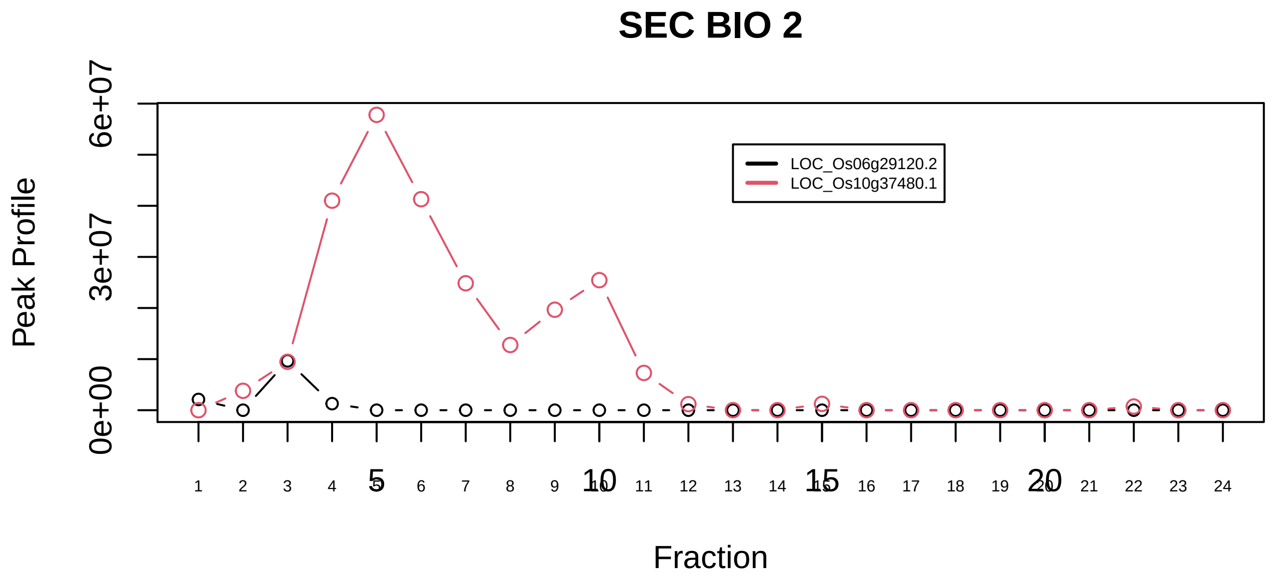

DLG5−MST1−MST2 complex|6716−−only_2_prots_before_clustering

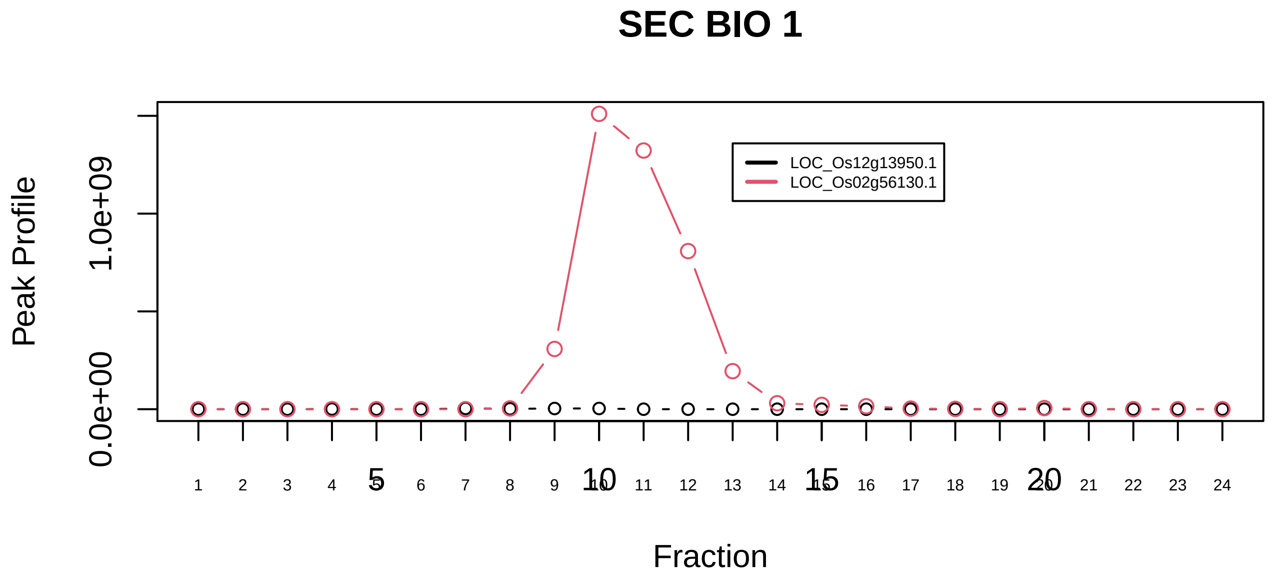

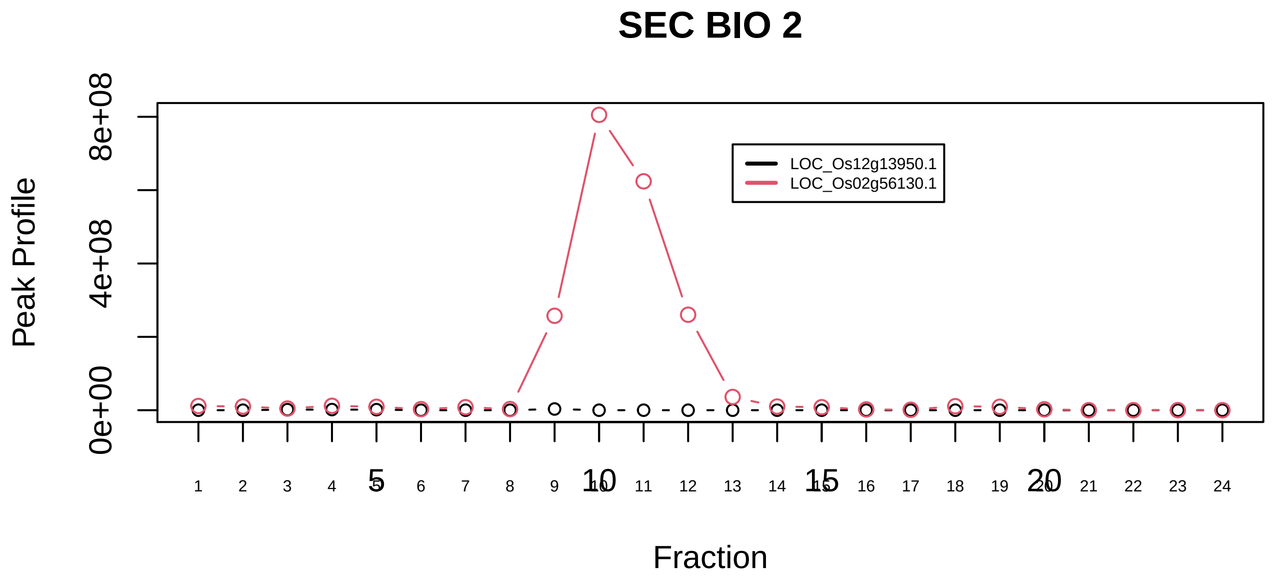

DNA synthesome complex (17 subunits)|1111−−sub_1

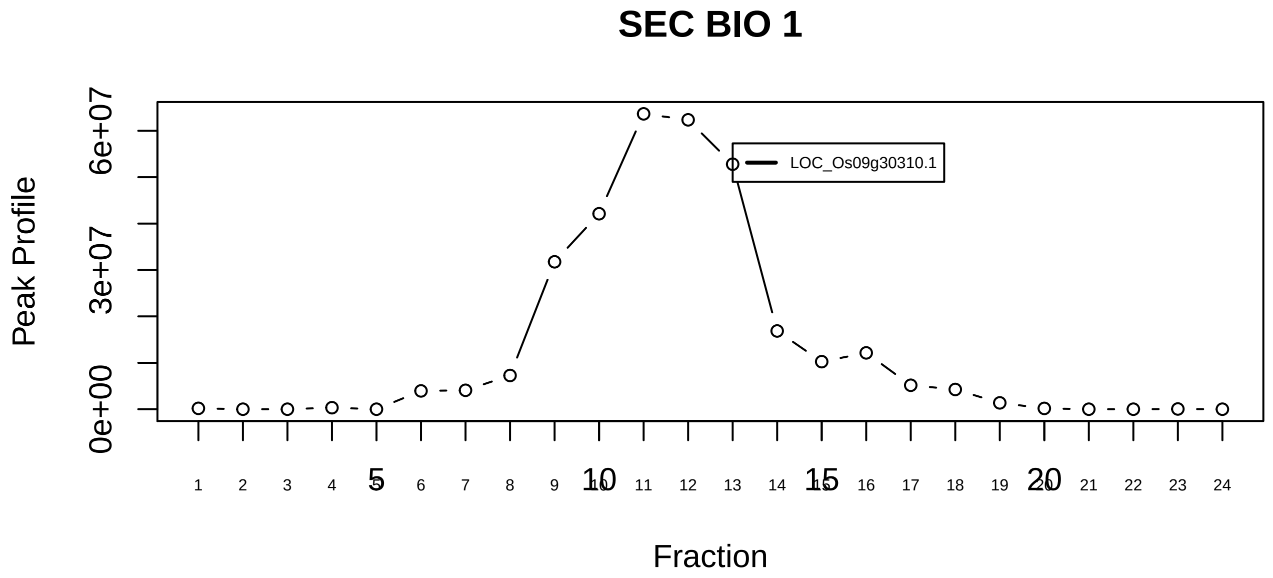

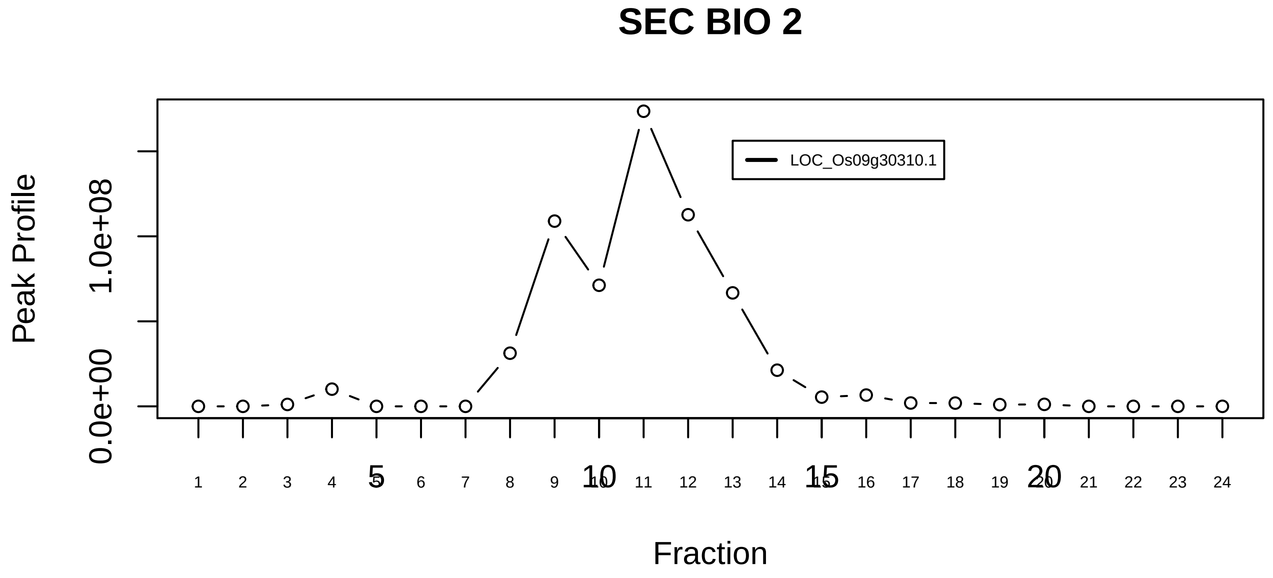

DNA synthesome complex (17 subunits)|1111−−singleton_1

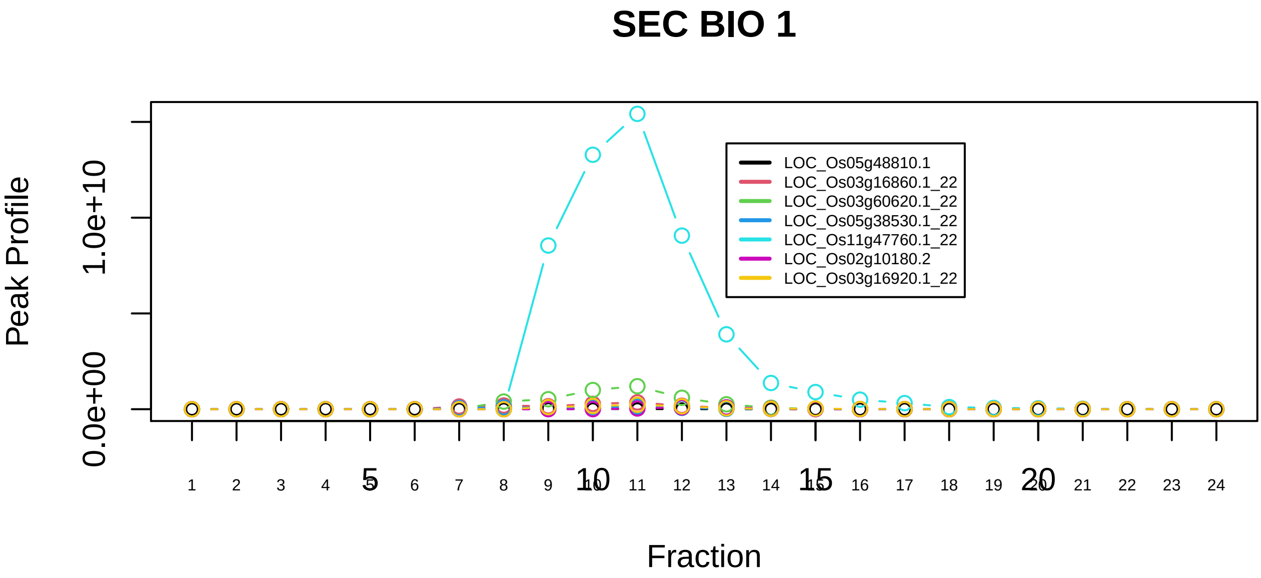

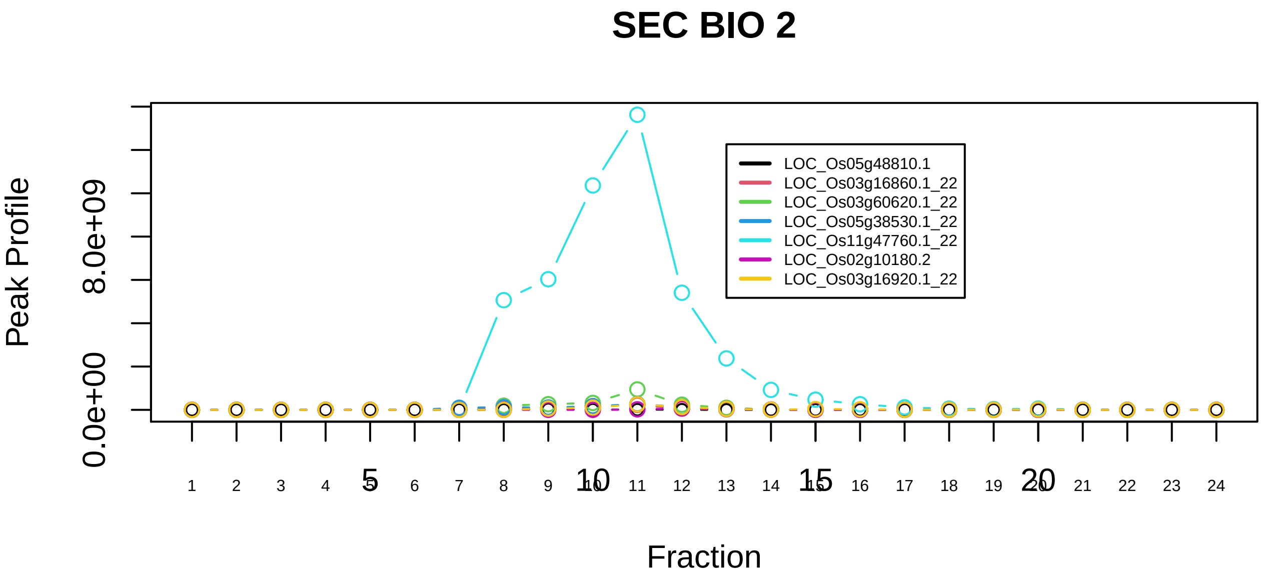

DNJC3−DNAJB1−HSPA8 complex|6396−−sub_1

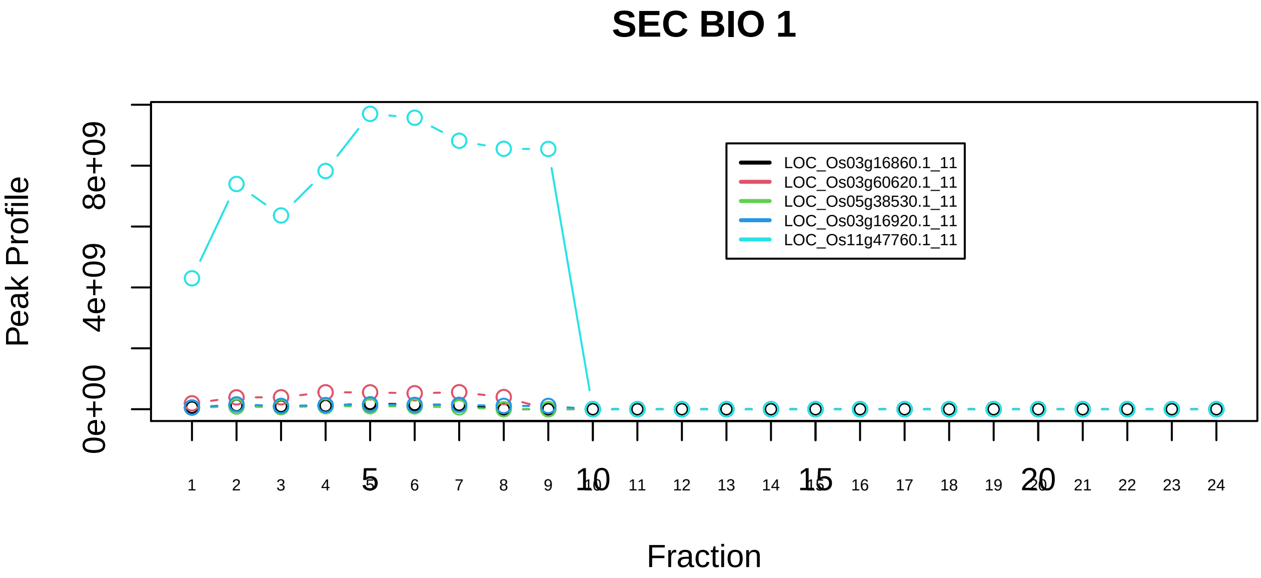

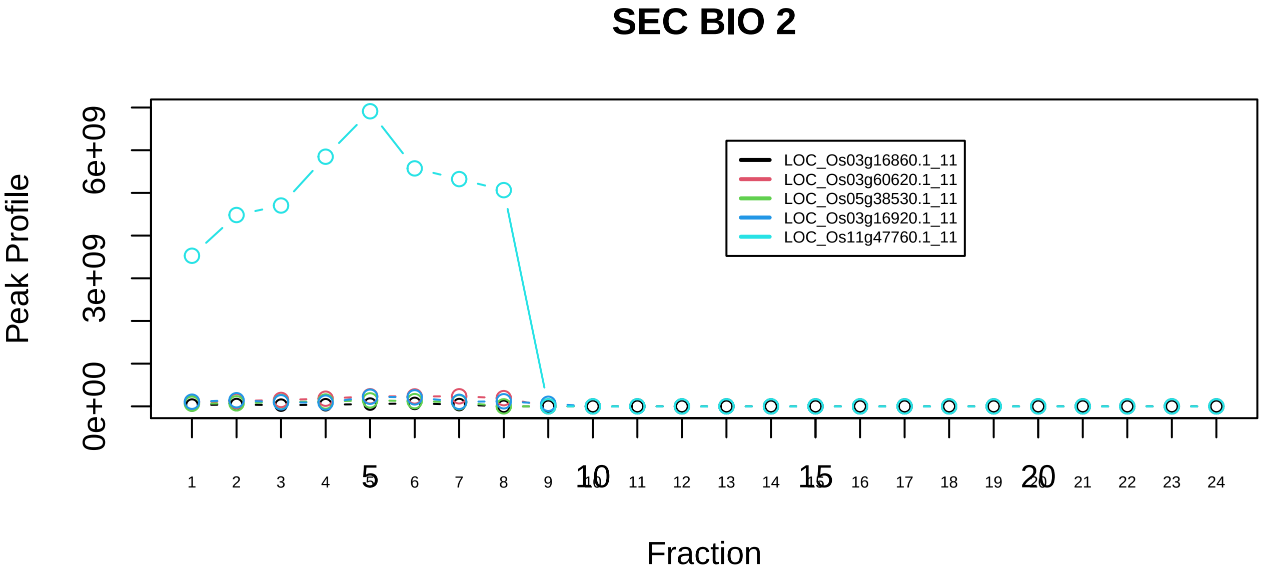

DNJC3−DNAJB1−HSPA8 complex|6396−−sub_orthoparalog_1

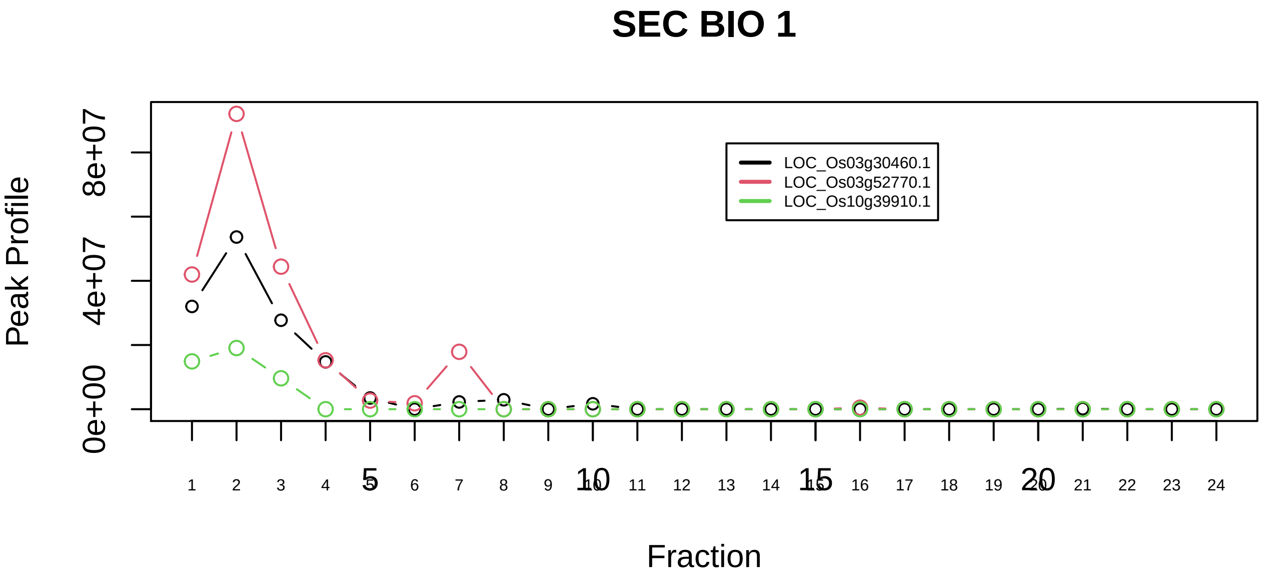

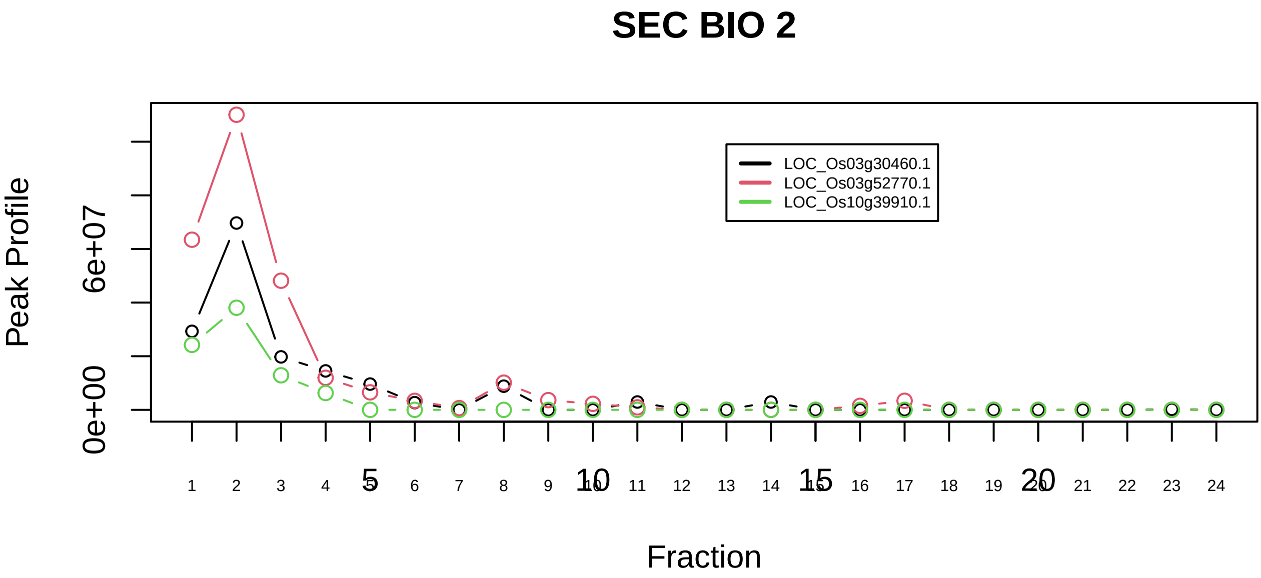

EARP complex|6459−−all_prots_before_clustering_GPFdist

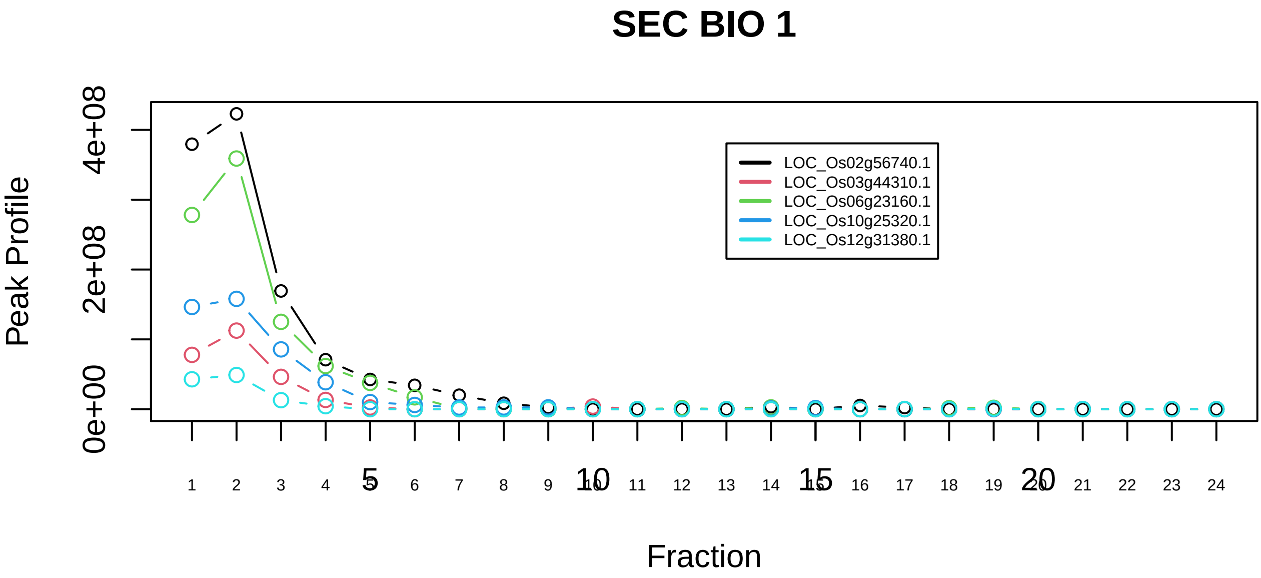

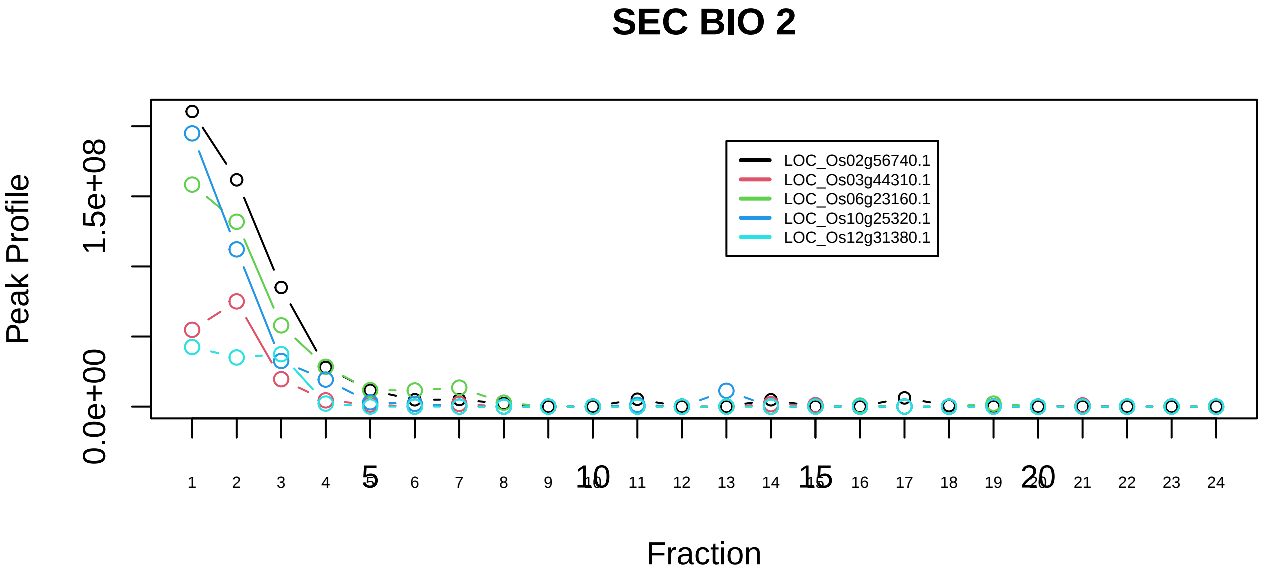

EIF2B1−EIF2B2−EIF2B3−EIF2B4−EIF2B5 complex|7292−−all_prots_before_clustering_sim

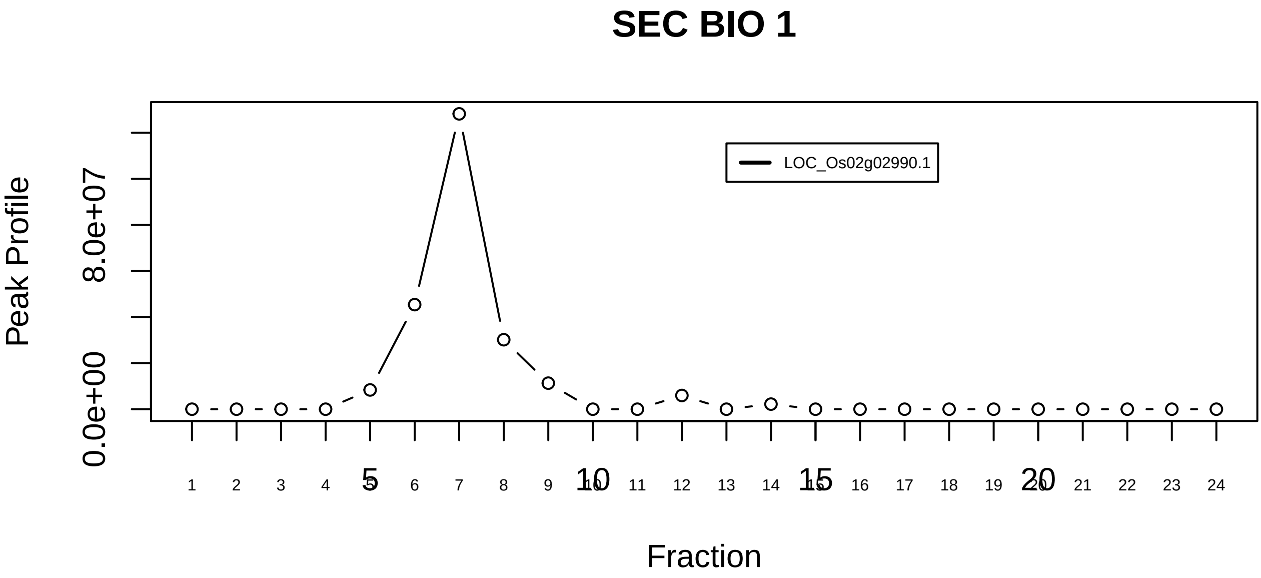

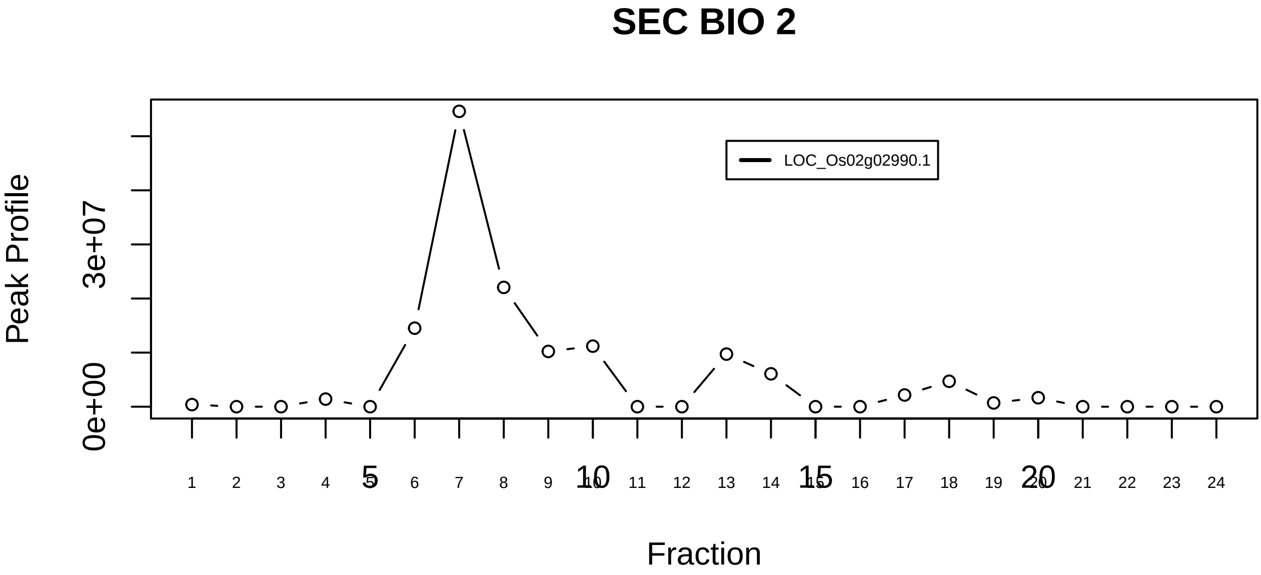

eIF3 complex (EIF3S6, EIF3S5, EIF3S4, EIF3S3, EIF3S6IP, EIF3S2, EIF3S9, EIF3S12, EIF3S10, EIF3S8, EIF3S1, EIF3S7, PCID1)|1097−−singleton_1

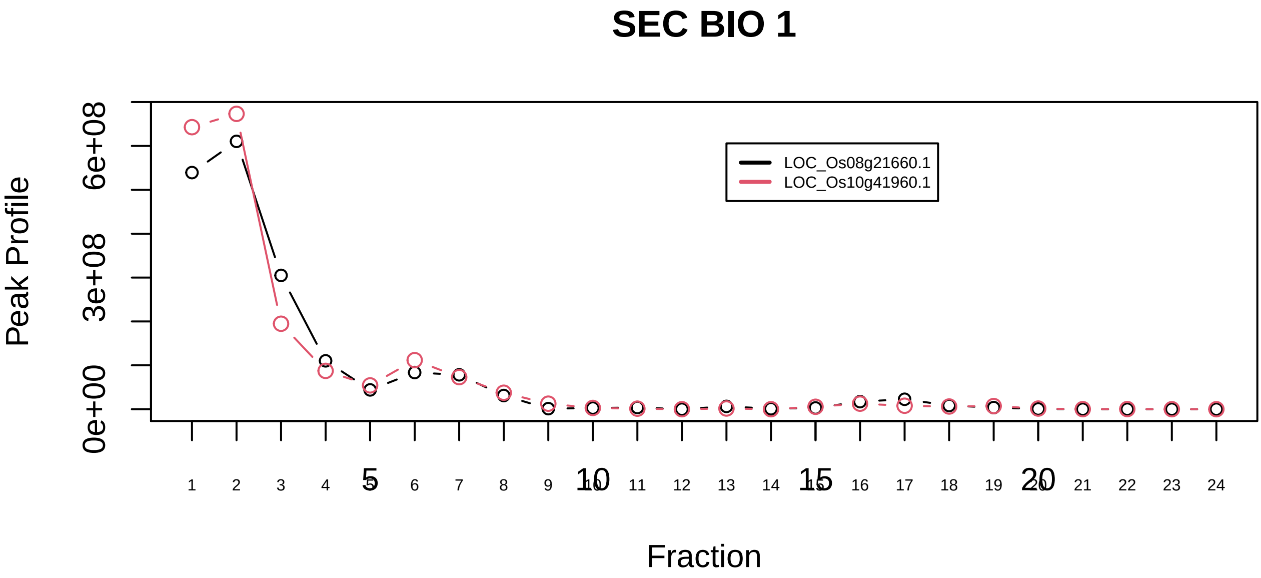

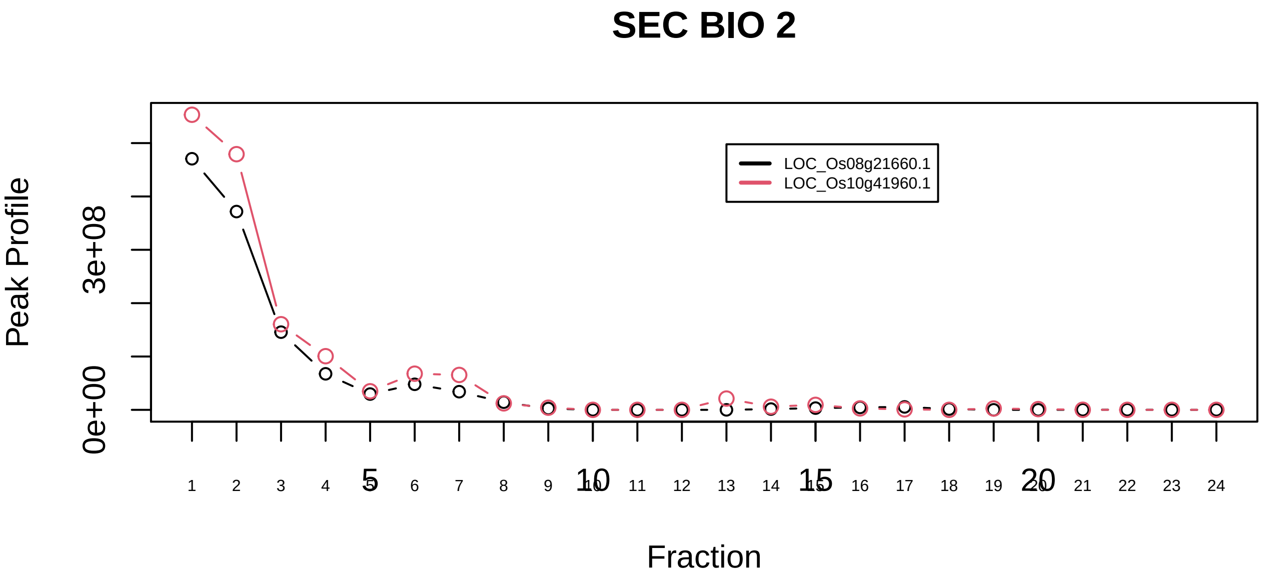

eIF3 complex (EIF3S6, EIF3S5, EIF3S4, EIF3S3, EIF3S6IP, EIF3S2, EIF3S9, EIF3S12, EIF3S10, EIF3S8, EIF3S1, EIF3S7, PCID1)|1097−−sub_1

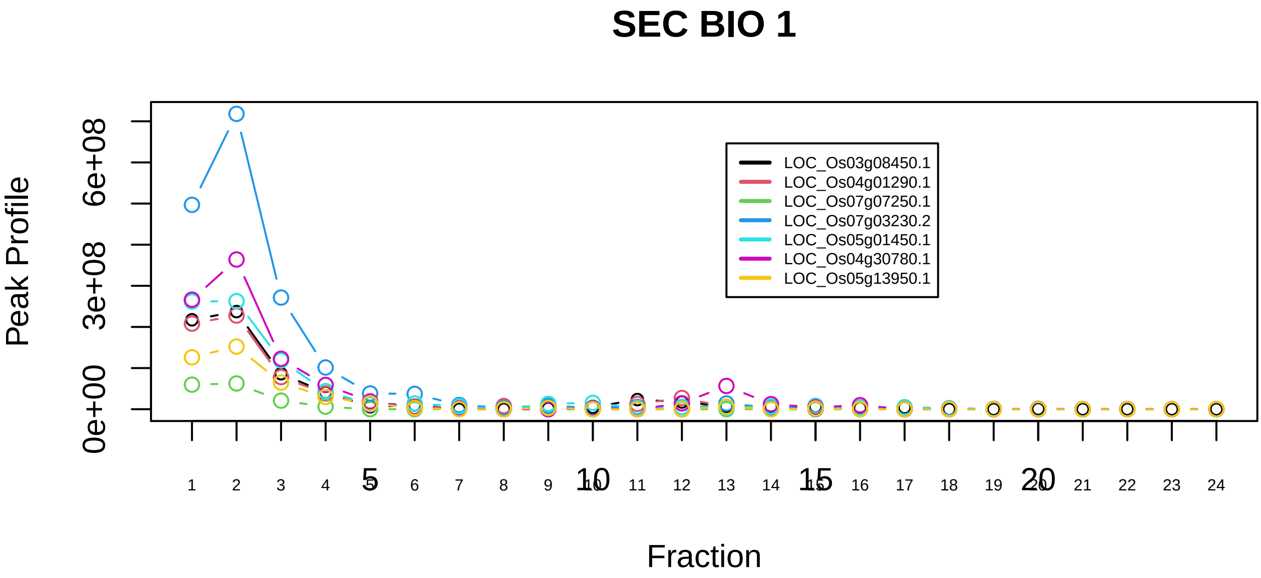

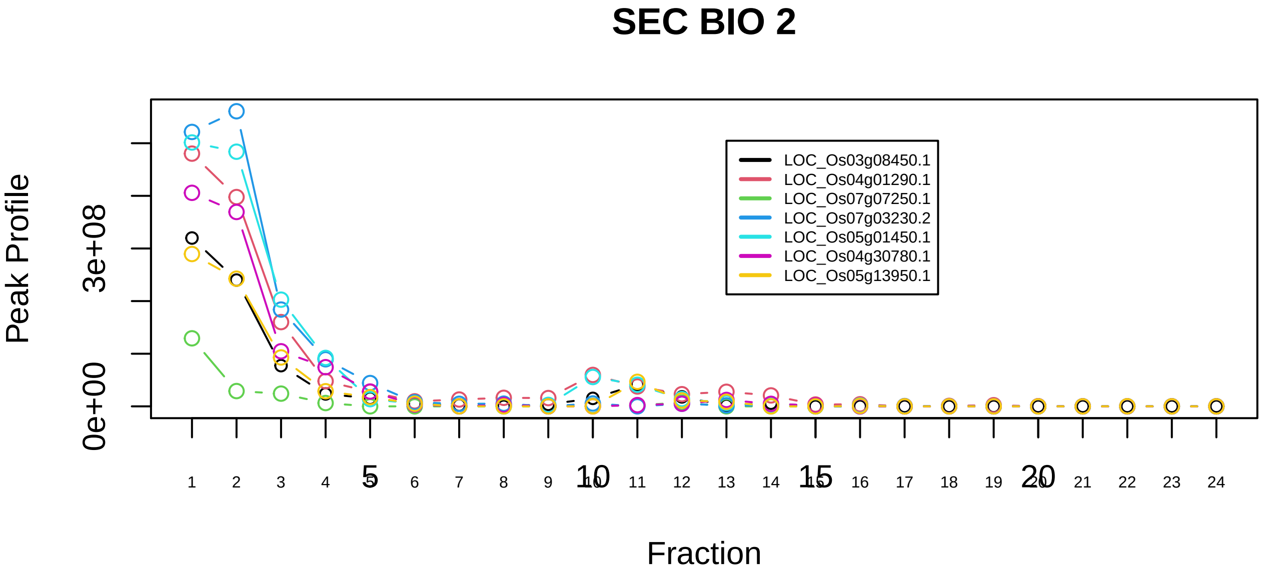

eIF3 complex (EIF3S6, EIF3S5, EIF3S4, EIF3S3, EIF3S6IP, EIF3S2, EIF3S9, EIF3S12, EIF3S10, EIF3S8, EIF3S1, EIF3S7, PCID1)|1097−−sub_2

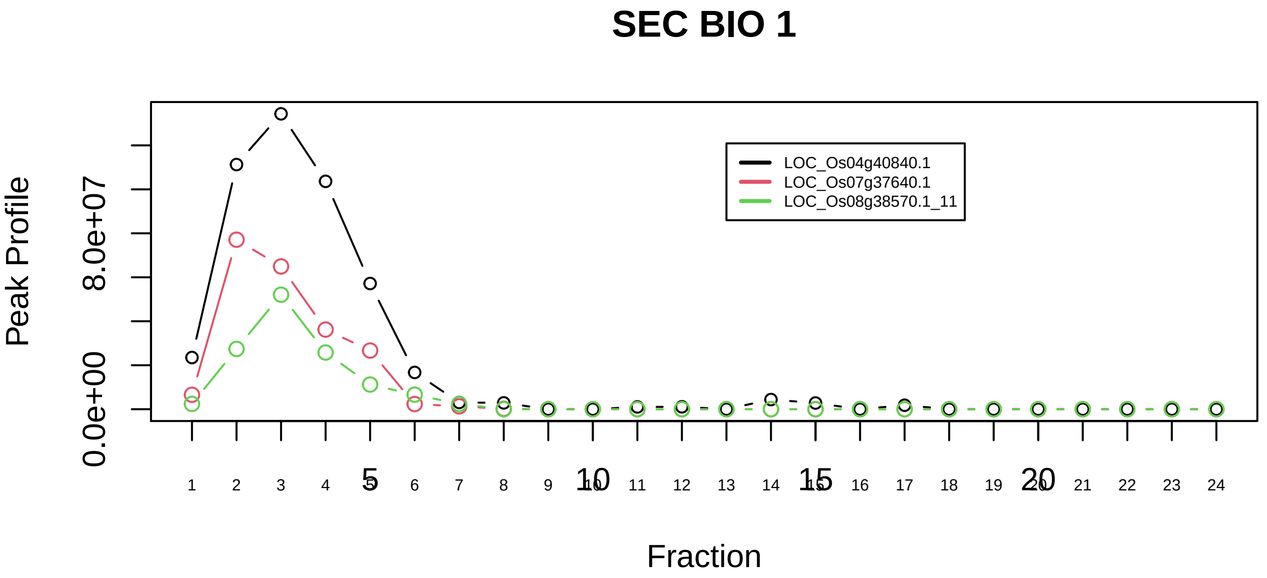

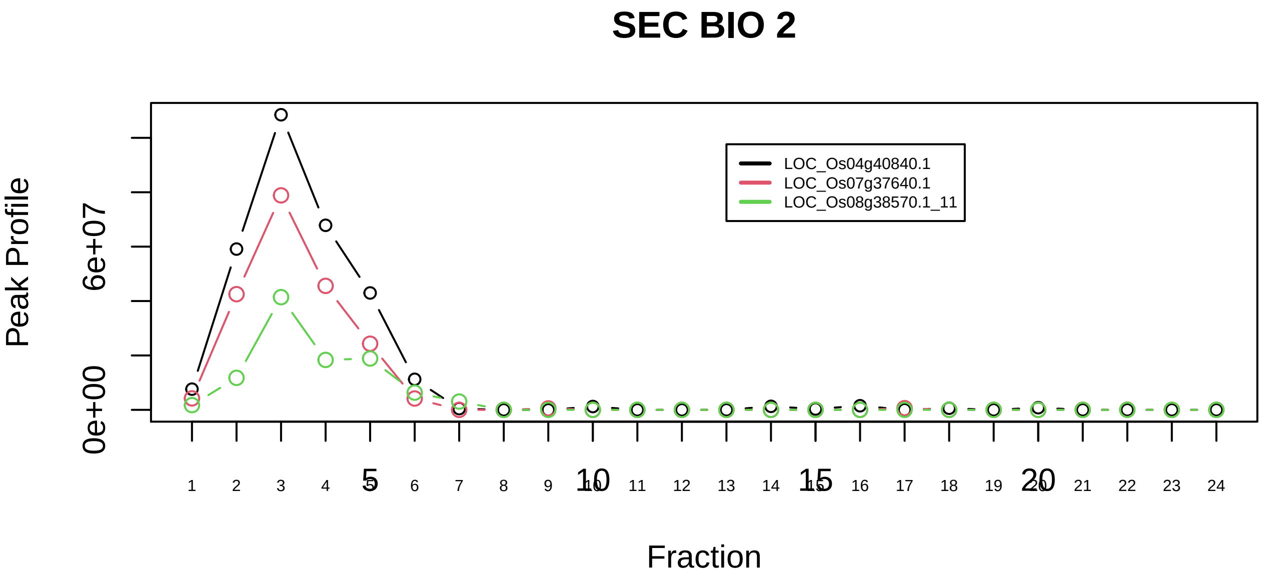

Elongator holo complex|1380−−all_prots_before_clustering_GPFdist

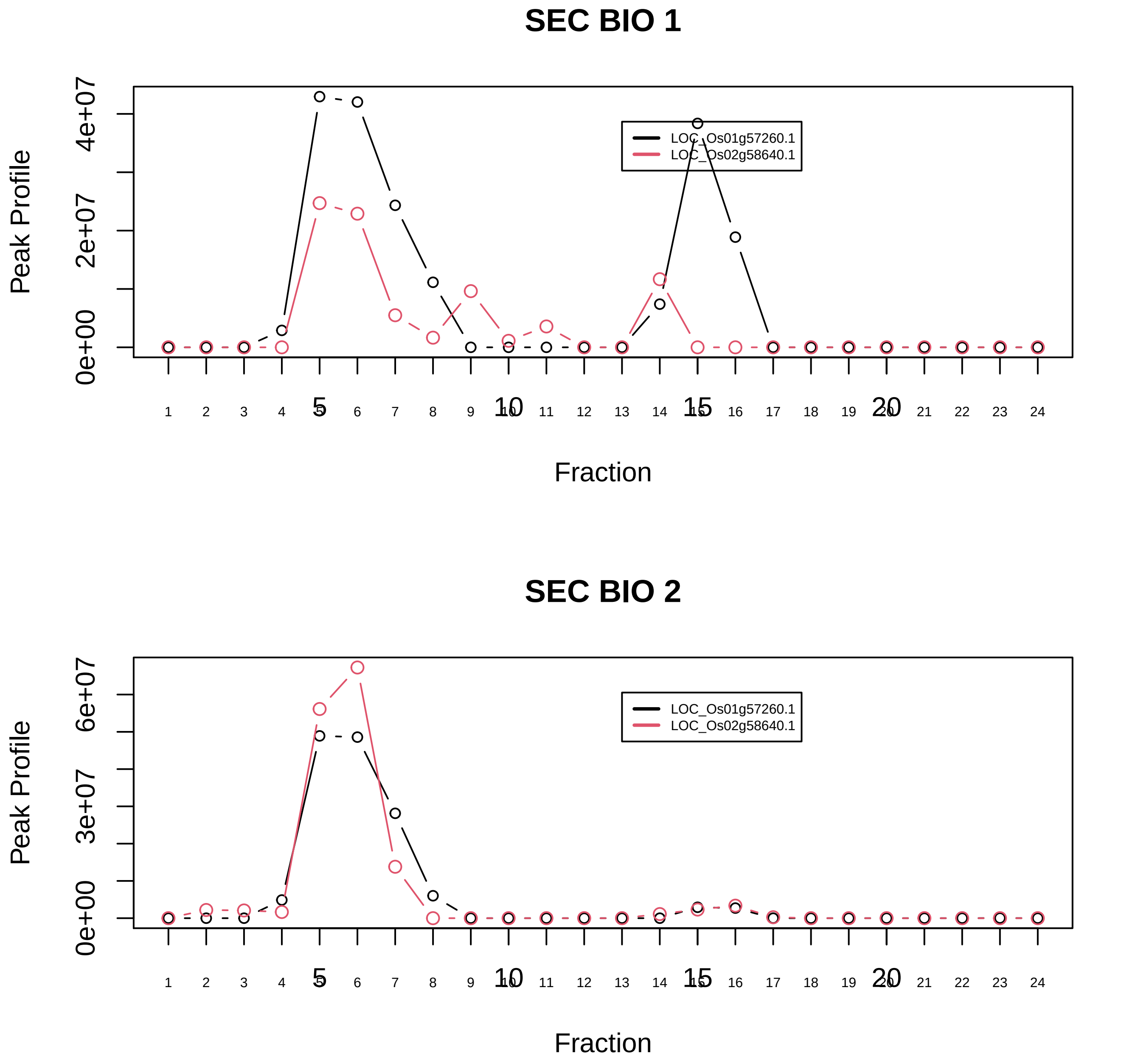

TSG101−VPS37B−VPS28 complex|1863−−only_2_prots_before_clustering

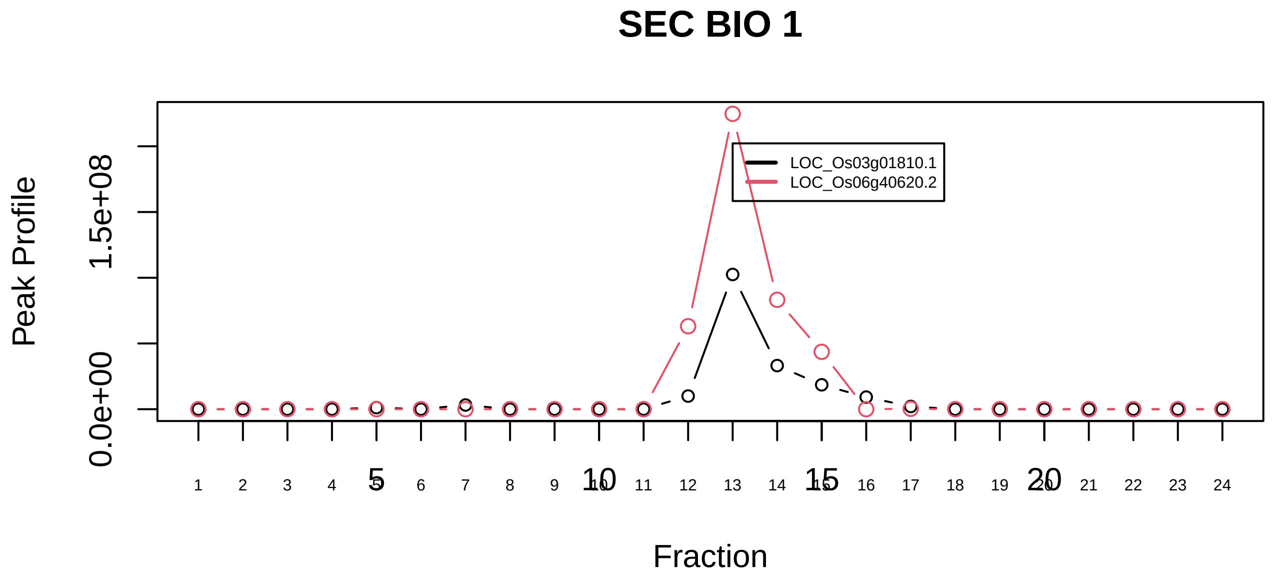

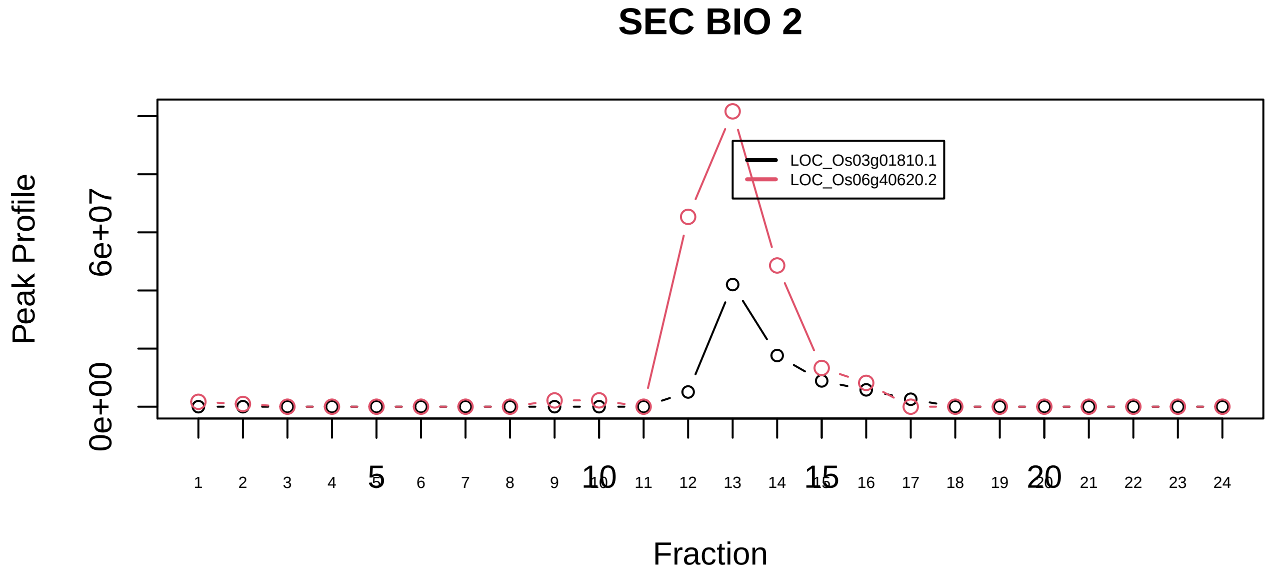

ESCRT−III complex|1186−−only_2_prots_before_clustering

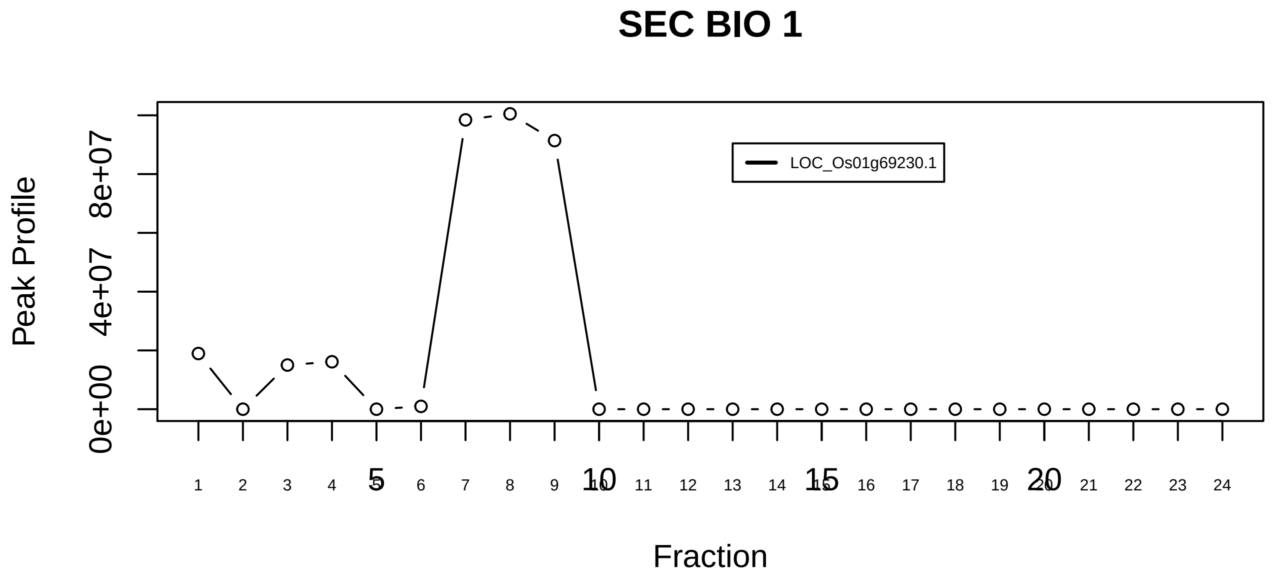

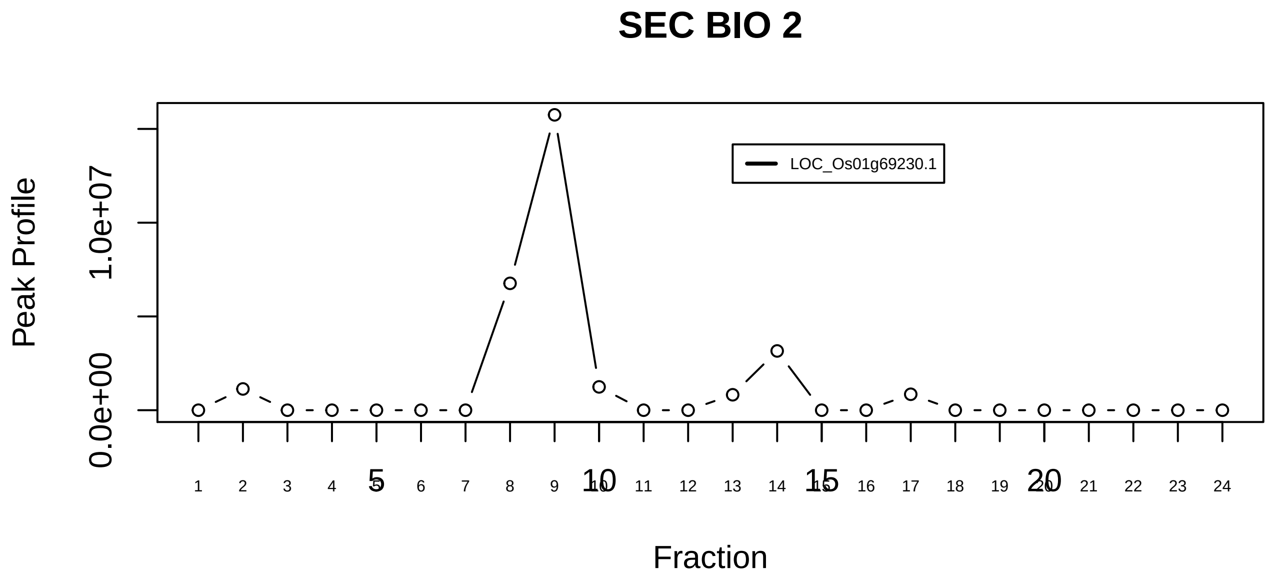

Exocyst Sec6/8 complex|89−−singleton_1

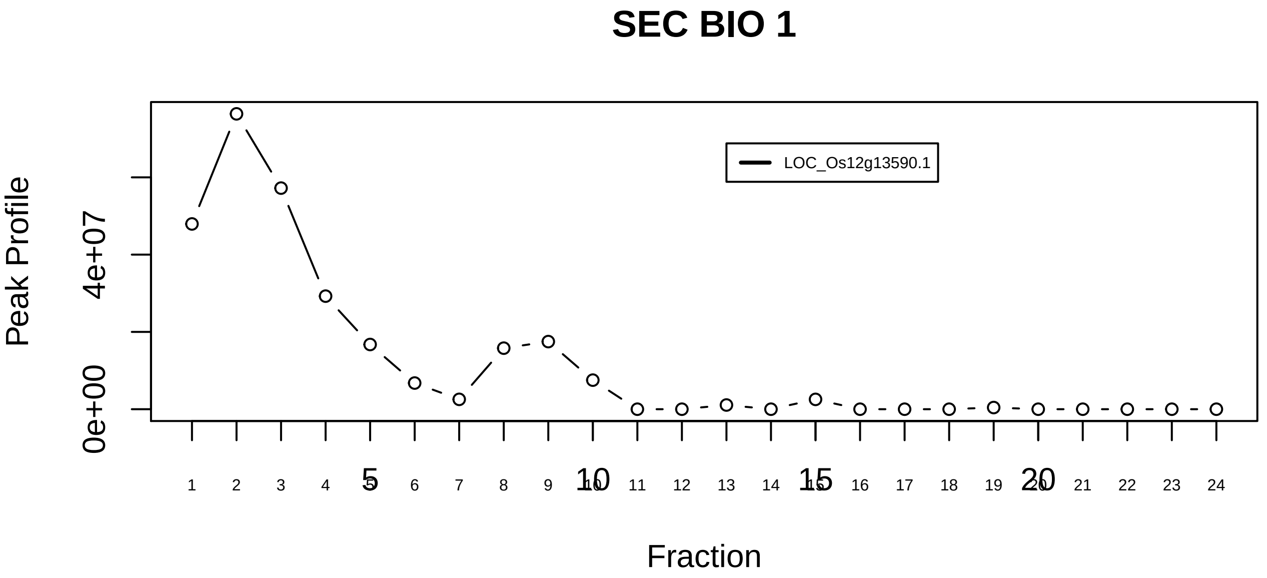

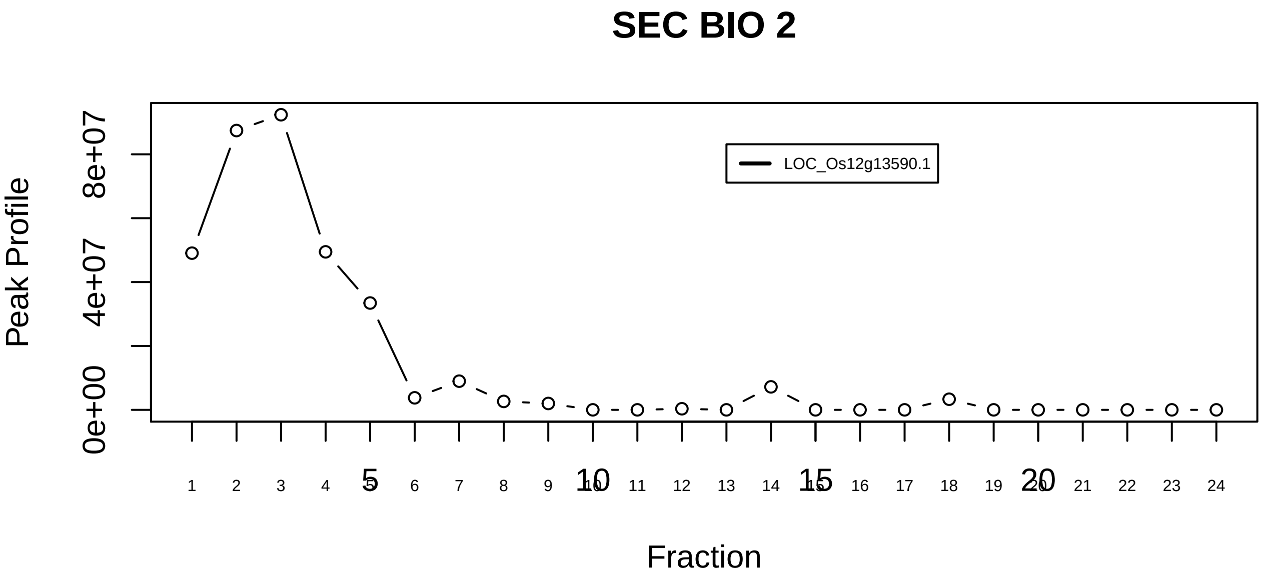

Exocyst Sec6/8 complex|89−−singleton_2

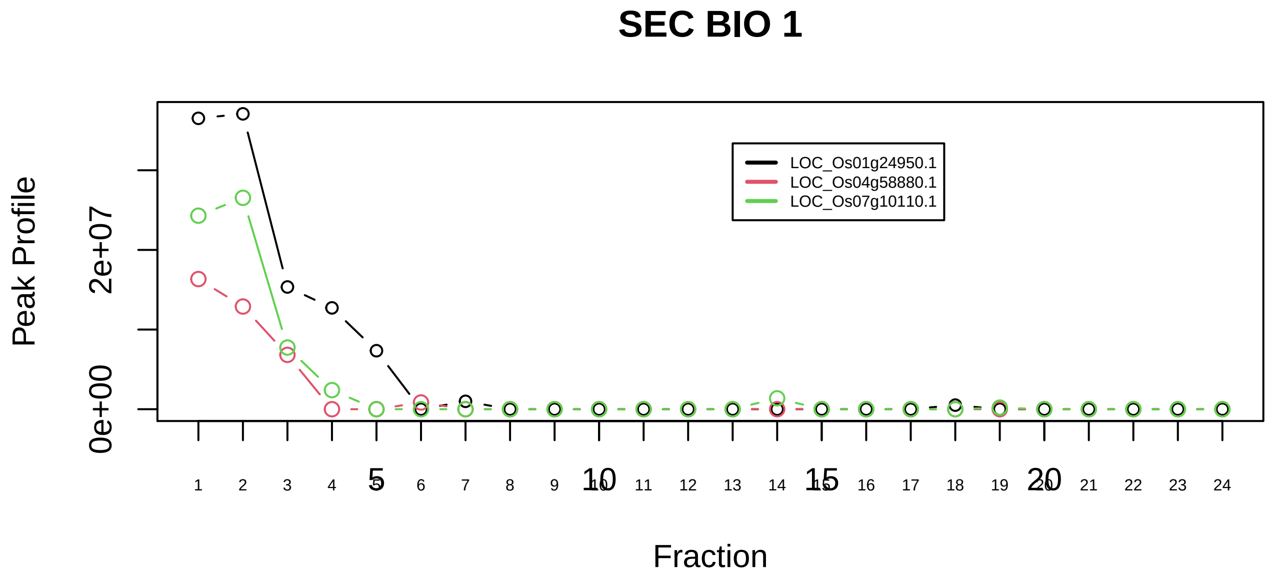

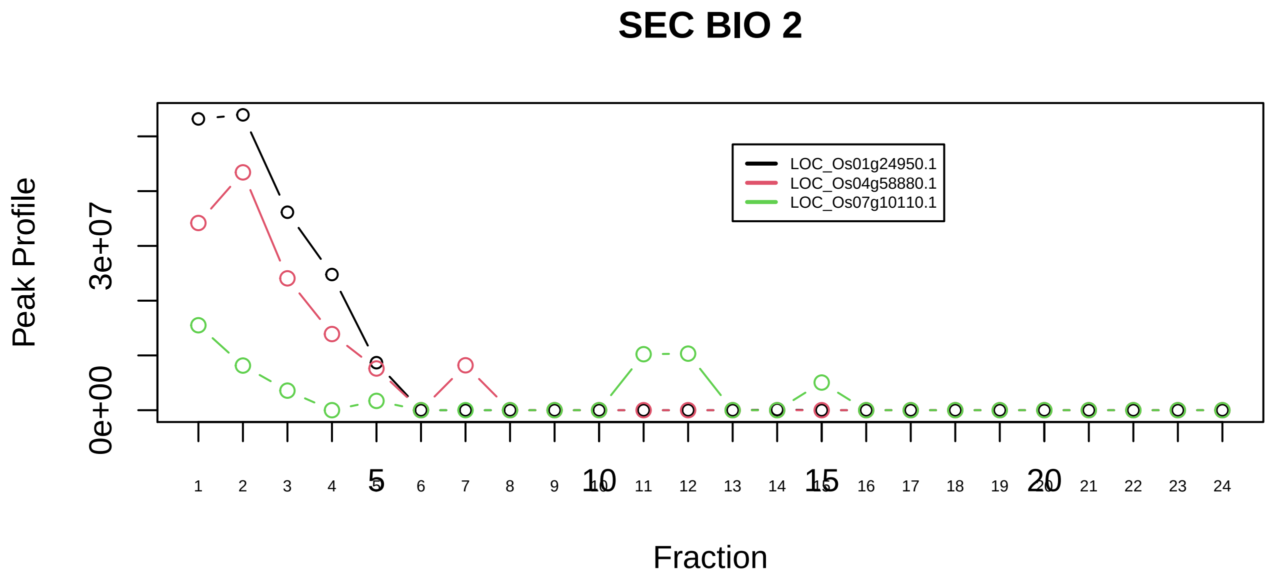

Exocyst Sec6/8 complex|89−−sub_1

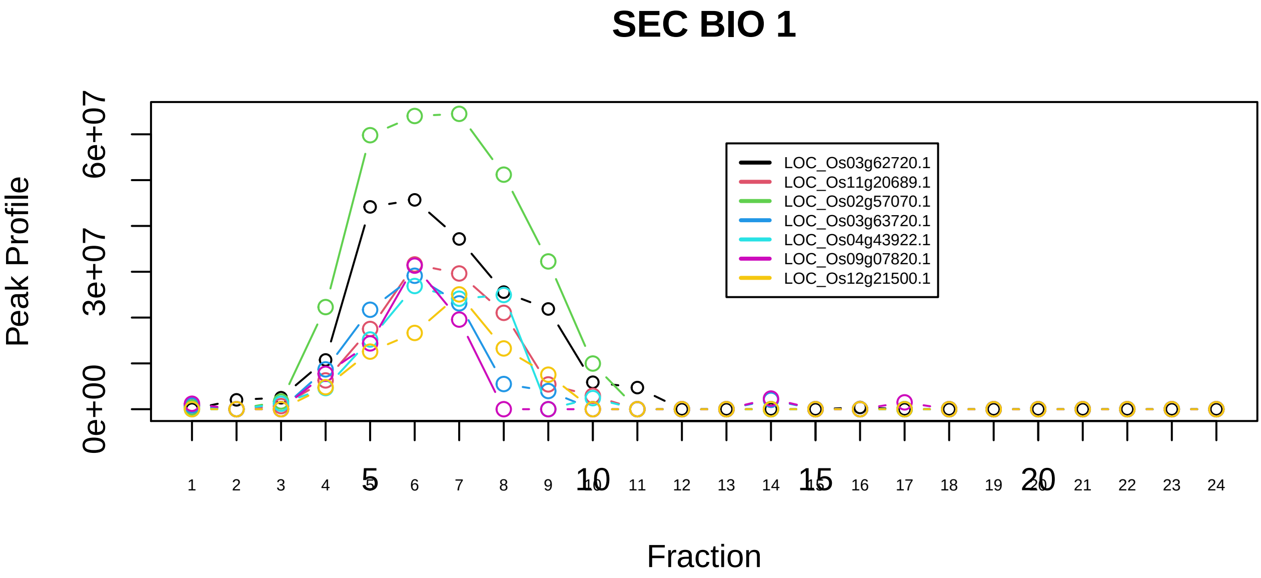

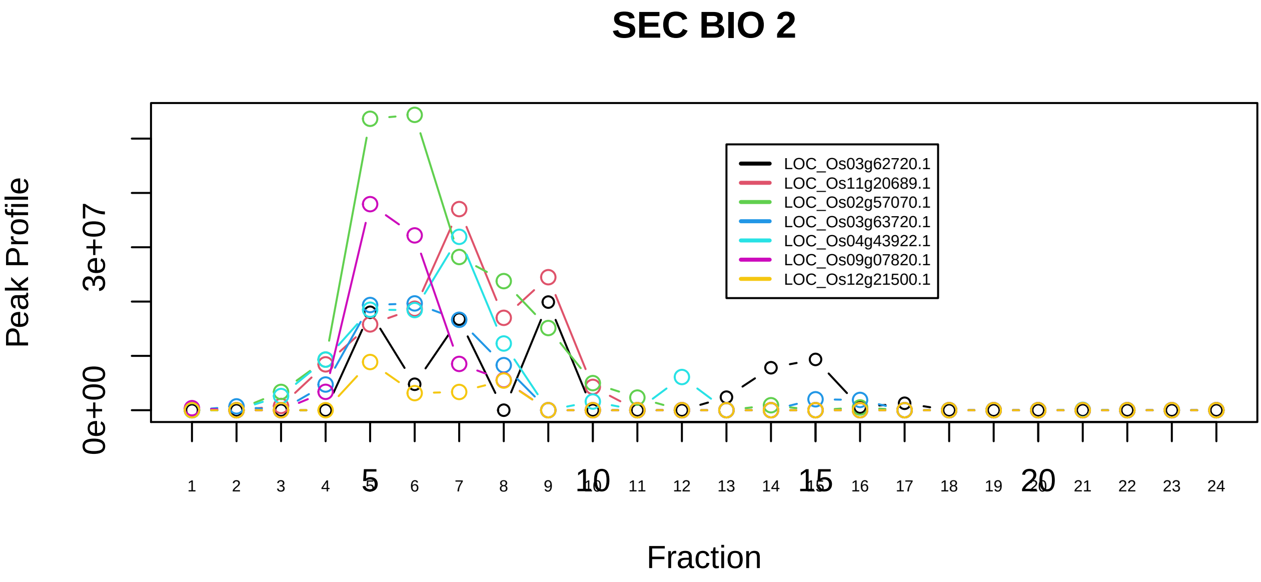

Exosome|789−−sub_1

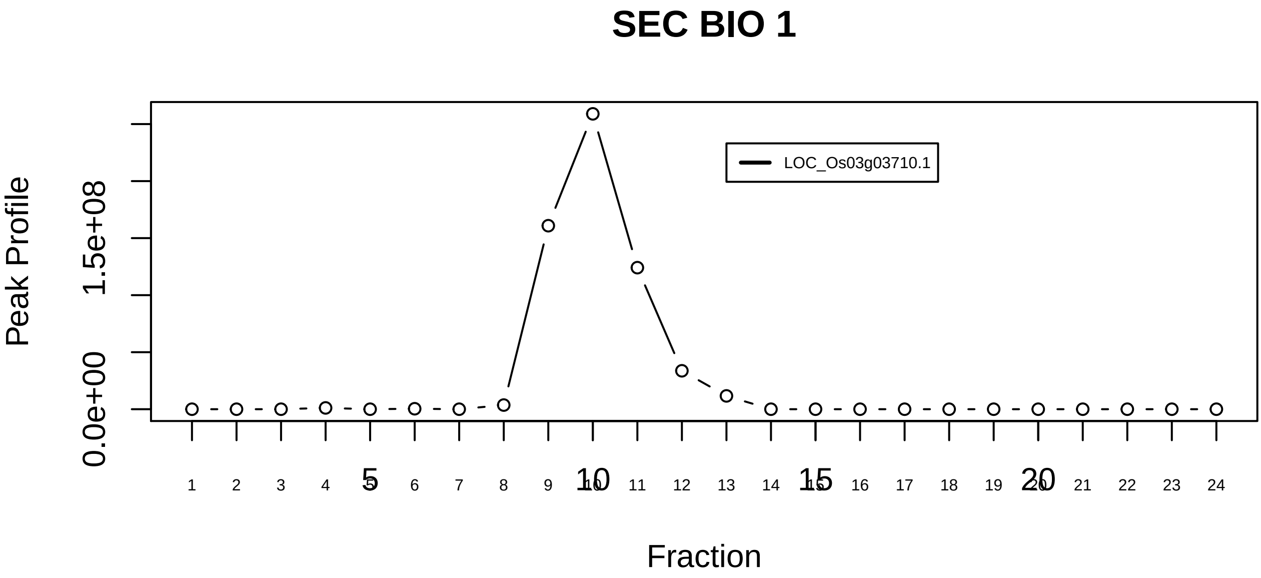

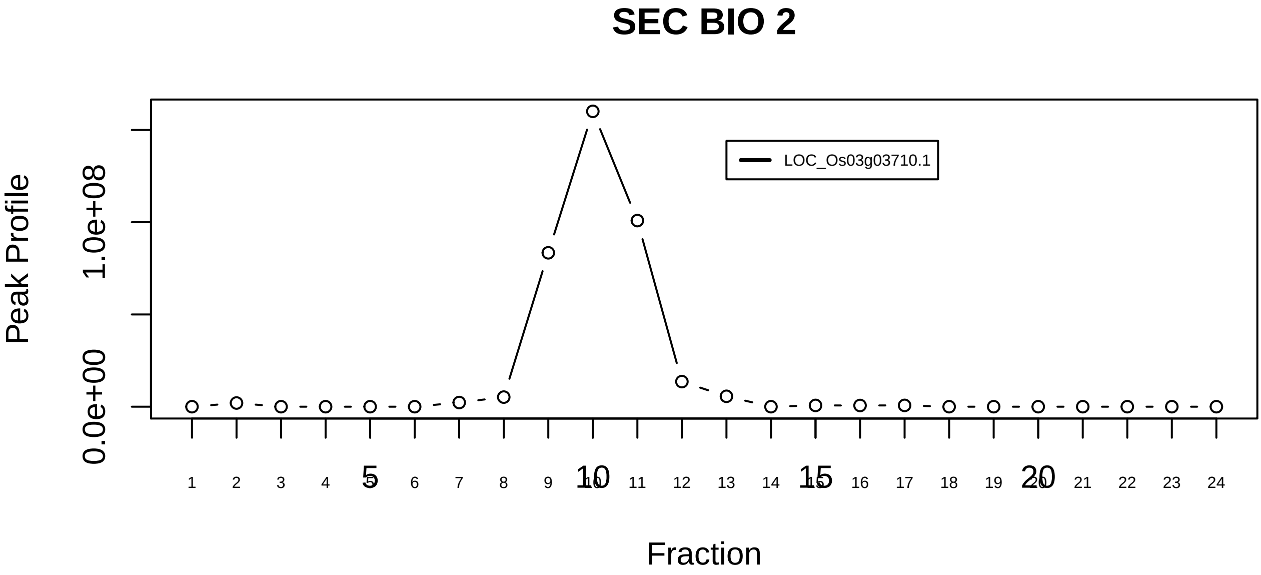

Exosome|789−−singleton_1

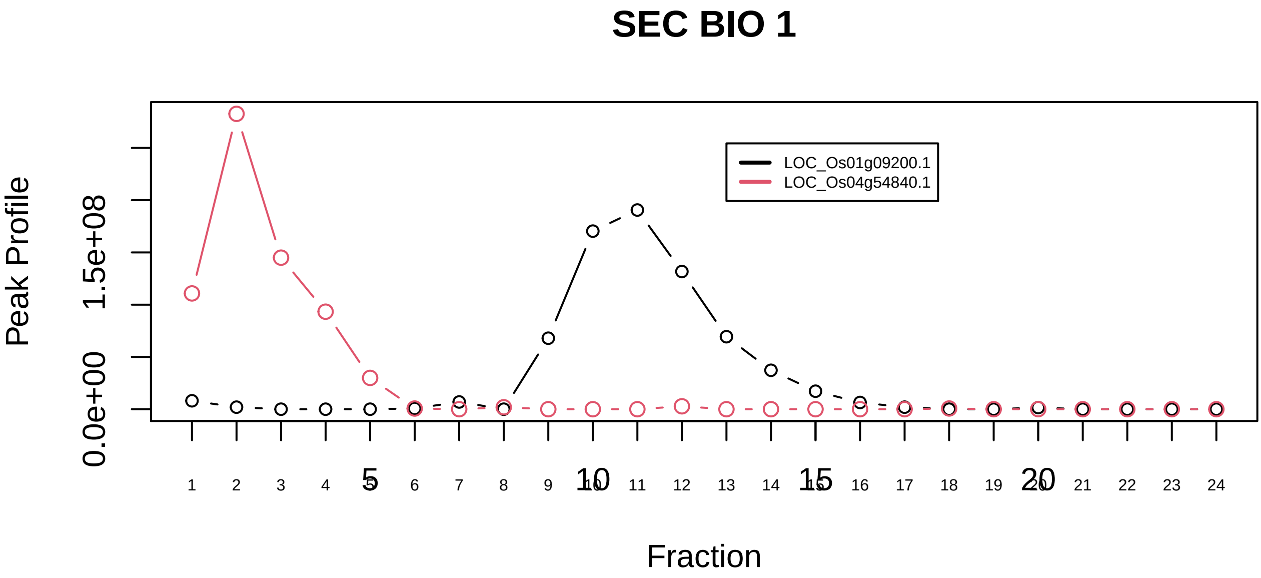

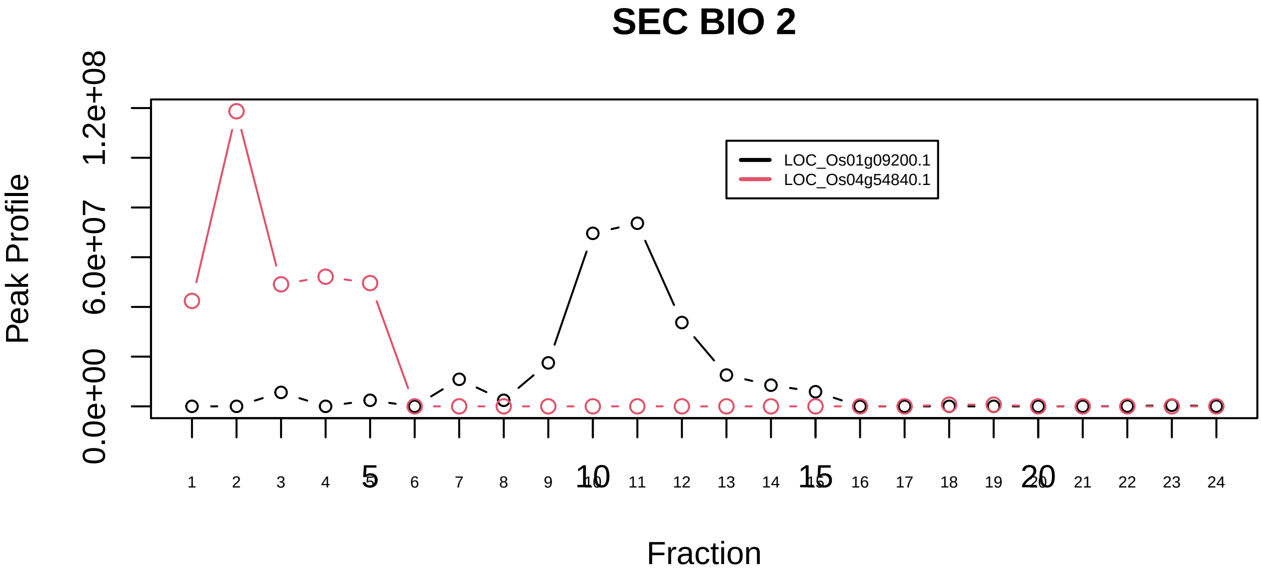

FCP1−associated protein complex|811−−sub_1

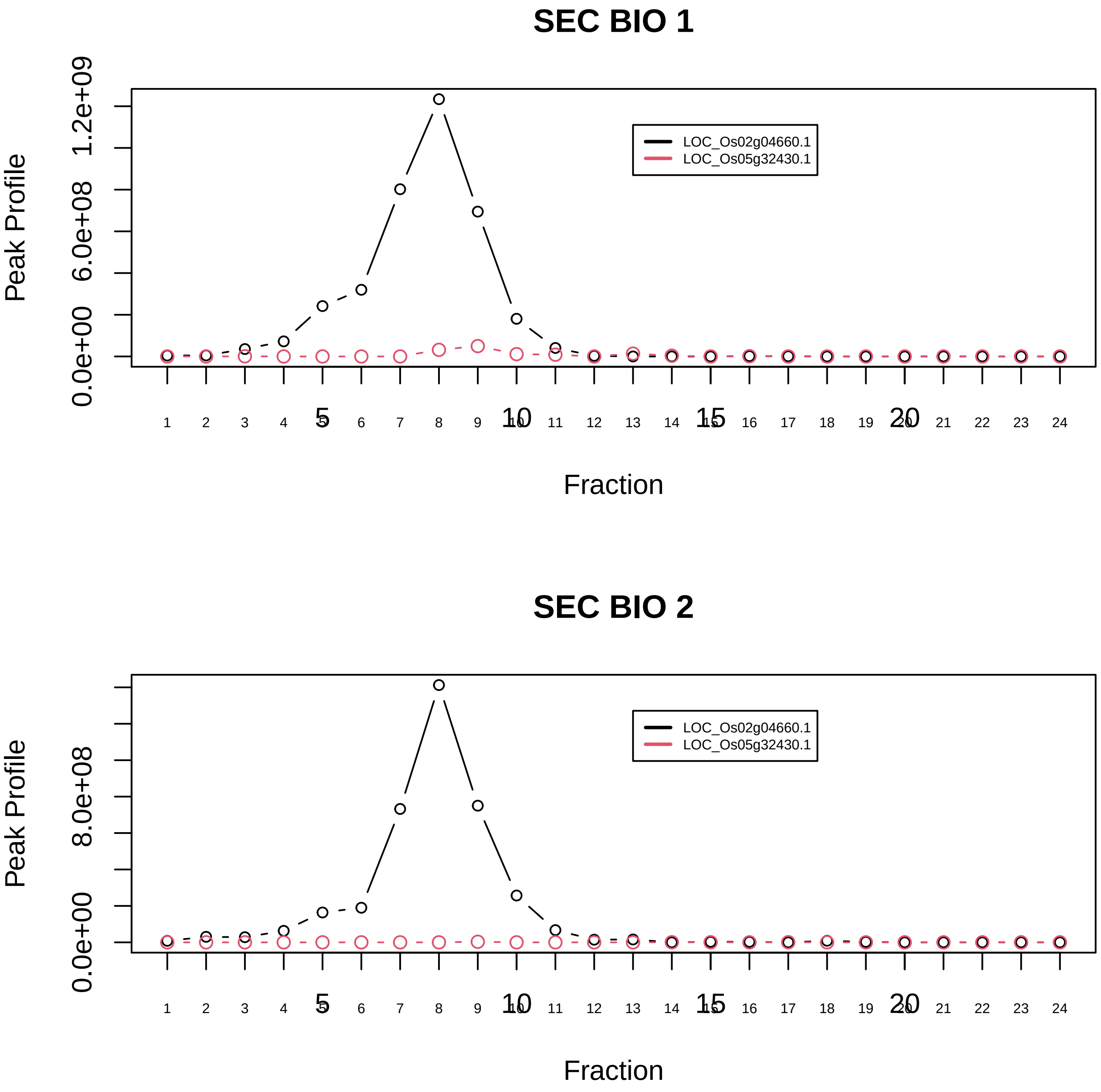

FCP1−associated protein complex|811−−sub_2

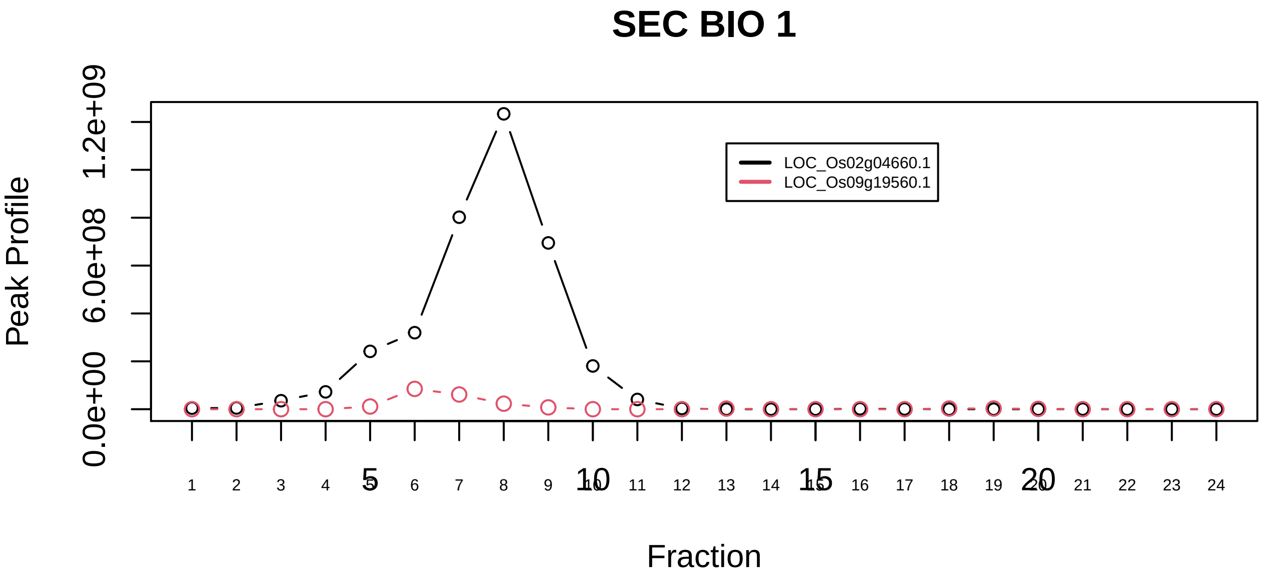

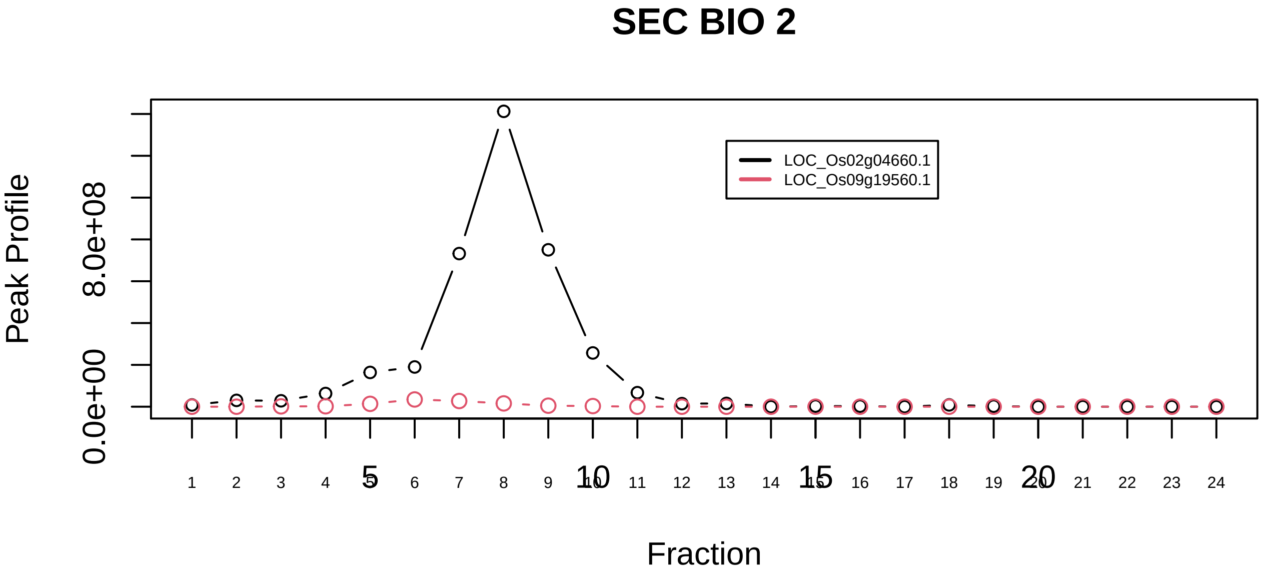

FIB−associated protein complex|1231−−sub_1

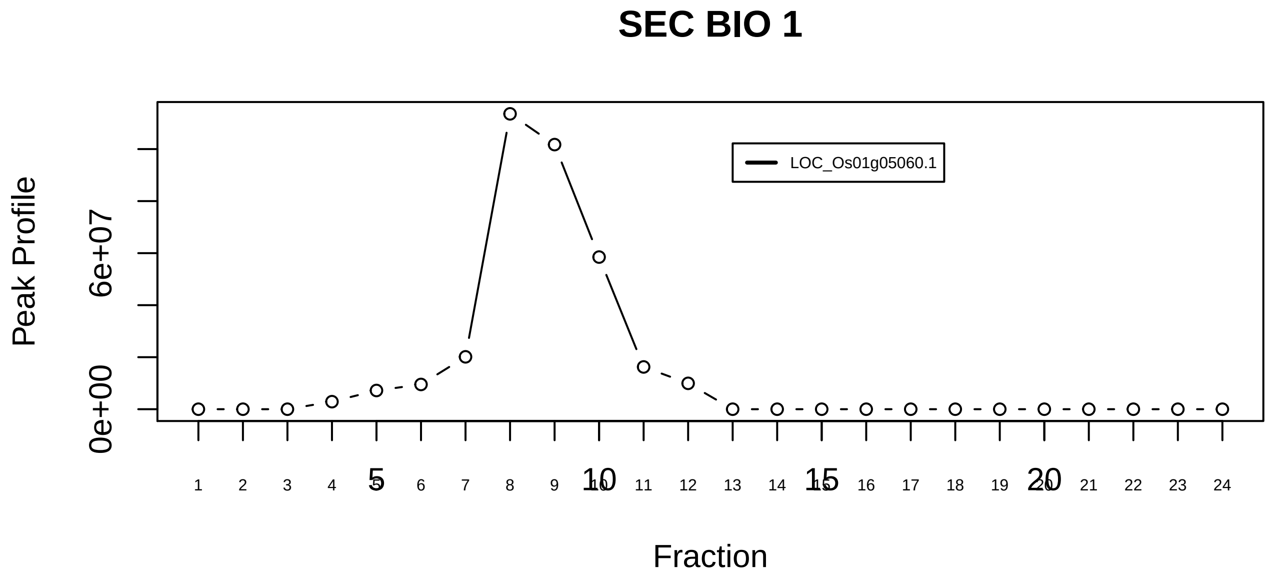

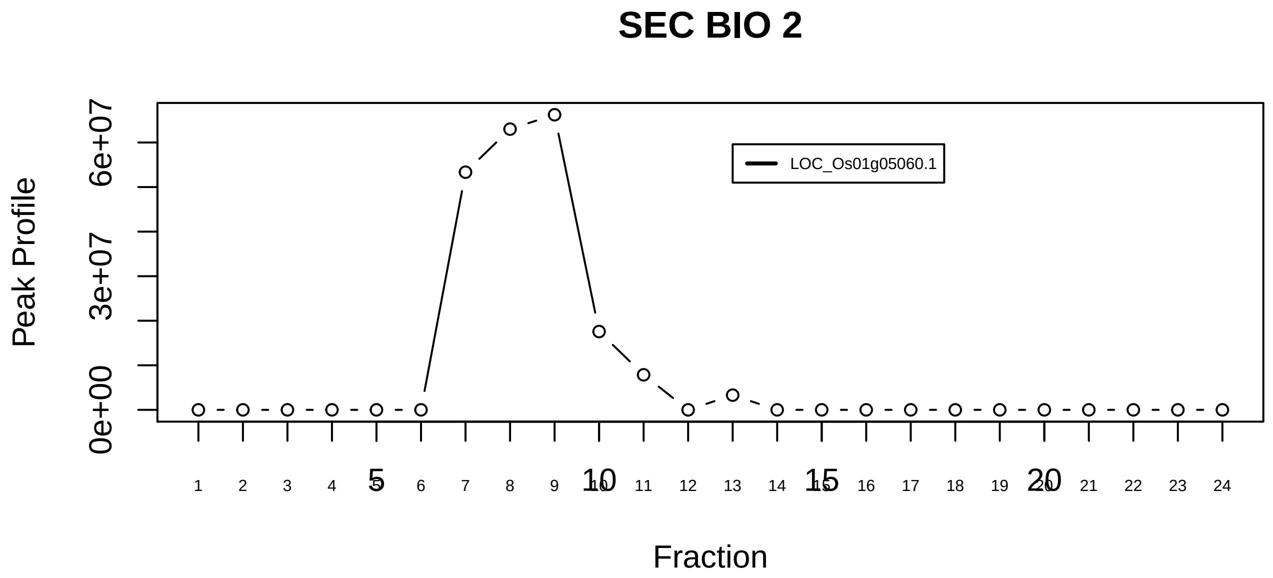

FIB−associated protein complex|1231−−singleton_1

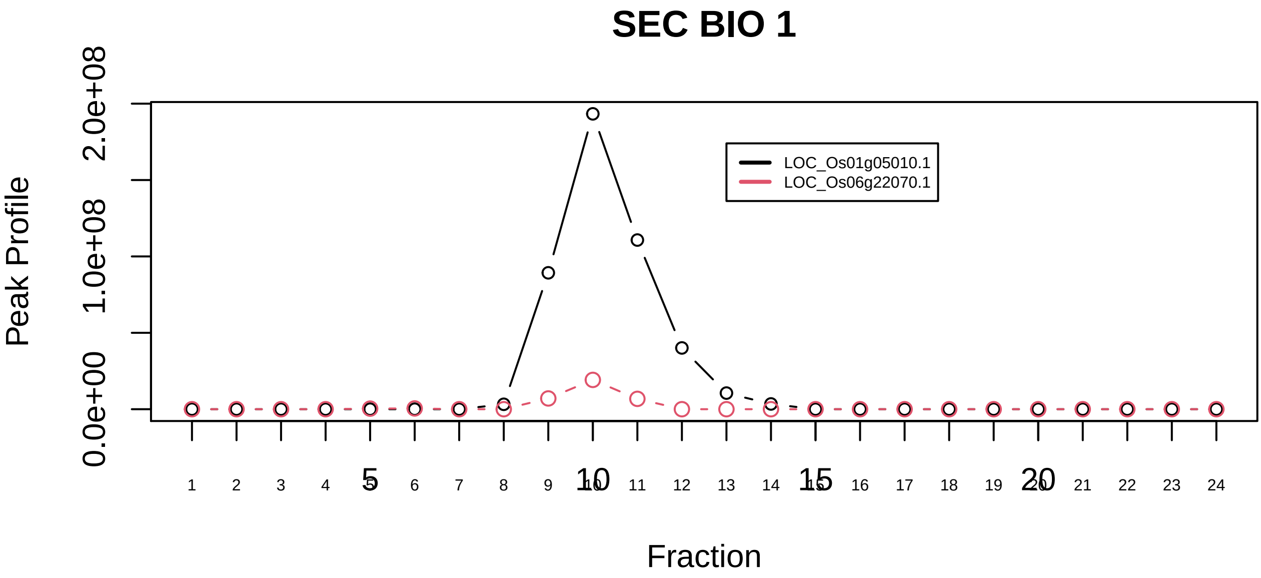

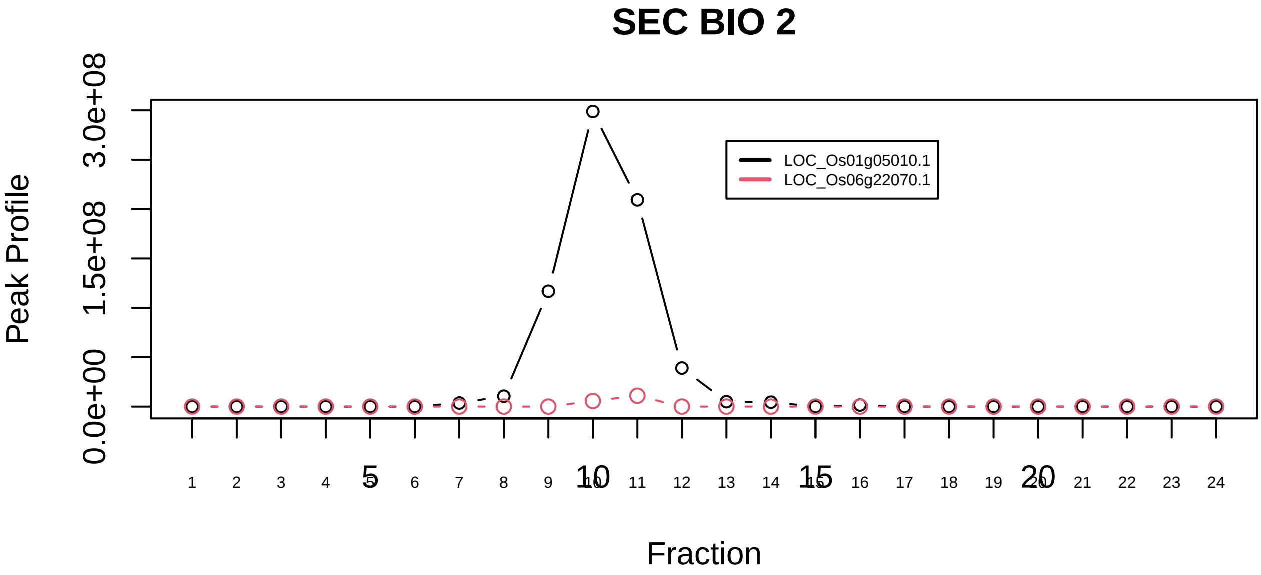

FIB−associated protein complex|1231−−sub_orthoparalog_1

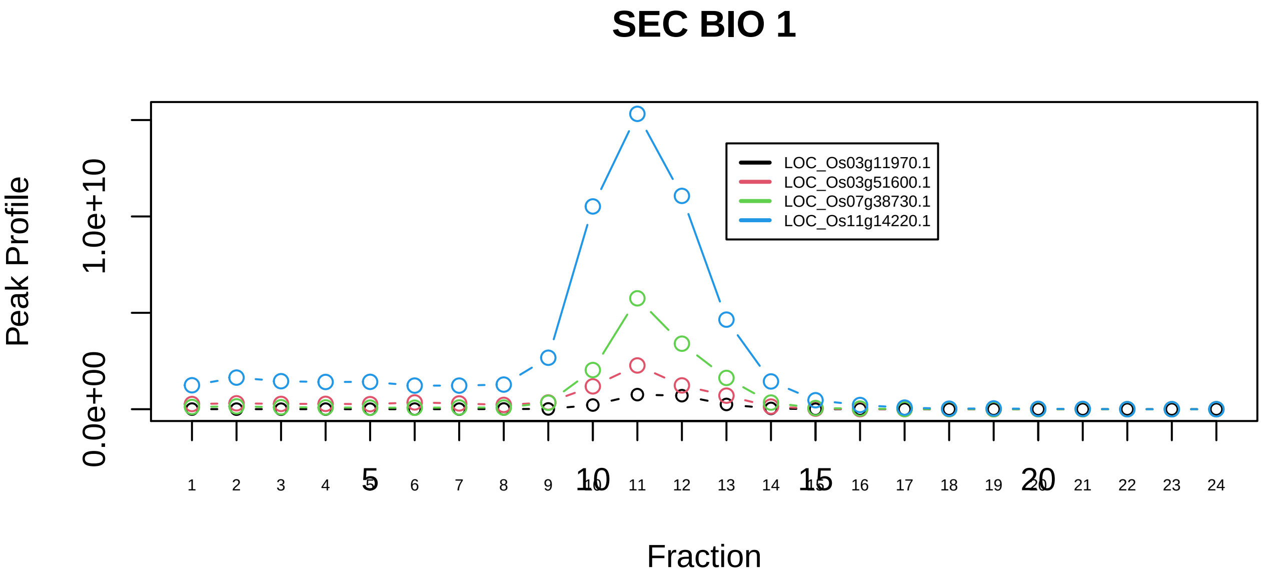

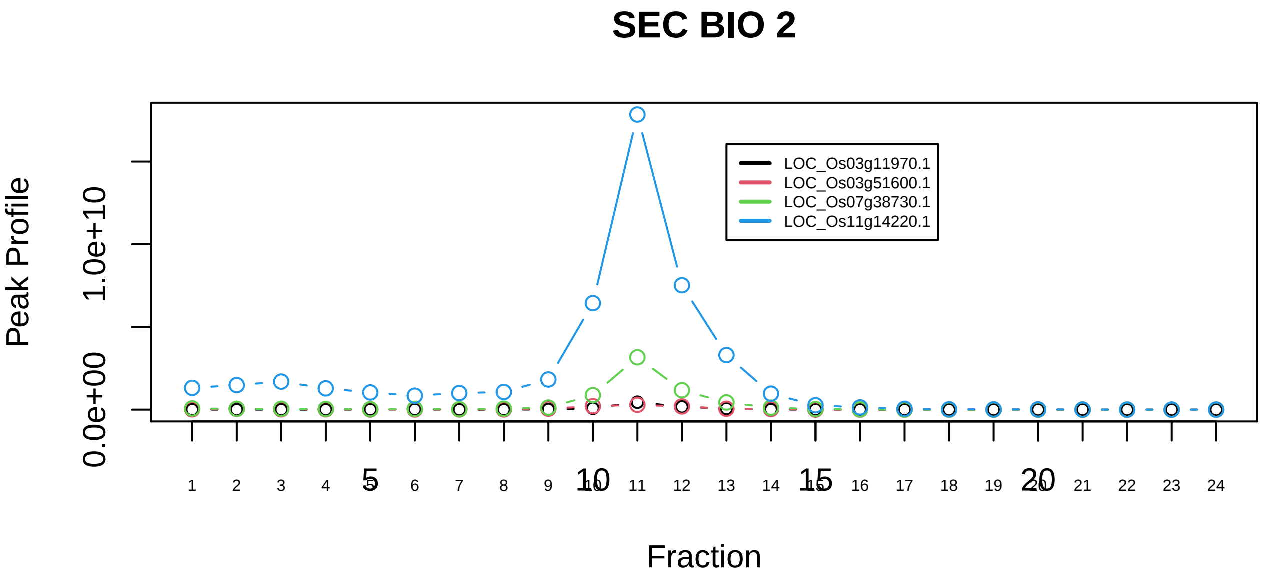

FIB−associated protein complex|1231−−sub_orthoparalog_2

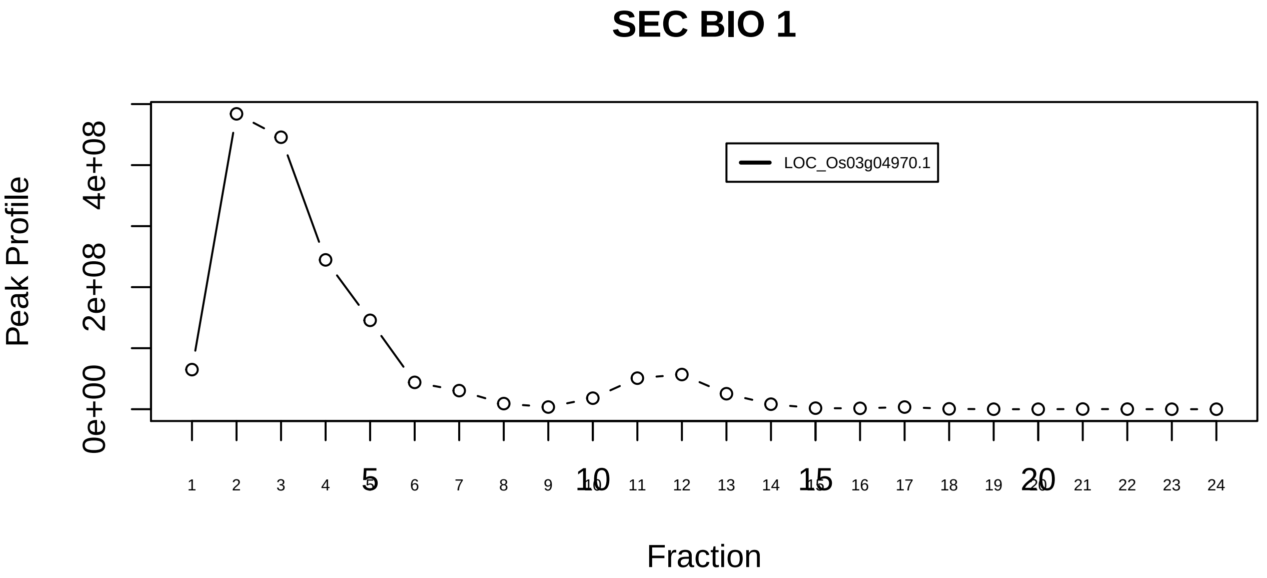

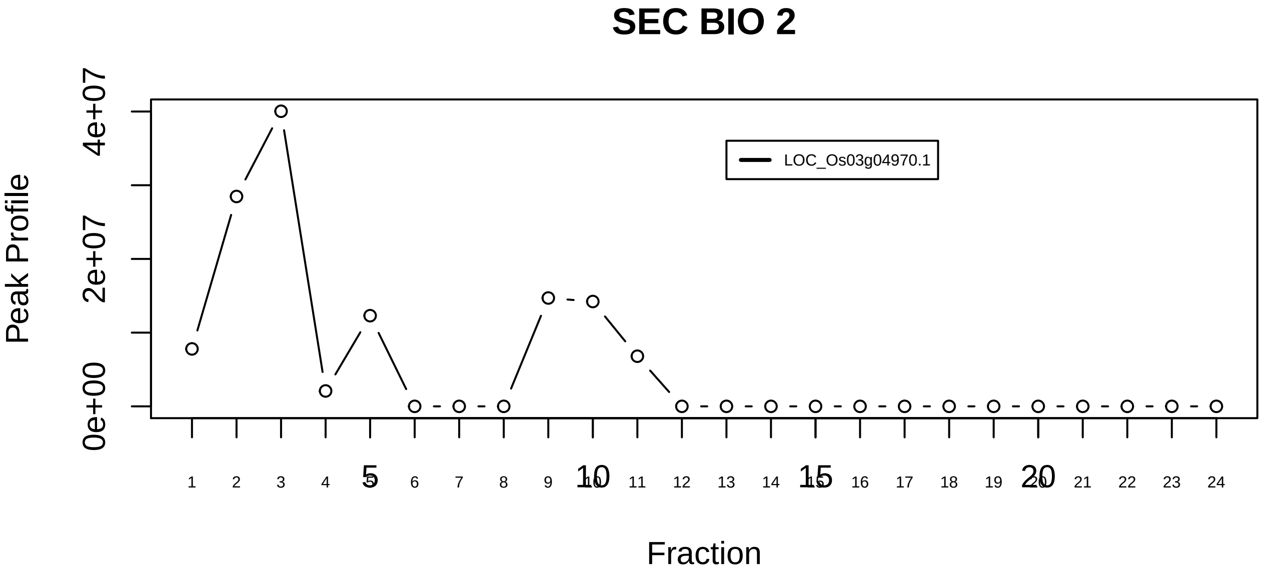

Frataxin complex|1094−−singleton_1

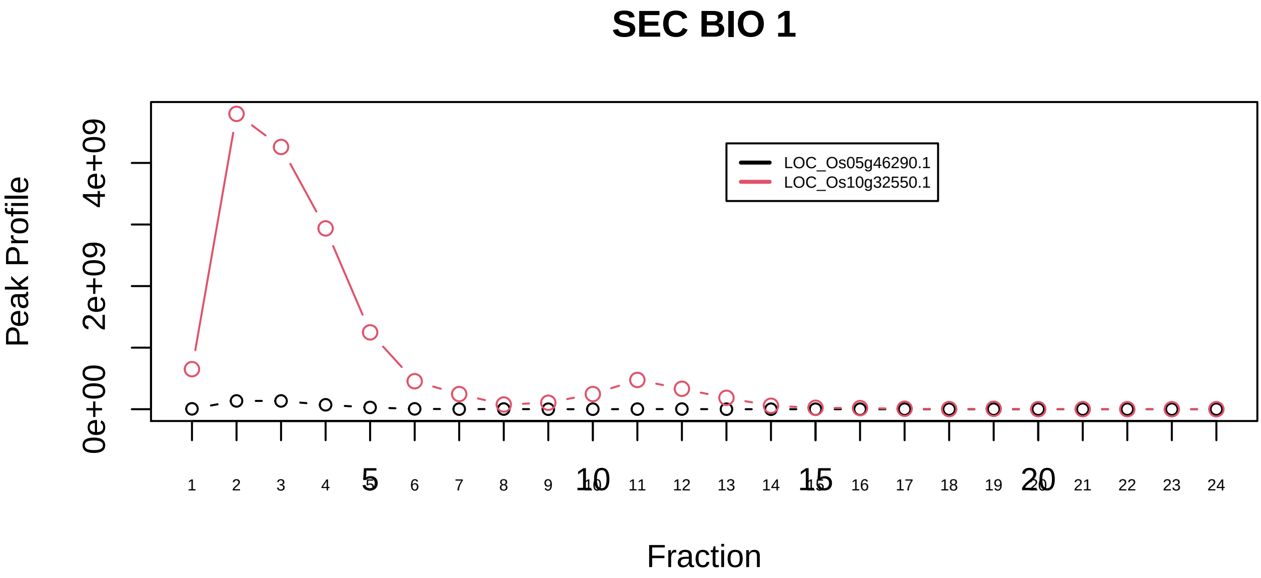

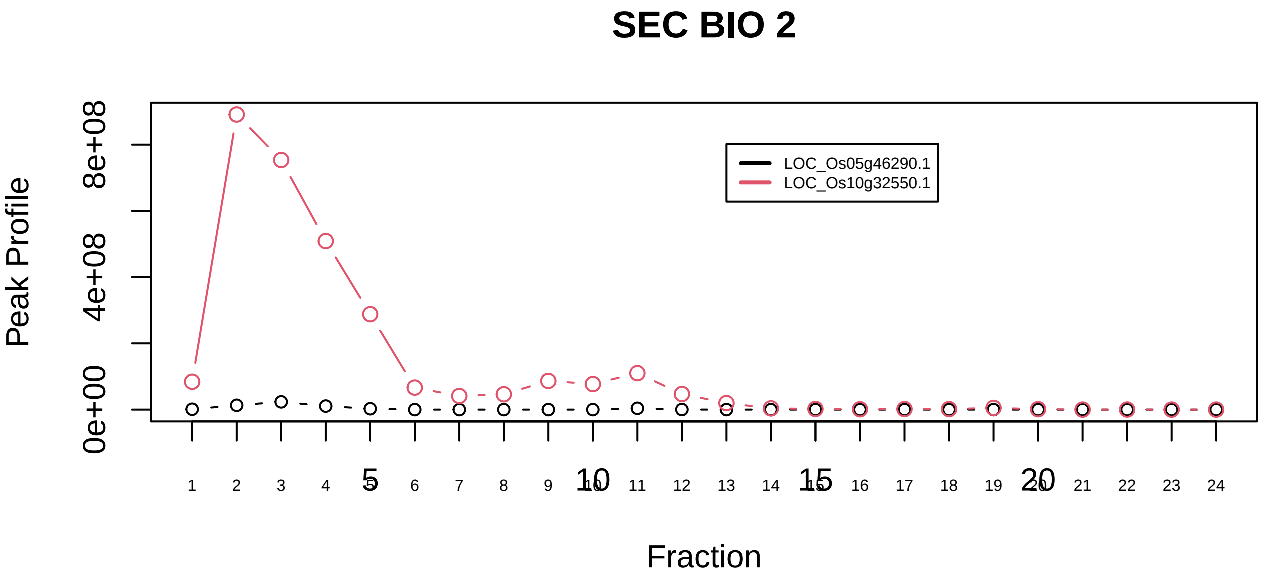

Frataxin complex|1094−−sub_orthoparalog_1

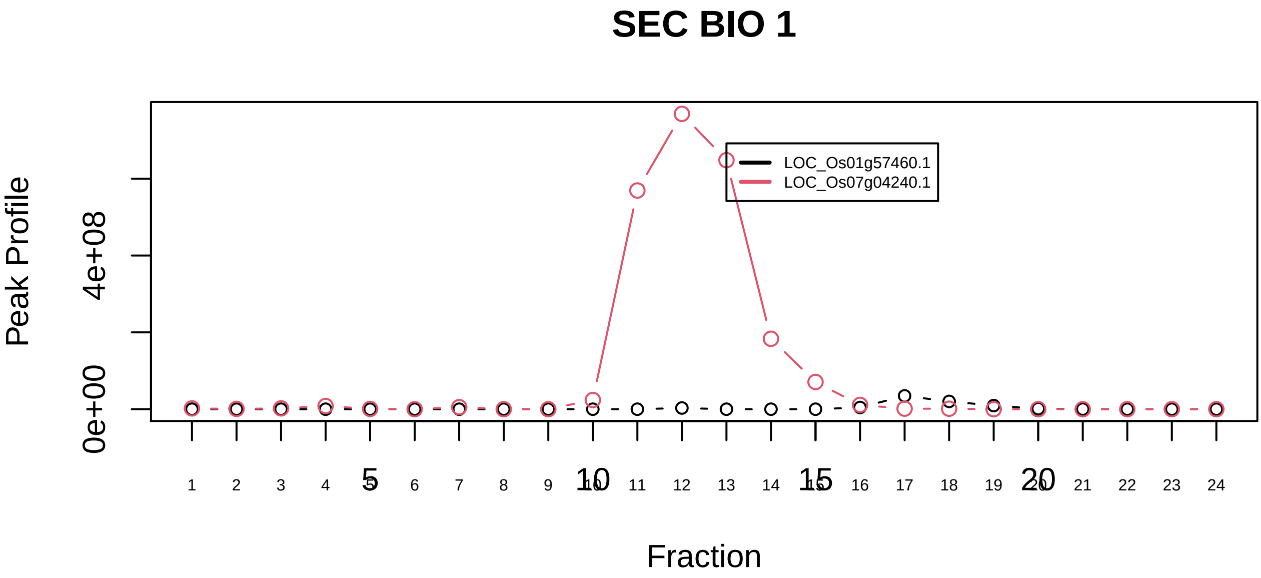

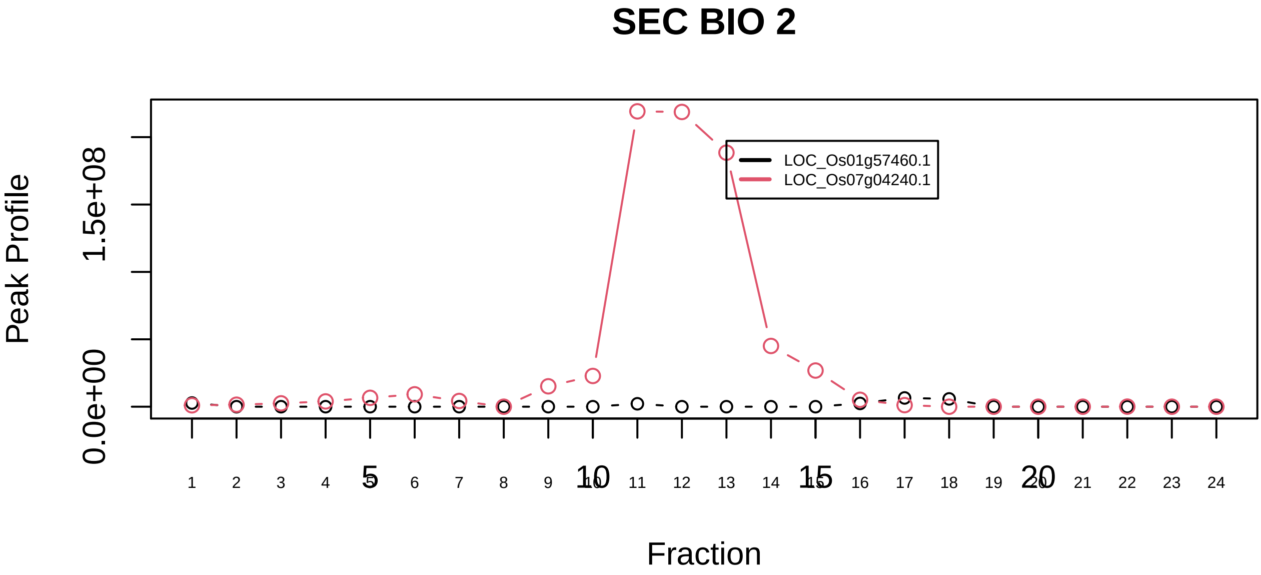

Frataxin complex|1094−−sub_1

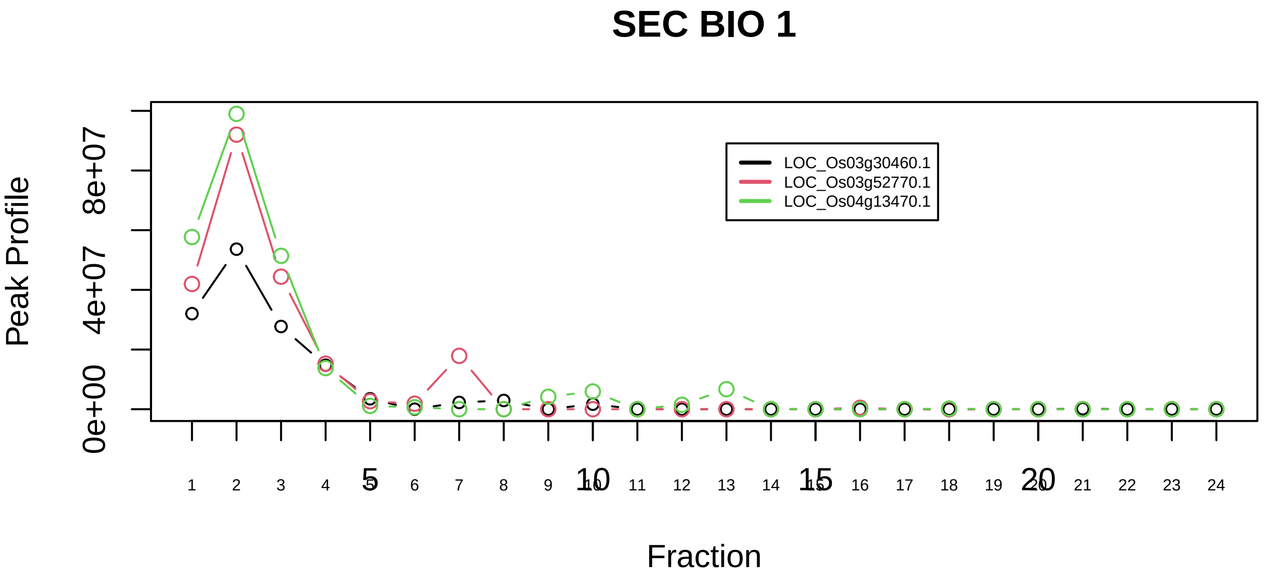

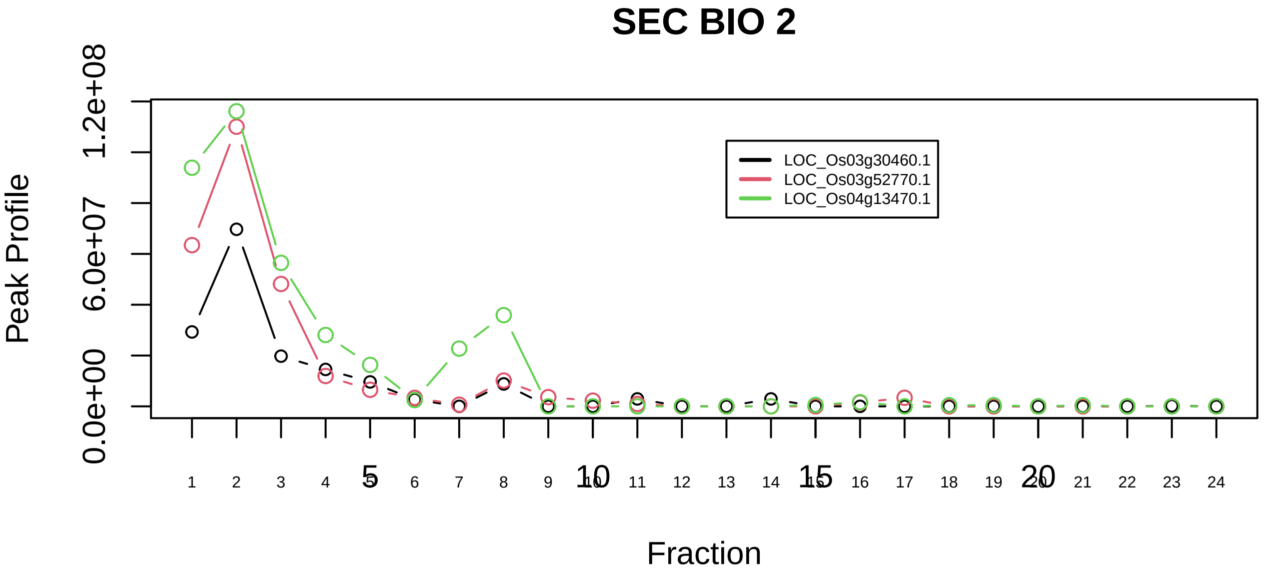

GARP complex|6463−−all_prots_before_clustering_GPFdist

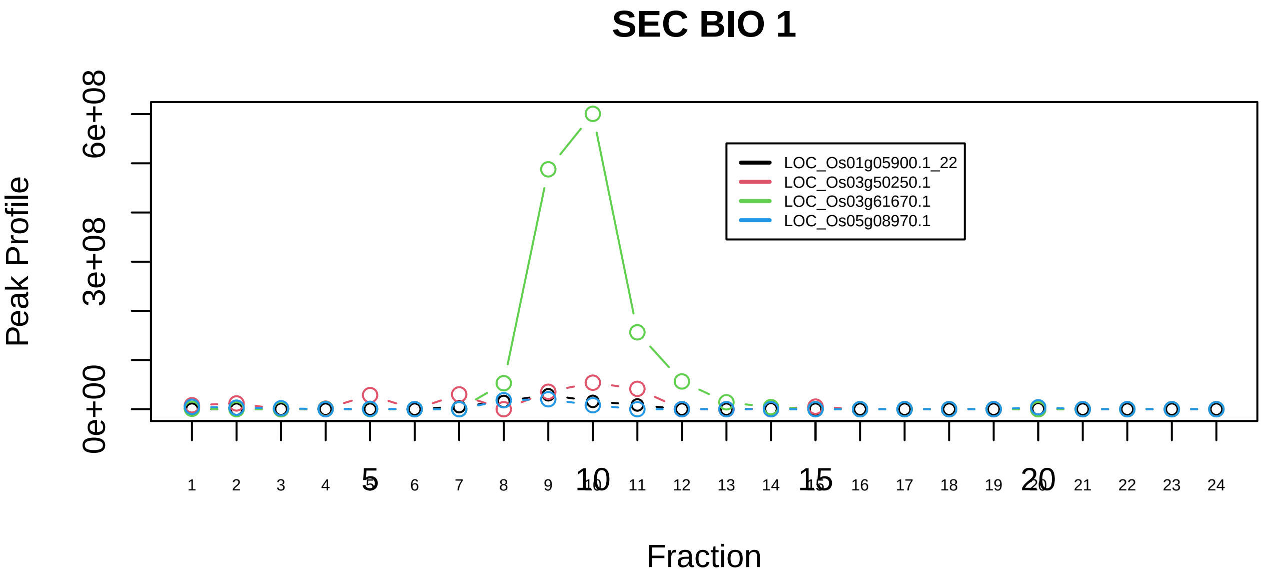

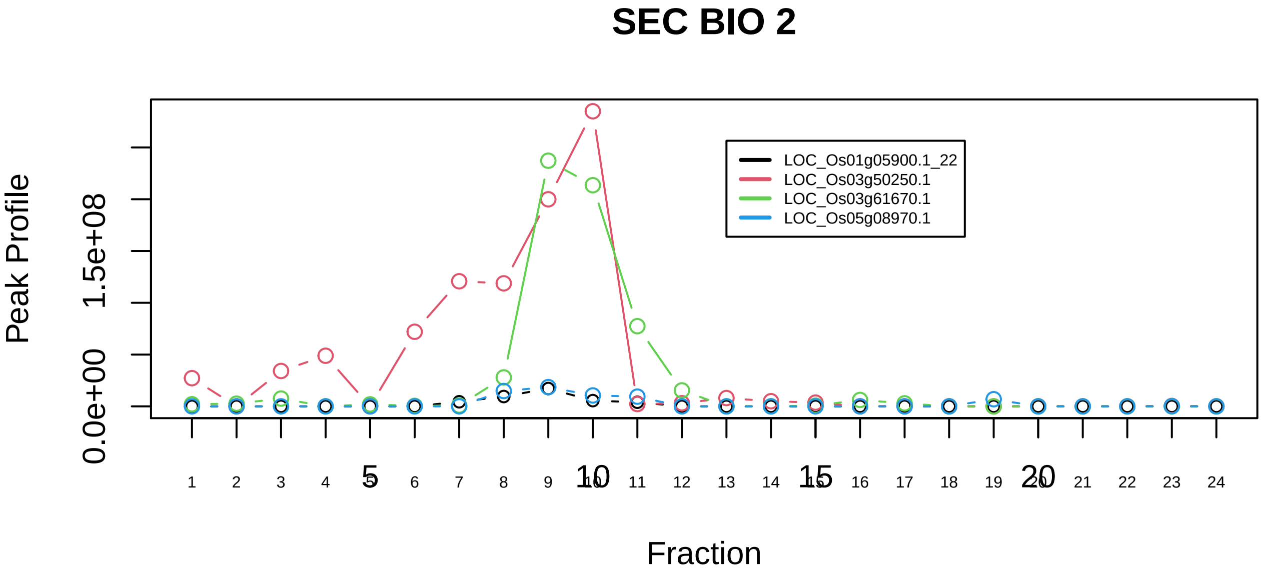

H2AX complex, isolated from cells without IR exposure|1223−−sub_1

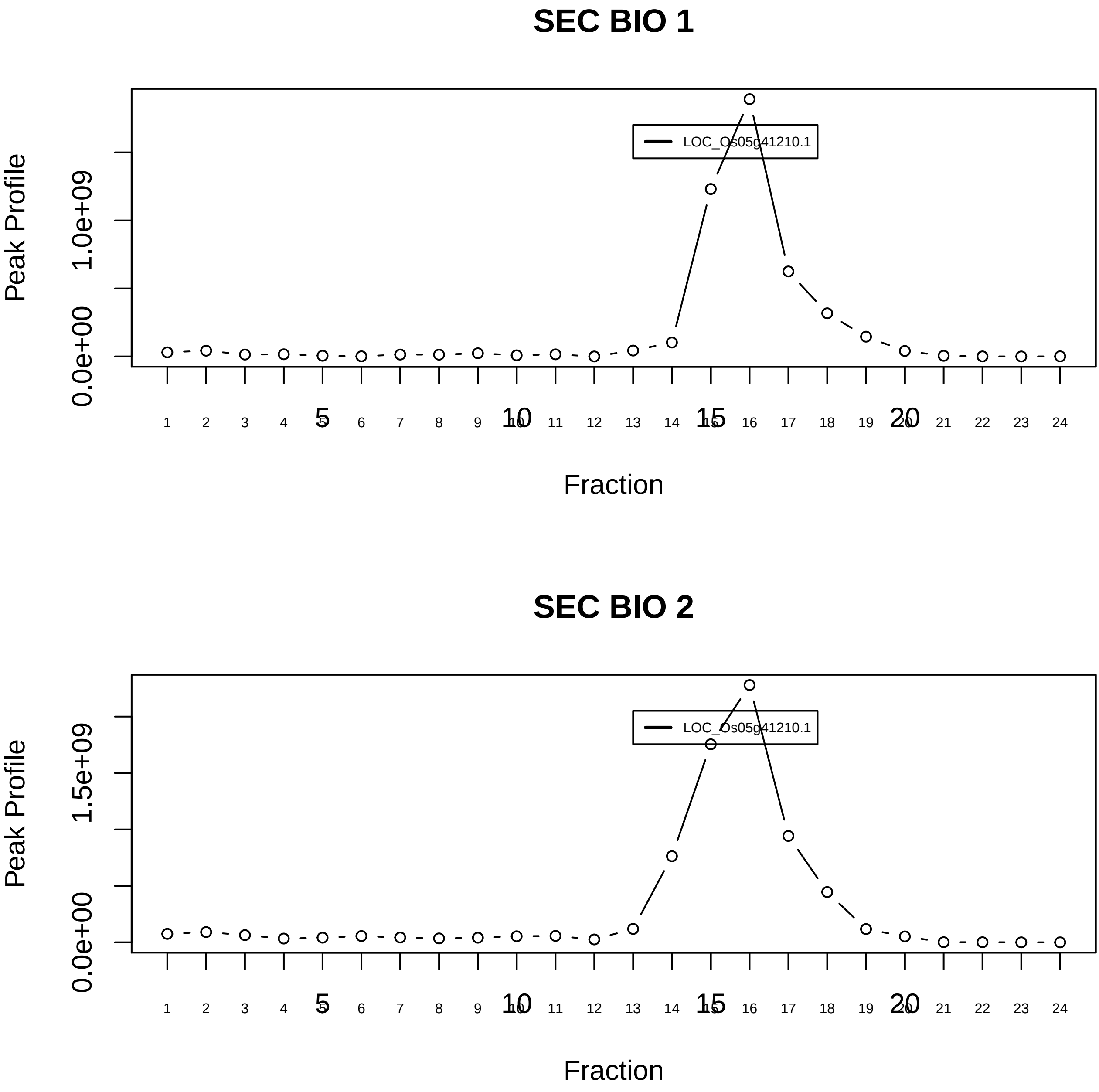

H2AX complex, isolated from cells without IR exposure|1223−−singleton_1

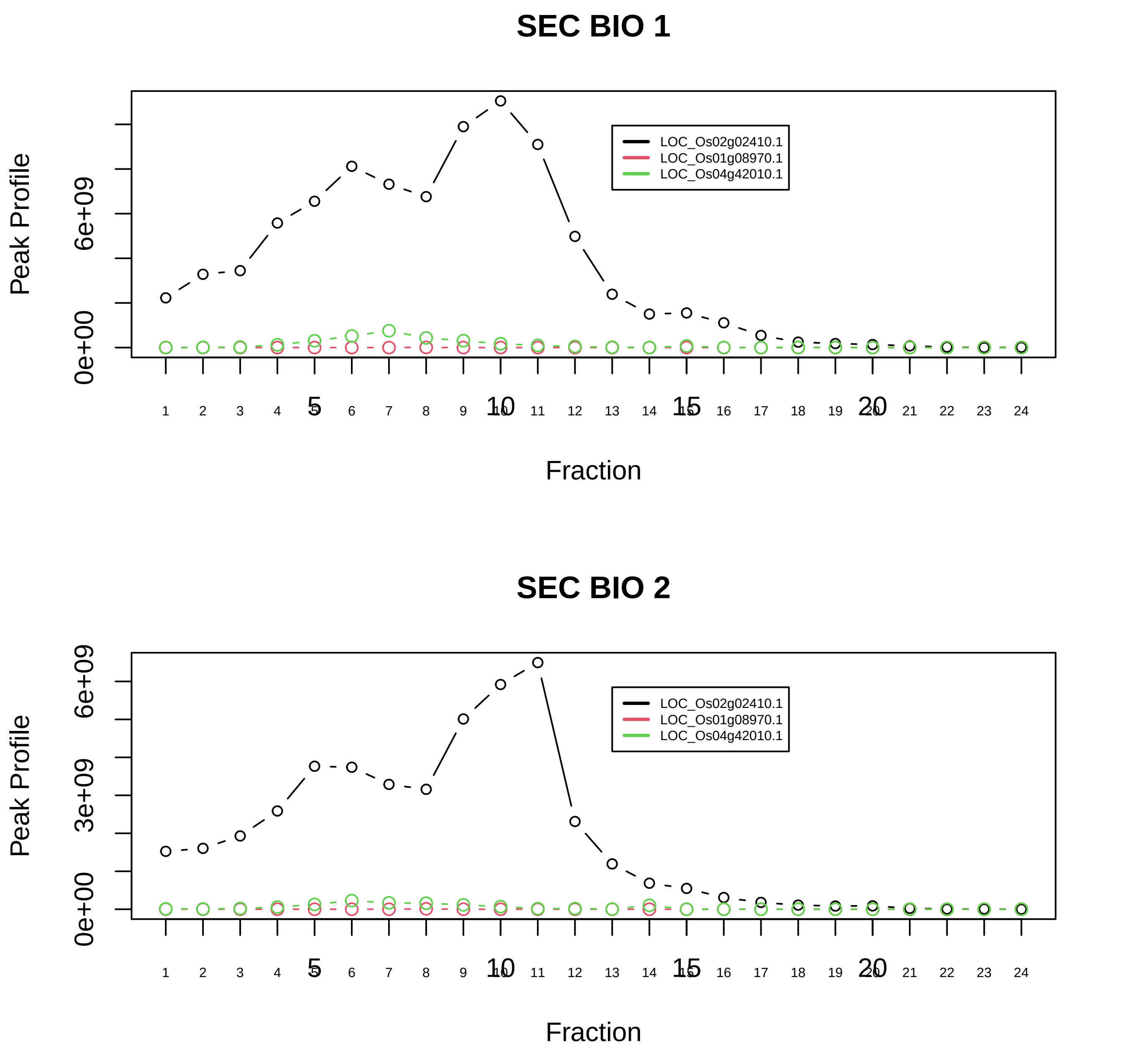

H2AX complex, isolated from cells without IR exposure|1223−−sub_2

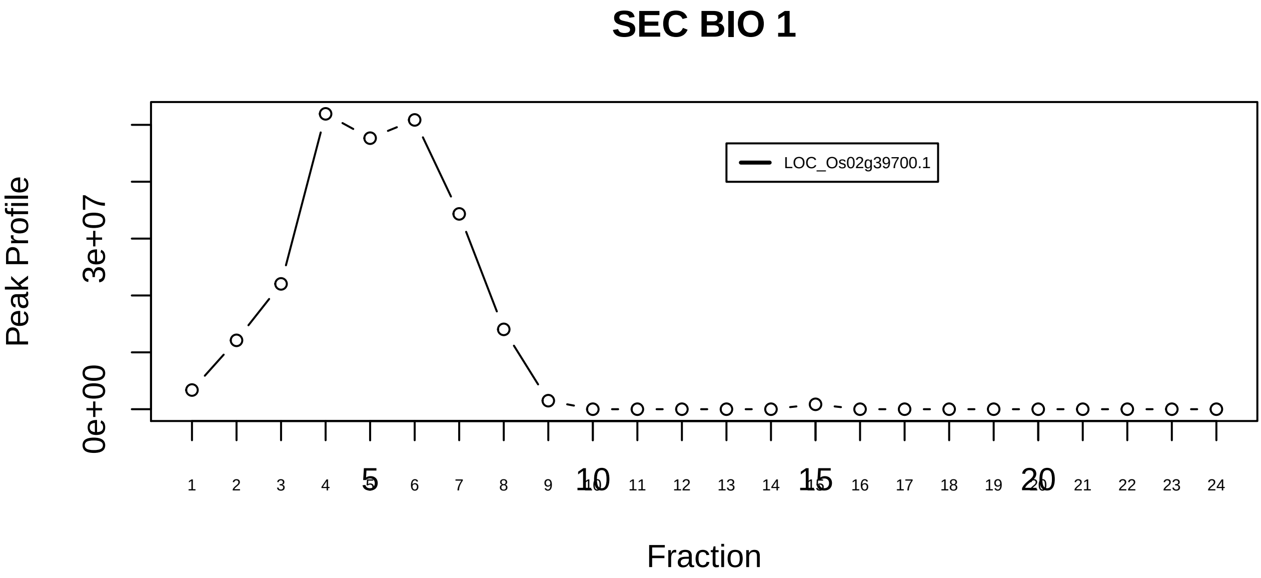

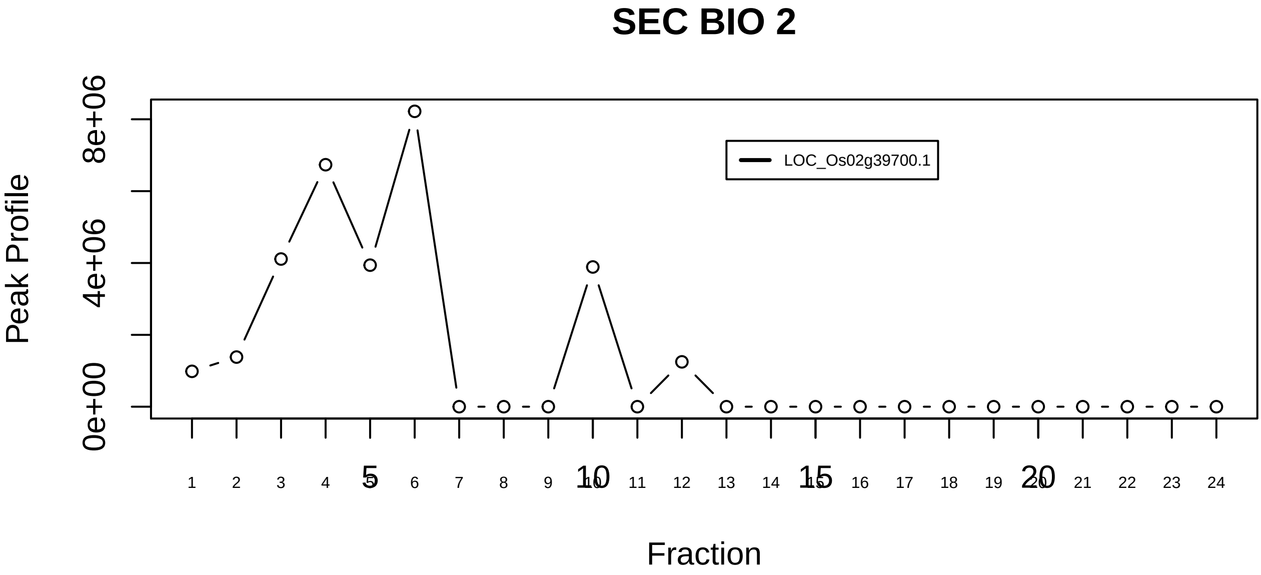

H2AX complex, isolated from cells without IR exposure|1223−−singleton_2

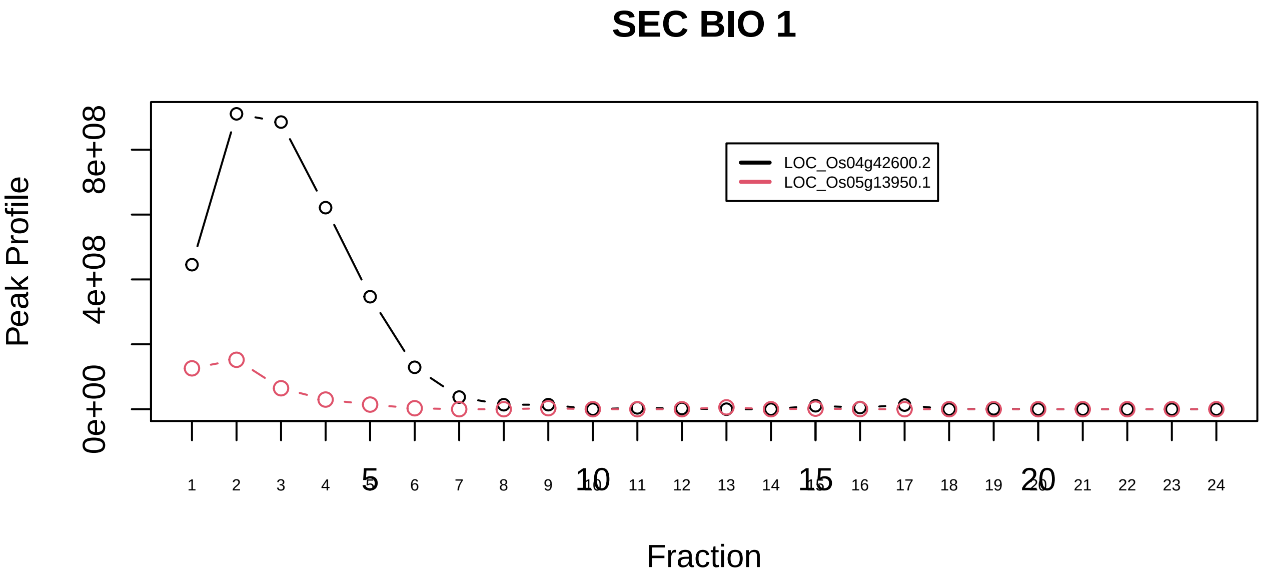

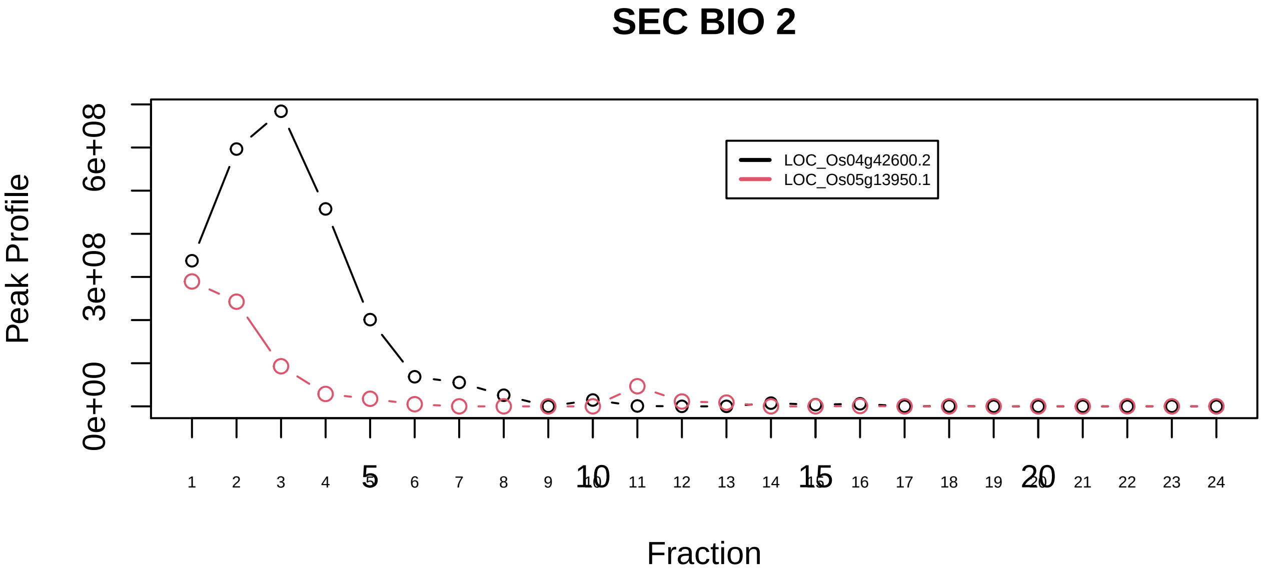

H2AX complex, isolated from cells without IR exposure|1223−−sub_3

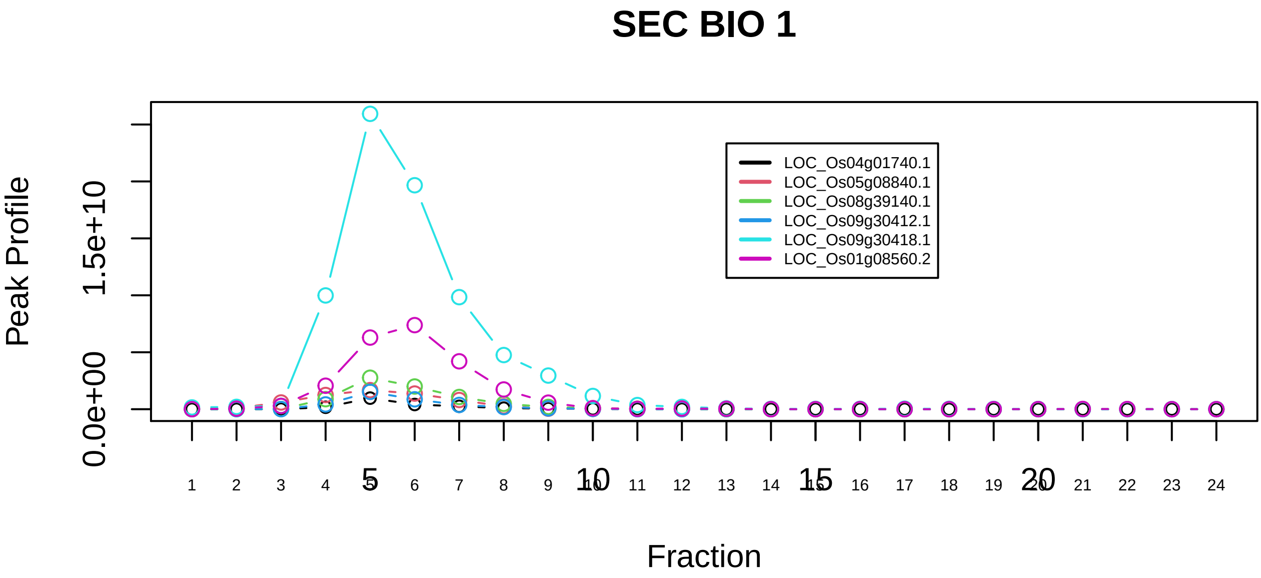

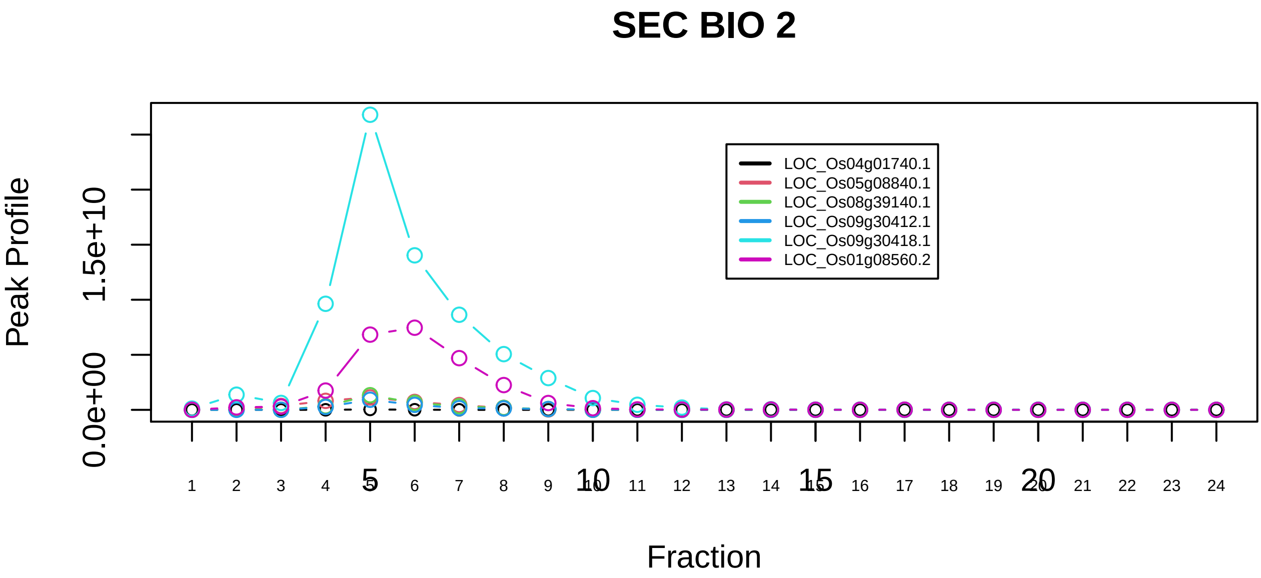

HCF−1 complex|2721−−sub_1

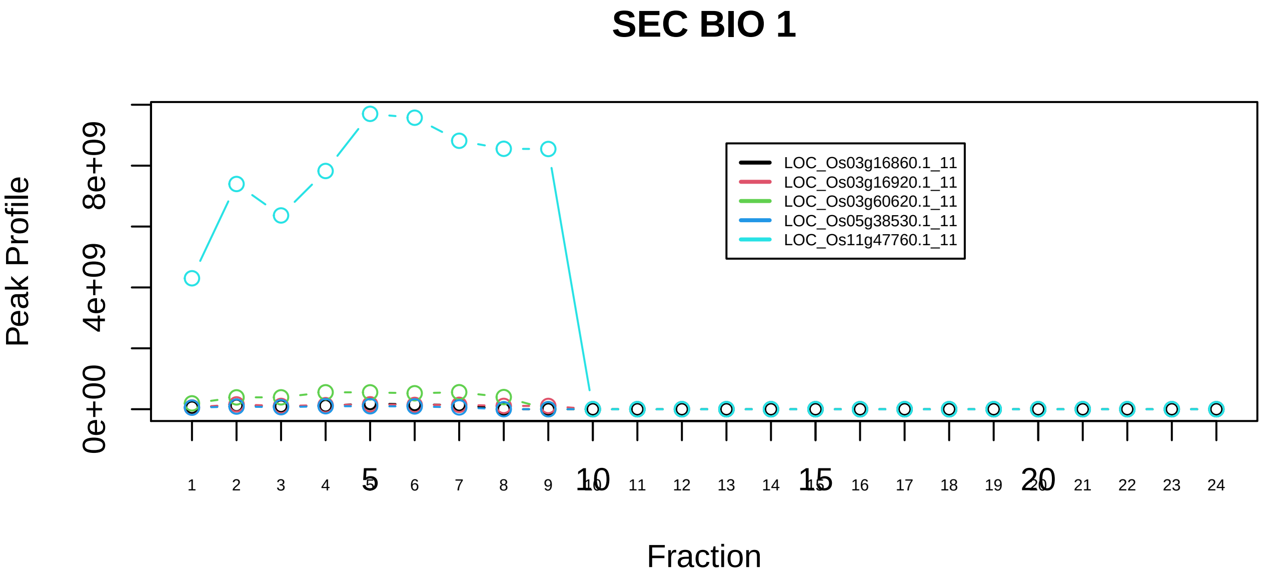

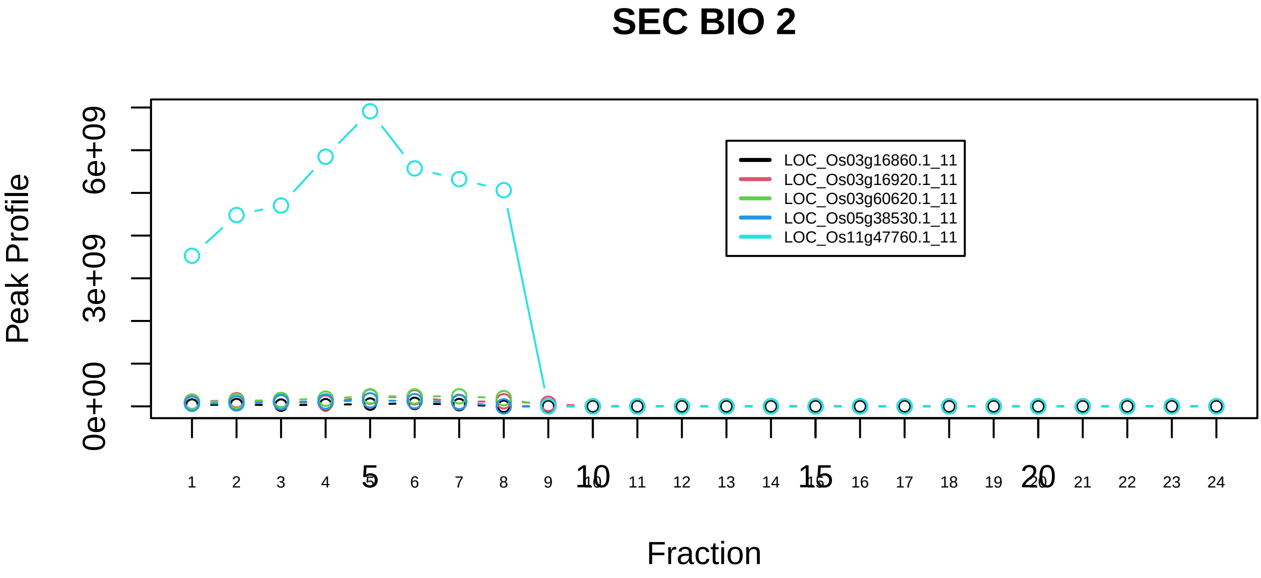

HCF−1 complex|2721−−sub_orthoparalog_1

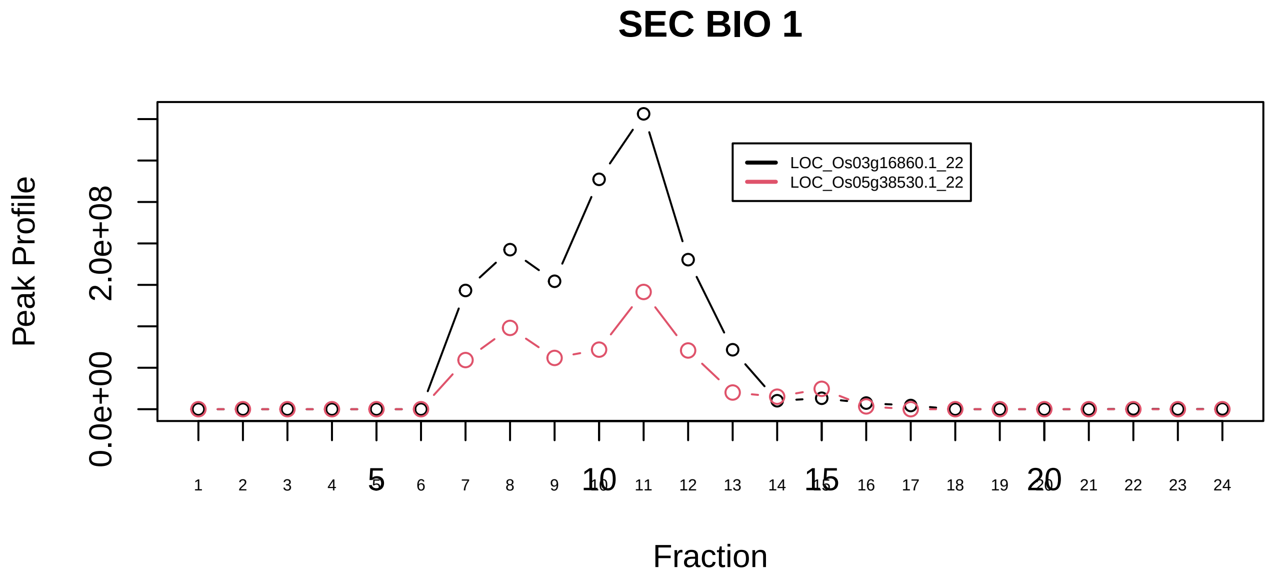

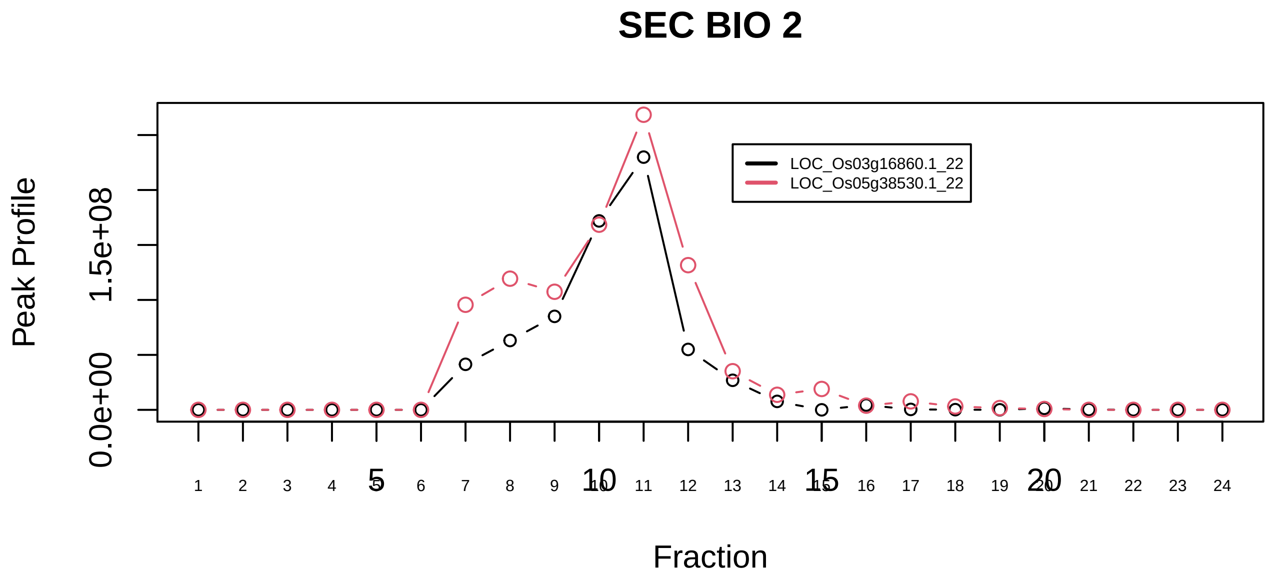

HCF−1 complex|2721−−sub_orthoparalog_2

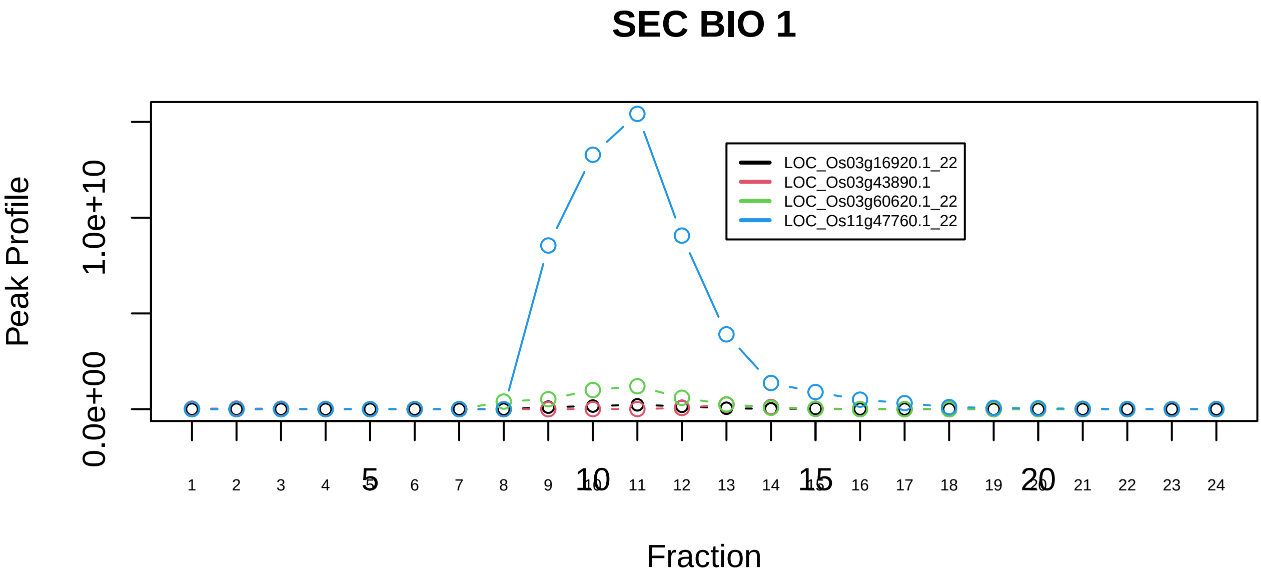

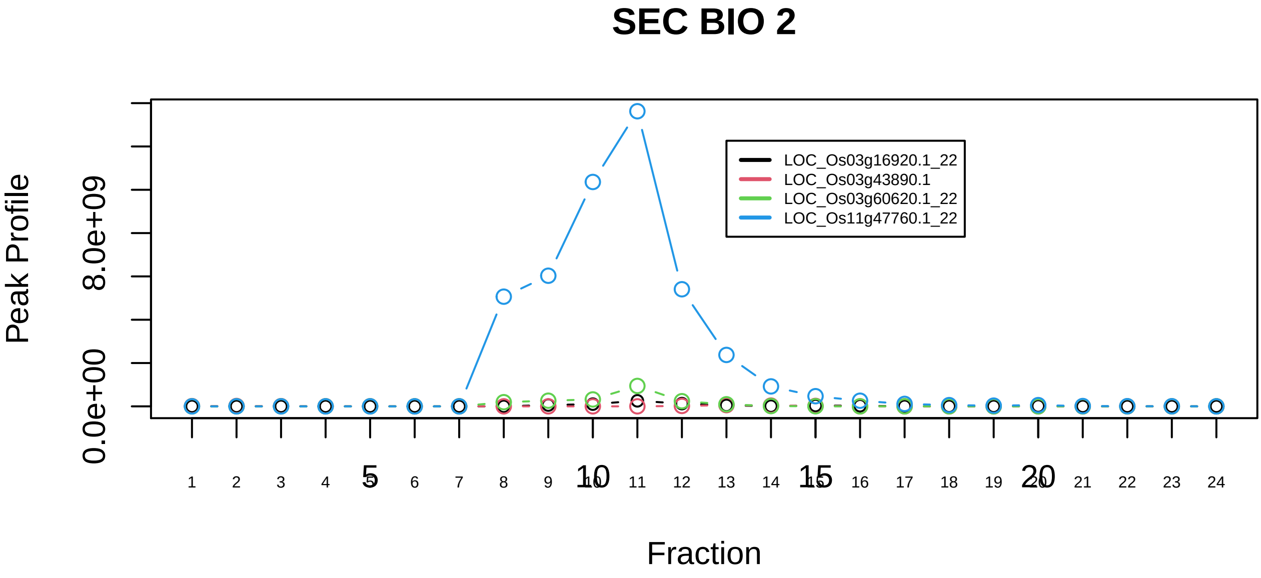

HCF−1 complex|2721−−sub_2

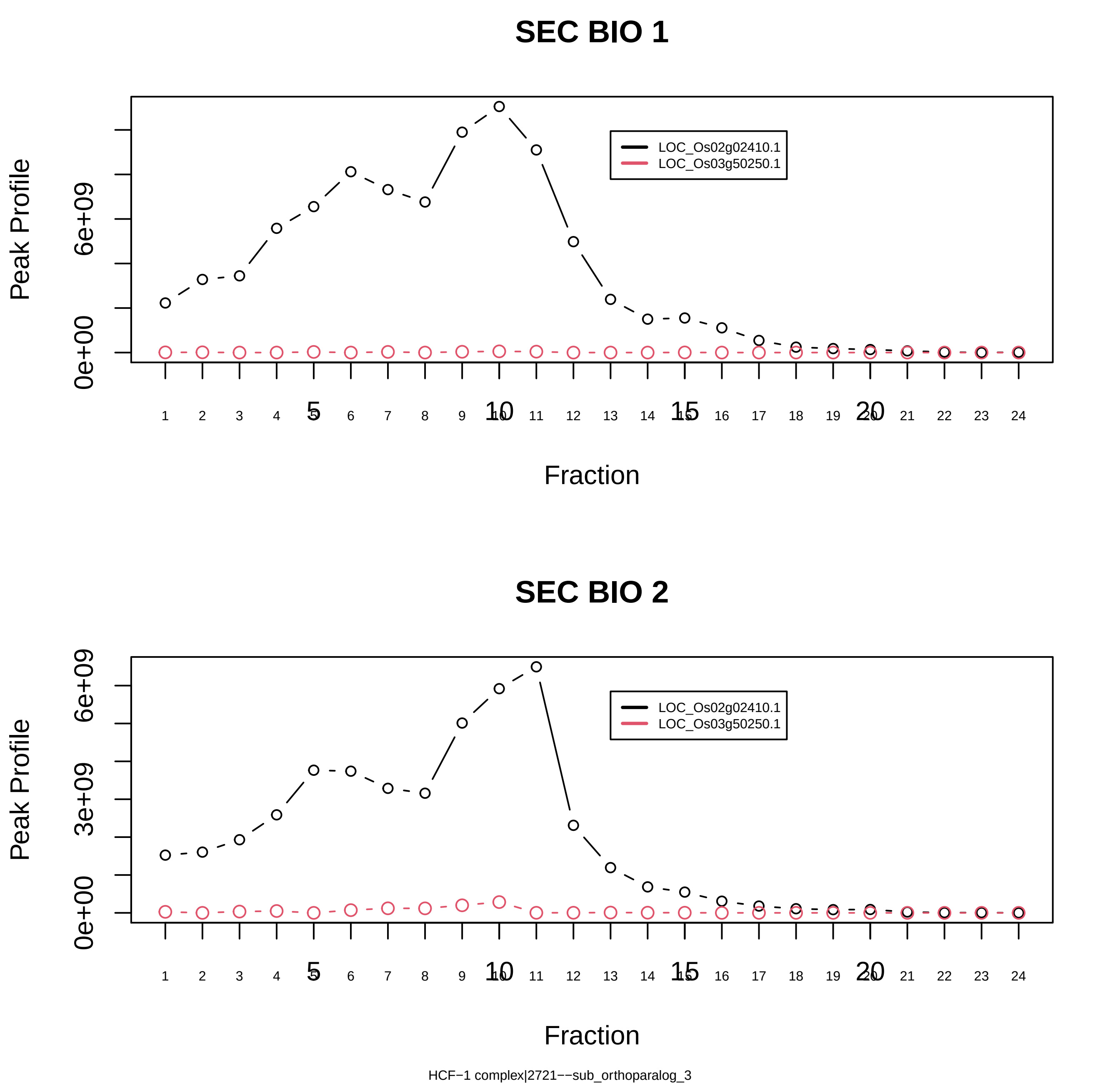
SEC BIO 1
Peak Profile
0e+00
6e+09
LOC_Os02g02410.1
LOC_Os03g50250.1
Fraction
SEC BIO 2
0e+00
3e+09
6e+09
Fraction
HCF−1 complex|2721−−sub_orthoparalog_3

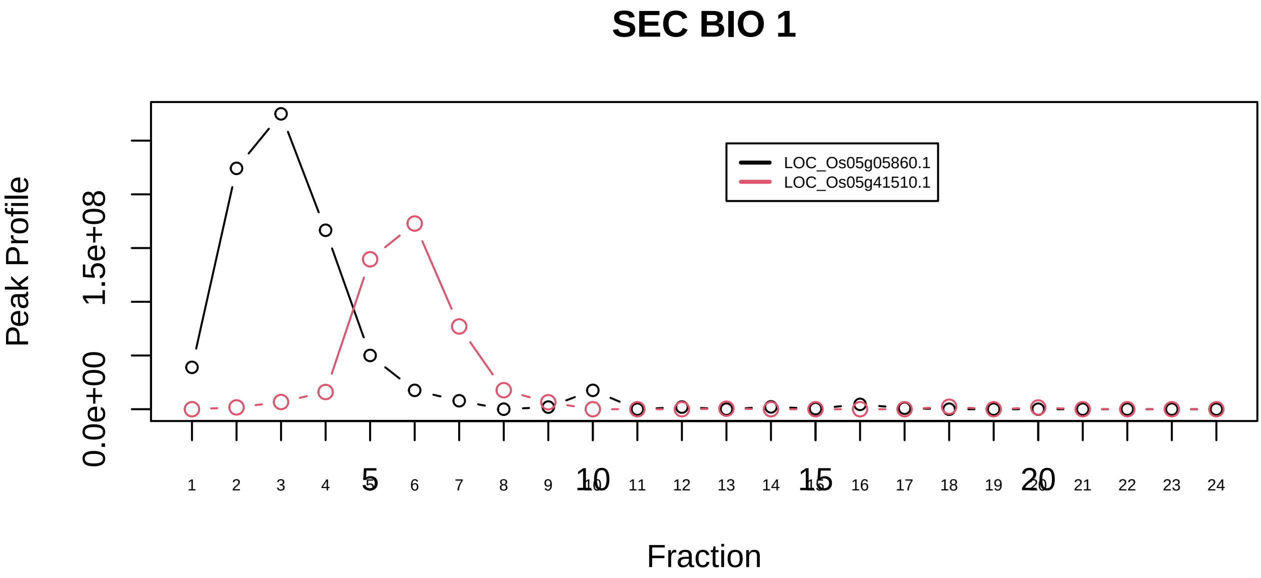

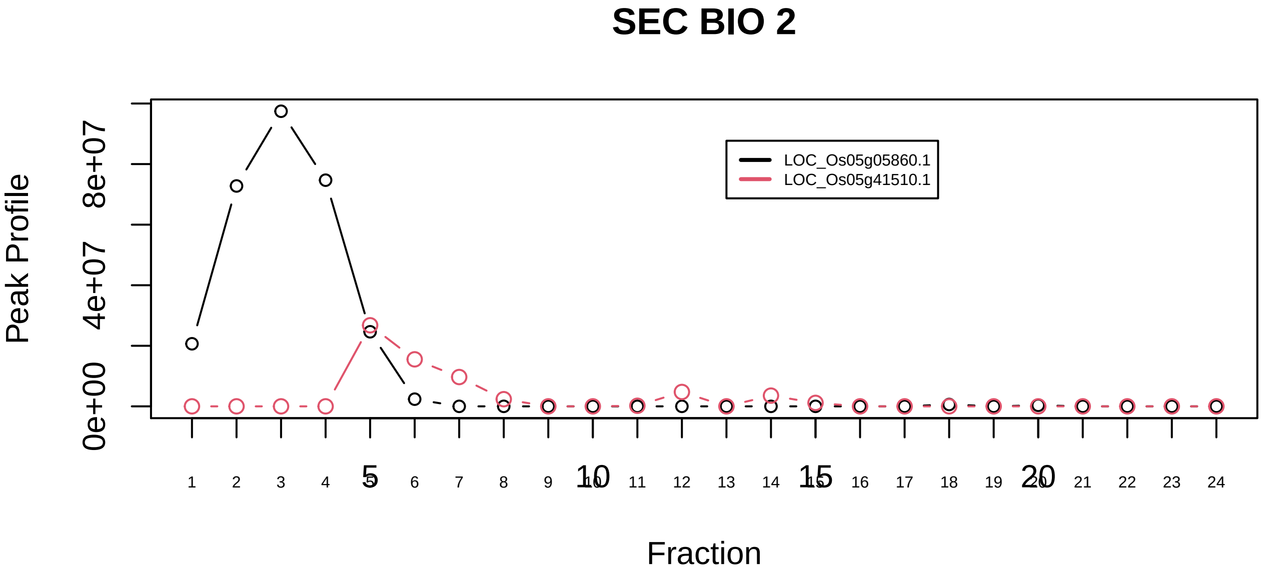

HES1 promoter−Notch enhancer complex|2639−−only_2_prots_before_clustering

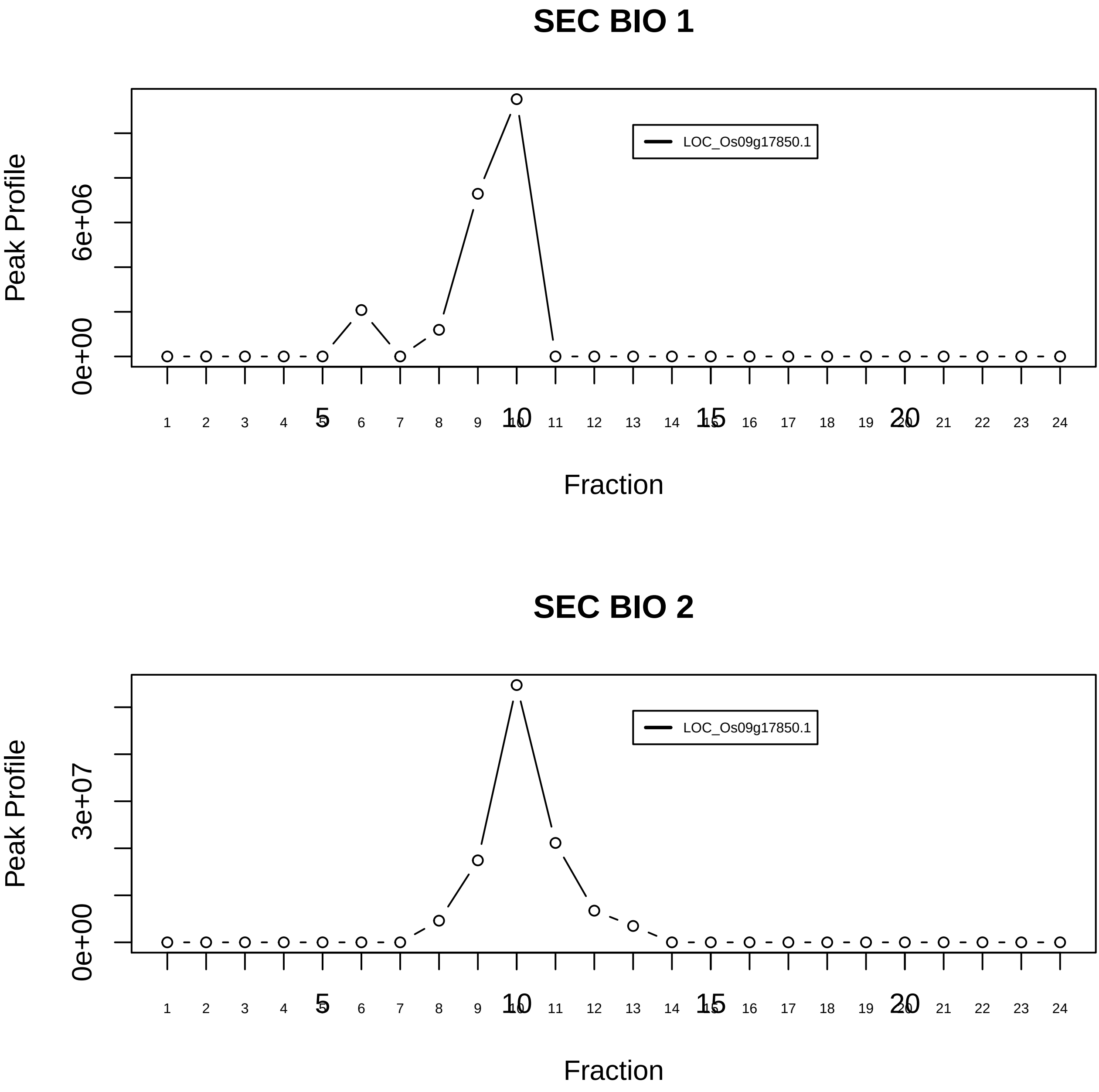

Histone H3.3 complex|1150−−singleton_1

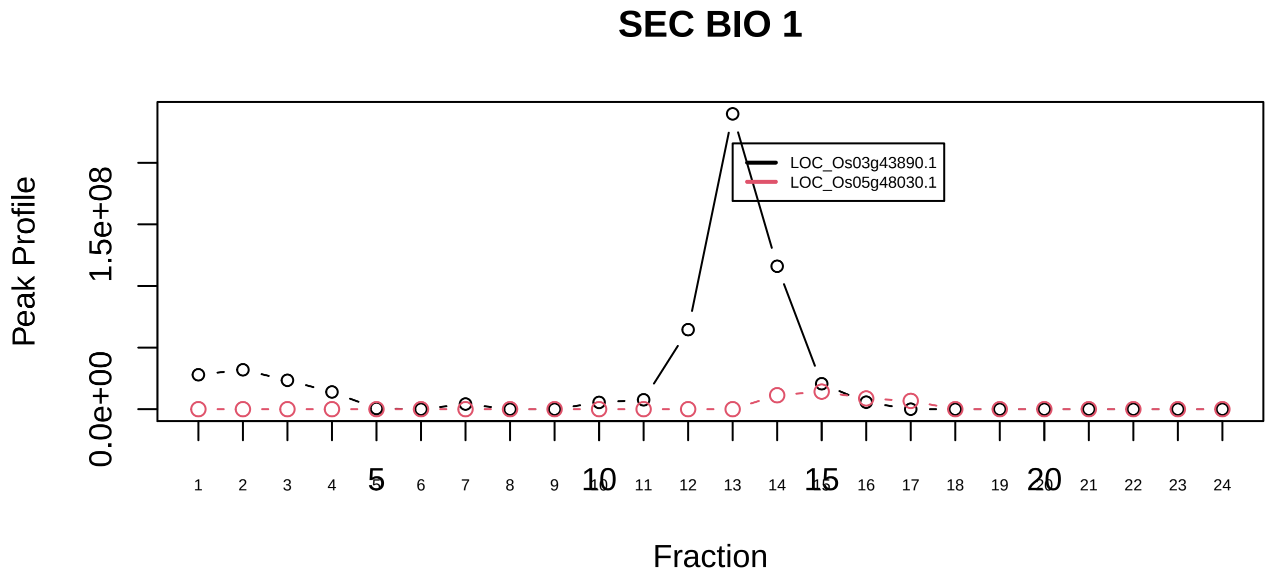

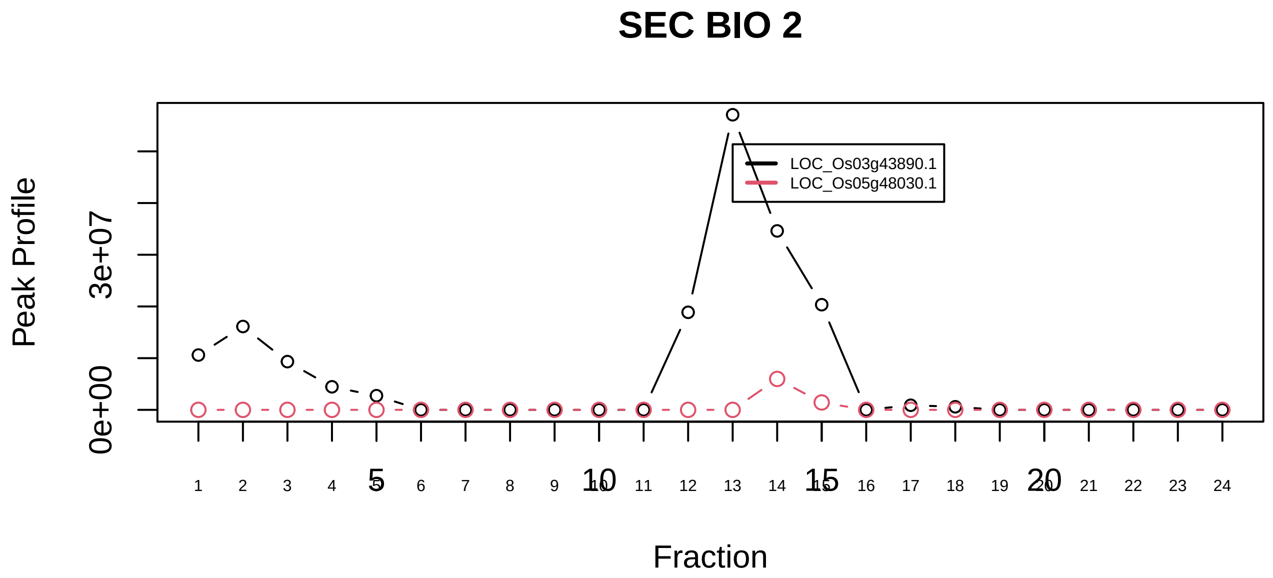

Histone H3.3 complex|1150−−sub_1

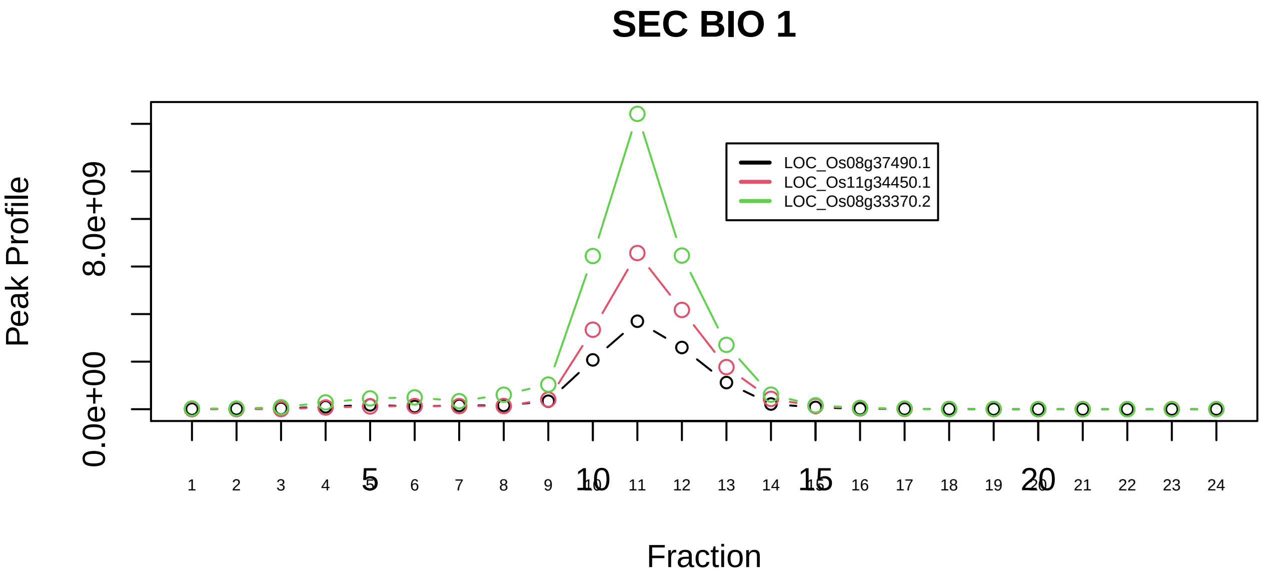

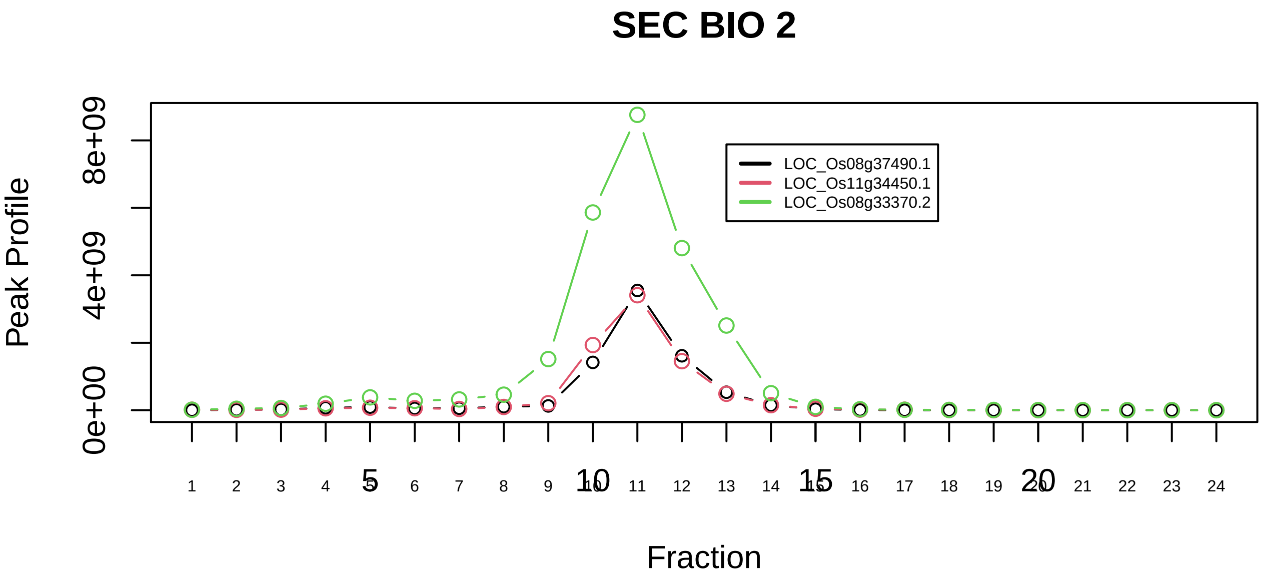

HSF1−YWHAE complex|2145−−sub_orthoparalog_1

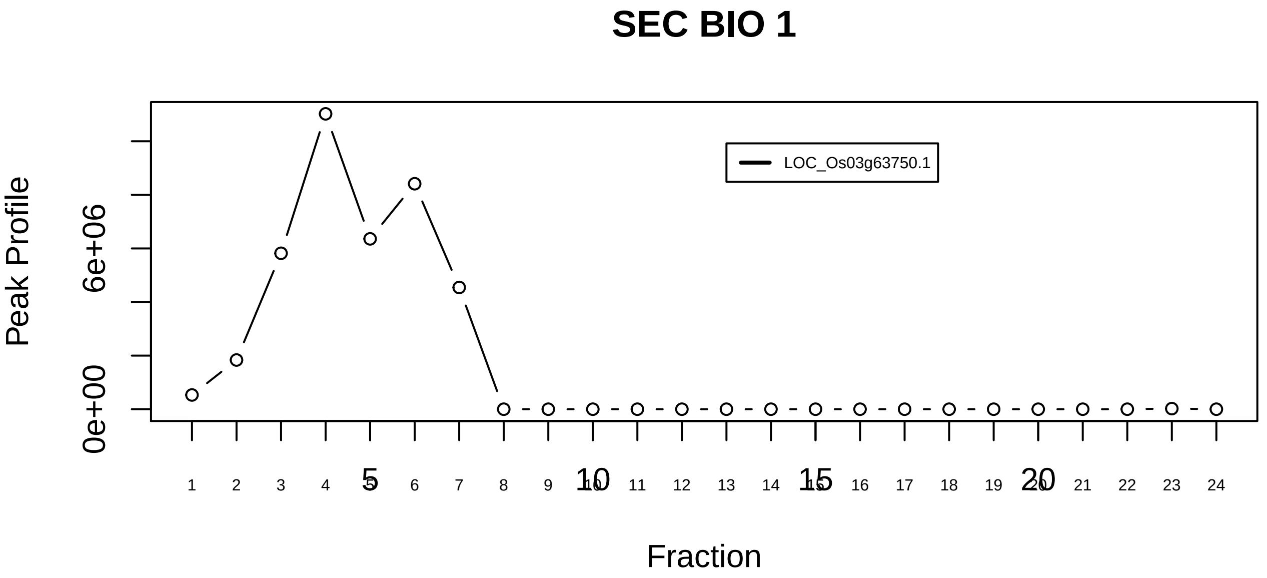

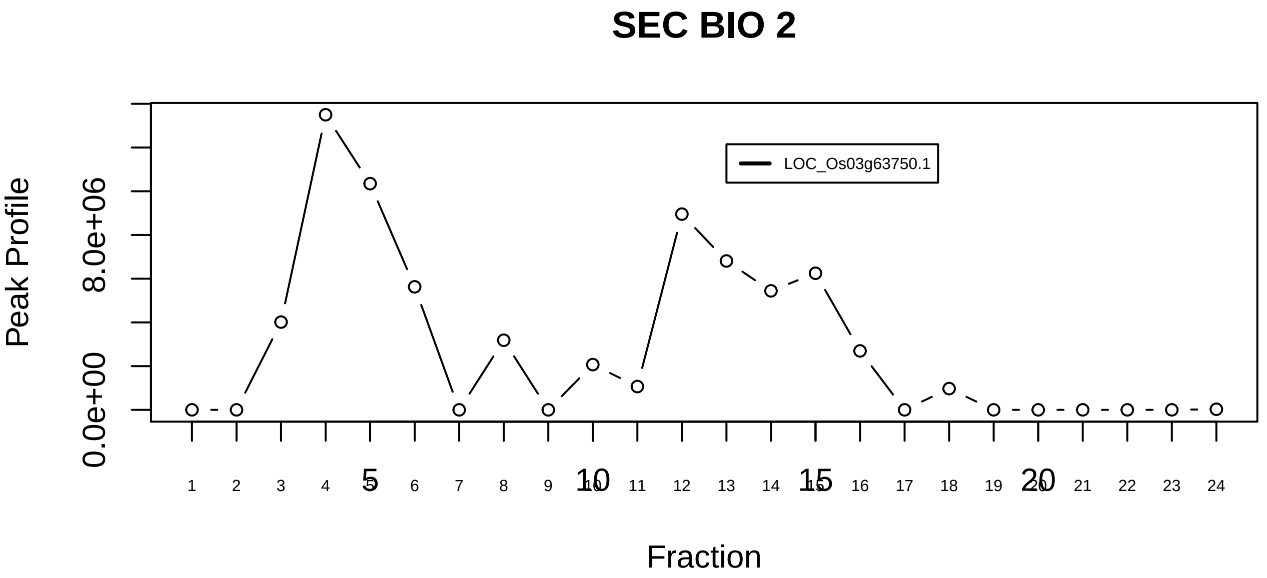

HSF1−YWHAE complex|2145−−singleton_1

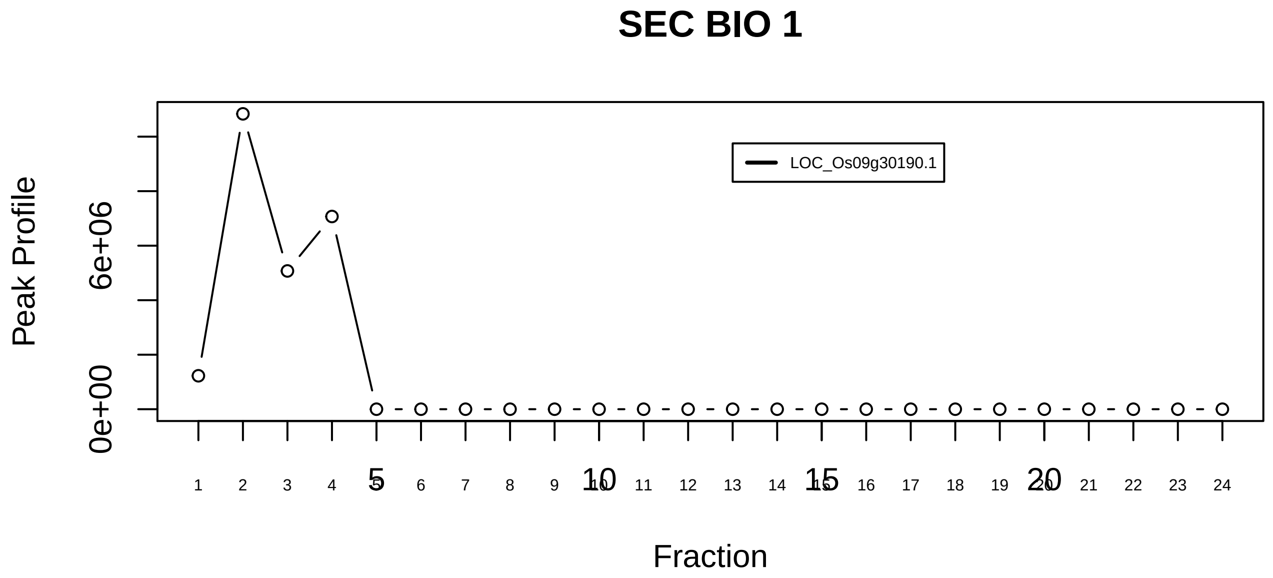

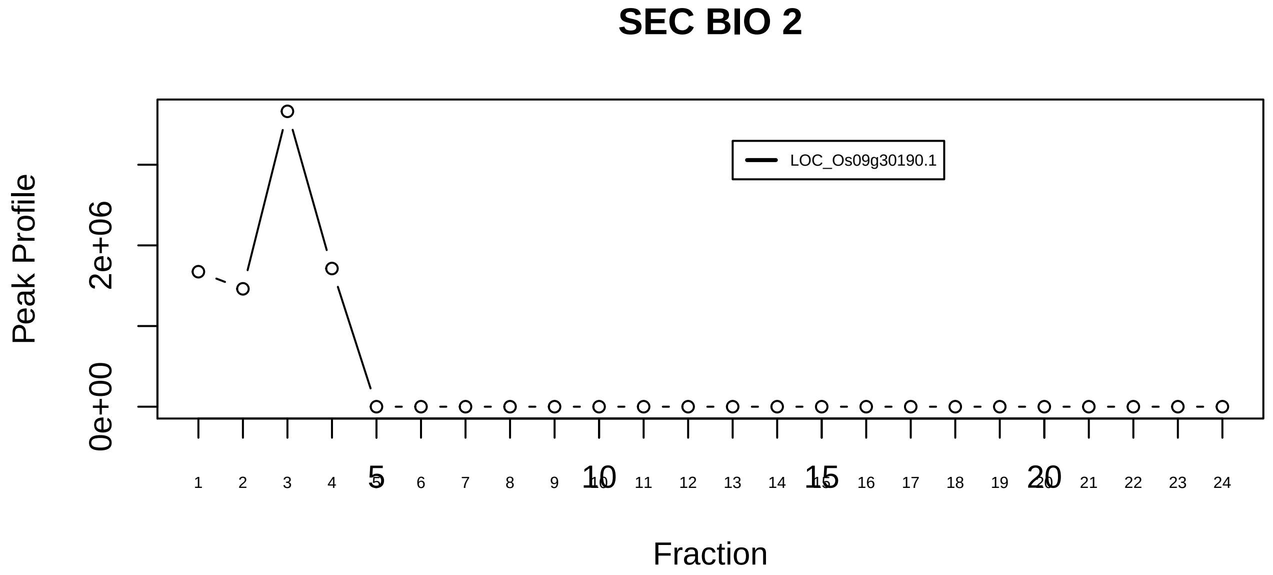

LRRK2−CHIP−HSP90 complex|5980−−singleton_1

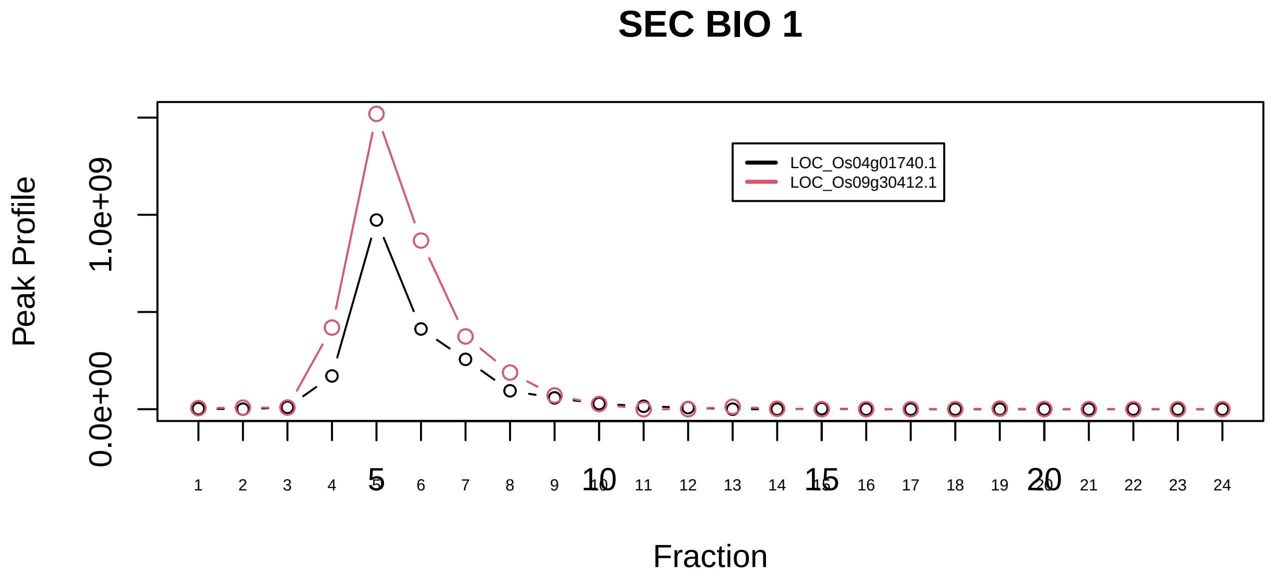

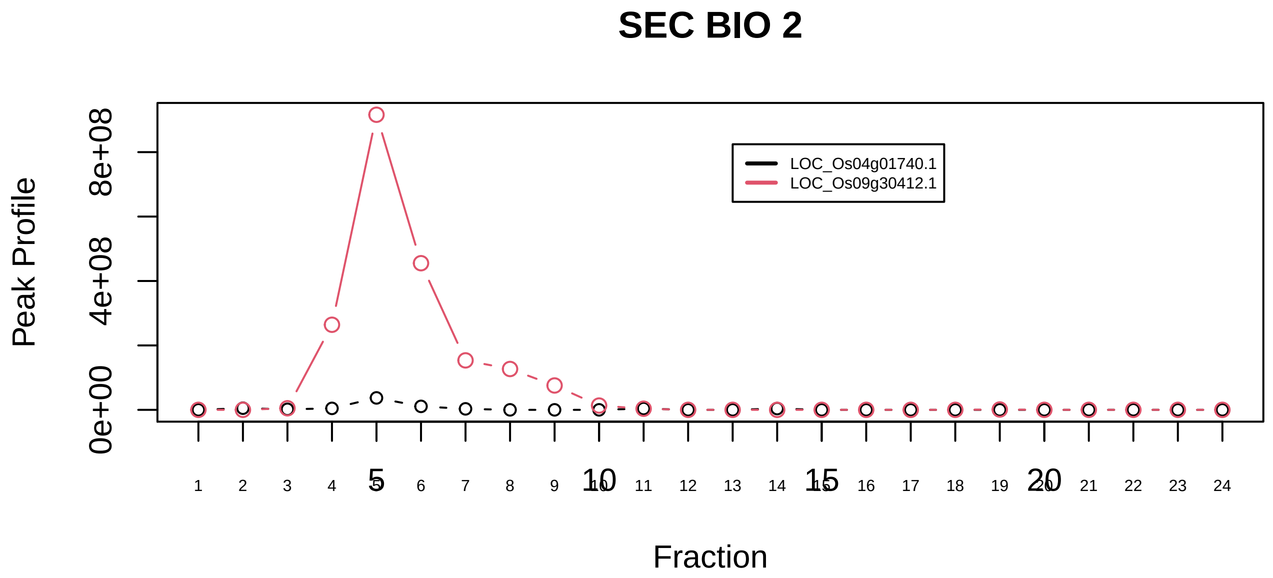

LRRK2−CHIP−HSP90 complex|5980−−sub_orthoparalog_1

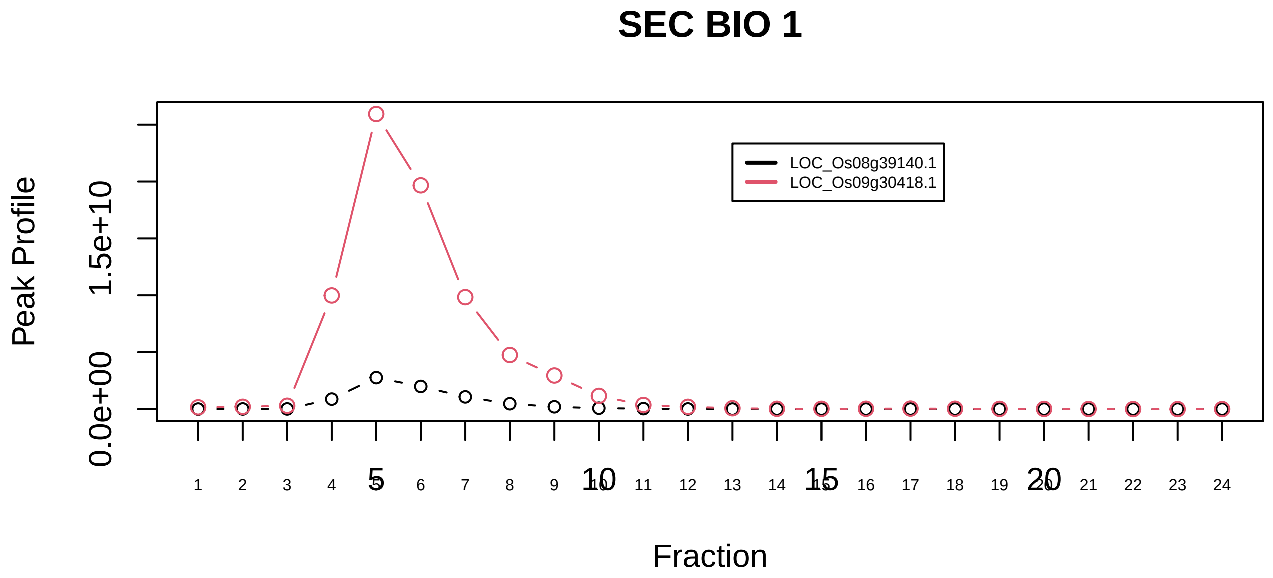

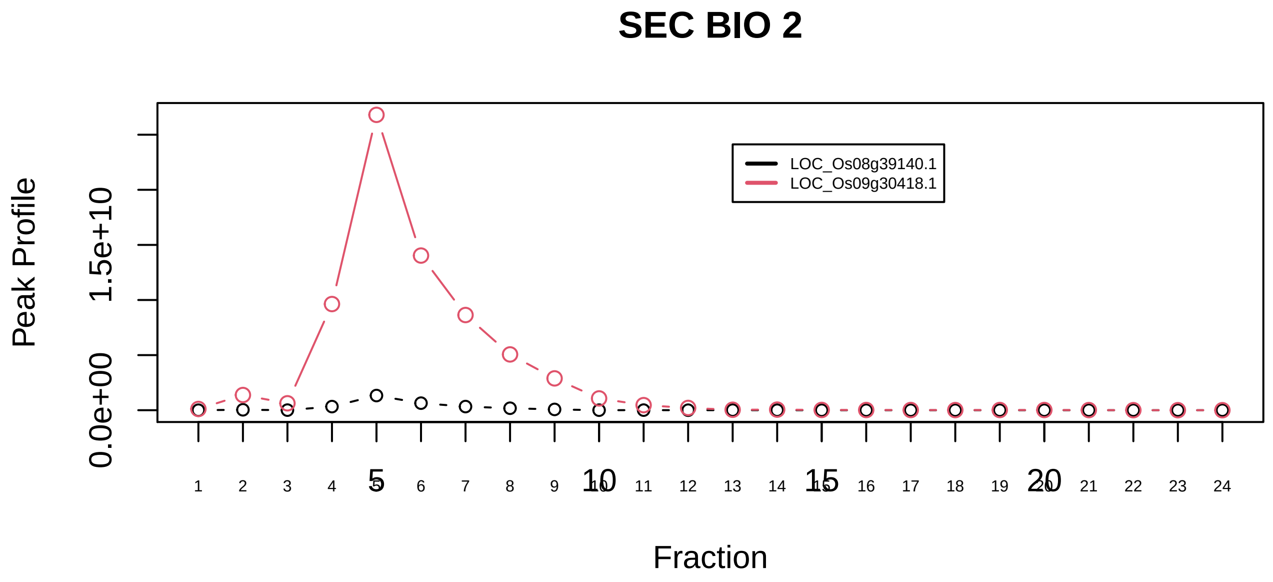

LRRK2−CHIP−HSP90 complex|5980−−sub_orthoparalog_2

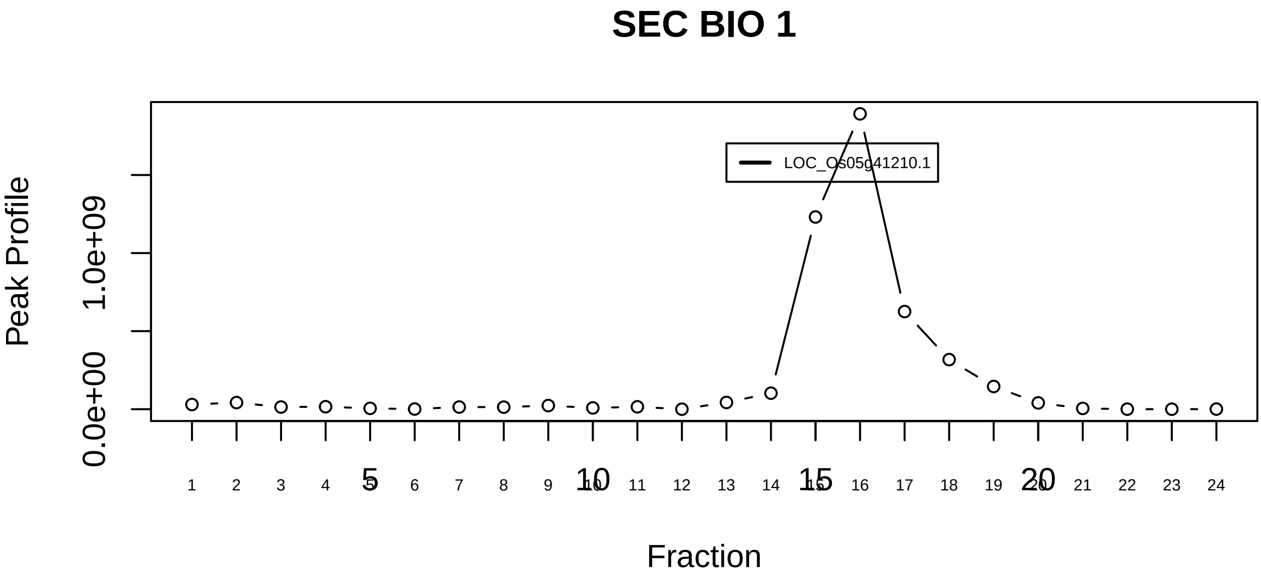

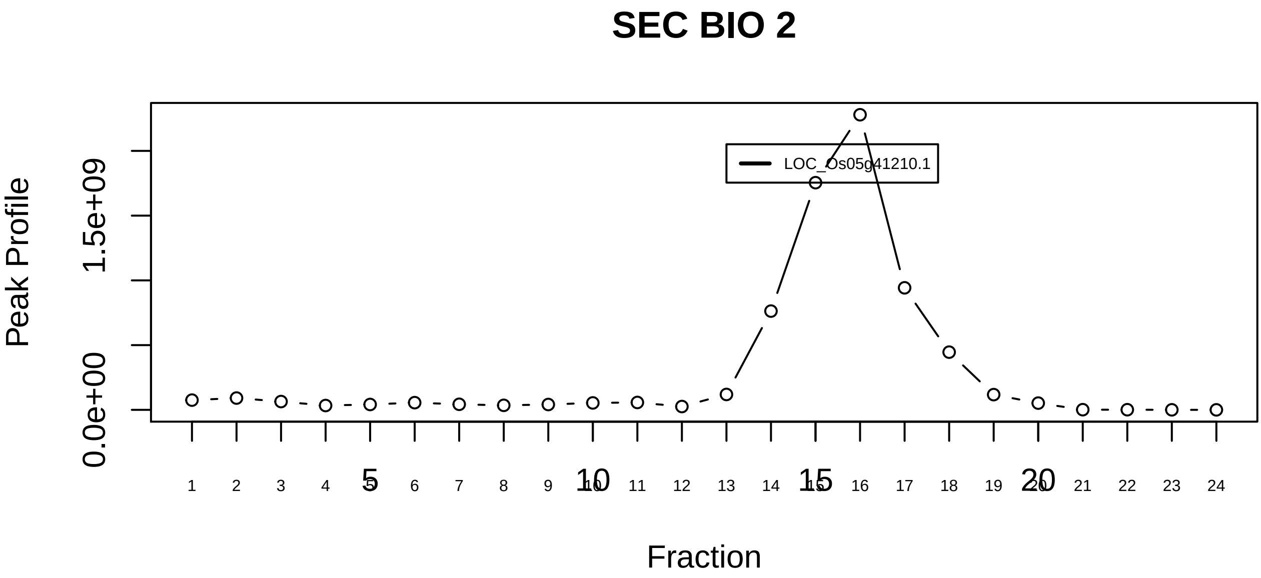

HSP90−FKBP38−CAM−Ca(2+) complex|4158−−singleton_1

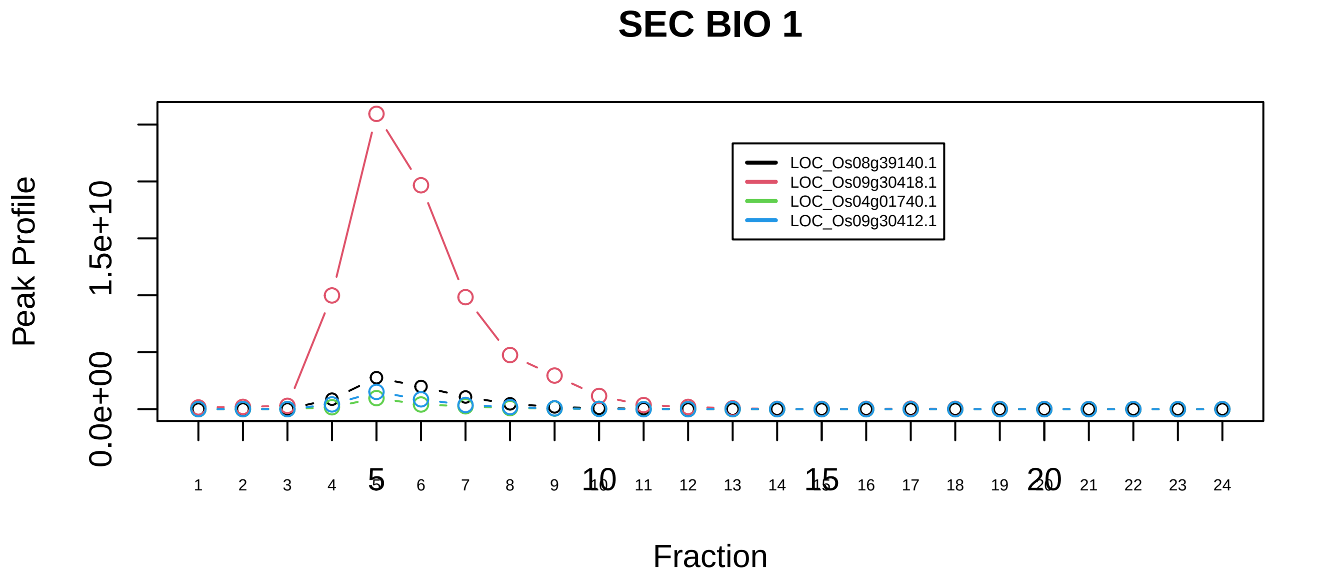

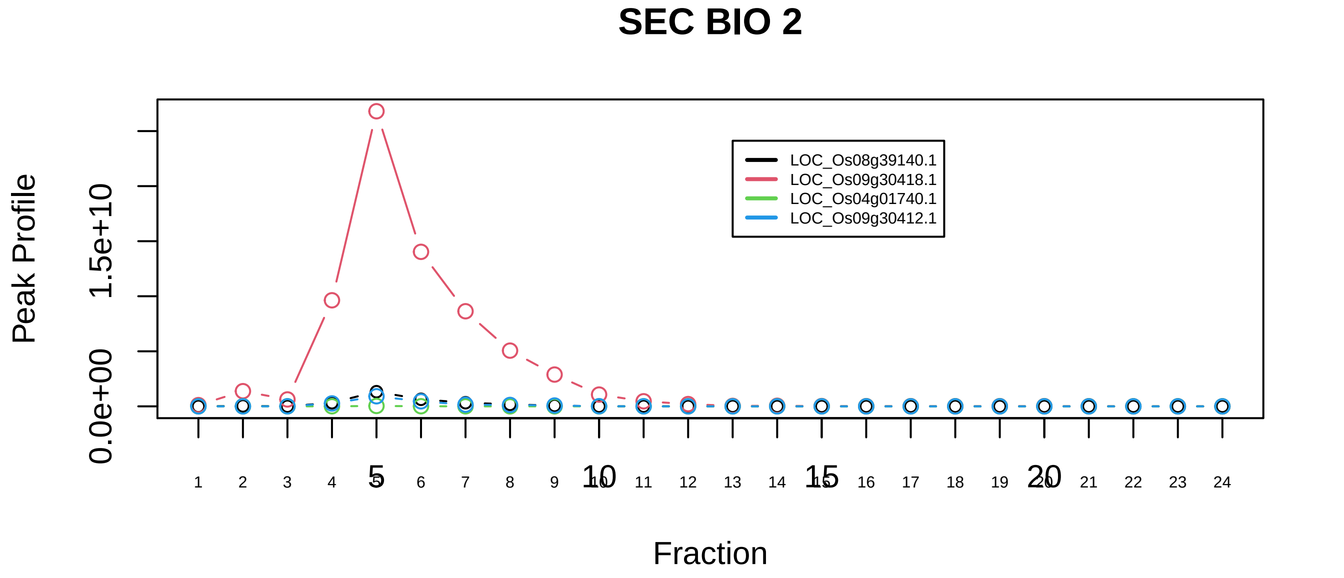

HSP90−FKBP38−CAM−Ca(2+) complex|4158−−sub_orthoparalog_1

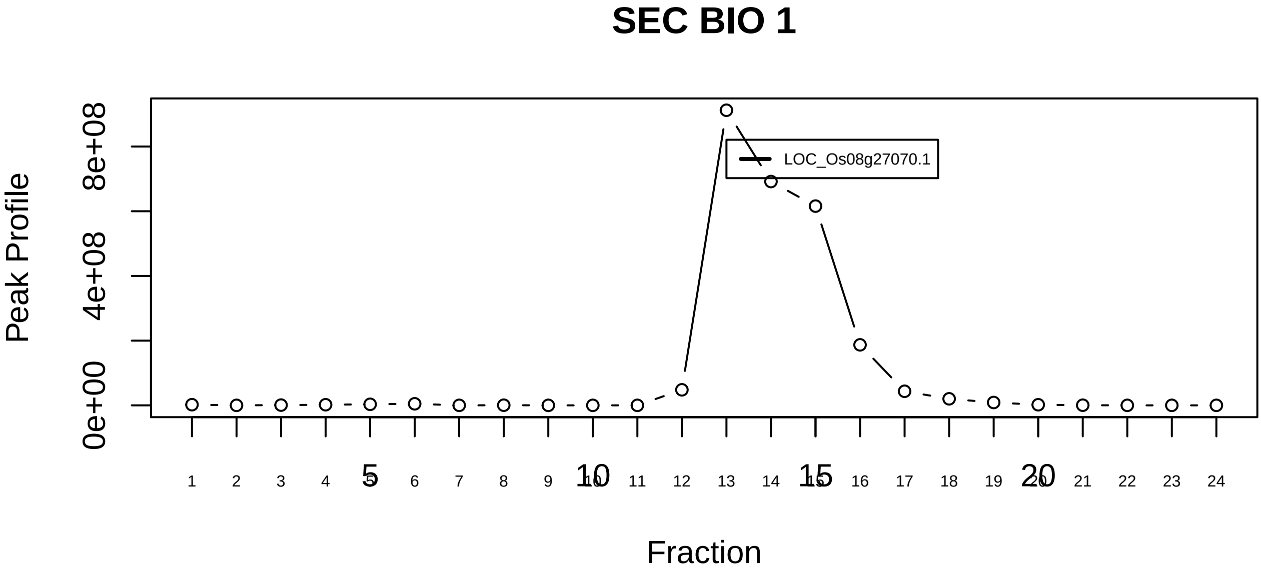

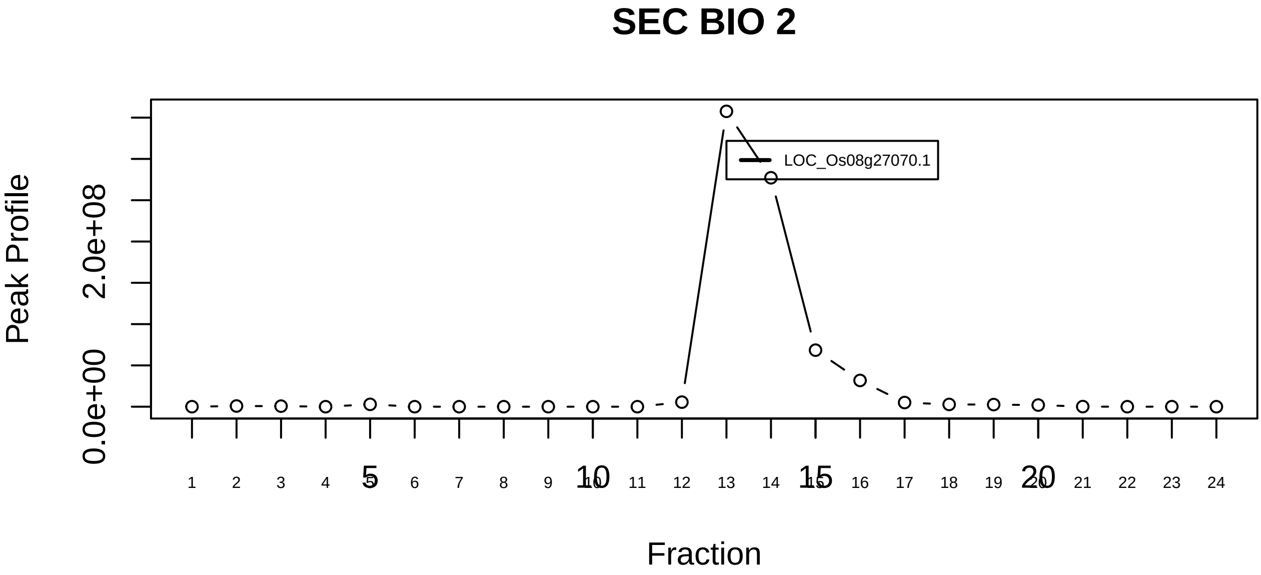

Hsp90−p23 complex|6591−−singleton_1

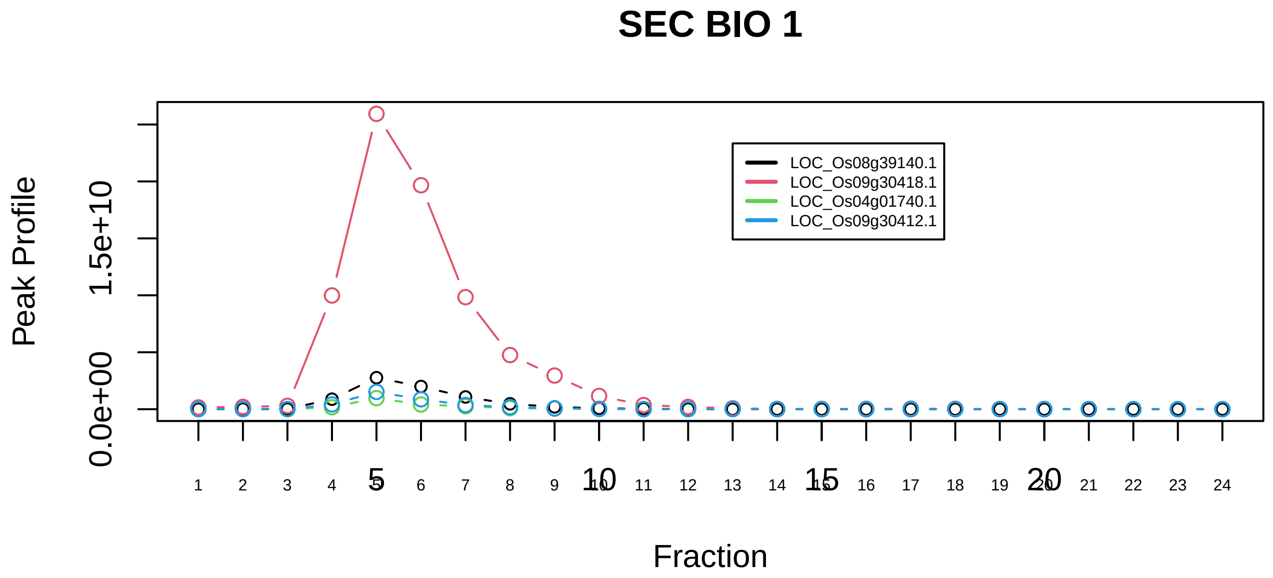

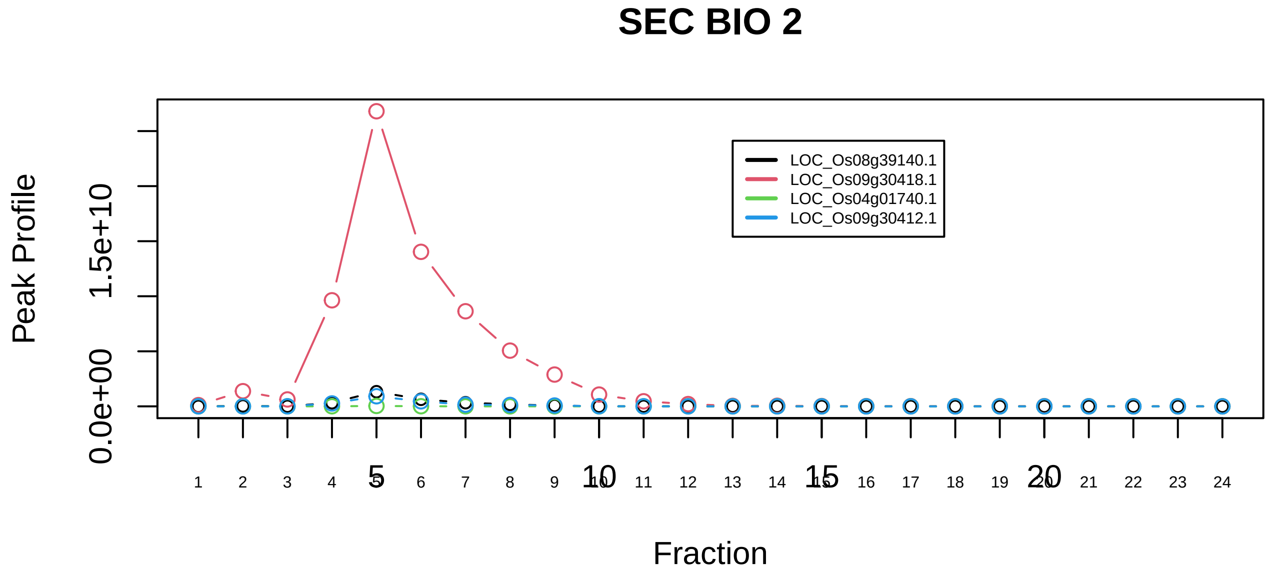

Hsp90−p23 complex|6591−−sub_orthoparalog_1

### SEC BIO 1

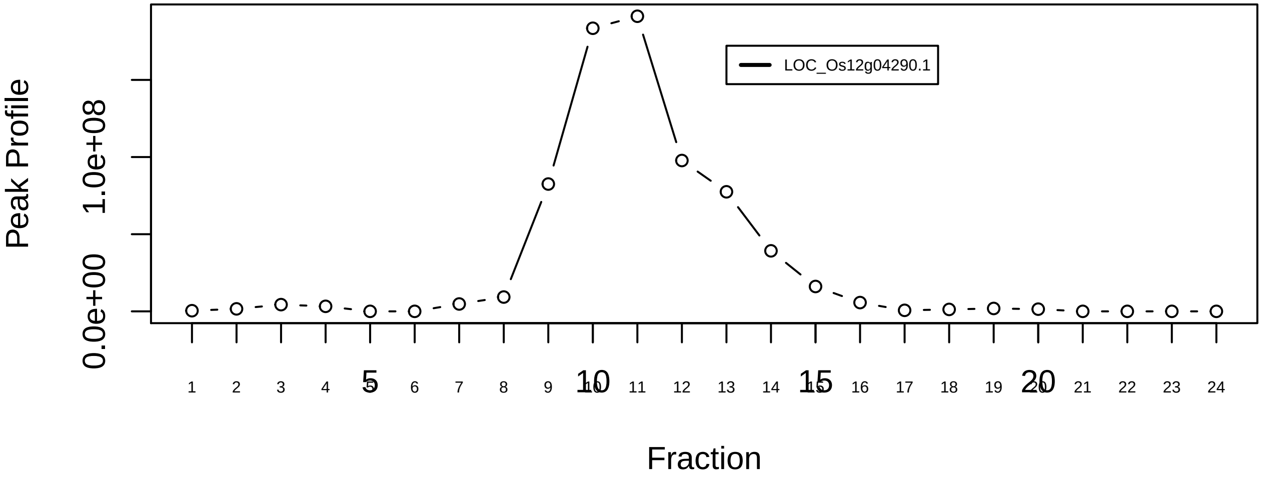

### SEC BIO 2

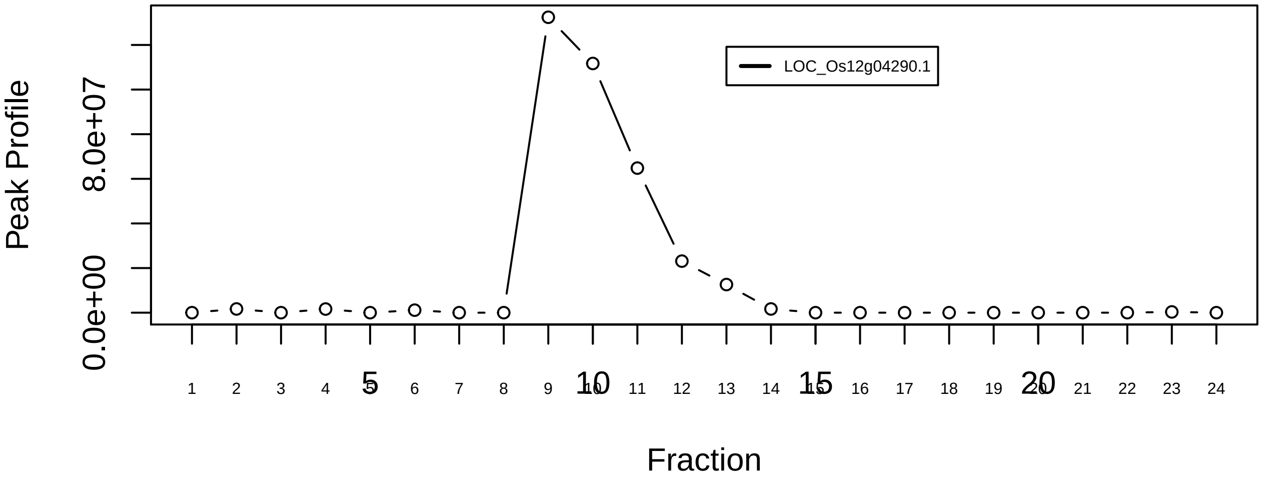

IGBP1−MID1−PPP2CA complex|7251−−singleton_1

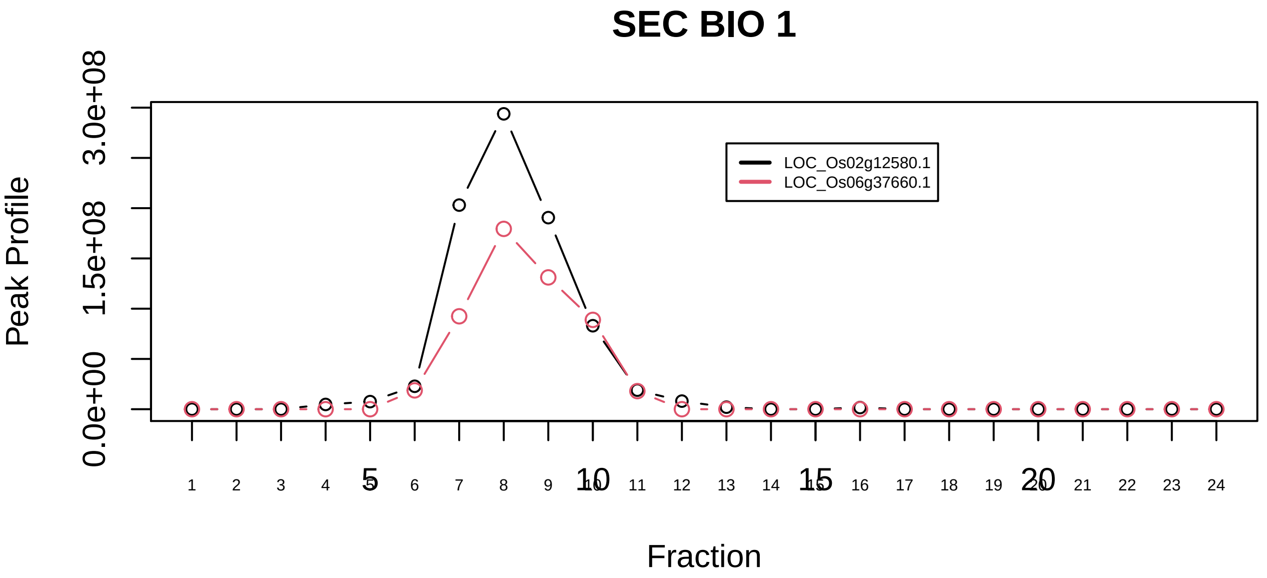

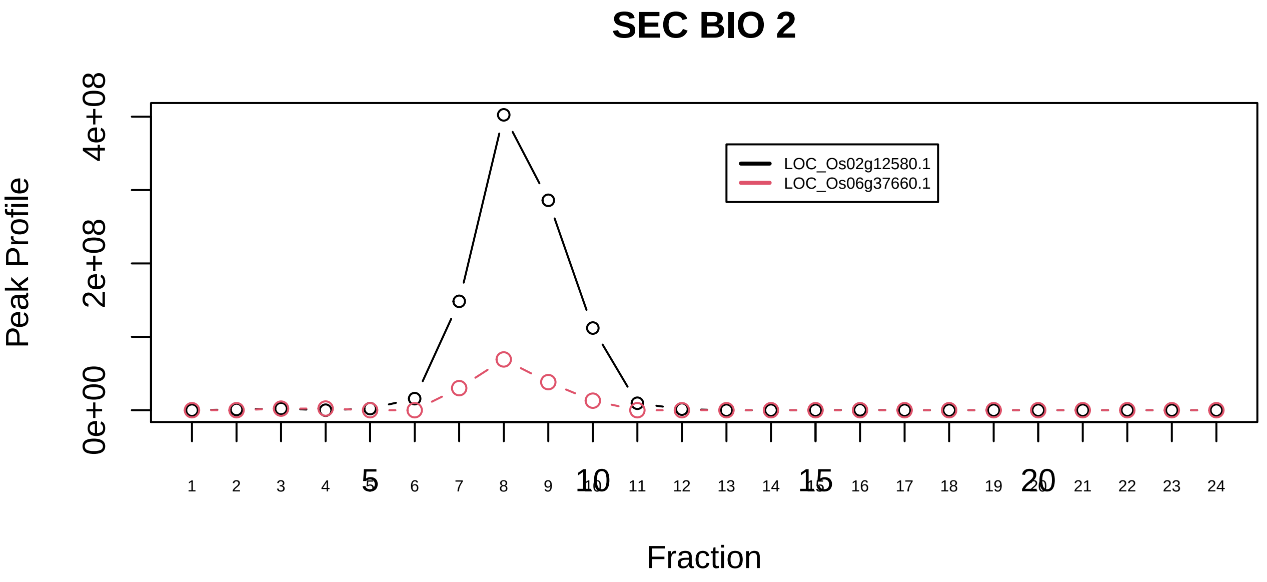

IGBP1−MID1−PPP2CA complex|7251−−sub_orthoparalog_1

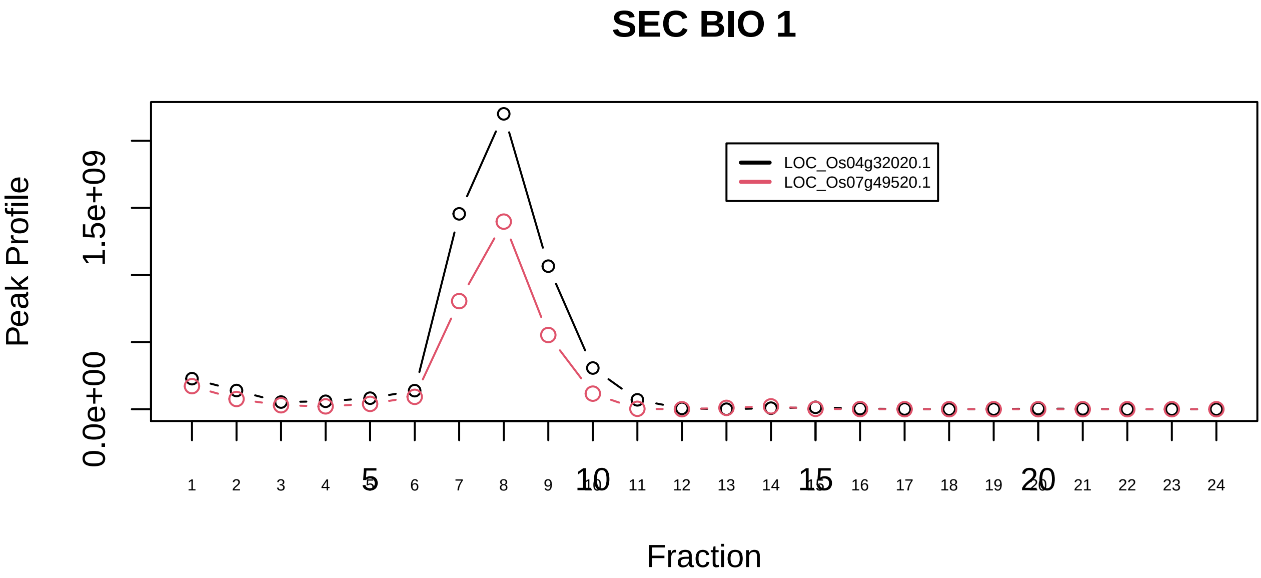

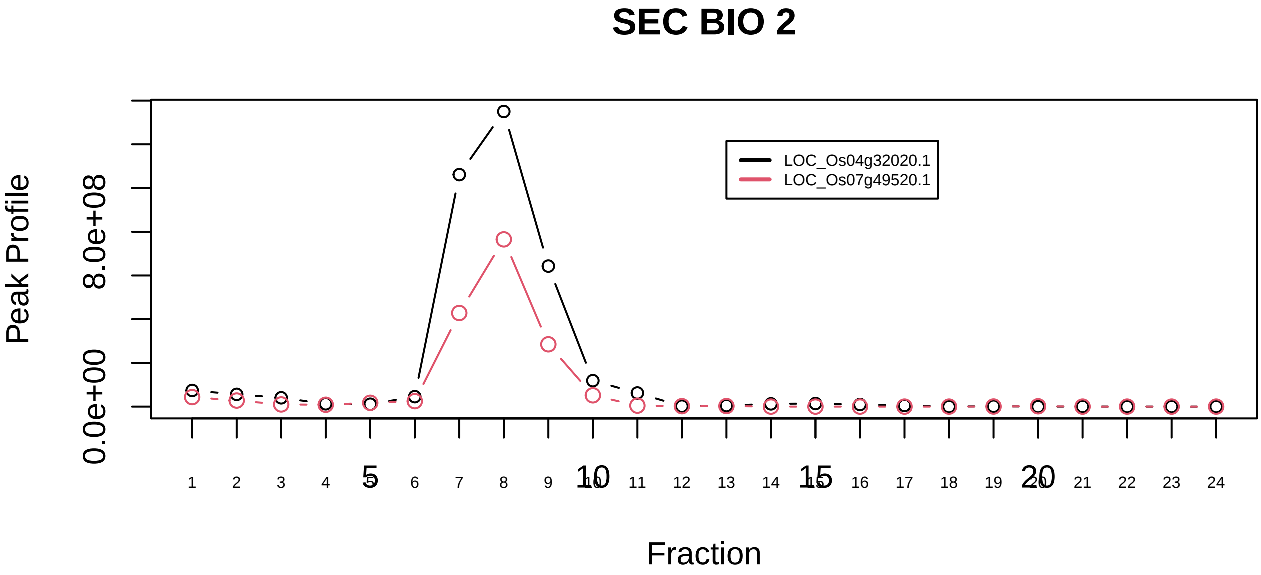

KAT2A−Oxoglutarate dehydrogenase complex|7267−−sub_orthoparalog_1

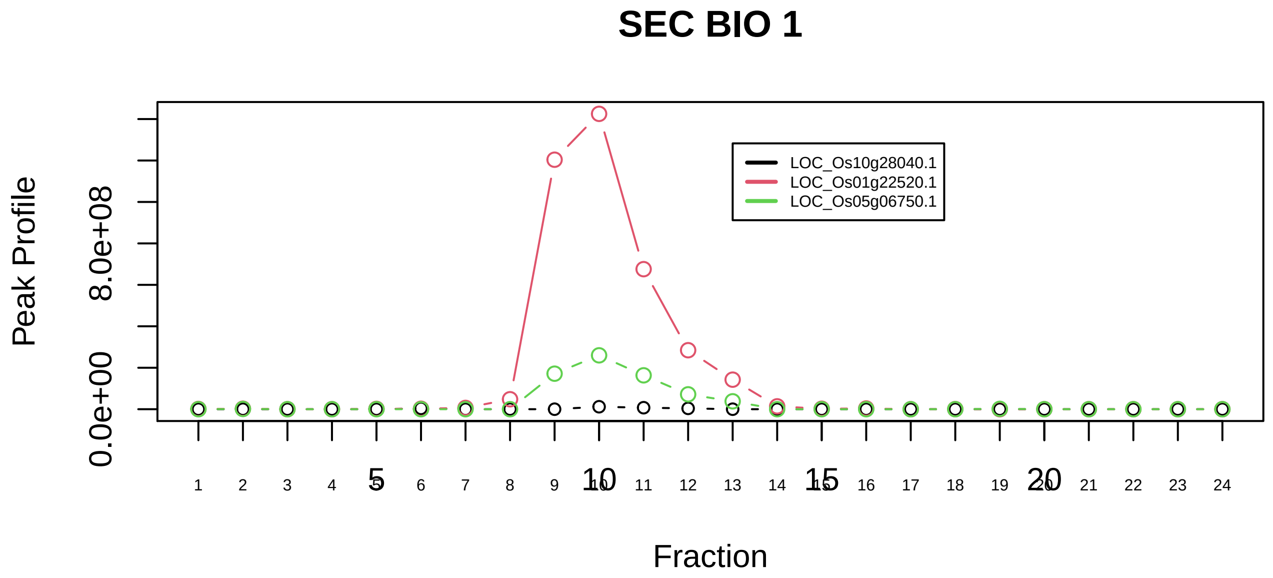

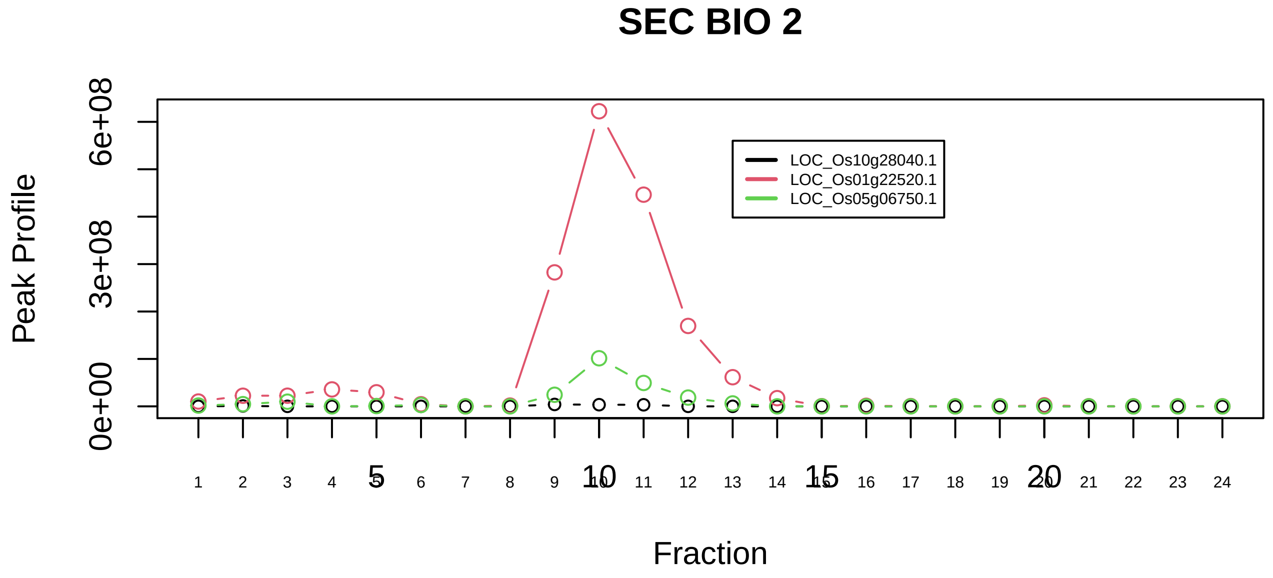

KAT2A−Oxoglutarate dehydrogenase complex|7267−−sub_1

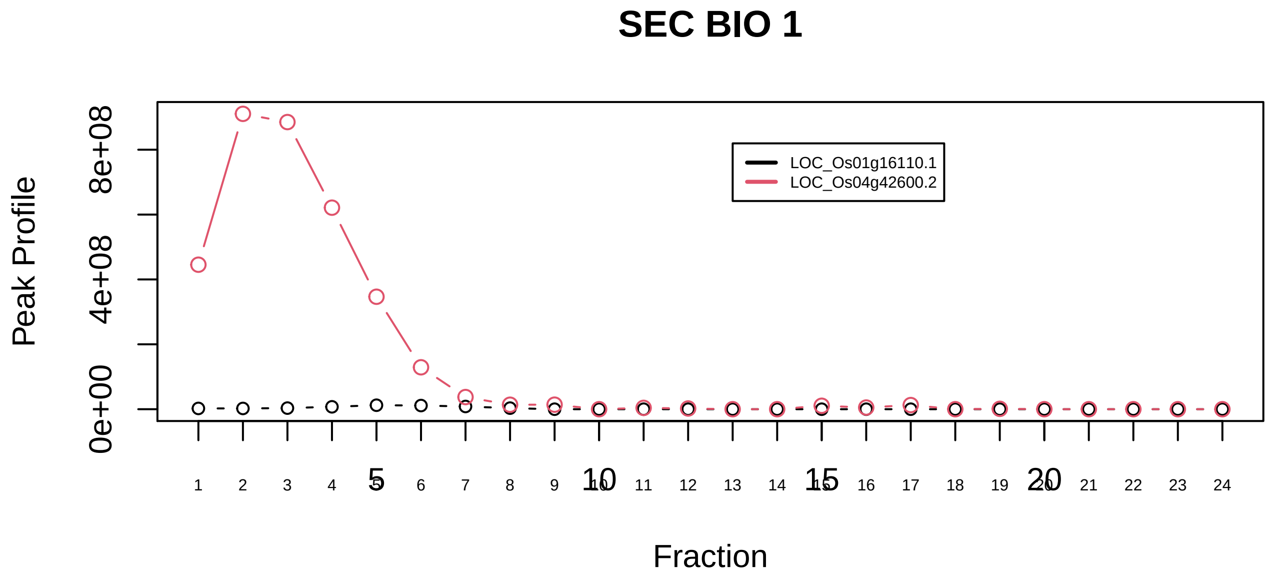

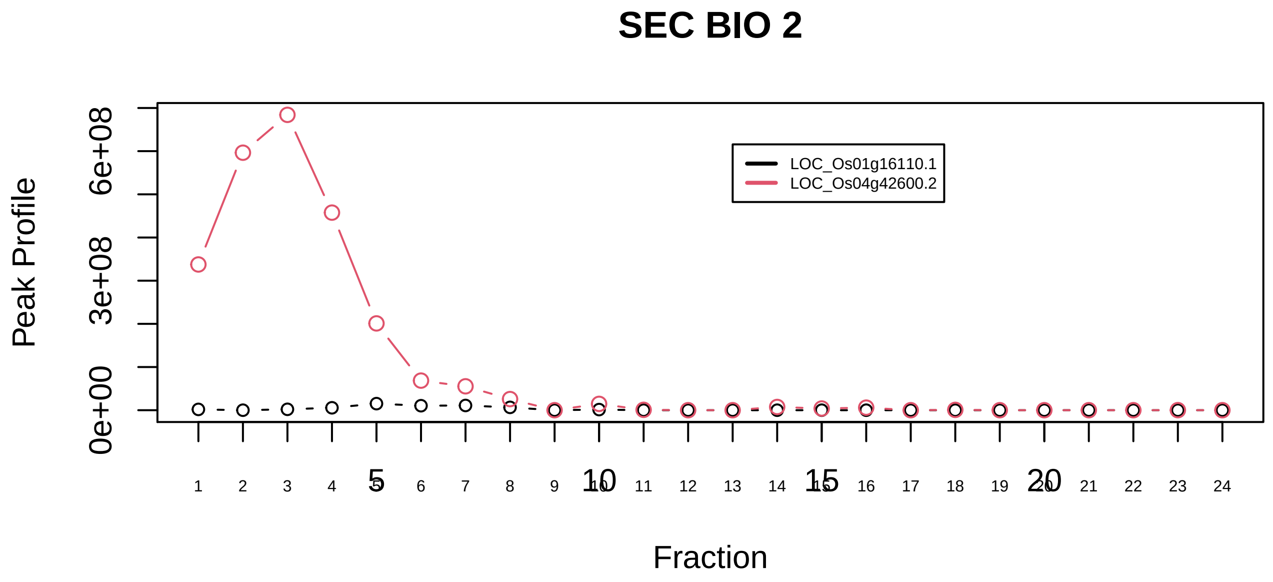

LARP1−PABPC1−RYDEN complex|7563−−only_2_prots_before_clustering

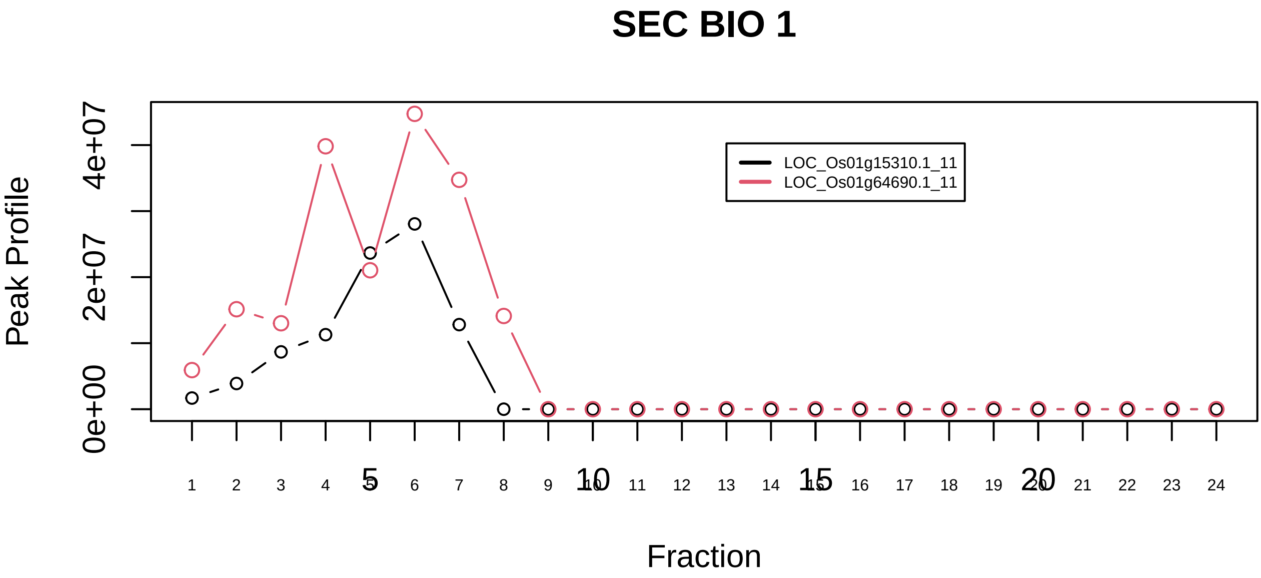

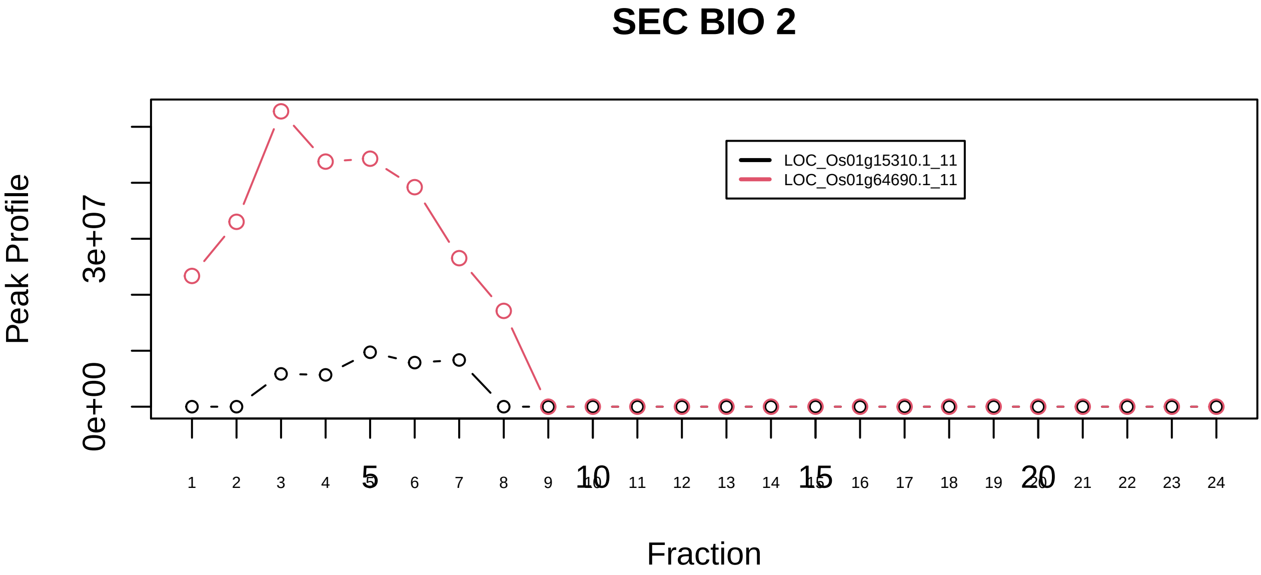

LSm1−7 complex|561−−sub_1

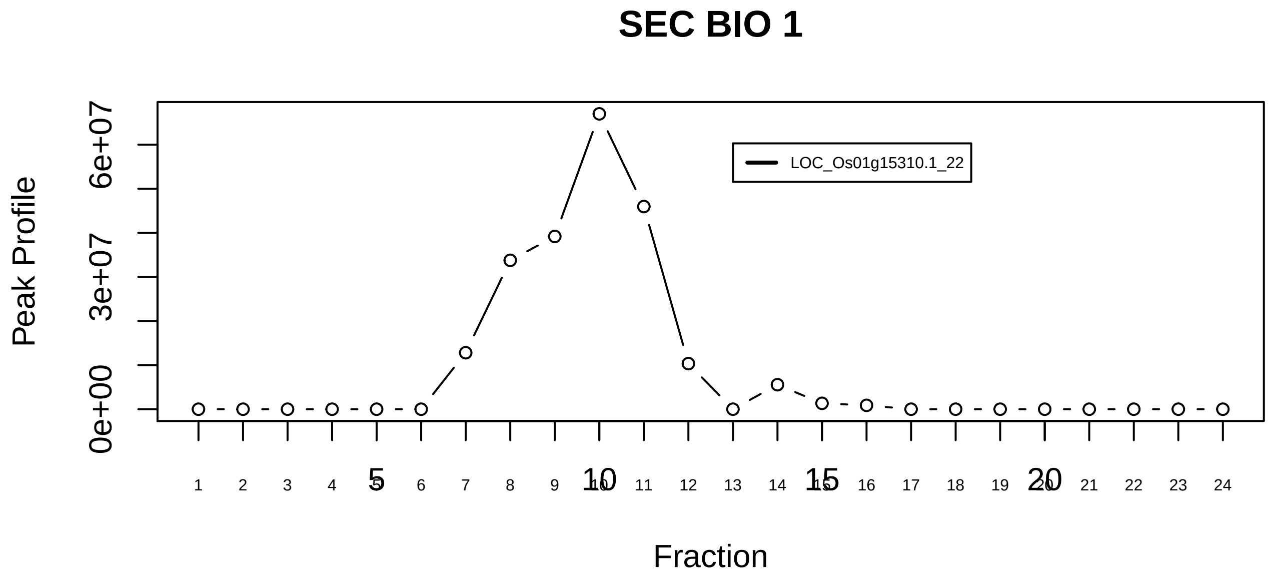

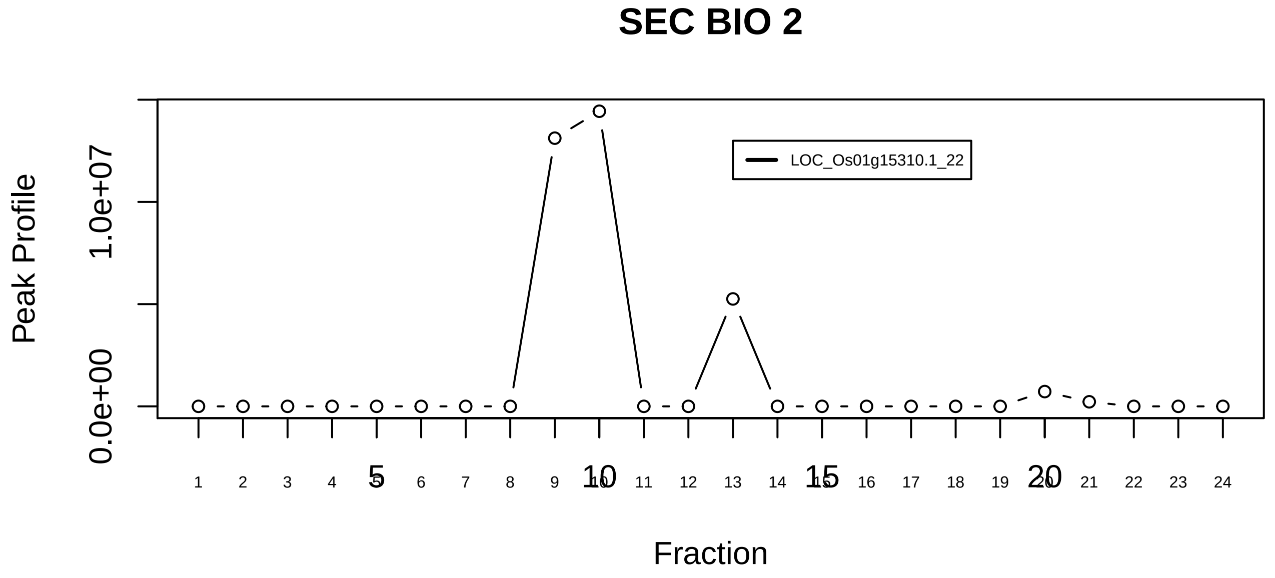

LSm1−7 complex|561−−singleton_1

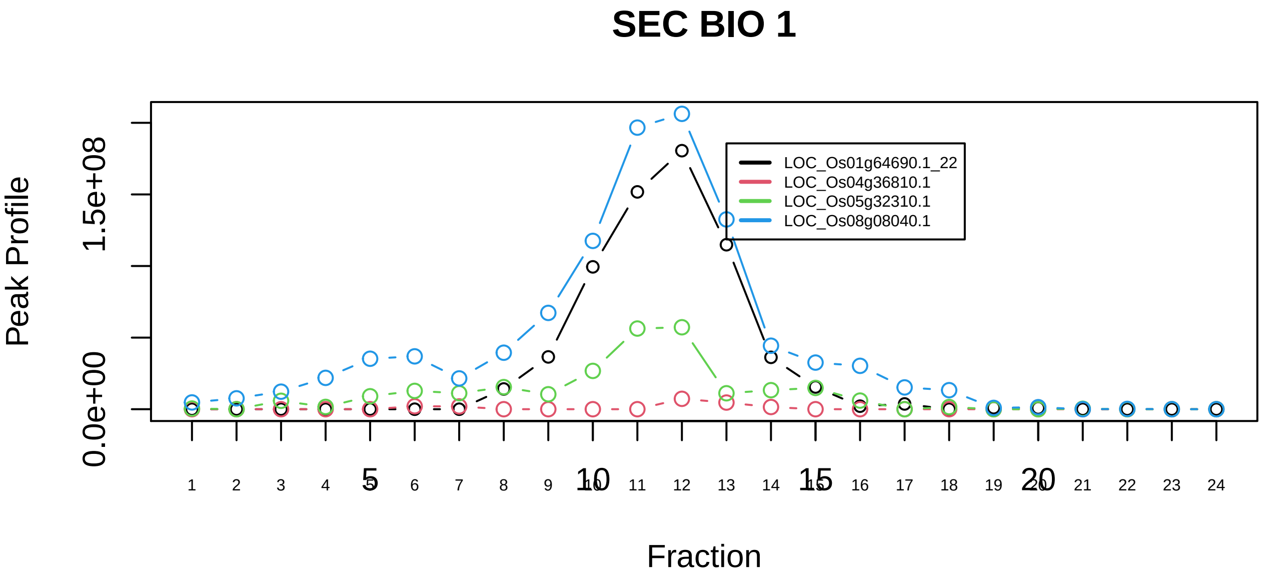

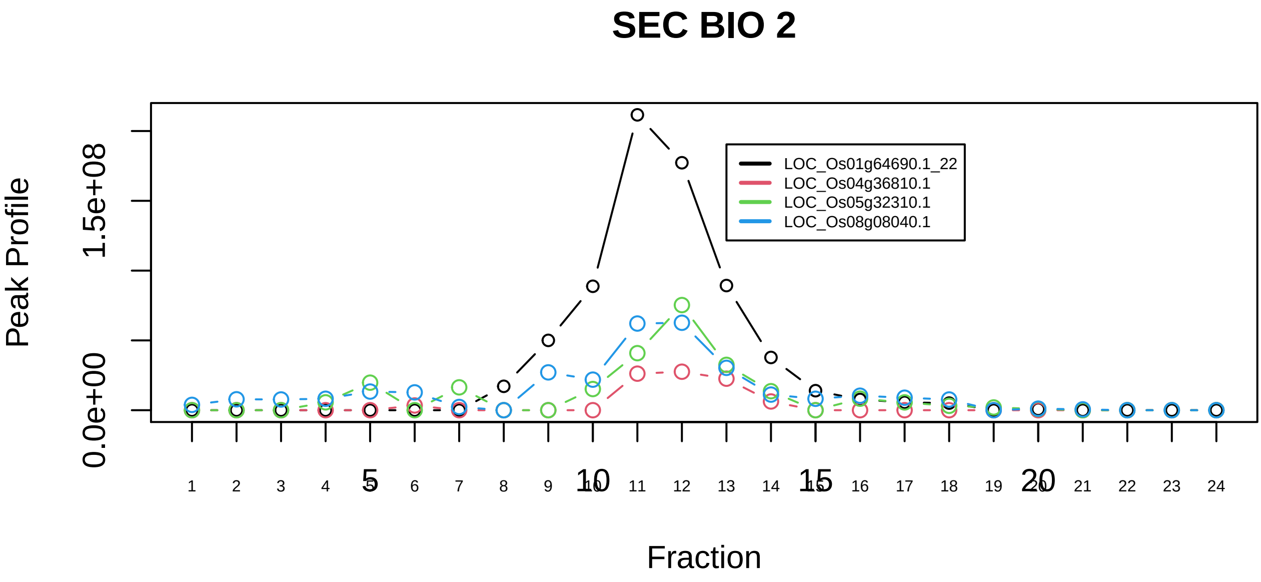

LSm1−7 complex|561−−sub_2

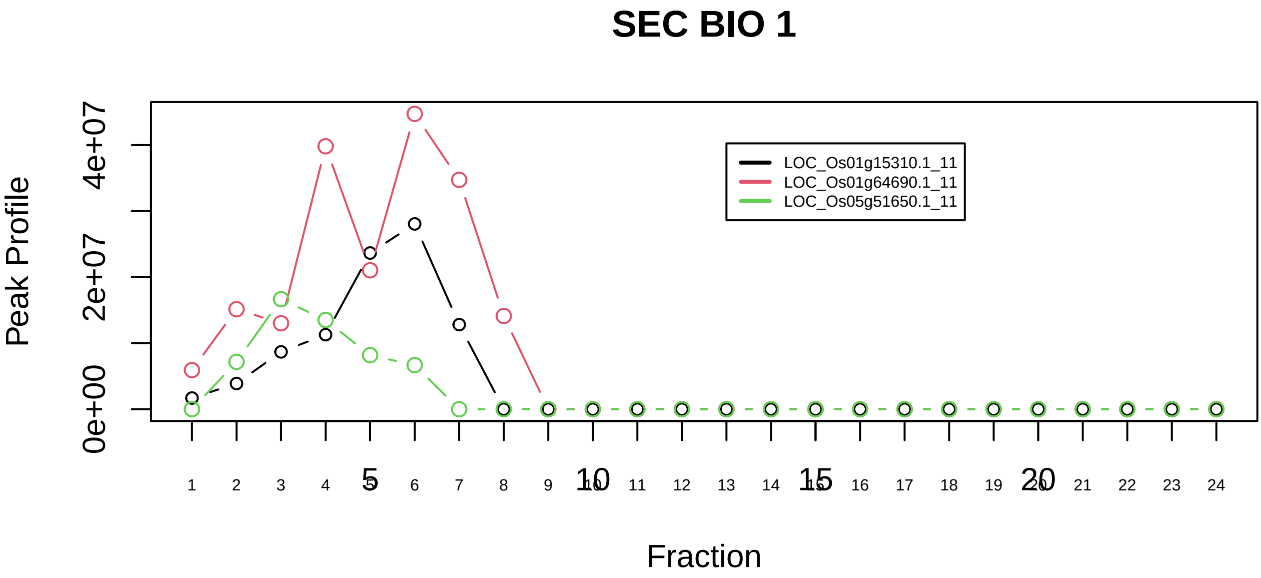

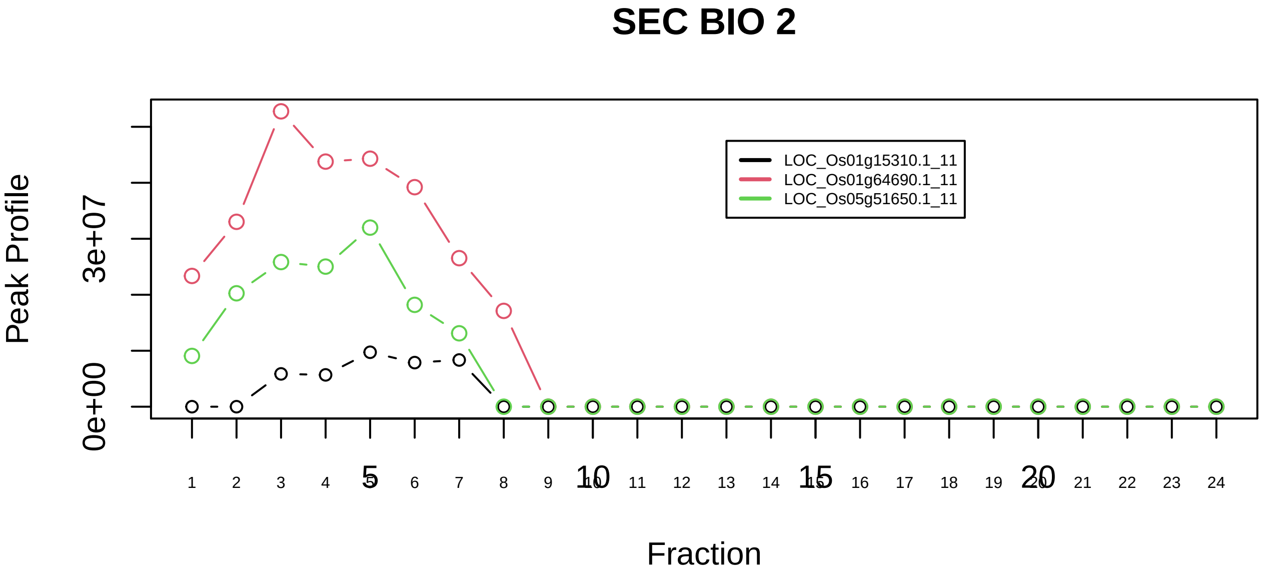

LSm2−8 complex|562−−sub_1

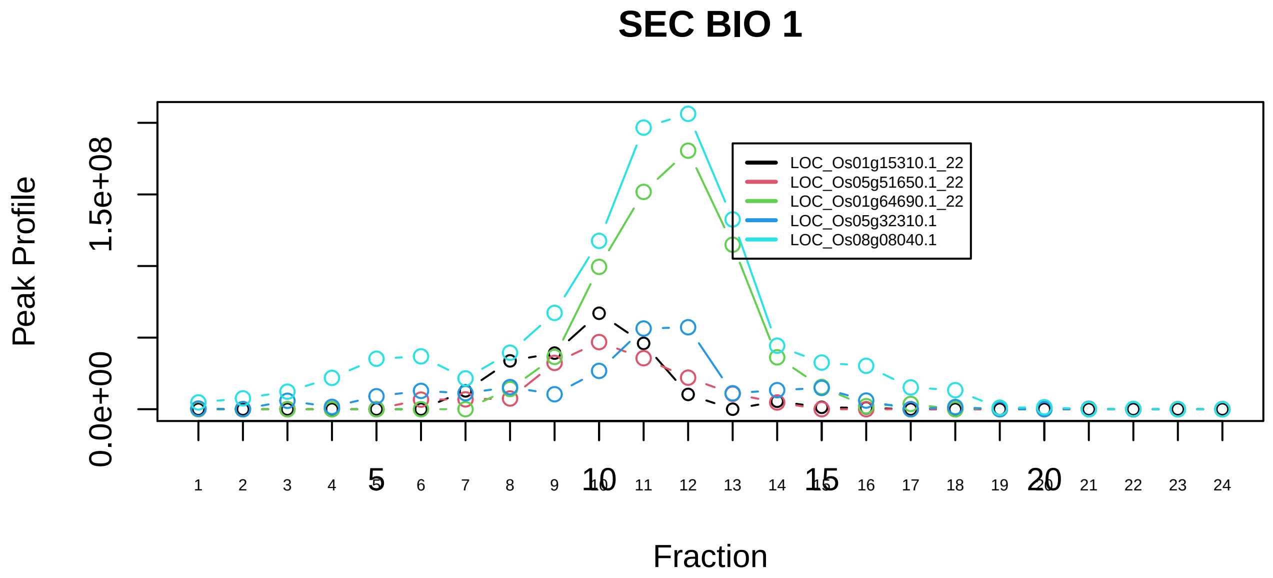

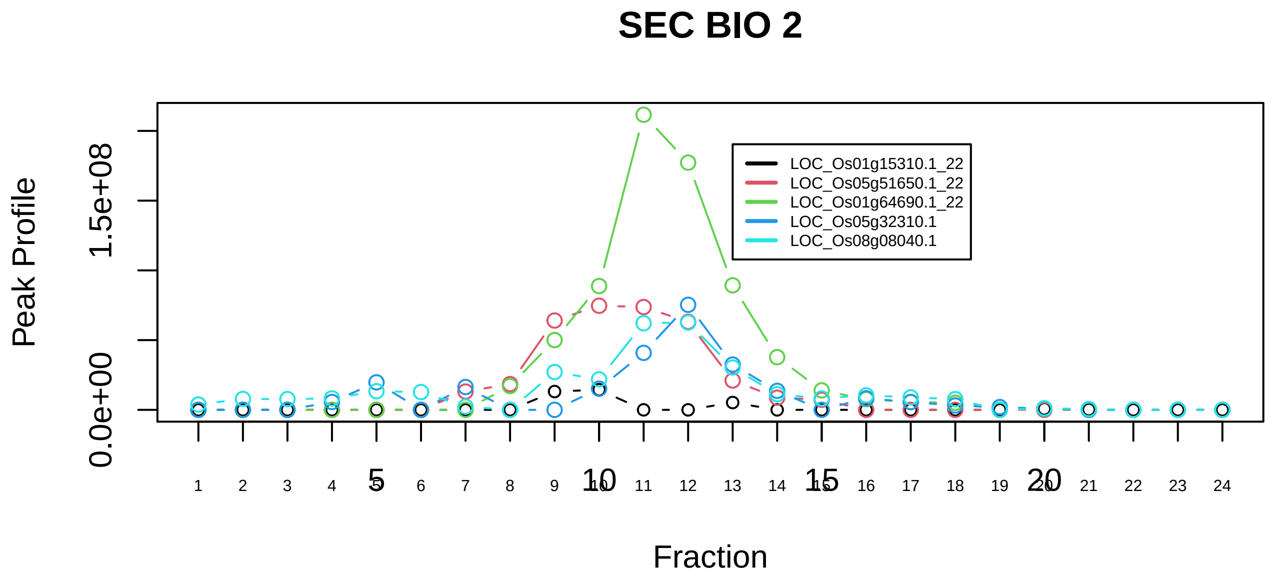

LSm2−8 complex|562−−sub_2

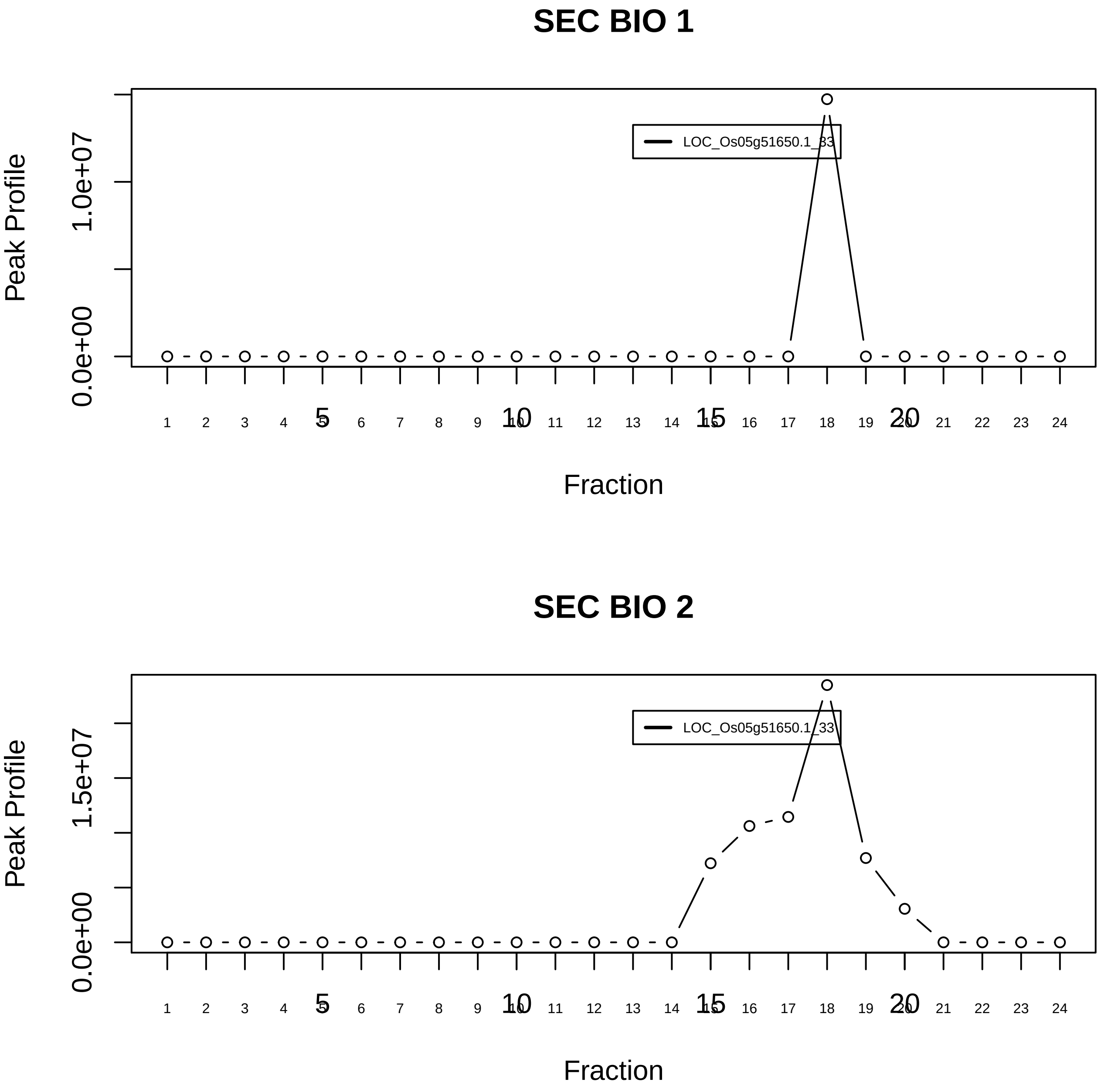

LSm2−8 complex|562−−singleton_1

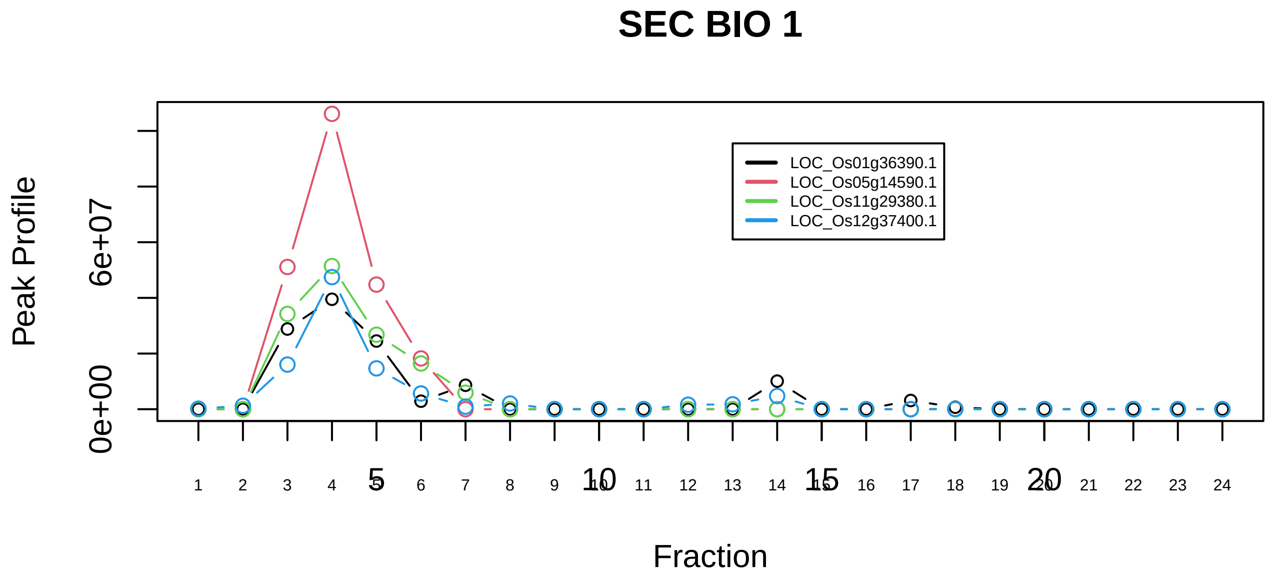

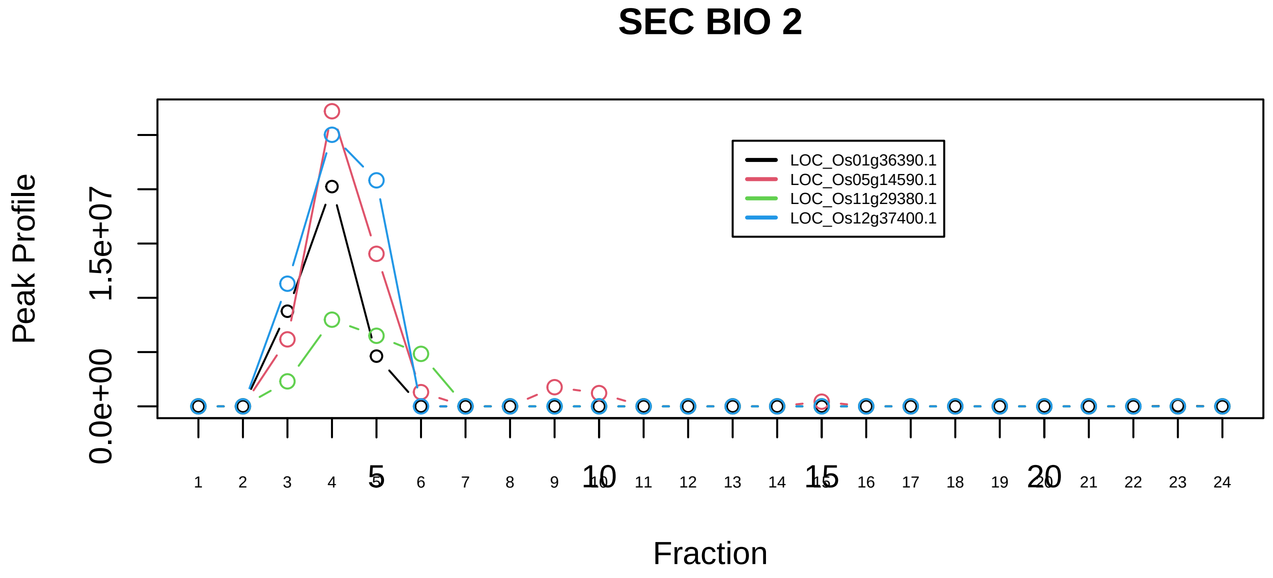

MCM2−MCM4−MCM6−MCM7 complex|2792−−all_prots_before_clustering_sim

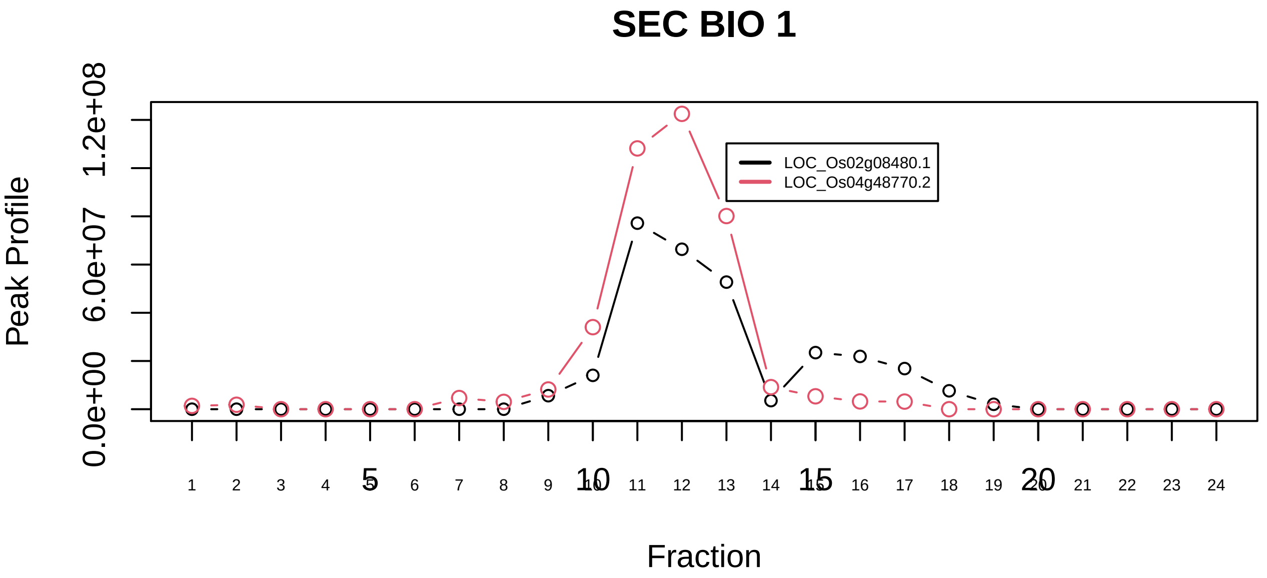

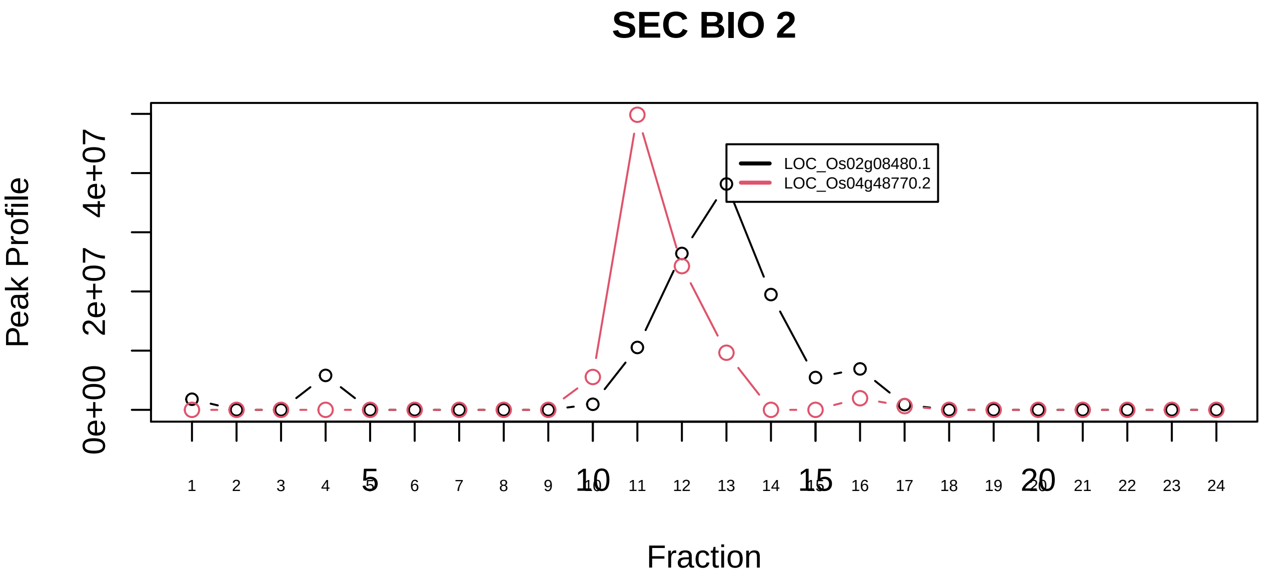

Membrane protein complex (VCP, UFD1L, SEC61B)|5685−−sub_orthoparalog_1

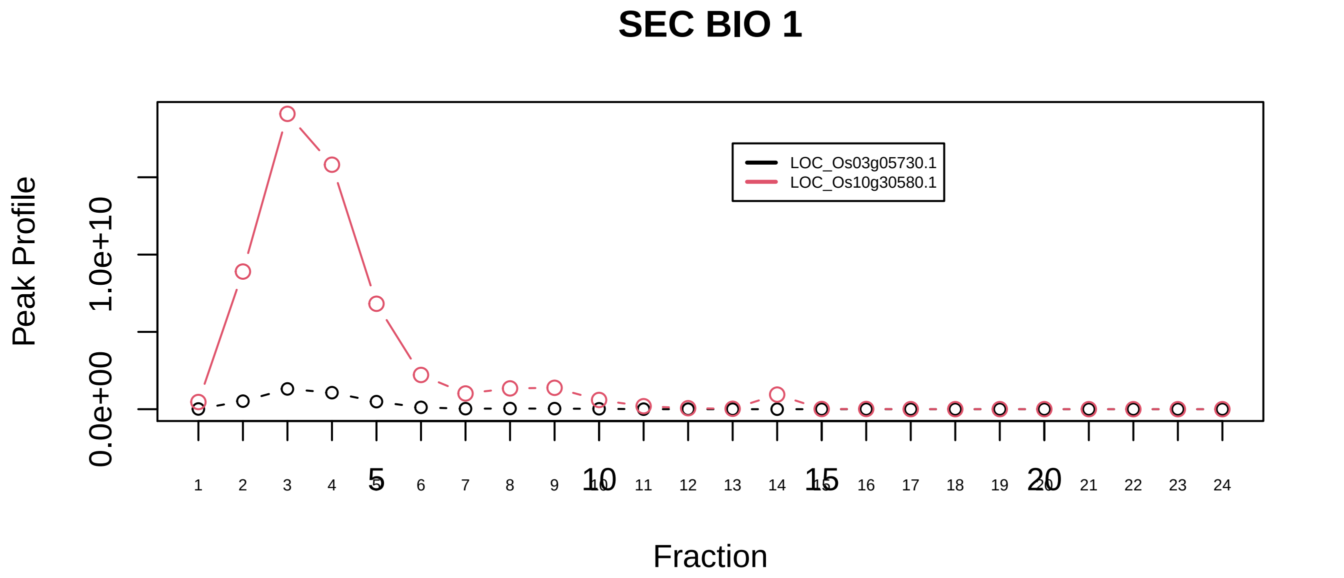

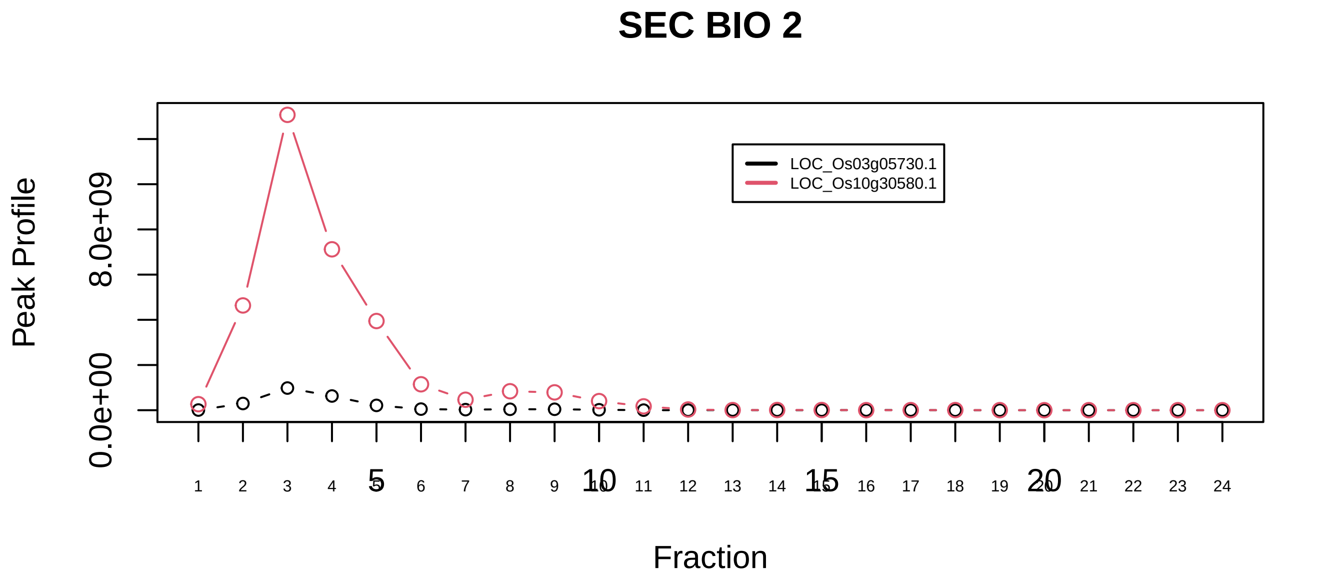

Membrane protein complex (VCP, UFD1L, SEC61B)|5685−−sub_orthoparalog_2

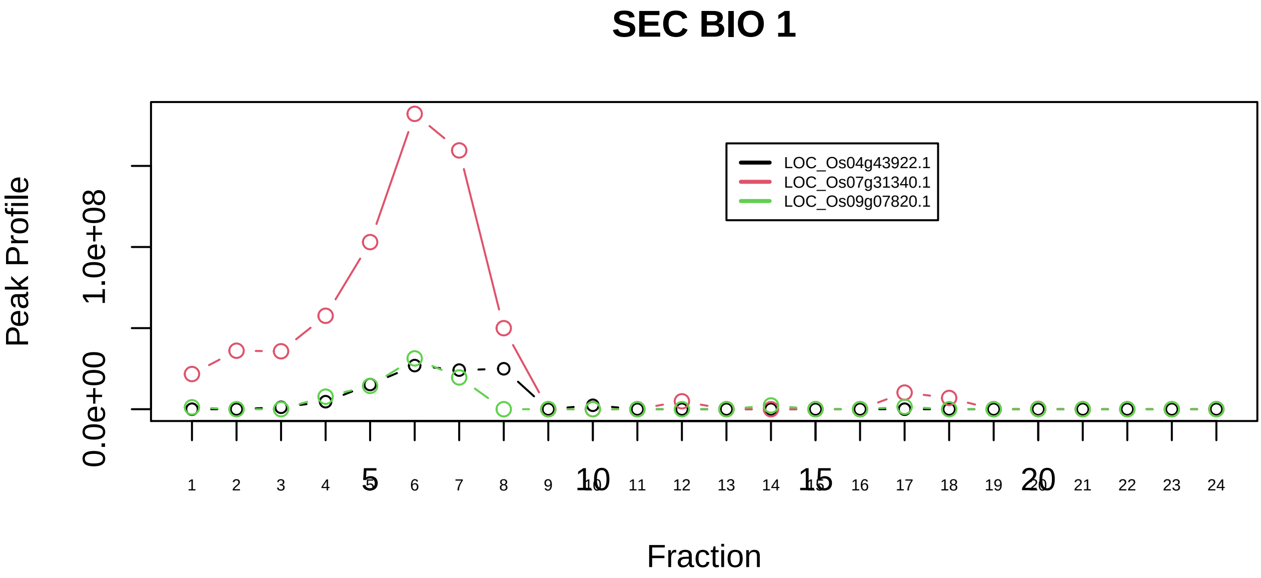

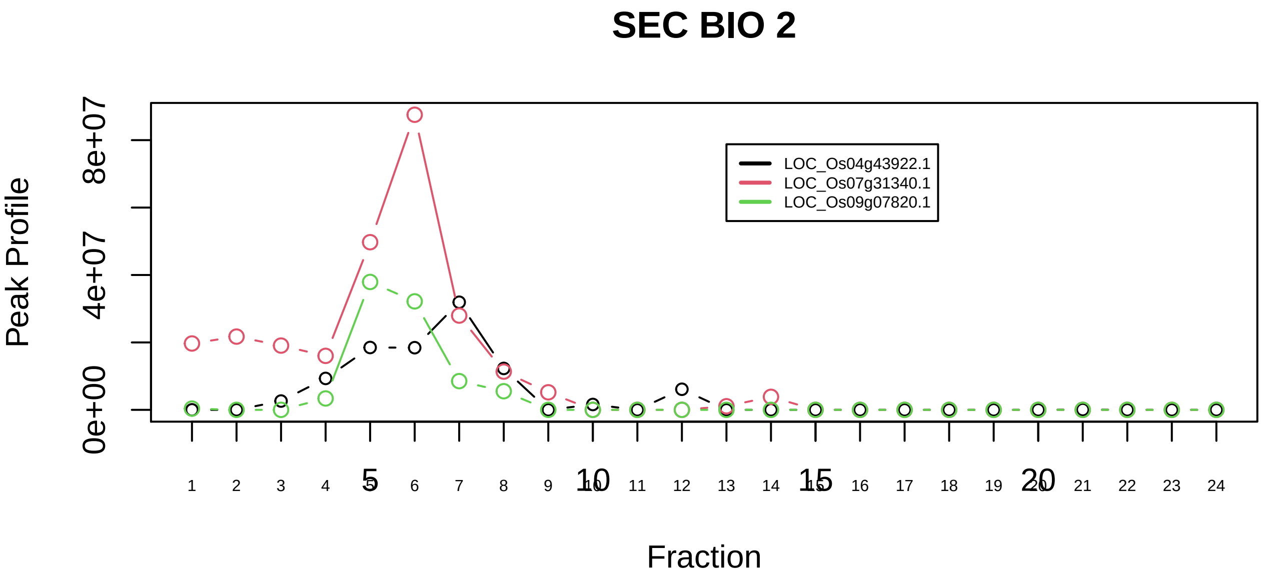

mRNA decay complex (UPF1, UPF2, UPF3B, DCP2, XRN1, XRN2, EXOSC2, EXOSC4, EXOSC10, PARN)|822−−sub_1

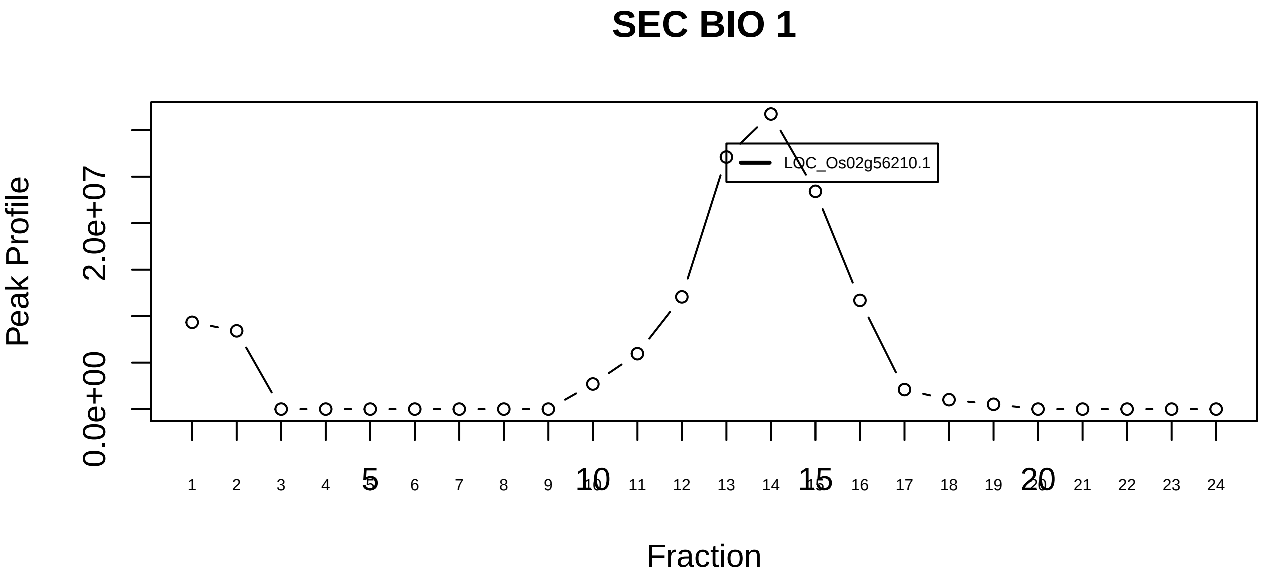

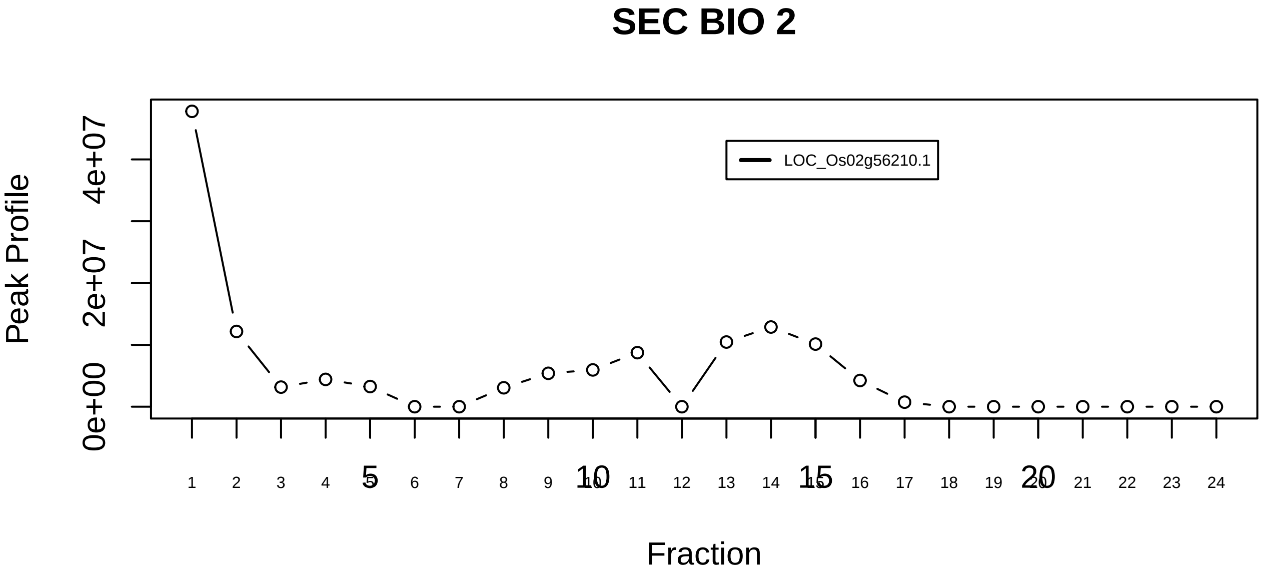

mRNA decay complex (UPF1, UPF2, UPF3B, DCP2, XRN1, XRN2, EXOSC2, EXOSC4, EXOSC10, PARN)|822−−singleton_1

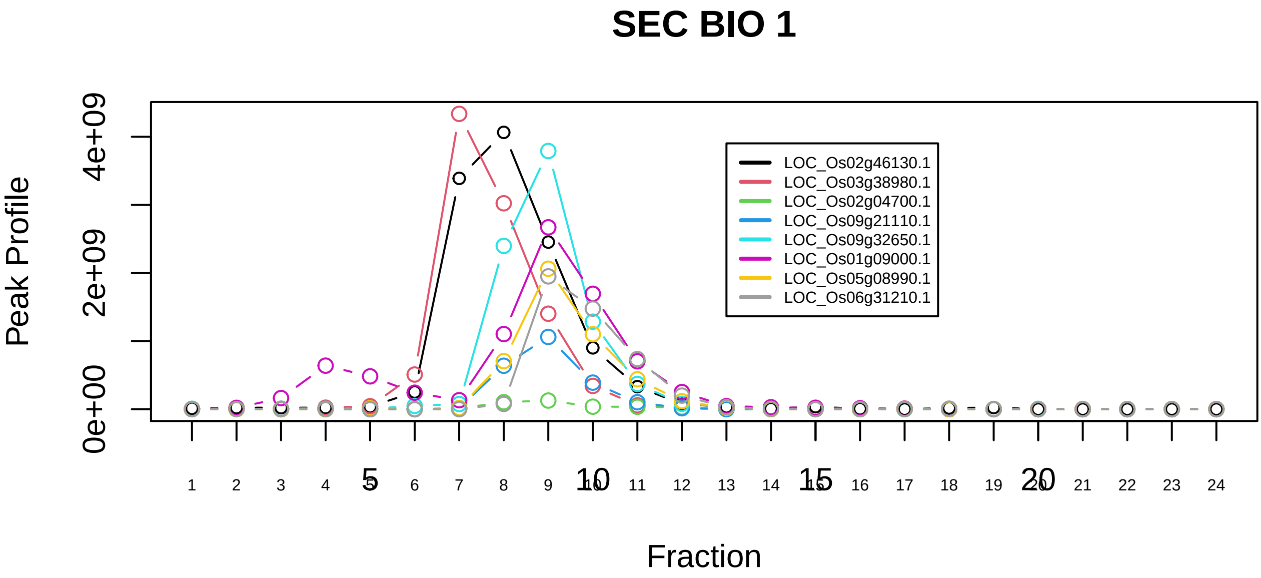

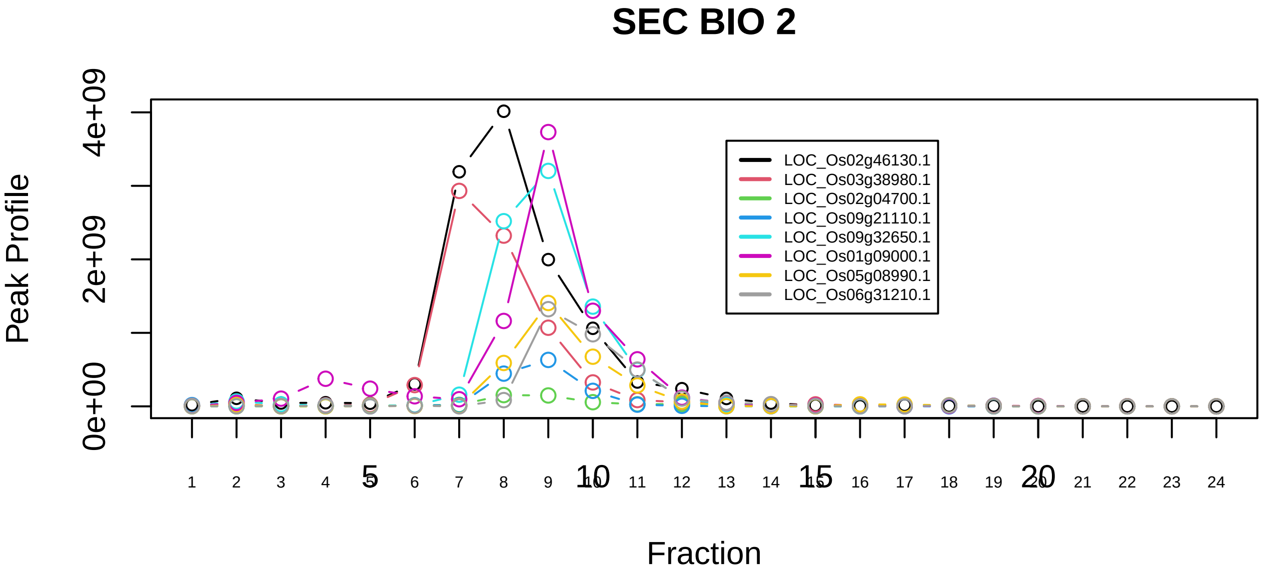

Multisynthetase complex|3040−−sub_1

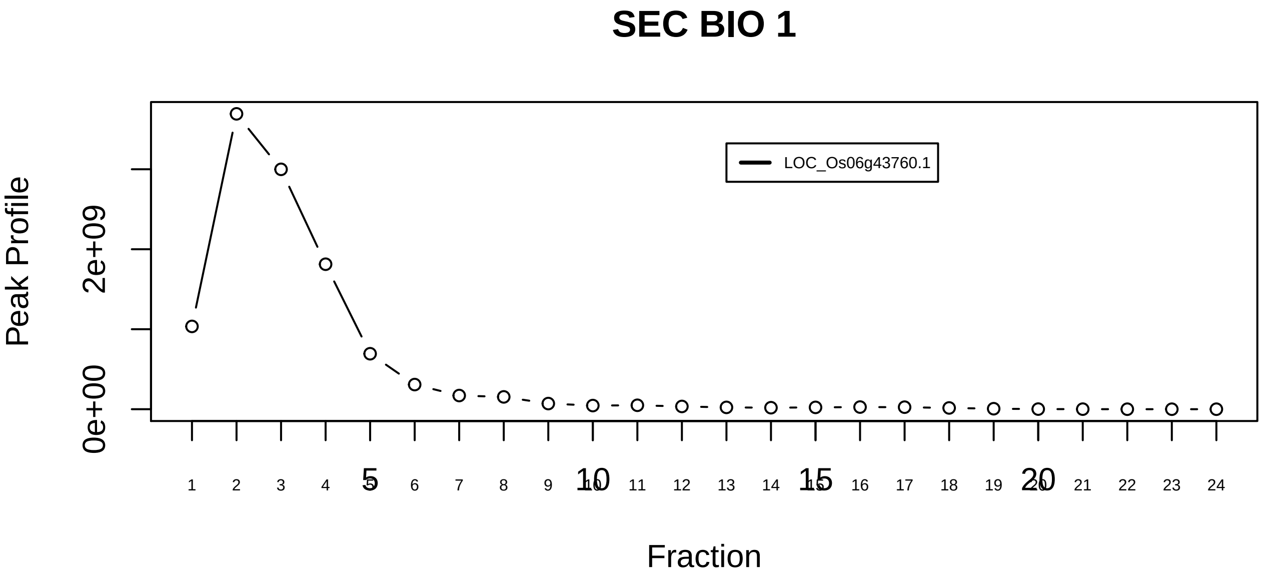

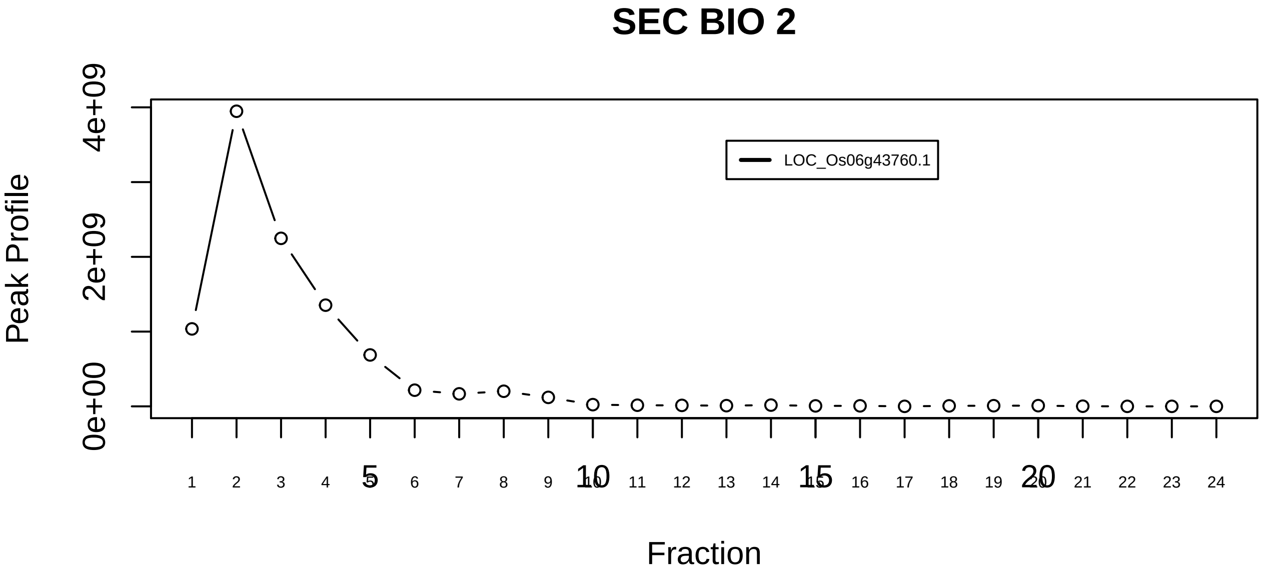

Multisynthetase complex|3040−−singleton_1

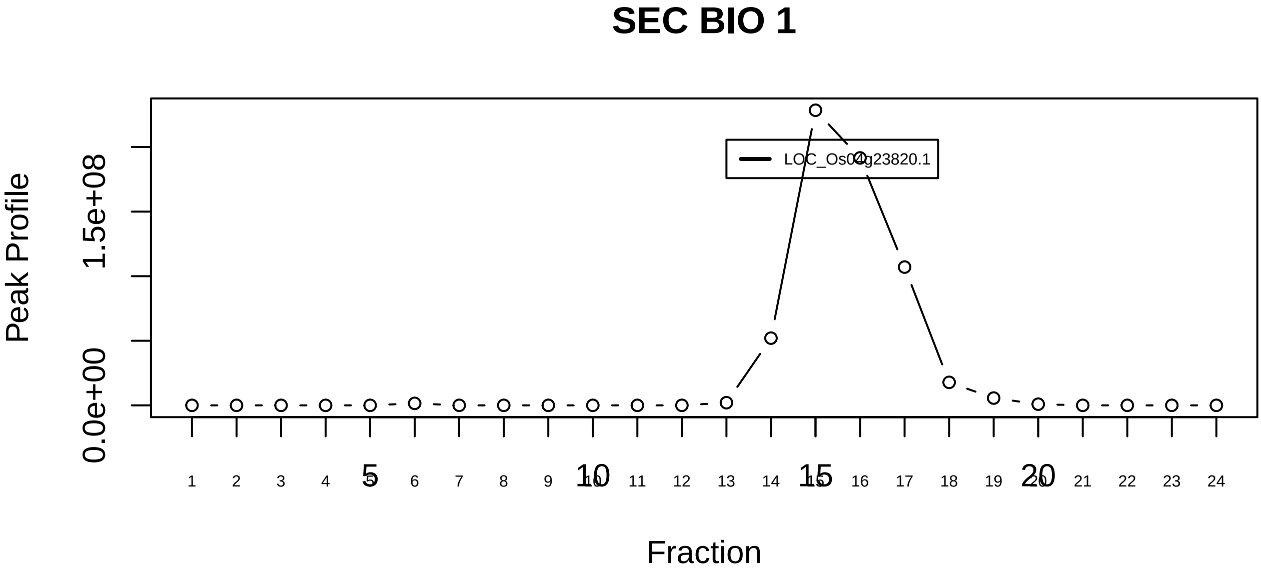

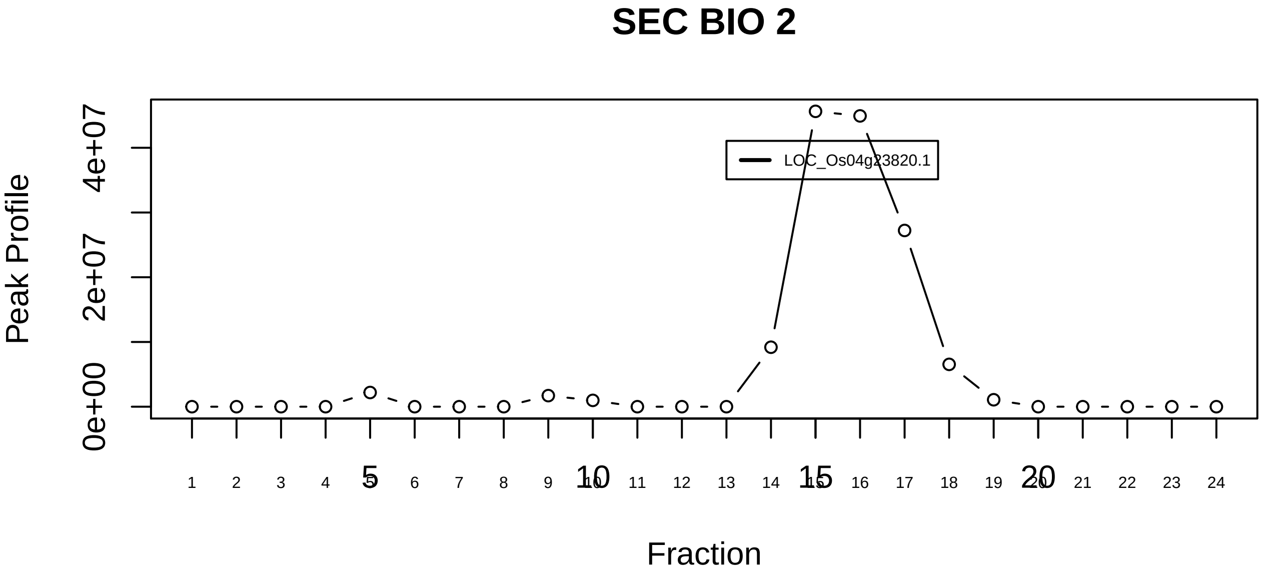

Multisynthetase complex|3040−−singleton_2

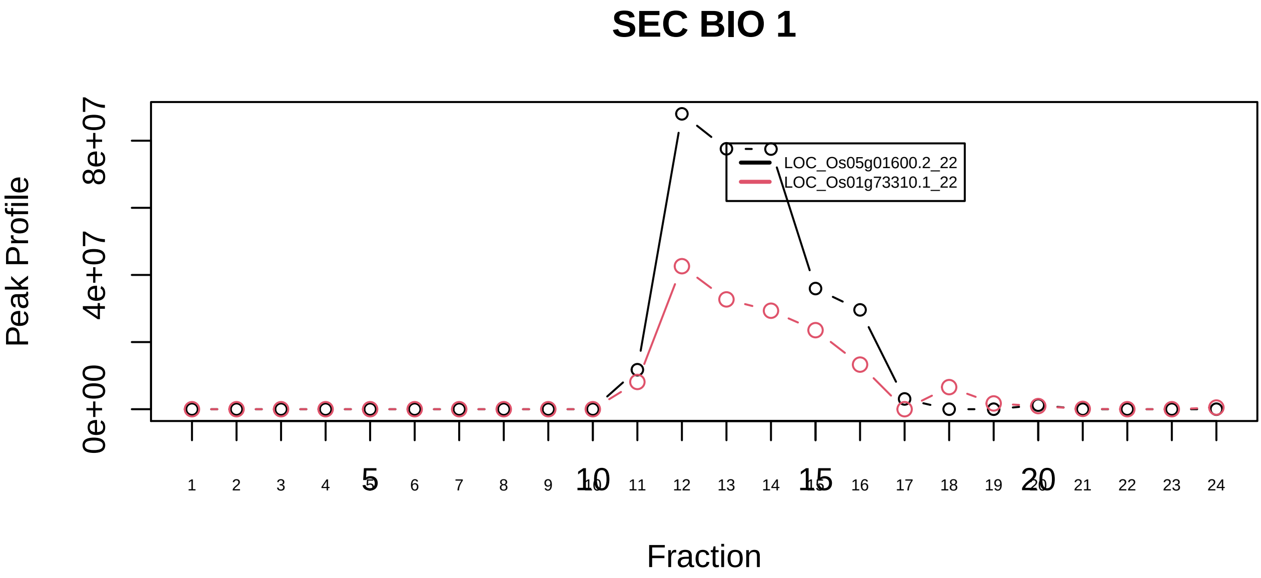

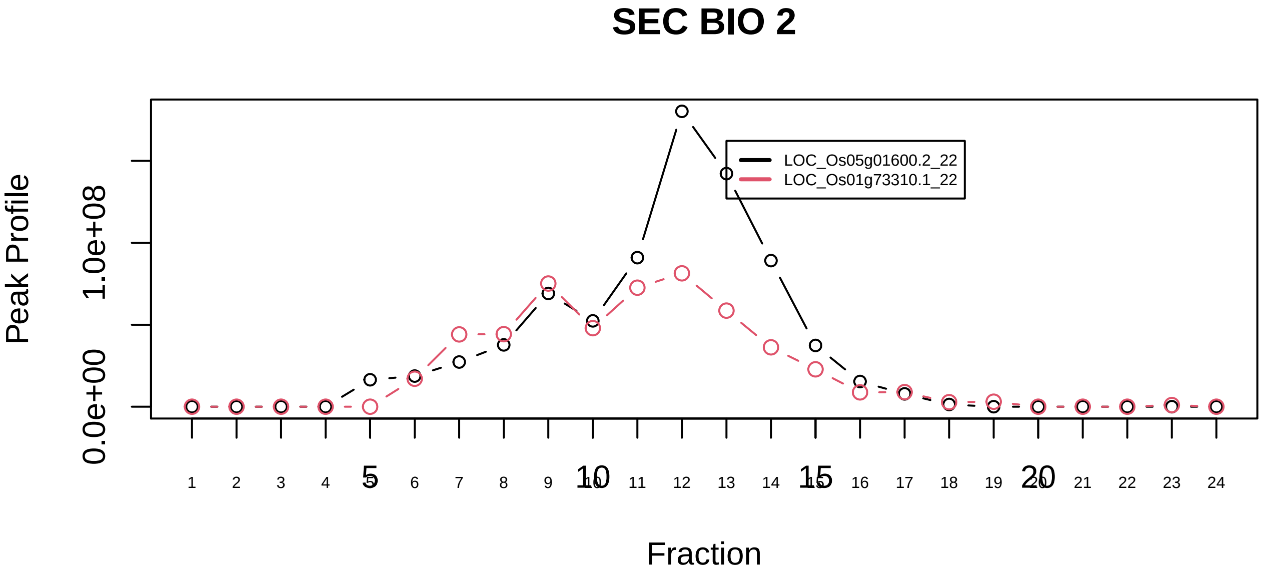

Nop56p−associated pre−rRNA complex|3055−−sub_1

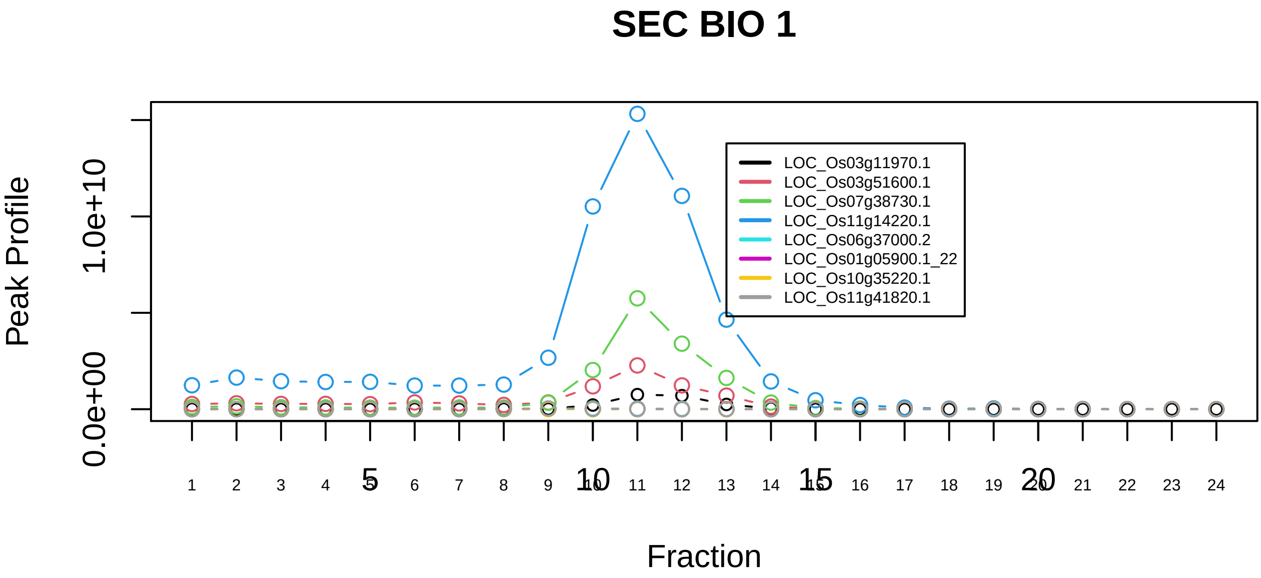

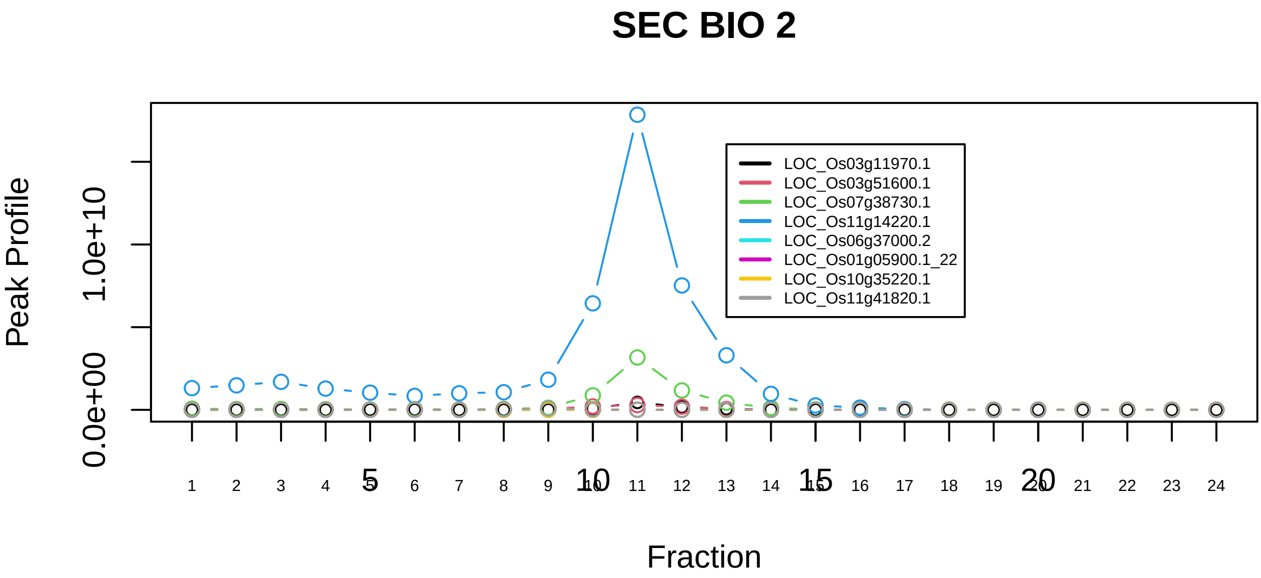

Nop56p−associated pre−rRNA complex|3055−−sub_2

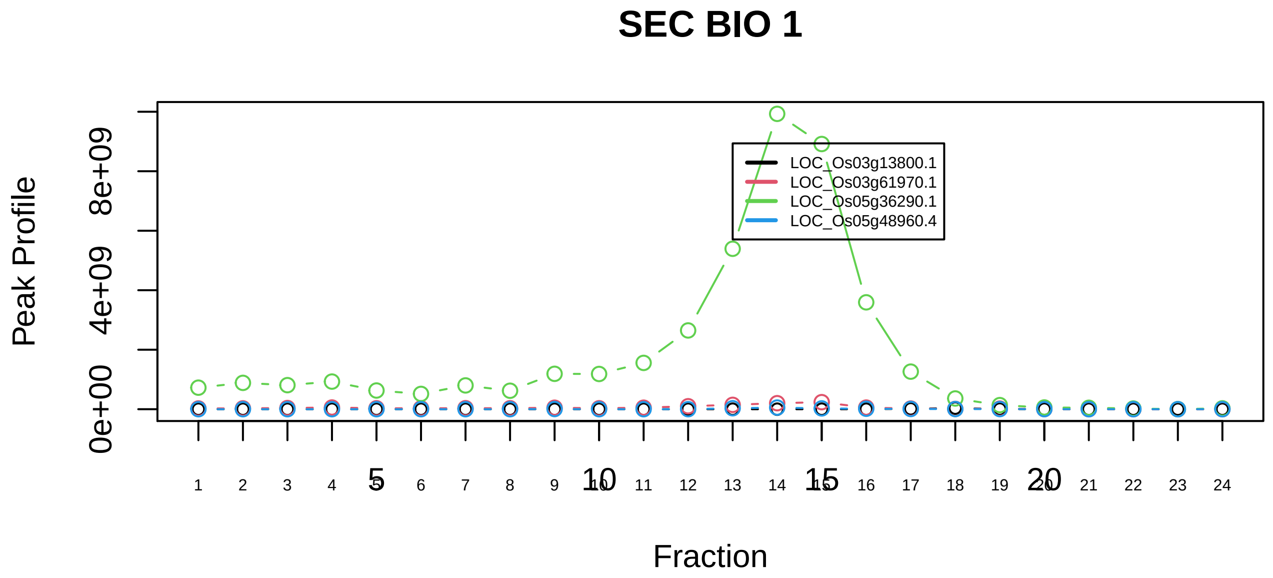

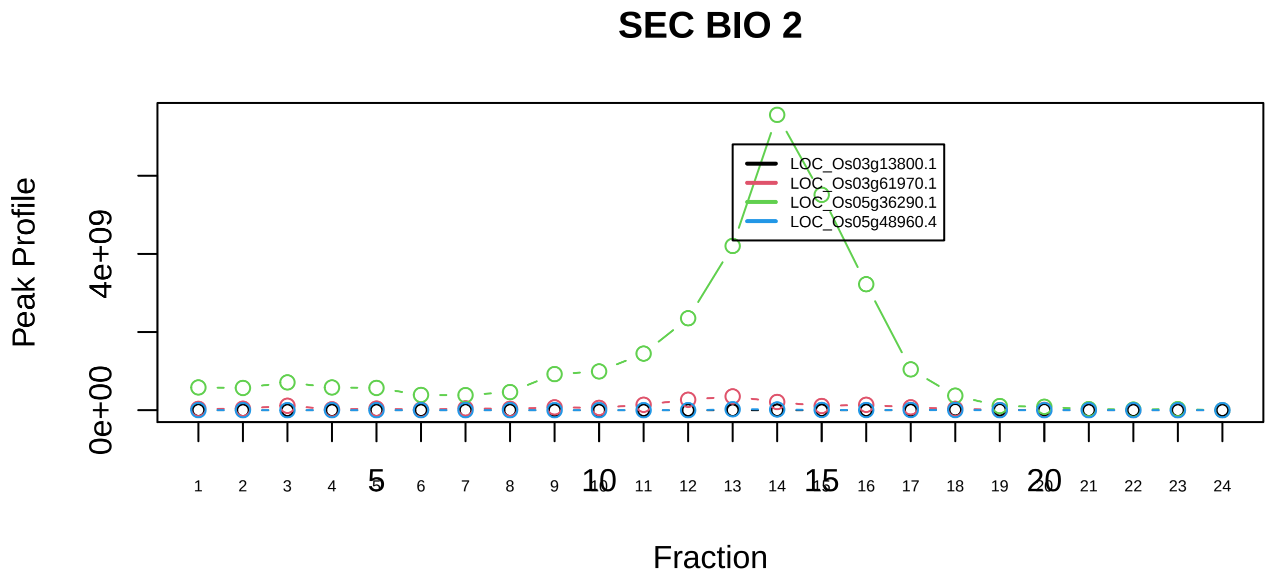

Nop56p−associated pre−rRNA complex|3055−−sub_3

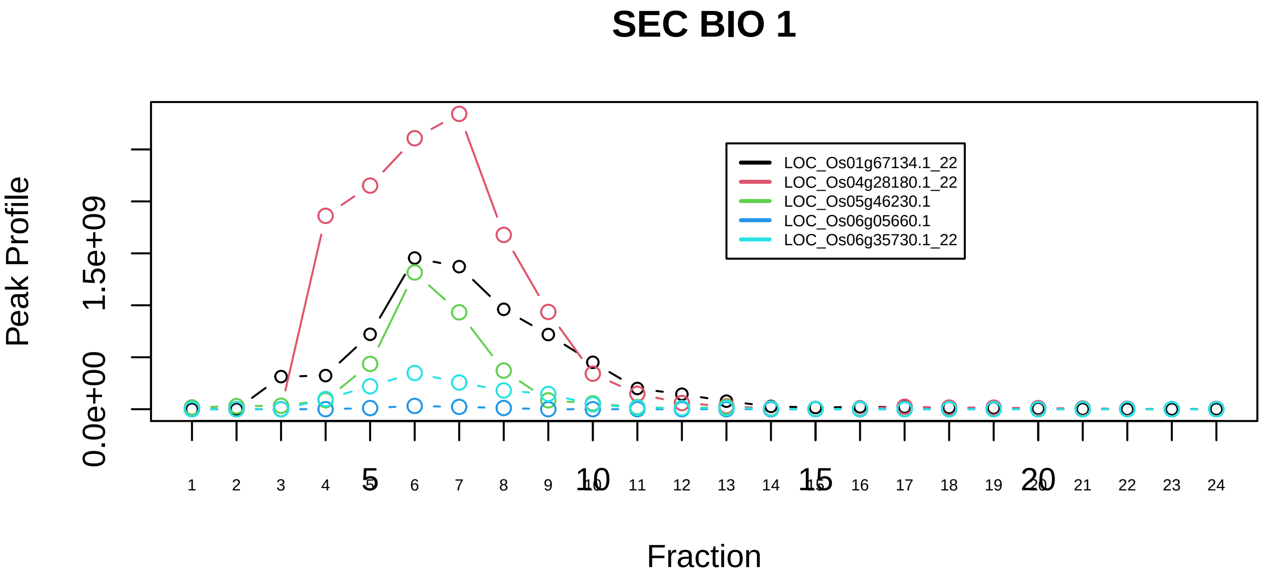

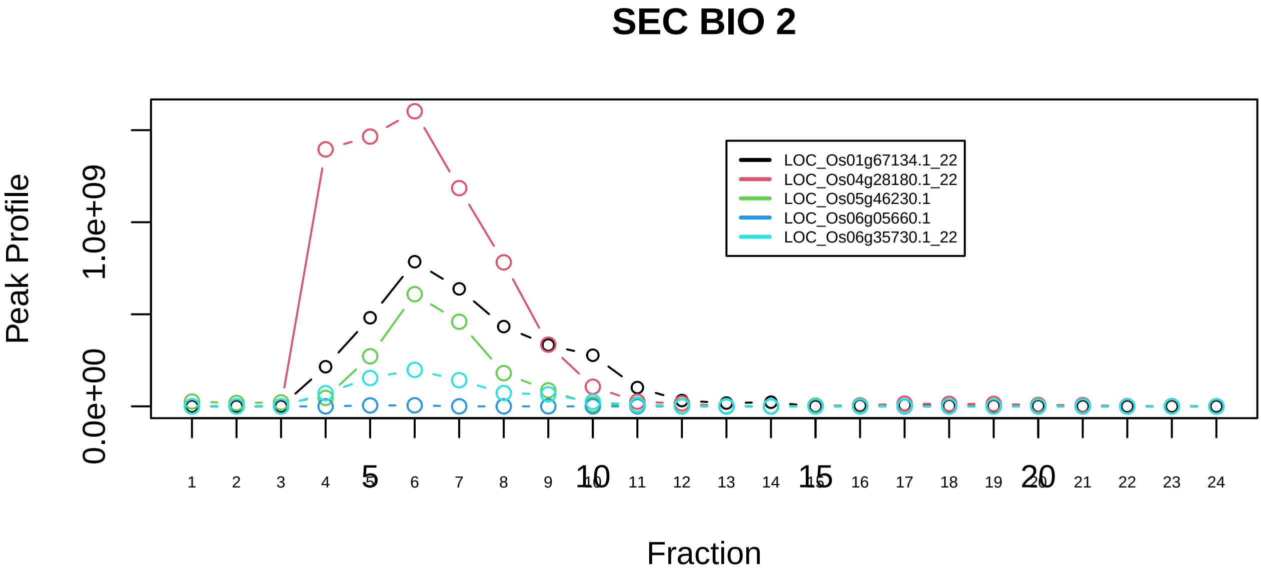

Nop56p−associated pre−rRNA complex|3055−−sub_4

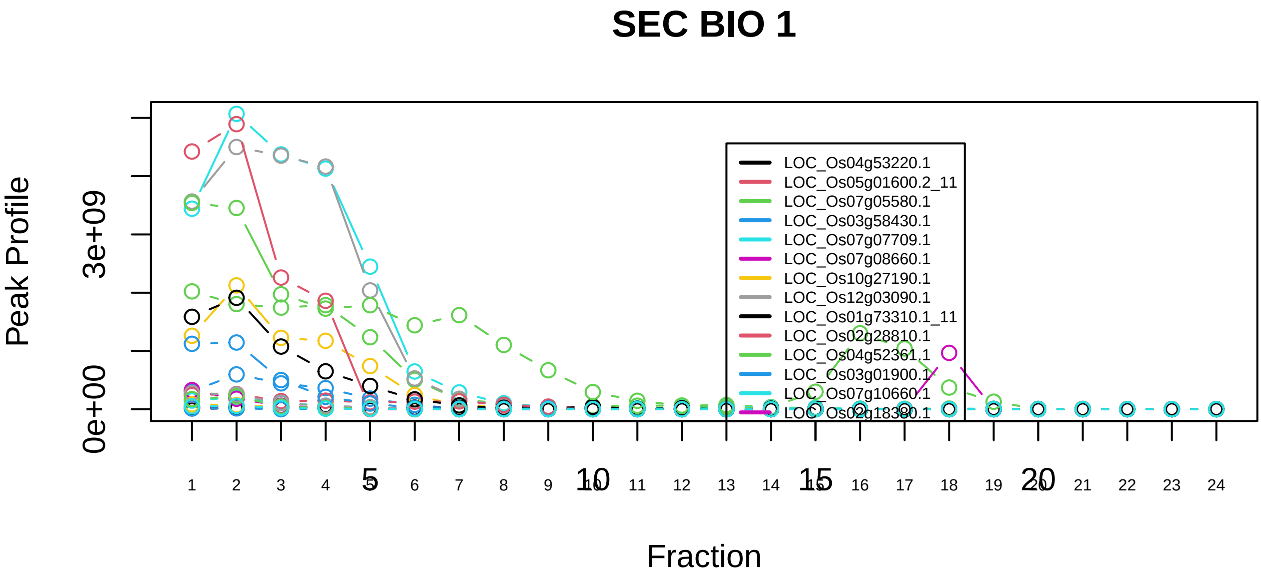

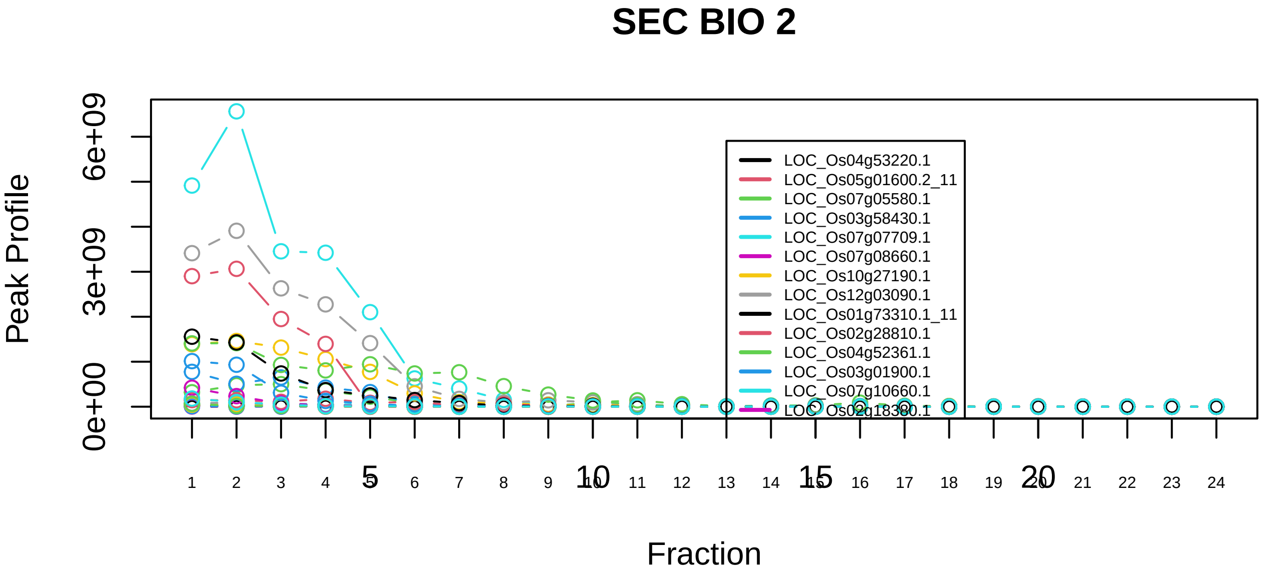

Nop56p−associated pre−rRNA complex|3055−−sub_5

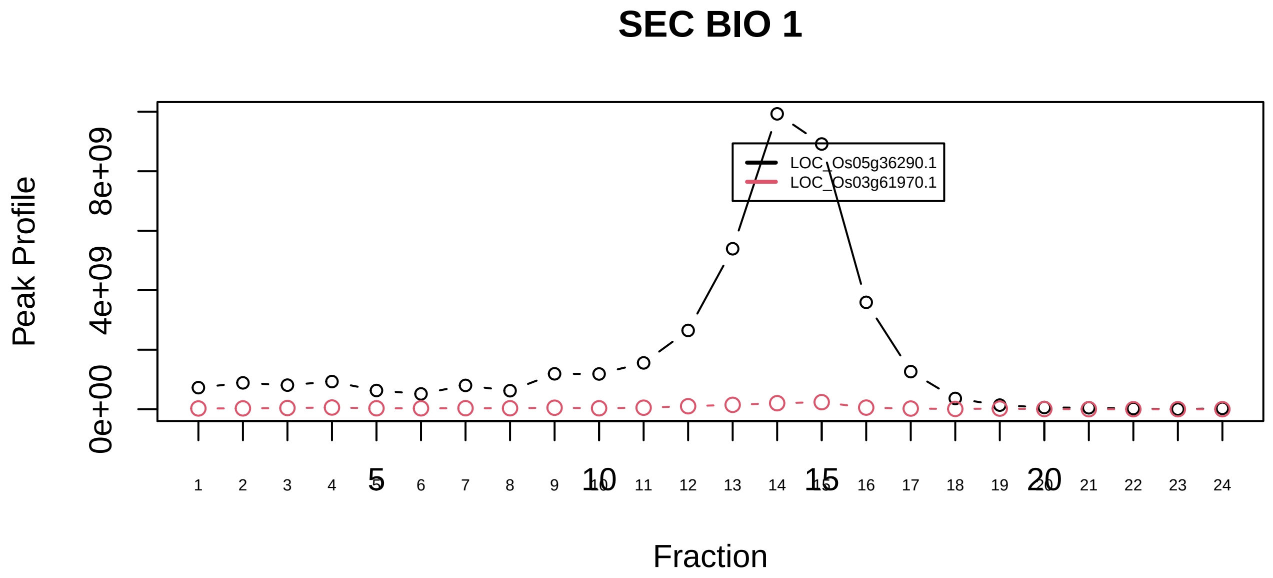

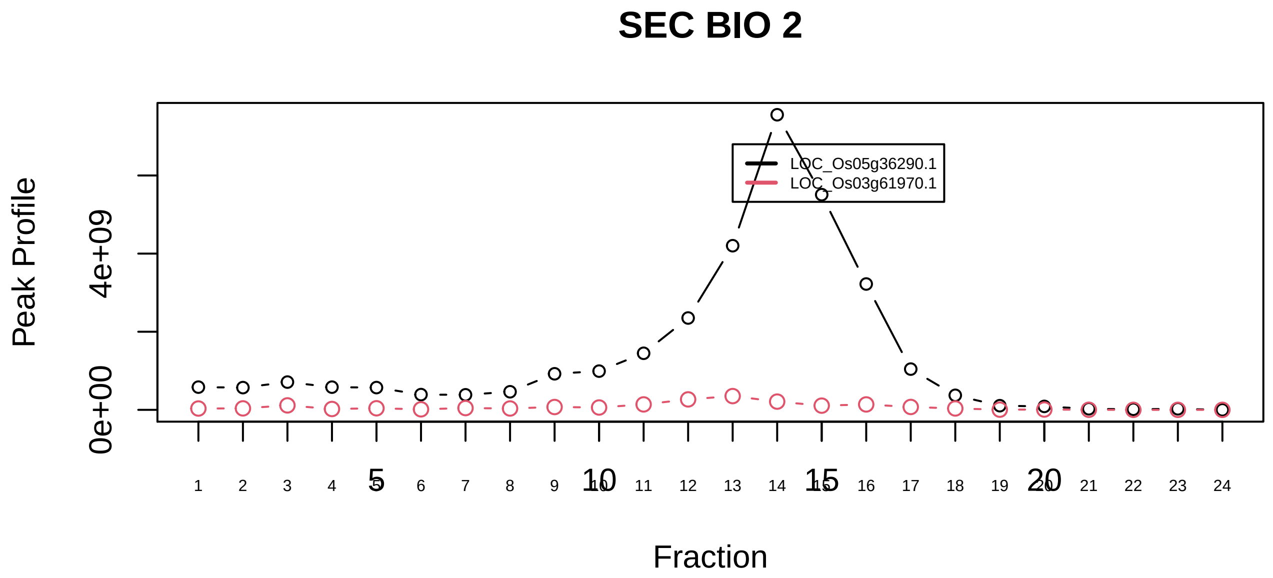

NUMAC complex (nucleosomal methylation activator complex)|86−−sub_orthoparalog_1

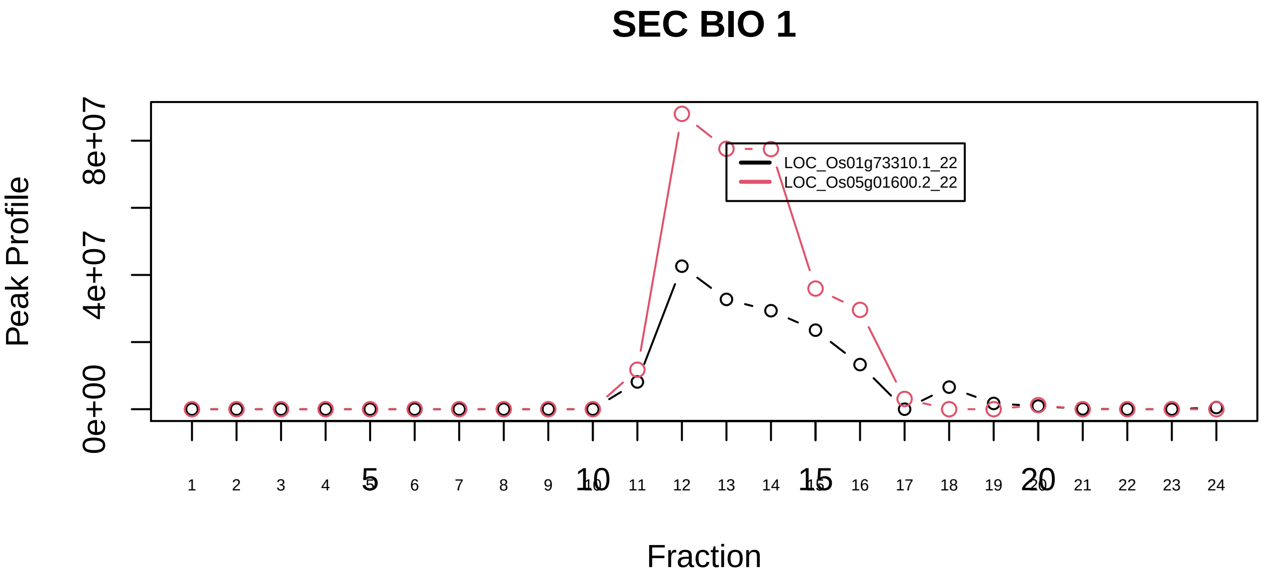

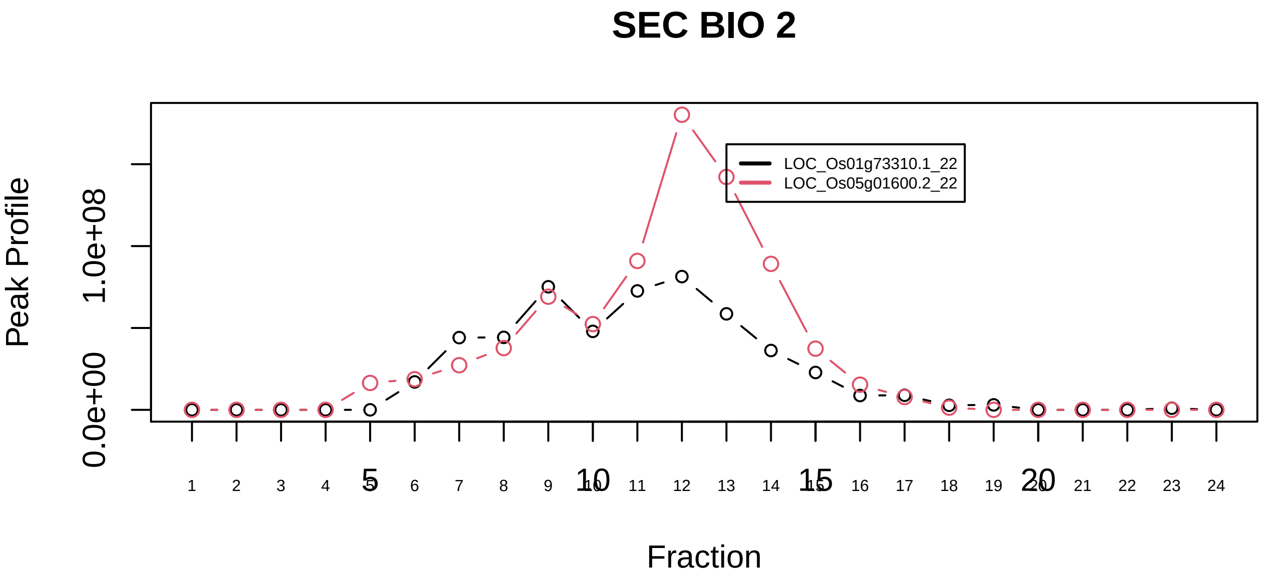

NUMAC complex (nucleosomal methylation activator complex)|86−−sub_orthoparalog_2

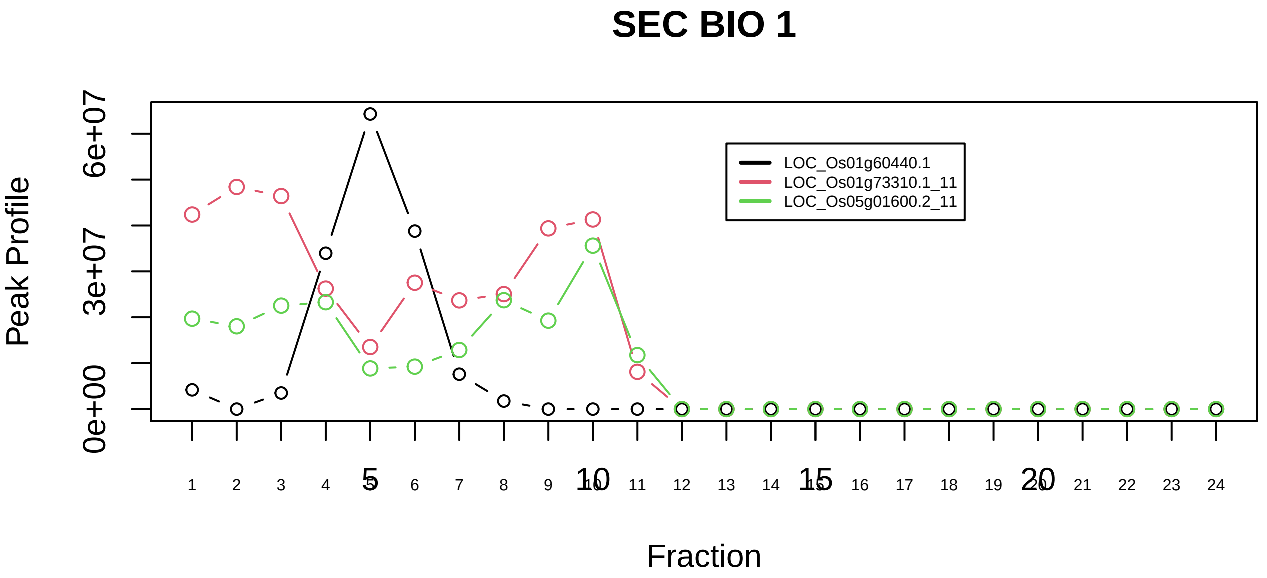

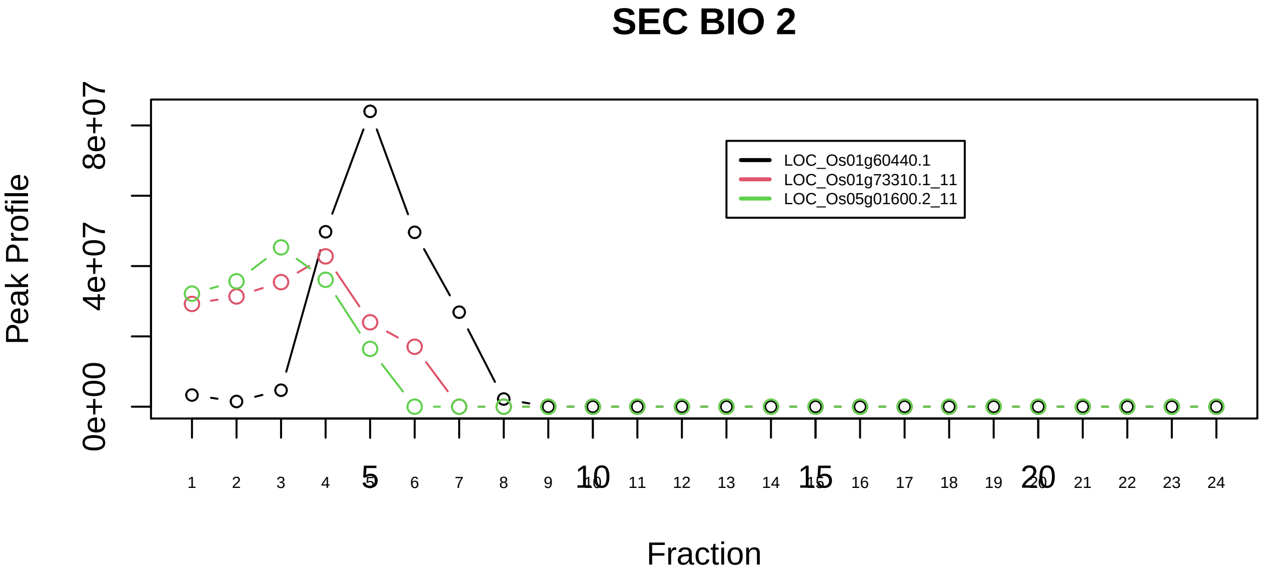

NUMAC complex (nucleosomal methylation activator complex)|86−−sub_1

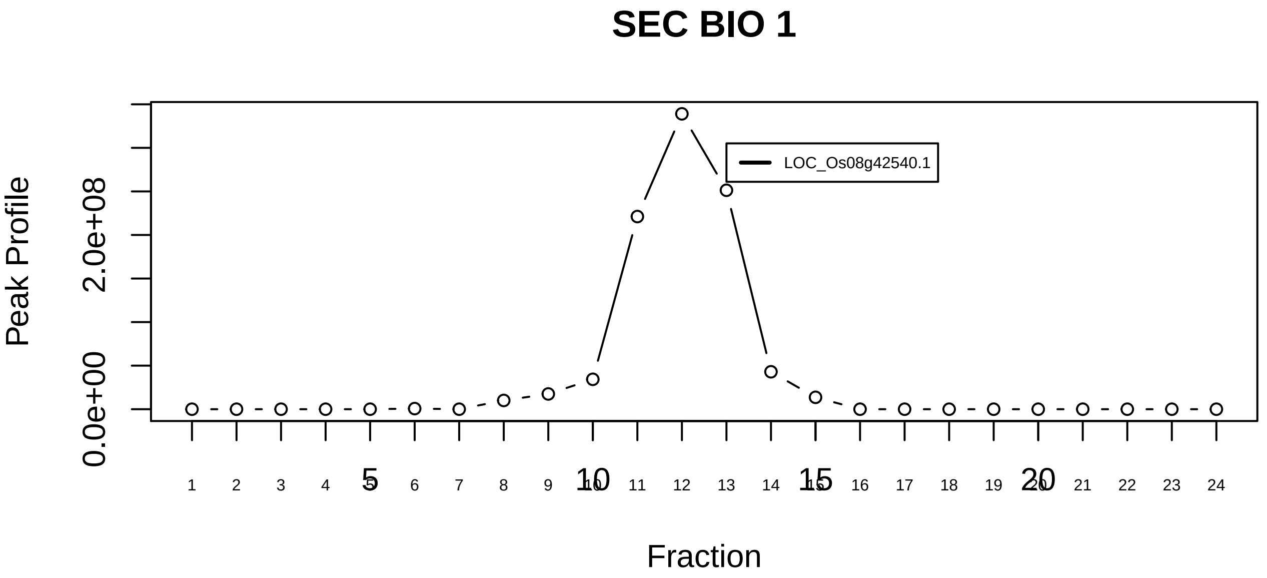

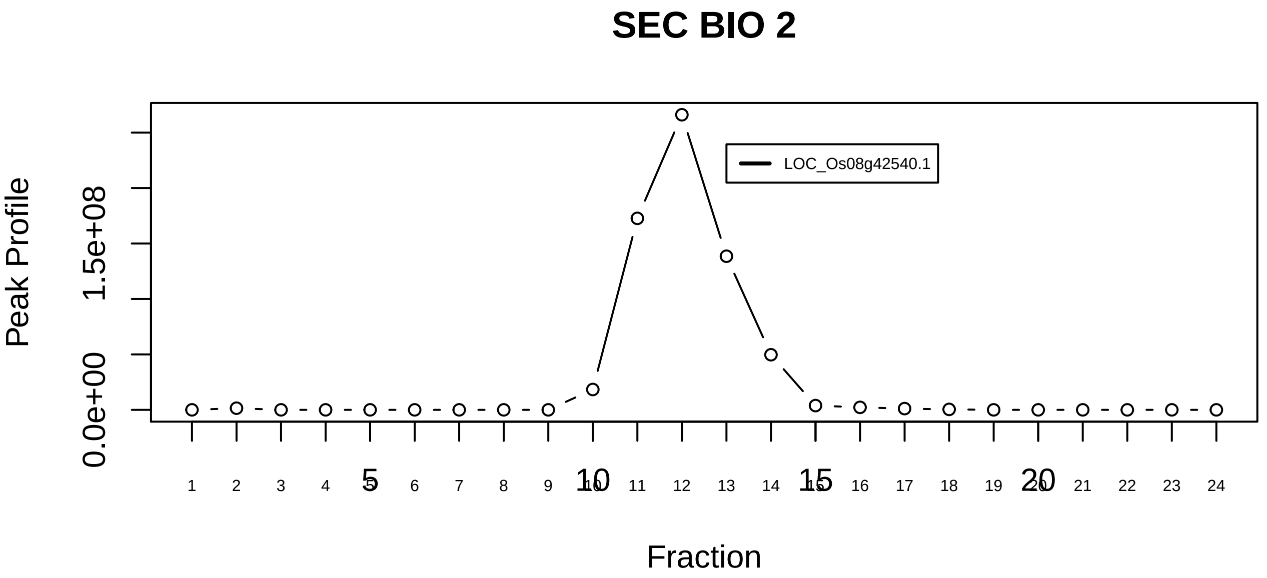

OTUB1−UBC13−MMS2 complex|7051−−singleton_1

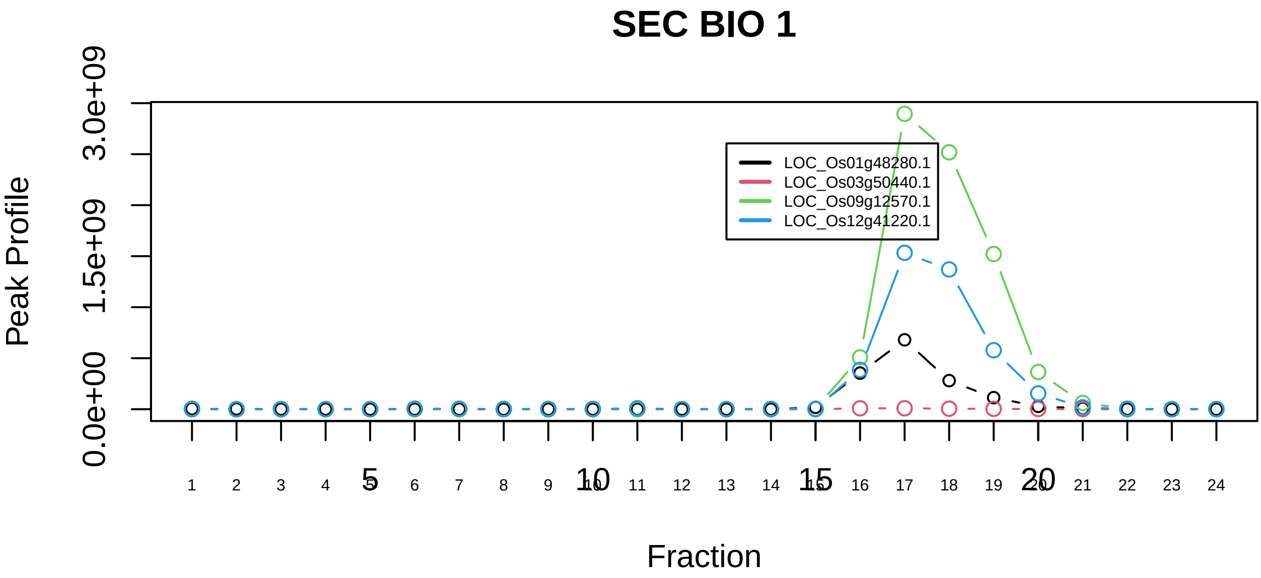

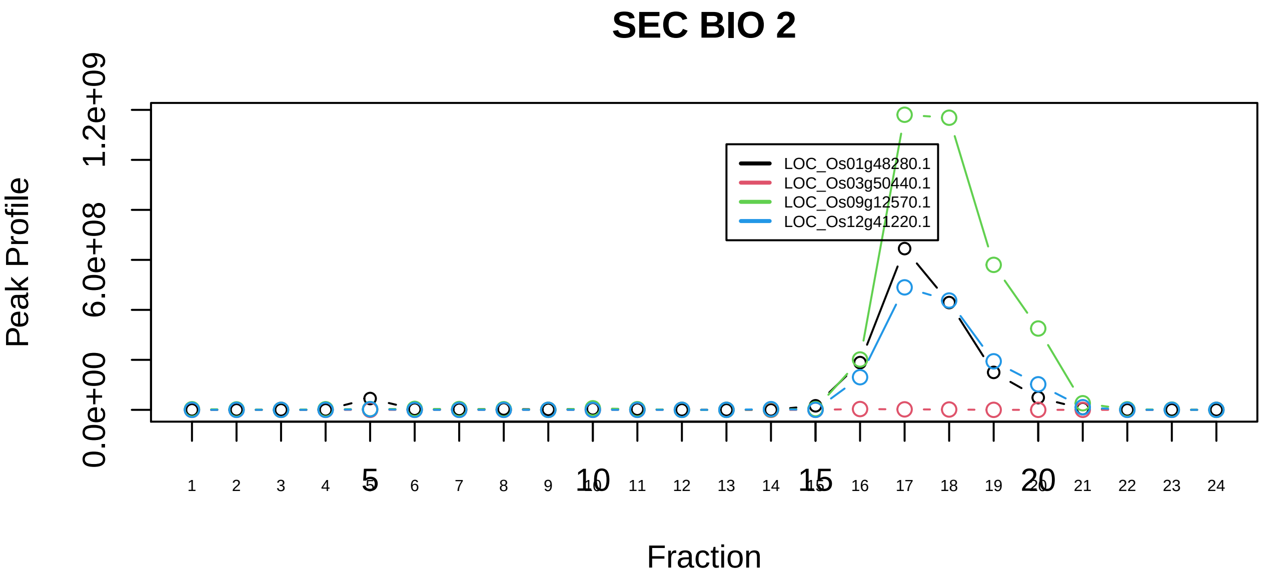

OTUB1−UBC13−MMS2 complex|7051−−sub_1

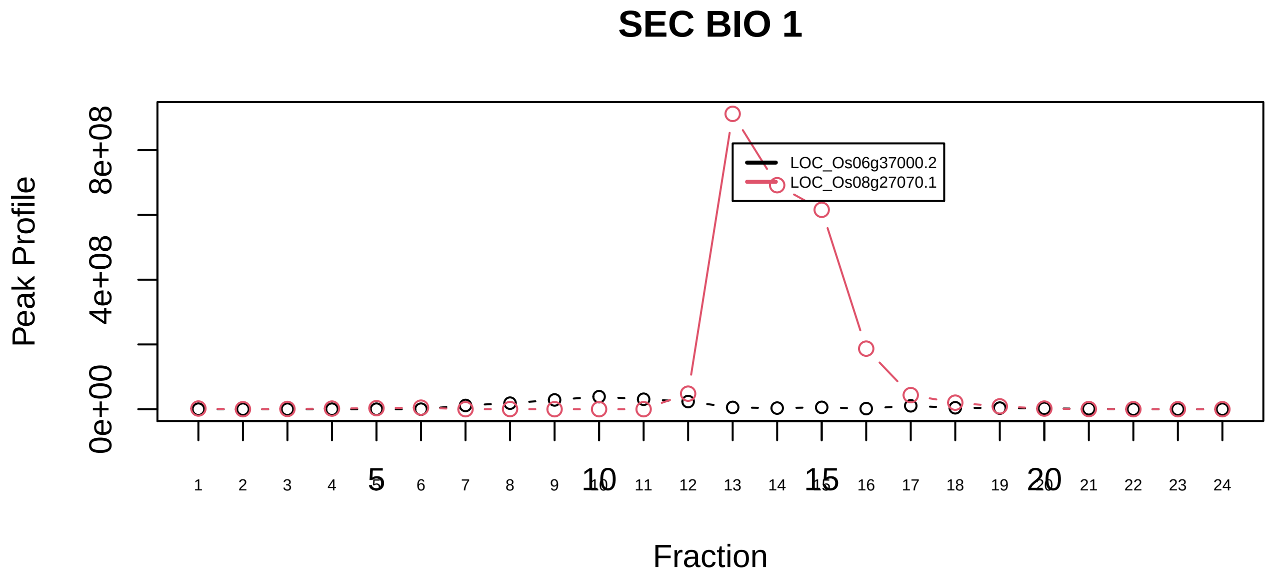

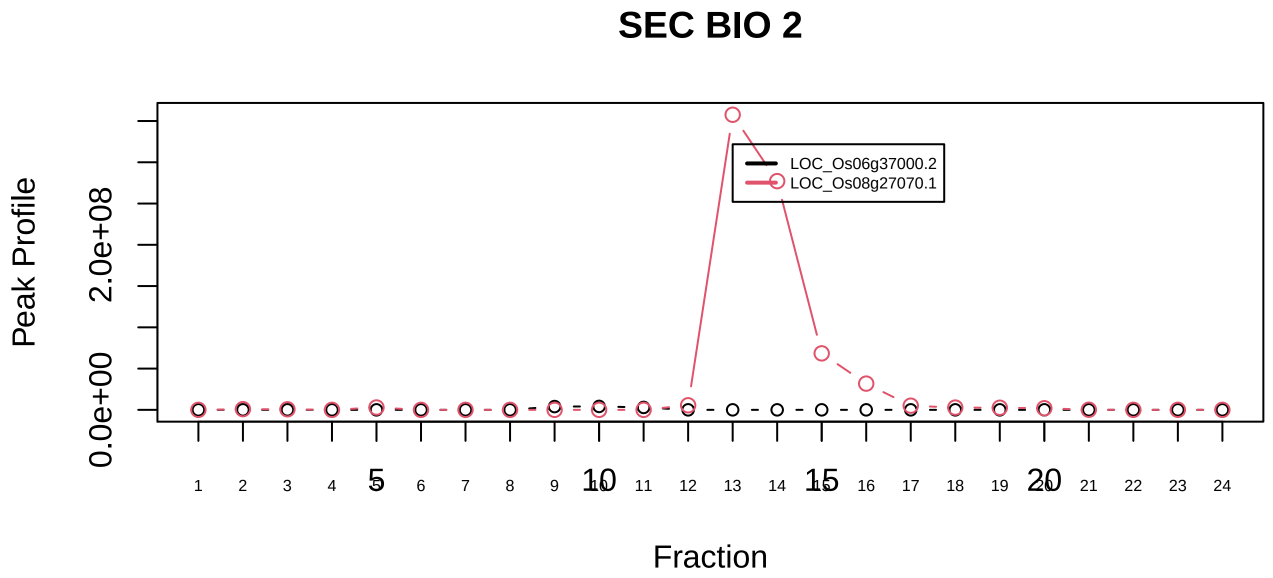

p23 protein complex|999−−only_2_prots_before_clustering

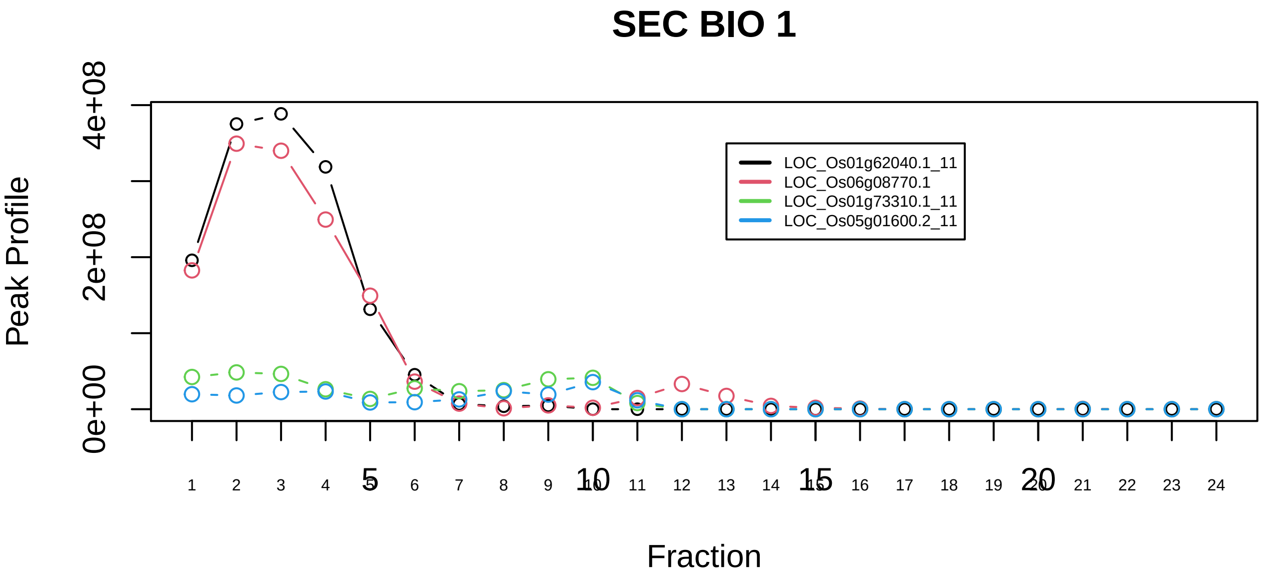

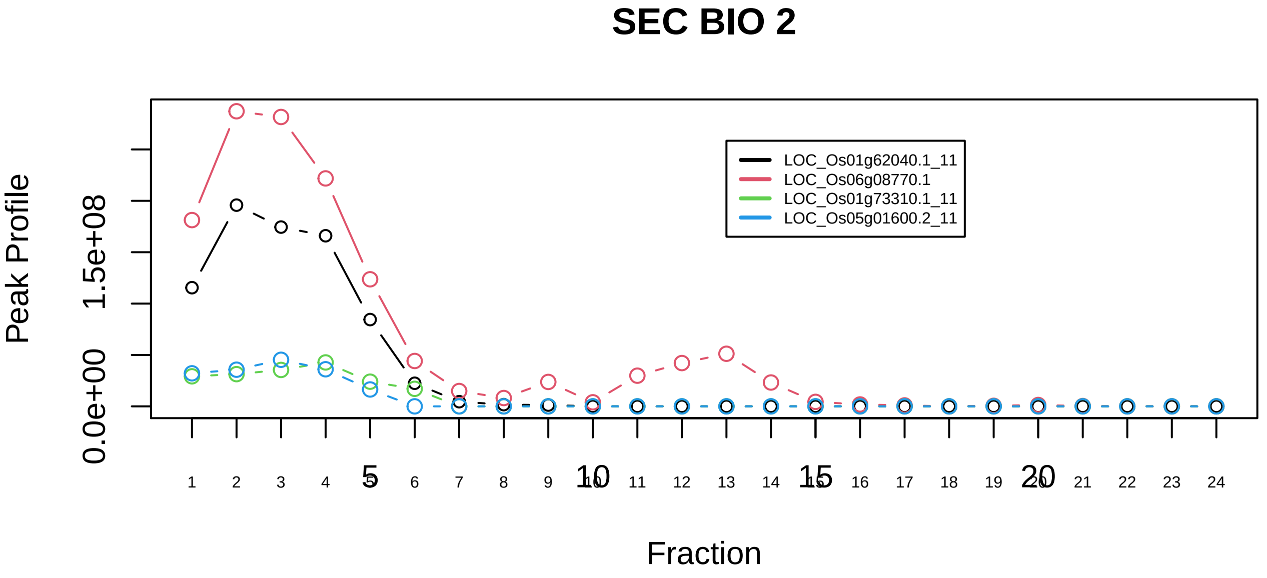

p400−associated complex|1166−−sub_1

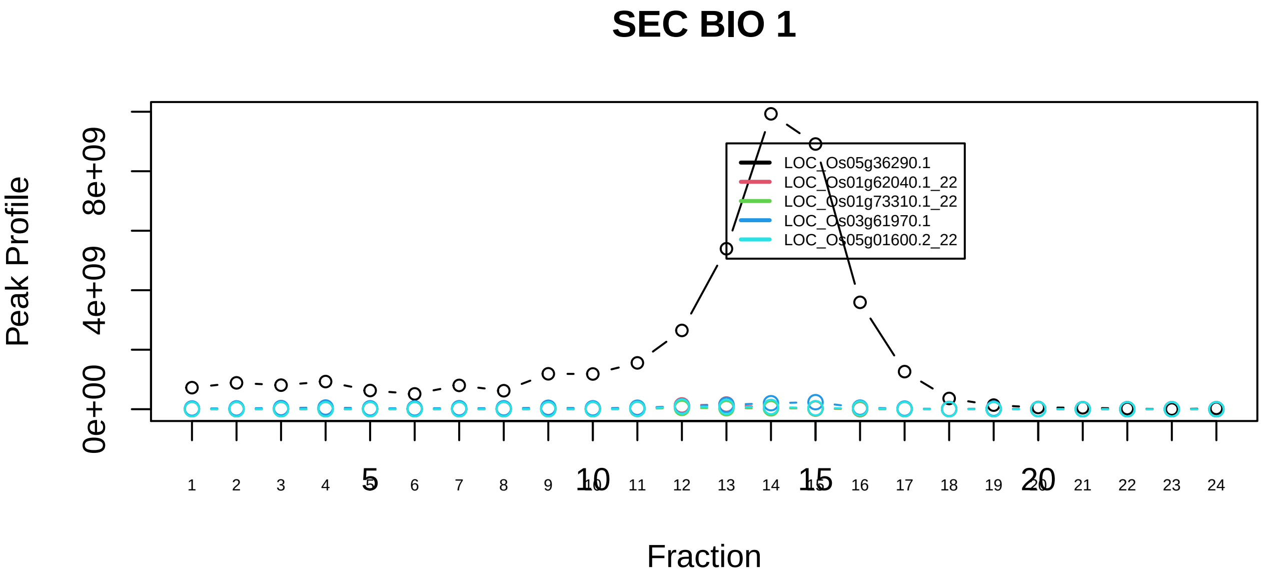

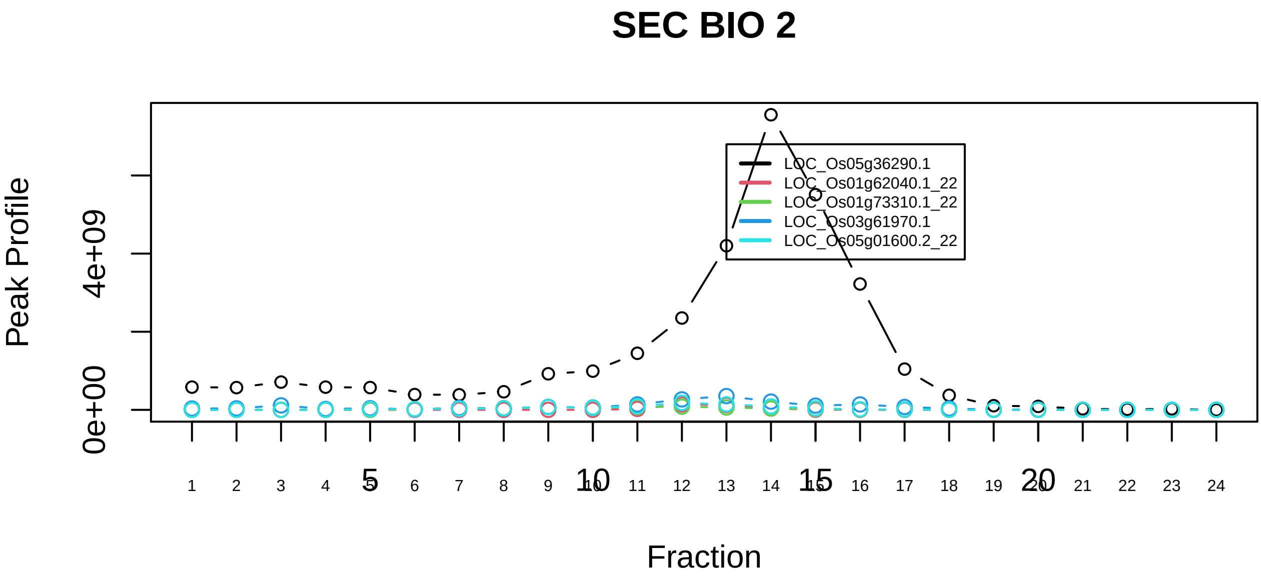

p400−associated complex|1166−−sub_2

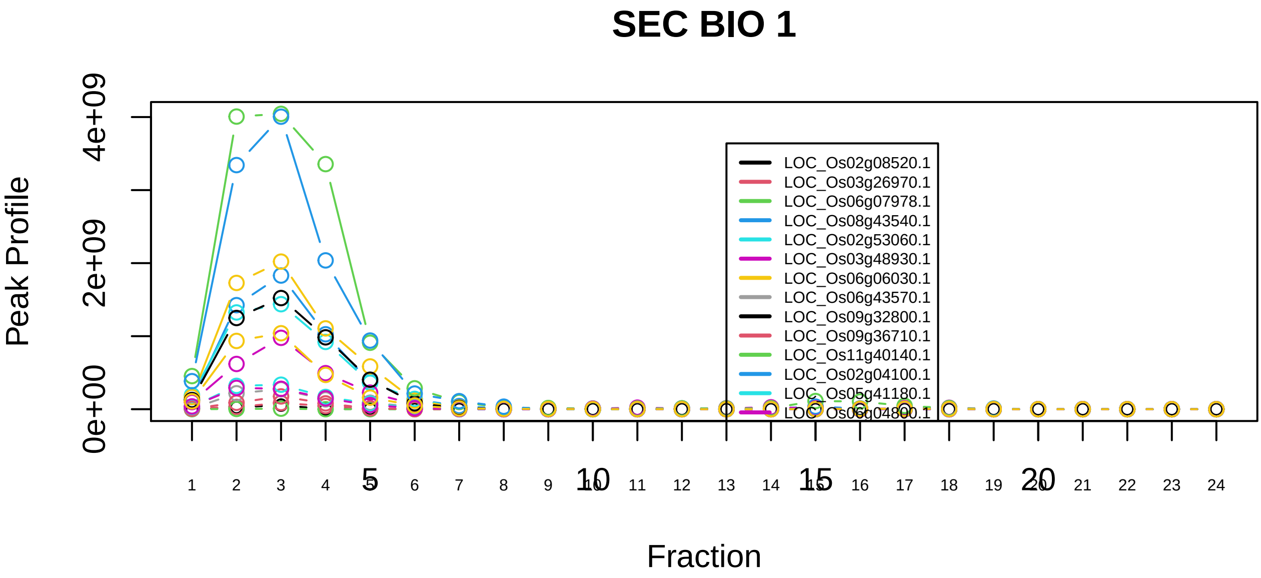

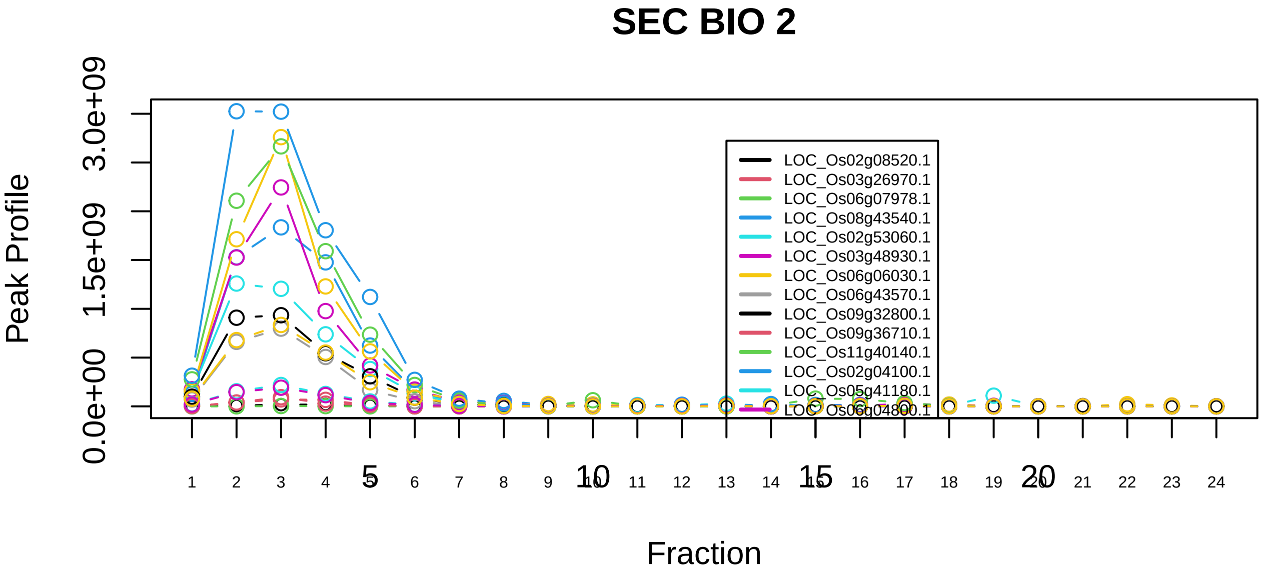

PA700−20S−PA28 complex|193−−sub_1

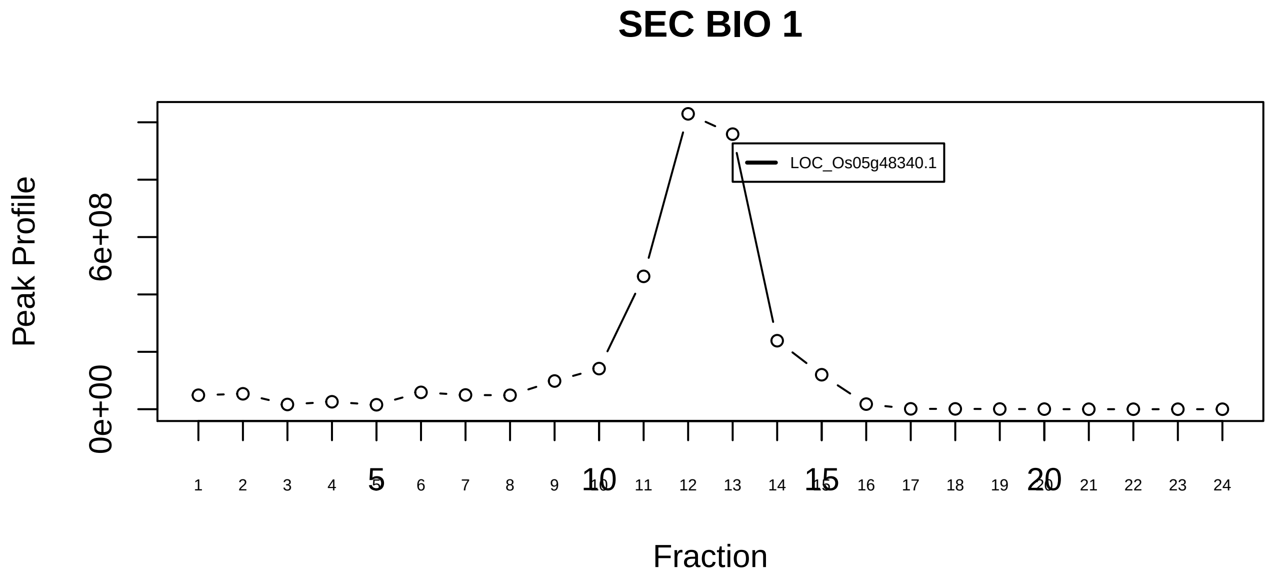

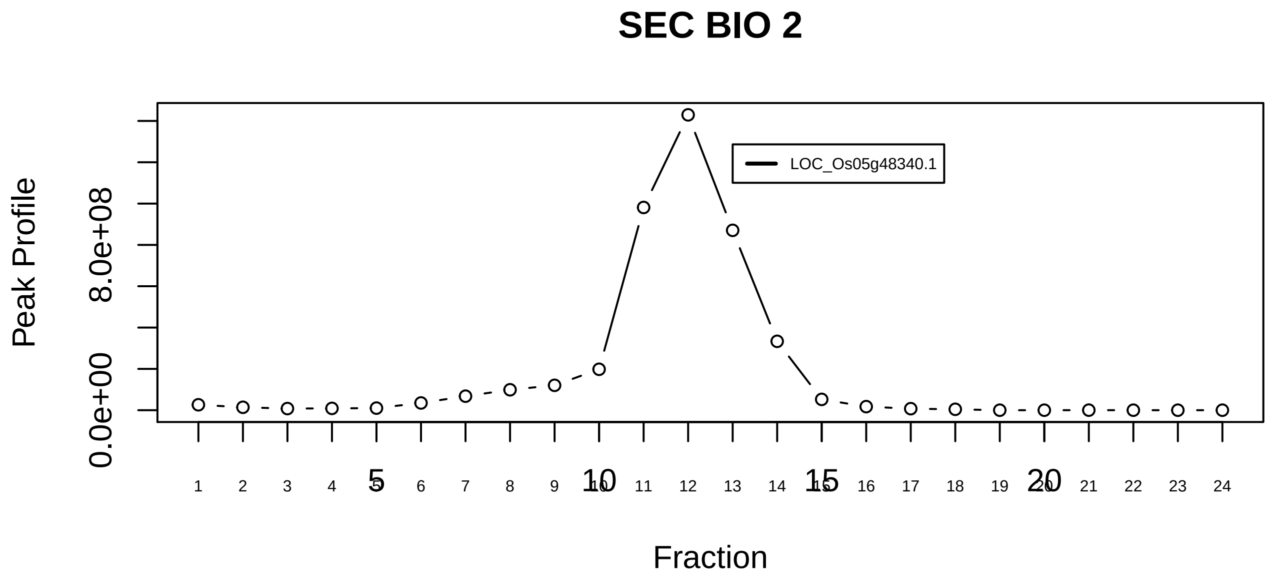

PA700−20S−PA28 complex|193−−singleton_1

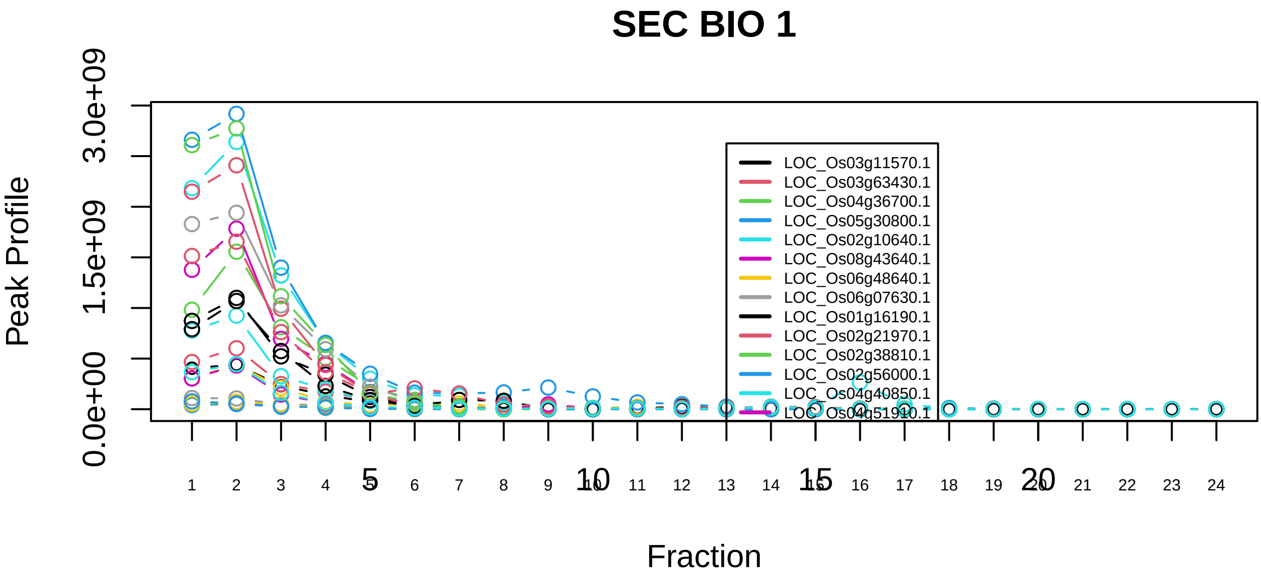

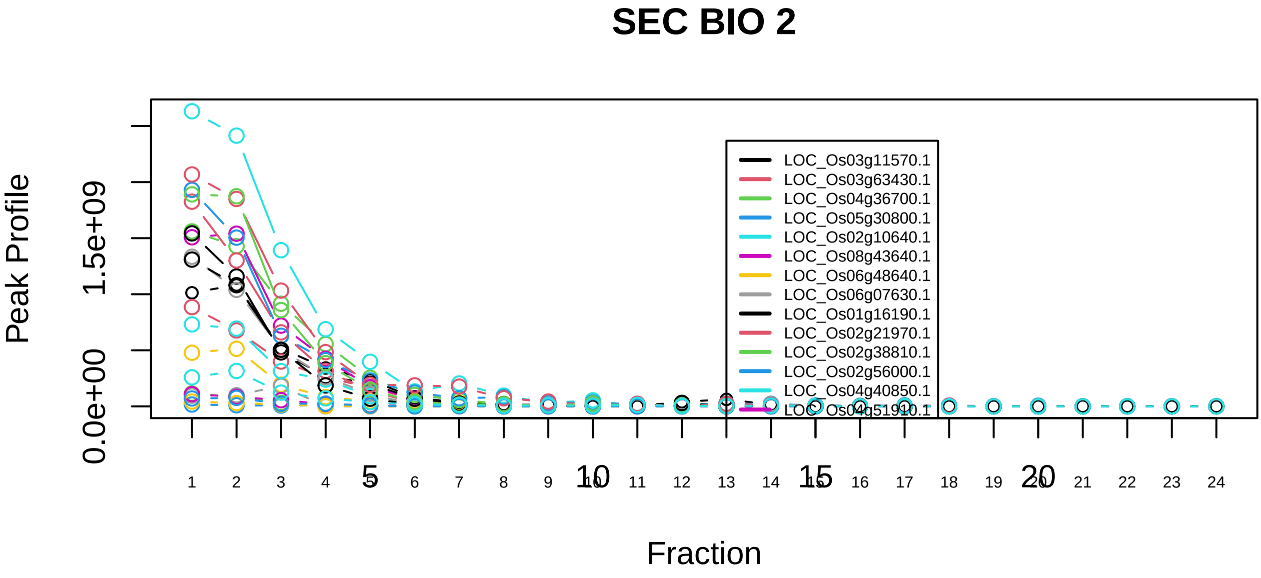

PA700−20S−PA28 complex|193−−sub_2

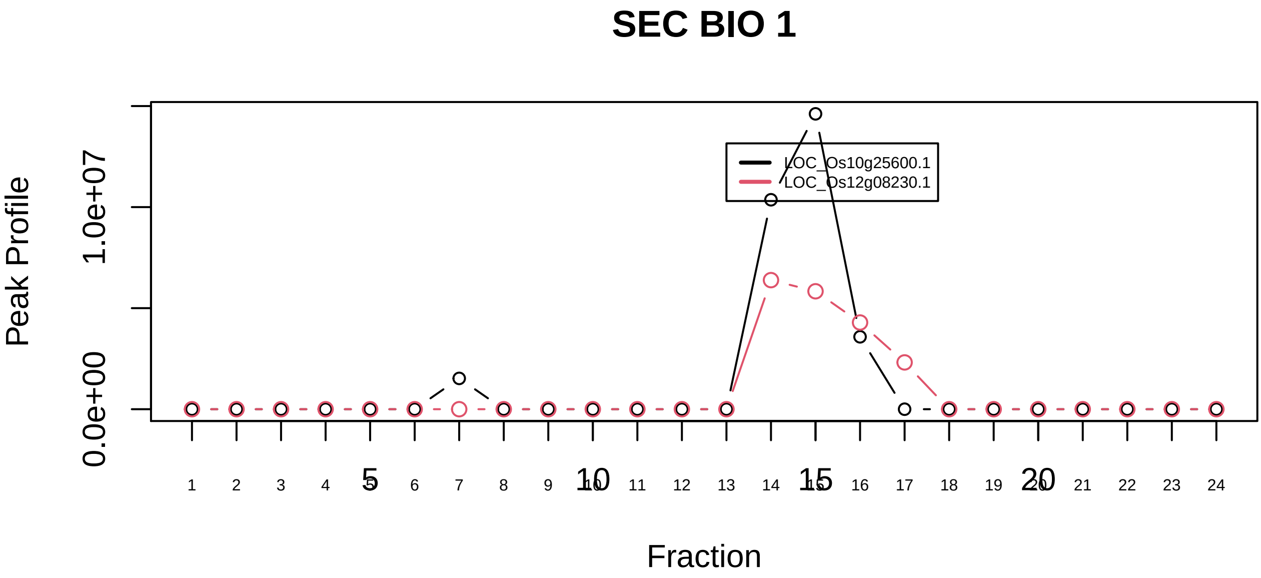

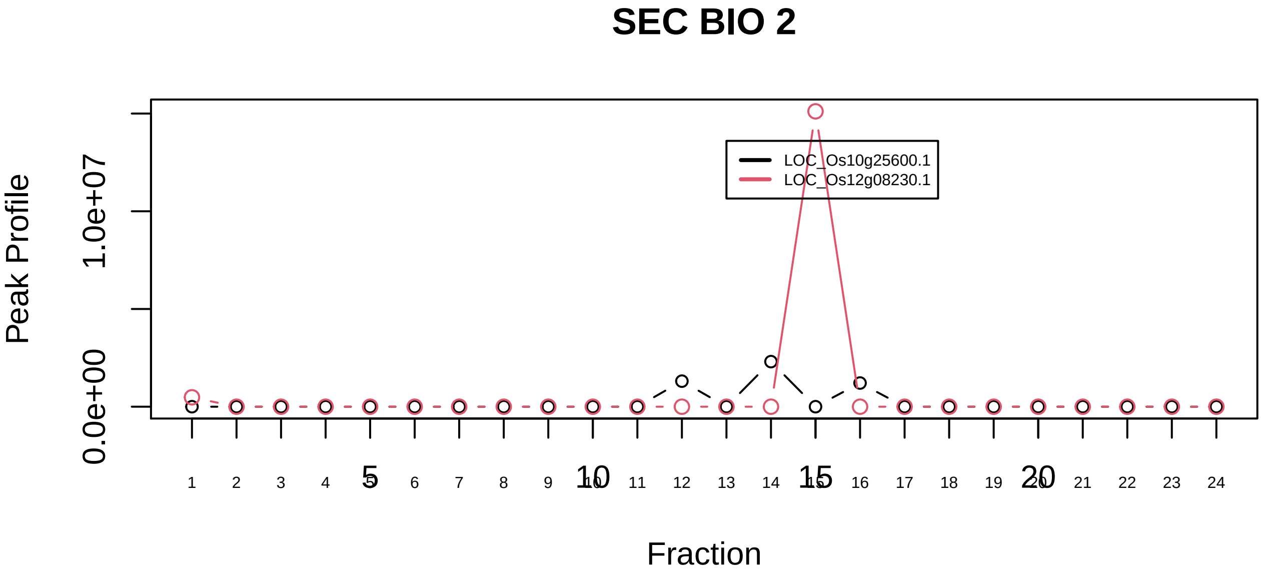

PAC3−PAC4 complex|3033−−only_2_prots_before_clustering

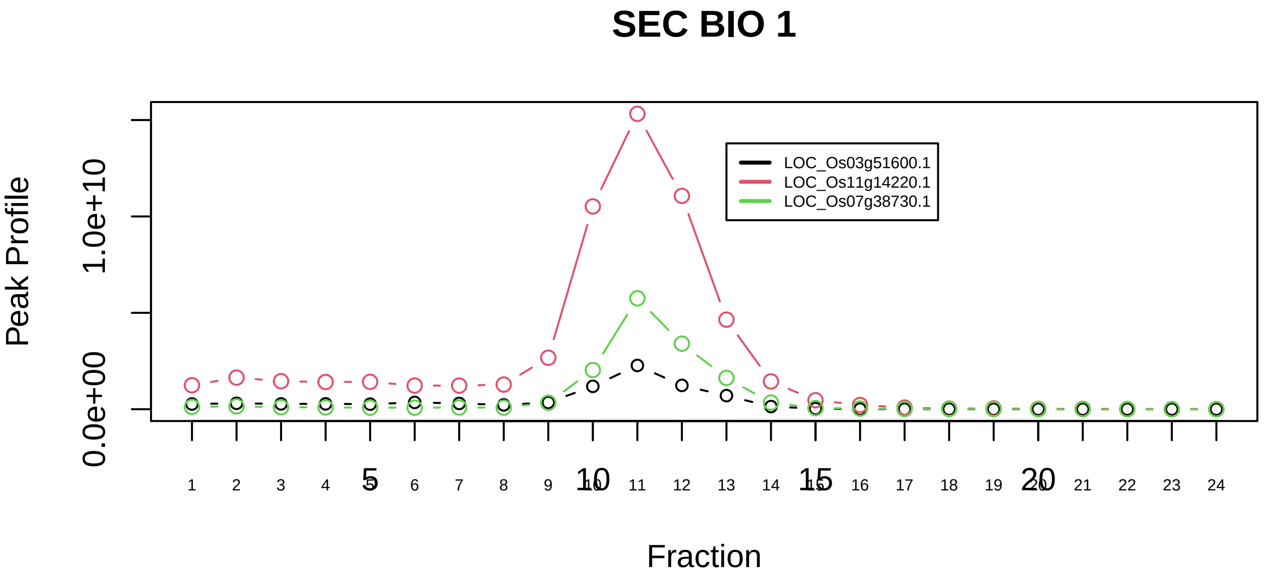

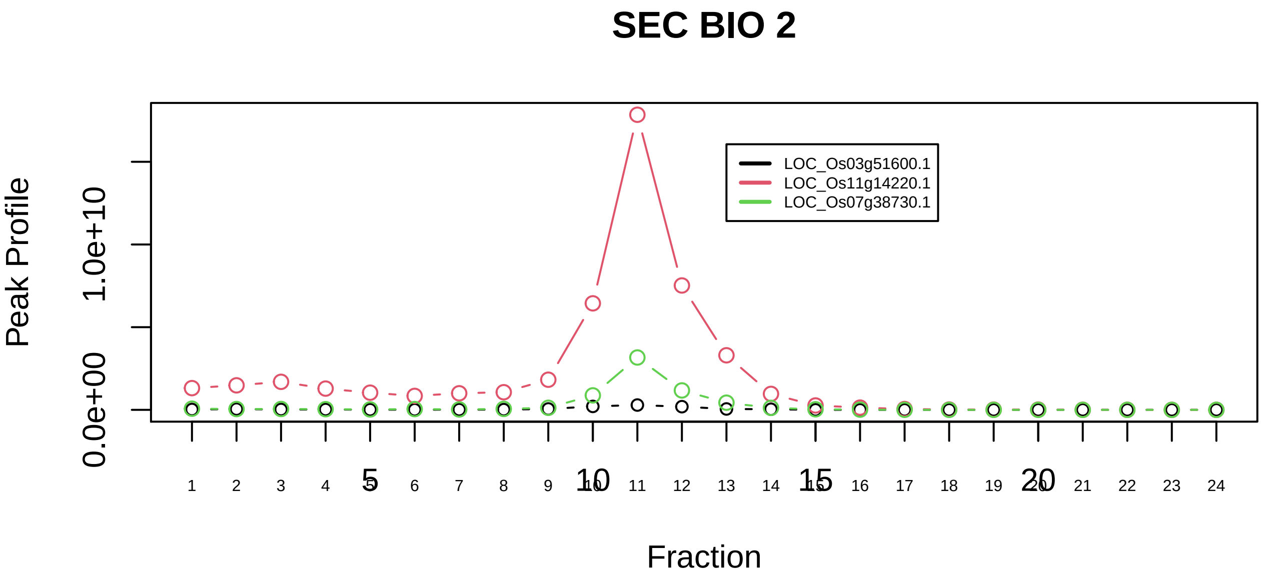

Parvulin−associated pre−rRNP complex|3047−−sub_orthoparalog_1

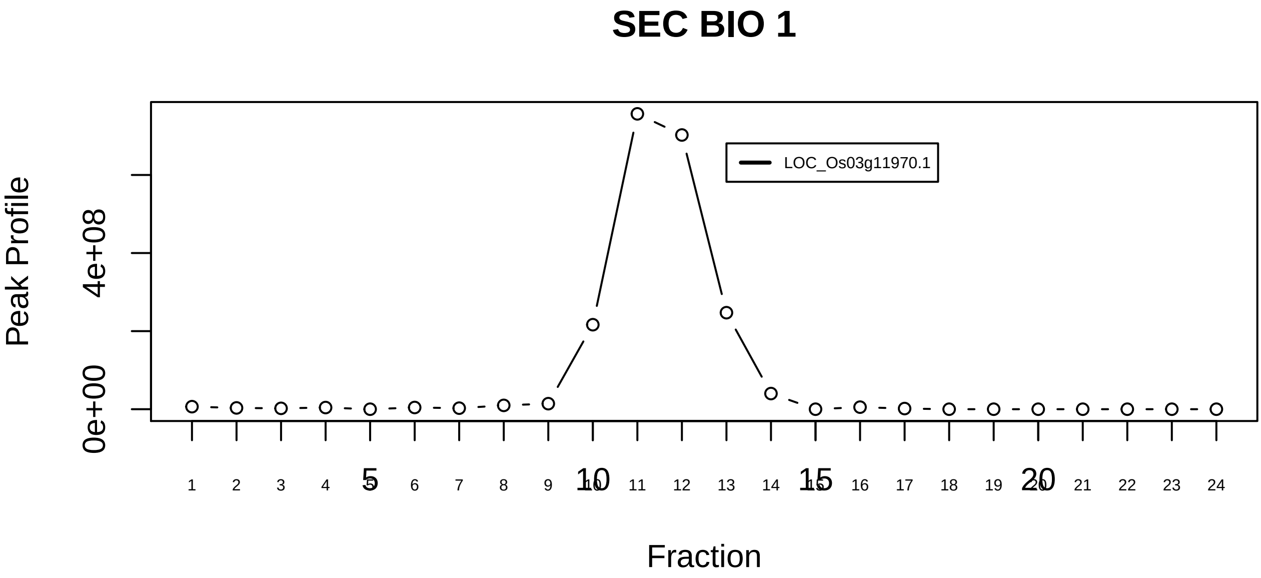

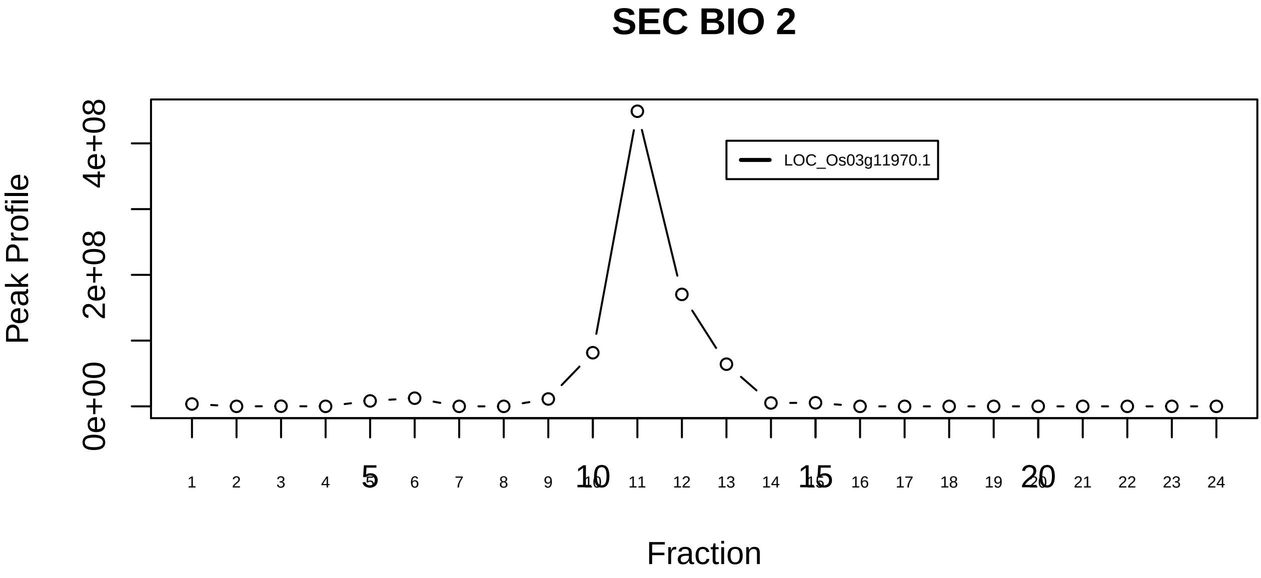

Parvulin−associated pre−rRNP complex|3047−−singleton_1

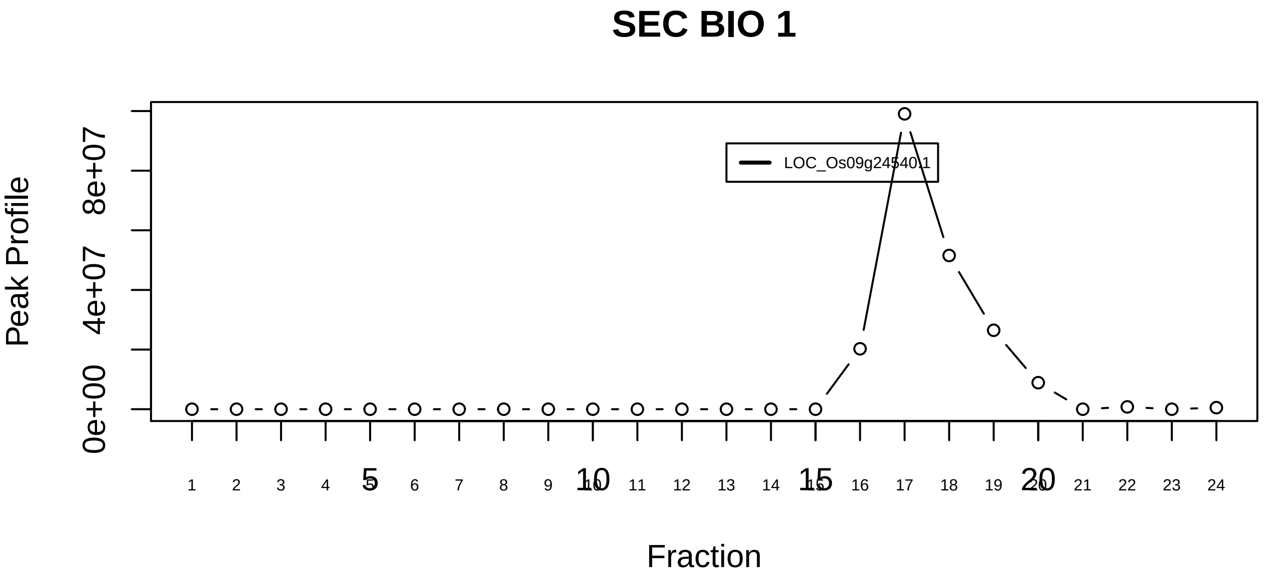

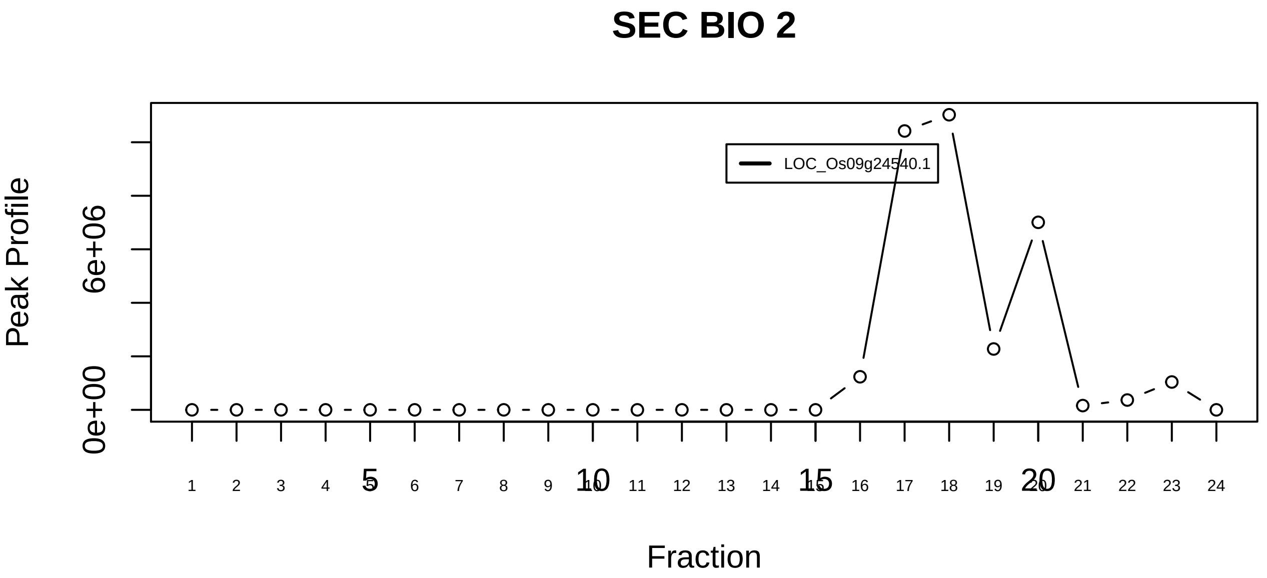

Parvulin−associated pre−rRNP complex|3047−−singleton_2

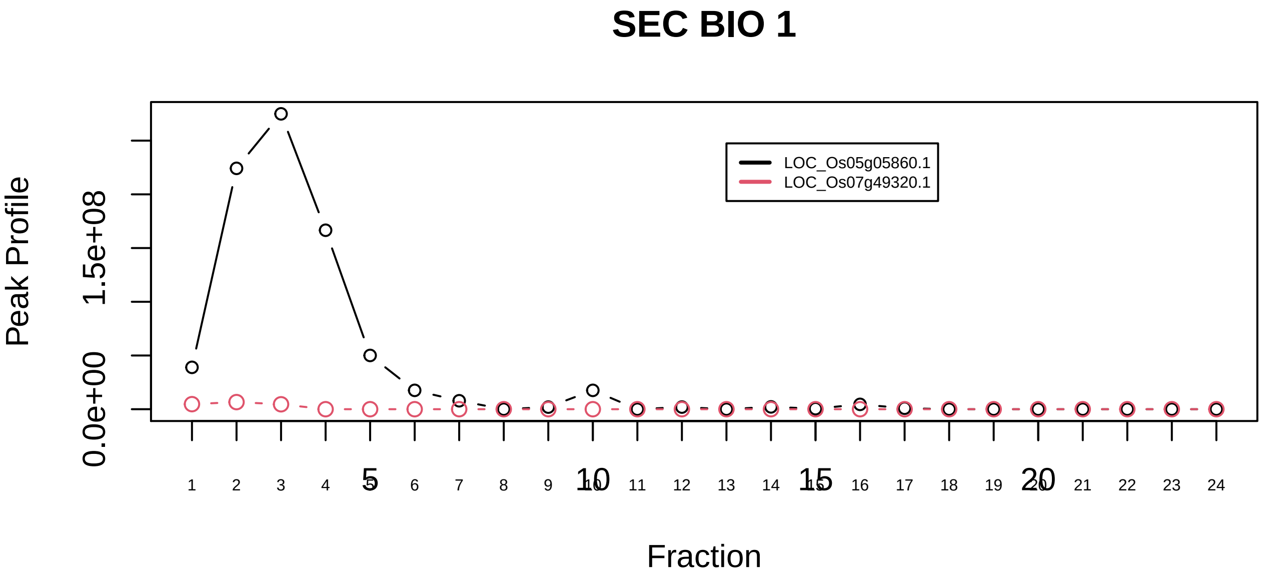

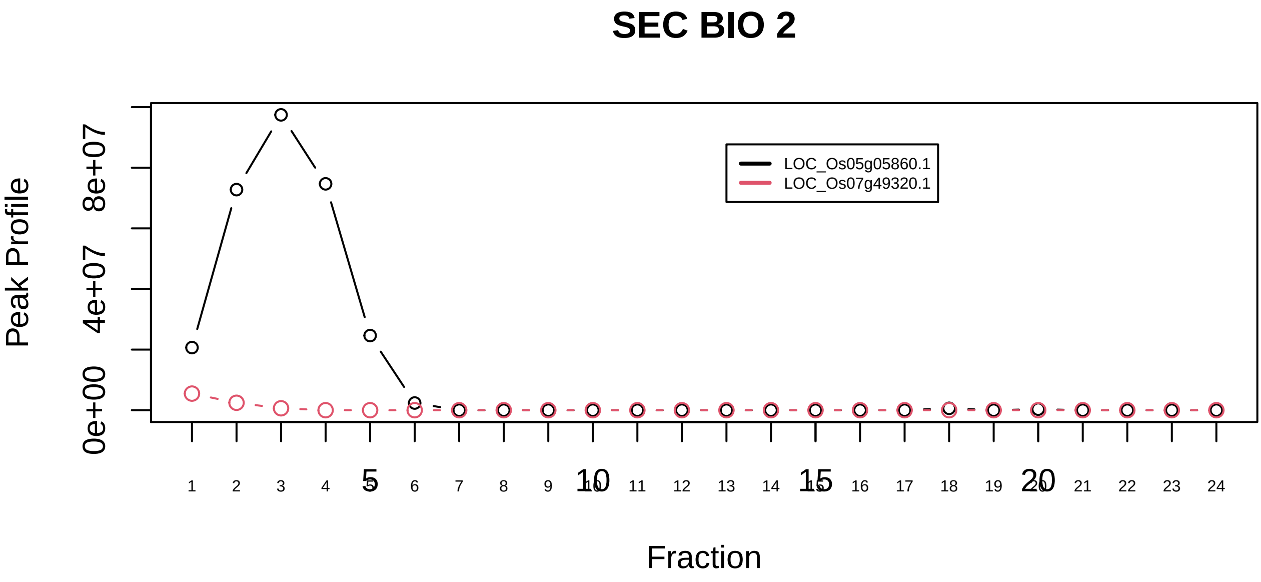

POL2R−SYMPK−SSU72 complex|6218−−only_2_prots_before_clustering

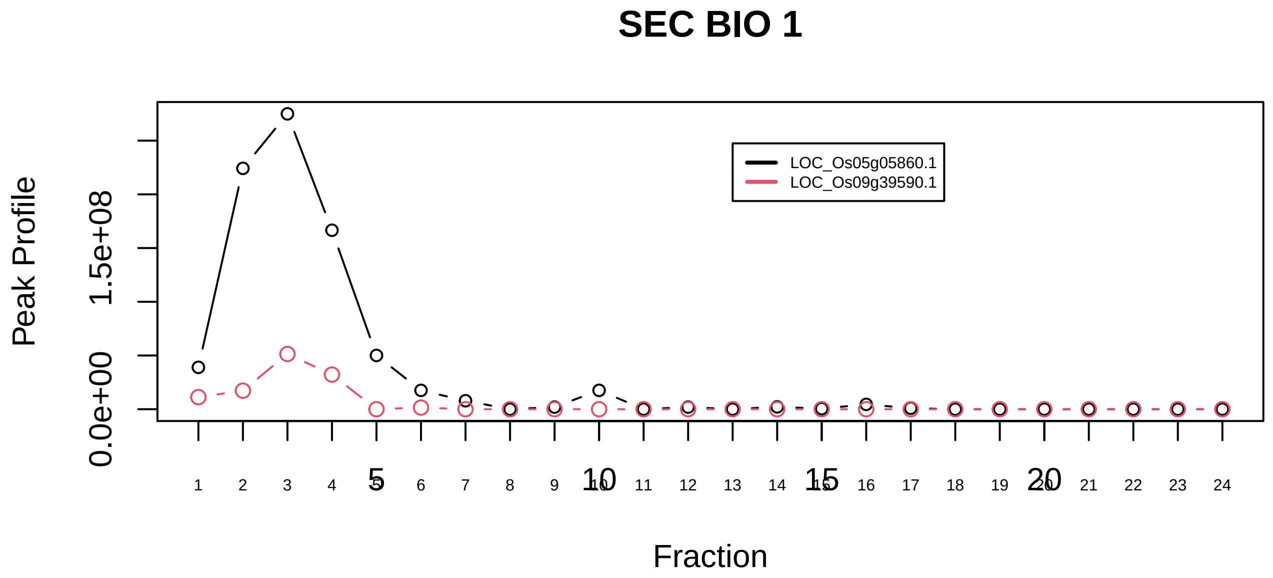

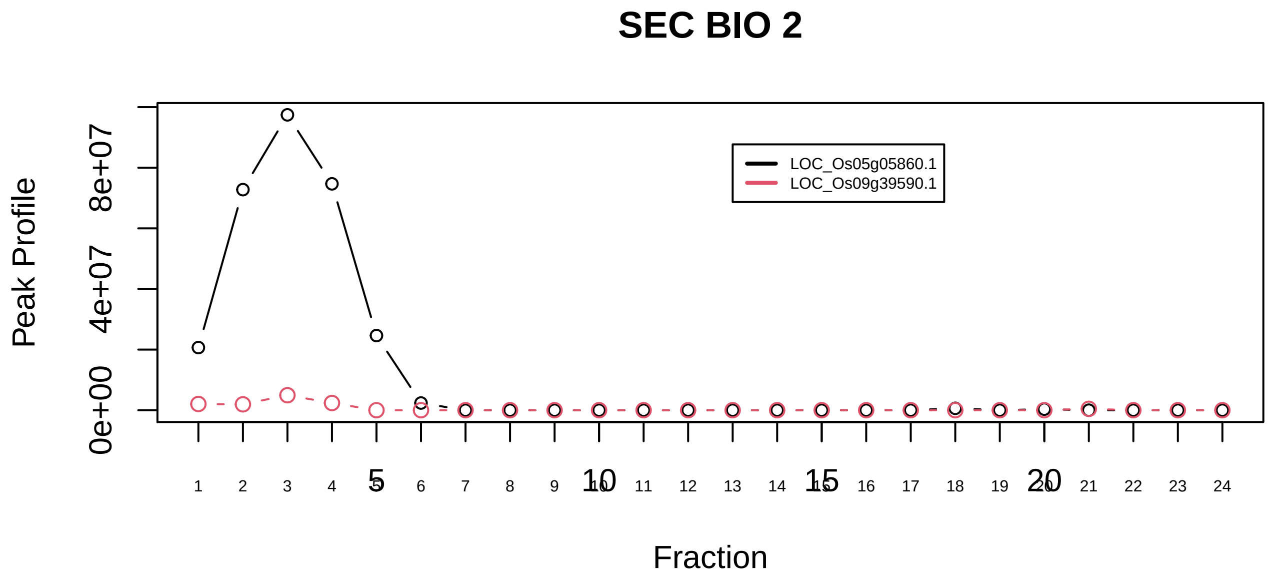

POLR2A−CCNT1−CDK9−NCL−LEM6−CPSF2 complex|2599−−only_2_prots_before_clustering

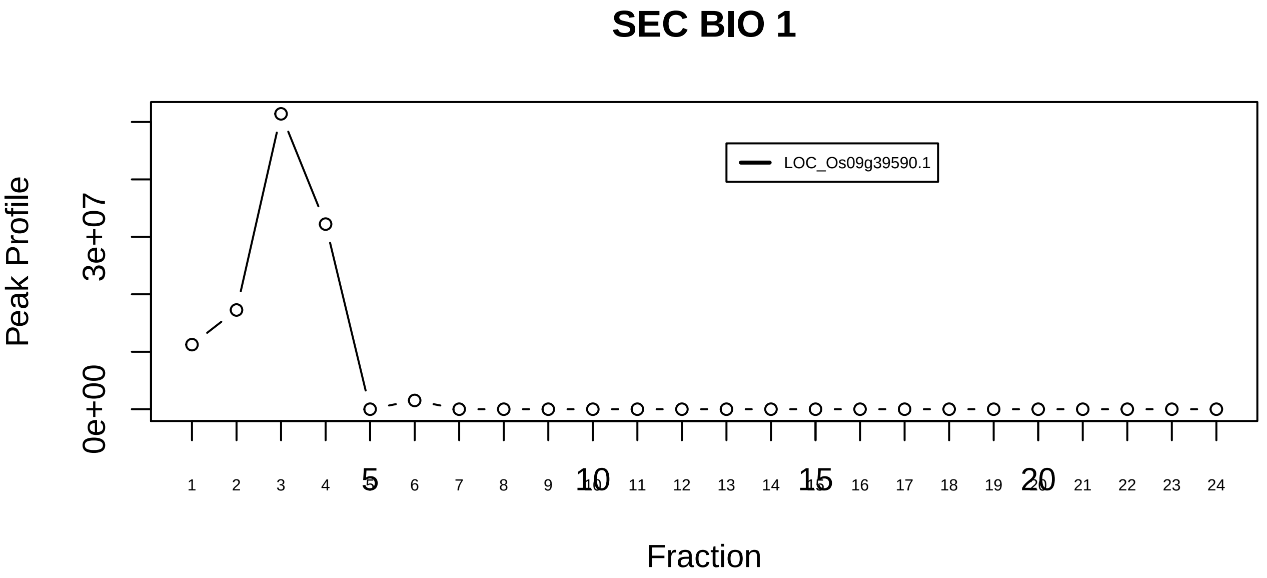

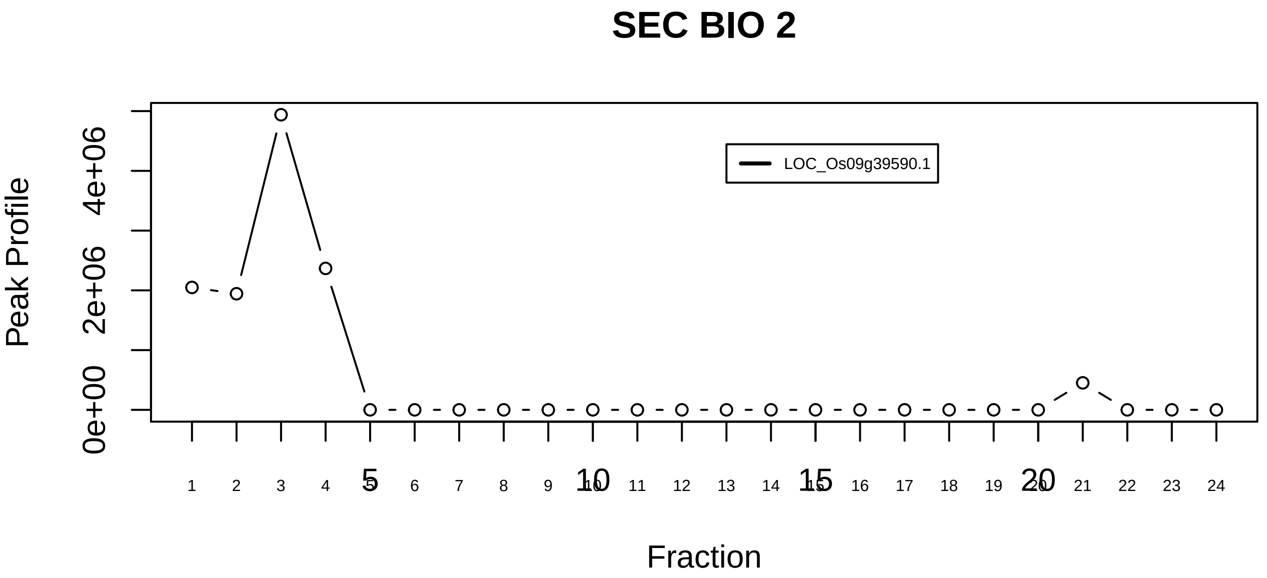

Polyadenylation complex (CSTF1, CSTF2, CSTF3, SYMPK CPSF1, CPSF2, CPSF3)|1147−−singleton_1

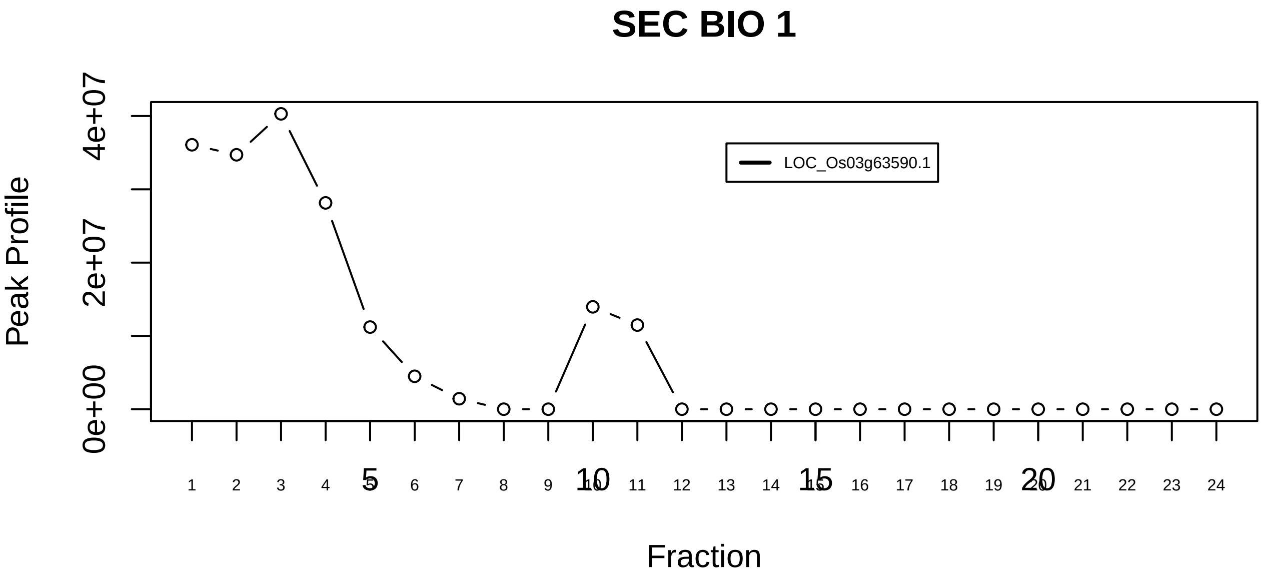

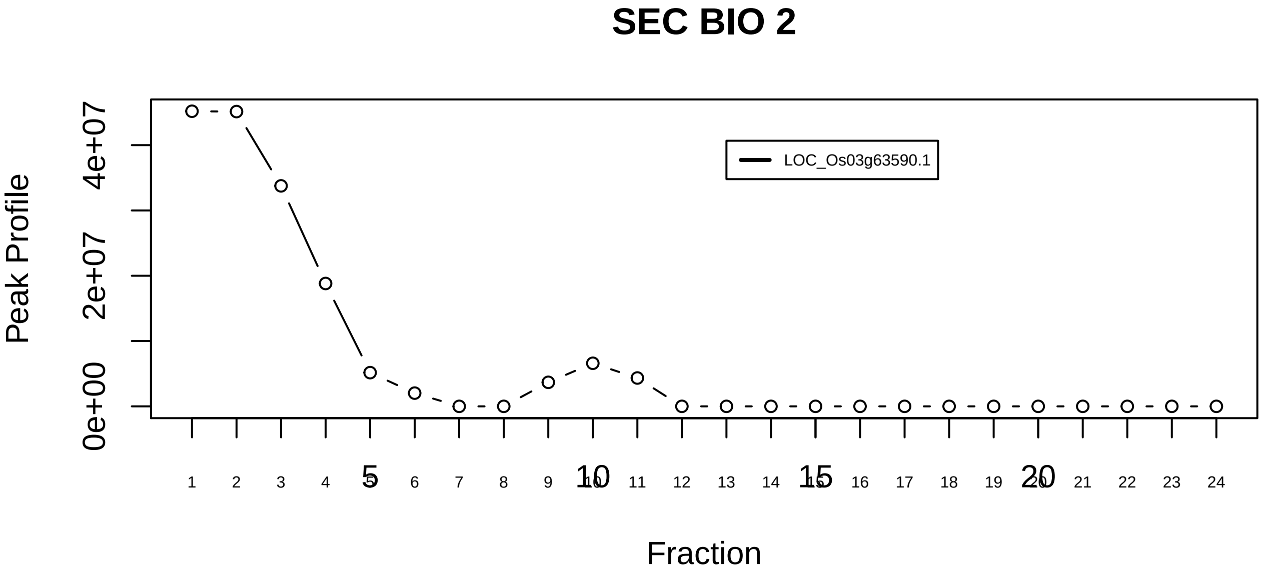

Polyadenylation complex (CSTF1, CSTF2, CSTF3, SYMPK CPSF1, CPSF2, CPSF3)|1147−−singleton_2

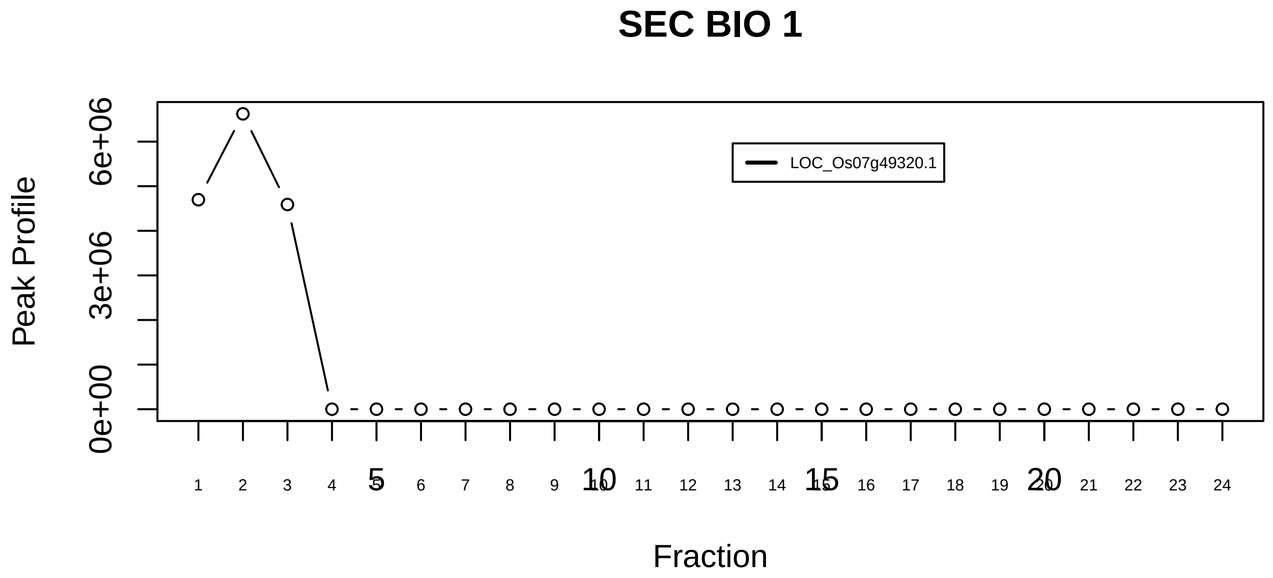

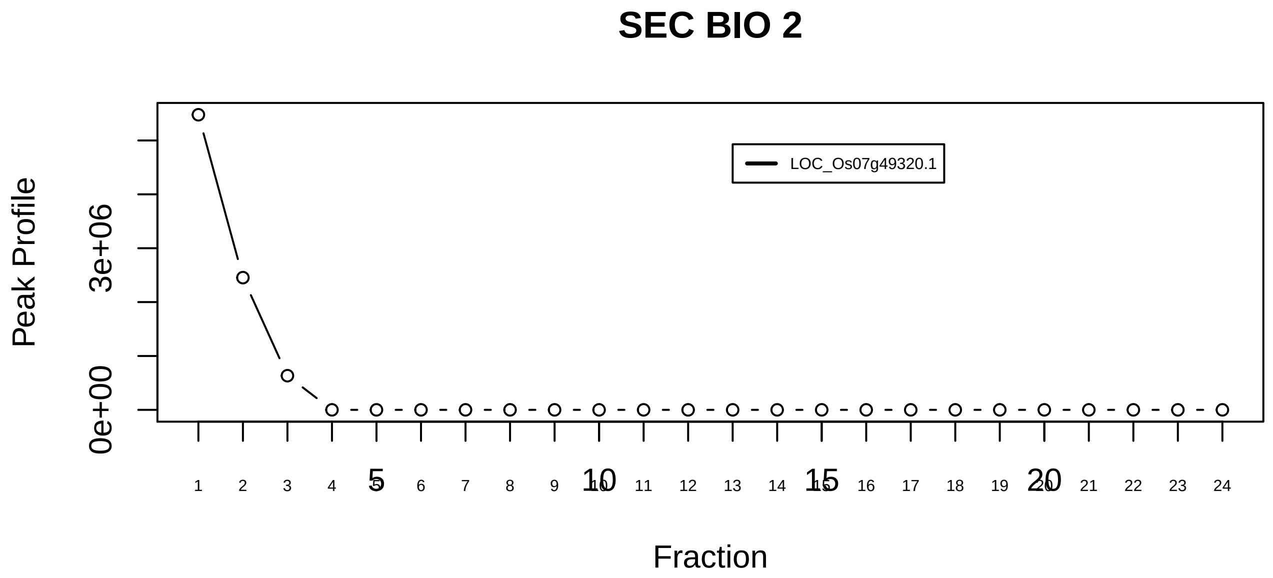

Polyadenylation complex (CSTF1, CSTF2, CSTF3, SYMPK CPSF1, CPSF2, CPSF3)|1147−−singleton_3

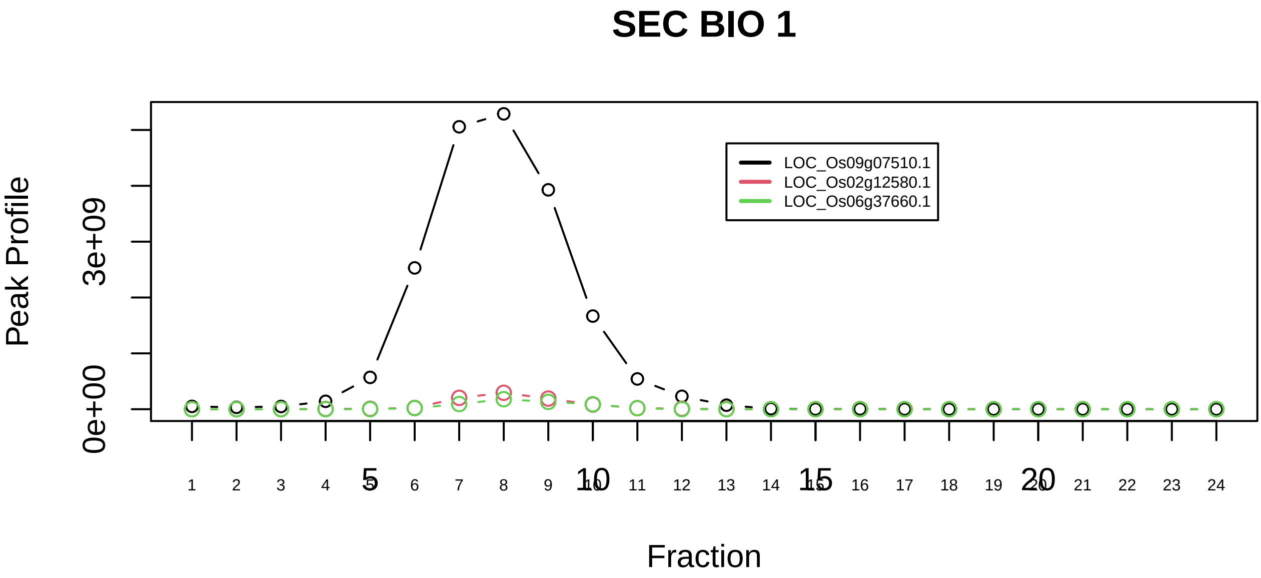

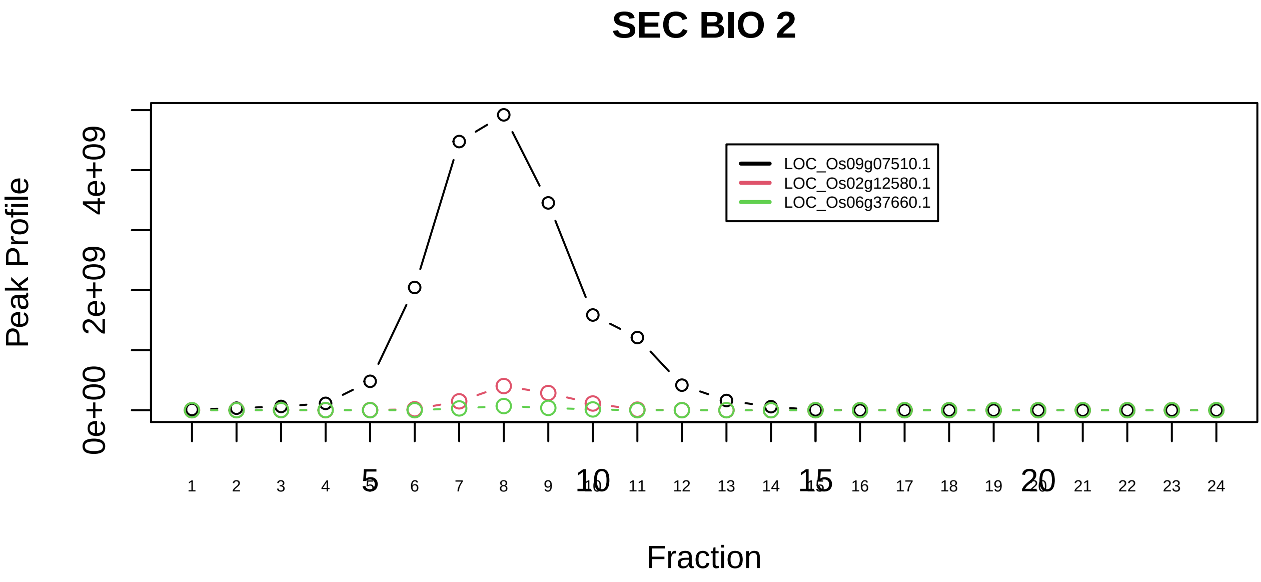

PPP2R1A−PPP2R1B−PPP2CA−PPME1−EIF4A1 complex|5876−−sub_1

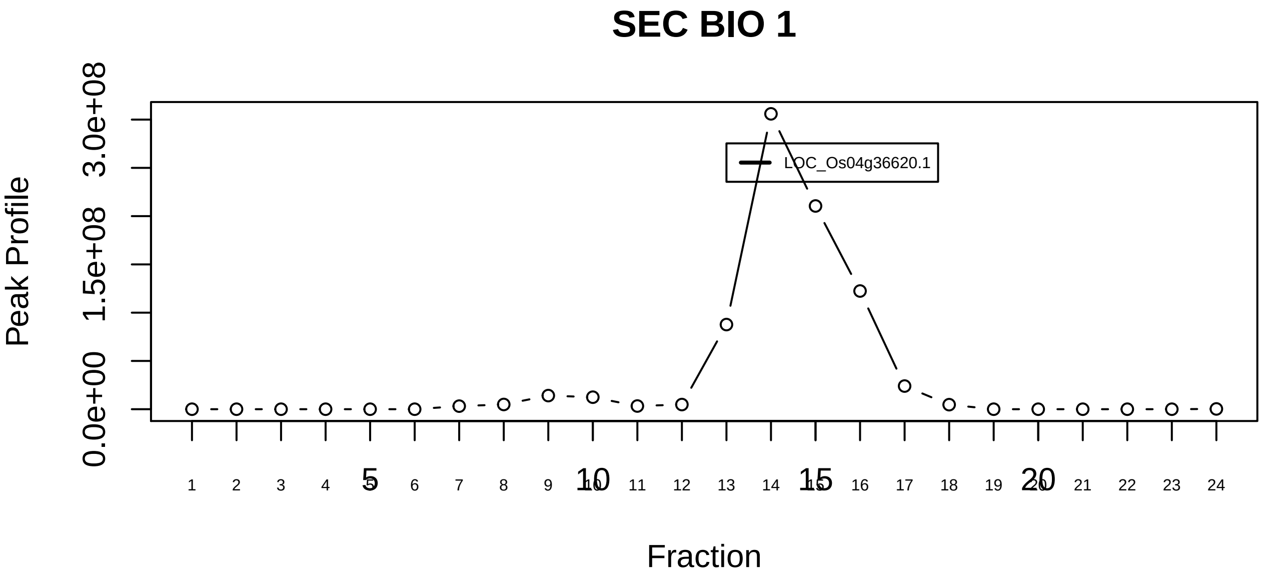

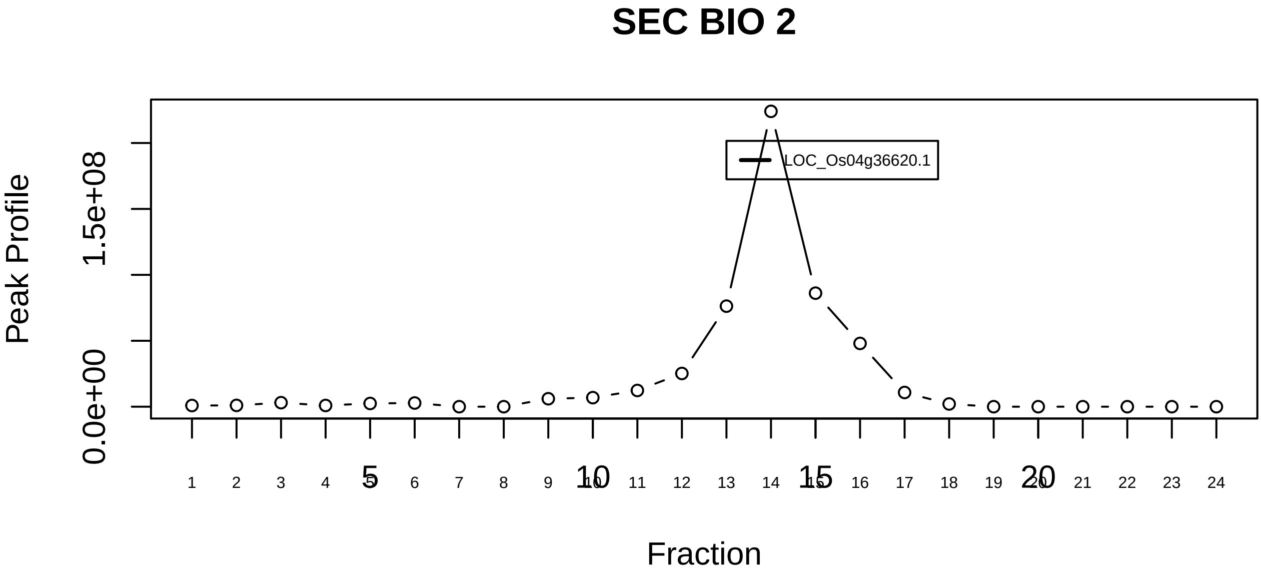

PPP2R1A−PPP2R1B−PPP2CA−PPME1−EIF4A1 complex|5876−−singleton_1

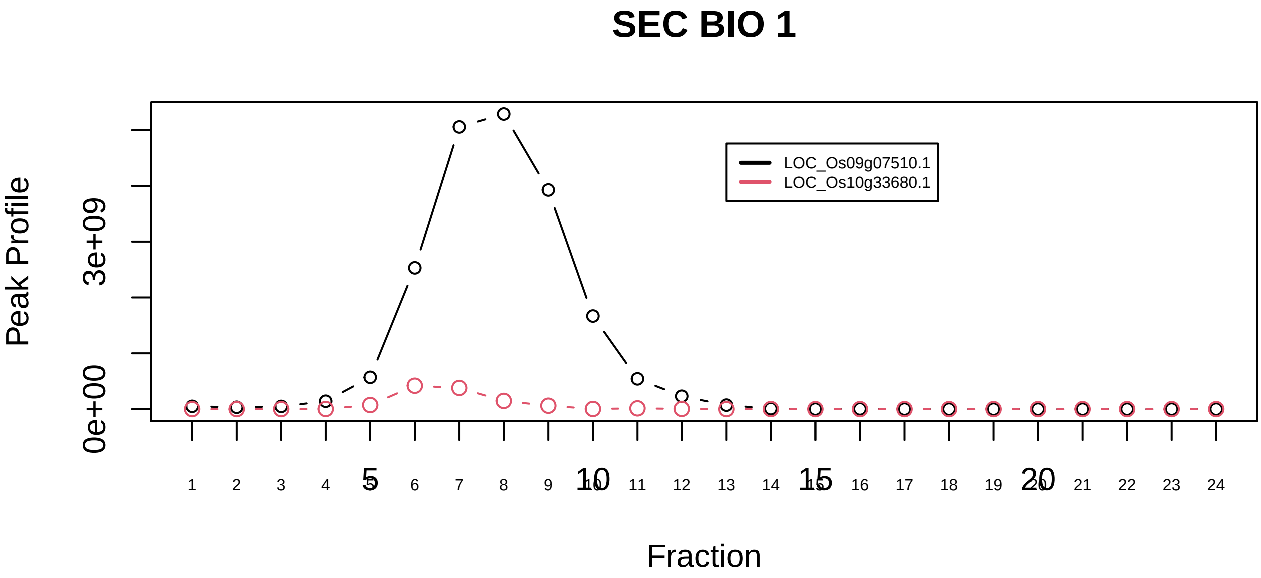

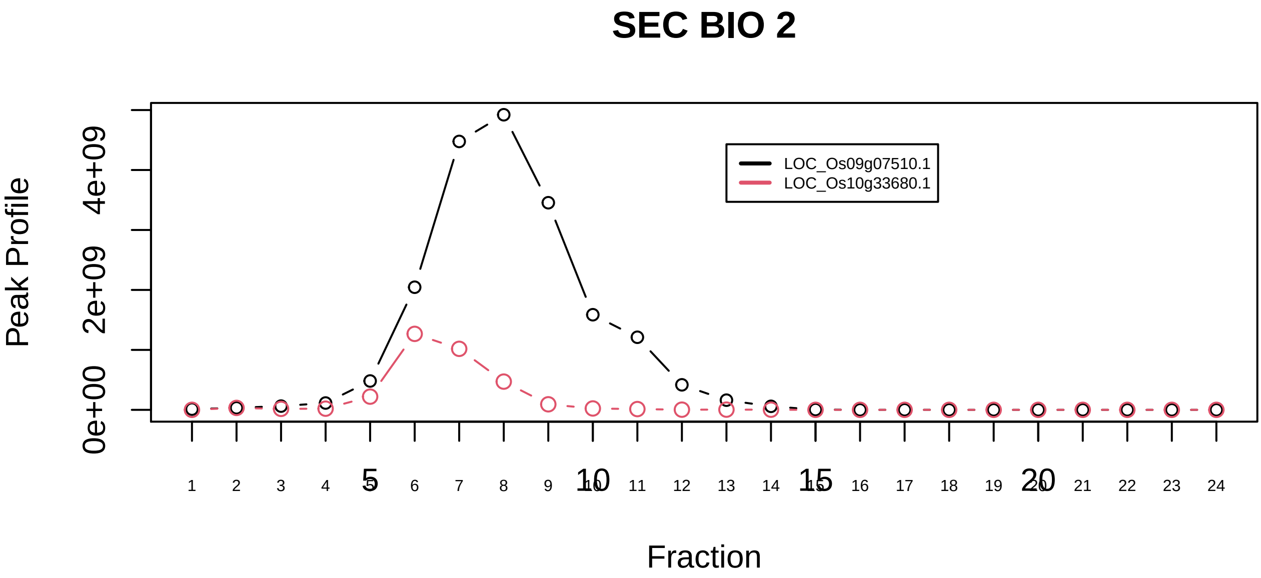

PPP2R1A−PPP2R3B complex|6180−−only_2_prots_before_clustering

### SEC BIO 1

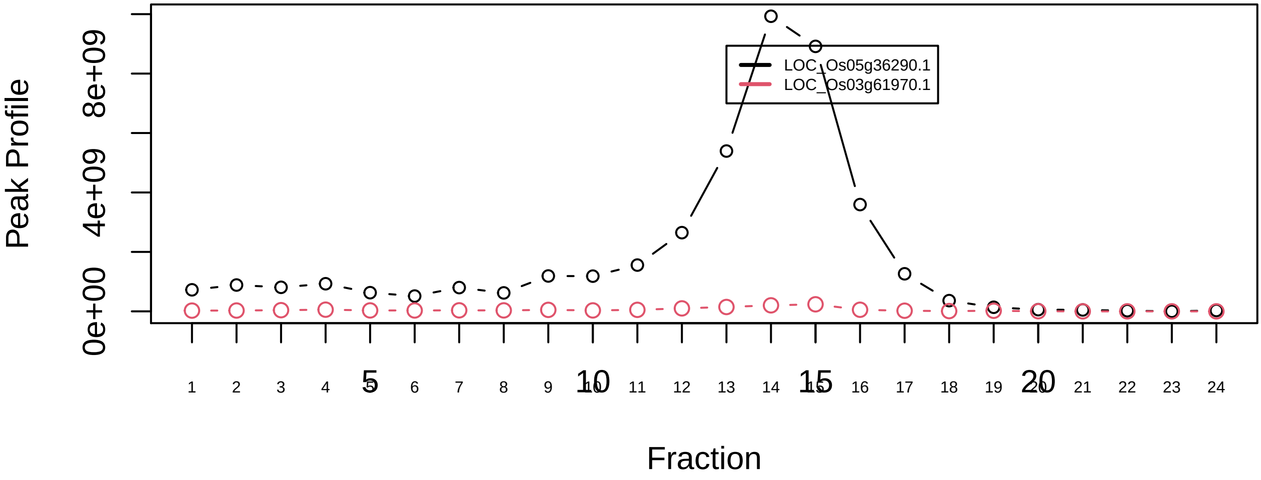

### SEC BIO 2

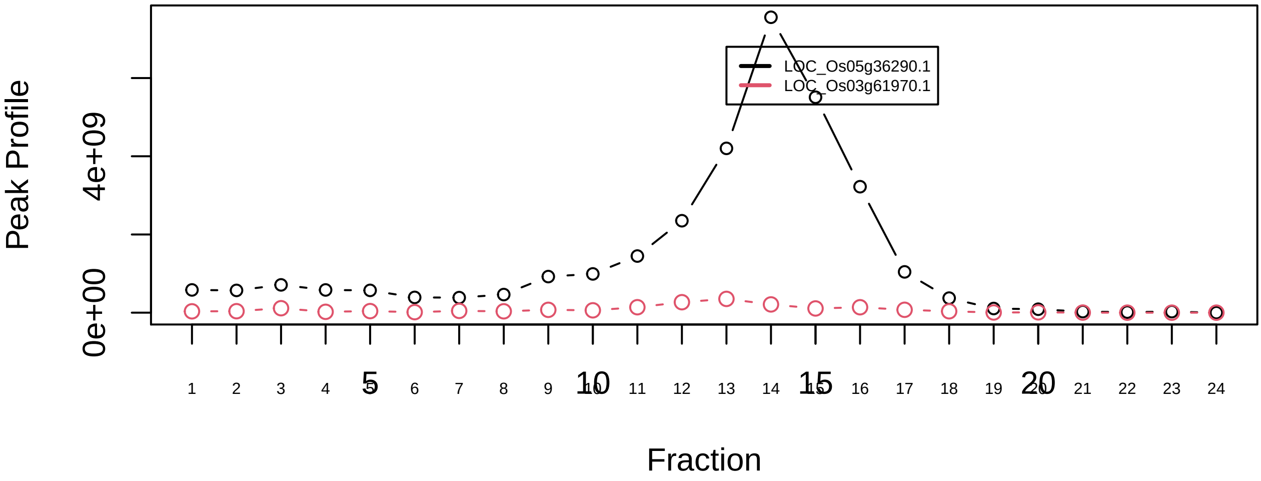

Pre−initiation complex (PIC)|5736−−sub_orthoparalog_1

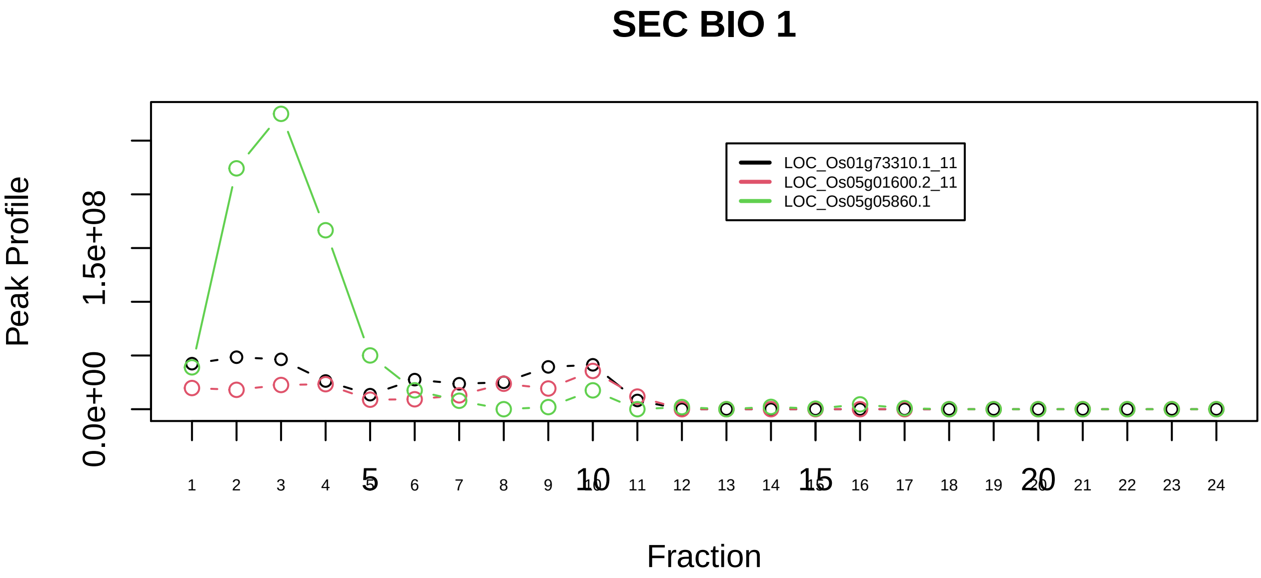

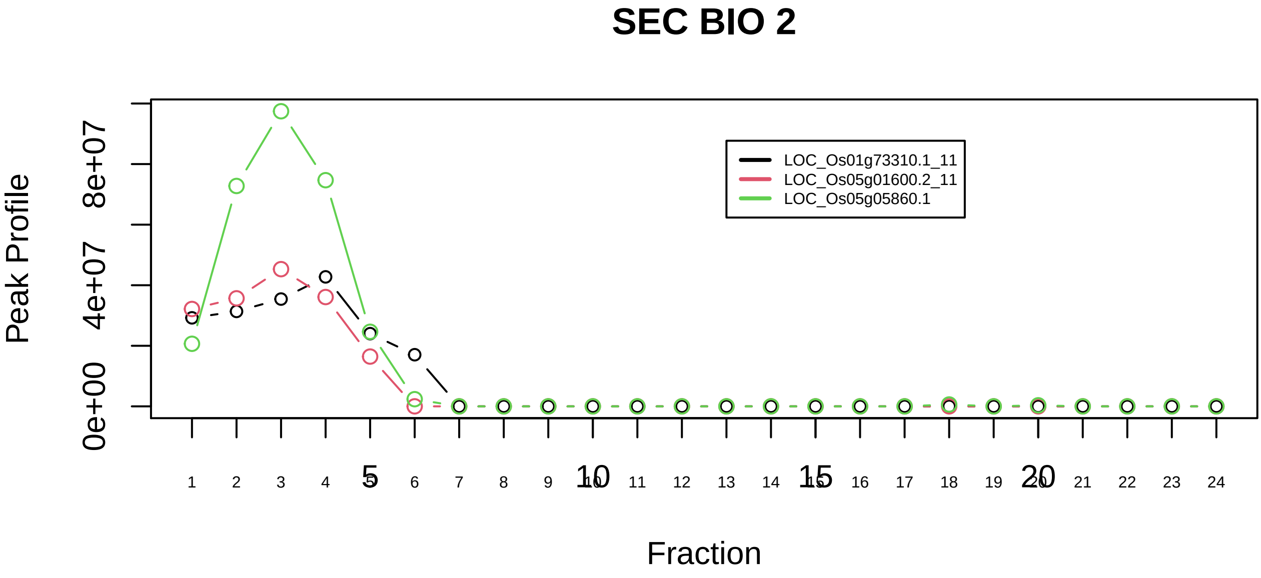

Pre−initiation complex (PIC)|5736−−sub_1

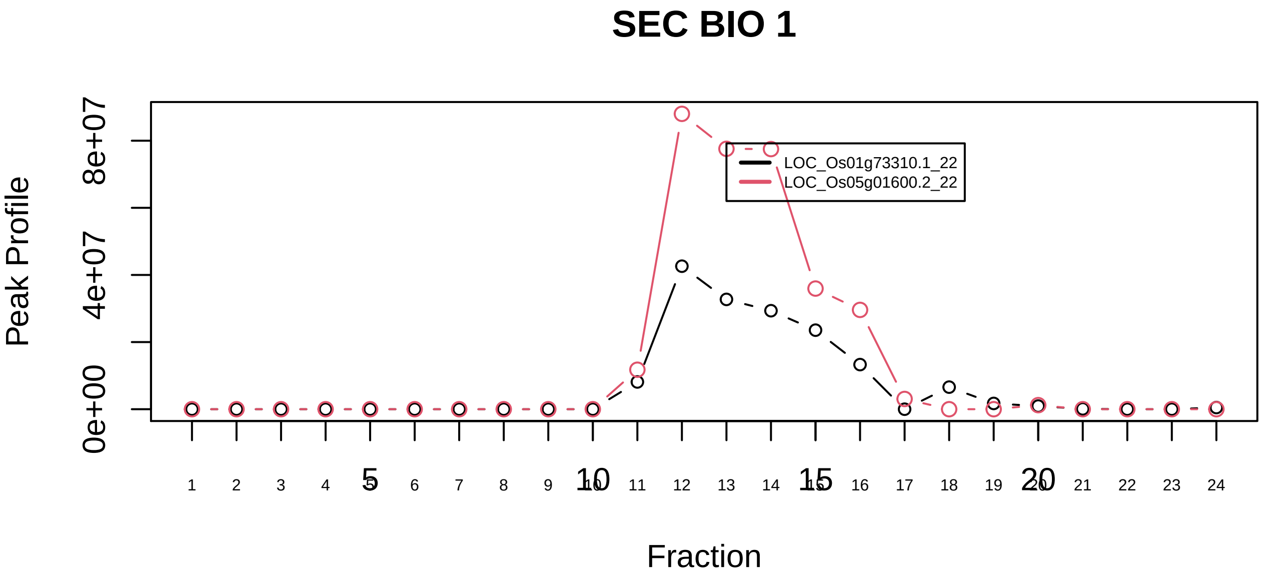

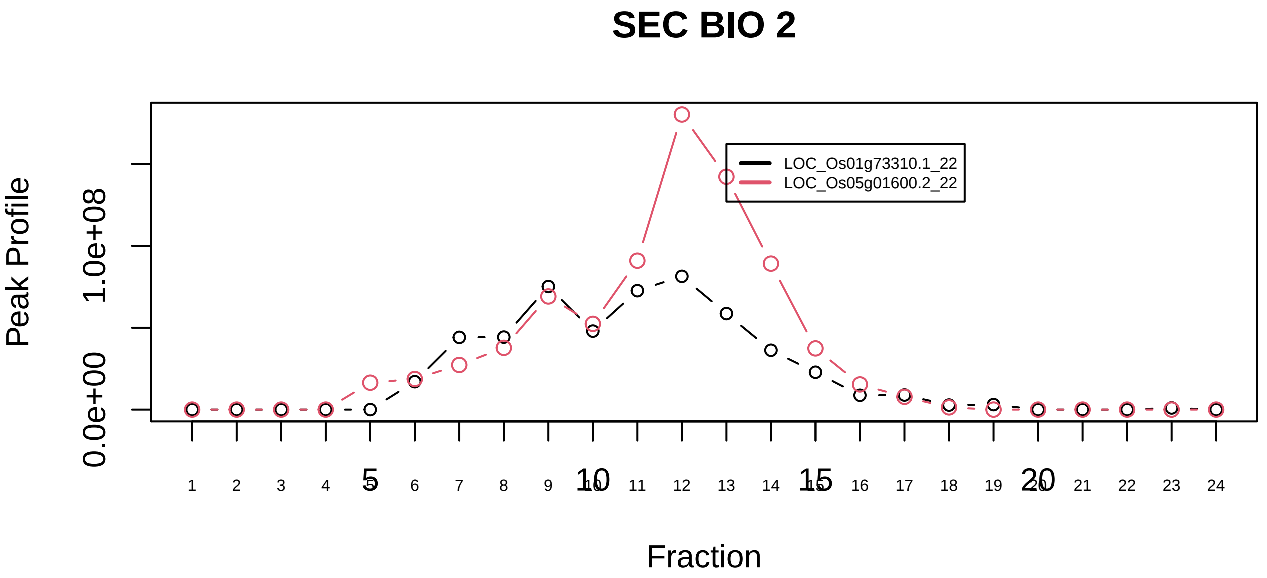

Pre−initiation complex (PIC)|5736−−sub_orthoparalog_2

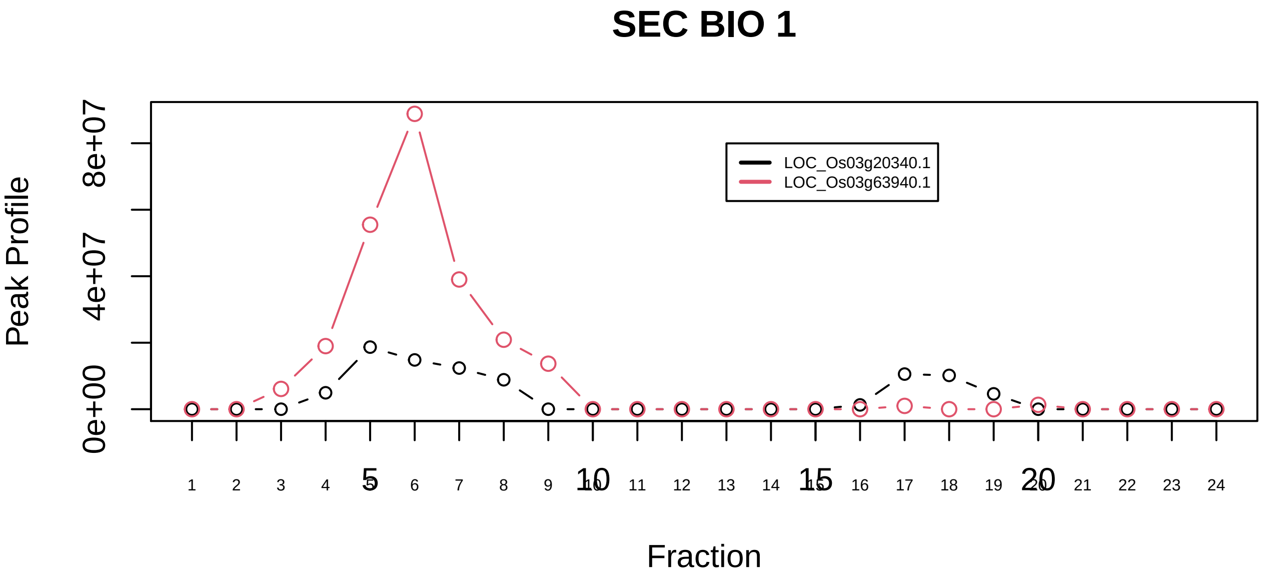

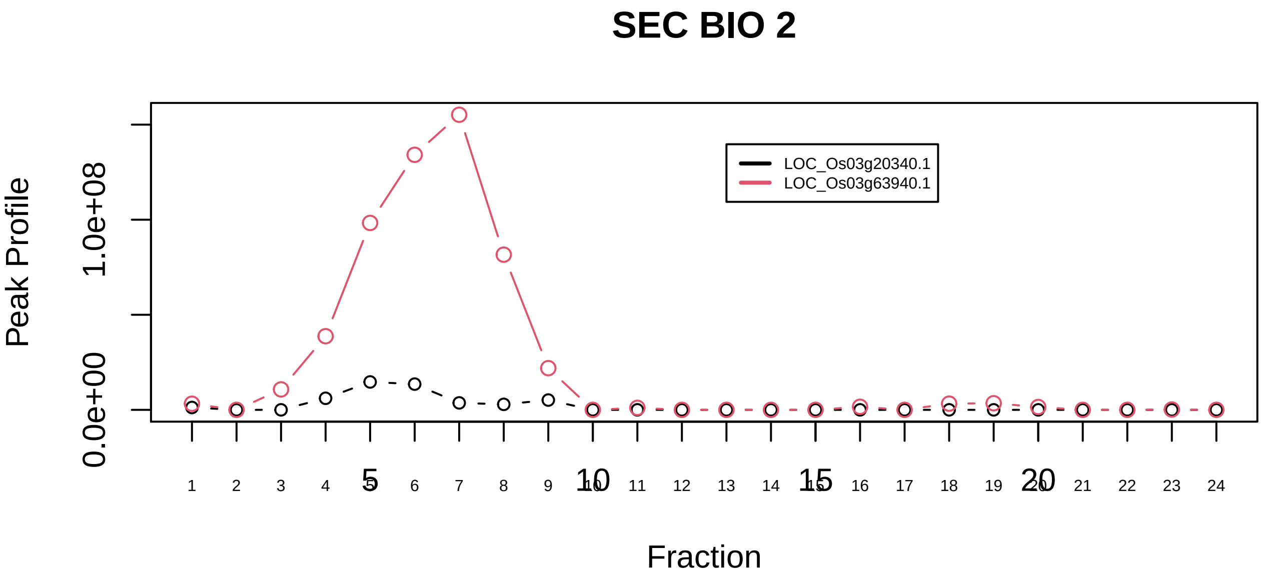

PRKAA2−PRKAB2−PRKAG3 complex|6153−−only_2_prots_before_clustering

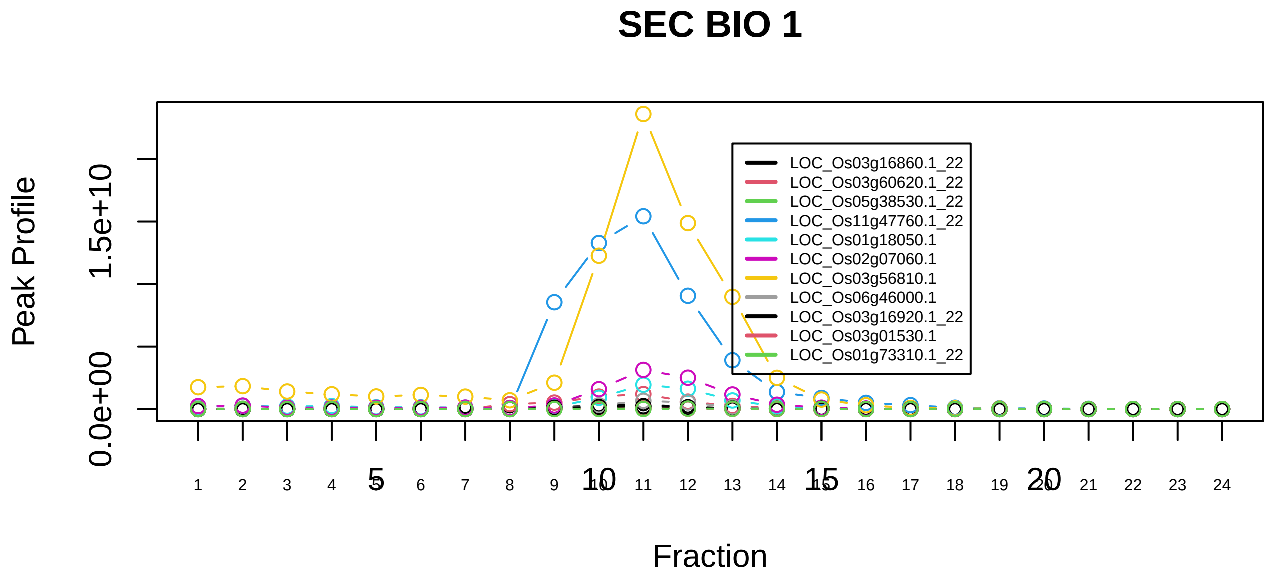

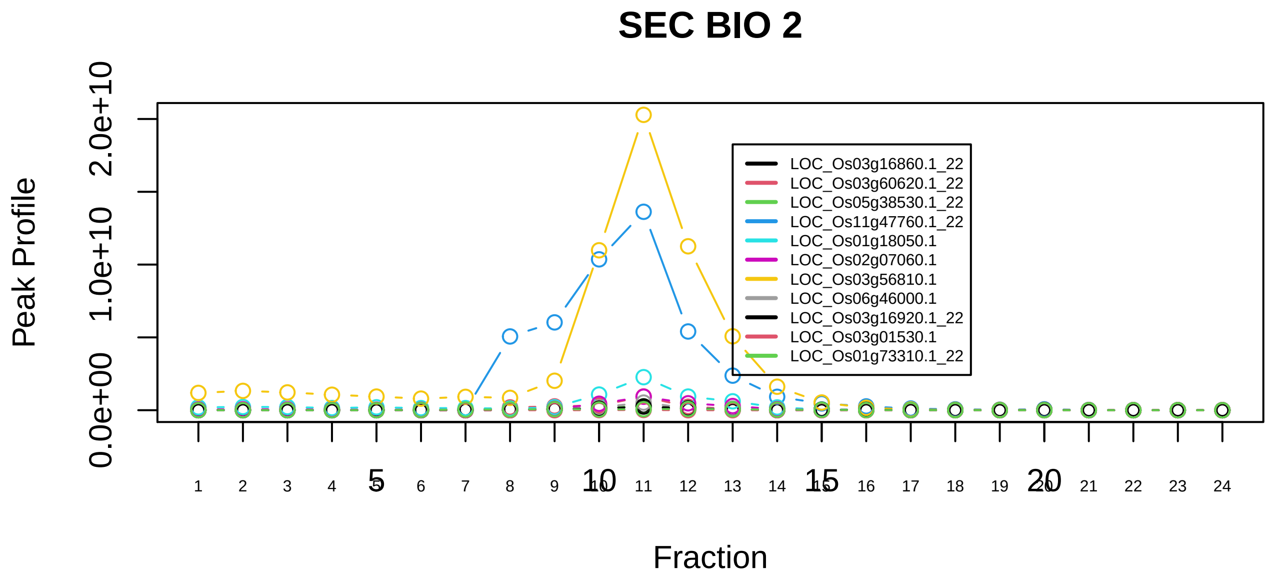

Profilin 1 complex|2837−−sub_1

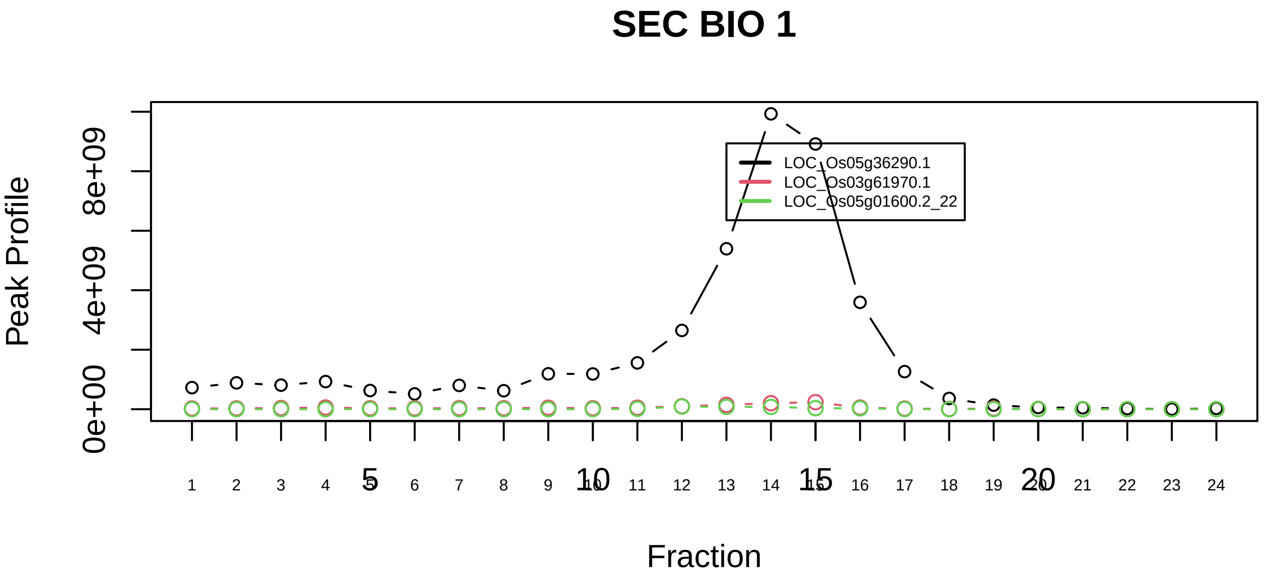

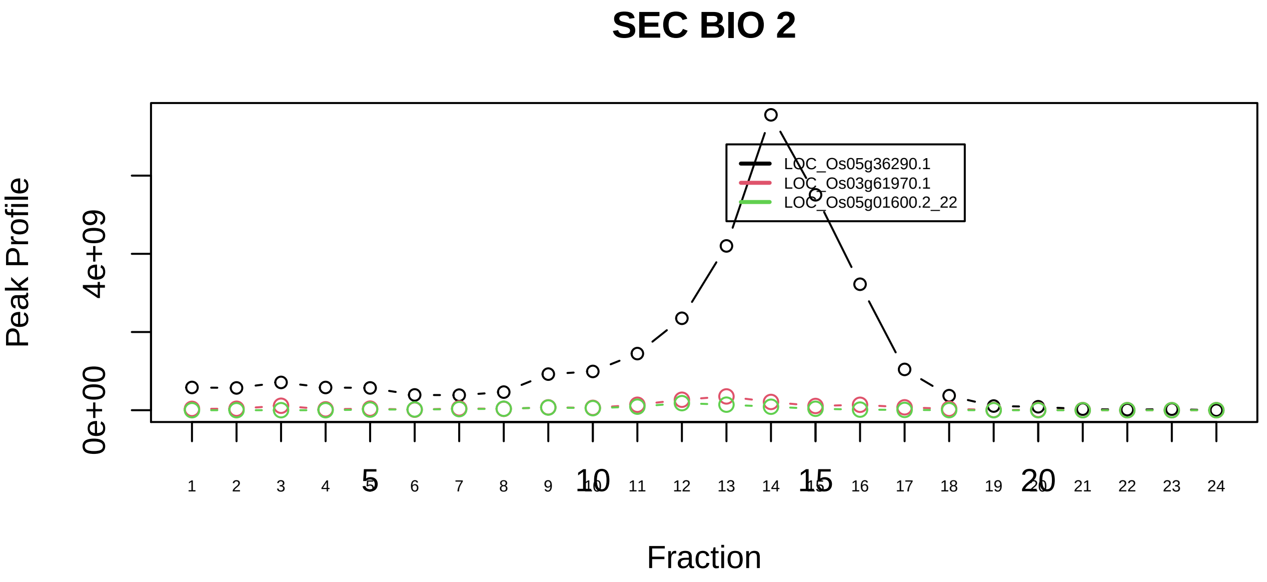

Profilin 1 complex|2837−−sub_orthoparalog_1

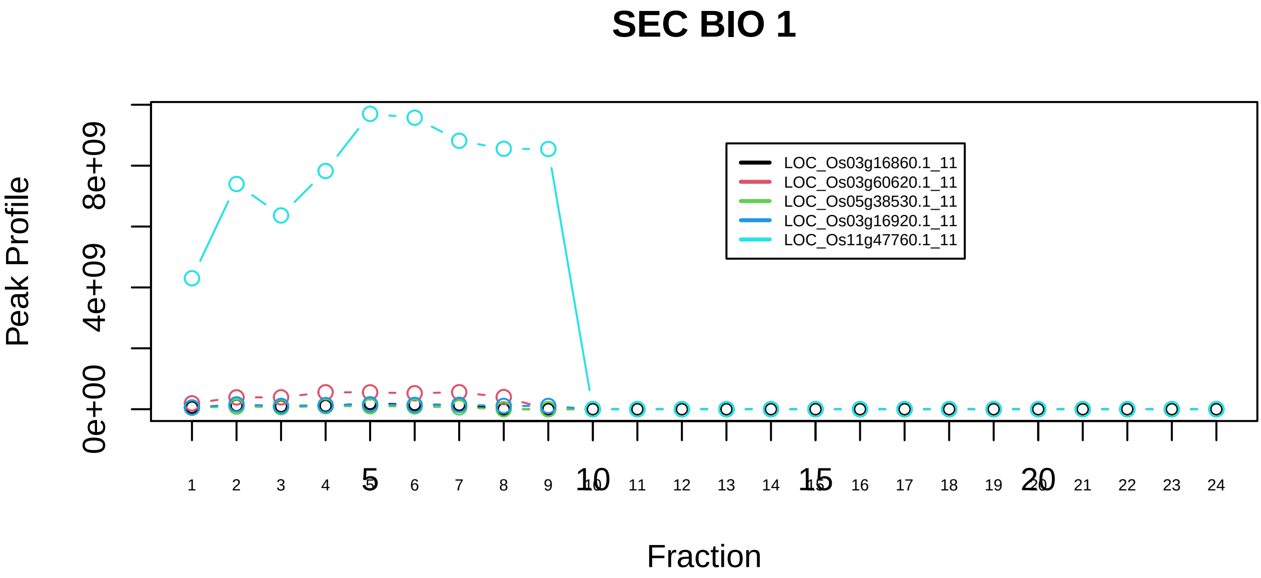

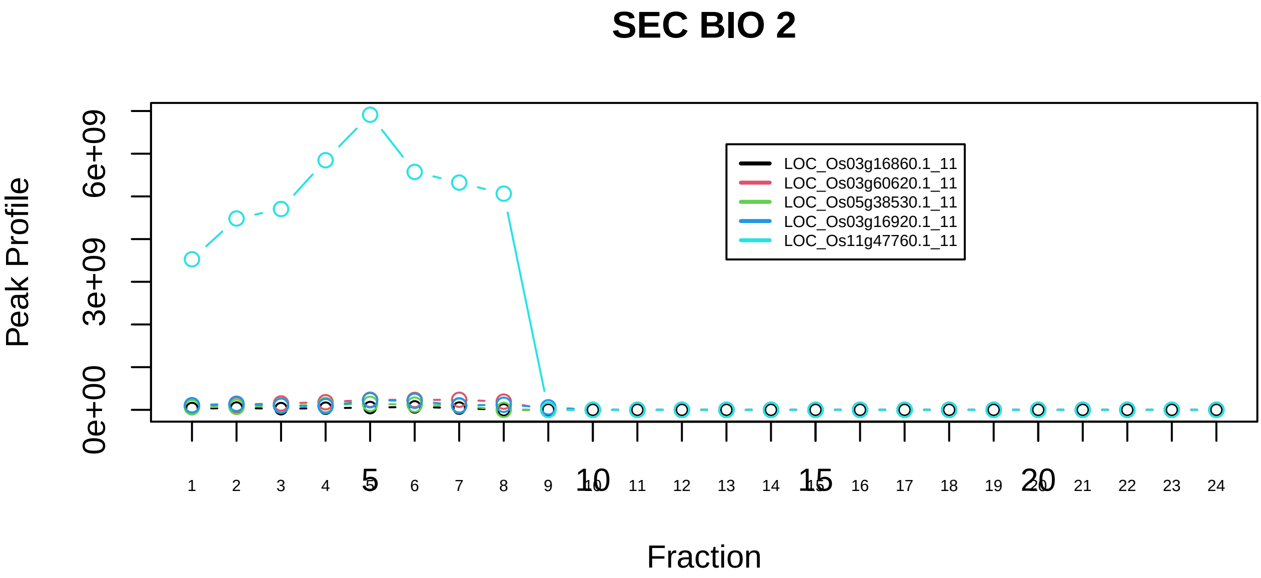

Profilin 1 complex|2837−−sub_orthoparalog_2

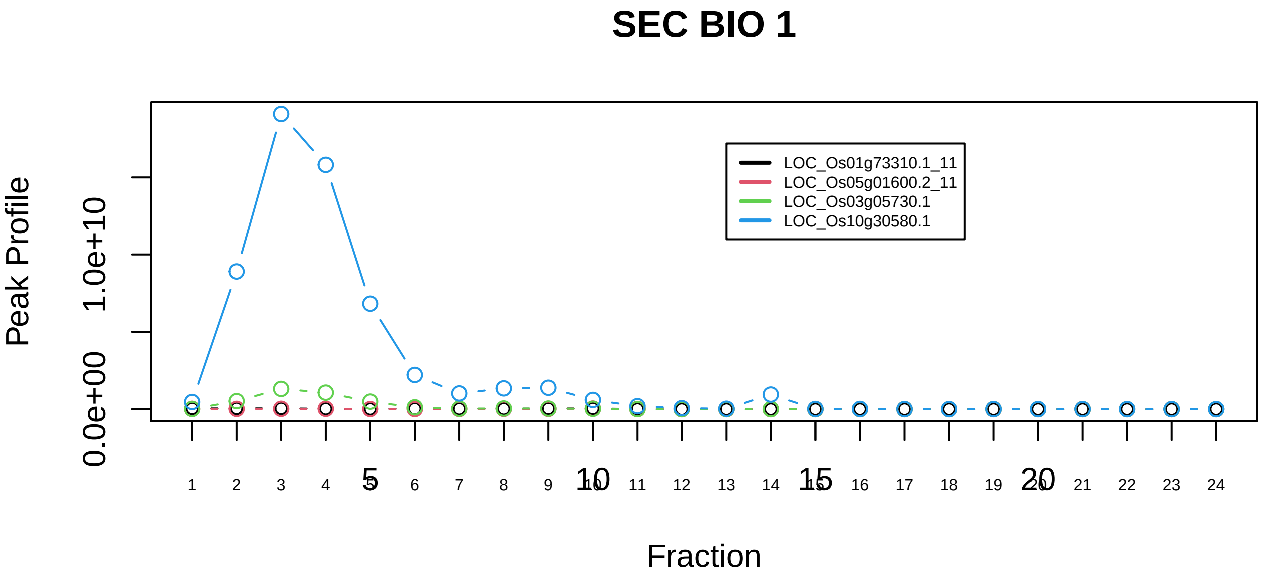

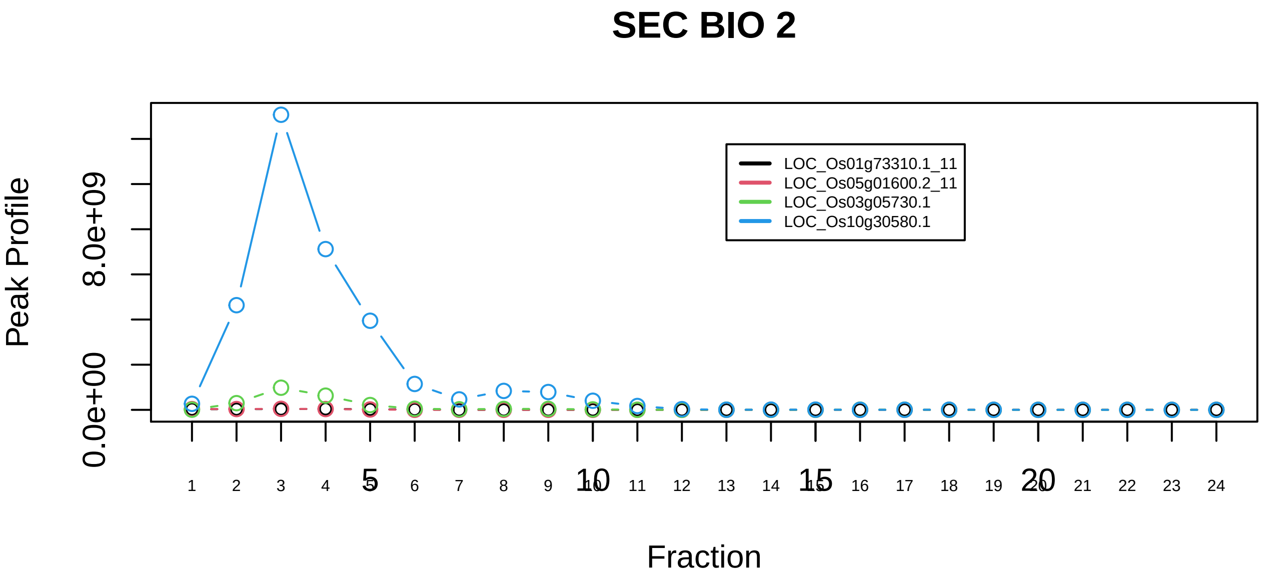

Profilin 1 complex|2837−−sub_2

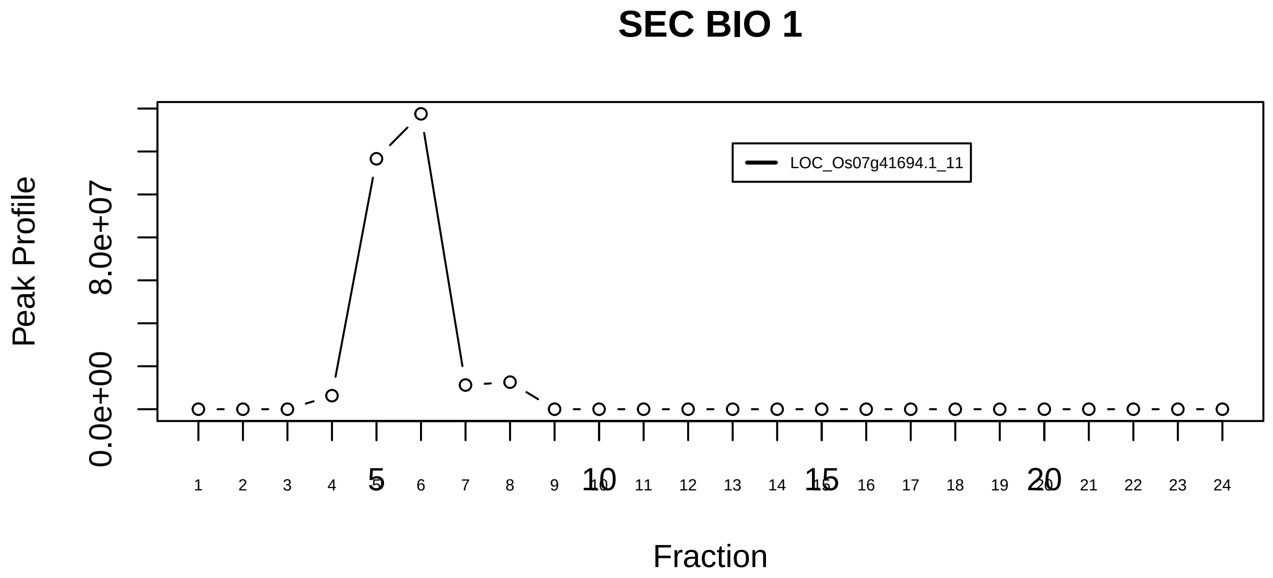

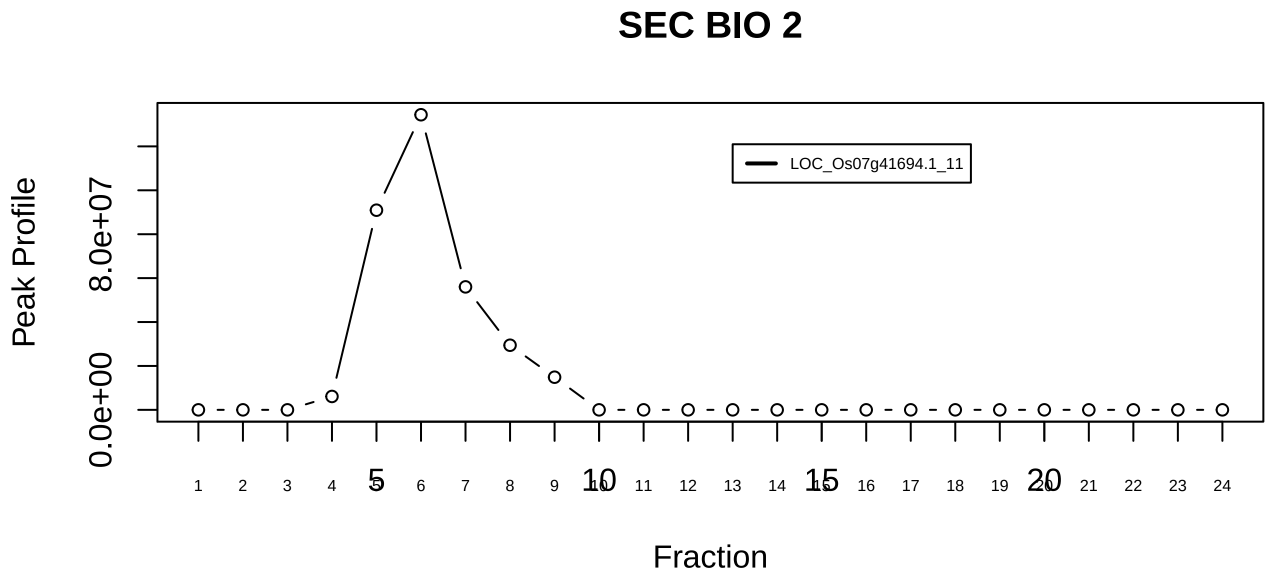

ProTalpha C8 complex|6988−−singleton_1

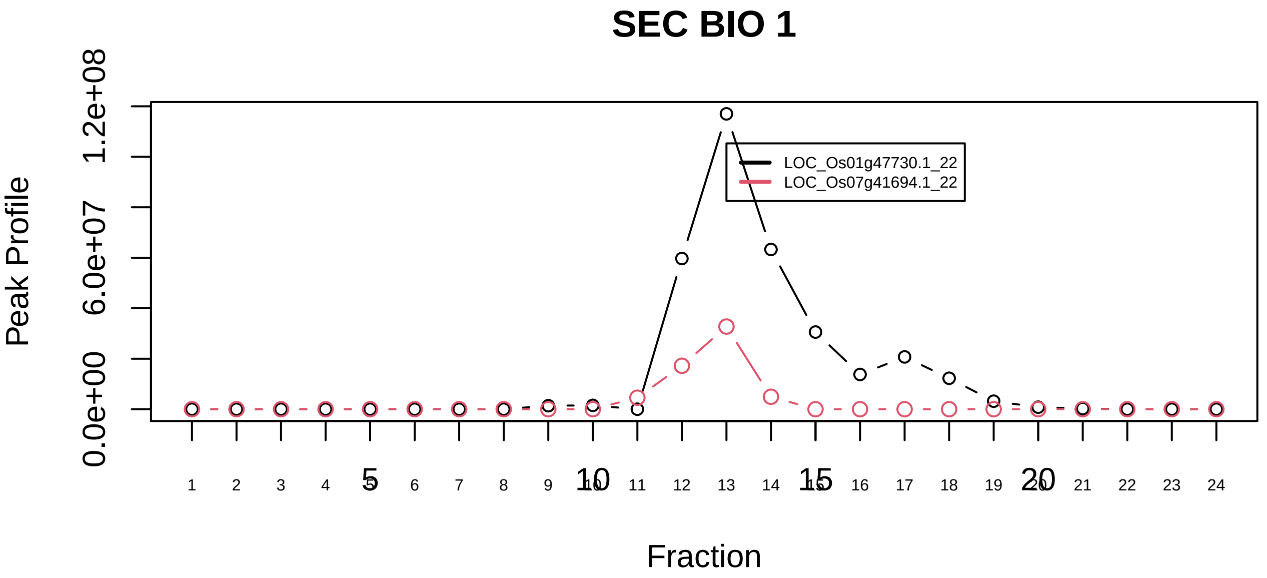

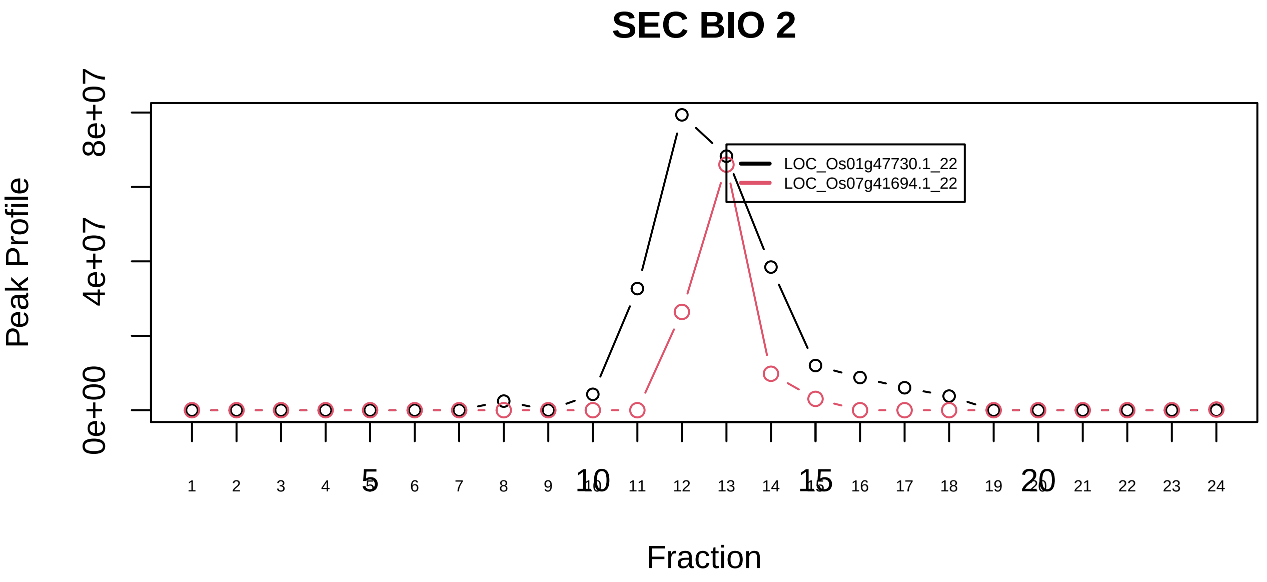

ProTalpha C8 complex|6988−−sub_1

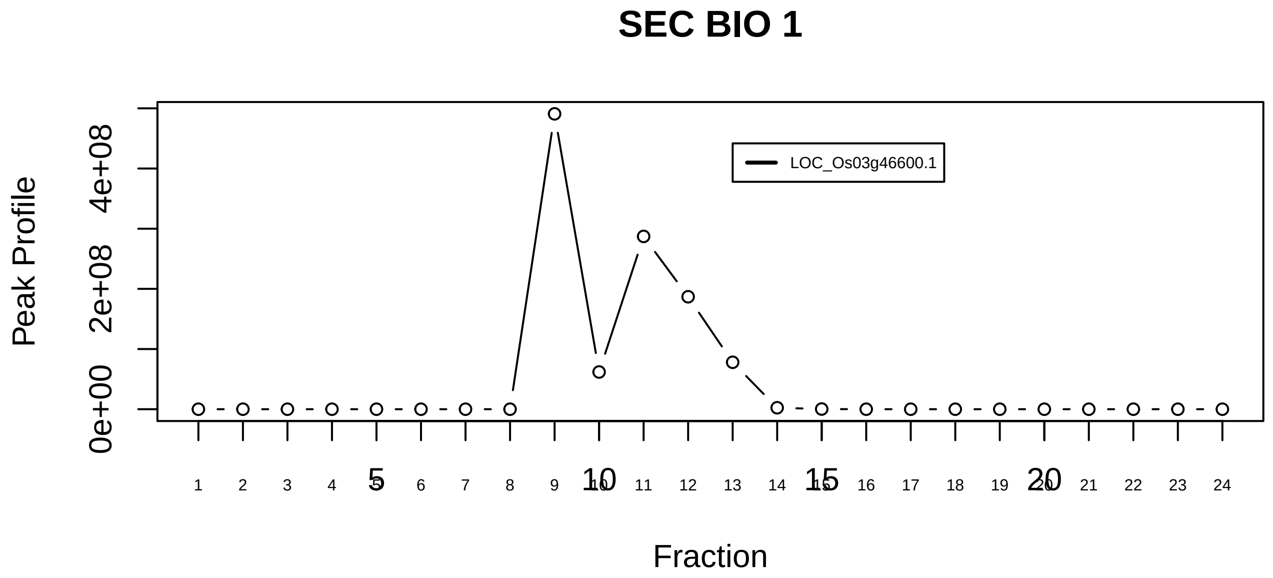

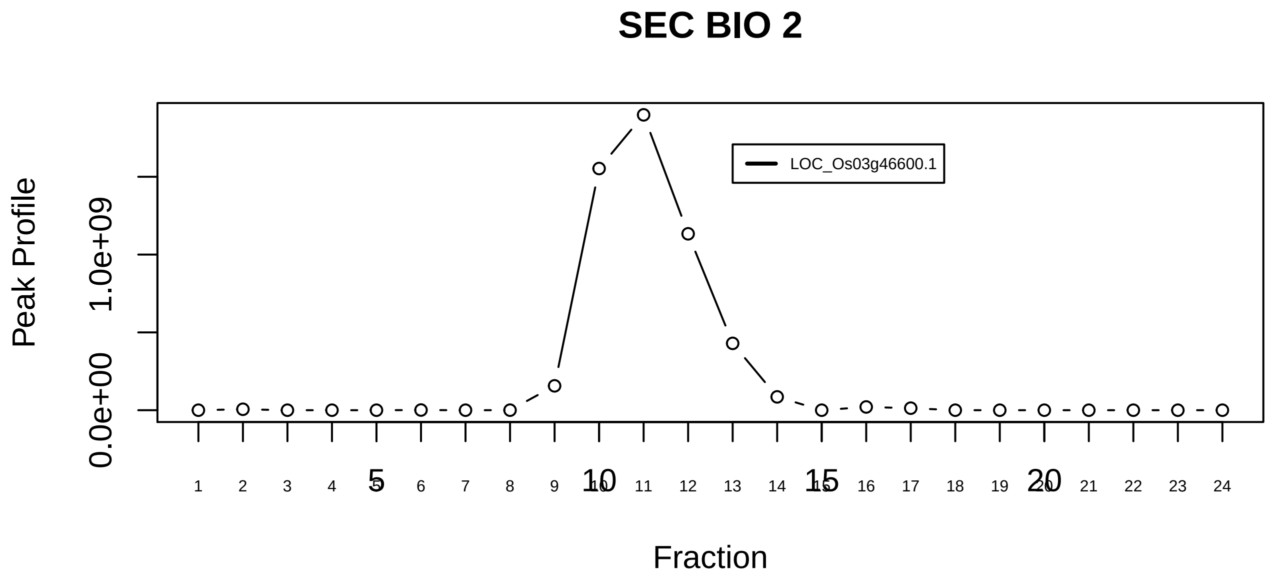

ProTalpha C8 complex|6988−−singleton_2

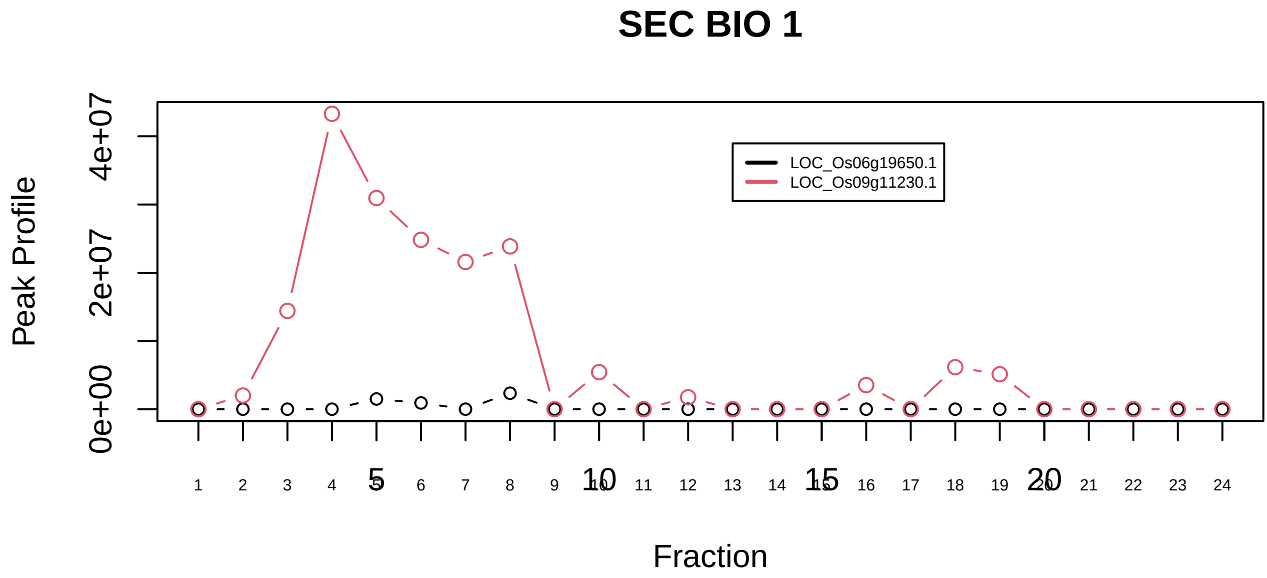

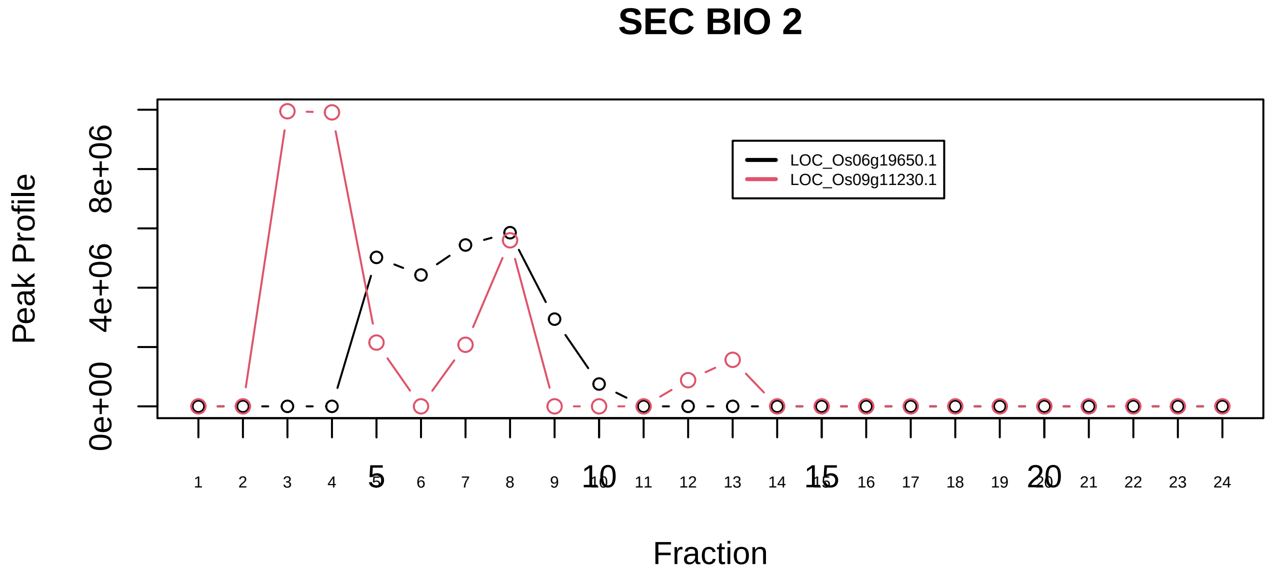

Protein phosphatase 4 complex|1245−−only_2_prots_before_clustering

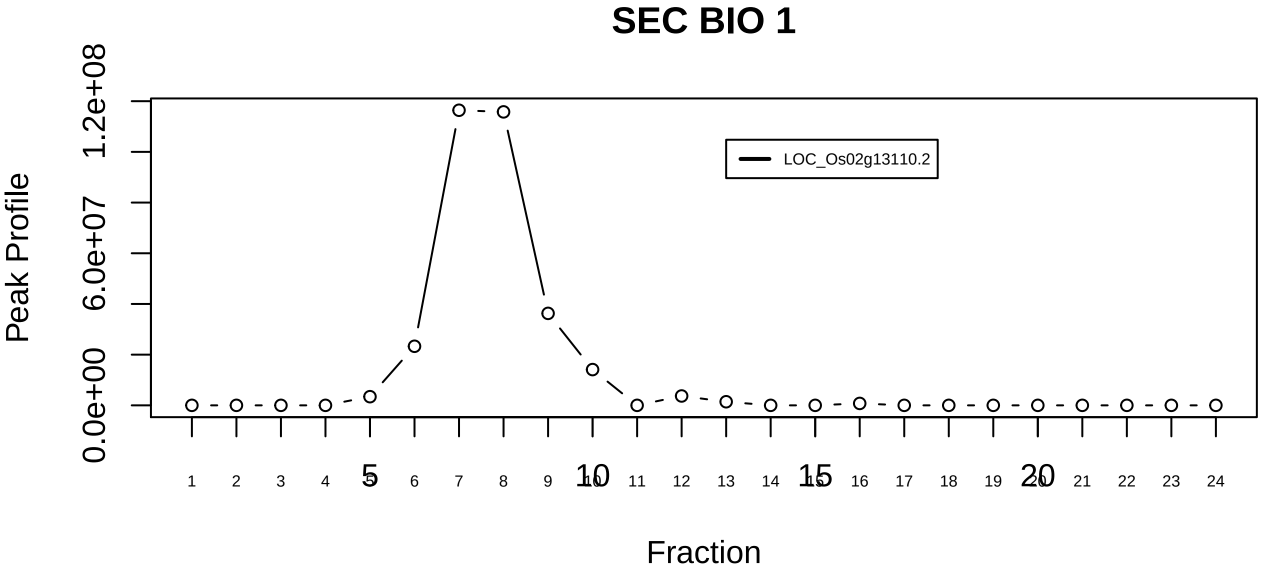

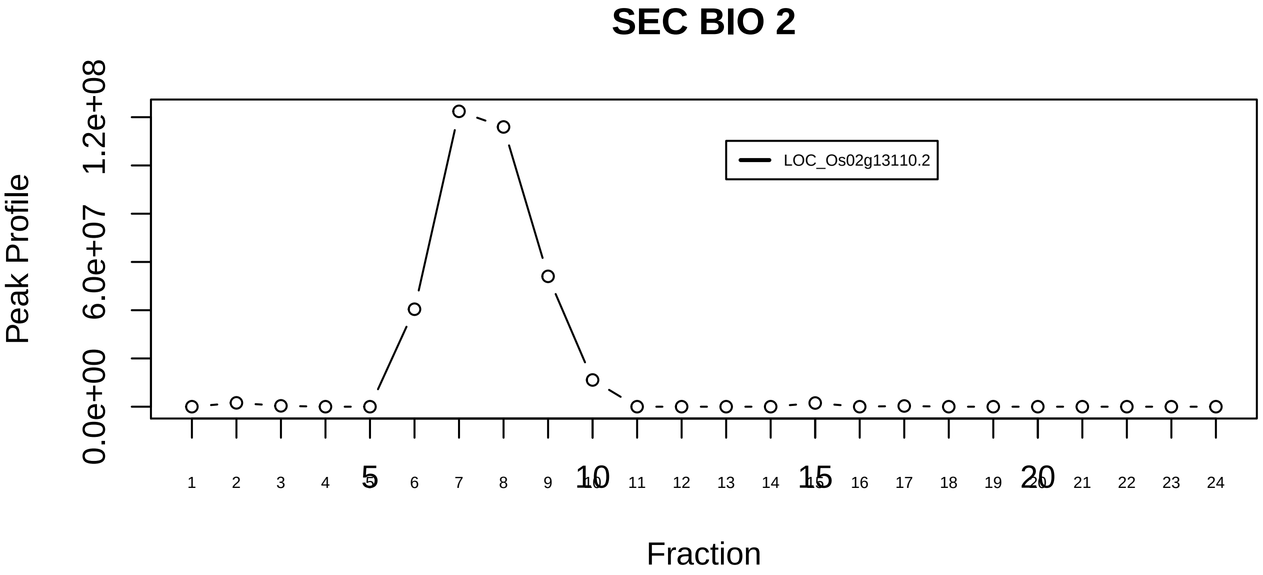

RAF1−PPP2−PIN1 complex|5211−−singleton_1

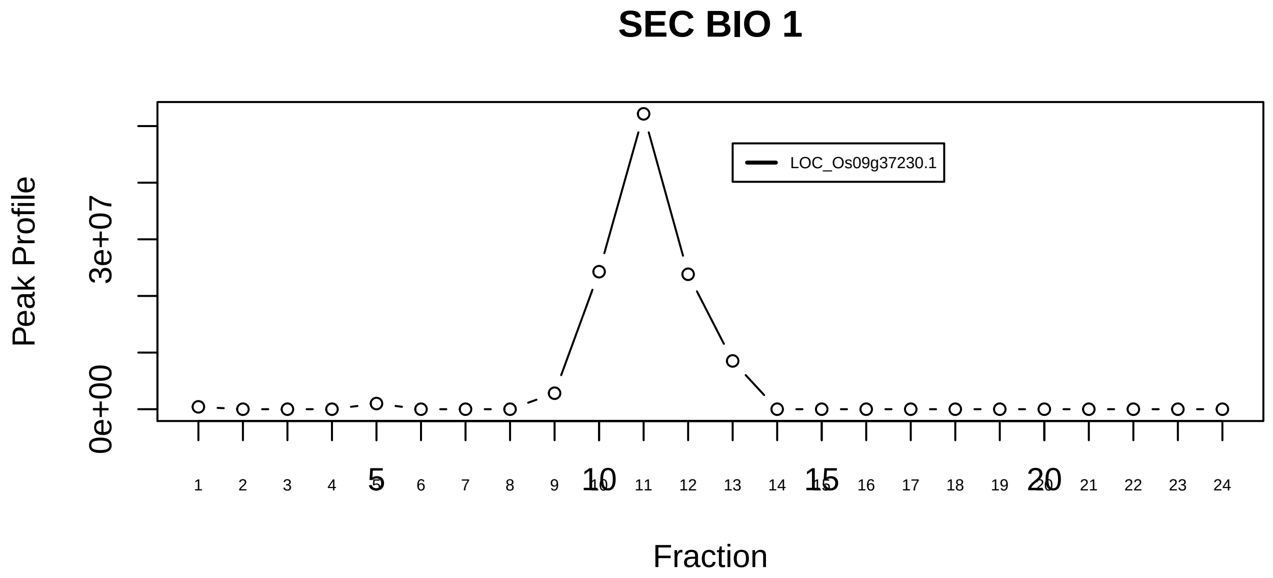

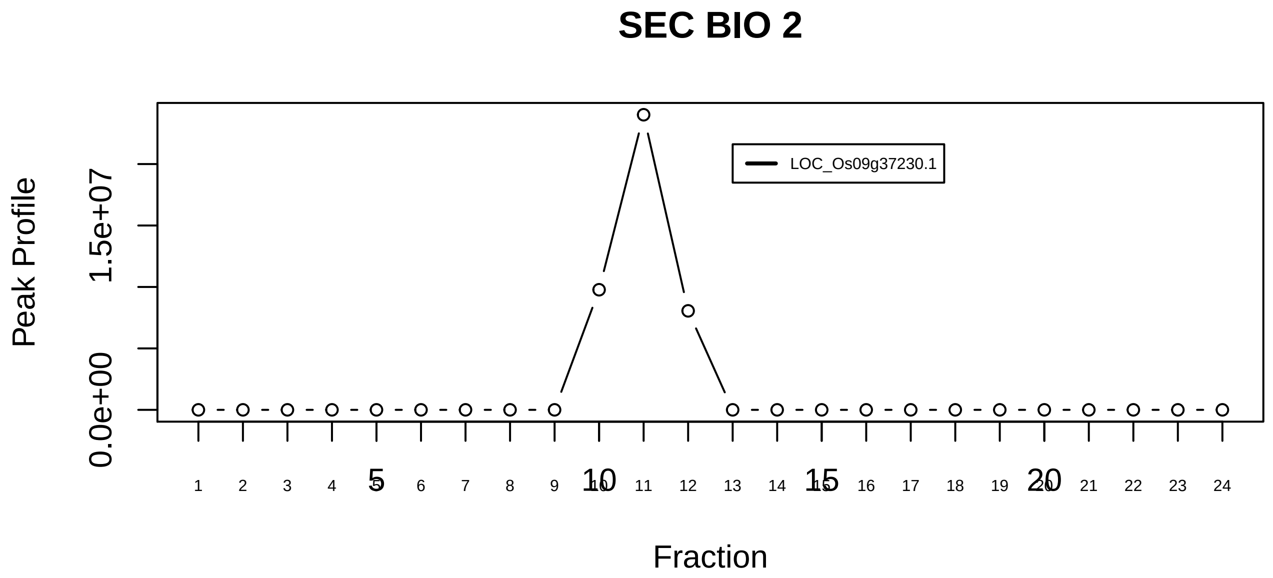

RAF1−PPP2−PIN1 complex|5211−−singleton_2

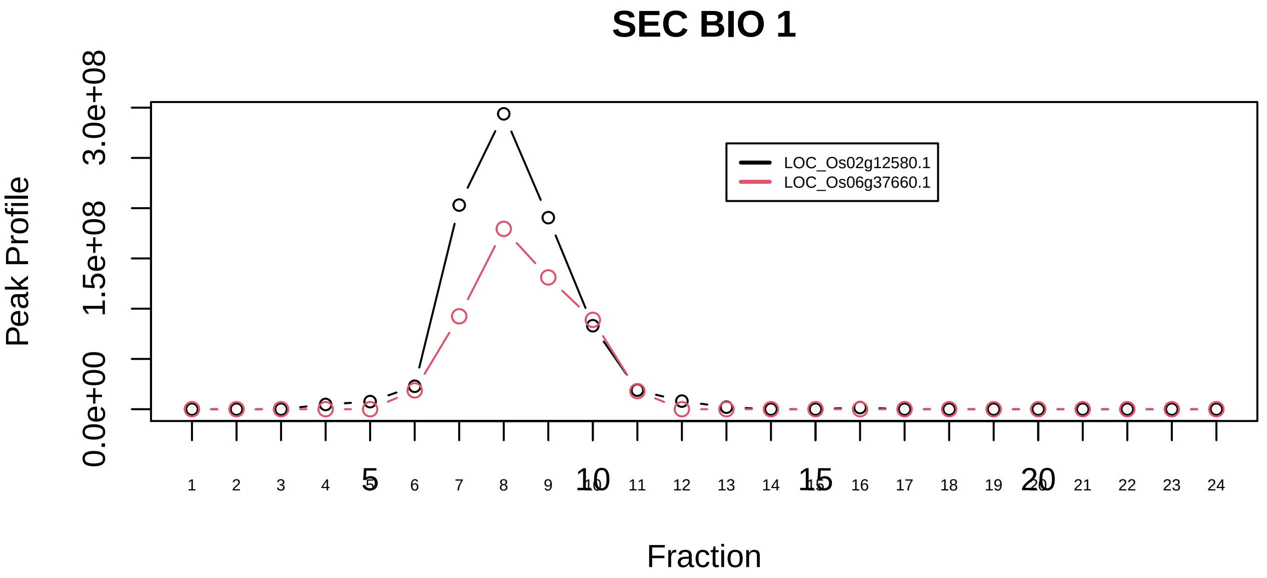

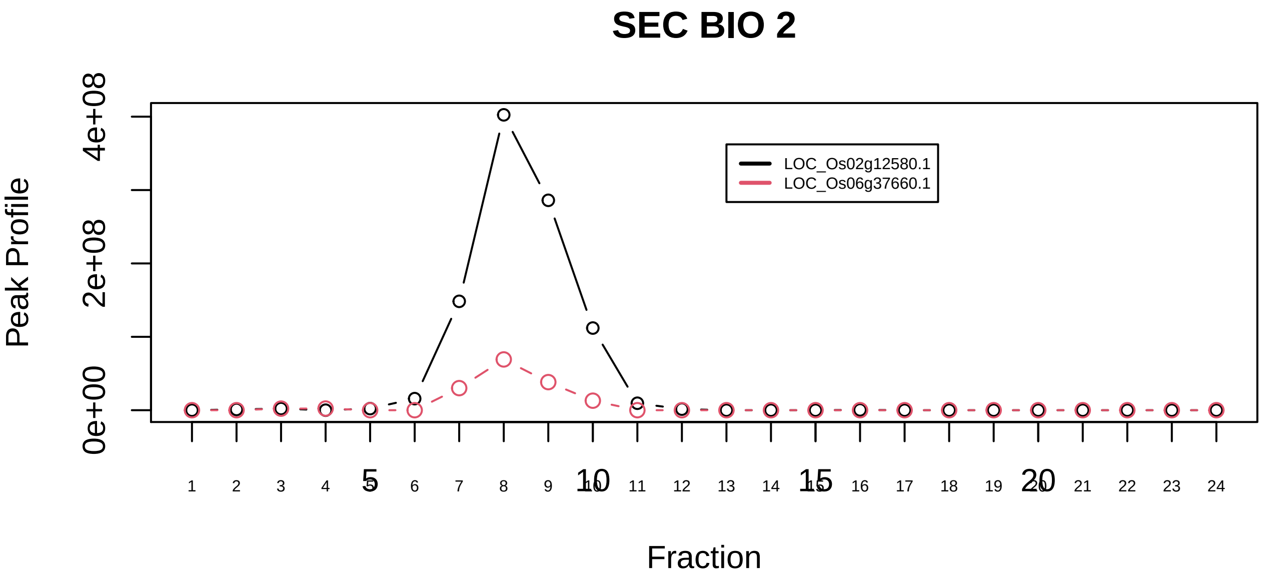

RAF1−PPP2−PIN1 complex|5211−−sub_orthoparalog_1

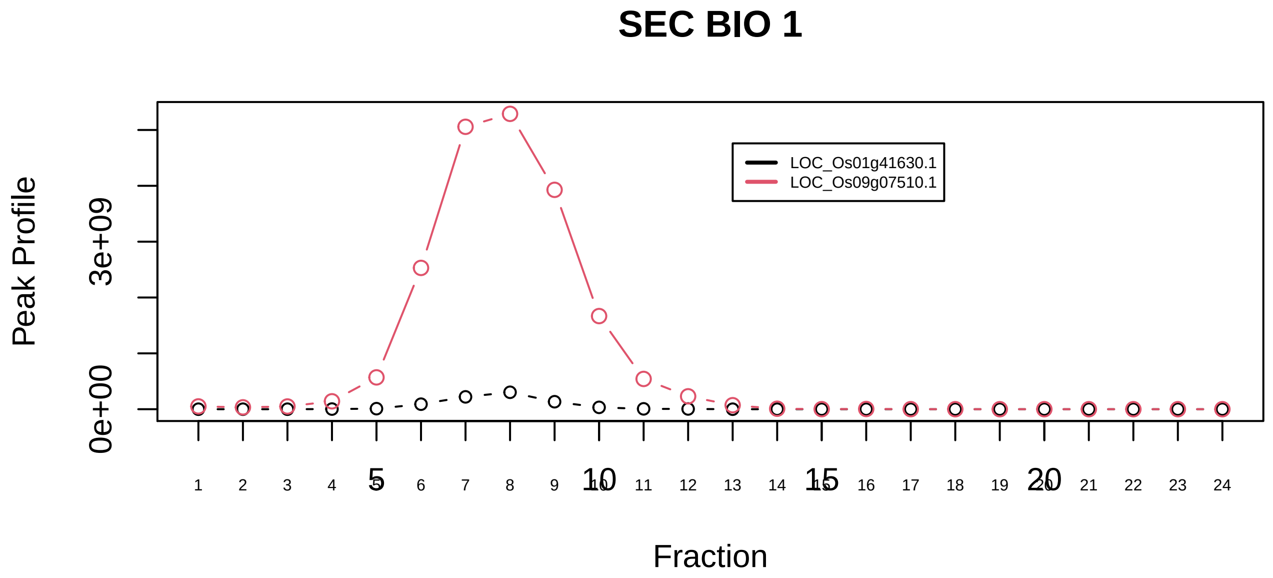

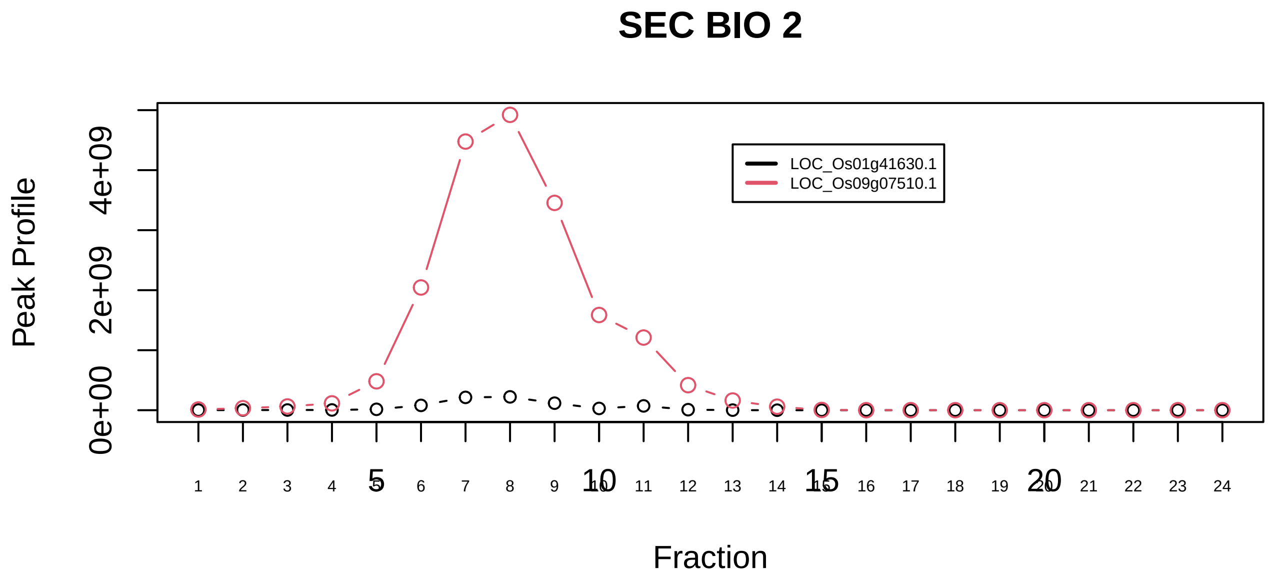

RAF1−PPP2−PIN1 complex|5211−−sub_1

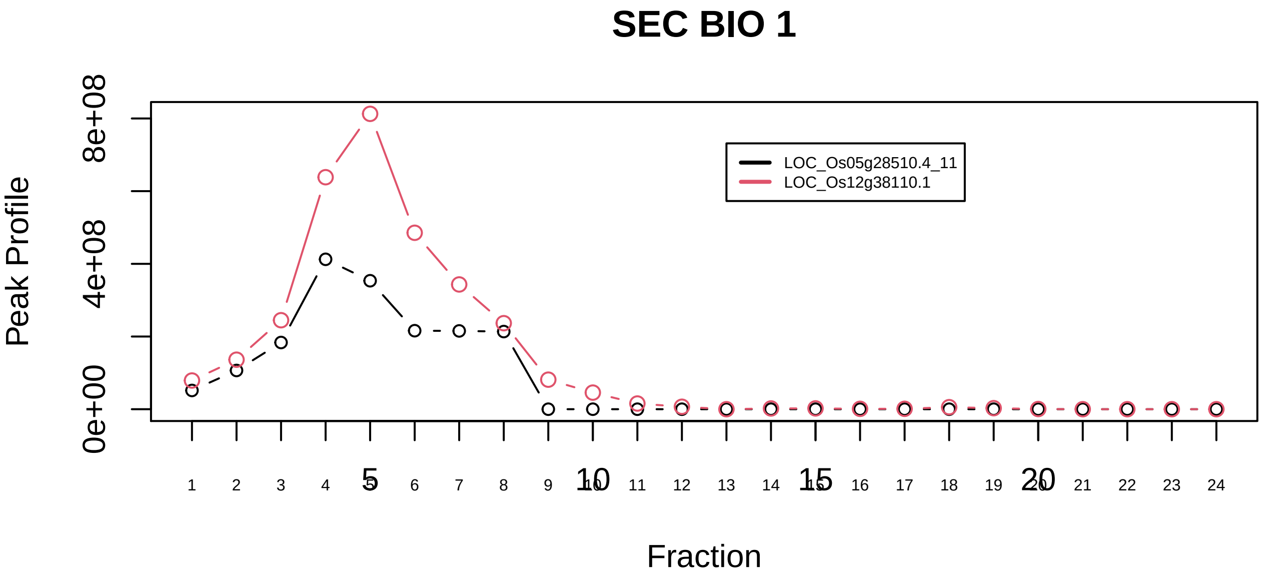

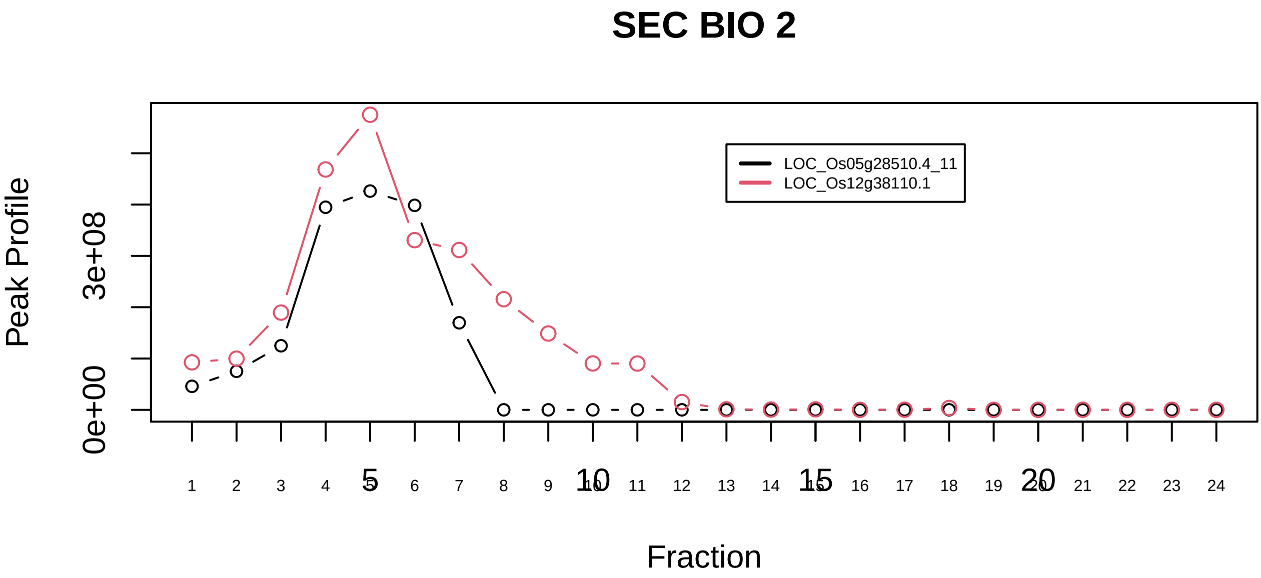

RANBP1−RAN−KPNB1 complex|1554−−sub_orthoparalog_1

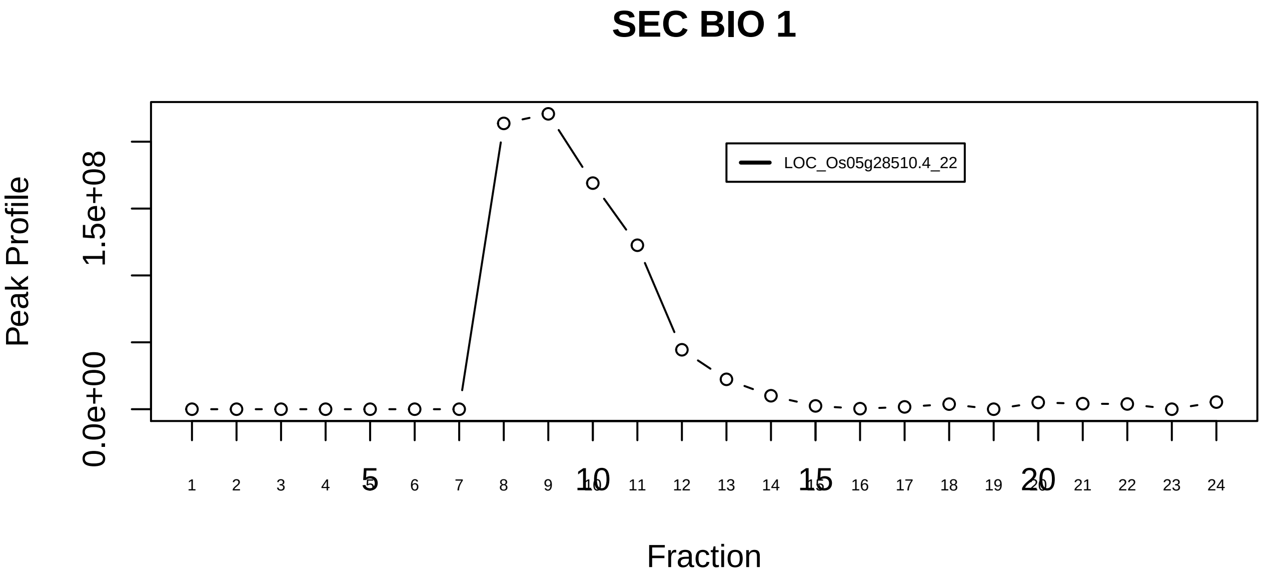

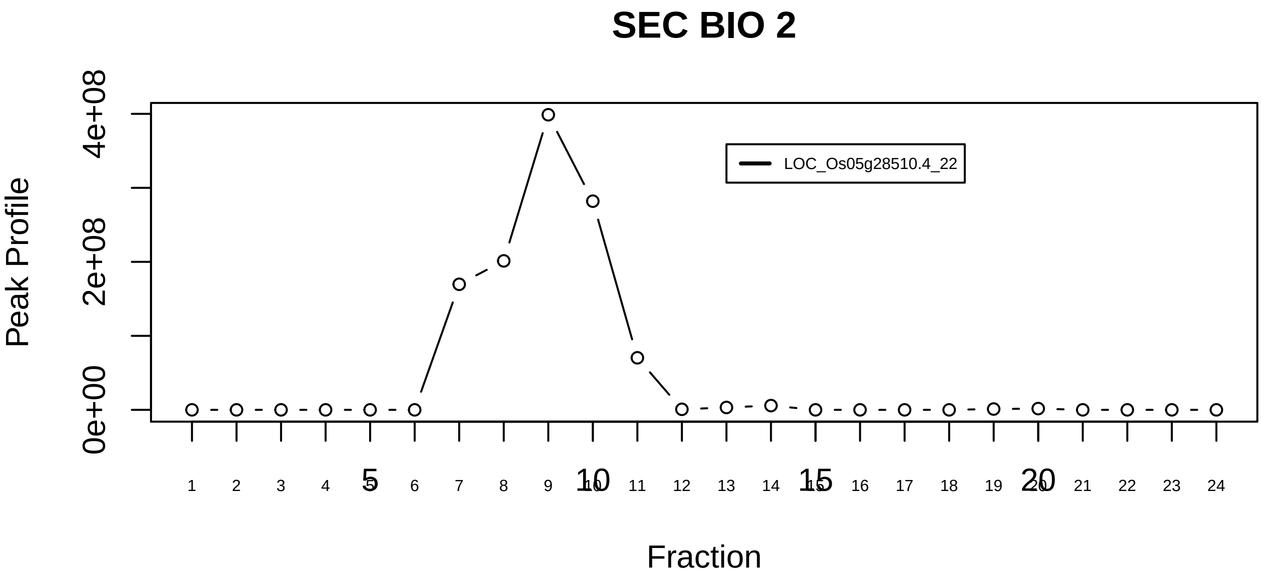

RANBP1−RAN−KPNB1 complex|1554−−singleton_1

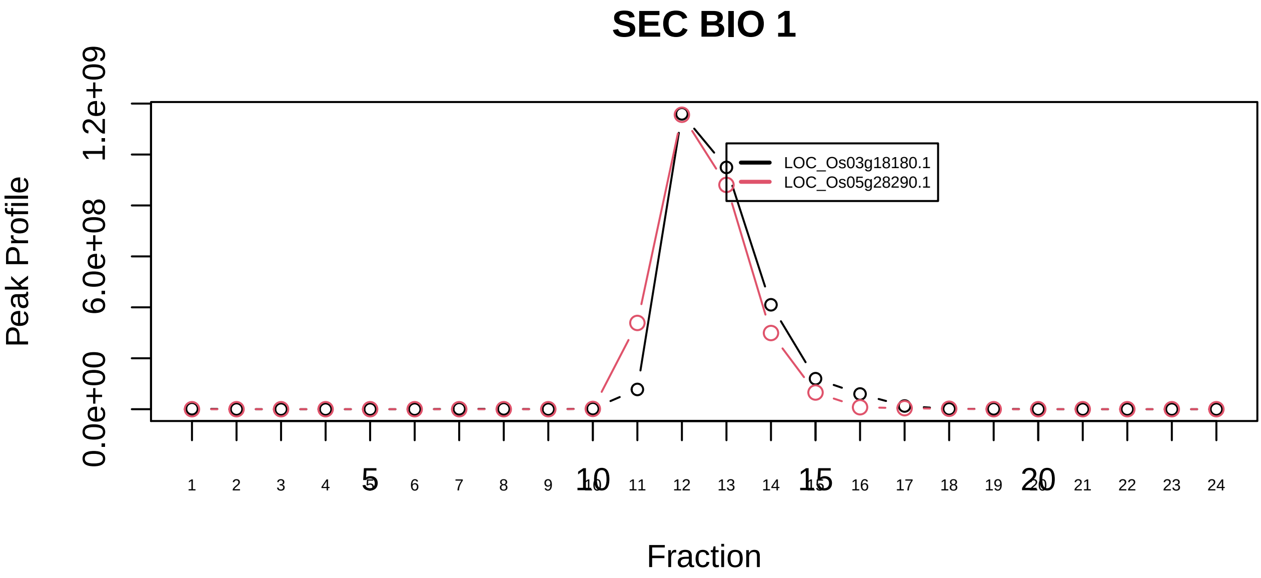

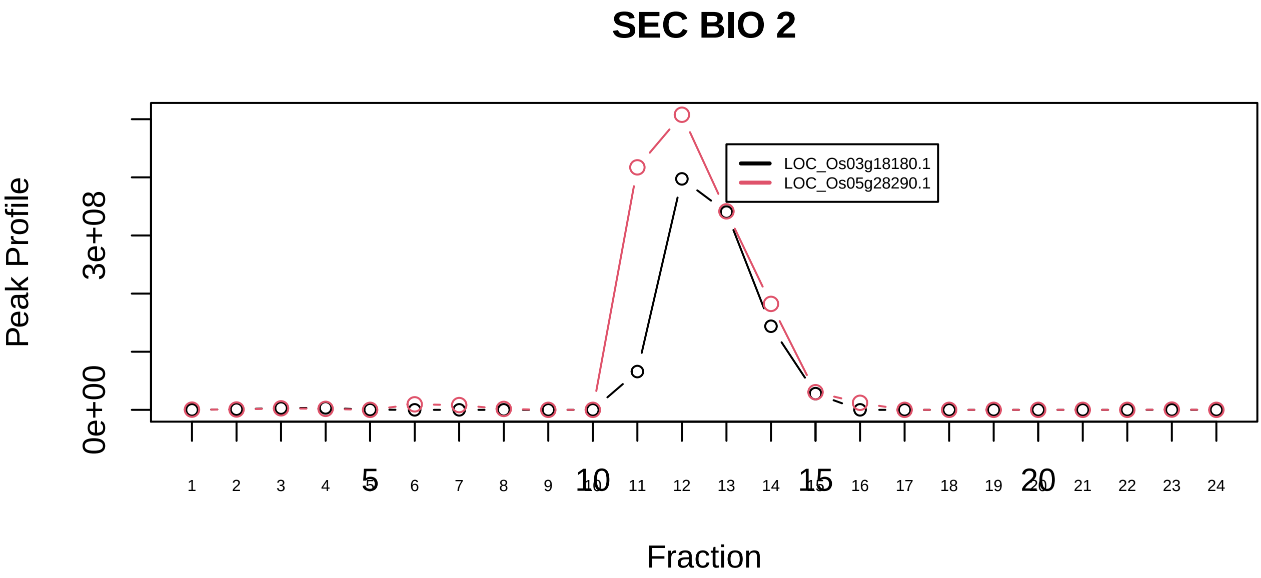

RANBP1−RAN−KPNB1 complex|1554−−sub_orthoparalog_2

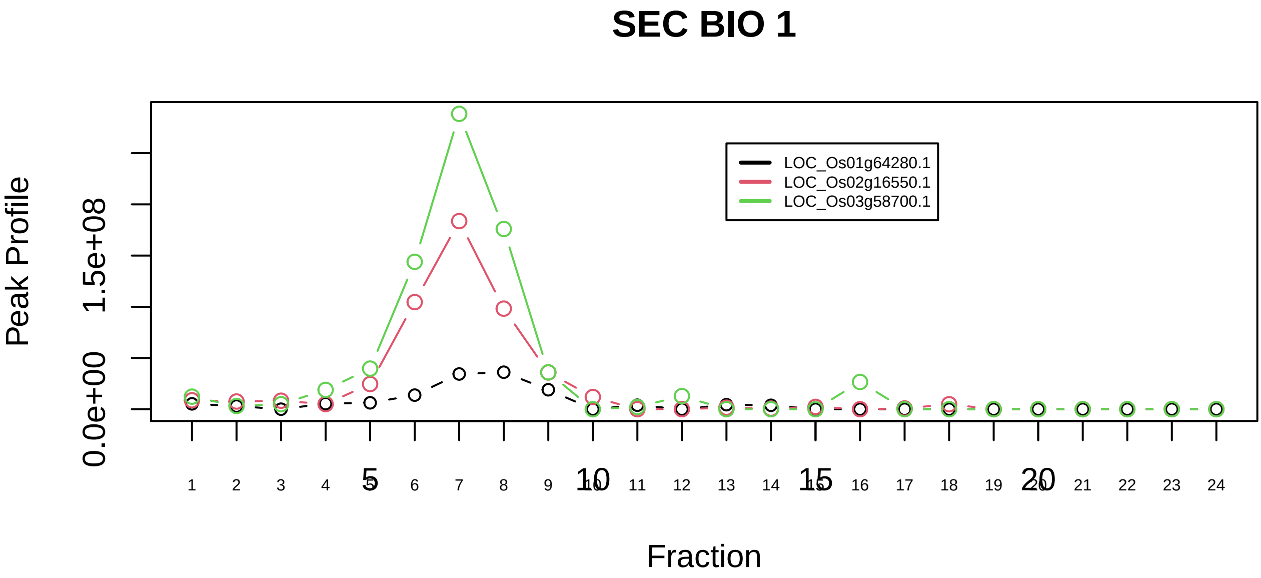

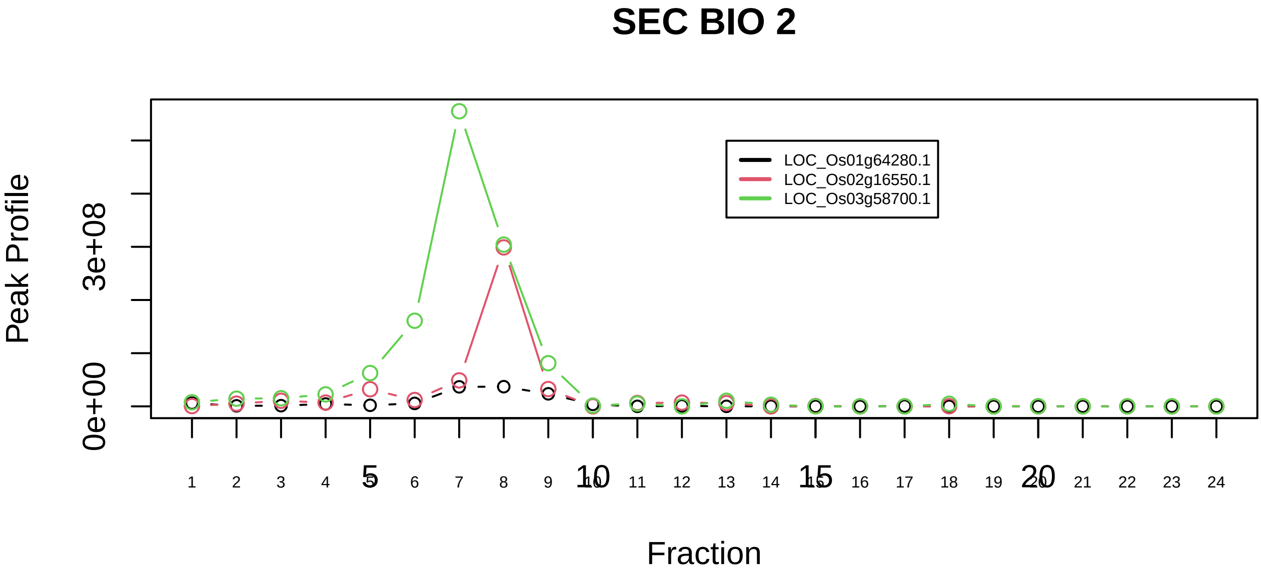

Retromer complex (SNX1, SNX2, VPS35, VPS29, VPS26B)|1060−−all_prots_before_clustering_sim

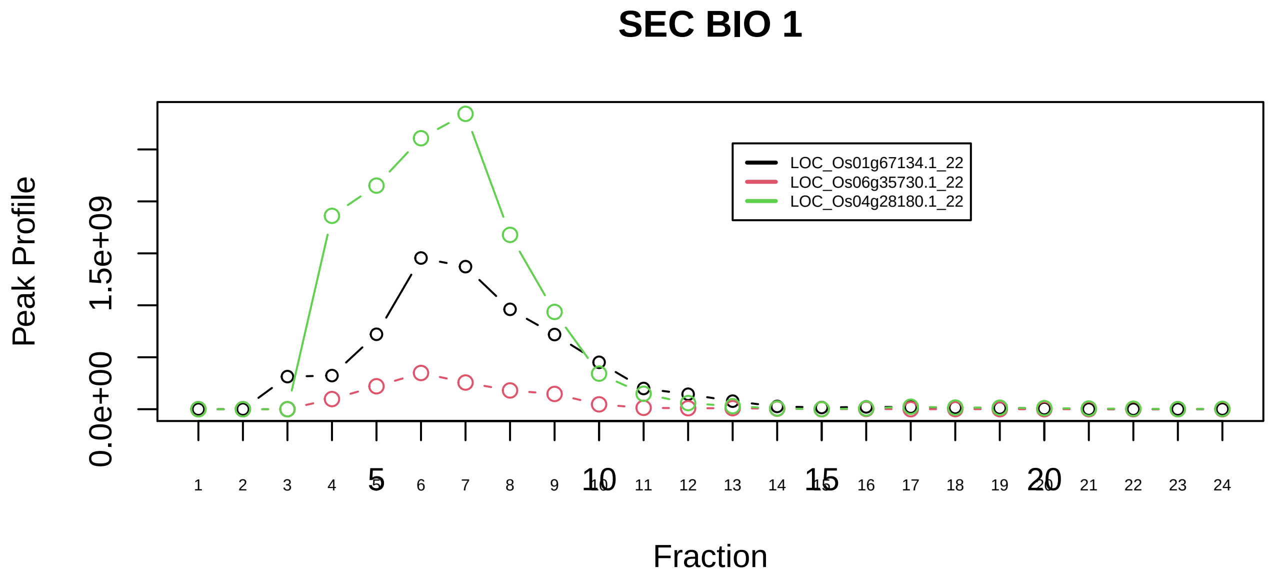

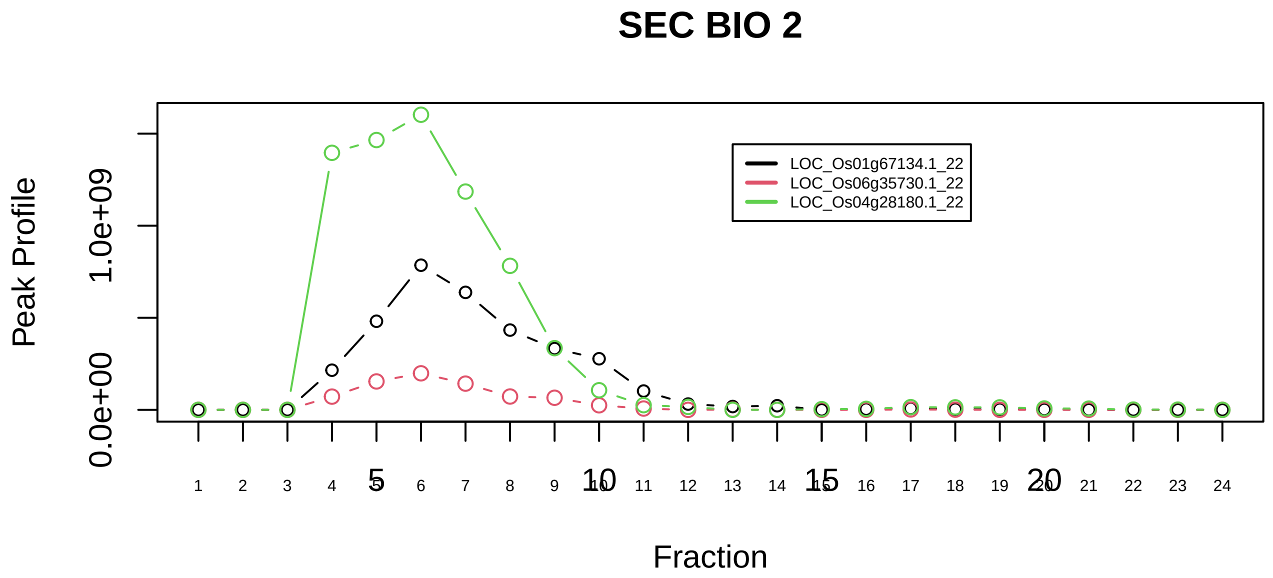

Ribosome, cytoplasmic|306−−sub_1

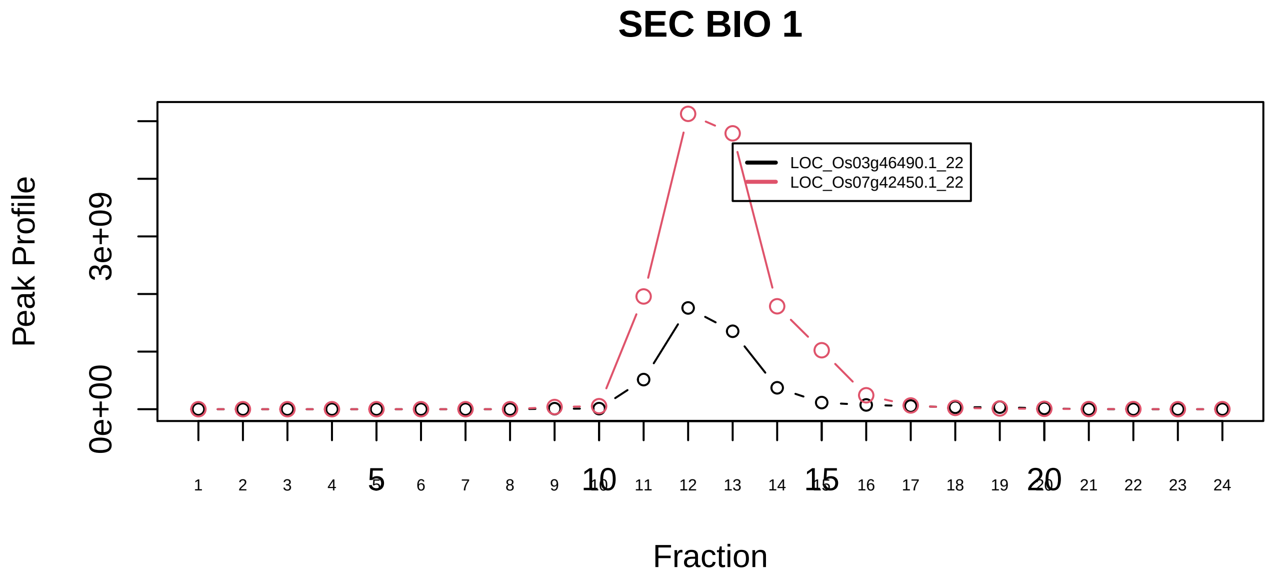

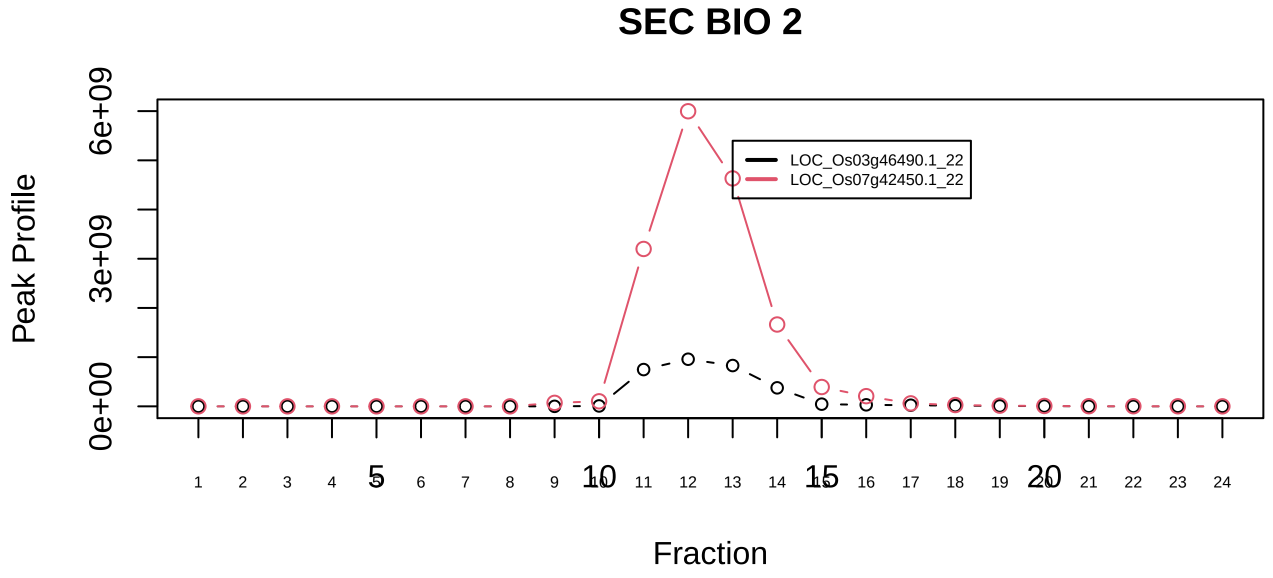

Ribosome, cytoplasmic|306−−sub_2

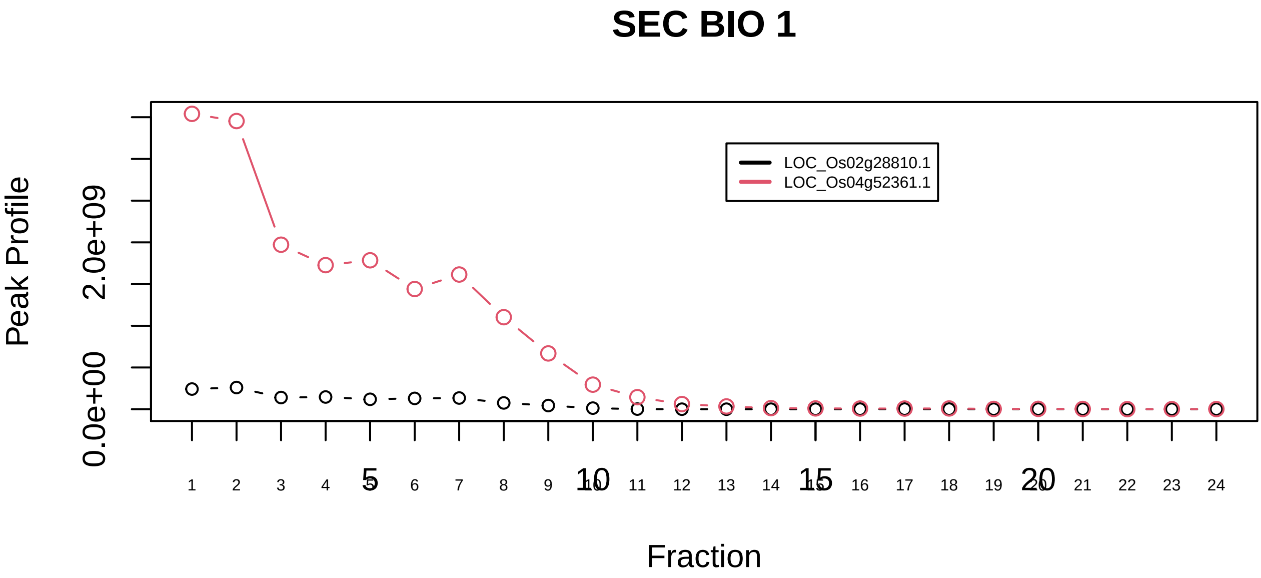

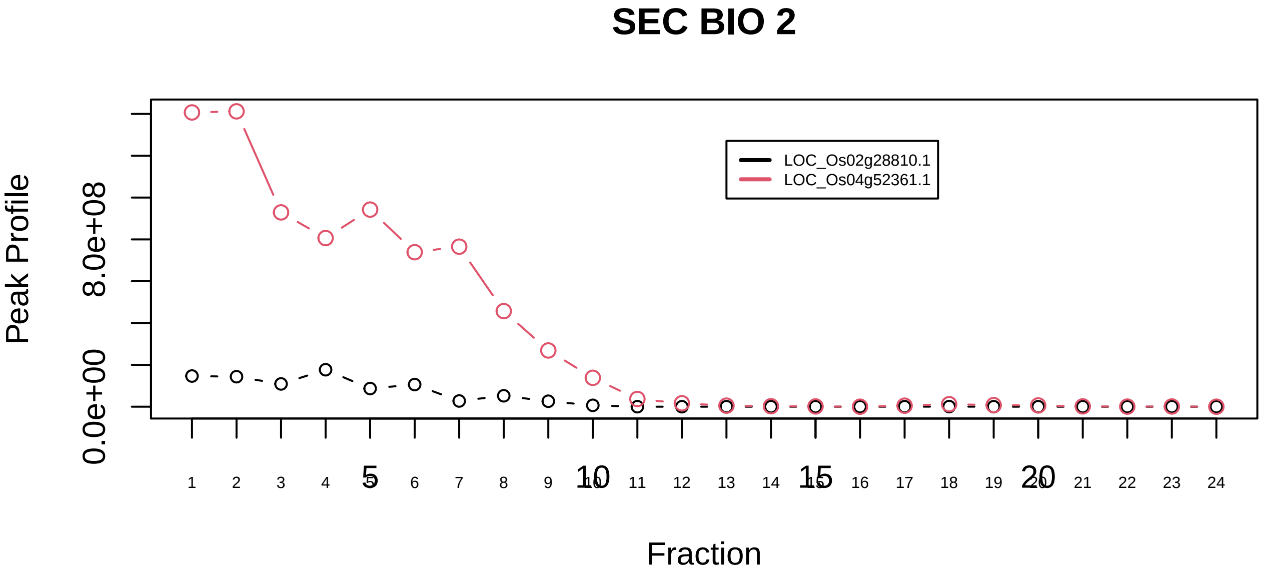

Ribosome, cytoplasmic|306−−sub_3

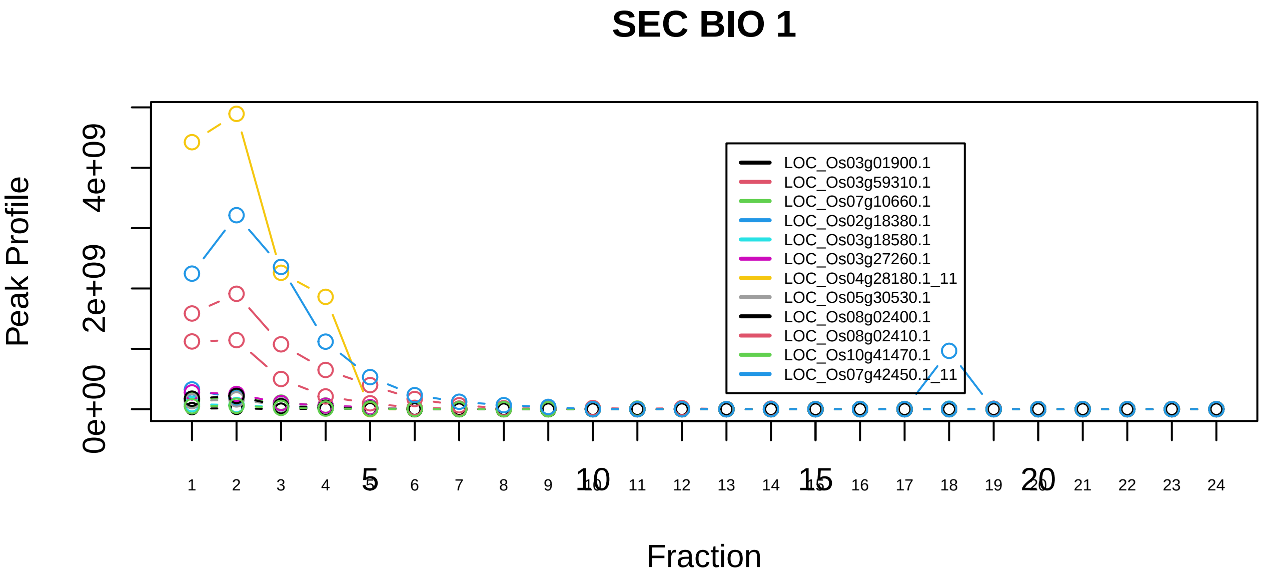

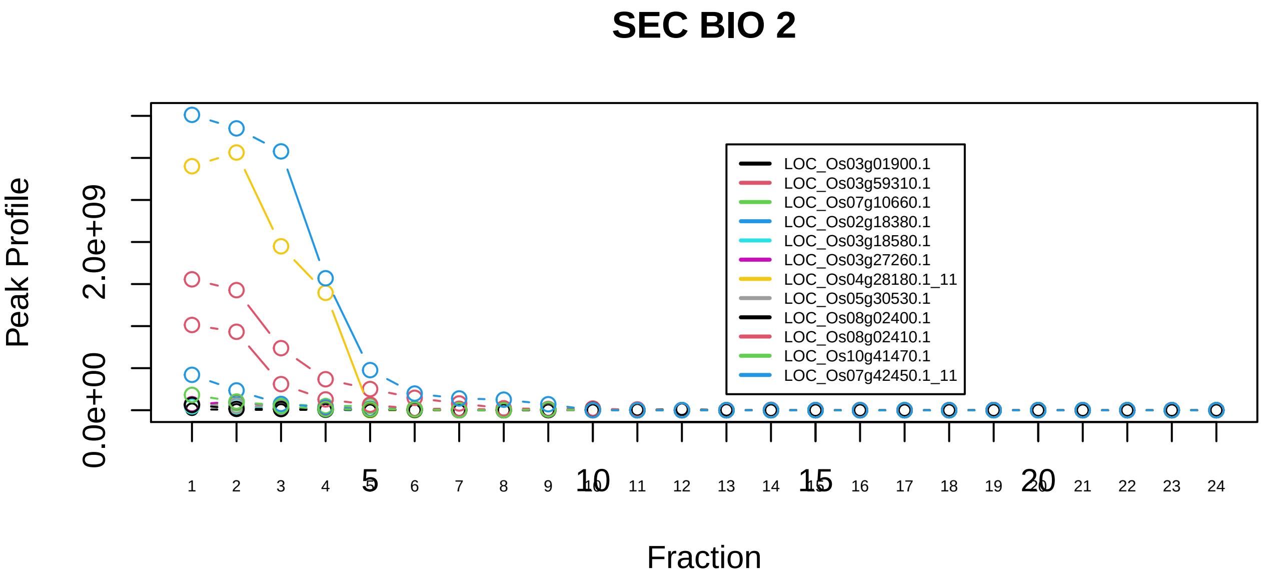

Ribosome, cytoplasmic|306−−sub_4

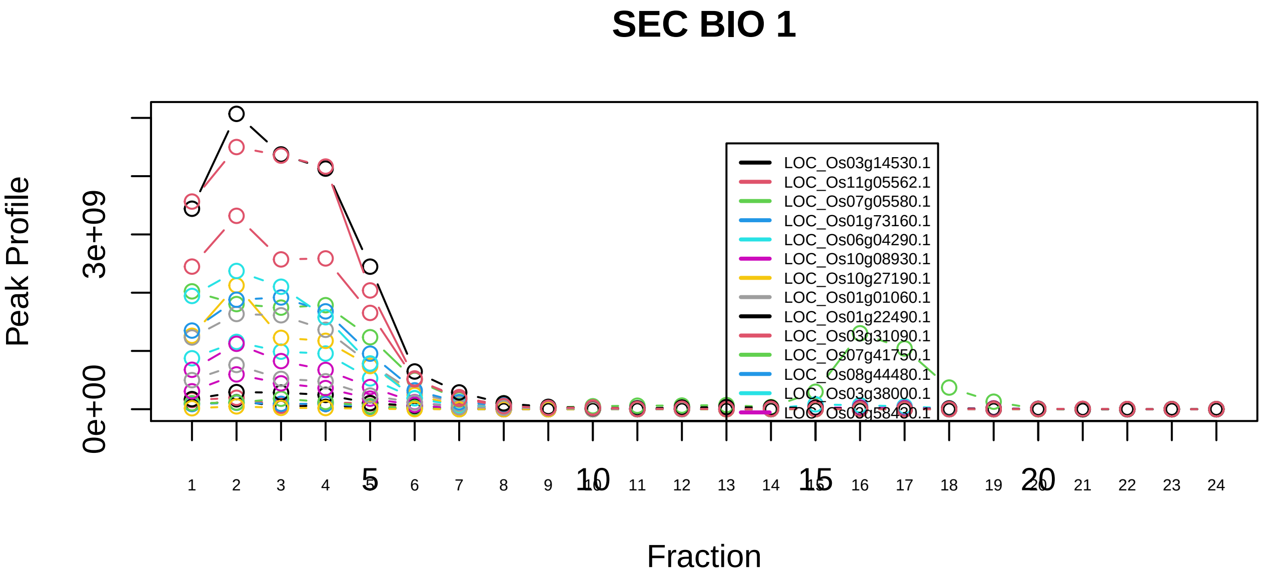

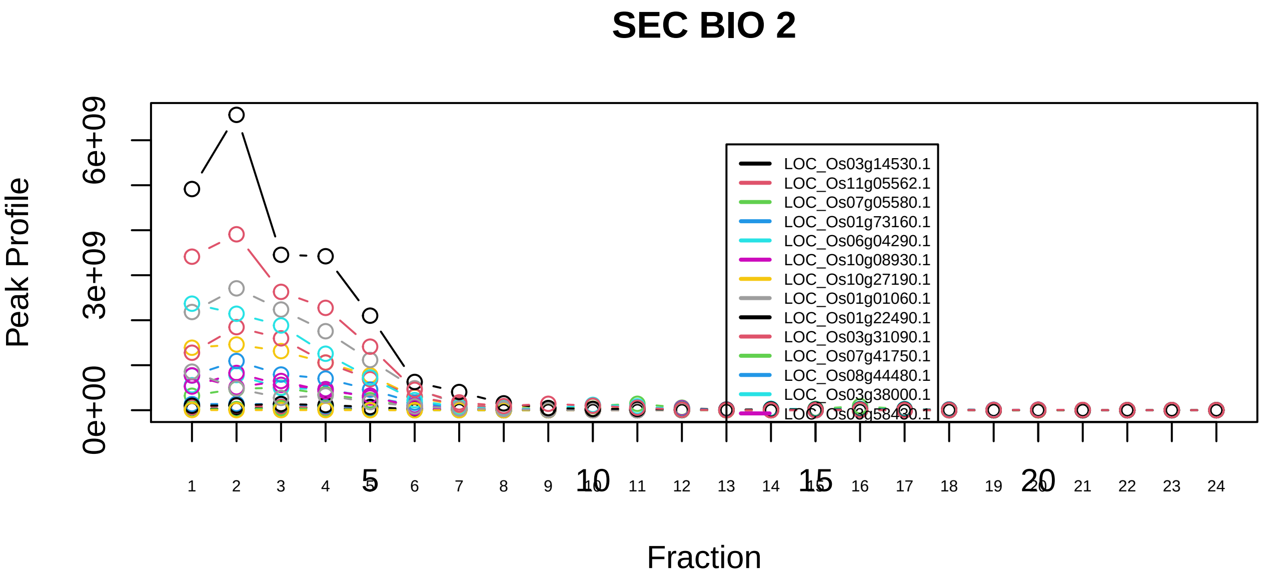

Ribosome, cytoplasmic|306−−sub_5

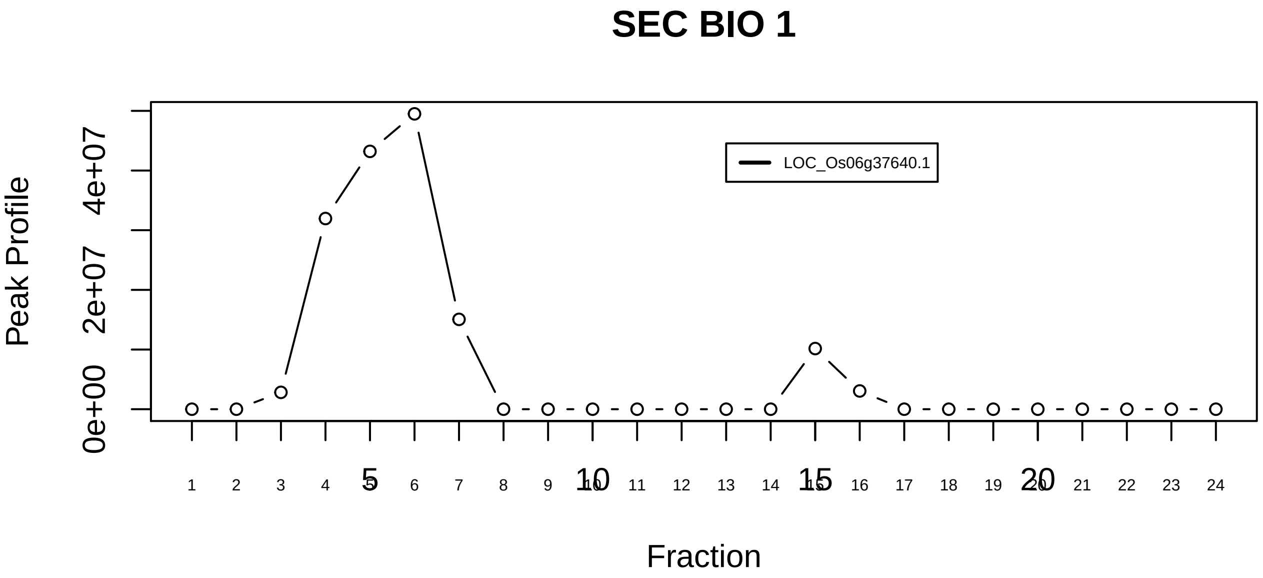

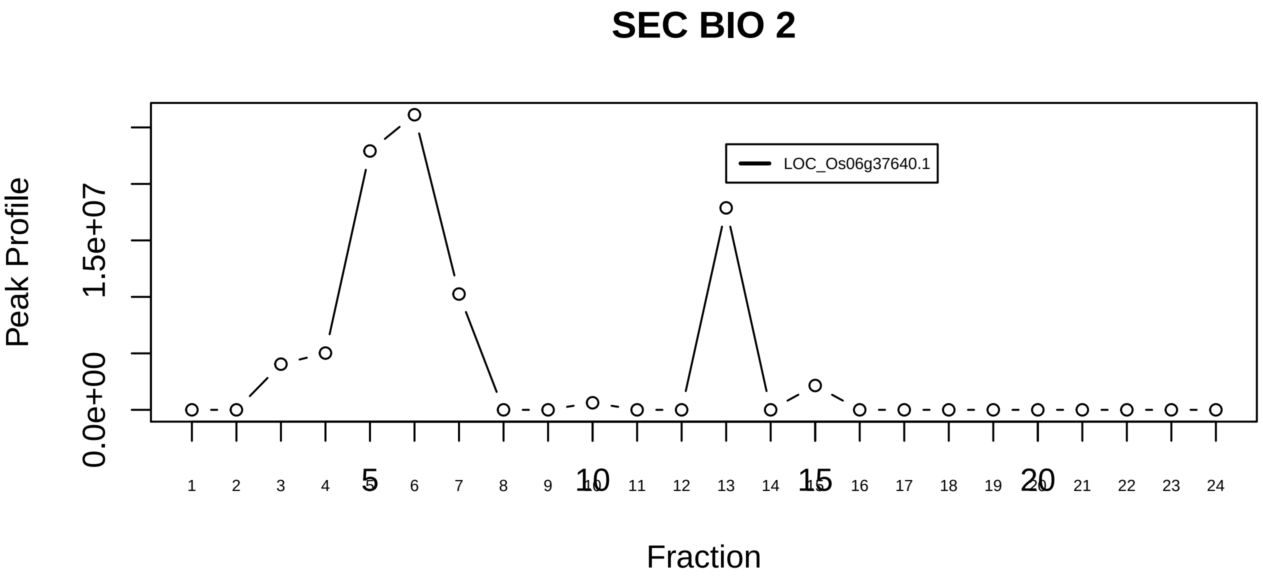

RNA polymerase II (RNAPII)|2685−−singleton_1

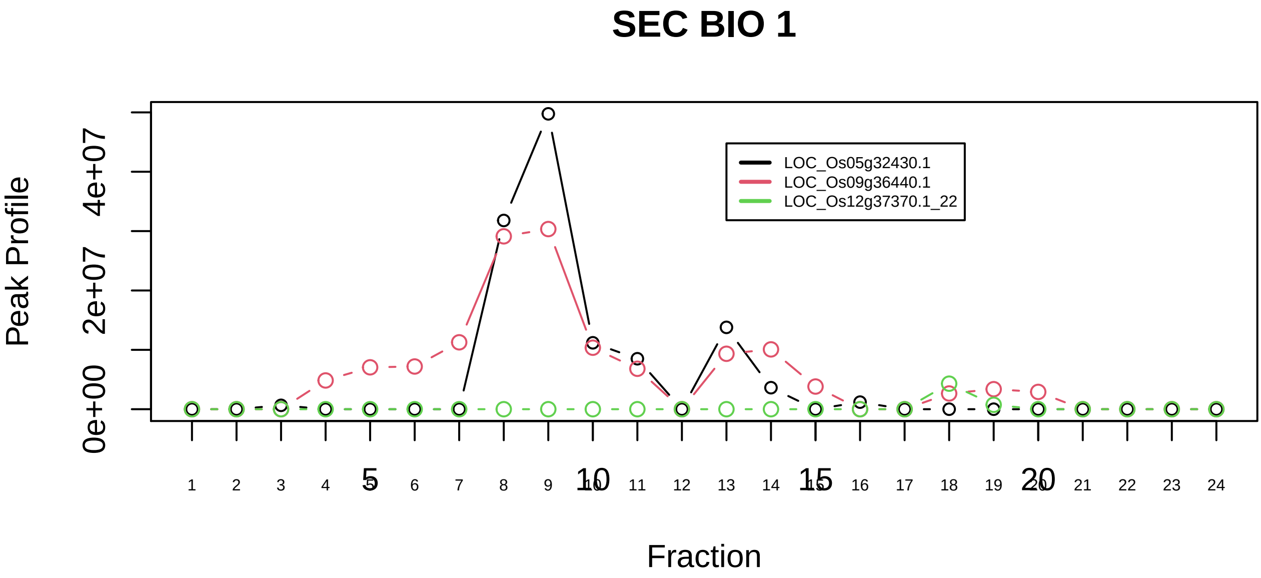

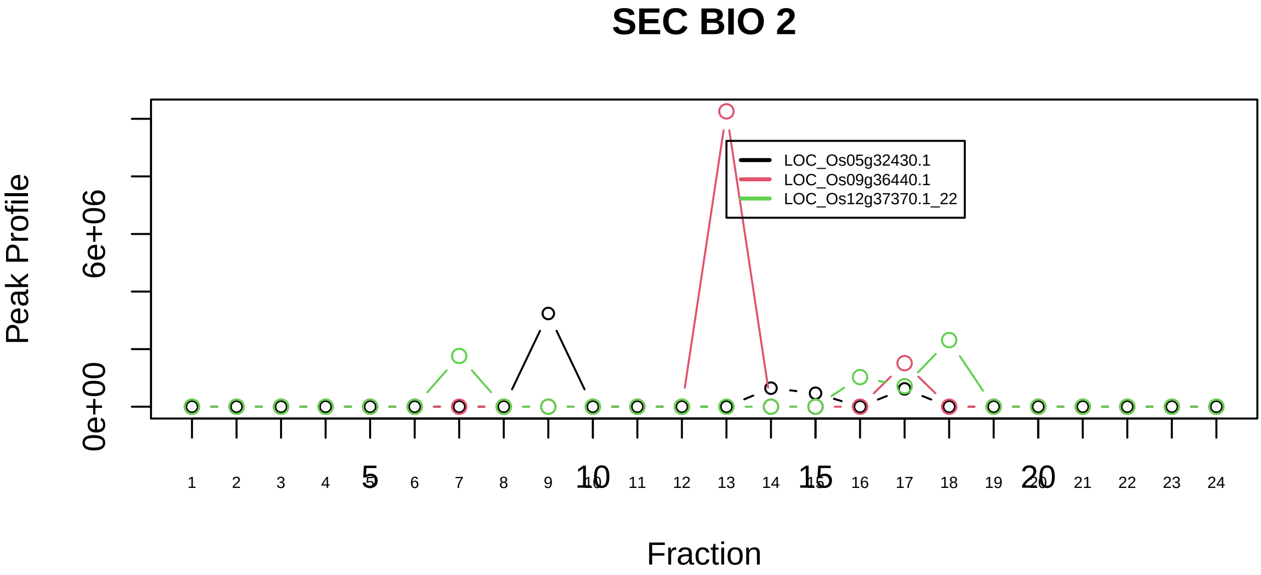

RNA polymerase II (RNAPII)|2685−−sub_1

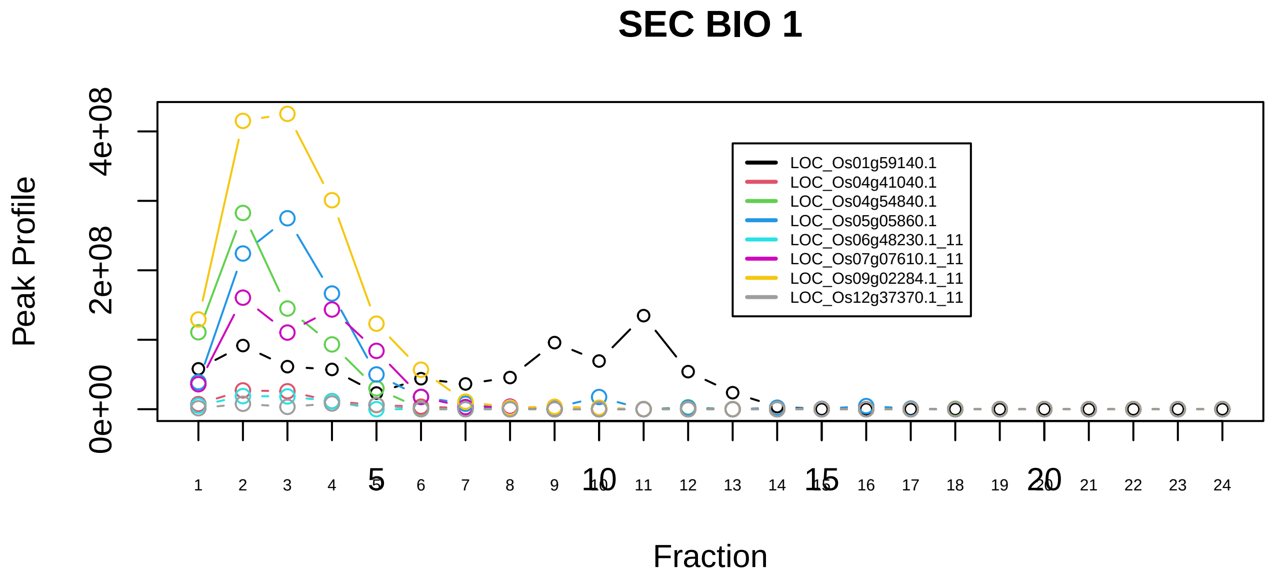

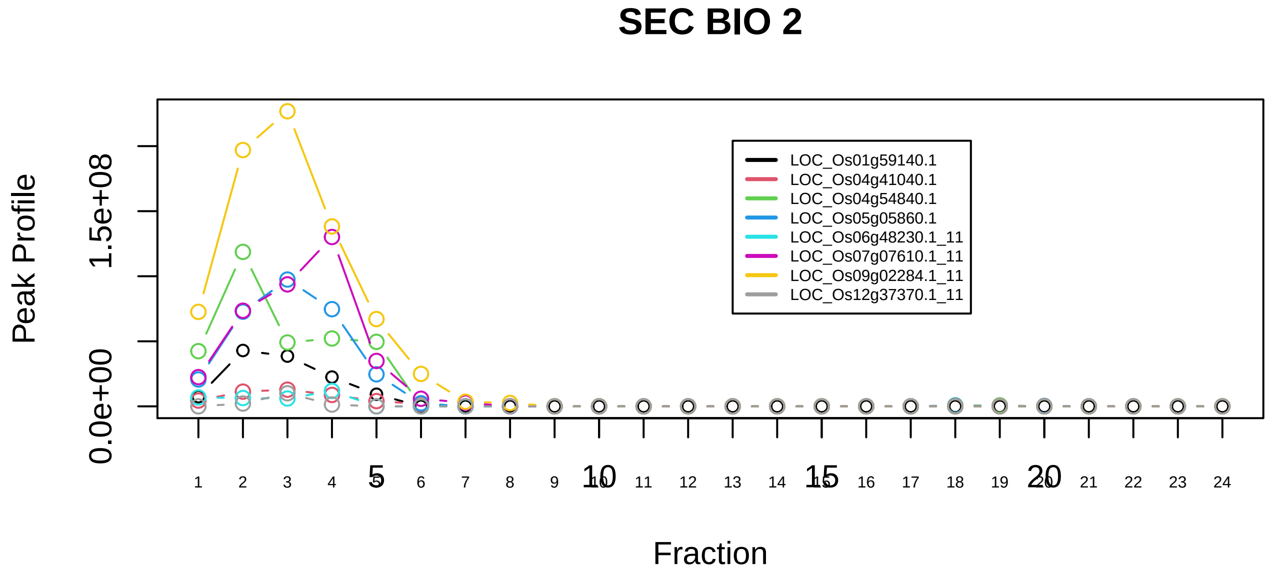

RNA polymerase II (RNAPII)|2685−−sub_2

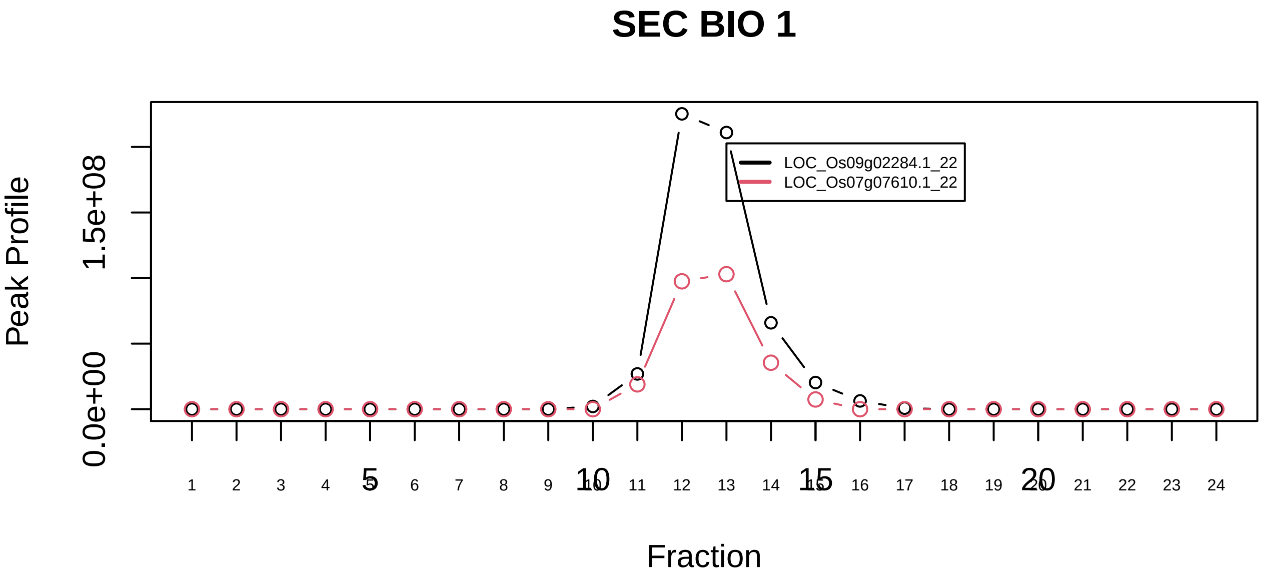

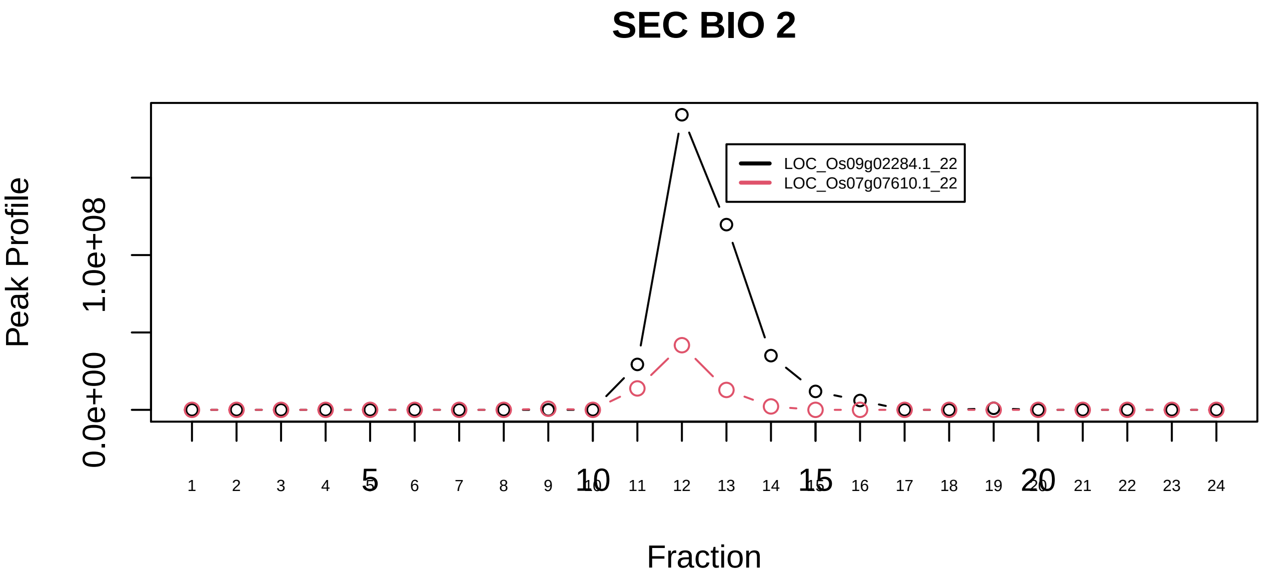

RNA polymerase II (RNAPII)|2685−−sub_3

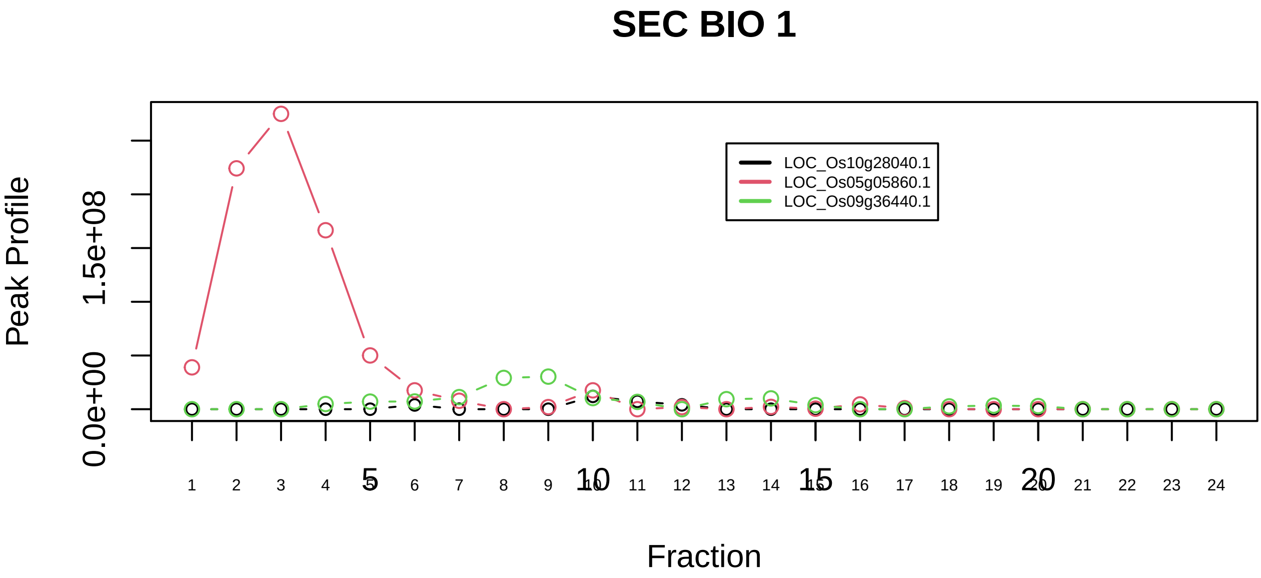

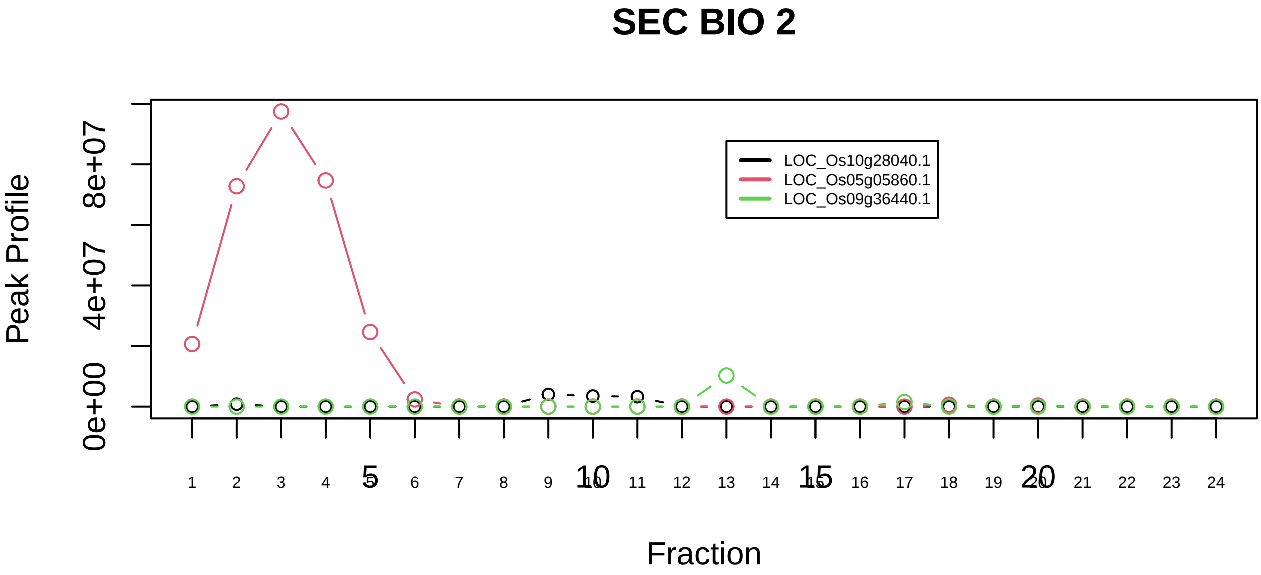

RNA polymerase II complex, chromatin structure modifying|3066−−sub_1

### SEC BIO 1

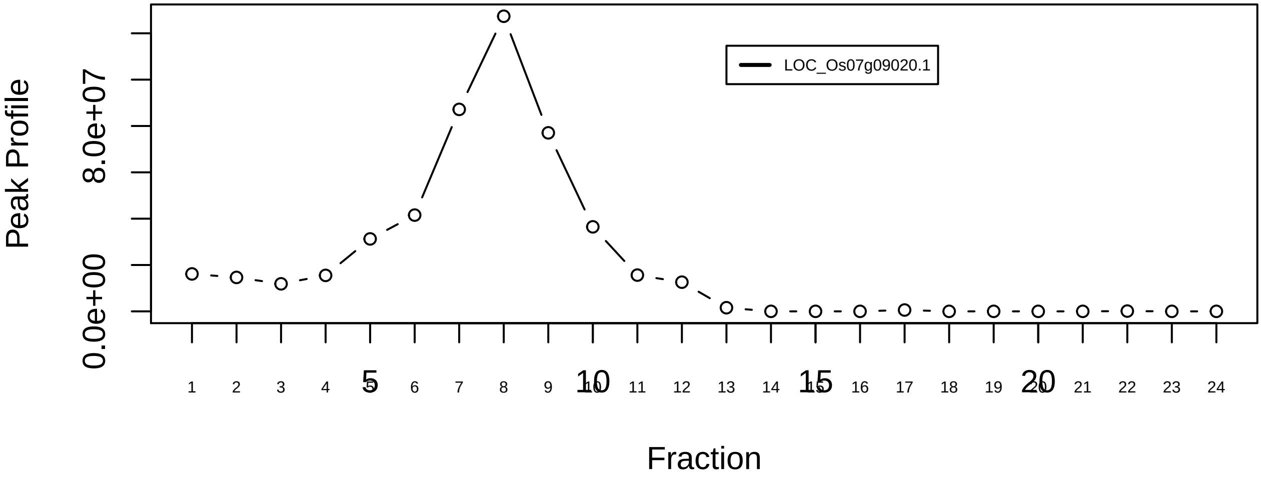

### SEC BIO 2

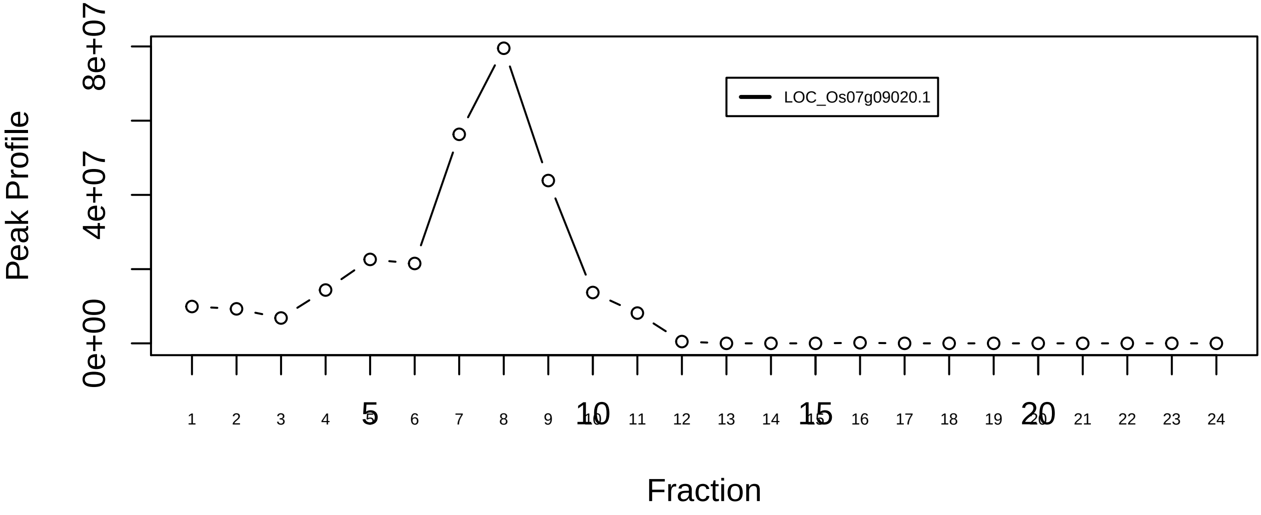

RNA−induced silencing complex, RISC|3032−−singleton_1

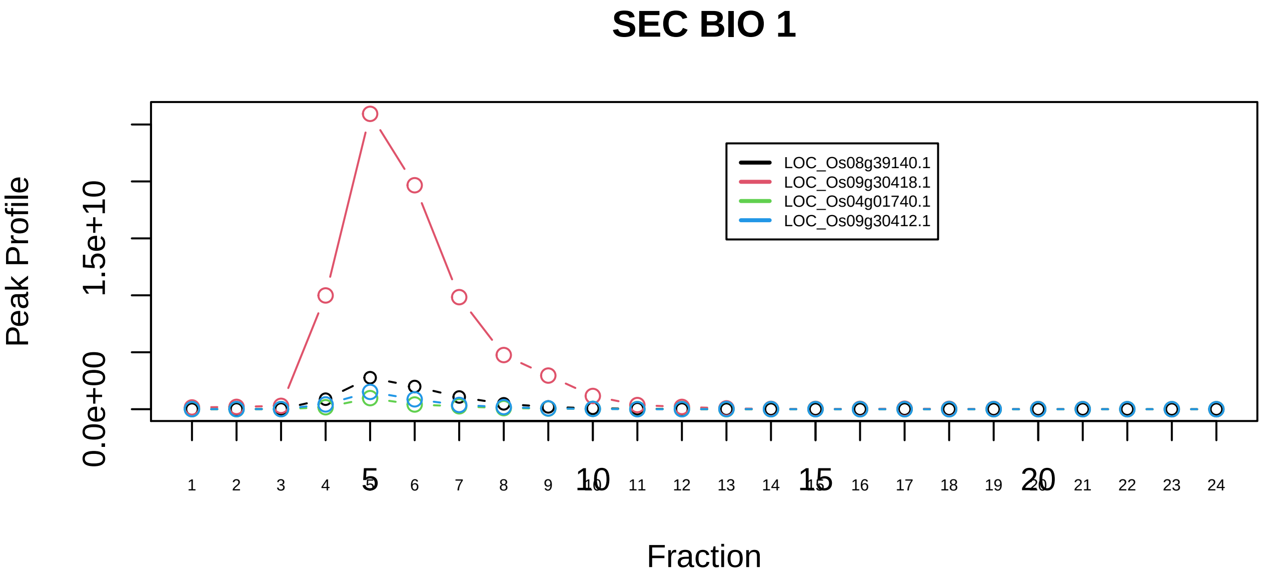

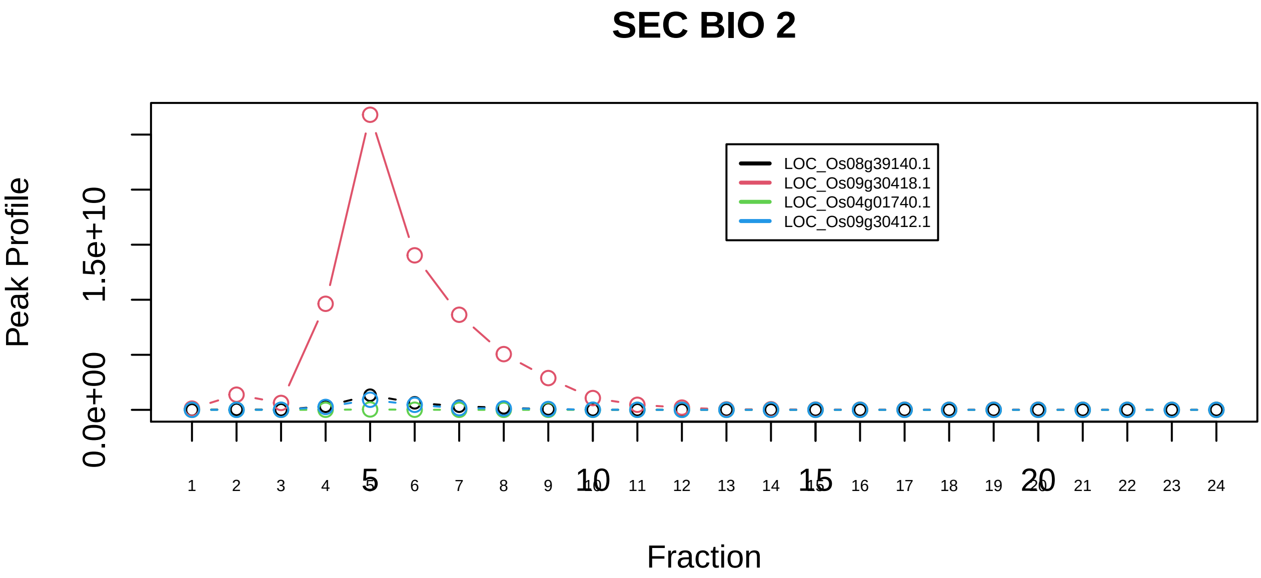

RNA−induced silencing complex, RISC|3032−−sub_orthoparalog_1

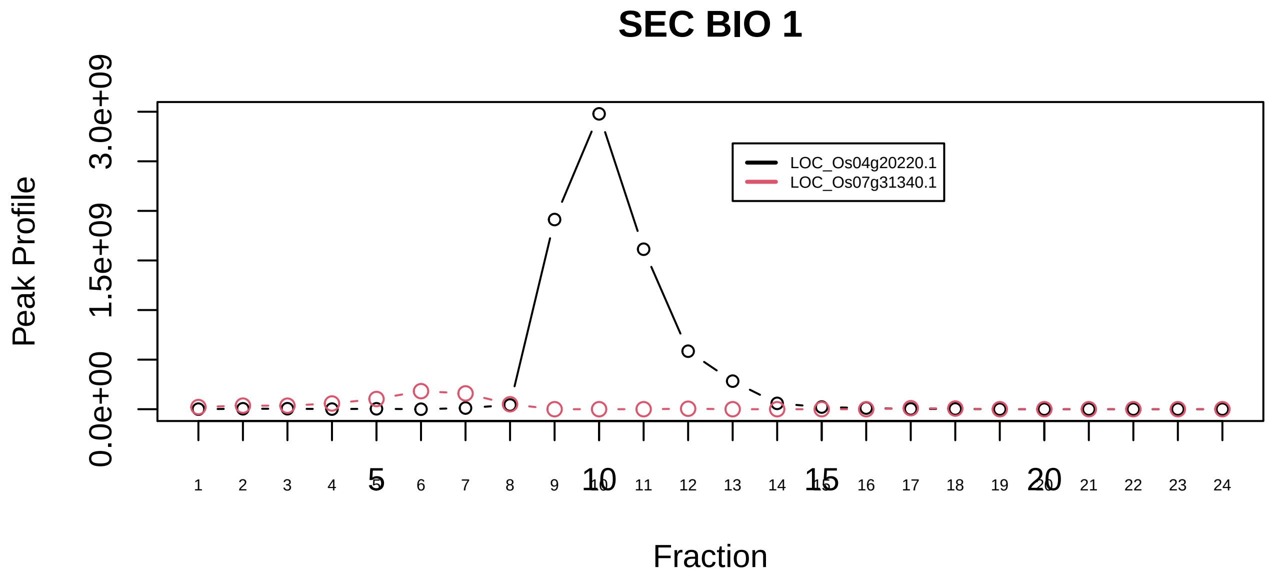

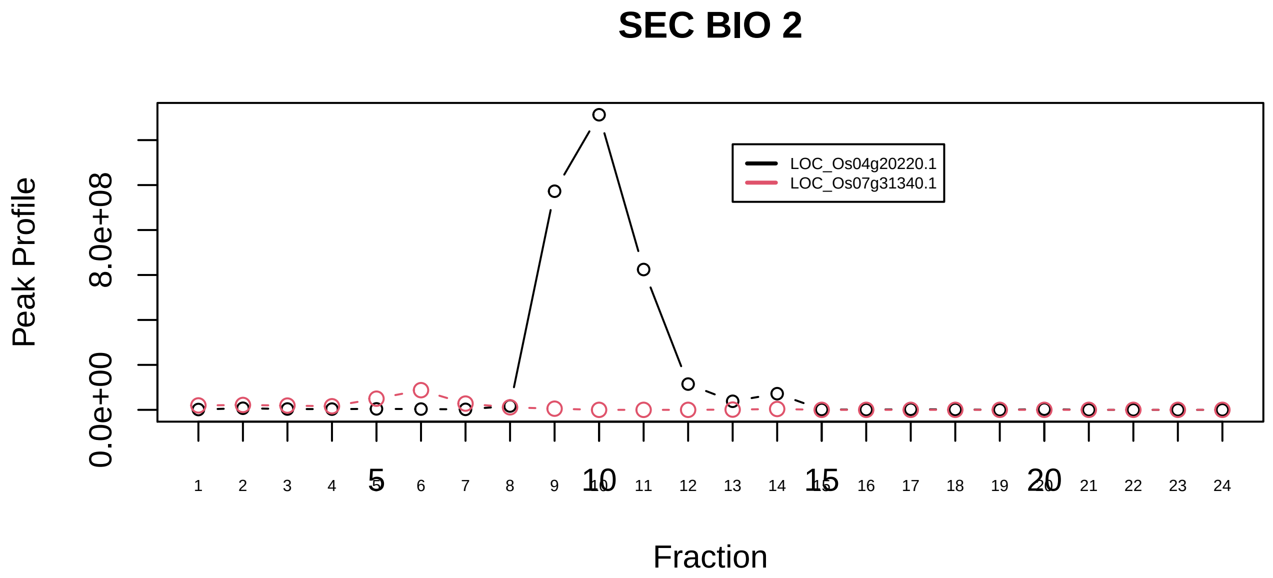

SMG−1−UPF−ERF1−ERF3 complex (SURF)|784−−only_2_prots_before_clustering

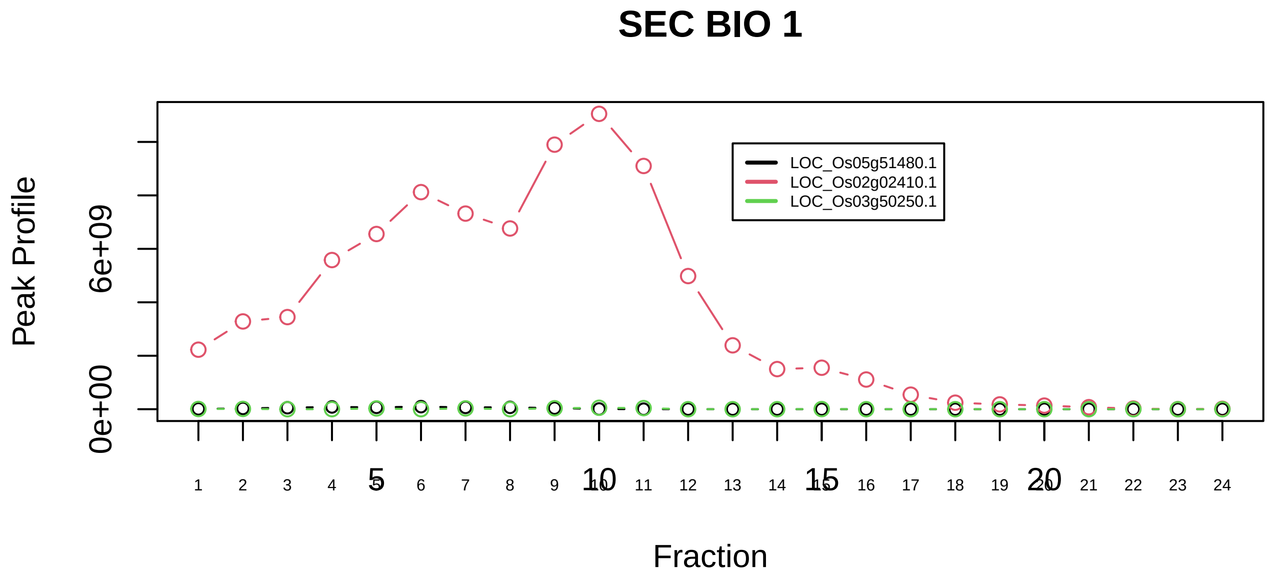

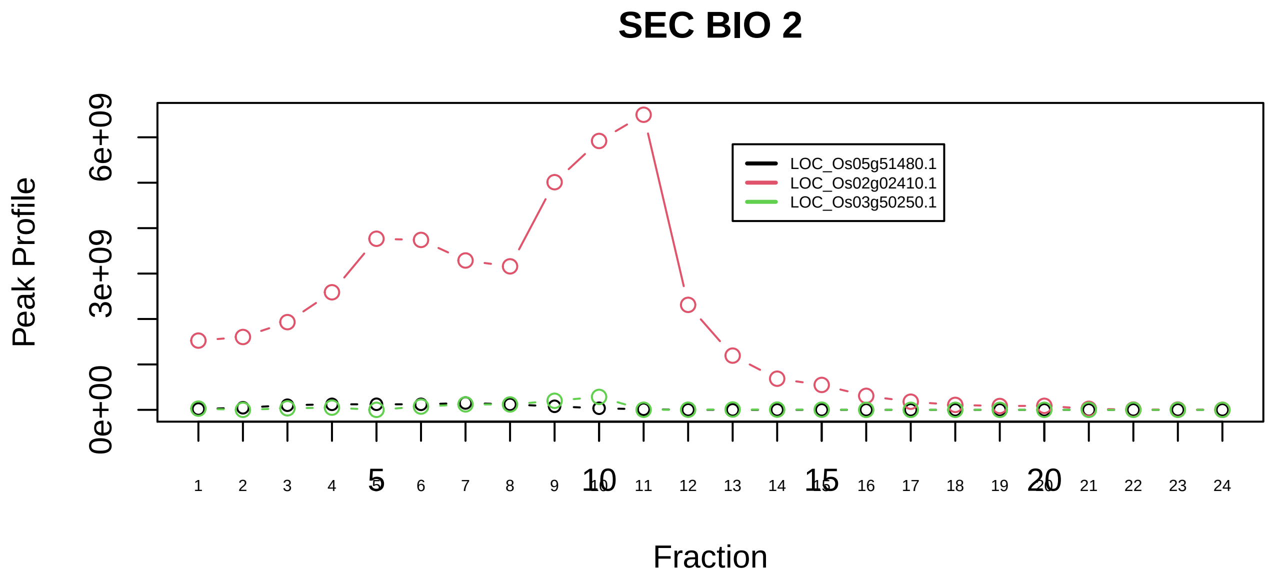

SNW1 complex|1335−−sub_1

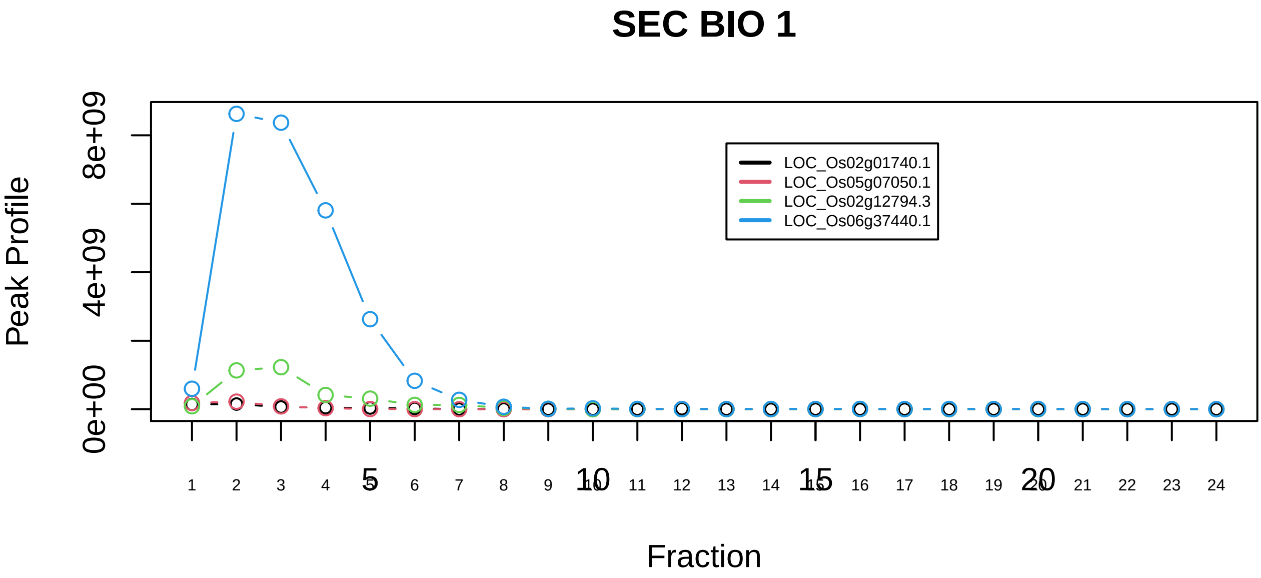

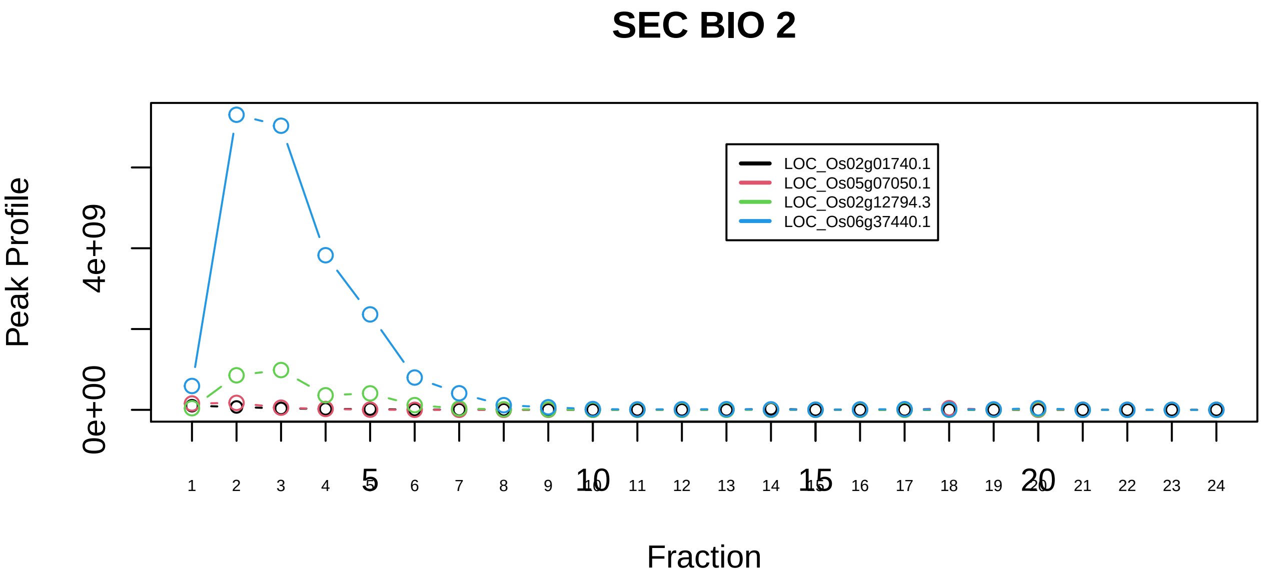

SNW1 complex|1335−−sub_2

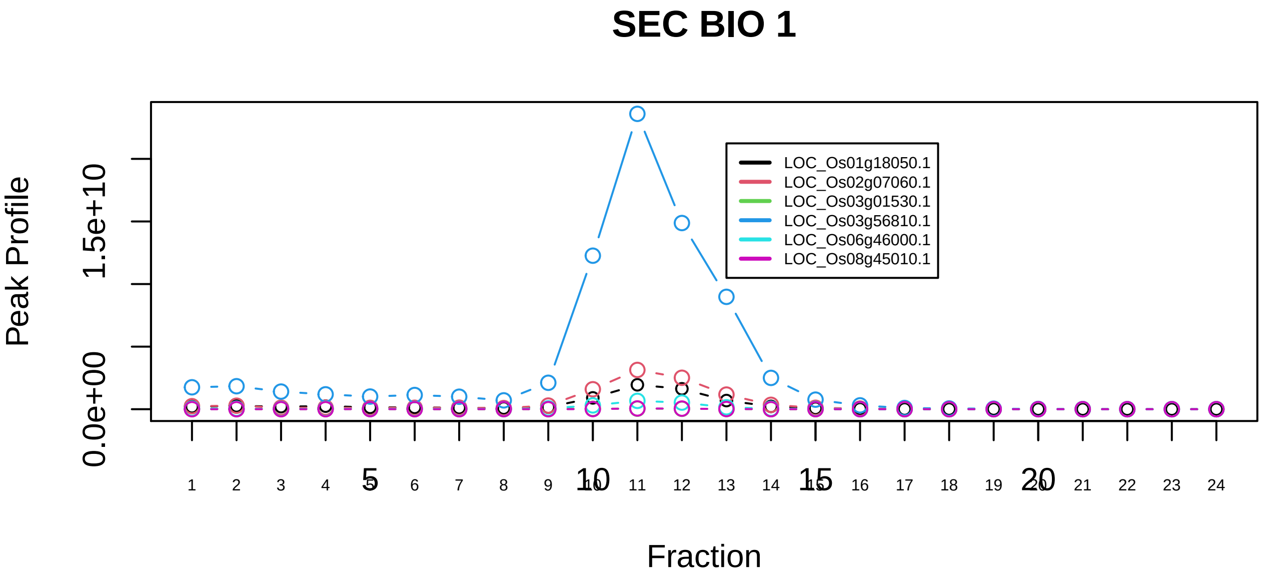

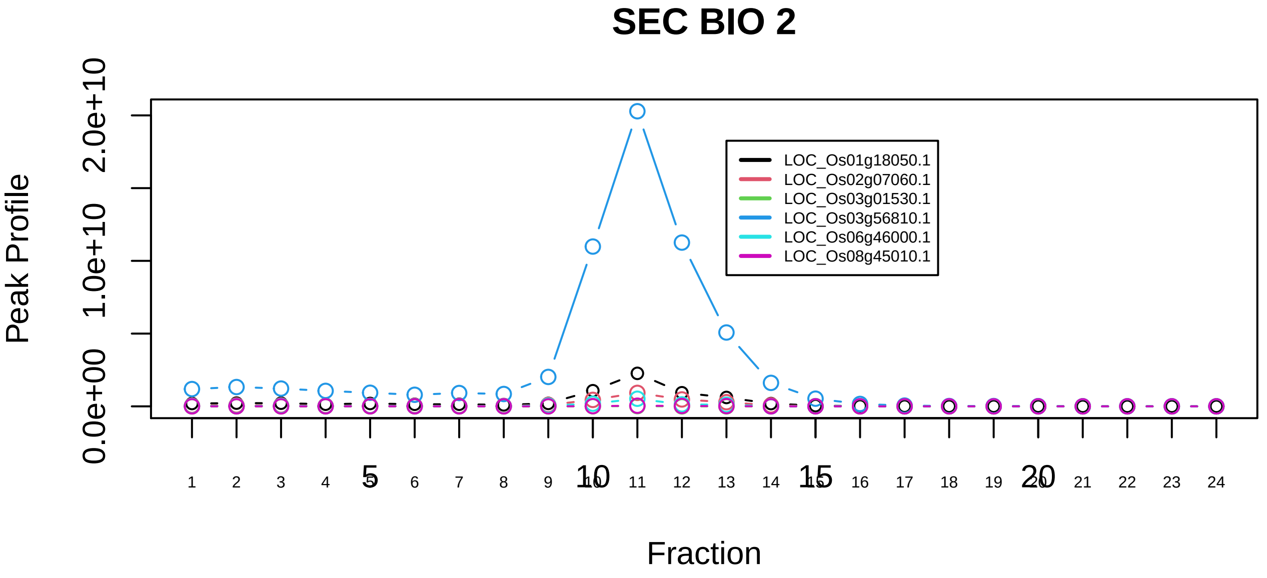

SNW1 complex|1335−−sub_3

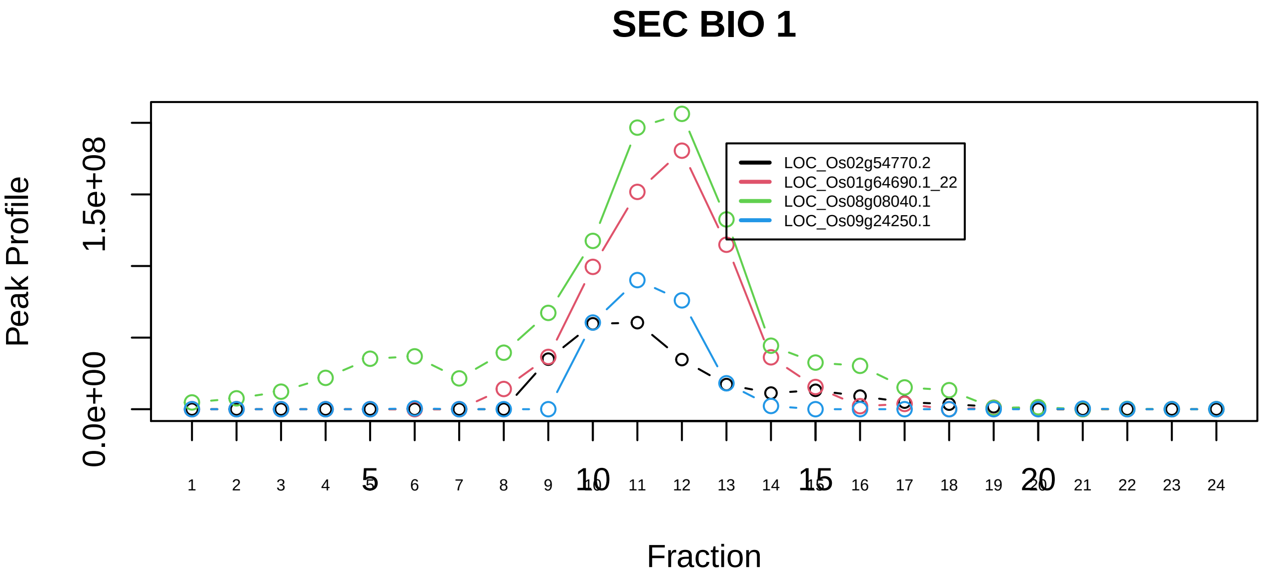

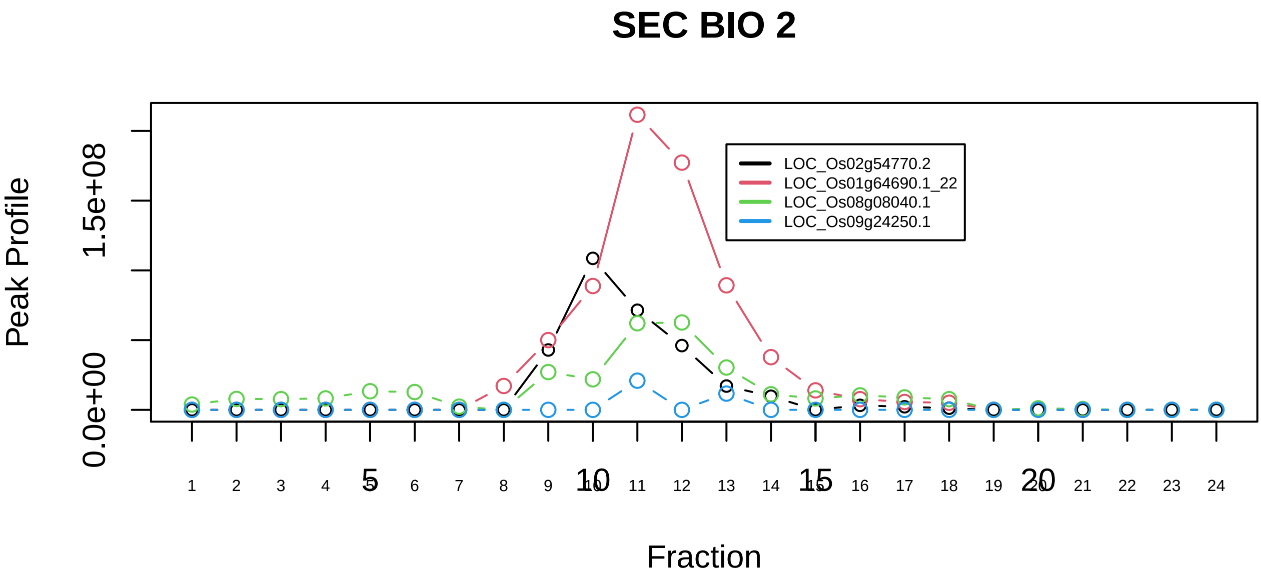

Spliceosome|351−−sub_1

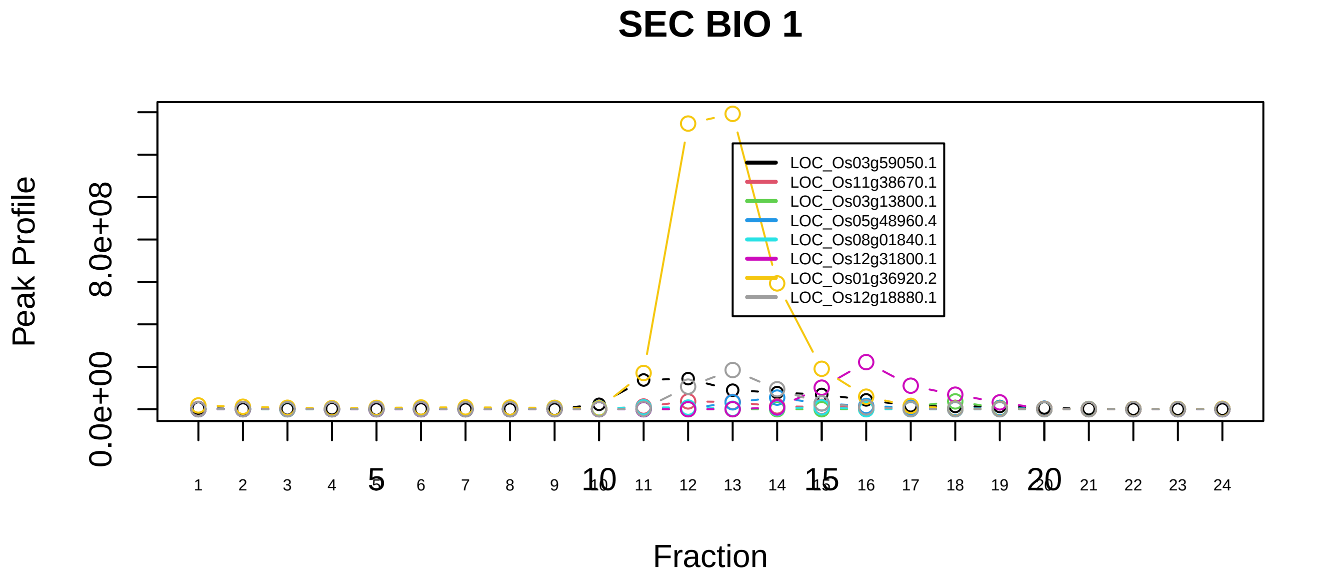

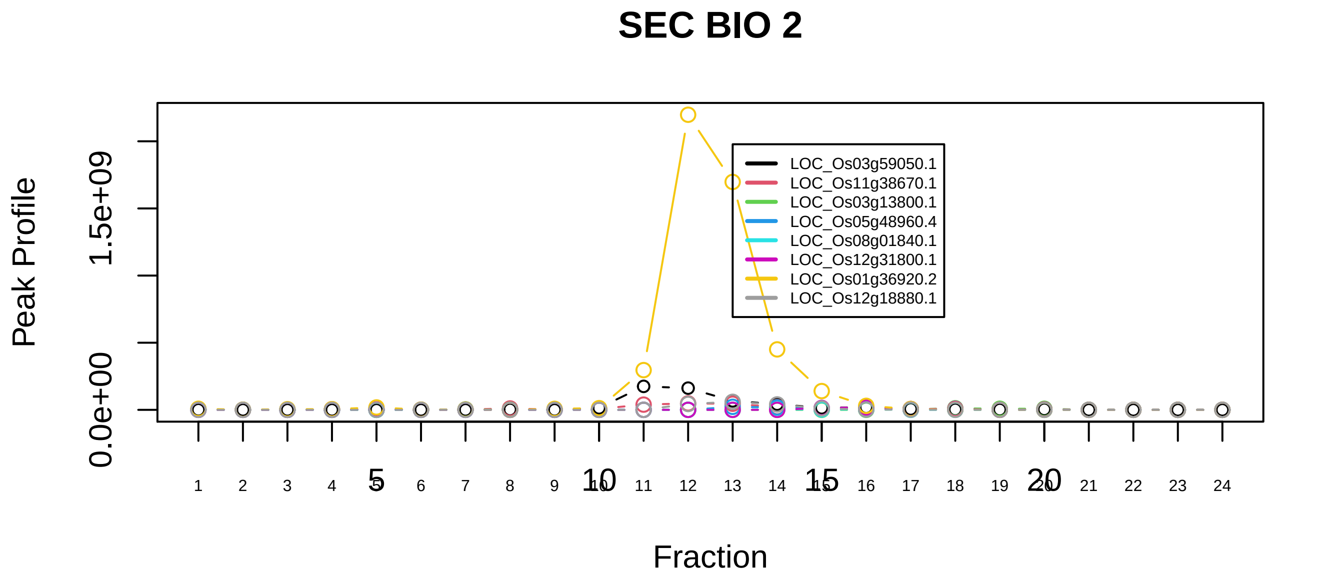

Spliceosome|351−−sub_2

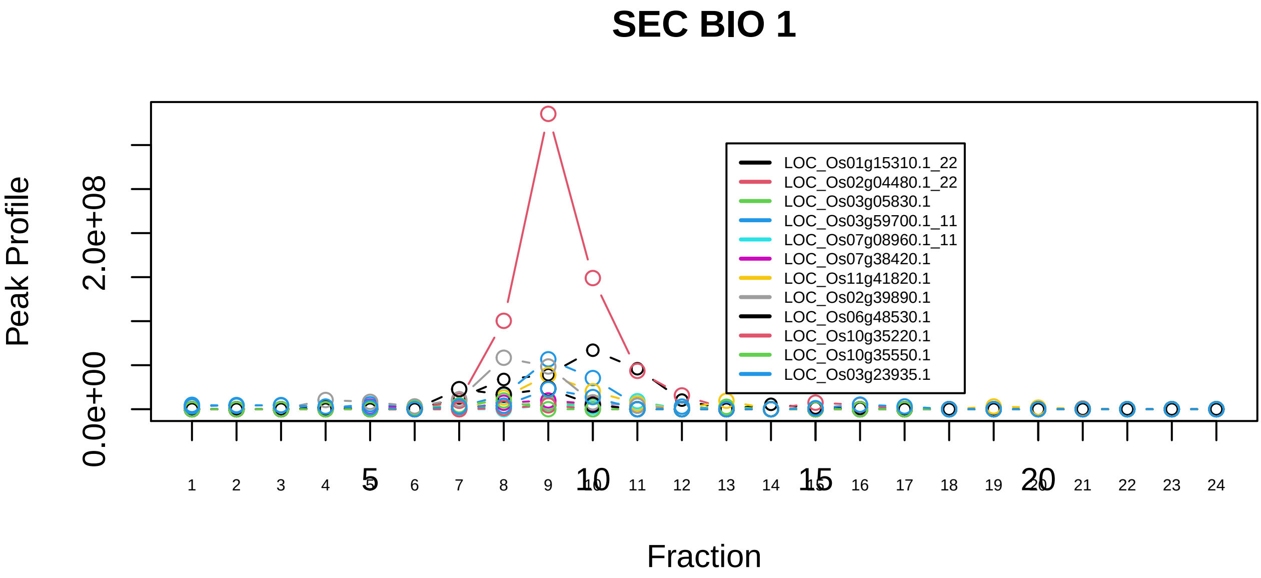

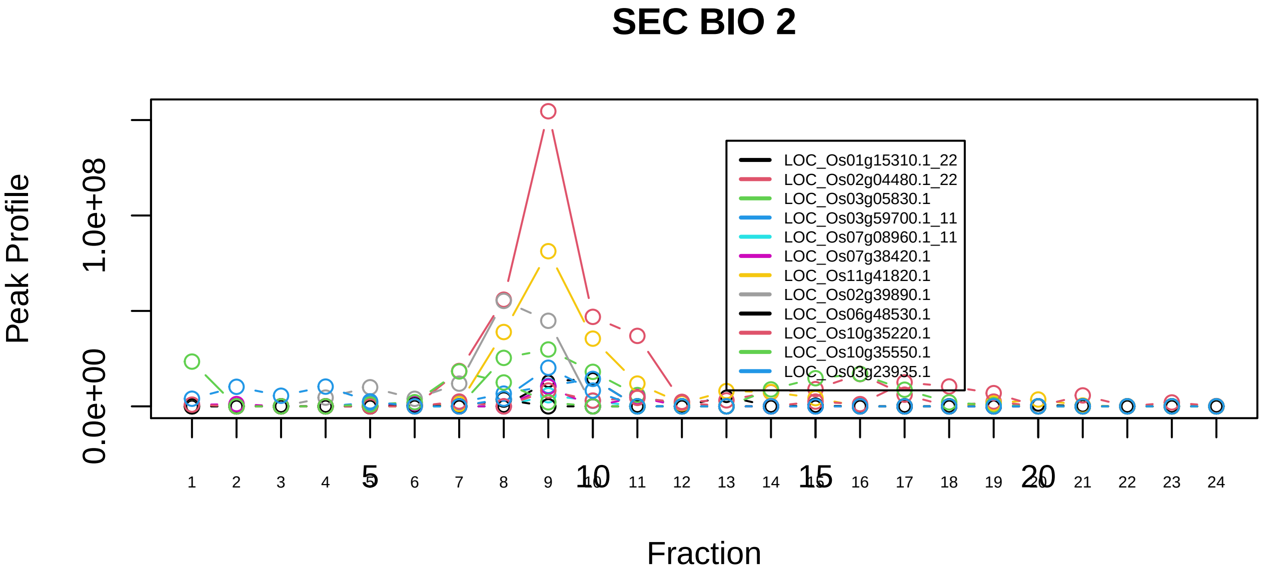

Spliceosome|351−−sub_3

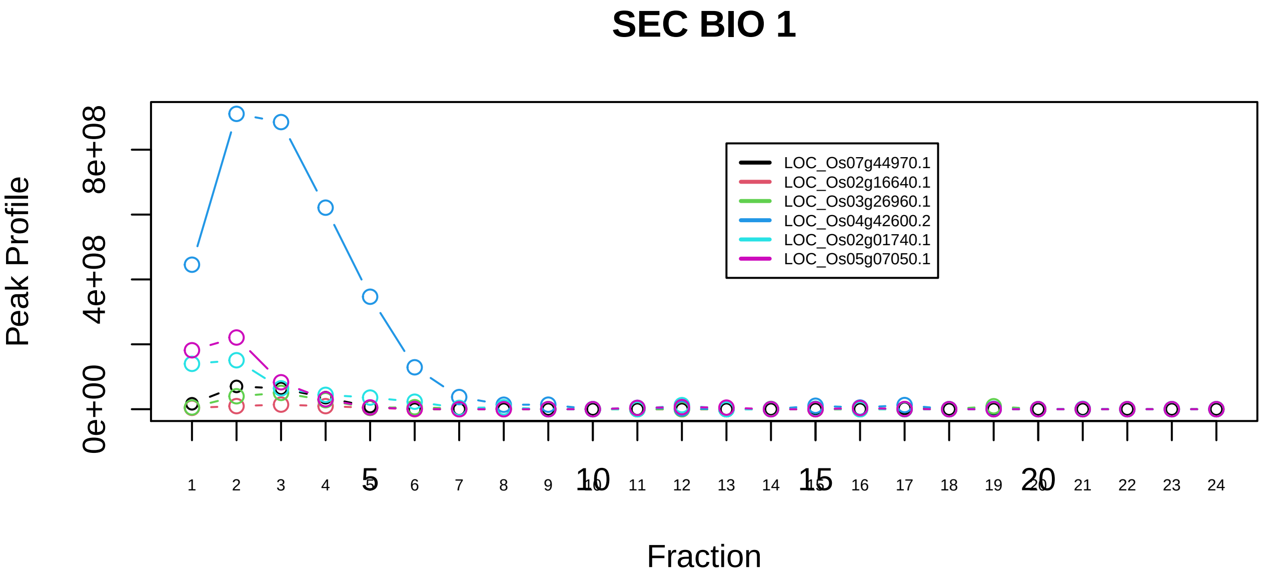

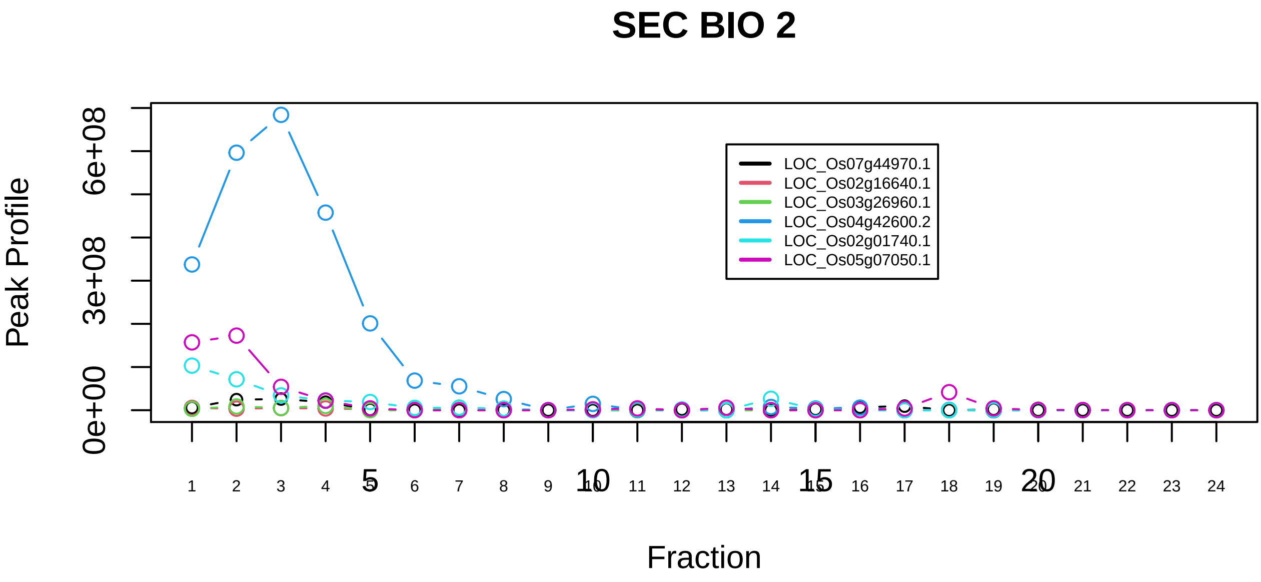

Spliceosome|351−−sub_4

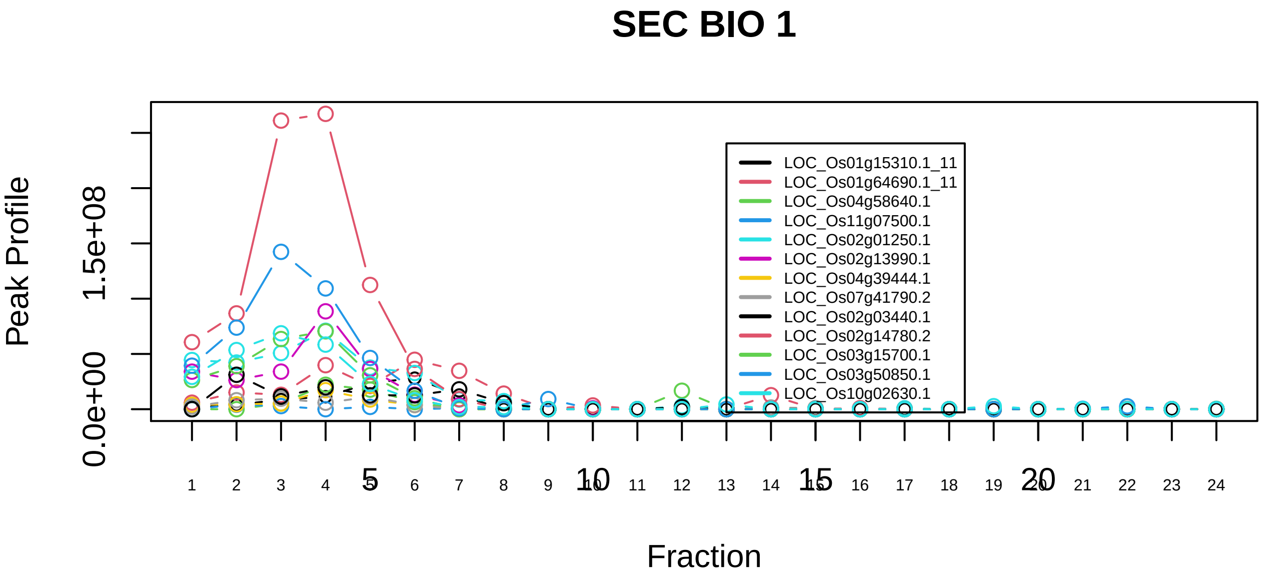

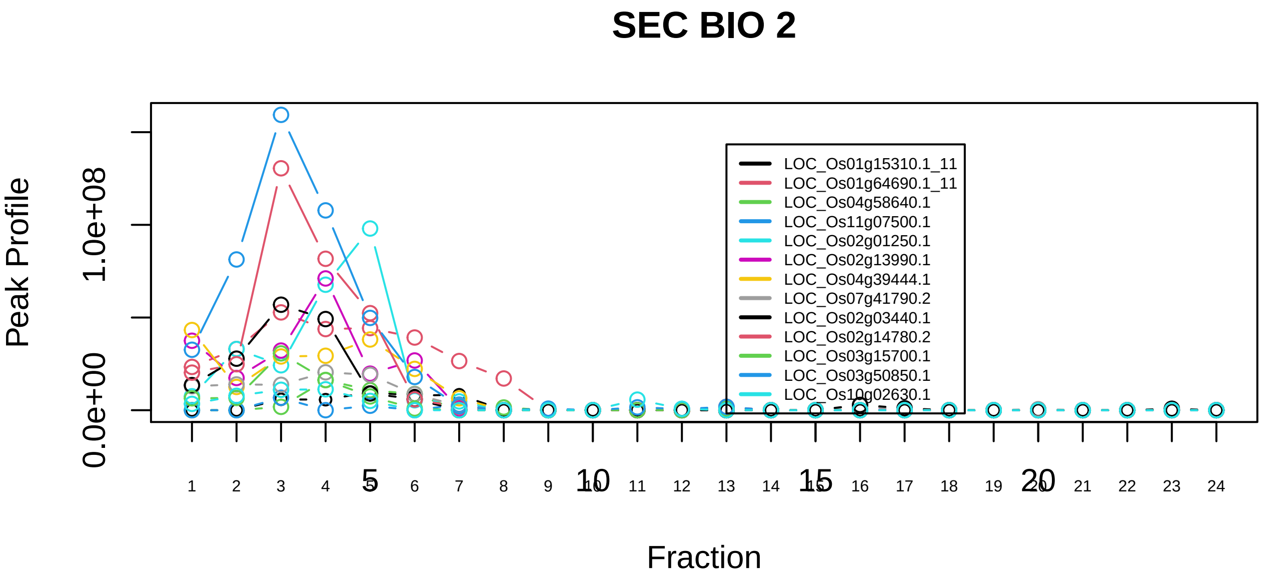

Spliceosome|351−−sub_5

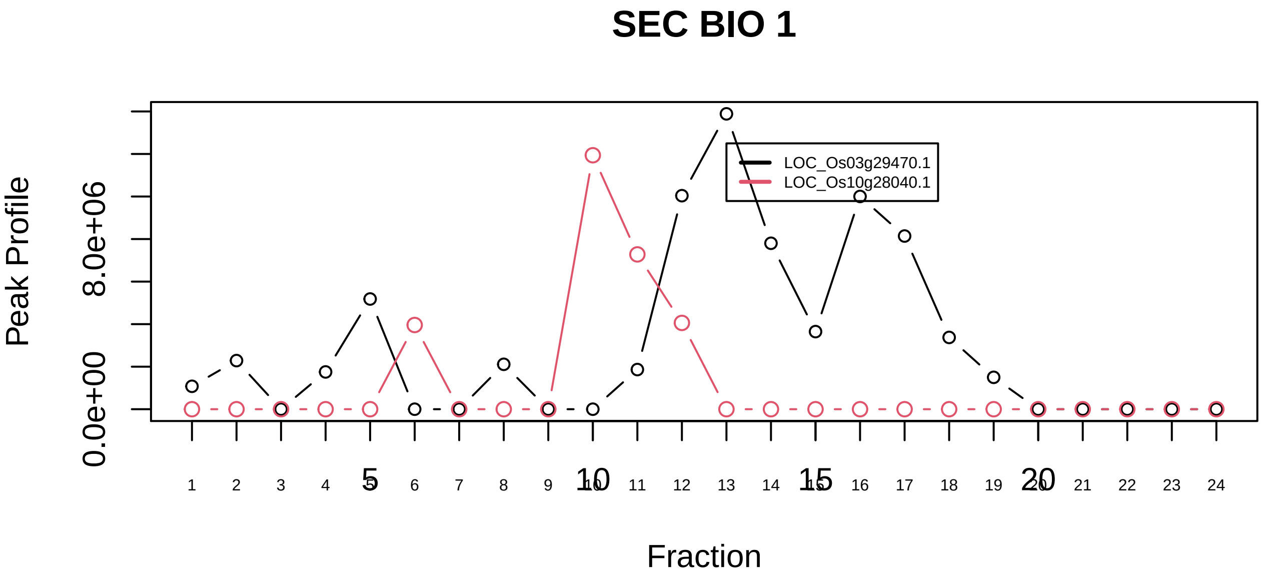

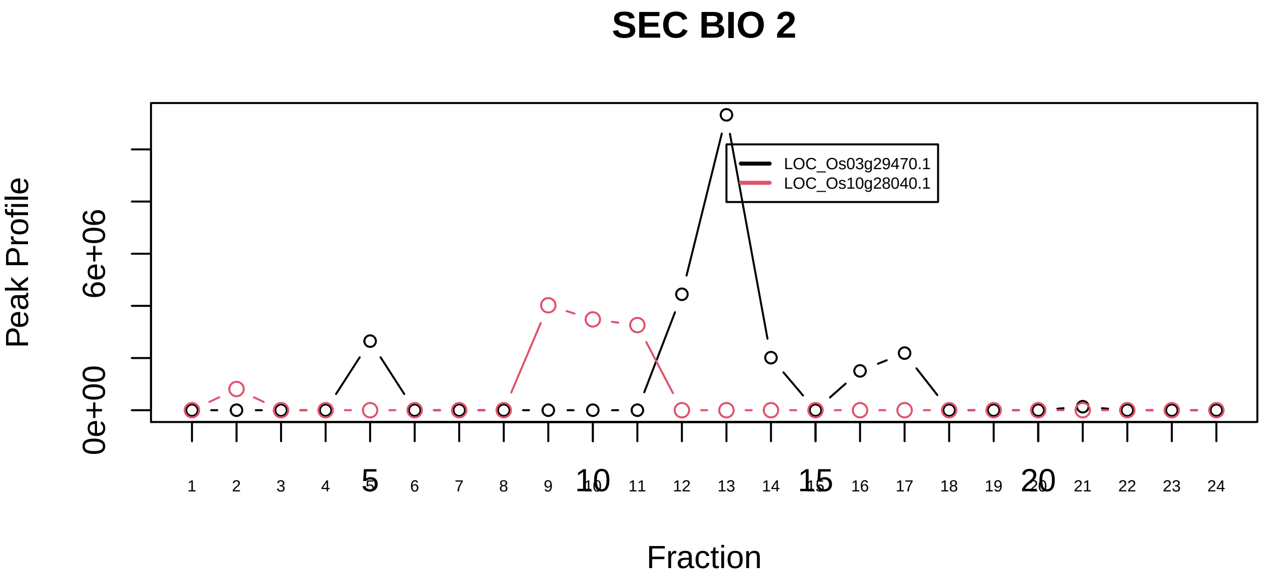

STAGA core complex|478−−only_2_prots_before_clustering

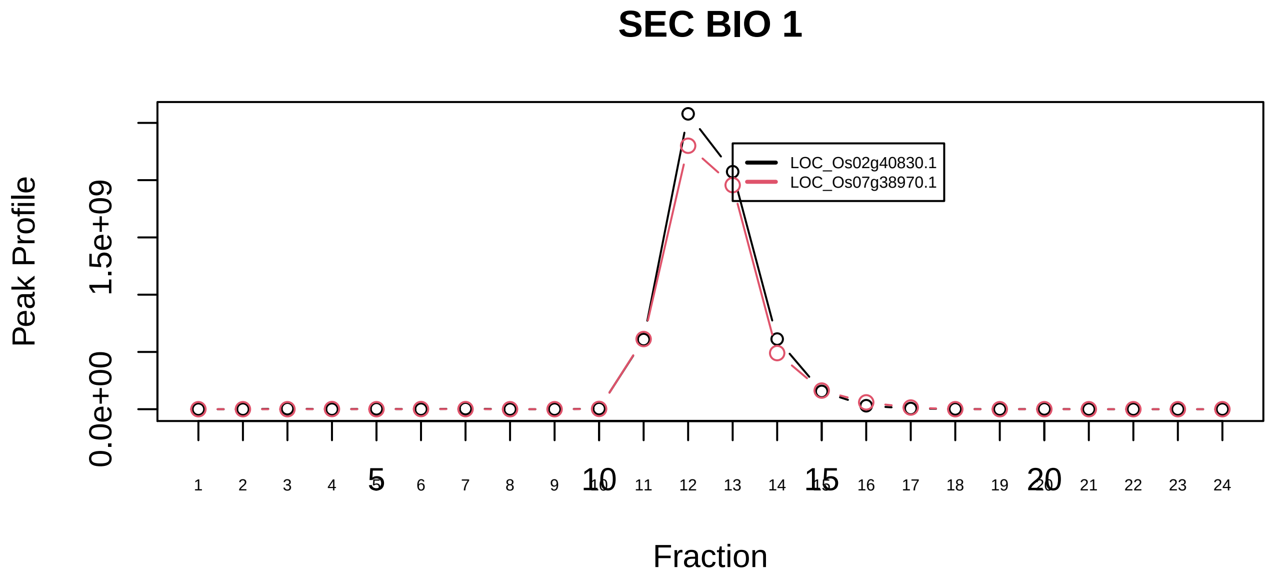

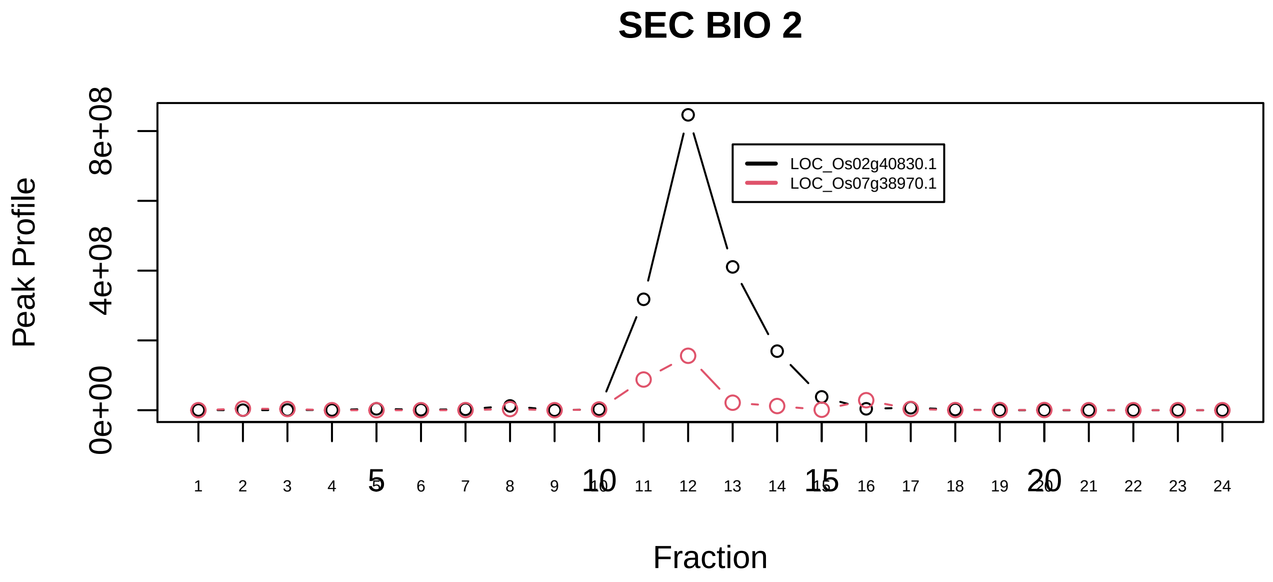

Succinyl−CoA synthetase, ADP−forming|394−−only_2_prots_before_clustering

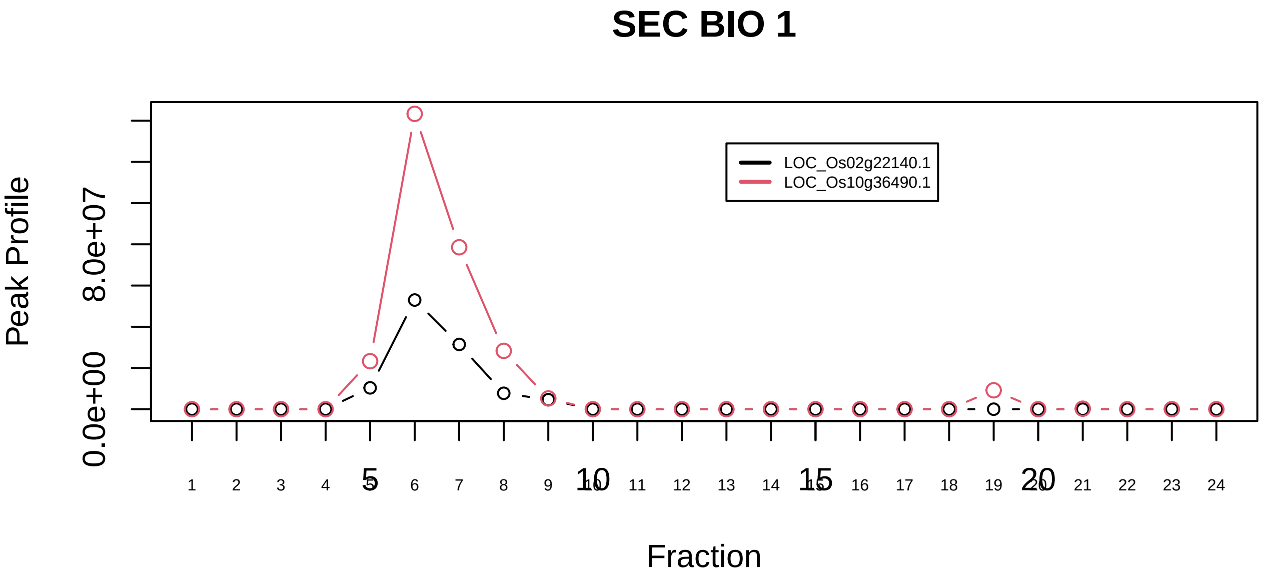

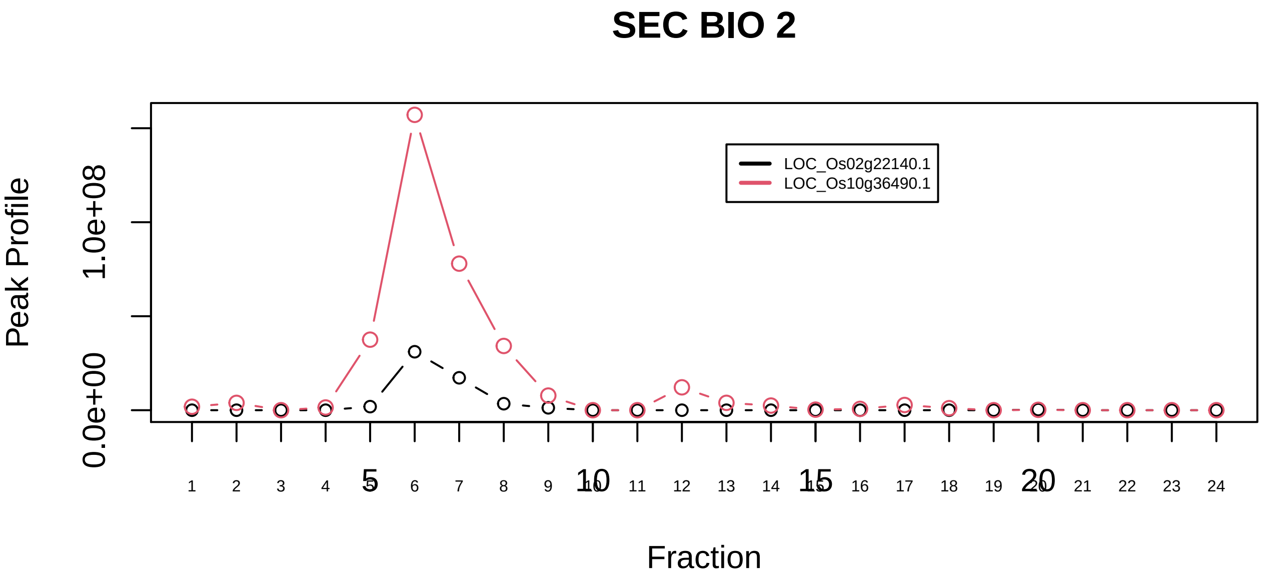

TBCD−ARL2−tubulin(beta)−TBCE complex|6715−−sub_1

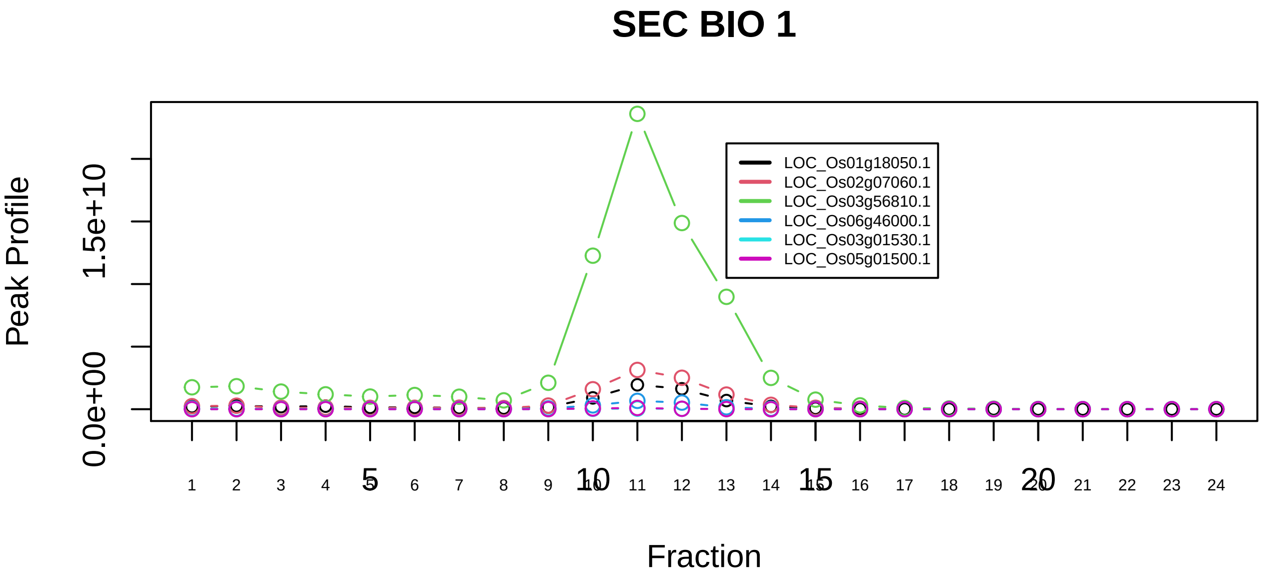

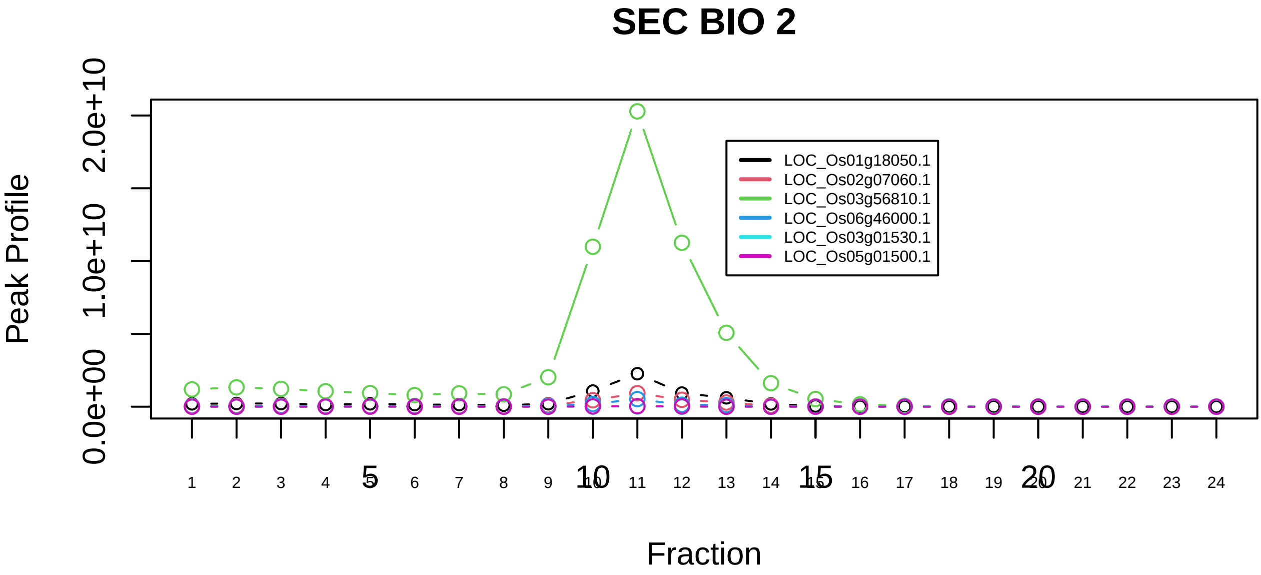

TBCD−ARL2−tubulin(beta)−TBCE complex|6715−−sub_2

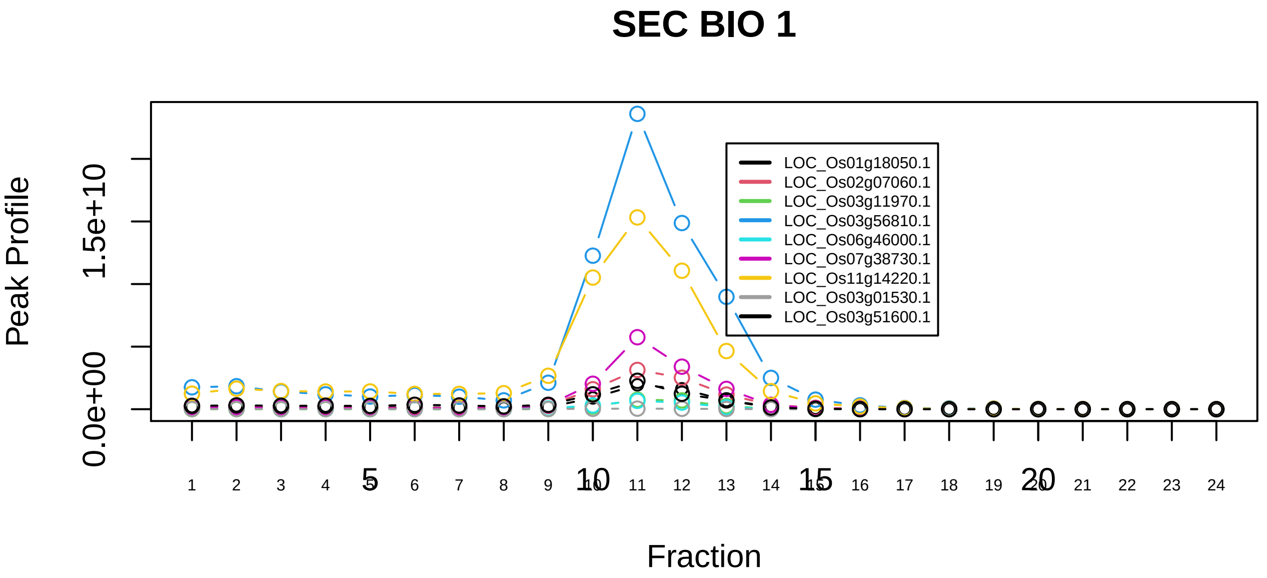

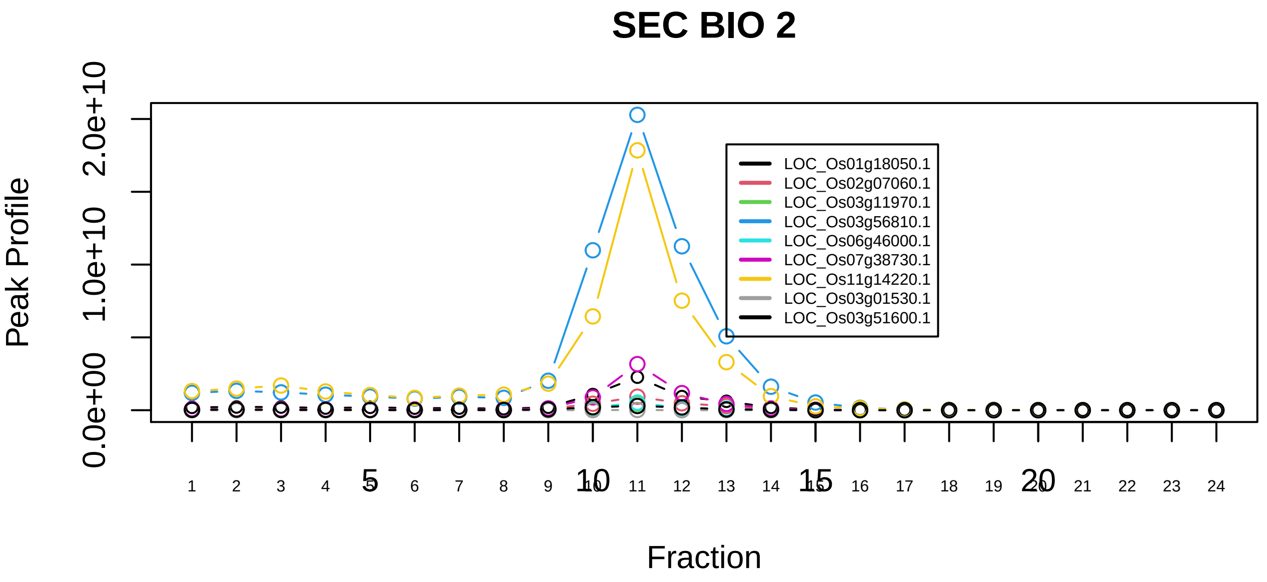

TBCD−tubulin(alpha)−tubulin(beta) complex|6703−−sub_1

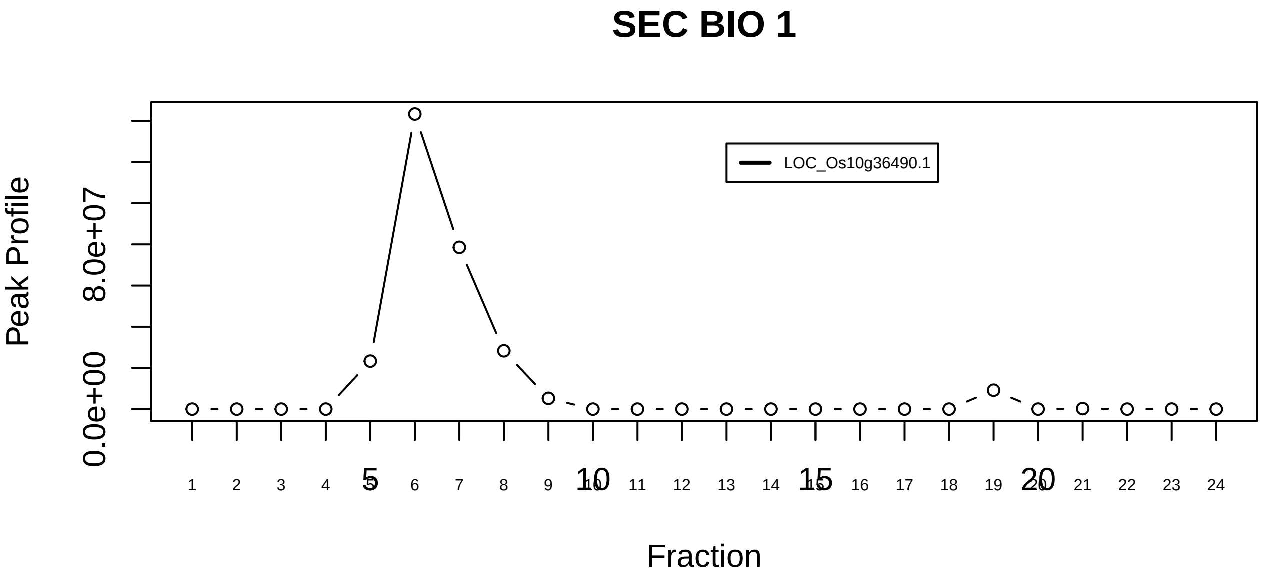

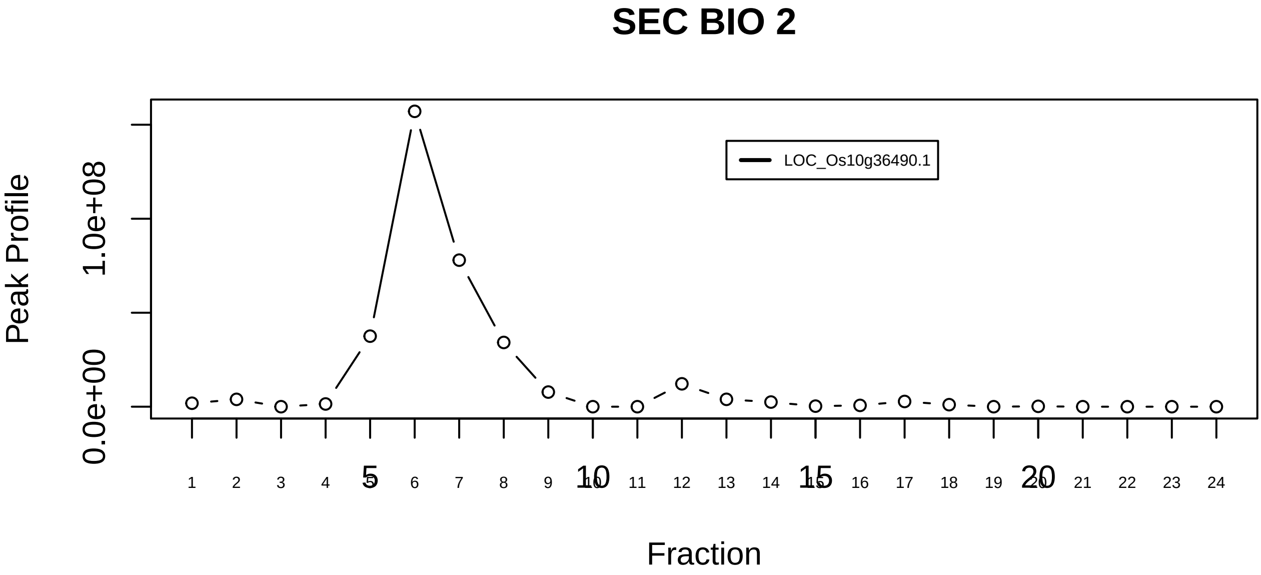

TBCD−tubulin(alpha)−tubulin(beta) complex|6703−−singleton_1

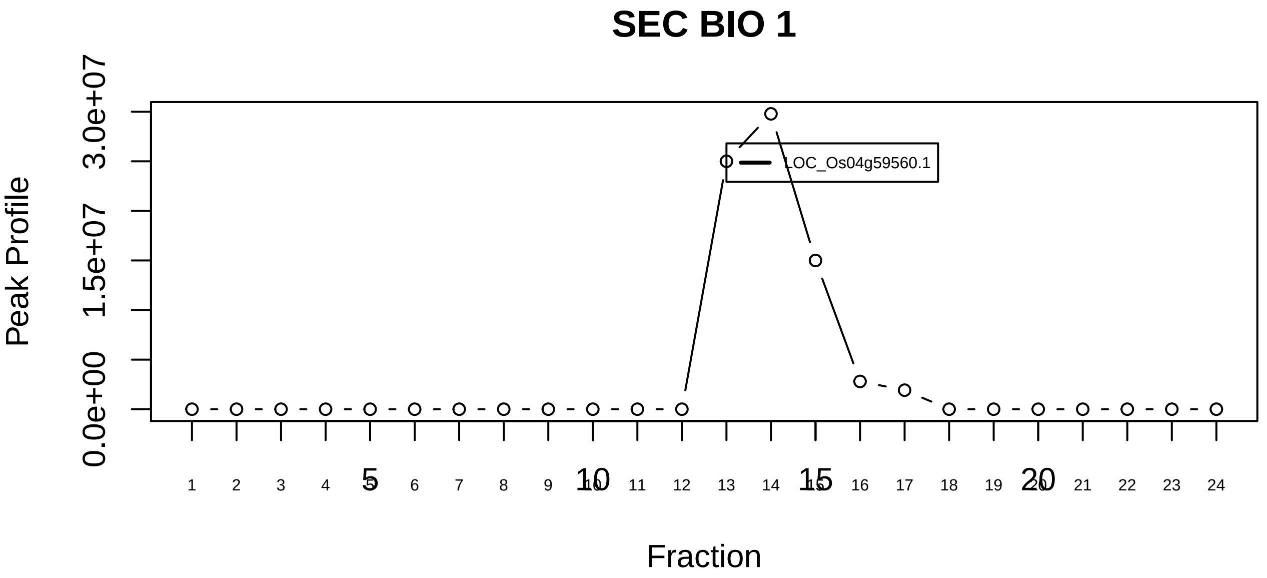

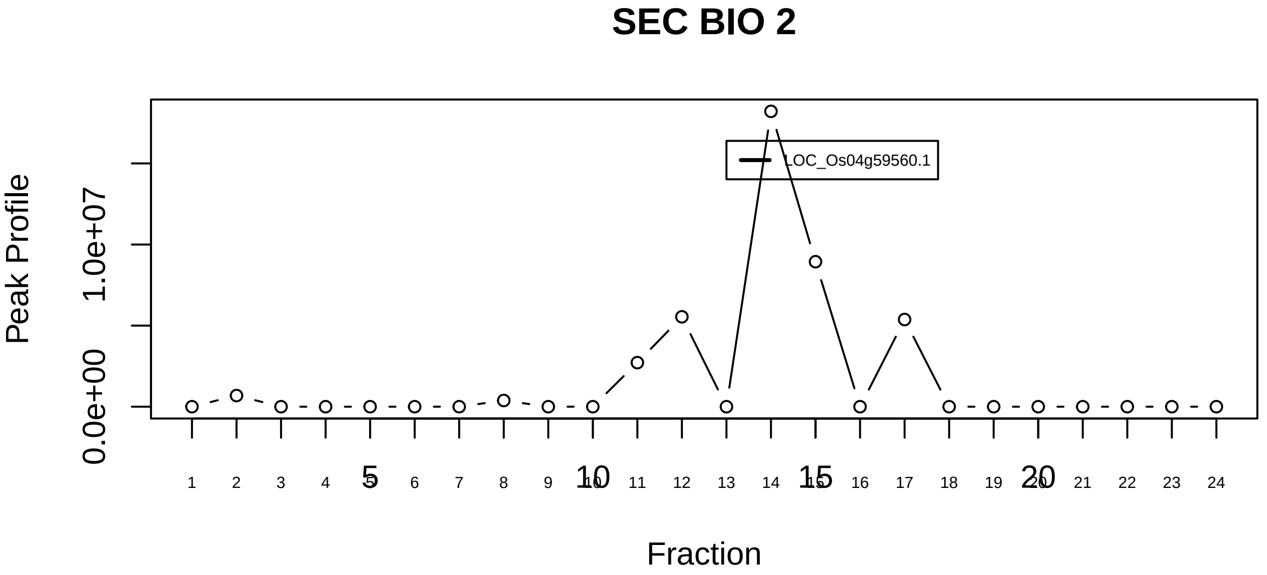

TBCE−TBCB−tubulin(alpha) complex|6714−−singleton_1

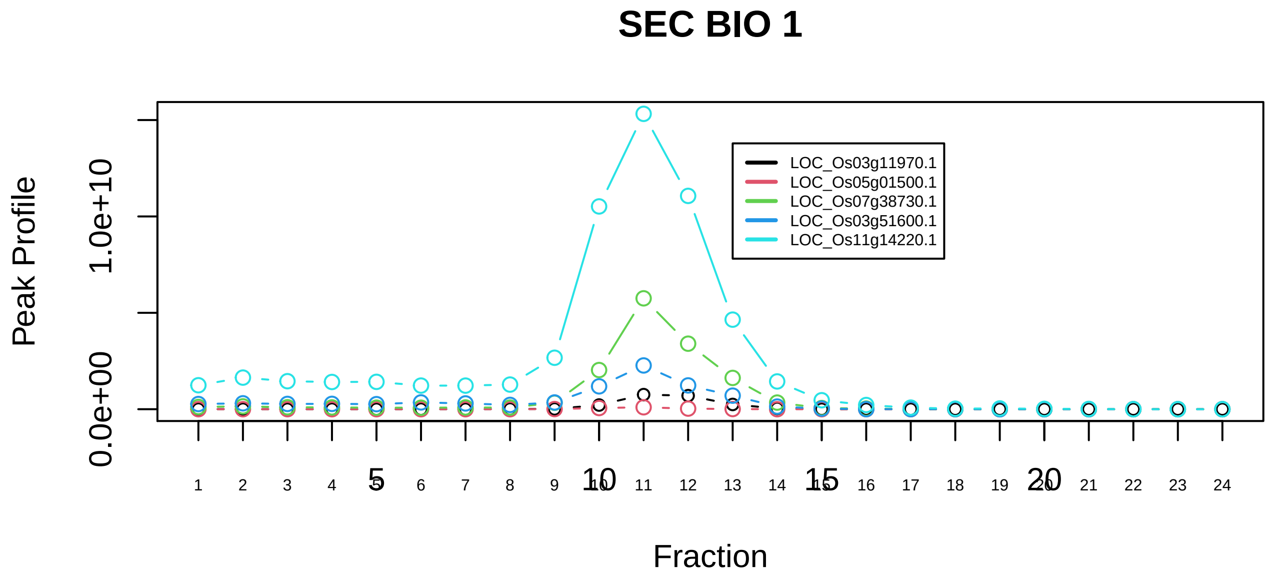

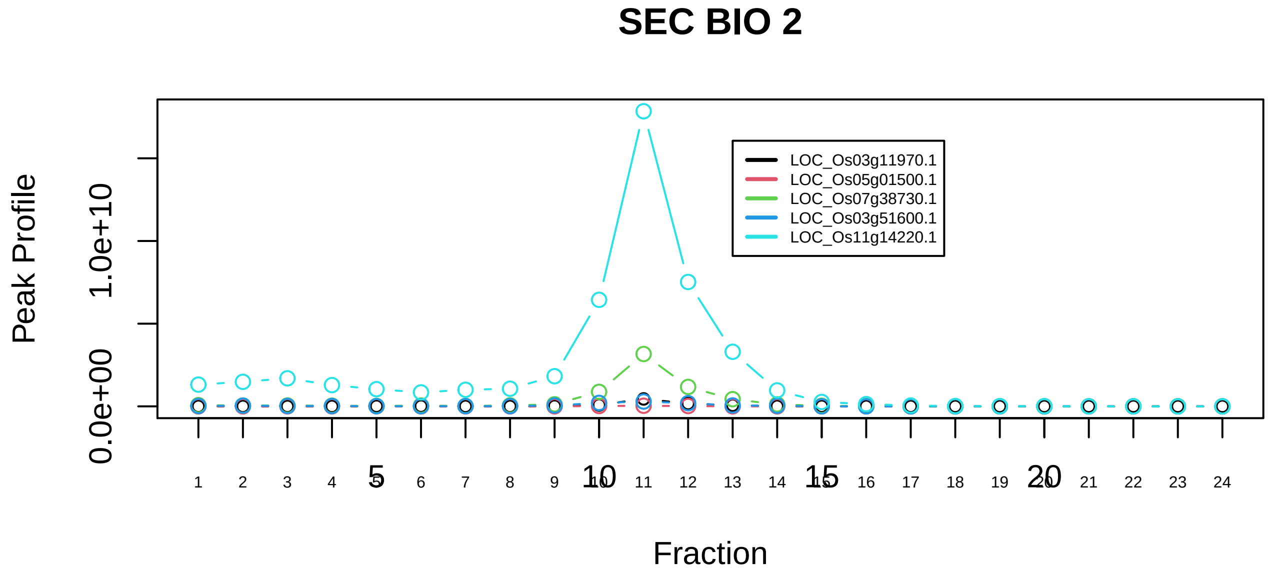

TBCE−TBCB−tubulin(alpha) complex|6714−−sub_1

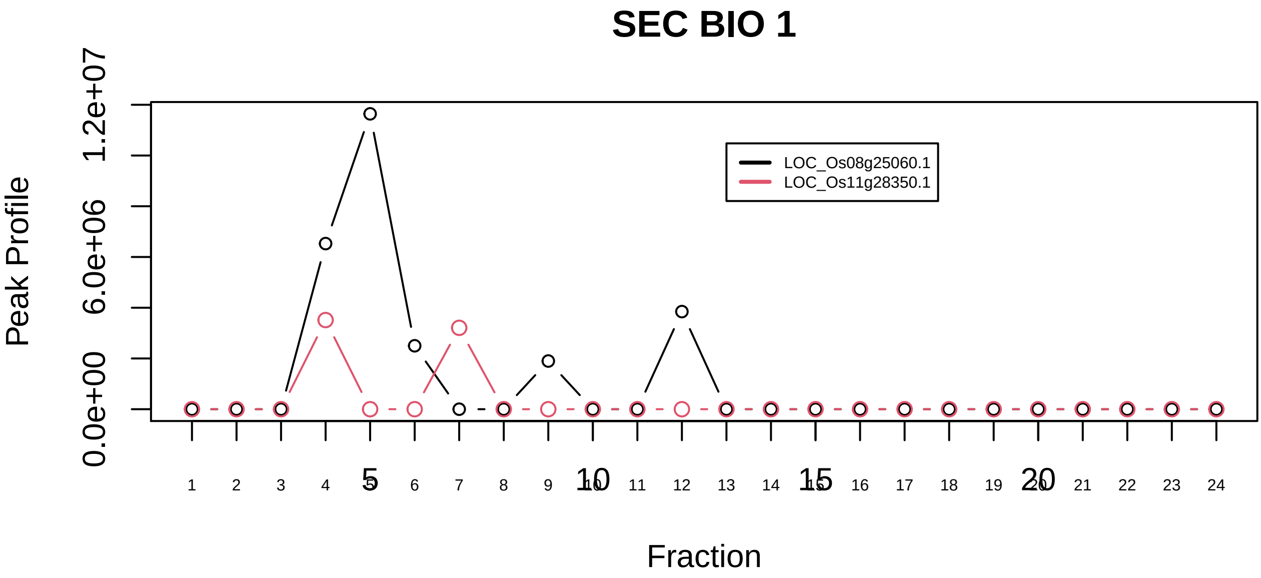

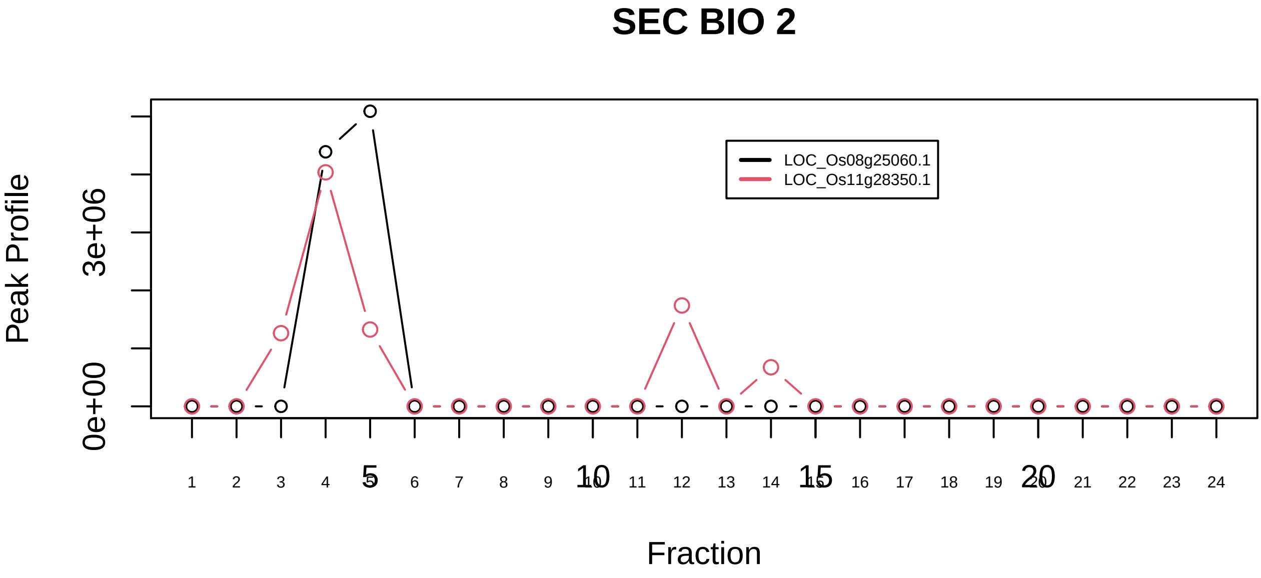

TFIIH transcription factor complex|107−−sub_1

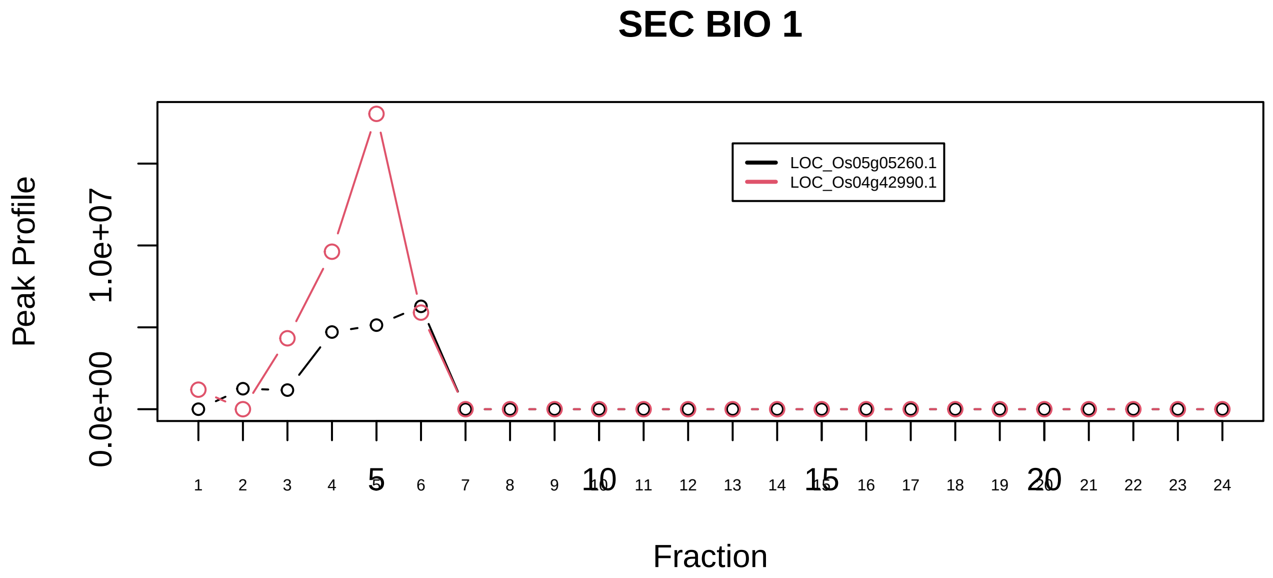

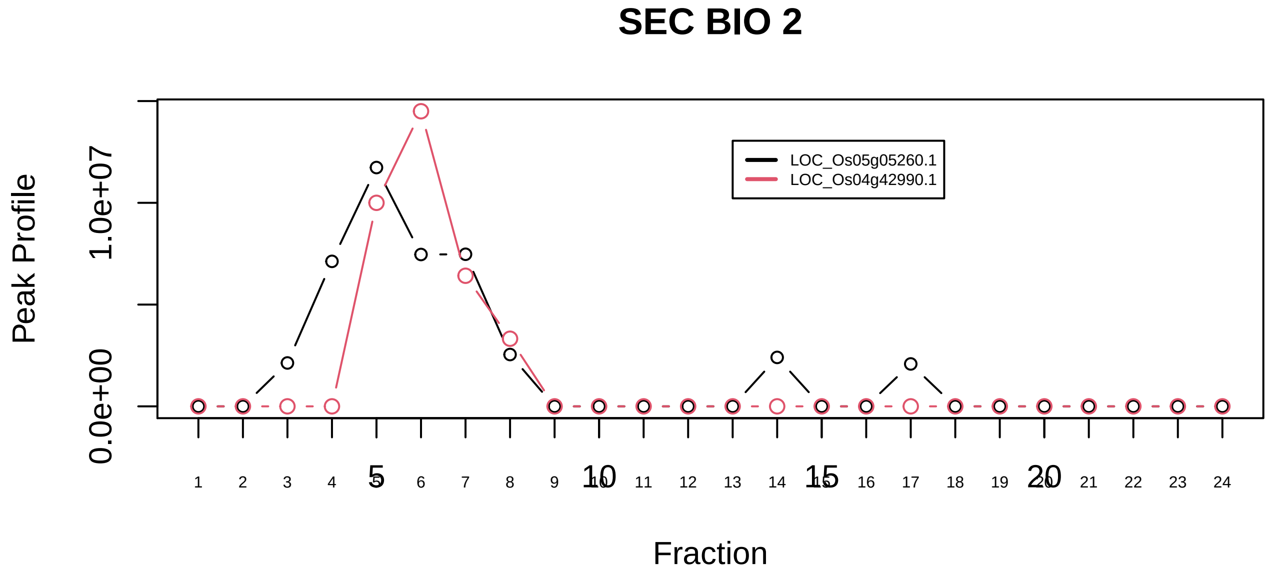

TFIIH transcription factor complex|107−−sub_2

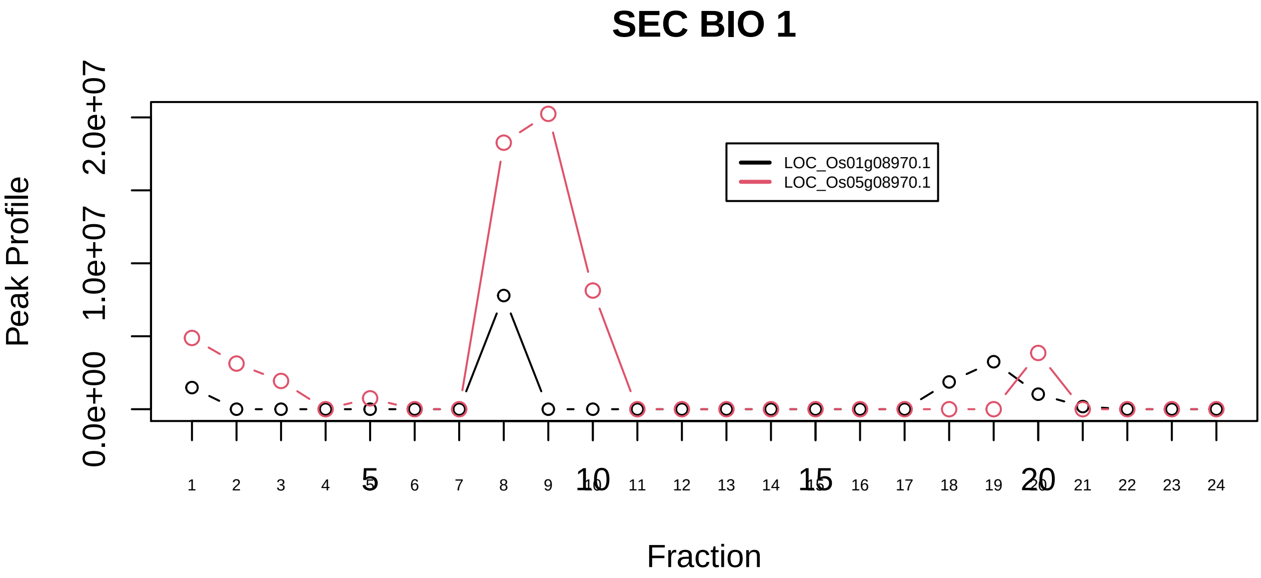

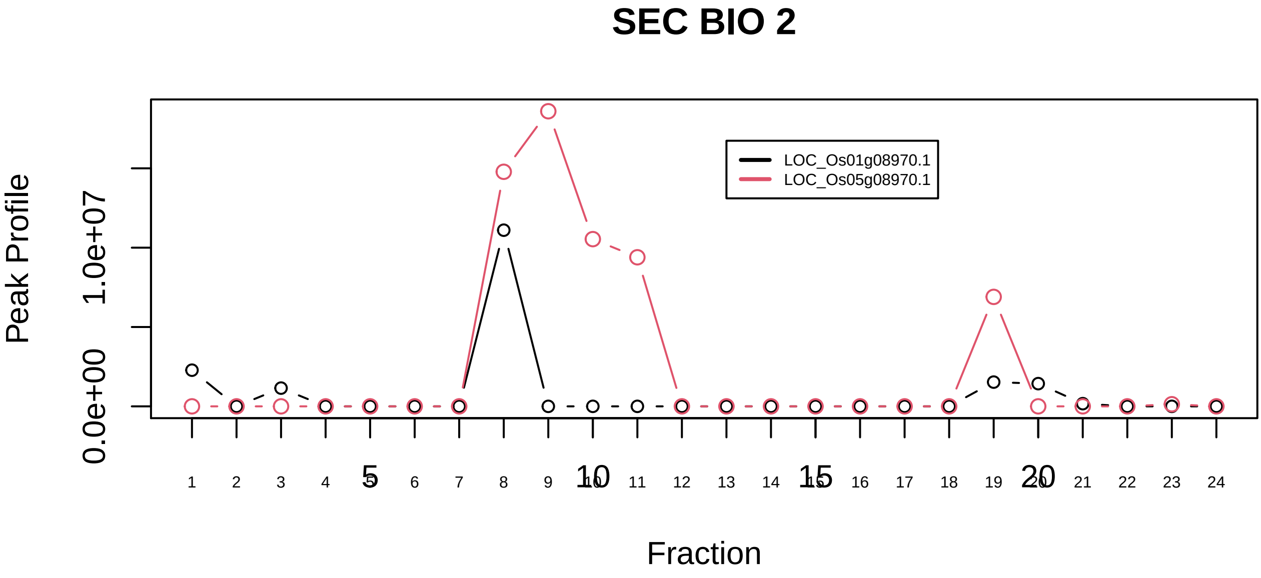

Toposome|924−−sub_orthoparalog_1

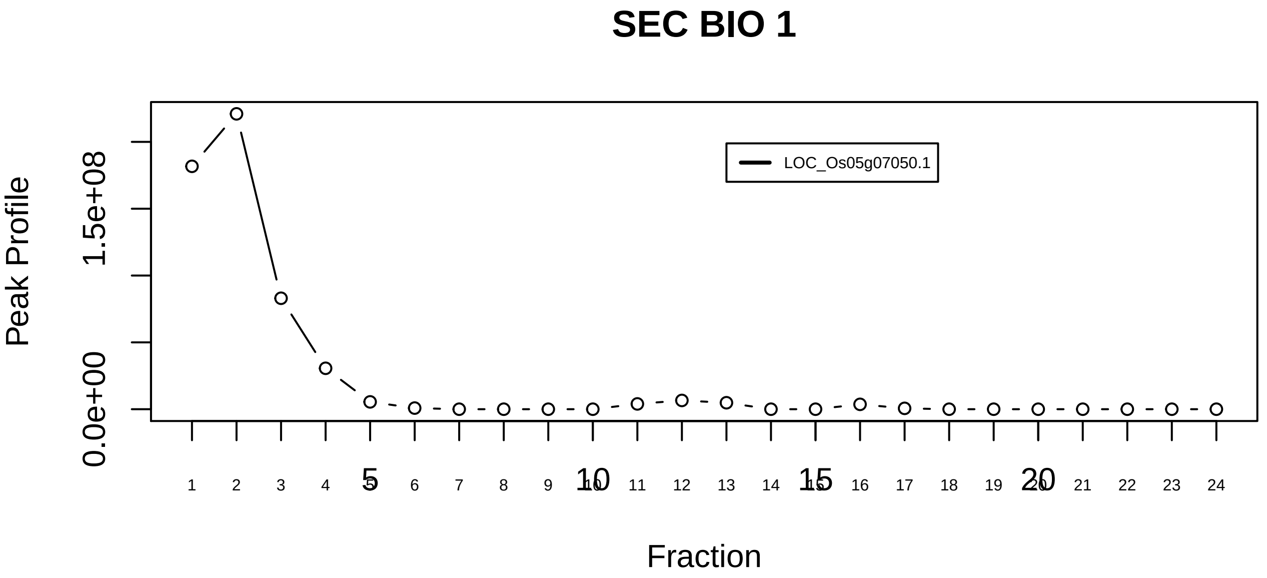

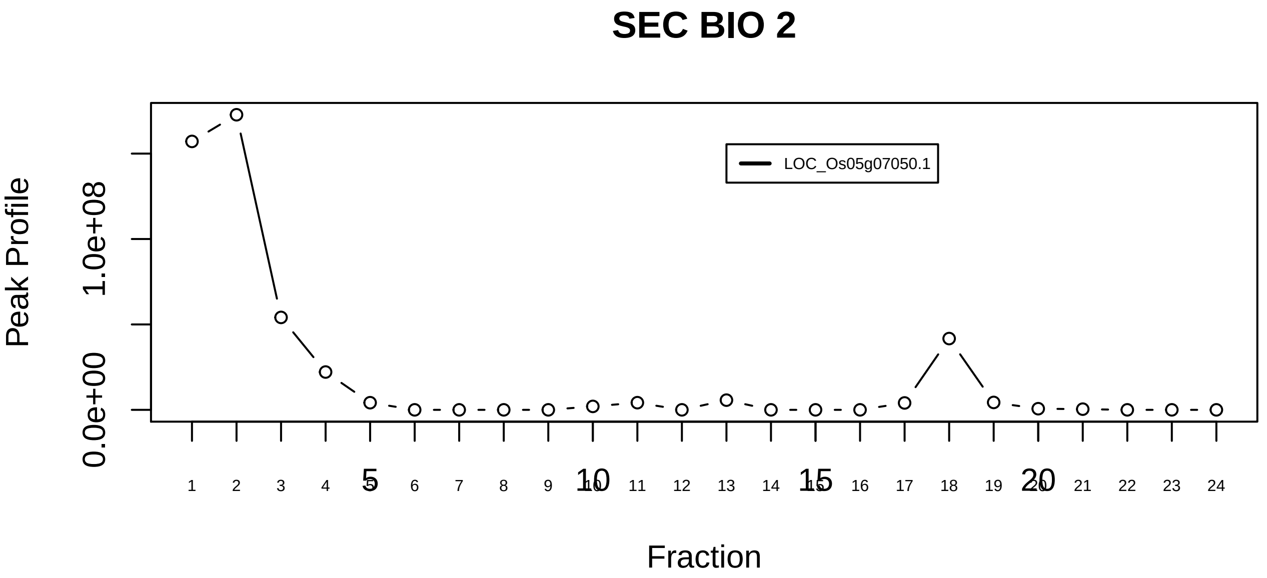

Toposome|924−−singleton_1

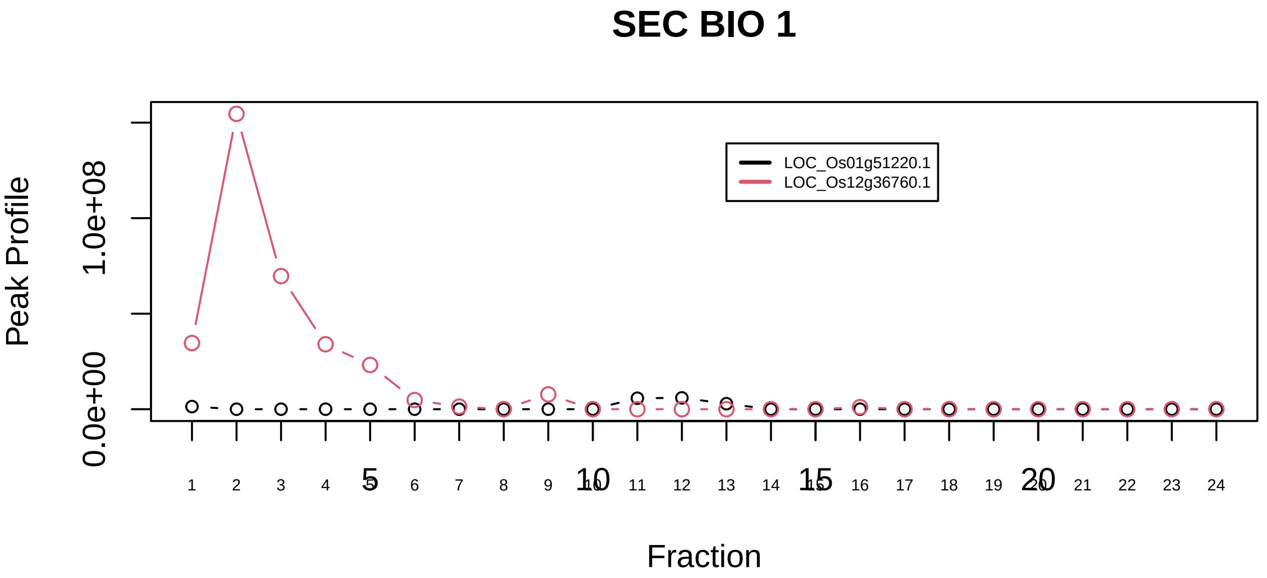

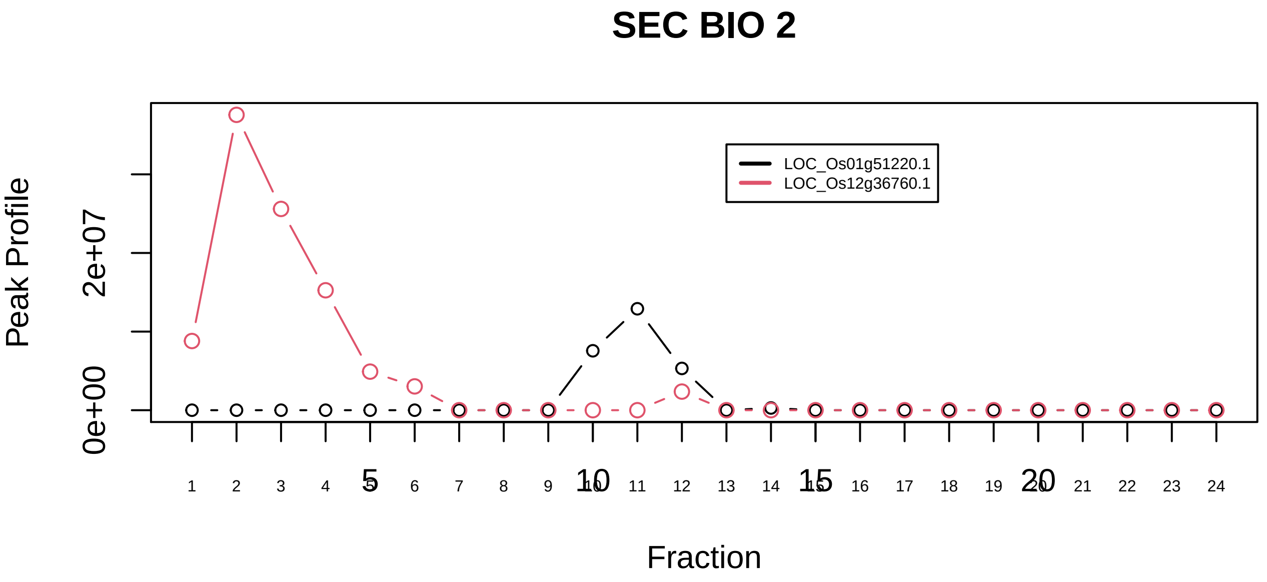

TRAPP complex|6468−−only_2_prots_before_clustering

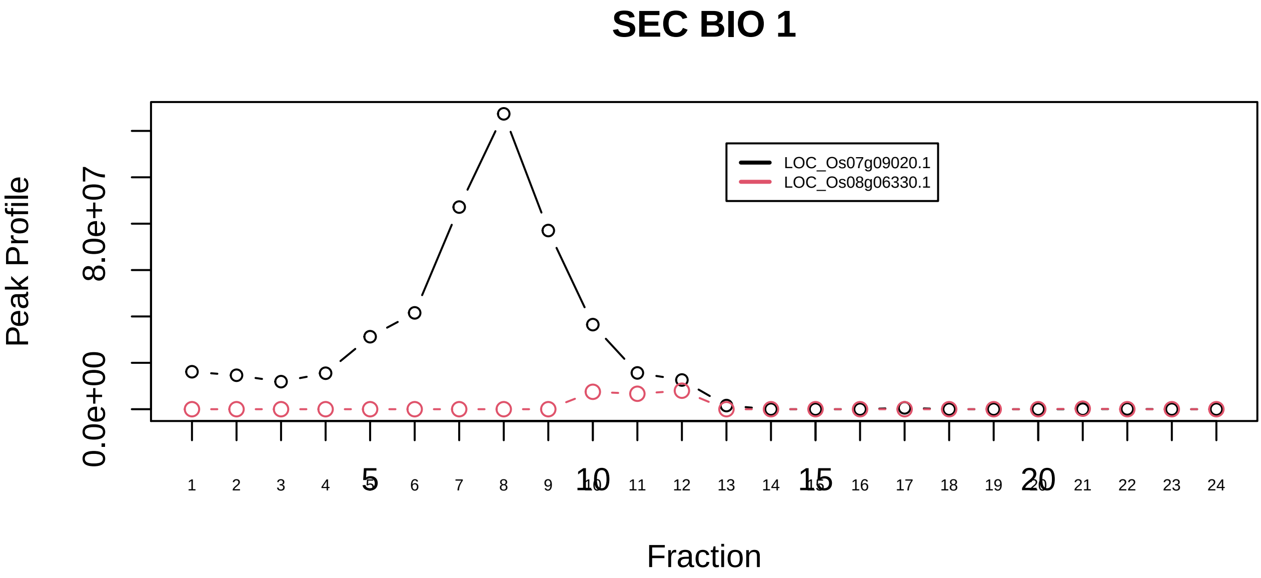

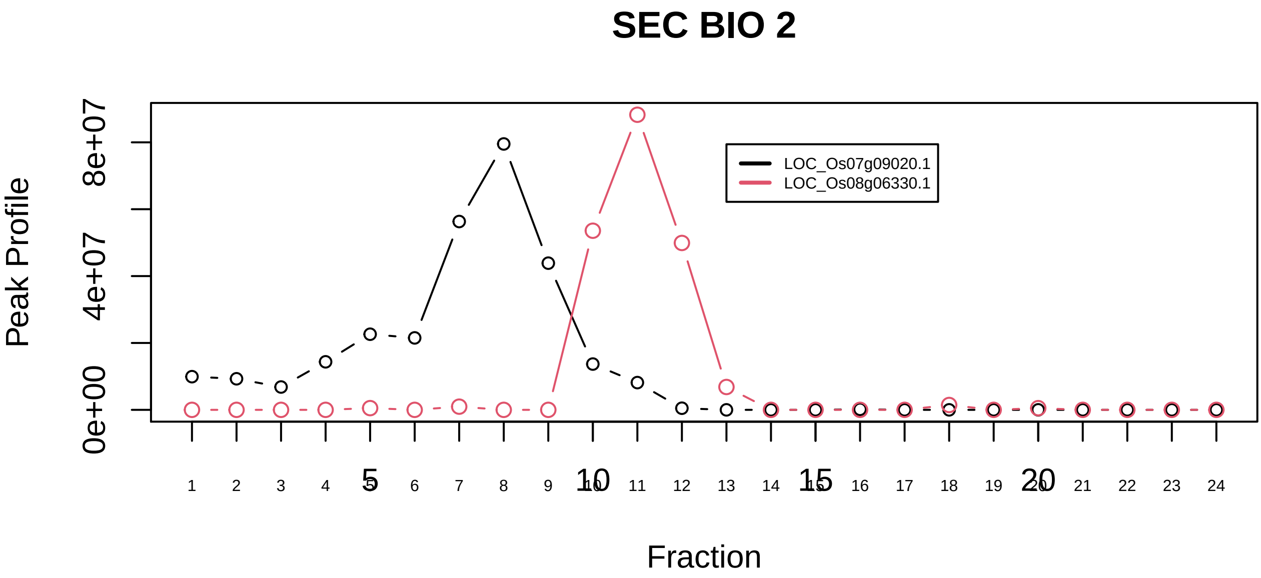

TTP−AGO4 complex|1386−−only_2_prots_before_clustering

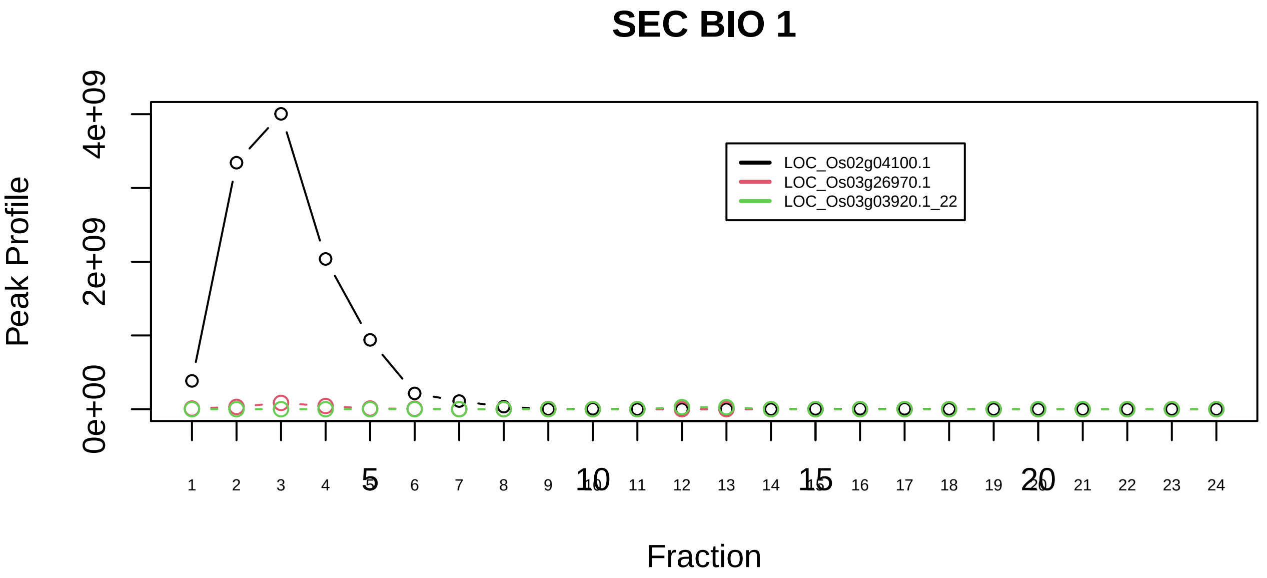

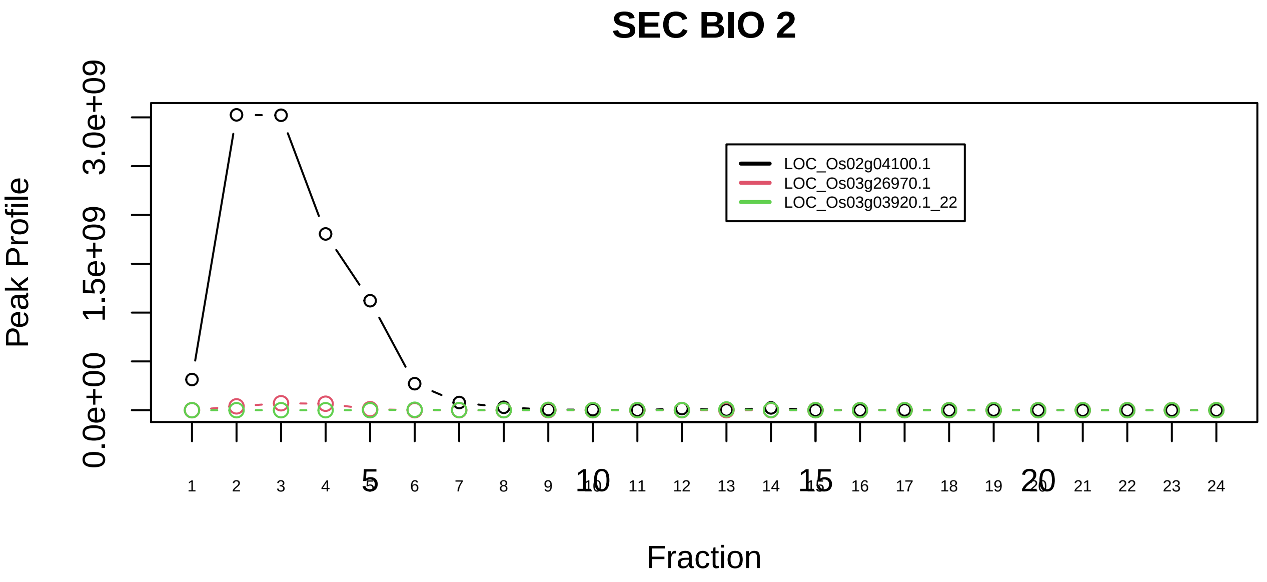

Ubiquilin−proteasome complex|5209−−sub_1

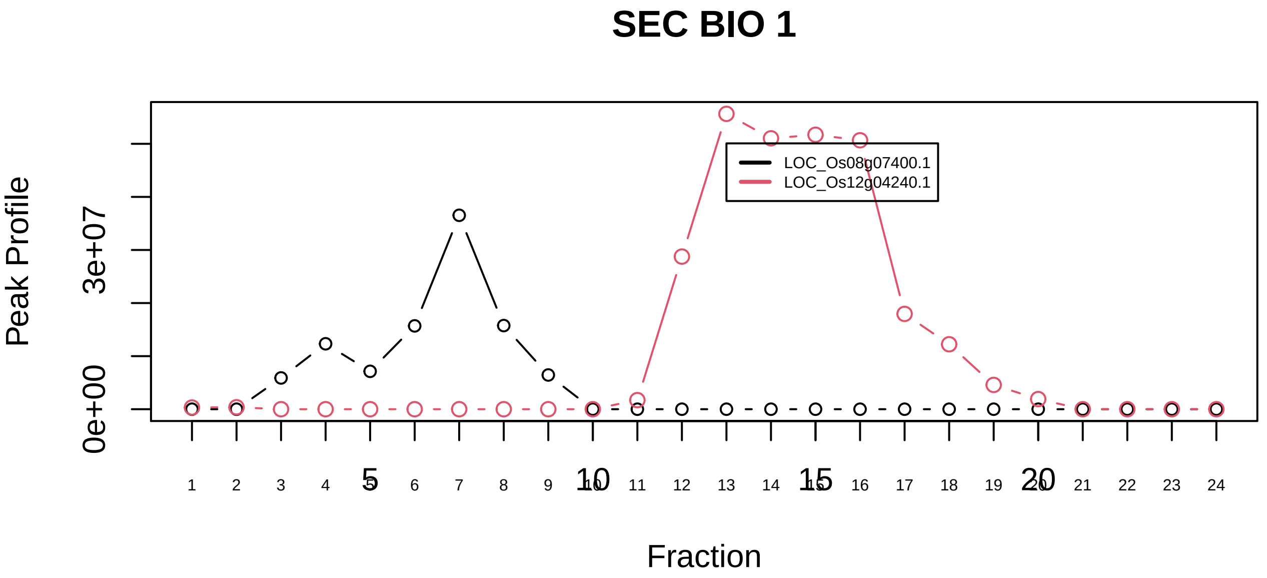

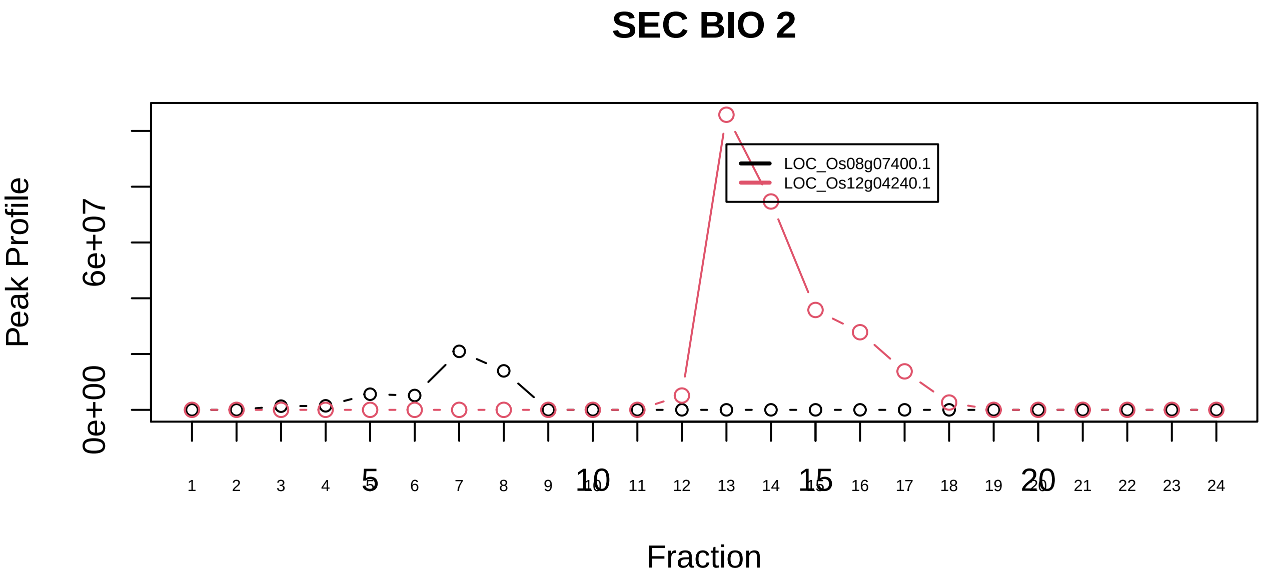

Ubiquitin E3 ligase (CUL3, KLHL12, PEF1, PDCD6)|7522−−only_2_prots_before_clustering

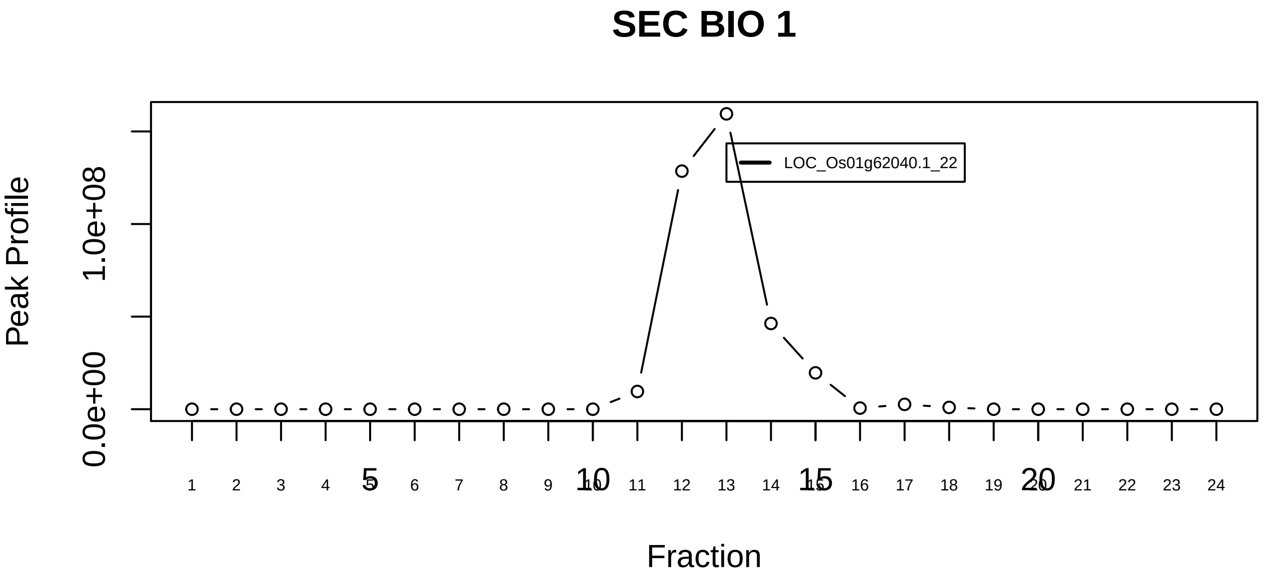

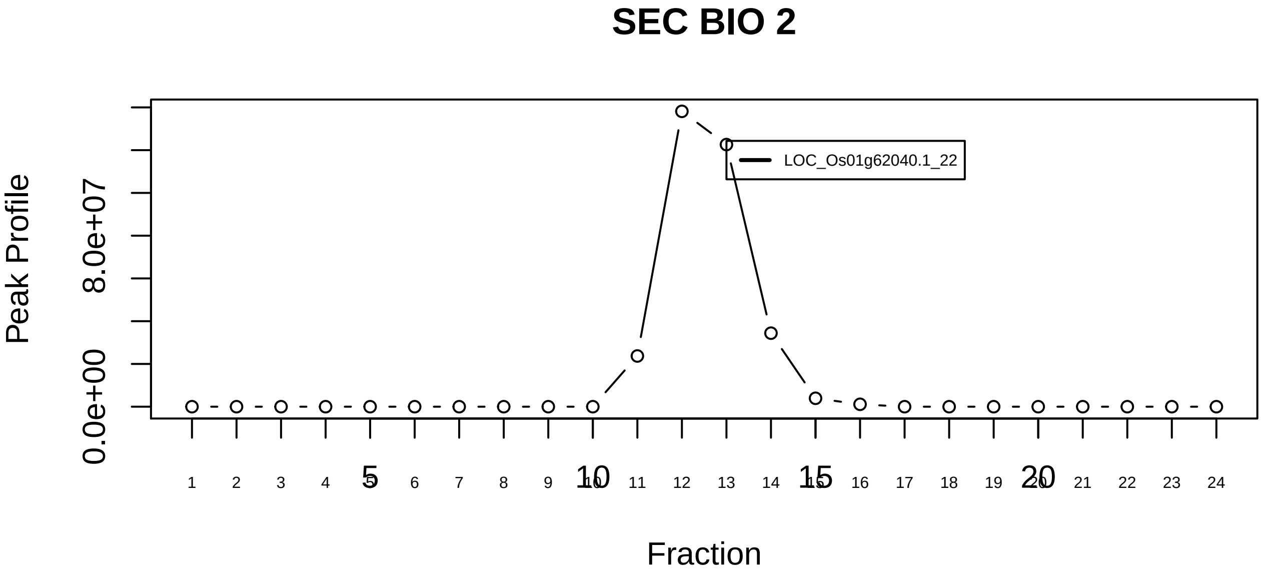

URI complex (Unconventional prefoldin RPB5 Interactor)|781−−singleton_1

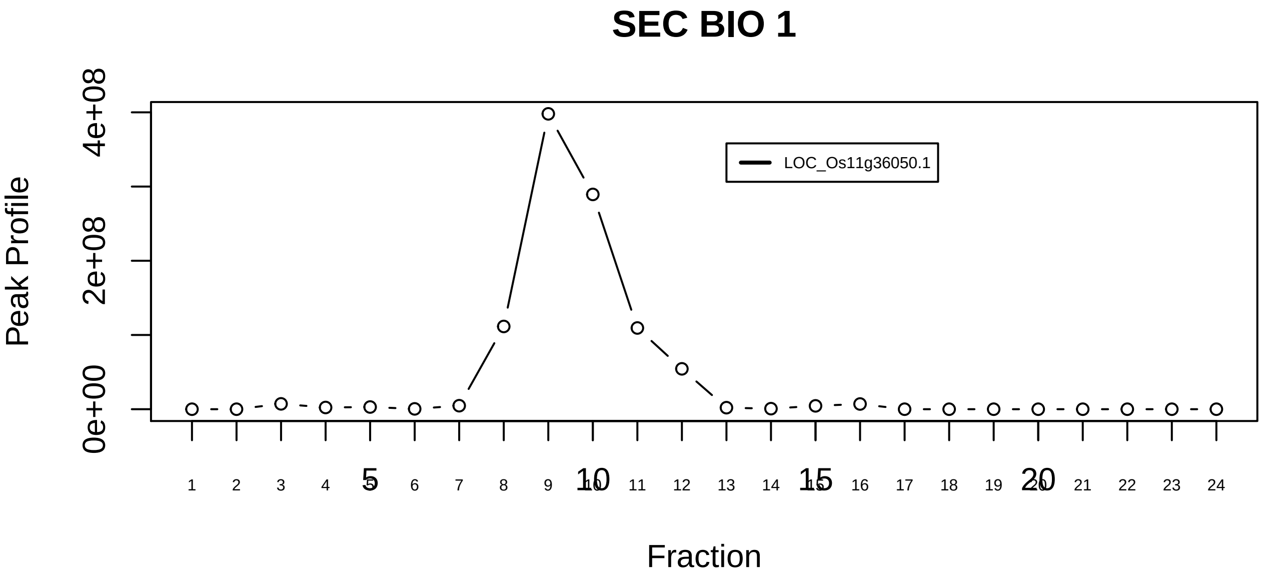

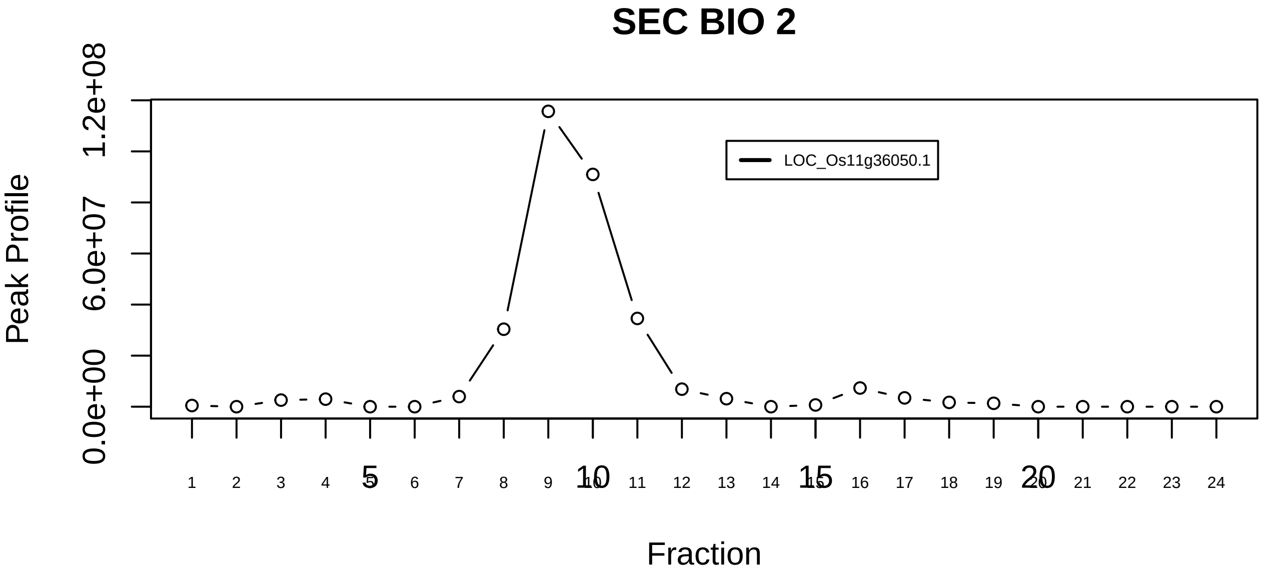

URI complex (Unconventional prefoldin RPB5 Interactor)|781−−singleton_2

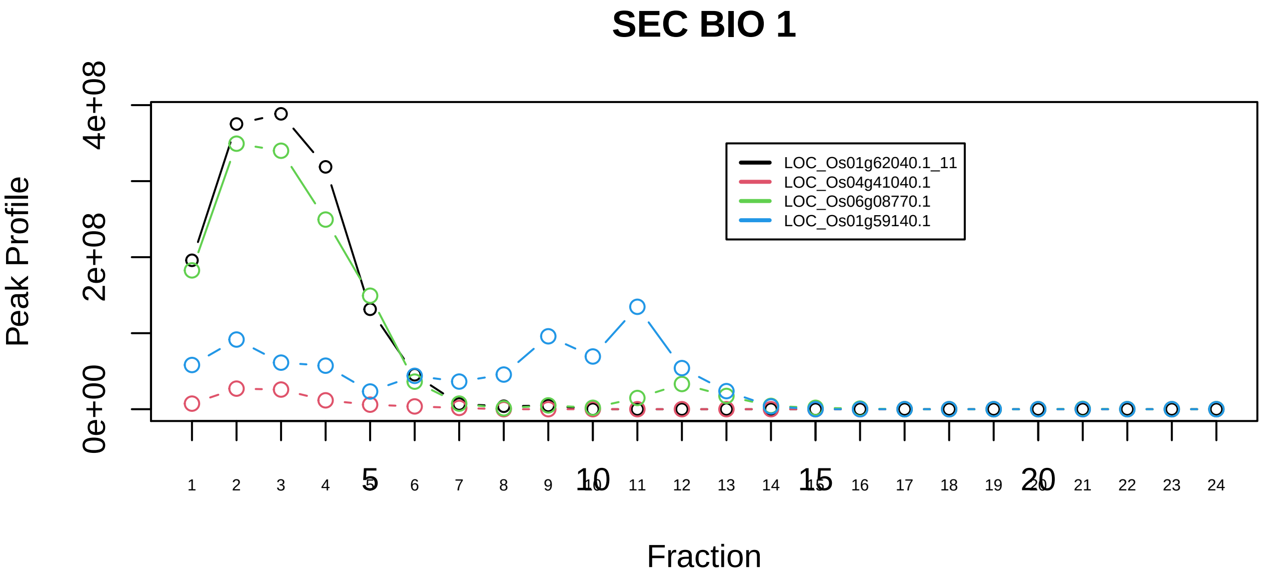

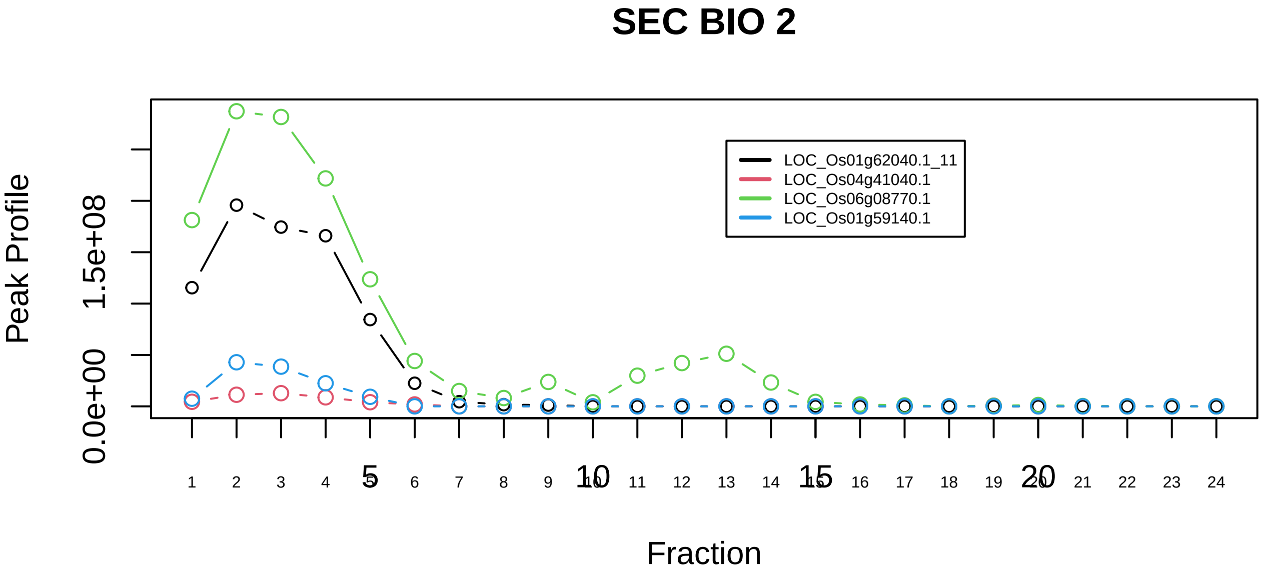

URI complex (Unconventional prefoldin RPB5 Interactor)|781−−sub_1
