## Supplemental Figure S4 Histograms of mean of pair-wised doverall from random simulations of rice orthocomplexes for "A machine learning-based approach to identify reliable gold standards for protein complex composition prediction"

**Supplemental Figure S4.** Histograms of mean of pair-wised $d_{overall}$ from a random simulations of rice orthocomplexes

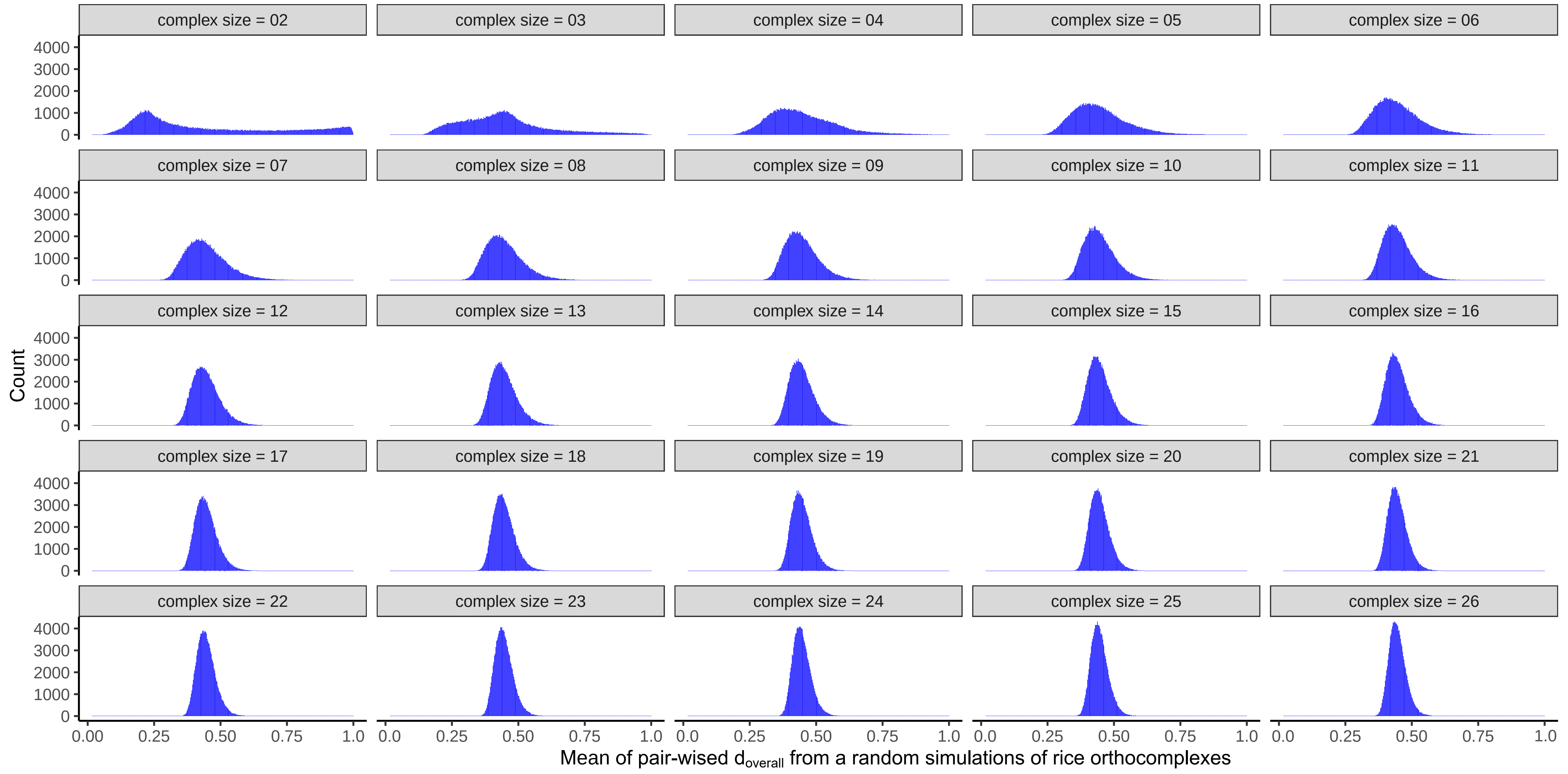

complex size = 02
complex size = 03
complex size = 04
complex size = 05
complex size = 06
complex size = 07
complex size = 08
complex size = 09
complex size = 10
complex size = 11
complex size = 12
complex size = 13
complex size = 14
complex size = 15
complex size = 16
complex size = 17
complex size = 18
complex size = 19
complex size = 20
complex size = 21
complex size = 22
complex size = 23
complex size = 24
complex size = 25
complex size = 26
Count
0
1000
2000
3000
4000
0.00
0.25
0.50
0.75
1.0
Mean of pair-wised $d_{overall}$ from a random simulations of rice orthocomplexes
