## Supplemental Figure S6 Arabidopsis subcomplex identification by small world analysis(Mapp vs Mcalc plots) for "A machine learning-based approach to identify reliable gold standards for protein complex composition prediction"

**Supplemental Figure S6.** Arabidopsis subcomplex identification by small world analysis ( $M_{app}$ vs $M_{calc}$ )

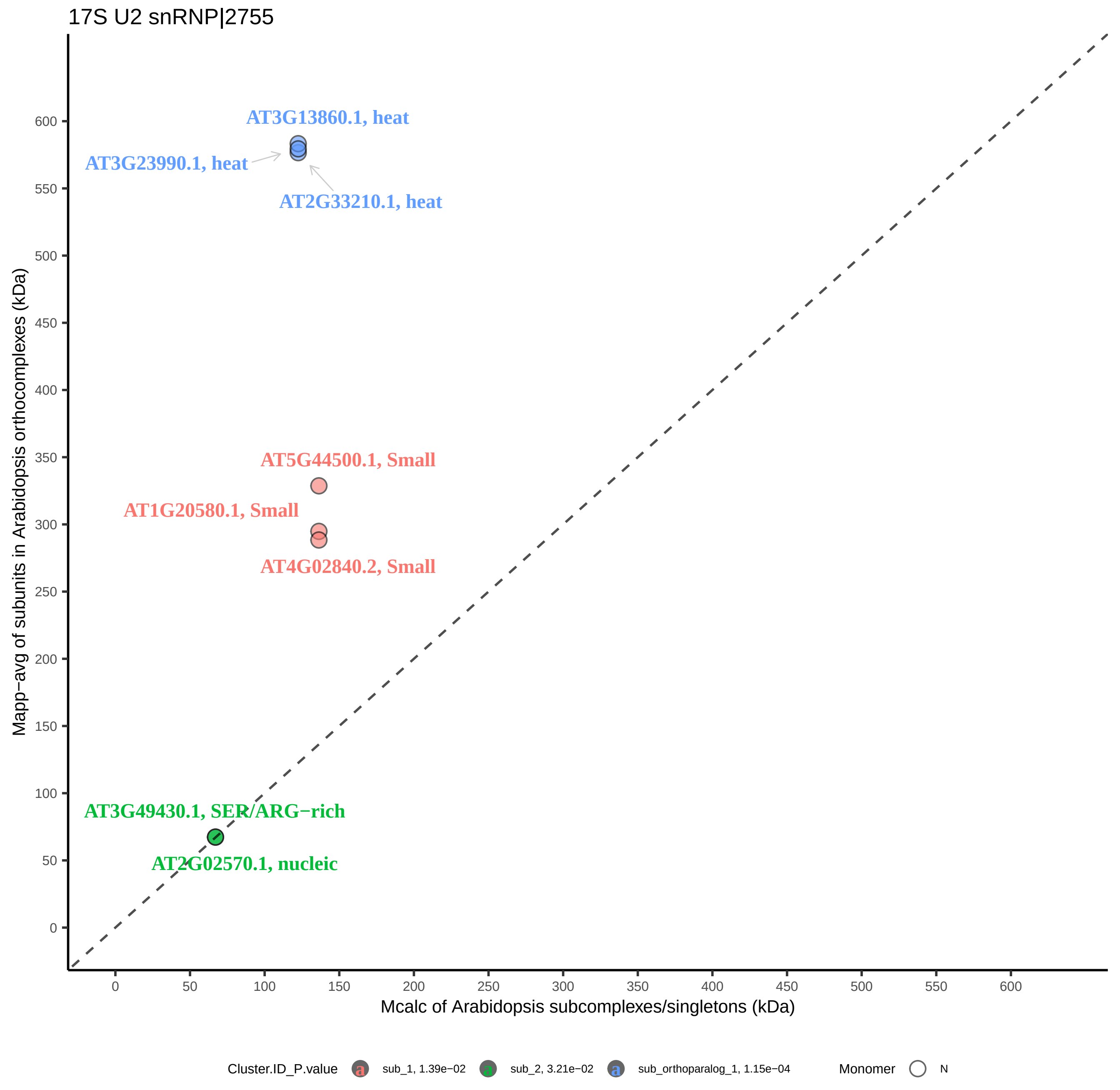

17S U2 snRNP|2755
AT3G13860.1, heat
AT3G23990.1, heat
AT2G33210.1, heat
AT5G44500.1, Small
AT1G20580.1, Small
AT4G02840.2, Small
AT3G49430.1, SER/ARG−rich
AT2G02570.1, nucleic
Mapp−avg of subunits in Arabidopsis orthocomplexes (kDa)
Mcalc of Arabidopsis subcomplexes/singletons (kDa)
0
50
100
150
200
250
300
350
400
450
500
550
600
Cluster.ID_P.value
sub_1, 1.39e−02
sub_2, 3.21e−02
sub_orthoparalog_1, 1.15e−04
Monomer
N

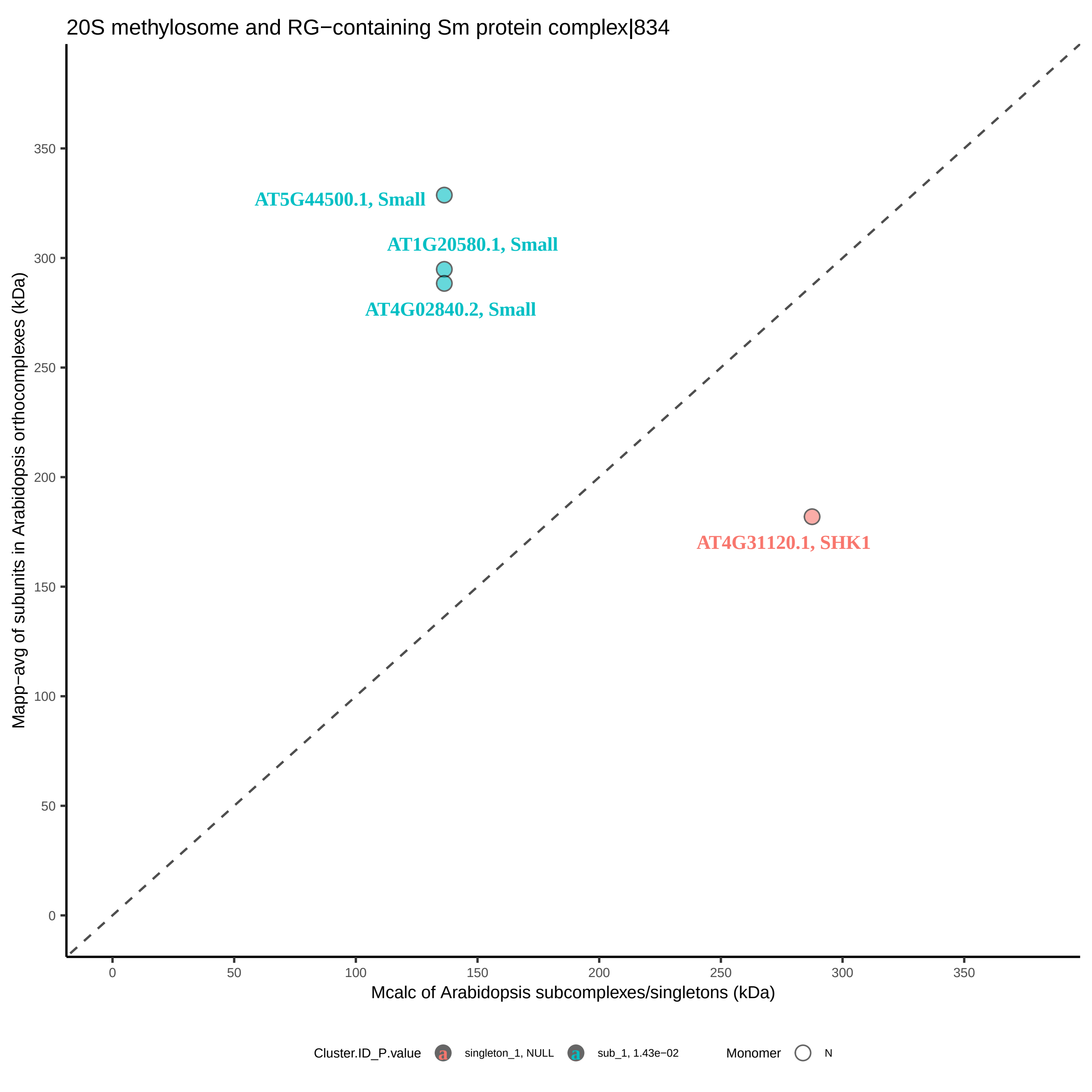
20S methylosome and RG−containing Sm protein complex|834
Mapp−avg of subunits in Arabidopsis orthocomplexes (kDa)
Mcalc of Arabidopsis subcomplexes/singletons (kDa)
AT5G44500.1, Small
AT1G20580.1, Small
AT4G02840.2, Small
AT4G31120.1, SHK1
0
50
100
150
200
250
300
350
Cluster.ID_P.value
singleton_1, NULL
sub_1, 1.43e−02
Monomer
N

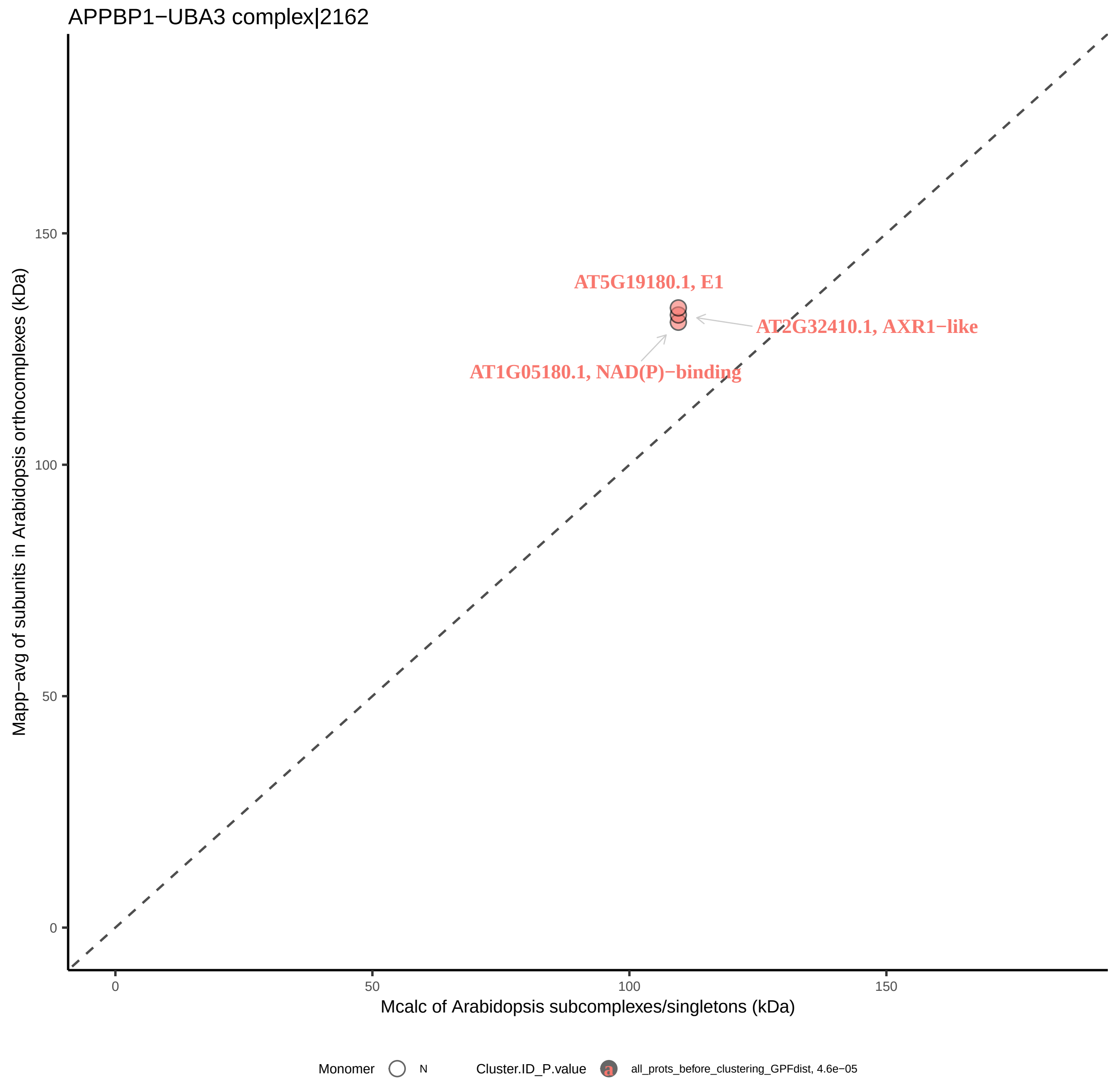
APPBP1−UBA3 complex|2162
Mapp−avg of subunits in Arabidopsis orthocomplexes (kDa)
Mcalc of Arabidopsis subcomplexes/singletons (kDa)
0
50
100
150
AT5G19180.1, E1
AT2G32410.1, AXR1−like
AT1G05180.1, NAD(P)−binding
Monomer
N
Cluster.ID_P.value
a
all_prots_before_clustering_GPFdist, 4.6e−05

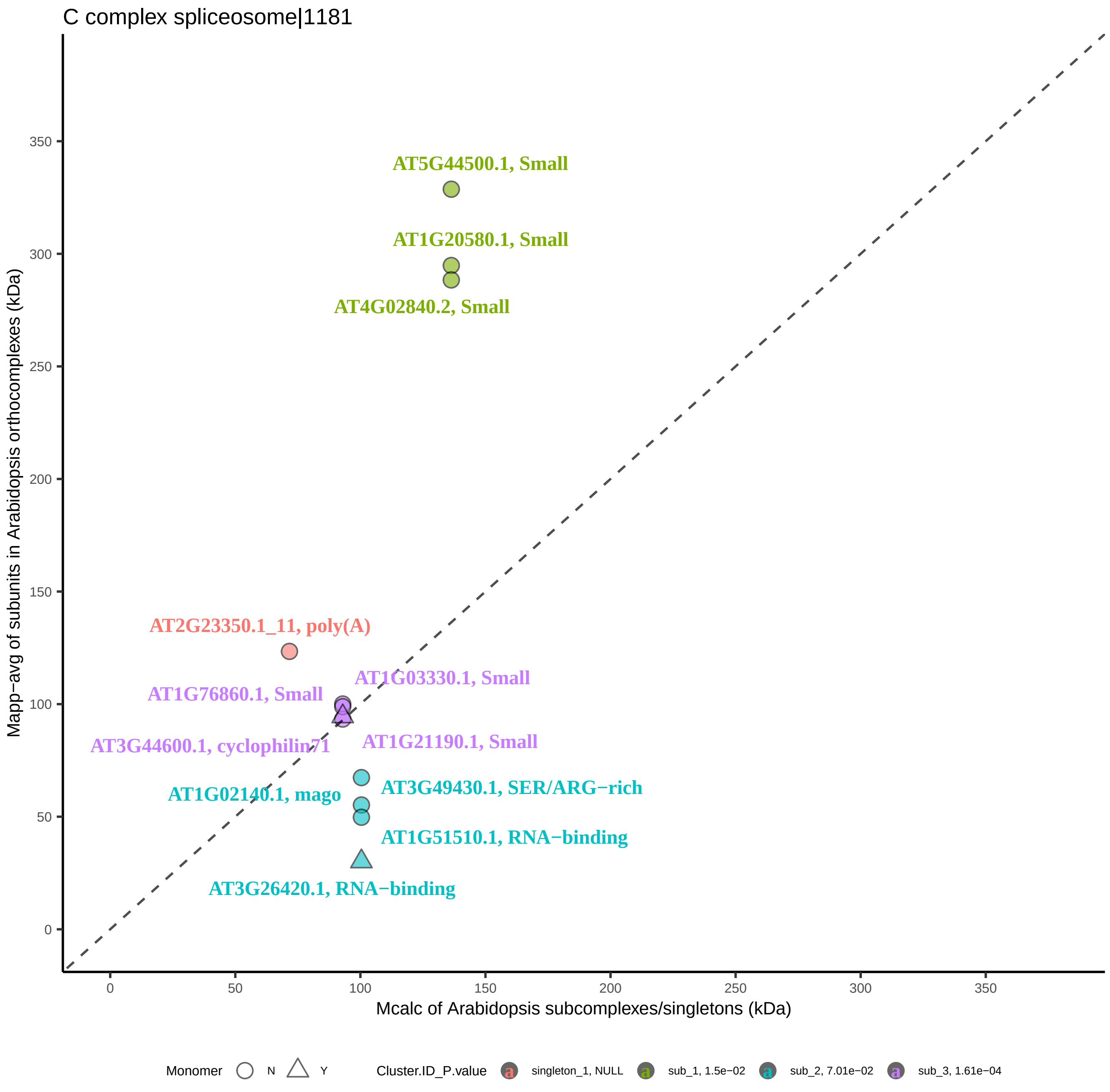
C complex spliceosome|1181
AT5G44500.1, Small
AT1G20580.1, Small
AT4G02840.2, Small
AT2G23350.1_11, poly(A)
AT1G03330.1, Small
AT1G76860.1, Small
AT3G44600.1, cyclophilin71
AT1G21190.1, Small
AT3G49430.1, SER/ARG−rich
AT1G02140.1, mago
AT1G51510.1, RNA−binding
AT3G26420.1, RNA−binding
Mapp−avg of subunits in Arabidopsis orthocomplexes (kDa)
Mcalc of Arabidopsis subcomplexes/singletons (kDa)
0
50
100
150
200
250
300
350
Monomer
N
Y
Cluster.ID_P.value
singleton_1, NULL
sub_1, 1.5e−02
sub_2, 7.01e−02
sub_3, 1.61e−04

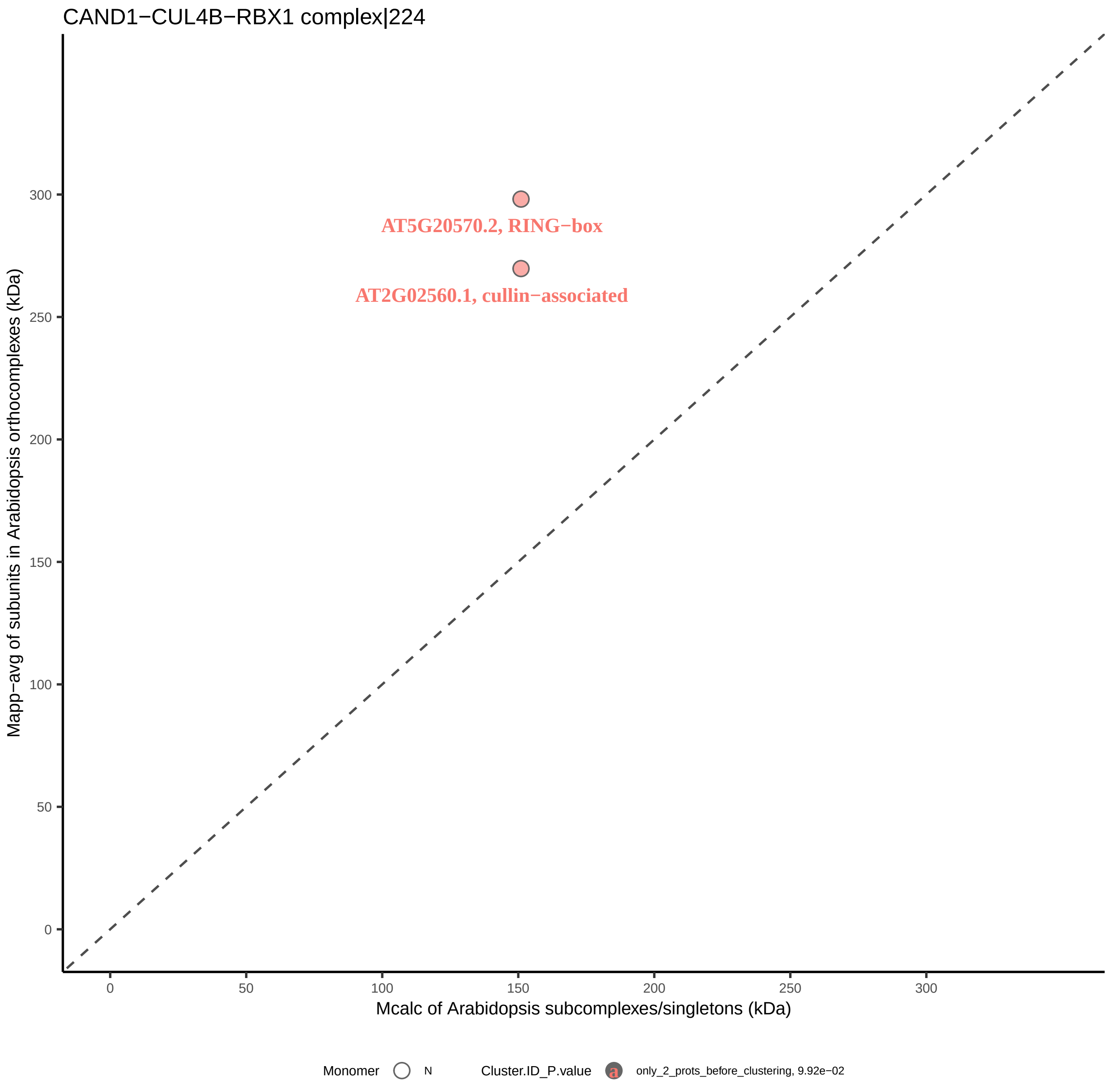

CAND1−CUL4B−RBX1 complex|224
Mapp−avg of subunits in Arabidopsis orthocomplexes (kDa)
300
250
200
150
100
50
0
AT5G20570.2, RING−box
AT2G02560.1, cullin−associated
0
50
100
150
200
250
300
Mcalc of Arabidopsis subcomplexes/singletons (kDa)
Monomer
N
Cluster.ID_P.value
a
only_2_prots_before_clustering, 9.92e−02

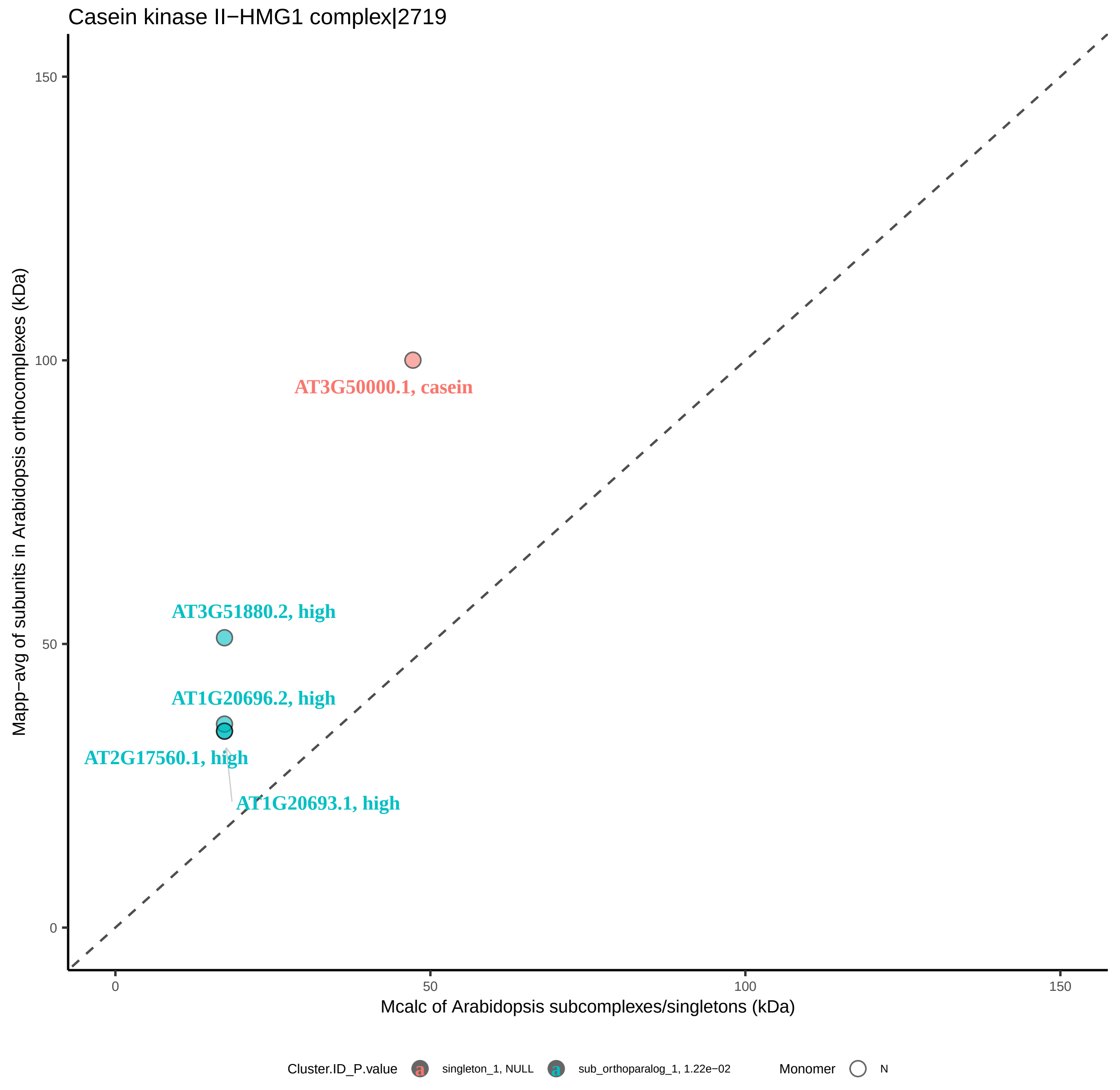

Casein kinase II−HMG1 complex|2719
Mapp−avg of subunits in Arabidopsis orthocomplexes (kDa)
Mcalc of Arabidopsis subcomplexes/singletons (kDa)
0
50
100
150
AT3G50000.1, casein
AT3G51880.2, high
AT1G20696.2, high
AT2G17560.1, high
AT1G20693.1, high
Cluster.ID_P.value
singleton_1, NULL
sub_orthoparalog_1, 1.22e−02
Monomer
N

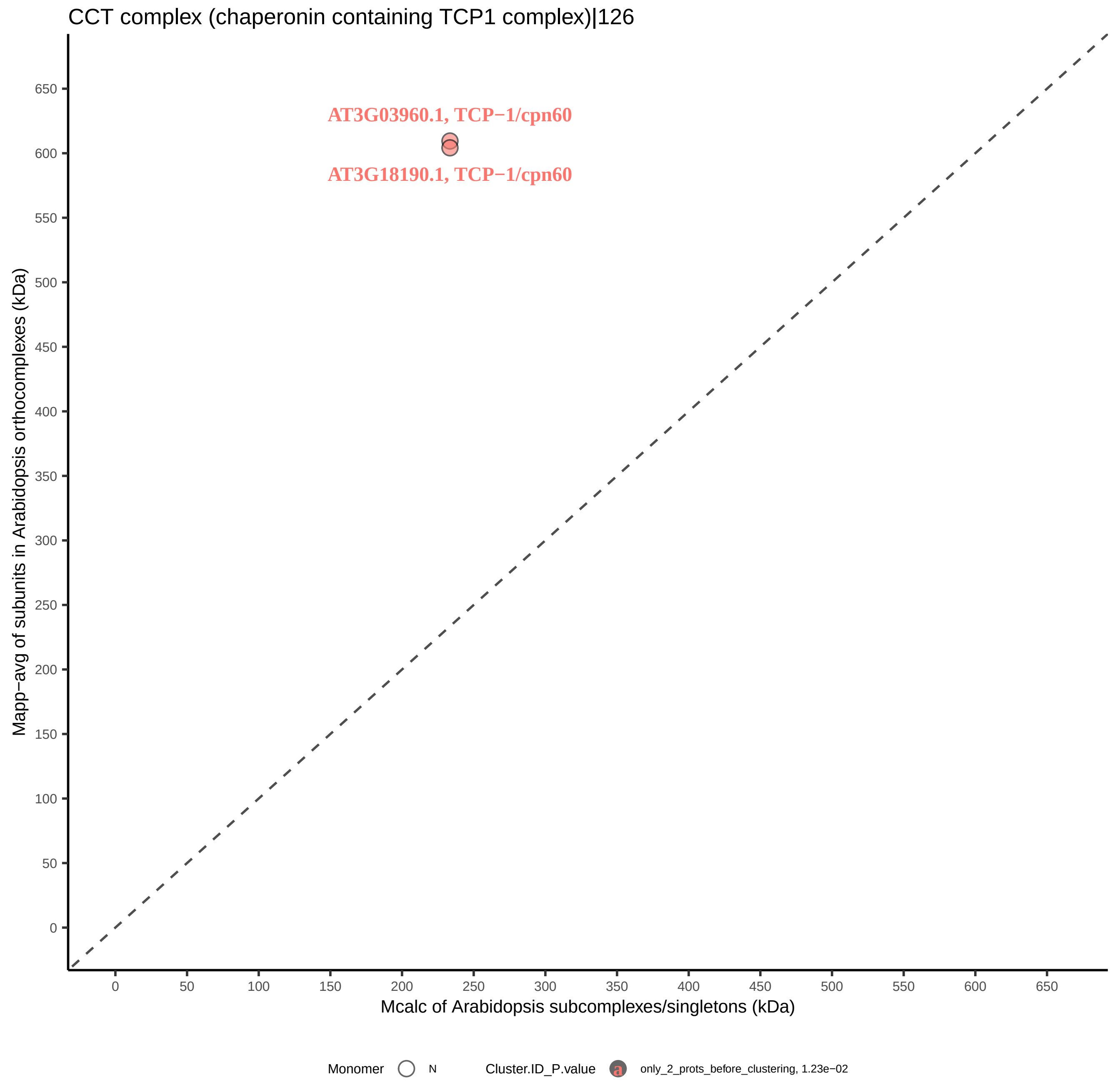
CCT complex (chaperonin containing TCP1 complex)|126
AT3G03960.1, TCP−1/cpn60
AT3G18190.1, TCP−1/cpn60
Mapp−avg of subunits in Arabidopsis orthocomplexes (kDa)
Mcalc of Arabidopsis subcomplexes/singletons (kDa)
0
50
100
150
200
250
300
350
400
450
500
550
600
650
Monomer
N
Cluster.ID_P.value
a
only_2_prots_before_clustering, 1.23e−02

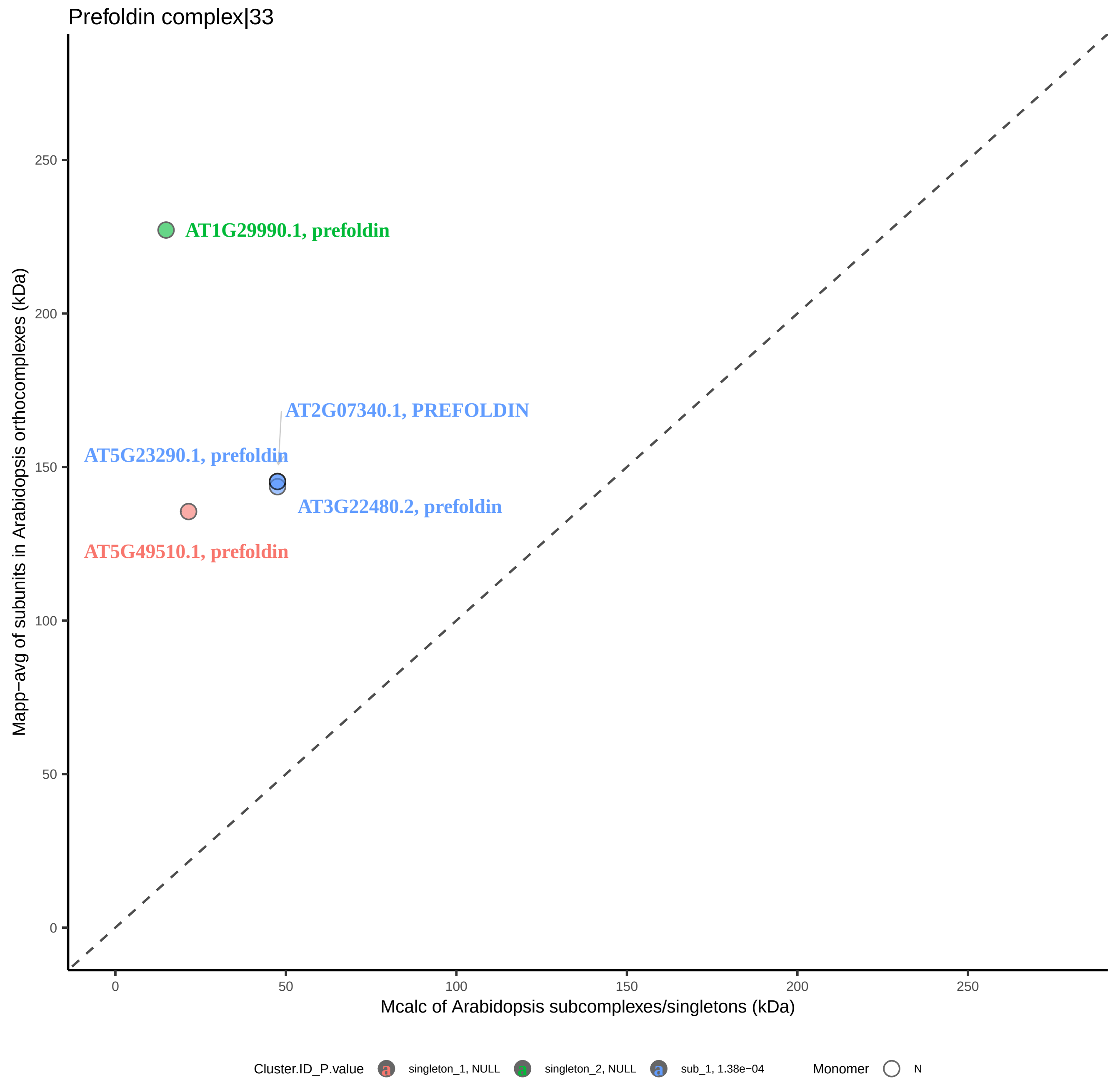

Prefoldin complex|33
AT1G29990.1, prefoldin
AT2G07340.1, PREFOLDIN
AT5G23290.1, prefoldin
AT3G22480.2, prefoldin
AT5G49510.1, prefoldin
Mapp−avg of subunits in Arabidopsis orthocomplexes (kDa)
Mcalc of Arabidopsis subcomplexes/singletons (kDa)
0
50
100
150
200
250
Cluster.ID_P.value
singleton_1, NULL
singleton_2, NULL
sub_1, 1.38e−04
Monomer
N

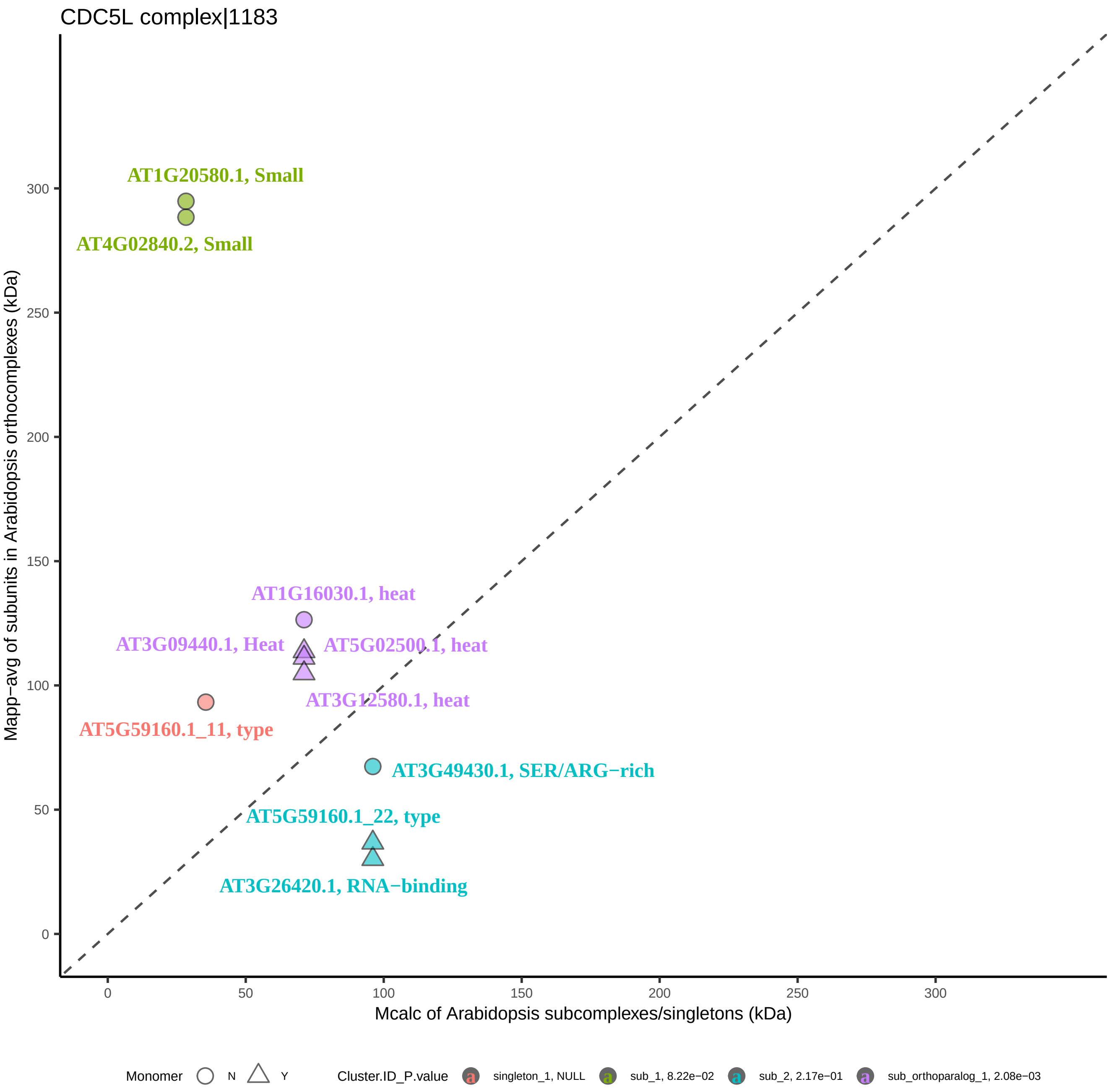

CDC5L complex|1183
AT1G20580.1, Small
AT4G02840.2, Small
AT1G16030.1, heat
AT3G09440.1, Heat
AT5G02500.1, heat
AT3G12580.1, heat
AT5G59160.1_11, type
AT3G49430.1, SER/ARG−rich
AT5G59160.1_22, type
AT3G26420.1, RNA−binding
Mapp−avg of subunits in Arabidopsis orthocomplexes (kDa)
Mcalc of Arabidopsis subcomplexes/singletons (kDa)
0
50
100
150
200
250
300
Monomer N Y
Cluster.ID_P.value
singleton_1, NULL
sub_1, 8.22e−02
sub_2, 2.17e−01
sub_orthoparalog_1, 2.08e−03

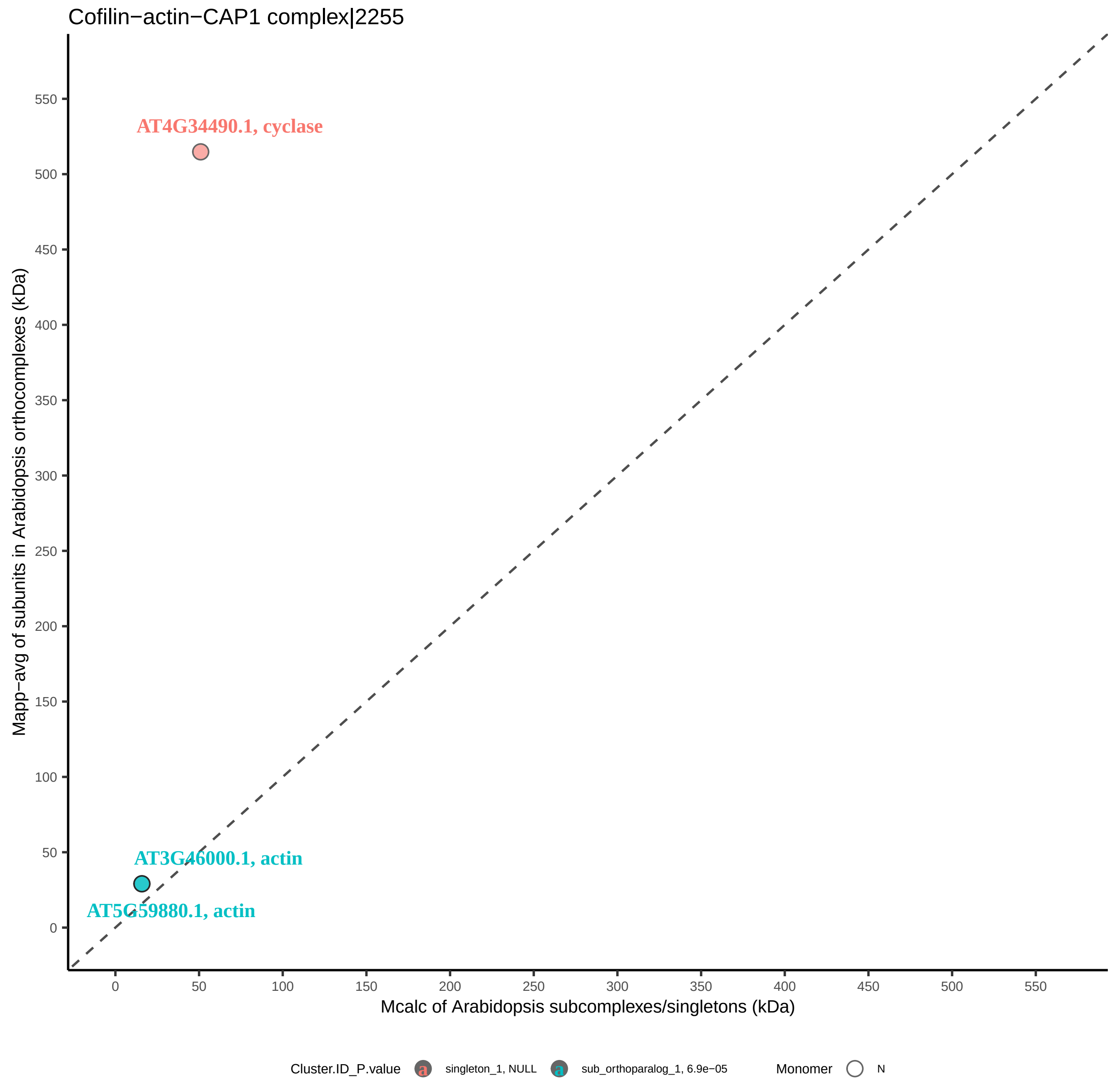

Cofilin−actin−CAP1 complex|2255
AT4G34490.1, cyclase
AT3G46000.1, actin
AT5G59880.1, actin
Mapp−avg of subunits in Arabidopsis orthocomplexes (kDa)
Mcalc of Arabidopsis subcomplexes/singletons (kDa)
0
50
100
150
200
250
300
350
400
450
500
550
Cluster.ID_P.value
singleton_1, NULL
sub_orthoparalog_1, 6.9e−05
Monomer
N

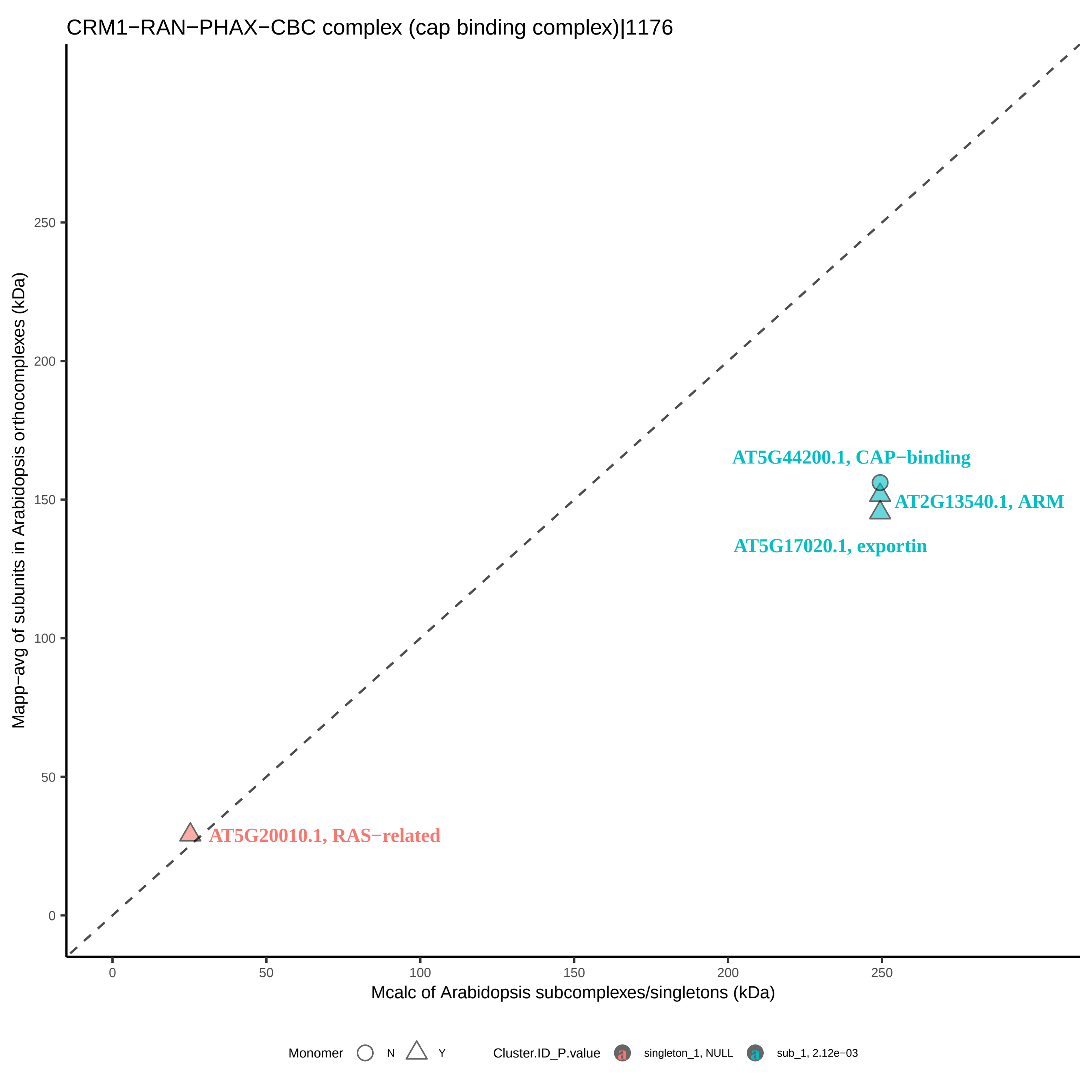

CRM1−RAN−PHAX−CBC complex (cap binding complex)|1176
Mapp−avg of subunits in Arabidopsis orthocomplexes (kDa)
Mcalc of Arabidopsis subcomplexes/singletons (kDa)
0
50
100
150
200
250
AT5G44200.1, CAP−binding
AT2G13540.1, ARM
AT5G17020.1, exportin
AT5G20010.1, RAS−related
Monomer
N
Y
Cluster.ID_P.value
singleton_1, NULL
sub_1, 2.12e−03

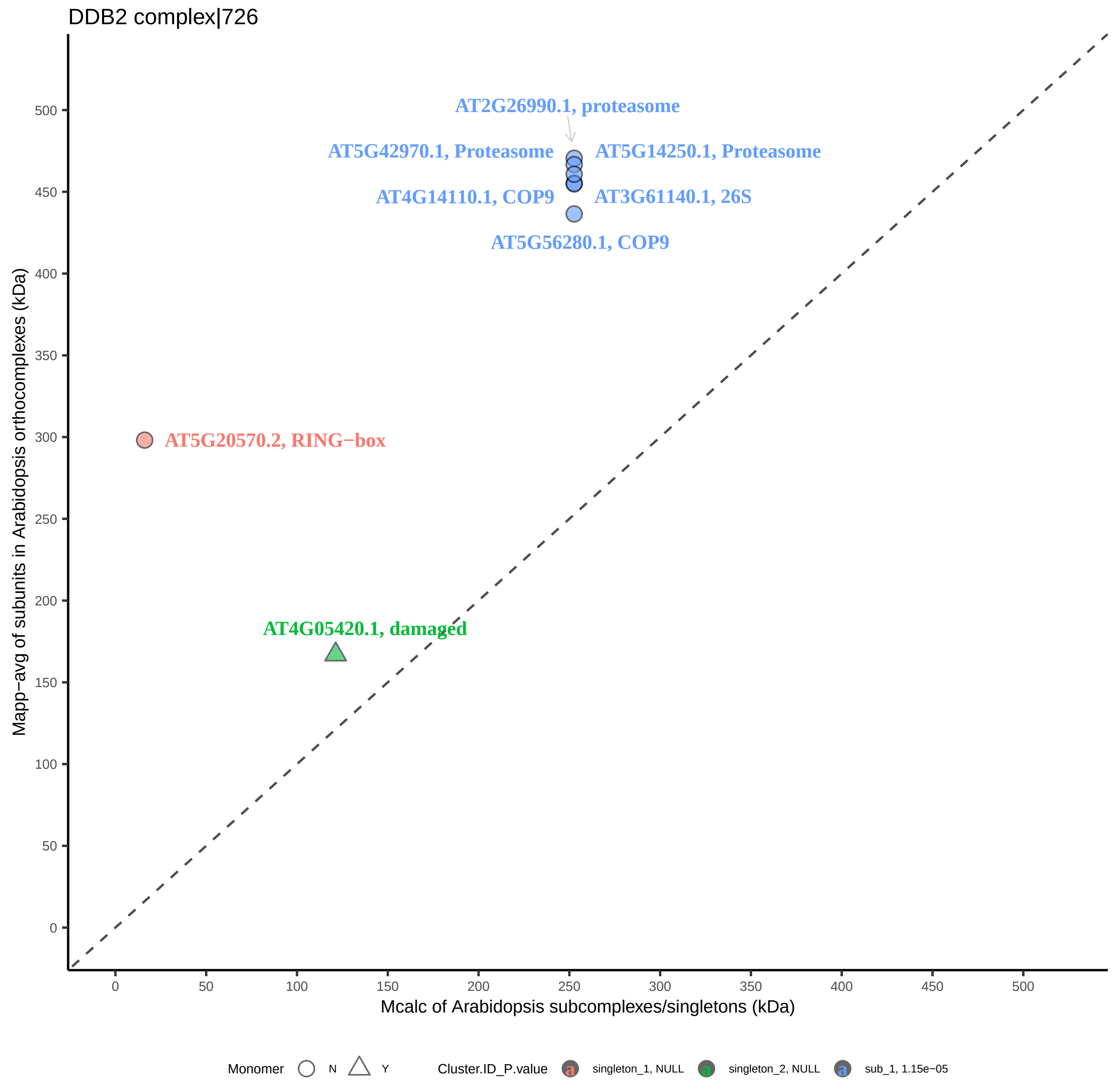
DDB2 complex|726
AT2G26990.1, proteasome
AT5G42970.1, Proteasome
AT5G14250.1, Proteasome
AT4G14110.1, COP9
AT3G61140.1, 26S
AT5G56280.1, COP9
AT5G20570.2, RING−box
AT4G05420.1, damaged
Mapp−avg of subunits in Arabidopsis orthocomplexes (kDa)
Mcalc of Arabidopsis subcomplexes/singletons (kDa)
0
50
100
150
200
250
300
350
400
450
500
Monomer
N
Y
Cluster.ID_P.value
singleton_1, NULL
singleton_2, NULL
sub_1, 1.15e−05

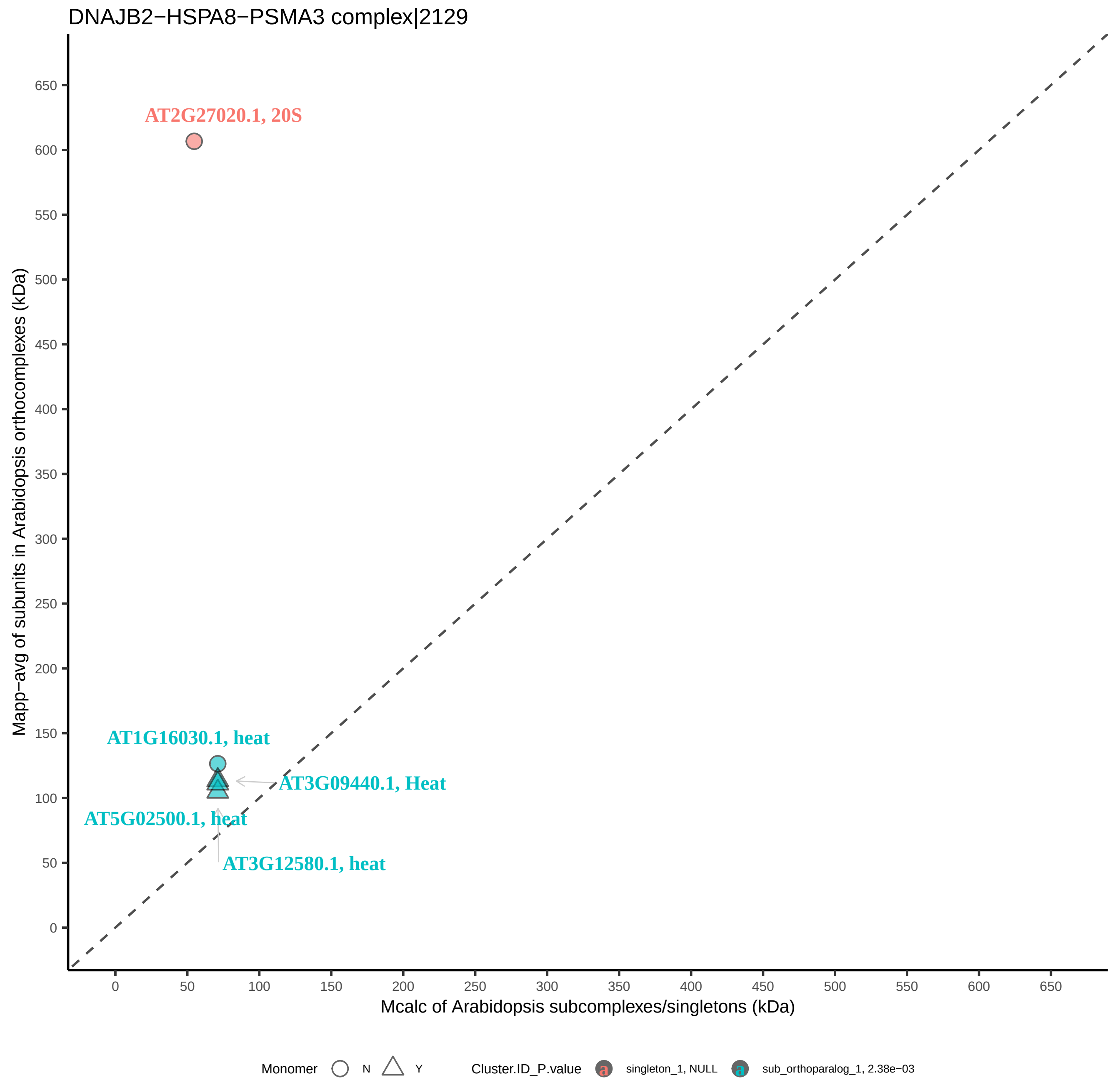

DNAJB2−HSPA8−PSMA3 complex|2129
AT2G27020.1, 20S
AT1G16030.1, heat
AT3G09440.1, Heat
AT5G02500.1, heat
AT3G12580.1, heat
Mapp−avg of subunits in Arabidopsis orthocomplexes (kDa)
Mcalc of Arabidopsis subcomplexes/singletons (kDa)
0
50
100
150
200
250
300
350
400
450
500
550
600
650
Monomer
N
Y
Cluster.ID_P.value
singleton_1, NULL
sub_orthoparalog_1, 2.38e−03

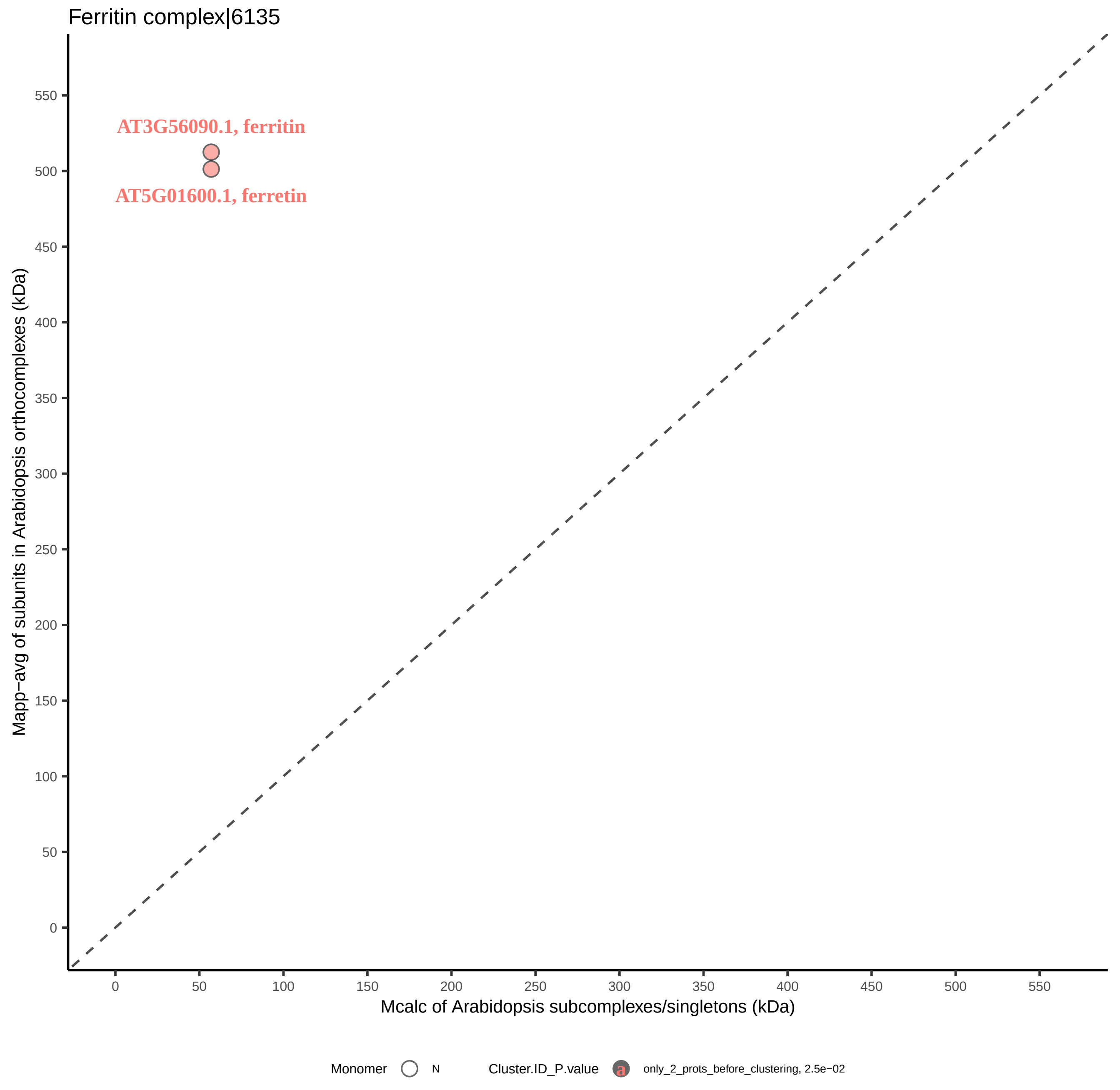
Ferritin complex|6135
AT3G56090.1, ferritin
AT5G01600.1, ferretin
Mapp−avg of subunits in Arabidopsis orthocomplexes (kDa)
Mcalc of Arabidopsis subcomplexes/singletons (kDa)
0
50
100
150
200
250
300
350
400
450
500
550
Monomer
N
Cluster.ID_P.value
a
only_2_prots_before_clustering, 2.5e−02

FIB−associated protein complex|1231

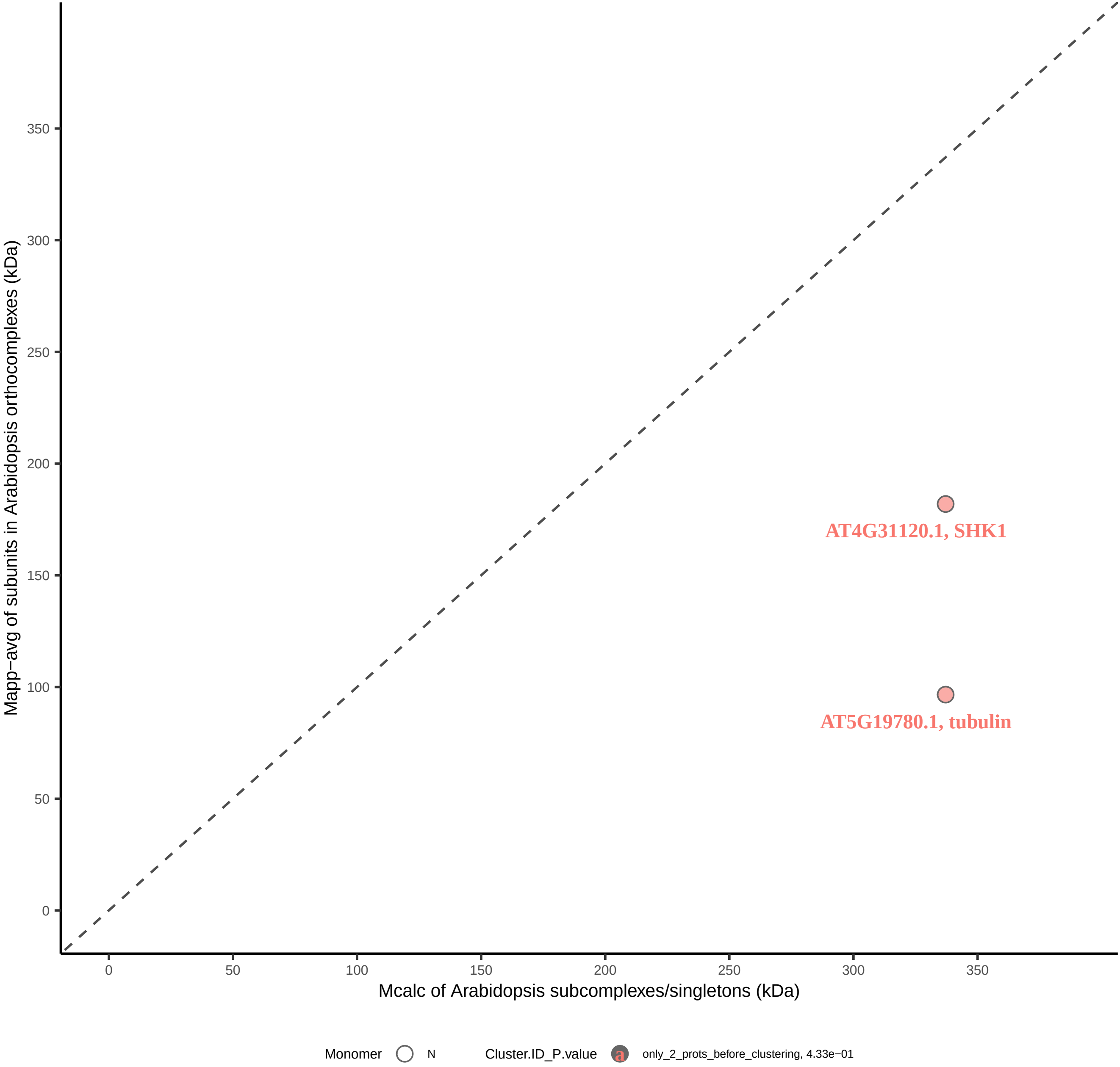

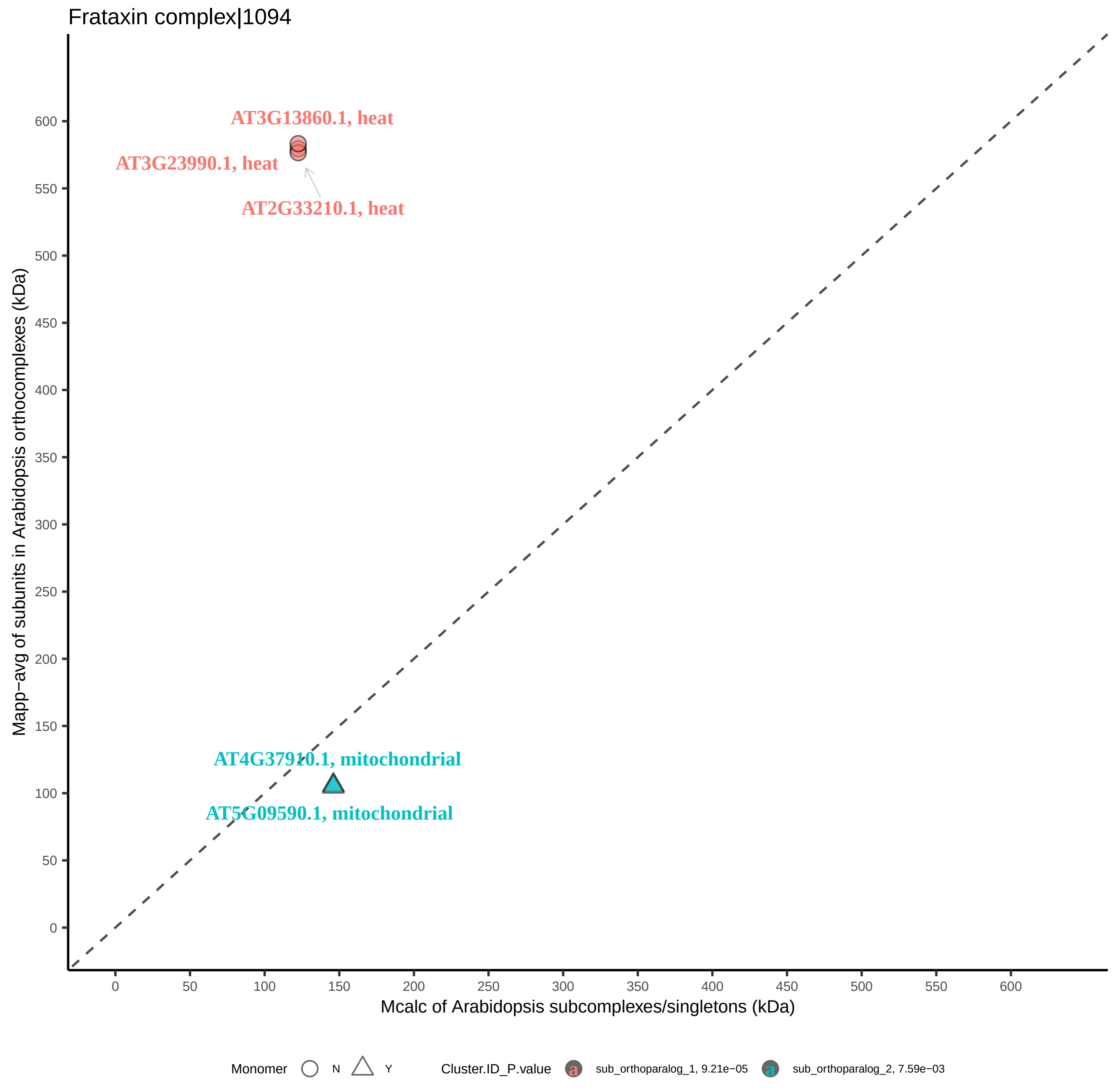
Frataxin complex|1094
AT3G13860.1, heat
AT3G23990.1, heat
AT2G33210.1, heat
AT4G37910.1, mitochondrial
AT5G09590.1, mitochondrial
Mapp−avg of subunits in Arabidopsis orthocomplexes (kDa)
Mcalc of Arabidopsis subcomplexes/singletons (kDa)
0
50
100
150
200
250
300
350
400
450
500
550
600
Monomer
N
Y
Cluster.ID_P.value
sub_orthoparalog_1, 9.21e−05
sub_orthoparalog_2, 7.59e−03

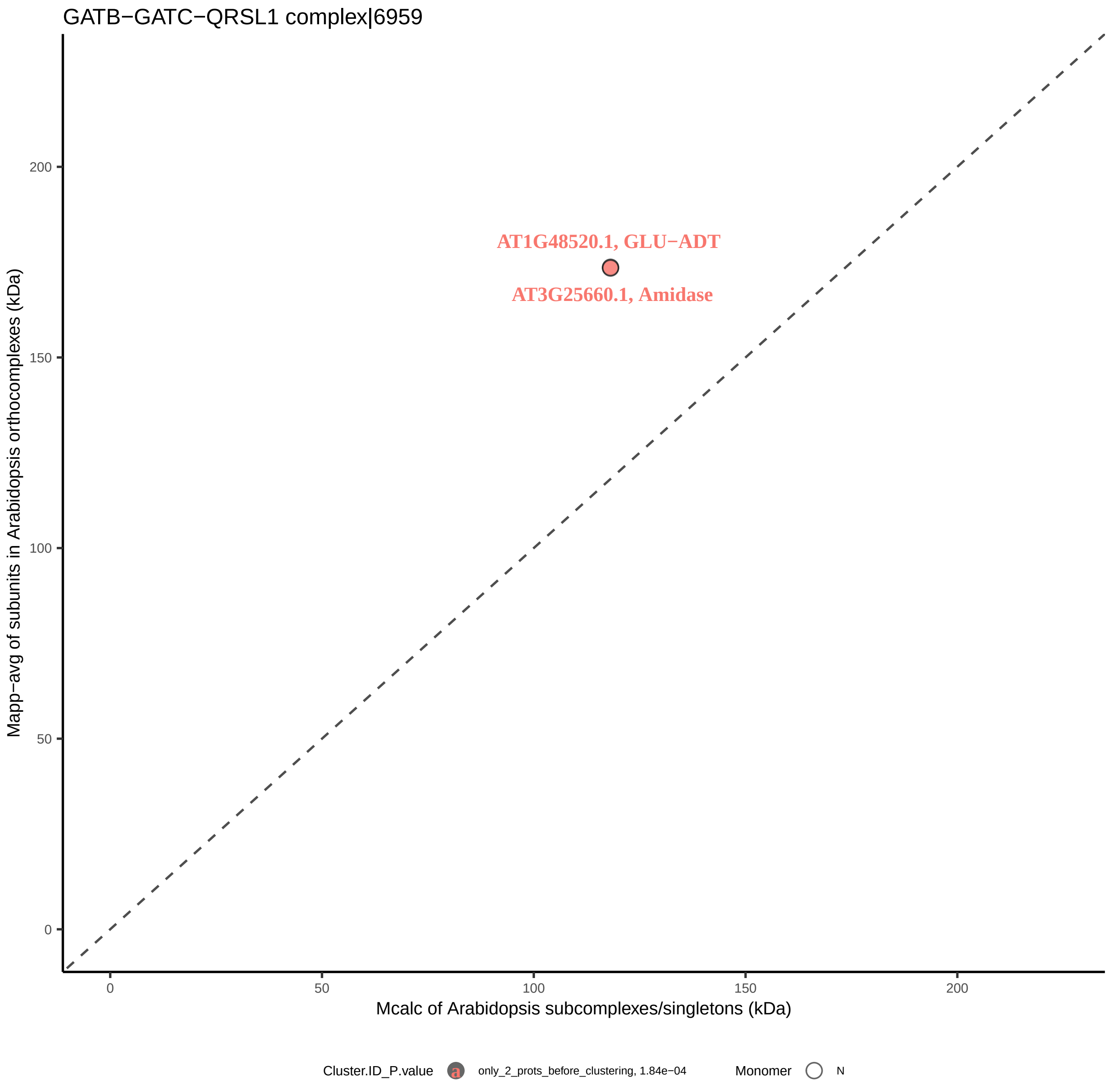

GATB−GATC−QRSL1 complex|6959
AT1G48520.1, GLU−ADT
AT3G25660.1, Amidase
Mapp−avg of subunits in Arabidopsis orthocomplexes (kDa)
Mcalc of Arabidopsis subcomplexes/singletons (kDa)
0
50
100
150
200
Cluster.ID_P.value
only_2_prots_before_clustering, 1.84e−04
Monomer
N

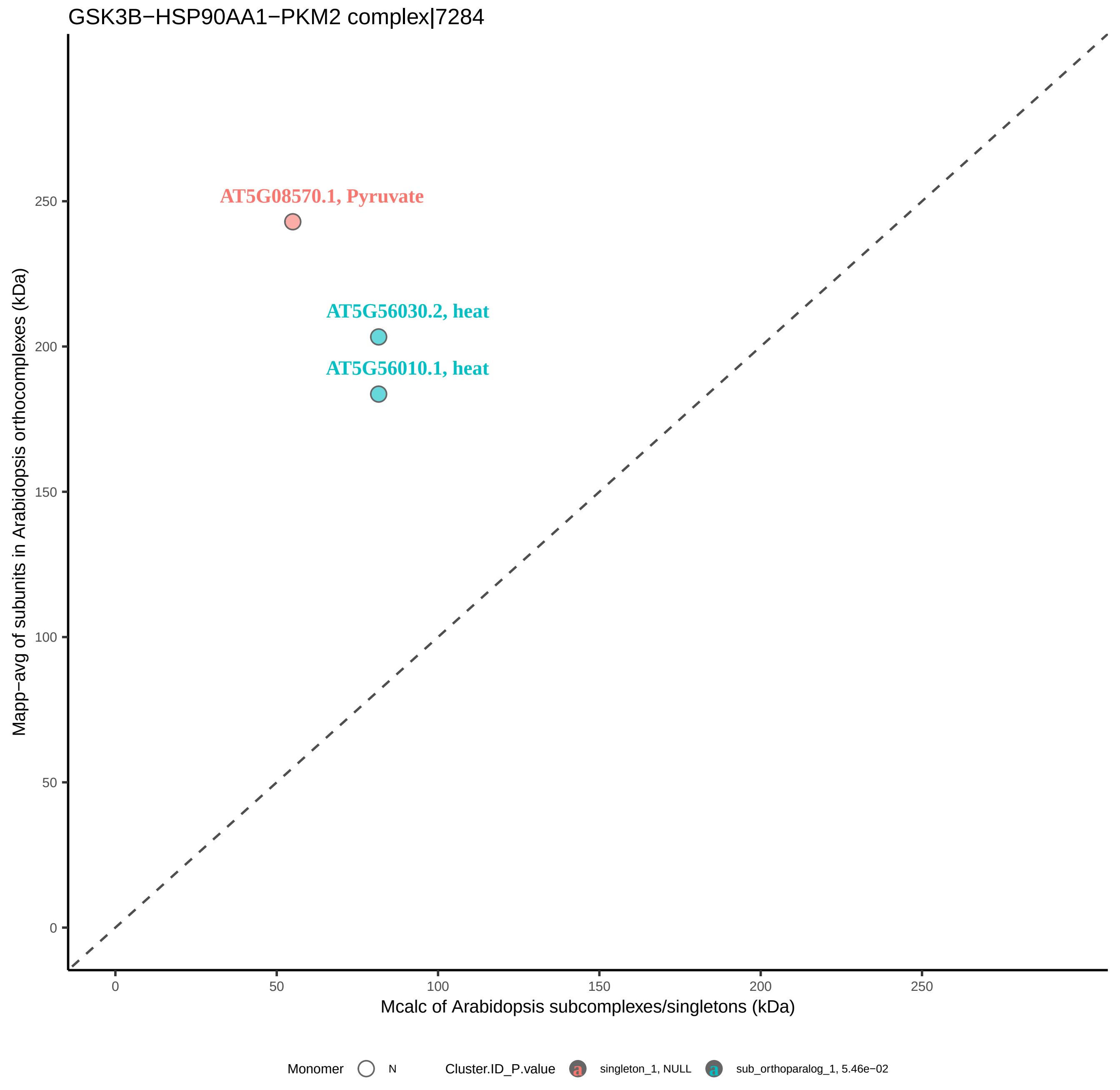
GSK3B−HSP90AA1−PKM2 complex|7284
AT5G08570.1, Pyruvate
AT5G56030.2, heat
AT5G56010.1, heat
Mapp−avg of subunits in Arabidopsis orthocomplexes (kDa)
Mcalc of Arabidopsis subcomplexes/singletons (kDa)
0
50
100
150
200
250
Monomer
N
Cluster.ID_P.value
singleton_1, NULL
sub_orthoparalog_1, 5.46e−02

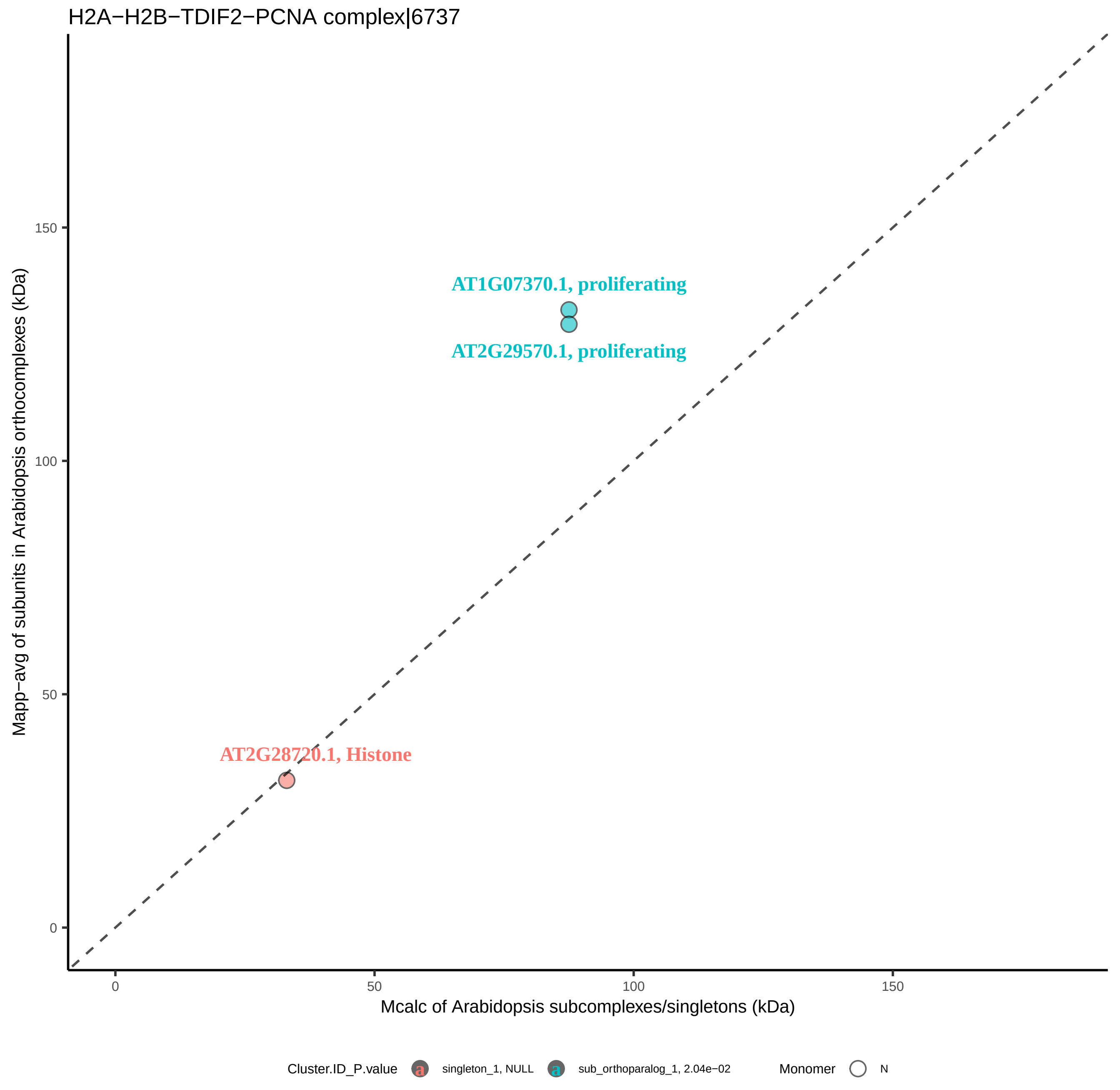
H2A−H2B−TDIF2−PCNA complex|6737
AT1G07370.1, proliferating
AT2G29570.1, proliferating
AT2G28720.1, Histone
Mapp−avg of subunits in Arabidopsis orthocomplexes (kDa)
Mcalc of Arabidopsis subcomplexes/singletons (kDa)
0
50
100
150
Cluster.ID_P.value
singleton_1, NULL
sub_orthoparalog_1, 2.04e−02
Monomer
N

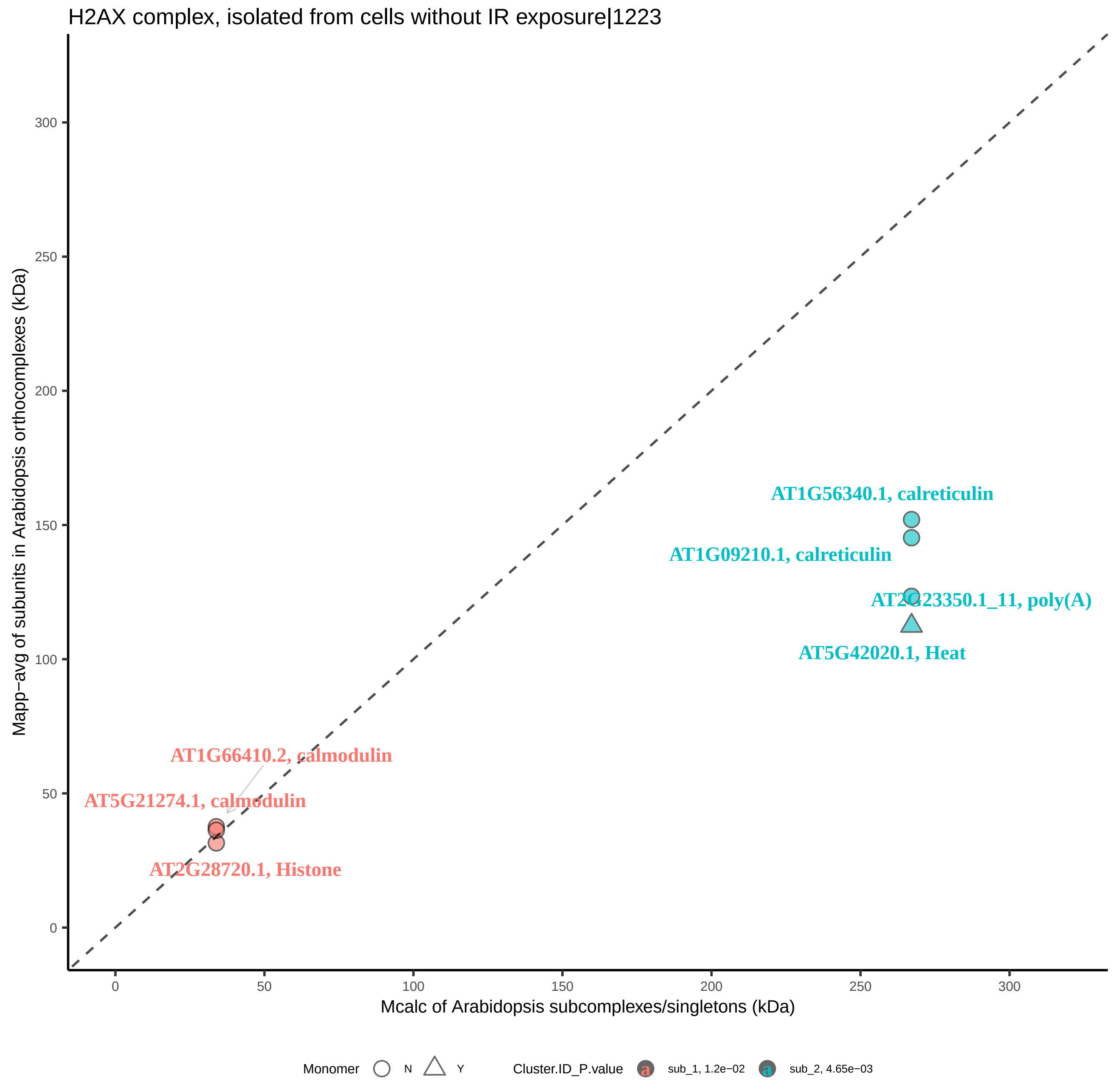

H2AX complex, isolated from cells without IR exposure|1223
Mapp−avg of subunits in Arabidopsis orthocomplexes (kDa)
Mcalc of Arabidopsis subcomplexes/singletons (kDa)
0
50
100
150
200
250
300
AT1G56340.1, calreticulin
AT1G09210.1, calreticulin
AT2G23350.1_11, poly(A)
AT5G42020.1, Heat
AT1G66410.2, calmodulin
AT5G21274.1, calmodulin
AT2G28720.1, Histone
Monomer
N
Y
Cluster.ID_P.value
sub_1, 1.2e−02
sub_2, 4.65e−03

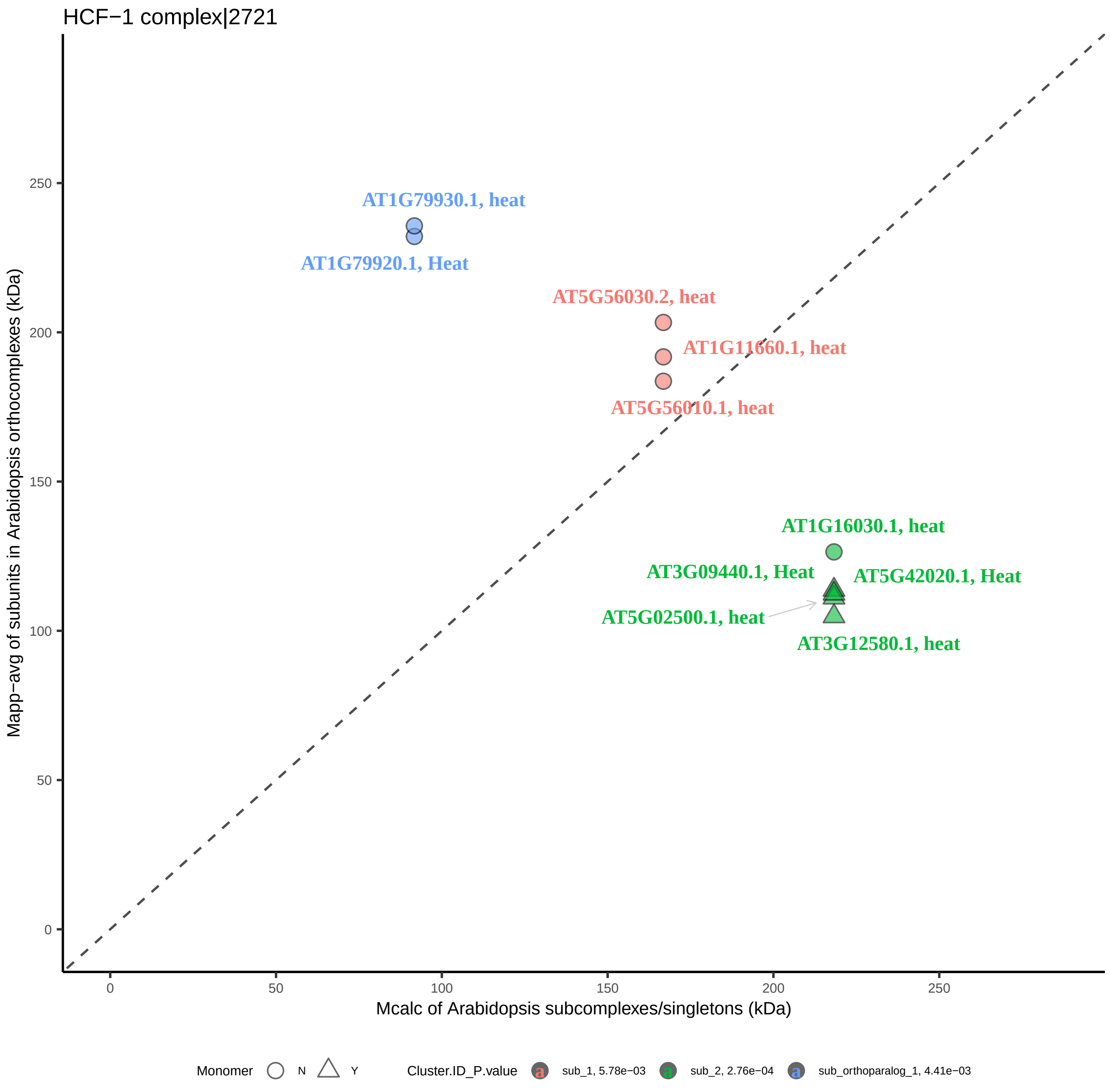
HCF−1 complex|2721
AT1G79930.1, heat
AT1G79920.1, Heat
AT5G56030.2, heat
AT1G11660.1, heat
AT5G56010.1, heat
AT1G16030.1, heat
AT3G09440.1, Heat
AT5G42020.1, Heat
AT5G02500.1, heat
AT3G12580.1, heat
Mapp−avg of subunits in Arabidopsis orthocomplexes (kDa)
Mcalc of Arabidopsis subcomplexes/singletons (kDa)
0
50
100
150
200
250
Monomer N Y
Cluster.ID_P.value
sub_1, 5.78e−03
sub_2, 2.76e−04
sub_orthoparalog_1, 4.41e−03

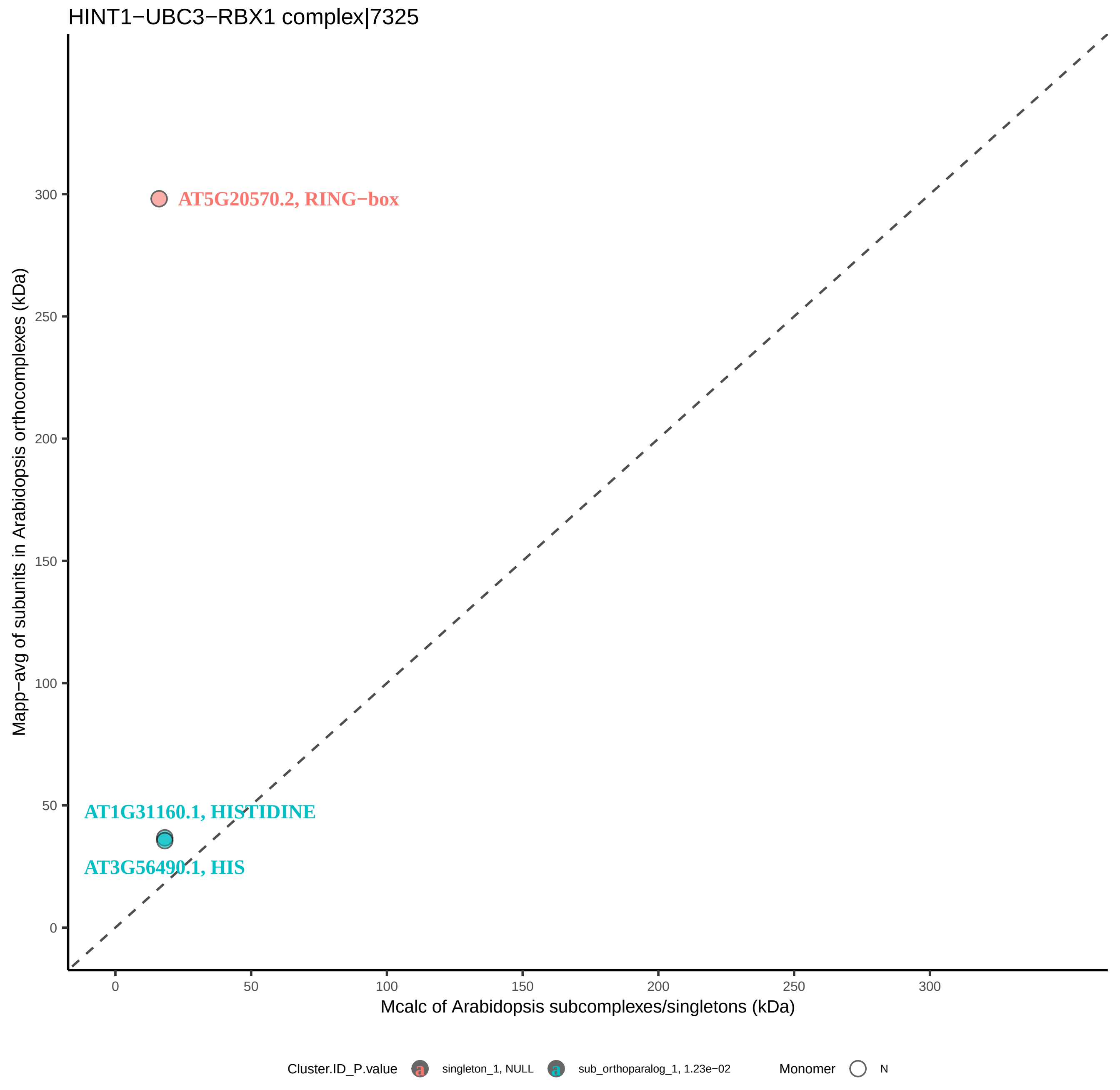
HINT1−UBC3−RBX1 complex|7325
AT5G20570.2, RING−box
AT1G31160.1, HISTIDINE
AT3G56490.1, HIS
Mapp−avg of subunits in Arabidopsis orthocomplexes (kDa)
Mcalc of Arabidopsis subcomplexes/singletons (kDa)
0
50
100
150
200
250
300
Cluster.ID_P.value
singleton_1, NULL
sub_orthoparalog_1, 1.23e−02
Monomer
N

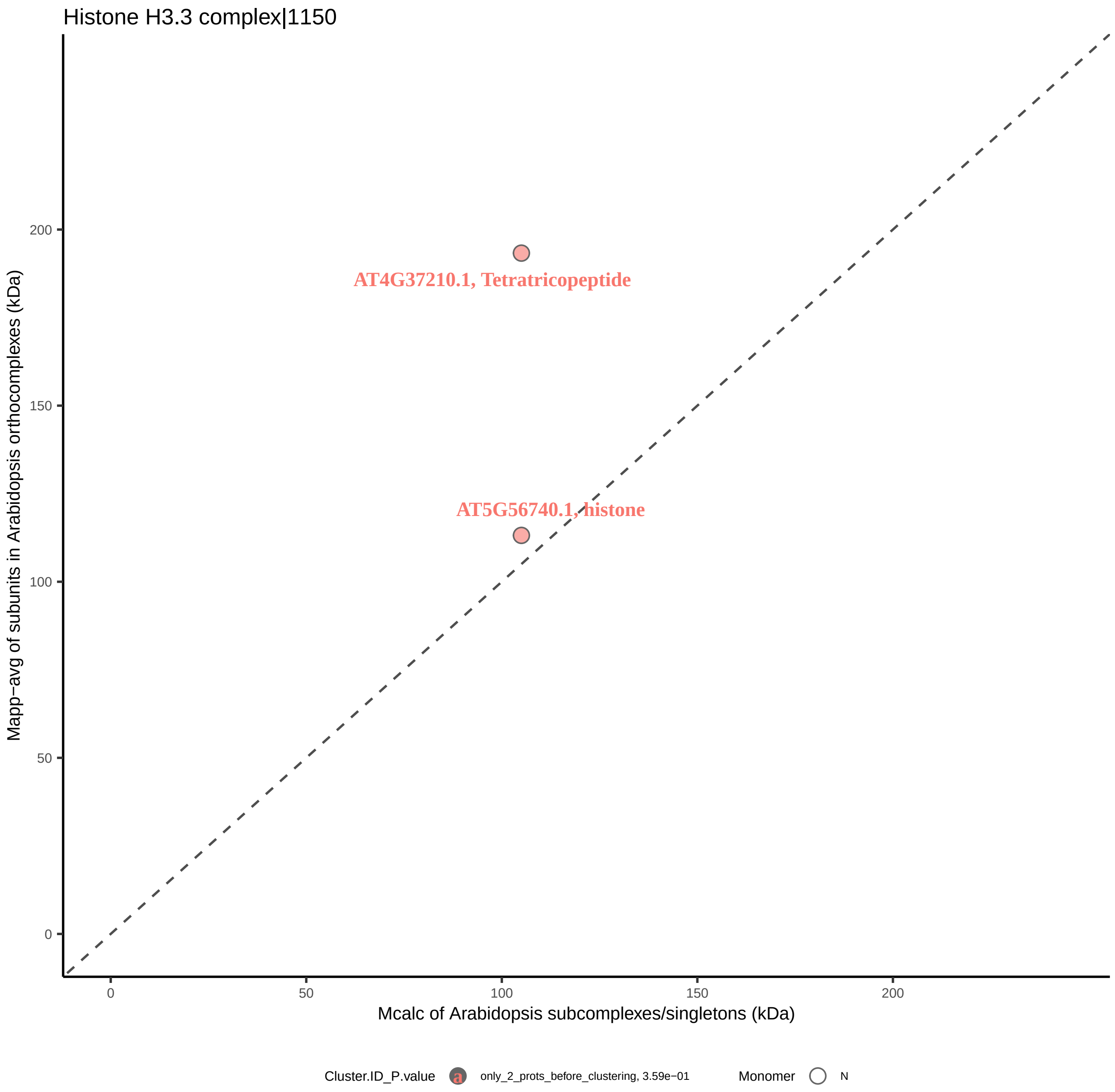

Histone H3.3 complex|1150
AT4G37210.1, Tetratricopeptide
AT5G56740.1, histone
Mapp−avg of subunits in Arabidopsis orthocomplexes (kDa)
Mcalc of Arabidopsis subcomplexes/singletons (kDa)
0
50
100
150
200
Cluster.ID_P.value
a
only_2_prots_before_clustering, 3.59e−01
Monomer
N

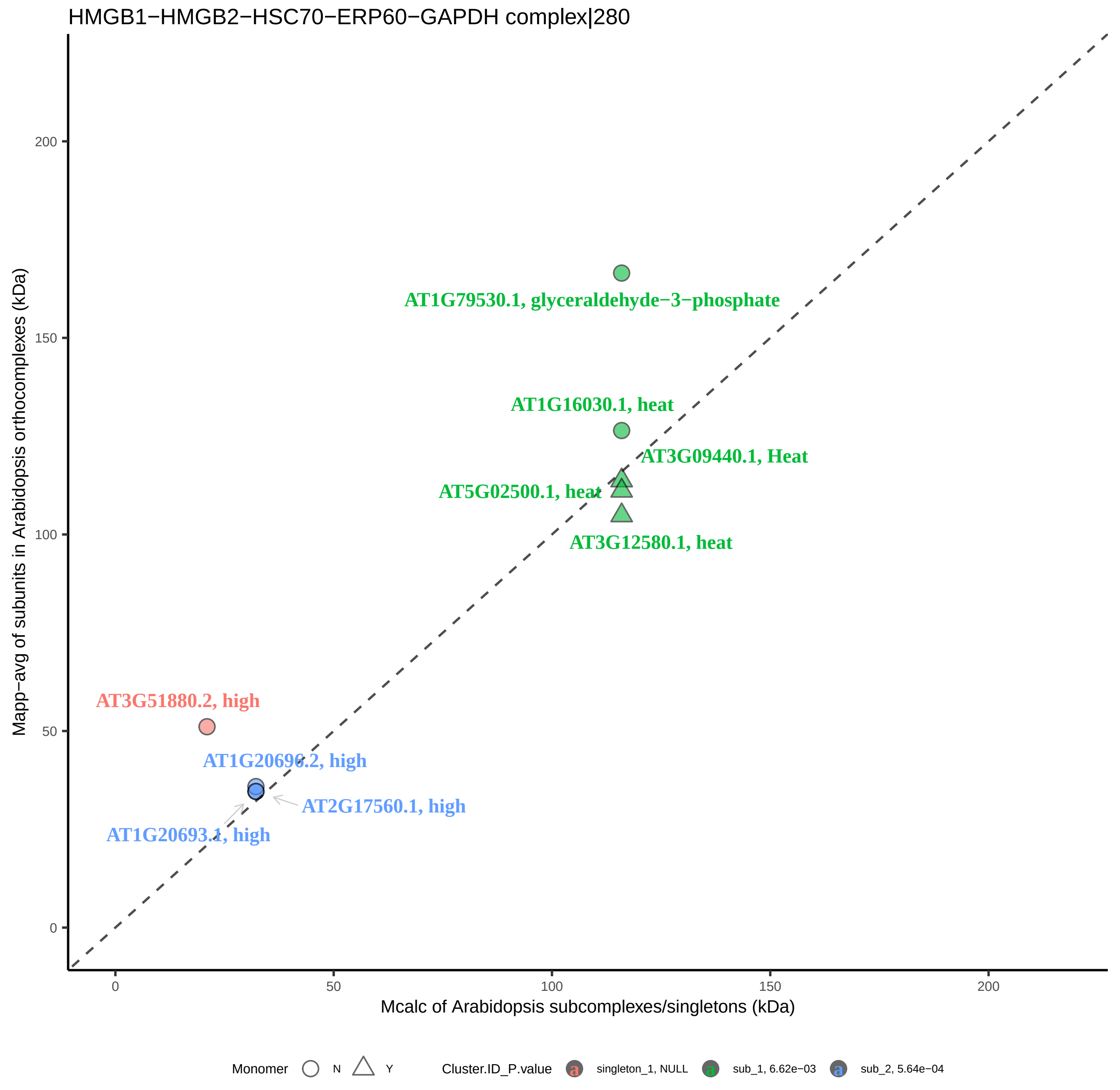

HMGB1−HMGB2−HSC70−ERP60−GAPDH complex|280
Mapp−avg of subunits in Arabidopsis orthocomplexes (kDa)
Mcalc of Arabidopsis subcomplexes/singletons (kDa)
0
50
100
150
200
AT1G79530.1, glyceraldehyde−3−phosphate
AT1G16030.1, heat
AT3G09440.1, Heat
AT5G02500.1, heat
AT3G12580.1, heat
AT3G51880.2, high
AT1G20696.2, high
AT2G17560.1, high
AT1G20693.1, high
Monomer
N
Y
Cluster.ID_P.value
singleton_1, NULL
sub_1, 6.62e−03
sub_2, 5.64e−04

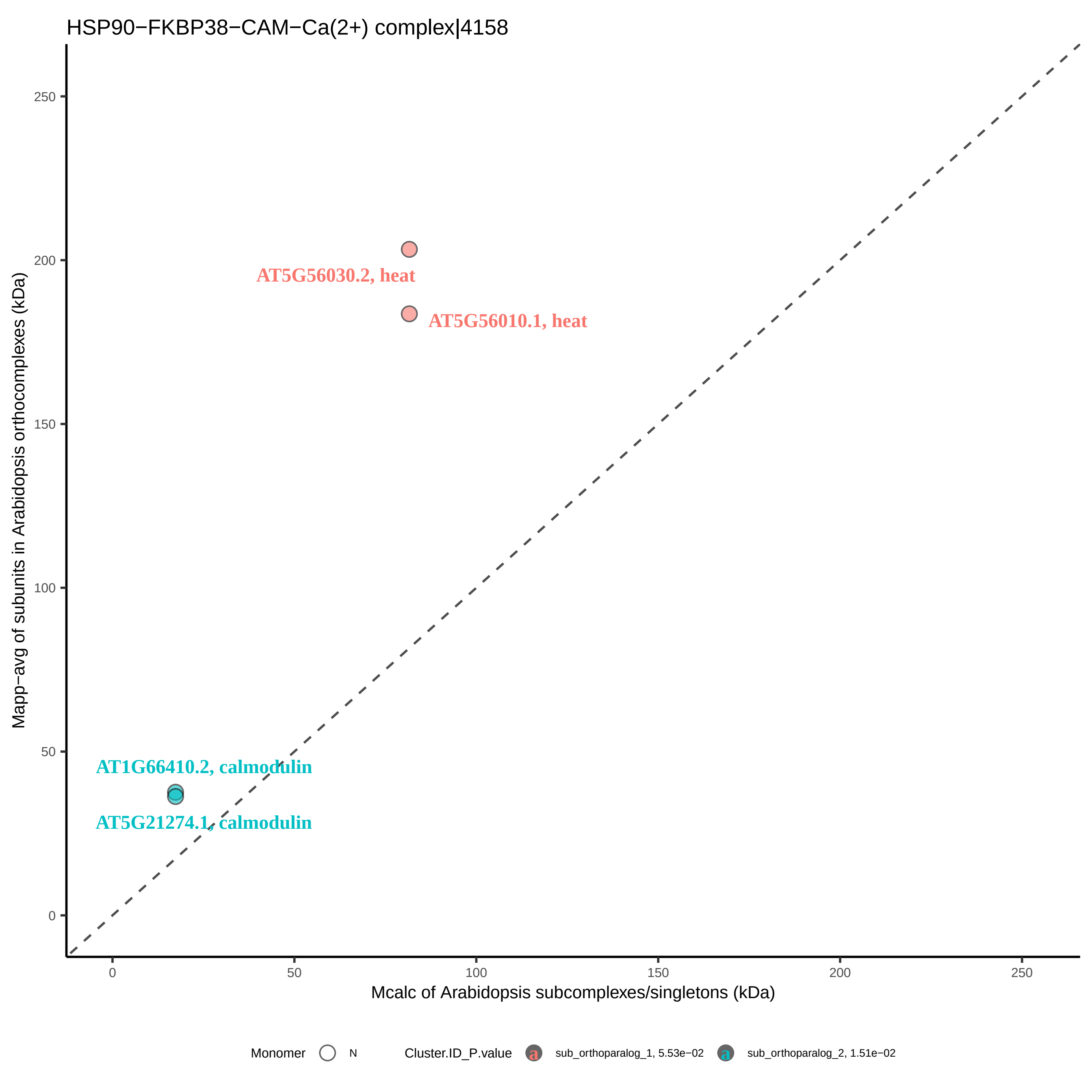
HSP90−FKBP38−CAM−Ca(2+) complex|4158
Mapp−avg of subunits in Arabidopsis orthocomplexes (kDa)
Mcalc of Arabidopsis subcomplexes/singletons (kDa)
0
50
100
150
200
250
AT5G56030.2, heat
AT5G56010.1, heat
AT1G66410.2, calmodulin
AT5G21274.1, calmodulin
Monomer
N
Cluster.ID_P.value
sub_orthoparalog_1, 5.53e−02
sub_orthoparalog_2, 1.51e−02

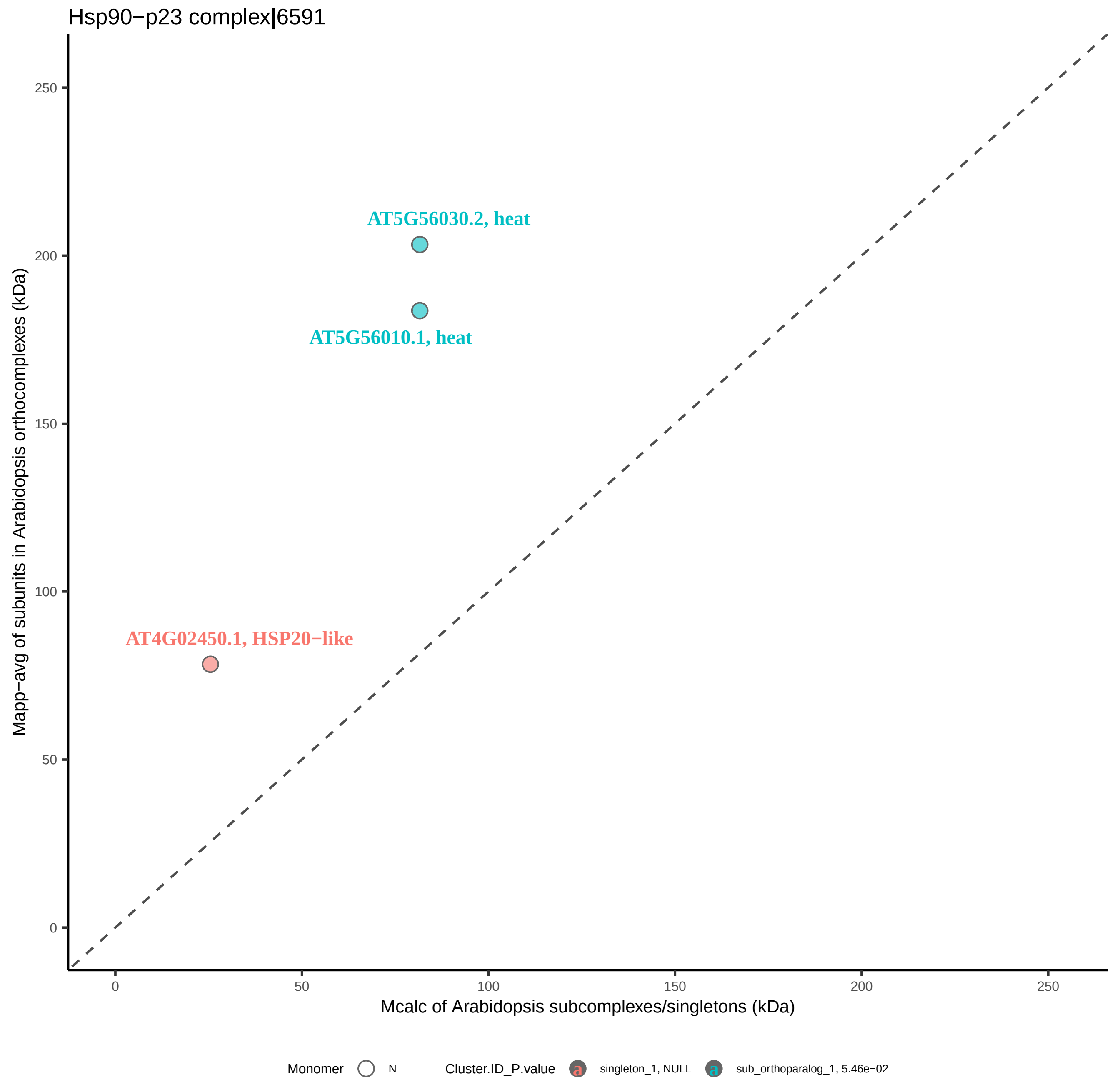

Hsp90−p23 complex|6591
AT5G56030.2, heat
AT5G56010.1, heat
AT4G02450.1, HSP20−like
Mapp−avg of subunits in Arabidopsis orthocomplexes (kDa)
Mcalc of Arabidopsis subcomplexes/singletons (kDa)
0
50
100
150
200
250
Monomer
N
Cluster.ID_P.value
singleton_1, NULL
sub_orthoparalog_1, 5.46e−02

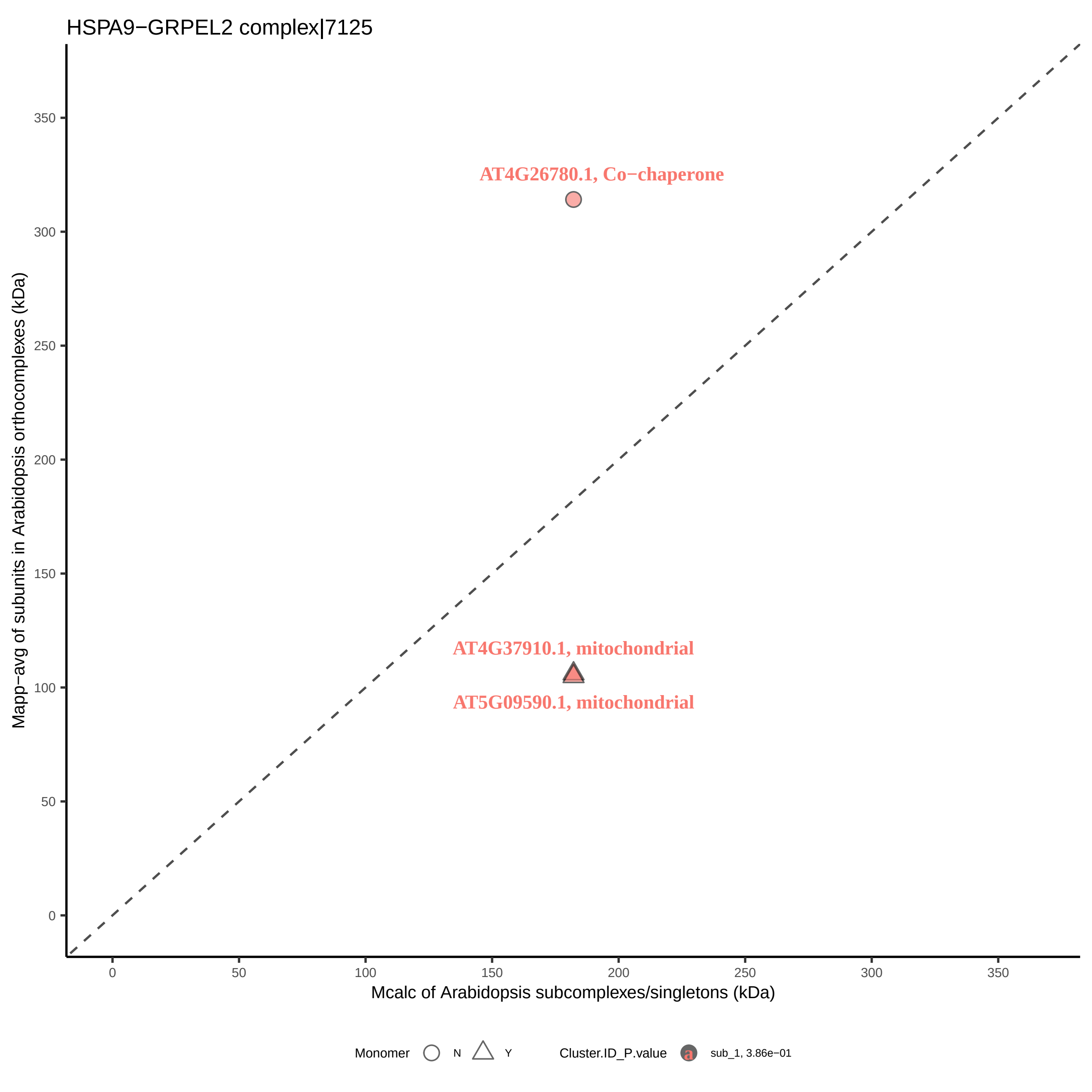

HSPA9−GRPEL2 complex|7125
AT4G26780.1, Co−chaperone
AT4G37910.1, mitochondrial
AT5G09590.1, mitochondrial
Mapp−avg of subunits in Arabidopsis orthocomplexes (kDa)
Mcalc of Arabidopsis subcomplexes/singletons (kDa)
0
50
100
150
200
250
300
350
Monomer
N
Y
Cluster.ID_P.value
sub_1, 3.86e−01

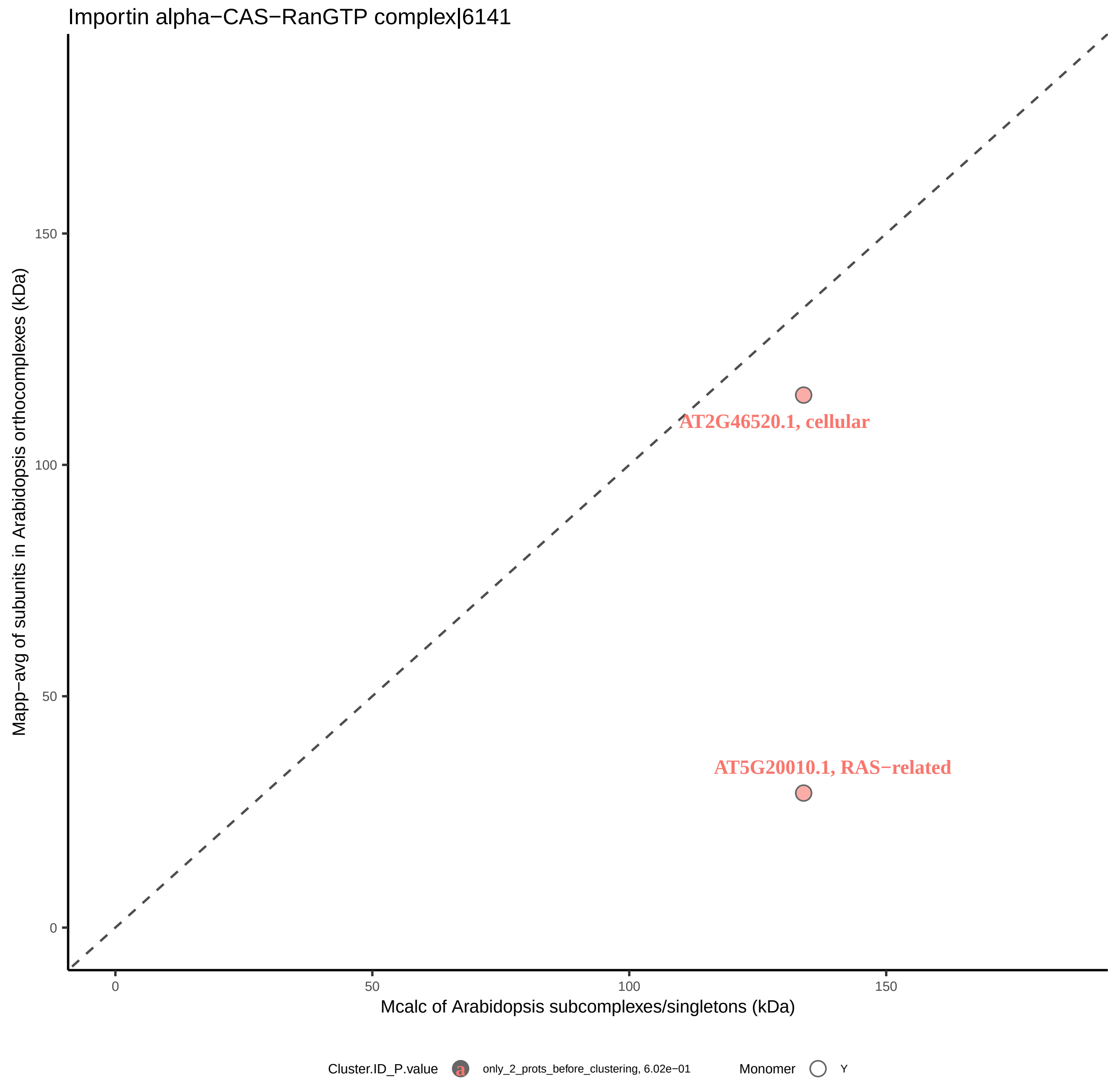
Importin alpha−CAS−RanGTP complex|6141
AT2G46520.1, cellular
AT5G20010.1, RAS−related
150
100
50
0
0
50
100
150
Mapp−avg of subunits in Arabidopsis orthocomplexes (kDa)
Mcalc of Arabidopsis subcomplexes/singletons (kDa)
Cluster.ID_P.value
a
only_2_prots_before_clustering, 6.02e−01
Monomer
Y

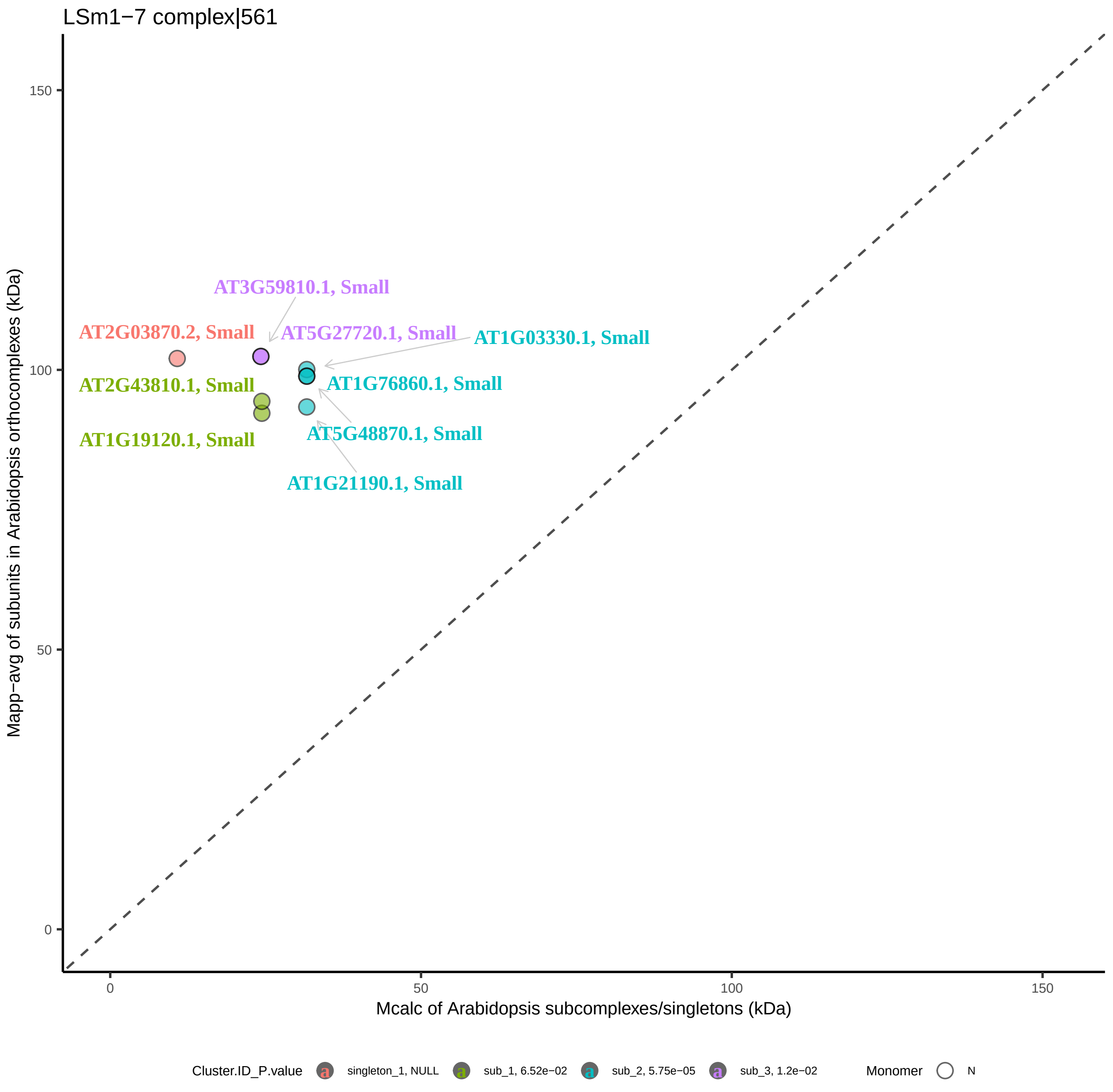

LSm1−7 complex|561
Mapp−avg of subunits in Arabidopsis orthocomplexes (kDa)
Mcalc of Arabidopsis subcomplexes/singletons (kDa)
0
50
100
150
AT3G59810.1, Small
AT2G03870.2, Small
AT5G27720.1, Small
AT1G03330.1, Small
AT2G43810.1, Small
AT1G76860.1, Small
AT5G48870.1, Small
AT1G19120.1, Small
AT1G21190.1, Small
Cluster.ID_P.value
singleton_1, NULL
sub_1, 6.52e−02
sub_2, 5.75e−05
sub_3, 1.2e−02
Monomer
N

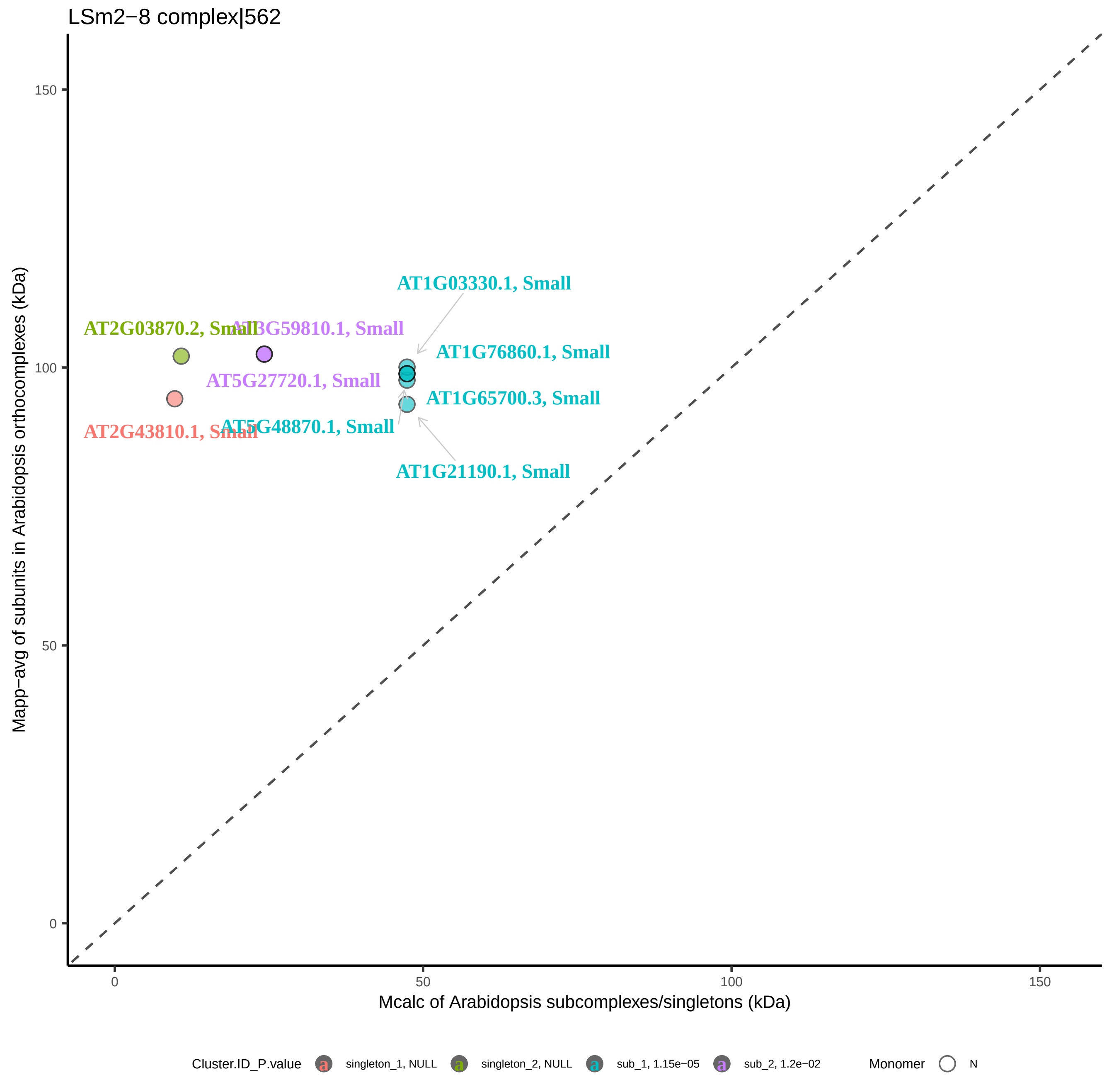

LSm2−8 complex|562
Mapp−avg of subunits in Arabidopsis orthocomplexes (kDa)
Mcalc of Arabidopsis subcomplexes/singletons (kDa)
0
50
100
150
AT1G03330.1, Small
AT2G03870.2, Small
AT3G59810.1, Small
AT1G76860.1, Small
AT5G27720.1, Small
AT1G65700.3, Small
AT2G43810.1, Small
AT5G48870.1, Small
AT1G21190.1, Small
Cluster.ID_P.value
singleton_1, NULL
singleton_2, NULL
sub_1, 1.15e−05
sub_2, 1.2e−02
Monomer
N

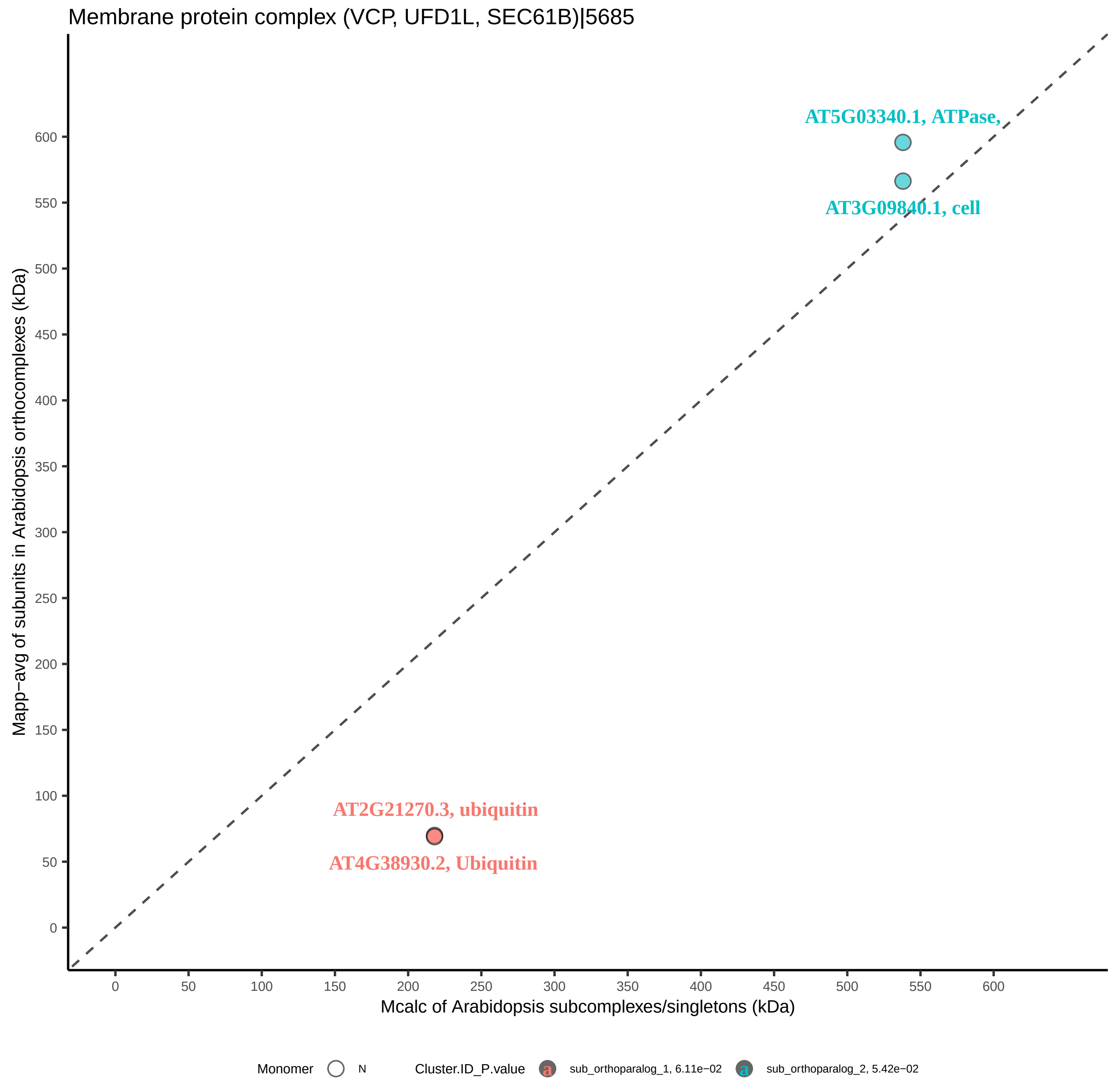
Membrane protein complex (VCP, UFD1L, SEC61B)|5685
AT5G03340.1, ATPase,
AT3G09840.1, cell
AT2G21270.3, ubiquitin
AT4G38930.2, Ubiquitin
Mapp−avg of subunits in Arabidopsis orthocomplexes (kDa)
Mcalc of Arabidopsis subcomplexes/singletons (kDa)
0
50
100
150
200
250
300
350
400
450
500
550
600
Monomer
N
Cluster.ID_P.value
sub_orthoparalog_1, 6.11e−02
sub_orthoparalog_2, 5.42e−02

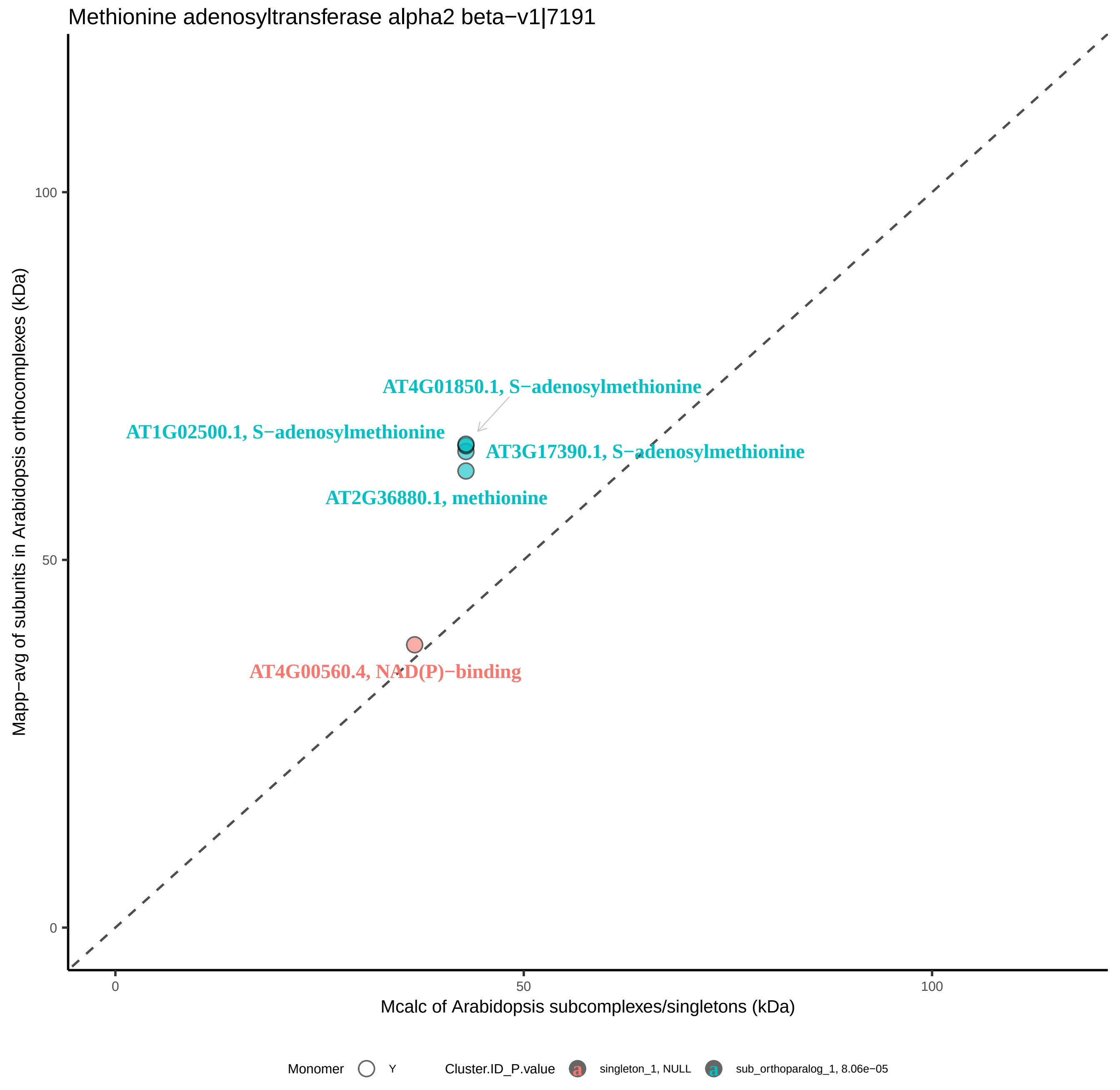

Methionine adenosyltransferase alpha2 beta−v1|7191
AT4G01850.1, S−adenosylmethionine
AT1G02500.1, S−adenosylmethionine
AT3G17390.1, S−adenosylmethionine
AT2G36880.1, methionine
AT4G00560.4, NAD(P)−binding
100
50
0
0
50
100
Mapp−avg of subunits in Arabidopsis orthocomplexes (kDa)
Mcalc of Arabidopsis subcomplexes/singletons (kDa)
Monomer
Y
Cluster.ID_P.value
singleton_1, NULL
sub_orthoparalog_1, 8.06e−05

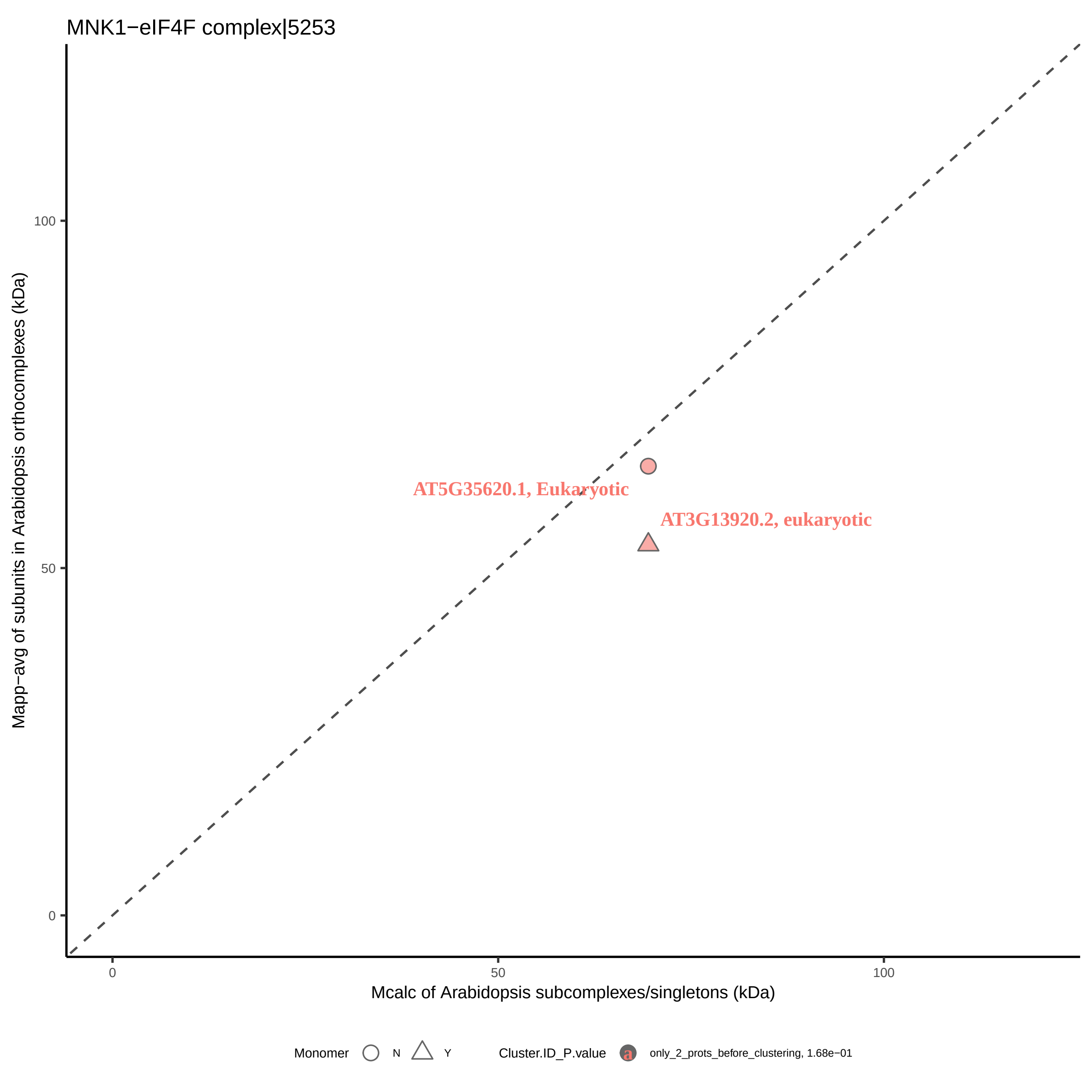
MNK1−eIF4F complex|5253
Mapp−avg of subunits in Arabidopsis orthocomplexes (kDa)
100
50
0
AT5G35620.1, Eukaryotic
AT3G13920.2, eukaryotic
0
50
100
Mcalc of Arabidopsis subcomplexes/singletons (kDa)
Monomer
N
Y
Cluster.ID_P.value
a
only_2_prots_before_clustering, 1.68e−01

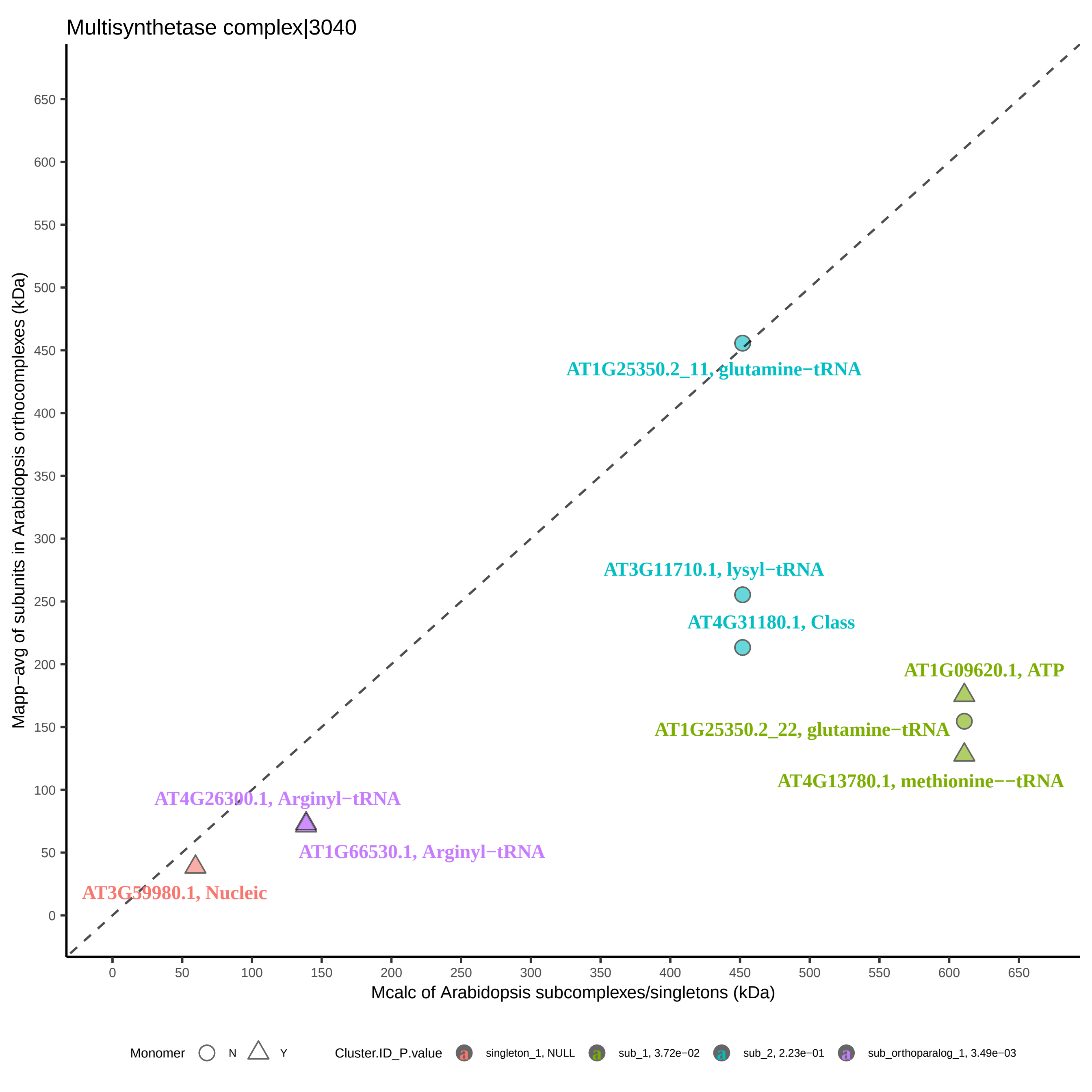

Multisynthetase complex|3040
Mapp−avg of subunits in Arabidopsis orthocomplexes (kDa)
Mcalc of Arabidopsis subcomplexes/singletons (kDa)
0
50
100
150
200
250
300
350
400
450
500
550
600
650
AT1G25350.2_11, glutamine−tRNA
AT3G11710.1, lysyl−tRNA
AT4G31180.1, Class
AT1G09620.1, ATP
AT1G25350.2_22, glutamine−tRNA
AT4G13780.1, methionine−−tRNA
AT4G26300.1, Arginyl−tRNA
AT1G66530.1, Arginyl−tRNA
AT3G59980.1, Nucleic
Monomer
N
Y
Cluster.ID_P.value
singleton_1, NULL
sub_1, 3.72e−02
sub_2, 2.23e−01
sub_orthoparalog_1, 3.49e−03

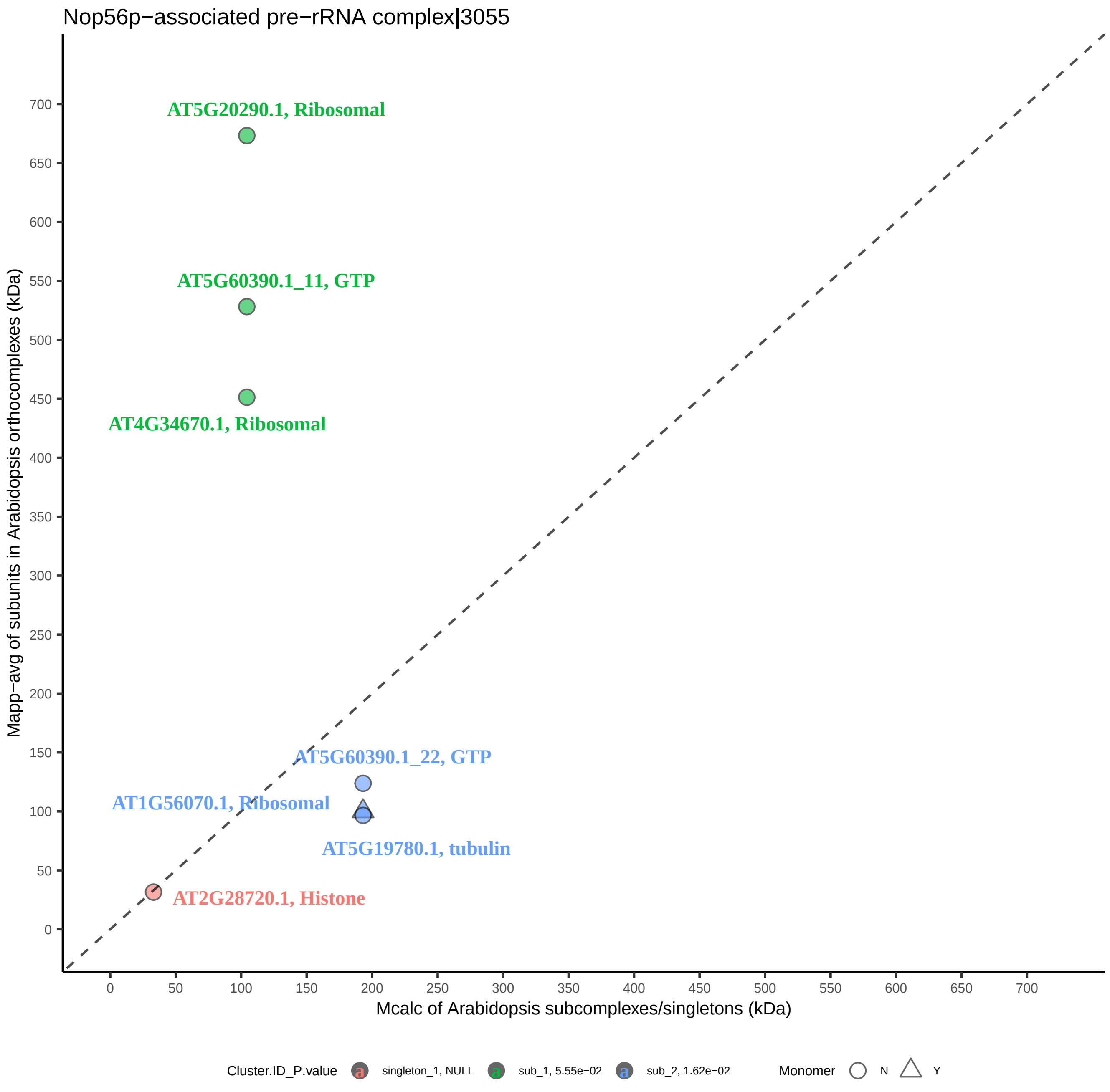

Nop56p−associated pre−rRNA complex|3055
AT5G20290.1, Ribosomal
AT5G60390.1_11, GTP
AT4G34670.1, Ribosomal
AT5G60390.1_22, GTP
AT1G56070.1, Ribosomal
AT5G19780.1, tubulin
AT2G28720.1, Histone
Mapp−avg of subunits in Arabidopsis orthocomplexes (kDa)
Mcalc of Arabidopsis subcomplexes/singletons (kDa)
0
50
100
150
200
250
300
350
400
450
500
550
600
650
700
Cluster.ID_P.value
singleton_1, NULL
sub_1, 5.55e−02
sub_2, 1.62e−02
Monomer
N
Y

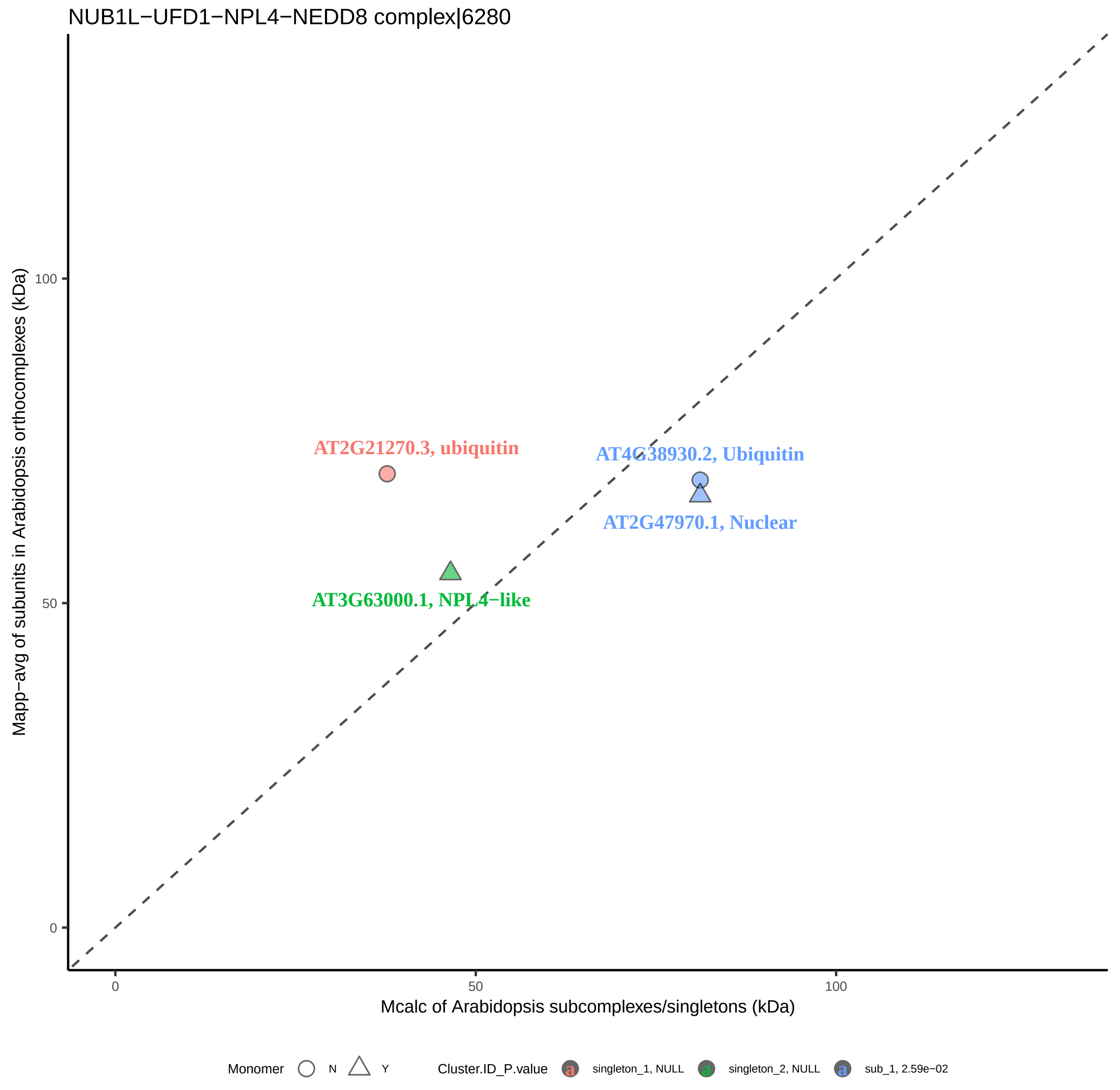

NUB1L−UFD1−NPL4−NEDD8 complex|6280
AT2G21270.3, ubiquitin
AT4G38930.2, Ubiquitin
AT2G47970.1, Nuclear
AT3G63000.1, NPL4−like
Mapp−avg of subunits in Arabidopsis orthocomplexes (kDa)
100
50
0
0
50
100
Mcalc of Arabidopsis subcomplexes/singletons (kDa)
Monomer
N
Y
Cluster.ID_P.value
singleton_1, NULL
singleton_2, NULL
sub_1, 2.59e−02

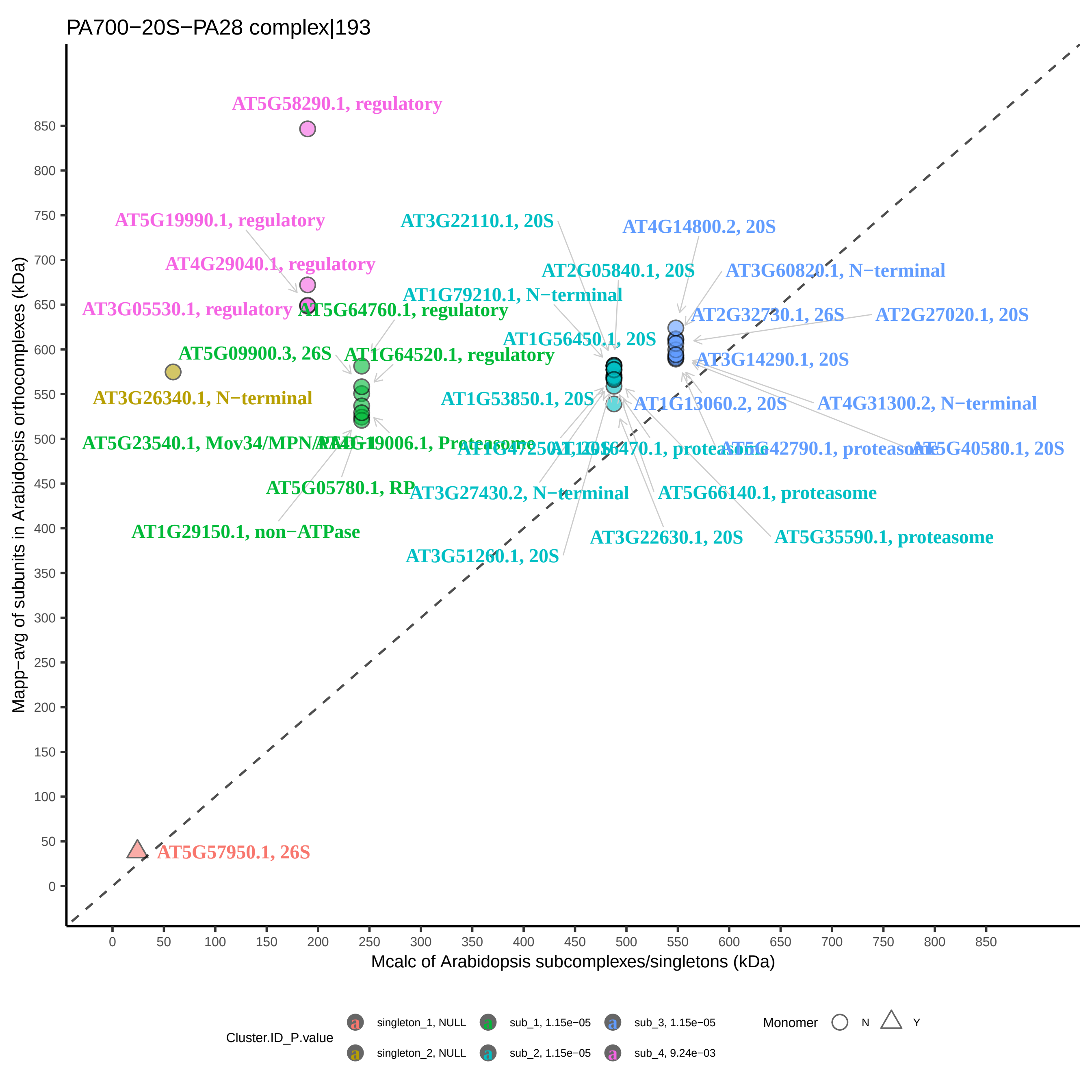

PA700−20S−PA28 complex|193
Mapp−avg of subunits in Arabidopsis orthocomplexes (kDa)
Mcalc of Arabidopsis subcomplexes/singletons (kDa)
0
50
100
150
200
250
300
350
400
450
500
550
600
650
700
750
800
850
AT5G58290.1, regulatory
AT5G19990.1, regulatory
AT4G29040.1, regulatory
AT3G05530.1, regulatory
AT5G64760.1, regulatory
AT5G09900.3, 26S
AT1G64520.1, regulatory
AT3G26340.1, N−terminal
AT5G05780.1, RP
AT1G29150.1, non−ATPase
AT3G22110.1, 20S
AT2G05840.1, 20S
AT1G79210.1, N−terminal
AT1G56450.1, 20S
AT1G53850.1, 20S
AT3G27430.2, N−terminal
AT5G66140.1, proteasome
AT3G22630.1, 20S
AT5G35590.1, proteasome
AT3G51260.1, 20S
AT4G14800.2, 20S
AT3G60820.1, N−terminal
AT2G32730.1, 26S
AT2G27020.1, 20S
AT3G14290.1, 20S
AT1G13060.2, 20S
AT4G31300.2, N−terminal
AT5G40580.1, 20S
AT5G57950.1, 26S
Cluster.ID_P.value
singleton_1, NULL
singleton_2, NULL
sub_1, 1.15e−05
sub_2, 1.15e−05
sub_3, 1.15e−05
sub_4, 9.24e−03
Monomer
N
Y

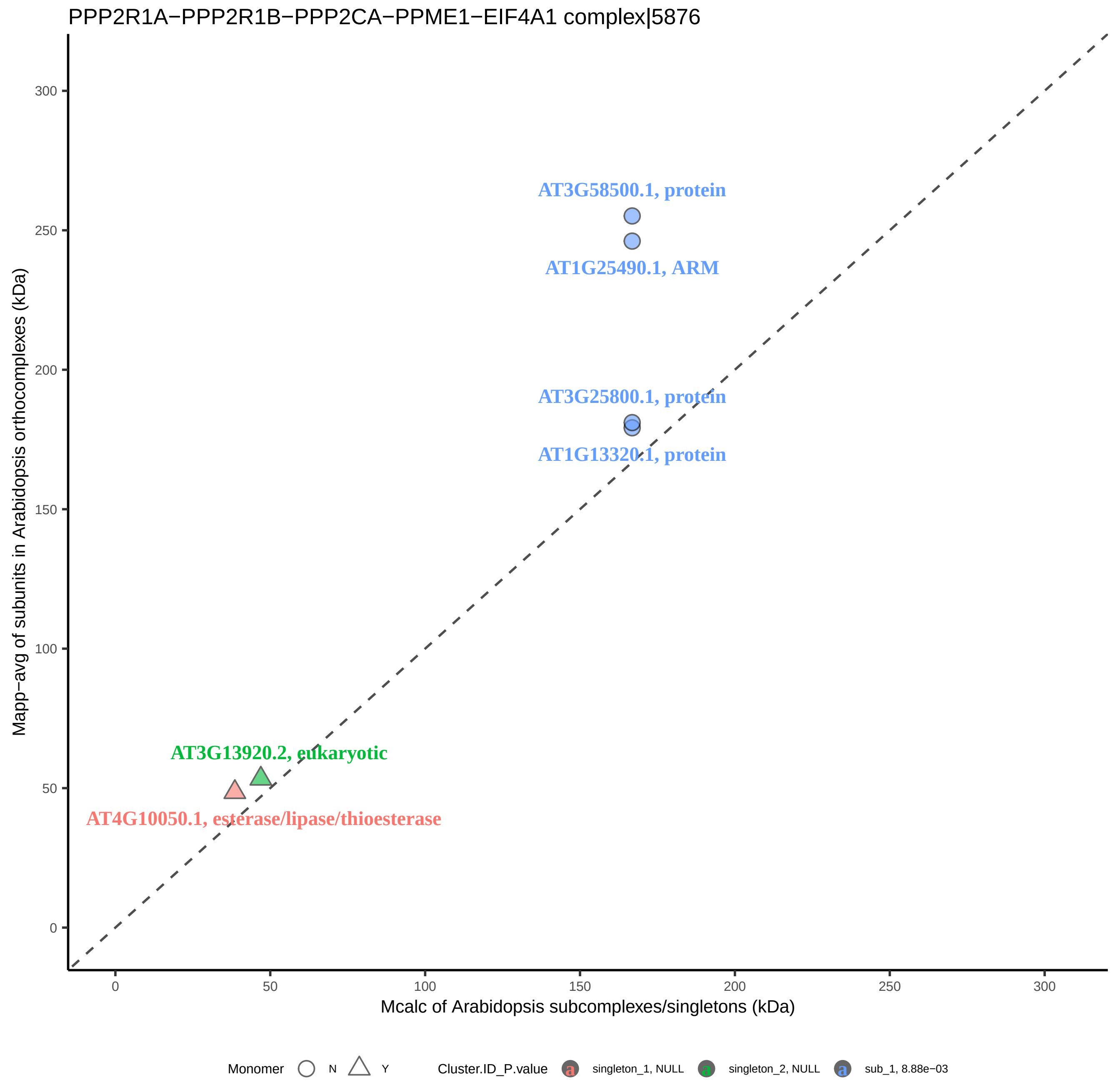

PPP2R1A−PPP2R1B−PPP2CA−PPME1−EIF4A1 complex|5876
AT3G58500.1, protein
AT1G25490.1, ARM
AT3G25800.1, protein
AT1G13320.1, protein
AT3G13920.2, eukaryotic
AT4G10050.1, esterase/lipase/thioesterase
Mapp−avg of subunits in Arabidopsis orthocomplexes (kDa)
Mcalc of Arabidopsis subcomplexes/singletons (kDa)
0
50
100
150
200
250
300
Monomer N Y
Cluster.ID_P.value singleton_1, NULL singleton_2, NULL sub_1, 8.88e−03

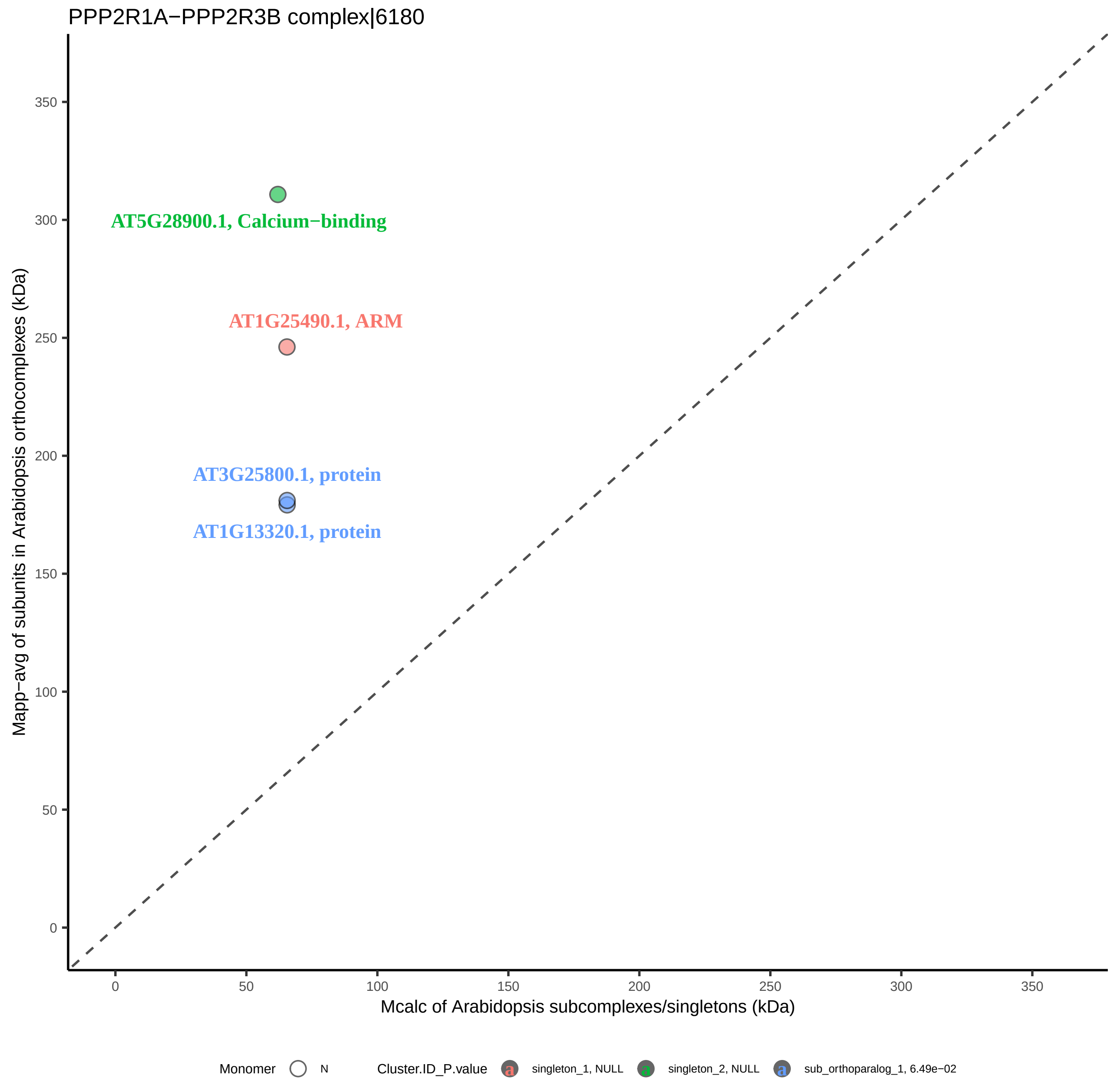

PPP2R1A−PPP2R3B complex|6180
AT5G28900.1, Calcium−binding
AT1G25490.1, ARM
AT3G25800.1, protein
AT1G13320.1, protein
Mapp−avg of subunits in Arabidopsis orthocomplexes (kDa)
Mcalc of Arabidopsis subcomplexes/singletons (kDa)
0
50
100
150
200
250
300
350
Monomer
N
Cluster.ID_P.value
singleton_1, NULL
singleton_2, NULL
sub_orthoparalog_1, 6.49e−02

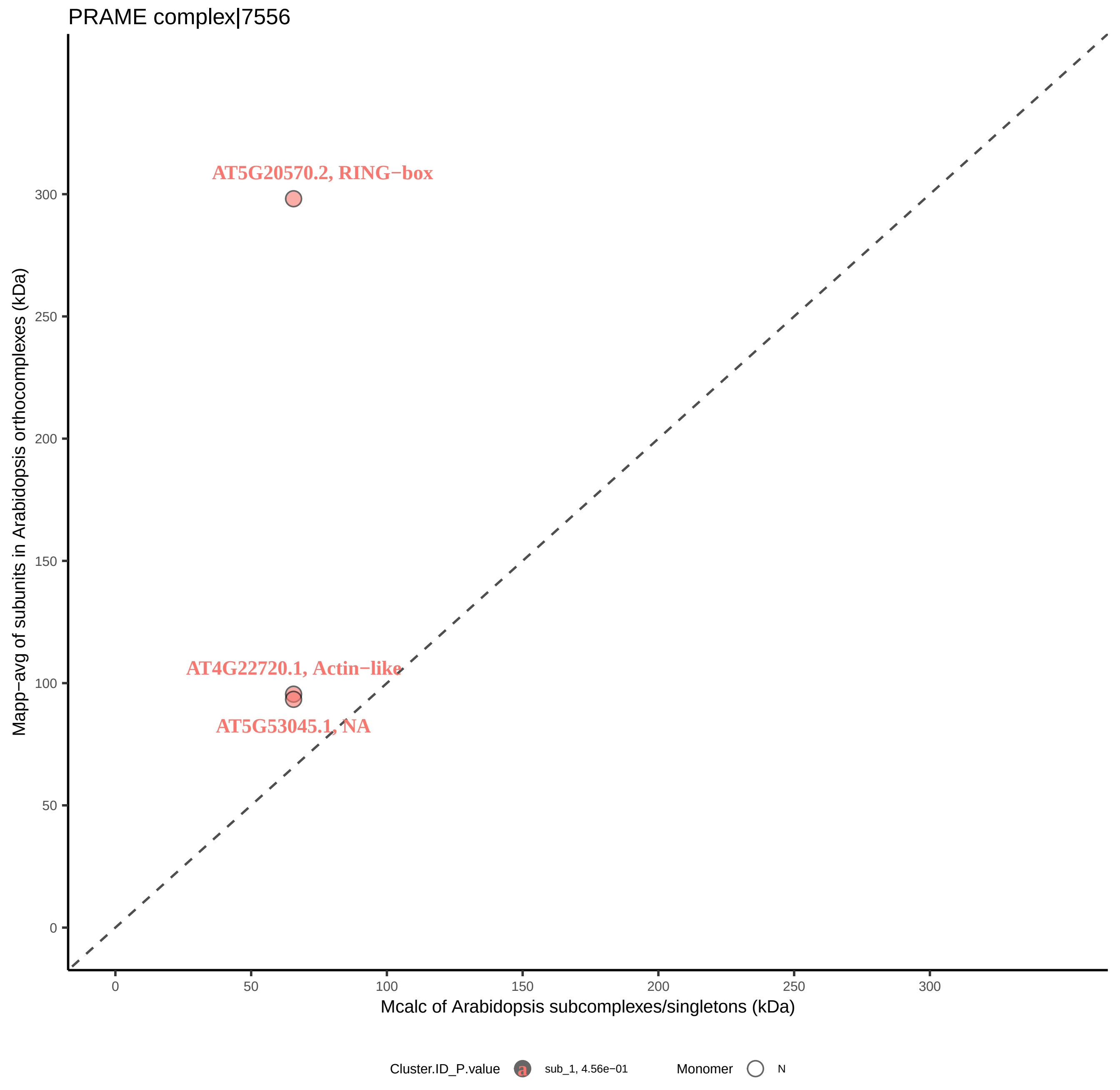

PRAME complex|7556
AT5G20570.2, RING−box
AT4G22720.1, Actin−like
AT5G53045.1, NA
Mapp−avg of subunits in Arabidopsis orthocomplexes (kDa)
Mcalc of Arabidopsis subcomplexes/singletons (kDa)
0
50
100
150
200
250
300
Cluster.ID_P.value
sub_1, 4.56e−01
Monomer
N

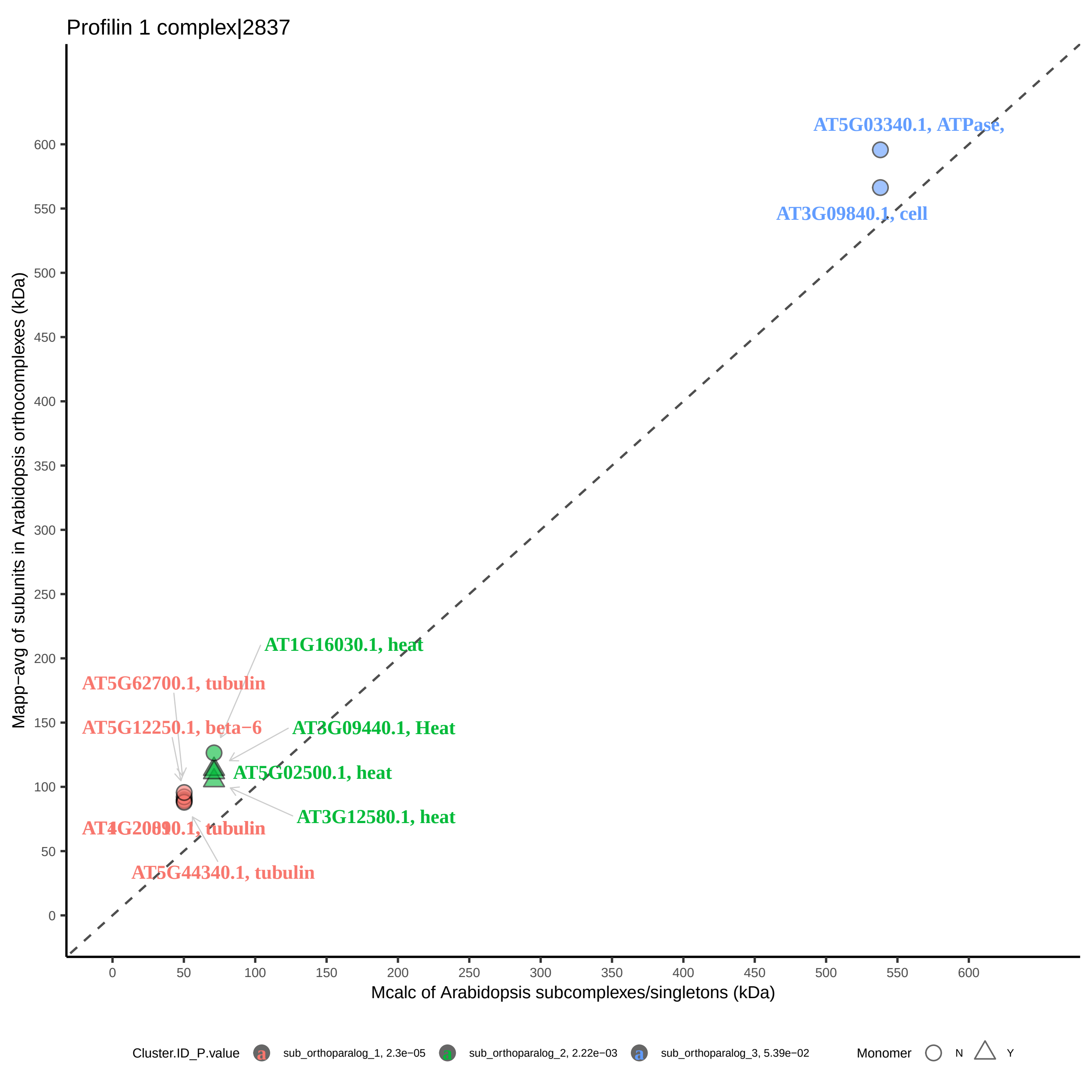
Profilin 1 complex|2837
AT5G03340.1, ATPase,
AT3G09840.1, cell
AT1G16030.1, heat
AT5G62700.1, tubulin
AT5G12250.1, beta−6
AT3G09440.1, Heat
AT5G02500.1, heat
AT3G12580.1, heat
AT5G44340.1, tubulin
Mapp−avg of subunits in Arabidopsis orthocomplexes (kDa)
Mcalc of Arabidopsis subcomplexes/singletons (kDa)
0
50
100
150
200
250
300
350
400
450
500
550
600
Cluster.ID_P.value
sub_orthoparalog_1, 2.3e−05
sub_orthoparalog_2, 2.22e−03
sub_orthoparalog_3, 5.39e−02
Monomer
N
Y

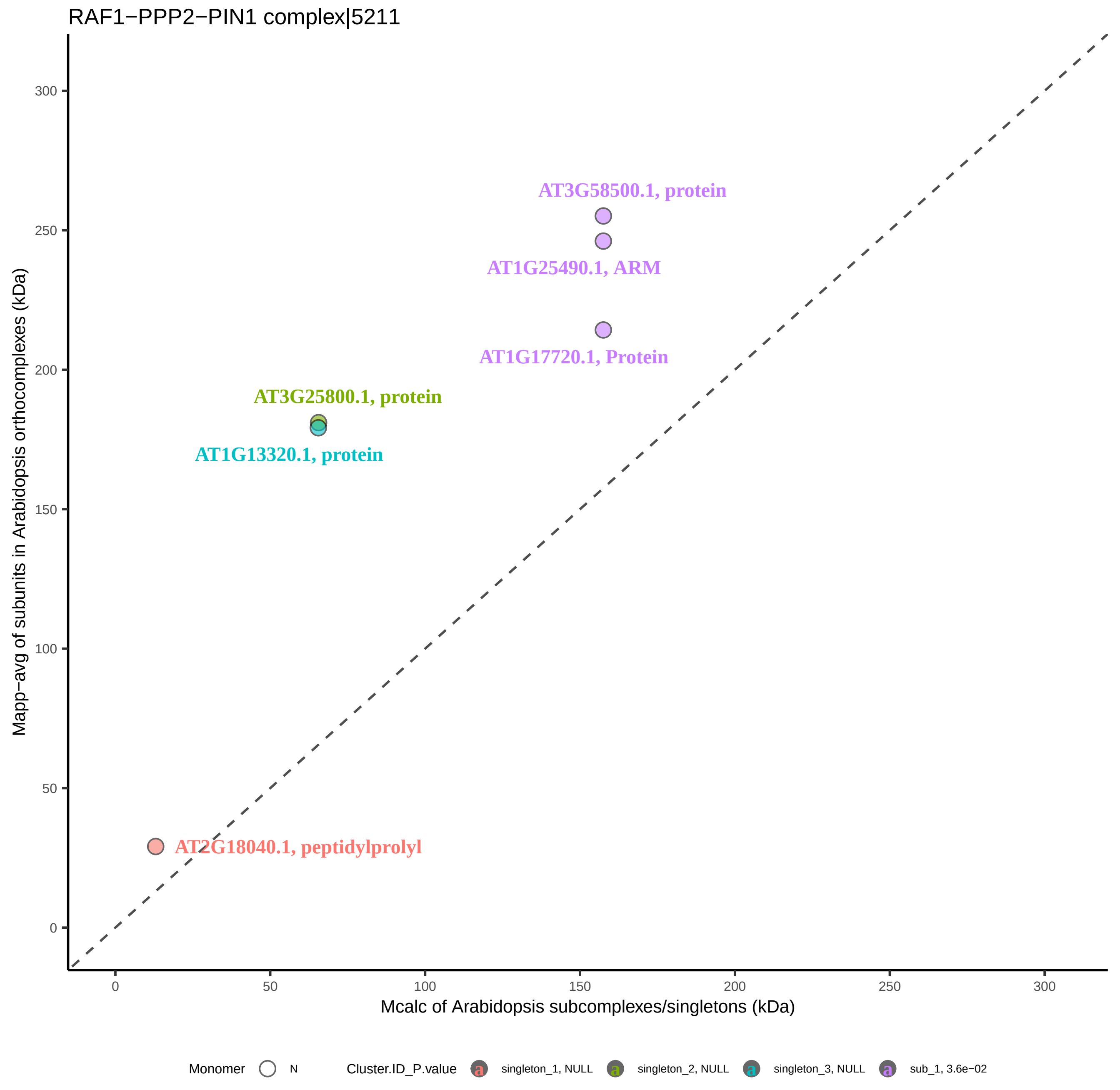
RAF1−PPP2−PIN1 complex|5211
AT3G58500.1, protein
AT1G25490.1, ARM
AT1G17720.1, Protein
AT3G25800.1, protein
AT1G13320.1, protein
AT2G18040.1, peptidylprolyl
Mapp−avg of subunits in Arabidopsis orthocomplexes (kDa)
Mcalc of Arabidopsis subcomplexes/singletons (kDa)
0
50
100
150
200
250
300
Monomer
N
Cluster.ID_P.value
singleton_1, NULL
singleton_2, NULL
singleton_3, NULL
sub_1, 3.6e−02

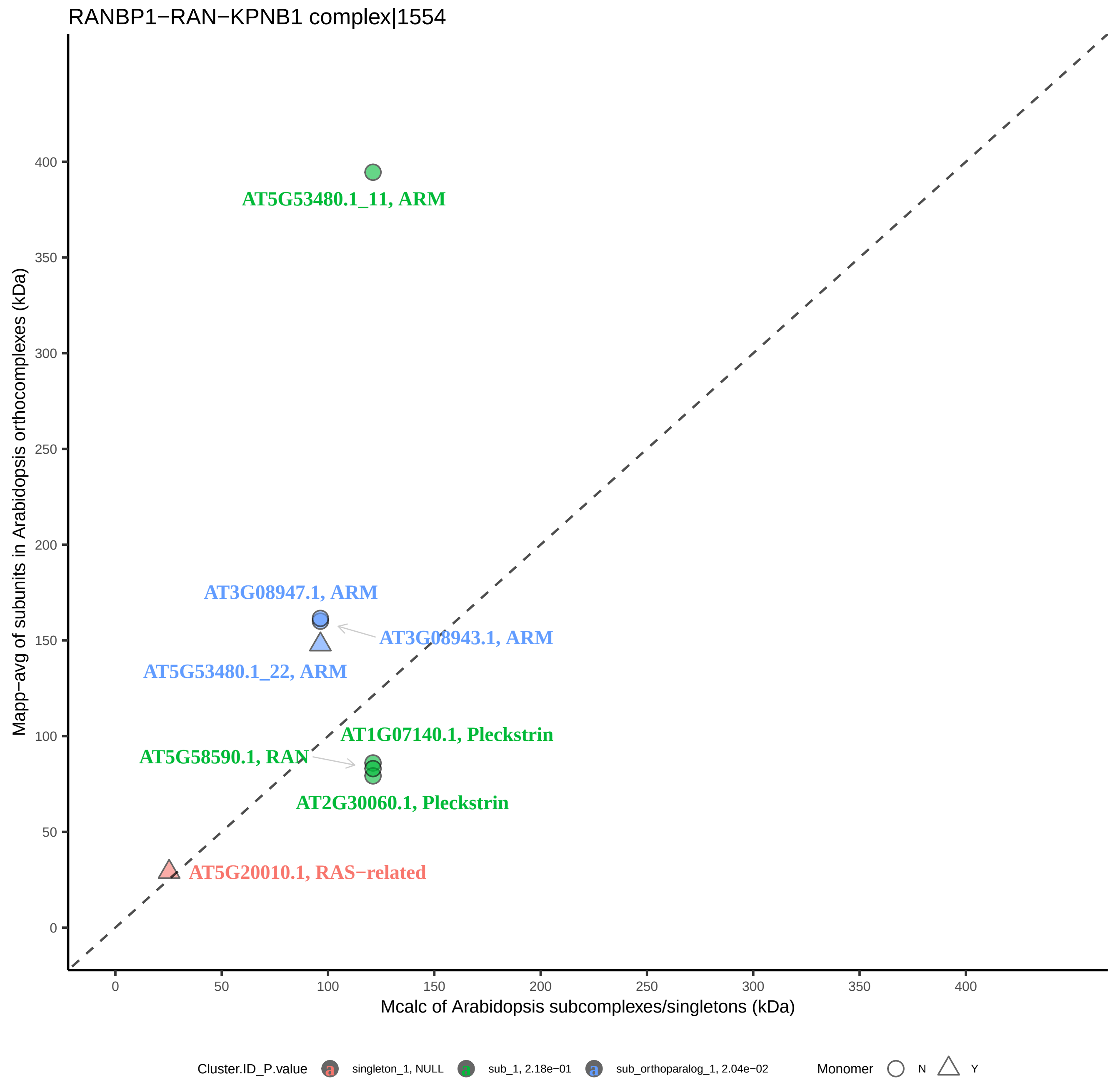

RANBP1−RAN−KPNB1 complex|1554
Mapp−avg of subunits in Arabidopsis orthocomplexes (kDa)
Mcalc of Arabidopsis subcomplexes/singletons (kDa)
0
50
100
150
200
250
300
350
400
AT5G53480.1_11, ARM
AT3G08947.1, ARM
AT3G08943.1, ARM
AT5G53480.1_22, ARM
AT1G07140.1, Pleckstrin
AT5G58590.1, RAN
AT2G30060.1, Pleckstrin
AT5G20010.1, RAS−related
Cluster.ID_P.value
singleton_1, NULL
sub_1, 2.18e−01
sub_orthoparalog_1, 2.04e−02
Monomer
N
Y

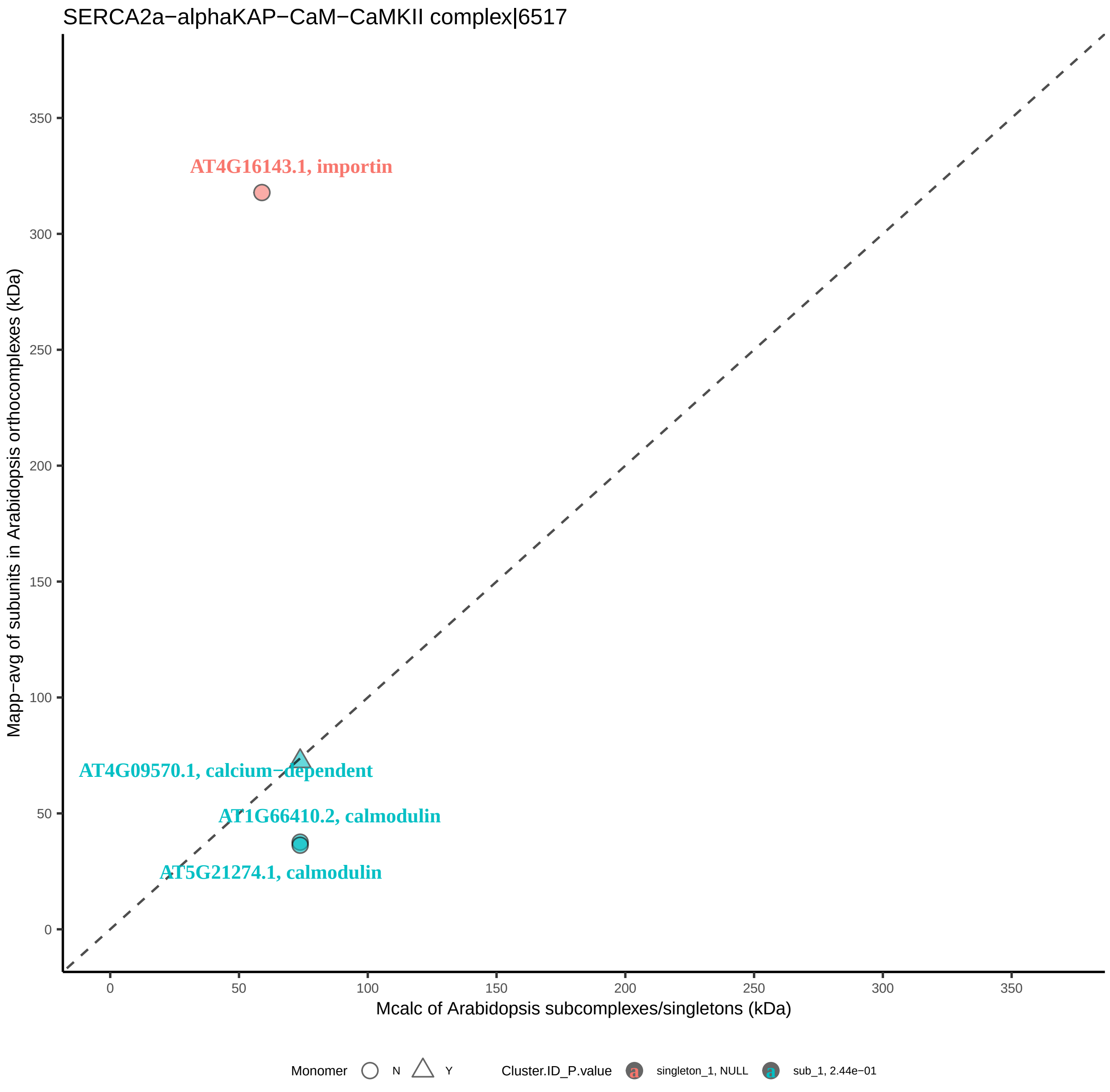
SERCA2a−alphaKAP−CaM−CaMKII complex|6517
Mapp−avg of subunits in Arabidopsis orthocomplexes (kDa)
Mcalc of Arabidopsis subcomplexes/singletons (kDa)
0
50
100
150
200
250
300
350
AT4G16143.1, importin
AT4G09570.1, calcium−dependent
AT1G66410.2, calmodulin
AT5G21274.1, calmodulin
Monomer
N
Y
Cluster.ID_P.value
singleton_1, NULL
sub_1, 2.44e−01

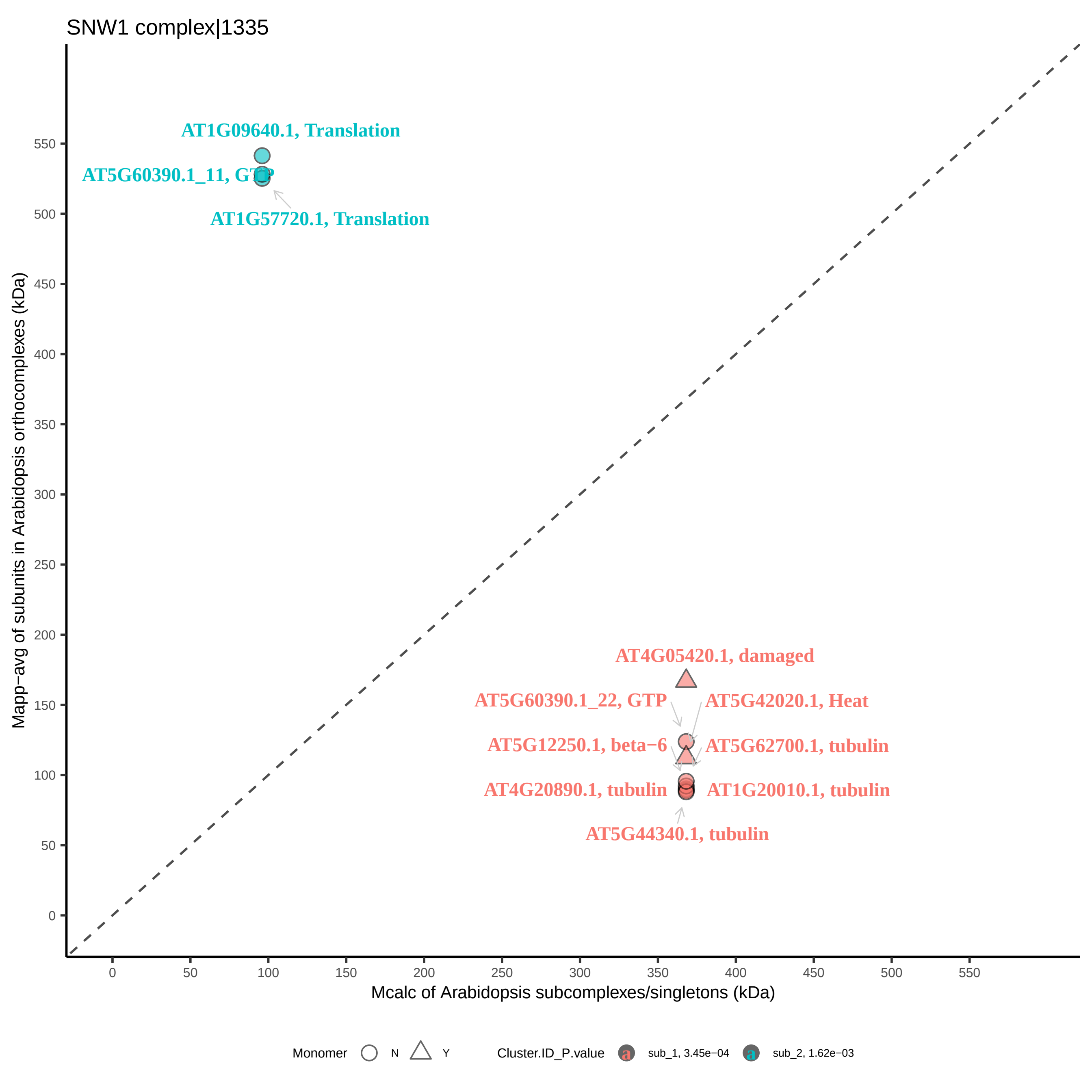

SNW1 complex|1335
AT1G09640.1, Translation
AT5G60390.1_11, GTP
AT1G57720.1, Translation
AT4G05420.1, damaged
AT5G60390.1_22, GTP
AT5G42020.1, Heat
AT5G12250.1, beta−6
AT5G62700.1, tubulin
AT4G20890.1, tubulin
AT1G20010.1, tubulin
AT5G44340.1, tubulin
Mapp−avg of subunits in Arabidopsis orthocomplexes (kDa)
Mcalc of Arabidopsis subcomplexes/singletons (kDa)
0
50
100
150
200
250
300
350
400
450
500
550
Monomer
N
Y
Cluster.ID_P.value
sub_1, 3.45e−04
sub_2, 1.62e−03

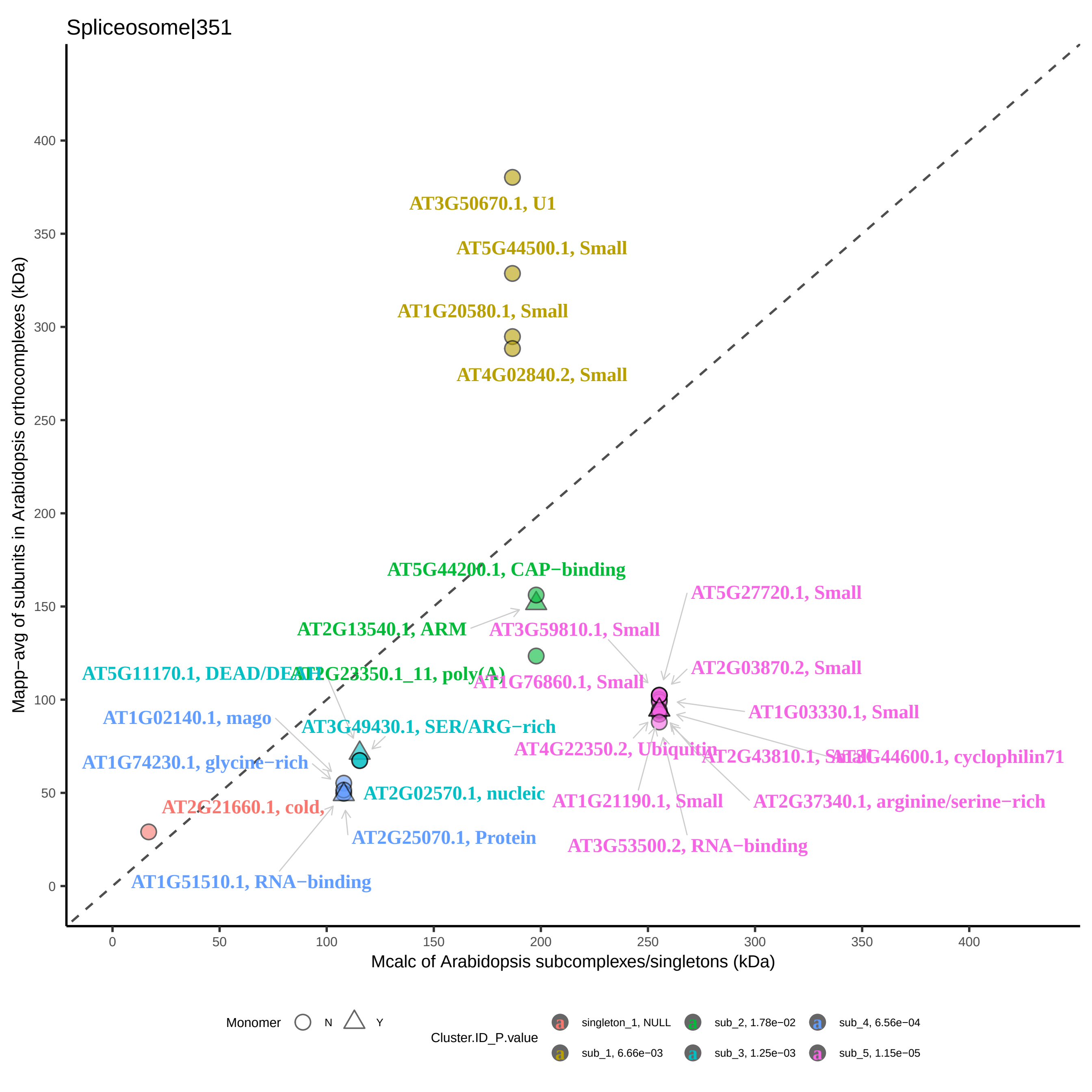
Spliceosome|351
Mapp−avg of subunits in Arabidopsis orthocomplexes (kDa)
Mcalc of Arabidopsis subcomplexes/singletons (kDa)
0
50
100
150
200
250
300
350
400
AT3G50670.1, U1
AT5G44500.1, Small
AT1G20580.1, Small
AT4G02840.2, Small
AT5G44200.1, CAP−binding
AT2G13540.1, ARM
AT3G59810.1, Small
AT5G27720.1, Small
AT2G03870.2, Small
AT1G76860.1, Small
AT1G03330.1, Small
AT5G11170.1, DEAD/DEAH
AT2G23350.1_11, poly(A)
AT1G02140.1, mago
AT3G49430.1, SER/ARG−rich
AT4G22350.2, Ubiquitin
AT2G43810.1, Small
AT3G44600.1, cyclophilin71
AT1G74230.1, glycine−rich
AT2G02570.1, nucleic
AT1G21190.1, Small
AT2G37340.1, arginine/serine−rich
AT2G21660.1, cold,
AT2G25070.1, Protein
AT3G53500.2, RNA−binding
AT1G51510.1, RNA−binding
Monomer
N
Y
Cluster.ID_P.value
singleton_1, NULL
sub_1, 6.66e−03
sub_2, 1.78e−02
sub_3, 1.25e−03
sub_4, 6.56e−04
sub_5, 1.15e−05

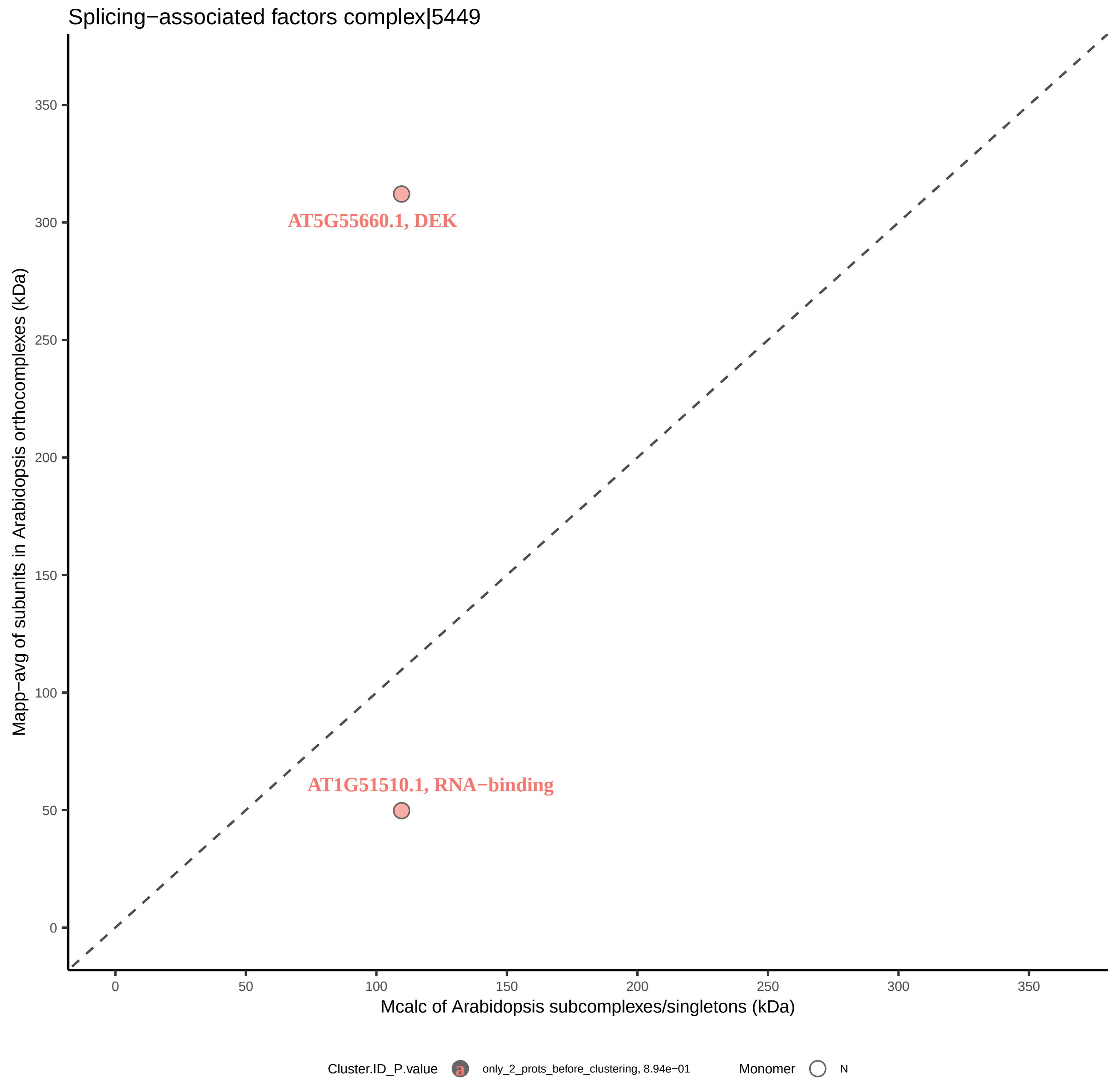

Splicing−associated factors complex|5449
AT5G55660.1, DEK
AT1G51510.1, RNA−binding
Mapp−avg of subunits in Arabidopsis orthocomplexes (kDa)
Mcalc of Arabidopsis subcomplexes/singletons (kDa)
0
50
100
150
200
250
300
350
Cluster.ID_P.value
only_2_prots_before_clustering, 8.94e−01
Monomer
N

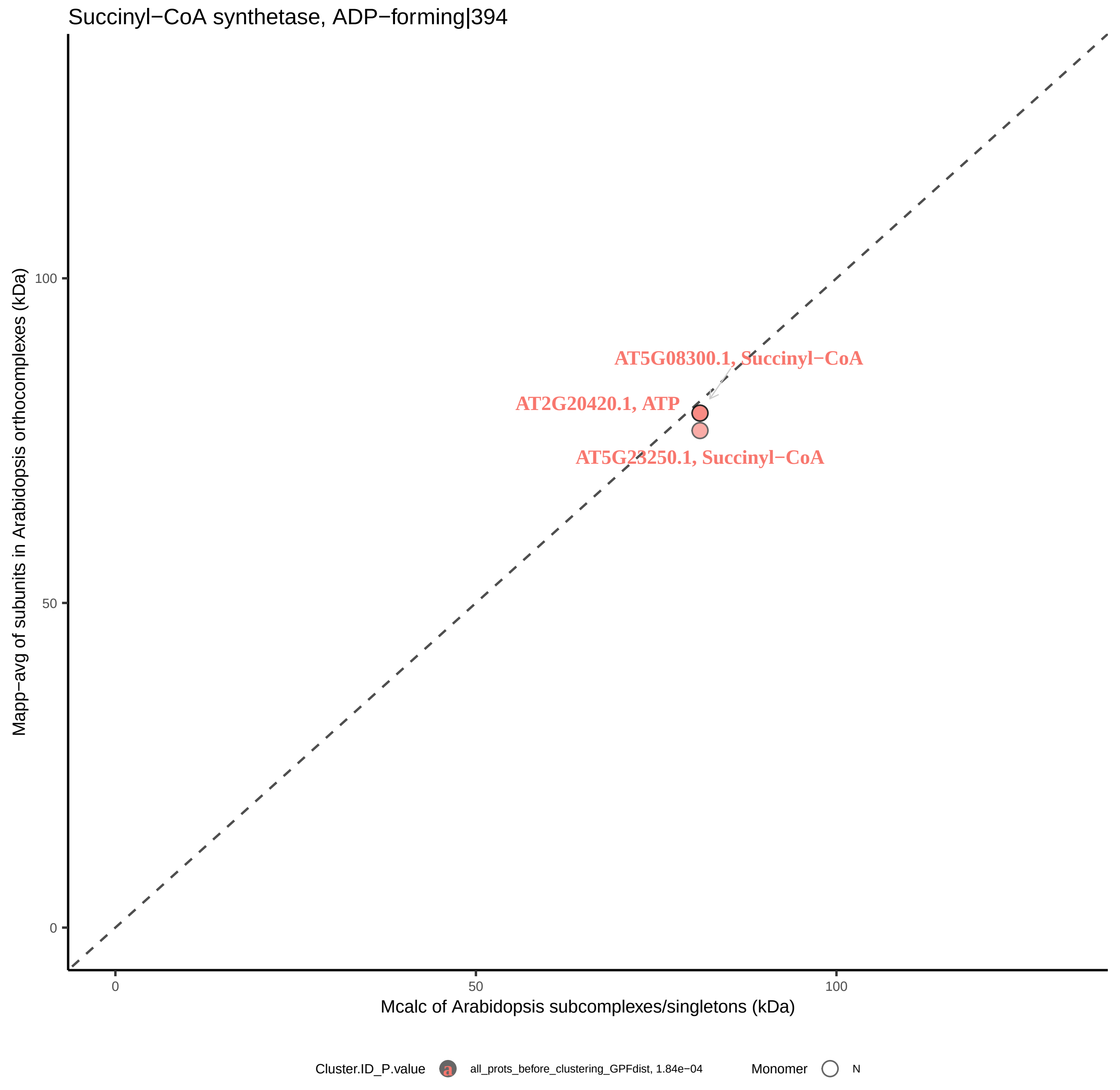

Succinyl−CoA synthetase, ADP−forming|394
Mapp−avg of subunits in Arabidopsis orthocomplexes (kDa)
100
50
0
AT5G08300.1, Succinyl−CoA
AT2G20420.1, ATP
AT5G23250.1, Succinyl−CoA
0
50
100
Mcalc of Arabidopsis subcomplexes/singletons (kDa)
Cluster.ID_P.value
a
all_prots_before_clustering_GPFdist, 1.84e−04
Monomer
N

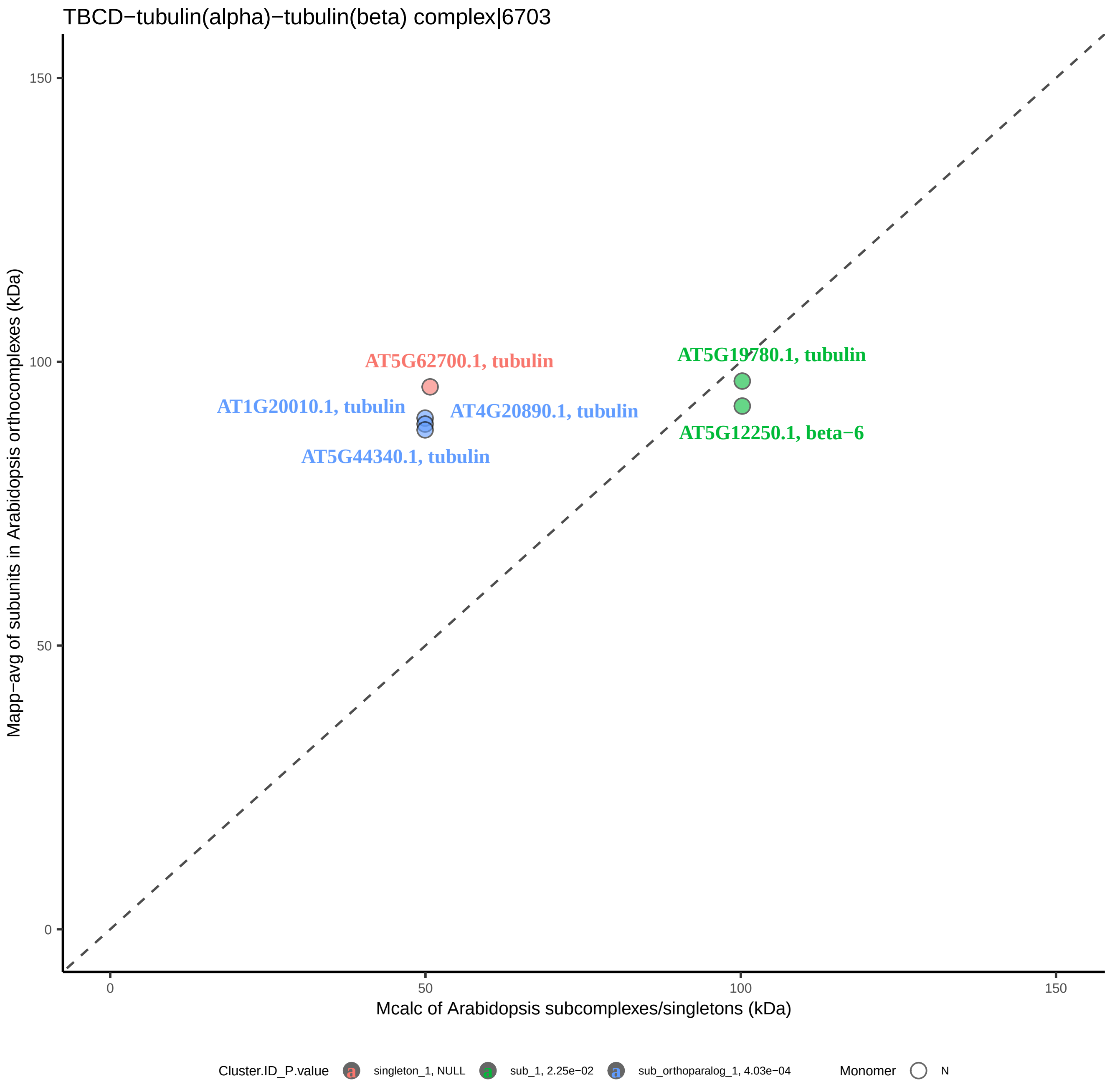
TBCD−tubulin(alpha)−tubulin(beta) complex|6703
Mapp−avg of subunits in Arabidopsis orthocomplexes (kDa)
Mcalc of Arabidopsis subcomplexes/singletons (kDa)
0
50
100
150
AT5G62700.1, tubulin
AT1G20010.1, tubulin
AT4G20890.1, tubulin
AT5G44340.1, tubulin
AT5G19780.1, tubulin
AT5G12250.1, beta−6
Cluster.ID_P.value
singleton_1, NULL
sub_1, 2.25e−02
sub_orthoparalog_1, 4.03e−04
Monomer
N

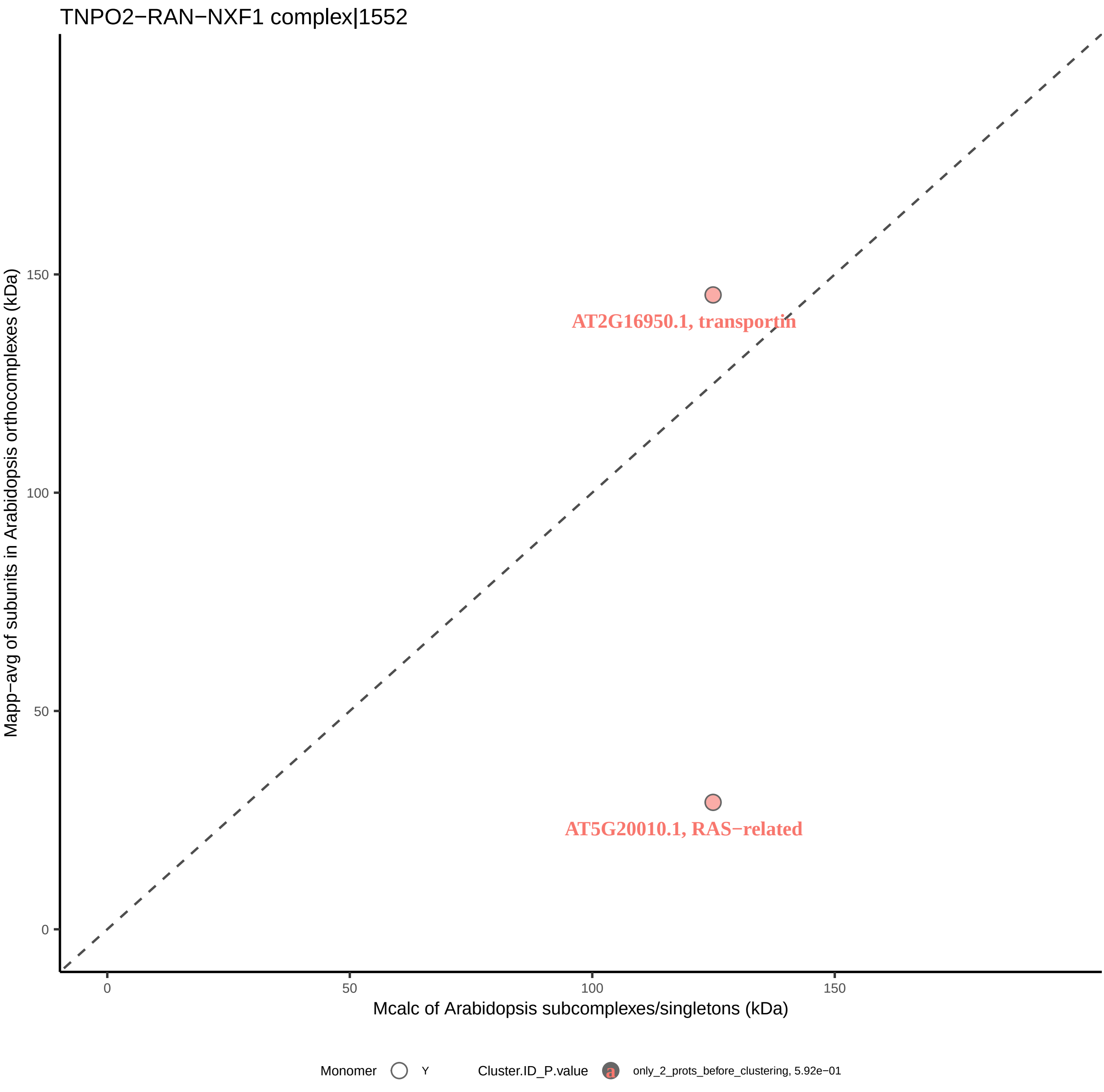
TNPO2−RAN−NXF1 complex|1552
Mapp−avg of subunits in Arabidopsis orthocomplexes (kDa)
150
100
50
0
AT2G16950.1, transportin
AT5G20010.1, RAS−related
0
50
100
150
Mcalc of Arabidopsis subcomplexes/singletons (kDa)
Monomer
Y
Cluster.ID_P.value
a
only_2_prots_before_clustering, 5.92e−01
