## Supplemental Figure S7 Arabidopsis protein SEC profiles illustrating small-world analysis results for "A machine learning-based approach to identify reliable gold standards for protein complex composition prediction"

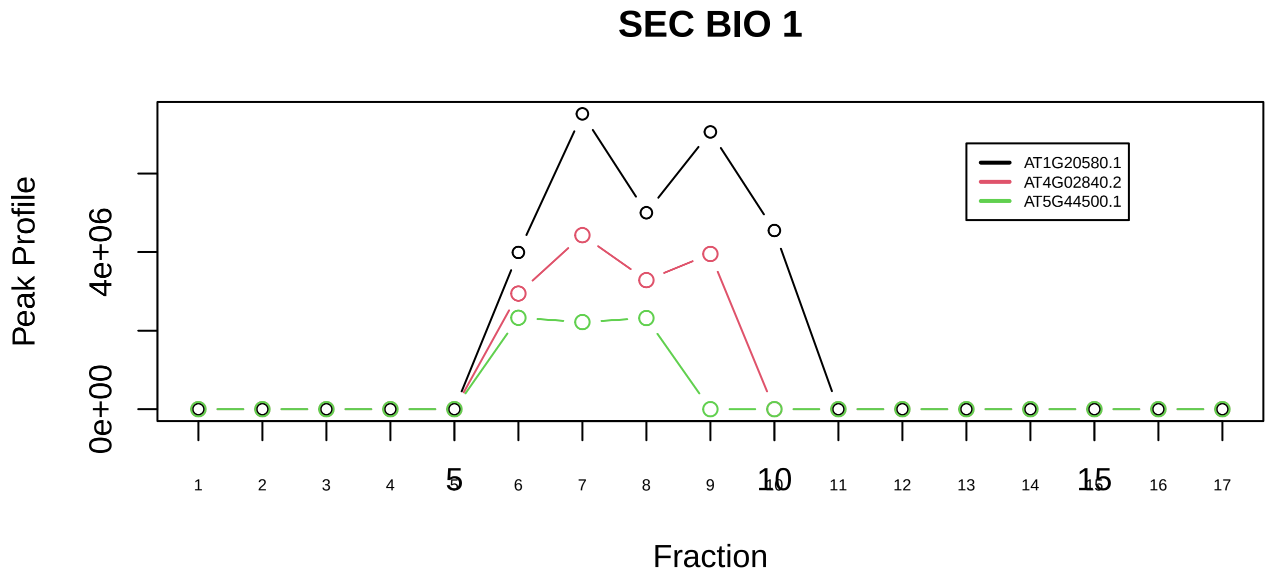

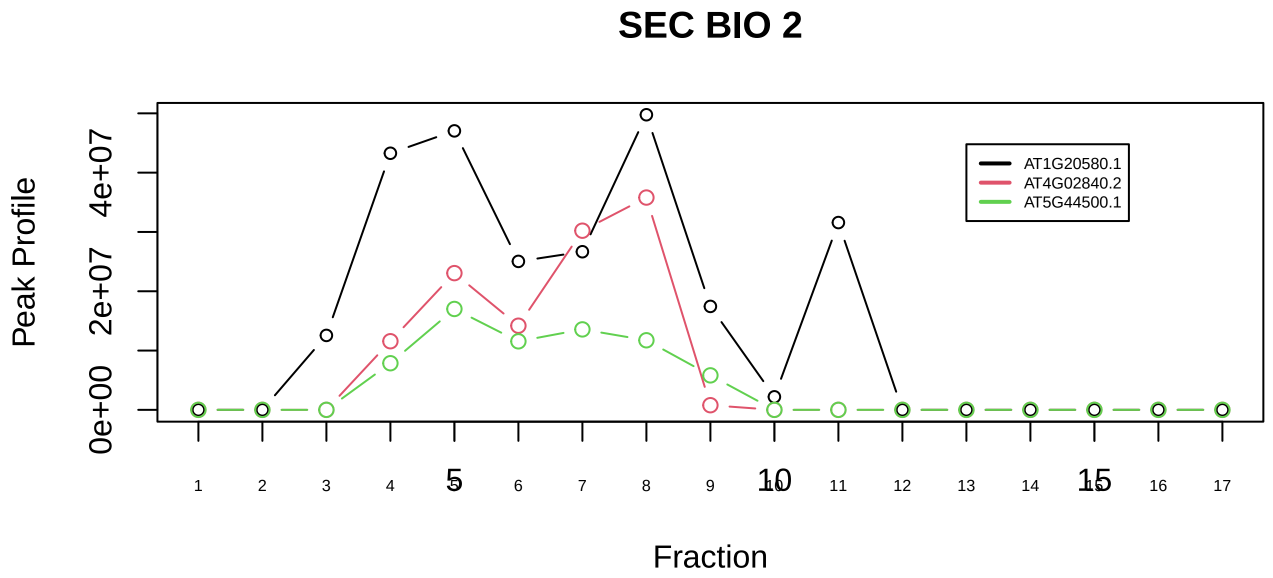

17S U2 snRNP|2755−−sub_1

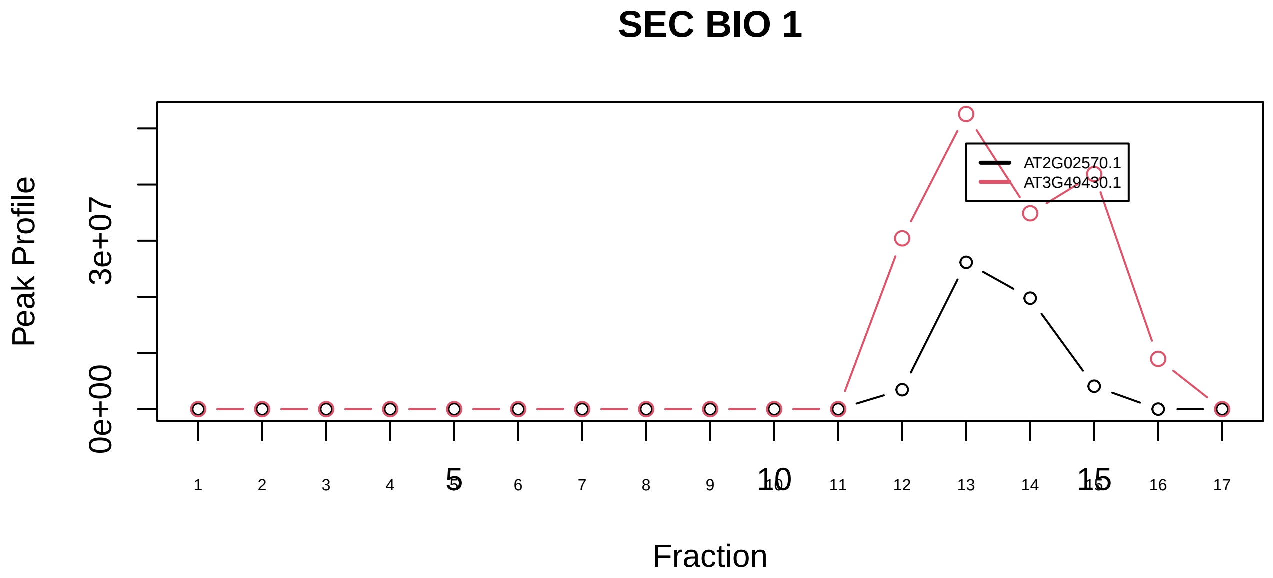

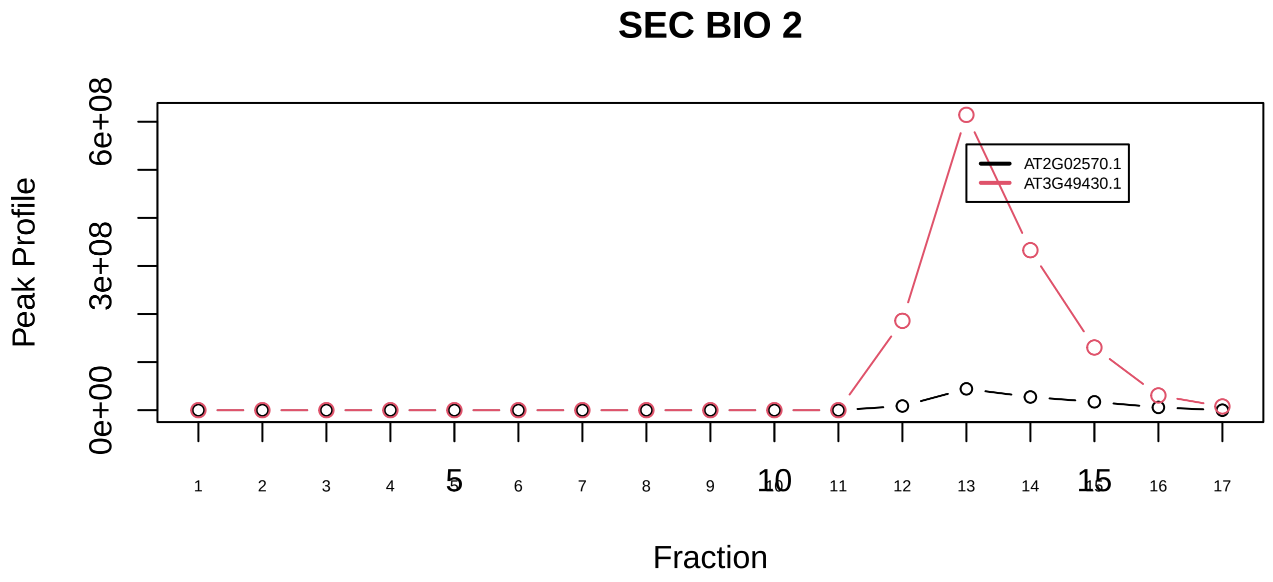

17S U2 snRNP|2755−−sub_2

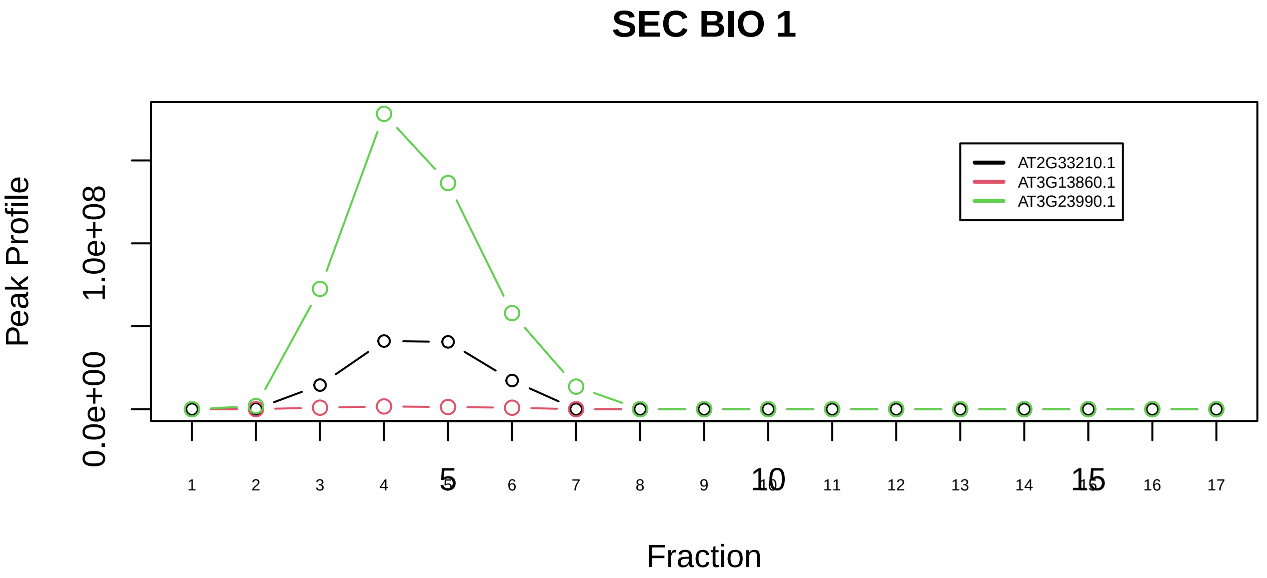

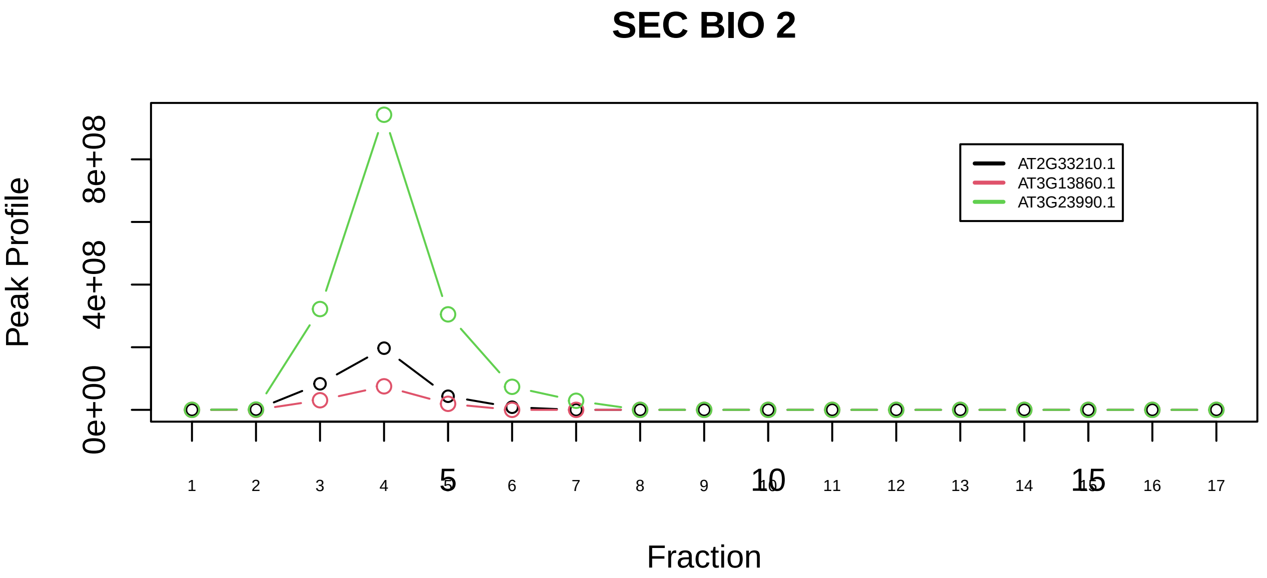

17S U2 snRNP|2755−−sub_orthoparalog_1

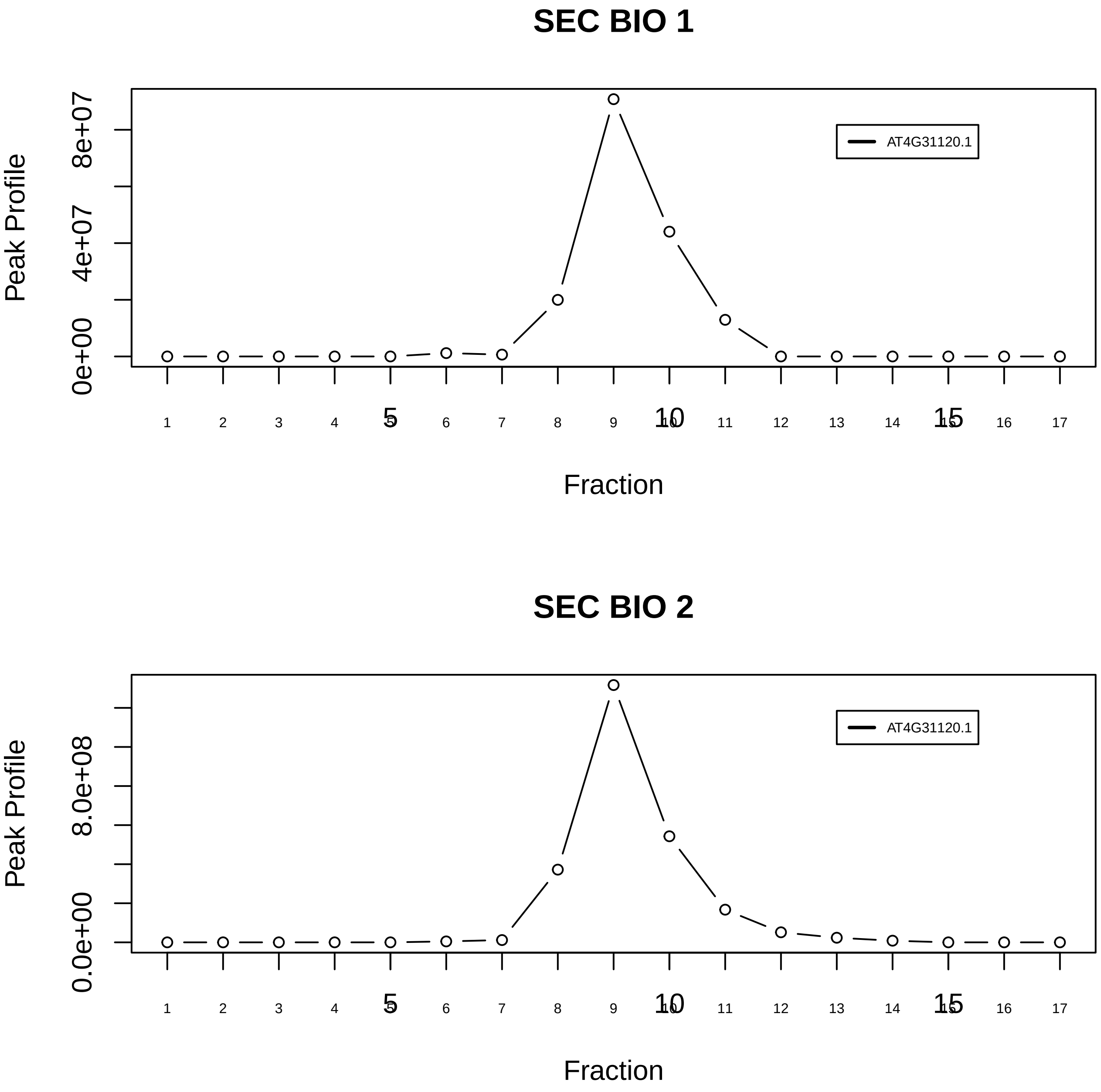

20S methylosome and RG−containing Sm protein complex|834−−singleton_1

### SEC BIO 1

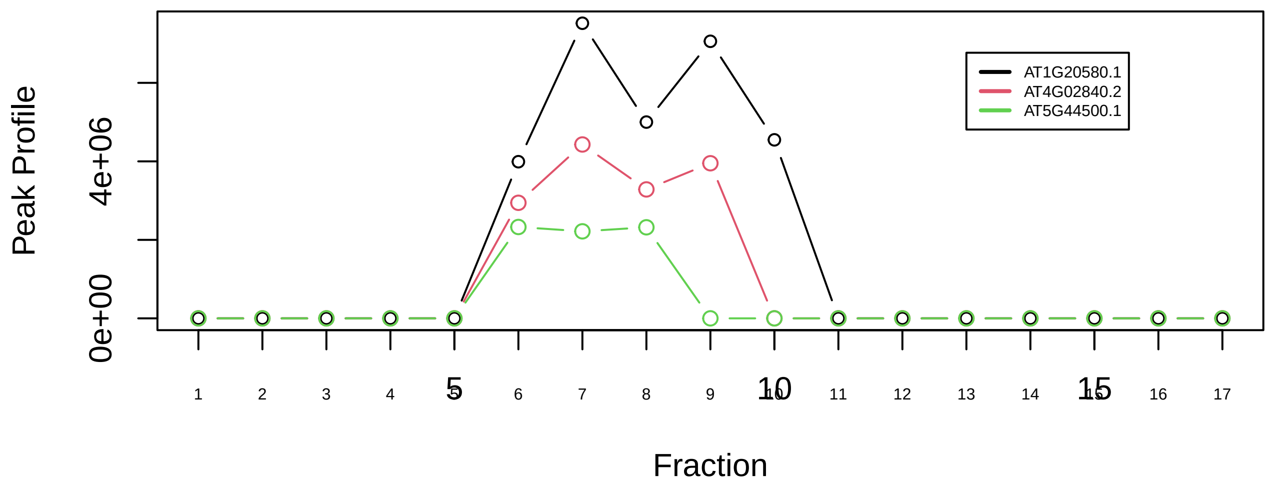

### SEC BIO 2

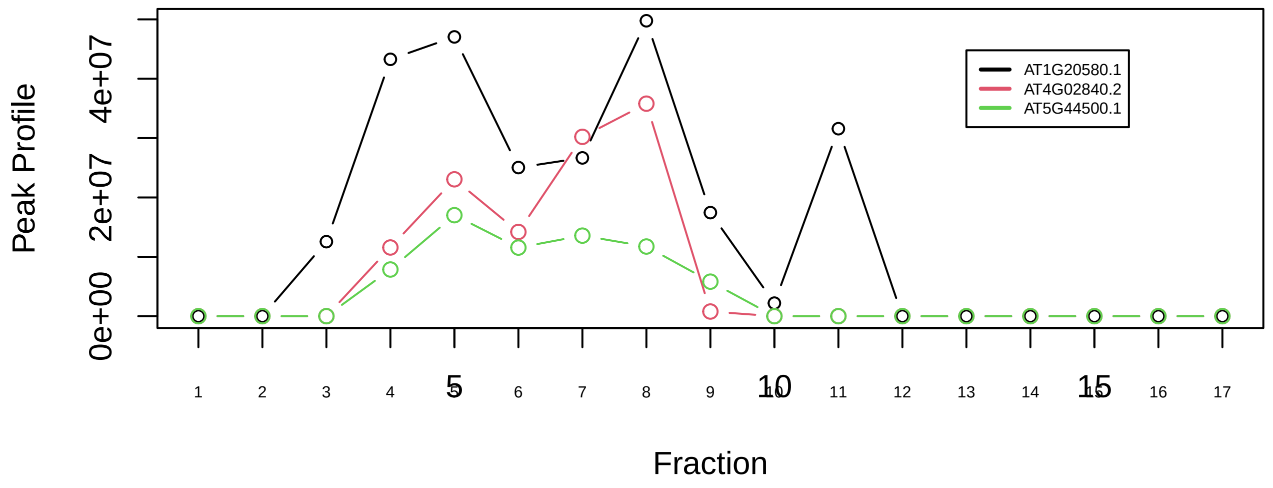

20S methylosome and RG−containing Sm protein complex|834−−sub_1

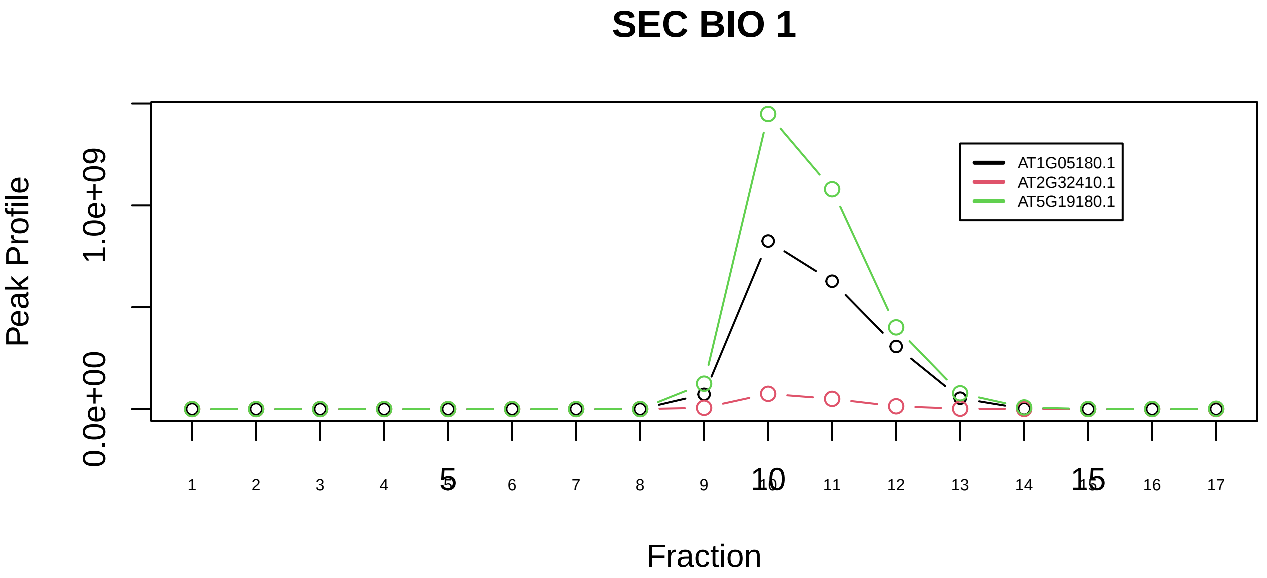

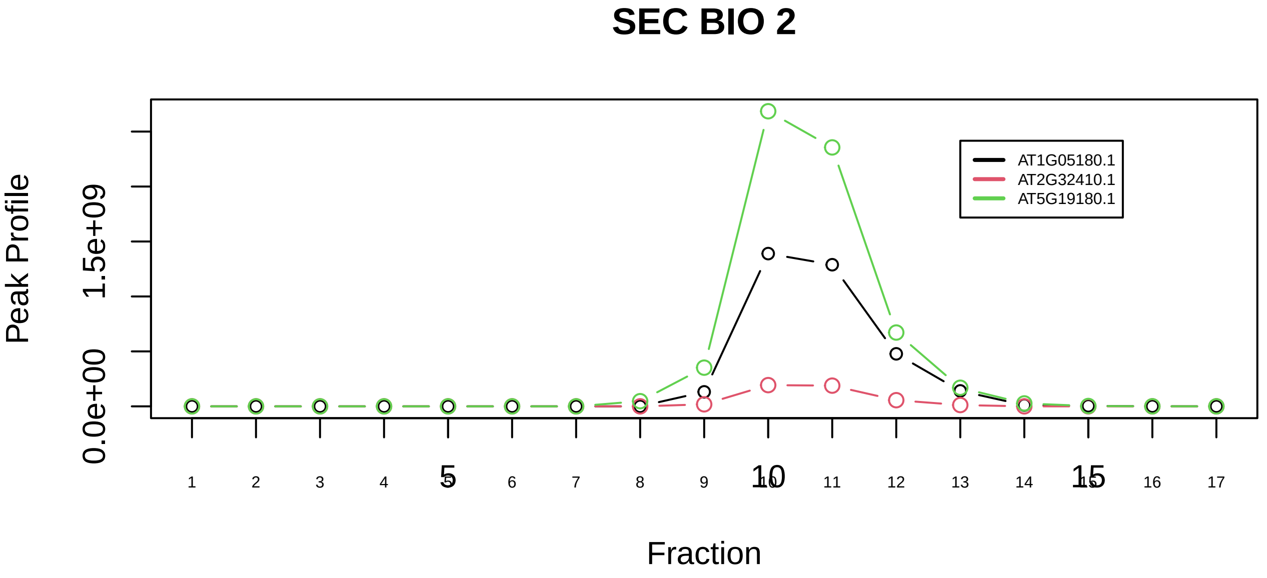

APPBP1−UBA3 complex|2162−−all_prots_before_clustering_GPFdist

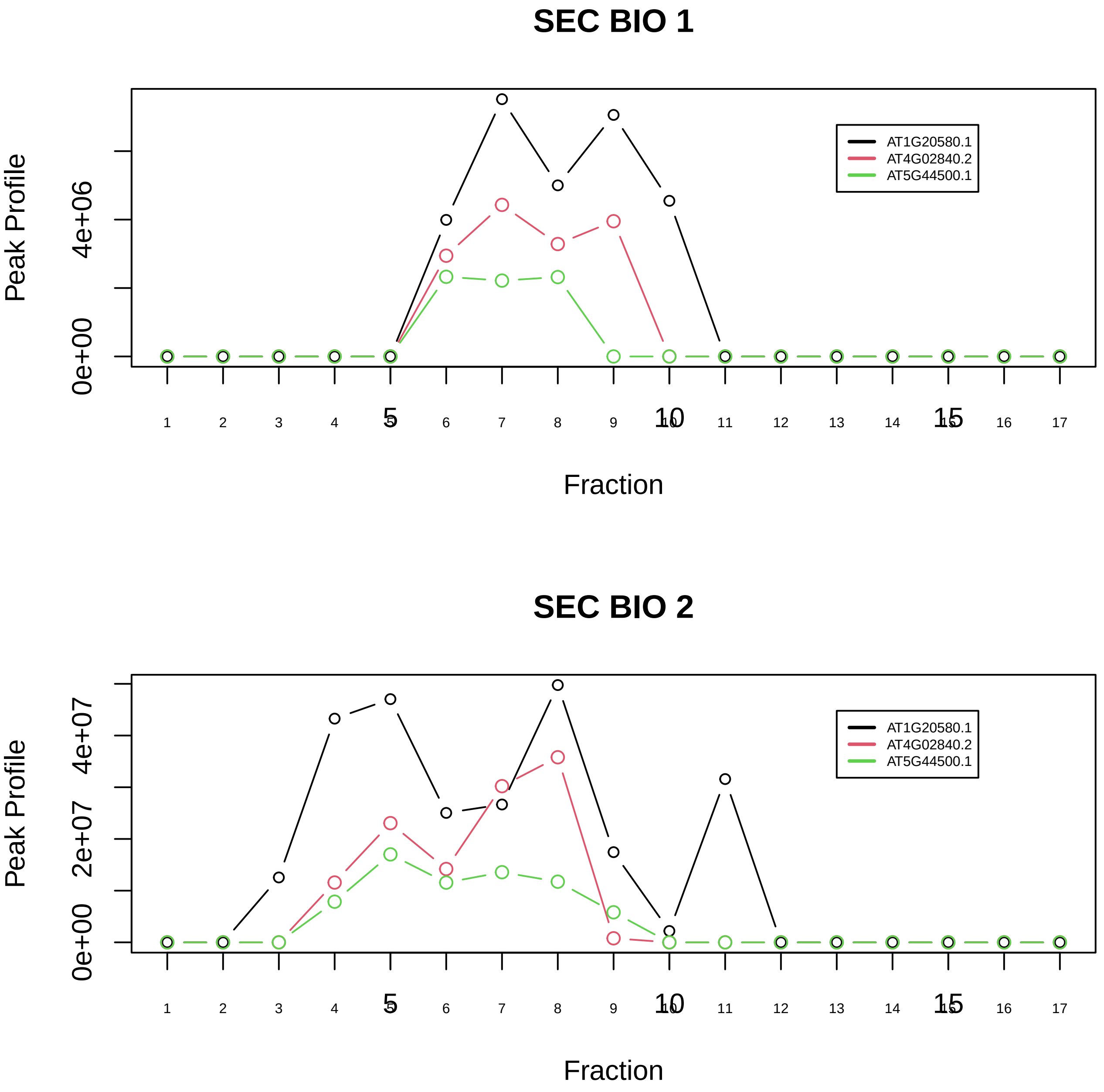

C complex spliceosome|1181−−sub_1

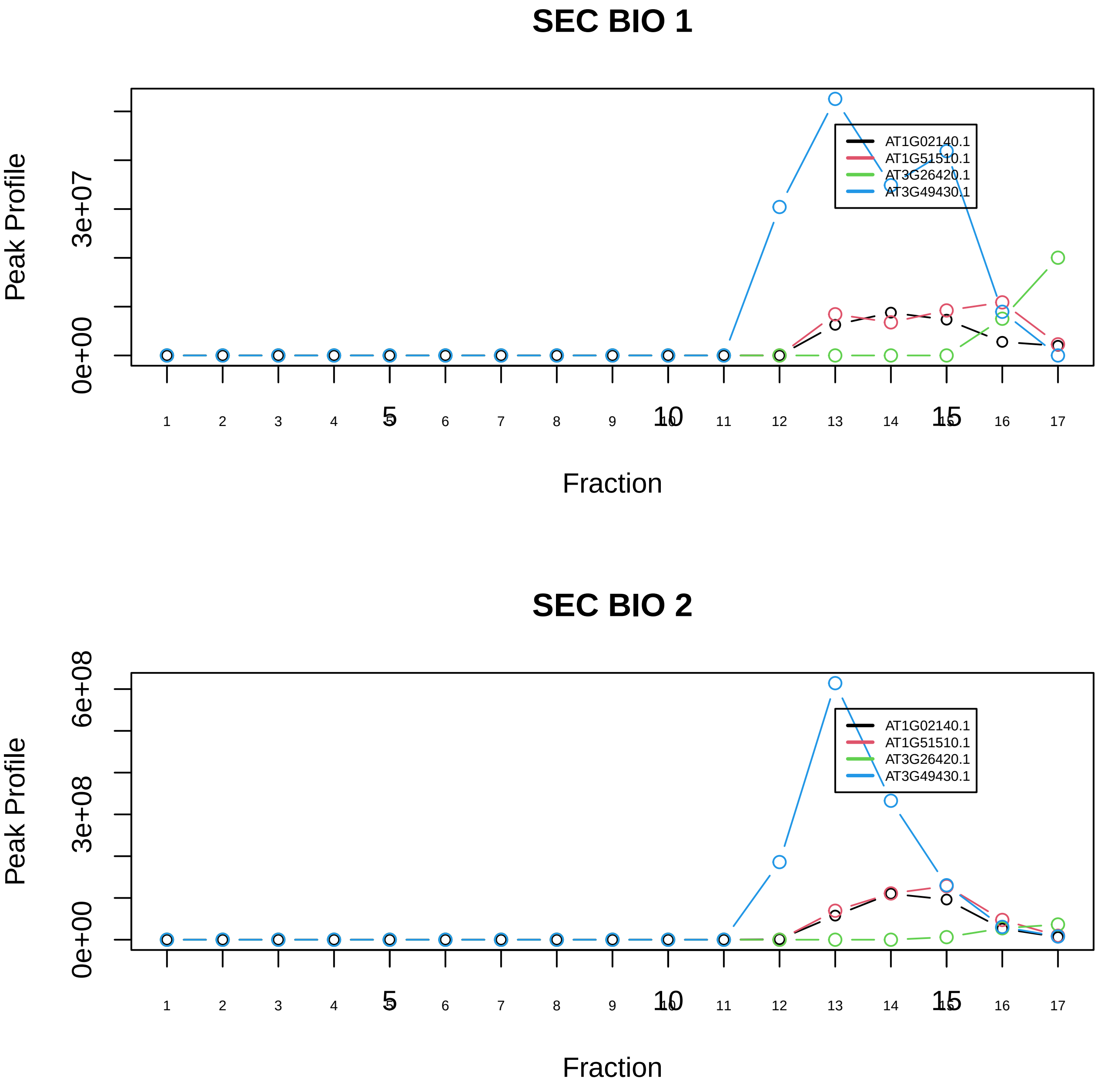

C complex spliceosome|1181−−sub_2

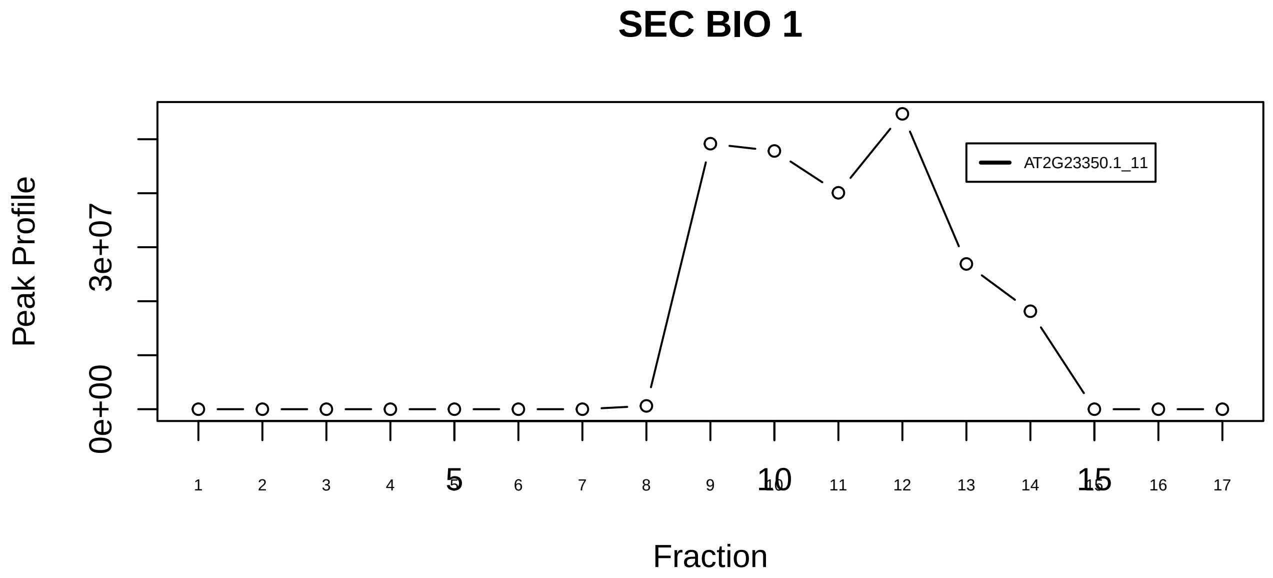

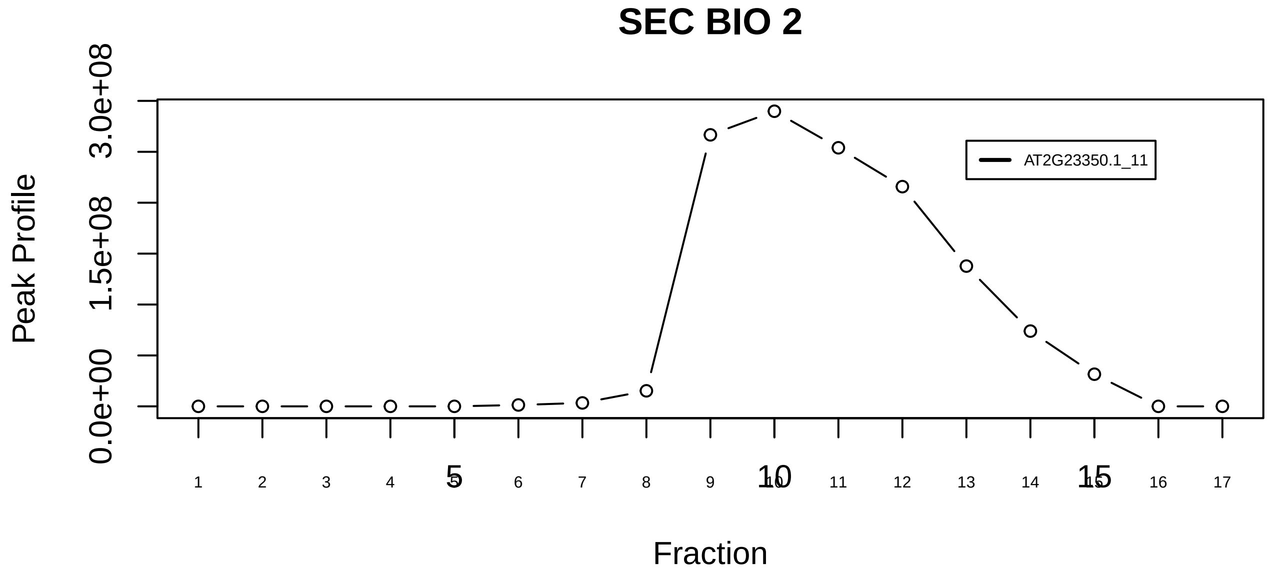

C complex spliceosome|1181−−singleton_1

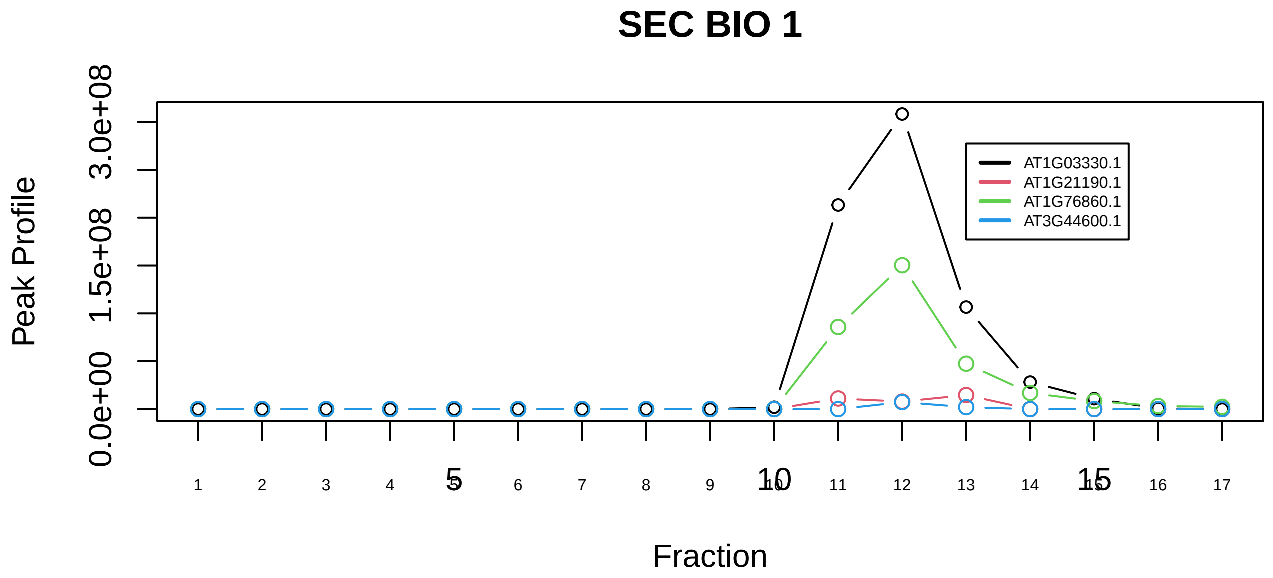

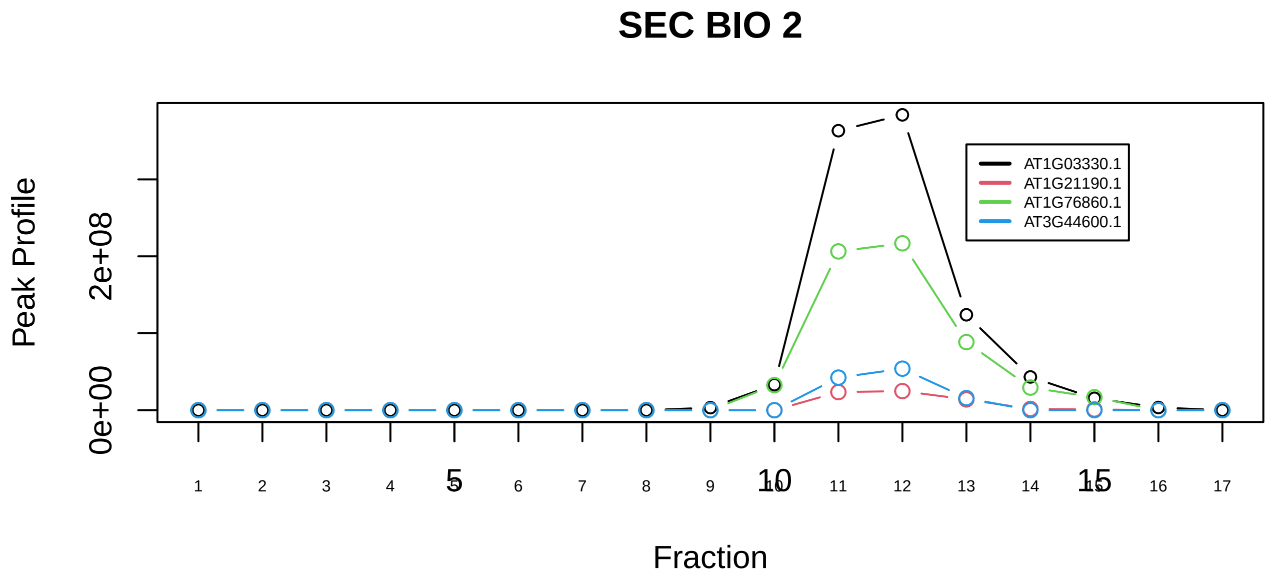

C complex spliceosome|1181−−sub_3

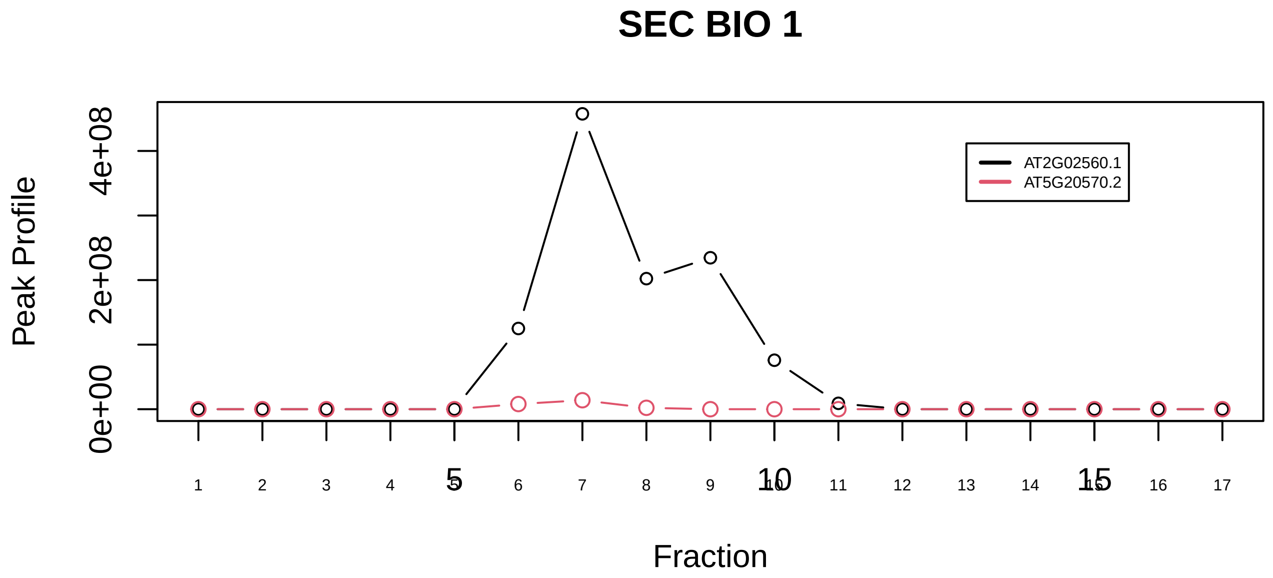

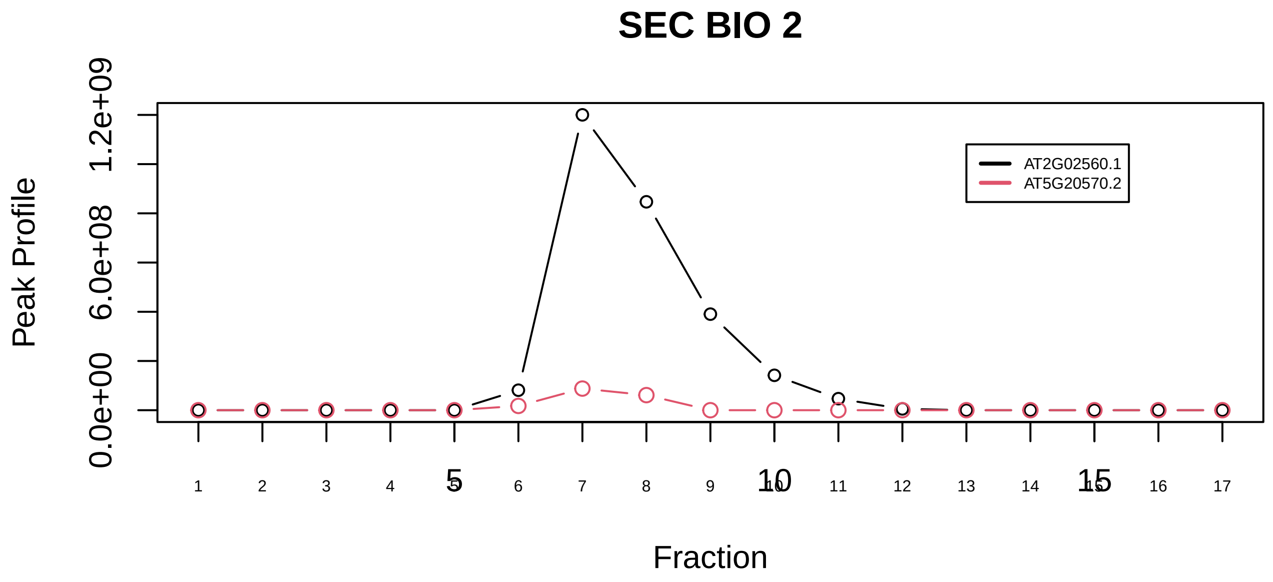

CAND1−CUL4B−RBX1 complex|224−−only_2_prots_before_clustering

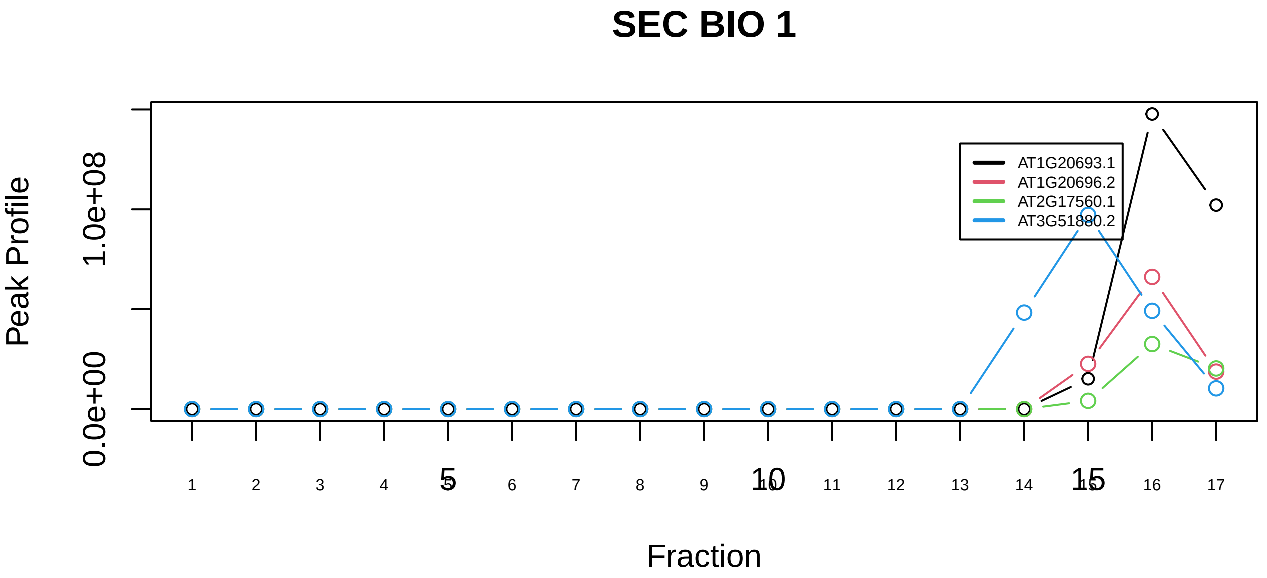

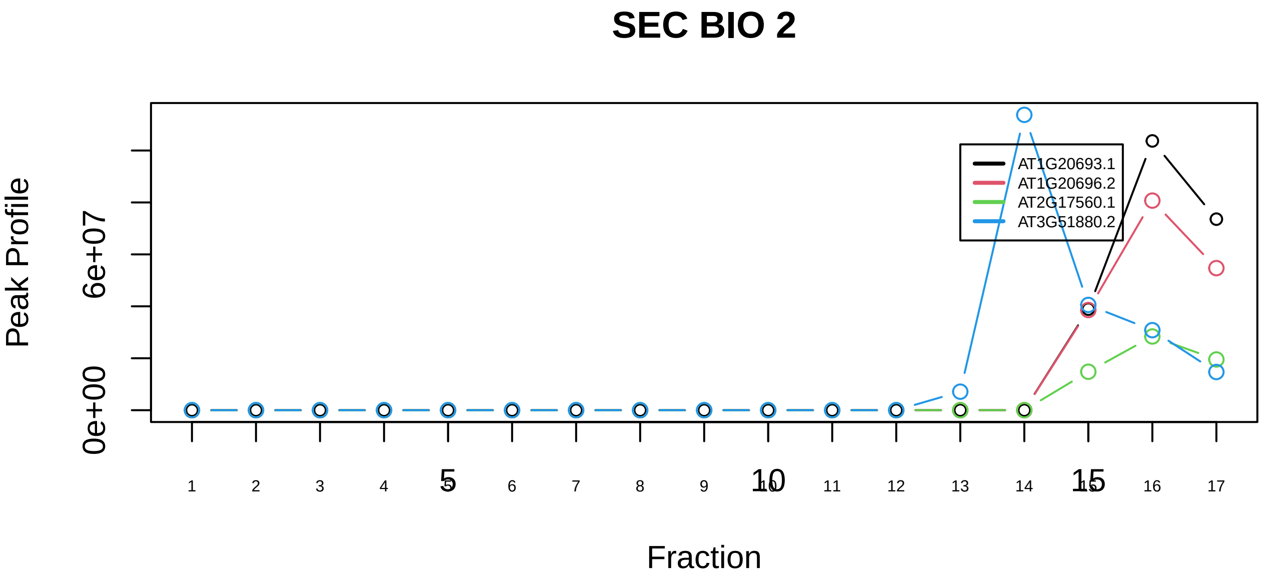

Casein kinase II−HMG1 complex|2719−−sub_orthoparalog_1

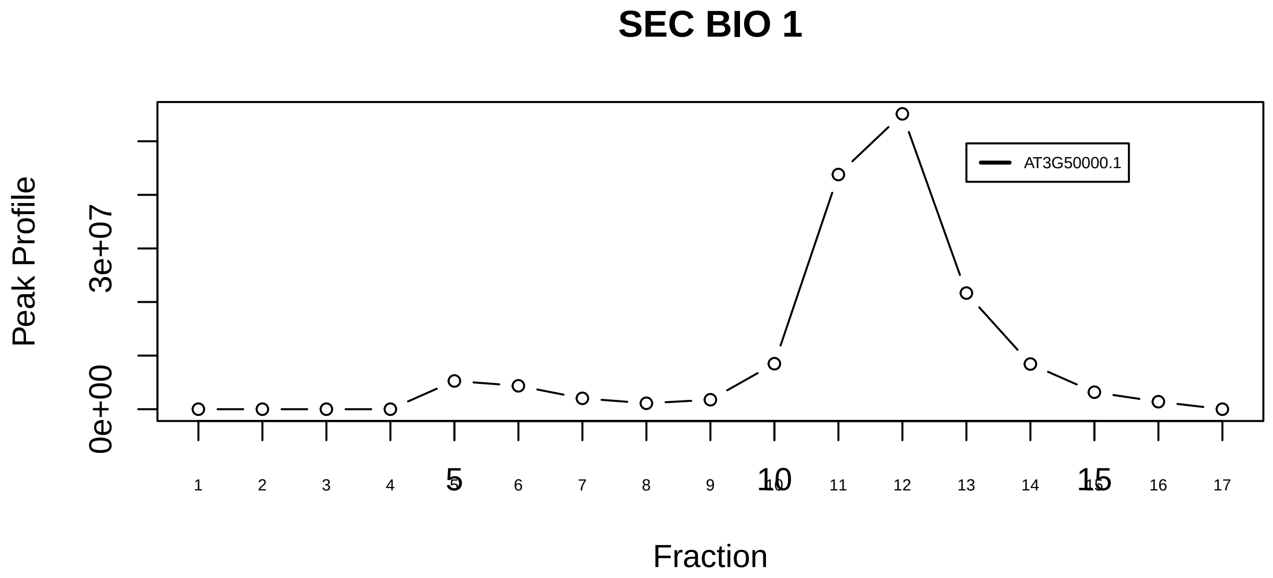

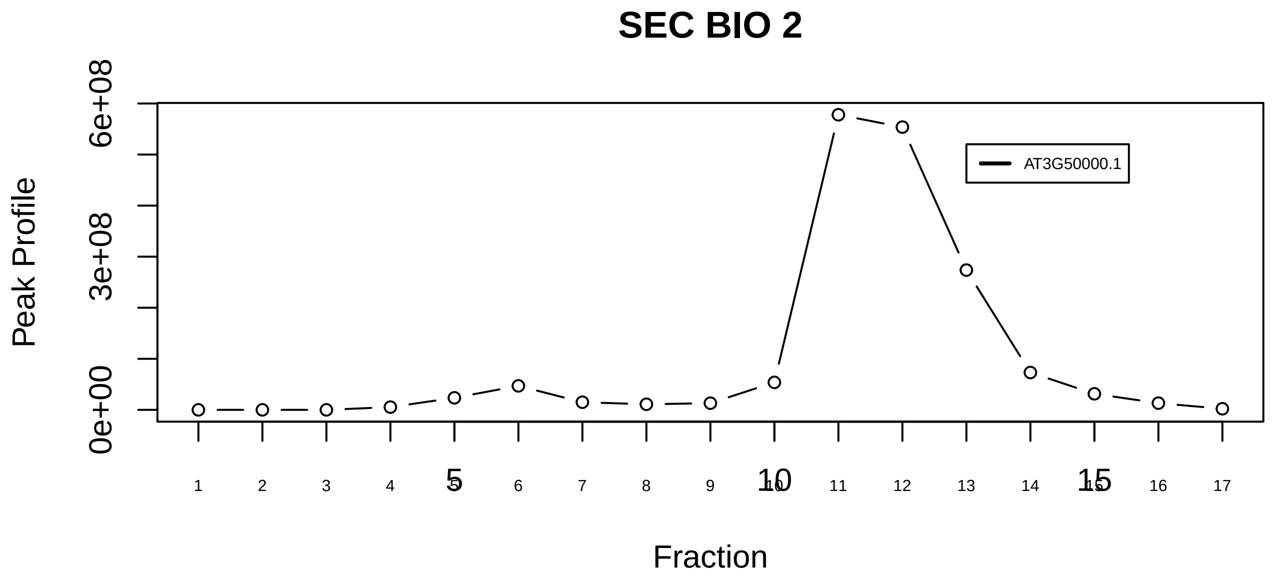

Casein kinase II−HMG1 complex|2719−−singleton_1

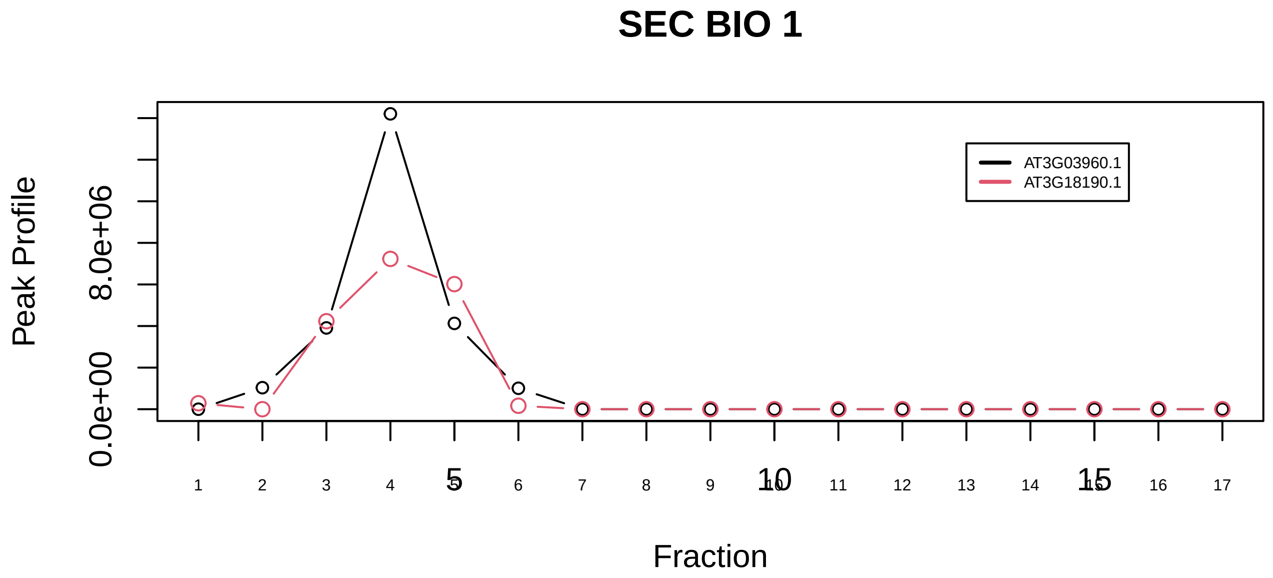

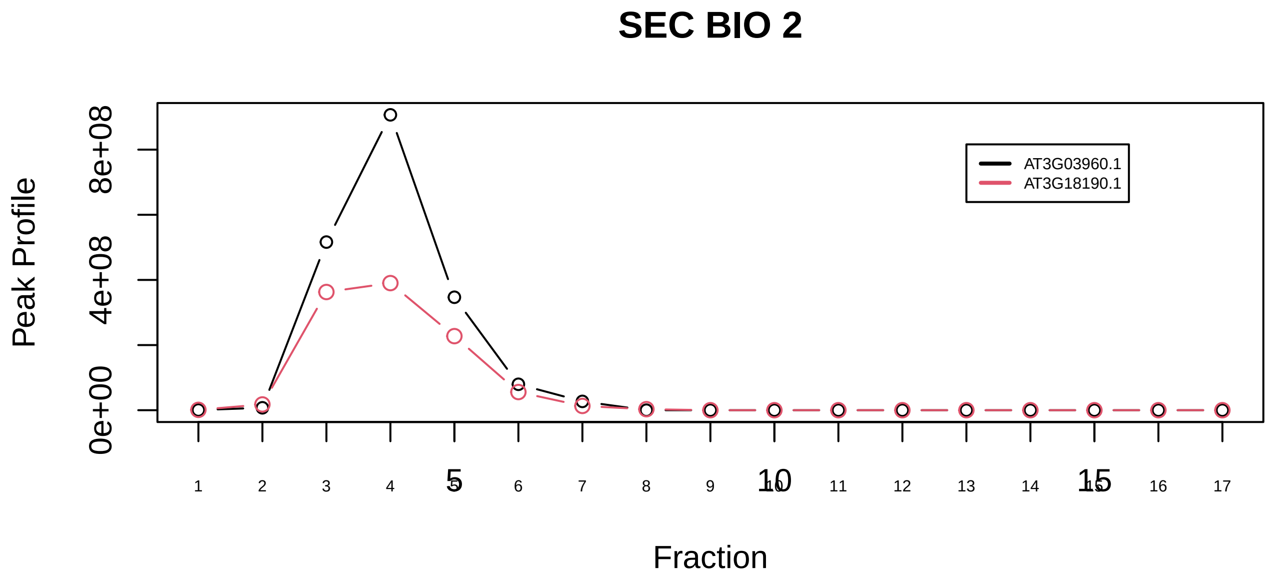

CCT complex (chaperonin containing TCP1 complex)|126−−only_2_prots_before_clustering

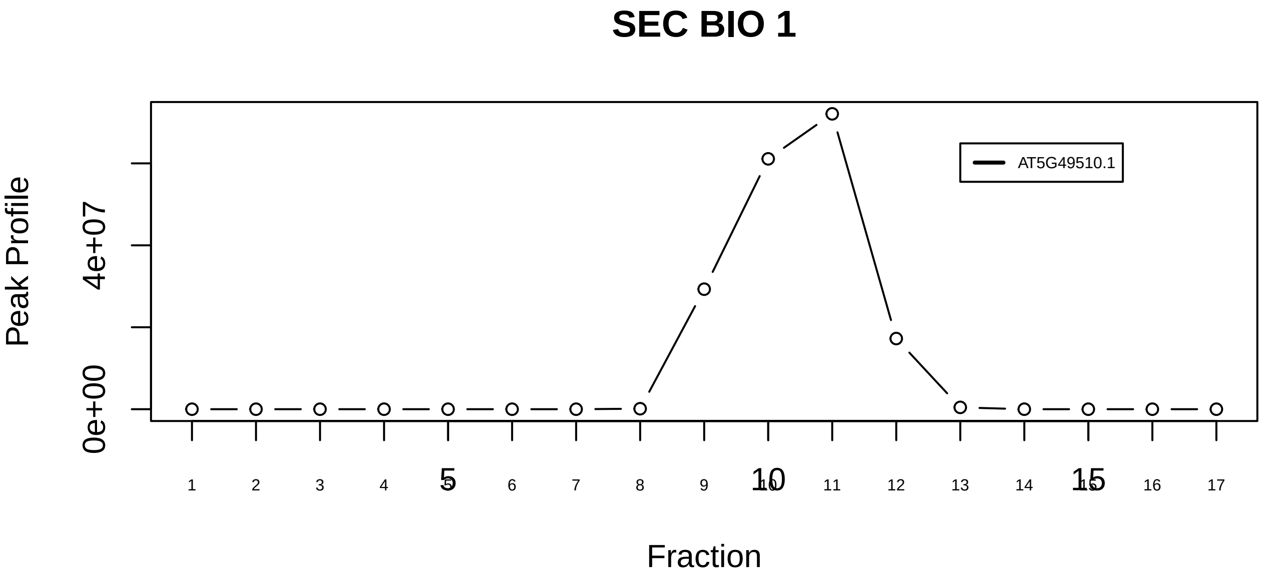

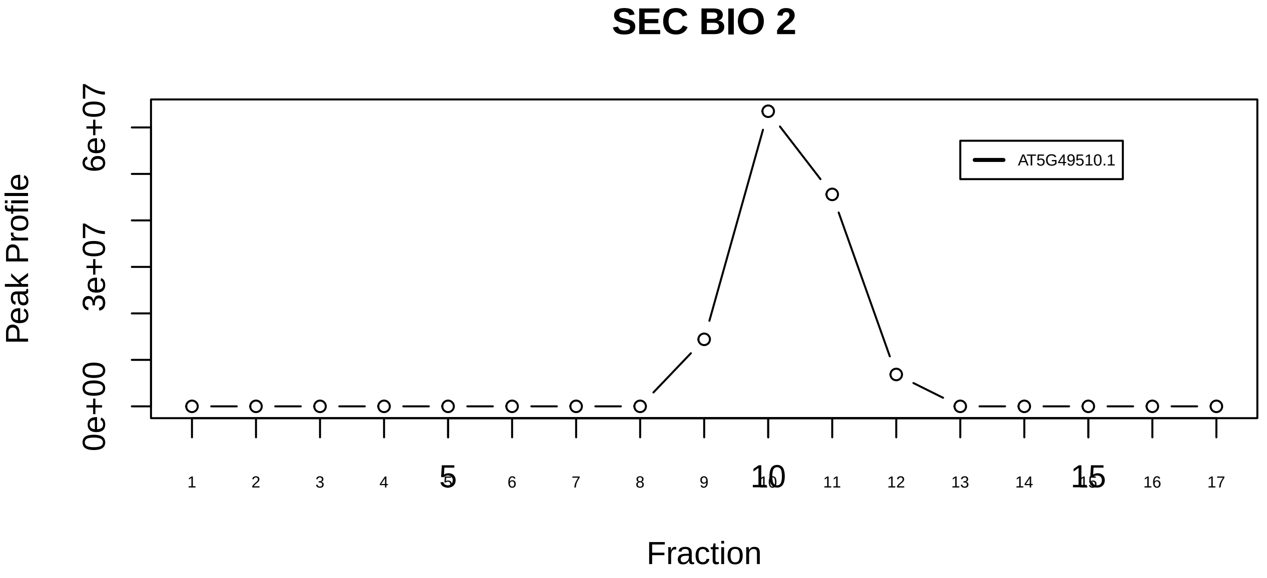

Prefoldin complex|33−−singleton_1

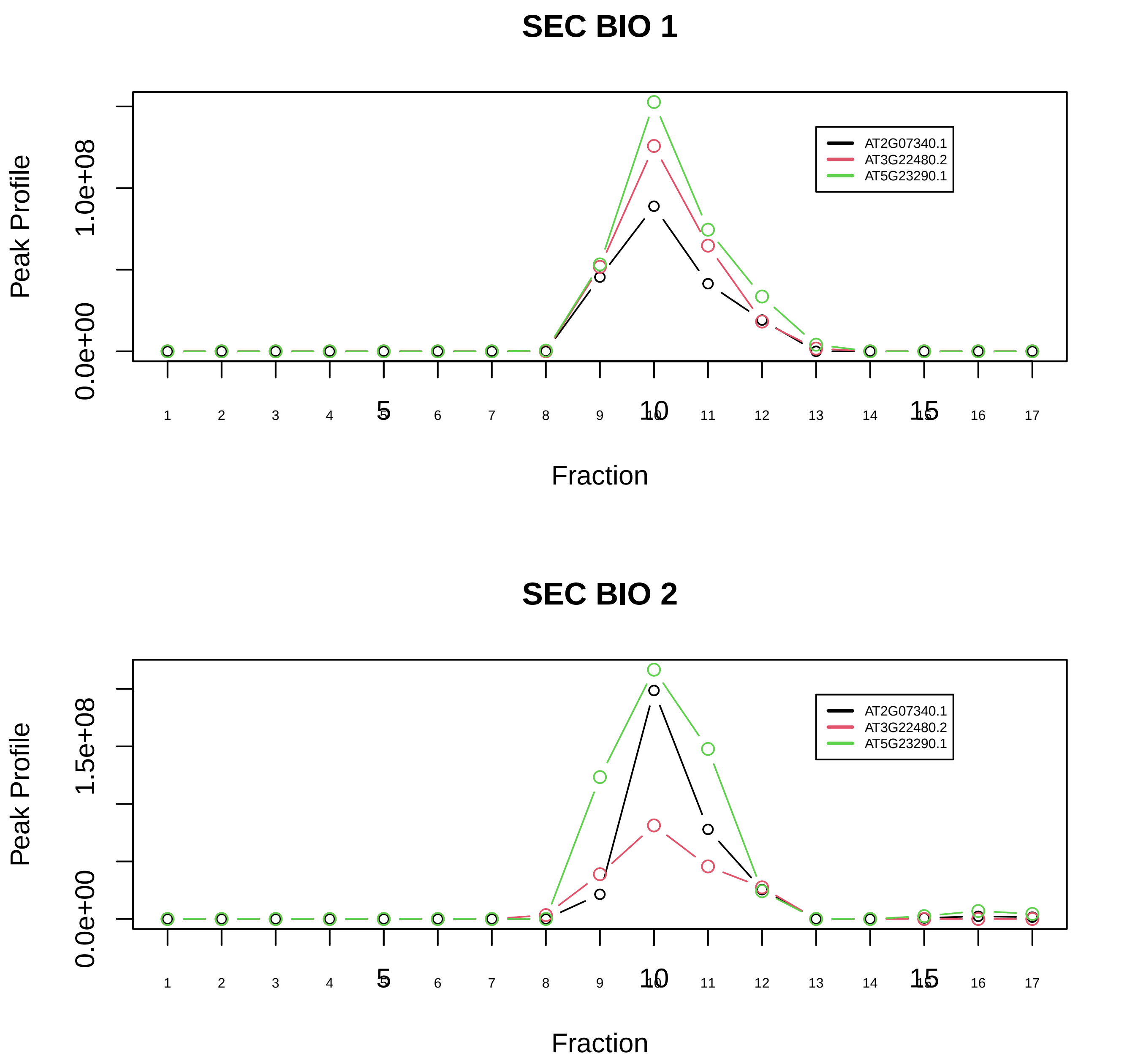

Prefoldin complex|33−−sub_1

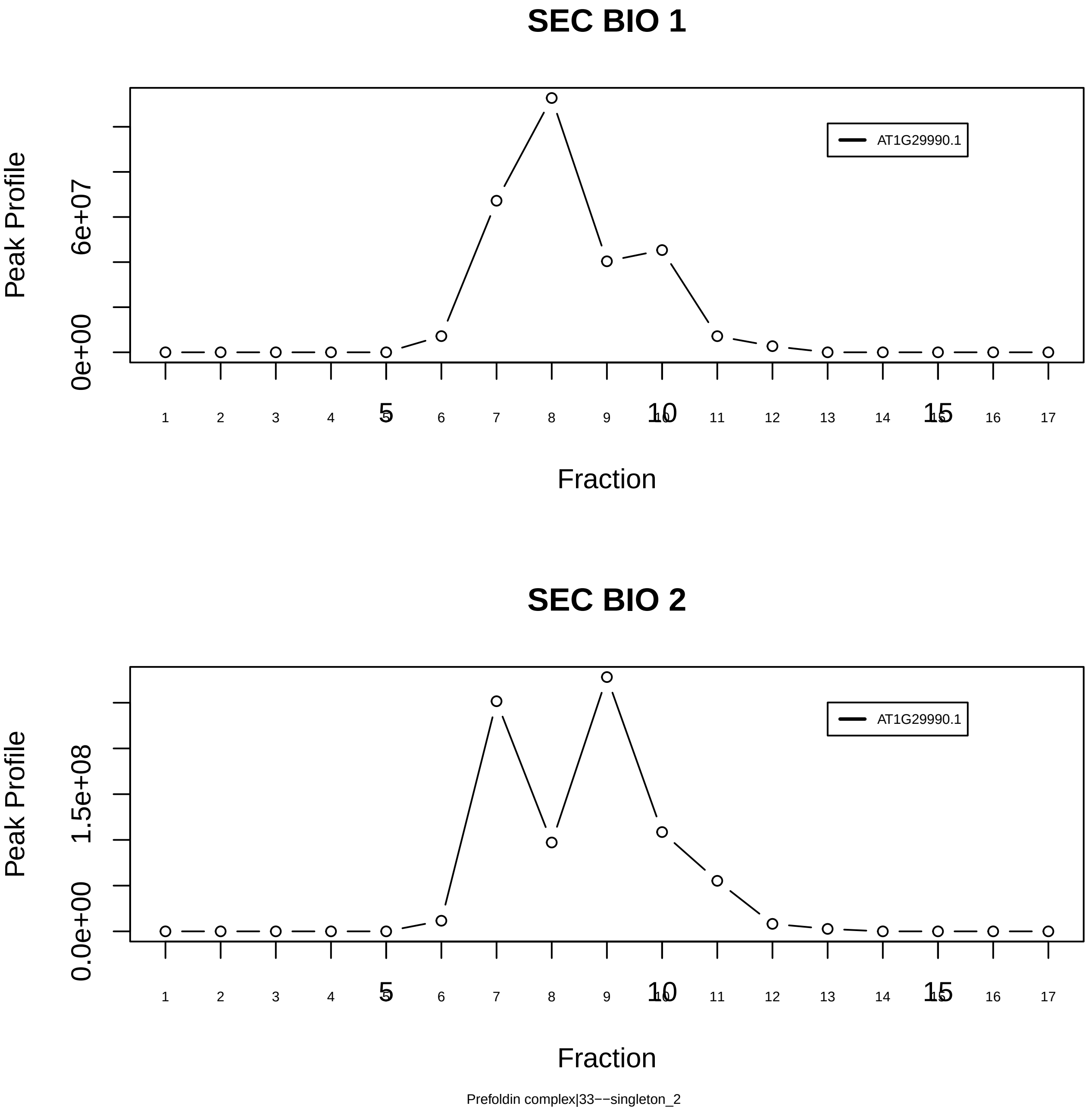
SEC BIO 1
AT1G29990.1
Peak Profile
0e+00
6e+07
Fraction
SEC BIO 2
AT1G29990.1
Peak Profile
0.0e+00
1.5e+08
Fraction
Prefoldin complex|33−−singleton_2

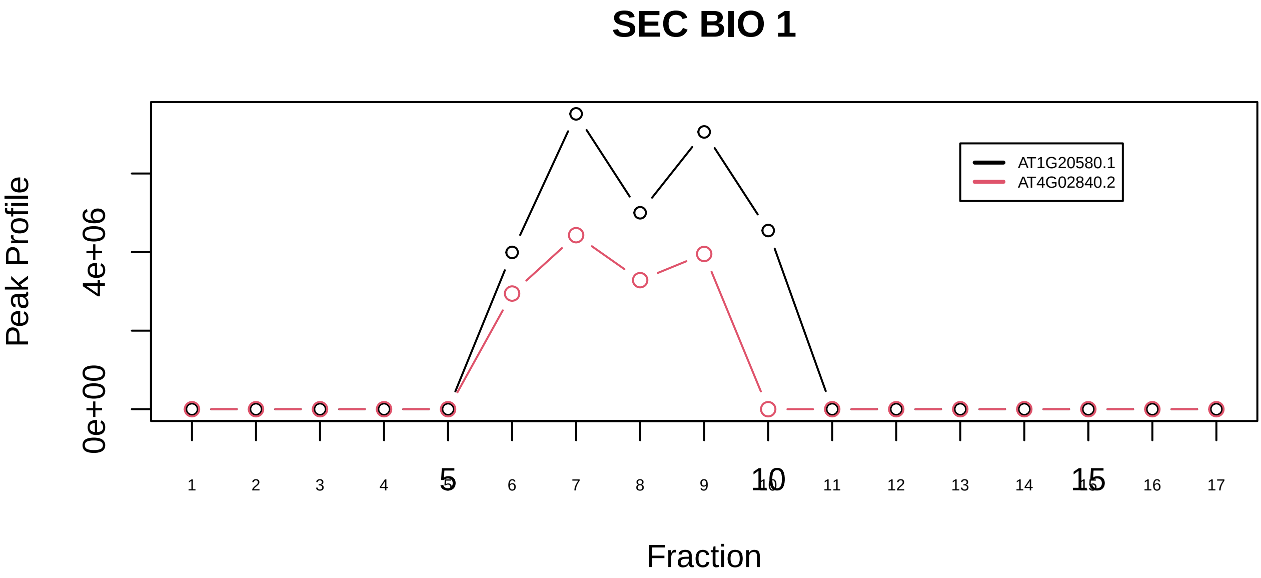

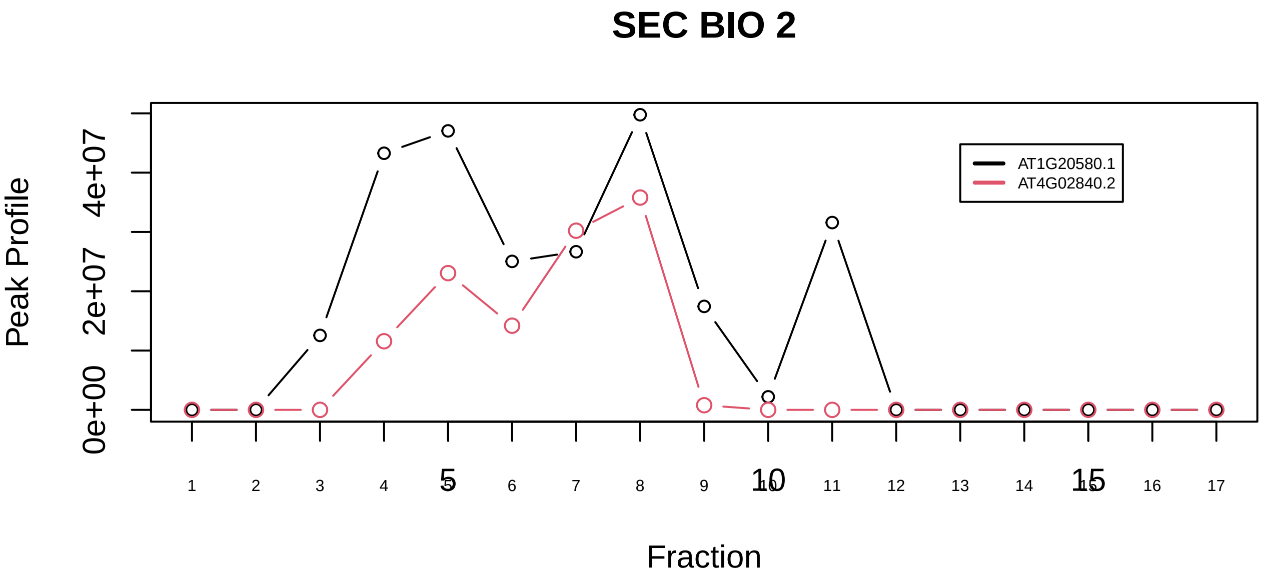

CDC5L complex|1183−−sub_1

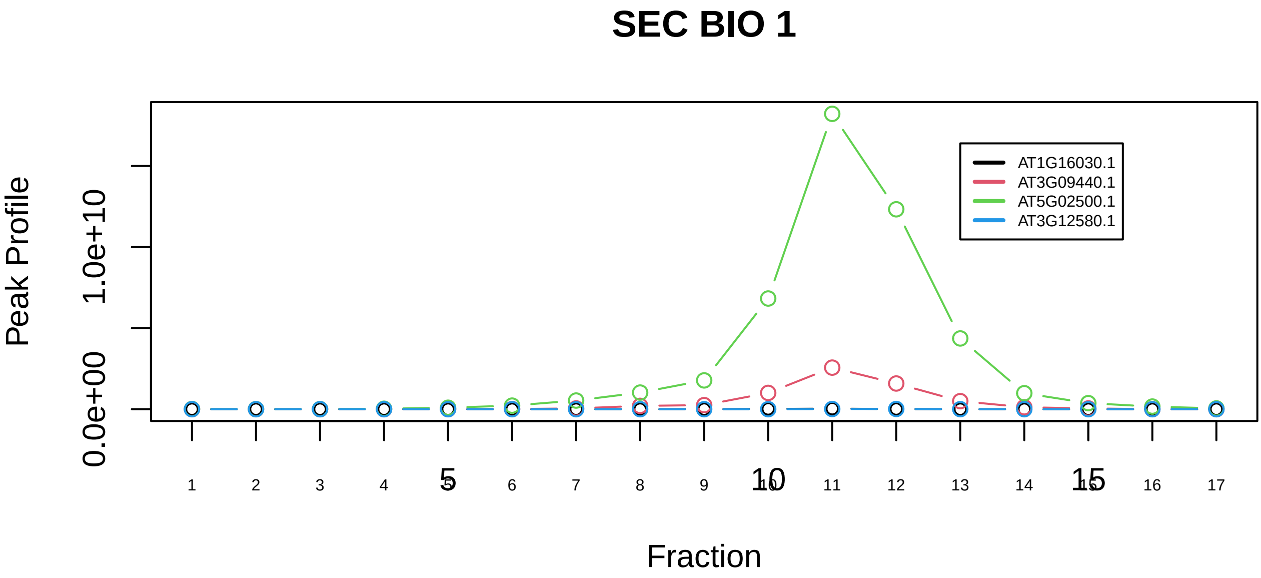

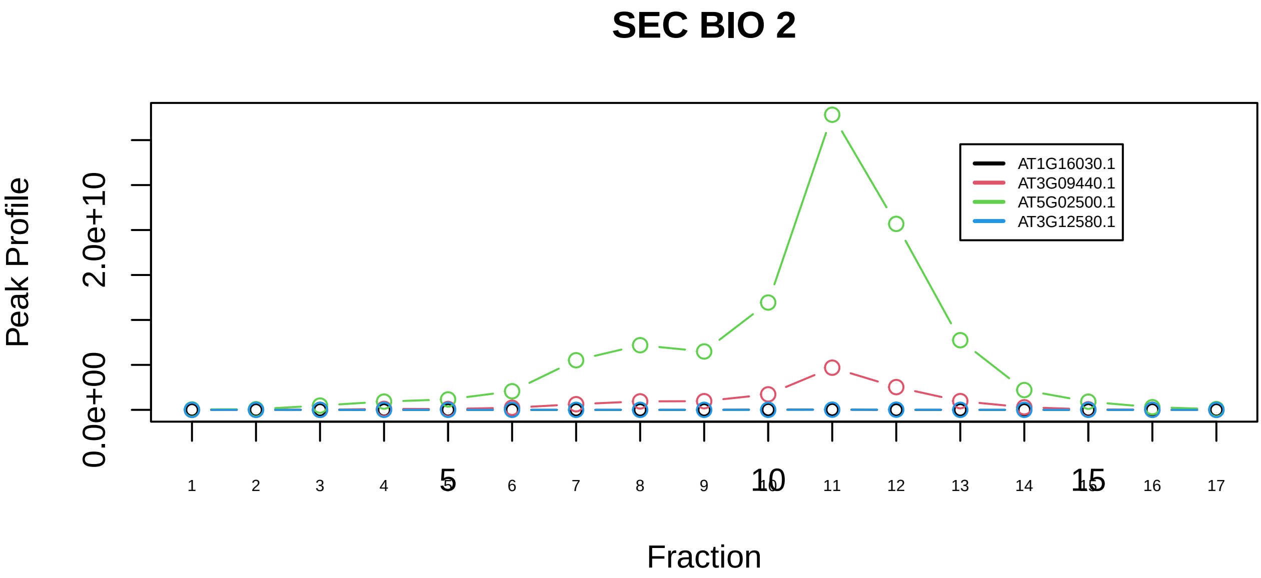

CDC5L complex|1183−−sub_orthoparalog_1

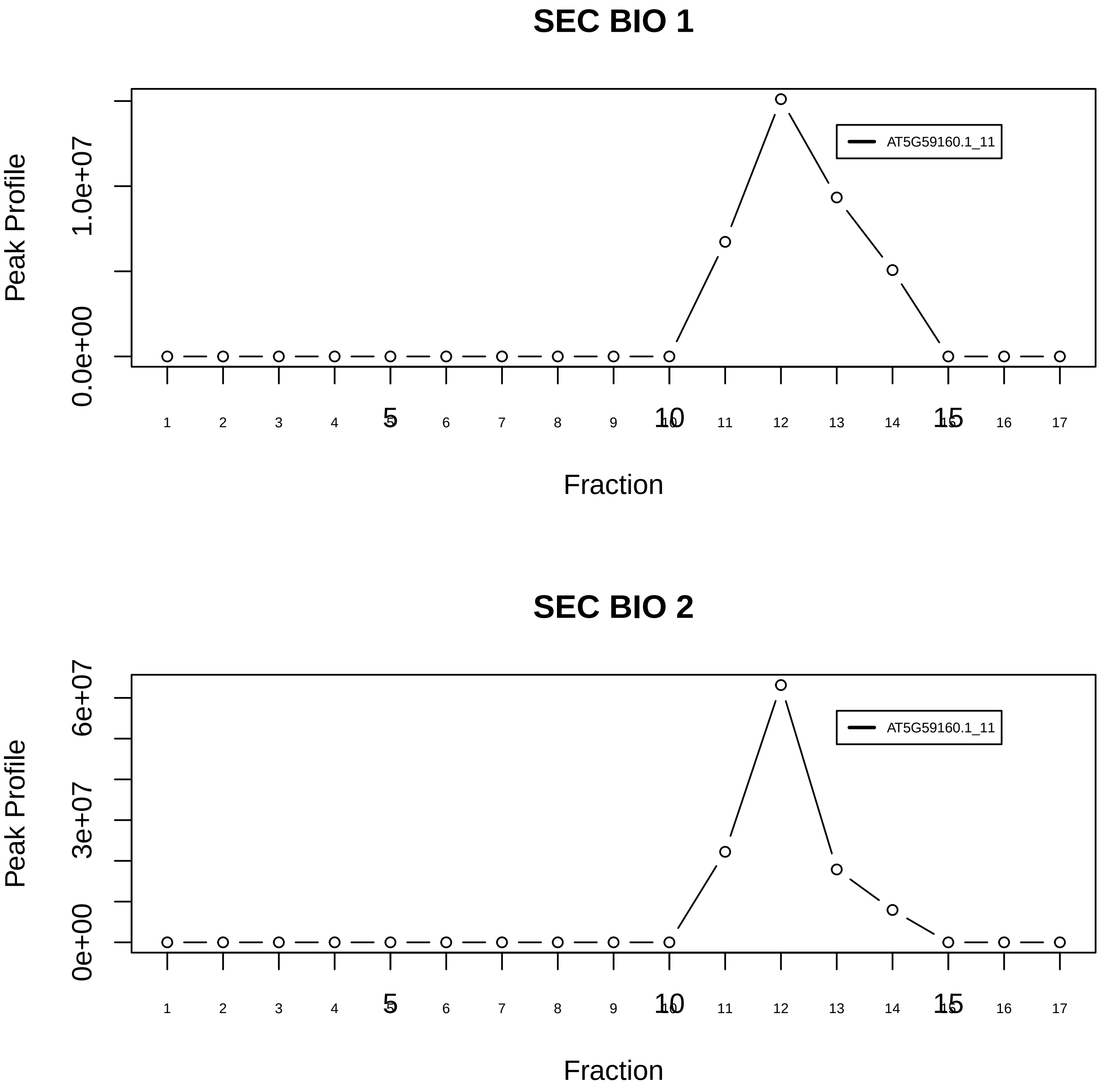

CDC5L complex|1183−−singleton_1

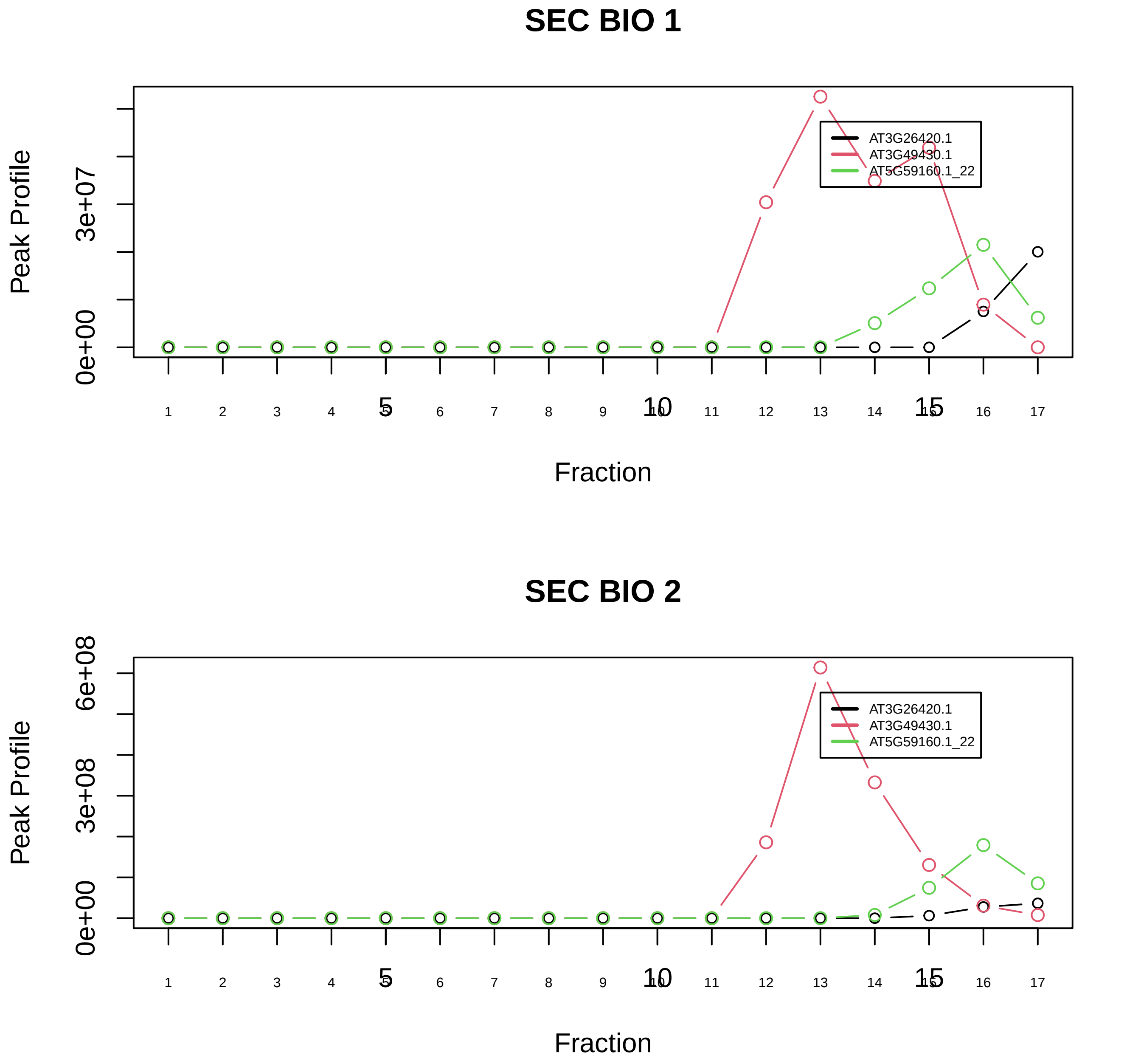

CDC5L complex|1183−−sub_2

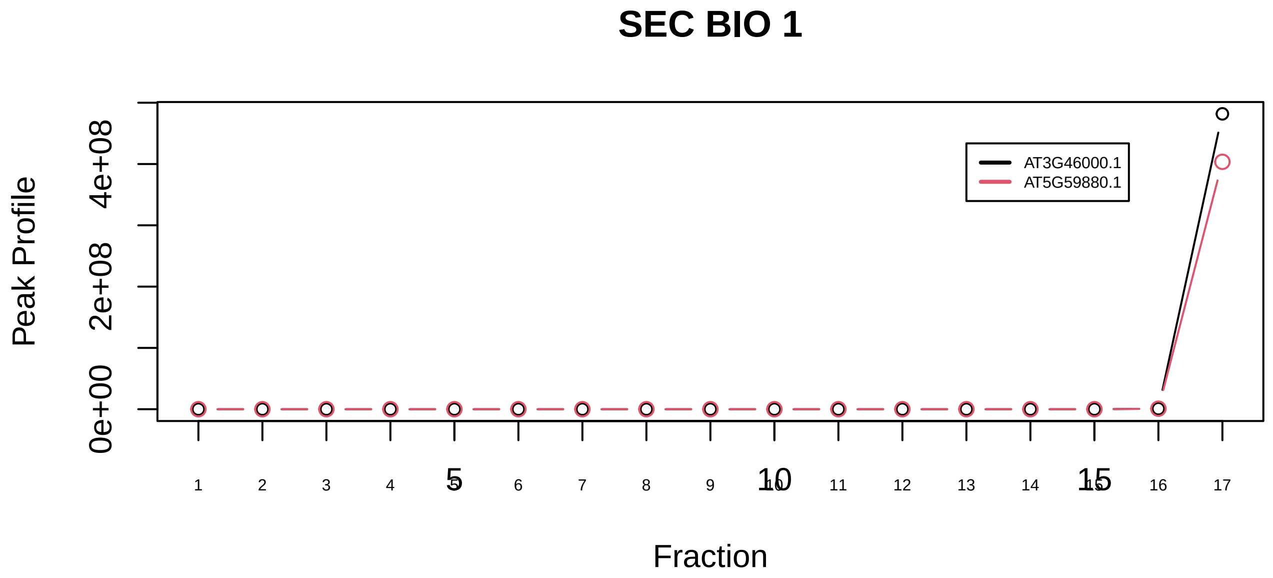

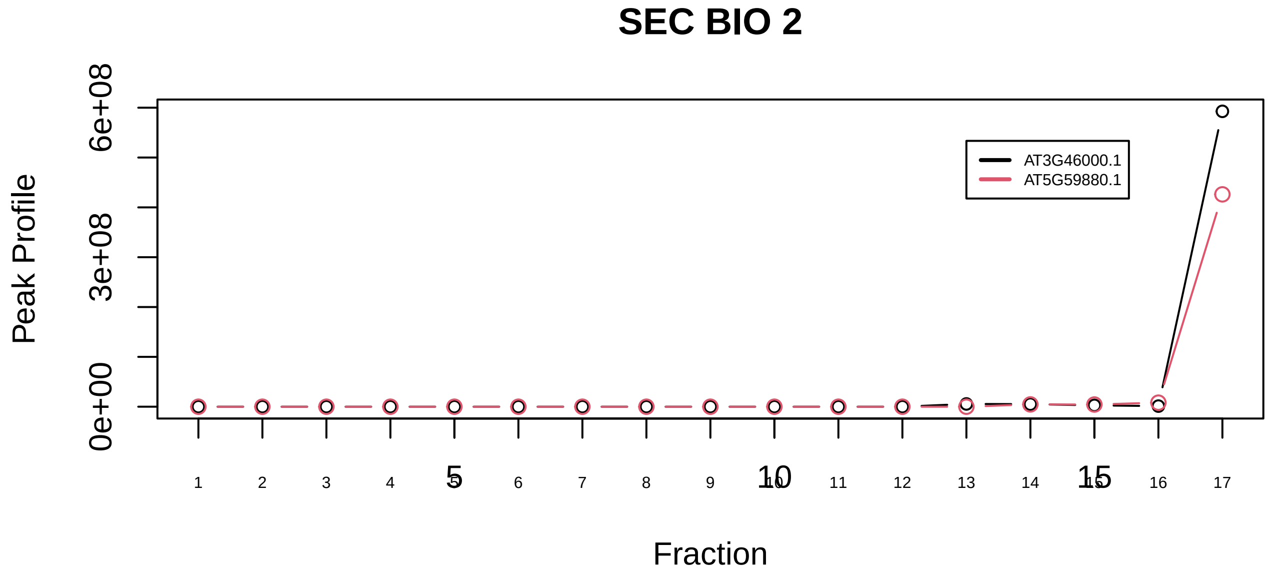

Cofilin−actin−CAP1 complex|2255−−sub_orthoparalog_1

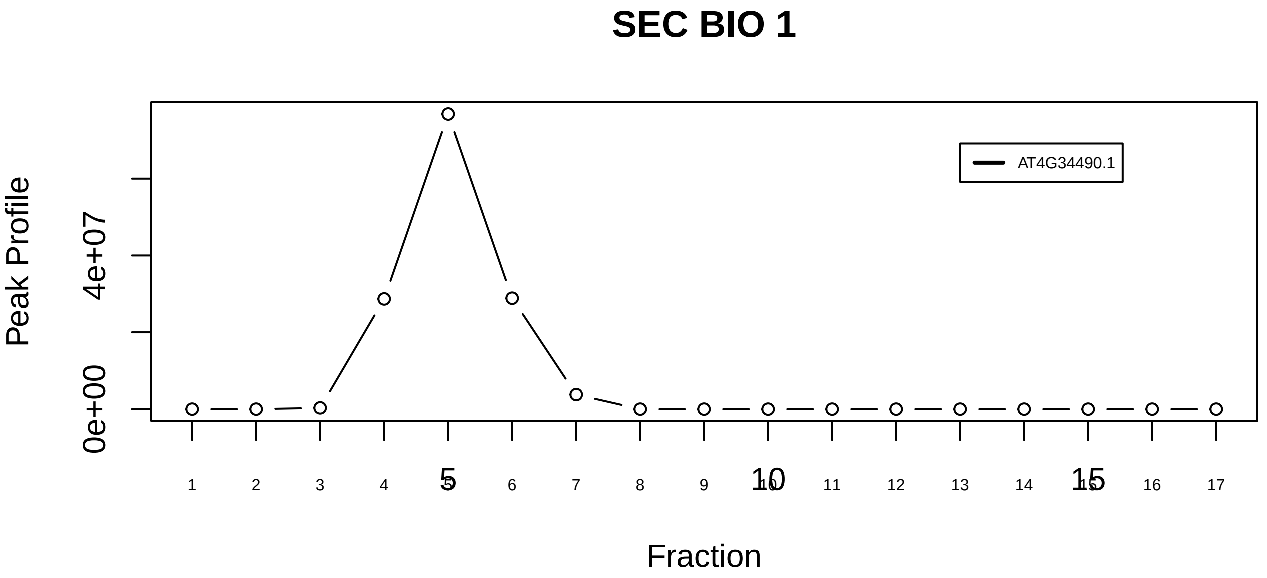

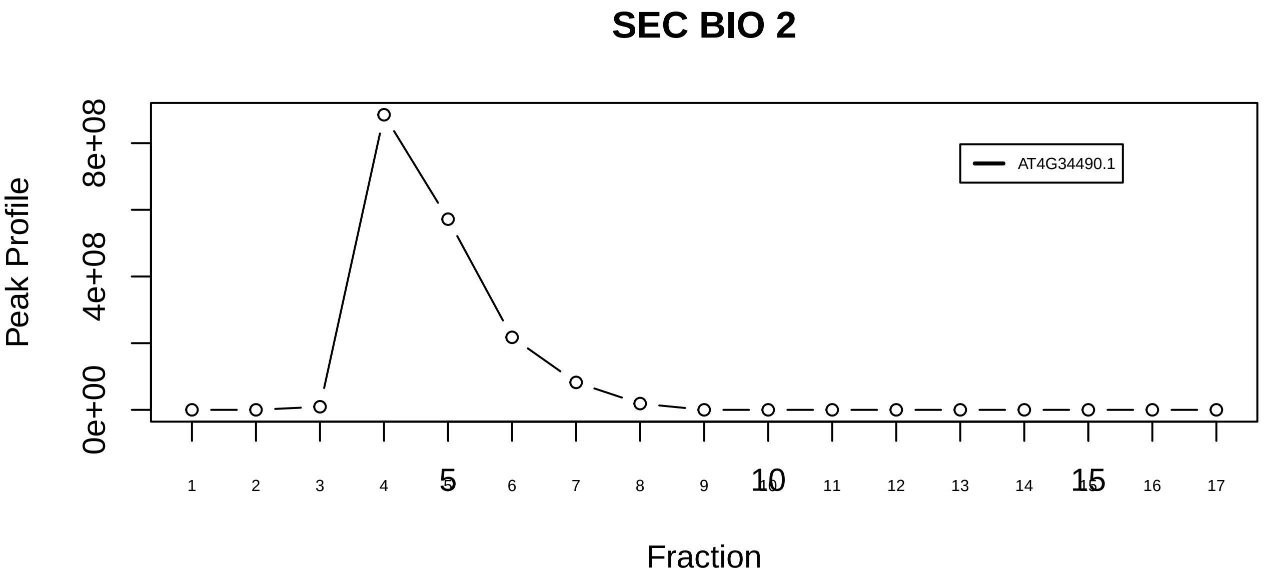

Cofilin−actin−CAP1 complex|2255−−singleton_1

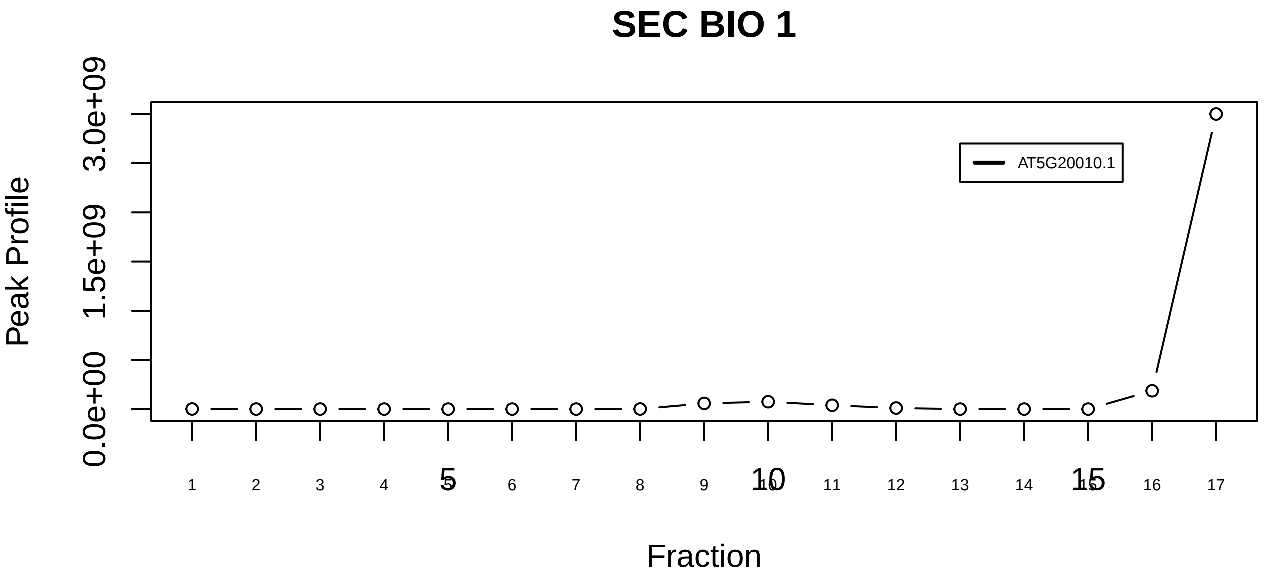

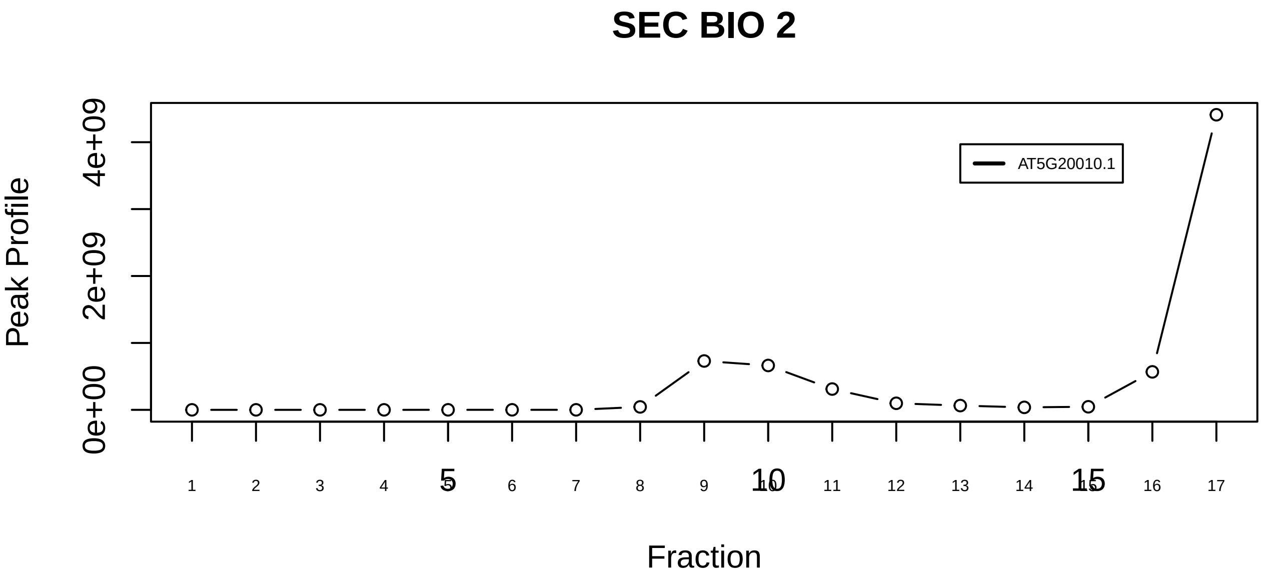

CRM1−RAN−PHAX−CBC complex (cap binding complex)|1176−−singleton_1

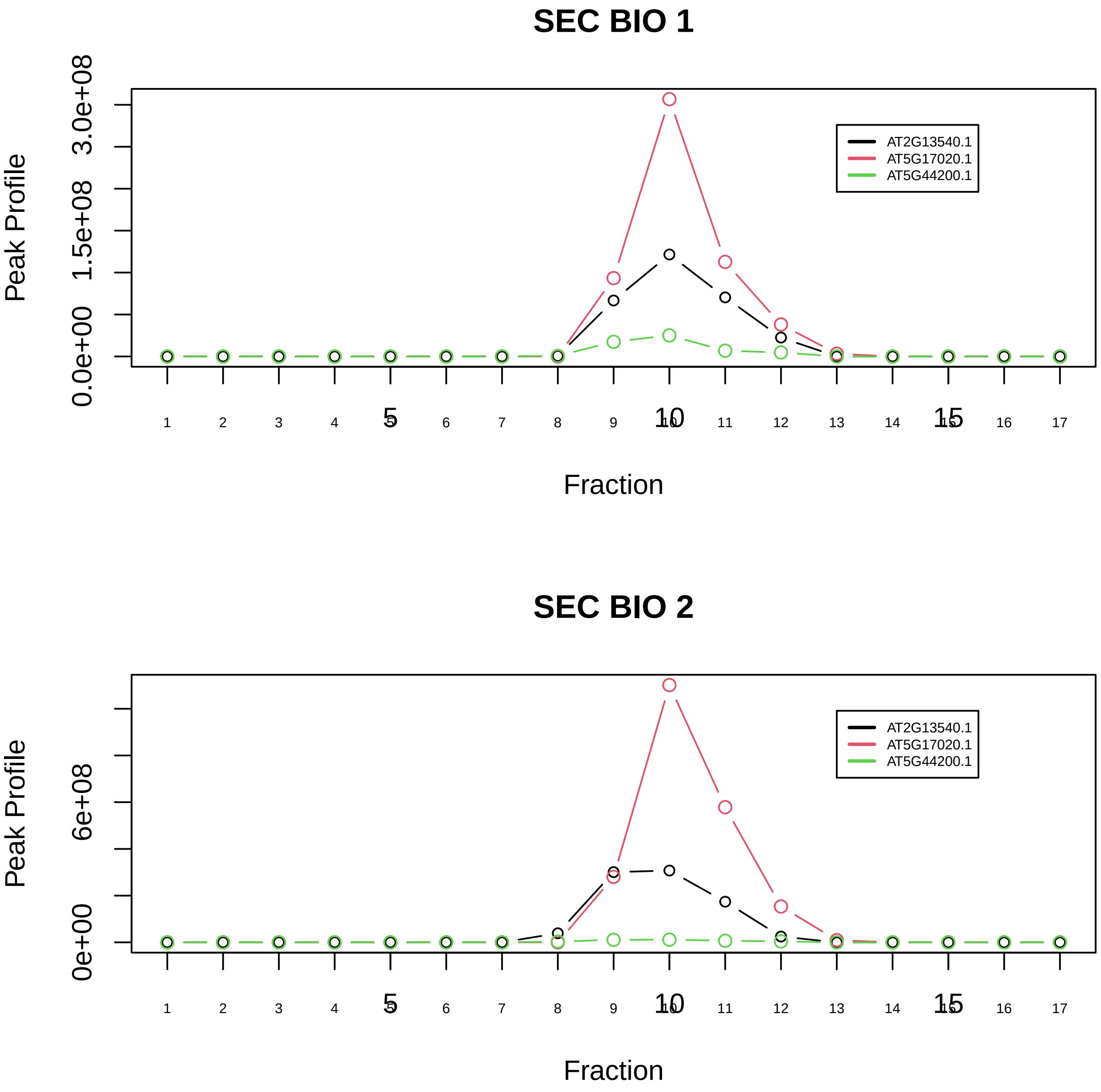

CRM1−RAN−PHAX−CBC complex (cap binding complex)|1176−−sub_1

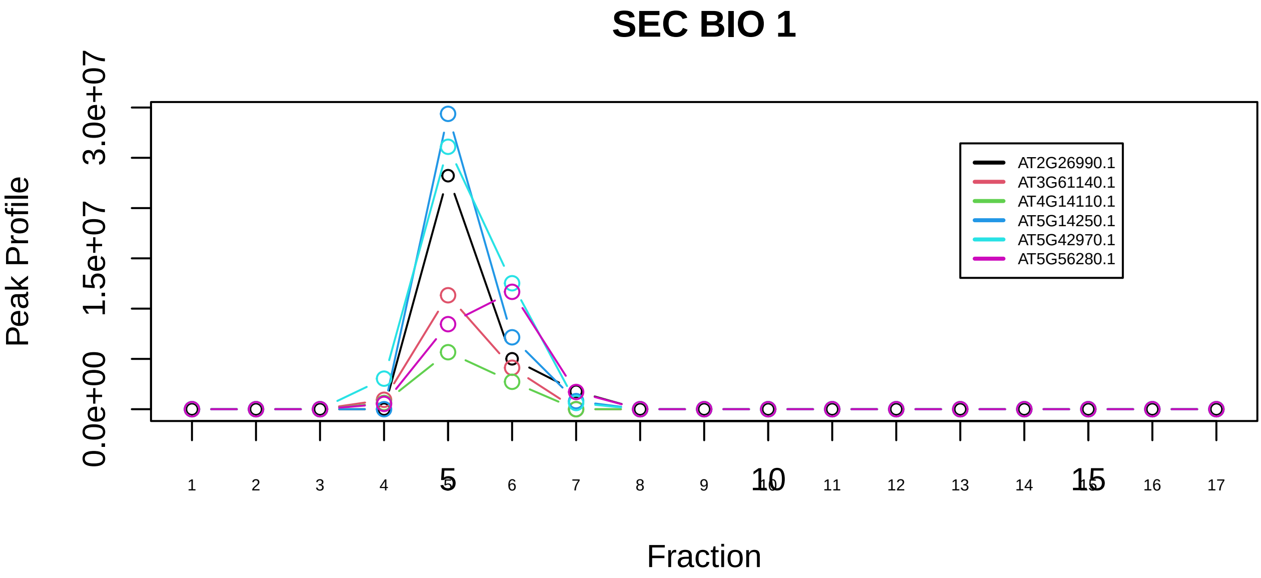

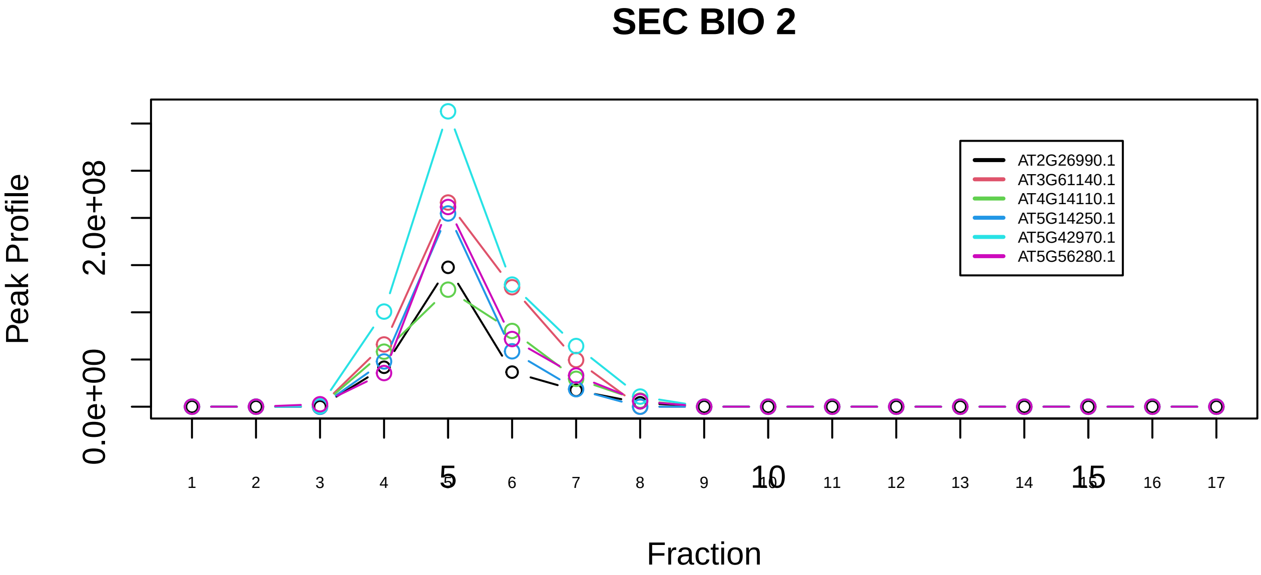

DDB2 complex|726−−sub_1

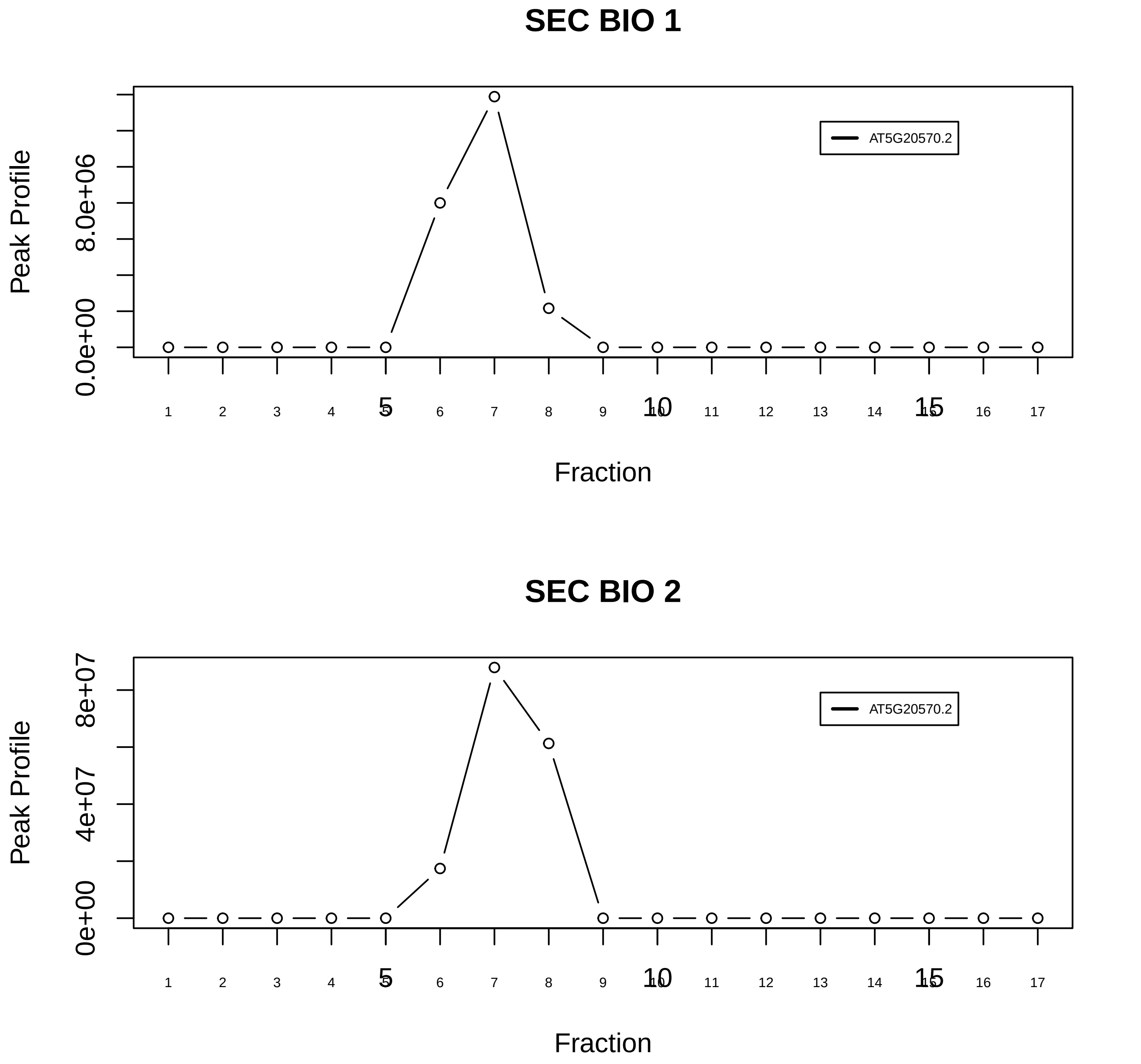

DDB2 complex|726−−singleton_1

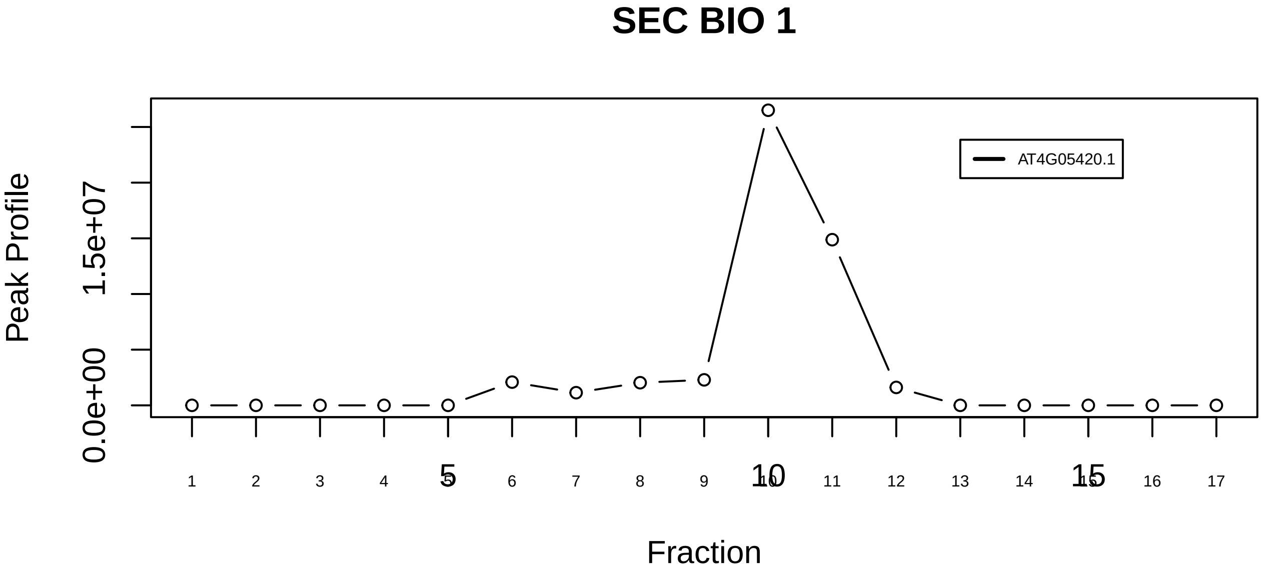

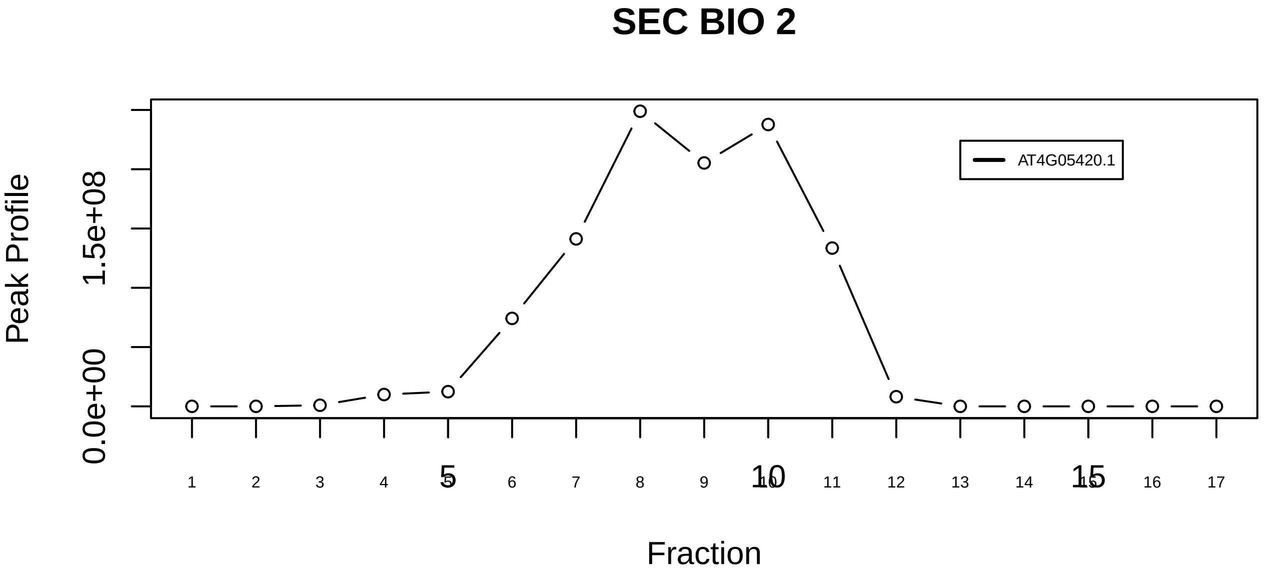

DDB2 complex|726−−singleton_2

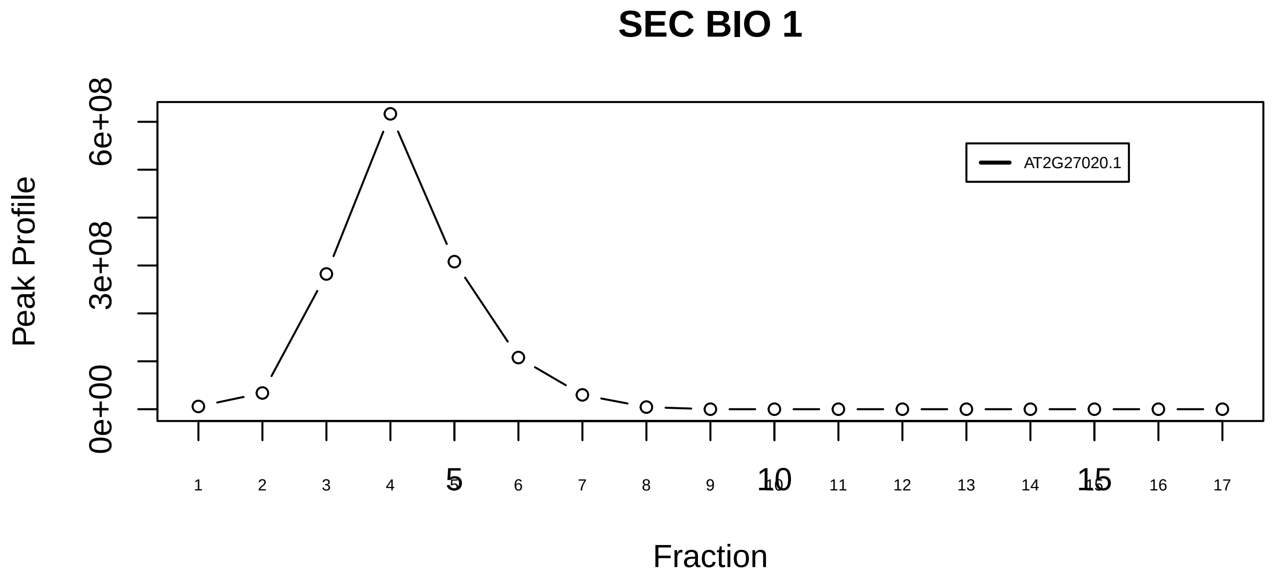

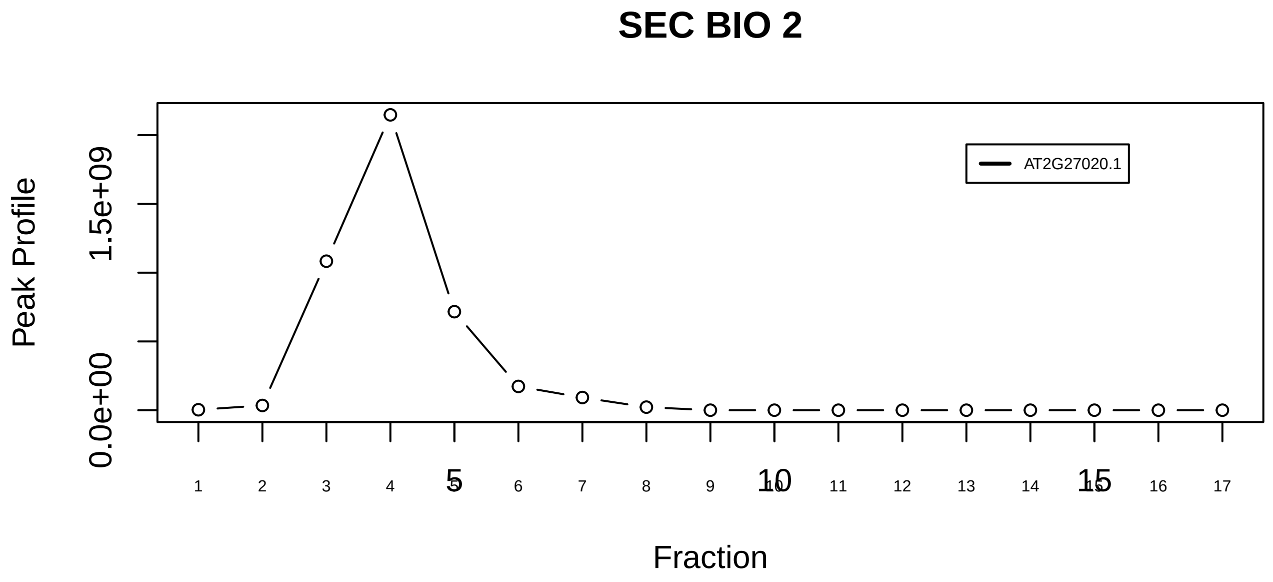

DNAJB2−HSPA8−PSMA3 complex|2129−−singleton_1

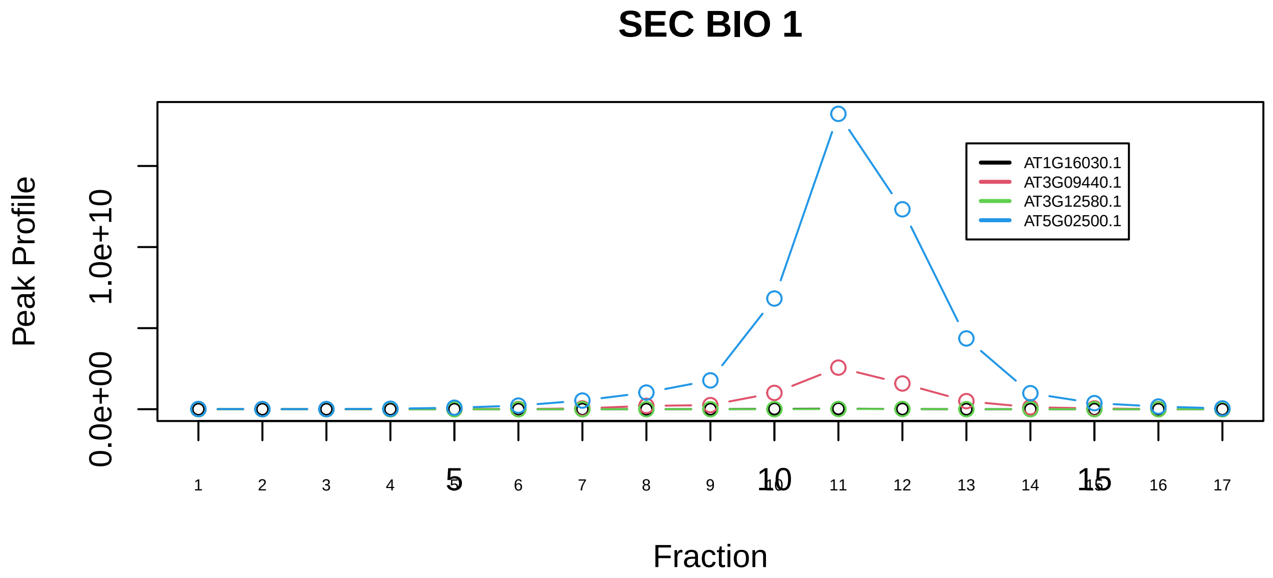

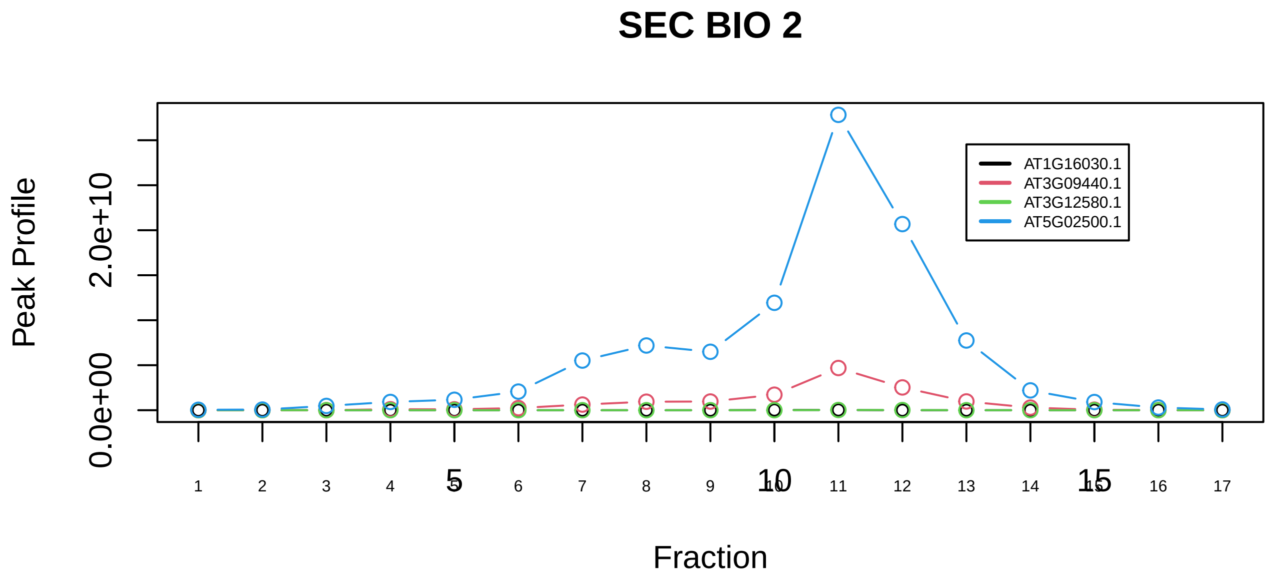

DNAJB2−HSPA8−PSMA3 complex|2129−−sub_orthoparalog_1

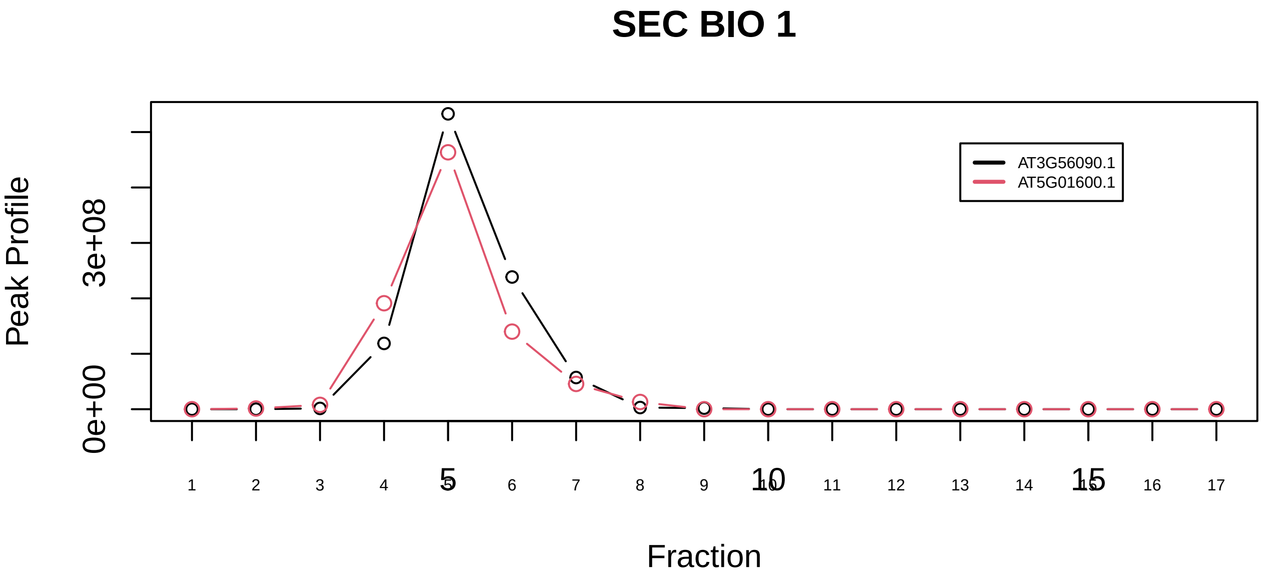

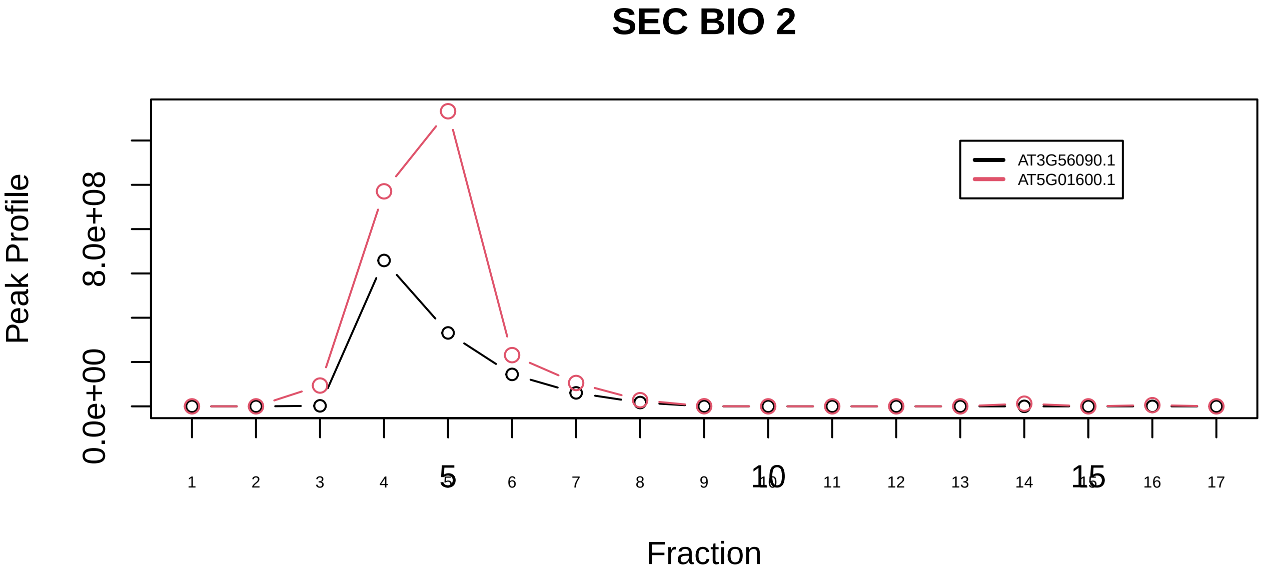

Ferritin complex|6135−−only_2_prots_before_clustering

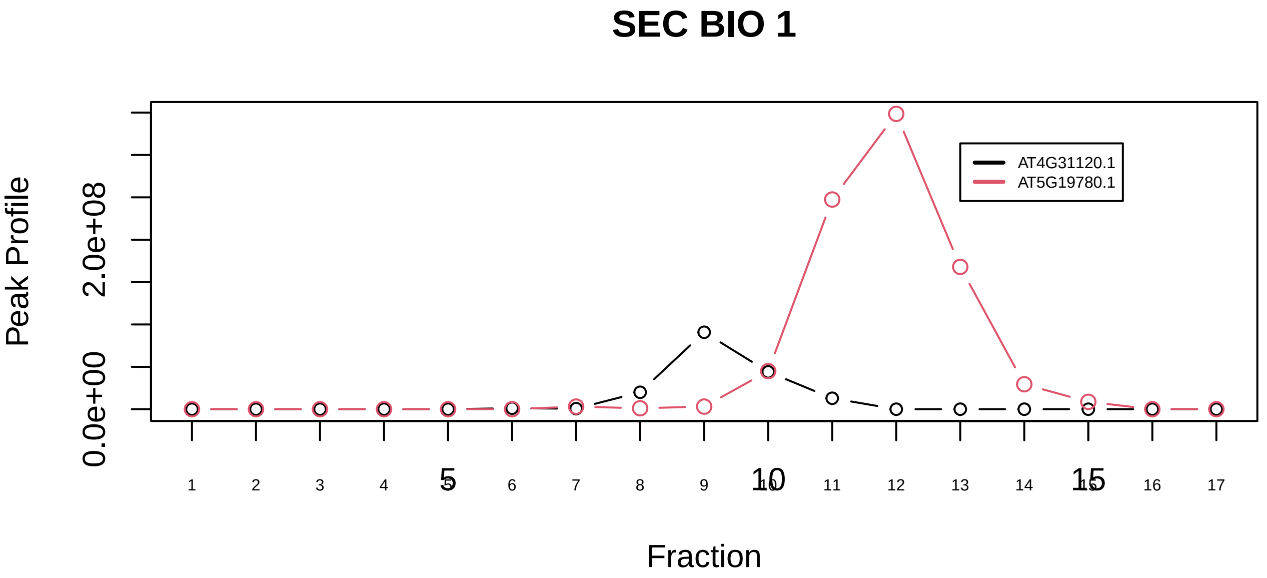

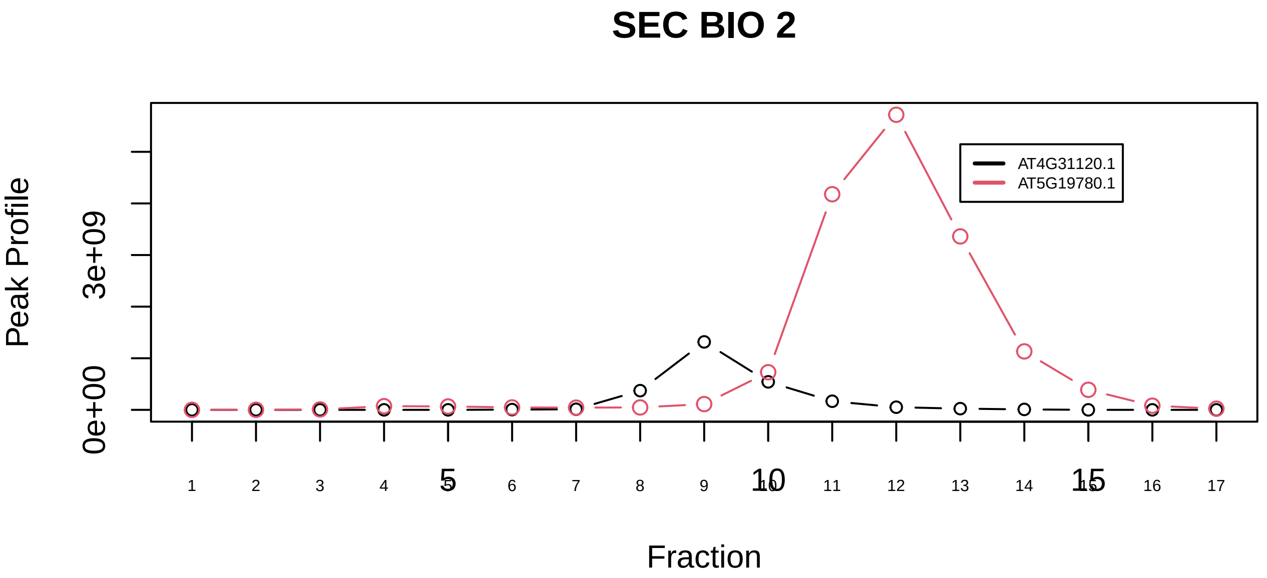

FIB−associated protein complex|1231−−only_2_prots_before_clustering

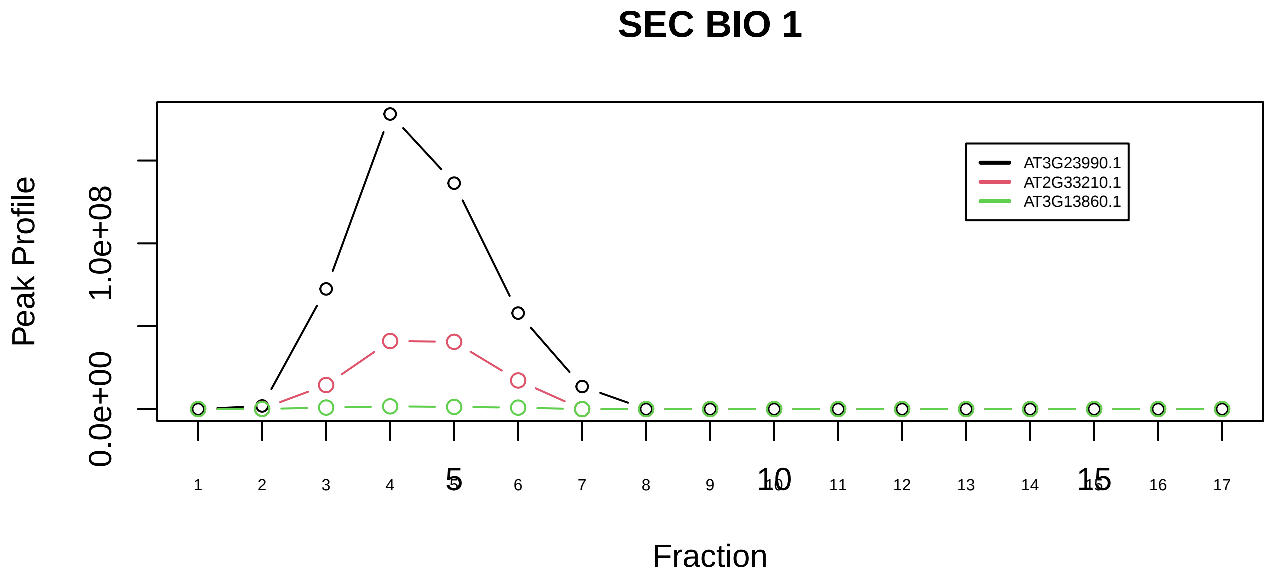

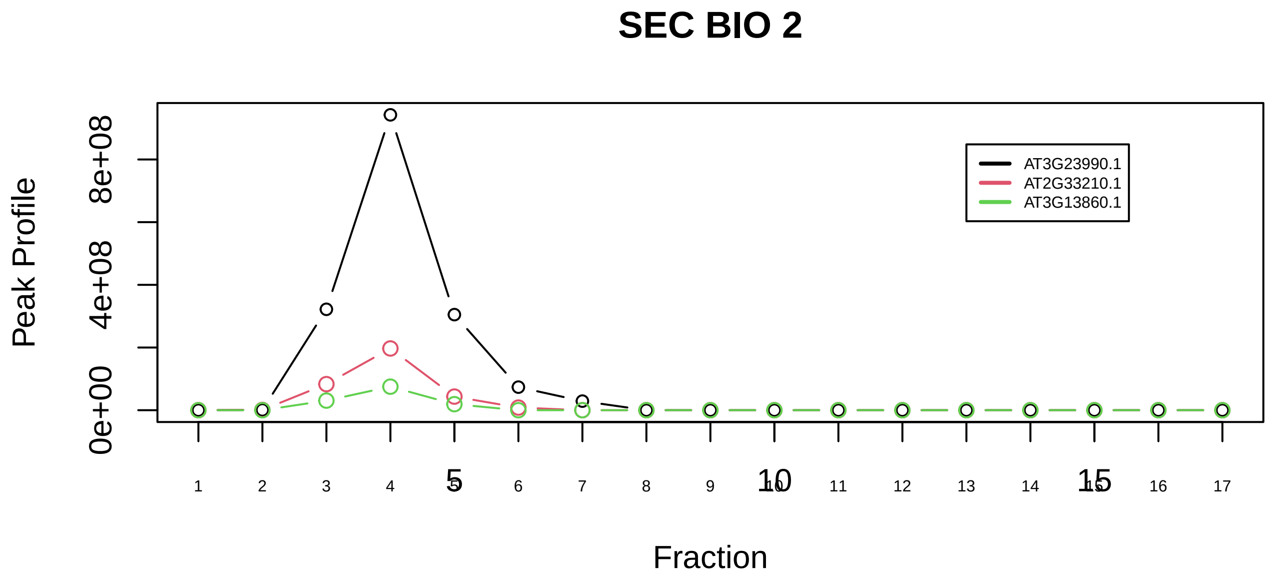

Frataxin complex|1094−−sub_orthoparalog_1

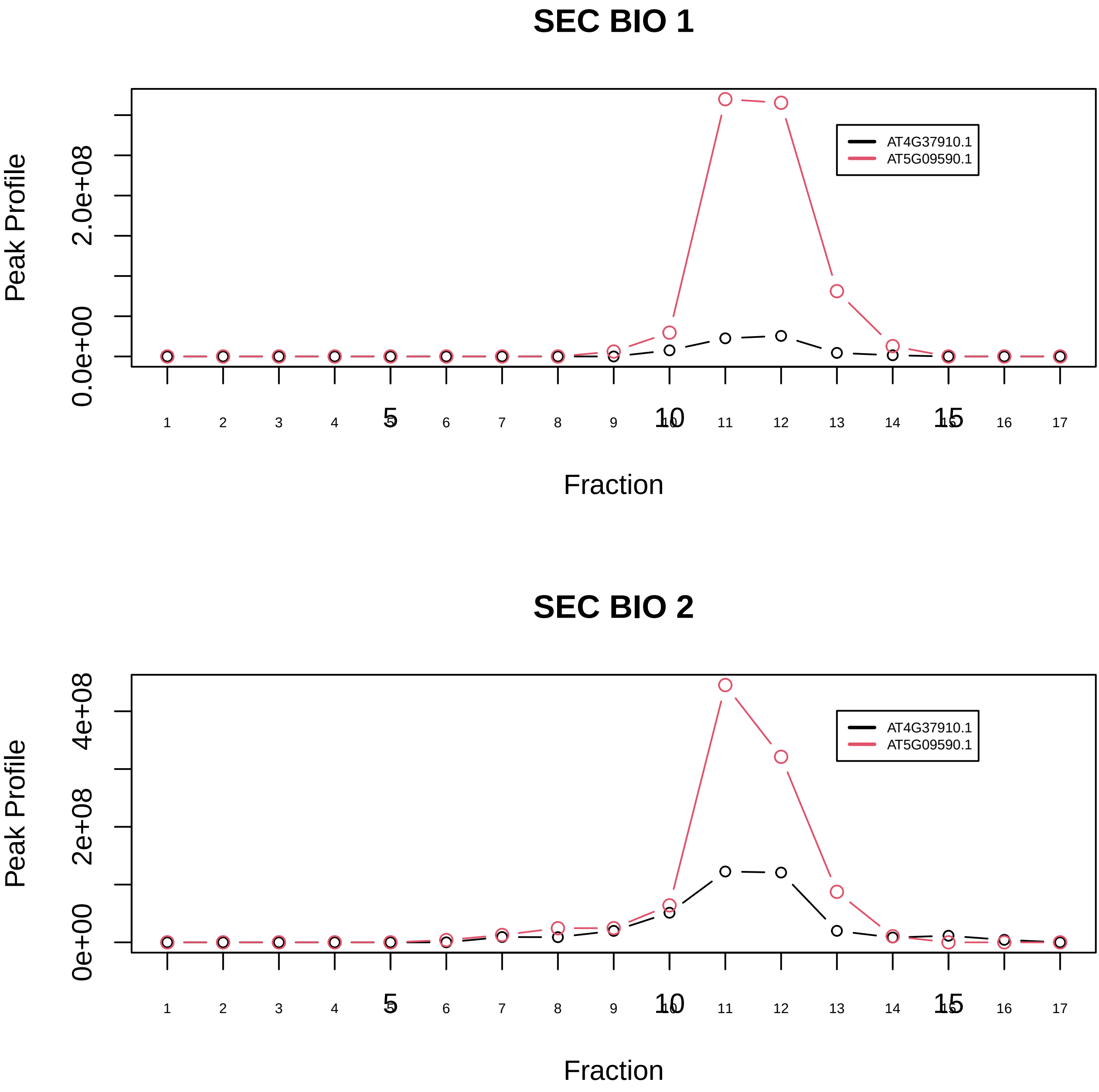

Frataxin complex|1094−−sub_orthoparalog_2

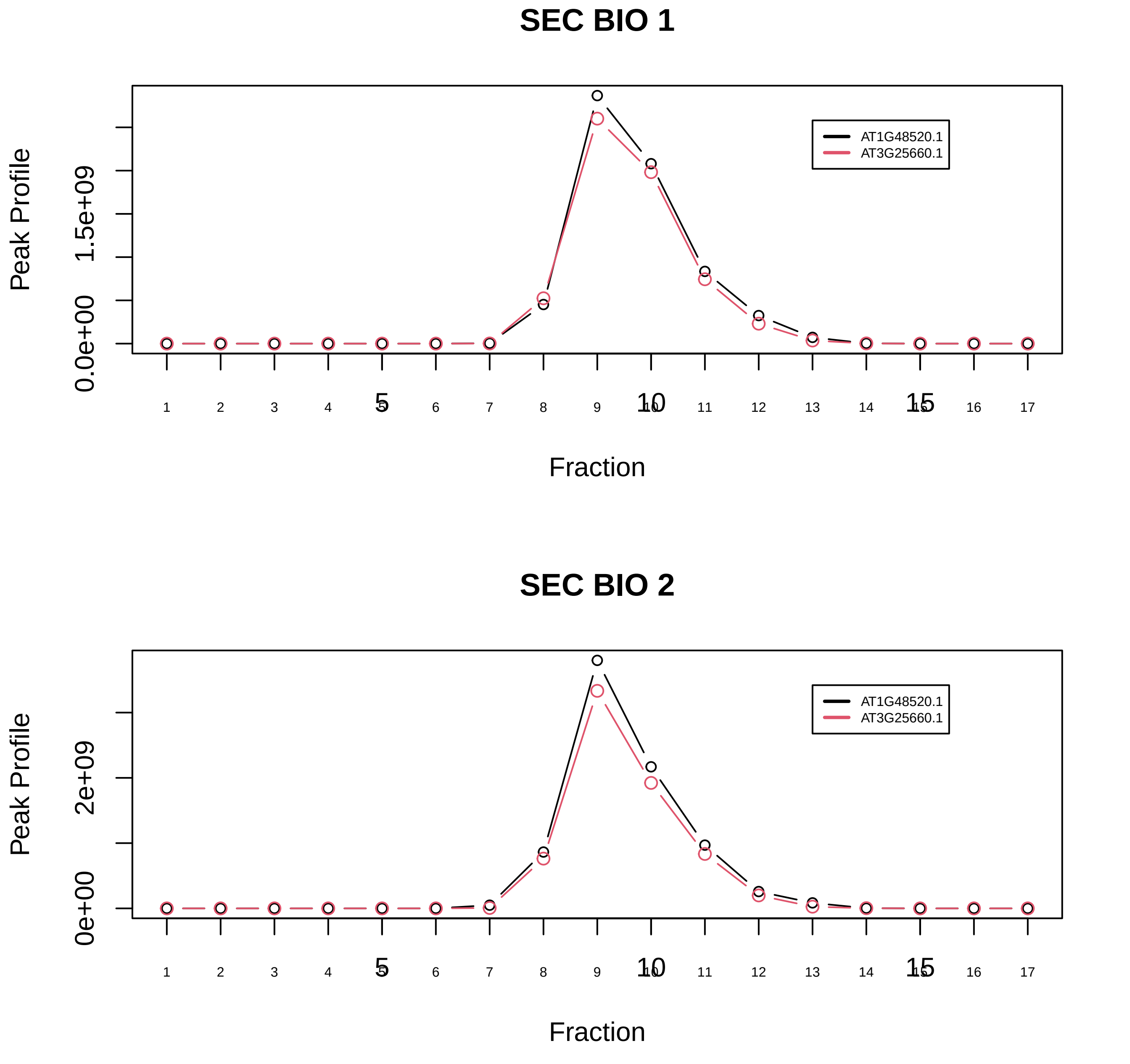

GATB−GATC−QRSL1 complex|6959−−only_2_prots_before_clustering

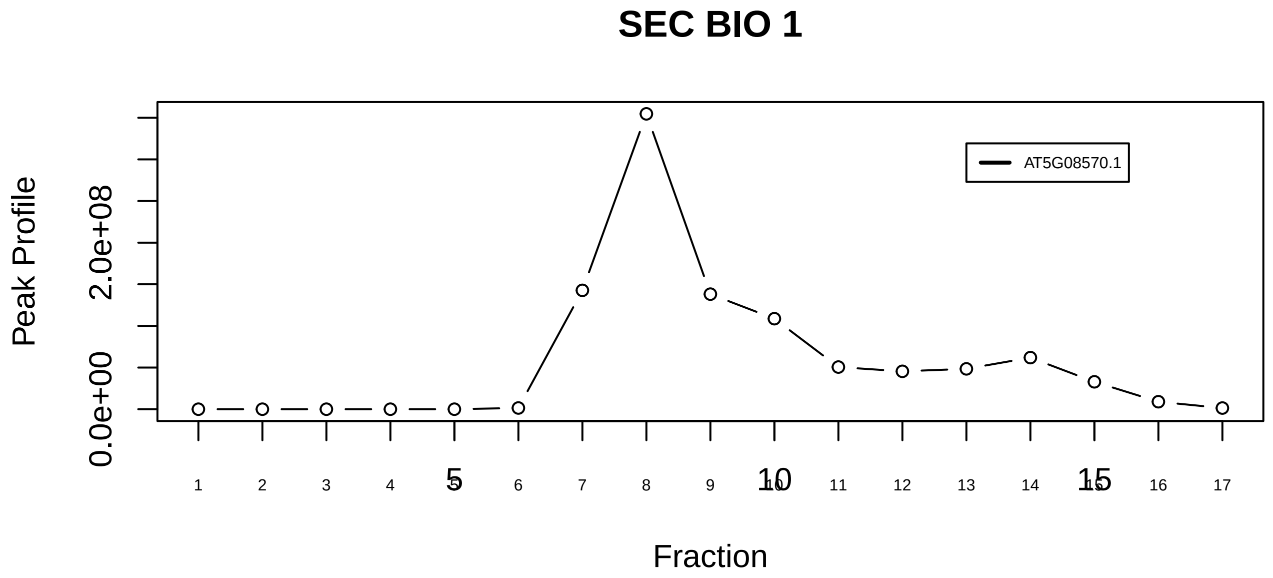

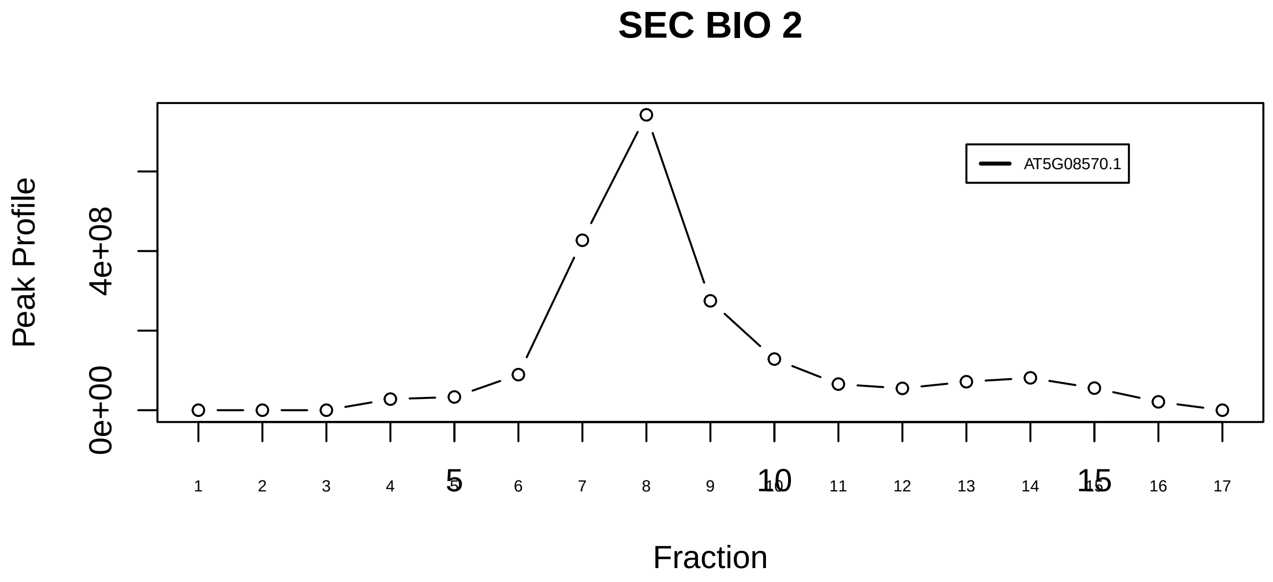

GSK3B−HSP90AA1−PKM2 complex|7284−−singleton_1

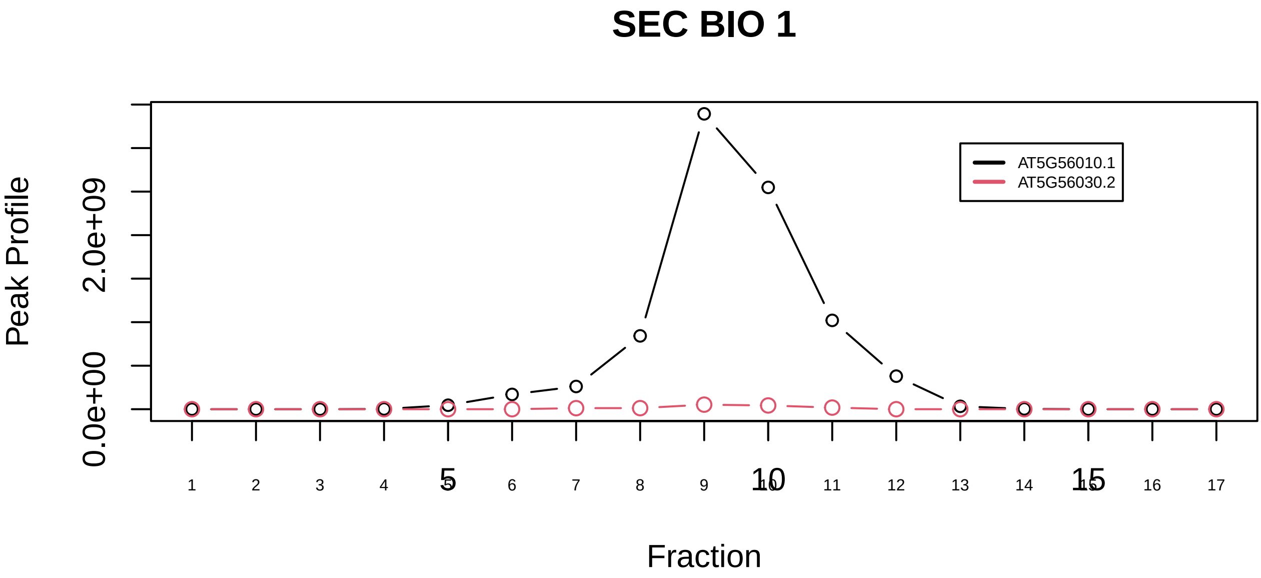

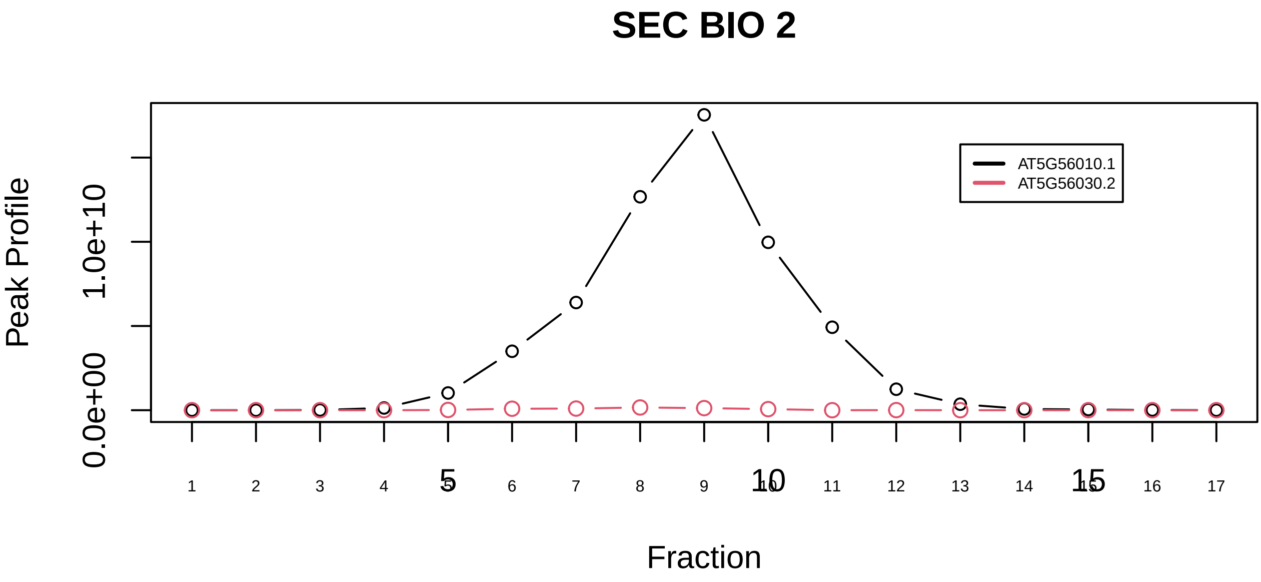

GSK3B−HSP90AA1−PKM2 complex|7284−−sub_orthoparalog_1

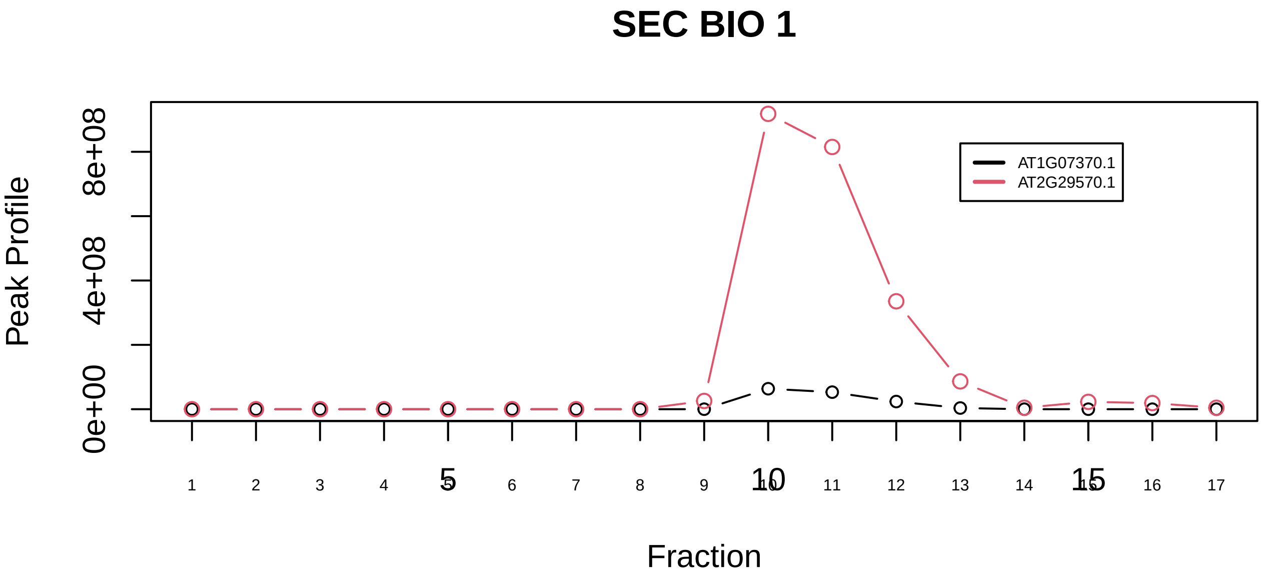

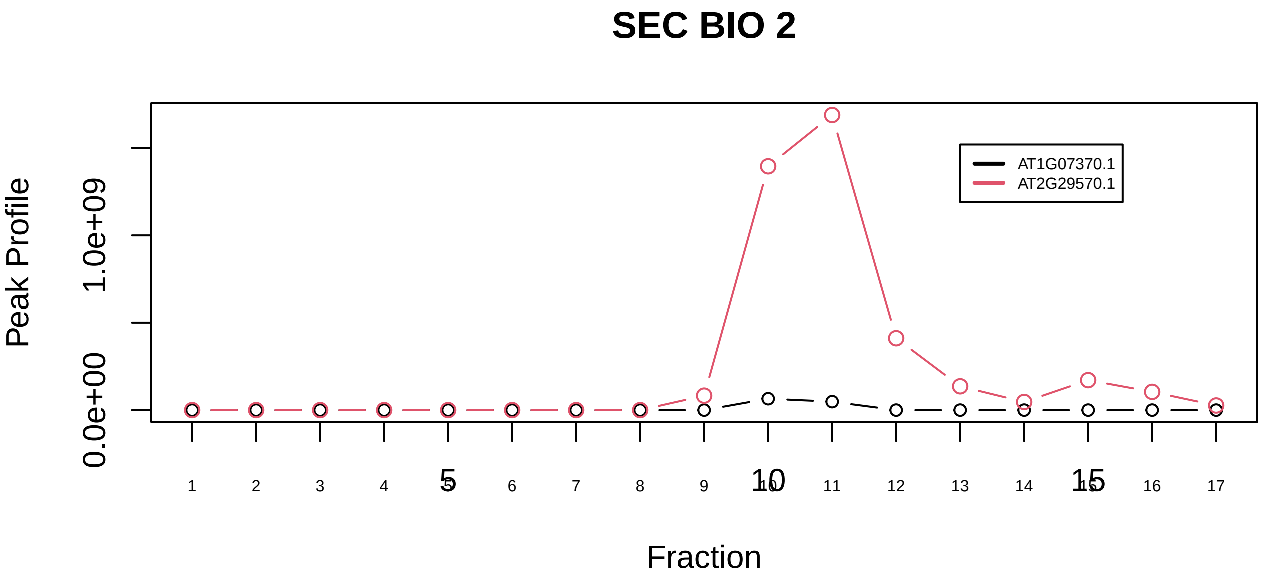

H2A−H2B−TDIF2−PCNA complex|6737−−sub_orthoparalog_1

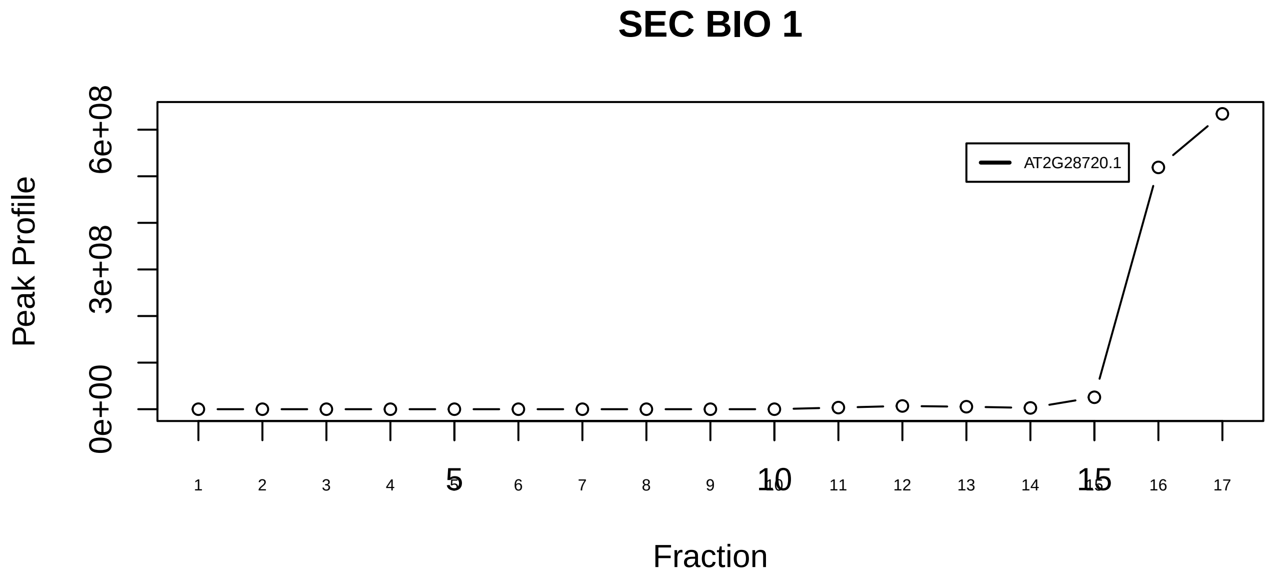

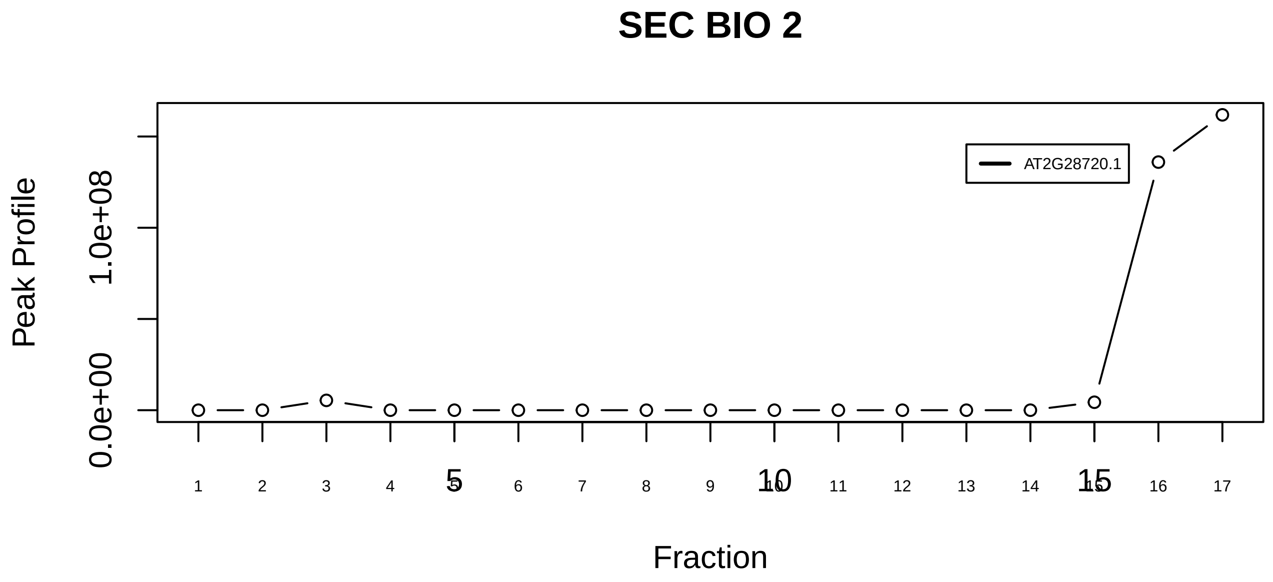

H2A−H2B−TDIF2−PCNA complex|6737−−singleton_1

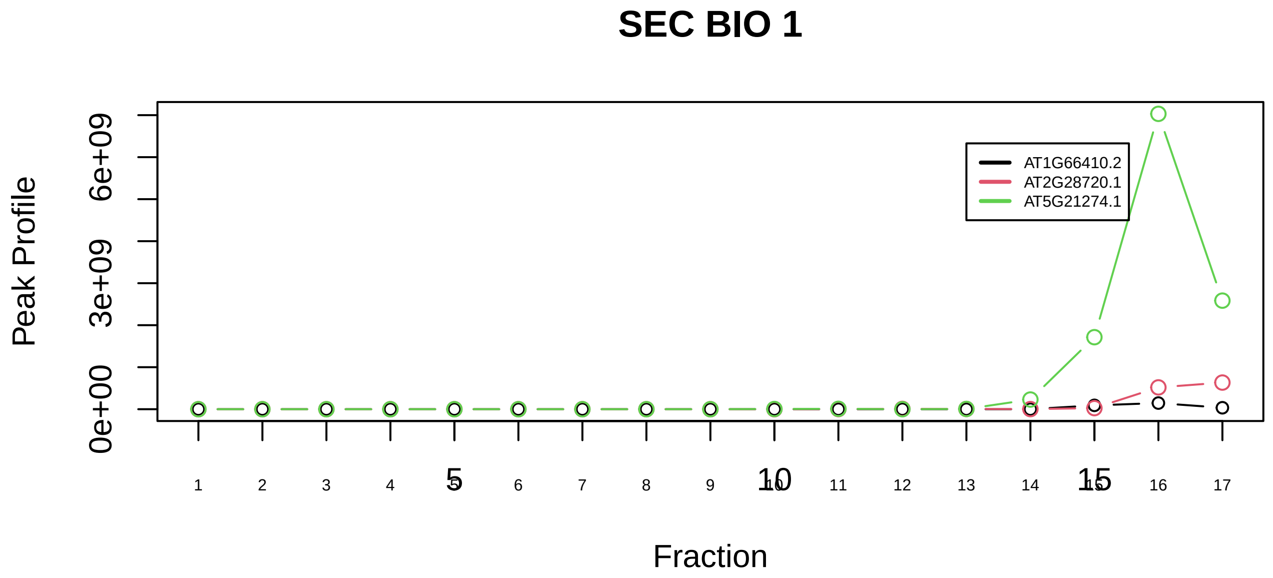

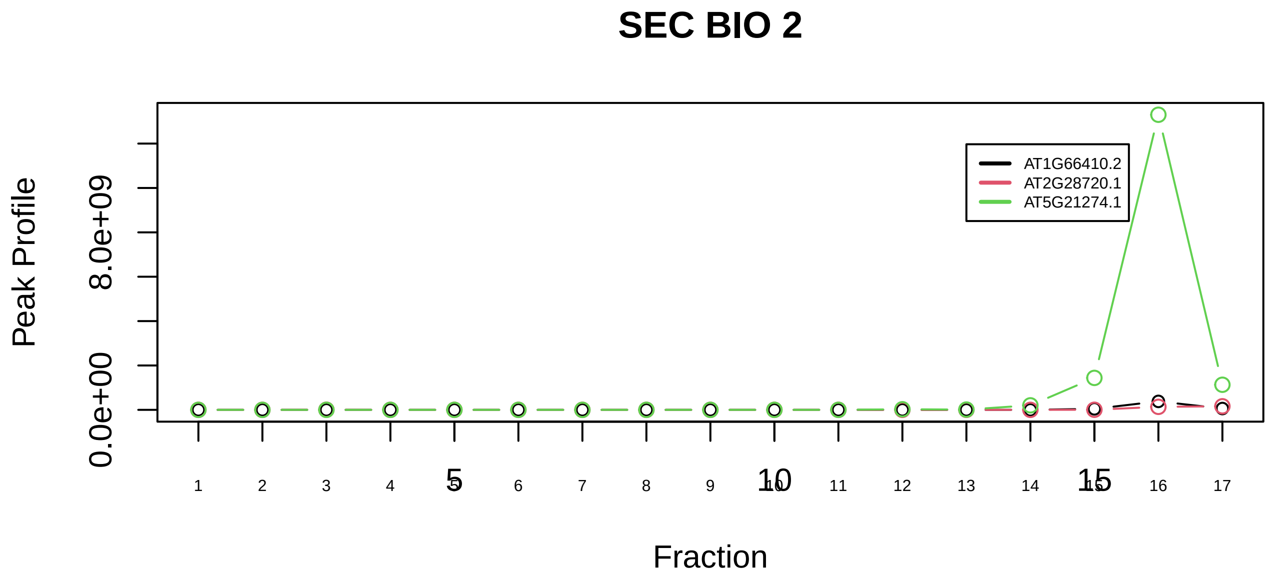

H2AX complex, isolated from cells without IR exposure|1223−−sub_1

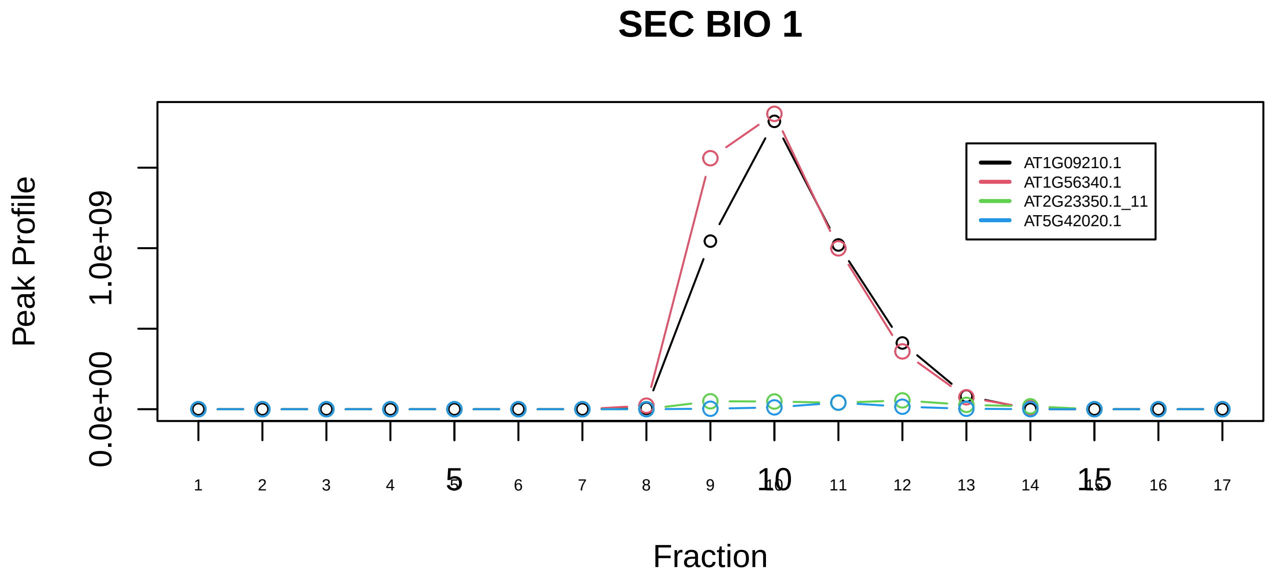

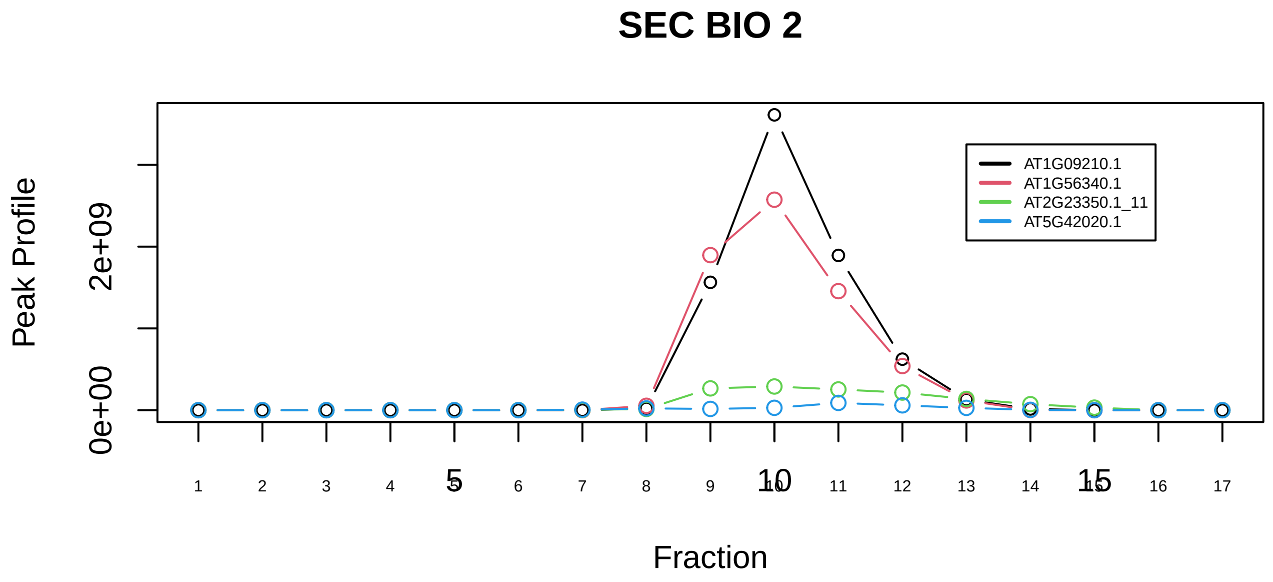

H2AX complex, isolated from cells without IR exposure|1223−−sub_2

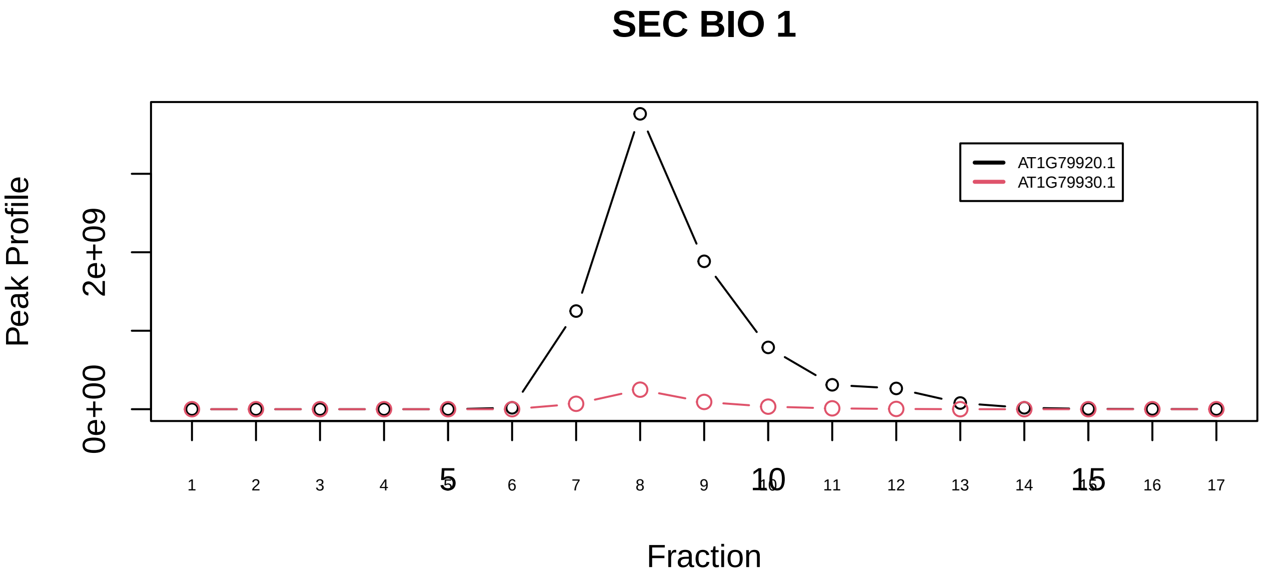

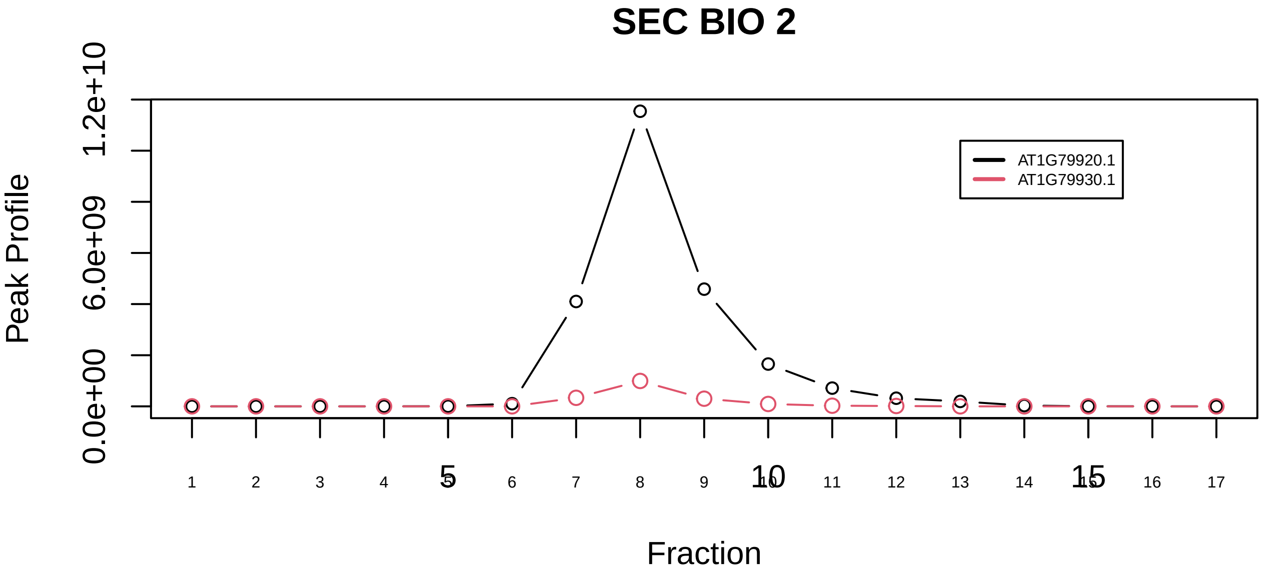

HCF−1 complex|2721−−sub_orthoparalog_1

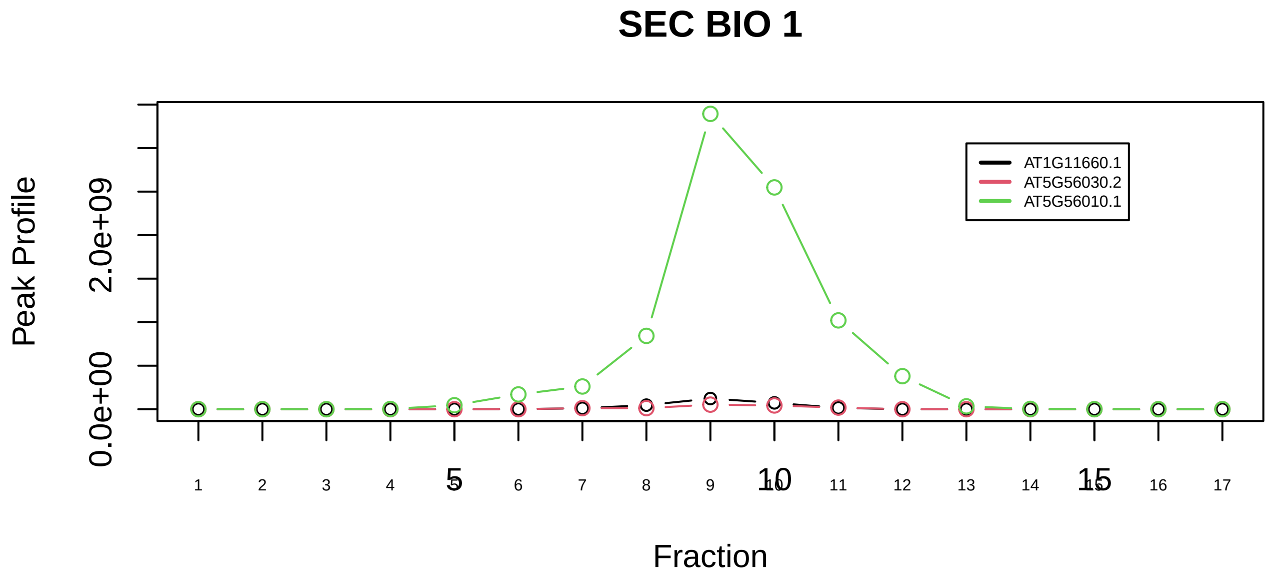

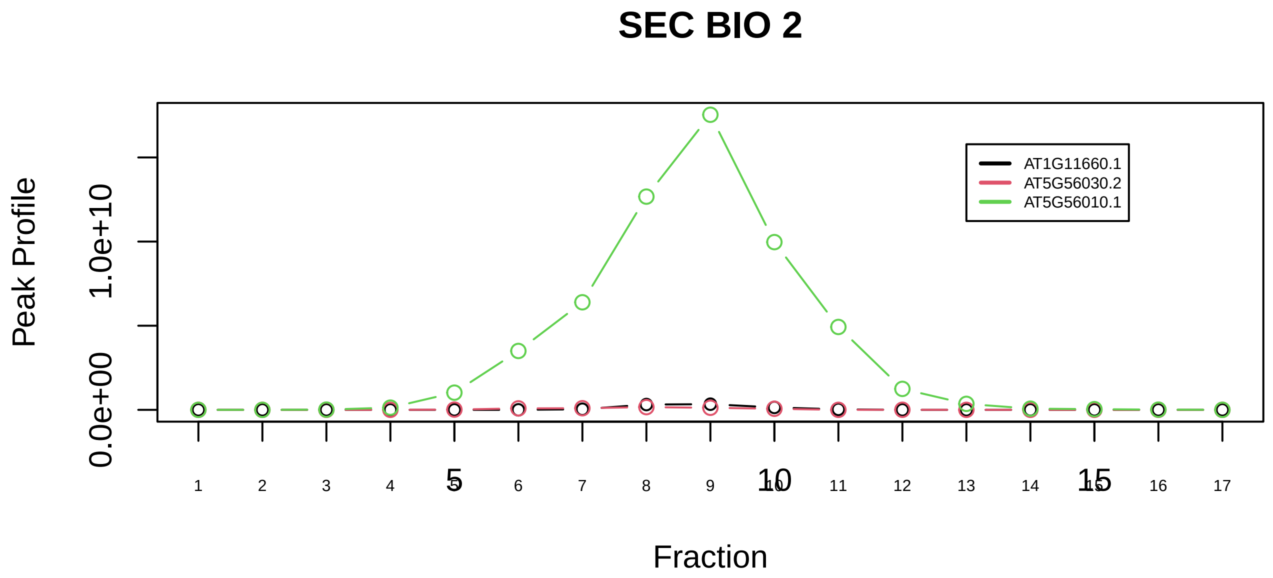

HCF−1 complex|2721−−sub_1

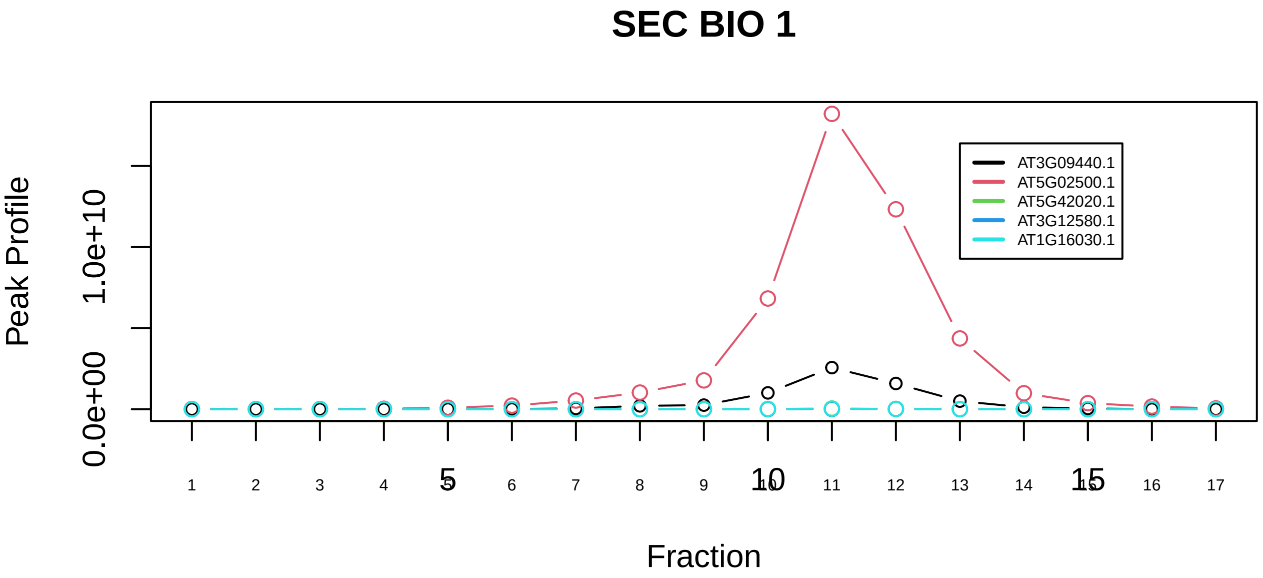

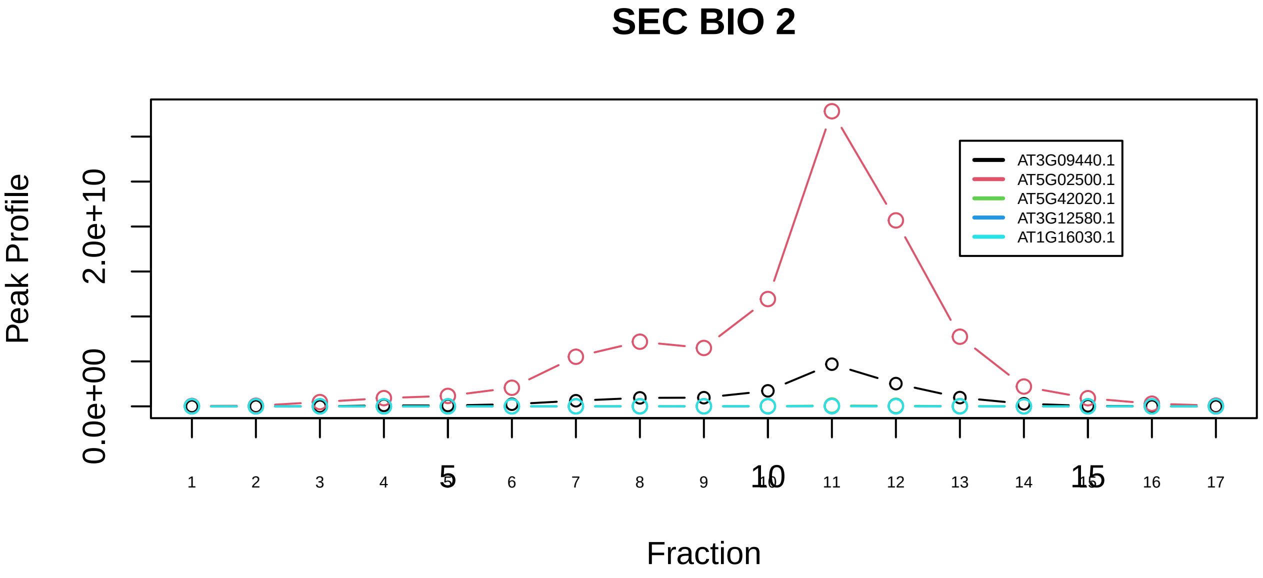

HCF−1 complex|2721−−sub_2

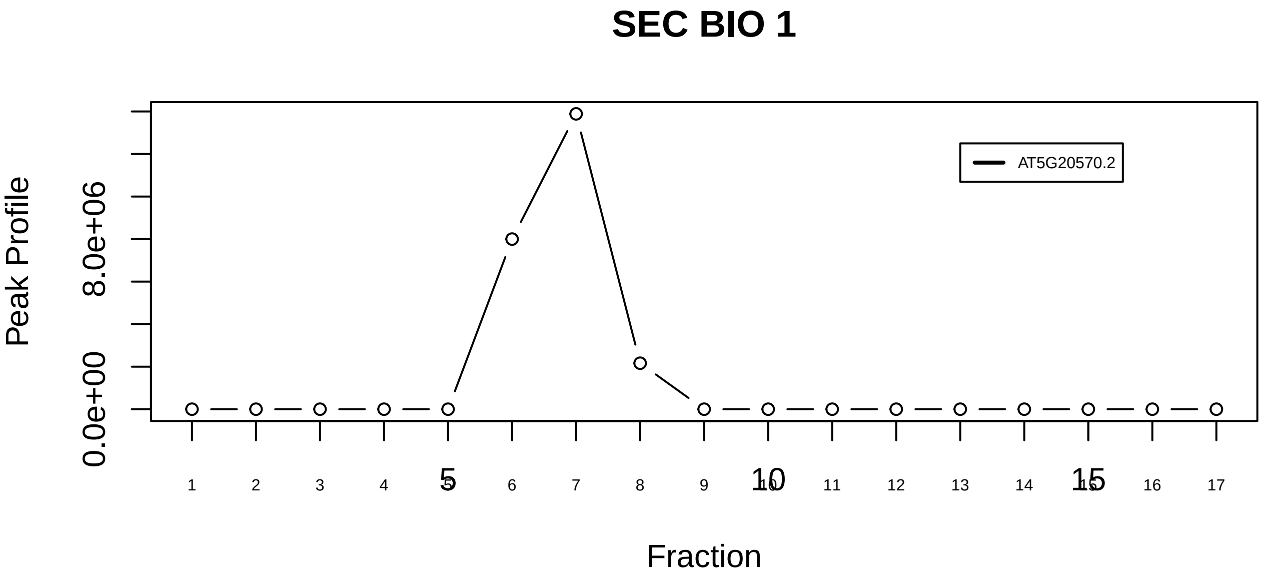

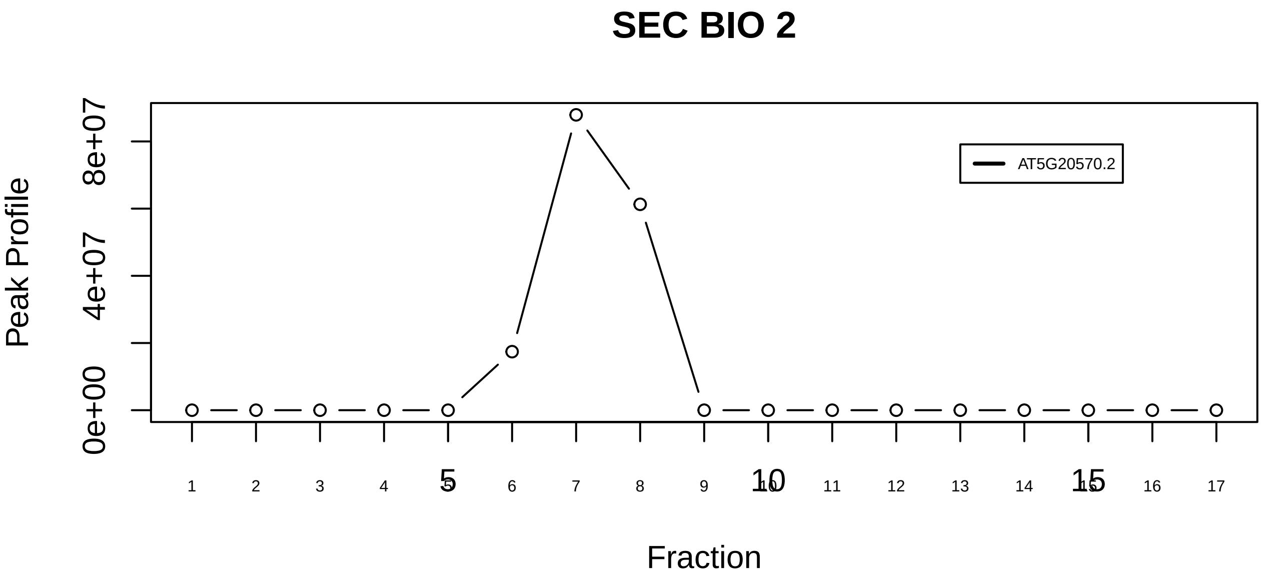

HINT1−UBC3−RBX1 complex|7325−−singleton_1

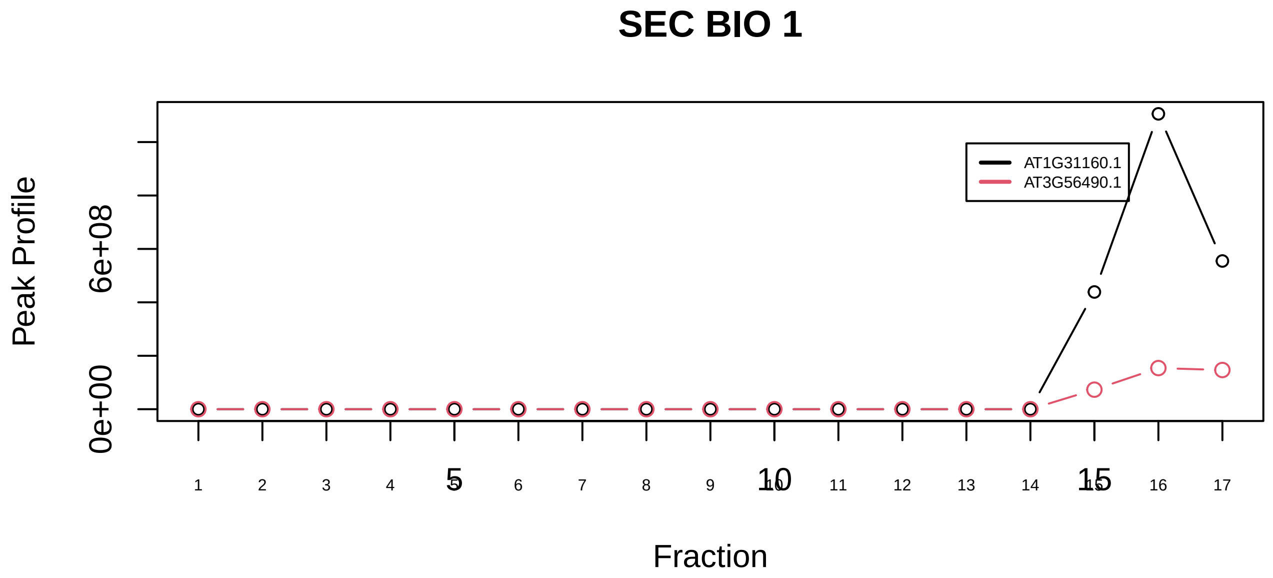

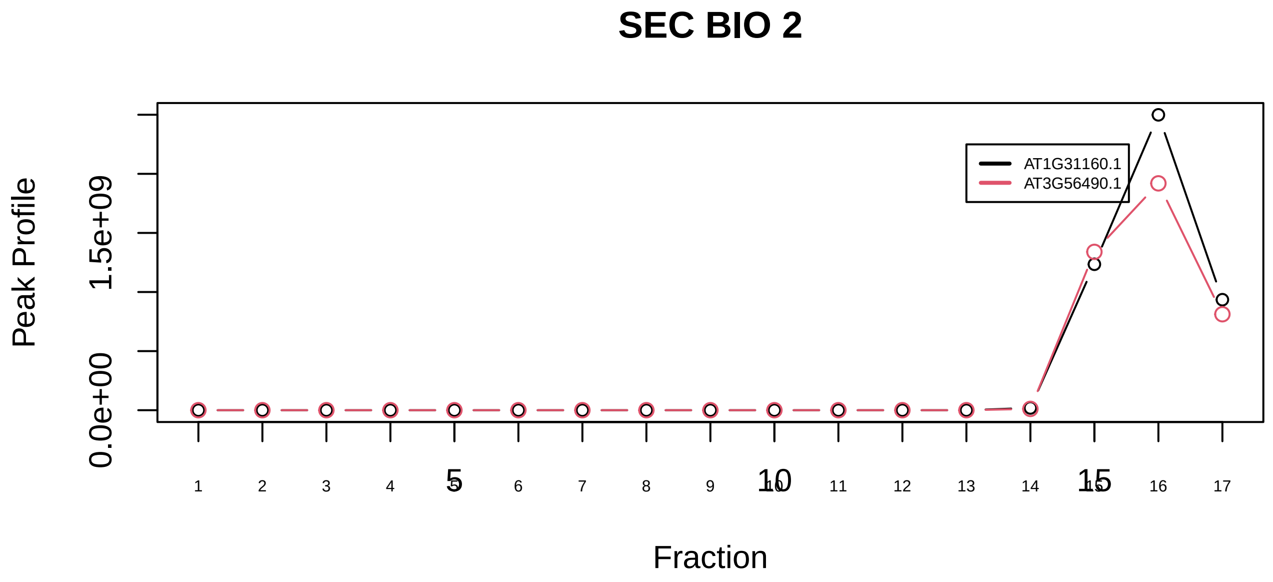

HINT1−UBC3−RBX1 complex|7325−−sub_orthoparalog_1

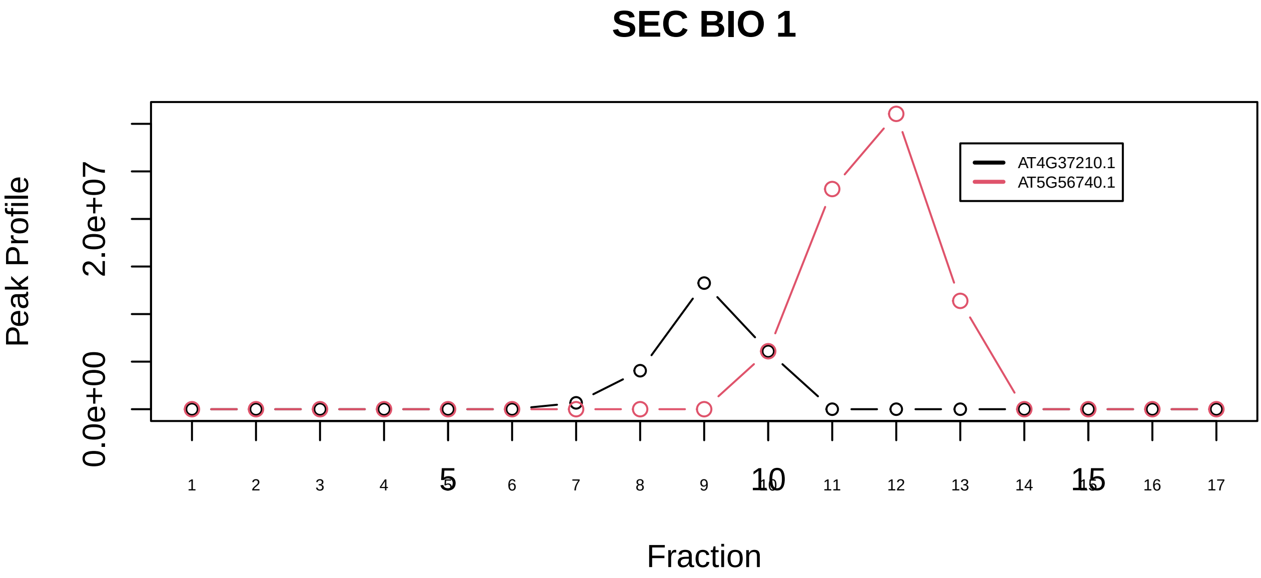

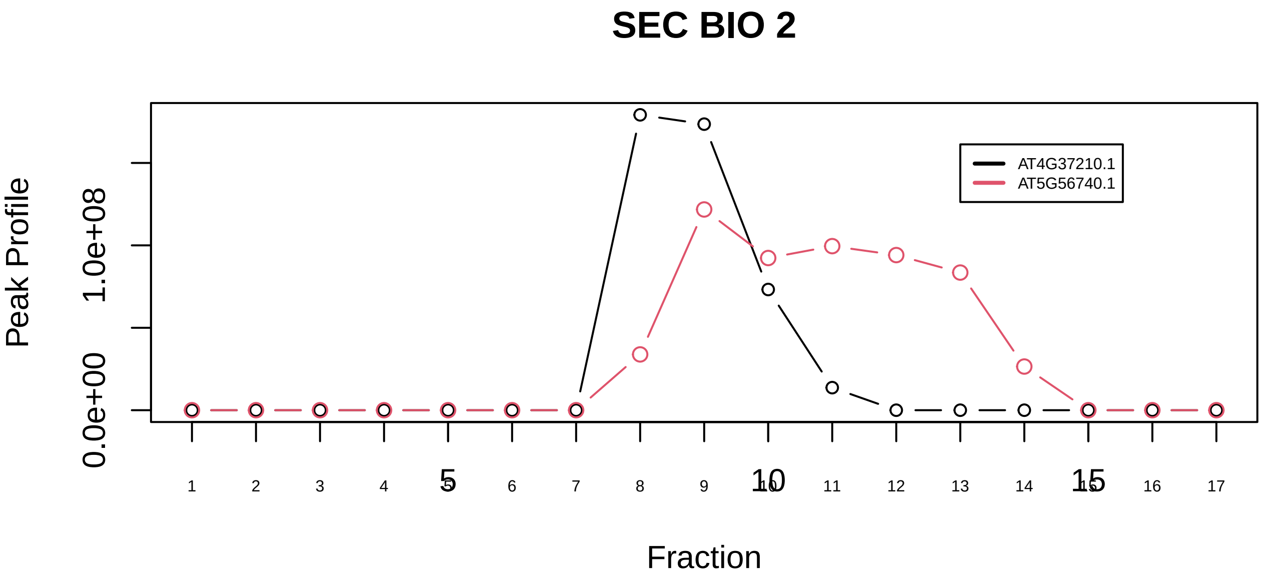

Histone H3.3 complex|1150−−only_2_prots_before_clustering

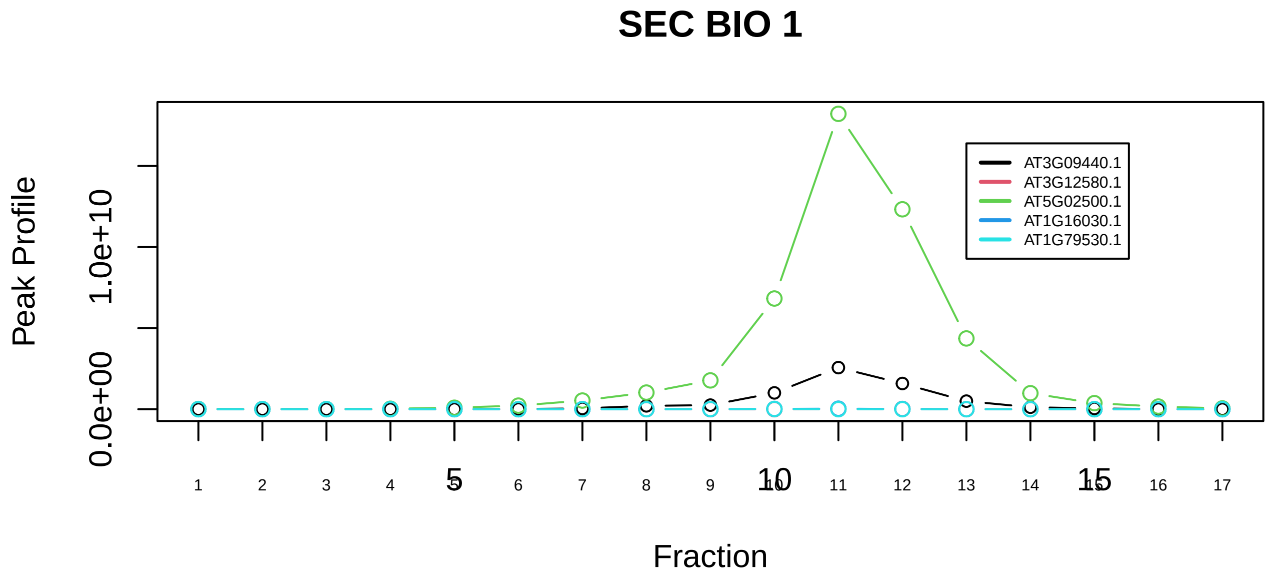

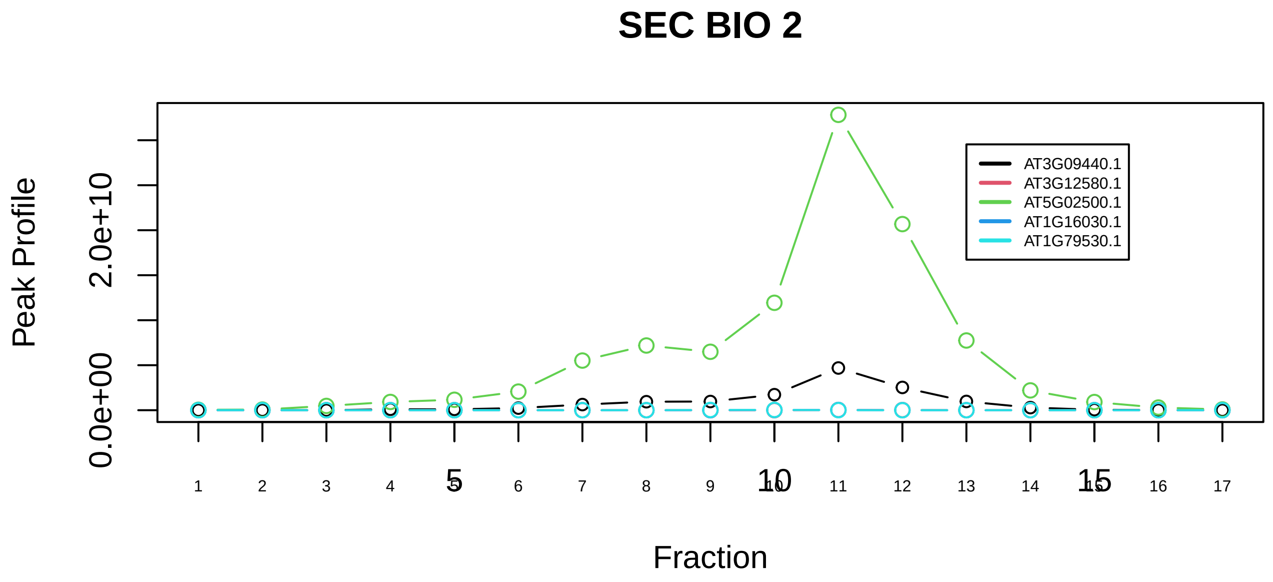

HMGB1−HMGB2−HSC70−ERP60−GAPDH complex|280−−sub_1

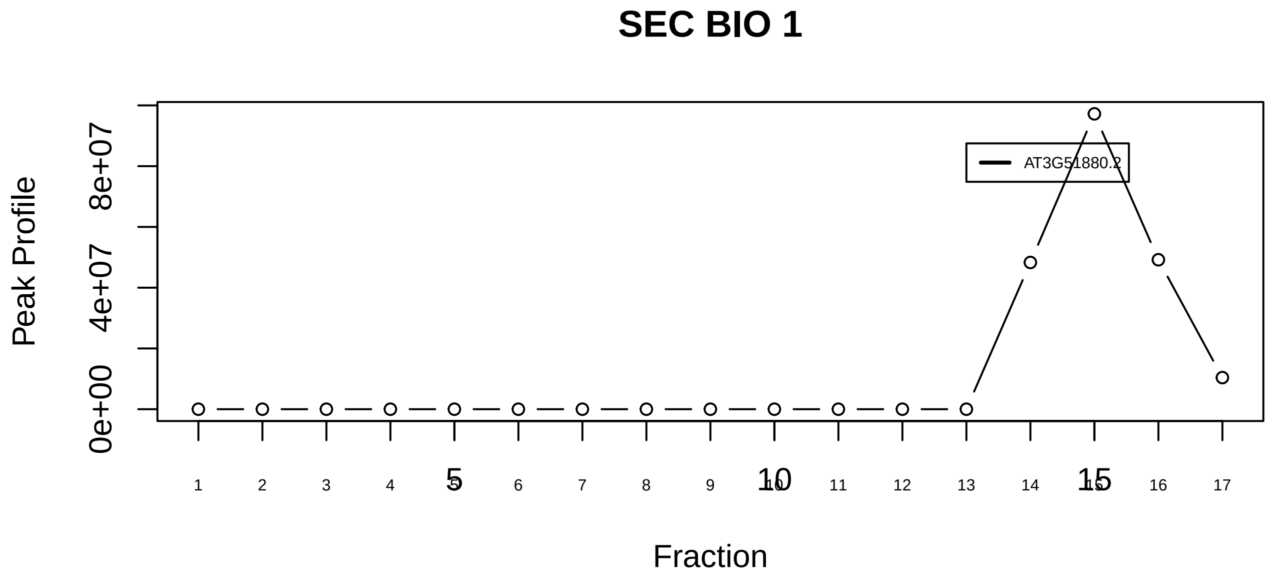

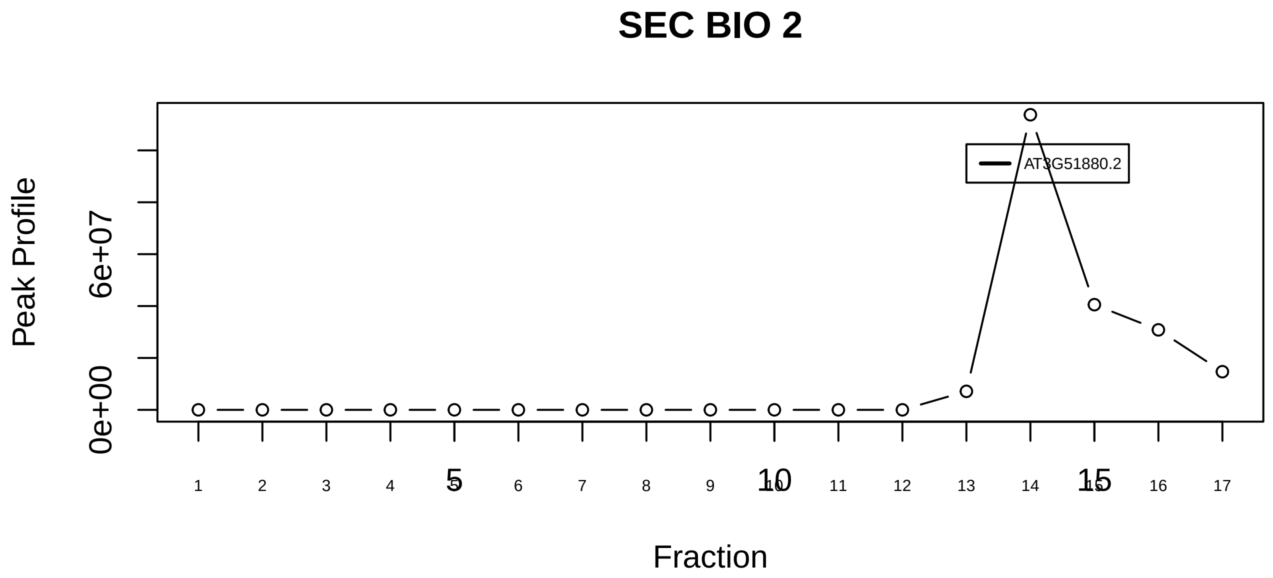

HMGB1−HMGB2−HSC70−ERP60−GAPDH complex|280−−singleton_1

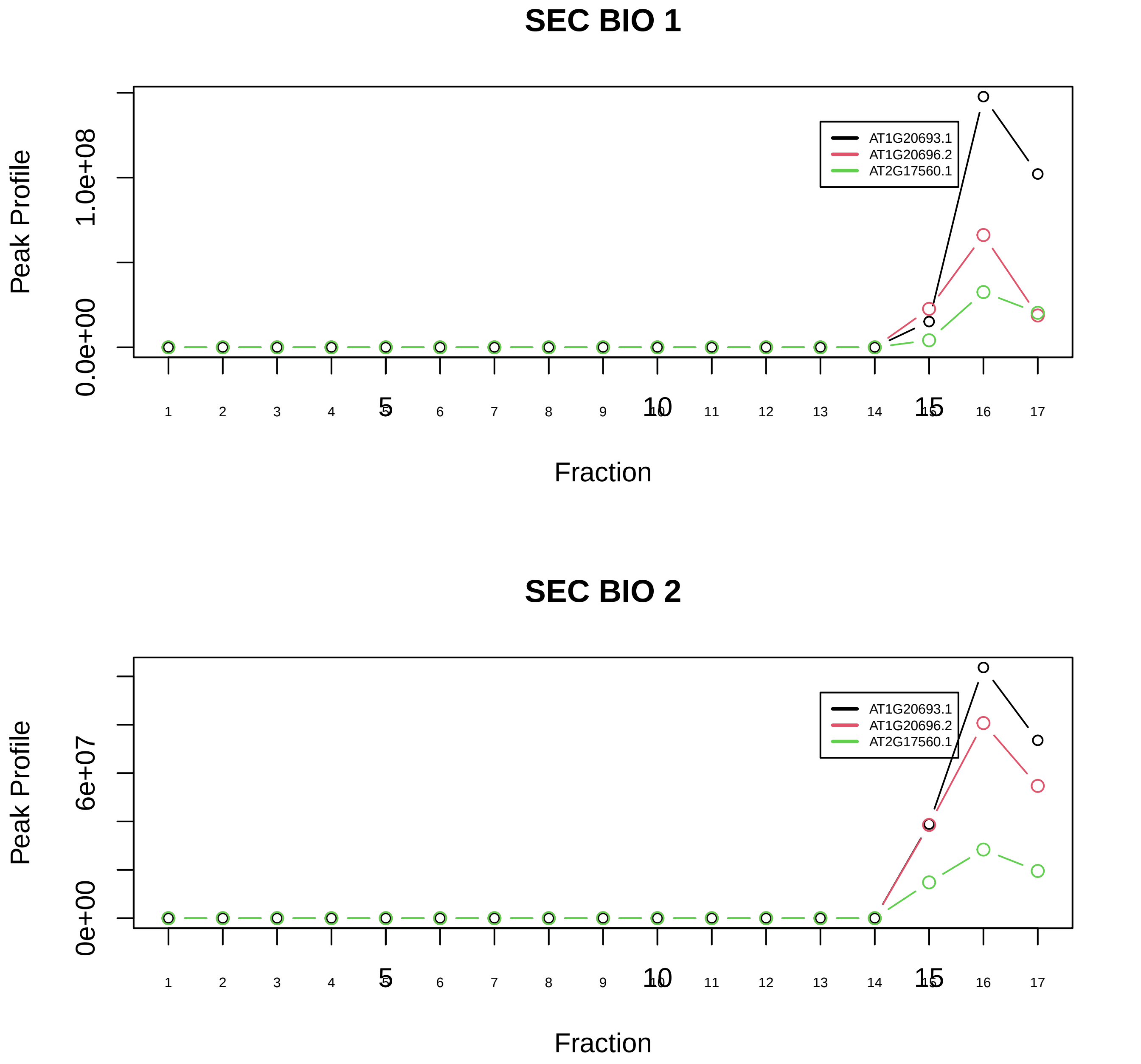

HMGB1−HMGB2−HSC70−ERP60−GAPDH complex|280−−sub_2

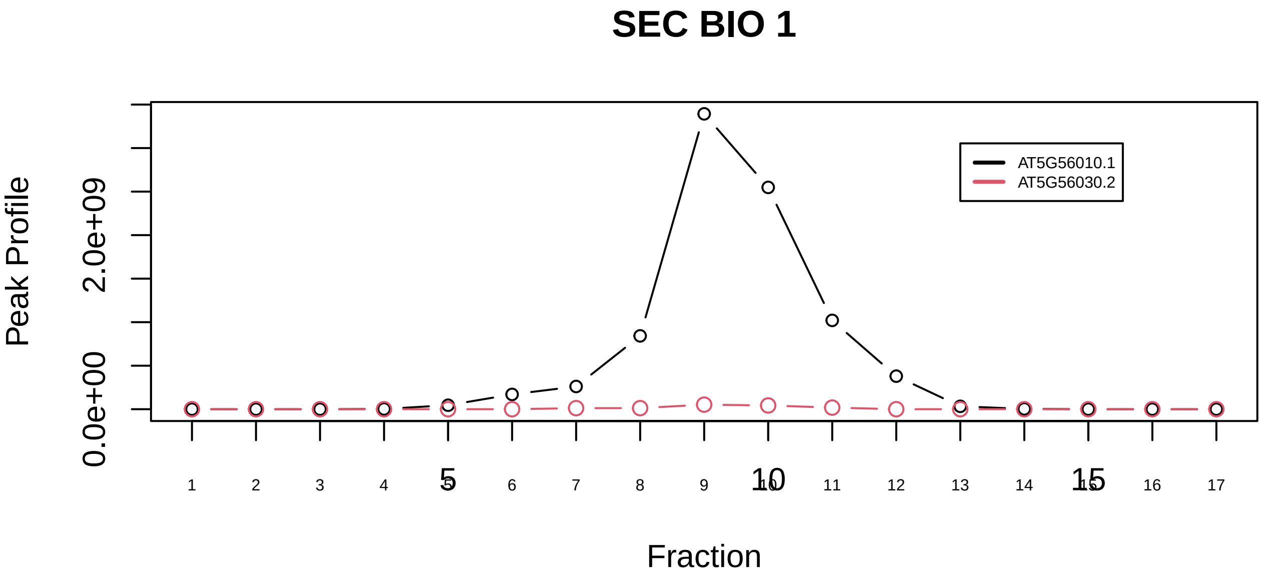

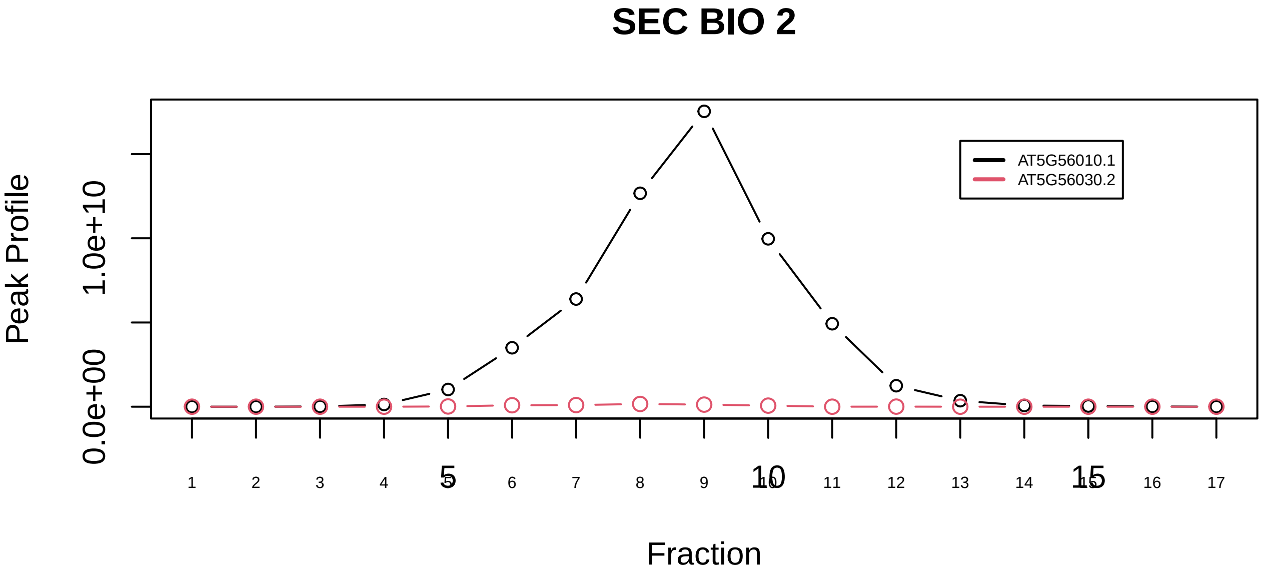

HSP90−FKBP38−CAM−Ca(2+) complex|4158−−sub_orthoparalog_1

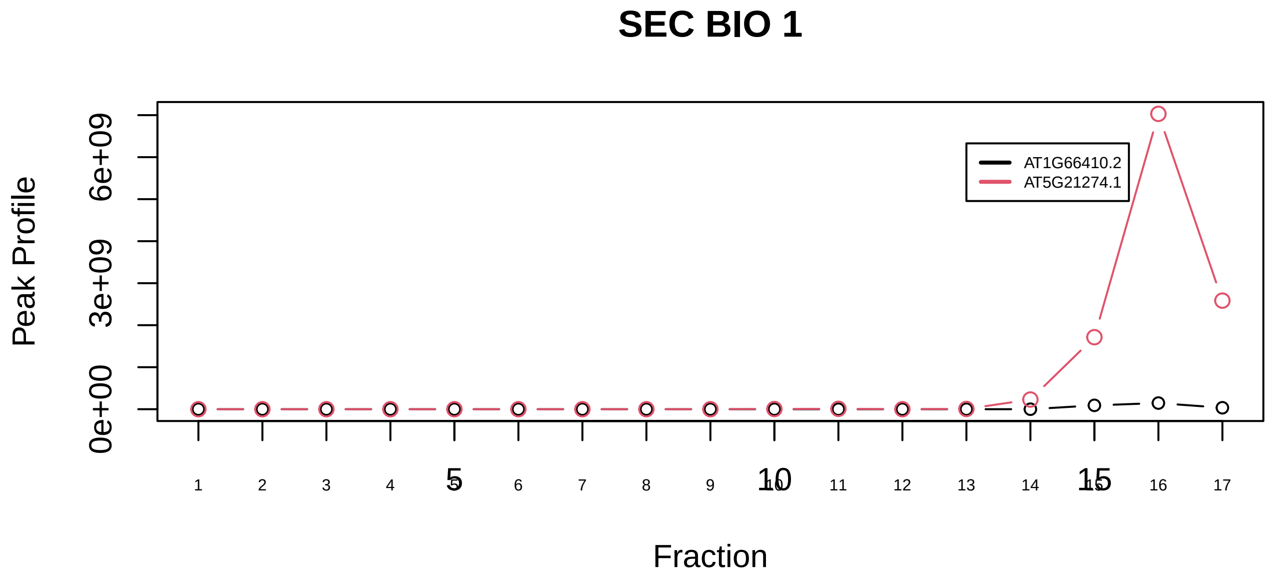

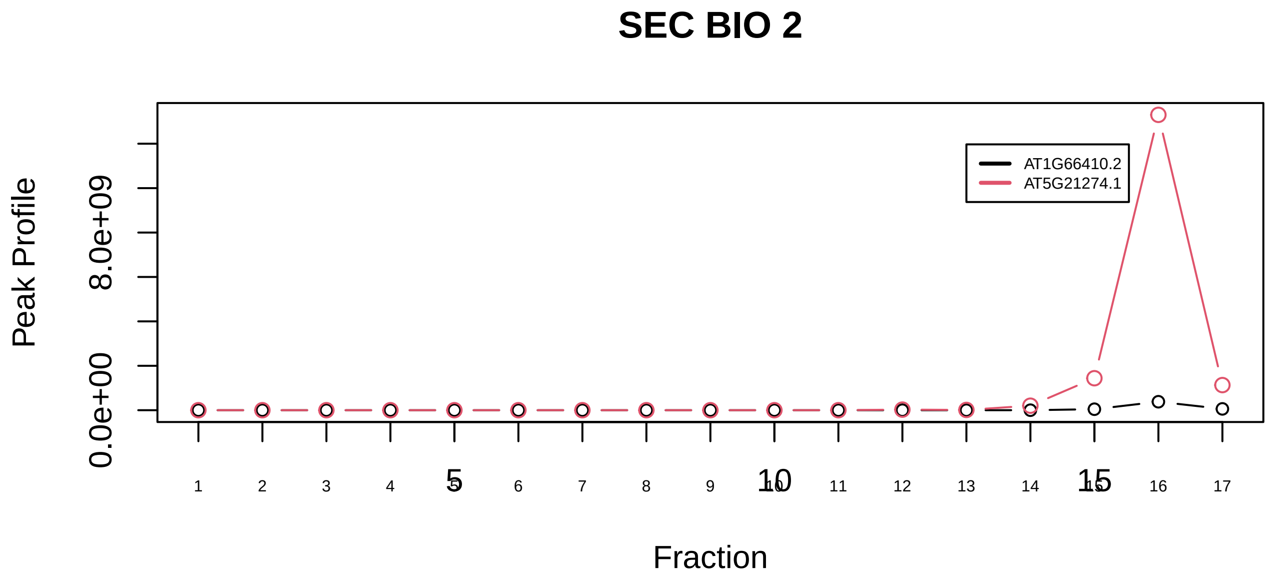

HSP90−FKBP38−CAM−Ca(2+) complex|4158−−sub_orthoparalog_2

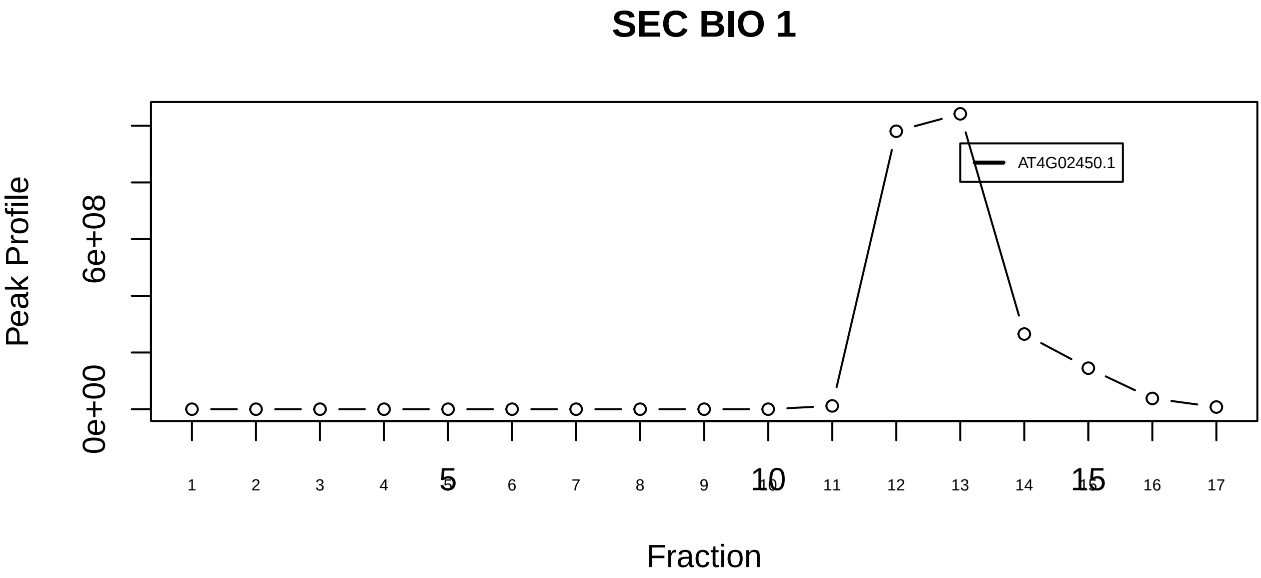

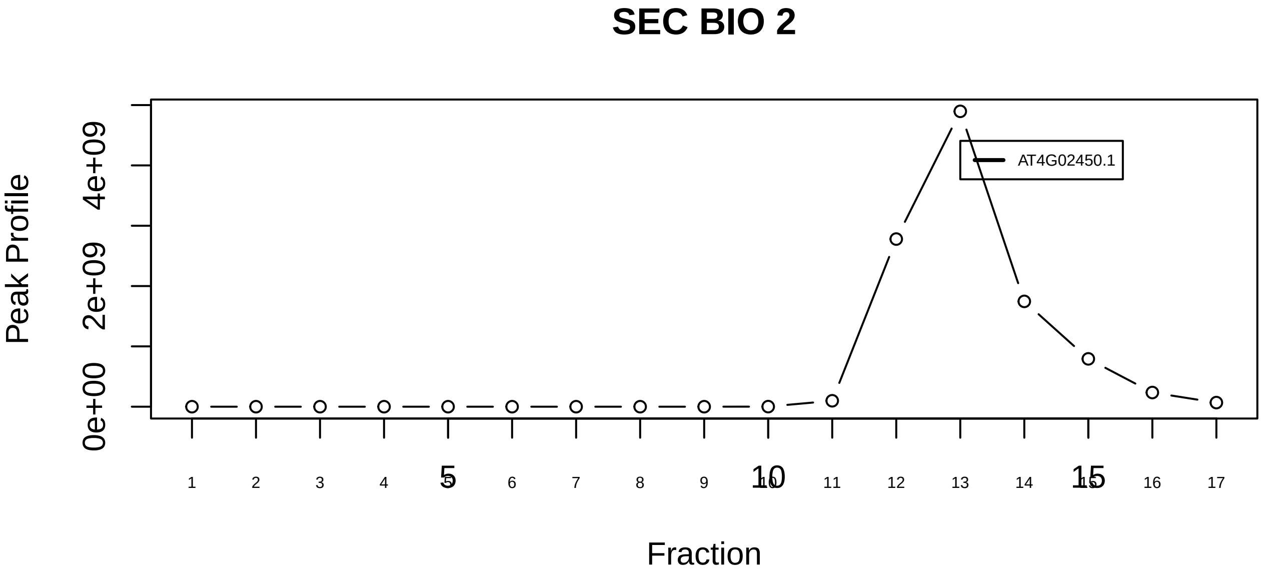

Hsp90−p23 complex|6591−−singleton_1

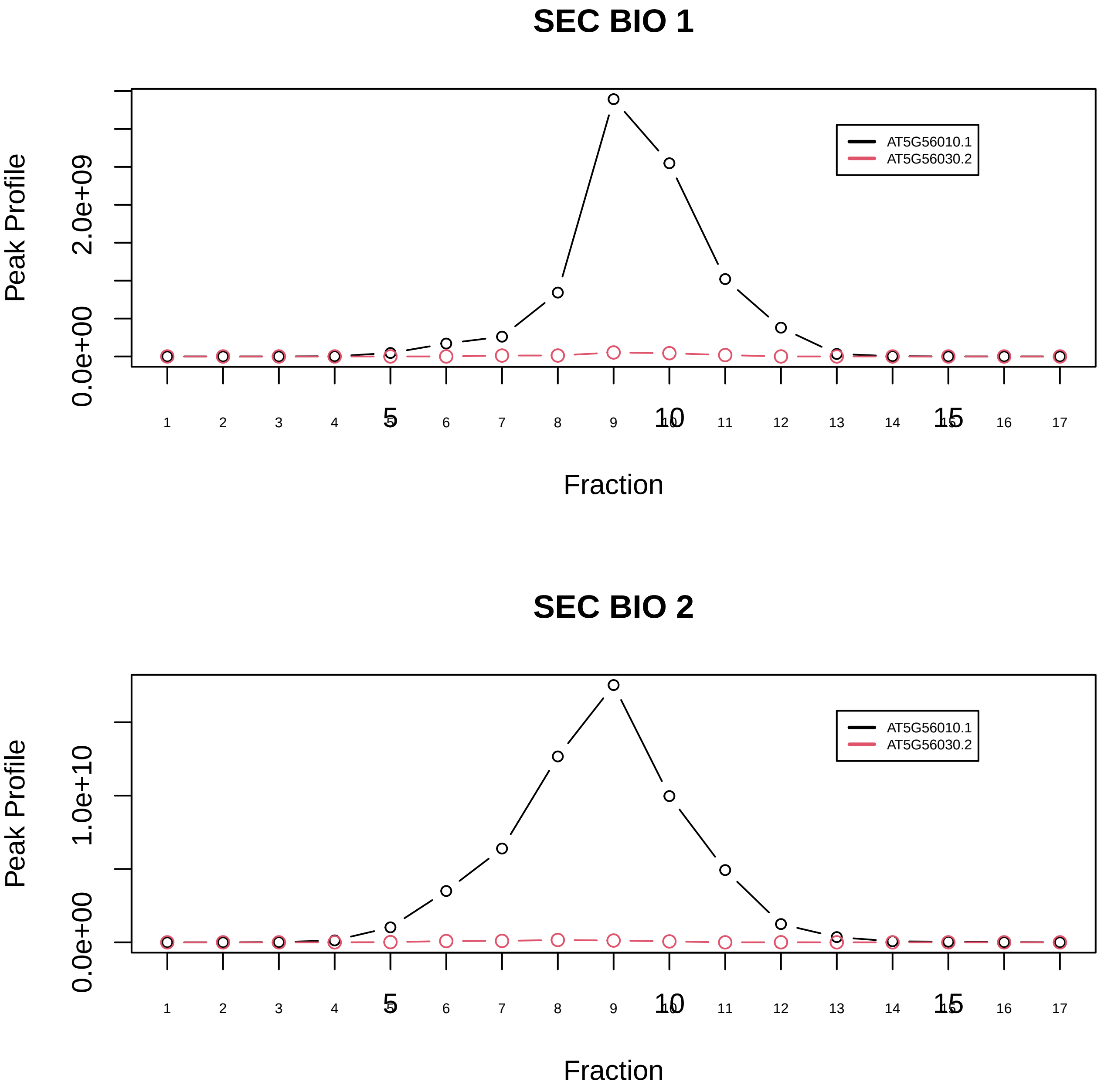

Hsp90−p23 complex|6591−−sub_orthoparalog_1

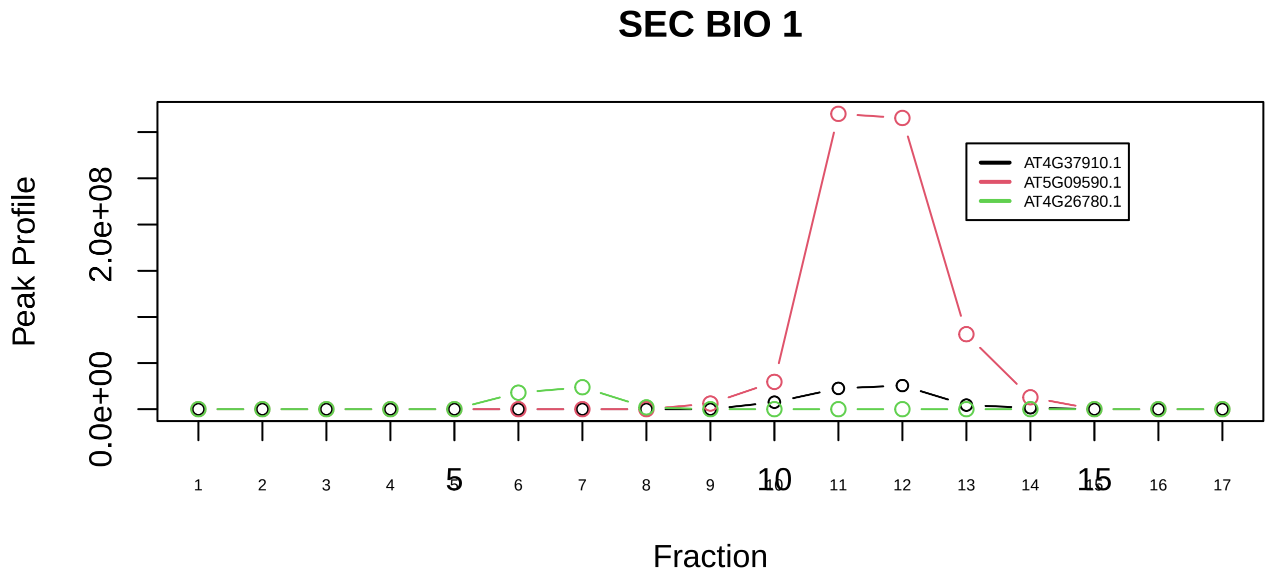

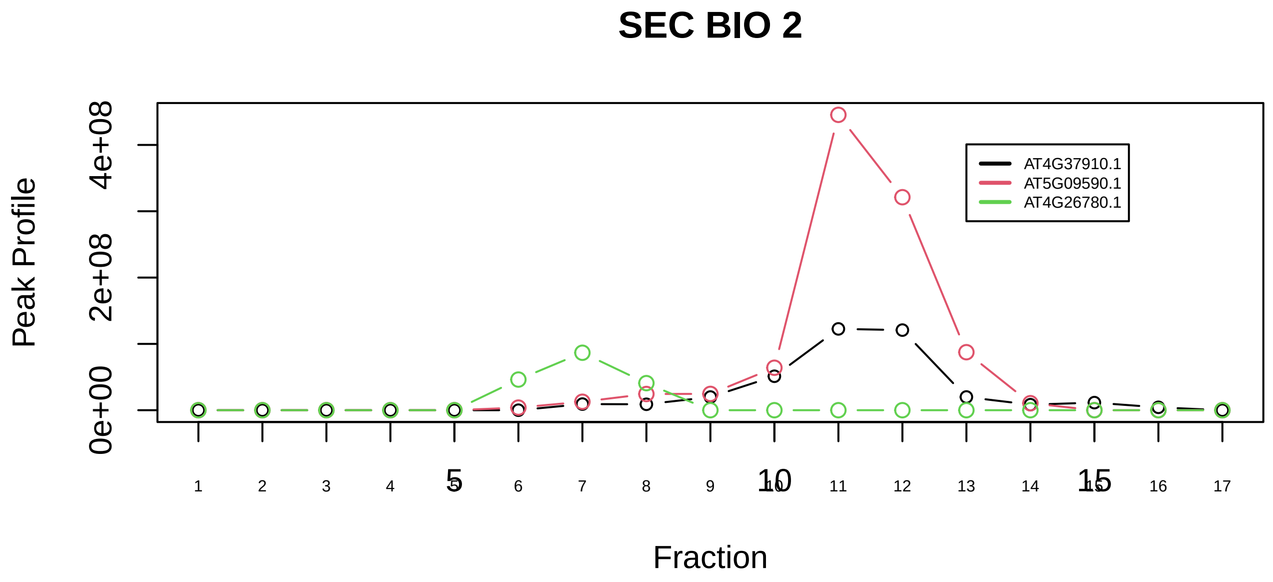

HSPA9−GRPEL2 complex|7125−−sub_1

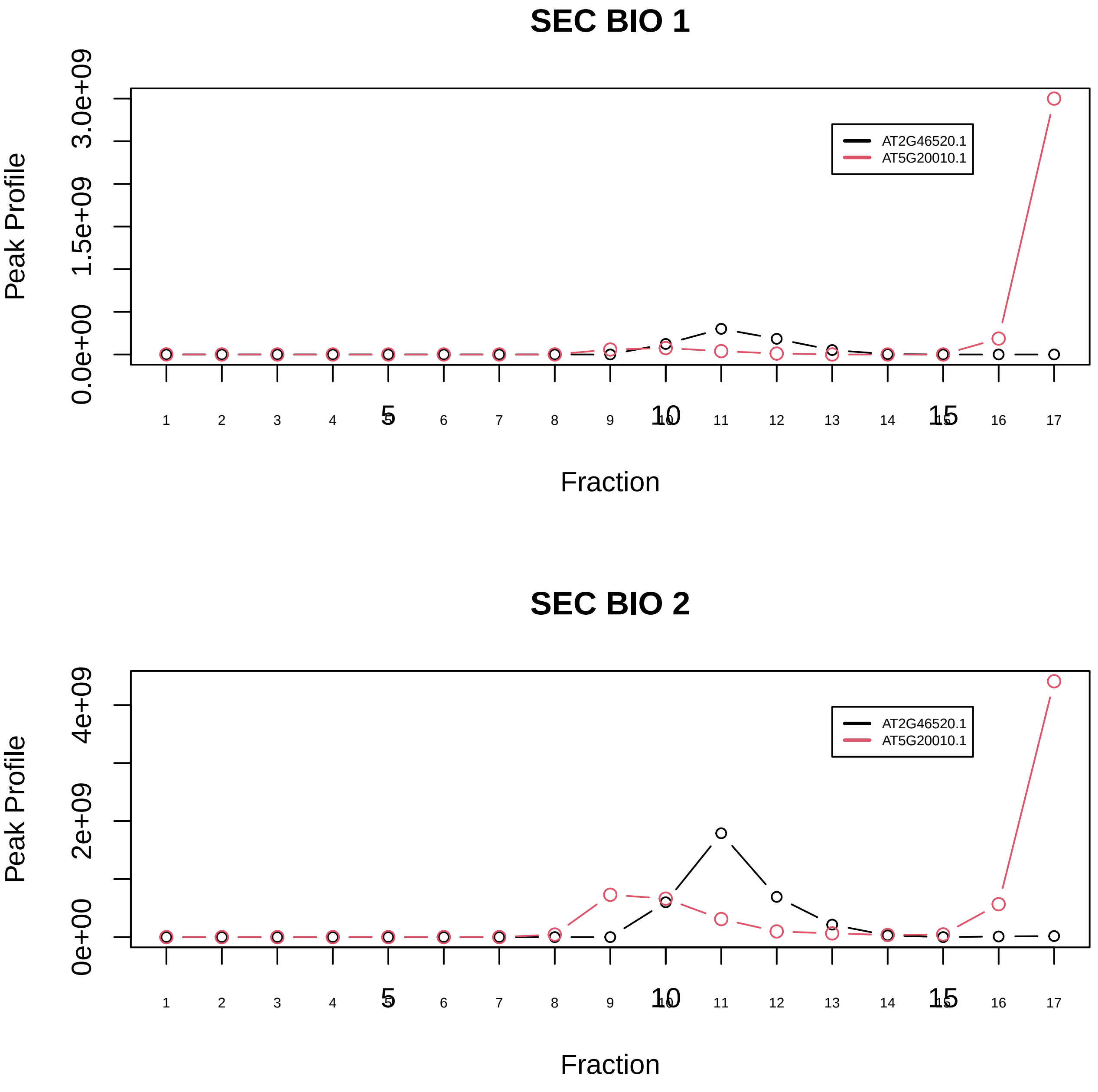

Importin alpha−CAS−RanGTP complex|6141−−only_2_prots_before_clustering

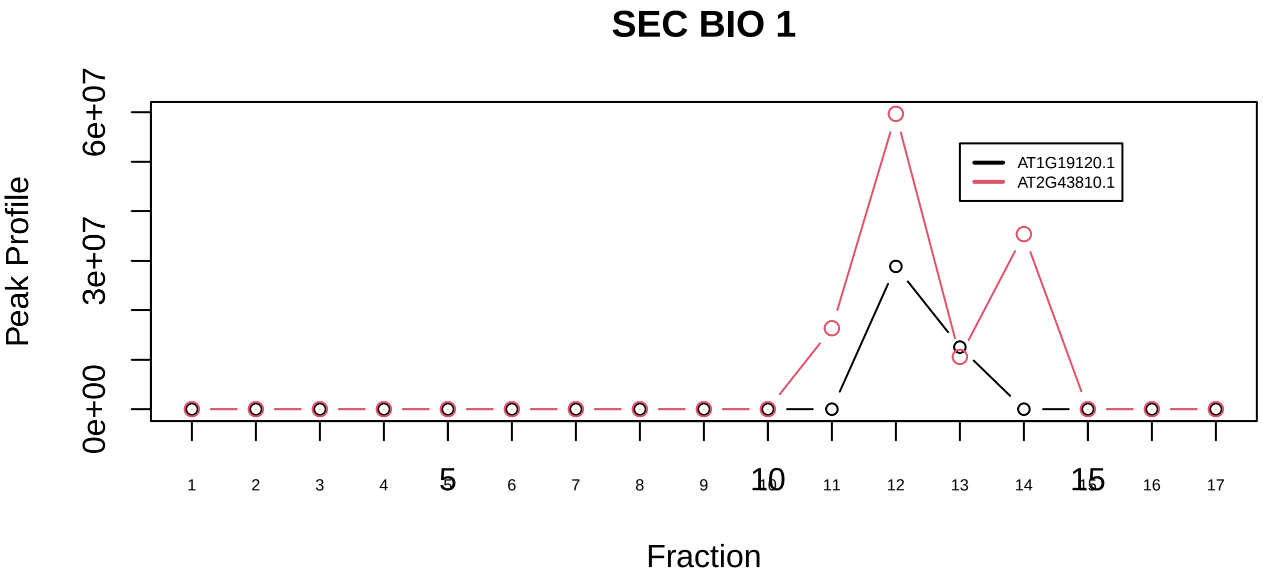

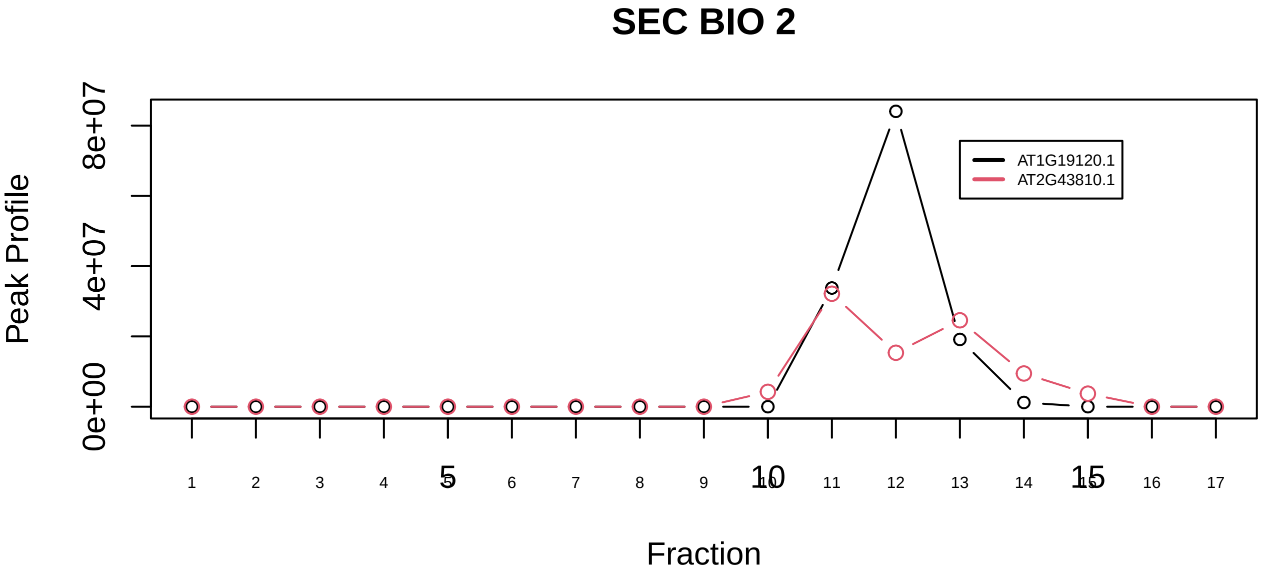

LSm1−7 complex|561−−sub_1

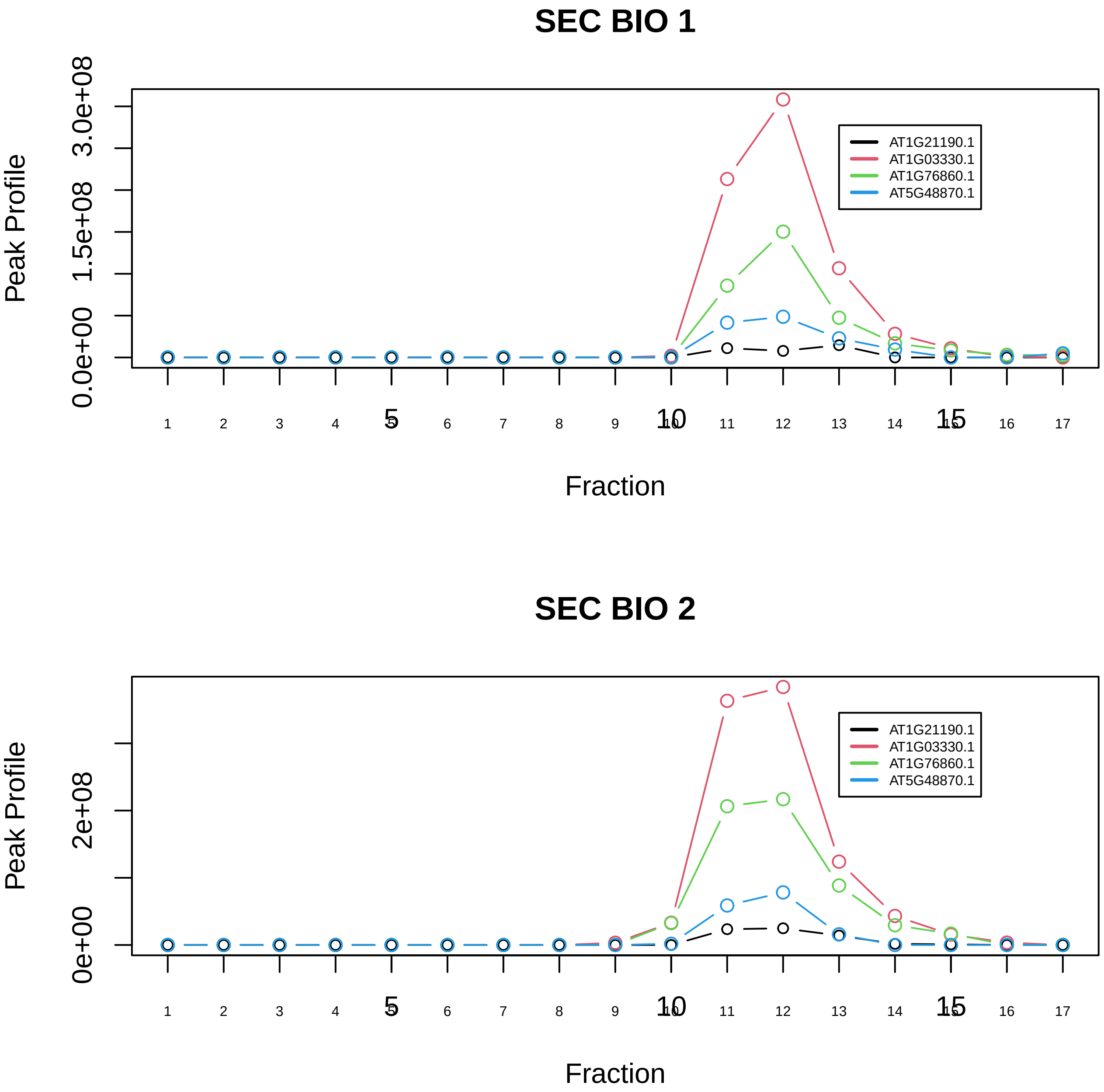

LSm1−7 complex|561−−sub_2

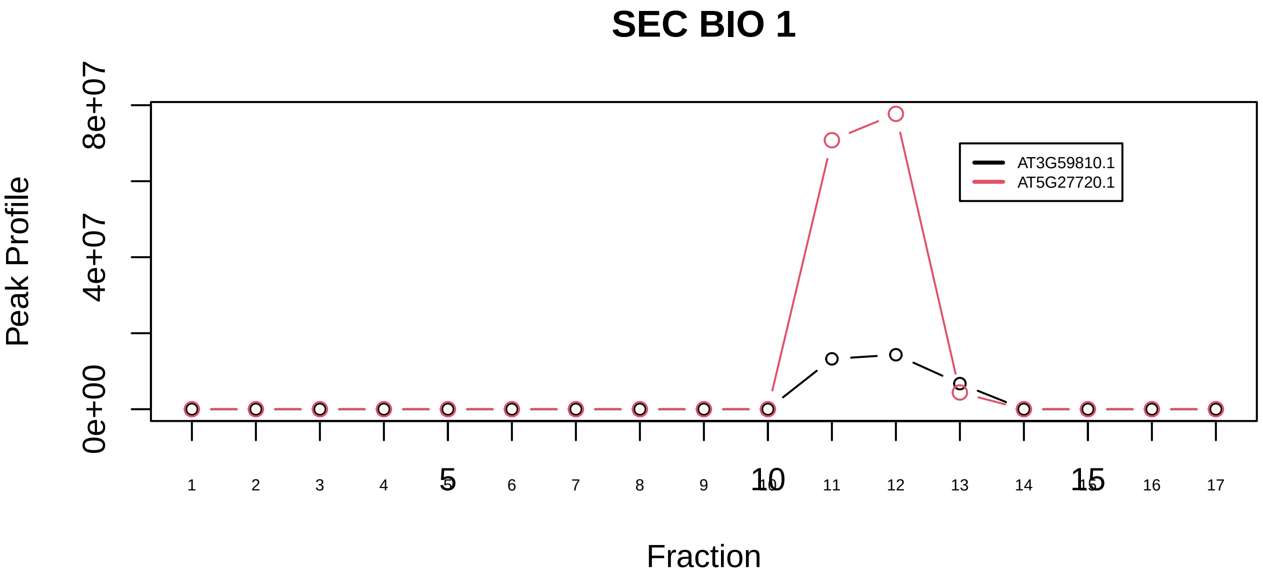

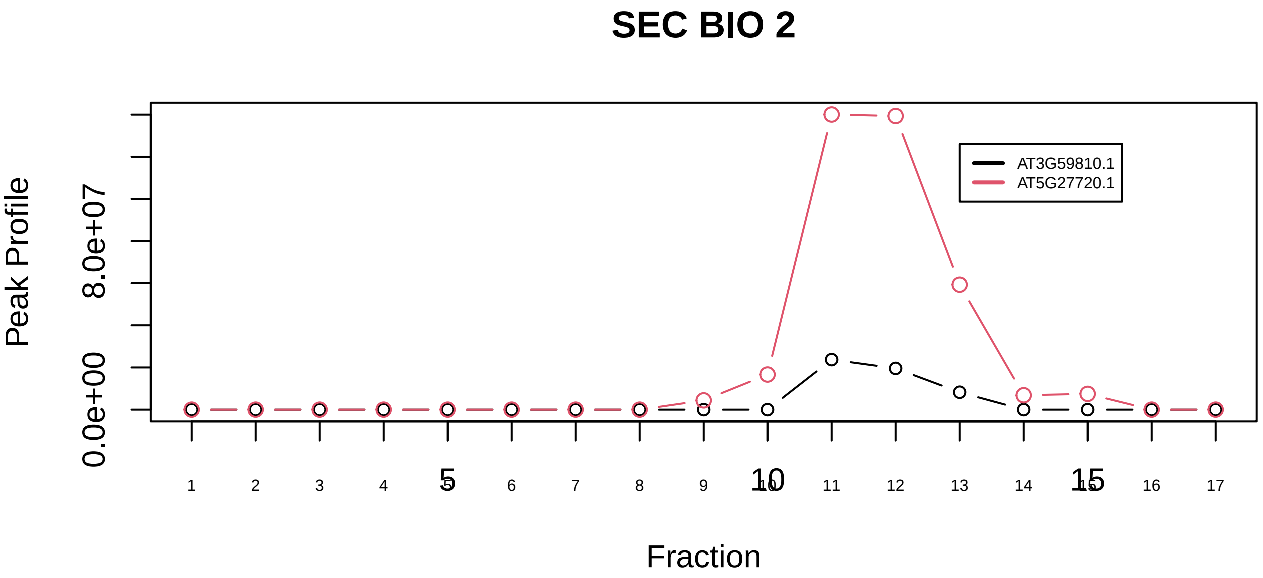

LSm1−7 complex|561−−sub_3

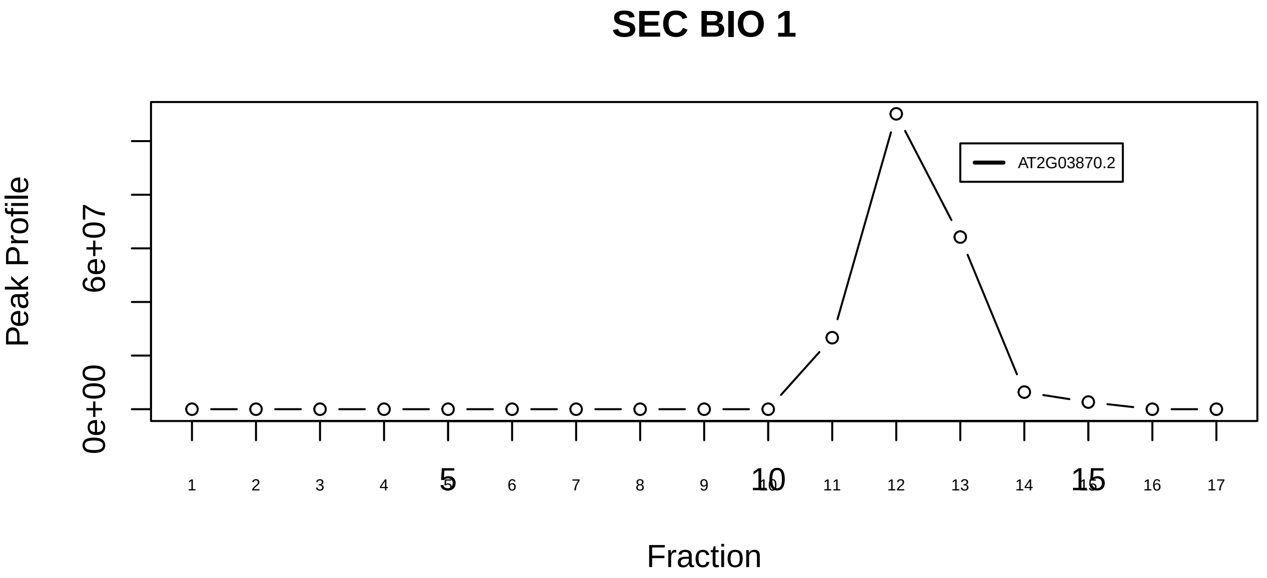

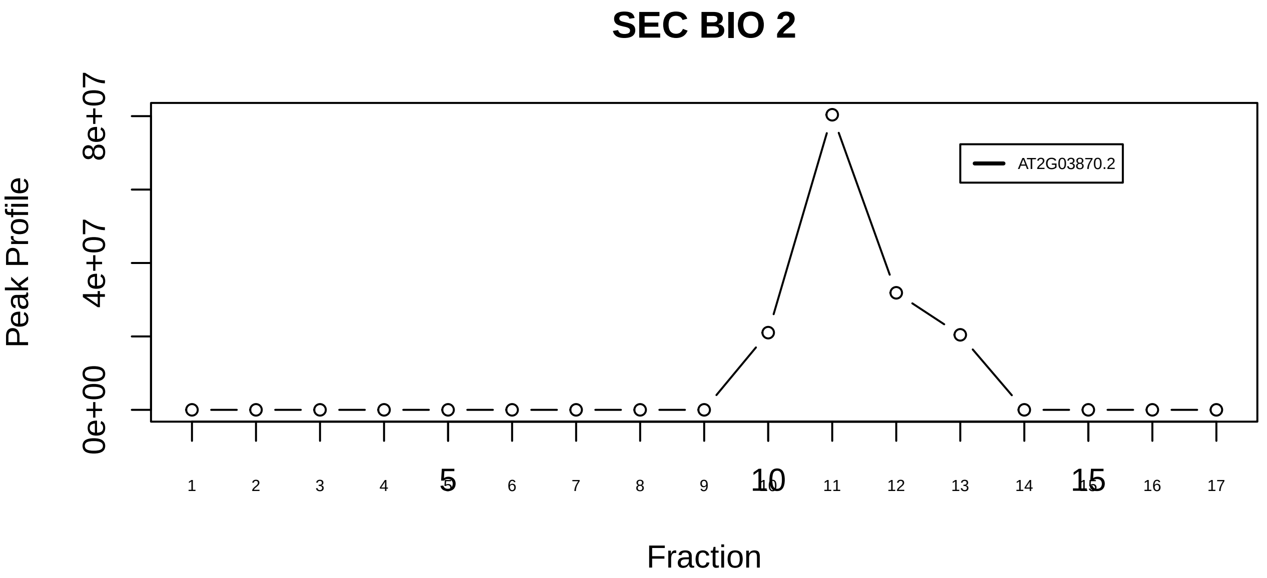

LSm1−7 complex|561−−singleton_1

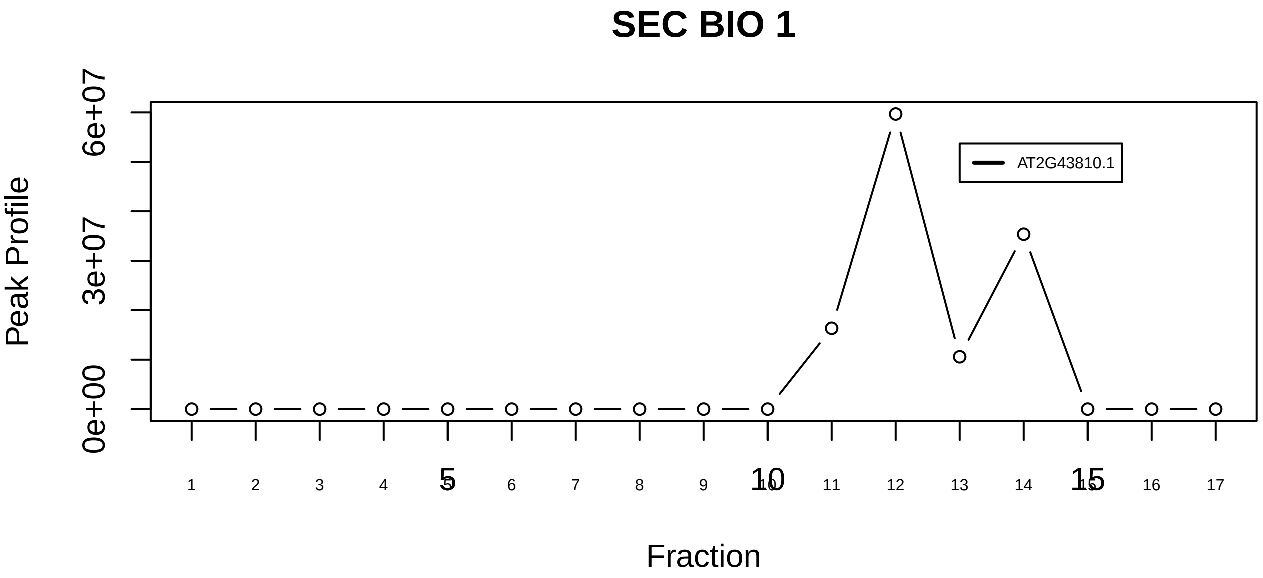

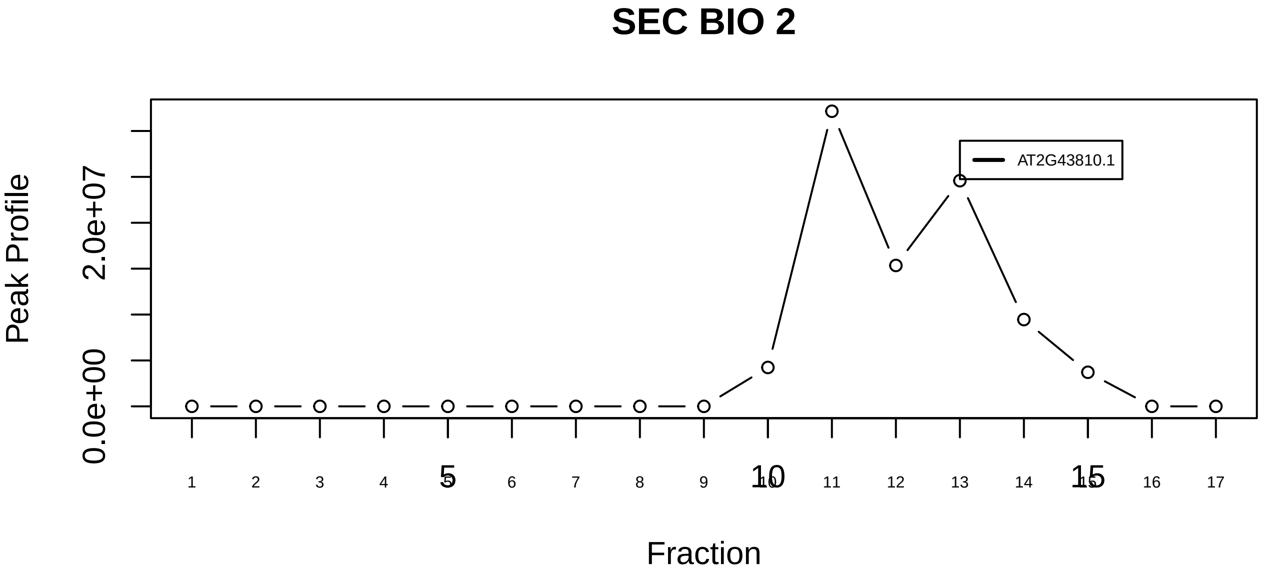

LSm2−8 complex|562−−singleton_1

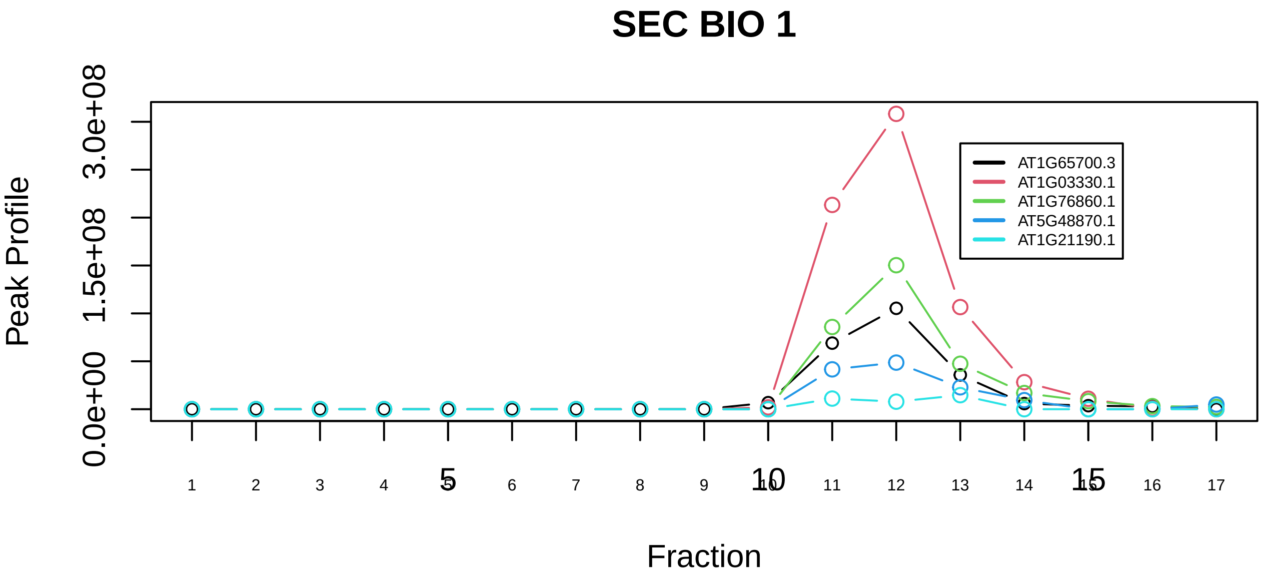

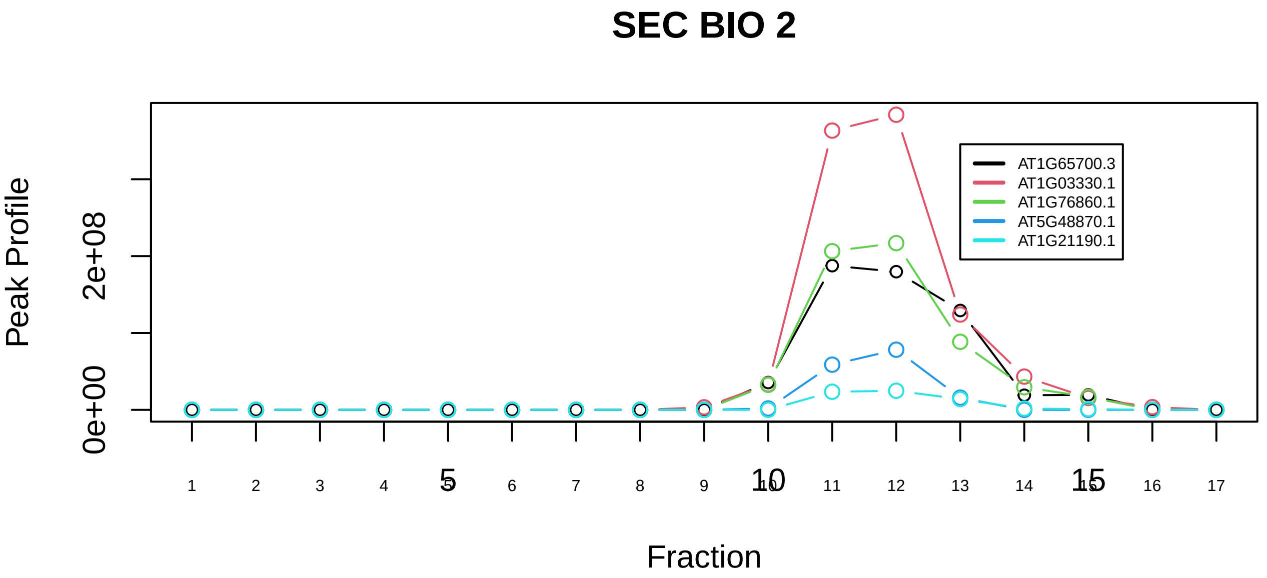

LSm2−8 complex|562−−sub_1

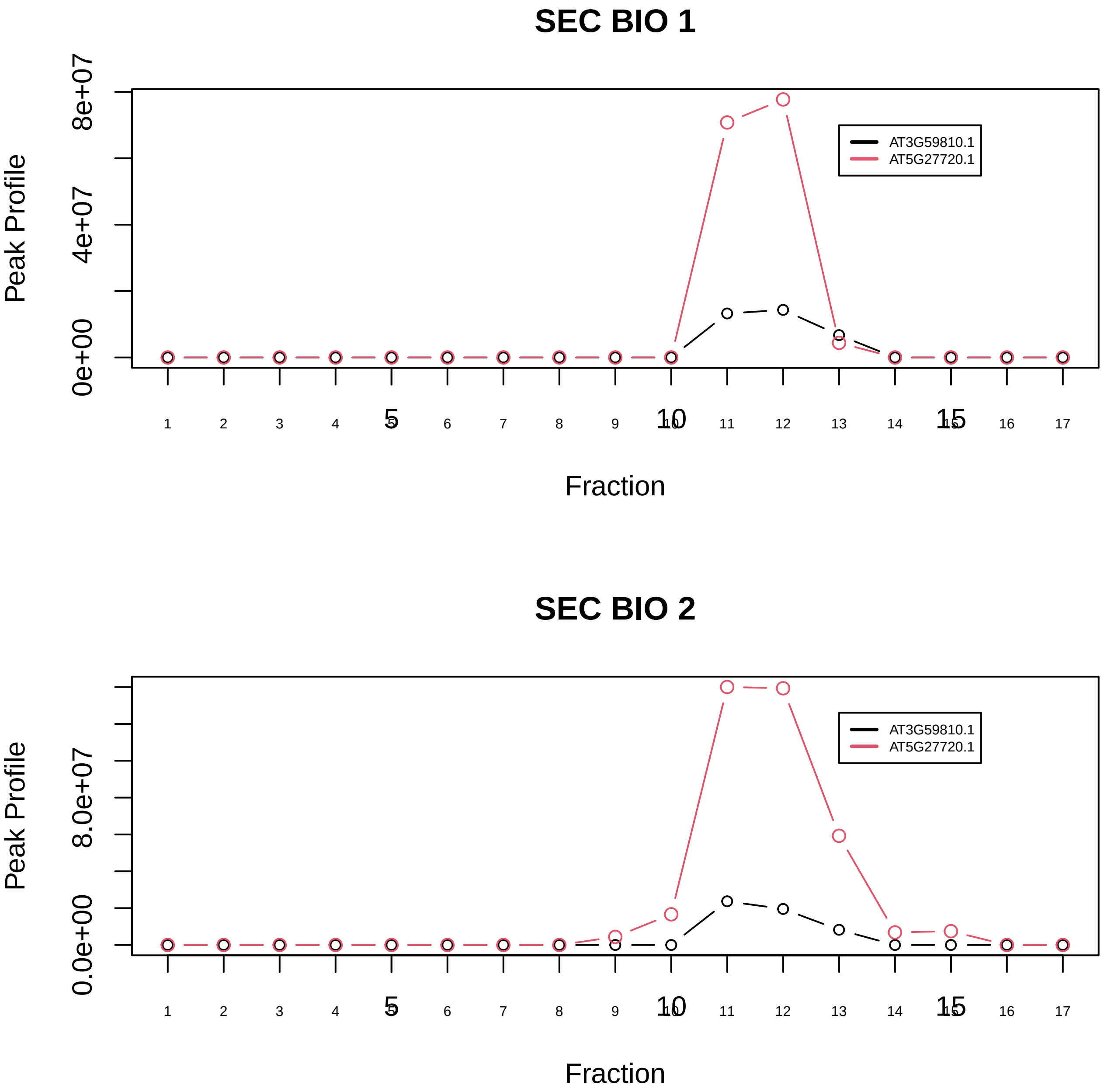

LSm2−8 complex|562−−sub_2

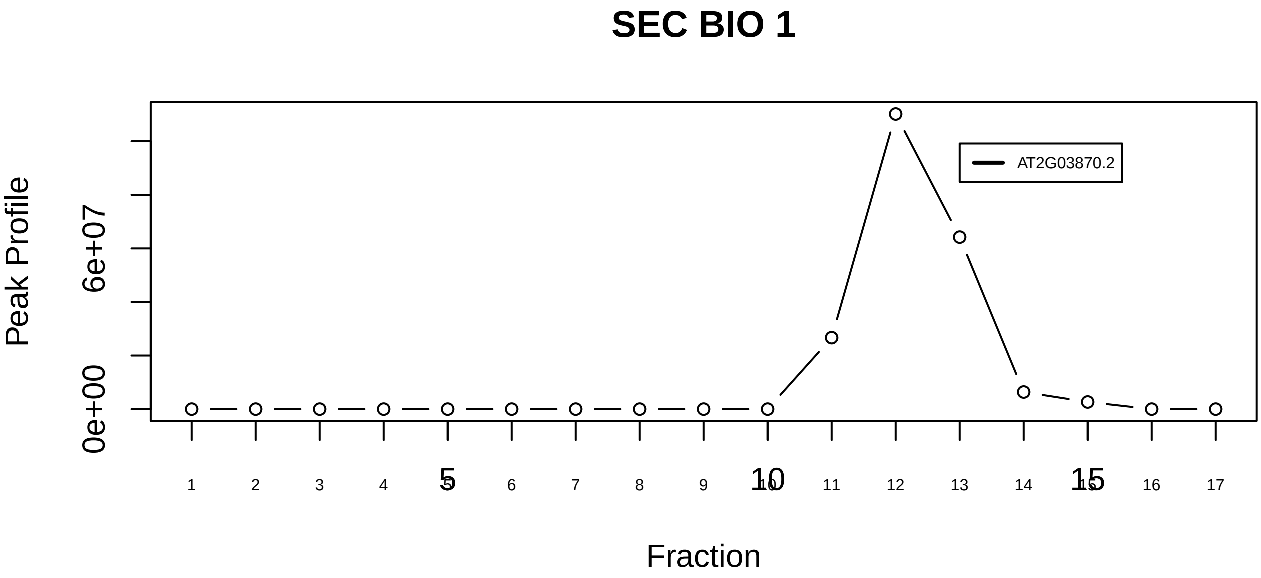

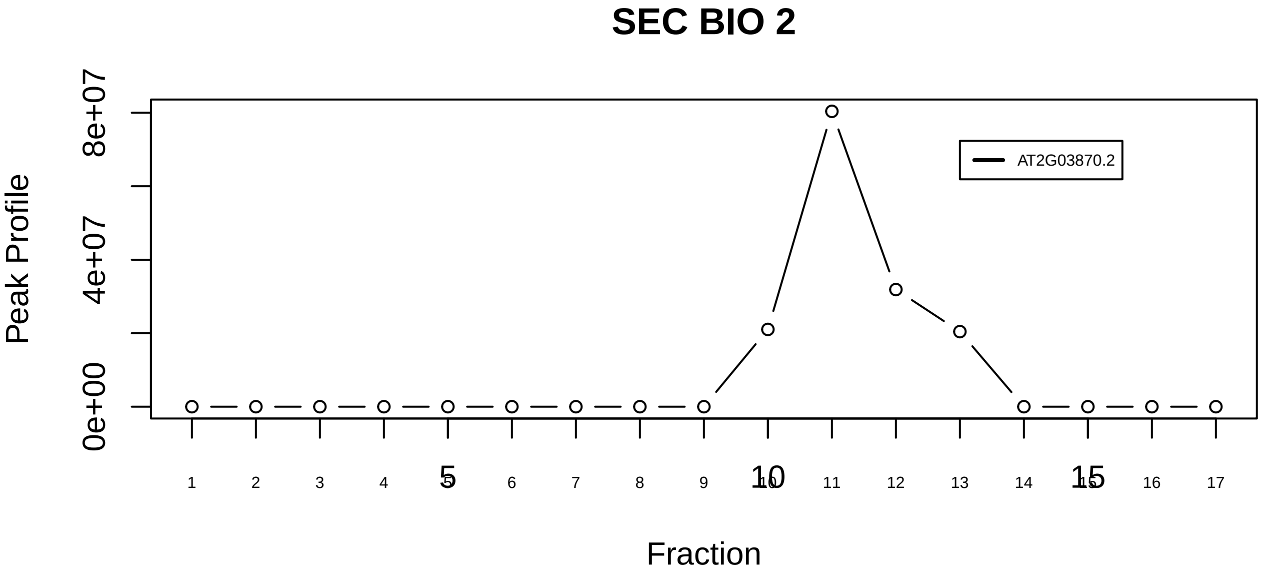

LSm2−8 complex|562−−singleton_2

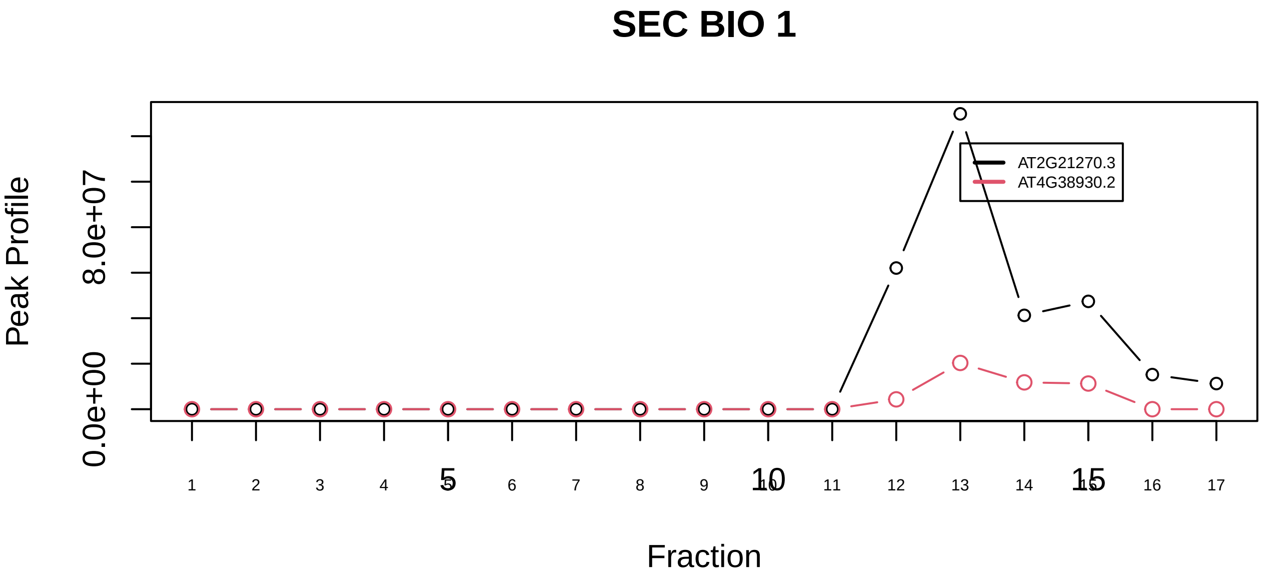

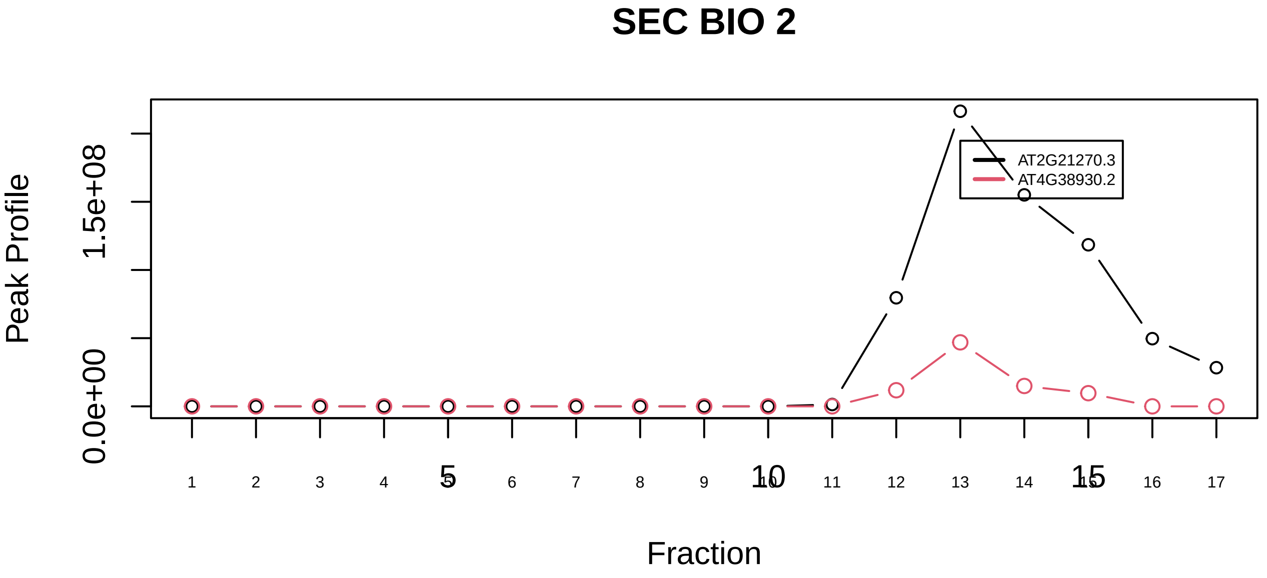

Membrane protein complex (VCP, UFD1L, SEC61B)|5685−−sub_orthoparalog_1

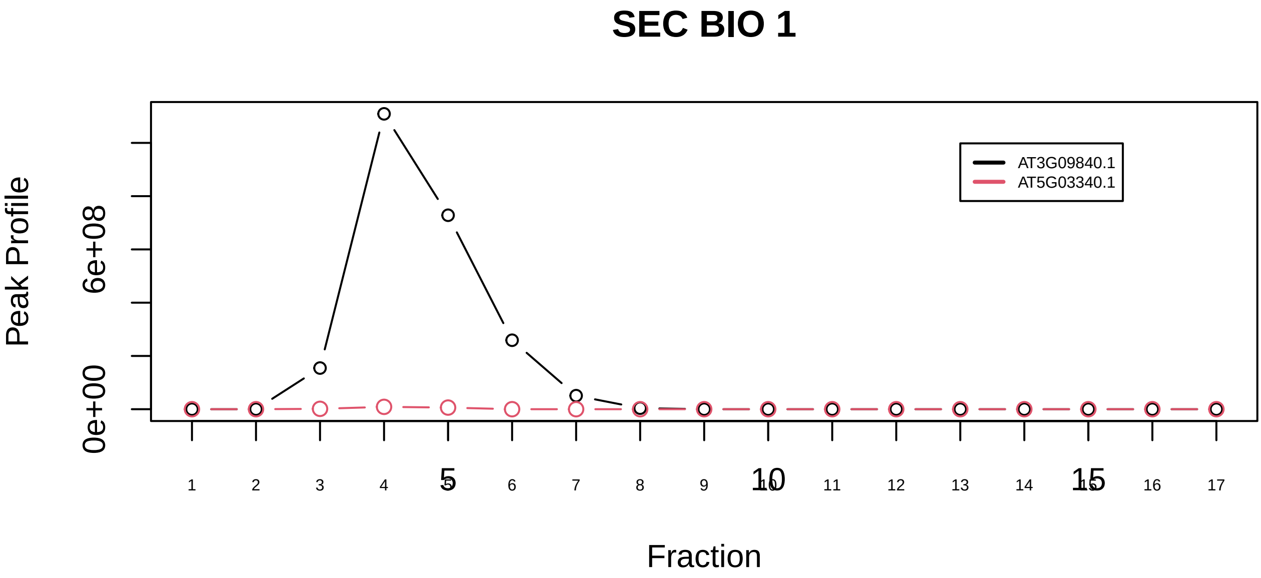

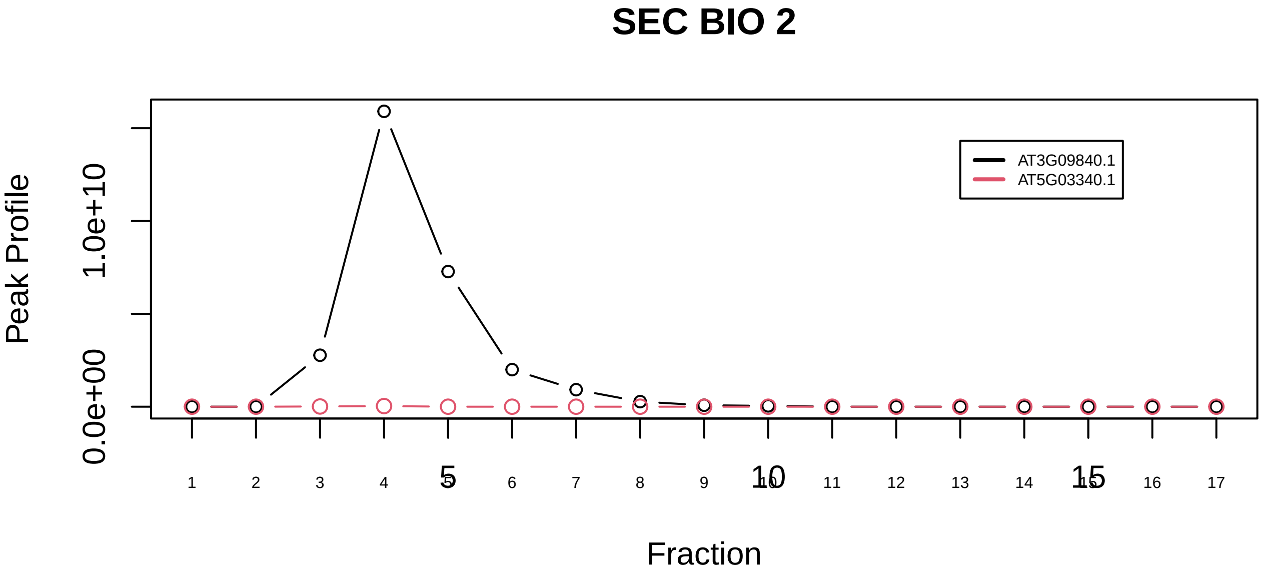

Membrane protein complex (VCP, UFD1L, SEC61B)|5685−−sub_orthoparalog_2

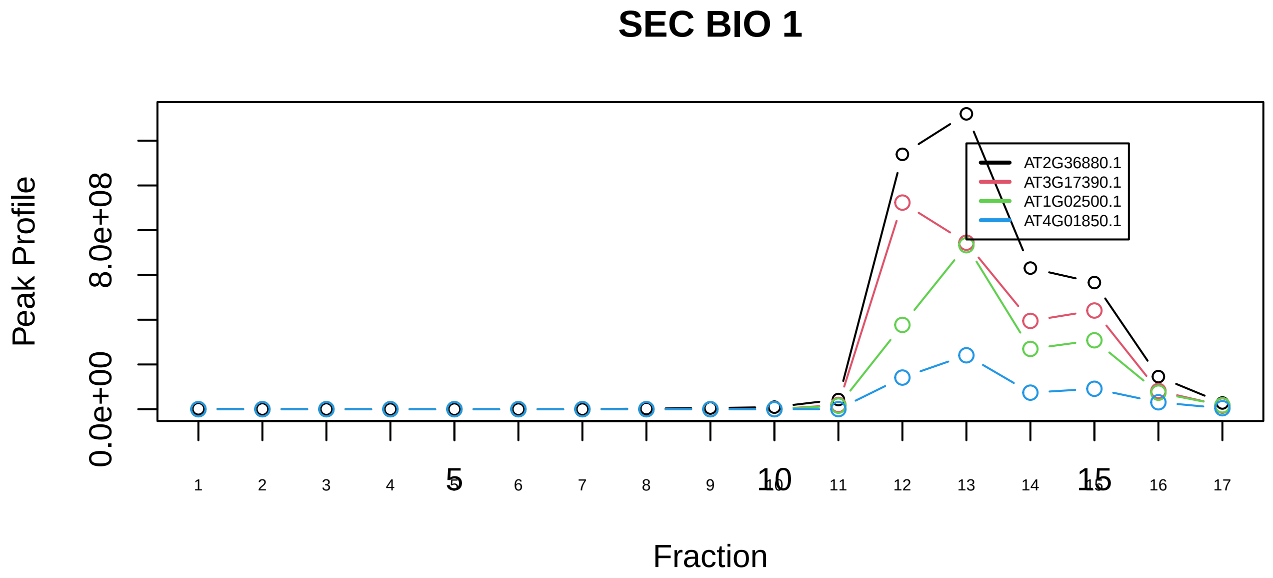

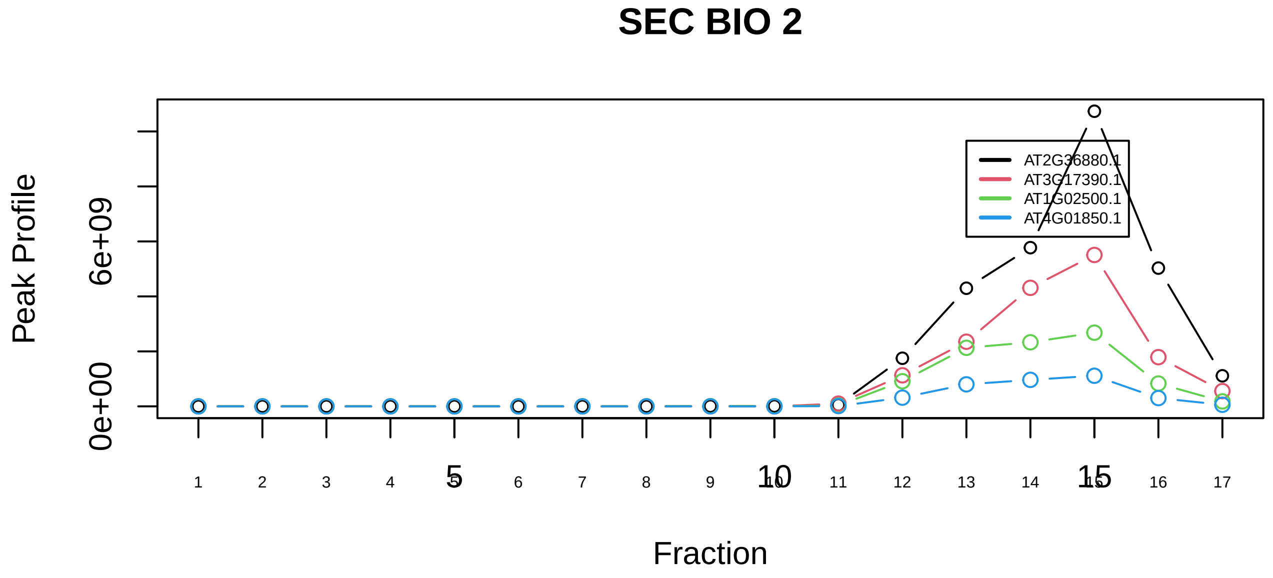

Methionine adenosyltransferase alpha2 beta−v1|7191−−sub_orthoparalog_1

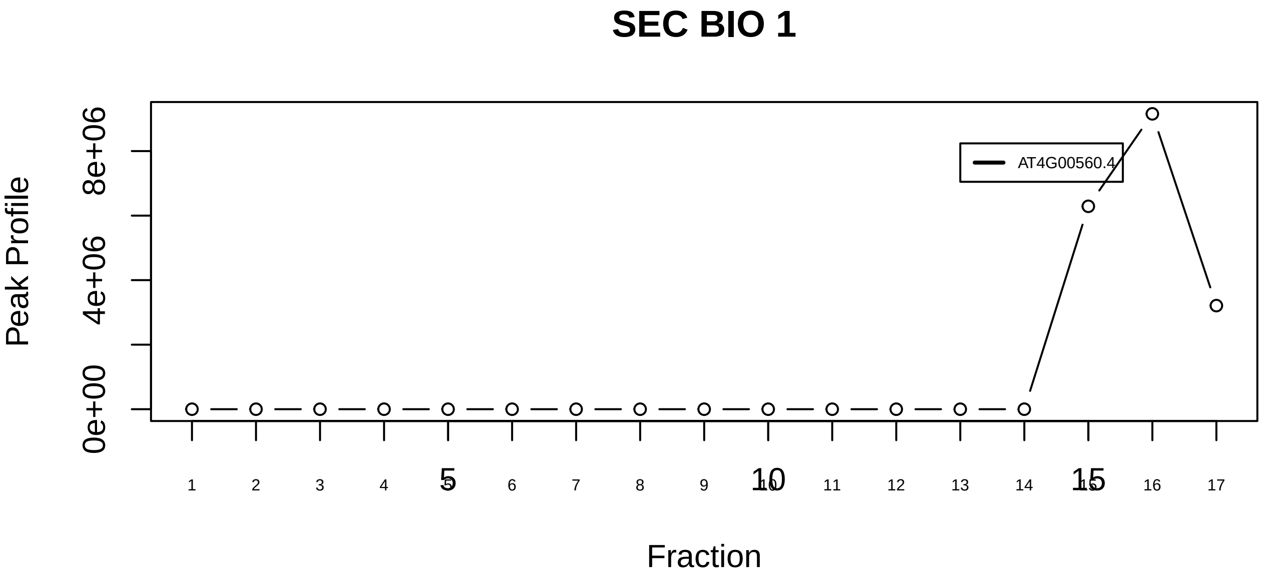

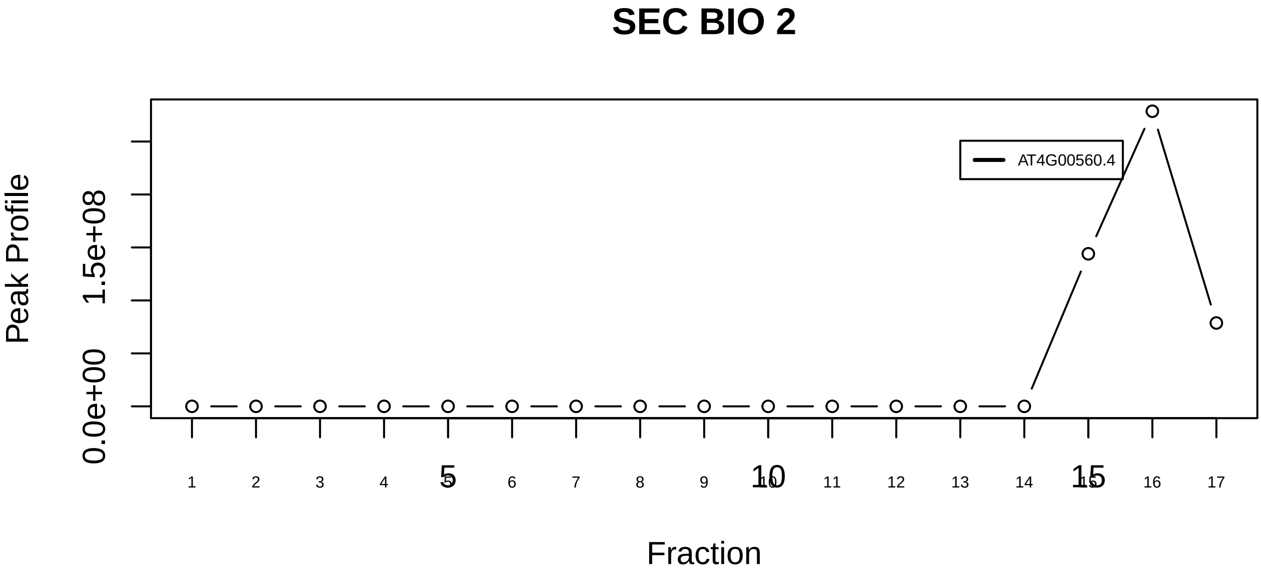

Methionine adenosyltransferase alpha2 beta−v1|7191−−singleton_1

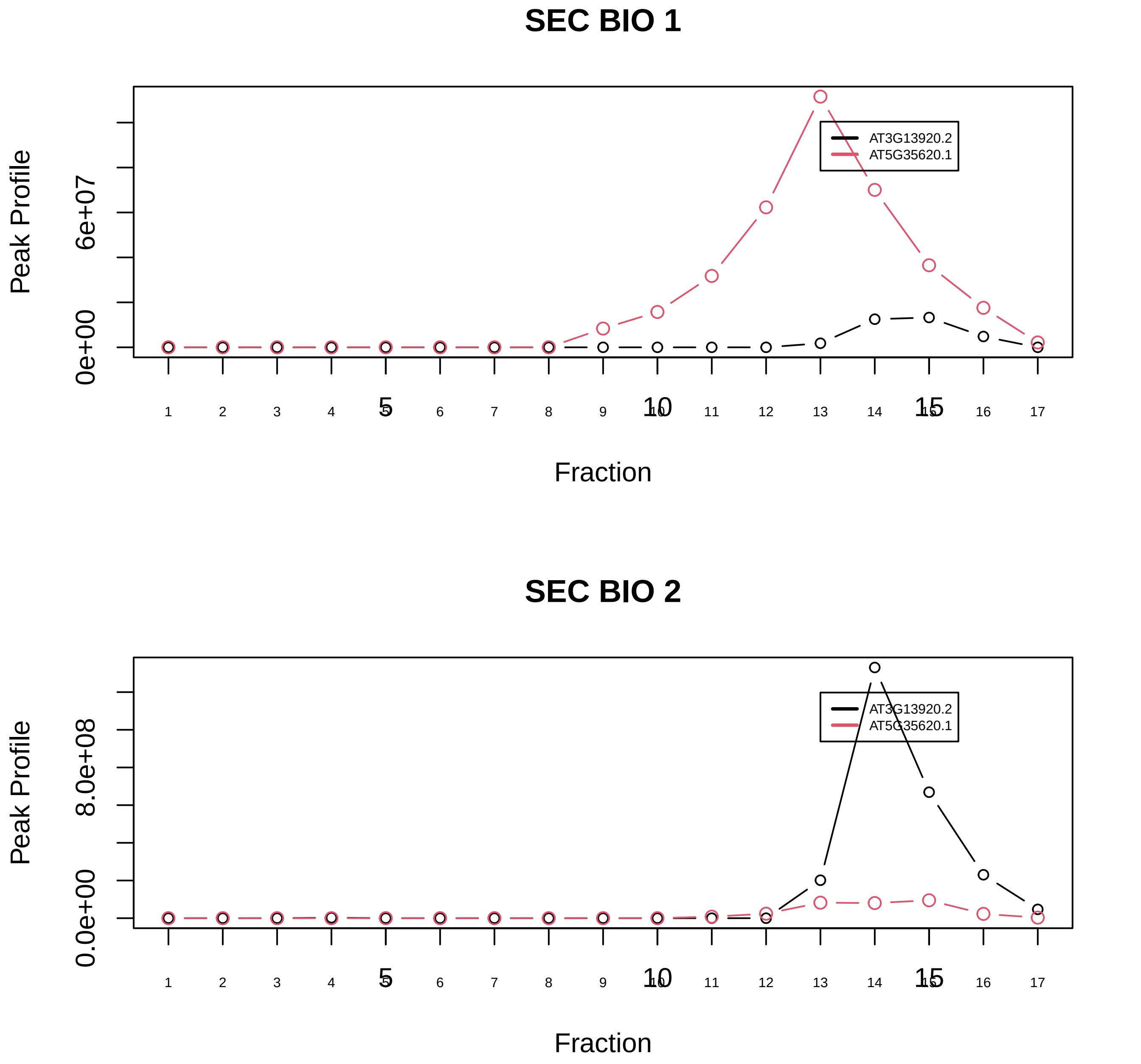

MNK1−eIF4F complex|5253−−only_2_prots_before_clustering

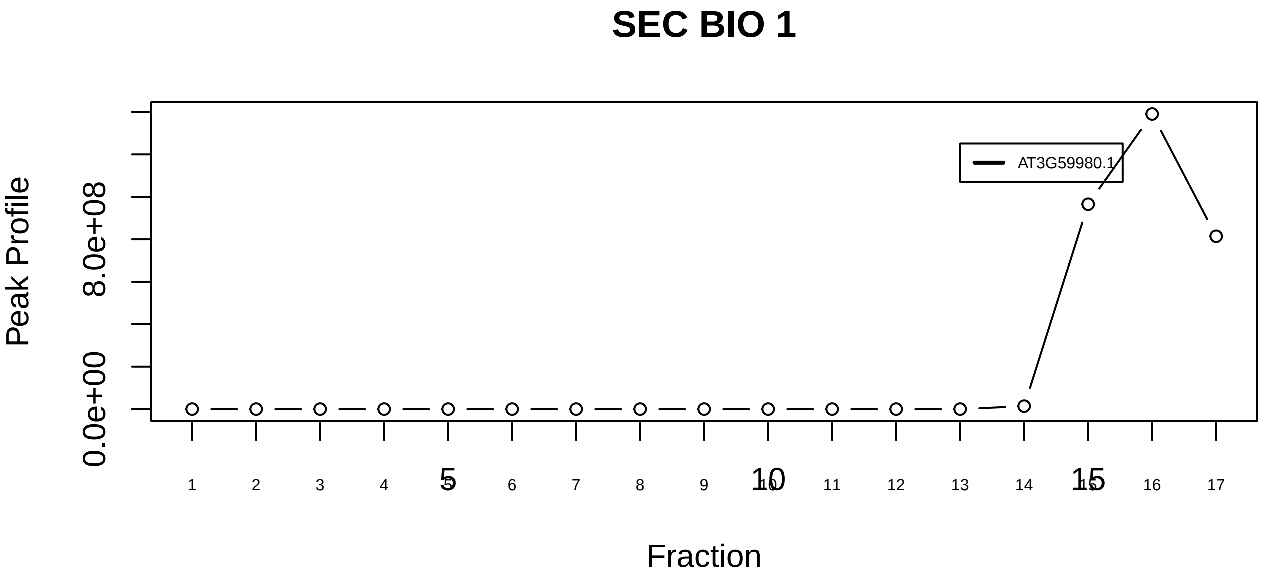

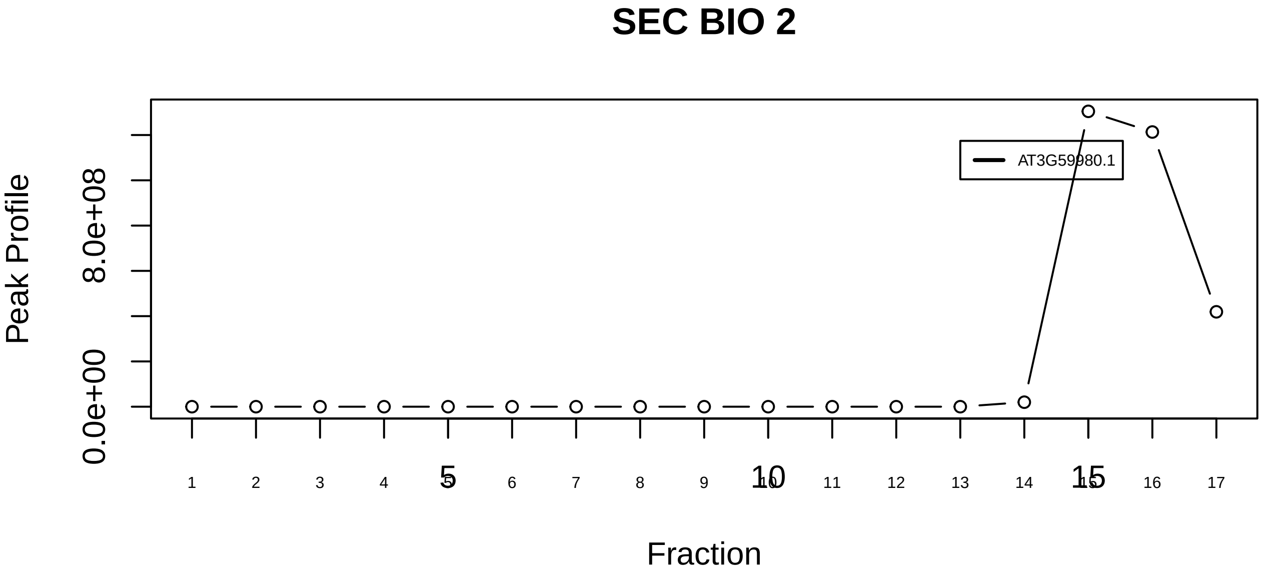

Multisynthetase complex|3040−−singleton_1

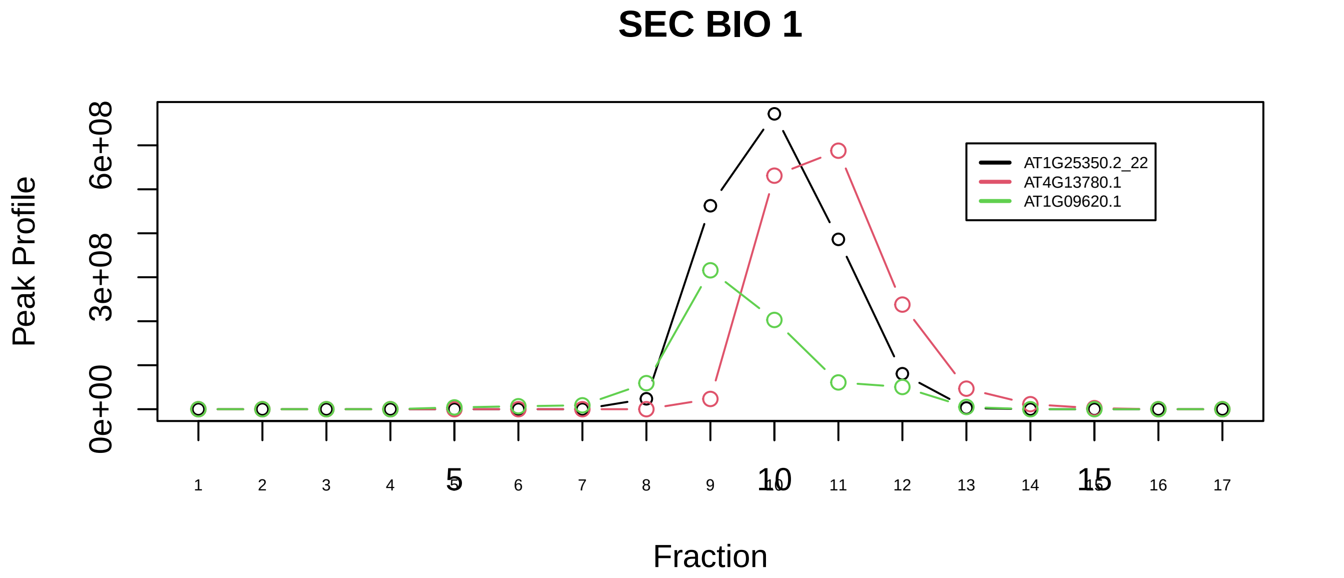

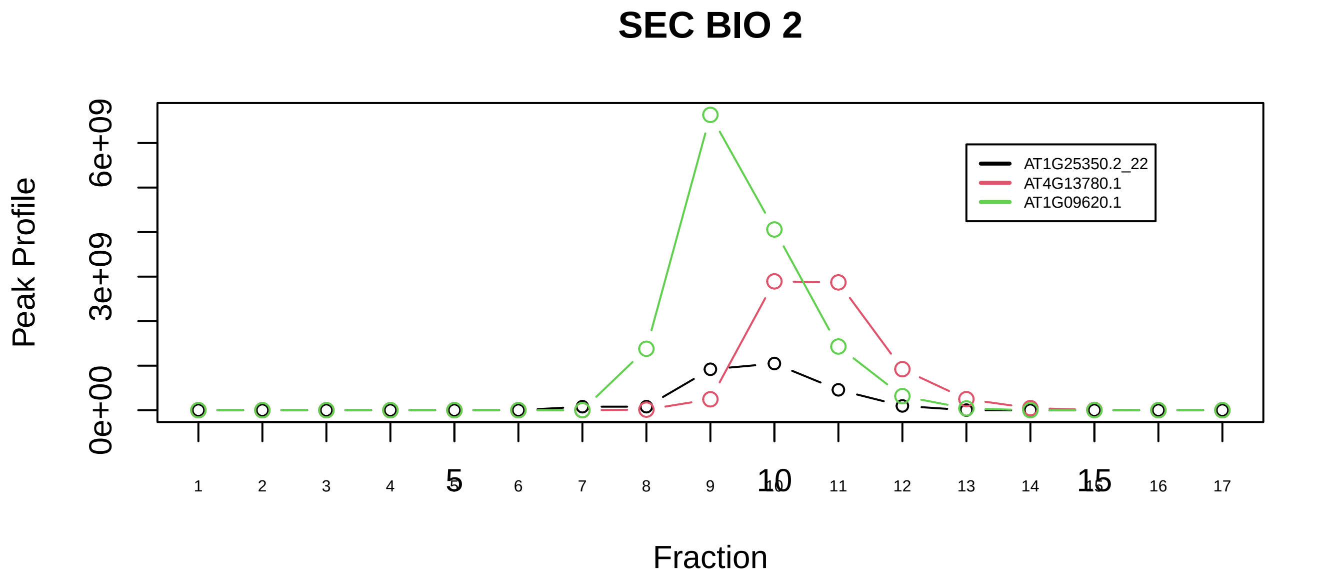

Multisynthetase complex|3040−−sub_1

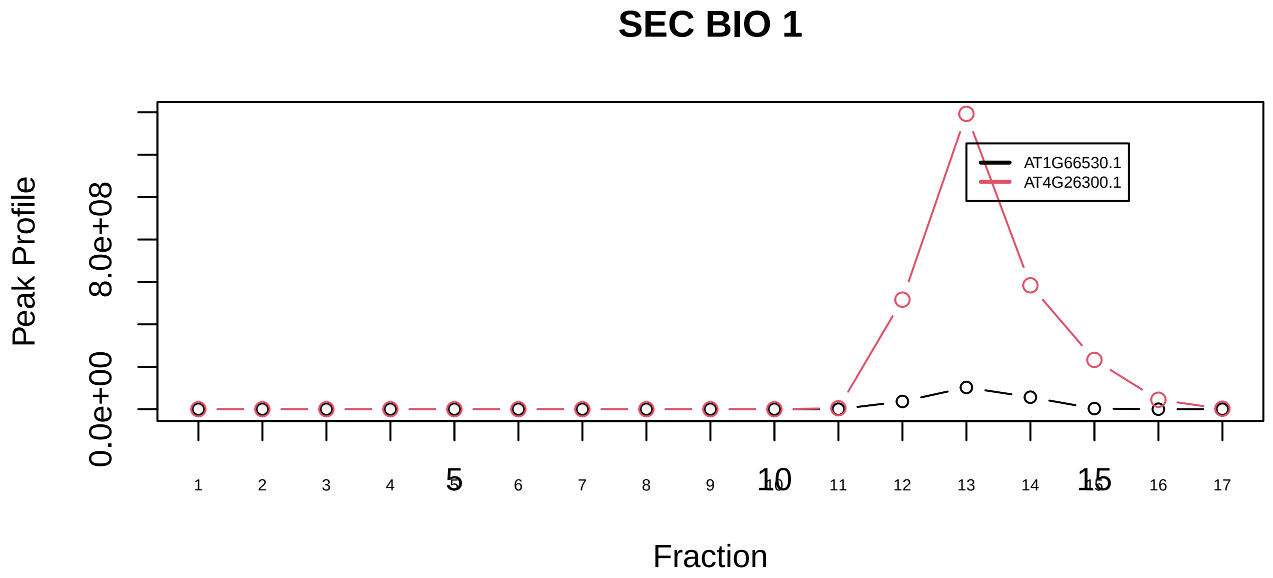

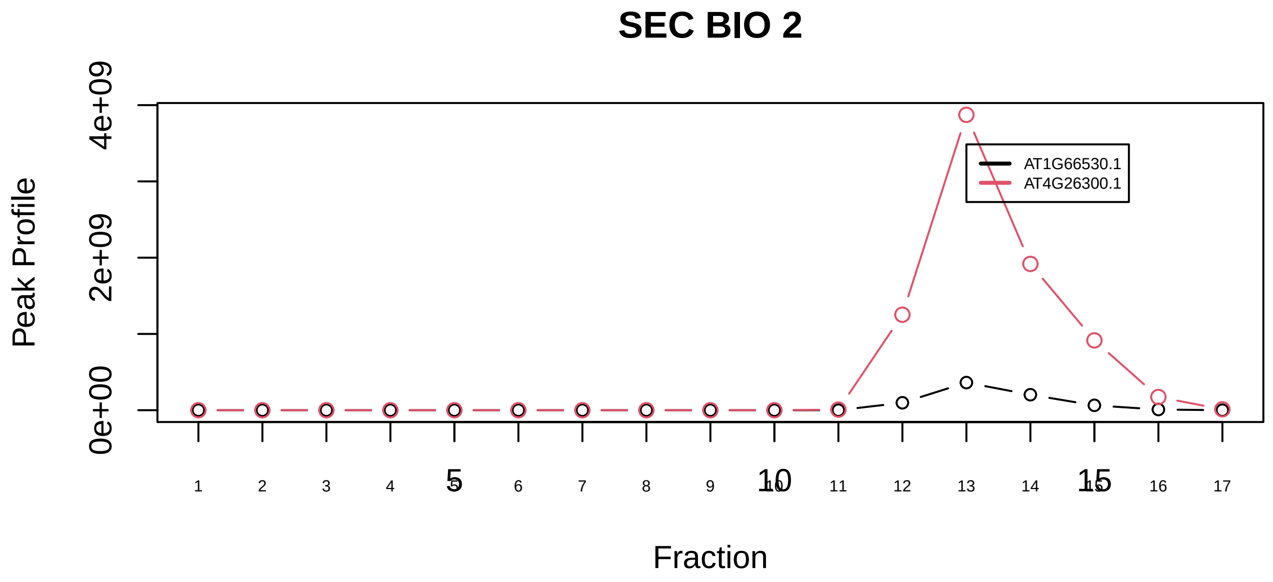

Multisynthetase complex|3040−−sub_orthoparalog_1

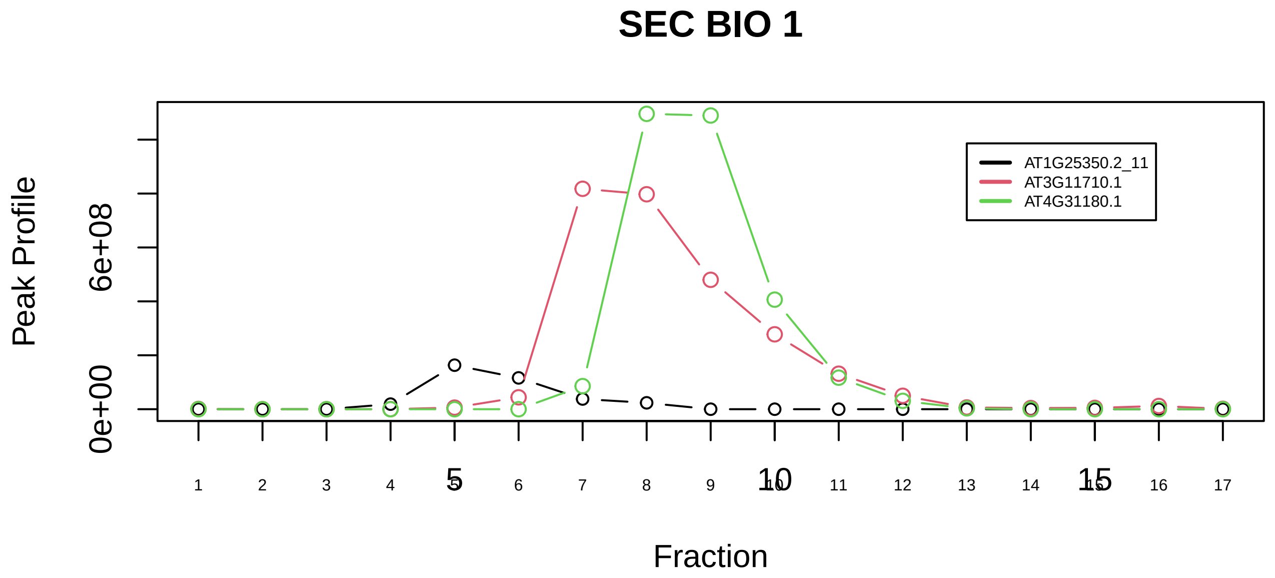

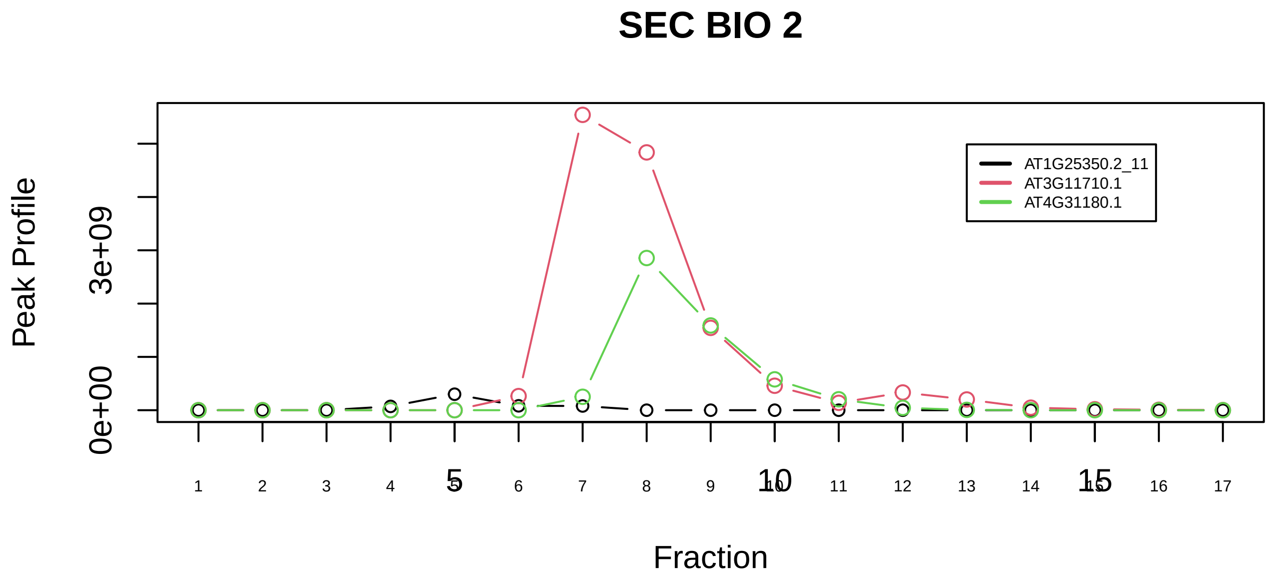

Multisynthetase complex|3040−−sub_2

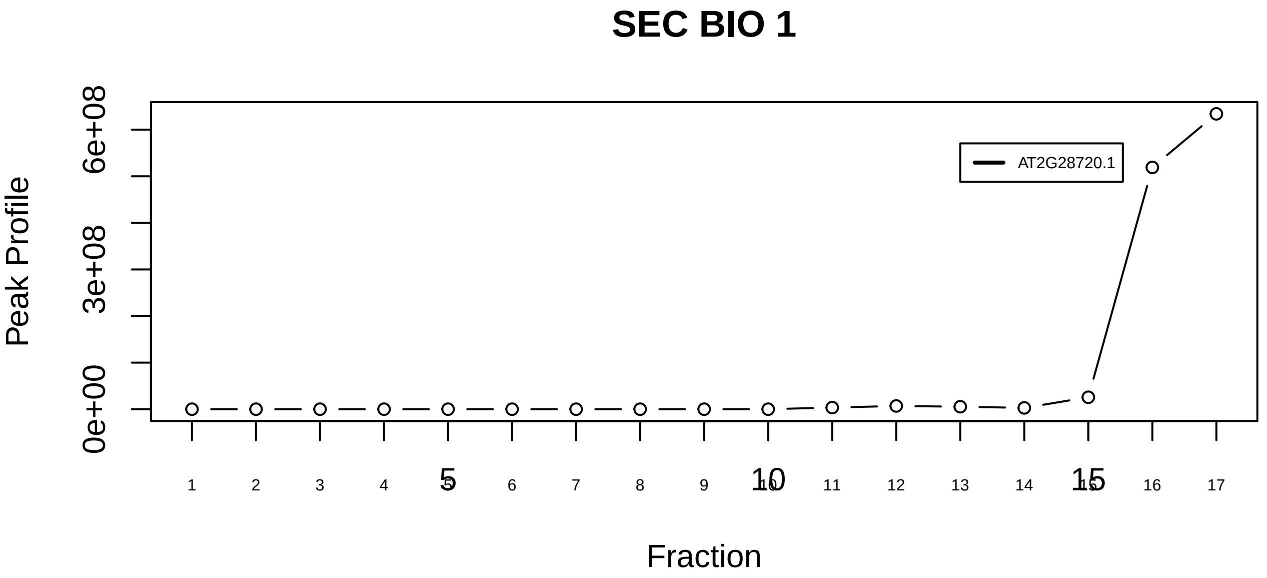

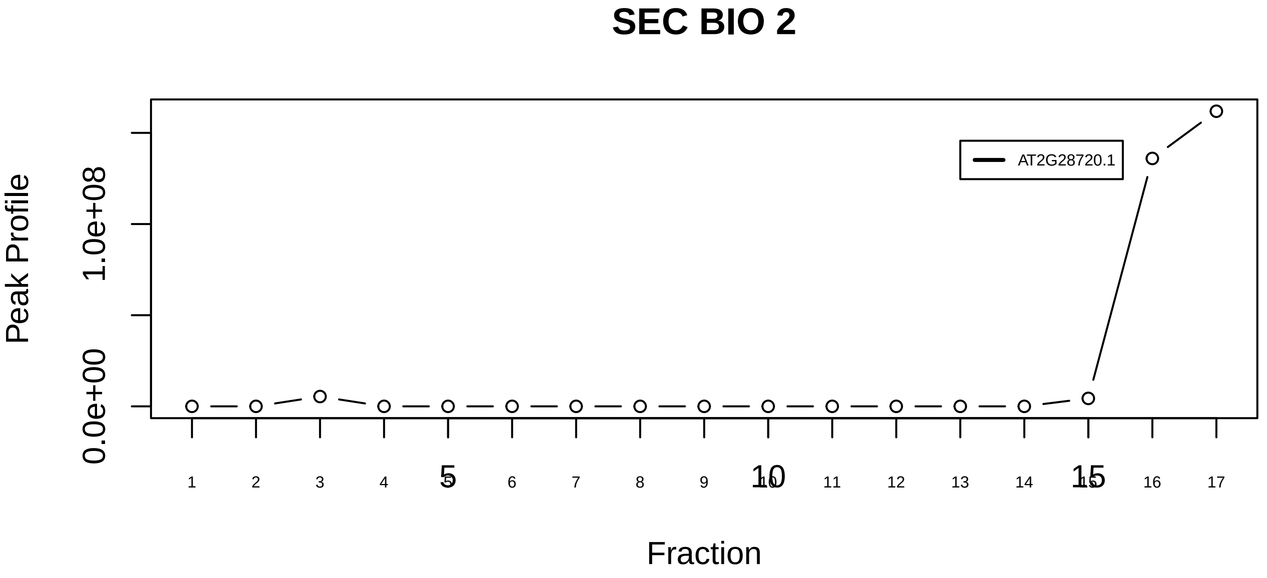

Nop56p−associated pre−rRNA complex|3055−−singleton_1

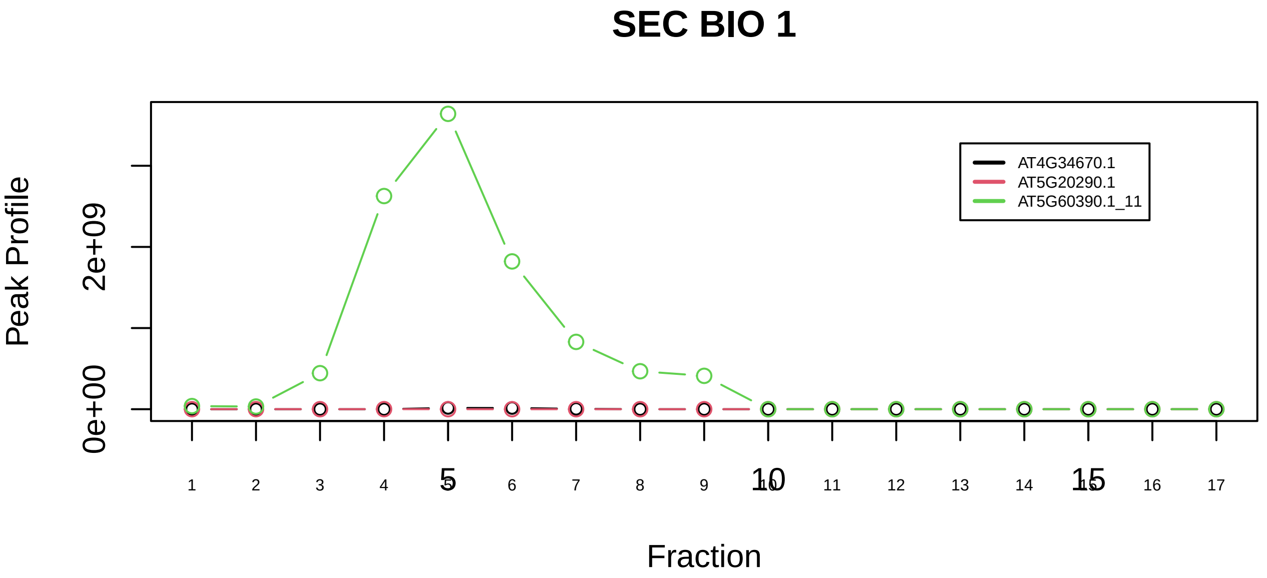

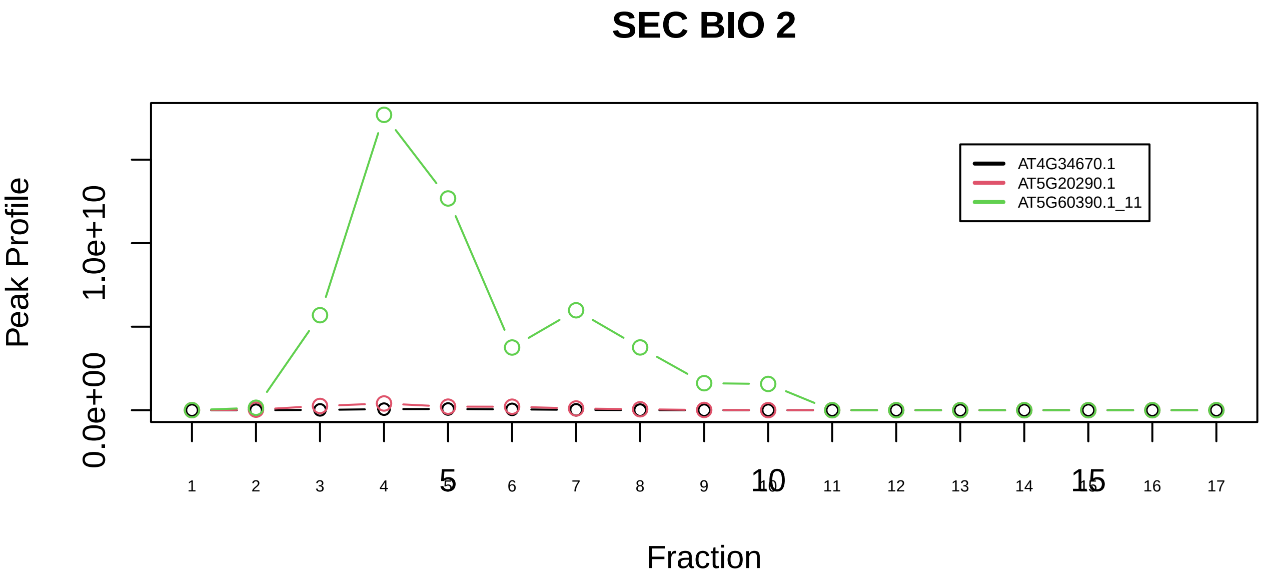

Nop56p−associated pre−rRNA complex|3055−−sub_1

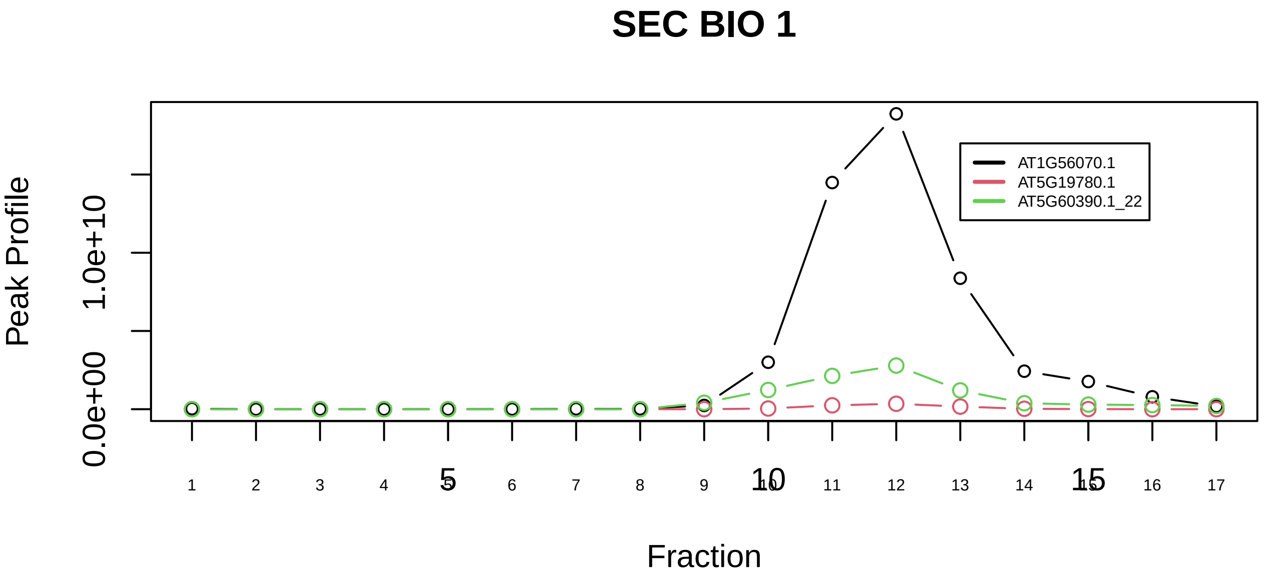

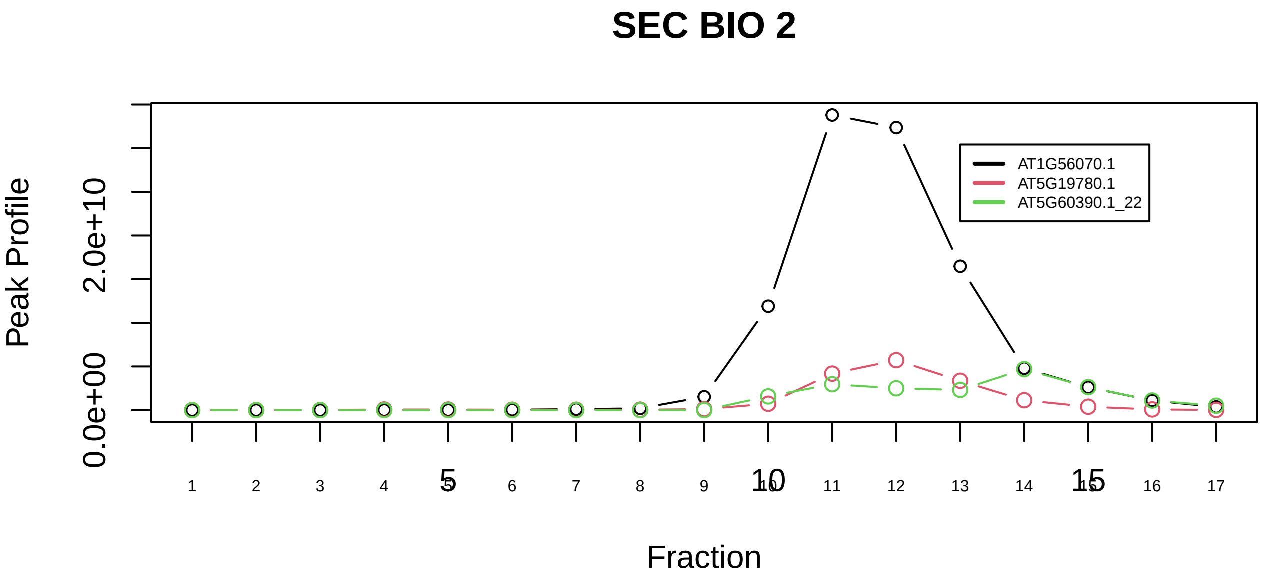

Nop56p−associated pre−rRNA complex|3055−−sub_2

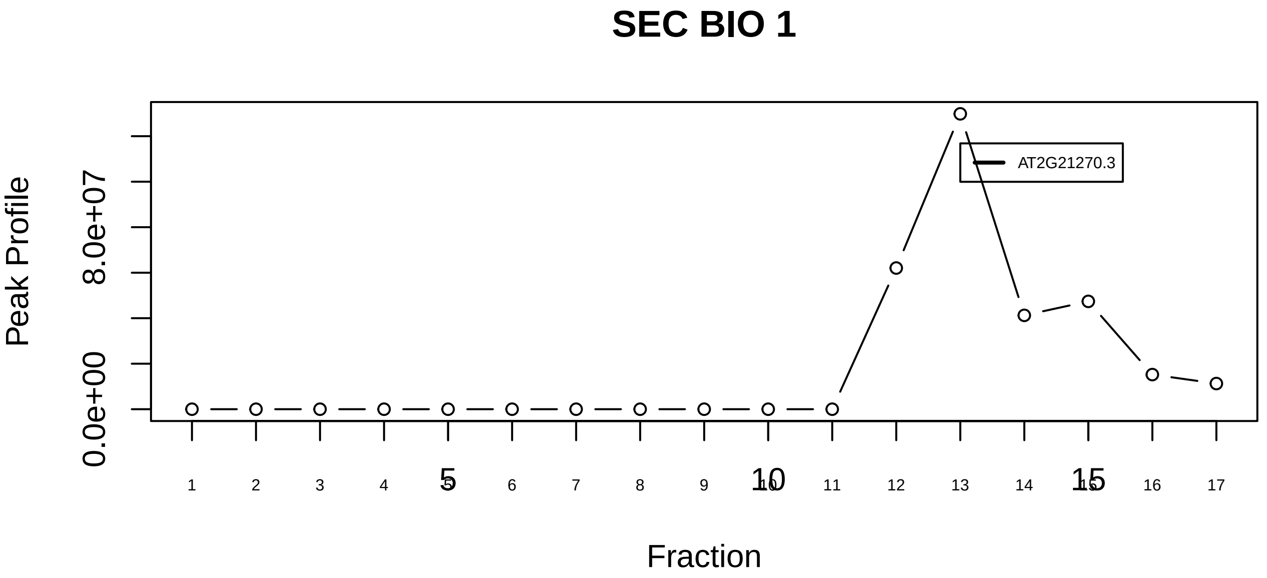

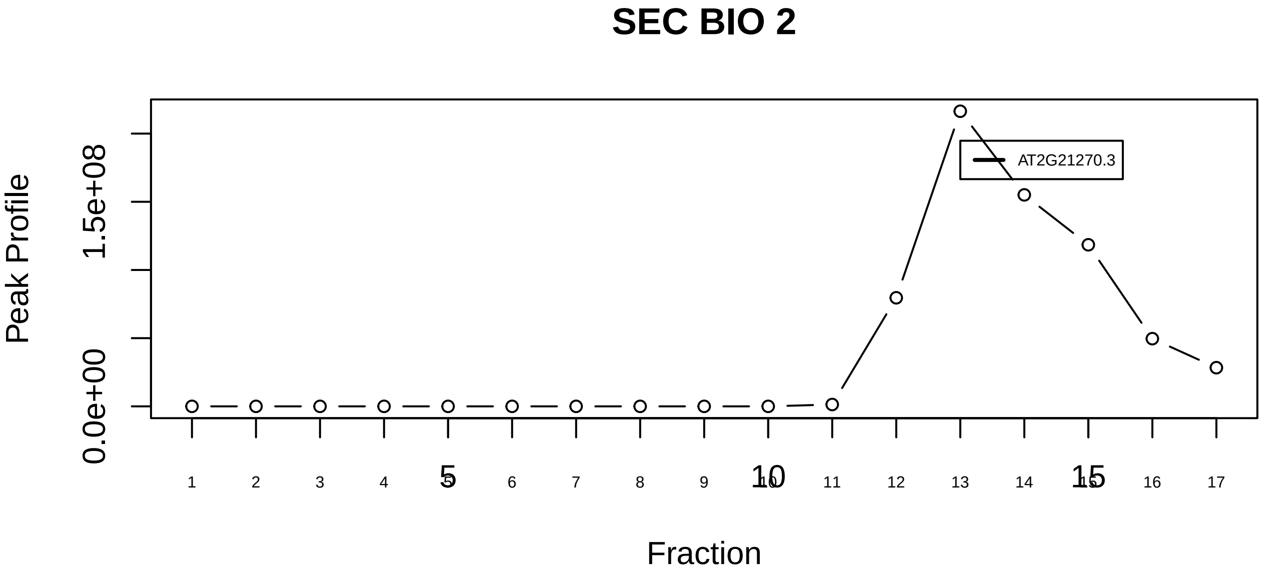

NUB1L−UFD1−NPL4−NEDD8 complex|6280−−singleton_1

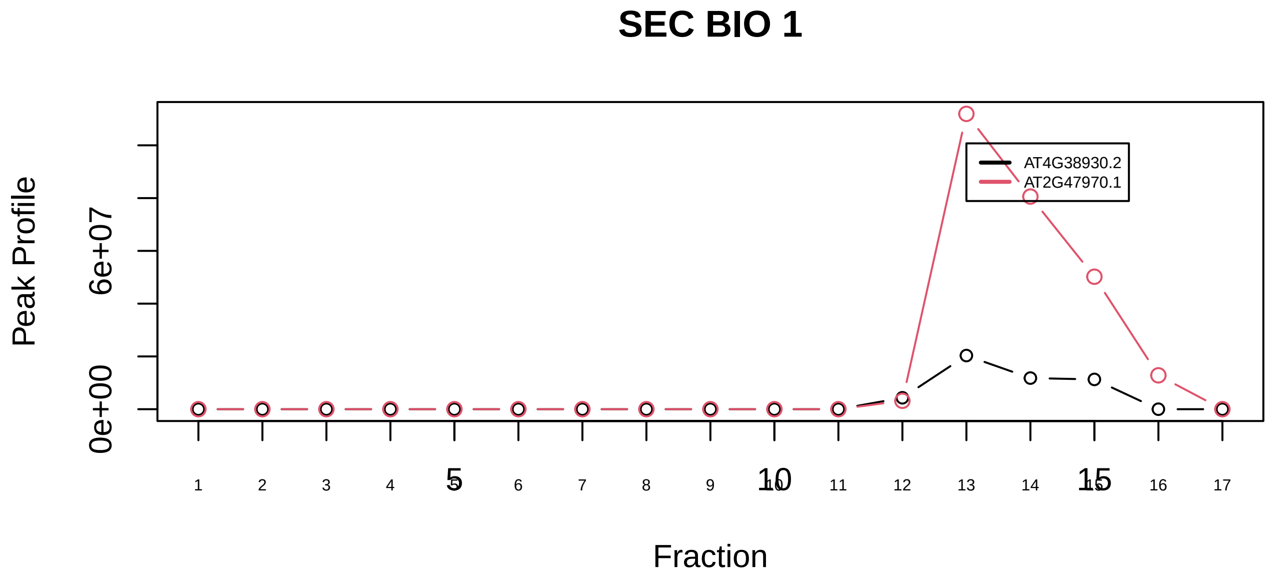

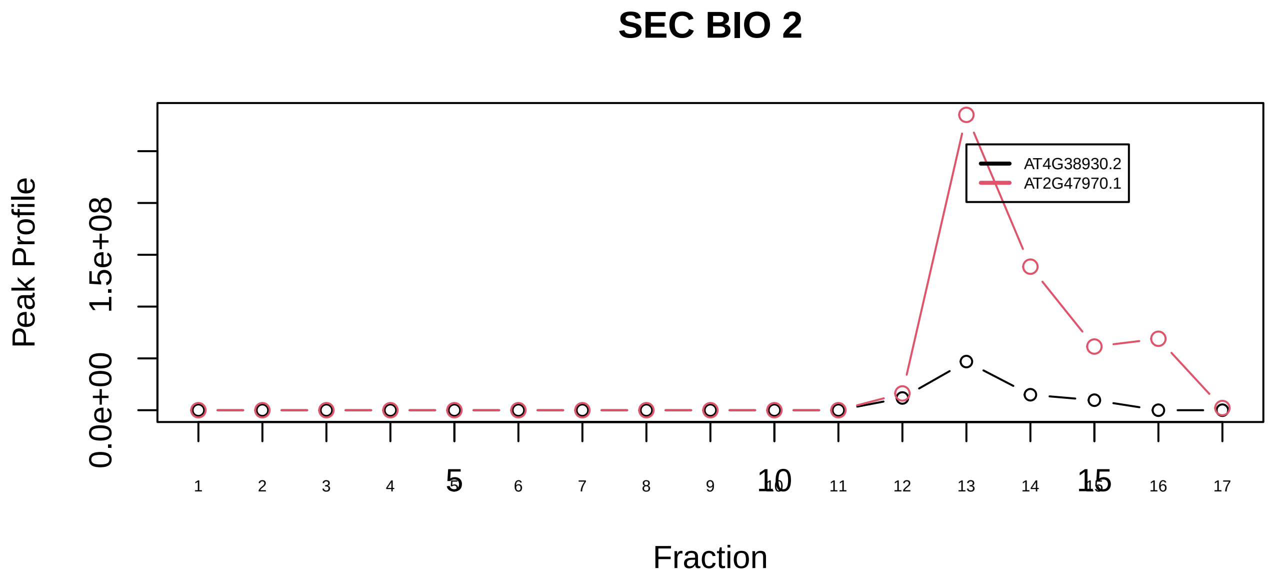

NUB1L−UFD1−NPL4−NEDD8 complex|6280−−sub_1

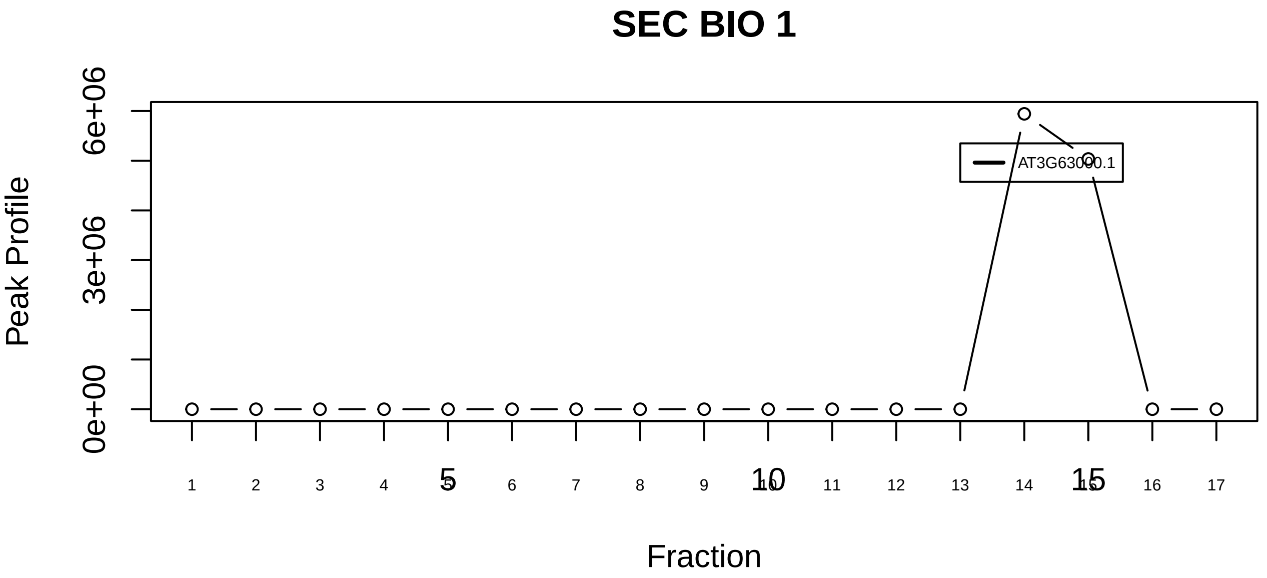

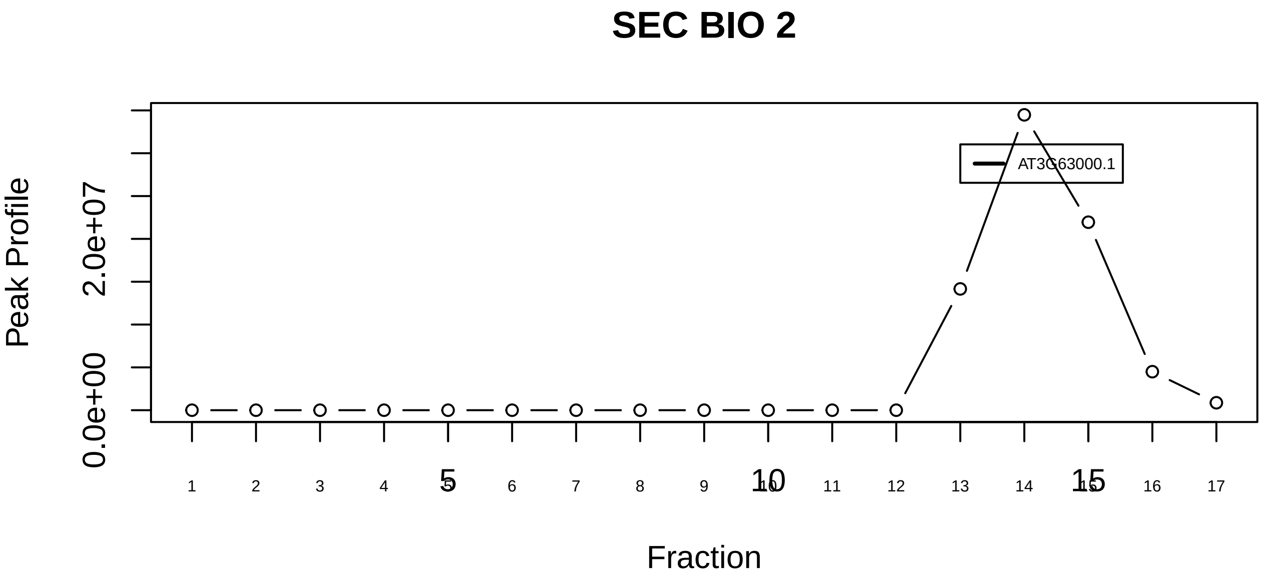

NUB1L−UFD1−NPL4−NEDD8 complex|6280−−singleton_2

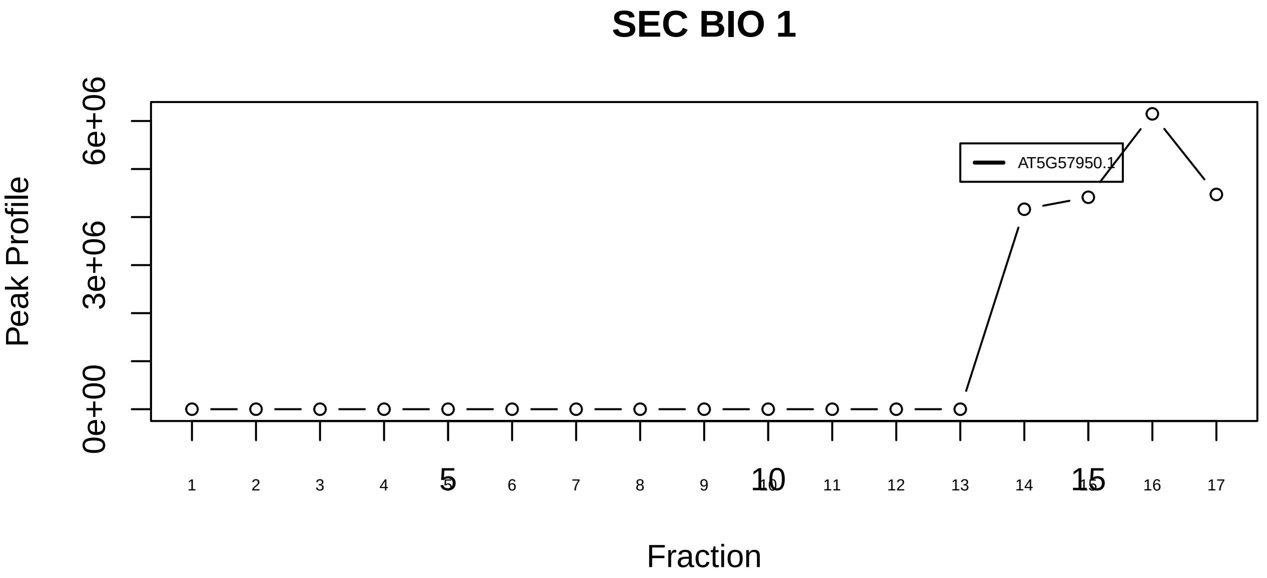

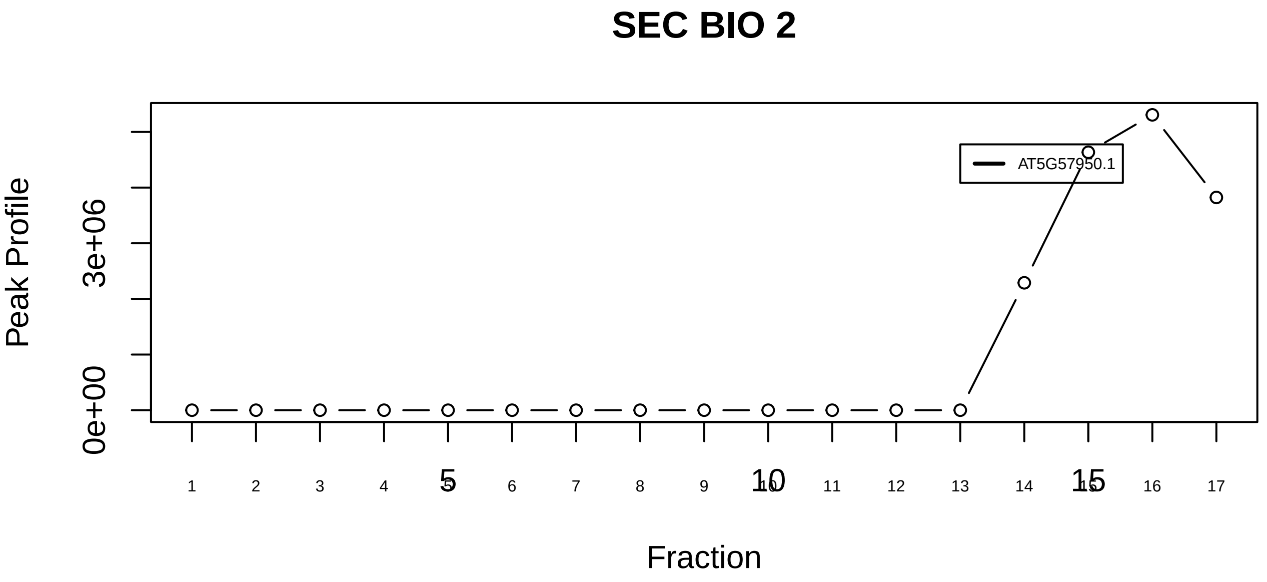

PA700−20S−PA28 complex|193−−singleton_1

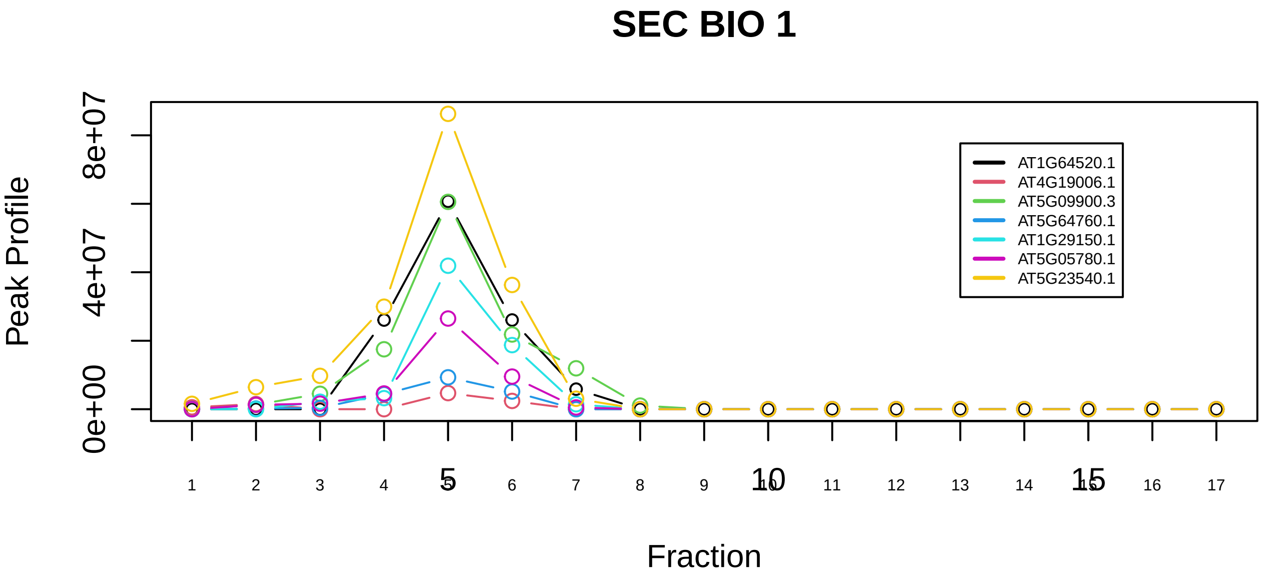

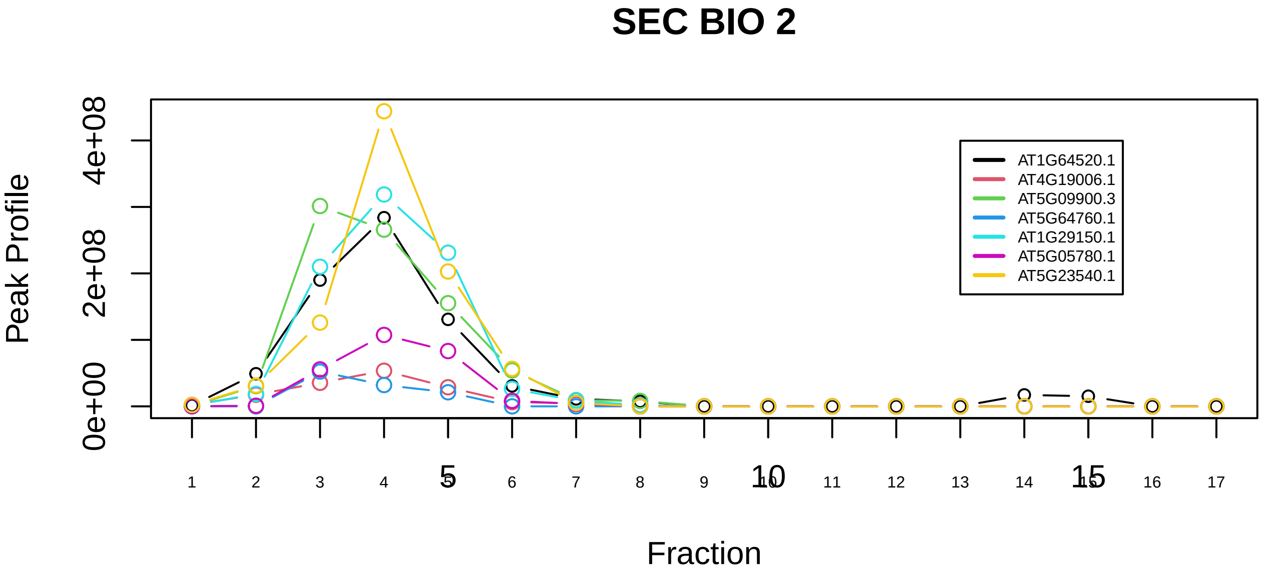

PA700−20S−PA28 complex|193−−sub_1

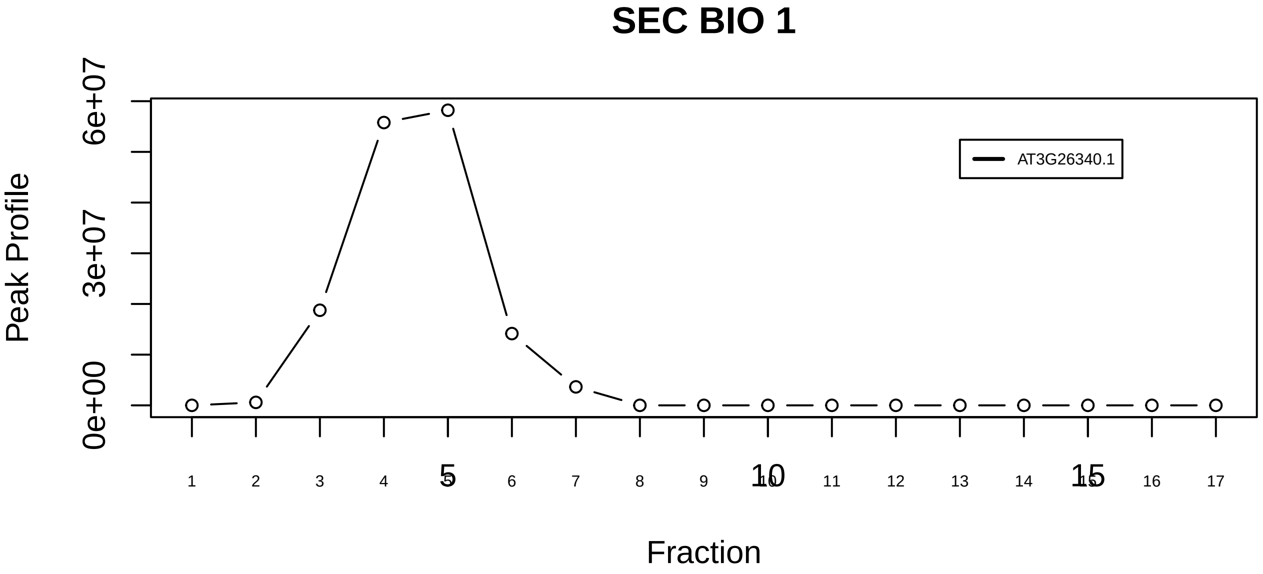

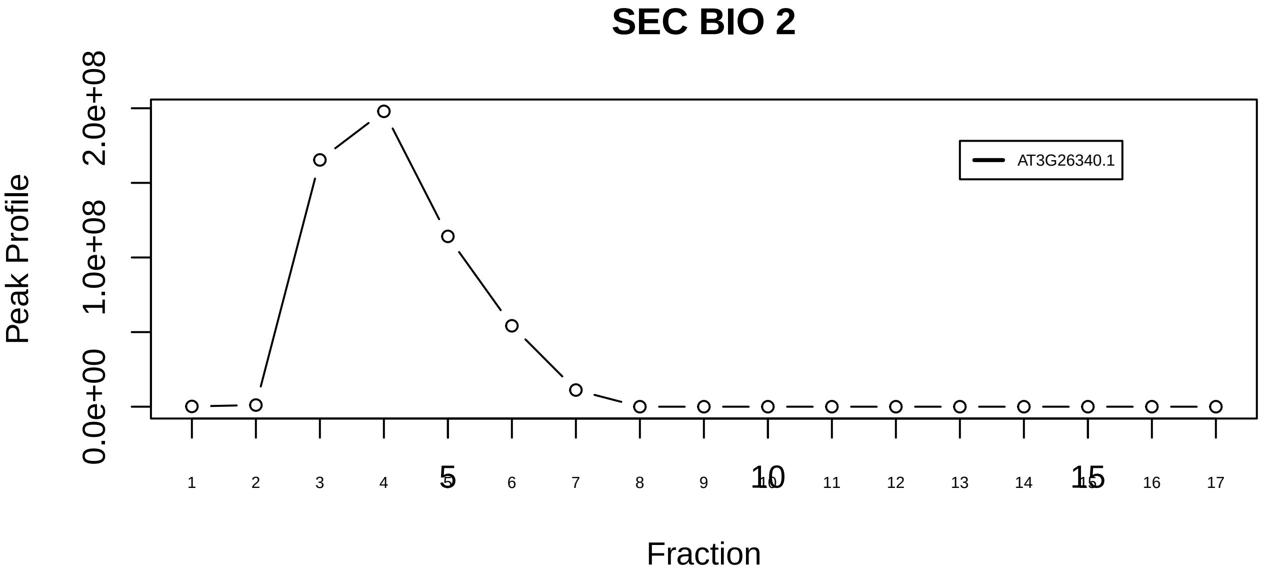

PA700−20S−PA28 complex|193−−singleton_2

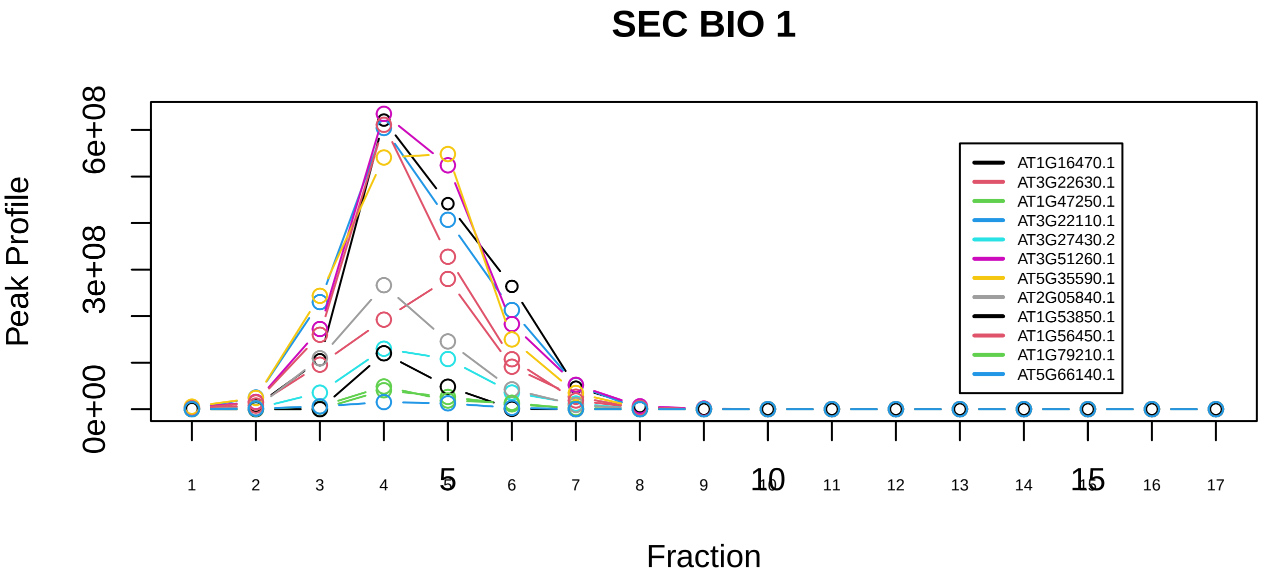

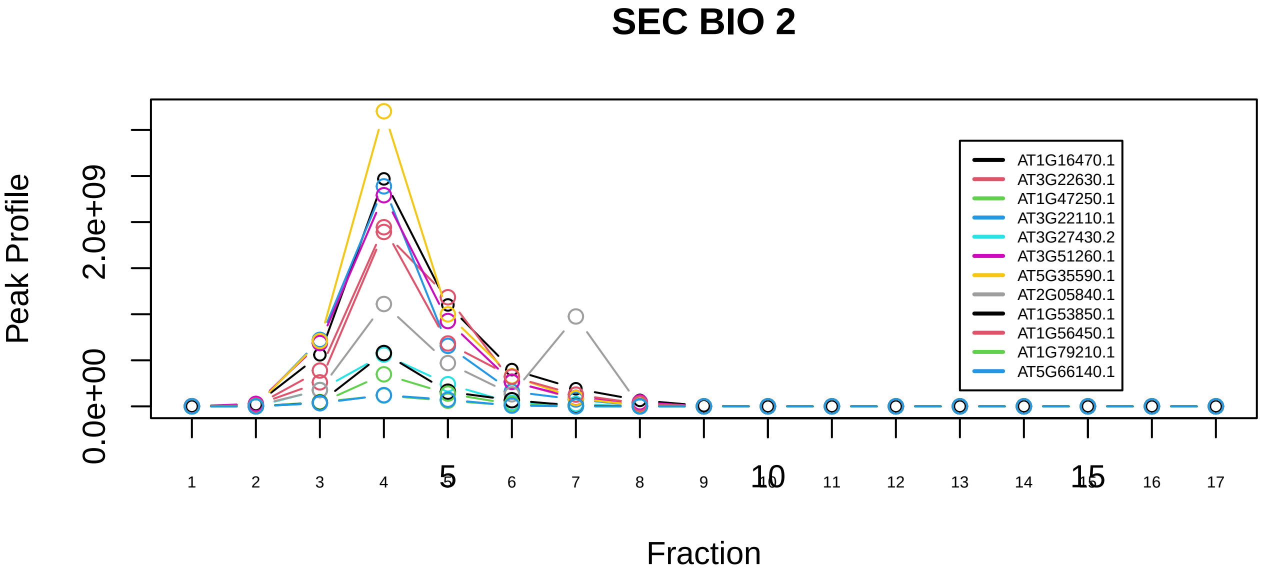

PA700−20S−PA28 complex|193−−sub_2

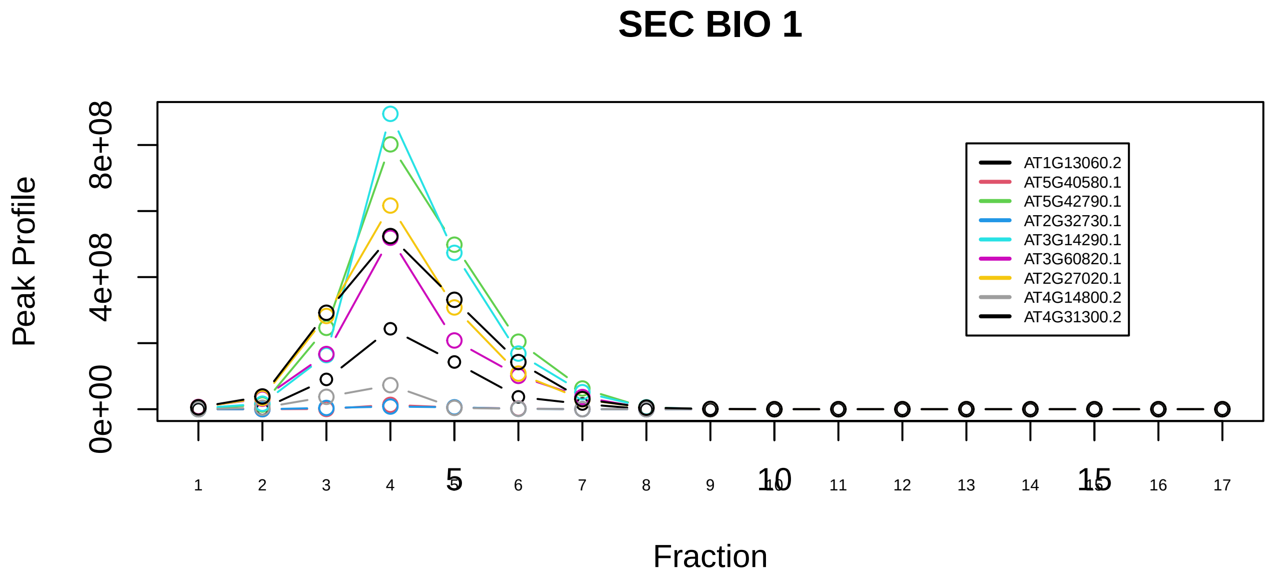

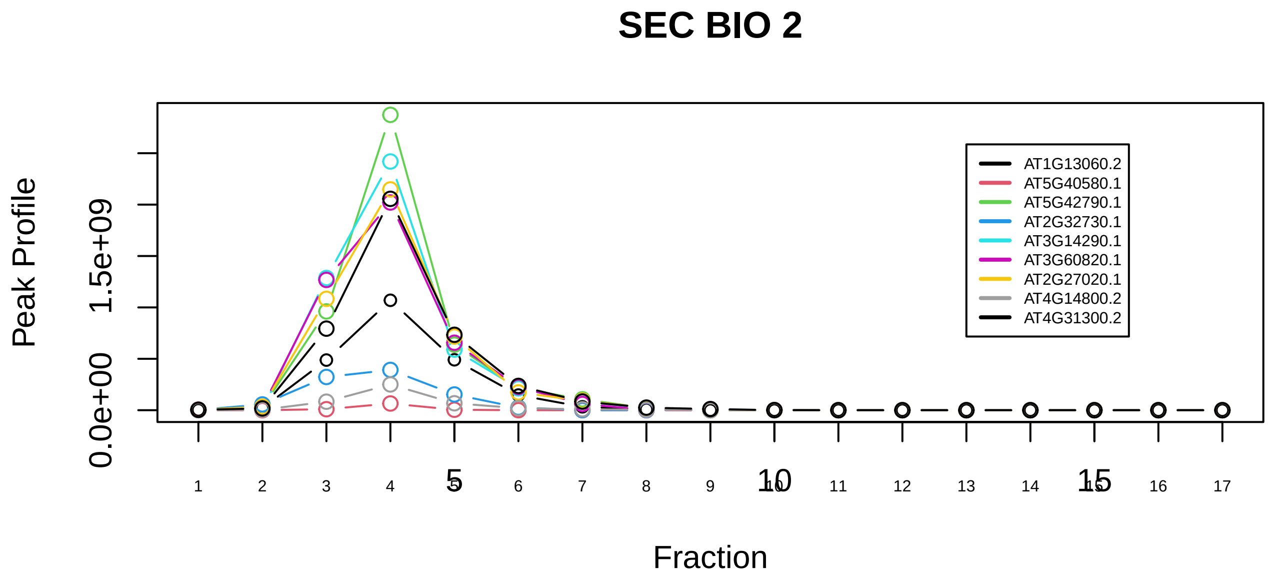

PA700−20S−PA28 complex|193−−sub_3

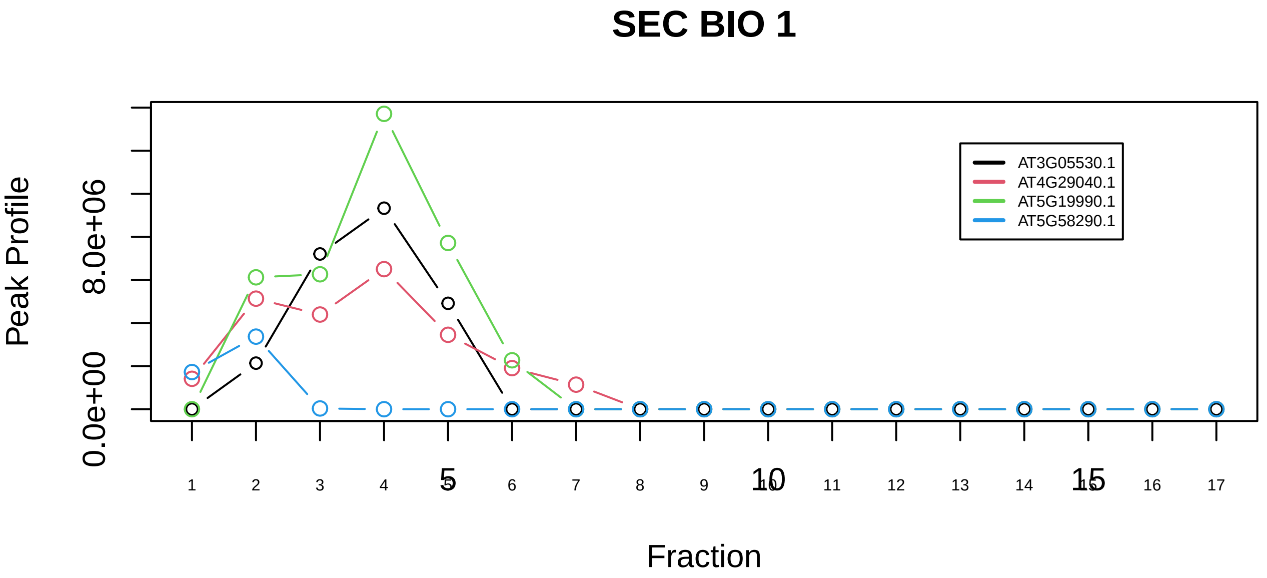

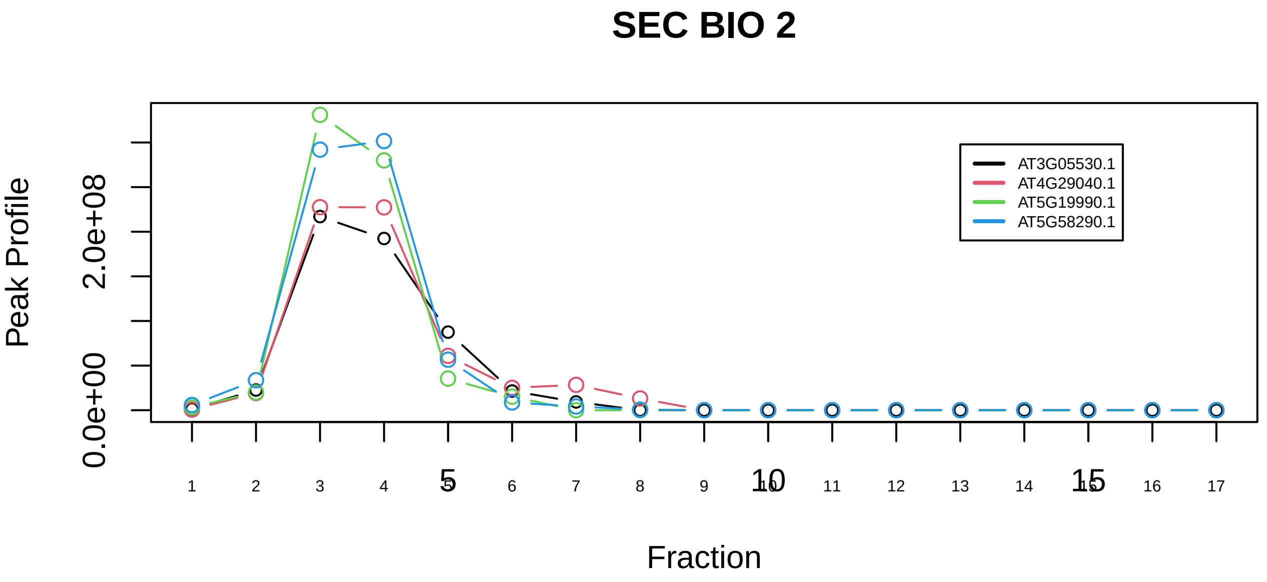

PA700−20S−PA28 complex|193−−sub_4

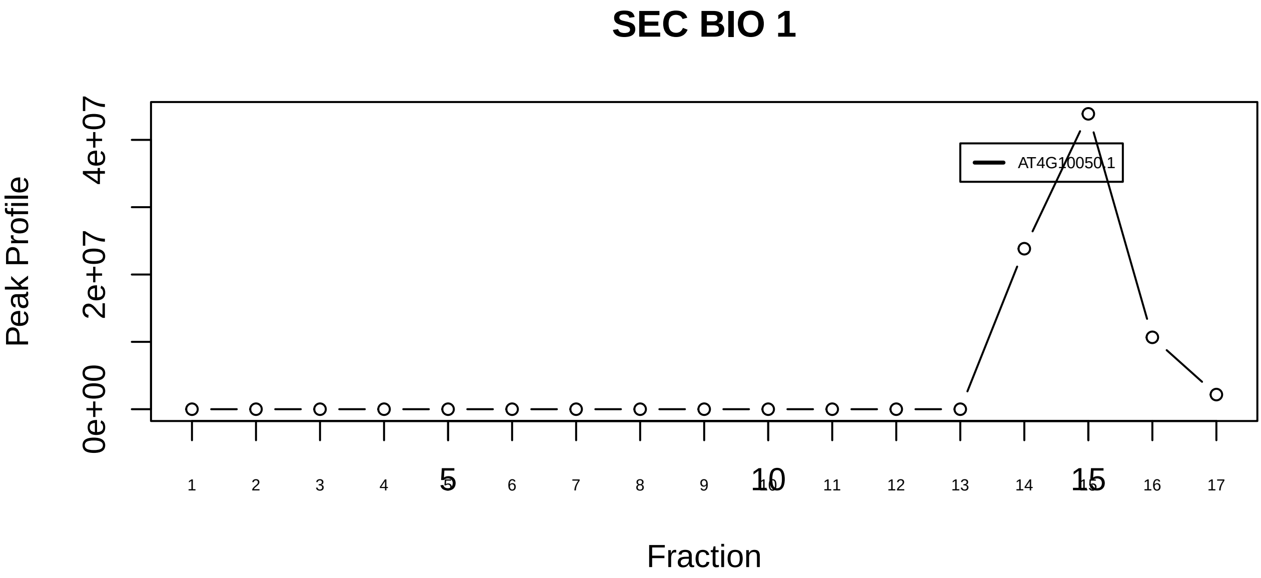

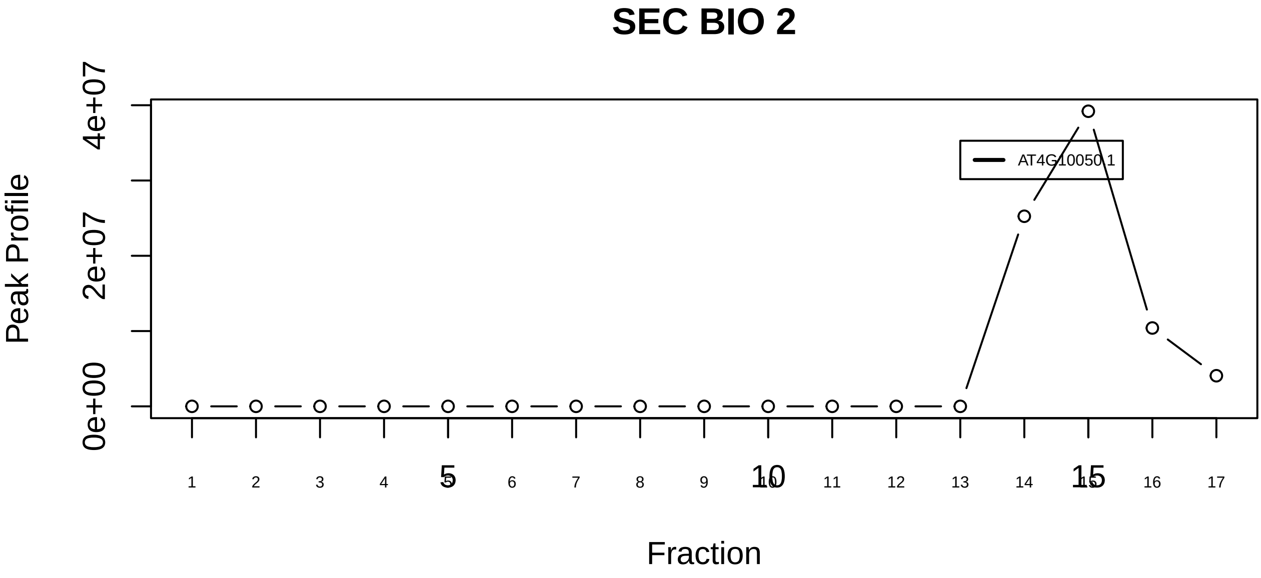

PPP2R1A−PPP2R1B−PPP2CA−PPME1−EIF4A1 complex|5876−−singleton_1

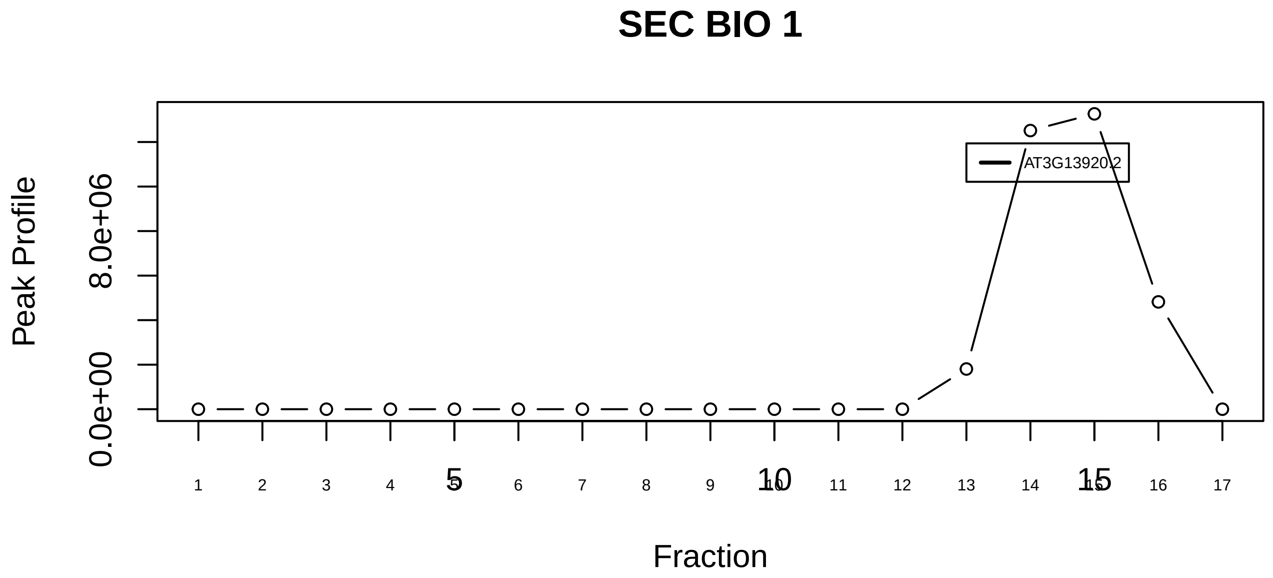

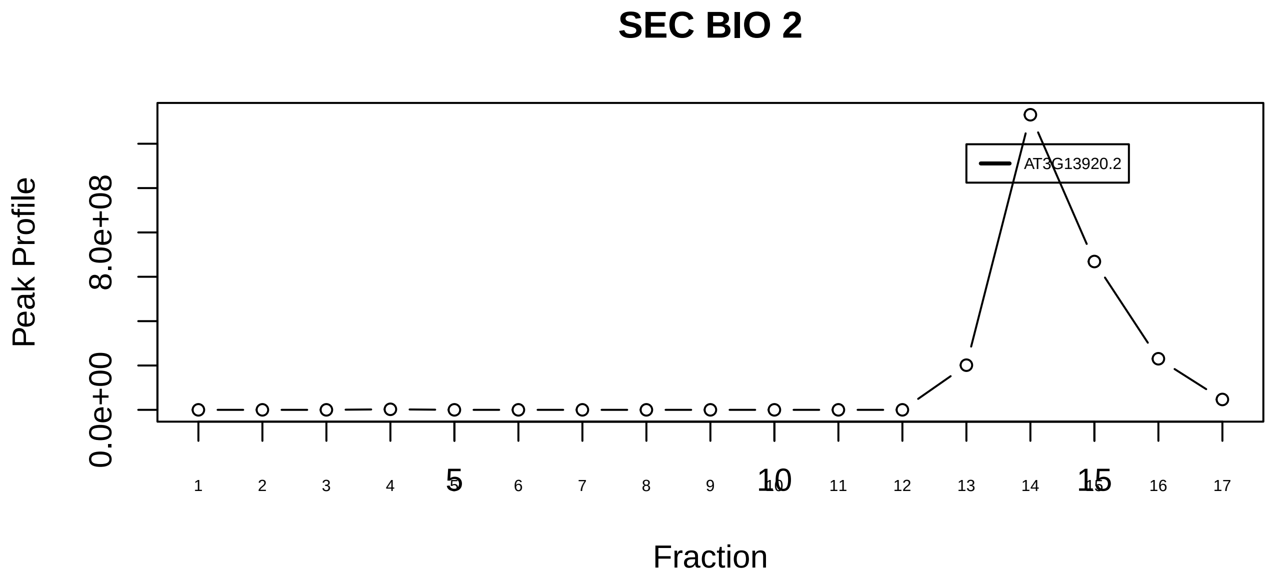

PPP2R1A−PPP2R1B−PPP2CA−PPME1−EIF4A1 complex|5876−−singleton_2

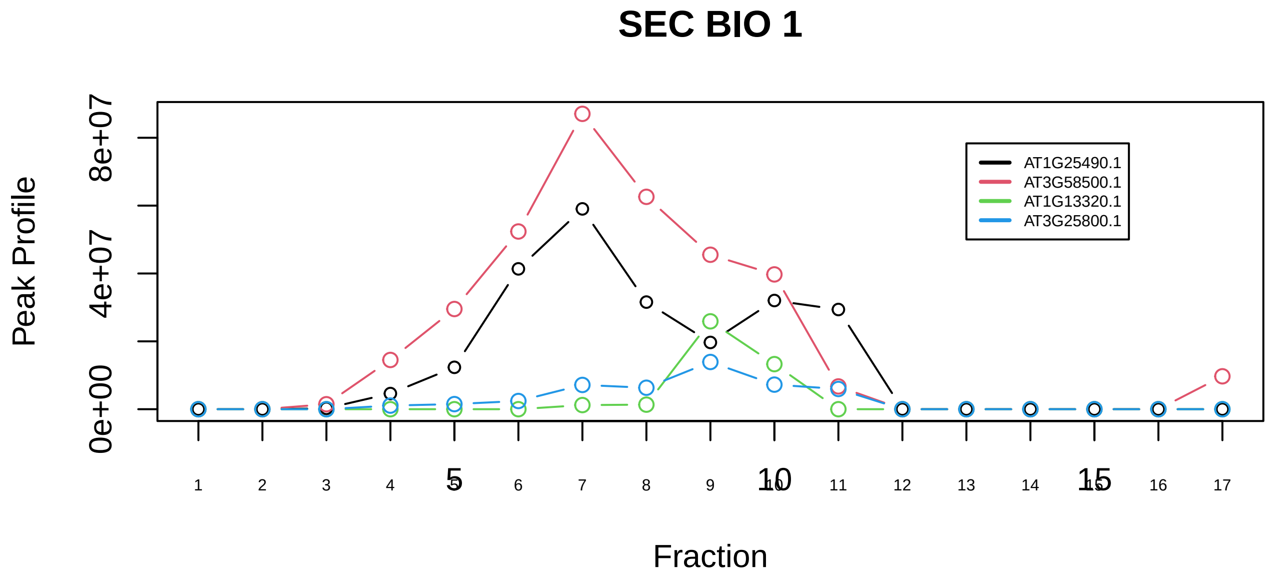

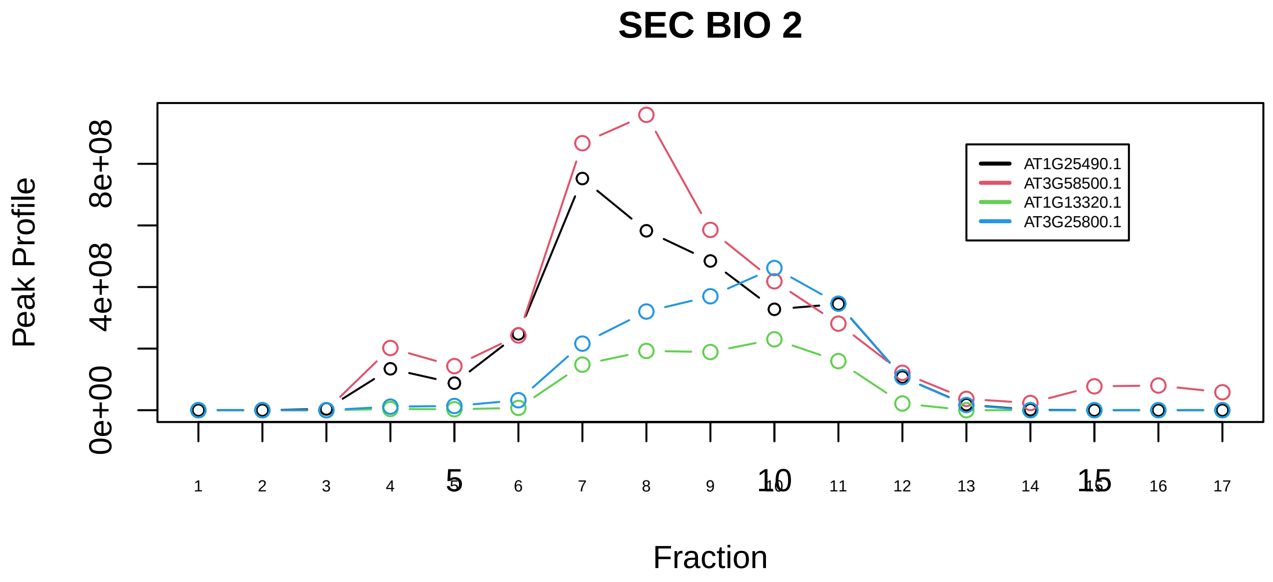

PPP2R1A−PPP2R1B−PPP2CA−PPME1−EIF4A1 complex|5876−−sub_1

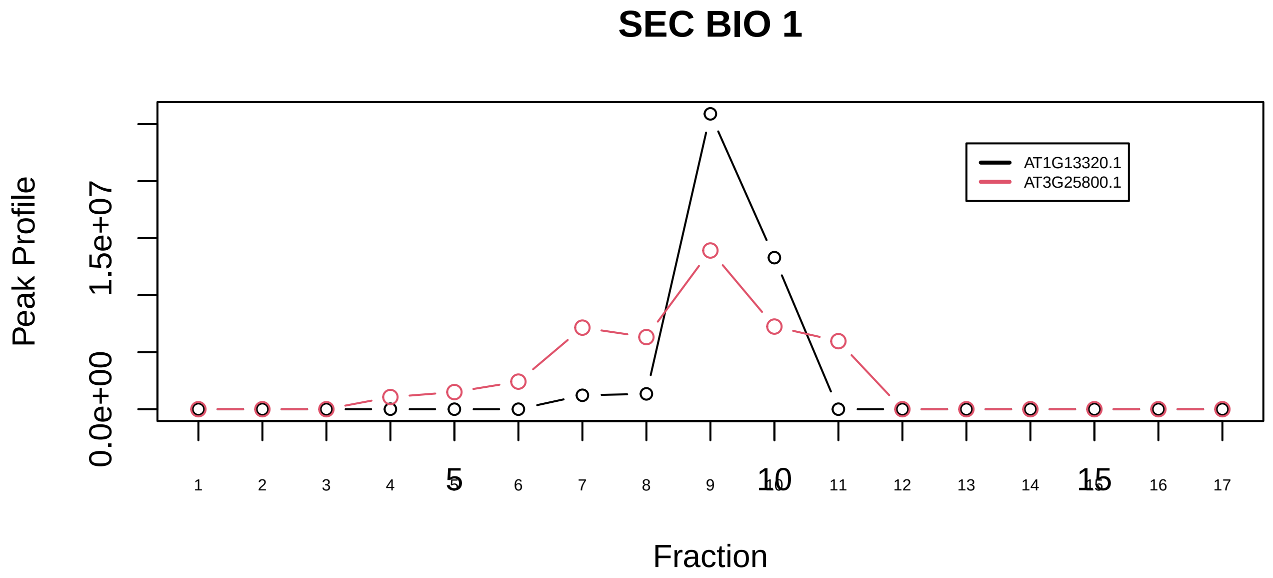

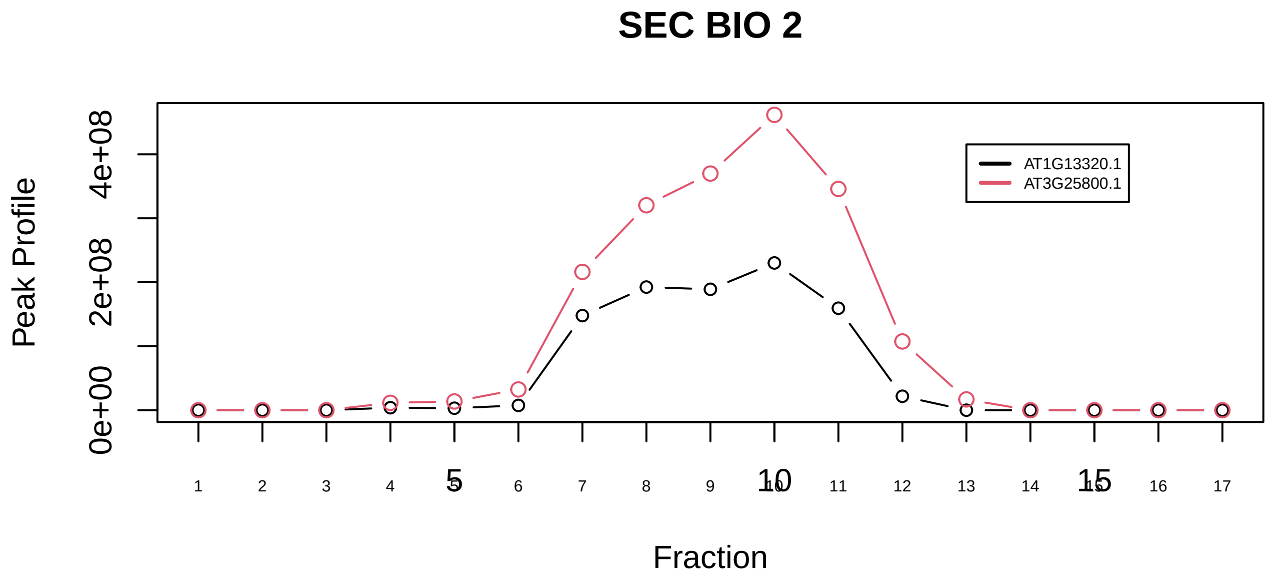

PPP2R1A−PPP2R3B complex|6180−−sub_orthoparalog_1

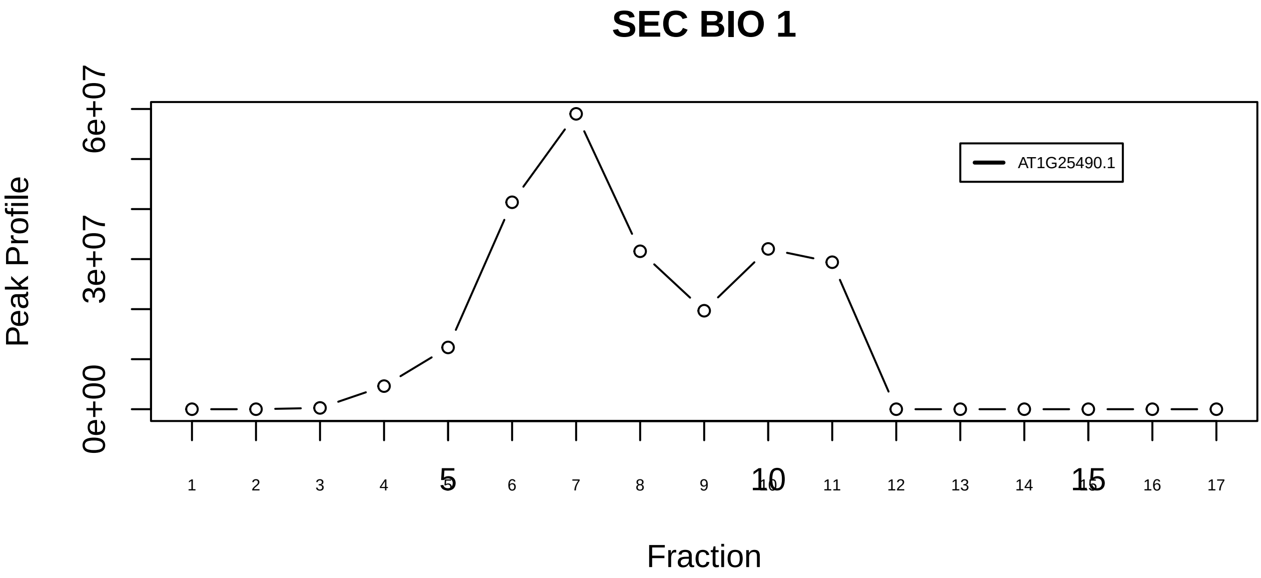

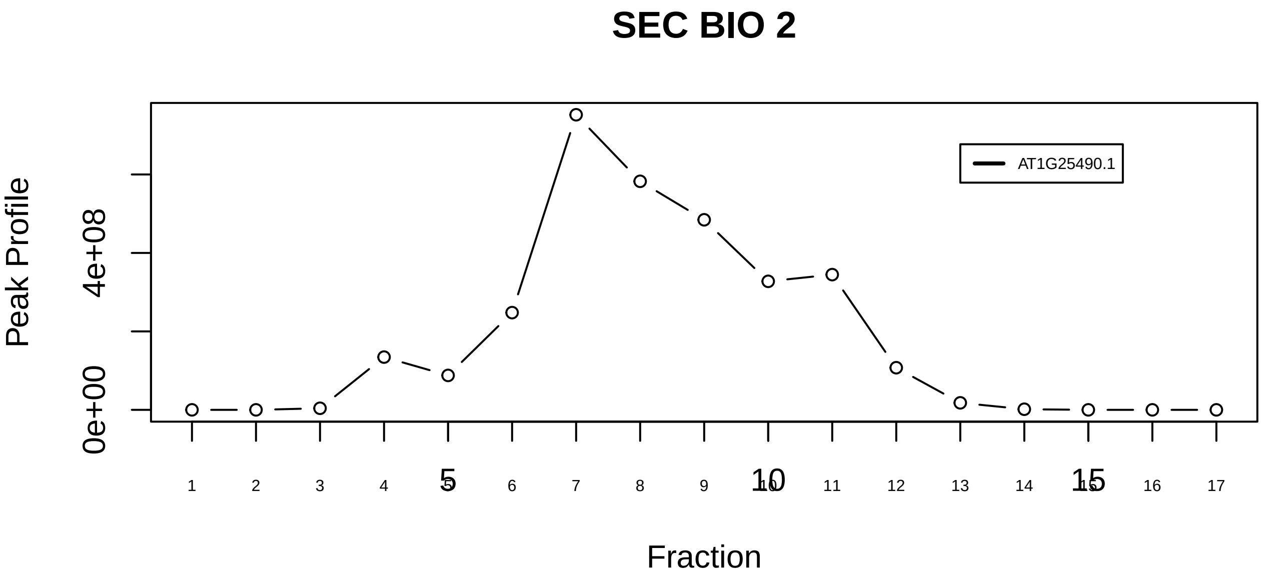

PPP2R1A−PPP2R3B complex|6180−−singleton_1

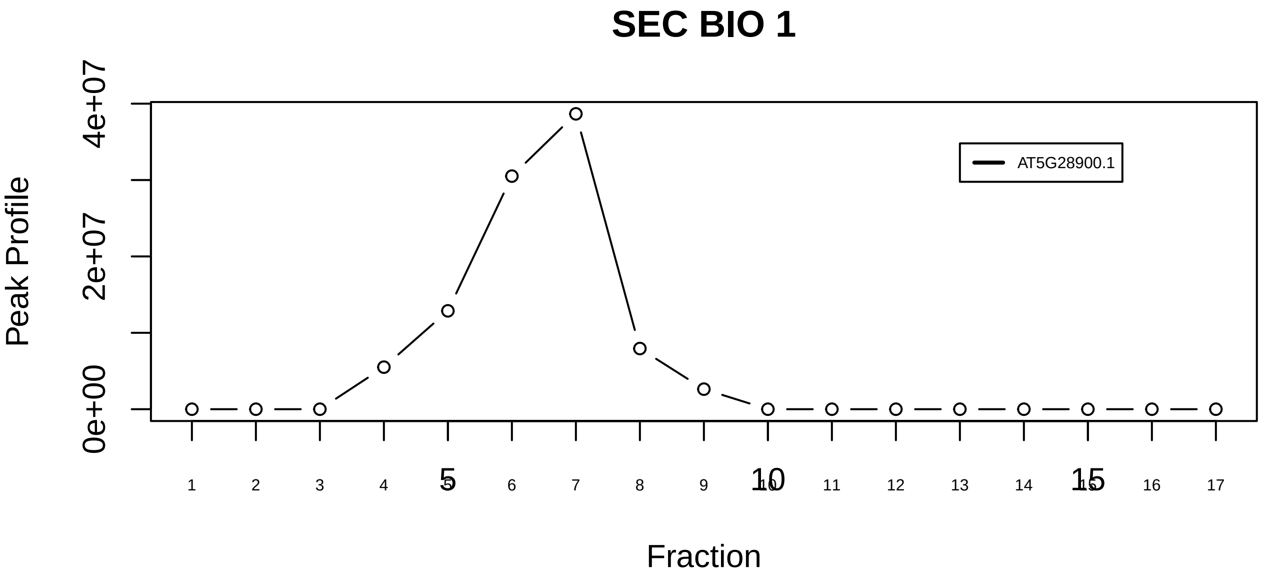

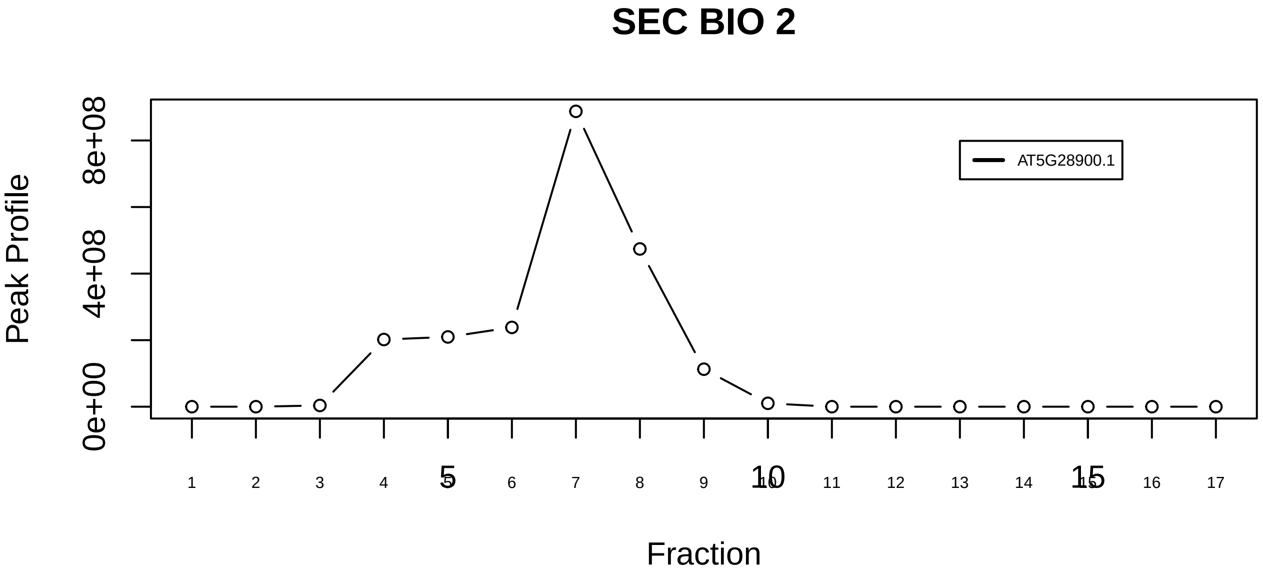

PPP2R1A−PPP2R3B complex|6180−−singleton_2

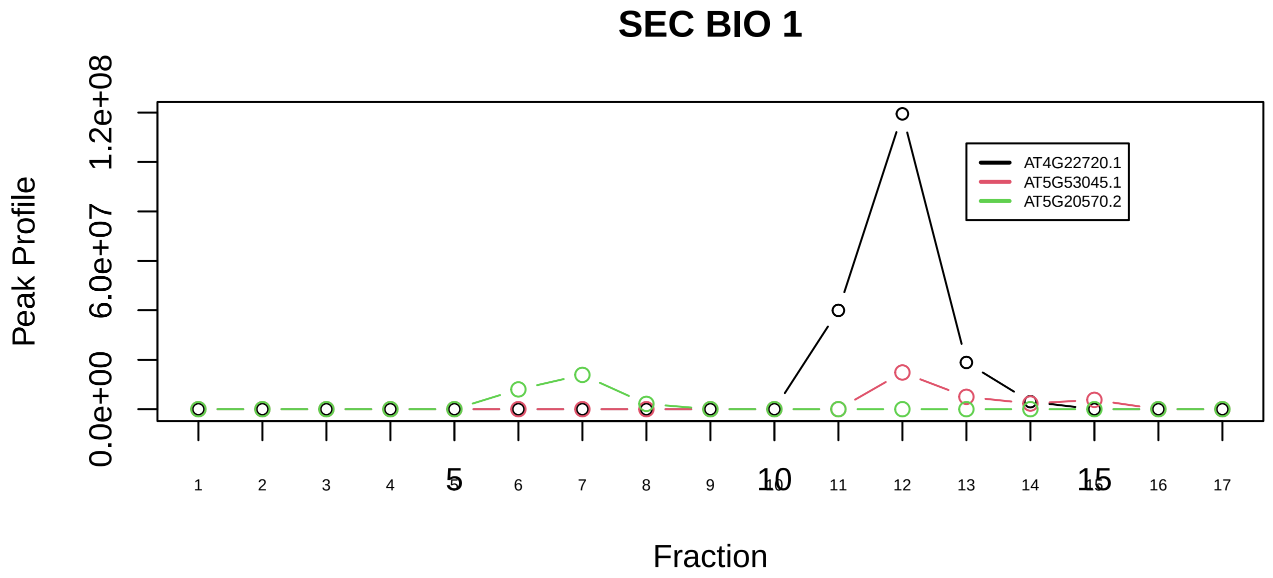

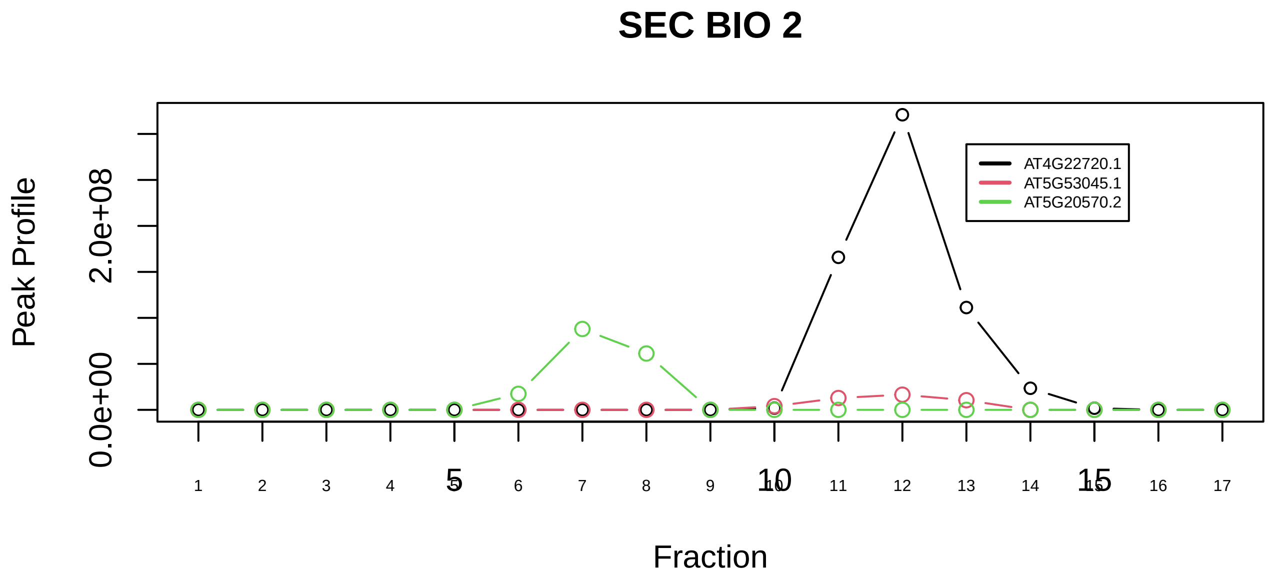

PRAME complex|7556−−sub_1

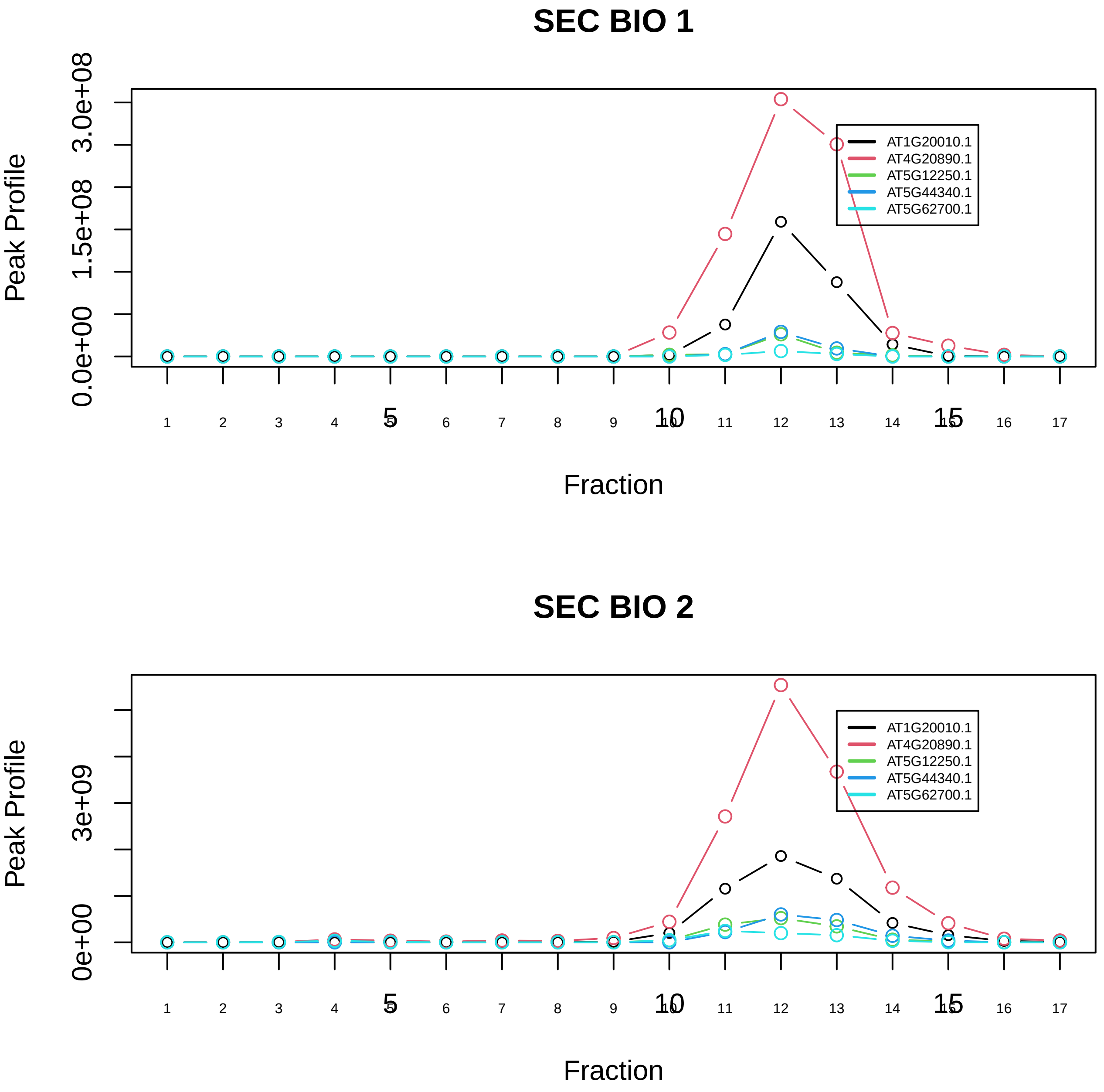

Profilin 1 complex|2837−−sub_orthoparalog_1

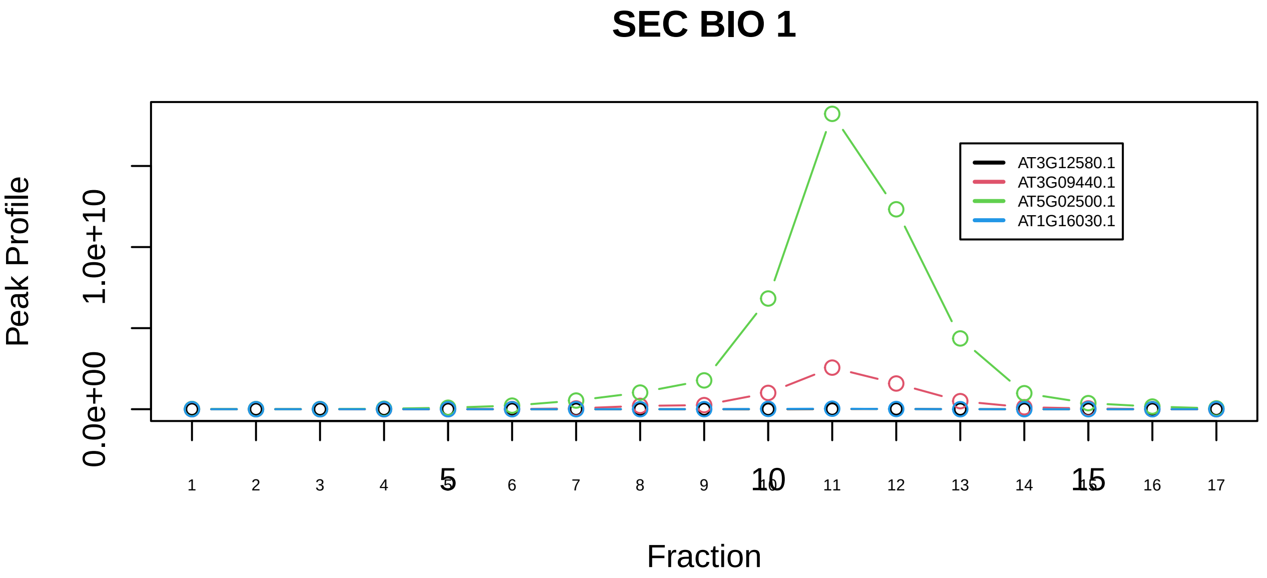

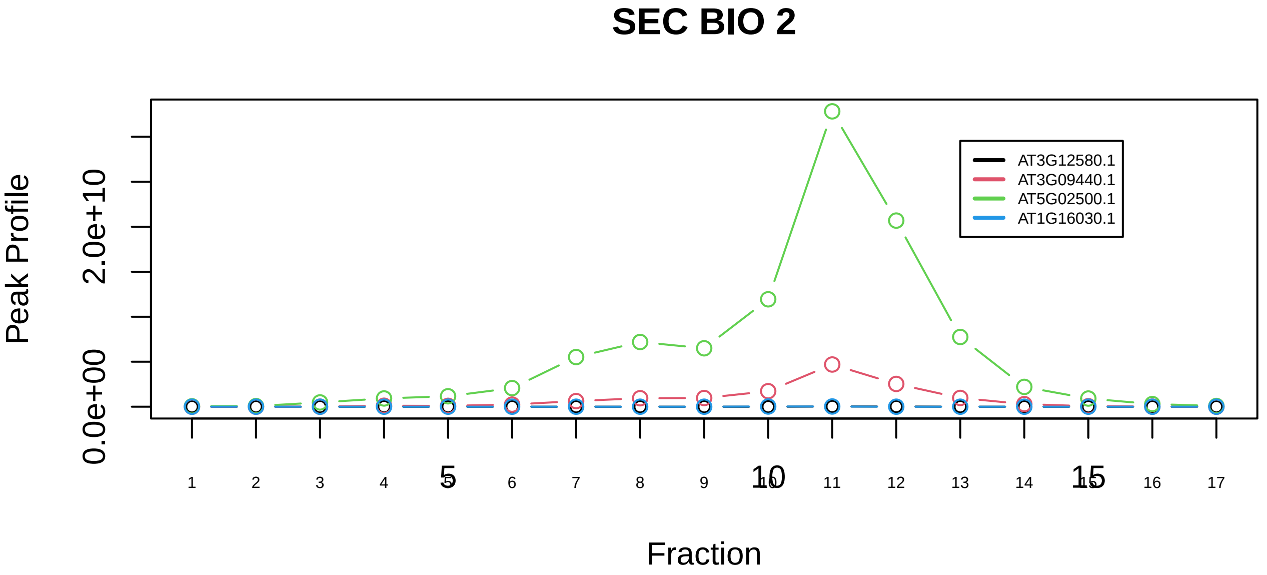

Profilin 1 complex|2837−−sub_orthoparalog_2

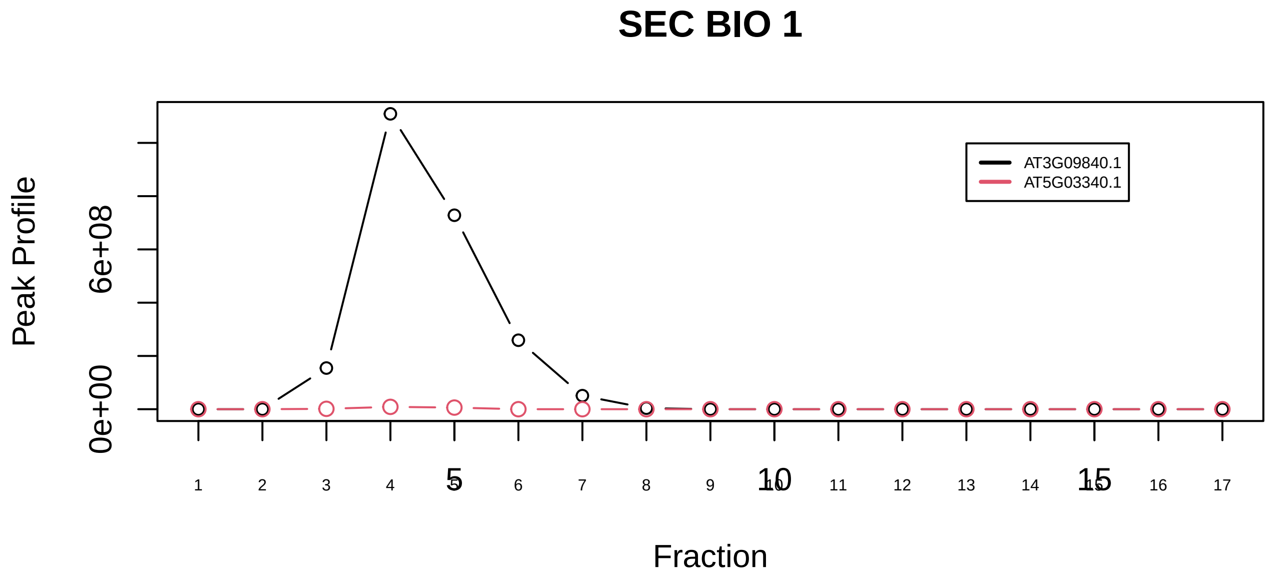

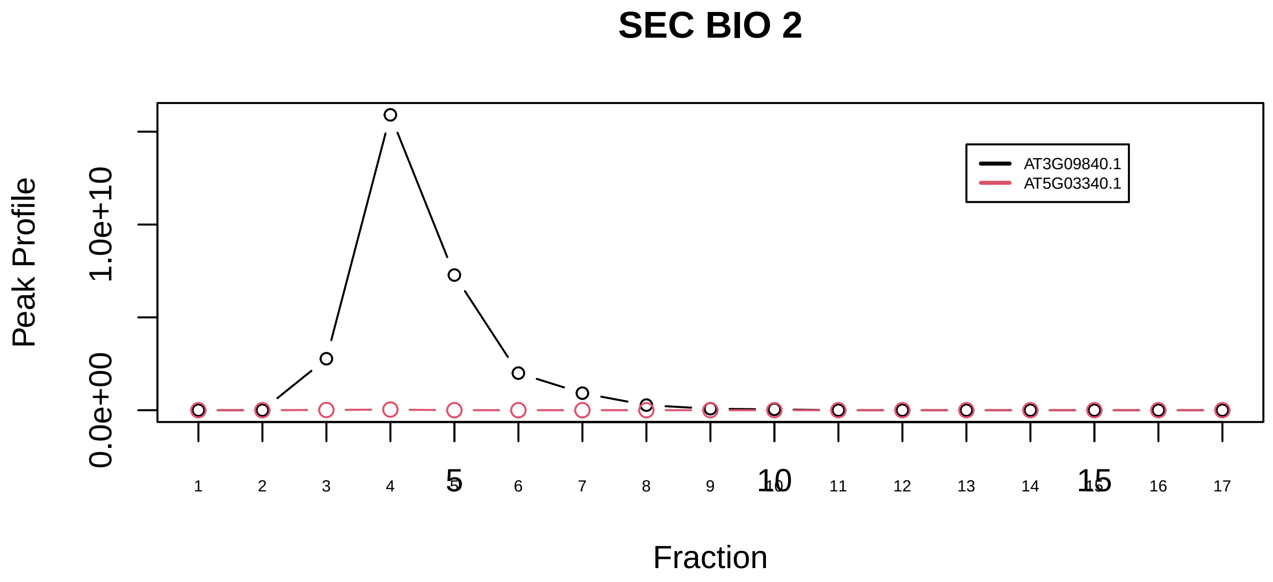

Profilin 1 complex|2837−−sub_orthoparalog_3

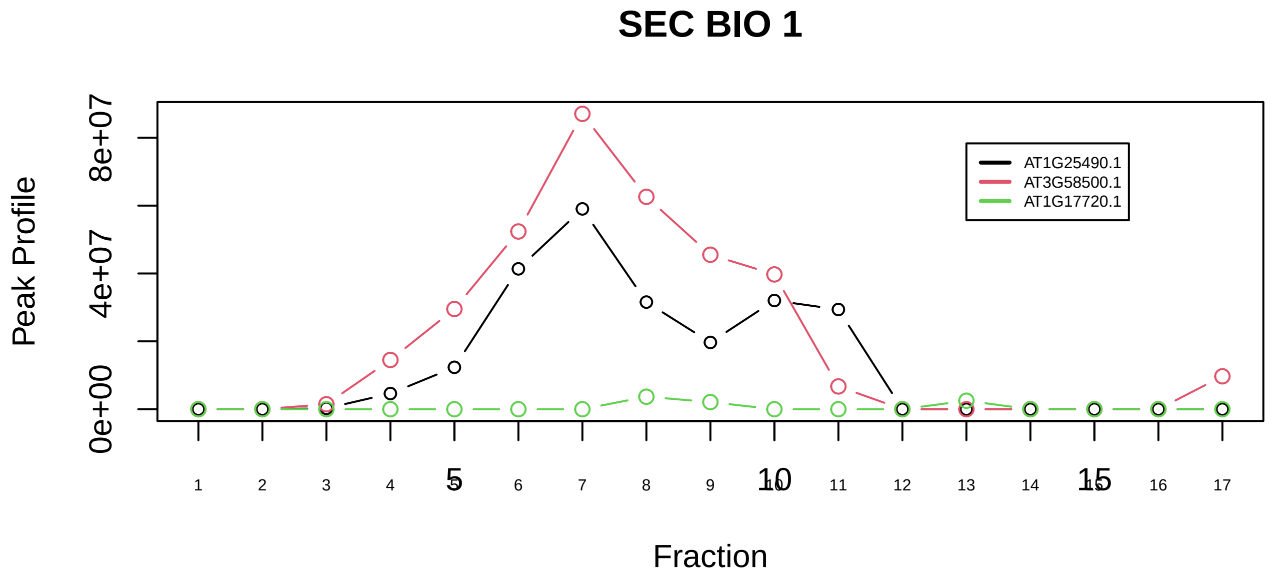

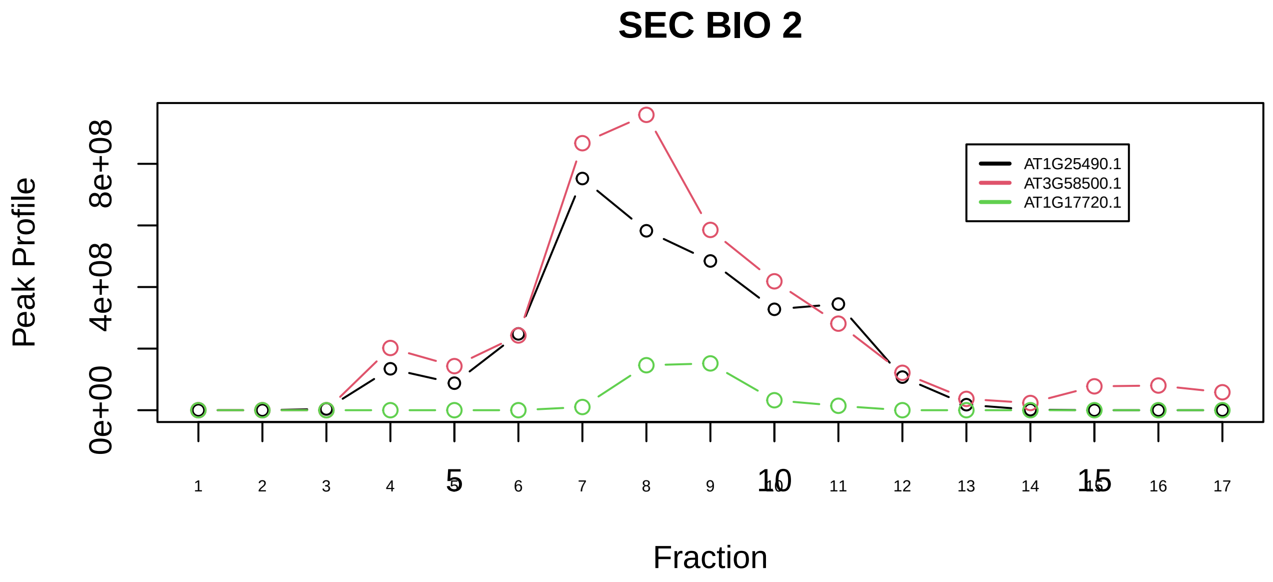

RAF1−PPP2−PIN1 complex|5211−−sub_1

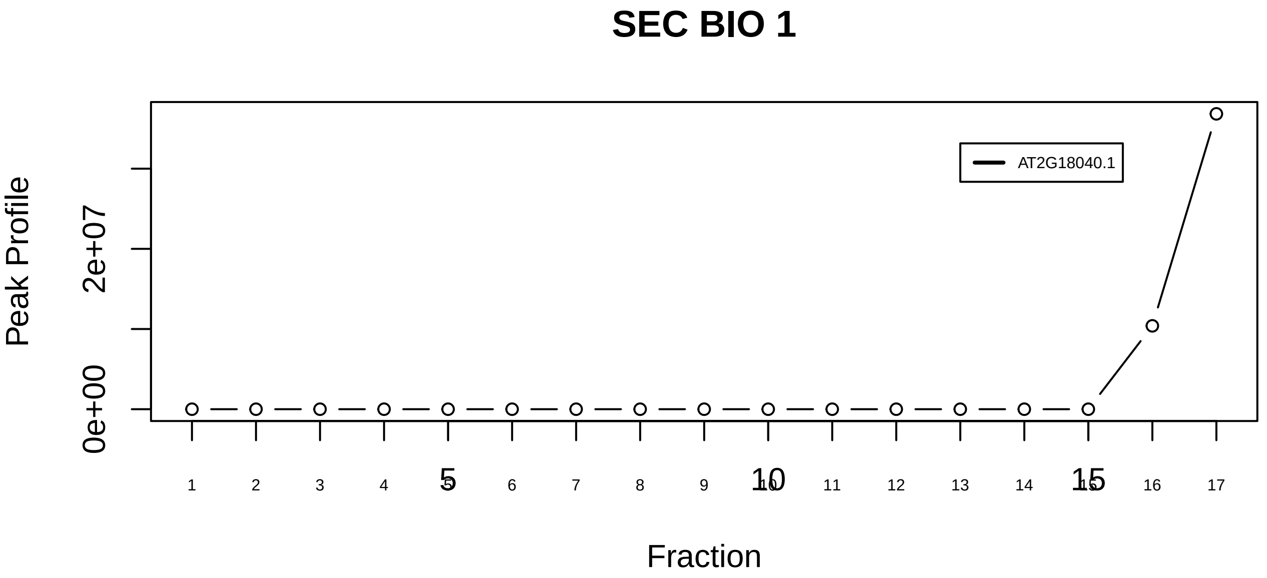

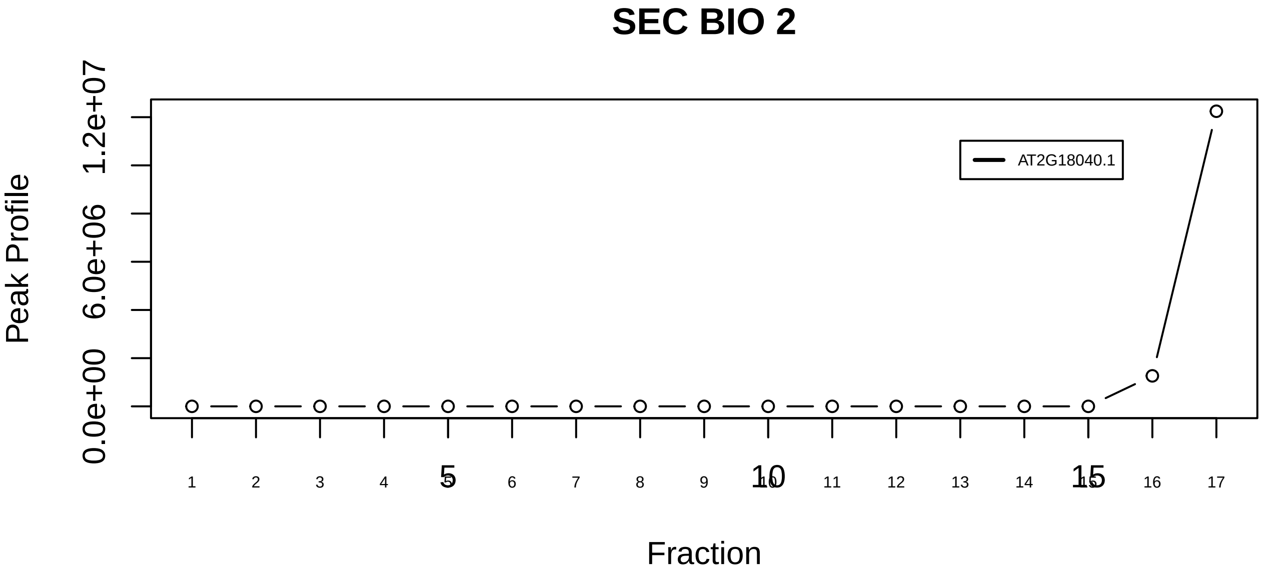

RAF1−PPP2−PIN1 complex|5211−−singleton_1

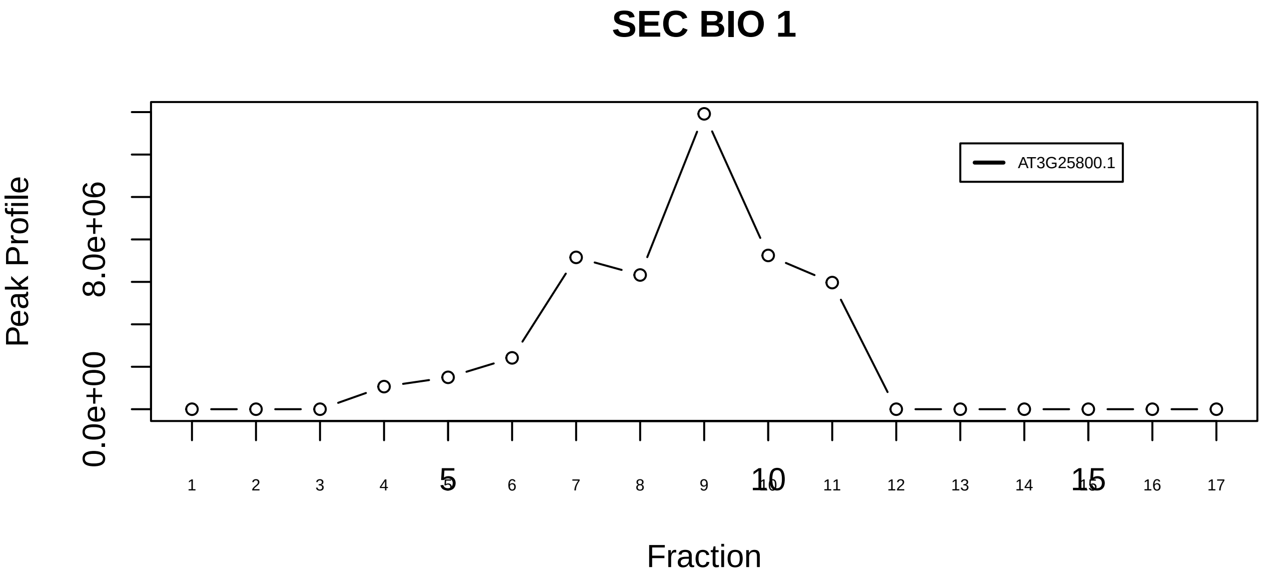

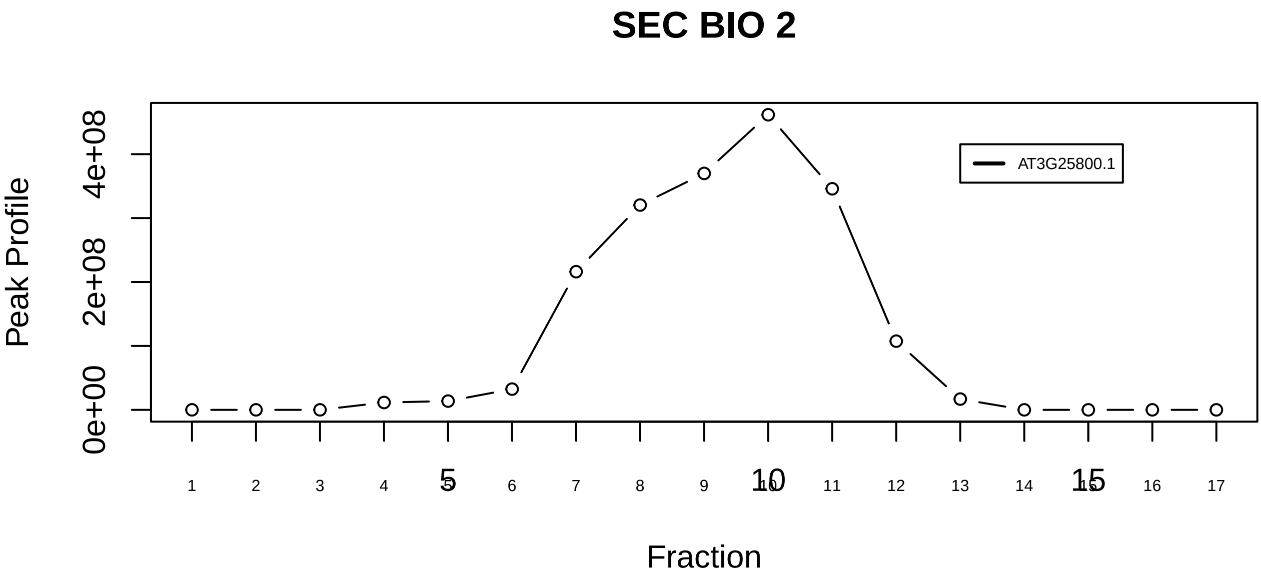

RAF1−PPP2−PIN1 complex|5211−−singleton_2

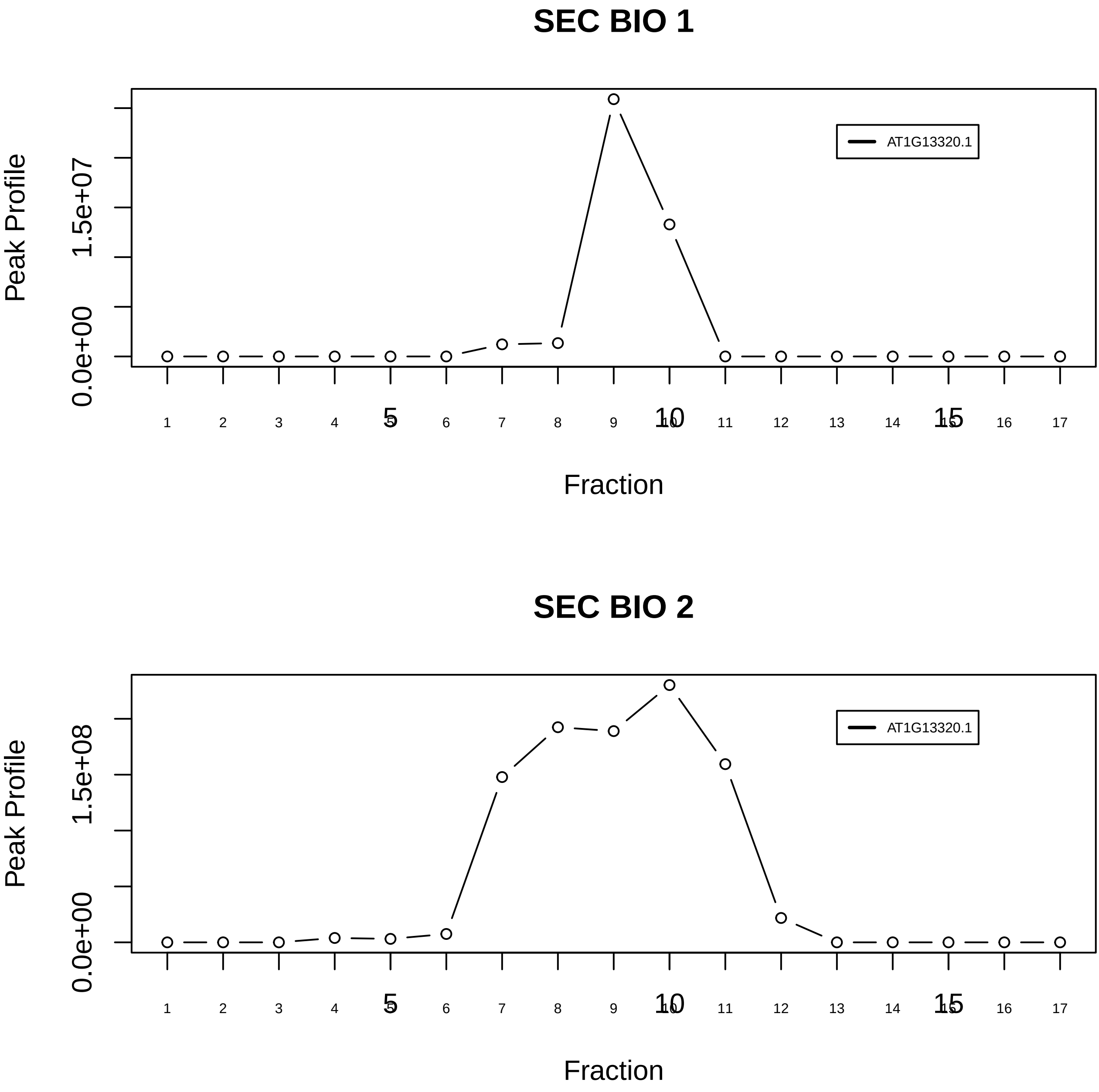

RAF1−PPP2−PIN1 complex|5211−−singleton_3

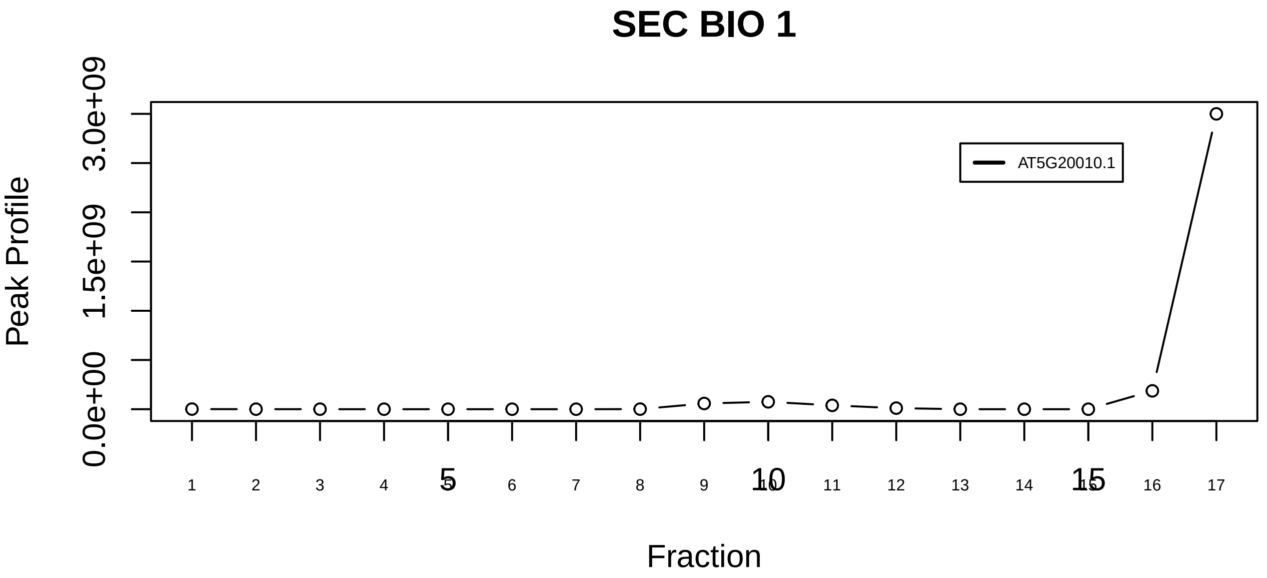

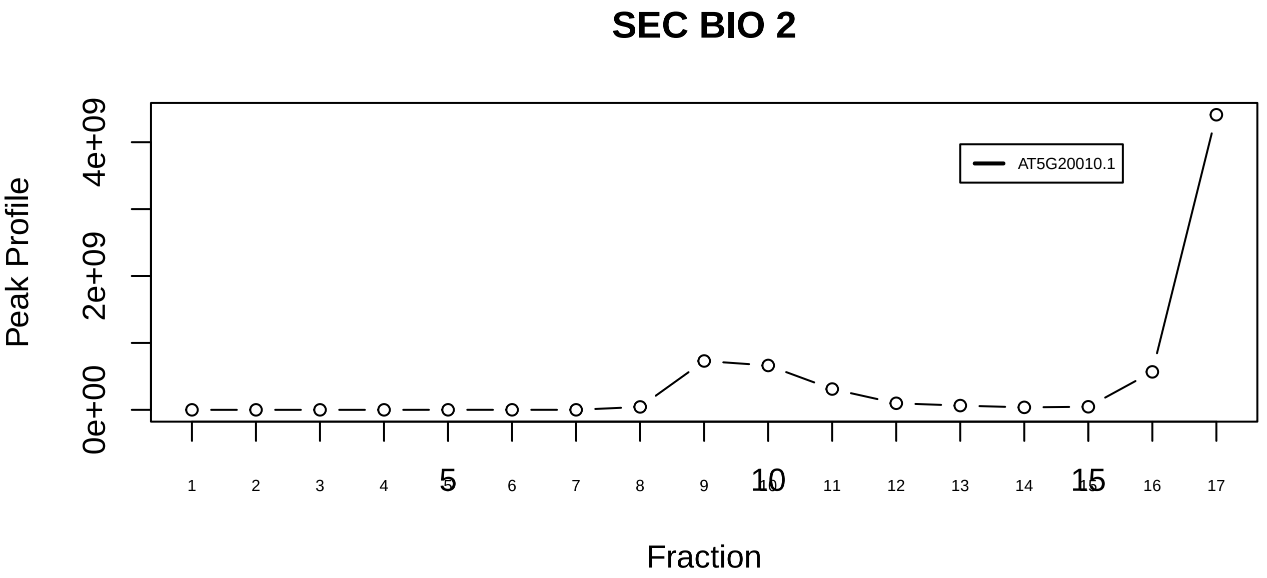

RANBP1−RAN−KPNB1 complex|1554−−singleton_1

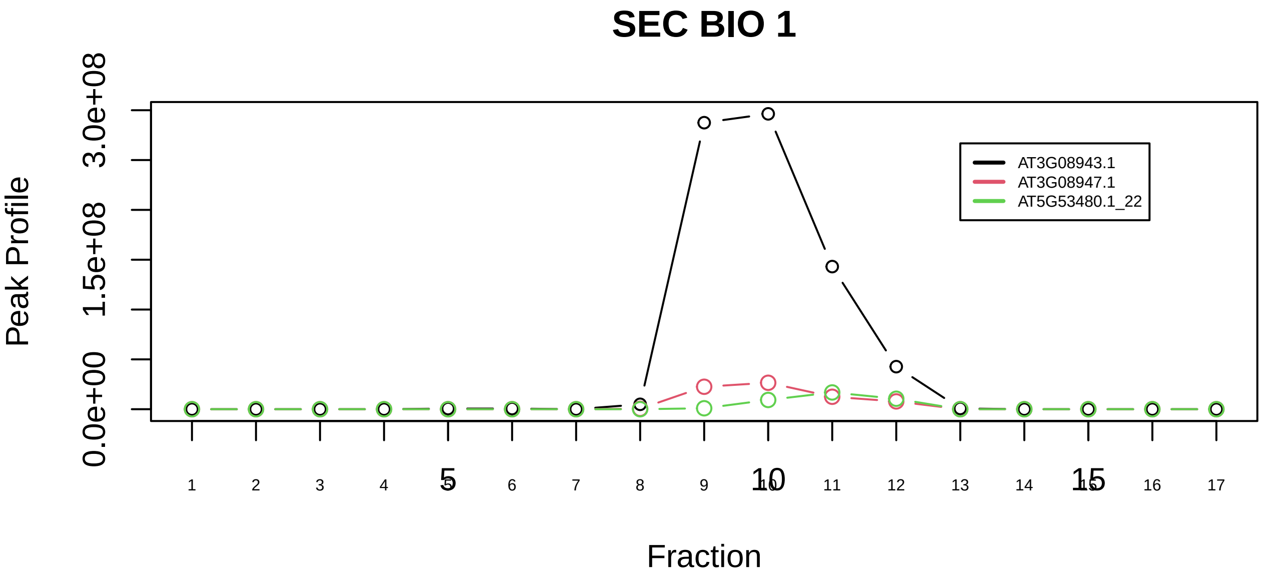

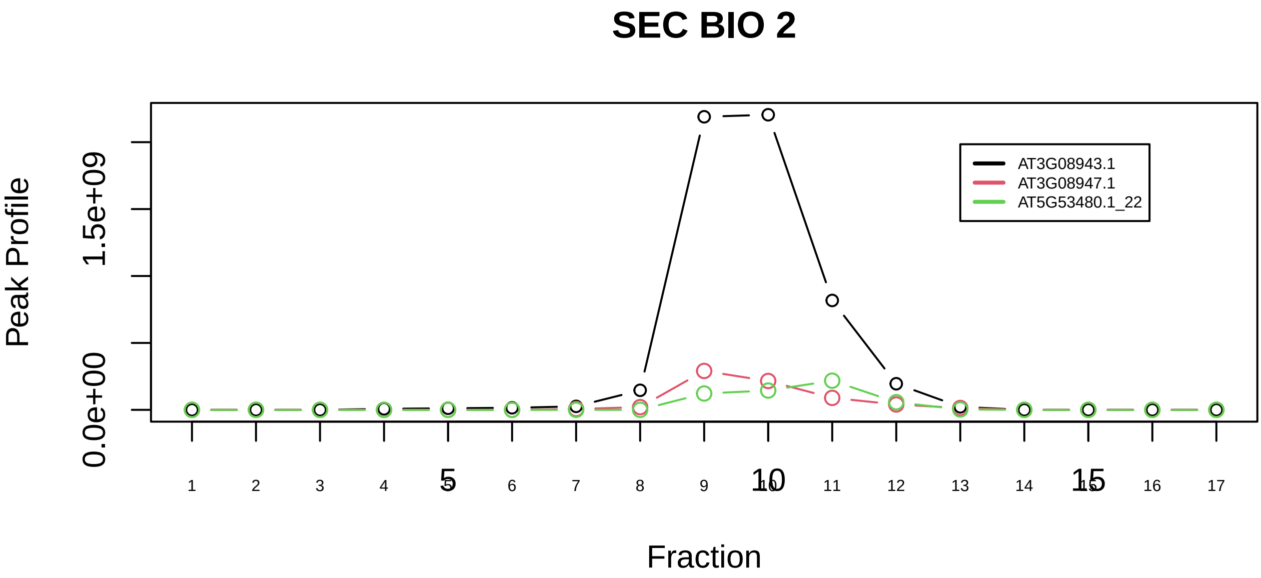

RANBP1−RAN−KPNB1 complex|1554−−sub_orthoparalog_1

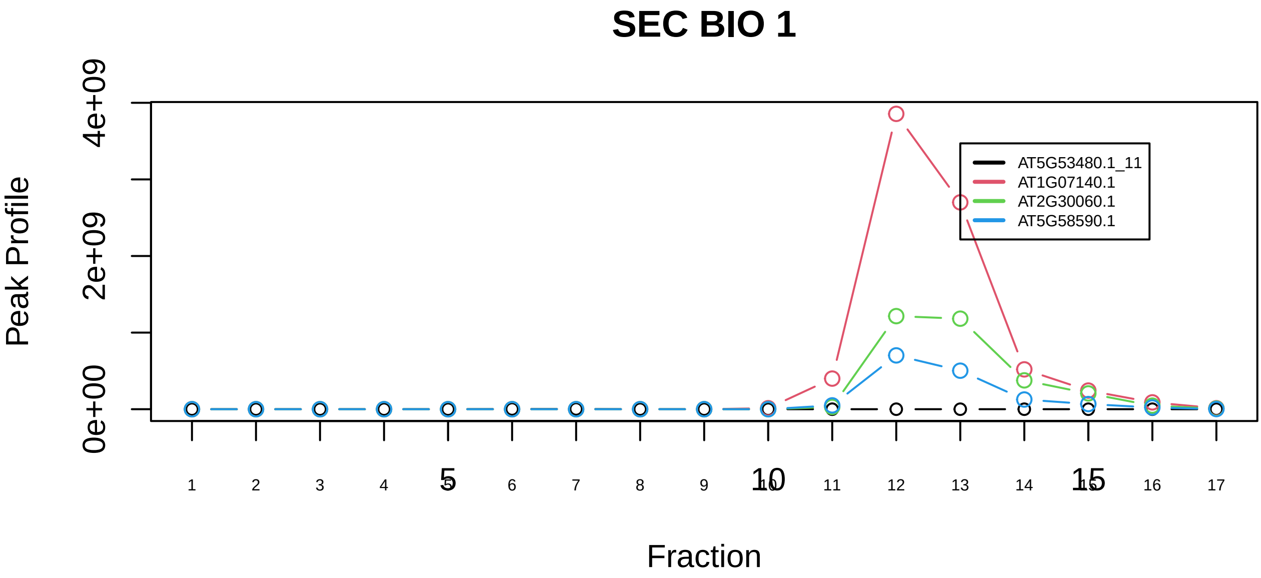

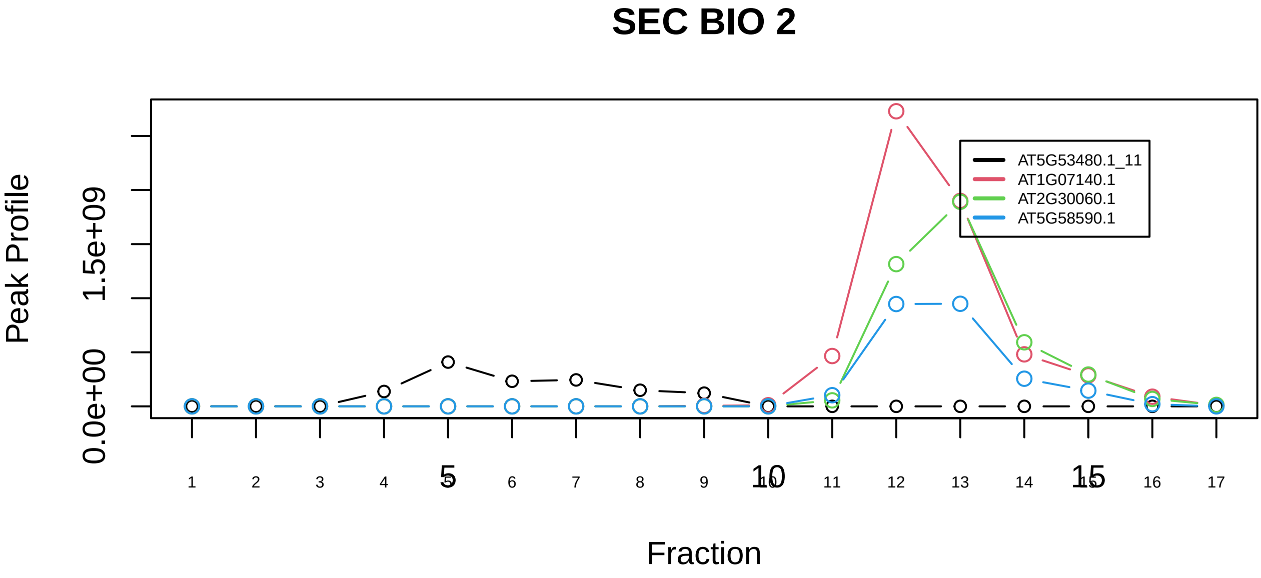

RANBP1−RAN−KPNB1 complex|1554−−sub_1

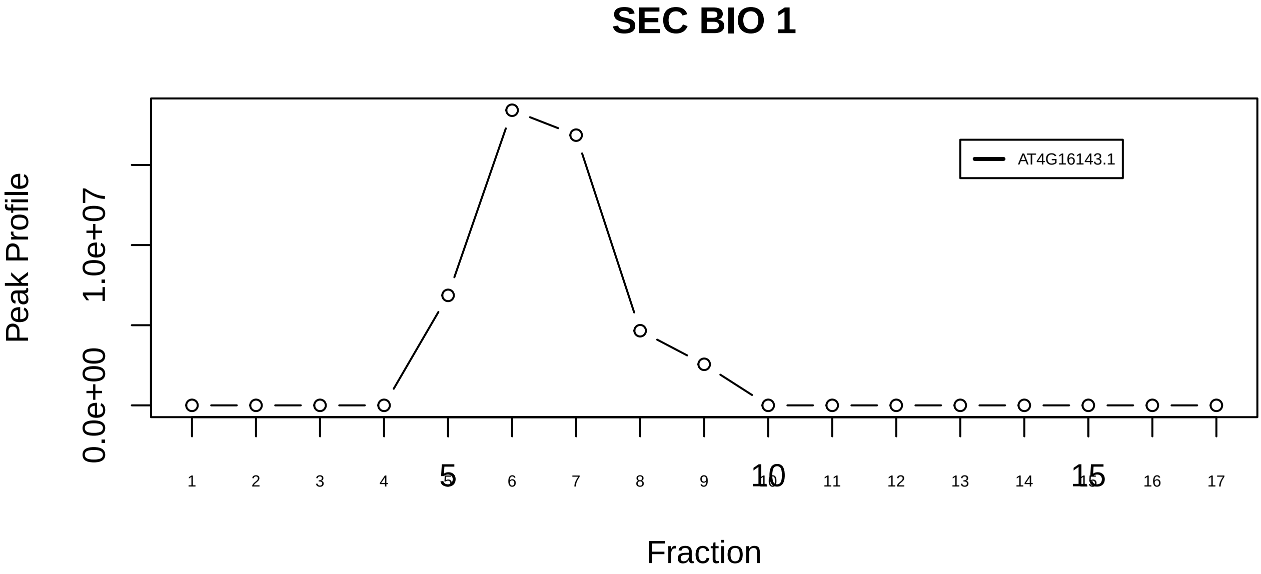

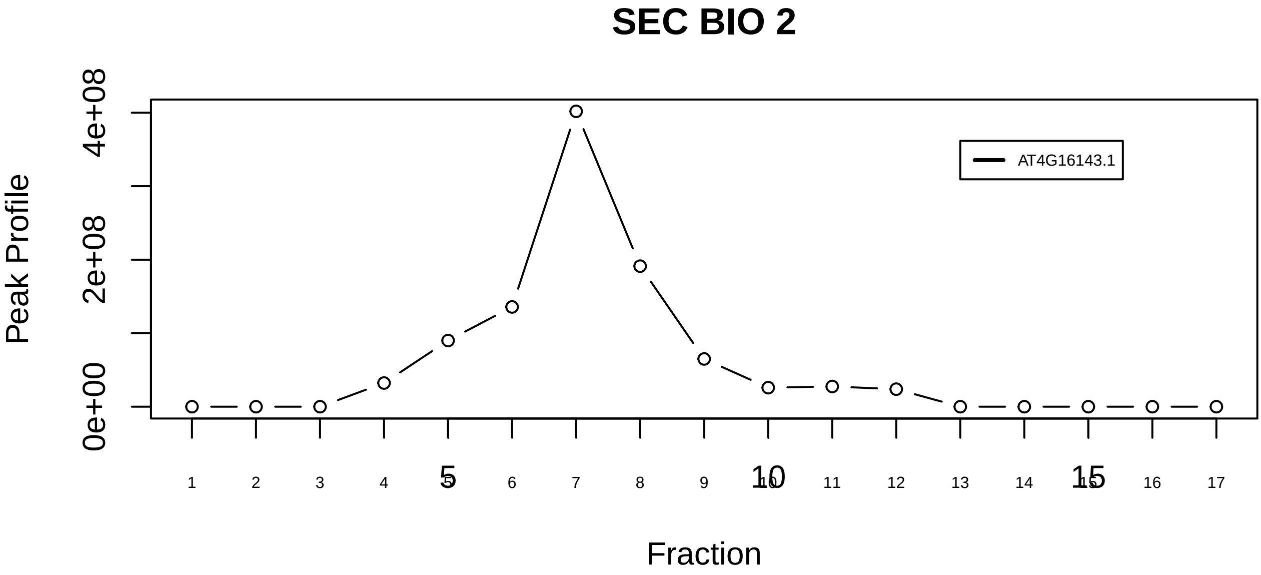

SERCA2a−alphaKAP−CaM−CaMKII complex|6517−−singleton_1

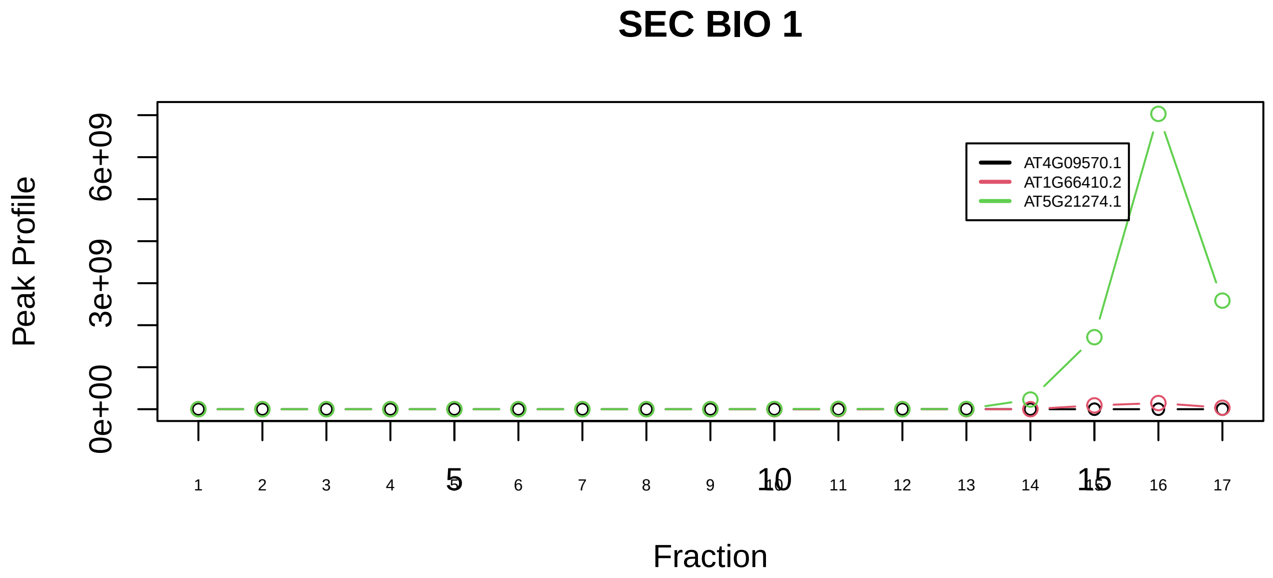

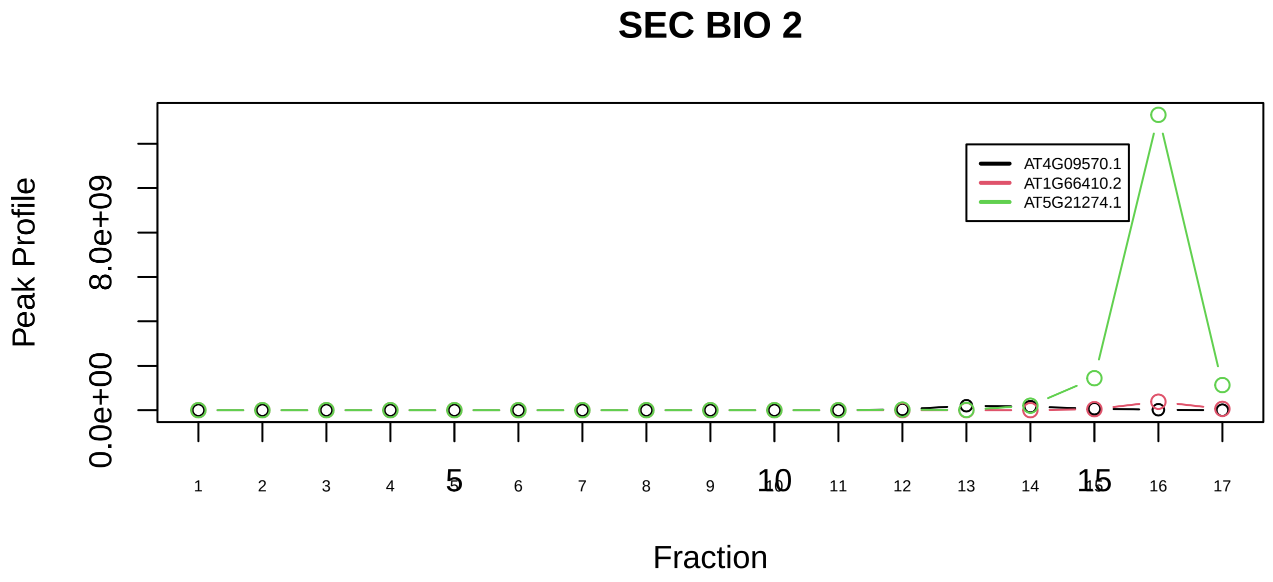

SERCA2a−alphaKAP−CaM−CaMKII complex|6517−−sub_1

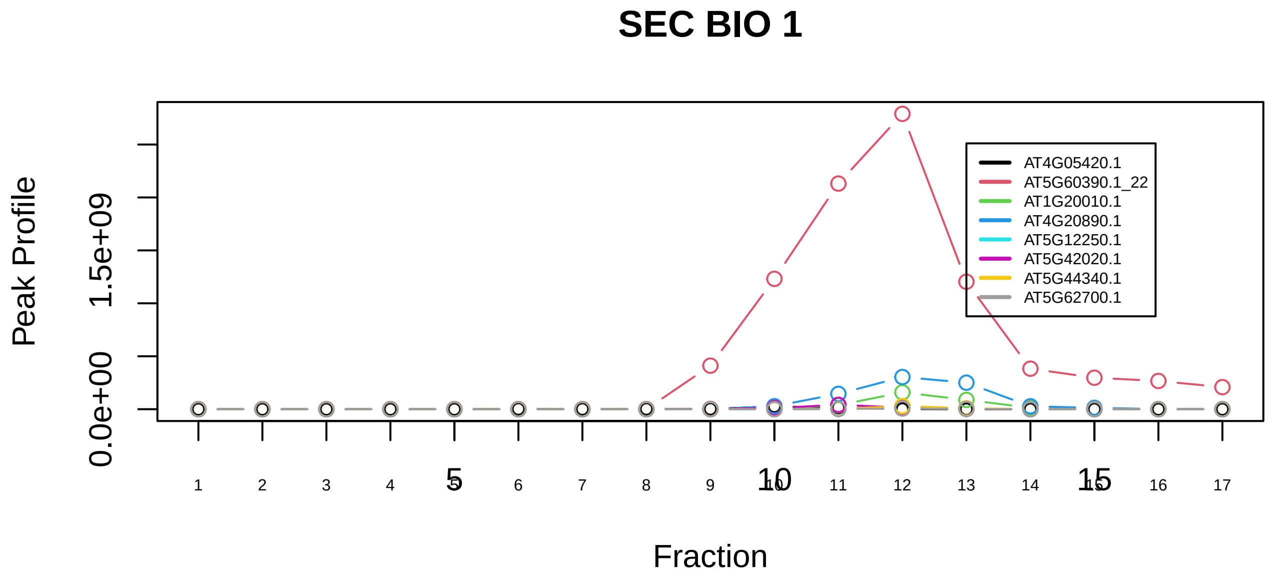

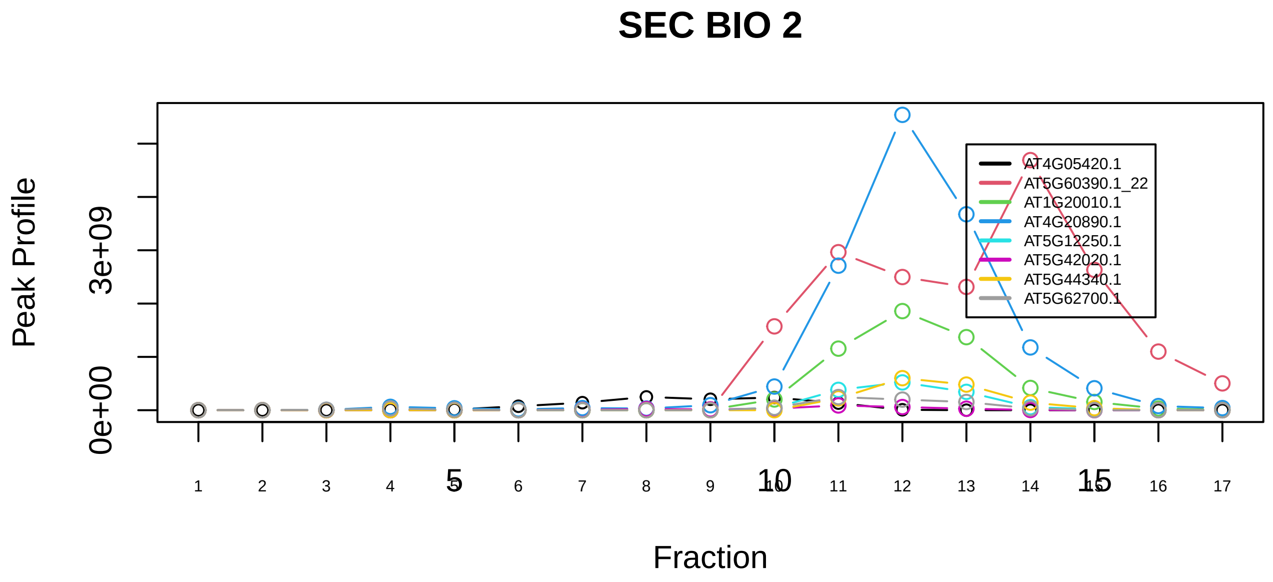

SNW1 complex|1335−−sub_1

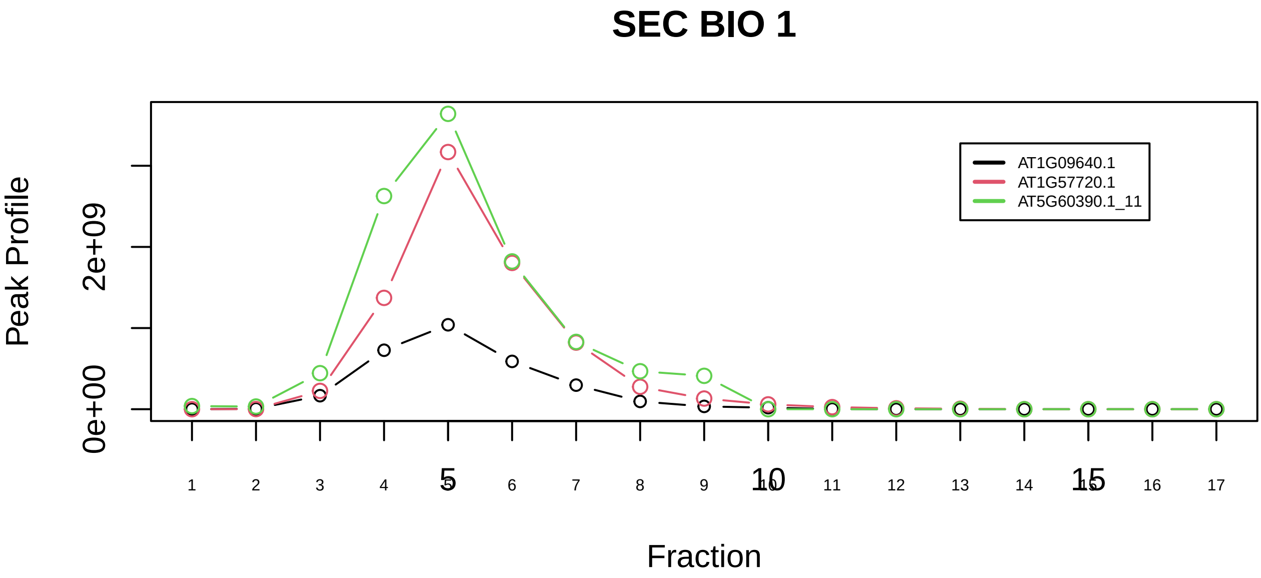

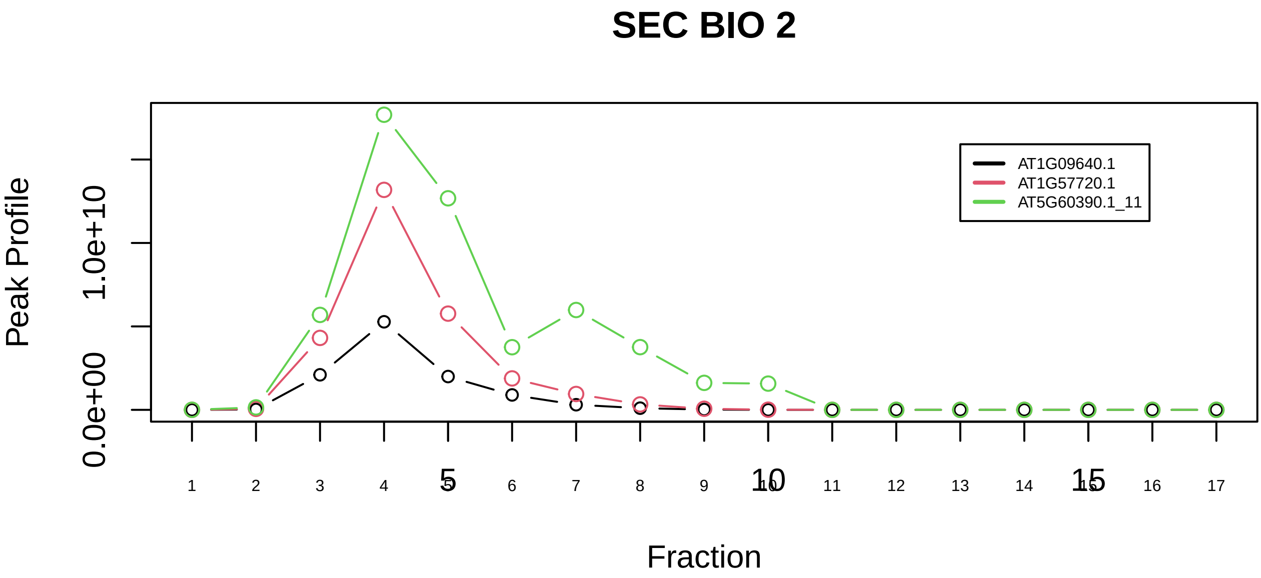

SNW1 complex|1335−−sub_2

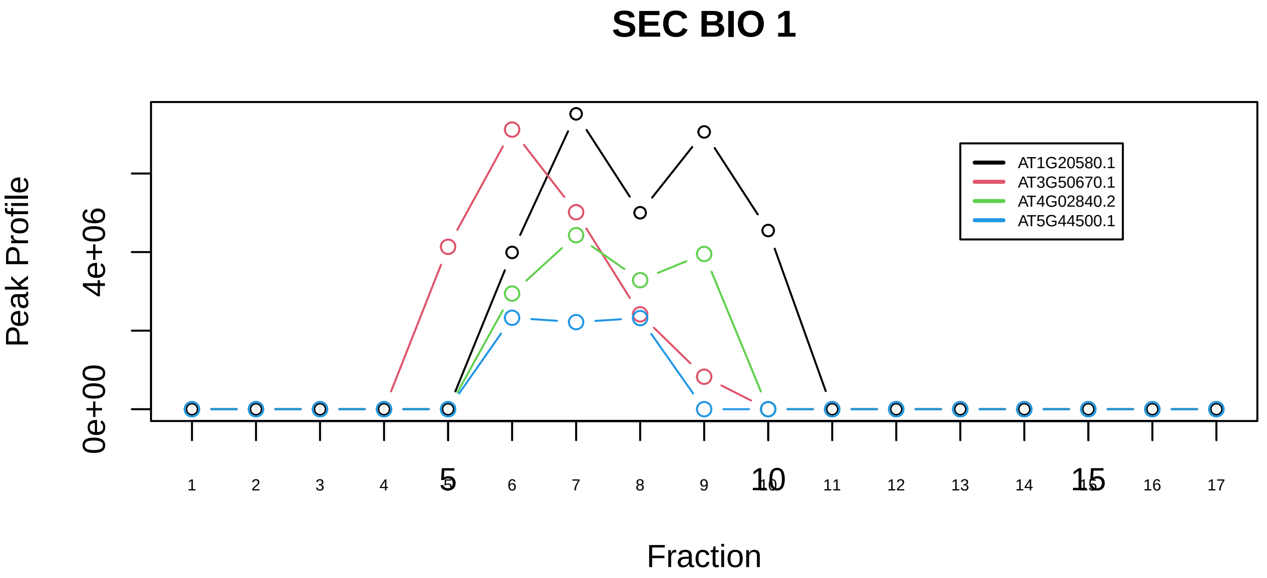

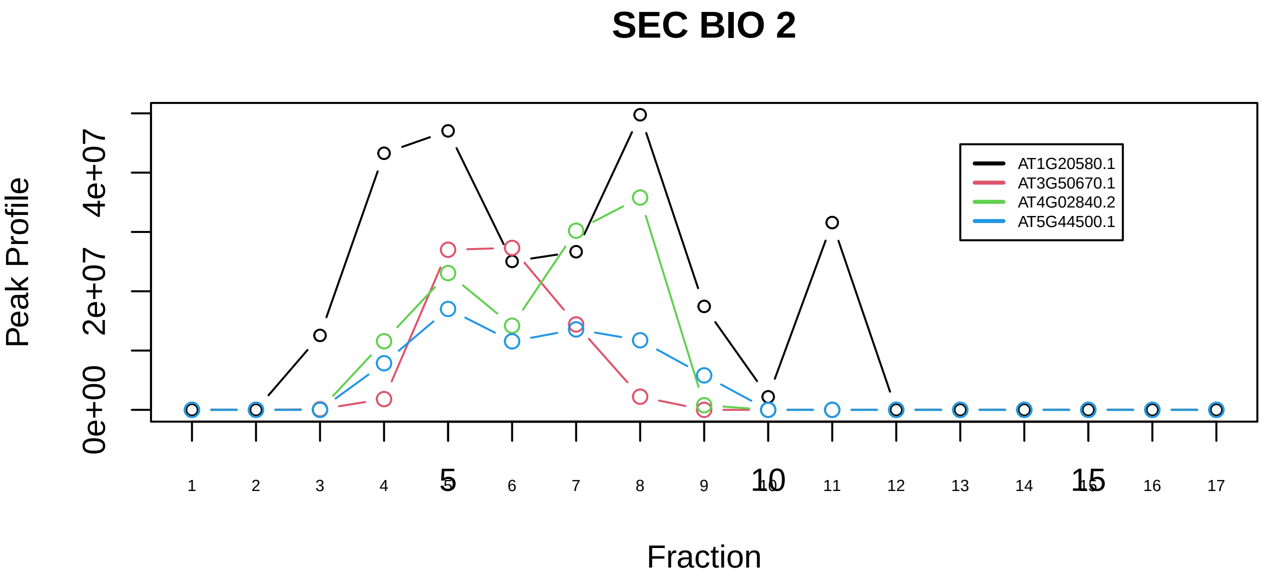

Spliceosome|351−−sub_1

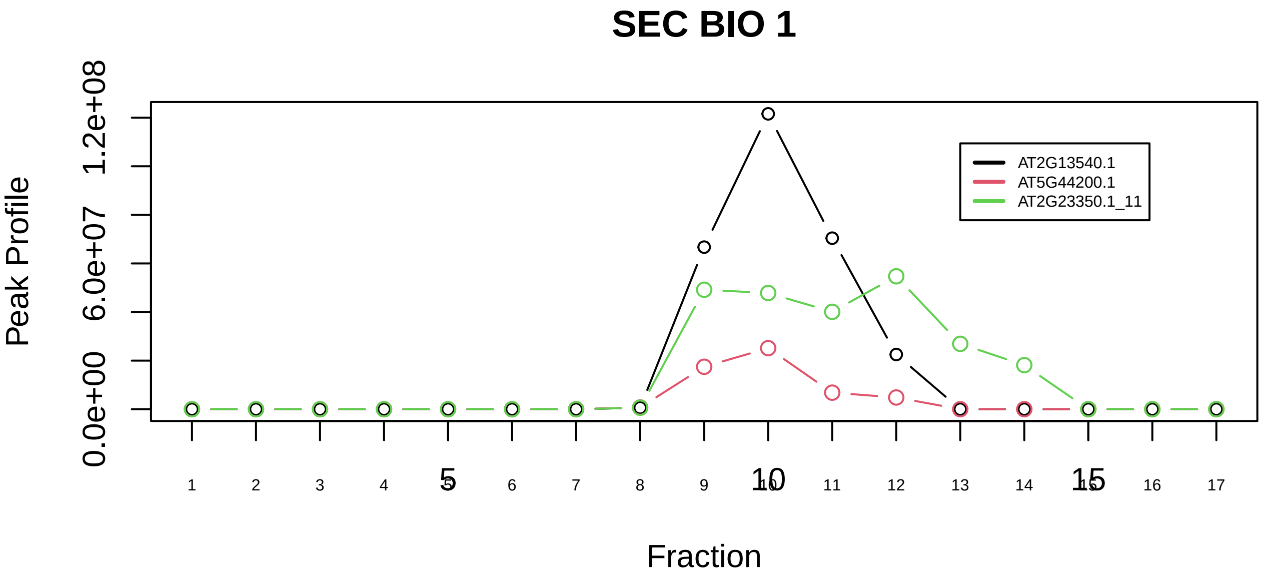

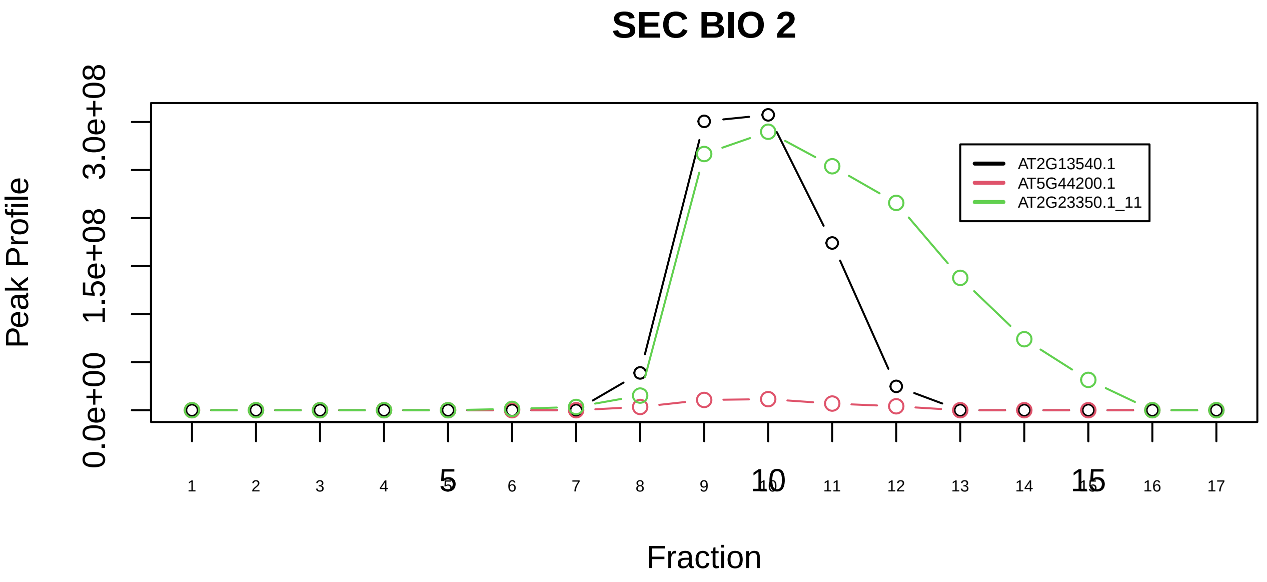

Spliceosome|351−−sub_2

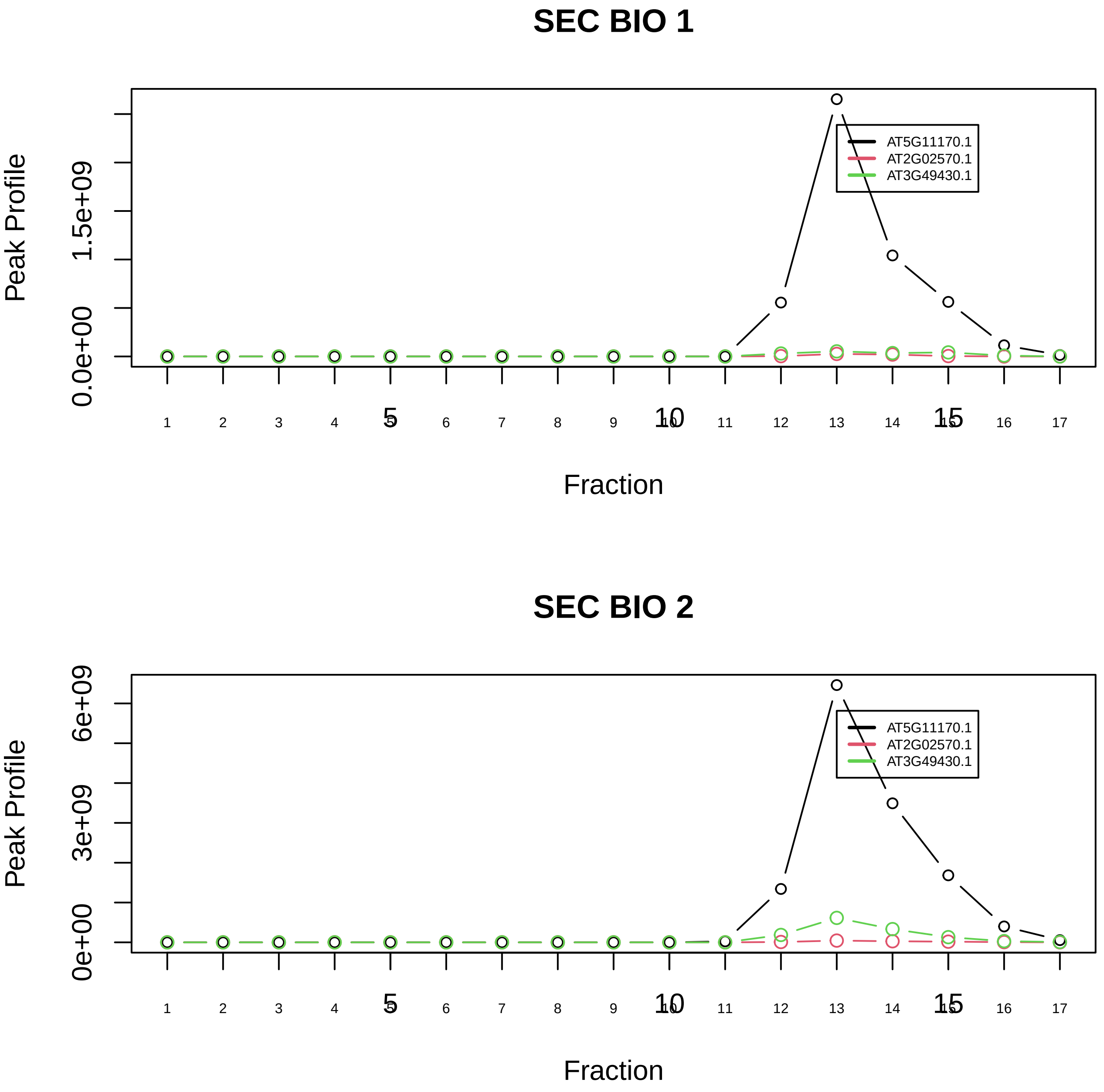

Spliceosome|351−−sub_3

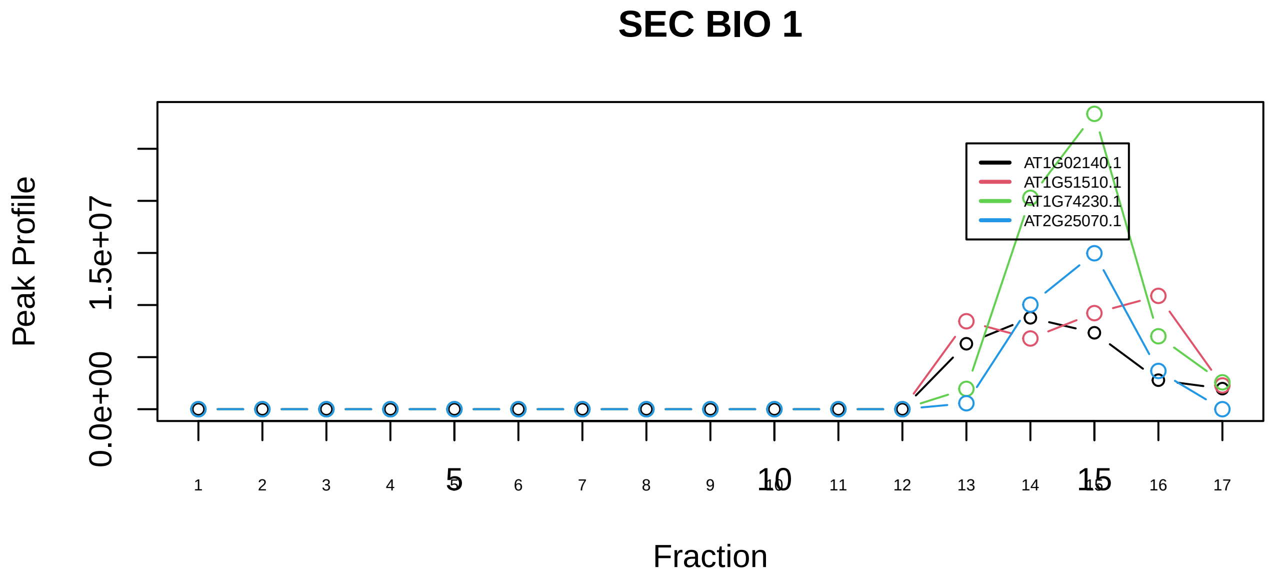

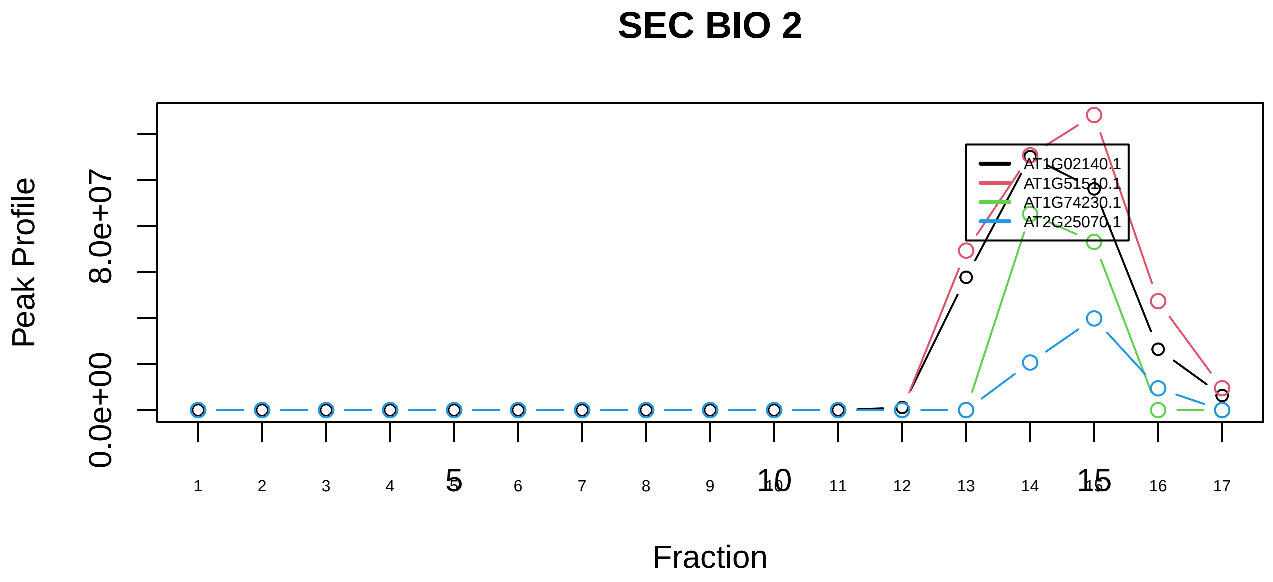

Spliceosome|351−−sub_4

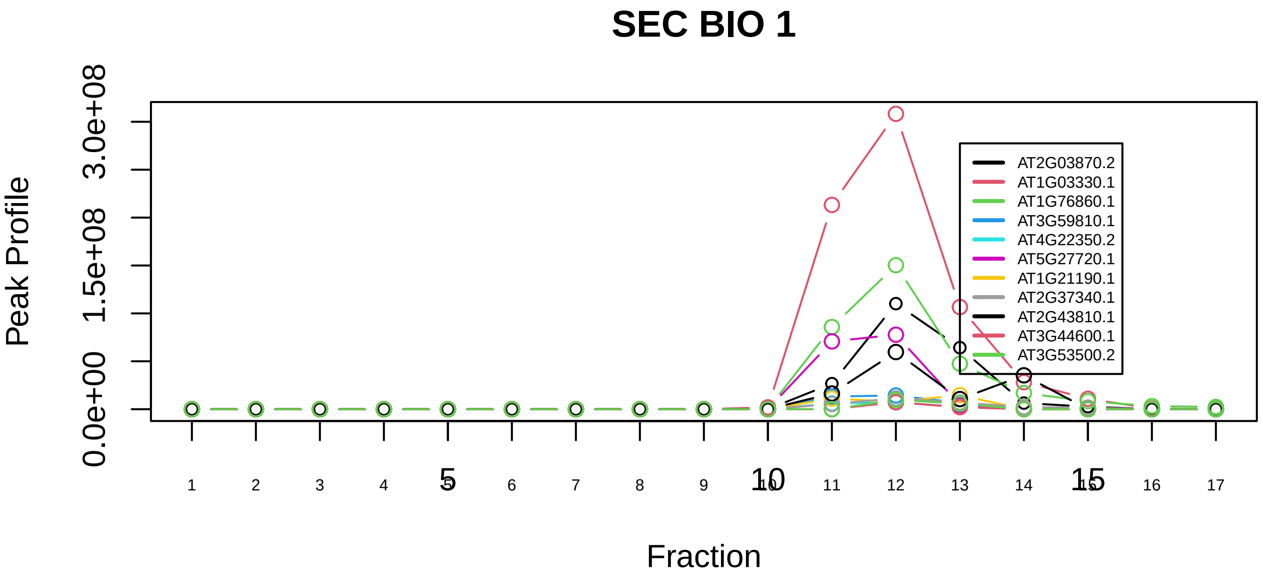

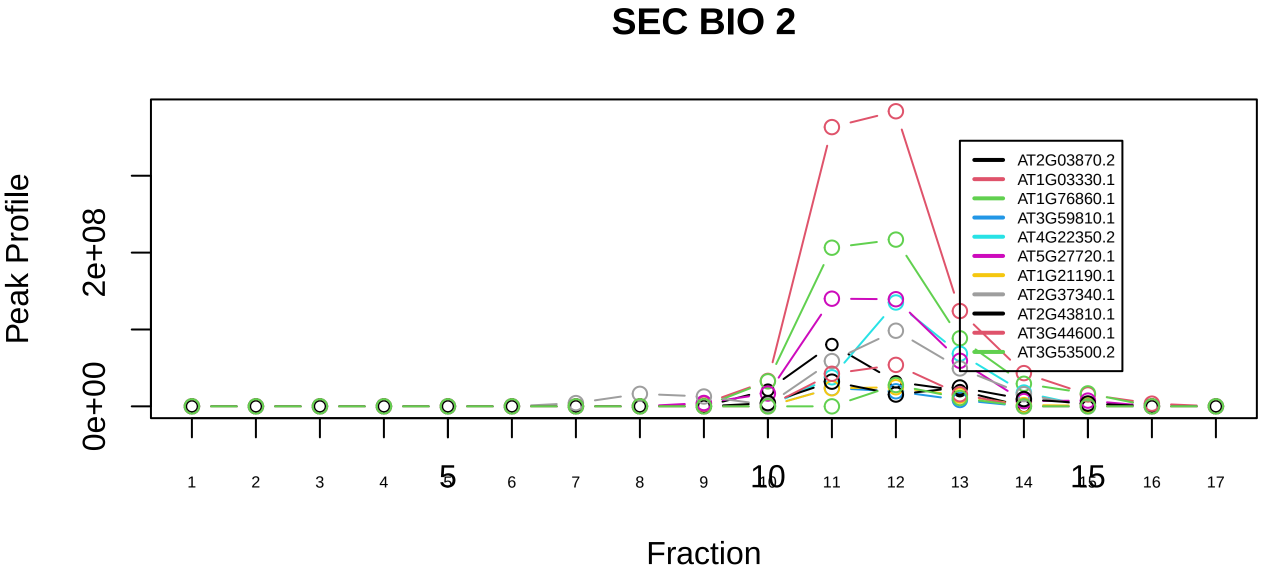

Spliceosome|351−−sub_5

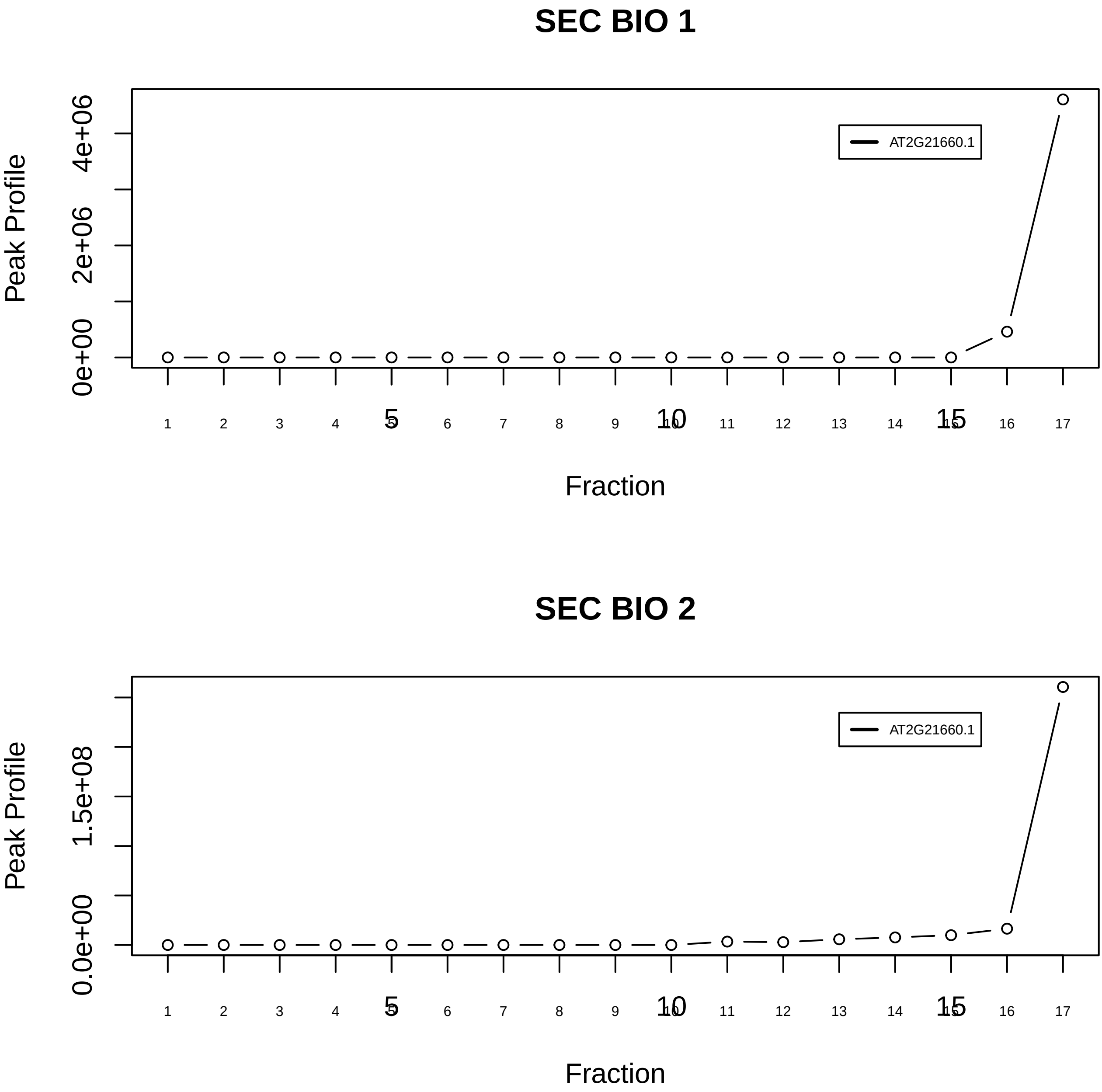

Spliceosome|351−−singleton_1

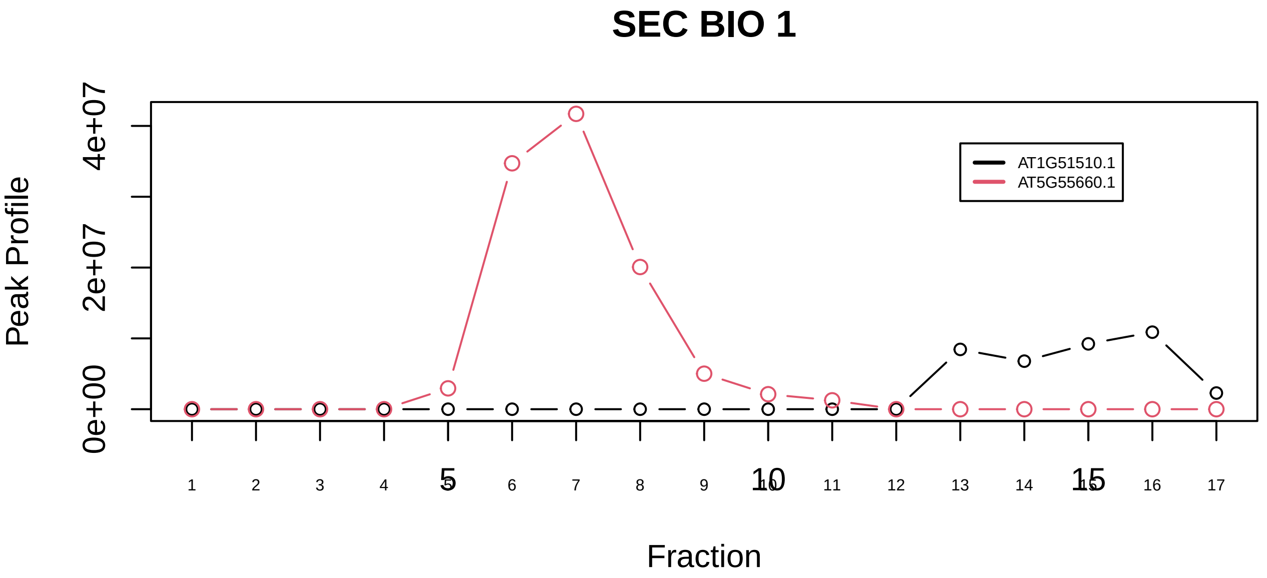

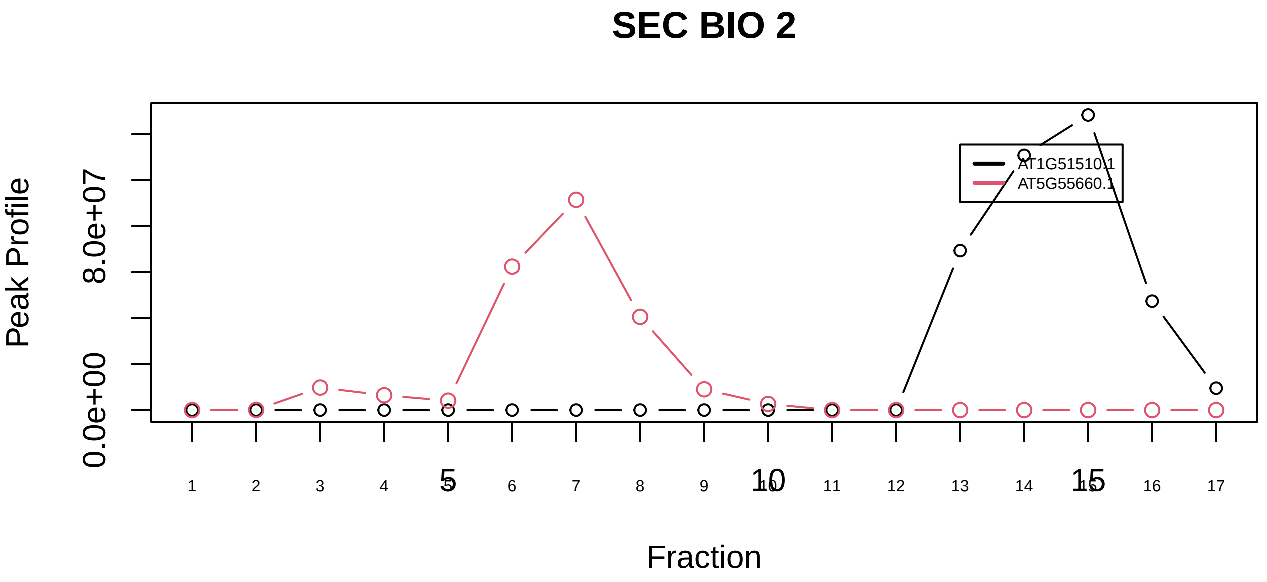

Splicing−associated factors complex|5449−−only_2_prots_before_clustering

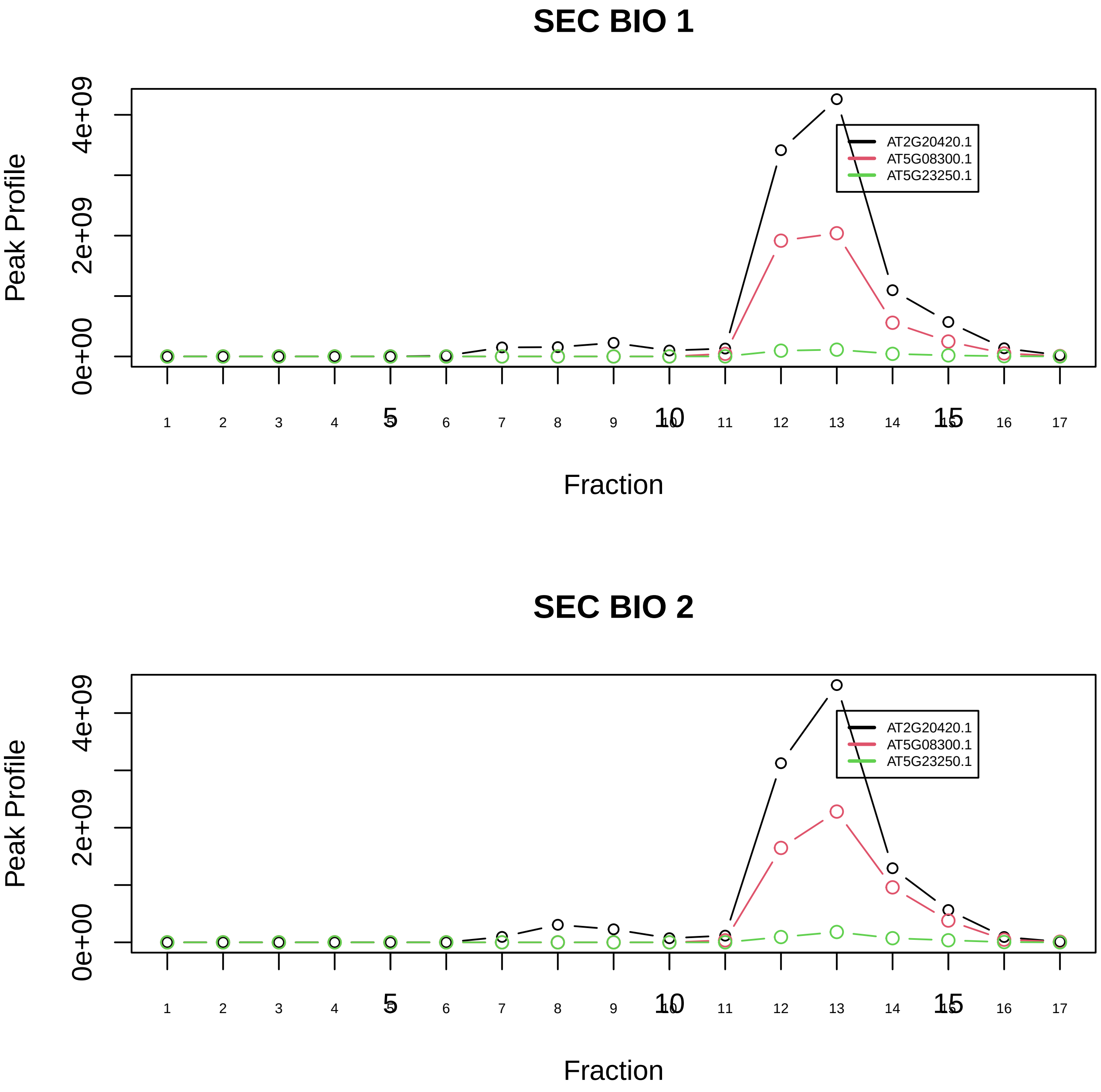

Succinyl−CoA synthetase, ADP−forming|394−−all_prots_before_clustering_GPFdist

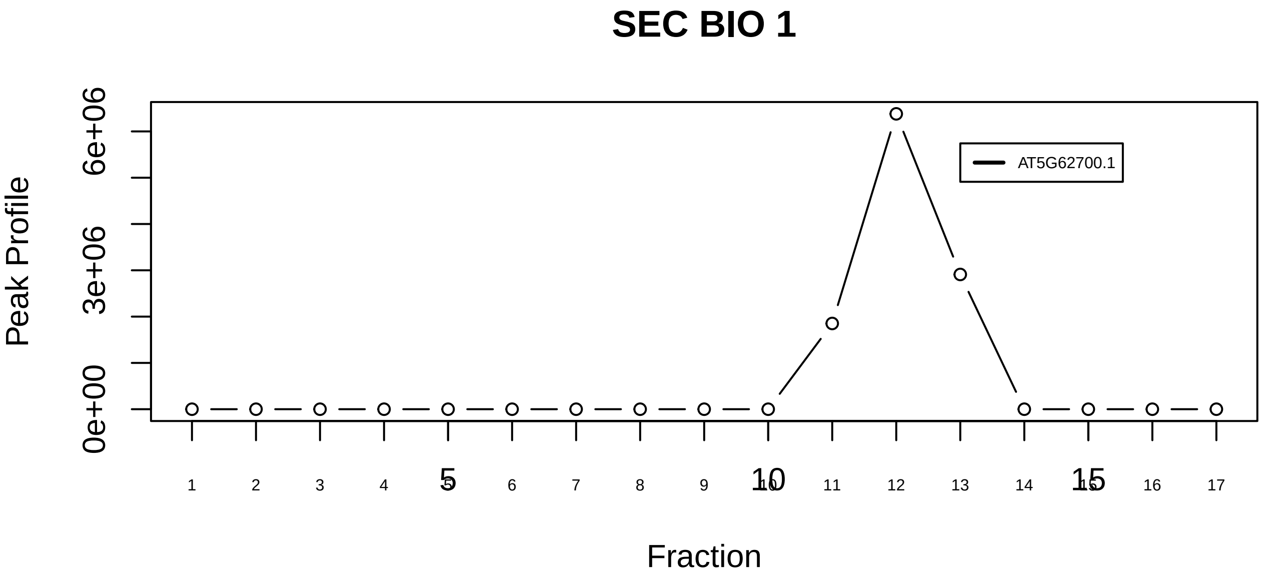

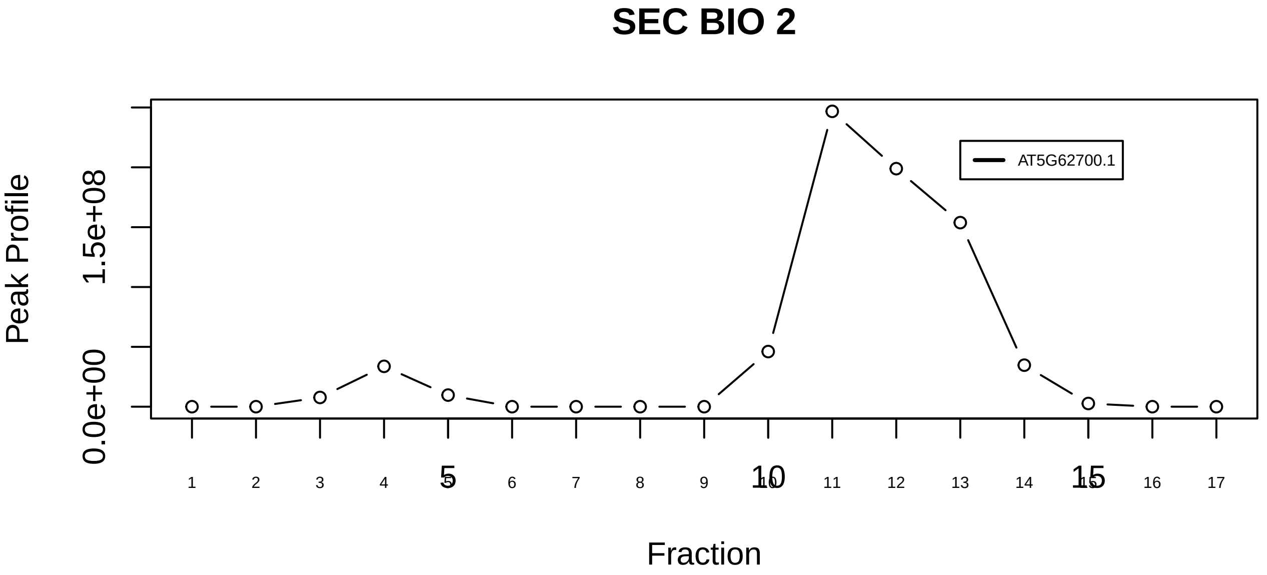

TBCD−tubulin(alpha)−tubulin(beta) complex|6703−−singleton_1

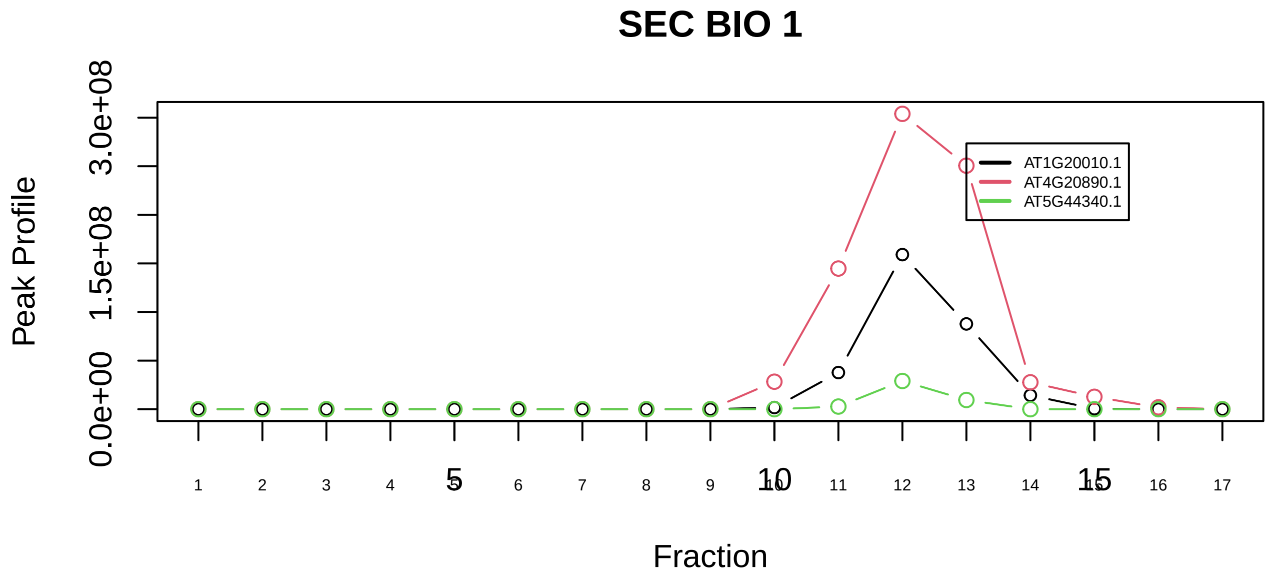

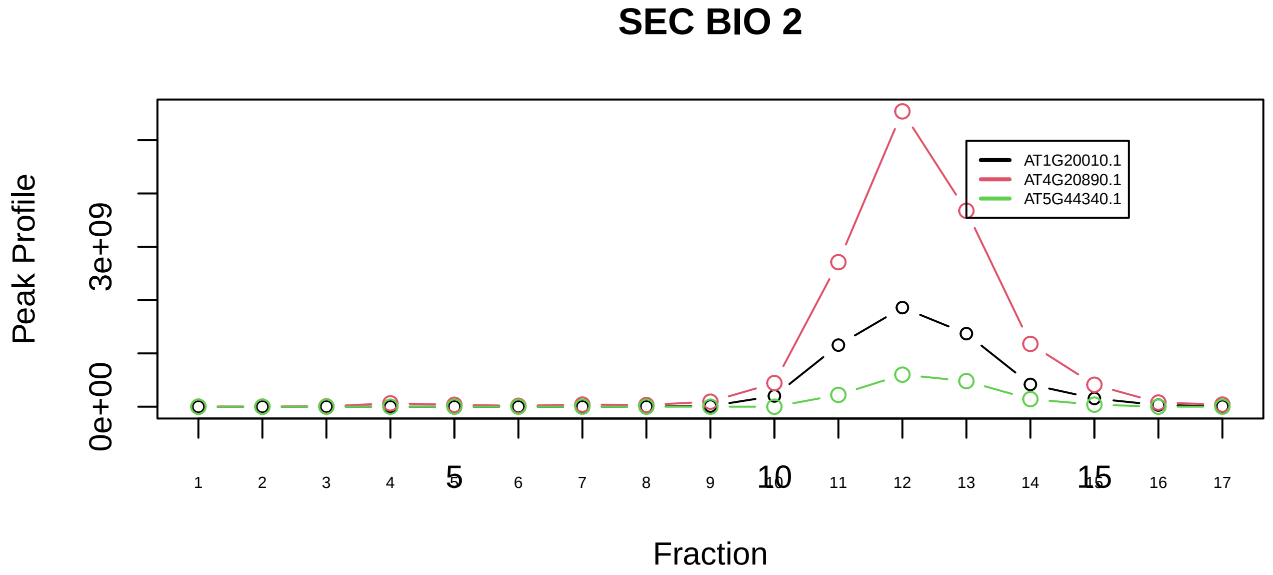

TBCD−tubulin(alpha)−tubulin(beta) complex|6703−−sub_orthoparalog_1

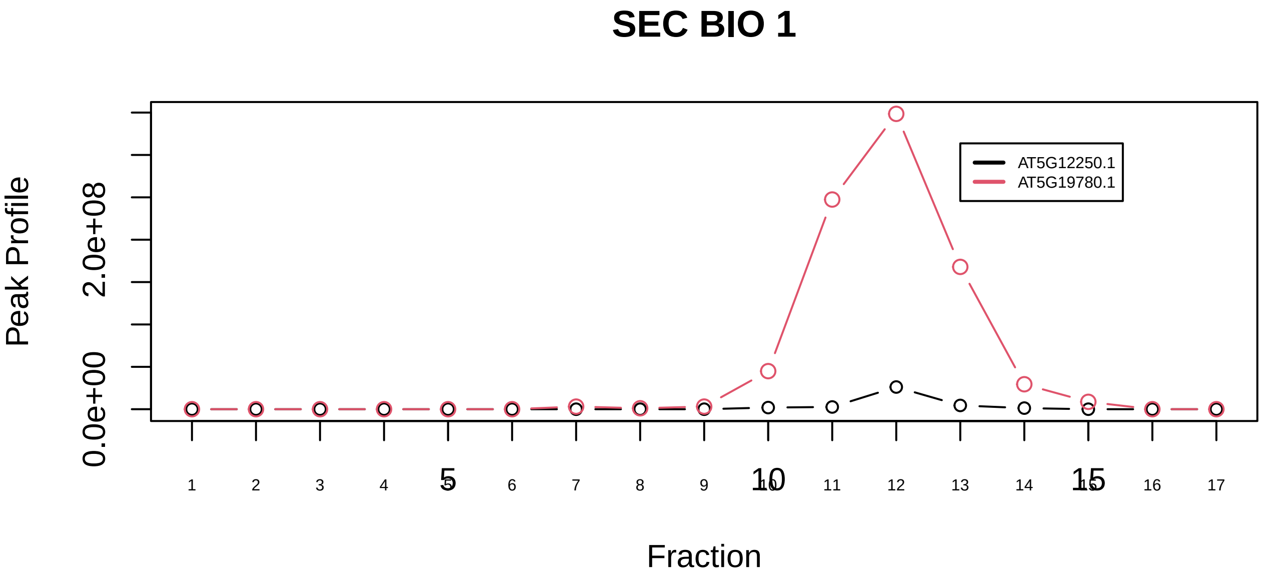

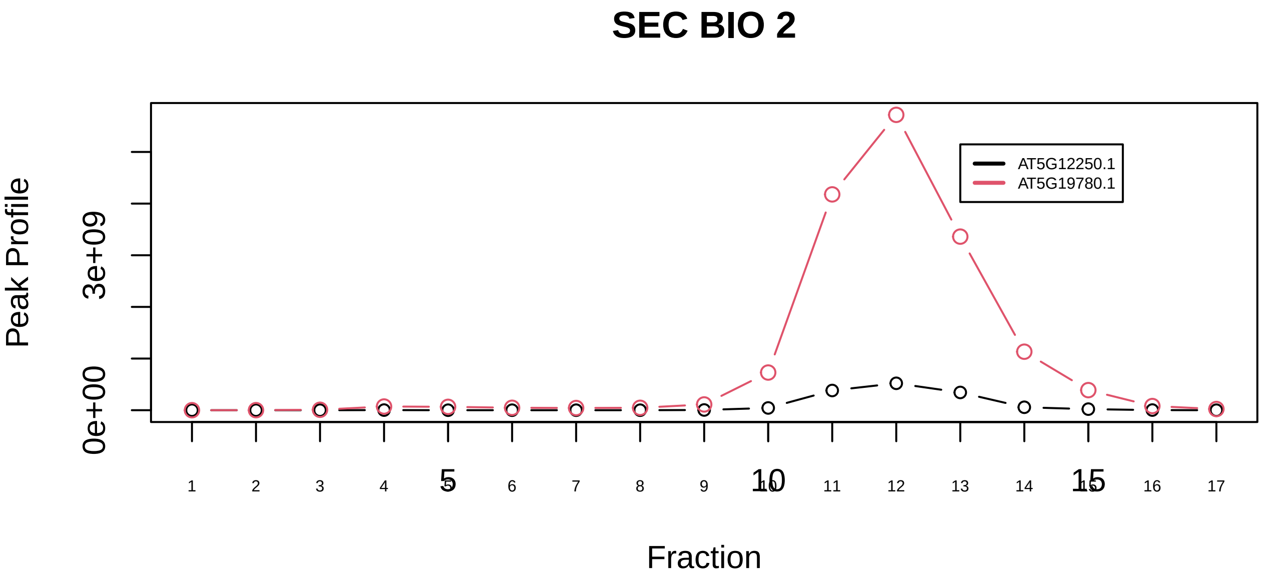

TBCD−tubulin(alpha)−tubulin(beta) complex|6703−−sub_1

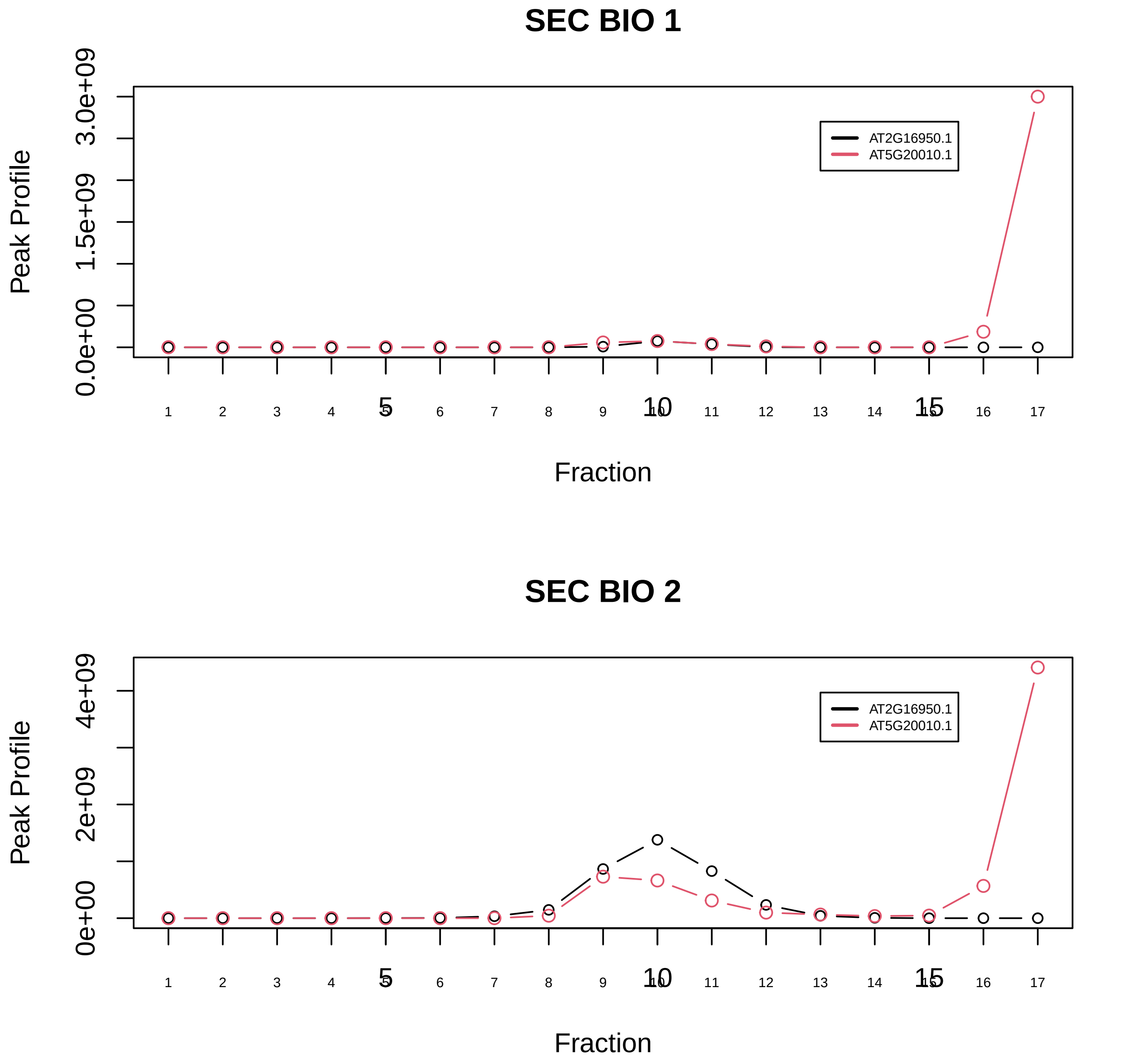

TNPO2−RAN−NXF1 complex|1552−−only_2_prots_before_clustering
